# Transcriptome analysis of *Plasmodium berghei* during exo-erythrocytic development

**DOI:** 10.1101/543207

**Authors:** Reto Caldelari, Sunil Dogga, Marc W. Schmid, Blandine Franke-Fayard, Chris J Janse, Dominique Soldati-Favre, Volker Heussler

## Abstract

The complex life cycle of malaria parasites requires well-orchestrated stage specific gene expression. In the vertebrate host the parasites grow and multiply by schizogony in two different environments: within erythrocytes and within hepatocytes. Whereas erythrocytic parasites are rather well-studied in this respect, relatively little is known about the exo-erythrocytic stages. In an attempt to fill this gap, we performed genome wide RNA-seq analyses of various exo-erythrocytic stages of *Plasmodium berghei* including sporozoites, samples from a time-course of liver stage development and detached cells, which contain infectious merozoites and represent the final step in exo-erythrocytic development. The analysis represents the completion of the transcriptome of the entire life cycle of *P. berghei* parasites with temporal detailed analysis of the liver stage allowing segmentation of the transcriptome across the progression of the life cycle. We have used these RNA-seq data from different developmental stages to cluster genes with similar expression profiles, in order to infer their functions. A comparison with published data of other parasite stages confirmed stage-specific gene expression and revealed numerous genes that are expressed differentially in blood and exo-erythrocytic stages. One of the most exo-erythrocytic stage-specific genes was PBANKA_1003900, which has previously been annotated as a “gametocyte specific protein”. The promoter of this gene drove high GFP expression in exo-erythrocytic stages, confirming its expression profile seen by RNA-seq. The comparative analysis of the genome wide mRNA expression profiles of erythrocytic and different exo-erythrocytic stages improves our understanding of gene regulation of *Plasmodium* parasites and can be used to model exo-erythrocytic stage metabolic networks and identify differences in metabolic processes during schizogony in erythrocytes and hepatocytes.

## Introduction

Malaria is a devastating disease caused by apicomplexan parasite *Plasmodium species*. Almost half of the world’s population is permanently at risk of malaria resulting in over 200 Million malaria cases worldwide mostly in African countries. There were more than 400,000 deaths in 2017 (1), majority of which were of children under the age of five.

The life cycle of *Plasmodium* parasites involves the injection of sporozoites into the vertebrate host during a blood meal of an infected female mosquito. For the rodent parasite *P. berghei* it has been shown that a proportion of injected sporozoites actively invade blood vessels and then are passively transported to the liver (2). After crossing the blood vessel endothelia in the liver to reach the parenchyma, the parasite transmigrates through several hepatocytes before it settles in one. While entering the ultimate host cell, the host plasma membrane invaginates forming a parasitophorous vacuole (PV) in which the parasite resides, develops and multiplies by exo-erythrocytic schizogony. The intracellular parasite extensively remodels the parasitophorous vacuole membrane (PVM), in particular by excluding or removing host cell proteins and incorporating parasite proteins (3).

Each exo-erythrocytic stage parasite (EEF: exo-erythrocytic form) generates tens of thousands of nuclei by the process of exo-erythrocytic schizogony. This rapid nuclear division is accompanied by growth and replication of organelles including the Golgi apparatus, endoplasmic reticulum, mitochondrion and apicoplast and by the vast expansion of the plasma membrane (4–6). Nuclei and organelles are eventually segregated into individual merozoites. Once EEF merozoites have completed their development, the PVM ruptures. This process requires an orchestrated action of multiple *Plasmodium* proteins such as lipases (e.g. PbPL (7)) and proteases (e.g. SUB1 (8,9) and possibly perforins (as shown for erythrocytic stage parasites (EF: erythrocytic form) (10–12). Upon rupture of the PVM, EEF merozoites disperse in the host cell cytoplasm and the host cell actin cytoskeleton collapses (13). *In vitro*, the final developmental stage of the EEF are detached cells (DC) and merosomes, host cell plasma membrane enclosed merozoites (14,15). *In vivo*, upon PVM rupture, infected cells become excluded from the liver tissue and merosomes are formed and pushed into the lumen of adjacent blood vessels. At tissue sites with small capillaries, merosomes rupture to release merozoites into the blood (16). Liberated merozoites immediately invade red blood cells (RBC), where they undergo repeated asexual reproduction cycles. In contrast to the tens of thousands merozoites generated by a single EEF, within erythrocytes the parasites produce only a limited number (12 to 32) merozoites by erythrocytic schizogony. Another difference between erythrocytic and exo-erythrocytic schizogony is that after rupture of the PVM the EEF merozoites can reside in the host cell cytoplasm for up to several hours, whereas EF merozoites are liberated from the PVM and the host cell plasma membrane almost simultaneously (17). Some EF will differentiate into male and female gametocytes. When these are ingested by a mosquito during a blood meal, they mature into macro-and micro-gametes and are liberated from the RBC. These sexual forms fuse to form zygotes and transform into motile ookinetes, which are able to cross the midgut epithelium of the mosquito to develop into oocysts, in which thousands of sporozoites are formed. After about 9-16 days (depending on the parasite species and the environmental temperature), sporozoites are liberated and invade the salivary glands of the mosquito (reviewed in (18)), whereupon they are then ready to be injected into a host during the next blood meal.

Studying the entire life cycle using human parasites under live or laboratory conditions is difficult due to ethical and safety reasons. However, the use of model organisms, such as the rodent malaria parasite *P. berghei is* experimentally tractable, to investigate the EEF development and fill gaps of knowledge that might also be relevant for human *Plasmodium* species. In addition, genetic manipulation of *P. berghei* is relatively easy and well established (19–21).

Many transcriptomic studies have been undertaken for different *Plasmodium* species, either by array or RNA-seq. **Table 1** lists transcriptomic data implemented in PlasmoDB (22). Recently, single cell transcriptomic profiles have also been published for EF of *P. falciparum* (23,24). However, of all of these studies, only two include the transcriptome of EEF parasites, one of which was done using the rodent parasite *Plasmodium yoelii* (25) and the other using *P. vivax* with emphasis on the dormant parasite stage, the hypnozoite (26). Since *P. berghei* is a widely used model in malaria research and quantitative data are missing for its EEF stages, we performed genome wide RNA-seq analyses of EEF development in a time course and compared expression data to already published data of gene-expression of other life cycle stages of *P. berghei*. In particular, we compared EEF merozoites originating from DC and EF merozoites (from *in vitro* cultivated schizonts). Remarkably, we found that their transcription profiles differ substantially and identified differences in metabolic processes during schizogony in erythrocytes and hepatocytes despite the fact that both types of merozoites infect RBC.

**Table 1:** Genome wide transcriptome data from studies implemented in PlasmoDB (release40; www.plasmodb.org). RNA-seq analyses, micro-array analyses and Single/DropSeq analyses are shown

**RNA-seq**
| ***Plasmodium* species** | **Life cycle stages** | **reference** |
| --- | --- | --- |
| *P. berghei* ANKA | 5 asexual and sexual stage transcriptomes | (31) |
| *P. berghei* ANKA | Female and male gametocyte | (80) |
| *P. chabaudi chabaudi* | Trophozoite transcriptomes after mosquito transmission or direct injection into mice | (81) |
| *P. yoelii yoelii* 17X | Salivary gland sporozoite transcriptomes: WT vs. Puf2-KO | (82) |
| *P. falciparum* 3D7 | NSR-seq Transcript Profiling of malaria-infected pregnant women and children | (83) |
| *P. falciparum* 3D7 | Polysomal and steady-*P. berghe* state asexual stage transcriptomes | (84) |
| *P. falciparum* 3D7 | Blood stage transcriptome (3D7) | (85) |
| *P. falciparum* 3D7 | Transcriptomes of 7 sexual and asexual life stages | (86) |
| *P. falciparum* 3D7 | Intraerythrocytic cycle transcriptome (3D7) | (Hoeijmakers et al.) NOT published |
| *P. falciparum* 3D7 | Strand specific transcriptomes of 4 life cycle stages | (86) |
| *P. falciparum* 3D7 | Transcriptome during intraerythrocytic development | (87) |
| *P. falciparum* 3D7 | Mosquito or cultured sporozoites and blood stage transcriptome (NF54) | (Hoffmann et al.) NOT published |
| *P. falciparum* 3D7 | Female and Male Gametocyte Transcriptomes | (32) |
| *P. falciparum* 3D7 | Ring, Oocyst and Sporozoite Transcriptomes | (88) |
| *P. vivax* P01 | Hypnozoite RNAseq | (26) |
| *P. vivax* P01 | Transcription profile of intraerythrocytic cycle | (89) |
| *P. cynomolgi* strain M | *P. cynomolgi* transcriptome of whole blood and bone marrow collected during 100-day infection of M. mulatta | (90) |

**Array**
| ***Plasmodium* species** | **Life cycle stages** | **reference** |
| --- | --- | --- |
| *P. berghei* ANKA | DOZI Mutant Transcript Profile | (33) |
| *P. berghei* ANKA | Transcript Profiling of Developmental Stages - High Producer (HP/HPE) | (91) |
| *P. berghei* ANKA | AP2-G2 knock out and WT expression profiles | (92) |
| *P. yoelii yoelii* 17X | Liver, mosquito and blood stage expression profiles | (25) |
| *P. yoelii yoelii* 17X | Life Cycle Stages | (93) |
| *P. falciparum* 3D7 | Pfal3D7 real-time transcription and decay | (Manuel Llinas) NOT published |
| *P. falciparum* 3D7 | Invasion pathway knockouts | (94) |
| *P. falciparum* 3D7 | Three Isogenic Lines w/ CQ Treatment: expression profiles | (95) |
| *P. falciparum* 3D7 | Life cycle expression data (3D7) | (96) |
| *P. falciparum* 3D7 | Sexually vs asexually committed schizont transcriptional profiles | (97) |
| *P. falciparum* 3D7 | Gametocyte stages I-V transcriptomes | (98) |
| *P. falciparum* 3D7 | Two Sir2 KO lines expression profiling | (99) |
| *P. falciparum* 3D7 | Asexual blood stage transcriptomes of clonal strains | (100) |
| *P. falciparum* 3D7 | Erythrocytic expression time series (3D7, DD2, HB3) | (101,102) |
| *P. falciparum* 3D7 | Microarray Expression from Patient Samples | (103) |
| *P. falciparum* 3D7 | mRNA Half life | (104) |
| *P. vivax* P01 | Intraerythrocytic developmental cycle of three isolates | (105) |
| *P. vivax* P01 | Sporozoite Expression Profiles | (106) |
| *P. knowlesi* strain H | Intraerythrocytic cycle expression profile: in vitro and ex vivo (Pkno PK1(A+)) | (107) |

Single cell / DropSeq
| ***Plasmodium* species** | **Life cycle stages** | **reference** |
| --- | --- | --- |
| *P. falciparum* 3D7 | DropSeq of single *P. falciparum* infected erythrocytes | (23) |
| *P. falciparum* 3D7 / *P. berghei* | Single cell RNA from *P. berghei* & *P. falciparum* infected erythrocytes | (24) |

## Results and Discussion

### High fidelity exo-erythrocytic stage RNA-seq data and sample selection criteria

We infected HeLa cells with *P. berghei* sporozoites that express mCherry under the control of a constitutive Hsp70 promoter, allowing the detection of fluorescent parasites in all developmental stages (7). Exo-erythrocytic form (EEF) parasites were isolated at different time-points of infection by FACS sorting (6 hours (h), 24 h, 48 h, 54 h and 60 h). At 69 h detached cells (DC) were collected from the culture medium supernatant. To preserve RNA integrity during FACS sorting of infected cells, cells were treated with RNAlater (27,28). In addition, sporozoite samples were generated by processing infected salivary glands of mosquitoes at day 20 post feeding. For each sample independent biological duplicates were collected. Following the isolation of RNA from infected HeLa cells and from infected salivary glands, libraries were sequenced with an Illumina HiSeq 2500 resulting in 34 to 61 million paired-end reads per sample (**Table S1**). After removal of low quality sequences, sequencing adapters and sequences arising from host RNA, reads were aligned to the *P. berghei* ANKA reference genome, resulting in around 0.23 to 21.4 million weighted alignments (in the following termed “hits”, see (29) and (30) for details) within genic regions (**Table S1**, raw counts are provided in **Table S2**). Samples collected 6 hours after infection were excluded from further analysis as only low amounts of hits were recovered (22,154 and 35,312 hits) in both biological replicates. The reason is most likely that, at this time-point of infection, parasite transcripts represent only a small fraction in comparison to host cell transcripts.

We identified 4475 transcribed genes (≥80 normalized read counts, corresponding on average to 20 RPKMs) in at least one developmental stage of the EEF. In a previously reported transcriptome analysis of *P. yoelii* EEF stages using array technology, about 2000 genes were detected (25). This exemplifies the higher sensitivity of the Next Generation Sequencing (NGS) compared to array technology (29).

A hierarchical clustering of the different EEF samples together with RNA-seq data of EF (rings, trophozoites and schizonts harvested 4, 16 and 22 hours after infection of RBC), as well as with RNA-seq data of gametocytes and ookinetes (31) was done. The replicates correlated (Spearman’s correlation) highly to each other and the different stages grouped well according to the host environment (exo-erythrocytic, erythrocytic) (**Fig. 1)**.

**Figure 1:**
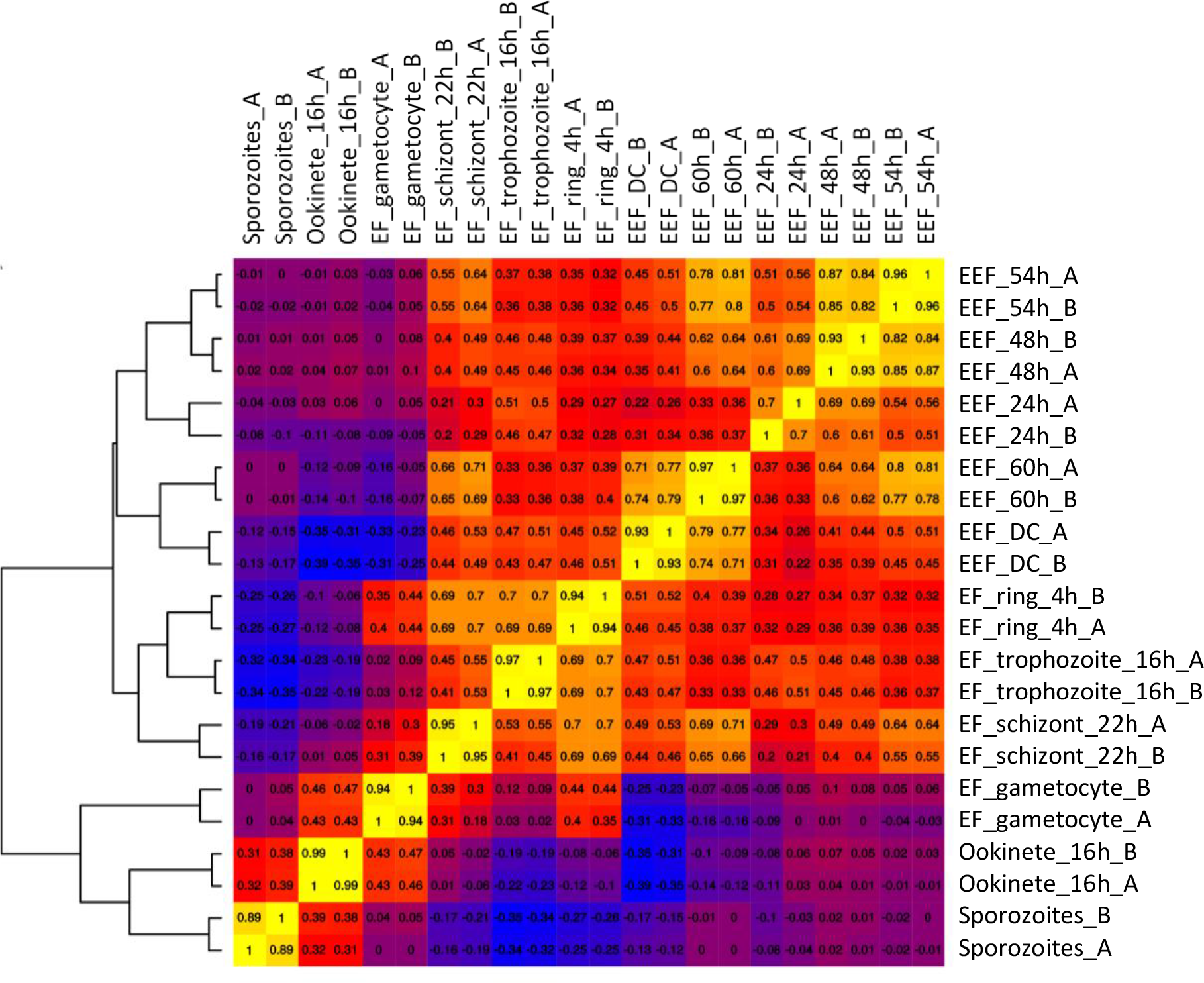
Clustering of samples of different life cycle stages based on genes with the highest overall high variance (90th percentile, Spearman correlation and hierarchical clustering). Stages are sporozoites, exo-erythrocytic (EEF) stages (DC are detached cells), erythrocytic (EF) stages (rings, trophozoites, schizonts, gametocytes) and ookinetes. _A, _B: Biological replicates. Heatmap was generated using normalized and log2(x+1)-transformed gene expression values (74). Heatmap drawn with the R-package gplots (108).

The analysis revealed that the mRNA expression profiles of the extracellular ookinetes and sporozoites cluster together, which might be due to the fact that both are motile stages that traverse mosquito host cells. Ookinetes and sporozoites are, however, markedly different from the profiles of EEF and asexual EF stages. Notably, the expression profile of gametocytes is different from both EEF and asexual EF stages but shows similarities to the ookinetes. This is not surprising as gametocytes and ookinetes are consecutive stages in the life cycle. In female gametocytes many different mRNAs are already produced that are translationally repressed and are only used during development of the zygote and ookinete (32–34). Among the EEF stages, the highest similarities of mRNA expression profiles were observed as expected for adjacent stages/timepoints (e.g., 48 h and 54 h). However, the early stages/timepoints (24 h, 48 h and 54 h) were fairly distinct from the later time-points (60 h and detached cells). The asexual EF stages showed a rather high degree of similarity among them. In fact, the mRNA expression profile of detached cells, containing the EEF merozoites, was rather distinct from the profile of EF merozoites, although both need to be prepared to invade RBC. It is noteworthy that proteomics data of *P. yoelii*, revealed that 90% of the proteins of late EEF were also detected in the early EF (25).

To verify the RNAseq data, we compared the expression pattern of selected genes to previously reported expression patterns. The expression pattern of genes coding for housekeeping proteins (GAPDH, actin 1 and alpha-tubulin 1; **Fig. S1**), putative proteases (SERA 1 to 5; **Fig. S2**), PVM proteins (EXP1, Exp2, UIS3 and UIS4; **Fig. S3**), sporozoite surface proteins (CSP and TRAP; **Fig. S4**), fatty acid biosynthesis enzymes (FabB/F, FabI, FabZ and FabG, **Fig. S5**) as well as merozoite surface proteins (MSP 1, 4/5, 7, 8, 9, 10; **Fig. S6**) were very similar to the already published data confirming the quality of the here presented RNA-seq data. For further information about the selected genes presented in **Fig. S1** to **Fig. S6**, the interested reader is referred to the supplementary information section.

An important aspect of the current RNA-seq analysis was that the expression profiles provide valuable information for the choice of promoters to drive expression of transgenes, such as fluorescent or luminescent reporter proteins. Previously, the promoters of the housekeeping genes heat shock protein (*hsp70*) and eukaryotic elongation factor 1α (e*ef1α*) have been used to drive expression of fluorescent reporters (7,35,36). According to our RNA-seq analysis, the *hsp70* promoter is a better choice for driving constitutive expression of reporters as *hsp70* mRNA exhibits a more uniform expression profile compared to e*ef1α* mRNA (**Fig. S7**).

We next aimed for a more detailed computational analysis of the exo-erythrocytic stage transcriptome and a comparison with other developmental stages.

### Gene co-expression network

To further explore the complexity of the parasite transcriptome, in particular the gene expression similarities among the different developmental stages, a gene co-expression network (GCN) was computed (37) and genes with similar expression patterns (“communities”)(38) were extracted and visualized (**Fig. 2**). With this analysis, we gained insight into similarly expressed genes in the different EEF and EF stages and into characteristic expression patterns within the entire transcriptome. We used these analyses to find functionally related genes based on similar expression patterns.

**Figure 2:**
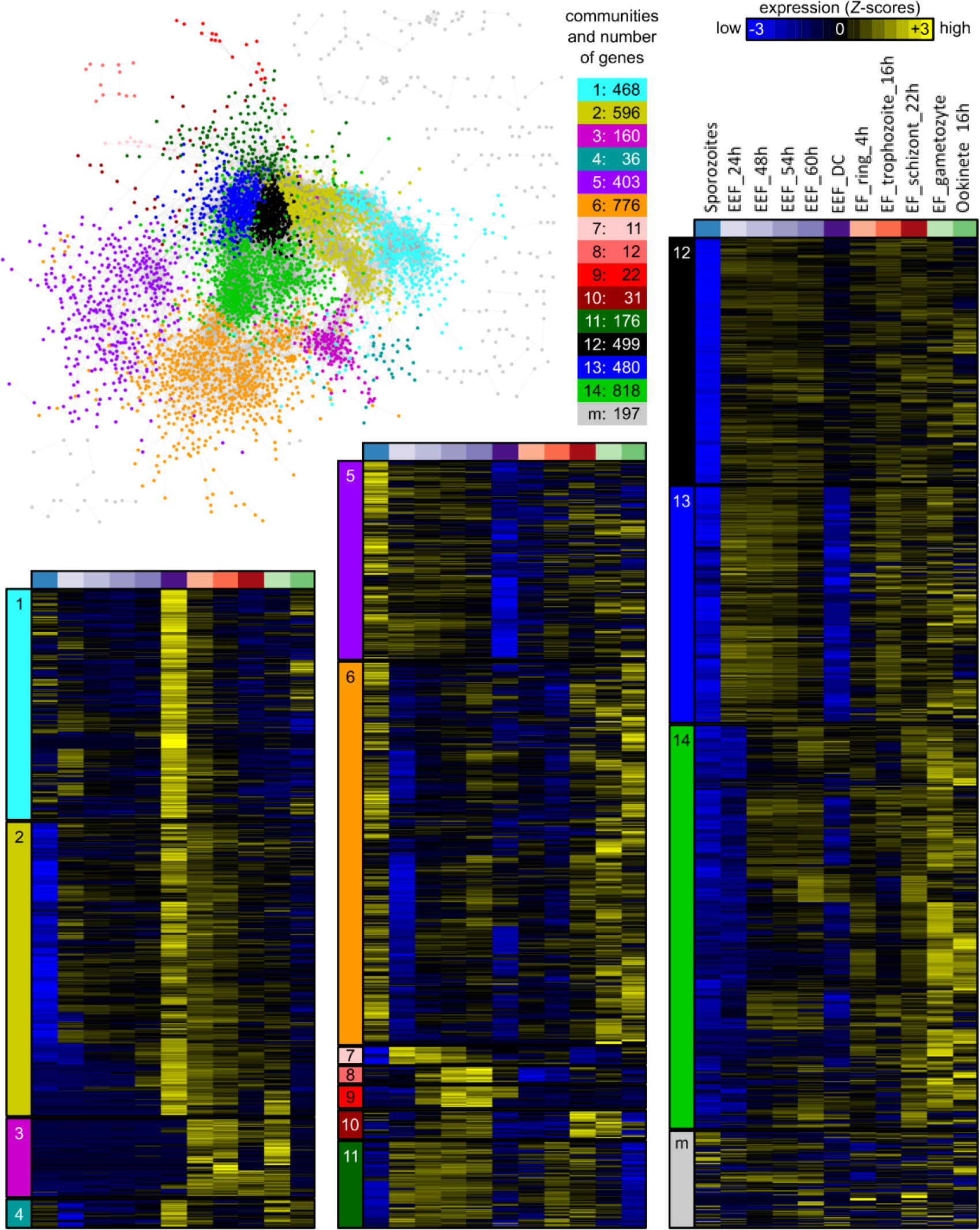
Gene co-expression network based on RNA-seq data of samples of different life cycle stages. Stages are sporozoites, exo-erythrocytic (EEF) stages (DC are detached cells), erythrocytic (EF) stages (rings, trophozoites, schizonts, gametocytes) and ookinetes. Each node represents a gene and each edge depicts a significant pairwise correlation. The network was visualized with Cytoscape (77) using the “prefuse force directed layout”. Nodes/genes are colored according to their membership in 14 communities and a ‘mixed’ (M) community (pool of communities with less than 11 genes per community), identified with a modularity optimization algorithm (38). For each community, a heatmap summarizes the expression patterns of all genes within the community. Expression values in the heatmaps correspond to gene-wise *Z*-scored of normalized and log2(x+1)-transformed count data averaged across the replicates.

The GCN analysis revealed 14 different communities comprised of 11 to 818 genes and a “mixed” community with 197 genes (a pool of communities with 10 or less genes per community) (**Fig. 2**, **Table S4**). Of a total of 5104 genes, 4675 genes are represented in the GCN. 429 genes did not meet the GCN criteria, as the expression pattern of each of them did not significantly correlate to the expression pattern of another gene throughout all stages. Interestingly, the GCN analysis indicated marked differences between EEF and EF. Surprisingly, even the transcriptome of EEF-derived and EF-derived merozoites (i.e., detached cells and late blood schizonts) were found to differ substantially. The 486 transcribed genes of community 1 were strongly expressed in DC, which contain EEF-derived merozoites and extended into the ring stage (initial phase of EF development). The 596 transcribed genes of community 2 were as well enriched in DC, but expression of these genes persisted longer during the EF (into trophozoite and partly into schizont stage) whereas expression was strongly reduced in the sporozoites. To functionally characterize the communities defined in the GCN, we tested for enrichment of gene ontology (GO) terms from the domain “Biological Process (BP)” (**Fig. 3**, **Table S3** for additional information). The first two communities in **Fig. 2** contain genes that are expressed specifically in DC and EF stages. The majority of the corresponding gene products were predicted to be involved in gene expression, ribosome biogenesis and transcription (Community 1, **Fig. 3**) or in mRNA processing and translation (Community 2, **Fig. 3**). All these functions are involved in DNA/RNA biology, in particular in (regulation of) gene expression. This is not surprising as invasive forms like the merozoites in DC prepare for the next growth phase after invasion and most likely need stored transcripts for a rapid protein synthesis after invasion of erythrocytes, comparable to storage of (repressed) transcripts in mature gametocytes and sporozoites (32–34,39–43). On the other hand, the 160 genes of community 3 were highly specific to the EF stages (**Fig. 3**). Conspicuously, this community has almost no GO-term annotations for biological processes (only 3 out of 160 genes were annotated). However, this community was highly enriched for small nucleolar RNAs (snoRNAs), PIR pseudogenes (*Plasmodium* interspersed repeat pseudo genes) and genes of the three large multigene families in rodent parasites, coding for PIR proteins, fam-a proteins and fam-b proteins. SnoRNAs are important components of ribosome biogenesis. They are non-coding RNAs with a diversity of function like pseudo-uridylation and 2’-O-methylation of RNAs or synthesis of telomeric DNA (44). PIR, fam-a and fam-b proteins are exported by EF stages into the cytoplasm of the host erythrocyte. The function of most of these proteins is unknown, although evidence has been presented for a role of PIRs in erythrocyte sequestration. Recently it has been shown that a subset of PIR, fam-a and fam-b proteins are also expressed in EEF stages (45).

**Figure 3:**
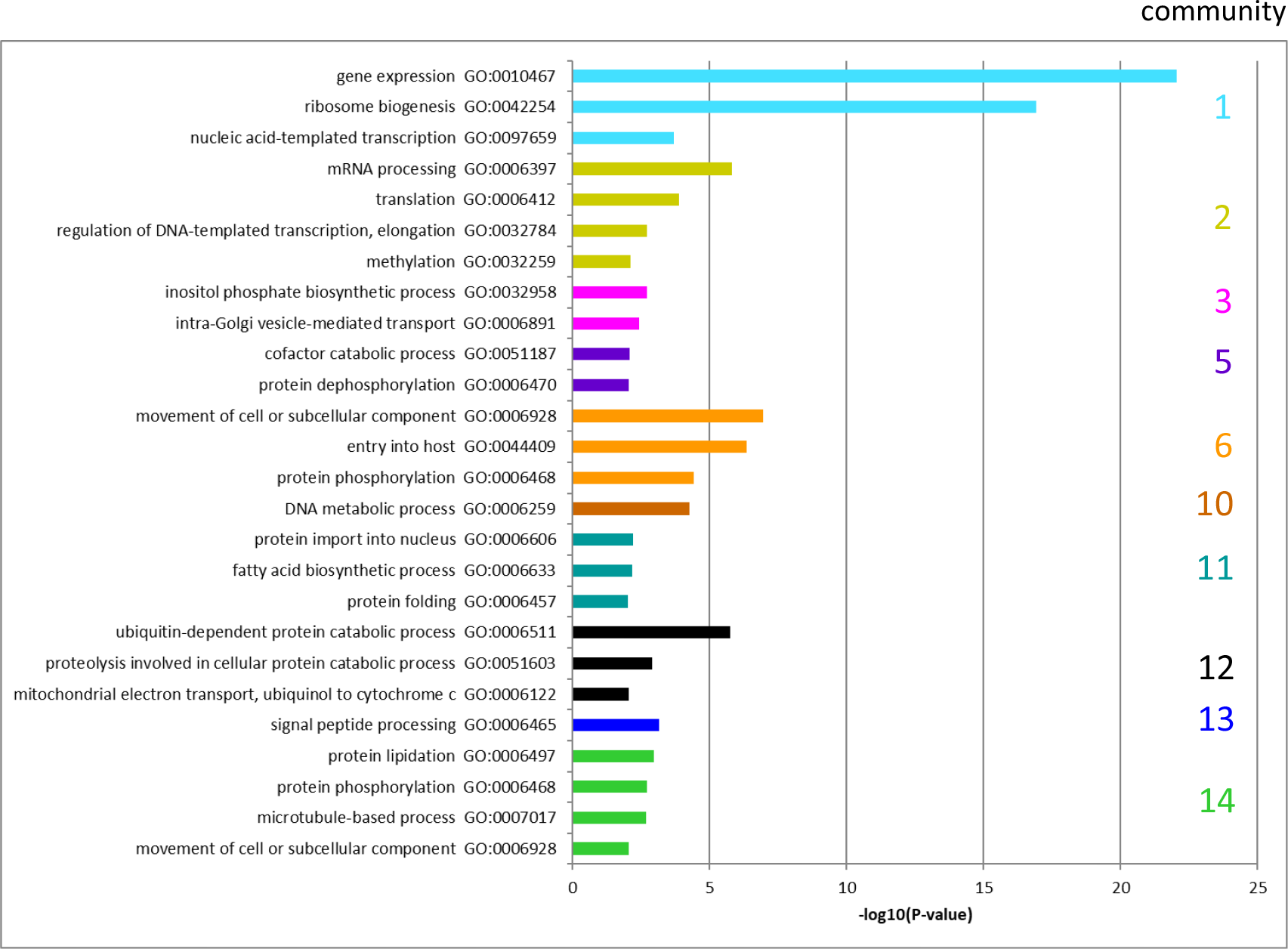
GO terms in GCN communities (expressed as −log10(P-value)). The colors refer to the different communities in Fig1 B. Only GO-terms with p-values < 0.01 were included. Communities containing less than 25 genes were ignored because of potential false significance (following recommendations in GeneSetEnrichmentAnalysis from the Broad Institute).

Community 6, consisting of 776 genes, was enriched for genes expressed in sporozoites, but also frequently expressed at elevated levels in gametocytes, ookinetes and schizonts. Not surprisingly, genes preferentially expressed in sporozoites, gametocytes and ookinetes (community 6, **Fig. 3**) were involved in host cell entry, host cell exit and parasite motility. In addition, genes that are involved in transport of subcellular components and DNA repair were present. Genes whose expression was found to be more specific to sporozoites, with persisting expression during the early EEF stages (community 5, **Fig. 3**), were involved in DNA synthesis and metabolic processes, consistent with the high multiplication observed following hepatocyte invasion by the sporozoites. Communities 7, 8 and 9 contain 45 genes in total, the expression of which were mostly specific to the developing EEF stages. According to guidelines of the Broad Institute on GeneSetEnrichmentAnalysis, small size communities should not be interpreted.

The expression of the 31 genes of community 10 was enriched in EF schizonts and in gametocytes, but these genes were also found well expressed in late EEF stages. Although the GO term ‘DNA metabolic process’ is listed for this community, it should be assessed with caution due to the reason mentioned above.

The 176 genes of community 11 were expressed during the developing EEF and EF stages, with a slight bias towards the developing EEF stages (85% of all genes in the community were on average expressed at a higher level in the EEF stages). In this community, 3 out of 9 genes of fatty acid biosynthesis have been identified as hits. This is in agreement with the high fatty acid usage of EEF stages to generate various parasite membranes (46). Apart from genes involved in fatty acid biosynthesis, it is very likely that genes identified in this community are involved in schizogony and merozoite development in both EEF and EF stages.

The remaining communities were mostly defined by genes with almost complete absence of expression in sporozoites (community 12; 499 genes), in sporozoites and DC (community 13; 480 genes) or in sporozoites, 24 h EEF stage and partly DC (community 14; 818 genes). However, whereas genes of community 12 and 13 were generally expressed throughout the EEF and the EF stages, genes of community 14 were more specific to gametocytes and ookinetes (**Fig. 2**). Sporozoites are not growing or proliferating and therefore it can be expected that in sporozoites, expression of genes involved in several metabolic processes, protein lipidation, phosphorylation and signal peptide processing is less pronounced than in other stages.

Altogether, we could identify 14 clearly defined communities and a pool of small communities (mix) with totally 4675 genes attributed (see supplemental **Table S4** for GeneIDs of the members of the communities and for genes excluded during the GCN analysis).

The generated GCN provides a first comprehensive overview of gene regulation in a *Plasmodium* parasite throughout EEF and EF development including several life cycle stages in the mosquito vector (ookinetes, sporozoites). The identification of clear communities of genes with comparable expression profiles may help identifying common signatures in the untranslated promotor regions that may be involved in regulation of gene expression.

### Differences in gene expression between developing exo-erythrocytic and erythrocytic parasites

To better elucidate the differences between EEF and EF stage parasites, we performed differential expression analyses. Firstly, the average gene expression of developing EEF stages (24 h, 48 h, 54 h and 60 h) was compared with developing EF stages (ring 4h, trophozoite 16 h. In this comparison, 299 genes were significantly upregulated in the EEF stages (LogFC ≥ 2, adjP ≤ 0.01) and 392 genes were significantly upregulated in the EF stages (**Fig. 4**; **Table S5**). GO-term enrichment (summarized in **Table 2)** revealed that genes preferentially expressed in the EEF stages were enriched in fatty acid biosynthesis (e.g. *fabB*/*fabF* in **Fig. 4**), entry into the host cell, leading strand elongation, tetrapyrrol biosynthesis and DNA replication. Considering that a single *P. berghei* EEF stage parasite generates more than 10,000 merozoites and an individual EF stage parasite only 12-18, an enrichment of expression in genes associated with fatty acid biosynthesis and DNA replication is expected and has already been confirmed in different studies (47,48) and reviewed in (49). In contrast, genes preferentially expressed in the EF stages were enriched for the GO terms for cell motility and intracellular, organellar transport (summarized as “movement of cell or subcellular components”), protein export into host cell cytoplasm, exit from host cell and pathogenesis. Enrichment of gene expression related to translocation of proteins (e.g. membrane associated histidine-rich protein 1: *mahrp1a* & *b* in **Fig. 4**) across the PVM is also required for parasite remodeling of the host RBC (50), including proteins transported to the surface of the RBC involved in RBC sequestration (50–52). Many *P. berghei* proteins are known that are transported into the host red blood cell (45,52) whereas only a few proteins have been identified that are transported into the host hepatocyte, for example, CSP (53,54) and LISP2 (55).

**Figure 4:**
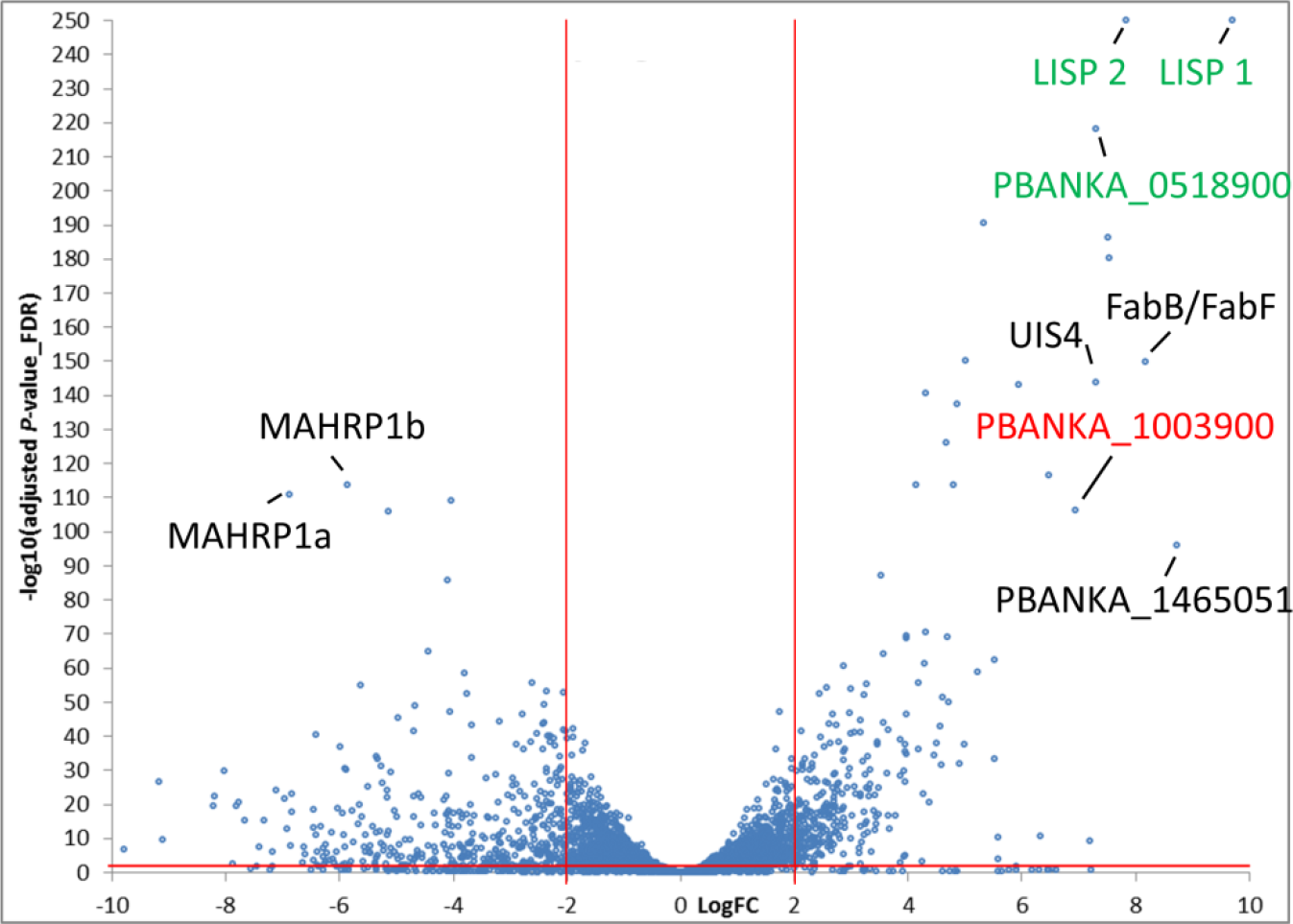
Volcano plot of differently expressed genes of developing exo-erythrocytic (EEF) stages (EEF_24h-60h) compared to developing erythrocytic stages (EF_ring, EF_trophozoites). In this analysis the DC and schizonts stages are not included. The graph shows LogFC values relative to FDR (-Log10(adjusted P-value). Positive LogFC values represent preferentially EEF stage expressed genes. Negative LogFC values represent preferentially EF stage genes. Genes in green and red show highest expression in liver stages compared to all other stages (including sporozoites, gametocytes and ookinetes).

**Table 2:**
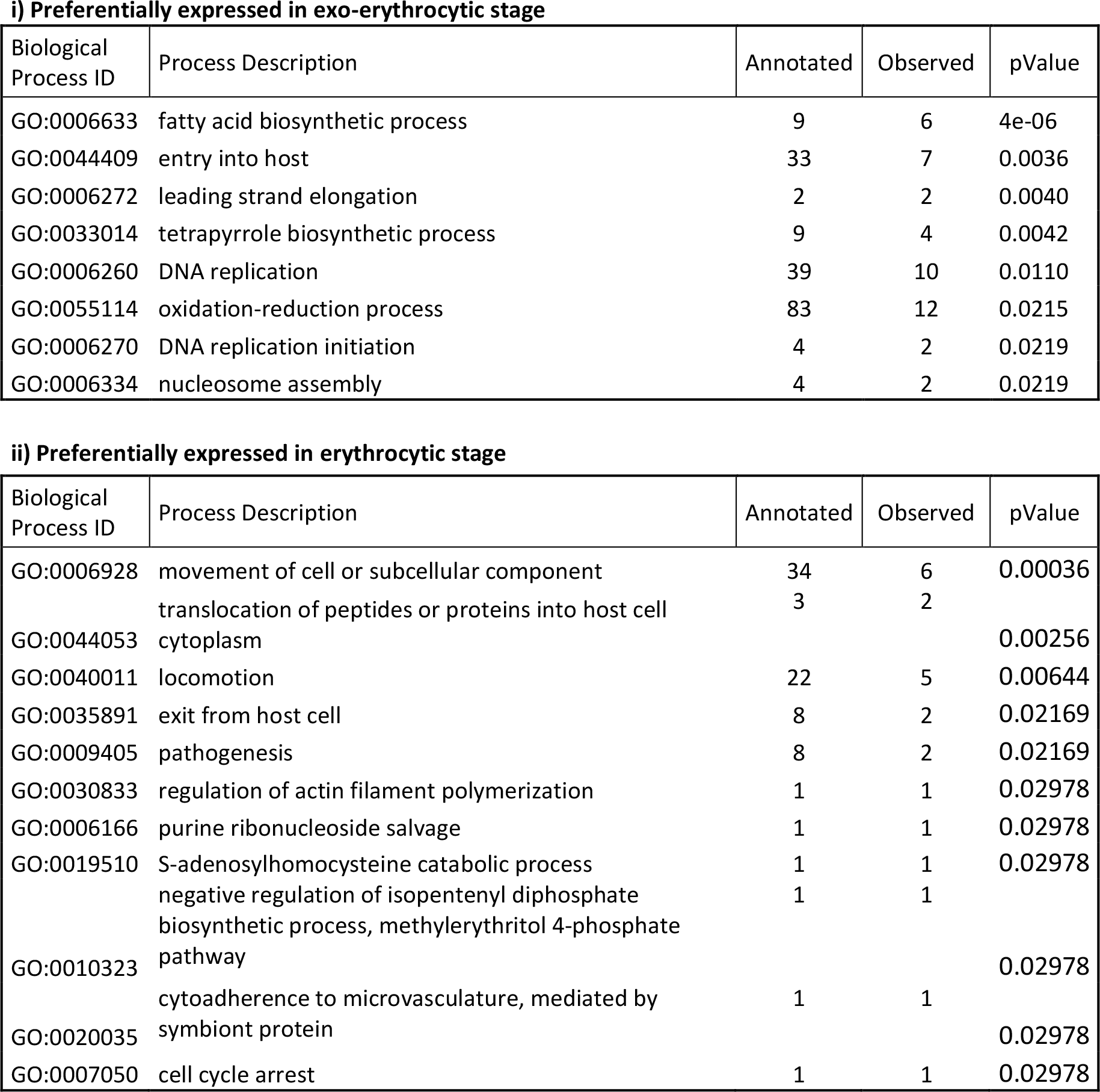
Gene ontology (GO) term annotation of genes preferentially expressed in i) EEF stages (24h, 48h, 54h and 60h) and ii) asexual EF stages (ring 4h, trophozoite 16h). The number of annotated genes (number genes per GO term present in the entire RNA-seq study) and observed genes (number of genes per GO term preferentially expressed in…) per Biological Process are listed. Only GO terms with p-values < 0.05 are shown.

### Exo-erythrocytic and erythrocytic merozoites: same but different

In contrast to the limited differences in gene expression seen during parasite development in hepatocytes and RBC, the differences between EEF and EF merozoites were much more pronounced (i.e. DC and the EF schizonts at 22 h) even though the 22h schizont sample was not entirely pure but also contained some immature schizonts and a small amount of gametocytes (31). 880 and 1275 genes were preferentially expressed in DC and in 22 h schizonts, respectively (**Fig. 5** and **Table S6**). Analysis of GO-term enrichment **(Table 3**) indicated clear differences between these sets. Genes preferentially expressed in the DC were found enriched for the GO-terms: gene expression, ribosome biogenesis, amide biosynthetic process, RNA biosynthetic process and mRNA splicing. In contrast, genes preferentially expressed in 22 h schizonts were enriched for the GO-terms: small GTPase-mediated signal transduction, DNA replication and recombination, protein localization and modification, and signal peptide processing. At first glance, it is rather surprising that merozoites derived from EEF and EF stage differ so markedly. However, it might reflect the fact that the mechanism of egress from their respective host cells is different. EF stage-derived merozoites almost simultaneously rupture the PVM and the plasma membrane of the RBC (17). On the other hand, EEF stage-derived merozoites are initially liberated from the PVM but can then stay for several hours in the host cell until they are extruded as merosomes, into a blood vessel (7,14) and eventually released in the fine capillaries of the lungs (16). Also, DC/merosomes do not form synchronously. *In vitro*, detached cell generation starts as early as 54 h, peaks at 65 h but continues until 69 h and even later. Since for the current study DC/merosomes were collected at 69 h to increase the yield, merozoites might be at different developmental and activation stages and thus might express different sets of genes. In the next section, we focus in more detail on genes involved in the egress of EEF stage-derived merozoites in comparison to EF stage-derived merozoites.

**Figure 5:**
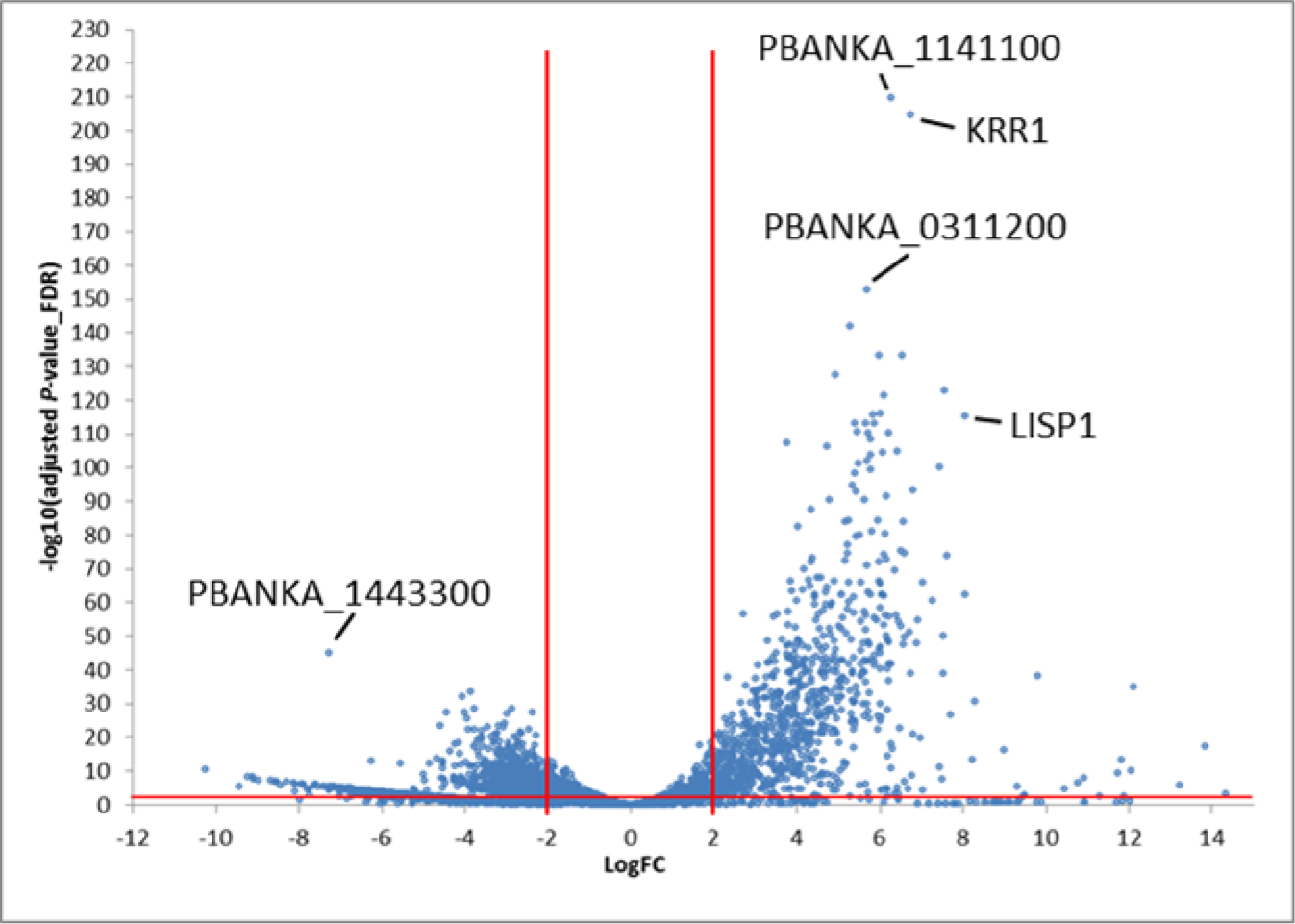
Volcano plot of differently expressed genes of detached cells (DC) compared to EF schizonts (sample 22h). Positive LogFC values represent preferentially DC expressed genes. Negative LogFC values represent preferentially EF schizont expressed genes. The graph shows LogFC values relative to FDR (-Log10(adjusted P-value).

**Table 3:** Gene ontology (GO) term annotation of genes preferentially expressed in i) EEF merozoites (DC) and ii) EF merozoites (22h schizonts). The number of annotated genes (number genes per GO term present in the entire RNA-seq study) and observed genes (number of genes per GO term preferentially expressed in…) per Biological Process are listed. Only GO terms with p-values < 0.05 are shown.

**i) Preferentially expressed in exo-erythrocytic stage merozoites**
| Biological Process ID | Process Description | Annotated | Observed | pValue |
| --- | --- | --- | --- | --- |
| GO:0010467 | gene expression | 462 | 181 | 3.0e-22 |
| GO:0042254 | ribosome biogenesis | 63 | 47 | 4.6e-20 |
| GO:0043604 | amide biosynthetic process | 219 | 98 | 1.1e-17 |
| GO:0032774 | RNA biosynthetic process | 103 | 35 | 0.00017 |
| GO:0017183 | peptidyl-diphthamide biosynthetic process from peptidyl-histidine | 5 | 4 | 0.00700 |
| GO:0000398 | mRNA splicing, via spliceosome | 43 | 15 | 0.01718 |
| GO:0008295 | spermidine biosynthetic process | 2 | 2 | 0.04105 |
| GO:0008612 | peptidyl-lysine modification to peptidyl-hypusine | 2 | 2 | 0.04105 |

**ii) Preferentially expressed in erythrocytic stage merozoites**
| Biological Process ID | Process Description | Annotated | Observed | pValue |
| --- | --- | --- | --- | --- |
| GO:0007264 | small GTPase mediated signal transduction | 26 | 14 | 0.0014 |
| GO:0006260 | DNA replication | 39 | 20 | 0.0049 |
| GO:0045184 | establishment of protein localization | 93 | 34 | 0.0060 |
| GO:0007017 | microtubule-based process | 35 | 15 | 0.0075 |
| GO:0000041 | transition metal ion transport | 7 | 6 | 0.0151 |
| GO:0006465 | signal peptide processing | 5 | 4 | 0.0153 |
| GO:0006848 | pyruvate transport | 3 | 3 | 0.0154 |
| GO:0006497 | protein lipidation | 21 | 10 | 0.0196 |
| GO:0006511 | ubiquitin-dependent protein catabolic process | 38 | 16 | 0.0248 |
| GO:0043248 | proteasome assembly | 8 | 5 | 0.0267 |
| GO:0006310 | DNA recombination | 9 | 6 | 0.0366 |
| GO:0006468 | protein phosphorylation | 82 | 28 | 0.0437 |

### Comparison of mRNA expression patterns at the end of exo-erythrocytic and erythrocytic stage development

Given that EF merozoite egress within minutes upon PVM rupture, whereas EEF merozoites remain in the hepatocyte cytoplasm for up to several hours, we searched for genes that might be i) necessary for the rapid egress specifically from RBC, ii) required for survival in the hepatocyte cytoplasm and iii) specific for both developmental stages (i.e. necessary for PVM rupture in RBC and hepatocytes). Interestingly, 74 genes were upregulated in 22h EF schizonts in comparison to EEF samples (LogFC ≥ 2, adjP ≤ 0.01, **Table S7**), but only four of these were upregulated when compared to all other stages (**Table S8**). These four genes code for the following proteins: i) the high mobility group protein B1 (PBANKA_0601900) of which the orthologue in *P. falciparum* has been shown to potently activate pro-inflammatory cytokines, suggesting a role in triggering host inflammatory immune responses (56); ii) a protein of the PIR multigene family (PBANKA_0500781); the function of PIRs is not known but these proteins are believed to be involved in antigenic variation and evasion from host immune responses (57); iii) a conserved *Plasmodium* protein of unknown function (PBANKA_0915200); iv) MSP9 (Merozoite surface protein 9, PBANKA_1443300). The orthologue of MSP9 in *P. falciparum* has been shown to bind to erythrocyte band 3 protein and to form a complex with MSP1 (58,59). *P. falciparum* merozoites are able to infect RBC via two different invasion mechanisms: one is sialic acid-dependent, involving MSP9 and the other one is MSP9 and sialic acid-independent. Interestingly MSP9 at a transcriptional level is barely expressed in DC and also hardly in other developmental EEF stages, which might be indicative that EEF stage-derived merozoites may not invade via the sialic acid-dependent mechanism. Attempts to knock out *P. berghei* MSP9 were not successful (60). In *P. yoelii*, MSP9 has been found in the erythrocyte cytoplasm (61) and might, in addition to invasion, also be involved in egress of merozoites from RBC.

Thereafter, we searched for genes that are specifically upregulated in DC when compared to all other stages and identified 293 genes (**Table S9**). It is conceivable that most of these are involved in the EEF stage-specific egress, in parasite survival in the dying host hepatocyte or in early RBC remodeling. As discussed earlier, it is plausible that transcripts are translationally suppressed in merozoites until the next stage in the life cycle, like has been described for mature gametocytes and sporozoites (33,62). Among the highly upregulated transcripts, *sbp1* (skeleton binding protein 1) (PBANKA_1101300), *mahrp1a* and *mahrp1b* (PBANKA_1145800 and PBANKA_1145900) were identified. These gene products are all involved in the trafficking of exported proteins from the parasite to the surface of the RBC (50). Knock out of MAHRP1a or SBP1 reduced the sequestration of infected cells (50). It is plausible that EEF stage-derived merozoites express high levels of these mRNAs in order to sufficiently express these proteins immediately after infection of RBC. Plausibly, this may allow the infected RBC to be efficiently sequestered in the periphery to avoid immediate clearance by the spleen. Why expression of *sbp1* and *mahrp1a* and *mahrp1b* appears to be lower in EF stage merozoites than in EEF stage merozoites is unknown. One reason might be, that in course of EF stage infection, the spleen is heavily remodeled (63–65) and allows for passage of infected RBC, making an efficient sequestration and thus a pronounced expression of sequestration ligands on the surface of the infected RBC less necessary (50). The parasite might thus be less dependent on intense trafficking of sequestration ligands. Another reason might be that MAHRP1a and SBP1 are already expressed during ring and trophozoite stage and thus less transcript would be needed to fulfill the same function as in EEF.

### Genes predominantly expressed in exo-erythrocytic stages

One of the goals of this study was to identify EEF stage-specific expressed genes. The EEF data were therefore compared with data from all other stages. The comparison revealed 5 highly specifically expressed transcripts for EEF stages with a LogFC >6 and adjP <0.01. (**Fig. 6**). The genes *lisp1* (PBANKA_1024600) *and lisp2* (PBANKA_1003000) have been previously reported to be expressed exclusively during EEF stage development (65–67). This is clearly reflected by our RNA-seq analysis where substantial *lisp1* and *lisp2* mRNA levels were only found in EEF stages, from 24hpi to DC. Along with *lisp1* and *lisp2* we identified two conserved *Plasmodium* genes (PBANKA_0518900 and PBANKA_0519500), one of which is annotated as membrane protein, although there is no transmembrane domain other than the signal peptide (PBANKA_0518900). The fifth gene in this EEF-specific group of genes is PBANKA_1003900. An averaged logFC of PBANKA_1003900 from later stages compared to non-EEF stages was similarly high as for lisp2 and lisp1 (**Table 4**). PBANKA_1003900 is a syntenic ortholog of *P. falciparum* sexual stage-specific protein precursor (Pfs16; PF3D7_0406200) which is expressed early during of development of *P. falciparum* gametocytes (68). *P. berghei* parasites expressing an mCherry-tagged PBANKA_1003900 provided experimental evidence that this gene is also expressed in gametocytes and was therefore annotated as a gametocyte specific protein (69). However, substantial PBANKA_1003900 transcript levels were only detected in EEF stages. We generated a transgenic parasite line expressing GFP under the control of the promoter of PBANKA_1003900 (PBANKA_1003900^GFP^) and analyzed GFP expression by fluorescence microscopy in the different EEF and EF stages. We could not detect GFP in any of the EF stages, including gametocytes. In contrast, GFP expression was detected by fluorescence microscopy of infected HeLa cells fixed at different time points (**Fig. 7**; 24h, 48h, 54h, 60h). At 24h of EEF stage development no or only very weak GFP-fluorescence was detectable. At 48h of EEF stage development, the signal was readily visible and at 54h and 60h post infection the fluorescent signal was profoundly intense. When performing live cell imaging, we observed the first signals at 30h post infection (**Movie 1**, starting from 30hpi). From 45h onwards the protein was substantially expressed confirming the results obtained from fixed cells. These fluorescence patterns (fixed and live) nicely confirmed our RNA-seq data during EEF stage development. Interestingly, analyses of gene-deletion mutants lacking PBANKA_1003900 demonstrated that the gene is not essential at any developmental stage throughout the complete parasite life cycle ((69) and own unpublished data). According to the expression profile deduced from the RNA seq analysis and also confirmed by the promoter analyses, we propose to revise the annotation of this gene “gametocyte specific protein” and to rename it as “liver specific protein 3 (LISP3)”.

**Figure 6:**
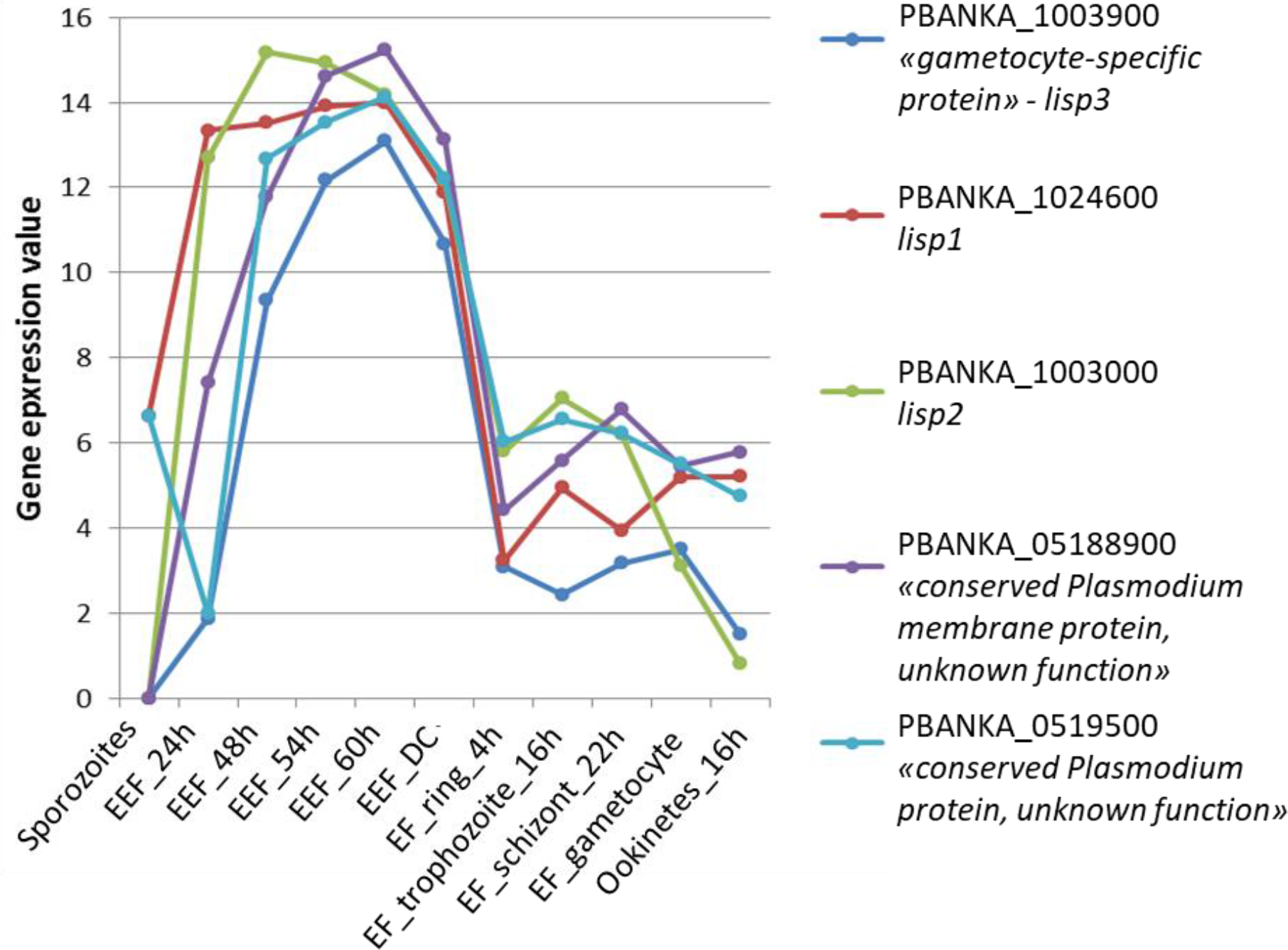
Expression profiles of the top 5 genes predominantly expressed in exo-erythrocytic stages compared to all other stages. Gene expression values corresponding to normalized and log2(x+1)-transformed read counts. The data were normalized with DESeq2 (with default parameters)(74).

**Figure 7:**
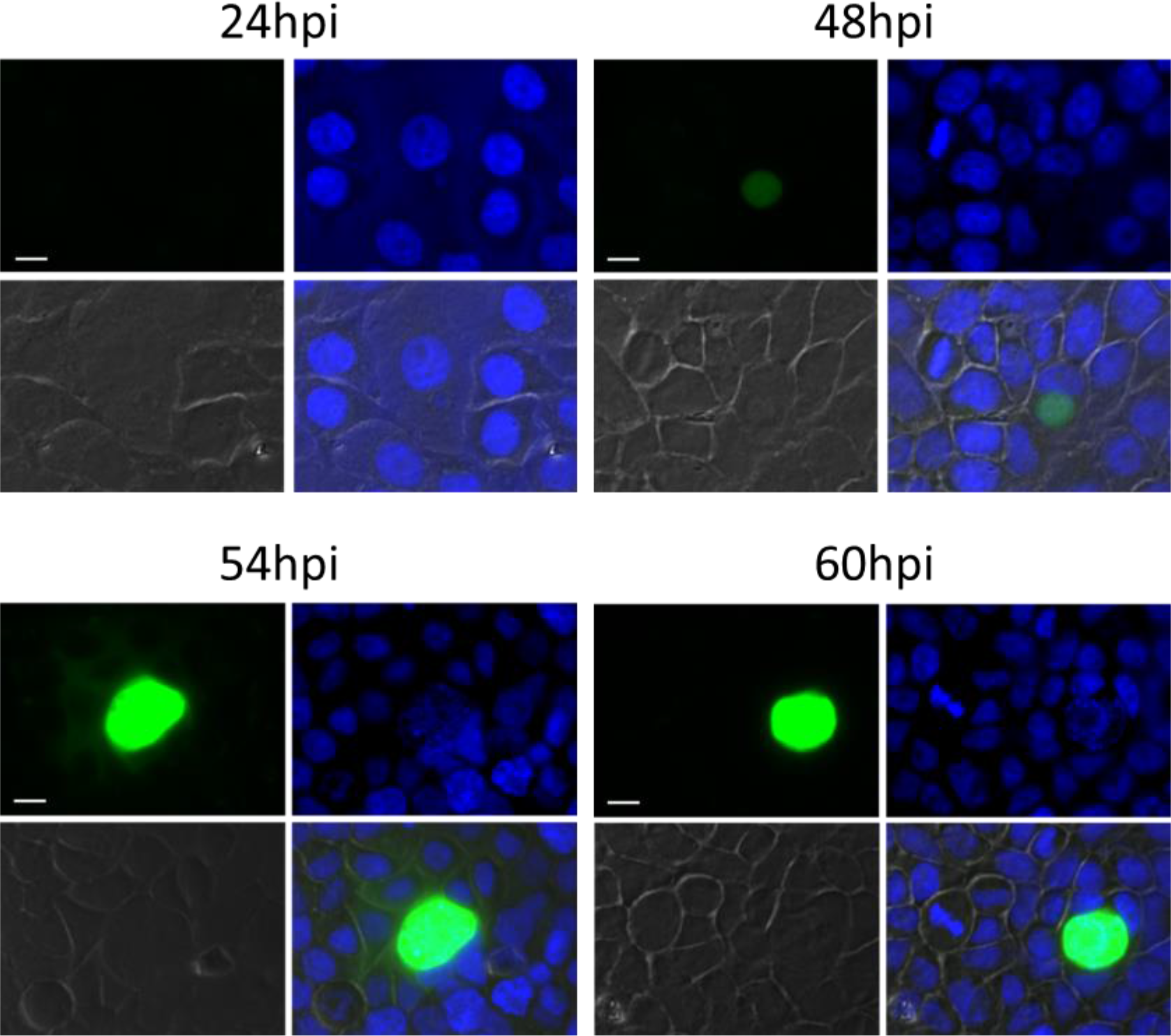
GFP expression during EEF stage development of the transgenic line 300 expressing GFP under control of the promoter region of PBANKA_1003900 (PBANKA_1003900^GFP^). Infected HeLa cells were fixed with 4%PFA/PBS at indicated times post infection with sporozoites. Nuclei were stained with Hoechst. Microscopy-settings (e.g. exposure time) were kept the same for all samples (bar 10µm).

**Table 4:** LogFC and FDR (adjP) values of EEF-specific genes (EEF_48h - DC compared to other stages). LogFC is indicated as mean of EEF late stages by single comparisons to each other stage. FDR (adjP) values are presented as Min and Max values of the different comparisons.

|  |  | 48h-DC compared to other stages |  |  |
| --- | --- | --- | --- | --- |
|  |  | mean LogFC | Min adjP | Max adjP |
| liver specific protein 2 (*lisp2*) | PBANKA_1003000 | 9.946 | 2.26E-228 | 1.22E-18 |
| "gametocyte-specific protein" (*lisp3*) | PBANKA_1003900 | 8.730 | 9.44E-133 | 2.17E-10 |
| conserved Plasmodium membrane protein, unknown function | PBANKA_0518900 | 8.697 | 3.36E-175 | 9.55E-17 |
| liver specific protein 1 (*lisp1*) | PBANKA_1024600 | 8.475 | 1.21E-189 | 1.24E-38 |
| conserved Plasmodium protein, unknown function | PBANKA_0519500 | 7.035 | 7.81E-64 | 2.28E-21 |

## Conclusion

We present here the first time-course transcriptome of the exo-erythrocytic stages of *P. berghei* by RNA-seq. This allowed an extended comparative gene transcription analysis of the exo-erythrocytic stages with published genome wide transcription analyses of erythrocytic stages, both asexual and sexual erythrocytic stages as well as ookinetes. This offers a comprehensive overview of gene transcription throughout most of the life cycle and allows a better understanding of gene regulation in different life cycle stages. In particular, the transcriptome of these different life cycle stages provide an invaluable tool for systems biology approaches to model metabolic pathways that are essential in different steps of the *Plasmodium* life cycle. In a first attempt to analyze transcription profiles during the *Plasmodium* life cycle, we have segmented almost 4700 genes into 14 distinct communities based on their different expression profile and attributed gene ontology terms to the individual communities. A more detailed analysis of the promoter regions of the genes in the communities with similar expression profiles could help identifying common DNA domains that might support our further understanding of regulation of gene expression in *Plasmodium*. We believe that the RNA-seq data provided here are of great interest for fundamental research questions with respect to parasite biology as well as for applied research aiming to identify new protein targets for vaccine and drug development.

## Acknowledgments

FACS sorting was performed at the Cytometry Laboratory Core facility of the University of Bern; we are thankful for the kind support by Stefan Müller. RNA-sequencing was performed at the iGE3 genomics platform of the University of Geneva

## Funding

DSF and VH were supported by the RTD grant MalarX (grant number: 51RTPO_151032), within SystemsX.ch (the Swiss Initiative for Systems Biology http://www.systemsx.ch/projects/research-technology-and-development-projects/malarx/)

## Experimental Procedures

### Mice, Parasites, Infections

Experiments were performed in accordance with the guidelines of the Swiss Act on animal protection (TSchG) and approved by the animal experimentation commission of the canton Bern (authorization numbers BE109/13 and BE132/16). The generation of the parasite line expressing GFP under the promoter of PBANKA_1003900 was approved by the Animal Experiments Committee of the Leiden University Medical Center (DEC 12042). The Dutch Experiments on Animal Act is established under European guidelines (EU directive no. 86/609/EEC regarding the Protection of Animals used for Experimental and Other Scientific Purposes).

BALB/c mice used for mosquito infections were between 6-10 weeks of age and were purchased from the Central animal facility at University of Bern, Harlan (Horst, the Netherlands) or Janvier Labs (Le Genest Saint Isle, France).

For RNA work: Mice were infected by intraperitoneal injection of blood stabilates of marker-free *P. berghei* ANKA expressing mCherry under the control of hsp70 regulatory sequences (PbmCherry_hsp70_) (7). At parasitemia of ~4%, the mouse was bled and 40 μl of infected blood was injected intravenously into phenylhydrazine treated mice (200ul of 6mg/ml in PBS, 2-3 days before). At day 3 to 4 after infection, each mouse with at least 7% parasitemia was anaesthetized with Ketasol/Xylasol and exposed to ~150 female *Anopheles stephensi* mosquitos (which were sugar starved for 5h). Mosquitoes were kept at 20.5°C and 80% humidity. From day 16-26 post infection salivary glands of infected mosquitos (sorted by fluorescence stereomicroscope *Olympus SZX10/ U-HGLGPS*) were dissected into serum free IMDM (*Iscove’s Modified Dulbecco’s Medium*, Sigma-aldrich). Sporozoites were liberated from the glands and were used to infect confluent confluent HeLa cells (per time point 10 wells of 96well plates were seeded wit 40’000 cells/well the day before). Each well was infected with ~ 20’000 PbmCherry_hsp70_ sporozoites for 6h. The cells were detached by the use of accutase (Innovative Cell Technology), pooled and the equivalent of 10 wells were seeded in 25cm2 cell culture flask. 1/6^th^ of the cells was washed once with PBS and pelleted by centrifugation (2min 100g). The pellet was loosened by flicking and the cells were resuspended with 250ul of RNAlater and stored at 4°C till all time points were harvested. It has been reported that RFP in contrast to GFP is preserved in RNAlater treated cells (28).

Media of the cultured cells were changed at 24hpi and 48hpi. At the respective time points the cells were detached from the surface by the use of accutase, washed once with PBS and then resuspended in 250ul RNAlater and stored at 4°C. With the use of RNAlater we could shorten the time in unnatural status not being in the incubator in adequate environment and medium) down to 10minutes compared to 1-2 hours in case of sorting fresh cells.

To generate transgenic parasites expressing *gfp* under control of the promoter of PBANKA_1003900 (PBANKA_1003900^GFP^), construct PbGFPcon vector was used (35). First, the PBANKA_1003900 promoter (1.7 kb) was amplified using primers GCTCTACCAATTTTGTGTCAC and GGATCCTTAAAAATTAATTTTGTATAAAATCG and cloned into pCR2.1-TOPO vector (Invitrogen) and sequenced. Then the *P. berghei elongation factor-1α* promoter of PbGFPcon was exchanged for the PBANKA_1003900 promoter (EcoRV from pCR2.1-TOPO vector /BamHI) and the *gfp* gene was re-introduced in the correct orientation as a BamHI fragment. Finally the construct was used to transfect the reference wild type *P. berghei* ANKA parasite line (cl15cy1 (ANKAwt))(19) to generate line 300 (PBANKA_1003900^GFP^). Transfection with episomal construct and positive selection of transfected parasites with pyrimethamine was performed as described previously(19).

For Microscopy work: confluent HeLa cells in 96well plate (40’000cells seeded the day before) were infected with ~20’000 PBANKA_1003900^GFP^ sporozoites. Cells were washed and detached 2hours post infection using accutase and seeded on glass covers in 24wells. At the indicated time points cells were fixed with 4%PFA in PBS for 10 min, washed with PBS and kept at 4°C. Nuclei were stained with 1microM Hoechst 33342 for 20 minutes, embedded with Dako-mounting medium. Fluorescent microscopy pictures were taken on a Leica DM5500B. Signals were photographed using same exposure settings.

For live cell imaging, infected cells were seeded onto glass bottom dishes (35-20-1.5-N, Cellvis, Mountain View). Live cell microscopy was performed with a Leica DMI6000B epifluorescence microscope equipped with a SOLA-SE-II light source starting at 30hpi.

### FACS sorting

Cells kept in RNAlater were FACS sorted on a BD FACSARIA III, FACSflow was used as sheath fluid. A 561nm laser was used in combination of 610/20nm filter detect the infected cells. To obtain maximal purity we sorted using the 4-way-purity mode with 100 microns nozzle. The sorted cells were collected into RNAlater (500ul in Eppedorf tube). 100’000 non-infected cells at each time point were sorted as negative controls.

### RNA isolation, Library preparation, Sequencing

Prior to isolation of RNA by ReliaPrep™ RNA Cell Miniprep System RNAlater was removed from the cells by adding an equal volume of ddH_2_O to the cells. The cells were then centrifuged (200g, 2 minutes). The RNA from the pelleted cells was extracted according to the manufacturer’s protocol and kept at −80°C. RNA extraction and Illumina m RNA-sequencing were performed in duplicates. Following RNA isolation, total RNA was quantified with a Qubit Fluorometer (Life Technologies). Quality of the extracted RNA was checked by the RNA integrity number (RIN), measured using an Agilent 2100 BioAnalyser (Agilent Technologies). The SMARTer™ Ultra Low RNA kit from Clontech was used for the reverse transcription and cDNA amplification according to manufacturer’s specifications, starting with 10 ng of total RNA as input. The Nextera XT kit (Illumina, San Diego, CA, USA) was used for cDNA libraries preparation using 200 pg of cDNA. Poly A selection was applied to get rid of ribosomal RNA. Library molarity and quality was assessed with the Qubit and Tapestation using a DNA High sensitivity chip (Agilent Technologies). The cDNA libraries were pooled and loaded at 12.5 pM, multiplexed on the lanes of HiSeq Rapid PE v2 Flow cells for generating paired reads of 100 bases on an Illumina HiSeq 2500 sequencer (Illumina, San Diego, CA, USA).

### Data processing

Short reads generated in this study were deposited at the European Nucleotide Archive (http://www.ebi.ac.uk/ena/) are accessible through the accession number PRJEB23770 (Secondary study accession number: ERP105548).” Publicly available data was obtained from SRA (SRP027529, ERS092084 and ERS092085). All reads were quality-checked with FastQC (bioinformatics.babraham.ac.uk/projects/fastqc). For the publicly available data, illumina adaptor sequences and low-quality reads were removed with TrimGalore (version 0.4.1 with the parameter --illumina, www.bioinformatics.babraham.ac.uk/projects/trim_galore). For the data generated in this study, Nextera transposase sequences and low quality reads were removed with Trimmomatic (version 0.33 with the parameters ILLUMINACLIP:adapters/NexteraPE-PE.fa:2:30:10 LEADING:3 TRAILING:3 SLIDINGWINDOW:5:30 MINLEN:50; (70). Low complexity reads were removed with fqtrim (version 0.9.4; (71). For paired-end reads, if only one end was removed, the remaining read end was treated as single-end read. To remove potential contamination with host RNA, all reads were aligned to the human genome (ensembl82) with Bowtie2 (version, 2.2.5; (72). Single-end reads and read pairs with none of the ends aligning to the human genome were kept and aligned to the *P. berghei* ANKA reference genome (PlasmoDB Release 33) with Subread (i.e. subjunc, version 1.4.6-p5; (73) allowing up to 10 alignments per read (options: − n 20 −m 5 −B 10 −H -all Junctions, always in single-end mode, i.e., ignoring the reverse read-end of paired-end reads). Count tables were generated with Rcount (30) with an allocation distance of 100 bp for calculating the weights of the reads with multiple alignments and a minimal number of 5 hits. Count tables are available in supplemental **Table S2.**

### Differential expression

Variation in gene expression was analyzed with a general linear model in R with the package DESeq2 (version 1.16.1; (74) according to a design with a single factor comprising all different experimental groups. Specific groups were compared with linear contrasts and *P*-values were adjusted for multiple testing (75), (i.e., false discovery rate). Genes with an adjusted *P*-value (FDR) below 0.01 and a minimal logFC of 2 were considered to be differentially expressed. Normalized gene expression data for plotting and clustering was likewise obtained with DESeq2 (version 1.16.1; (74).

### Gene co-expression network

To identify groups of genes with similar expression patterns across the life cycle of *P. berghei*, we constructed a gene co-expression network (GCN. We therefore calculated an adjacency matrix with pairwise Pearson correlation coefficients, applied Fisher’s z-transformation and tested each pairwise correlation coefficient for being significantly bigger than zero (as described in (37)). *P*-values were adjusted for multiple testing (75) and correlation coefficients with an adjusted *P*-value below 0.001 were identified as significant. The significant pairwise correlation coefficients were then used to construct the GCN. To resolve the community structure of the GCN, we used a modularity optimization algorithm (38) implemented by the function “cluster_louvain” in the R package “igraph” (version 1.0.1; (76) Communities with less than 11 genes were collapsed into a single “mixed” community (70 communities with a total of 197 genes). The network was visualized with Cytoscape (version 3.5.1, “prefuse force directed layout”; (77) The GeneIDs per community are listed in **Table S4.**

### Gene ontology enrichment

To functionally characterize the network communities or genes found to be differentially expressed, we tested for enrichment of gene ontology (GO) terms with topGO (version 3.4.1 in conjunction with the GO annotation available from PlasmoDB (22). Analysis was based on gene counts (genes in the set of interest compared to all annotated genes) using the “weight” algorithm with Fisher’s exact test (both implemented in topGO). A term was identified as significant if the *P*-value was below 0.05.

### Enrichment of selected gene groups

To test for enrichment of a specific group of genes (e.g., “merozoite invasion genes” from within a gene set of interest compared to all genes annotated with any of the tested groups, we used Fisher’s exact test (two-by-two contingency table). *P*-values were adjusted for multiple testing (75) and groups with an adjusted *P*-value (FDR) below 0.05 were identified as significant.

