## Supplementary Info for "Transcriptome analysis of *Plasmodium berghei* during exo-erythrocytic development"

### Supplementary Information (Text, Figures and Table Legends)

In the following section individual genes are discussed and RNA-seq expression profiles compared with already published mRNA or protein expression.

i) House-keeping genes (**Fig. S1**): GAPDH is often used as control in RT-PCR and Western blot analyses to normalize the expression of a gene (1). GAPDH is a glycolysis enzyme considered to be constitutively expressed in all metabolically active life cycle stages. The RNA-seq profile shows very low levels in sporozoites and reduced levels in detached cells (DC), containing infectious merozoites. Considering that both sporozoites and merozoites are prepared for invasion and are neither growing nor replicating it can be expected that several metabolic processes are less active and glycolysis might be such a process. It has recently been shown that GAPDH resides on the sporozoite surface and interacts with CD68 on Kupffer cells for cell traversal through them (2,3).

The cytoskeleton proteins actin and tubulin are also often used as controls for RT-PCR and Western blot analyses (4,5). We found that *actin I* and *alpha tubulin 1* are constitutively expressed in all life cycle stages tested.

ii) Genes encoding Serine-repeat antigens (**Fig. S2**)

Previously reported RT-PCR-based analyses of the putative proteases SERA1-5 suggested distinct expression profiles for the different members of this gene family (4). *sera5* was predominantly expressed in sporozoites, whereas all other *sera* mRNAs were either not expressed or expressed less during this stage but were found upregulated during the liver and blood stages (4,6,7). Our RNA-seq analysis showed a very similar expression profile as that described for this gene family during liver stage development with high expression of *sera1 to sera4* during EEF and EF_schizont and low expression during sporozoite, ookinete stage and EF_gametocyte.

iii) Genes encoding parasitophorous vacuole membrane (PVM) proteins (**Fig. S3**)

Exported protein 1 (EXP1), expressed in blood and liver stages (8–10) is widely used as a marker for the PVM in immunofluorescence assays. According to its localization in the PVM in intracellular stages, high level of expression is expected in intracellular liver and blood stages and a lower expression in the extracellular (motile) stages, such as ookinetes and sporozoites. Our RNA-seq shows indeed nearly absence in sporozoites and ookinetes and high expression in developing blood and liver stages.

The two ETRAM proteins, Upregulated in infective sporozoites 3 (UIS3) and 4 (UIS4) are PVM proteins of EEF stage (11,12). We found that in particular *uis4* is strongly upregulated in sporozoites but already at 24 hours post infection, the expression level in the liver drops more than 50fold. This decrease in expression of *uis4* is in line with the published data (13)

v) genes encoding sporozoite surface proteins (**Fig. S4**)

Sporozoite-specific protein expression has been reported for *csp* and *trap* (14,15). Our analysis clearly confirms the published stage-specific mRNA profile for both genes. Although there is some basal expression in other stages, dramatically higher levels of *csp* and *trap* mRNA levels were detected in the sporozoites.

v) Genes encoding fatty acid biosynthesis enzymes (**Fig. S5**)

It is well-established that fatty acid biosynthesis is essential for successful completion of liver stage development of rodent malaria parasites (Vaughan et al., 2009; Yu et al., 2008). In *P. yoelii*, the transcription profile of the four genes coding for enzymes related to fatty acid biosynthesis (*fabB/F, fabI, fabZ, fabG*) exhibit a very strong upregulation during liver stage development (Vaughan et al., 2009). We have analyzed the same genes in *P. berghei* and found upregulated expression for all four genes during liver stage development. As in *P. yoelii*, *P. berghei fabB/F* and *fabI* appear to be significantly transcribed already during the sporozoite stage.

vi) Genes encoding merozoite surface proteins (**Fig. S6**)

Merozoite surface protein 1 (MSP1) has been detected by immunofluorescence analysis from mid to late liver stages and similar levels of MSP1 expression was detected in blood stage and liver stage merozoites (18). The RNA-seq data confirm this expression pattern, in that MSP1 expression is increasing towards late liver stage parasites. In early blood stages, MSP1 expression is high but then drops sharply in trophozoites to increase again in blood stage schizonts when merozoites are formed. The other MSPs have in common to be upregulated towards late liver stage and blood stage schizonts and except MSP8 that drops in expression at blood trophozoite stage.

vii) Selected genes for promoter driven reporter proteins (**Fig. S7**)

Heat shock protein 70 (hsp70) is highly expressed in all life cycle stages and the hsp70 promoter has been used to constitutively express reporter proteins in *P. berghei* (19,20). In fact, it is considered superior to the eef1α promoter that was thought to be constitutively active and has already been used to generate *P. berghei* reporter cell lines (21–24). Our RNA-seq data confirm the expected profile of both hsp70 and eef1α. However, whereas eef1α was found hardly expressed in sporozoites, hsp70 shows a more evenly expression throughout the life cycle and thus is better suited for the generation of reporter parasite lines.

### Supplemental Figures

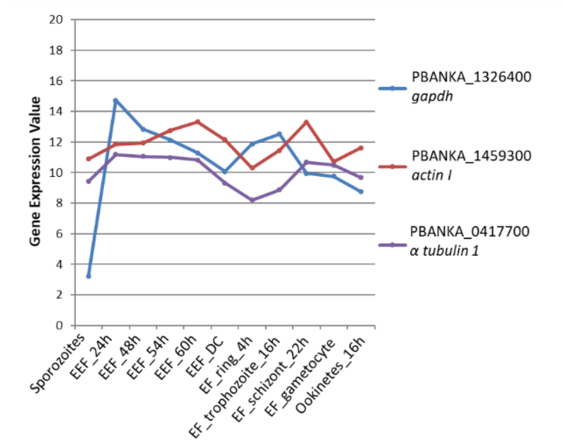


**Figure S1:** RNA expression profiles of 3 housekeeping genes (*gapdh, actinI, tubulin1*). Gene expression values corresponding to normalized and log2(x+1)-transformed read counts. The data were normalized with DESeq2 (with default parameters)(25).

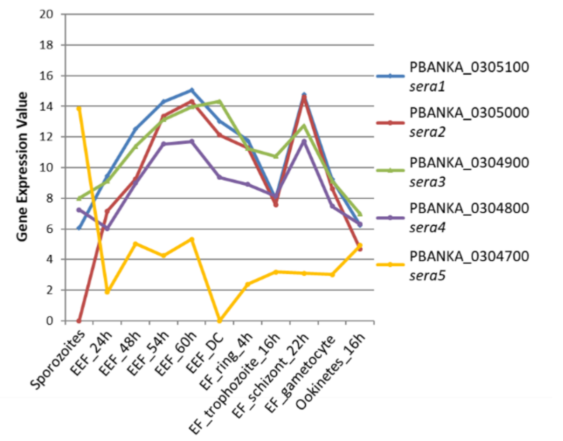


**Figure S2:** RNA expression profiles of 5 genes encoding serine-repeat antigens, serine-type proteases (SERA1-5): Gene expression values corresponding to normalized and log2(x+1)-transformed read counts. The data were normalized with DESeq2 (with default parameters)(25).

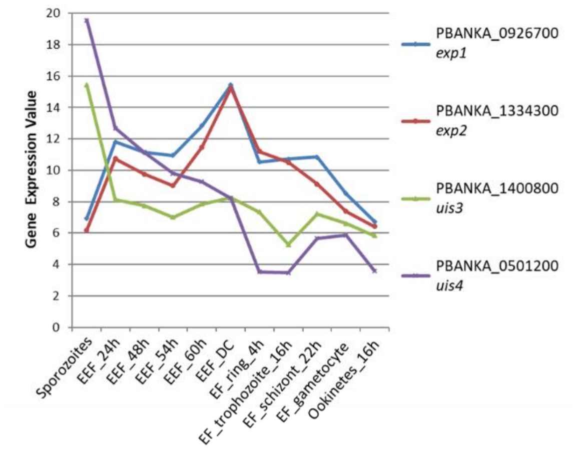


**Figure S3**: RNA expression profiles of 5 genes encoding proteins of the parasitophorous vacuole membrane (Exported protein 1, Exported protein 2, UIS3, UIS4). Gene expression values corresponding to normalized and log2(x+1)-transformed read counts. The data were normalized with DESeq2 (with default parameters)(25).

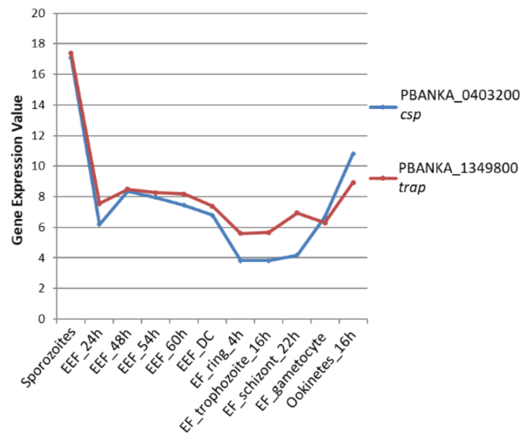


**Figure S4:** RNA expression profiles of genes encoding 2 sporozoite surface proteins (CSP and TRAP). Gene expression values corresponding to normalized and log2(x+1)-transformed read counts. The data were normalized with DESeq2 (with default parameters)(25).

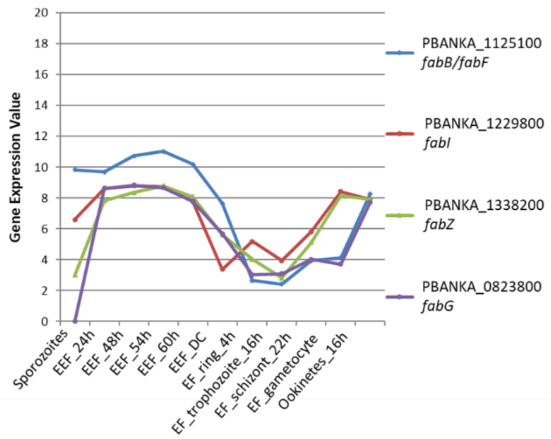


**Figure S5:** RNA expression profiles of 4 genes encoding enzymes involved in fatty acid biosynthesis (FabB/F, FabI, FabZ, FabG). Gene expression values corresponding to normalized and log2(x+1)-transformed read counts. The data were normalized with DESeq2 (with default parameters)(25).

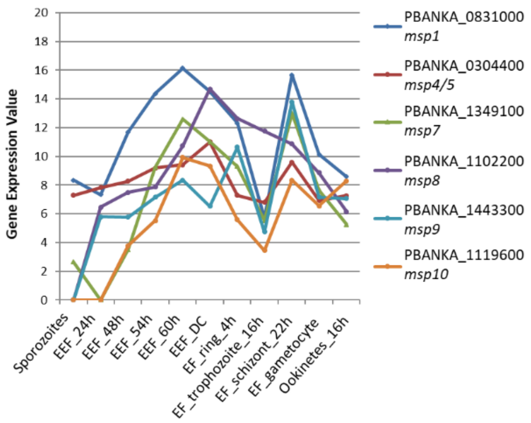


**Figure S6:** RNA expression profiles of 6 genes encoding merozoite surface proteins (MSP). Gene expression values corresponding to normalized and log2(x+1)-transformed read counts. The data were normalized with DESeq2 (with default parameters)(25).

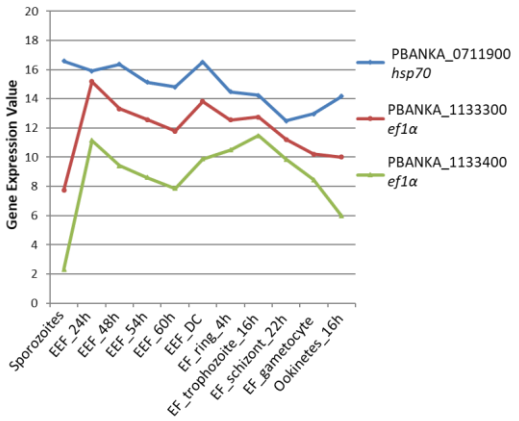


**Figure S7:** RNA expression profiles of 3 genes whose promoter regions have been used to drive expression of fluorescent/luminescent reporter proteins (HSP70, two genes for EF1α). Gene expression values corresponding to normalized and log2(x+1)-transformed read counts. The data were normalized with DESeq2 (with default parameters)(25).

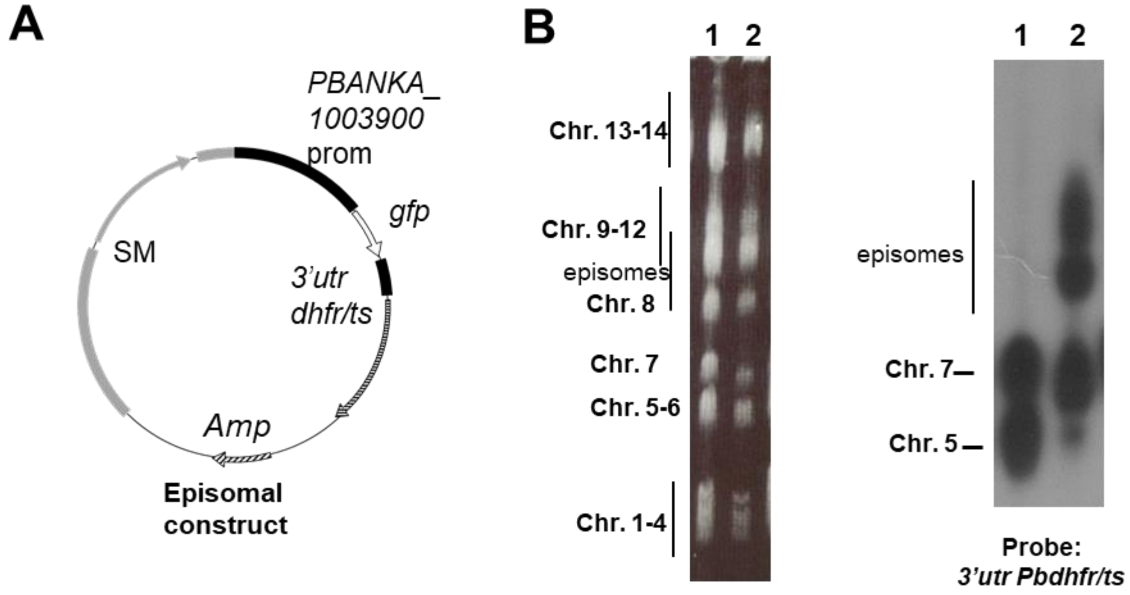


**Figure S8:** Generation and genotyping of parasites expressing *gfp* under control of the promoter of PBANKA_1003900 (PBANKA_1003900$^{GFP}$). **(A)** Schematic representation of the plasmid used to generate parasites expressing *gfp* under control of the PBANKA_1003900 promoter. The construct contains the *Toxoplasma gondii* dihydrofolate reductase - thymidylate synthase (TgDHFR-TS) selectable marker cassette (SM: grey boxes and arrow) and the *gfp* expression cassette under control of the 1,7 kb of the PBANKA_1003900 promoter region (black boxes and white arrow). (**B)** Southern analysis of pulsed field gel-separated chromosomes confirmed episomal transfection of construct in line 300 (lane 2). Right panel: chromosomes separated by pulsed field gel electrophoresis (FIGE) of two *P. berghei* mutant lines (line 300 and control line 299). Chromosomes are visualized by ethidium bromide staining of the gels. Left panel: separated chromosomes were hybridized with 3'utr Pb*dhfr/ts* recognizing the selectable marker and the GFP-expression cassettes of the introduced episomes and the 3'*utr* of the endogenous *Pbdhfr/ts* gene on chromosome 7. Lane 2 (exp 300) shows presence of episomal copies of the plasmid expressing GFP under control of the PBANKA_1003900 promoter and lane 1 show hybridization pattern of control line 299 with a construct integrated in chromosome 5.

**Movie:** GFP expression under the control of PBANKA_1003900 promoter (PBANKA_1003900$^{GFP}$) during EEF stage development. *P.berghei* PBANKA_1003900$^{GFP}$ sporozoites were monitored by live-cell time-lapse fluorescence microscopy starting at 30hpi of HeLa cells with 1h time interval between frames.

### Supplementary Table Legends

**Table S1:** RNA-seq data of different life cycle stages of *P. berghei* used in this study. Sporozoites, Exo-erythrocytic forms (EEF) at different time points (DC are detached cells), Erythrocytic forms (EF) (rings, trophozoites, schizonts, gametocytes) and ookinetes. A, B: Biological replicates. The sum of counts in Genes unique and in Genes Multireads were used for analysis.

**Table S2:** RNA-seq analysis: Raw sequencing counts per gene in the different life cycle stages as described in Table S1 (data generated by this study and archived Data SRP027529, ERS092084, and ERS092085)

**Table S3:** Gene ontology (GO) term annotation of genes of the individual communities in the GCN. The different communities are numbered 1-14 and indicated as different colours. Biological Process (BP), Cellular Component (CC), Molecular Function (MF). The number of annotated and observed genes per GO term are listed. Only GO terms with p-values < 0.05 are shown.

**Table S4:** *P. berghei* geneIDs of genes of the different communities in the GCN (1-14 and 'mixed') and geneIDs of genes that did not meet the GCN criteria.

**Table S5:** Genes preferentially expressed in developing EEF stages compared to developing EF stages. For each gene the mean expression (logBasemean), the log(fold change) (logFC), the p-value (pVal) and the adjusted p-value (adjP) is listed. Genes with an adjP < 0.01 and logFC > 2 were considered to be differentially expressed.

**Table S6:** Genes preferentially expressed in detached cells (DC) compared to erythrocytic schizonts. For each gene the mean expression (logBasemean), the log(fold change) (logFC), the p-value (pVal) and the adjusted p-value (adjP) is listed. Genes with an adjP < 0.01 and logFC > 2 were considered to be differentially expressed.

**Table S7:** Genes preferentially expressed in blood schizonts (22h) compared to EEF stages. Gene expression values (DESeq2 normalized) are listed for the individual genes in the different stages.

**Table S8:** Genes preferentially expressed in blood schizonts (22h) compared to all other stages. Gene expression values (DESeq2 normalized) are listed for the individual genes in the different stages.

**Table S9:** Genes preferentially expressed in detached cells (DC) compared to all other stages. Gene expression values (DESeq2 normalized) are listed for the individual genes in the different stages
