## Supplementary material for "Transcriptome analysis of *Plasmodium berghei* during exo-erythrocytic development": Table S1

| SampleID | Sample Name | Number of reads | | | | | Submitted by |
|---|---|---|---|---|---|---|---|
| | | at beginning | after trimming | after trimming and contamination removal | unique in Genes | multireads in Genes | |
| sporozoites_A | sporozoites_A | 37500288 | 17921395 | 26064163 | 646692 | 40468.51 | our study |
| sporozoites_B | sporozoites_B | 37931914 | 18251389 | 26329859 | 900938 | 50430.14 | our study |
| 6h_A | EEF_6h_A | 47689140 | 19115693 | 3111445 | 30236 | 5076.01 | our study |
| 6h_B | EEF_6h_B | 34165157 | 15481184 | 2166154 | 19796 | 2358.01 | our study |
| 24h_A | EEF_24h_A | 56359997 | 21243366 | 5452779 | 743431 | 80087.9 | our study |
| 24h_B | EEF_24h_B | 35009572 | 16240105 | 2847741 | 204577 | 25909.03 | our study |
| 48h_A | EEF_48h_A | 59525690 | 22716021 | 7150927 | 1594854 | 201381.56 | our study |
| 48h_B | EEF_48h_B | 37452654 | 17363067 | 3855098 | 797923 | 74255.13 | our study |
| 54h_A | EEF_54h_A | 57934160 | 22780746 | 8324943 | 2291368 | 258570.85 | our study |
| 54h_B | EEF_54h_B | 37874942 | 17463231 | 4045986 | 947547 | 70101.23 | our study |
| 60h_A | EEF_60h_A | 61838703 | 24241278 | 6873698 | 1526072 | 149856.75 | our study |
| 60h_B | EEF_60h_B | 34975120 | 16076252 | 3699030 | 835920 | 58091.45 | our study |
| DC_A | EEF_DC_A | 60324191 | 22452075 | 6631295 | 1762414 | 245726.87 | our study |
| DC_B | EEF_DC_B | 36918489 | 16800740 | 3975889 | 1199311 | 86103.86 | our study |
| SRR935544 | EF_ring_4h_A | 30755432 | 27171960 | 25775403 | 8580890 | 1688858.88 | RADBOUD UNIVERSITY NIJMEGEN |
| SRR935549 | EF_ring_4h_B | 20644266 | 14828351 | 14361601 | 5113160 | 206140.56 | RADBOUD UNIVERSITY NIJMEGEN |
| SRR935553 | EF_trophozoite_16h_A | 20126017 | 15544235 | 15505960 | 3174804 | 203762.83 | RADBOUD UNIVERSITY NIJMEGEN |
| SRR935554 | EF_trophozoite_16h_B | 26378106 | 23769936 | 23604036 | 13771565 | 541988.9 | RADBOUD UNIVERSITY NIJMEGEN |
| SRR935555 | EF_schizont_22h_B | 22664938 | 11273435 | 11271955 | 2463391 | 61668.18 | RADBOUD UNIVERSITY NIJMEGEN |
| SRR935557 | EF_schizont_22h_A | 24127263 | 22535424 | 22459281 | 14387523 | 510439.12 | RADBOUD UNIVERSITY NIJMEGEN |
| SRR935558 | EF_gametocyte_A | 31951285 | 30942518 | 30861381 | 14577361 | 849699.98 | RADBOUD UNIVERSITY NIJMEGEN |
| SRR935559 | EF_gametocyte_B | 26051565 | 23164642 | 23127054 | 14836047 | 322399.14 | RADBOUD UNIVERSITY NIJMEGEN |
| ERR435801 | ookinete_16h_A | 3889541 | 3842631 | 3858639 | 2777705 | 12730.94 | The Wellcome Trust Sanger Institute |
| ERR435802 | ookinete_16h_B | 29246226 | 28856086 | 29023126 | 21301968 | 132116.31 | The Wellcome Trust Sanger Institute |
