## Supplementary material for "Transcriptome analysis of *Plasmodium berghei* during exo-erythrocytic development": Table S2

| Table S2 | sporozoites_A | sporozoites_B | EEF_24h_A | EEF_24h_B | EEF_48h_A | EEF_48h_B | EEF_54h_A | EEF_54h_B | EEF_60h_A | EEF_60h_B | EEF_DC_A | EEF_DC_B | EF_ring_4h_A | EF_ring_4h_B | EF_trophozoite_16h_A | EF_trophozoite_16h_B | EF_schizont_22h_A | EF_schizont_22h_B | EF_gametocyte_A | EF_gametocyte_B | ookinete_16h_A | ookinete_16h_B |
|---|---|---|---|---|---|---|---|---|---|---|---|---|---|---|---|---|---|---|---|---|---|---|
| PBANKA_0100061 | 18 | 16 | 5 | 0 | 43 | 23 | 39 | 16 | 123 | 80 | 531 | 351 | 223 | 169 | 38 | 156 | 140 | 63 | 350 | 184 | 9 | 53 |
| PBANKA_0100200 | 8 | 0 | 20 | 6 | 195 | 142 | 189 | 79 | 125 | 81 | 325 | 205 | 2462 | 938 | 1079 | 4602 | 970 | 174 | 2009 | 1129 | 51 | 492 |
| PBANKA_0100700 | 0 | 0 | 11 | 0 | 85 | 89 | 201 | 156 | 1796 | 1579 | 14310 | 14001 | 9093 | 6054 | 2330 | 6242 | 4843 | 1616 | 1498 | 695 | 21 | 331 |
| PBANKA_0100800 | 77 | 88 | 72 | 22 | 177 | 85 | 242 | 110 | 131 | 97 | 355 | 213 | 1181 | 1473 | 331 | 2307 | 1914 | 433 | 7030 | 14666 | 305 | 3069 |
| PBANKA_0100900 | 134 | 163 | 358 | 110 | 865 | 393 | 770 | 321 | 559 | 304 | 5480 | 3801 | 2488 | 3380 | 723 | 4317 | 1506 | 223 | 1883 | 2181 | 214 | 1639 |
| PBANKA_0101000 | 21 | 21 | 128 | 41 | 202 | 128 | 197 | 96 | 63 | 31 | 44 | 23 | 843 | 474 | 576 | 1722 | 563 | 95 | 61 | 55 | 0 | 32 |
| PBANKA_0101100 | 16 | 31 | 78 | 5 | 309 | 150 | 390 | 109 | 135 | 56 | 11 | 5 | 94 | 37 | 63 | 106 | 197 | 28 | 292 | 142 | 13 | 117 |
| PBANKA_0101200 | 60 | 98 | 168 | 52 | 374 | 167 | 348 | 137 | 157 | 67 | 11 | 6 | 1238 | 387 | 535 | 2497 | 1775 | 210 | 281 | 239 | 255 | 1584 |
| PBANKA_0101300 | 17 | 18 | 55 | 17 | 142 | 97 | 170 | 60 | 70 | 30 | 0 | 11 | 624 | 144 | 298 | 1000 | 362 | 55 | 82 | 56 | 26 | 304 |
| PBANKA_0101400 | 122 | 210 | 185 | 40 | 559 | 335 | 1150 | 458 | 561 | 313 | 72 | 44 | 1673 | 1114 | 954 | 4638 | 5189 | 854 | 1610 | 2966 | 420 | 3093 |
| PBANKA_0101500 | 0 | 7 | 19 | 5 | 68 | 31 | 100 | 51 | 75 | 42 | 25 | 15 | 933 | 536 | 130 | 558 | 739 | 142 | 17416 | 14037 | 93 | 820 |
| PBANKA_0101600 | 23 | 21 | 30 | 5 | 194 | 76 | 858 | 302 | 364 | 141 | 50 | 18 | 522 | 676 | 159 | 738 | 654 | 116 | 216 | 155 | 30 | 123 |
| PBANKA_0101700 | 37 | 23 | 97 | 11 | 206 | 84 | 289 | 92 | 121 | 43 | 25 | 7 | 918 | 402 | 259 | 1384 | 1136 | 166 | 1838 | 1096 | 123 | 808 |
| PBANKA_0101800 | 165 | 298 | 98 | 18 | 261 | 132 | 284 | 92 | 169 | 60 | 191 | 71 | 3665 | 1755 | 545 | 3512 | 1535 | 331 | 15782 | 11557 | 683 | 5245 |
| PBANKA_0101900 | 21 | 48 | 114 | 32 | 459 | 231 | 700 | 266 | 305 | 148 | 35 | 13 | 567 | 183 | 304 | 1459 | 2763 | 362 | 908 | 712 | 49 | 457 |
| PBANKA_0102000 | 34 | 61 | 90 | 27 | 278 | 184 | 395 | 104 | 128 | 88 | 22 | 6 | 268 | 55 | 344 | 899 | 915 | 175 | 185 | 108 | 83 | 614 |
| PBANKA_0102100 | 17 | 40 | 106 | 34 | 215 | 132 | 253 | 118 | 124 | 43 | 75 | 65 | 1615 | 815 | 214 | 1084 | 819 | 153 | 3190 | 2982 | 159 | 1223 |
| PBANKA_0102200 | 9 | 25 | 225 | 48 | 1058 | 299 | 1200 | 344 | 430 | 159 | 933 | 552 | 2466 | 3591 | 892 | 5713 | 1490 | 163 | 1445 | 2031 | 157 | 847 |
| PBANKA_0102300 | 34 | 32 | 30 | 16 | 257 | 187 | 355 | 197 | 93 | 40 | 15 | 21 | 204 | 71 | 442 | 1026 | 640 | 140 | 1162 | 506 | 102 | 777 |
| PBANKA_0102400 | 0 | 0 | 5 | 0 | 72 | 49 | 152 | 53 | 89 | 28 | 19 | 0 | 120 | 118 | 265 | 864 | 525 | 74 | 800 | 2722 | 51 | 350 |
| PBANKA_0102500 | 0 | 0 | 55 | 10 | 180 | 85 | 179 | 53 | 84 | 33 | 25 | 12 | 249 | 50 | 224 | 633 | 358 | 90 | 764 | 395 | 43 | 252 |
| PBANKA_0102600 | 13 | 14 | 53 | 5 | 690 | 338 | 1504 | 592 | 813 | 340 | 68 | 49 | 1422 | 676 | 156 | 555 | 9892 | 1174 | 3185 | 3250 | 2070 | 11638 |
| PBANKA_0102800 | 0 | 0 | 74 | 10 | 153 | 55 | 326 | 126 | 217 | 82 | 26 | 8 | 1063 | 917 | 660 | 4199 | 2225 | 281 | 1780 | 1473 | 199 | 1358 |
| PBANKA_0103000 | 0 | 0 | 25 | 6 | 70 | 27 | 74 | 20 | 48 | 8 | 0 | 0 | 54 | 8 | 31 | 71 | 100 | 53 | 245 | 83 | 37 | 168 |
| PBANKA_0103100 | 0 | 0 | 51 | 7 | 146 | 73 | 216 | 72 | 72 | 36 | 20 | 7 | 592 | 212 | 314 | 1278 | 778 | 122 | 618 | 376 | 169 | 1045 |
| PBANKA_0103300 | 66 | 90 | 583 | 105 | 1247 | 425 | 1080 | 362 | 636 | 239 | 1452 | 836 | 7777 | 15906 | 5272 | 30483 | 8071 | 1172 | 4246 | 9439 | 389 | 2413 |
| PBANKA_0103400 | 0 | 0 | 26 | 14 | 131 | 62 | 98 | 23 | 33 | 13 | 0 | 0 | 312 | 73 | 225 | 861 | 414 | 63 | 902 | 562 | 268 | 2021 |
| PBANKA_0103500 | 5 | 25 | 58 | 6 | 140 | 43 | 152 | 71 | 108 | 52 | 7 | 0 | 611 | 244 | 197 | 821 | 1039 | 195 | 619 | 465 | 299 | 1748 |
| PBANKA_0103700 | 0 | 0 | 12 | 0 | 34 | 18 | 25 | 16 | 15 | 6 | 0 | 0 | 140 | 16 | 95 | 383 | 113 | 12 | 293 | 135 | 11 | 73 |
| PBANKA_0103800 | 17 | 22 | 78 | 22 | 160 | 74 | 134 | 63 | 37 | 21 | 9 | 0 | 430 | 40 | 391 | 1300 | 347 | 46 | 249 | 116 | 216 | 1318 |
| PBANKA_0103900 | 375 | 397 | 142 | 40 | 621 | 270 | 849 | 335 | 499 | 199 | 234 | 145 | 4750 | 2090 | 927 | 5129 | 3040 | 555 | 4328 | 2850 | 970 | 6869 |
| PBANKA_0104000 | 46 | 51 | 92 | 15 | 139 | 75 | 159 | 107 | 49 | 56 | 8 | 0 | 279 | 67 | 147 | 664 | 559 | 91 | 1596 | 761 | 197 | 1702 |
| PBANKA_0104100 | 16 | 26 | 86 | 21 | 266 | 170 | 357 | 175 | 219 | 117 | 23 | 28 | 241 | 93 | 105 | 210 | 596 | 160 | 2681 | 1034 | 109 | 1020 |
| PBANKA_0104600 | 0 | 7 | 161 | 40 | 414 | 166 | 350 | 154 | 527 | 285 | 207 | 89 | 832 | 783 | 2594 | 12250 | 6708 | 1142 | 2750 | 8504 | 1882 | 8234 |
| PBANKA_0104700 | 0 | 0 | 152 | 50 | 301 | 140 | 271 | 136 | 80 | 55 | 14 | 6 | 352 | 98 | 491 | 636 | 844 | 407 | 2203 | 718 | 143 | 1183 |
| PBANKA_0104800 | 10 | 19 | 18 | 8 | 76 | 37 | 154 | 63 | 247 | 122 | 1568 | 893 | 2422 | 2145 | 404 | 2293 | 2792 | 576 | 1122 | 1247 | 85 | 651 |
| PBANKA_0104900 | 12 | 19 | 174 | 42 | 448 | 187 | 495 | 227 | 138 | 86 | 68 | 49 | 7337 | 3428 | 1820 | 10787 | 3119 | 375 | 2384 | 3647 | 341 | 2787 |
| PBANKA_0105000 | 100 | 154 | 46 | 9 | 119 | 70 | 135 | 92 | 71 | 66 | 0 | 8 | 300 | 76 | 104 | 232 | 282 | 143 | 501 | 202 | 156 | 677 |
| PBANKA_0105200 | 0 | 15 | 60 | 10 | 302 | 133 | 826 | 368 | 414 | 268 | 259 | 140 | 2049 | 1368 | 601 | 2416 | 2001 | 295 | 343 | 306 | 361 | 2232 |
| PBANKA_0105300 | 108 | 175 | 238 | 51 | 648 | 268 | 1004 | 369 | 541 | 260 | 550 | 414 | 2318 | 3459 | 1462 | 8004 | 3584 | 482 | 1212 | 3434 | 5423 | 37755 |
| PBANKA_0105400 | 21 | 37 | 254 | 64 | 922 | 509 | 1416 | 594 | 656 | 316 | 91 | 86 | 3631 | 1259 | 2314 | 10878 | 9706 | 1185 | 2169 | 2070 | 416 | 2891 |
| PBANKA_0105500 | 7 | 5 | 297 | 62 | 354 | 169 | 346 | 107 | 77 | 46 | 38 | 41 | 1160 | 499 | 513 | 2021 | 616 | 86 | 486 | 249 | 339 | 2040 |
| PBANKA_0105600 | 0 | 0 | 317 | 75 | 505 | 161 | 406 | 139 | 170 | 69 | 206 | 99 | 7907 | 14282 | 3512 | 25268 | 5280 | 546 | 868 | 5035 | 143 | 1224 |
| PBANKA_0105700 | 0 | 0 | 30 | 11 | 151 | 76 | 138 | 73 | 47 | 32 | 9 | 0 | 62 | 21 | 22 | 105 | 61 | 18 | 749 | 421 | 203 | 892 |
| PBANKA_0105800 | 0 | 0 | 165 | 26 | 159 | 53 | 148 | 34 | 54 | 10 | 12 | 0 | 336 | 71 | 219 | 684 | 1048 | 166 | 1741 | 1068 | 201 | 1315 |
| PBANKA_0105900 | 0 | 0 | 191 | 45 | 229 | 107 | 201 | 73 | 59 | 29 | 7 | 0 | 259 | 66 | 160 | 461 | 890 | 177 | 1305 | 701 | 68 | 422 |
| PBANKA_0106000 | 199 | 247 | 77 | 8 | 363 | 99 | 533 | 151 | 361 | 124 | 54 | 19 | 672 | 154 | 360 | 1442 | 1981 | 434 | 1449 | 746 | 212 | 982 |
| PBANKA_0106100 | 30 | 27 | 29 | 0 | 111 | 45 | 135 | 39 | 42 | 16 | 20 | 0 | 287 | 102 | 220 | 916 | 527 | 120 | 977 | 529 | 131 | 550 |
| PBANKA_0106200 | 0 | 0 | 13 | 0 | 165 | 84 | 707 | 312 | 318 | 176 | 28 | 15 | 437 | 163 | 63 | 269 | 854 | 154 | 3288 | 2148 | 166 | 1065 |
| PBANKA_0106300 | 59 | 88 | 35 | 8 | 385 | 182 | 1707 | 725 | 1335 | 625 | 170 | 77 | 2781 | 3470 | 172 | 475 | 25272 | 4462 | 30522 | 95751 | 490 | 3693 |
| PBANKA_0106400 | 10 | 11 | 64 | 18 | 578 | 216 | 577 | 193 | 222 | 83 | 51 | 14 | 746 | 233 | 260 | 1435 | 794 | 201 | 230 | 254 | 1407 | 8316 |
| PBANKA_0106500 | 0 | 0 | 85 | 16 | 213 | 49 | 193 | 63 | 99 | 33 | 30 | 9 | 1404 | 712 | 1045 | 5624 | 1108 | 110 | 347 | 548 | 1887 | 13250 |
| PBANKA_0106600 | 0 | 0 | 119 | 28 | 156 | 85 | 180 | 91 | 40 | 25 | 5 | 0 | 346 | 62 | 368 | 664 | 377 | 59 | 180 | 90 | 144 | 962 |
| PBANKA_0106700 | 0 | 0 | 167 | 40 | 318 | 142 | 337 | 92 | 156 | 78 | 46 | 17 | 950 | 764 | 1432 | 8244 | 3420 | 387 | 870 | 1361 | 950 | 6186 |
| PBANKA_0106800 | 0 | 9 | 14 | 0 | 23 | 0 | 22 | 9 | 13 | 6 | 0 | 0 | 131 | 21 | 100 | 253 | 222 | 57 | 397 | 238 | 60 | 394 |
| PBANKA_0106900 | 12 | 11 | 67 | 13 | 311 | 92 | 356 | 120 | 168 | 70 | 36 | 17 | 1357 | 493 | 812 | 3356 | 2263 | 460 | 2116 | 1638 | 91 | 693 |
| PBANKA_0106950 | 0 | 0 | 63 | 12 | 139 | 65 | 220 | 73 | 94 | 33 | 23 | 18 | 567 | 177 | 224 | 820 | 520 | 125 | 536 | 174 | 299 | 1636 |
| PBANKA_0107000 | 0 | 0 | 49 | 10 | 93 | 56 | 120 | 62 | 41 | 30 | 6 | 0 | 235 | 63 | 162 | 383 | 673 | 126 | 1054 | 358 | 573 | 4251 |
| PBANKA_0107100 | 0 | 0 | 258 | 79 | 412 | 258 | 542 | 264 | 133 | 93 | 20 | 7 | 1113 | 140 | 1370 | 3972 | 1749 | 265 | 889 | 382 | 66 | 566 |
| PBANKA_0107200 | 0 | 0 | 6 | 0 | 14 | 6 | 49 | 14 | 35 | 15 | 0 | 0 | 335 | 82 | 27 | 213 | 348 | 47 | 3071 | 1561 | 576 | 3309 |
| PBANKA_0107300 | 30 | 43 | 788 | 213 | 830 | 422 | 758 | 321 | 212 | 132 | 35 | 13 | 3058 | 629 | 3015 | 12020 | 2509 | 308 | 1197 | 837 | 227 | 1845 |
| PBANKA_0107400 | 149 | 307 | 75 | 32 | 203 | 123 | 410 | 190 | 185 | 107 | 53 | 24 | 518 | 129 | 765 | 3172 | 2052 | 281 | 1104 | 890 | 186 | 1743 |
| PBANKA_0107500 | 8 | 23 | 54 | 26 | 254 | 92 | 295 | 117 | 123 | 58 | 24 | 5 | 496 | 228 | 333 | 1166 | 1048 | 208 | 889 | 622 | 120 | 1013 |
| PBANKA_0107600 | 36 | 53 | 922 | 227 | 1982 | 624 | 3002 | 993 | 1712 | 619 | 863 | 440 | 2418 | 4546 | 2215 | 10526 | 17951 | 1880 | 1170 | 3381 | 307 | 2155 |
| PBANKA_0107700 | 58 | 55 | 101 | 35 | 112 | 84 | 78 | 45 | 104 | 92 | 1102 | 1165 | 2325 | 1012 | 494 | 2142 | 315 | 86 | 2023 | 1027 | 674 | 5098 |
| PBANKA_0107800 | 30 | 23 | 45 | 12 | 154 | 70 | 149 | 44 | 50 | 22 | 30 | 39 | 260 | 97 | 58 | 176 | 123 | 34 | 972 | 399 | 340 | 2257 |
| PBANKA_0107900 | 251 | 411 | 40 | 13 | 186 | 87 | 350 | 135 | 371 | 158 | 106 | 40 | 1477 | 667 | 261 | 1515 | 4850 | 881 | 2019 | 2156 | 1723 | 9202 |
| PBANKA_0108000 | 221 | 297 | 74 | 13 | 220 | 69 | 718 | 216 | 728 | 333 | 178 | 77 | 620 | 279 | 243 | 1122 | 1708 | 342 | 871 | 467 | 83 | 632 |
| PBANKA_0108200 | 5 | 0 | 50 | 11 | 229 | 80 | 680 | 247 | 646 | 299 | 653 | 417 | 2914 | 2979 | 693 | 4350 | 6384 | 1068 | 1430 | 2292 | 113 | 769 |
| PBANKA_0108400 | 0 | 7 | 104 | 30 | 465 | 110 | 745 | 242 | 613 | 173 | 127 | 31 | 1628 | 512 | 1034 | 5581 | 7468 | 1021 | 2598 | 3560 | 328 | 1891 |
| PBANKA_0108500 | 0 | 7 | 89 | 11 | 133 | 54 | 140 | 47 | 54 | 30 | 11 | 0 | 1110 | 496 | 304 | 1953 | 529 | 79 | 371 | 282 | 168 | 1198 |
| PBANKA_0108600 | 49 | 96 | 99 | 23 | 217 | 115 | 285 | 141 | 164 | 82 | 94 | 34 | 2773 | 1522 | 1089 | 5064 | 2296 | 364 | 697 | 1027 | 293 | 1484 |
| PBANKA_0108700 | 12 | 17 | 303 | 71 | 663 | 209 | 555 | 189 | 201 | 105 | 904 | 547 | 3757 | 5168 | 2179 | 15723 | 5091 | 489 | 3780 | 12796 | 1653 | 9860 |
| PBANKA_0108800 | 117 | 226 | 137 | 38 | 307 | 164 | 1032 | 719 | 1046 | 765 | 155 | 125 | 931 | 224 | 170 | 222 | 6863 | 2334 | 5480 | 1727 | 145 | 1220 |
| PBANKA_0108900 | 0 | 0 | 113 | 43 | 67 | 49 | 134 | 107 | 82 | 57 | 8 | 12 | 487 | 123 | 369 | 1167 | 179 | 49 | 139 | 68 | 0 | 38 |
| PBANKA_0109000 | 0 | 0 | 29 | 9 | 95 | 50 | 261 | 108 | 122 | 64 | 8 | 8 | 167 | 52 | 169 | 878 | 308 | 19 | 49 | 30 | 20 | 109 |
| PBANKA_0109100 | 40 | 48 | 511 | 182 | 1482 | 769 | 3710 | 1588 | 1689 | 1071 | 150 | 126 | 3401 | 850 | 3383 | 14508 | 4794 | 568 | 1875 | 1384 | 1518 | 13012 |
| PBANKA_0109200 | 10 | 14 | 213 | 76 | 671 | 255 | 825 | 280 | 414 | 180 | 1407 | 905 | 1611 | 3451 | 1331 | 8122 | 5310 | 660 | 617 | 3188 | 1032 | 7020 |
| PBANKA_0109300 | 0 | 0 | 39 | 5 | 199 | 98 | 338 | 123 | 135 | 56 | 12 | 8 | 528 | 143 | 108 | 474 | 1511 | 442 | 792 | 518 | 85 | 687 |
| PBANKA_0109400 | 0 | 0 | 24 | 0 | 60 | 37 | 95 | 68 | 25 | 28 | 0 | 0 | 67 | 11 | 114 | 252 | 242 | 59 | 153 | 51 | 11 | 82 |
| PBANKA_0109500 | 14 | 10 | 269 | 83 | 515 | 290 | 452 | 228 | 115 | 101 | 14 | 5 | 542 | 86 | 478 | 1654 | 789 | 167 | 752 | 351 | 83 | 548 |
| PBANKA_0109600 | 10 | 18 | 175 | 52 | 314 | 200 | 447 | 229 | 159 | 94 | 27 | 30 | 655 | 116 | 520 | 1857 | 1948 | 317 | 1328 | 804 | 436 | 3440 |

| | | | | | | | | | | | | | | | | | | | | | | |
|---|---|---|---|---|---|---|---|---|---|---|---|---|---|---|---|---|---|---|---|---|---|---|
| PBANKA_0109700 | 74 | 90 | 144 | 28 | 330 | 190 | 538 | 214 | 221 | 109 | 398 | 378 | 1888 | 1229 | 698 | 3116 | 2497 | 372 | 2807 | 2641 | 96 | 594 |
| PBANKA_0110000 | 57 | 50 | 163 | 37 | 111 | 68 | 134 | 35 | 46 | 28 | 302 | 199 | 2177 | 709 | 904 | 1066 | 460 | 198 | 486 | 136 | 26 | 139 |
| PBANKA_0110100 | 575 | 1041 | 196 | 40 | 1288 | 528 | 2819 | 1062 | 1547 | 734 | 235 | 114 | 1451 | 2179 | 685 | 3516 | 6854 | 833 | 2798 | 6097 | 596 | 4661 |
| PBANKA_0110200 | 0 | 0 | 50 | 12 | 118 | 47 | 205 | 84 | 76 | 43 | 6 | 0 | 549 | 178 | 176 | 704 | 370 | 130 | 865 | 473 | 69 | 695 |
| PBANKA_0110400 | 12 | 11 | 232 | 52 | 255 | 119 | 402 | 150 | 173 | 83 | 182 | 111 | 3161 | 1532 | 915 | 3413 | 923 | 165 | 1263 | 722 | 302 | 2291 |
| PBANKA_0110500 | 100 | 119 | 108 | 35 | 314 | 180 | 698 | 286 | 1136 | 619 | 188 | 152 | 1321 | 2670 | 846 | 4956 | 33258 | 3913 | 4796 | 38464 | 213 | 1248 |
| PBANKA_0110600 | 0 | 0 | 11 | 0 | 19 | 5 | 18 | 17 | 31 | 10 | 0 | 0 | 83 | 12 | 24 | 86 | 358 | 100 | 274 | 160 | 0 | 43 |
| PBANKA_0110700 | 0 | 0 | 20 | 0 | 63 | 14 | 50 | 20 | 41 | 19 | 0 | 0 | 138 | 36 | 38 | 172 | 553 | 148 | 2522 | 949 | 61 | 314 |
| PBANKA_0110900 | 0 | 0 | 93 | 20 | 173 | 80 | 117 | 49 | 55 | 40 | 58 | 45 | 498 | 369 | 210 | 1061 | 230 | 31 | 2682 | 1681 | 625 | 4438 |
| PBANKA_0111000 | 8 | 9 | 9 | 0 | 105 | 42 | 633 | 323 | 1455 | 1029 | 166 | 112 | 571 | 240 | 82 | 222 | 4394 | 1510 | 245 | 170 | 382 | 2440 |
| PBANKA_0111100 | 116 | 141 | 56 | 14 | 141 | 84 | 151 | 73 | 208 | 107 | 177 | 104 | 442 | 269 | 139 | 738 | 595 | 160 | 308 | 206 | 909 | 7051 |
| PBANKA_0111200 | 119 | 193 | 399 | 113 | 449 | 212 | 476 | 177 | 570 | 266 | 3046 | 2212 | 4544 | 3700 | 813 | 4865 | 1355 | 272 | 2012 | 1906 | 1017 | 7588 |
| PBANKA_0111300 | 12 | 9 | 208 | 65 | 223 | 145 | 207 | 110 | 49 | 34 | 7 | 11 | 103 | 21 | 509 | 928 | 239 | 65 | 605 | 168 | 47 | 277 |
| PBANKA_0111400 | 35 | 49 | 21 | 10 | 57 | 33 | 84 | 40 | 12 | 30 | 7 | 13 | 282 | 61 | 127 | 150 | 183 | 76 | 613 | 260 | 36 | 185 |
| PBANKA_0111500 | 0 | 0 | 10 | 0 | 35 | 0 | 21 | 15 | 8 | 12 | 0 | 0 | 114 | 14 | 47 | 121 | 136 | 47 | 309 | 136 | 12 | 81 |
| PBANKA_0111600 | 0 | 0 | 15 | 0 | 43 | 14 | 183 | 72 | 859 | 449 | 162 | 41 | 897 | 305 | 55 | 242 | 10167 | 2493 | 853 | 775 | 716 | 4826 |
| PBANKA_0111700 | 0 | 7 | 79 | 28 | 201 | 151 | 303 | 137 | 116 | 61 | 18 | 7 | 419 | 78 | 231 | 849 | 1068 | 228 | 2200 | 1480 | 139 | 911 |
| PBANKA_0111800 | 128 | 136 | 101 | 31 | 333 | 152 | 454 | 171 | 377 | 220 | 105 | 45 | 1488 | 709 | 577 | 2762 | 1818 | 342 | 1159 | 1060 | 338 | 1690 |
| PBANKA_0111900 | 92 | 145 | 37 | 9 | 90 | 38 | 123 | 30 | 188 | 119 | 59 | 19 | 763 | 302 | 255 | 1177 | 3503 | 911 | 1141 | 1202 | 346 | 1665 |
| PBANKA_0112000 | 17 | 34 | 125 | 26 | 241 | 95 | 258 | 113 | 158 | 103 | 63 | 26 | 1666 | 748 | 438 | 1930 | 1614 | 275 | 752 | 605 | 890 | 4247 |
| PBANKA_0112100 | 28 | 43 | 13 | 0 | 235 | 76 | 1870 | 726 | 2692 | 1133 | 340 | 106 | 2383 | 2454 | 122 | 395 | 86054 | 8896 | 1443 | 5884 | 165 | 1109 |
| PBANKA_0112200 | 18 | 10 | 13 | 12 | 293 | 207 | 517 | 156 | 5570 | 3875 | 2048 | 1073 | 825 | 1805 | 101 | 322 | 19804 | 3371 | 261 | 1743 | 1635 | 10936 |
| PBANKA_0112300 | 29 | 44 | 26 | 8 | 126 | 69 | 254 | 104 | 328 | 149 | 56 | 27 | 391 | 267 | 207 | 1065 | 4876 | 890 | 510 | 1101 | 640 | 4833 |
| PBANKA_0112400 | 313 | 500 | 66 | 23 | 228 | 100 | 246 | 111 | 183 | 112 | 95 | 57 | 1220 | 1030 | 310 | 1416 | 885 | 236 | 603 | 537 | 935 | 5742 |
| PBANKA_0112500 | 143 | 209 | 29 | 7 | 216 | 117 | 382 | 132 | 376 | 266 | 940 | 696 | 1387 | 918 | 308 | 1559 | 893 | 237 | 479 | 419 | 105 | 761 |
| PBANKA_0112600 | 0 | 0 | 62 | 12 | 223 | 114 | 371 | 138 | 228 | 121 | 1718 | 1179 | 3862 | 2686 | 1322 | 6977 | 2955 | 287 | 313 | 363 | 11 | 125 |
| PBANKA_0112701 | 0 | 0 | 9 | 0 | 0 | 0 | 23 | 15 | 26 | 12 | 86 | 35 | 1473 | 1014 | 431 | 1247 | 806 | 195 | 46 | 20 | 0 | 19 |
| PBANKA_0200400 | 38 | 32 | 7 | 6 | 43 | 30 | 63 | 41 | 225 | 152 | 2063 | 1734 | 1531 | 962 | 152 | 752 | 792 | 327 | 769 | 684 | 10 | 131 |
| PBANKA_0200600 | 7 | 24 | 13 | 0 | 77 | 45 | 111 | 21 | 53 | 17 | 291 | 157 | 591 | 718 | 222 | 892 | 650 | 86 | 1081 | 744 | 50 | 708 |
| PBANKA_0200700 | 0 | 0 | 13 | 0 | 11 | 9 | 10 | 0 | 19 | 0 | 0 | 0 | 1135 | 162 | 141 | 329 | 59 | 24 | 70 | 21 | 5 | 34 |
| PBANKA_0201051 | 12 | 18 | 6 | 0 | 11 | 0 | 39 | 12 | 20 | 15 | 0 | 0 | 34 | 79 | 100 | 318 | 380 | 49 | 107 | 294 | 157 | 1106 |
| PBANKA_0201340 | 0 | 0 | 9 | 0 | 74 | 36 | 83 | 52 | 23 | 16 | 8 | 0 | 1051 | 215 | 532 | 2537 | 1427 | 195 | 743 | 518 | 0 | 59 |
| PBANKA_0201500 | 0 | 0 | 14 | 0 | 33 | 20 | 40 | 17 | 134 | 89 | 1946 | 1767 | 3737 | 1981 | 1211 | 4230 | 3223 | 504 | 229 | 222 | 12 | 98 |
| PBANKA_0201600 | 0 | 0 | 15 | 14 | 210 | 141 | 919 | 629 | 2160 | 1839 | 7676 | 6896 | 15145 | 6389 | 8223 | 18692 | 28823 | 7851 | 3752 | 1649 | 332 | 3631 |
| PBANKA_0201800 | 0 | 0 | 72 | 34 | 136 | 62 | 107 | 34 | 29 | 14 | 7 | 0 | 298 | 31 | 581 | 2250 | 264 | 30 | 359 | 216 | 84 | 537 |
| PBANKA_0201900 | 0 | 0 | 19 | 6 | 26 | 13 | 29 | 8 | 8 | 5 | 0 | 0 | 118 | 20 | 135 | 464 | 136 | 33 | 449 | 254 | 40 | 261 |
| PBANKA_0202000 | 6 | 6 | 172 | 53 | 300 | 130 | 235 | 92 | 91 | 28 | 22 | 0 | 2897 | 716 | 732 | 4169 | 947 | 227 | 2233 | 1266 | 398 | 2087 |
| PBANKA_0202100 | 0 | 0 | 18 | 0 | 29 | 16 | 43 | 14 | 8 | 9 | 0 | 0 | 88 | 12 | 30 | 121 | 39 | 8 | 80 | 23 | 10 | 44 |
| PBANKA_0202200 | 0 | 0 | 43 | 7 | 87 | 47 | 105 | 30 | 45 | 13 | 8 | 0 | 473 | 52 | 179 | 826 | 208 | 45 | 474 | 188 | 44 | 221 |
| PBANKA_0202300 | 0 | 0 | 126 | 30 | 103 | 67 | 127 | 57 | 34 | 31 | 0 | 0 | 1484 | 474 | 354 | 1299 | 508 | 140 | 1335 | 429 | 43 | 358 |
| PBANKA_0202400 | 14 | 30 | 39 | 11 | 127 | 54 | 333 | 108 | 140 | 78 | 17 | 6 | 462 | 151 | 91 | 338 | 2238 | 362 | 688 | 795 | 597 | 5205 |
| PBANKA_0202500 | 15 | 32 | 50 | 9 | 260 | 83 | 406 | 178 | 165 | 70 | 211 | 145 | 1414 | 512 | 290 | 869 | 4797 | 734 | 6062 | 4283 | 110 | 739 |
| PBANKA_0202600 | 0 | 0 | 19 | 6 | 21 | 10 | 17 | 5 | 7 | 10 | 0 | 0 | 35 | 12 | 22 | 56 | 5 | 0 | 151 | 37 | 9 | 61 |
| PBANKA_0202700 | 28 | 38 | 15 | 0 | 34 | 18 | 41 | 24 | 54 | 19 | 12 | 0 | 452 | 198 | 223 | 1021 | 1061 | 249 | 3013 | 3412 | 105 | 863 |
| PBANKA_0202800 | 24 | 30 | 67 | 15 | 51 | 28 | 45 | 13 | 34 | 7 | 23 | 13 | 384 | 147 | 173 | 581 | 148 | 43 | 326 | 100 | 100 | 586 |
| PBANKA_0202900 | 143 | 261 | 138 | 35 | 226 | 116 | 182 | 81 | 164 | 109 | 293 | 257 | 1350 | 758 | 444 | 1918 | 651 | 149 | 1216 | 709 | 891 | 6631 |
| PBANKA_0203000 | 301 | 525 | 227 | 76 | 1137 | 698 | 1610 | 790 | 442 | 207 | 54 | 36 | 3515 | 724 | 3217 | 8764 | 12218 | 2104 | 7052 | 2776 | 329 | 3202 |
| PBANKA_0203100 | 228 | 372 | 171 | 36 | 355 | 147 | 438 | 191 | 317 | 166 | 167 | 74 | 2818 | 1873 | 1156 | 5426 | 5707 | 898 | 2266 | 2767 | 373 | 2438 |
| PBANKA_0203200 | 0 | 0 | 78 | 16 | 261 | 68 | 328 | 102 | 198 | 96 | 1052 | 563 | 1796 | 2128 | 710 | 4339 | 3228 | 488 | 1032 | 1620 | 67 | 598 |
| PBANKA_0203300 | 0 | 5 | 144 | 30 | 292 | 102 | 272 | 127 | 114 | 60 | 187 | 118 | 657 | 436 | 195 | 712 | 439 | 74 | 307 | 138 | 87 | 622 |
| PBANKA_0203400 | 6 | 5 | 30 | 0 | 32 | 23 | 24 | 9 | 16 | 6 | 0 | 0 | 152 | 15 | 111 | 447 | 226 | 57 | 329 | 206 | 811 | 5027 |
| PBANKA_0203500 | 34 | 39 | 133 | 35 | 294 | 140 | 345 | 125 | 116 | 63 | 13 | 0 | 1099 | 280 | 495 | 3004 | 1661 | 284 | 1845 | 1026 | 1472 | 9485 |
| PBANKA_0203600 | 0 | 0 | 22 | 5 | 55 | 21 | 74 | 15 | 20 | 17 | 0 | 0 | 570 | 253 | 150 | 529 | 413 | 197 | 11844 | 5886 | 588 | 4268 |
| PBANKA_0203650 | 11 | 29 | 112 | 28 | 178 | 172 | 220 | 98 | 108 | 76 | 5 | 0 | 122 | 11 | 189 | 421 | 363 | 105 | 215 | 121 | 113 | 411 |
| PBANKA_0203750 | 60 | 92 | 14 | 0 | 108 | 35 | 158 | 55 | 172 | 71 | 9 | 0 | 270 | 57 | 112 | 266 | 1253 | 334 | 1360 | 629 | 159 | 1106 |
| PBANKA_0203800 | 20 | 21 | 133 | 25 | 249 | 85 | 281 | 95 | 178 | 92 | 252 | 184 | 2354 | 1850 | 774 | 4815 | 2127 | 337 | 1000 | 1142 | 85 | 531 |
| PBANKA_0203900 | 10 | 19 | 35 | 0 | 38 | 22 | 85 | 34 | 98 | 60 | 195 | 90 | 958 | 444 | 213 | 746 | 1104 | 327 | 1152 | 587 | 70 | 322 |
| PBANKA_0204000 | 0 | 0 | 47 | 18 | 37 | 25 | 28 | 0 | 0 | 0 | 0 | 0 | 186 | 31 | 140 | 581 | 47 | 14 | 147 | 40 | 16 | 76 |
| PBANKA_0204100 | 5 | 17 | 96 | 15 | 83 | 46 | 82 | 42 | 28 | 19 | 217 | 143 | 1068 | 442 | 387 | 1778 | 475 | 65 | 612 | 319 | 503 | 3986 |
| PBANKA_0204200 | 0 | 0 | 66 | 15 | 142 | 55 | 175 | 78 | 46 | 18 | 6 | 5 | 285 | 76 | 190 | 352 | 456 | 109 | 1547 | 517 | 111 | 647 |
| PBANKA_0204300 | 0 | 0 | 16 | 0 | 57 | 25 | 87 | 17 | 19 | 15 | 7 | 0 | 139 | 19 | 40 | 116 | 98 | 36 | 867 | 386 | 22 | 114 |
| PBANKA_0204400 | 0 | 0 | 50 | 15 | 127 | 69 | 111 | 46 | 41 | 12 | 5 | 6 | 875 | 234 | 301 | 1259 | 522 | 158 | 1692 | 1003 | 105 | 899 |
| PBANKA_0204500 | 5 | 17 | 25 | 9 | 53 | 21 | 81 | 28 | 50 | 15 | 9 | 0 | 699 | 377 | 107 | 231 | 1072 | 519 | 14183 | 13847 | 599 | 5817 |
| PBANKA_0204600 | 15 | 18 | 10 | 0 | 73 | 27 | 111 | 60 | 1240 | 1183 | 276 | 183 | 503 | 272 | 13 | 39 | 984 | 898 | 2222 | 1067 | 4106 | 39438 |
| PBANKA_0204700 | 29 | 33 | 97 | 20 | 171 | 80 | 164 | 68 | 64 | 19 | 20 | 0 | 691 | 286 | 222 | 896 | 593 | 142 | 691 | 467 | 622 | 4025 |
| PBANKA_0204800 | 276 | 485 | 158 | 48 | 552 | 213 | 702 | 288 | 314 | 177 | 535 | 312 | 4637 | 3149 | 935 | 6215 | 5637 | 693 | 1860 | 2511 | 2163 | 13088 |
| PBANKA_0204900 | 0 | 0 | 41 | 12 | 194 | 70 | 223 | 98 | 67 | 51 | 116 | 51 | 379 | 99 | 239 | 793 | 524 | 158 | 499 | 231 | 91 | 370 |
| PBANKA_0205000 | 26 | 30 | 313 | 81 | 183 | 129 | 199 | 96 | 580 | 390 | 8030 | 6329 | 10300 | 11395 | 423 | 2784 | 2006 | 544 | 1890 | 2514 | 251 | 1998 |
| PBANKA_0205100 | 0 | 0 | 75 | 16 | 350 | 162 | 416 | 163 | 145 | 118 | 356 | 274 | 775 | 387 | 122 | 439 | 272 | 85 | 610 | 262 | 86 | 402 |
| PBANKA_0205200 | 340 | 438 | 489 | 153 | 1065 | 444 | 1574 | 563 | 954 | 399 | 1111 | 756 | 7160 | 10475 | 3630 | 24448 | 26535 | 2683 | 4088 | 9178 | 2656 | 18901 |
| PBANKA_0205300 | 0 | 5 | 57 | 9 | 176 | 74 | 178 | 67 | 58 | 25 | 12 | 20 | 546 | 155 | 338 | 1582 | 581 | 79 | 680 | 452 | 102 | 919 |
| PBANKA_0205400 | 0 | 0 | 205 | 48 | 451 | 280 | 598 | 277 | 160 | 126 | 20 | 8 | 597 | 145 | 673 | 2309 | 1230 | 179 | 588 | 255 | 189 | 1589 |
| PBANKA_0205500 | 6 | 0 | 32 | 6 | 69 | 20 | 47 | 11 | 63 | 22 | 28 | 11 | 659 | 195 | 92 | 253 | 125 | 47 | 213 | 129 | 56 | 345 |
| PBANKA_0205600 | 11 | 11 | 71 | 14 | 134 | 56 | 118 | 39 | 90 | 31 | 28 | 23 | 797 | 305 | 395 | 1925 | 436 | 94 | 600 | 732 | 100 | 674 |
| PBANKA_0205700 | 6 | 0 | 38 | 0 | 46 | 17 | 31 | 0 | 11 | 7 | 0 | 0 | 39 | 22 | 86 | 603 | 113 | 14 | 37 | 41 | 73 | 483 |
| PBANKA_0205800 | 3750 | 5495 | 57 | 8 | 362 | 149 | 451 | 180 | 488 | 260 | 510 | 248 | 1541 | 1833 | 268 | 2418 | 2283 | 483 | 11699 | 36351 | 2503 | 14464 |
| PBANKA_0205900 | 678 | 684 | 160 | 41 | 475 | 163 | 618 | 198 | 804 | 384 | 286 | 131 | 7505 | 3857 | 1526 | 8253 | 9772 | 1767 | 2544 | 2708 | 338 | 2544 |
| PBANKA_0206000 | 0 | 0 | 47 | 23 | 167 | 85 | 285 | 110 | 121 | 76 | 8 | 0 | 473 | 101 | 390 | 1504 | 2483 | 412 | 1934 | 1223 | 105 | 757 |
| PBANKA_0206100 | 45 | 54 | 74 | 15 | 230 | 104 | 245 | 104 | 115 | 63 | 23 | 6 | 136 | 51 | 165 | 611 | 271 | 73 | 433 | 204 | 180 | 1051 |
| PBANKA_0206200 | 27 | 52 | 60 | 8 | 203 | 62 | 291 | 105 | 110 | 43 | 16 | 0 | 341 | 73 | 458 | 2441 | 614 | 120 | 1234 | 735 | 102 | 541 |
| PBANKA_0206300 | 6 | 6 | 56 | 17 | 216 | 143 | 564 | 389 | 277 | 306 | 45 | 17 | 1064 | 147 | 257 | 350 | 2206 | 632 | 1987 | 820 | 78 | 403 |
| PBANKA_0206400 | 0 | 0 | 15 | 7 | 141 | 61 | 221 | 85 | 131 | 58 | 13 | 0 | 320 | 113 | 163 | 552 | 688 | 171 | 1102 | 395 | 138 | 1077 |
| PBANKA_0206500 | 0 | 0 | 80 | 16 | 372 | 293 | 941 | 457 | 507 | 267 | 105 | 51 | 301 | 77 | 206 | 520 | 682 | 263 | 550 | 244 | 31 | 264 |
| PBANKA_0206600 | 12 | 25 | 185 | 42 | 311 | 161 | 302 | 121 | 349 | 214 | 2147 | 1307 | 3373 | 4948 | 907 | 5235 | 2360 | 436 | 448 | 1624 | 296 | 1902 |
| PBANKA_0206800 | 5 | 0 | 49 | 7 | 186 | 77 | 294 | 96 | 158 | 62 | 85 | 30 | 689 | 503 | 414 | 2021 | 4112 | 429 | 1930 | 3078 | 603 | 3974 |
| PBANKA_0206900 | 0 | 0 | 44 | 15 | 75 | 34 | 28 | 18 | 8 | 7 | 141 | 125 | 322 | 200 | 110 | 476 | 47 | 6 | 129 | 101 | 9 | 105 |
| PBANKA_0207000 | 20 | 47 | 298 | 75 | 715 | 358 | 1023 | 345 | 476 | 196 | 231 | 102 | 3358 | 1286 | 2382 | 10017 | 7182 | 1039 | 4206 | 3673 | 526 | 4665 |
| PBANKA_0207200 | 20 | 11 | 53 | 13 | 198 | 91 | 243 | 84 | 69 | 29 | 21 | 16 | 1167 | 466 | 485 | 1677 | 557 | 141 | 1524 | 619 | 66 | 652 |
| PBANKA_0207300 | 32 | 40 | 65 | 15 | 285 | 110 | 329 | 127 | 166 | 72 | 23 | 8 | 668 | 332 | 211 | 639 | 2704 | 388 | 537 | 646 | 633 | 3147 |
| PBANKA_0207400 | 0 | 0 | 37 | 0 | 131 | 69 | 251 | 91 | 89 | 48 | 19 | 0 | 330 | 69 | 155 | 659 | 782 | 153 | 1091 | 617 | 83 | 597 |

| | | | | | | | | | | | | | | | | | | | | | | |
|---|---|---|---|---|---|---|---|---|---|---|---|---|---|---|---|---|---|---|---|---|---|---|
| PBANKA_0207500 | 66 | 126 | 121 | 33 | 541 | 258 | 1118 | 455 | 624 | 262 | 78 | 42 | 1046 | 432 | 199 | 558 | 7168 | 1052 | 2054 | 1972 | 131 | 1089 |
| PBANKA_0207600 | 82 | 129 | 170 | 45 | 1448 | 653 | 2190 | 768 | 918 | 413 | 215 | 73 | 374 | 244 | 486 | 2041 | 1250 | 214 | 2339 | 4736 | 2763 | 21616 |
| PBANKA_0207700 | 6 | 7 | 23 | 26 | 223 | 96 | 296 | 76 | 114 | 47 | 72 | 75 | 509 | 398 | 533 | 2811 | 900 | 130 | 578 | 900 | 1273 | 7741 |
| PBANKA_0207800 | 0 | 5 | 80 | 13 | 168 | 83 | 216 | 94 | 83 | 51 | 10 | 5 | 339 | 35 | 269 | 1049 | 394 | 112 | 166 | 61 | 7 | 69 |
| PBANKA_0208000 | 0 | 0 | 35 | 0 | 119 | 51 | 361 | 211 | 148 | 93 | 16 | 6 | 460 | 77 | 79 | 243 | 2443 | 459 | 1235 | 568 | 90 | 725 |
| PBANKA_0208100 | 46 | 57 | 172 | 46 | 301 | 160 | 384 | 131 | 142 | 55 | 8 | 9 | 2078 | 681 | 730 | 2764 | 1351 | 242 | 1595 | 933 | 490 | 3461 |
| PBANKA_0208200 | 626 | 614 | 142 | 38 | 303 | 168 | 396 | 199 | 158 | 90 | 21 | 10 | 434 | 121 | 190 | 706 | 1002 | 304 | 530 | 314 | 125 | 738 |
| PBANKA_0208300 | 476 | 539 | 573 | 135 | 836 | 564 | 924 | 318 | 307 | 152 | 12 | 18 | 640 | 130 | 317 | 921 | 2479 | 439 | 1175 | 724 | 78 | 578 |
| PBANKA_0208400 | 235 | 192 | 32 | 11 | 531 | 178 | 531 | 132 | 165 | 70 | 26 | 15 | 212 | 63 | 39 | 205 | 206 | 72 | 2106 | 1493 | 433 | 3346 |
| PBANKA_0208500 | 0 | 0 | 102 | 16 | 212 | 96 | 156 | 44 | 81 | 22 | 25 | 12 | 524 | 313 | 430 | 2161 | 316 | 52 | 1707 | 567 | 123 | 849 |
| PBANKA_0208600 | 0 | 0 | 91 | 20 | 133 | 26 | 122 | 40 | 48 | 14 | 18 | 5 | 548 | 160 | 277 | 1206 | 301 | 38 | 725 | 418 | 34 | 186 |
| PBANKA_0208700 | 0 | 0 | 233 | 37 | 331 | 153 | 334 | 105 | 80 | 43 | 93 | 64 | 3077 | 1362 | 1077 | 4283 | 1241 | 206 | 1027 | 770 | 44 | 337 |
| PBANKA_0208800 | 10 | 10 | 189 | 43 | 1683 | 492 | 5600 | 1862 | 4417 | 1547 | 4306 | 2097 | 7427 | 18708 | 4920 | 27994 | 39726 | 5658 | 2186 | 9864 | 1292 | 5815 |
| PBANKA_0209100 | 1473 | 1445 | 15 | 0 | 83 | 45 | 316 | 162 | 208 | 99 | 35 | 13 | 186 | 44 | 0 | 8 | 825 | 183 | 327 | 195 | 696 | 3469 |
| PBANKA_0209200 | 137 | 95 | 54 | 18 | 213 | 109 | 336 | 142 | 138 | 80 | 10 | 17 | 841 | 217 | 529 | 1885 | 1645 | 289 | 1222 | 631 | 250 | 1546 |
| PBANKA_0209300 | 0 | 5 | 12 | 0 | 409 | 217 | 664 | 411 | 348 | 216 | 38 | 45 | 709 | 242 | 19 | 60 | 1099 | 324 | 10213 | 5008 | 1129 | 7224 |
| PBANKA_0209400 | 13 | 12 | 39 | 6 | 212 | 78 | 216 | 121 | 122 | 79 | 13 | 8 | 512 | 124 | 434 | 1421 | 504 | 128 | 2168 | 729 | 77 | 240 |
| PBANKA_0209500 | 70 | 99 | 187 | 30 | 249 | 90 | 192 | 74 | 189 | 122 | 93 | 61 | 3547 | 2894 | 995 | 5646 | 1808 | 384 | 1947 | 4050 | 410 | 2152 |
| PBANKA_0209600 | 15 | 20 | 8 | 0 | 107 | 56 | 103 | 41 | 107 | 26 | 32 | 15 | 237 | 64 | 238 | 907 | 446 | 84 | 1398 | 637 | 123 | 635 |
| PBANKA_0209700 | 561 | 881 | 13 | 0 | 206 | 76 | 631 | 307 | 637 | 429 | 112 | 72 | 215 | 184 | 212 | 916 | 545 | 121 | 1321 | 5483 | 8538 | 57067 |
| PBANKA_0209800 | 71 | 108 | 40 | 0 | 136 | 41 | 207 | 63 | 166 | 66 | 76 | 31 | 357 | 292 | 166 | 821 | 570 | 73 | 191 | 342 | 1526 | 10341 |
| PBANKA_0209900 | 24 | 26 | 367 | 84 | 606 | 311 | 512 | 145 | 153 | 91 | 33 | 12 | 569 | 177 | 396 | 1572 | 400 | 50 | 271 | 169 | 104 | 667 |
| PBANKA_0210000 | 0 | 11 | 93 | 16 | 239 | 101 | 483 | 166 | 283 | 130 | 58 | 19 | 1662 | 484 | 631 | 3774 | 3897 | 506 | 2892 | 2726 | 308 | 1635 |
| PBANKA_0210100 | 17 | 12 | 8 | 0 | 92 | 62 | 169 | 89 | 96 | 62 | 11 | 7 | 116 | 54 | 8 | 25 | 715 | 120 | 726 | 473 | 83 | 550 |
| PBANKA_0210200 | 13 | 12 | 442 | 93 | 699 | 232 | 581 | 173 | 240 | 97 | 457 | 336 | 3870 | 2570 | 1272 | 5852 | 1551 | 239 | 5499 | 2821 | 650 | 3991 |
| PBANKA_0210400 | 221 | 410 | 122 | 33 | 561 | 222 | 992 | 470 | 651 | 290 | 160 | 54 | 1444 | 625 | 737 | 3627 | 6676 | 907 | 3507 | 5242 | 789 | 4410 |
| PBANKA_0210500 | 197 | 256 | 281 | 62 | 937 | 300 | 2457 | 957 | 1635 | 737 | 472 | 185 | 1019 | 1055 | 1367 | 4853 | 6329 | 911 | 1431 | 2393 | 76 | 427 |
| PBANKA_0210600 | 288 | 558 | 642 | 145 | 1721 | 548 | 1469 | 526 | 642 | 252 | 304 | 177 | 3450 | 3928 | 6777 | 27500 | 9093 | 1271 | 4581 | 7661 | 563 | 4393 |
| PBANKA_0210700 | 0 | 0 | 27 | 5 | 123 | 41 | 172 | 62 | 75 | 43 | 16 | 6 | 434 | 231 | 503 | 3784 | 3730 | 387 | 916 | 1820 | 63 | 509 |
| PBANKA_0210800 | 26 | 34 | 70 | 20 | 291 | 123 | 775 | 317 | 262 | 133 | 31 | 9 | 1123 | 364 | 453 | 1805 | 1526 | 324 | 319 | 186 | 59 | 400 |
| PBANKA_0210900 | 20 | 37 | 77 | 23 | 181 | 68 | 150 | 74 | 47 | 21 | 18 | 7 | 153 | 72 | 75 | 216 | 58 | 38 | 38 | 16 | 0 | 21 |
| PBANKA_0211000 | 43 | 38 | 89 | 20 | 153 | 62 | 167 | 52 | 81 | 50 | 48 | 17 | 982 | 430 | 230 | 1140 | 414 | 86 | 674 | 394 | 63 | 433 |
| PBANKA_0211100 | 58 | 99 | 48 | 9 | 179 | 53 | 267 | 81 | 110 | 34 | 39 | 20 | 599 | 323 | 236 | 1034 | 1653 | 257 | 355 | 252 | 158 | 901 |
| PBANKA_0211200 | 40 | 49 | 36 | 0 | 102 | 48 | 149 | 57 | 90 | 50 | 255 | 174 | 488 | 399 | 272 | 1282 | 829 | 131 | 504 | 870 | 151 | 1189 |
| PBANKA_0211300 | 1744 | 2892 | 130 | 23 | 245 | 128 | 329 | 149 | 163 | 109 | 39 | 19 | 945 | 351 | 460 | 1807 | 1373 | 229 | 949 | 882 | 814 | 6690 |
| PBANKA_0211400 | 10 | 9 | 81 | 8 | 276 | 105 | 286 | 129 | 99 | 41 | 25 | 0 | 466 | 91 | 576 | 2250 | 903 | 150 | 733 | 411 | 292 | 1341 |
| PBANKA_0211500 | 0 | 0 | 309 | 108 | 456 | 400 | 652 | 384 | 191 | 133 | 18 | 8 | 1064 | 168 | 1242 | 3861 | 1681 | 263 | 606 | 303 | 521 | 4790 |
| PBANKA_0211600 | 0 | 0 | 71 | 19 | 223 | 74 | 318 | 130 | 119 | 50 | 38 | 8 | 984 | 414 | 396 | 1904 | 1717 | 190 | 1545 | 1432 | 214 | 1644 |
| PBANKA_0211700 | 18 | 24 | 31 | 5 | 76 | 33 | 175 | 64 | 88 | 37 | 16 | 5 | 300 | 113 | 132 | 789 | 291 | 41 | 976 | 1211 | 58 | 353 |
| PBANKA_0211800 | 0 | 0 | 18 | 0 | 139 | 40 | 203 | 49 | 91 | 30 | 12 | 9 | 689 | 149 | 287 | 1528 | 819 | 133 | 496 | 301 | 50 | 346 |
| PBANKA_0211900 | 6 | 0 | 162 | 51 | 449 | 171 | 650 | 259 | 268 | 116 | 45 | 13 | 1723 | 923 | 964 | 4120 | 3950 | 560 | 1181 | 1205 | 125 | 921 |
| PBANKA_0212000 | 16 | 35 | 70 | 7 | 128 | 42 | 171 | 61 | 451 | 184 | 416 | 178 | 1020 | 1167 | 149 | 917 | 1721 | 463 | 810 | 729 | 113 | 537 |
| PBANKA_0212100 | 7 | 20 | 987 | 238 | 1375 | 793 | 1349 | 545 | 388 | 241 | 596 | 515 | 9955 | 3617 | 2565 | 11331 | 4406 | 590 | 3299 | 2606 | 349 | 2605 |
| PBANKA_0212200 | 127 | 176 | 285 | 52 | 1001 | 388 | 1704 | 617 | 835 | 340 | 97 | 44 | 948 | 806 | 2227 | 8566 | 9720 | 1108 | 980 | 3087 | 264 | 1993 |
| PBANKA_0212300 | 0 | 0 | 27 | 0 | 140 | 46 | 178 | 60 | 62 | 27 | 14 | 5 | 446 | 102 | 210 | 1104 | 705 | 101 | 836 | 747 | 63 | 506 |
| PBANKA_0212400 | 7 | 0 | 93 | 17 | 429 | 226 | 1000 | 467 | 331 | 169 | 48 | 12 | 377 | 64 | 472 | 1483 | 516 | 108 | 689 | 491 | 103 | 852 |
| PBANKA_0212500 | 9 | 8 | 39 | 5 | 37 | 11 | 36 | 18 | 11 | 0 | 0 | 0 | 91 | 0 | 74 | 292 | 98 | 29 | 612 | 245 | 39 | 287 |
| PBANKA_0212700 | 283 | 367 | 104 | 22 | 615 | 221 | 876 | 306 | 377 | 163 | 88 | 28 | 783 | 591 | 1343 | 6857 | 2097 | 354 | 2815 | 10005 | 924 | 5855 |
| PBANKA_0212800 | 14 | 15 | 6 | 6 | 111 | 49 | 234 | 85 | 368 | 213 | 35 | 19 | 283 | 105 | 74 | 334 | 2714 | 725 | 1716 | 1092 | 109 | 648 |
| PBANKA_0212900 | 0 | 0 | 66 | 12 | 231 | 75 | 645 | 265 | 426 | 209 | 51 | 21 | 371 | 91 | 475 | 2257 | 1390 | 301 | 662 | 554 | 553 | 3708 |
| PBANKA_0213000 | 162 | 159 | 95 | 27 | 365 | 195 | 389 | 179 | 174 | 87 | 457 | 281 | 1887 | 1273 | 706 | 3418 | 1344 | 202 | 1824 | 1002 | 111 | 633 |
| PBANKA_0213100 | 21 | 14 | 34 | 11 | 58 | 40 | 90 | 28 | 29 | 26 | 81 | 51 | 177 | 93 | 87 | 289 | 141 | 21 | 125 | 88 | 0 | 25 |
| PBANKA_0213200 | 39 | 97 | 323 | 115 | 1400 | 622 | 2242 | 783 | 1139 | 445 | 222 | 65 | 1360 | 1490 | 1712 | 6687 | 22238 | 2299 | 2223 | 9704 | 219 | 1883 |
| PBANKA_0213300 | 0 | 0 | 69 | 16 | 213 | 113 | 169 | 97 | 50 | 28 | 7 | 0 | 746 | 160 | 140 | 511 | 295 | 42 | 233 | 123 | 0 | 27 |
| PBANKA_0213400 | 205 | 318 | 285 | 37 | 358 | 151 | 348 | 119 | 210 | 117 | 1917 | 1351 | 3141 | 3137 | 599 | 3146 | 1012 | 134 | 417 | 595 | 876 | 6182 |
| PBANKA_0213500 | 215 | 305 | 172 | 46 | 147 | 57 | 82 | 38 | 120 | 61 | 1212 | 887 | 1882 | 1563 | 238 | 1446 | 341 | 87 | 1745 | 1600 | 365 | 2420 |
| PBANKA_0213600 | 328 | 487 | 74 | 30 | 443 | 229 | 455 | 159 | 388 | 254 | 188 | 97 | 1119 | 919 | 398 | 2119 | 2643 | 408 | 1023 | 1830 | 127 | 979 |
| PBANKA_0213700 | 19 | 27 | 43 | 12 | 96 | 55 | 114 | 52 | 38 | 33 | 7 | 0 | 685 | 170 | 176 | 940 | 496 | 52 | 1330 | 1269 | 105 | 383 |
| PBANKA_0213800 | 8 | 13 | 80 | 17 | 251 | 134 | 311 | 109 | 180 | 92 | 26 | 14 | 332 | 113 | 68 | 269 | 1042 | 343 | 3941 | 3391 | 278 | 2017 |
| PBANKA_0214100 | 25 | 27 | 10 | 5 | 31 | 18 | 99 | 54 | 61 | 41 | 14 | 0 | 148 | 35 | 58 | 152 | 24 | 0 | 81 | 21 | 0 | 9 |
| PBANKA_0214200 | 119 | 237 | 214 | 53 | 513 | 271 | 2026 | 923 | 1179 | 506 | 96 | 67 | 1525 | 764 | 870 | 4208 | 851 | 111 | 541 | 826 | 26 | 241 |
| PBANKA_0214300 | 15 | 18 | 145 | 43 | 624 | 379 | 869 | 275 | 396 | 152 | 42 | 27 | 1872 | 927 | 701 | 2544 | 1868 | 355 | 1204 | 760 | 42 | 345 |
| PBANKA_0214400 | 1153 | 1735 | 451 | 150 | 1337 | 898 | 1431 | 501 | 1266 | 785 | 374 | 163 | 1500 | 1079 | 2535 | 16369 | 10095 | 1392 | 1940 | 3936 | 154 | 1422 |
| PBANKA_0214500 | 0 | 0 | 11 | 0 | 83 | 82 | 227 | 80 | 157 | 130 | 43 | 18 | 263 | 63 | 98 | 245 | 626 | 301 | 973 | 303 | 13 | 101 |
| PBANKA_0214700 | 0 | 0 | 95 | 35 | 272 | 83 | 472 | 174 | 547 | 273 | 1880 | 1335 | 9858 | 7520 | 3151 | 15811 | 15984 | 2652 | 4385 | 3879 | 109 | 846 |
| PBANKA_0214800 | 6 | 5 | 91 | 24 | 248 | 155 | 370 | 148 | 174 | 133 | 1951 | 1690 | 4697 | 2598 | 455 | 1464 | 1464 | 377 | 5863 | 2762 | 46 | 338 |
| PBANKA_0214900 | 0 | 0 | 33 | 0 | 115 | 110 | 143 | 47 | 40 | 35 | 120 | 79 | 234 | 287 | 75 | 150 | 119 | 31 | 45 | 19 | 10 | 45 |
| PBANKA_0214951 | 0 | 0 | 7 | 0 | 47 | 12 | 39 | 5 | 6 | 11 | 0 | 0 | 107 | 19 | 66 | 356 | 160 | 28 | 15 | 15 | 0 | 19 |
| PBANKA_0300700 | 254 | 426 | 63 | 24 | 161 | 80 | 254 | 98 | 258 | 116 | 65 | 62 | 1313 | 1066 | 309 | 1820 | 8447 | 1233 | 2512 | 4929 | 1009 | 6537 |
| PBANKA_0300800 | 28 | 47 | 43 | 0 | 172 | 67 | 180 | 83 | 71 | 33 | 12 | 8 | 189 | 66 | 185 | 857 | 517 | 119 | 653 | 387 | 36 | 353 |
| PBANKA_0300900 | 0 | 9 | 34 | 0 | 70 | 28 | 77 | 27 | 29 | 20 | 0 | 0 | 32 | 14 | 58 | 182 | 99 | 22 | 196 | 83 | 7 | 69 |
| PBANKA_0301000 | 5 | 9 | 6 | 0 | 93 | 37 | 178 | 87 | 474 | 243 | 202 | 109 | 3297 | 4605 | 1495 | 11155 | 19383 | 3678 | 4340 | 27787 | 474 | 1787 |
| PBANKA_0301100 | 0 | 0 | 72 | 18 | 89 | 40 | 166 | 75 | 107 | 50 | 19 | 11 | 2269 | 800 | 482 | 2609 | 3022 | 500 | 4958 | 4784 | 139 | 1003 |
| PBANKA_0301200 | 0 | 0 | 42 | 5 | 104 | 41 | 162 | 68 | 69 | 34 | 12 | 0 | 212 | 60 | 132 | 393 | 575 | 153 | 1227 | 368 | 19 | 177 |
| PBANKA_0301300 | 31 | 31 | 102 | 55 | 161 | 116 | 175 | 77 | 52 | 42 | 6 | 6 | 1686 | 315 | 558 | 1278 | 1978 | 481 | 19672 | 5531 | 313 | 2792 |
| PBANKA_0301500 | 15 | 20 | 115 | 33 | 105 | 47 | 89 | 36 | 122 | 105 | 1718 | 1236 | 1622 | 2069 | 270 | 2029 | 466 | 72 | 274 | 618 | 204 | 1572 |
| PBANKA_0301600 | 23 | 37 | 20 | 0 | 48 | 43 | 65 | 23 | 29 | 15 | 7 | 0 | 1985 | 726 | 70 | 291 | 771 | 330 | 49722 | 34246 | 1545 | 10201 |
| PBANKA_0301700 | 0 | 0 | 18 | 0 | 89 | 37 | 157 | 63 | 58 | 37 | 7 | 0 | 190 | 61 | 209 | 669 | 398 | 65 | 1247 | 878 | 109 | 817 |
| PBANKA_0301900 | 107 | 178 | 61 | 17 | 329 | 86 | 802 | 260 | 794 | 288 | 257 | 123 | 1385 | 1310 | 516 | 3140 | 8366 | 1203 | 1706 | 2876 | 1060 | 6240 |
| PBANKA_0302000 | 0 | 0 | 110 | 14 | 295 | 102 | 448 | 165 | 338 | 179 | 375 | 243 | 2399 | 863 | 466 | 1427 | 1750 | 922 | 2547 | 1620 | 123 | 1218 |
| PBANKA_0302100 | 14 | 16 | 20 | 0 | 94 | 18 | 113 | 43 | 53 | 20 | 24 | 12 | 767 | 611 | 263 | 1818 | 1377 | 212 | 1936 | 3268 | 1449 | 7238 |
| PBANKA_0302300 | 13 | 25 | 95 | 21 | 67 | 38 | 76 | 31 | 50 | 14 | 312 | 191 | 1089 | 884 | 204 | 872 | 295 | 42 | 719 | 466 | 162 | 1021 |
| PBANKA_0302400 | 0 | 0 | 161 | 29 | 117 | 41 | 77 | 41 | 59 | 30 | 14 | 0 | 1060 | 527 | 270 | 1675 | 853 | 180 | 1597 | 497 | 133 | 690 |
| PBANKA_0302500 | 461 | 574 | 394 | 79 | 783 | 458 | 996 | 478 | 436 | 297 | 58 | 60 | 5516 | 1144 | 2272 | 8155 | 7669 | 1226 | 2938 | 1479 | 2156 | 14892 |
| PBANKA_0302600 | 134 | 155 | 30 | 12 | 49 | 32 | 71 | 31 | 53 | 10 | 9 | 0 | 355 | 52 | 87 | 486 | 143 | 63 | 5051 | 2154 | 102 | 733 |
| PBANKA_0302700 | 7 | 10 | 35 | 19 | 79 | 24 | 65 | 31 | 19 | 0 | 8 | 0 | 161 | 26 | 227 | 1005 | 129 | 34 | 399 | 243 | 93 | 666 |
| PBANKA_0302800 | 13 | 13 | 22 | 0 | 53 | 37 | 40 | 18 | 56 | 32 | 27 | 0 | 185 | 71 | 144 | 402 | 351 | 78 | 1022 | 969 | 15 | 113 |
| PBANKA_0303100 | 0 | 0 | 131 | 13 | 288 | 120 | 381 | 123 | 143 | 55 | 21 | 13 | 1373 | 413 | 576 | 2458 | 939 | 167 | 868 | 472 | 219 | 1410 |
| PBANKA_0303200 | 0 | 0 | 69 | 8 | 104 | 61 | 172 | 43 | 68 | 36 | 11 | 32 | 393 | 131 | 190 | 615 | 484 | 84 | 209 | 98 | 87 | 543 |
| PBANKA_0303300 | 0 | 0 | 26 | 0 | 101 | 49 | 162 | 65 | 78 | 34 | 19 | 0 | 658 | 130 | 353 | 1727 | 1774 | 400 | 3960 | 1752 | 80 | 518 |

| | | | | | | | | | | | | | | | | | | | | | | |
|---|---|---|---|---|---|---|---|---|---|---|---|---|---|---|---|---|---|---|---|---|---|---|
| PBANKA_0303400 | 18 | 36 | 58 | 15 | 183 | 114 | 556 | 404 | 419 | 200 | 39 | 27 | 685 | 139 | 295 | 738 | 1006 | 409 | 1197 | 406 | 71 | 448 |
| PBANKA_0303500 | 0 | 0 | 11 | 0 | 26 | 20 | 91 | 37 | 48 | 22 | 6 | 0 | 155 | 46 | 24 | 150 | 418 | 82 | 1814 | 1048 | 150 | 1202 |
| PBANKA_0303600 | 5 | 24 | 702 | 207 | 2204 | 1359 | 3230 | 1140 | 1341 | 715 | 184 | 81 | 5966 | 1560 | 3777 | 12227 | 10167 | 1872 | 3064 | 1751 | 2018 | 14715 |
| PBANKA_0303700 | 128 | 208 | 462 | 84 | 1672 | 557 | 1705 | 536 | 773 | 257 | 215 | 74 | 1608 | 1314 | 1377 | 6914 | 9326 | 1144 | 2960 | 5892 | 616 | 4696 |
| PBANKA_0303800 | 111 | 192 | 57 | 14 | 217 | 82 | 243 | 150 | 114 | 66 | 36 | 10 | 289 | 74 | 87 | 217 | 657 | 268 | 1336 | 445 | 120 | 846 |
| PBANKA_0303900 | 765 | 1042 | 12 | 0 | 77 | 44 | 126 | 35 | 503 | 241 | 116 | 74 | 537 | 221 | 108 | 415 | 2073 | 604 | 3004 | 2368 | 1723 | 15891 |
| PBANKA_0304000 | 117 | 157 | 78 | 24 | 259 | 99 | 132 | 63 | 124 | 55 | 64 | 40 | 998 | 413 | 805 | 3567 | 1070 | 160 | 1303 | 859 | 290 | 1752 |
| PBANKA_0304100 | 233 | 227 | 30 | 0 | 86 | 22 | 103 | 31 | 92 | 30 | 367 | 207 | 595 | 568 | 306 | 1602 | 733 | 120 | 1304 | 2118 | 1912 | 14164 |
| PBANKA_0304200 | 0 | 5 | 17 | 8 | 50 | 26 | 27 | 10 | 6 | 0 | 23 | 13 | 237 | 141 | 80 | 259 | 36 | 13 | 113 | 95 | 29 | 180 |
| PBANKA_0304300 | 20 | 27 | 76 | 14 | 166 | 112 | 150 | 42 | 97 | 38 | 1229 | 810 | 458 | 400 | 194 | 1074 | 157 | 26 | 178 | 142 | 179 | 1392 |
| PBANKA_0304400 | 27 | 29 | 82 | 24 | 281 | 169 | 632 | 383 | 454 | 248 | 560 | 349 | 949 | 271 | 286 | 625 | 3423 | 1204 | 730 | 321 | 88 | 854 |
| PBANKA_0304500 | 313 | 583 | 47 | 8 | 328 | 96 | 1165 | 588 | 1662 | 863 | 409 | 131 | 1213 | 741 | 393 | 2232 | 6072 | 1504 | 2736 | 5036 | 3095 | 19611 |
| PBANKA_0304600 | 7 | 14 | 9 | 0 | 35 | 18 | 35 | 12 | 16 | 5 | 0 | 0 | 245 | 40 | 88 | 232 | 51 | 26 | 379 | 147 | 39 | 188 |
| PBANKA_0304700 | 2140 | 3361 | 5 | 0 | 25 | 20 | 12 | 19 | 28 | 13 | 0 | 0 | 21 | 9 | 23 | 41 | 27 | 14 | 34 | 24 | 15 | 179 |
| PBANKA_0304800 | 24 | 31 | 20 | 8 | 600 | 211 | 3930 | 1639 | 2322 | 1153 | 199 | 102 | 3005 | 819 | 645 | 1861 | 24601 | 3216 | 641 | 780 | 46 | 415 |
| PBANKA_0304900 | 38 | 55 | 187 | 63 | 2333 | 1551 | 11096 | 5258 | 10177 | 6170 | 5619 | 3505 | 10685 | 5926 | 3153 | 13905 | 43798 | 7247 | 2390 | 2132 | 78 | 638 |
| PBANKA_0305000 | 0 | 0 | 39 | 20 | 594 | 314 | 13517 | 5893 | 13525 | 7534 | 1420 | 663 | 16148 | 4193 | 338 | 1611 | 1E+05 | 28884 | 1401 | 1798 | 9 | 210 |
| PBANKA_0305100 | 10 | 14 | 215 | 90 | 5153 | 3371 | 25749 | 11548 | 21890 | 12772 | 2552 | 1278 | 22387 | 5855 | 698 | 1344 | 2E+05 | 29106 | 2492 | 2325 | 27 | 626 |
| PBANKA_0305200 | 0 | 7 | 45 | 7 | 297 | 138 | 1056 | 371 | 869 | 373 | 92 | 41 | 898 | 321 | 1004 | 4396 | 3185 | 465 | 629 | 289 | 968 | 5591 |
| PBANKA_0305300 | 112 | 140 | 165 | 29 | 140 | 54 | 121 | 54 | 126 | 59 | 1087 | 701 | 1276 | 786 | 219 | 889 | 237 | 42 | 496 | 388 | 284 | 1657 |
| PBANKA_0305400 | 115 | 151 | 192 | 19 | 497 | 141 | 612 | 194 | 442 | 203 | 253 | 123 | 1548 | 1451 | 566 | 2808 | 2306 | 427 | 1158 | 873 | 343 | 2009 |
| PBANKA_0305500 | 90 | 101 | 60 | 13 | 227 | 72 | 413 | 131 | 257 | 89 | 32 | 8 | 401 | 124 | 220 | 1182 | 542 | 188 | 904 | 825 | 167 | 1271 |
| PBANKA_0305600 | 0 | 0 | 118 | 26 | 430 | 190 | 440 | 171 | 109 | 64 | 18 | 11 | 47 | 12 | 101 | 225 | 245 | 54 | 112 | 55 | 0 | 40 |
| PBANKA_0305700 | 38 | 52 | 22 | 0 | 187 | 78 | 311 | 104 | 137 | 81 | 27 | 11 | 675 | 186 | 466 | 2255 | 3361 | 455 | 1175 | 825 | 478 | 2326 |
| PBANKA_0305800 | 5 | 5 | 246 | 56 | 404 | 192 | 384 | 134 | 78 | 54 | 26 | 22 | 459 | 156 | 226 | 586 | 285 | 48 | 451 | 224 | 111 | 834 |
| PBANKA_0305900 | 0 | 0 | 63 | 11 | 200 | 107 | 223 | 70 | 67 | 31 | 12 | 0 | 657 | 212 | 253 | 789 | 286 | 51 | 1733 | 733 | 170 | 1003 |
| PBANKA_0306000 | 251 | 243 | 302 | 55 | 698 | 314 | 860 | 254 | 672 | 228 | 241 | 150 | 543 | 150 | 192 | 101 | 228 | 164 | 2953 | 668 | 87 | 645 |
| PBANKA_0306100 | 13 | 14 | 13 | 0 | 76 | 42 | 75 | 25 | 75 | 34 | 21 | 12 | 254 | 147 | 82 | 203 | 820 | 120 | 2020 | 3206 | 35 | 270 |
| PBANKA_0306200 | 19 | 39 | 63 | 14 | 252 | 96 | 330 | 160 | 198 | 71 | 42 | 29 | 1852 | 720 | 568 | 3052 | 890 | 168 | 766 | 549 | 320 | 2913 |
| PBANKA_0306300 | 0 | 0 | 57 | 12 | 132 | 66 | 181 | 131 | 93 | 89 | 25 | 15 | 565 | 320 | 334 | 989 | 497 | 108 | 220 | 134 | 253 | 1481 |
| PBANKA_0306400 | 10 | 10 | 16 | 0 | 44 | 29 | 50 | 34 | 28 | 17 | 8 | 0 | 468 | 161 | 187 | 515 | 122 | 57 | 432 | 198 | 50 | 319 |
| PBANKA_0306500 | 36 | 27 | 79 | 11 | 177 | 98 | 231 | 99 | 48 | 40 | 12 | 0 | 399 | 75 | 610 | 1672 | 920 | 201 | 1348 | 359 | 99 | 792 |
| PBANKA_0306600 | 86 | 120 | 16 | 0 | 61 | 26 | 63 | 25 | 45 | 24 | 7 | 6 | 363 | 144 | 11 | 24 | 656 | 117 | 4477 | 2776 | 1658 | 9763 |
| PBANKA_0306700 | 0 | 0 | 35 | 0 | 152 | 84 | 1118 | 505 | 852 | 413 | 58 | 30 | 1168 | 291 | 40 | 133 | 9726 | 1902 | 383 | 339 | 128 | 775 |
| PBANKA_0306800 | 43 | 73 | 786 | 226 | 850 | 522 | 896 | 438 | 287 | 139 | 516 | 410 | 10863 | 3742 | 3822 | 17067 | 5478 | 665 | 2350 | 1701 | 2774 | 22352 |
| PBANKA_0306900 | 0 | 0 | 38 | 6 | 79 | 61 | 183 | 99 | 86 | 51 | 27 | 13 | 771 | 143 | 312 | 762 | 1313 | 333 | 4382 | 1312 | 295 | 2151 |
| PBANKA_0307000 | 99 | 102 | 1011 | 351 | 754 | 448 | 791 | 476 | 251 | 213 | 1208 | 1097 | 8244 | 3315 | 1728 | 5474 | 2461 | 483 | 1747 | 695 | 63 | 594 |
| PBANKA_0307100 | 0 | 0 | 297 | 60 | 862 | 308 | 1482 | 501 | 618 | 279 | 107 | 33 | 1797 | 1228 | 1875 | 8024 | 15303 | 1686 | 2611 | 8574 | 156 | 1476 |
| PBANKA_0307200 | 18 | 25 | 21 | 10 | 137 | 102 | 189 | 92 | 50 | 48 | 18 | 7 | 361 | 91 | 290 | 1530 | 720 | 170 | 2955 | 1185 | 77 | 576 |
| PBANKA_0307300 | 0 | 0 | 19 | 6 | 53 | 36 | 58 | 28 | 12 | 11 | 0 | 0 | 37 | 11 | 101 | 309 | 110 | 14 | 291 | 65 | 8 | 68 |
| PBANKA_0307400 | 182 | 248 | 100 | 28 | 297 | 95 | 333 | 104 | 190 | 111 | 77 | 39 | 1518 | 655 | 376 | 2107 | 1632 | 520 | 1288 | 1012 | 527 | 2885 |
| PBANKA_0307500 | 19 | 24 | 21 | 0 | 55 | 23 | 81 | 42 | 392 | 294 | 130 | 53 | 453 | 209 | 43 | 177 | 2328 | 725 | 383 | 328 | 87 | 531 |
| PBANKA_0307600 | 21 | 21 | 23 | 5 | 51 | 20 | 81 | 24 | 120 | 48 | 21 | 5 | 872 | 277 | 172 | 546 | 982 | 331 | 10463 | 6367 | 280 | 2161 |
| PBANKA_0307700 | 28 | 33 | 10 | 0 | 57 | 31 | 83 | 16 | 40 | 14 | 0 | 5 | 424 | 170 | 119 | 505 | 536 | 121 | 332 | 97 | 8 | 41 |
| PBANKA_0307800 | 101 | 146 | 198 | 38 | 287 | 156 | 271 | 121 | 163 | 113 | 716 | 637 | 3366 | 1731 | 770 | 3490 | 3069 | 519 | 9808 | 6977 | 358 | 2322 |
| PBANKA_0307900 | 57 | 47 | 55 | 5 | 68 | 33 | 63 | 24 | 19 | 0 | 0 | 0 | 632 | 247 | 300 | 1707 | 288 | 53 | 667 | 459 | 17 | 99 |
| PBANKA_0308000 | 9 | 6 | 66 | 0 | 79 | 28 | 86 | 26 | 45 | 34 | 8 | 0 | 247 | 82 | 246 | 898 | 1174 | 186 | 621 | 259 | 101 | 681 |
| PBANKA_0308100 | 0 | 0 | 102 | 26 | 219 | 105 | 221 | 112 | 51 | 41 | 12 | 7 | 450 | 92 | 339 | 1129 | 562 | 104 | 300 | 141 | 24 | 139 |
| PBANKA_0308200 | 0 | 0 | 80 | 19 | 279 | 152 | 218 | 84 | 41 | 29 | 0 | 0 | 26 | 6 | 18 | 24 | 43 | 16 | 319 | 135 | 23 | 205 |
| PBANKA_0308300 | 245 | 342 | 174 | 38 | 660 | 237 | 916 | 279 | 764 | 368 | 117 | 51 | 819 | 609 | 954 | 4737 | 10984 | 2057 | 2317 | 5969 | 368 | 2012 |
| PBANKA_0308400 | 0 | 0 | 27 | 8 | 45 | 20 | 40 | 12 | 20 | 7 | 5 | 0 | 39 | 24 | 105 | 398 | 110 | 24 | 111 | 63 | 323 | 2029 |
| PBANKA_0308500 | 46 | 61 | 173 | 24 | 228 | 51 | 197 | 67 | 94 | 34 | 357 | 182 | 2680 | 1708 | 523 | 3122 | 612 | 80 | 1115 | 996 | 685 | 3834 |
| PBANKA_0308600 | 75 | 147 | 851 | 274 | 771 | 478 | 661 | 325 | 316 | 239 | 463 | 347 | 4472 | 2425 | 3036 | 10559 | 2673 | 544 | 809 | 797 | 351 | 2470 |
| PBANKA_0308700 | 46 | 53 | 68 | 12 | 102 | 33 | 126 | 32 | 93 | 37 | 28 | 0 | 84 | 41 | 119 | 372 | 263 | 47 | 779 | 287 | 61 | 365 |
| PBANKA_0308800 | 19 | 17 | 19 | 0 | 105 | 76 | 240 | 96 | 158 | 99 | 28 | 18 | 245 | 50 | 221 | 787 | 786 | 166 | 641 | 276 | 235 | 1536 |
| PBANKA_0308900 | 0 | 0 | 54 | 6 | 136 | 71 | 372 | 139 | 263 | 138 | 51 | 54 | 695 | 405 | 397 | 2314 | 2529 | 339 | 517 | 544 | 599 | 3845 |
| PBANKA_0309000 | 0 | 0 | 72 | 25 | 175 | 60 | 153 | 60 | 46 | 30 | 11 | 0 | 435 | 138 | 296 | 1317 | 651 | 140 | 508 | 293 | 46 | 341 |
| PBANKA_0309100 | 36 | 37 | 280 | 98 | 410 | 256 | 474 | 217 | 183 | 105 | 269 | 210 | 3677 | 1690 | 709 | 3360 | 2304 | 353 | 799 | 726 | 187 | 1465 |
| PBANKA_0309200 | 212 | 282 | 255 | 55 | 725 | 218 | 1340 | 398 | 868 | 315 | 270 | 113 | 2128 | 2055 | 1326 | 5888 | 7485 | 1035 | 1983 | 3015 | 1076 | 6002 |
| PBANKA_0309400 | 116 | 129 | 574 | 155 | 1393 | 603 | 2062 | 783 | 1022 | 440 | 981 | 522 | 2710 | 5344 | 2352 | 13539 | 17386 | 1531 | 810 | 2981 | 1323 | 7216 |
| PBANKA_0309500 | 412 | 498 | 28 | 9 | 87 | 82 | 91 | 54 | 242 | 217 | 57 | 44 | 158 | 63 | 39 | 65 | 423 | 249 | 191 | 143 | 99 | 630 |
| PBANKA_0309600 | 33 | 39 | 66 | 19 | 149 | 70 | 207 | 96 | 85 | 51 | 21 | 0 | 177 | 25 | 255 | 847 | 412 | 103 | 231 | 86 | 21 | 185 |
| PBANKA_0309700 | 9 | 16 | 16 | 0 | 154 | 59 | 187 | 64 | 64 | 46 | 24 | 14 | 548 | 167 | 384 | 1428 | 749 | 143 | 692 | 450 | 40 | 239 |
| PBANKA_0309800 | 0 | 0 | 21 | 5 | 82 | 47 | 155 | 50 | 77 | 28 | 27 | 9 | 449 | 83 | 252 | 1029 | 958 | 138 | 574 | 323 | 23 | 113 |
| PBANKA_0309900 | 0 | 0 | 34 | 0 | 111 | 28 | 64 | 25 | 22 | 12 | 9 | 0 | 168 | 44 | 108 | 478 | 127 | 23 | 173 | 95 | 14 | 60 |
| PBANKA_0310000 | 18 | 18 | 362 | 102 | 543 | 319 | 607 | 253 | 227 | 114 | 187 | 192 | 4583 | 1977 | 1193 | 5046 | 2529 | 373 | 1283 | 876 | 1667 | 12157 |
| PBANKA_0310100 | 11 | 10 | 73 | 15 | 142 | 89 | 137 | 75 | 48 | 29 | 0 | 0 | 147 | 16 | 209 | 478 | 224 | 51 | 512 | 217 | 114 | 1015 |
| PBANKA_0310200 | 0 | 0 | 19 | 8 | 48 | 26 | 49 | 19 | 15 | 5 | 0 | 0 | 84 | 9 | 111 | 108 | 117 | 27 | 131 | 48 | 49 | 197 |
| PBANKA_0310300 | 10 | 8 | 65 | 11 | 176 | 96 | 186 | 75 | 46 | 33 | 12 | 0 | 255 | 58 | 238 | 622 | 545 | 161 | 2326 | 1156 | 134 | 967 |
| PBANKA_0310400 | 10 | 16 | 51 | 0 | 120 | 54 | 118 | 24 | 33 | 19 | 21 | 0 | 725 | 232 | 339 | 1209 | 335 | 104 | 538 | 239 | 160 | 1066 |
| PBANKA_0310500 | 8 | 0 | 14 | 0 | 63 | 28 | 45 | 15 | 124 | 65 | 270 | 123 | 890 | 566 | 71 | 306 | 1363 | 262 | 3077 | 3843 | 336 | 1755 |
| PBANKA_0310600 | 140 | 190 | 159 | 57 | 157 | 91 | 160 | 62 | 52 | 26 | 64 | 25 | 601 | 447 | 169 | 696 | 381 | 49 | 150 | 78 | 1490 | 9879 |
| PBANKA_0310800 | 11 | 8 | 66 | 19 | 218 | 111 | 213 | 68 | 123 | 52 | 67 | 35 | 1837 | 1029 | 520 | 3477 | 1664 | 230 | 1044 | 992 | 231 | 1777 |
| PBANKA_0310900 | 91 | 115 | 717 | 194 | 866 | 484 | 718 | 272 | 279 | 159 | 215 | 115 | 4045 | 1606 | 2422 | 9378 | 2838 | 481 | 2605 | 1744 | 359 | 3013 |
| PBANKA_0311000 | 147 | 277 | 443 | 139 | 1392 | 739 | 2195 | 832 | 1207 | 559 | 818 | 481 | 5440 | 2649 | 2672 | 9035 | 10281 | 1589 | 2737 | 1854 | 1179 | 10026 |
| PBANKA_0311100 | 29 | 30 | 105 | 29 | 232 | 115 | 196 | 97 | 83 | 57 | 24 | 29 | 348 | 60 | 371 | 1289 | 625 | 102 | 847 | 321 | 147 | 1100 |
| PBANKA_0311200 | 74 | 107 | 115 | 28 | 191 | 70 | 180 | 74 | 110 | 66 | 1580 | 1064 | 3772 | 2647 | 520 | 2543 | 766 | 121 | 933 | 884 | 223 | 1753 |
| PBANKA_0311400 | 571 | 998 | 14 | 0 | 21 | 10 | 49 | 24 | 603 | 345 | 190 | 54 | 523 | 635 | 26 | 125 | 3158 | 627 | 3975 | 11197 | 4999 | 32797 |
| PBANKA_0311600 | 7 | 13 | 69 | 25 | 290 | 106 | 216 | 86 | 722 | 399 | 366 | 201 | 8648 | 4576 | 1728 | 9497 | 5991 | 1790 | 2622 | 2811 | 197 | 1297 |
| PBANKA_0311700 | 8 | 13 | 488 | 114 | 767 | 391 | 1026 | 398 | 1162 | 597 | 227 | 119 | 1155 | 1147 | 699 | 3642 | 20042 | 3552 | 260 | 1162 | 105 | 652 |
| PBANKA_0311800 | 833 | 1038 | 352 | 80 | 818 | 424 | 701 | 249 | 1891 | 1119 | 12741 | 8932 | 12064 | 8482 | 687 | 3401 | 5499 | 1992 | 1586 | 1169 | 43 | 426 |
| PBANKA_0311900 | 38 | 44 | 121 | 45 | 293 | 185 | 514 | 164 | 233 | 122 | 40 | 18 | 404 | 107 | 108 | 354 | 1028 | 239 | 645 | 326 | 513 | 3074 |
| PBANKA_0312100 | 0 | 0 | 26 | 0 | 203 | 87 | 385 | 170 | 254 | 134 | 98 | 52 | 1441 | 452 | 618 | 2993 | 4861 | 931 | 2415 | 2508 | 532 | 4395 |
| PBANKA_0312200 | 0 | 0 | 47 | 9 | 157 | 58 | 178 | 74 | 79 | 34 | 51 | 44 | 892 | 405 | 348 | 1528 | 767 | 111 | 1416 | 857 | 189 | 1433 |
| PBANKA_0312300 | 66 | 69 | 66 | 16 | 161 | 74 | 300 | 145 | 112 | 64 | 86 | 81 | 1993 | 998 | 364 | 2119 | 2233 | 360 | 1952 | 1153 | 236 | 1528 |
| PBANKA_0312400 | 186 | 300 | 281 | 123 | 914 | 588 | 1603 | 716 | 704 | 319 | 1139 | 851 | 3074 | 1785 | 1368 | 6277 | 4287 | 456 | 2187 | 2454 | 583 | 6450 |
| PBANKA_0312500 | 12 | 35 | 87 | 20 | 283 | 124 | 509 | 167 | 253 | 148 | 32 | 17 | 912 | 270 | 66 | 217 | 4910 | 616 | 2946 | 1712 | 447 | 4082 |
| PBANKA_0312700 | 33 | 41 | 38 | 13 | 49 | 47 | 59 | 27 | 80 | 53 | 13 | 8 | 1197 | 1995 | 309 | 1143 | 1791 | 733 | 17489 | 71490 | 250 | 1801 |
| PBANKA_0312800 | 0 | 0 | 59 | 23 | 136 | 65 | 138 | 44 | 61 | 38 | 25 | 27 | 781 | 455 | 506 | 2488 | 582 | 120 | 639 | 417 | 866 | 5903 |
| PBANKA_0312900 | 0 | 7 | 13 | 0 | 73 | 28 | 80 | 35 | 32 | 17 | 10 | 0 | 412 | 195 | 63 | 287 | 208 | 33 | 58 | 31 | 23 | 189 |

| | | | | | | | | | | | | | | | | | | | | | | |
|---|---|---|---|---|---|---|---|---|---|---|---|---|---|---|---|---|---|---|---|---|---|---|
| PBANKA_0313000 | 11 | 22 | 59 | 7 | 180 | 80 | 187 | 60 | 68 | 34 | 8 | 11 | 419 | 247 | 163 | 851 | 383 | 50 | 912 | 378 | 503 | 3503 |
| PBANKA_0313100 | 8 | 13 | 24 | 0 | 236 | 98 | 249 | 100 | 88 | 32 | 17 | 0 | 897 | 285 | 349 | 1753 | 798 | 83 | 555 | 472 | 347 | 2201 |
| PBANKA_0313200 | 0 | 0 | 5 | 0 | 62 | 25 | 117 | 42 | 90 | 40 | 16 | 0 | 237 | 227 | 132 | 587 | 929 | 129 | 388 | 547 | 955 | 5511 |
| PBANKA_0313300 | 8 | 0 | 41 | 10 | 269 | 130 | 515 | 234 | 568 | 285 | 156 | 79 | 2988 | 1156 | 421 | 2989 | 11820 | 1325 | 3866 | 3087 | 4123 | 31568 |
| PBANKA_0313400 | 30 | 56 | 11 | 0 | 310 | 114 | 334 | 138 | 883 | 429 | 472 | 231 | 1048 | 3010 | 173 | 1280 | 17551 | 4008 | 447 | 5018 | 860 | 4574 |
| PBANKA_0313700 | 0 | 0 | 5 | 0 | 0 | 0 | 25 | 11 | 25 | 8 | 6 | 0 | 167 | 46 | 12 | 46 | 458 | 118 | 2389 | 1322 | 97 | 730 |
| PBANKA_0313800 | 0 | 8 | 417 | 102 | 823 | 348 | 1249 | 554 | 420 | 210 | 64 | 32 | 1714 | 592 | 2247 | 11092 | 10599 | 1221 | 4162 | 6718 | 1014 | 8702 |
| PBANKA_0313900 | 38 | 24 | 56 | 8 | 152 | 50 | 107 | 33 | 83 | 37 | 18 | 6 | 483 | 187 | 388 | 1950 | 758 | 156 | 1570 | 610 | 262 | 1328 |
| PBANKA_0314000 | 40 | 19 | 6 | 0 | 19 | 9 | 62 | 38 | 47 | 15 | 0 | 0 | 130 | 30 | 55 | 132 | 236 | 72 | 117 | 42 | 103 | 393 |
| PBANKA_0314100 | 78 | 92 | 58 | 15 | 123 | 75 | 108 | 55 | 79 | 49 | 17 | 11 | 639 | 170 | 235 | 810 | 748 | 129 | 939 | 584 | 287 | 2177 |
| PBANKA_0314200 | 71 | 99 | 46 | 11 | 223 | 141 | 608 | 232 | 741 | 368 | 98 | 44 | 1987 | 465 | 443 | 1334 | 13072 | 2899 | 3038 | 2515 | 187 | 1508 |
| PBANKA_0314300 | 0 | 5 | 77 | 16 | 249 | 71 | 332 | 85 | 260 | 130 | 205 | 134 | 1238 | 767 | 380 | 1908 | 1882 | 349 | 1708 | 831 | 850 | 5155 |
| PBANKA_0314400 | 0 | 0 | 61 | 17 | 107 | 49 | 77 | 22 | 26 | 7 | 22 | 20 | 635 | 144 | 153 | 864 | 125 | 22 | 751 | 327 | 353 | 2459 |
| PBANKA_0314500 | 21 | 13 | 1144 | 326 | 973 | 642 | 972 | 503 | 276 | 193 | 925 | 699 | 6485 | 2662 | 2032 | 5006 | 1357 | 287 | 1841 | 699 | 114 | 1092 |
| PBANKA_0314600 | 0 | 6 | 36 | 6 | 108 | 67 | 145 | 65 | 65 | 39 | 43 | 35 | 1349 | 503 | 559 | 2155 | 578 | 158 | 1135 | 532 | 78 | 589 |
| PBANKA_0314700 | 0 | 0 | 125 | 31 | 342 | 172 | 459 | 177 | 163 | 65 | 19 | 11 | 403 | 99 | 411 | 1530 | 1481 | 257 | 2776 | 1446 | 443 | 4107 |
| PBANKA_0314900 | 0 | 0 | 170 | 29 | 246 | 97 | 161 | 34 | 82 | 32 | 32 | 8 | 4093 | 2401 | 530 | 3757 | 484 | 116 | 1317 | 1282 | 619 | 4063 |
| PBANKA_0315000 | 0 | 0 | 89 | 23 | 56 | 33 | 47 | 11 | 23 | 6 | 6 | 0 | 1135 | 301 | 460 | 1529 | 250 | 52 | 69 | 59 | 21 | 99 |
| PBANKA_0315100 | 38 | 41 | 90 | 16 | 82 | 49 | 126 | 48 | 119 | 37 | 523 | 317 | 1834 | 1132 | 254 | 1013 | 808 | 219 | 743 | 428 | 687 | 3910 |
| PBANKA_0315200 | 0 | 0 | 111 | 20 | 131 | 73 | 118 | 70 | 46 | 27 | 44 | 13 | 763 | 207 | 310 | 843 | 374 | 61 | 205 | 138 | 25 | 159 |
| PBANKA_0315300 | 0 | 7 | 61 | 19 | 242 | 65 | 338 | 94 | 389 | 171 | 375 | 179 | 1080 | 1283 | 507 | 2743 | 3112 | 514 | 1045 | 2222 | 255 | 1474 |
| PBANKA_0315400 | 74 | 126 | 99 | 14 | 150 | 72 | 205 | 94 | 171 | 67 | 646 | 441 | 2184 | 2521 | 533 | 2547 | 1537 | 280 | 760 | 844 | 407 | 2405 |
| PBANKA_0315410 | 5 | 6 | 143 | 37 | 239 | 105 | 245 | 91 | 99 | 17 | 40 | 15 | 555 | 184 | 236 | 610 | 417 | 81 | 503 | 217 | 107 | 742 |
| PBANKA_0315500 | 18 | 21 | 49 | 5 | 81 | 37 | 71 | 26 | 39 | 16 | 0 | 0 | 451 | 114 | 77 | 559 | 160 | 33 | 872 | 664 | 56 | 395 |
| PBANKA_0315600 | 244 | 284 | 537 | 204 | 416 | 306 | 443 | 330 | 197 | 197 | 1144 | 936 | 2661 | 1290 | 1074 | 2145 | 1220 | 410 | 637 | 253 | 52 | 372 |
| PBANKA_0315700 | 76 | 92 | 15 | 0 | 102 | 52 | 125 | 60 | 33 | 19 | 0 | 0 | 217 | 65 | 104 | 212 | 331 | 181 | 477 | 126 | 48 | 393 |
| PBANKA_0315800 | 18 | 32 | 62 | 16 | 160 | 131 | 534 | 206 | 143 | 83 | 16 | 16 | 467 | 81 | 233 | 366 | 1011 | 412 | 4112 | 1665 | 184 | 985 |
| PBANKA_0315900 | 5 | 11 | 161 | 30 | 296 | 136 | 319 | 116 | 103 | 53 | 12 | 11 | 984 | 117 | 970 | 4458 | 1372 | 271 | 1243 | 661 | 37 | 307 |
| PBANKA_0316000 | 281 | 391 | 184 | 26 | 597 | 292 | 1332 | 459 | 801 | 284 | 98 | 40 | 1853 | 416 | 773 | 2719 | 8329 | 1385 | 6865 | 3755 | 200 | 1338 |
| PBANKA_0316100 | 26 | 31 | 18 | 5 | 138 | 65 | 261 | 108 | 314 | 189 | 799 | 440 | 2775 | 912 | 818 | 2008 | 1569 | 676 | 1015 | 433 | 22 | 169 |
| PBANKA_0316200 | 108 | 109 | 5 | 117 | 191 | 224 | 431 | 494 | 6388 | 5242 | 1E+05 | 1E+05 | 19719 | 54646 | 9210 | 81638 | 14774 | 1120 | 6899 | 20928 | 903 | 7314 |
| PBANKA_0316700 | 0 | 0 | 6 | 0 | 54 | 26 | 89 | 31 | 12 | 14 | 0 | 0 | 552 | 27 | 288 | 1354 | 351 | 63 | 204 | 92 | 15 | 78 |
| PBANKA_0316821 | 0 | 0 | 21 | 10 | 125 | 54 | 94 | 35 | 25 | 18 | 45 | 55 | 100 | 65 | 18 | 86 | 97 | 10 | 32 | 22 | 7 | 44 |
| PBANKA_0316841 | 0 | 0 | 8 | 0 | 103 | 74 | 139 | 46 | 34 | 24 | 233 | 176 | 1450 | 1762 | 185 | 1063 | 289 | 31 | 69 | 60 | 0 | 27 |
| PBANKA_0316981 | 0 | 0 | 14 | 0 | 81 | 34 | 120 | 34 | 375 | 245 | 138 | 53 | 155 | 646 | 171 | 926 | 2254 | 819 | 158 | 616 | 32 | 136 |
| PBANKA_0317061 | 0 | 0 | 10 | 0 | 55 | 43 | 32 | 15 | 19 | 7 | 154 | 121 | 395 | 732 | 52 | 151 | 135 | 34 | 37 | 34 | 0 | 21 |
| PBANKA_0317141 | 10 | 18 | 12 | 0 | 40 | 30 | 56 | 20 | 11 | 18 | 132 | 104 | 320 | 135 | 84 | 246 | 223 | 60 | 460 | 268 | 18 | 145 |
| PBANKA_0400900 | 0 | 0 | 31 | 7 | 72 | 19 | 47 | 18 | 9 | 8 | 0 | 0 | 485 | 155 | 167 | 950 | 243 | 43 | 1188 | 972 | 86 | 760 |
| PBANKA_0401000 | 0 | 0 | 72 | 25 | 194 | 107 | 244 | 98 | 97 | 37 | 186 | 96 | 3636 | 2165 | 718 | 4591 | 2884 | 339 | 1801 | 1592 | 185 | 1543 |
| PBANKA_0401100 | 24 | 45 | 279 | 70 | 831 | 457 | 1006 | 378 | 350 | 147 | 1027 | 631 | 2784 | 4695 | 1959 | 10163 | 5198 | 650 | 3099 | 6643 | 1534 | 11702 |
| PBANKA_0401200 | 0 | 0 | 100 | 12 | 321 | 119 | 380 | 124 | 167 | 60 | 20 | 0 | 382 | 184 | 602 | 2271 | 1930 | 299 | 1074 | 1229 | 125 | 931 |
| PBANKA_0401300 | 0 | 0 | 62 | 9 | 98 | 62 | 85 | 47 | 31 | 29 | 15 | 0 | 806 | 110 | 232 | 896 | 483 | 102 | 939 | 433 | 51 | 301 |
| PBANKA_0401400 | 138 | 92 | 43 | 12 | 121 | 49 | 193 | 86 | 138 | 50 | 38 | 18 | 2139 | 879 | 265 | 2117 | 2022 | 267 | 3033 | 1740 | 247 | 1493 |
| PBANKA_0401500 | 43 | 23 | 274 | 68 | 887 | 432 | 1467 | 565 | 643 | 348 | 70 | 28 | 3021 | 820 | 1915 | 10095 | 12320 | 1436 | 2924 | 3552 | 136 | 1164 |
| PBANKA_0401600 | 13 | 26 | 131 | 17 | 273 | 121 | 370 | 124 | 129 | 52 | 24 | 9 | 2047 | 961 | 919 | 5006 | 2610 | 294 | 907 | 1038 | 115 | 799 |
| PBANKA_0401700 | 7 | 12 | 127 | 9 | 197 | 69 | 284 | 100 | 97 | 43 | 48 | 22 | 2190 | 1651 | 443 | 2754 | 1308 | 134 | 1245 | 1218 | 299 | 1864 |
| PBANKA_0401800 | 28 | 56 | 244 | 71 | 742 | 340 | 1142 | 468 | 597 | 355 | 137 | 53 | 3420 | 2125 | 1794 | 7469 | 6765 | 1193 | 1793 | 2386 | 399 | 2723 |
| PBANKA_0401900 | 55 | 68 | 110 | 26 | 197 | 90 | 186 | 82 | 61 | 30 | 335 | 220 | 1787 | 745 | 314 | 1909 | 1101 | 144 | 1590 | 1124 | 176 | 1394 |
| PBANKA_0402100 | 9 | 20 | 126 | 16 | 453 | 198 | 699 | 240 | 336 | 121 | 49 | 7 | 825 | 567 | 815 | 3317 | 7294 | 879 | 1999 | 3252 | 1170 | 6376 |
| PBANKA_0402200 | 0 | 5 | 33 | 5 | 159 | 84 | 213 | 78 | 52 | 23 | 0 | 0 | 53 | 10 | 61 | 216 | 38 | 12 | 52 | 14 | 31 | 232 |
| PBANKA_0402300 | 8 | 8 | 44 | 17 | 169 | 108 | 278 | 123 | 158 | 85 | 27 | 6 | 413 | 152 | 124 | 302 | 2593 | 350 | 821 | 782 | 188 | 1665 |
| PBANKA_0402400 | 0 | 9 | 18 | 7 | 62 | 58 | 226 | 90 | 121 | 55 | 19 | 14 | 346 | 137 | 29 | 44 | 2052 | 456 | 4847 | 1952 | 29 | 271 |
| PBANKA_0402600 | 5 | 0 | 10 | 0 | 30 | 15 | 25 | 19 | 55 | 32 | 11 | 10 | 56 | 37 | 0 | 7 | 139 | 41 | 855 | 788 | 118 | 968 |
| PBANKA_0402800 | 0 | 6 | 30 | 18 | 165 | 99 | 194 | 114 | 427 | 307 | 464 | 367 | 2653 | 1368 | 277 | 1564 | 1476 | 616 | 1345 | 1011 | 107 | 900 |
| PBANKA_0402900 | 0 | 10 | 129 | 27 | 315 | 142 | 335 | 104 | 153 | 64 | 18 | 20 | 1624 | 571 | 533 | 2470 | 1216 | 183 | 2457 | 1432 | 1463 | 9454 |
| PBANKA_0403000 | 14 | 6 | 785 | 185 | 545 | 372 | 621 | 347 | 221 | 185 | 728 | 670 | 5377 | 2055 | 1161 | 3527 | 1573 | 346 | 1021 | 451 | 76 | 542 |
| PBANKA_0403100 | 11 | 17 | 56 | 13 | 112 | 58 | 134 | 69 | 31 | 27 | 0 | 0 | 277 | 39 | 152 | 464 | 427 | 90 | 841 | 341 | 803 | 4418 |
| PBANKA_0403200 | 21380 | 29519 | 18 | 11 | 328 | 160 | 351 | 125 | 105 | 69 | 31 | 18 | 91 | 20 | 26 | 99 | 92 | 22 | 479 | 347 | 1024 | 10098 |
| PBANKA_0403300 | 235 | 235 | 33 | 5 | 161 | 45 | 145 | 34 | 75 | 23 | 21 | 11 | 340 | 95 | 227 | 989 | 302 | 43 | 2789 | 864 | 403 | 2088 |
| PBANKA_0403400 | 15 | 12 | 26 | 0 | 26 | 22 | 39 | 14 | 10 | 12 | 36 | 17 | 775 | 375 | 150 | 969 | 328 | 26 | 296 | 269 | 101 | 739 |
| PBANKA_0403500 | 7 | 9 | 45 | 17 | 90 | 40 | 100 | 59 | 30 | 15 | 12 | 0 | 372 | 46 | 170 | 649 | 567 | 100 | 82 | 37 | 130 | 944 |
| PBANKA_0403600 | 78 | 139 | 186 | 35 | 730 | 285 | 1680 | 629 | 1007 | 396 | 279 | 129 | 1945 | 3130 | 2336 | 12353 | 11486 | 1242 | 3823 | 14192 | 5515 | 37279 |
| PBANKA_0403700 | 6 | 17 | 135 | 58 | 315 | 131 | 354 | 136 | 474 | 214 | 542 | 401 | 1671 | 2186 | 376 | 2256 | 6001 | 1419 | 3552 | 5917 | 1899 | 12118 |
| PBANKA_0403800 | 121 | 229 | 141 | 43 | 332 | 181 | 352 | 155 | 247 | 167 | 204 | 137 | 6997 | 2960 | 2459 | 13830 | 4051 | 705 | 2515 | 2127 | 515 | 3684 |
| PBANKA_0403900 | 109 | 92 | 69 | 30 | 224 | 151 | 276 | 109 | 103 | 87 | 26 | 12 | 350 | 141 | 172 | 830 | 495 | 104 | 629 | 291 | 101 | 414 |
| PBANKA_0404000 | 140 | 159 | 295 | 59 | 1012 | 304 | 1353 | 416 | 1065 | 430 | 3013 | 1371 | 3147 | 3934 | 2263 | 11746 | 6560 | 808 | 2393 | 3867 | 286 | 1909 |
| PBANKA_0404100 | 0 | 0 | 30 | 0 | 140 | 54 | 368 | 172 | 211 | 100 | 34 | 14 | 875 | 206 | 255 | 743 | 4408 | 669 | 2153 | 1186 | 106 | 966 |
| PBANKA_0404200 | 0 | 0 | 121 | 43 | 232 | 209 | 471 | 290 | 166 | 89 | 22 | 14 | 953 | 330 | 565 | 1944 | 1044 | 315 | 796 | 382 | 45 | 281 |
| PBANKA_0404400 | 0 | 0 | 100 | 54 | 230 | 173 | 465 | 247 | 179 | 192 | 25 | 17 | 281 | 75 | 326 | 546 | 1360 | 492 | 852 | 304 | 272 | 2558 |
| PBANKA_0404500 | 253 | 313 | 101 | 20 | 220 | 97 | 262 | 145 | 173 | 60 | 301 | 208 | 1921 | 915 | 416 | 1721 | 1757 | 338 | 562 | 339 | 152 | 1146 |
| PBANKA_0404600 | 269 | 361 | 350 | 83 | 503 | 280 | 381 | 157 | 126 | 91 | 93 | 51 | 1917 | 407 | 775 | 3107 | 504 | 83 | 1193 | 563 | 355 | 2326 |
| PBANKA_0404700 | 173 | 169 | 237 | 57 | 330 | 282 | 401 | 328 | 215 | 118 | 25 | 17 | 265 | 36 | 542 | 451 | 522 | 283 | 715 | 335 | 142 | 698 |
| PBANKA_0404800 | 271 | 319 | 246 | 68 | 1434 | 720 | 3544 | 1514 | 1746 | 879 | 170 | 111 | 3287 | 741 | 1261 | 4424 | 17774 | 2793 | 3187 | 2750 | 1126 | 8777 |
| PBANKA_0404900 | 0 | 0 | 81 | 19 | 219 | 139 | 381 | 215 | 126 | 73 | 26 | 12 | 594 | 99 | 377 | 1399 | 1443 | 242 | 1607 | 848 | 502 | 3984 |
| PBANKA_0405000 | 0 | 0 | 28 | 9 | 105 | 40 | 148 | 69 | 62 | 36 | 0 | 5 | 81 | 12 | 82 | 213 | 258 | 78 | 173 | 63 | 21 | 114 |
| PBANKA_0405100 | 0 | 0 | 29 | 18 | 80 | 78 | 111 | 88 | 59 | 42 | 9 | 0 | 330 | 139 | 120 | 374 | 445 | 113 | 537 | 236 | 87 | 484 |
| PBANKA_0405200 | 0 | 0 | 943 | 202 | 820 | 565 | 949 | 354 | 297 | 108 | 49 | 12 | 6405 | 1729 | 3869 | 20025 | 4388 | 477 | 1914 | 1454 | 482 | 4435 |
| PBANKA_0405300 | 0 | 0 | 154 | 79 | 135 | 126 | 116 | 102 | 52 | 50 | 434 | 446 | 8953 | 1679 | 2621 | 5686 | 1497 | 461 | 1986 | 588 | 154 | 1549 |
| PBANKA_0405400 | 7 | 13 | 1564 | 474 | 934 | 704 | 1160 | 673 | 347 | 269 | 1533 | 1378 | 9466 | 3473 | 2616 | 9819 | 2458 | 397 | 2699 | 1374 | 79 | 771 |
| PBANKA_0405500 | 6 | 10 | 1812 | 395 | 1023 | 558 | 962 | 403 | 292 | 148 | 2301 | 1724 | 28364 | 12607 | 3972 | 21944 | 7095 | 745 | 3153 | 2346 | 553 | 4268 |
| PBANKA_0405600 | 0 | 0 | 9 | 0 | 53 | 5 | 42 | 6 | 12 | 0 | 12 | 5 | 324 | 145 | 33 | 160 | 136 | 40 | 2091 | 1017 | 396 | 2474 |
| PBANKA_0405700 | 16 | 21 | 55 | 0 | 46 | 28 | 49 | 25 | 29 | 16 | 10 | 0 | 557 | 121 | 139 | 609 | 399 | 112 | 2124 | 777 | 165 | 921 |
| PBANKA_0405800 | 0 | 0 | 50 | 16 | 147 | 52 | 223 | 113 | 137 | 74 | 15 | 0 | 431 | 83 | 138 | 353 | 1443 | 310 | 970 | 523 | 329 | 2084 |
| PBANKA_0405900 | 0 | 5 | 115 | 25 | 237 | 90 | 369 | 107 | 143 | 46 | 11 | 6 | 352 | 112 | 228 | 684 | 1330 | 271 | 759 | 484 | 384 | 2423 |
| PBANKA_0406000 | 29 | 56 | 499 | 128 | 1462 | 831 | 1867 | 728 | 968 | 542 | 169 | 66 | 2253 | 875 | 859 | 3954 | 14190 | 1872 | 7756 | 6165 | 149 | 1233 |
| PBANKA_0406100 | 14 | 24 | 219 | 52 | 305 | 154 | 299 | 114 | 119 | 60 | 24 | 11 | 1029 | 381 | 603 | 3216 | 2926 | 428 | 784 | 558 | 182 | 783 |
| PBANKA_0406200 | 51 | 62 | 45 | 10 | 116 | 49 | 96 | 48 | 133 | 88 | 20 | 9 | 255 | 302 | 173 | 929 | 1009 | 211 | 813 | 4197 | 341 | 1408 |
| PBANKA_0406300 | 17 | 34 | 38 | 6 | 124 | 66 | 106 | 48 | 35 | 22 | 7 | 0 | 258 | 74 | 390 | 1707 | 787 | 118 | 1117 | 496 | 14 | 148 |
| PBANKA_0406500 | 0 | 0 | 629 | 158 | 825 | 555 | 876 | 396 | 241 | 147 | 28 | 16 | 2520 | 809 | 2155 | 8176 | 1663 | 254 | 1404 | 856 | 65 | 438 |
| PBANKA_0406600 | 0 | 11 | 11 | 0 | 53 | 42 | 70 | 32 | 411 | 241 | 164 | 82 | 286 | 99 | 85 | 365 | 930 | 393 | 268 | 224 | 41 | 238 |

| | | | | | | | | | | | | | | | | | | | | | | |
|---|---|---|---|---|---|---|---|---|---|---|---|---|---|---|---|---|---|---|---|---|---|---|
| PBANKA_0406700 | 0 | 0 | 114 | 25 | 98 | 65 | 62 | 41 | 16 | 8 | 0 | 5 | 800 | 159 | 288 | 1069 | 285 | 51 | 308 | 118 | 10 | 70 |
| PBANKA_0406800 | 0 | 0 | 158 | 28 | 211 | 141 | 294 | 130 | 81 | 35 | 28 | 25 | 1394 | 721 | 503 | 1787 | 1040 | 182 | 505 | 331 | 48 | 409 |
| PBANKA_0406900 | 115 | 179 | 12 | 0 | 57 | 26 | 71 | 21 | 318 | 236 | 144 | 59 | 156 | 49 | 16 | 27 | 555 | 206 | 660 | 478 | 2762 | 18070 |
| PBANKA_0407000 | 9 | 12 | 48 | 19 | 99 | 49 | 83 | 33 | 29 | 15 | 9 | 0 | 65 | 14 | 186 | 420 | 103 | 40 | 306 | 118 | 83 | 536 |
| PBANKA_0407100 | 7 | 10 | 151 | 47 | 454 | 234 | 596 | 216 | 239 | 111 | 112 | 58 | 2215 | 1154 | 1057 | 5125 | 2583 | 357 | 1423 | 1237 | 99 | 657 |
| PBANKA_0407200 | 78 | 132 | 34 | 8 | 145 | 50 | 121 | 53 | 44 | 31 | 22 | 0 | 398 | 96 | 152 | 700 | 291 | 69 | 2586 | 1800 | 81 | 497 |
| PBANKA_0407400 | 30 | 53 | 28 | 12 | 468 | 158 | 1372 | 470 | 628 | 231 | 110 | 28 | 652 | 639 | 124 | 392 | 6354 | 749 | 1882 | 5148 | 3471 | 23564 |
| PBANKA_0407500 | 0 | 0 | 104 | 23 | 161 | 64 | 155 | 78 | 80 | 35 | 26 | 9 | 1113 | 698 | 246 | 914 | 257 | 46 | 820 | 499 | 143 | 839 |
| PBANKA_0407600 | 0 | 0 | 421 | 122 | 564 | 345 | 610 | 221 | 205 | 124 | 631 | 421 | 3808 | 1429 | 1518 | 5457 | 1329 | 230 | 2078 | 1259 | 776 | 6697 |
| PBANKA_0407700 | 39 | 53 | 1445 | 378 | 1235 | 577 | 1240 | 512 | 473 | 264 | 2164 | 1281 | 12887 | 3831 | 2706 | 10886 | 4439 | 831 | 3292 | 1546 | 113 | 1082 |
| PBANKA_0407800 | 0 | 0 | 33 | 7 | 79 | 34 | 84 | 49 | 63 | 14 | 14 | 17 | 1378 | 589 | 253 | 1474 | 856 | 136 | 1154 | 1009 | 163 | 1065 |
| PBANKA_0407900 | 0 | 0 | 22 | 10 | 89 | 39 | 142 | 63 | 69 | 48 | 9 | 13 | 648 | 139 | 180 | 428 | 580 | 159 | 747 | 259 | 87 | 579 |
| PBANKA_0408000 | 0 | 0 | 111 | 21 | 257 | 129 | 226 | 83 | 113 | 66 | 22 | 0 | 219 | 92 | 497 | 2872 | 776 | 86 | 884 | 640 | 173 | 1012 |
| PBANKA_0408100 | 0 | 0 | 96 | 12 | 204 | 91 | 155 | 50 | 78 | 34 | 15 | 0 | 71 | 19 | 157 | 545 | 202 | 47 | 512 | 437 | 201 | 1172 |
| PBANKA_0408300 | 246 | 297 | 268 | 47 | 658 | 166 | 546 | 175 | 719 | 355 | 1074 | 537 | 2463 | 6922 | 2369 | 12023 | 4515 | 962 | 6121 | 13691 | 1871 | 11547 |
| PBANKA_0408400 | 90 | 79 | 133 | 30 | 603 | 260 | 1009 | 302 | 386 | 142 | 358 | 255 | 2763 | 1917 | 875 | 4722 | 1751 | 328 | 5942 | 6114 | 312 | 2536 |
| PBANKA_0408500 | 11 | 0 | 13 | 0 | 194 | 74 | 379 | 166 | 237 | 83 | 21 | 16 | 766 | 316 | 263 | 894 | 5382 | 768 | 375 | 284 | 24 | 170 |
| PBANKA_0408600 | 62 | 55 | 233 | 73 | 414 | 222 | 453 | 187 | 111 | 74 | 83 | 25 | 1801 | 567 | 593 | 2108 | 684 | 140 | 776 | 281 | 204 | 1124 |
| PBANKA_0408700 | 27 | 40 | 52 | 14 | 81 | 34 | 111 | 46 | 108 | 52 | 22 | 19 | 571 | 189 | 169 | 631 | 566 | 168 | 712 | 315 | 124 | 715 |
| PBANKA_0408800 | 36 | 41 | 58 | 14 | 81 | 40 | 159 | 50 | 117 | 48 | 25 | 12 | 811 | 323 | 286 | 1062 | 852 | 213 | 814 | 361 | 98 | 728 |
| PBANKA_0408900 | 170 | 271 | 36 | 10 | 149 | 50 | 218 | 95 | 146 | 75 | 17 | 9 | 1372 | 412 | 286 | 1397 | 3459 | 593 | 1456 | 1223 | 1400 | 9647 |
| PBANKA_0409000 | 14 | 5 | 21 | 0 | 39 | 17 | 50 | 12 | 13 | 9 | 0 | 5 | 461 | 281 | 313 | 1317 | 462 | 62 | 156 | 122 | 843 | 5964 |
| PBANKA_0409100 | 0 | 0 | 152 | 42 | 261 | 132 | 217 | 102 | 137 | 40 | 32 | 14 | 2508 | 1768 | 1306 | 7486 | 2333 | 189 | 413 | 596 | 12 | 74 |
| PBANKA_0409200 | 0 | 0 | 77 | 16 | 243 | 64 | 268 | 83 | 99 | 38 | 24 | 8 | 1289 | 534 | 591 | 2499 | 1604 | 306 | 1060 | 654 | 43 | 265 |
| PBANKA_0409300 | 8 | 12 | 254 | 100 | 715 | 367 | 702 | 289 | 262 | 159 | 71 | 25 | 4178 | 1655 | 1514 | 10062 | 2538 | 360 | 5161 | 4383 | 227 | 1853 |
| PBANKA_0409400 | 0 | 0 | 28 | 15 | 218 | 117 | 207 | 86 | 170 | 102 | 60 | 28 | 708 | 1473 | 4213 | 23497 | 6381 | 871 | 3334 | 23825 | 3589 | 29185 |
| PBANKA_0409500 | 0 | 0 | 14 | 0 | 23 | 15 | 25 | 0 | 11 | 6 | 89 | 65 | 655 | 148 | 318 | 899 | 151 | 52 | 975 | 454 | 11 | 58 |
| PBANKA_0409600 | 11 | 10 | 213 | 52 | 428 | 212 | 377 | 108 | 110 | 50 | 24 | 0 | 172 | 46 | 410 | 1691 | 387 | 67 | 1375 | 621 | 207 | 1294 |
| PBANKA_0409800 | 21 | 35 | 321 | 68 | 632 | 344 | 978 | 464 | 443 | 218 | 39 | 49 | 967 | 336 | 988 | 3563 | 6461 | 945 | 1934 | 2112 | 231 | 2045 |
| PBANKA_0409900 | 0 | 0 | 24 | 9 | 38 | 35 | 61 | 32 | 26 | 14 | 0 | 0 | 295 | 35 | 103 | 359 | 200 | 41 | 2335 | 542 | 110 | 638 |
| PBANKA_0410000 | 0 | 0 | 46 | 28 | 133 | 100 | 155 | 83 | 51 | 24 | 7 | 0 | 138 | 28 | 225 | 704 | 219 | 59 | 345 | 110 | 42 | 212 |
| PBANKA_0410100 | 0 | 0 | 31 | 8 | 141 | 53 | 170 | 52 | 81 | 29 | 5 | 0 | 658 | 102 | 275 | 1376 | 777 | 157 | 1368 | 894 | 51 | 320 |
| PBANKA_0410200 | 0 | 0 | 13 | 0 | 77 | 20 | 104 | 29 | 45 | 12 | 9 | 0 | 379 | 114 | 167 | 1021 | 1141 | 156 | 2195 | 2011 | 22 | 169 |
| PBANKA_0410300 | 0 | 0 | 142 | 19 | 391 | 157 | 618 | 193 | 175 | 92 | 18 | 8 | 1572 | 429 | 776 | 3560 | 3104 | 435 | 2042 | 959 | 261 | 1297 |
| PBANKA_0410400 | 0 | 0 | 75 | 15 | 186 | 107 | 340 | 128 | 153 | 74 | 30 | 6 | 730 | 198 | 54 | 246 | 2885 | 524 | 2485 | 1842 | 545 | 4881 |
| PBANKA_0410500 | 0 | 7 | 133 | 33 | 302 | 144 | 467 | 143 | 721 | 430 | 177 | 62 | 3307 | 1558 | 922 | 3356 | 4932 | 1485 | 1437 | 1003 | 741 | 5368 |
| PBANKA_0410600 | 75 | 102 | 709 | 229 | 637 | 286 | 783 | 364 | 235 | 170 | 1107 | 693 | 10722 | 4904 | 1444 | 7592 | 2376 | 340 | 1336 | 673 | 164 | 874 |
| PBANKA_0410700 | 0 | 6 | 35 | 9 | 123 | 63 | 148 | 76 | 45 | 26 | 10 | 0 | 248 | 75 | 163 | 607 | 461 | 82 | 441 | 270 | 34 | 210 |
| PBANKA_0410800 | 0 | 0 | 269 | 49 | 432 | 160 | 417 | 161 | 127 | 48 | 23 | 0 | 1915 | 742 | 806 | 3270 | 1559 | 214 | 302 | 273 | 28 | 222 |
| PBANKA_0410900 | 0 | 0 | 39 | 14 | 89 | 47 | 85 | 34 | 27 | 16 | 5 | 0 | 287 | 67 | 143 | 692 | 211 | 44 | 293 | 160 | 28 | 184 |
| PBANKA_0411000 | 0 | 0 | 56 | 13 | 186 | 60 | 156 | 48 | 65 | 19 | 12 | 0 | 54 | 7 | 73 | 386 | 136 | 9 | 74 | 58 | 9 | 26 |
| PBANKA_0411100 | 0 | 8 | 130 | 27 | 265 | 120 | 208 | 108 | 128 | 54 | 44 | 27 | 1512 | 717 | 479 | 2346 | 936 | 152 | 703 | 467 | 31 | 191 |
| PBANKA_0411200 | 0 | 12 | 111 | 18 | 213 | 80 | 190 | 61 | 104 | 32 | 30 | 13 | 511 | 295 | 257 | 1292 | 462 | 92 | 643 | 397 | 13 | 80 |
| PBANKA_0411300 | 117 | 236 | 26 | 10 | 100 | 53 | 127 | 66 | 40 | 29 | 10 | 0 | 575 | 151 | 119 | 462 | 451 | 127 | 4019 | 2427 | 247 | 2146 |
| PBANKA_0411400 | 0 | 6 | 50 | 14 | 151 | 96 | 156 | 76 | 48 | 22 | 7 | 0 | 354 | 41 | 335 | 1027 | 619 | 124 | 630 | 238 | 29 | 183 |
| PBANKA_0411500 | 7 | 0 | 127 | 30 | 245 | 108 | 261 | 109 | 90 | 72 | 10 | 0 | 602 | 142 | 426 | 1840 | 1968 | 280 | 1096 | 880 | 62 | 482 |
| PBANKA_0411600 | 98 | 143 | 145 | 22 | 385 | 101 | 375 | 105 | 216 | 62 | 51 | 11 | 650 | 214 | 509 | 2380 | 2841 | 522 | 1944 | 1524 | 127 | 759 |
| PBANKA_0411700 | 15 | 10 | 83 | 23 | 297 | 130 | 291 | 146 | 126 | 67 | 22 | 15 | 1412 | 403 | 805 | 2254 | 1810 | 454 | 1649 | 763 | 257 | 1842 |
| PBANKA_0411800 | 0 | 0 | 25 | 0 | 113 | 78 | 142 | 73 | 43 | 30 | 33 | 28 | 250 | 83 | 151 | 446 | 339 | 67 | 191 | 101 | 20 | 118 |
| PBANKA_0411900 | 0 | 0 | 23 | 8 | 101 | 36 | 125 | 36 | 50 | 21 | 5 | 0 | 234 | 45 | 225 | 1149 | 747 | 134 | 485 | 260 | 7 | 41 |
| PBANKA_0411950 | 0 | 13 | 94 | 19 | 252 | 86 | 286 | 98 | 104 | 60 | 20 | 0 | 698 | 269 | 469 | 1839 | 2090 | 338 | 1795 | 1182 | 73 | 645 |
| PBANKA_0412000 | 7 | 0 | 43 | 5 | 97 | 40 | 129 | 44 | 53 | 25 | 19 | 0 | 113 | 60 | 112 | 504 | 637 | 73 | 78 | 63 | 7 | 54 |
| PBANKA_0412100 | 32 | 17 | 50 | 8 | 90 | 60 | 134 | 53 | 54 | 34 | 86 | 72 | 1176 | 359 | 241 | 1355 | 1148 | 160 | 1768 | 860 | 55 | 466 |
| PBANKA_0412200 | 203 | 281 | 100 | 35 | 135 | 52 | 91 | 53 | 58 | 43 | 537 | 317 | 1440 | 554 | 146 | 711 | 503 | 122 | 838 | 343 | 52 | 335 |
| PBANKA_0412300 | 42 | 41 | 19 | 5 | 58 | 21 | 85 | 29 | 56 | 44 | 109 | 43 | 1023 | 544 | 172 | 876 | 1553 | 226 | 1126 | 976 | 136 | 900 |
| PBANKA_0412400 | 834 | 1257 | 168 | 49 | 469 | 186 | 422 | 113 | 134 | 44 | 41 | 17 | 31 | 60 | 20 | 62 | 57 | 11 | 65 | 64 | 107 | 902 |
| PBANKA_0412500 | 542 | 609 | 21 | 10 | 171 | 80 | 154 | 53 | 104 | 43 | 14 | 0 | 288 | 98 | 70 | 264 | 2190 | 388 | 958 | 702 | 65 | 328 |
| PBANKA_0412600 | 171 | 162 | 21 | 7 | 164 | 60 | 230 | 61 | 104 | 43 | 13 | 0 | 149 | 20 | 81 | 282 | 490 | 108 | 835 | 255 | 48 | 254 |
| PBANKA_0412700 | 13 | 8 | 29 | 23 | 81 | 33 | 96 | 27 | 19 | 7 | 0 | 0 | 72 | 9 | 114 | 320 | 153 | 30 | 537 | 229 | 30 | 194 |
| PBANKA_0412800 | 8 | 8 | 172 | 57 | 329 | 266 | 282 | 164 | 97 | 48 | 376 | 387 | 1040 | 449 | 470 | 1701 | 572 | 144 | 1041 | 554 | 83 | 887 |
| PBANKA_0412900 | 36 | 48 | 5 | 0 | 16 | 11 | 30 | 21 | 33 | 29 | 38 | 24 | 409 | 316 | 54 | 212 | 212 | 74 | 3167 | 4431 | 1E+05 | 1E+06 |
| PBANKA_0413000 | 37 | 39 | 9 | 0 | 216 | 65 | 597 | 208 | 390 | 162 | 39 | 27 | 237 | 124 | 66 | 206 | 410 | 122 | 1244 | 1279 | 199 | 1238 |
| PBANKA_0413200 | 60 | 87 | 164 | 31 | 246 | 124 | 368 | 162 | 151 | 71 | 22 | 11 | 1690 | 502 | 336 | 1307 | 2179 | 353 | 498 | 255 | 69 | 437 |
| PBANKA_0413300 | 0 | 0 | 59 | 14 | 120 | 44 | 111 | 50 | 52 | 17 | 11 | 0 | 136 | 48 | 398 | 1244 | 345 | 67 | 856 | 440 | 239 | 1640 |
| PBANKA_0413500 | 126 | 147 | 154 | 23 | 390 | 183 | 563 | 234 | 226 | 92 | 87 | 34 | 760 | 365 | 308 | 906 | 1411 | 408 | 10180 | 6644 | 527 | 3274 |
| PBANKA_0413600 | 53 | 51 | 83 | 15 | 152 | 54 | 120 | 48 | 61 | 19 | 113 | 54 | 763 | 490 | 265 | 934 | 368 | 63 | 187 | 97 | 24 | 171 |
| PBANKA_0413700 | 155 | 190 | 18 | 6 | 47 | 27 | 35 | 21 | 31 | 21 | 8 | 0 | 206 | 44 | 41 | 165 | 127 | 39 | 312 | 116 | 14 | 106 |
| PBANKA_0413800 | 0 | 0 | 49 | 11 | 109 | 55 | 97 | 54 | 70 | 27 | 8 | 5 | 64 | 28 | 73 | 343 | 83 | 20 | 404 | 278 | 129 | 901 |
| PBANKA_0413900 | 0 | 0 | 60 | 8 | 137 | 63 | 167 | 59 | 80 | 37 | 0 | 6 | 221 | 57 | 126 | 786 | 310 | 45 | 219 | 153 | 82 | 580 |
| PBANKA_0414000 | 51 | 55 | 98 | 37 | 355 | 165 | 404 | 151 | 195 | 101 | 21 | 18 | 742 | 332 | 836 | 3628 | 2258 | 377 | 1601 | 1744 | 138 | 874 |
| PBANKA_0414100 | 253 | 356 | 617 | 165 | 838 | 451 | 933 | 453 | 340 | 135 | 53 | 23 | 1651 | 440 | 812 | 3071 | 2831 | 501 | 4694 | 1985 | 706 | 4815 |
| PBANKA_0414200 | 0 | 9 | 186 | 46 | 262 | 125 | 299 | 93 | 137 | 55 | 26 | 7 | 224 | 87 | 1140 | 5437 | 1636 | 293 | 315 | 570 | 1035 | 6589 |
| PBANKA_0414300 | 50 | 38 | 53 | 16 | 171 | 67 | 269 | 116 | 156 | 102 | 25 | 13 | 573 | 157 | 123 | 418 | 3425 | 618 | 1549 | 1207 | 2904 | 18818 |
| PBANKA_0414400 | 648 | 786 | 191 | 58 | 423 | 300 | 635 | 266 | 267 | 136 | 21 | 32 | 3062 | 735 | 1064 | 2954 | 4325 | 857 | 559 | 308 | 239 | 1902 |
| PBANKA_0414500 | 7 | 6 | 36 | 11 | 131 | 84 | 168 | 78 | 77 | 23 | 27 | 8 | 534 | 115 | 366 | 1169 | 1466 | 279 | 595 | 331 | 68 | 633 |
| PBANKA_0414600 | 32 | 52 | 916 | 246 | 742 | 372 | 845 | 470 | 336 | 214 | 1247 | 915 | 4542 | 1926 | 1798 | 3280 | 1352 | 443 | 1026 | 496 | 170 | 1419 |
| PBANKA_0414700 | 81 | 124 | 168 | 34 | 253 | 72 | 222 | 92 | 406 | 211 | 3636 | 2140 | 3733 | 3691 | 515 | 4845 | 1784 | 293 | 1824 | 2058 | 1413 | 9728 |
| PBANKA_0414800 | 301 | 394 | 250 | 87 | 347 | 228 | 293 | 100 | 128 | 36 | 60 | 43 | 1071 | 354 | 410 | 1314 | 227 | 56 | 410 | 278 | 328 | 2010 |
| PBANKA_0414900 | 350 | 474 | 7 | 0 | 75 | 20 | 71 | 28 | 48 | 27 | 19 | 7 | 385 | 233 | 90 | 434 | 383 | 125 | 1064 | 1658 | 618 | 3558 |
| PBANKA_0415000 | 8 | 10 | 92 | 25 | 295 | 152 | 400 | 189 | 138 | 70 | 28 | 10 | 443 | 92 | 215 | 834 | 1172 | 253 | 685 | 418 | 409 | 3514 |
| PBANKA_0415100 | 6 | 7 | 22 | 5 | 124 | 32 | 223 | 74 | 77 | 29 | 22 | 6 | 446 | 58 | 87 | 466 | 1572 | 262 | 1542 | 411 | 263 | 1955 |
| PBANKA_0415200 | 53 | 110 | 23 | 7 | 151 | 93 | 231 | 109 | 105 | 70 | 22 | 7 | 342 | 54 | 76 | 329 | 1092 | 227 | 646 | 390 | 25 | 198 |
| PBANKA_0415300 | 128 | 206 | 170 | 24 | 419 | 141 | 472 | 171 | 232 | 97 | 63 | 21 | 1350 | 974 | 668 | 4215 | 3609 | 631 | 938 | 2322 | 342 | 2319 |
| PBANKA_0415400 | 8 | 41 | 96 | 15 | 266 | 84 | 364 | 168 | 327 | 119 | 244 | 105 | 1885 | 966 | 561 | 2368 | 4759 | 782 | 1989 | 1434 | 2182 | 12227 |
| PBANKA_0415500 | 11 | 10 | 118 | 43 | 528 | 256 | 790 | 335 | 251 | 98 | 30 | 28 | 1082 | 489 | 1324 | 6481 | 3909 | 496 | 947 | 1241 | 174 | 1184 |
| PBANKA_0415600 | 71 | 89 | 66 | 15 | 217 | 88 | 240 | 104 | 185 | 78 | 770 | 428 | 1950 | 2250 | 653 | 3684 | 2053 | 372 | 1656 | 2133 | 118 | 780 |
| PBANKA_0415700 | 198 | 255 | 87 | 30 | 224 | 154 | 363 | 184 | 199 | 124 | 48 | 43 | 535 | 215 | 201 | 779 | 1161 | 283 | 2472 | 1314 | 322 | 1739 |
| PBANKA_0415900 | 0 | 0 | 10 | 0 | 23 | 16 | 199 | 85 | 235 | 134 | 23 | 14 | 277 | 73 | 39 | 199 | 2018 | 485 | 385 | 206 | 69 | 588 |
| PBANKA_0416000 | 0 | 0 | 25 | 12 | 1314 | 924 | 6206 | 3405 | 3288 | 2148 | 248 | 135 | 8282 | 1995 | 5984 | 17679 | 58476 | 9918 | 1187 | 978 | 68 | 622 |
| PBANKA_0416100 | 28 | 19 | 19 | 0 | 131 | 45 | 303 | 85 | 385 | 150 | 140 | 38 | 850 | 1206 | 379 | 1639 | 3971 | 702 | 3462 | 21894 | 241 | 1734 |

| | | | | | | | | | | | | | | | | | | | | | | |
|---|---|---|---|---|---|---|---|---|---|---|---|---|---|---|---|---|---|---|---|---|---|---|
| PBANKA_0416200 | 134 | 150 | 143 | 47 | 429 | 148 | 377 | 147 | 300 | 130 | 982 | 607 | 2320 | 2089 | 874 | 4422 | 1395 | 263 | 1627 | 1511 | 560 | 3525 |
| PBANKA_0416300 | 620 | 705 | 351 | 119 | 1213 | 750 | 1835 | 808 | 870 | 461 | 290 | 127 | 2339 | 1637 | 618 | 3873 | 5997 | 644 | 1520 | 2493 | 798 | 5841 |
| PBANKA_0416400 | 15 | 22 | 78 | 10 | 171 | 85 | 213 | 84 | 76 | 33 | 8 | 0 | 182 | 62 | 182 | 573 | 674 | 95 | 768 | 348 | 118 | 871 |
| PBANKA_0416500 | 32 | 42 | 384 | 112 | 1131 | 516 | 1222 | 440 | 385 | 150 | 49 | 28 | 1039 | 533 | 3487 | 16122 | 7710 | 806 | 1817 | 2540 | 430 | 3279 |
| PBANKA_0416600 | 91 | 168 | 201 | 50 | 403 | 214 | 700 | 318 | 242 | 124 | 23 | 20 | 772 | 228 | 698 | 2275 | 3450 | 511 | 3867 | 2436 | 34 | 377 |
| PBANKA_0416700 | 0 | 0 | 92 | 7 | 251 | 144 | 345 | 185 | 106 | 86 | 15 | 10 | 956 | 163 | 274 | 942 | 4147 | 545 | 503 | 161 | 9 | 90 |
| PBANKA_0416800 | 19 | 27 | 243 | 71 | 564 | 310 | 748 | 278 | 331 | 142 | 87 | 26 | 3331 | 2360 | 1203 | 6572 | 5707 | 635 | 1169 | 1411 | 154 | 1158 |
| PBANKA_0416900 | 0 | 0 | 35 | 10 | 52 | 30 | 46 | 32 | 19 | 12 | 0 | 0 | 451 | 44 | 232 | 511 | 175 | 42 | 190 | 86 | 0 | 49 |
| PBANKA_0417000 | 0 | 0 | 58 | 6 | 60 | 29 | 77 | 37 | 13 | 7 | 0 | 0 | 717 | 145 | 241 | 1310 | 407 | 78 | 1197 | 741 | 15 | 128 |
| PBANKA_0417300 | 0 | 0 | 160 | 27 | 268 | 136 | 269 | 115 | 121 | 60 | 25 | 10 | 4040 | 1487 | 1132 | 6512 | 2289 | 276 | 2072 | 1736 | 120 | 875 |
| PBANKA_0417400 | 29 | 61 | 152 | 32 | 169 | 104 | 244 | 155 | 79 | 46 | 9 | 0 | 721 | 205 | 256 | 970 | 3230 | 507 | 3065 | 1989 | 233 | 1991 |
| PBANKA_0417500 | 78 | 113 | 1126 | 377 | 1133 | 663 | 937 | 520 | 282 | 247 | 2458 | 1999 | 8218 | 3310 | 1719 | 6767 | 2549 | 446 | 1503 | 998 | 155 | 1155 |
| PBANKA_0417700 | 111 | 137 | 696 | 300 | 1569 | 1428 | 2413 | 1227 | 1045 | 771 | 214 | 88 | 1847 | 492 | 1153 | 2801 | 10888 | 1700 | 7489 | 4451 | 403 | 5260 |
| PBANKA_0417800 | 249 | 436 | 24 | 9 | 716 | 253 | 2200 | 1062 | 1998 | 1006 | 340 | 176 | 1642 | 1493 | 51 | 213 | 32877 | 4946 | 1315 | 4461 | 251 | 1189 |
| PBANKA_0417900 | 137 | 228 | 176 | 26 | 553 | 310 | 959 | 399 | 326 | 149 | 31 | 40 | 1527 | 1206 | 588 | 2804 | 3488 | 529 | 696 | 730 | 274 | 2048 |
| PBANKA_0418000 | 8 | 16 | 732 | 141 | 1254 | 418 | 1202 | 350 | 364 | 147 | 1349 | 803 | 5840 | 10373 | 4420 | 25956 | 6277 | 535 | 640 | 3266 | 194 | 1576 |
| PBANKA_0418100 | 21 | 39 | 67 | 20 | 685 | 291 | 1628 | 618 | 943 | 434 | 162 | 30 | 817 | 760 | 957 | 3061 | 14523 | 2280 | 696 | 2963 | 66 | 452 |
| PBANKA_0418200 | 0 | 0 | 148 | 37 | 402 | 349 | 527 | 481 | 251 | 159 | 21 | 15 | 905 | 210 | 674 | 1530 | 1910 | 493 | 856 | 376 | 50 | 341 |
| PBANKA_0418300 | 20 | 27 | 49 | 18 | 126 | 57 | 110 | 67 | 82 | 44 | 35 | 30 | 2215 | 706 | 351 | 1548 | 2126 | 589 | 3345 | 1251 | 147 | 1317 |
| PBANKA_0418400 | 0 | 0 | 21 | 0 | 57 | 23 | 52 | 16 | 20 | 13 | 13 | 6 | 139 | 24 | 93 | 297 | 237 | 60 | 196 | 93 | 0 | 5 |
| PBANKA_0501051 | 0 | 0 | 6 | 0 | 32 | 21 | 35 | 31 | 24 | 0 | 90 | 87 | 639 | 711 | 87 | 196 | 263 | 54 | 53 | 43 | 0 | 7 |
| PBANKA_0501200 | 1E+05 | 2E+05 | 2058 | 813 | 1941 | 1336 | 1112 | 514 | 367 | 254 | 72 | 58 | 73 | 16 | 19 | 80 | 400 | 43 | 241 | 219 | 7 | 55 |
| PBANKA_0501300 | 427 | 433 | 190 | 59 | 897 | 491 | 911 | 349 | 299 | 140 | 54 | 16 | 826 | 194 | 1112 | 3525 | 3897 | 610 | 4400 | 1925 | 36 | 200 |
| PBANKA_0501500 | 0 | 5 | 101 | 19 | 266 | 174 | 373 | 129 | 217 | 98 | 32 | 15 | 339 | 75 | 549 | 1861 | 1465 | 356 | 540 | 298 | 20 | 99 |
| PBANKA_0501600 | 469 | 694 | 126 | 19 | 324 | 174 | 422 | 144 | 509 | 235 | 88 | 47 | 112 | 82 | 18 | 48 | 1656 | 306 | 323 | 469 | 563 | 3909 |
| PBANKA_0501700 | 21 | 27 | 7 | 0 | 90 | 67 | 282 | 182 | 154 | 131 | 46 | 19 | 112 | 31 | 12 | 61 | 364 | 71 | 1465 | 779 | 97 | 690 |
| PBANKA_0501800 | 146 | 160 | 35 | 11 | 203 | 127 | 250 | 129 | 637 | 570 | 222 | 139 | 248 | 116 | 17 | 46 | 507 | 454 | 438 | 243 | 628 | 4598 |
| PBANKA_0501900 | 22 | 28 | 61 | 11 | 181 | 82 | 245 | 80 | 80 | 44 | 21 | 0 | 1107 | 658 | 526 | 3159 | 2038 | 264 | 410 | 401 | 105 | 757 |
| PBANKA_0502000 | 19 | 28 | 41 | 6 | 69 | 25 | 73 | 47 | 66 | 25 | 7 | 6 | 899 | 562 | 210 | 673 | 415 | 175 | 14763 | 18078 | 44 | 314 |
| PBANKA_0502100 | 0 | 0 | 144 | 33 | 170 | 90 | 163 | 52 | 47 | 12 | 13 | 6 | 783 | 339 | 410 | 2267 | 527 | 68 | 595 | 719 | 34 | 207 |
| PBANKA_0502200 | 0 | 0 | 218 | 36 | 497 | 249 | 681 | 229 | 213 | 99 | 22 | 27 | 1169 | 354 | 1511 | 6299 | 2624 | 392 | 862 | 675 | 44 | 450 |
| PBANKA_0502300 | 43 | 80 | 87 | 24 | 174 | 91 | 221 | 69 | 84 | 35 | 10 | 0 | 1356 | 518 | 463 | 2499 | 1251 | 151 | 163 | 184 | 11 | 73 |
| PBANKA_0502400 | 8 | 16 | 20 | 0 | 35 | 17 | 57 | 15 | 22 | 0 | 5 | 0 | 408 | 126 | 101 | 582 | 332 | 77 | 1263 | 378 | 5 | 22 |
| PBANKA_0502500 | 143 | 247 | 33 | 8 | 247 | 104 | 818 | 308 | 1216 | 535 | 444 | 229 | 2190 | 2149 | 286 | 1193 | 20840 | 3571 | 18345 | 46616 | 481 | 3654 |
| PBANKA_0502600 | 38 | 80 | 110 | 30 | 474 | 232 | 1198 | 490 | 545 | 260 | 69 | 21 | 2029 | 632 | 717 | 3660 | 6481 | 841 | 813 | 762 | 259 | 1970 |
| PBANKA_0502650 | 6 | 10 | 16 | 5 | 21 | 22 | 29 | 13 | 25 | 9 | 0 | 5 | 289 | 160 | 7 | 29 | 92 | 84 | 8987 | 4393 | 93 | 1028 |
| PBANKA_0502700 | 50 | 38 | 106 | 68 | 190 | 179 | 248 | 83 | 77 | 100 | 277 | 423 | 405 | 169 | 289 | 765 | 203 | 63 | 960 | 301 | 6 | 62 |
| PBANKA_0502800 | 22 | 23 | 112 | 18 | 275 | 87 | 278 | 114 | 237 | 118 | 345 | 172 | 1703 | 3809 | 519 | 3205 | 2474 | 359 | 1851 | 7651 | 1435 | 8233 |
| PBANKA_0502900 | 81 | 108 | 34 | 16 | 109 | 60 | 171 | 38 | 111 | 42 | 349 | 187 | 3212 | 2202 | 381 | 2212 | 1564 | 264 | 2218 | 1455 | 171 | 1241 |
| PBANKA_0503000 | 8 | 9 | 98 | 22 | 180 | 92 | 134 | 59 | 73 | 21 | 99 | 67 | 130 | 151 | 123 | 530 | 237 | 47 | 530 | 551 | 97 | 781 |
| PBANKA_0503100 | 10 | 24 | 59 | 14 | 99 | 37 | 125 | 52 | 156 | 99 | 31 | 21 | 639 | 213 | 164 | 644 | 275 | 78 | 1411 | 717 | 104 | 612 |
| PBANKA_0503200 | 193 | 346 | 313 | 62 | 1223 | 476 | 2121 | 738 | 1129 | 471 | 273 | 117 | 4036 | 4497 | 2396 | 12760 | 17651 | 2615 | 2542 | 5653 | 840 | 6966 |
| PBANKA_0503300 | 0 | 0 | 17 | 0 | 66 | 31 | 245 | 110 | 864 | 408 | 163 | 101 | 1313 | 1548 | 226 | 1092 | 7721 | 1775 | 13782 | 56340 | 423 | 3054 |
| PBANKA_0503500 | 26 | 41 | 171 | 49 | 428 | 176 | 427 | 173 | 221 | 98 | 61 | 32 | 915 | 1223 | 76 | 378 | 2298 | 662 | 15953 | 34279 | 510 | 3334 |
| PBANKA_0503600 | 5 | 6 | 946 | 238 | 545 | 334 | 497 | 413 | 201 | 159 | 840 | 706 | 4271 | 1916 | 1104 | 3213 | 1007 | 272 | 1071 | 507 | 95 | 627 |
| PBANKA_0503700 | 73 | 80 | 48 | 15 | 329 | 192 | 410 | 157 | 154 | 73 | 17 | 8 | 417 | 158 | 901 | 3409 | 1628 | 322 | 1194 | 895 | 1728 | 10203 |
| PBANKA_0503800 | 23 | 38 | 15 | 0 | 50 | 32 | 60 | 24 | 56 | 21 | 16 | 8 | 176 | 136 | 112 | 441 | 543 | 147 | 3636 | 5692 | 2938 | 18851 |
| PBANKA_0503900 | 16 | 20 | 5 | 0 | 99 | 34 | 286 | 119 | 153 | 85 | 20 | 6 | 319 | 113 | 90 | 346 | 2432 | 372 | 555 | 542 | 45 | 341 |
| PBANKA_0504000 | 34 | 38 | 148 | 19 | 230 | 80 | 201 | 75 | 112 | 20 | 19 | 8 | 1539 | 710 | 778 | 4026 | 1162 | 192 | 5017 | 5328 | 708 | 4705 |
| PBANKA_0504100 | 8 | 0 | 10 | 0 | 16 | 18 | 29 | 5 | 79 | 39 | 10 | 6 | 104 | 25 | 22 | 110 | 922 | 278 | 783 | 332 | 69 | 550 |
| PBANKA_0504200 | 12 | 15 | 68 | 21 | 65 | 40 | 58 | 19 | 80 | 32 | 390 | 283 | 1934 | 1596 | 321 | 1986 | 916 | 211 | 723 | 679 | 60 | 273 |
| PBANKA_0504500 | 0 | 0 | 62 | 15 | 153 | 67 | 167 | 59 | 57 | 26 | 12 | 0 | 408 | 132 | 214 | 1039 | 966 | 207 | 954 | 1469 | 956 | 6876 |
| PBANKA_0504600 | 0 | 0 | 84 | 17 | 127 | 66 | 162 | 58 | 51 | 23 | 17 | 7 | 649 | 252 | 364 | 2222 | 917 | 112 | 134 | 74 | 308 | 1765 |
| PBANKA_0504700 | 0 | 0 | 117 | 27 | 265 | 122 | 323 | 115 | 120 | 85 | 29 | 12 | 585 | 356 | 427 | 2559 | 1702 | 136 | 121 | 99 | 143 | 833 |
| PBANKA_0504800 | 13 | 23 | 89 | 8 | 174 | 68 | 247 | 86 | 89 | 35 | 34 | 31 | 1673 | 1091 | 1045 | 4863 | 3493 | 436 | 3495 | 2851 | 357 | 2126 |
| PBANKA_0504900 | 20 | 22 | 229 | 33 | 205 | 81 | 142 | 46 | 140 | 57 | 404 | 181 | 2246 | 2094 | 758 | 3475 | 1092 | 222 | 542 | 680 | 157 | 1244 |
| PBANKA_0505000 | 0 | 0 | 143 | 47 | 743 | 377 | 800 | 426 | 208 | 140 | 19 | 0 | 41 | 25 | 18 | 50 | 121 | 21 | 236 | 294 | 525 | 3670 |
| PBANKA_0505100 | 0 | 0 | 225 | 109 | 346 | 267 | 794 | 489 | 427 | 319 | 99 | 56 | 1426 | 643 | 771 | 1787 | 3411 | 830 | 1119 | 678 | 96 | 962 |
| PBANKA_0505200 | 0 | 0 | 129 | 40 | 299 | 166 | 517 | 220 | 170 | 93 | 26 | 16 | 780 | 136 | 493 | 1813 | 2518 | 450 | 1783 | 703 | 1243 | 9366 |
| PBANKA_0505300 | 79 | 121 | 133 | 26 | 358 | 171 | 544 | 186 | 186 | 85 | 62 | 22 | 1383 | 431 | 523 | 2657 | 2562 | 339 | 729 | 600 | 214 | 1387 |
| PBANKA_0505400 | 7 | 12 | 34 | 6 | 186 | 61 | 319 | 109 | 157 | 78 | 54 | 26 | 1240 | 376 | 342 | 1714 | 4095 | 663 | 4842 | 4628 | 1188 | 7959 |
| PBANKA_0505500 | 0 | 0 | 38 | 0 | 92 | 32 | 190 | 58 | 81 | 28 | 11 | 0 | 1054 | 155 | 243 | 794 | 931 | 234 | 2361 | 735 | 264 | 1644 |
| PBANKA_0505600 | 0 | 0 | 102 | 34 | 337 | 200 | 462 | 196 | 199 | 103 | 37 | 16 | 1302 | 260 | 699 | 1988 | 2319 | 466 | 534 | 255 | 135 | 1086 |
| PBANKA_0505700 | 113 | 156 | 238 | 49 | 472 | 192 | 487 | 175 | 191 | 129 | 1624 | 1009 | 1660 | 1229 | 458 | 2078 | 1351 | 260 | 2407 | 2276 | 1865 | 11688 |
| PBANKA_0505800 | 15 | 12 | 35 | 7 | 158 | 93 | 174 | 71 | 61 | 41 | 201 | 143 | 693 | 272 | 413 | 1085 | 906 | 191 | 3564 | 1636 | 197 | 1210 |
| PBANKA_0505900 | 11 | 19 | 148 | 18 | 481 | 186 | 631 | 195 | 2990 | 1288 | 2526 | 1161 | 3499 | 10012 | 3579 | 17718 | 26053 | 4957 | 3061 | 11549 | 120 | 772 |
| PBANKA_0506000 | 192 | 368 | 239 | 70 | 541 | 232 | 956 | 387 | 740 | 396 | 145 | 66 | 1278 | 1167 | 743 | 3765 | 25667 | 3193 | 738 | 3528 | 6137 | 40315 |
| PBANKA_0506100 | 66 | 115 | 655 | 145 | 1094 | 415 | 2250 | 789 | 1197 | 489 | 981 | 544 | 3413 | 3174 | 3998 | 22422 | 45828 | 4384 | 994 | 4859 | 5684 | 37682 |
| PBANKA_0506200 | 1204 | 1530 | 696 | 175 | 1807 | 699 | 1472 | 415 | 744 | 281 | 4154 | 2139 | 2535 | 3675 | 435 | 2416 | 3178 | 428 | 994 | 1129 | 887 | 6545 |
| PBANKA_0506300 | 22 | 23 | 50 | 9 | 55 | 27 | 67 | 28 | 44 | 33 | 289 | 215 | 572 | 460 | 180 | 668 | 204 | 46 | 853 | 485 | 153 | 908 |
| PBANKA_0506400 | 0 | 0 | 95 | 29 | 162 | 87 | 224 | 91 | 64 | 19 | 17 | 6 | 2083 | 669 | 545 | 2205 | 2023 | 262 | 554 | 258 | 9 | 55 |
| PBANKA_0506500 | 325 | 296 | 154 | 50 | 338 | 213 | 296 | 137 | 84 | 44 | 9 | 7 | 703 | 205 | 302 | 655 | 381 | 110 | 955 | 394 | 34 | 185 |
| PBANKA_0506600 | 45 | 45 | 385 | 141 | 572 | 365 | 934 | 434 | 393 | 225 | 54 | 27 | 2869 | 828 | 1977 | 13020 | 12612 | 1178 | 12154 | 15357 | 133 | 1454 |
| PBANKA_0506700 | 413 | 580 | 100 | 22 | 328 | 283 | 371 | 138 | 88 | 64 | 5 | 15 | 255 | 121 | 495 | 1952 | 819 | 188 | 3068 | 1918 | 145 | 1274 |
| PBANKA_0506800 | 227 | 239 | 156 | 56 | 211 | 105 | 330 | 152 | 170 | 66 | 103 | 110 | 595 | 829 | 751 | 5018 | 553 | 103 | 670 | 1736 | 169 | 1230 |
| PBANKA_0506900 | 80 | 119 | 20 | 0 | 75 | 44 | 119 | 72 | 469 | 322 | 87 | 50 | 276 | 172 | 15 | 31 | 2557 | 770 | 2759 | 1956 | 602 | 5051 |
| PBANKA_0507000 | 39 | 52 | 106 | 43 | 414 | 214 | 507 | 223 | 320 | 140 | 94 | 25 | 323 | 101 | 486 | 2216 | 3205 | 518 | 1486 | 1560 | 53 | 383 |
| PBANKA_0507100 | 0 | 0 | 87 | 31 | 127 | 153 | 205 | 156 | 83 | 82 | 11 | 6 | 128 | 15 | 188 | 273 | 432 | 105 | 343 | 108 | 18 | 137 |
| PBANKA_0507200 | 163 | 145 | 15 | 0 | 102 | 30 | 139 | 28 | 72 | 49 | 0 | 0 | 184 | 34 | 85 | 293 | 552 | 162 | 504 | 230 | 392 | 2386 |
| PBANKA_0507300 | 30 | 21 | 147 | 36 | 485 | 185 | 592 | 228 | 242 | 89 | 65 | 5 | 630 | 666 | 1124 | 5516 | 4299 | 572 | 2948 | 15491 | 272 | 2195 |
| PBANKA_0507400 | 0 | 0 | 91 | 21 | 154 | 75 | 113 | 56 | 37 | 19 | 6 | 7 | 573 | 157 | 388 | 1959 | 563 | 64 | 521 | 307 | 291 | 2103 |
| PBANKA_0507500 | 0 | 0 | 12 | 6 | 18 | 9 | 36 | 12 | 6 | 0 | 0 | 0 | 56 | 6 | 78 | 234 | 104 | 21 | 298 | 94 | 0 | 9 |
| PBANKA_0507600 | 18 | 38 | 51 | 15 | 123 | 92 | 121 | 74 | 46 | 22 | 6 | 0 | 38 | 0 | 62 | 211 | 113 | 30 | 169 | 99 | 29 | 150 |
| PBANKA_0507700 | 0 | 7 | 27 | 0 | 59 | 17 | 50 | 20 | 12 | 7 | 0 | 0 | 98 | 10 | 53 | 153 | 136 | 36 | 417 | 225 | 19 | 100 |
| PBANKA_0507800 | 0 | 0 | 15 | 0 | 21 | 15 | 35 | 26 | 18 | 11 | 5 | 0 | 790 | 290 | 94 | 547 | 960 | 136 | 5053 | 5013 | 21 | 162 |
| PBANKA_0508100 | 838 | 994 | 1076 | 240 | 2562 | 1144 | 3055 | 1273 | 3977 | 1901 | 15445 | 7776 | 12718 | 21635 | 7299 | 48120 | 43633 | 5113 | 5276 | 17478 | 1982 | 15129 |
| PBANKA_0508200 | 16 | 15 | 66 | 12 | 170 | 70 | 160 | 50 | 70 | 21 | 38 | 12 | 1292 | 526 | 394 | 1713 | 1449 | 212 | 304 | 147 | 124 | 824 |
| PBANKA_0508300 | 694 | 973 | 55 | 20 | 338 | 119 | 273 | 136 | 129 | 69 | 34 | 15 | 261 | 188 | 83 | 420 | 1322 | 169 | 793 | 2791 | 10359 | 66538 |
| PBANKA_0508400 | 44 | 60 | 47 | 10 | 120 | 55 | 199 | 95 | 103 | 40 | 5 | 6 | 297 | 49 | 157 | 476 | 708 | 162 | 436 | 214 | 14 | 157 |
| PBANKA_0508500 | 24 | 49 | 43 | 18 | 151 | 79 | 184 | 87 | 50 | 45 | 0 | 5 | 409 | 51 | 263 | 880 | 929 | 170 | 983 | 444 | 29 | 273 |

| | | | | | | | | | | | | | | | | | | | | | | |
|---|---|---|---|---|---|---|---|---|---|---|---|---|---|---|---|---|---|---|---|---|---|---|
| PBANKA_0508600 | 0 | 0 | 39 | 6 | 129 | 75 | 192 | 60 | 63 | 37 | 12 | 0 | 637 | 218 | 239 | 750 | 1281 | 251 | 3444 | 3555 | 312 | 2060 |
| PBANKA_0508700 | 19 | 19 | 21 | 7 | 75 | 22 | 60 | 25 | 29 | 8 | 0 | 0 | 215 | 59 | 128 | 324 | 338 | 85 | 803 | 267 | 184 | 1067 |
| PBANKA_0508800 | 0 | 0 | 48 | 23 | 107 | 63 | 102 | 64 | 54 | 32 | 11 | 0 | 599 | 183 | 106 | 468 | 235 | 49 | 802 | 367 | 450 | 2486 |
| PBANKA_0508900 | 0 | 0 | 87 | 18 | 270 | 115 | 347 | 101 | 99 | 53 | 21 | 0 | 1270 | 427 | 370 | 2369 | 1652 | 229 | 1037 | 663 | 1113 | 7722 |
| PBANKA_0509000 | 0 | 0 | 22 | 7 | 163 | 83 | 230 | 79 | 155 | 90 | 1709 | 1428 | 10962 | 5008 | 4051 | 19296 | 6244 | 885 | 6508 | 4866 | 276 | 2564 |
| PBANKA_0509100 | 8 | 17 | 61 | 26 | 156 | 91 | 136 | 48 | 97 | 71 | 51 | 20 | 666 | 327 | 556 | 2632 | 498 | 129 | 793 | 660 | 371 | 2335 |
| PBANKA_0509200 | 38 | 49 | 71 | 0 | 220 | 112 | 386 | 210 | 305 | 143 | 1356 | 798 | 6041 | 4307 | 1491 | 8100 | 7002 | 1095 | 4543 | 4122 | 1154 | 7900 |
| PBANKA_0509300 | 5 | 10 | 58 | 6 | 295 | 172 | 475 | 183 | 211 | 98 | 53 | 31 | 664 | 217 | 535 | 1657 | 3191 | 730 | 3414 | 2306 | 1169 | 9503 |
| PBANKA_0509400 | 0 | 0 | 26 | 12 | 76 | 41 | 108 | 26 | 23 | 8 | 10 | 0 | 96 | 36 | 210 | 1019 | 204 | 32 | 307 | 154 | 76 | 469 |
| PBANKA_0509500 | 221 | 355 | 313 | 68 | 707 | 330 | 728 | 282 | 238 | 107 | 44 | 29 | 1645 | 1013 | 1164 | 5667 | 3292 | 440 | 4804 | 5780 | 3071 | 16375 |
| PBANKA_0509600 | 12 | 47 | 10 | 0 | 47 | 26 | 119 | 46 | 379 | 182 | 62 | 28 | 771 | 162 | 68 | 193 | 4727 | 1650 | 796 | 387 | 261 | 1365 |
| PBANKA_0509700 | 5 | 8 | 11 | 0 | 126 | 29 | 136 | 50 | 207 | 72 | 96 | 32 | 447 | 430 | 266 | 1452 | 3542 | 563 | 2437 | 6950 | 35 | 245 |
| PBANKA_0509800 | 0 | 0 | 23 | 7 | 61 | 26 | 79 | 26 | 13 | 11 | 0 | 0 | 101 | 19 | 200 | 701 | 215 | 35 | 409 | 208 | 10 | 70 |
| PBANKA_0509900 | 5 | 0 | 59 | 0 | 197 | 86 | 204 | 65 | 65 | 40 | 14 | 0 | 218 | 57 | 519 | 1869 | 672 | 149 | 1183 | 634 | 71 | 522 |
| PBANKA_0510000 | 0 | 0 | 19 | 0 | 33 | 27 | 69 | 34 | 29 | 29 | 0 | 0 | 80 | 21 | 26 | 87 | 296 | 117 | 465 | 188 | 24 | 186 |
| PBANKA_0510100 | 27 | 59 | 17 | 0 | 82 | 49 | 433 | 158 | 852 | 403 | 104 | 63 | 851 | 749 | 127 | 340 | 10389 | 2361 | 2929 | 4628 | 1040 | 5600 |
| PBANKA_0510200 | 15 | 8 | 137 | 29 | 307 | 88 | 295 | 81 | 161 | 65 | 133 | 71 | 8626 | 3730 | 1754 | 11966 | 2918 | 823 | 5206 | 3935 | 395 | 2400 |
| PBANKA_0510300 | 0 | 0 | 20 | 5 | 103 | 32 | 71 | 31 | 22 | 15 | 0 | 0 | 600 | 262 | 84 | 443 | 171 | 116 | 14330 | 6929 | 130 | 752 |
| PBANKA_0510400 | 9 | 0 | 134 | 8 | 145 | 71 | 119 | 38 | 45 | 21 | 10 | 0 | 349 | 69 | 384 | 1370 | 153 | 75 | 3132 | 1379 | 1727 | 10523 |
| PBANKA_0510500 | 0 | 0 | 53 | 9 | 403 | 175 | 464 | 132 | 96 | 47 | 33 | 9 | 125 | 85 | 98 | 463 | 130 | 38 | 818 | 772 | 609 | 3595 |
| PBANKA_0510600 | 0 | 0 | 5 | 0 | 23 | 9 | 34 | 14 | 22 | 8 | 0 | 0 | 240 | 57 | 26 | 59 | 158 | 42 | 2038 | 1140 | 99 | 736 |
| PBANKA_0510700 | 453 | 499 | 15 | 0 | 15 | 9 | 14 | 6 | 8 | 0 | 0 | 0 | 15 | 0 | 9 | 26 | 65 | 7 | 5 | 8 | 479 | 2692 |
| PBANKA_0510800 | 794 | 989 | 23 | 5 | 87 | 25 | 211 | 53 | 851 | 436 | 352 | 151 | 876 | 1789 | 199 | 1032 | 10576 | 2877 | 823 | 4062 | 1071 | 5584 |
| PBANKA_0510900 | 163 | 166 | 1952 | 664 | 1287 | 1285 | 1356 | 777 | 456 | 269 | 2365 | 2336 | 10391 | 6591 | 2807 | 10055 | 2444 | 370 | 2141 | 1192 | 1335 | 11474 |
| PBANKA_0511000 | 0 | 6 | 159 | 38 | 161 | 91 | 123 | 38 | 39 | 22 | 12 | 7 | 94 | 56 | 26 | 65 | 65 | 12 | 33 | 9 | 1127 | 7419 |
| PBANKA_0511100 | 0 | 0 | 86 | 9 | 328 | 90 | 486 | 126 | 210 | 90 | 27 | 17 | 127 | 96 | 98 | 251 | 2769 | 337 | 224 | 405 | 388 | 2637 |
| PBANKA_0511200 | 53 | 69 | 251 | 44 | 443 | 187 | 538 | 221 | 329 | 151 | 1532 | 1069 | 4220 | 5444 | 828 | 4707 | 2504 | 640 | 1118 | 2909 | 411 | 2772 |
| PBANKA_0511300 | 10 | 12 | 68 | 28 | 171 | 86 | 139 | 72 | 64 | 29 | 68 | 22 | 760 | 491 | 267 | 950 | 512 | 149 | 607 | 433 | 383 | 2890 |
| PBANKA_0511400 | 1041 | 1505 | 123 | 26 | 285 | 120 | 1418 | 826 | 3096 | 2162 | 2413 | 1938 | 10134 | 8679 | 294 | 1335 | 82106 | 21609 | 3965 | 5976 | 458 | 2755 |
| PBANKA_0511500 | 107 | 116 | 124 | 34 | 167 | 164 | 307 | 179 | 98 | 68 | 11 | 7 | 646 | 262 | 268 | 1114 | 1062 | 150 | 433 | 283 | 50 | 362 |
| PBANKA_0511600 | 0 | 0 | 8 | 0 | 30 | 0 | 39 | 11 | 12 | 6 | 0 | 0 | 101 | 24 | 34 | 158 | 102 | 18 | 20 | 12 | 6 | 39 |
| PBANKA_0511700 | 76 | 70 | 25 | 7 | 59 | 54 | 170 | 66 | 88 | 58 | 12 | 0 | 696 | 173 | 130 | 291 | 476 | 234 | 6805 | 2674 | 831 | 3915 |
| PBANKA_0511800 | 0 | 0 | 23 | 8 | 131 | 84 | 335 | 183 | 169 | 120 | 18 | 13 | 481 | 109 | 97 | 259 | 1525 | 447 | 1095 | 507 | 44 | 318 |
| PBANKA_0511900 | 12 | 10 | 2922 | 927 | 2010 | 1211 | 2110 | 918 | 771 | 455 | 3629 | 2654 | 21272 | 11106 | 4718 | 19792 | 6278 | 830 | 2660 | 1955 | 351 | 2964 |
| PBANKA_0512100 | 12 | 10 | 102 | 25 | 120 | 64 | 85 | 29 | 87 | 44 | 47 | 22 | 1138 | 646 | 204 | 715 | 284 | 69 | 327 | 232 | 230 | 1329 |
| PBANKA_0512200 | 52 | 84 | 67 | 10 | 160 | 69 | 189 | 41 | 66 | 15 | 30 | 14 | 712 | 631 | 112 | 544 | 283 | 74 | 2798 | 1161 | 335 | 1975 |
| PBANKA_0512300 | 0 | 0 | 14 | 0 | 71 | 25 | 125 | 43 | 50 | 24 | 10 | 0 | 149 | 30 | 123 | 471 | 493 | 41 | 593 | 294 | 152 | 893 |
| PBANKA_0512400 | 32 | 24 | 142 | 22 | 166 | 65 | 116 | 38 | 103 | 65 | 1309 | 1002 | 2045 | 2165 | 322 | 1670 | 401 | 98 | 1871 | 1526 | 614 | 3851 |
| PBANKA_0512500 | 20 | 11 | 121 | 31 | 85 | 39 | 80 | 29 | 91 | 55 | 1090 | 699 | 2133 | 2136 | 152 | 926 | 283 | 84 | 2837 | 2588 | 1116 | 7812 |
| PBANKA_0512600 | 70 | 87 | 115 | 27 | 256 | 91 | 386 | 127 | 231 | 114 | 161 | 92 | 1787 | 1030 | 367 | 2518 | 4353 | 581 | 1623 | 1866 | 332 | 1341 |
| PBANKA_0512700 | 36 | 34 | 13 | 5 | 110 | 49 | 262 | 158 | 265 | 214 | 43 | 19 | 826 | 160 | 177 | 457 | 2699 | 834 | 2060 | 1075 | 93 | 485 |
| PBANKA_0512800 | 365 | 468 | 25 | 23 | 56 | 81 | 205 | 72 | 2189 | 1639 | 399 | 266 | 1174 | 712 | 273 | 1285 | 4317 | 1947 | 4495 | 2491 | 263 | 3379 |
| PBANKA_0512900 | 17 | 14 | 80 | 30 | 333 | 219 | 563 | 286 | 309 | 248 | 45 | 18 | 297 | 56 | 297 | 616 | 1142 | 432 | 669 | 215 | 26 | 469 |
| PBANKA_0513100 | 10 | 12 | 38 | 10 | 139 | 42 | 167 | 62 | 144 | 66 | 21 | 12 | 330 | 210 | 199 | 858 | 399 | 80 | 592 | 476 | 664 | 3478 |
| PBANKA_0513200 | 115 | 130 | 50 | 19 | 79 | 20 | 105 | 29 | 70 | 31 | 163 | 99 | 1575 | 734 | 277 | 1253 | 270 | 63 | 631 | 352 | 383 | 2444 |
| PBANKA_0513300 | 51 | 61 | 37 | 5 | 86 | 34 | 115 | 78 | 103 | 62 | 85 | 27 | 537 | 415 | 305 | 1332 | 244 | 61 | 339 | 273 | 481 | 2303 |
| PBANKA_0513400 | 6 | 8 | 24 | 7 | 74 | 31 | 154 | 47 | 232 | 78 | 48 | 14 | 1594 | 436 | 475 | 2360 | 2954 | 616 | 678 | 461 | 93 | 645 |
| PBANKA_0513500 | 0 | 0 | 5 | 0 | 0 | 0 | 7 | 0 | 0 | 0 | 0 | 0 | 44 | 0 | 32 | 200 | 38 | 8 | 433 | 171 | 7 | 39 |
| PBANKA_0513600 | 85 | 92 | 472 | 119 | 272 | 139 | 212 | 105 | 79 | 47 | 28 | 13 | 4607 | 693 | 5171 | 28790 | 1250 | 380 | 2098 | 1261 | 95 | 661 |
| PBANKA_0513800 | 117 | 85 | 85 | 27 | 272 | 125 | 555 | 224 | 308 | 168 | 21 | 25 | 997 | 296 | 450 | 1837 | 3371 | 499 | 814 | 571 | 430 | 3708 |
| PBANKA_0513900 | 30 | 65 | 128 | 32 | 427 | 159 | 3389 | 1616 | 3553 | 1795 | 418 | 174 | 1761 | 883 | 342 | 1790 | 21475 | 4039 | 7084 | 10814 | 1047 | 7439 |
| PBANKA_0514000 | 0 | 8 | 37 | 11 | 84 | 44 | 70 | 40 | 35 | 30 | 282 | 203 | 308 | 109 | 116 | 329 | 200 | 75 | 517 | 153 | 72 | 453 |
| PBANKA_0514100 | 19 | 21 | 218 | 43 | 575 | 233 | 691 | 250 | 438 | 230 | 2000 | 1212 | 5608 | 3096 | 1469 | 6690 | 3659 | 523 | 1668 | 1450 | 660 | 5051 |
| PBANKA_0514200 | 244 | 312 | 6 | 0 | 78 | 41 | 130 | 49 | 356 | 281 | 85 | 54 | 262 | 91 | 41 | 125 | 1540 | 412 | 534 | 307 | 99 | 698 |
| PBANKA_0514300 | 160 | 94 | 83 | 22 | 412 | 120 | 509 | 172 | 438 | 163 | 984 | 469 | 2622 | 3092 | 494 | 4902 | 2978 | 407 | 14200 | 26640 | 1107 | 9161 |
| PBANKA_0514400 | 0 | 0 | 11 | 0 | 47 | 22 | 128 | 63 | 81 | 25 | 13 | 0 | 148 | 42 | 40 | 148 | 74 | 28 | 364 | 135 | 21 | 166 |
| PBANKA_0514500 | 0 | 0 | 64 | 12 | 132 | 76 | 287 | 131 | 102 | 39 | 10 | 5 | 1954 | 724 | 974 | 4814 | 2570 | 381 | 1535 | 1171 | 168 | 1153 |
| PBANKA_0514600 | 20 | 25 | 96 | 20 | 226 | 102 | 238 | 79 | 70 | 38 | 43 | 21 | 1094 | 597 | 351 | 1471 | 629 | 84 | 160 | 176 | 32 | 115 |
| PBANKA_0514700 | 20 | 16 | 86 | 15 | 127 | 78 | 105 | 42 | 42 | 17 | 11 | 7 | 729 | 347 | 220 | 1250 | 336 | 40 | 73 | 70 | 7 | 41 |
| PBANKA_0514800 | 45 | 70 | 13 | 0 | 46 | 32 | 85 | 39 | 55 | 43 | 9 | 0 | 391 | 162 | 100 | 664 | 950 | 229 | 1594 | 1055 | 58 | 382 |
| PBANKA_0515000 | 251 | 284 | 19 | 11 | 41 | 27 | 32 | 30 | 18 | 20 | 0 | 16 | 10662 | 3739 | 254 | 499 | 3064 | 2709 | 4E+05 | 2E+05 | 17643 | 2E+05 |
| PBANKA_0515100 | 0 | 0 | 19 | 0 | 50 | 22 | 47 | 17 | 22 | 11 | 0 | 0 | 117 | 22 | 82 | 316 | 202 | 26 | 321 | 62 | 31 | 194 |
| PBANKA_0515200 | 15 | 12 | 14 | 0 | 55 | 24 | 65 | 30 | 274 | 158 | 59 | 51 | 218 | 169 | 46 | 551 | 2294 | 363 | 297 | 572 | 55 | 569 |
| PBANKA_0515300 | 101 | 94 | 111 | 31 | 288 | 141 | 543 | 223 | 408 | 189 | 62 | 24 | 821 | 411 | 312 | 1529 | 8414 | 1074 | 1879 | 4549 | 1400 | 9664 |
| PBANKA_0515350 | 5 | 10 | 8 | 5 | 19 | 6 | 24 | 9 | 10 | 10 | 10 | 0 | 38 | 14 | 15 | 46 | 71 | 21 | 350 | 231 | 79 | 495 |
| PBANKA_0515400 | 203 | 272 | 87 | 22 | 321 | 122 | 464 | 185 | 246 | 86 | 58 | 11 | 1544 | 1066 | 375 | 3001 | 3758 | 326 | 3376 | 5505 | 548 | 4322 |
| PBANKA_0515500 | 43 | 46 | 55 | 18 | 139 | 88 | 87 | 47 | 31 | 38 | 21 | 17 | 211 | 63 | 123 | 227 | 387 | 137 | 123 | 59 | 68 | 380 |
| PBANKA_0515700 | 0 | 0 | 10 | 0 | 51 | 30 | 54 | 27 | 20 | 8 | 0 | 5 | 59 | 15 | 99 | 272 | 117 | 28 | 433 | 314 | 20 | 201 |
| PBANKA_0515800 | 6 | 9 | 103 | 22 | 196 | 82 | 191 | 64 | 98 | 31 | 103 | 67 | 1770 | 1001 | 421 | 1759 | 1022 | 170 | 690 | 295 | 95 | 571 |
| PBANKA_0515900 | 154 | 146 | 147 | 38 | 615 | 169 | 949 | 304 | 411 | 182 | 1105 | 652 | 4481 | 3966 | 1036 | 8073 | 9213 | 984 | 4741 | 7266 | 1361 | 8942 |
| PBANKA_0516100 | 10 | 5 | 72 | 10 | 98 | 46 | 97 | 33 | 38 | 19 | 19 | 8 | 462 | 150 | 406 | 1618 | 649 | 146 | 1402 | 1601 | 95 | 673 |
| PBANKA_0516200 | 0 | 0 | 196 | 46 | 581 | 378 | 878 | 417 | 280 | 183 | 67 | 16 | 633 | 135 | 611 | 2014 | 1853 | 356 | 929 | 538 | 60 | 422 |
| PBANKA_0516300 | 48 | 80 | 92 | 25 | 272 | 114 | 256 | 102 | 90 | 47 | 9 | 9 | 932 | 212 | 584 | 1761 | 670 | 146 | 1103 | 489 | 22 | 126 |
| PBANKA_0516400 | 8 | 0 | 29 | 5 | 81 | 30 | 110 | 33 | 55 | 14 | 22 | 22 | 371 | 133 | 131 | 587 | 232 | 75 | 1264 | 920 | 14 | 114 |
| PBANKA_0516500 | 73 | 104 | 134 | 45 | 402 | 173 | 694 | 366 | 346 | 223 | 164 | 79 | 3567 | 2397 | 690 | 3822 | 4471 | 565 | 1443 | 1317 | 210 | 1543 |
| PBANKA_0516600 | 0 | 0 | 96 | 18 | 186 | 92 | 394 | 173 | 252 | 168 | 48 | 26 | 846 | 164 | 349 | 1600 | 295 | 84 | 1604 | 623 | 21 | 163 |
| PBANKA_0516700 | 9 | 15 | 104 | 27 | 273 | 101 | 343 | 136 | 161 | 95 | 36 | 19 | 1034 | 875 | 563 | 2836 | 926 | 129 | 453 | 626 | 52 | 330 |
| PBANKA_0516800 | 0 | 0 | 63 | 22 | 217 | 108 | 256 | 94 | 89 | 38 | 8 | 0 | 726 | 125 | 442 | 1678 | 430 | 76 | 558 | 427 | 40 | 316 |
| PBANKA_0516900 | 1282 | 1948 | 934 | 259 | 1569 | 796 | 1536 | 460 | 1109 | 454 | 4422 | 2538 | 5142 | 4490 | 3199 | 12090 | 3919 | 747 | 2423 | 2775 | 3163 | 25834 |
| PBANKA_0517000 | 31 | 26 | 159 | 47 | 250 | 206 | 280 | 143 | 1716 | 1539 | 11986 | 11488 | 20683 | 10792 | 4522 | 14632 | 5331 | 1501 | 6139 | 2279 | 196 | 2211 |
| PBANKA_0517100 | 0 | 0 | 38 | 19 | 50 | 56 | 41 | 23 | 12 | 8 | 19 | 13 | 362 | 157 | 123 | 199 | 70 | 19 | 158 | 48 | 0 | 7 |
| PBANKA_0517200 | 0 | 0 | 130 | 21 | 89 | 41 | 74 | 39 | 6 | 5 | 47 | 28 | 2920 | 2137 | 487 | 2398 | 531 | 79 | 723 | 407 | 8 | 60 |
| PBANKA_0517300 | 0 | 0 | 87 | 19 | 197 | 101 | 241 | 73 | 94 | 42 | 85 | 48 | 1228 | 1227 | 455 | 2613 | 811 | 120 | 265 | 528 | 46 | 291 |
| PBANKA_0517400 | 132 | 164 | 197 | 59 | 799 | 483 | 841 | 350 | 353 | 171 | 901 | 533 | 2470 | 1969 | 1464 | 6721 | 7001 | 1080 | 1558 | 2427 | 166 | 1495 |
| PBANKA_0517500 | 26 | 32 | 261 | 74 | 266 | 136 | 257 | 93 | 220 | 130 | 1750 | 1544 | 5250 | 3342 | 537 | 2582 | 2157 | 402 | 2699 | 1360 | 139 | 1416 |
| PBANKA_0517700 | 107 | 183 | 15 | 0 | 514 | 255 | 976 | 534 | 379 | 220 | 60 | 27 | 577 | 103 | 244 | 745 | 386 | 74 | 1341 | 523 | 31 | 267 |
| PBANKA_0517800 | 0 | 0 | 63 | 32 | 396 | 168 | 508 | 196 | 213 | 84 | 19 | 11 | 1050 | 370 | 555 | 2478 | 1750 | 272 | 1386 | 807 | 65 | 501 |
| PBANKA_0517900 | 0 | 0 | 8 | 0 | 30 | 15 | 43 | 10 | 19 | 6 | 0 | 0 | 90 | 16 | 42 | 113 | 125 | 35 | 253 | 104 | 16 | 221 |
| PBANKA_0518000 | 0 | 0 | 30 | 6 | 91 | 42 | 105 | 56 | 59 | 17 | 8 | 6 | 250 | 100 | 136 | 670 | 549 | 87 | 1620 | 695 | 75 | 601 |
| PBANKA_0518200 | 0 | 0 | 206 | 83 | 456 | 336 | 459 | 220 | 121 | 67 | 16 | 22 | 157 | 41 | 584 | 2346 | 907 | 125 | 1289 | 666 | 85 | 998 |

| | | | | | | | | | | | | | | | | | | | | | | |
|---|---|---|---|---|---|---|---|---|---|---|---|---|---|---|---|---|---|---|---|---|---|---|
| PBANKA_0518300 | 0 | 0 | 60 | 19 | 237 | 93 | 405 | 107 | 124 | 49 | 19 | 5 | 452 | 99 | 516 | 2403 | 1458 | 248 | 1176 | 1341 | 18 | 103 |
| PBANKA_0518400 | 0 | 0 | 44 | 7 | 93 | 47 | 145 | 68 | 52 | 23 | 9 | 0 | 279 | 52 | 189 | 710 | 1003 | 164 | 572 | 335 | 6 | 85 |
| PBANKA_0518500 | 0 | 0 | 8 | 0 | 23 | 12 | 36 | 10 | 22 | 0 | 7 | 0 | 400 | 104 | 43 | 182 | 970 | 213 | 8425 | 4965 | 173 | 1665 |
| PBANKA_0518600 | 8 | 9 | 63 | 20 | 54 | 39 | 56 | 31 | 21 | 7 | 0 | 0 | 121 | 26 | 104 | 544 | 65 | 8 | 196 | 94 | 0 | 17 |
| PBANKA_0518700 | 111 | 182 | 773 | 308 | 1413 | 883 | 1652 | 573 | 516 | 302 | 1792 | 1694 | 6083 | 4109 | 1592 | 7008 | 3417 | 412 | 1714 | 1637 | 48 | 455 |
| PBANKA_0518800 | 0 | 0 | 171 | 54 | 349 | 160 | 675 | 245 | 398 | 187 | 233 | 101 | 1413 | 1661 | 589 | 3086 | 255 | 32 | 106 | 195 | 21 | 176 |
| PBANKA_0518900 | 0 | 0 | 65 | 17 | 3650 | 1698 | 29901 | 15205 | 24152 | 14636 | 2598 | 1436 | 93 | 48 | 83 | 393 | 564 | 145 | 122 | 243 | 30 | 309 |
| PBANKA_0519000 | 11 | 9 | 28 | 14 | 397 | 412 | 2298 | 1294 | 6325 | 5396 | 499 | 321 | 4322 | 6137 | 22 | 200 | 98680 | 2347 | 1608 | 3855 | 896 | 10281 |
| PBANKA_0519300 | 0 | 0 | 19 | 0 | 673 | 413 | 1706 | 1008 | 1811 | 1310 | 302 | 169 | 1345 | 520 | 7 | 14 | 14079 | 2942 | 381 | 523 | 41 | 434 |
| PBANKA_0519500 | 15 | 21 | 6 | 0 | 6253 | 3456 | 13981 | 7166 | 10883 | 7082 | 1354 | 777 | 652 | 68 | 149 | 862 | 408 | 91 | 149 | 213 | 16 | 134 |
| PBANKA_0519600 | 0 | 0 | 25 | 6 | 71 | 35 | 57 | 28 | 27 | 14 | 12 | 0 | 502 | 80 | 274 | 764 | 263 | 77 | 726 | 317 | 39 | 273 |
| PBANKA_0519700 | 0 | 0 | 62 | 14 | 114 | 108 | 116 | 60 | 29 | 15 | 0 | 0 | 722 | 142 | 503 | 1408 | 320 | 90 | 555 | 243 | 14 | 107 |
| PBANKA_0519800 | 0 | 0 | 81 | 28 | 477 | 321 | 842 | 547 | 331 | 208 | 44 | 18 | 1243 | 176 | 508 | 2020 | 3061 | 562 | 1923 | 947 | 66 | 464 |
| PBANKA_0519900 | 66 | 101 | 819 | 182 | 1391 | 574 | 1688 | 521 | 817 | 321 | 4608 | 2330 | 8393 | 10201 | 3217 | 15483 | 8700 | 1022 | 3846 | 5130 | 1557 | 11458 |
| PBANKA_0520000 | 0 | 0 | 270 | 79 | 557 | 357 | 711 | 251 | 247 | 84 | 33 | 20 | 613 | 147 | 450 | 1531 | 1123 | 170 | 644 | 341 | 48 | 364 |
| PBANKA_0520100 | 0 | 0 | 20 | 0 | 108 | 40 | 178 | 59 | 59 | 33 | 10 | 10 | 340 | 145 | 42 | 117 | 237 | 64 | 2097 | 1100 | 58 | 540 |
| PBANKA_0520200 | 29 | 18 | 236 | 72 | 378 | 249 | 404 | 193 | 147 | 65 | 538 | 488 | 4863 | 1932 | 943 | 4397 | 2674 | 352 | 4992 | 2829 | 148 | 1543 |
| PBANKA_0520300 | 14 | 5 | 44 | 0 | 220 | 79 | 234 | 89 | 81 | 52 | 12 | 7 | 595 | 197 | 352 | 1727 | 1883 | 389 | 569 | 392 | 195 | 1076 |
| PBANKA_0520400 | 0 | 6 | 37 | 5 | 125 | 85 | 444 | 185 | 422 | 234 | 52 | 29 | 688 | 234 | 96 | 468 | 4953 | 1131 | 1102 | 972 | 165 | 1569 |
| PBANKA_0520500 | 27 | 31 | 138 | 41 | 350 | 170 | 530 | 195 | 192 | 81 | 47 | 36 | 876 | 373 | 536 | 1885 | 1577 | 234 | 435 | 466 | 35 | 319 |
| PBANKA_0520600 | 0 | 0 | 93 | 37 | 123 | 63 | 190 | 112 | 81 | 38 | 33 | 25 | 577 | 293 | 286 | 539 | 357 | 73 | 93 | 159 | 8 | 59 |
| PBANKA_0520700 | 0 | 0 | 36 | 9 | 72 | 34 | 59 | 23 | 14 | 9 | 16 | 7 | 538 | 351 | 94 | 385 | 246 | 94 | 1754 | 1532 | 23 | 127 |
| PBANKA_0520800 | 0 | 0 | 20 | 7 | 116 | 48 | 142 | 67 | 76 | 34 | 10 | 14 | 531 | 131 | 126 | 636 | 473 | 148 | 467 | 202 | 24 | 167 |
| PBANKA_0520900 | 0 | 0 | 428 | 181 | 438 | 505 | 1130 | 576 | 561 | 392 | 739 | 561 | 3792 | 1146 | 1142 | 3592 | 6790 | 1317 | 1059 | 502 | 59 | 646 |
| PBANKA_0521000 | 0 | 0 | 113 | 24 | 432 | 186 | 456 | 205 | 202 | 91 | 23 | 10 | 972 | 731 | 726 | 3135 | 1445 | 216 | 436 | 314 | 33 | 191 |
| PBANKA_0521100 | 0 | 0 | 18 | 0 | 22 | 14 | 24 | 15 | 8 | 9 | 0 | 0 | 97 | 21 | 36 | 85 | 45 | 26 | 130 | 66 | 0 | 16 |
| PBANKA_0521221 | 8398 | 11787 | 10728 | 3046 | 20876 | 5107 | 23034 | 4493 | 13627 | 3394 | 12676 | 3227 | 9E+05 | 55408 | 70051 | 2E+05 | 1E+05 | 15211 | 2E+05 | 55562 | 771 | 11992 |
| PBANKA_0521241 | 0 | 0 | 338 | 136 | 38 | 26 | 35 | 10 | 23 | 12 | 0 | 0 | 198 | 28 | 1928 | 1865 | 380 | 102 | 241 | 21 | 12 | 140 |
| PBANKA_0521261 | 25021 | 28177 | 24791 | 9353 | 78405 | 26807 | 1E+05 | 25849 | 56853 | 22416 | 98194 | 28265 | 1E+05 | 13419 | 63100 | 82808 | 84185 | 11100 | 2E+05 | 72284 | 943 | 13641 |
| PBANKA_0521300 | 193 | 275 | 300 | 52 | 661 | 261 | 1106 | 396 | 1242 | 631 | 5999 | 3291 | 2582 | 10646 | 2613 | 14977 | 8336 | 1043 | 982 | 5069 | 289 | 1770 |
| PBANKA_0521400 | 42 | 59 | 109 | 22 | 204 | 70 | 206 | 74 | 138 | 54 | 661 | 388 | 2527 | 1851 | 384 | 2151 | 1023 | 168 | 771 | 832 | 115 | 794 |
| PBANKA_0521500 | 14 | 0 | 13 | 0 | 67 | 21 | 59 | 26 | 43 | 30 | 13 | 0 | 90 | 52 | 45 | 198 | 133 | 58 | 2319 | 1447 | 48 | 293 |
| PBANKA_0521600 | 11 | 17 | 128 | 15 | 209 | 67 | 251 | 79 | 85 | 34 | 19 | 6 | 810 | 217 | 721 | 3002 | 847 | 168 | 681 | 329 | 55 | 383 |
| PBANKA_0521700 | 6411 | 10023 | 503 | 153 | 776 | 353 | 737 | 261 | 3437 | 1766 | 2425 | 1199 | 3684 | 7535 | 1748 | 12176 | 6731 | 1661 | 2099 | 5208 | 3603 | 28112 |
| PBANKA_0521800 | 0 | 0 | 32 | 10 | 70 | 31 | 96 | 34 | 77 | 24 | 10 | 0 | 201 | 76 | 273 | 1309 | 546 | 67 | 389 | 221 | 15 | 116 |
| PBANKA_0521900 | 0 | 5 | 8 | 5 | 22 | 17 | 51 | 20 | 29 | 11 | 0 | 0 | 157 | 25 | 123 | 302 | 241 | 123 | 1062 | 407 | 73 | 378 |
| PBANKA_0522000 | 15 | 11 | 86 | 10 | 127 | 25 | 229 | 73 | 149 | 70 | 23 | 13 | 1503 | 317 | 383 | 2019 | 5981 | 1015 | 2361 | 2078 | 342 | 1989 |
| PBANKA_0522400 | 11 | 0 | 119 | 17 | 252 | 174 | 200 | 92 | 42 | 40 | 22 | 0 | 81 | 19 | 61 | 109 | 122 | 50 | 1356 | 590 | 459 | 3591 |
| PBANKA_0522500 | 0 | 0 | 76 | 5 | 102 | 35 | 106 | 24 | 32 | 20 | 22 | 10 | 1573 | 585 | 331 | 2074 | 1270 | 173 | 909 | 765 | 189 | 1162 |
| PBANKA_0522600 | 303 | 403 | 248 | 64 | 844 | 349 | 932 | 302 | 531 | 269 | 424 | 260 | 4303 | 6005 | 3441 | 14413 | 10100 | 1861 | 2438 | 8563 | 1367 | 9324 |
| PBANKA_0522700 | 0 | 0 | 56 | 17 | 405 | 372 | 827 | 500 | 262 | 253 | 37 | 19 | 3356 | 904 | 2763 | 9155 | 13768 | 2262 | 21045 | 9524 | 344 | 4051 |
| PBANKA_0522800 | 102 | 88 | 1601 | 416 | 935 | 586 | 1214 | 558 | 404 | 273 | 1876 | 1494 | 13397 | 3647 | 2545 | 9227 | 3925 | 793 | 2817 | 997 | 643 | 4260 |
| PBANKA_0522900 | 94 | 123 | 412 | 73 | 721 | 232 | 948 | 302 | 480 | 208 | 335 | 181 | 3391 | 7127 | 2695 | 13806 | 9381 | 1212 | 1304 | 3738 | 521 | 3023 |
| PBANKA_0523000 | 0 | 8 | 23 | 0 | 69 | 27 | 118 | 26 | 31 | 7 | 12 | 0 | 138 | 134 | 119 | 367 | 245 | 45 | 197 | 97 | 24 | 116 |
| PBANKA_0523100 | 28 | 46 | 68 | 16 | 185 | 106 | 227 | 86 | 80 | 46 | 132 | 87 | 1111 | 639 | 379 | 1931 | 732 | 76 | 543 | 386 | 73 | 668 |
| PBANKA_0523200 | 6 | 12 | 16 | 0 | 54 | 33 | 95 | 29 | 40 | 28 | 101 | 66 | 236 | 43 | 218 | 659 | 553 | 95 | 196 | 155 | 150 | 1025 |
| PBANKA_0523300 | 0 | 0 | 90 | 22 | 248 | 109 | 213 | 108 | 60 | 47 | 11 | 0 | 275 | 35 | 265 | 1052 | 572 | 109 | 586 | 350 | 14 | 129 |
| PBANKA_0523400 | 0 | 7 | 40 | 0 | 121 | 31 | 138 | 40 | 78 | 31 | 13 | 0 | 548 | 185 | 359 | 1505 | 1538 | 301 | 700 | 575 | 102 | 571 |
| PBANKA_0523500 | 0 | 0 | 41 | 8 | 66 | 34 | 120 | 16 | 45 | 17 | 0 | 0 | 211 | 45 | 146 | 434 | 250 | 49 | 160 | 103 | 8 | 66 |
| PBANKA_0523600 | 0 | 0 | 71 | 8 | 154 | 76 | 179 | 60 | 41 | 20 | 0 | 0 | 433 | 90 | 136 | 814 | 499 | 96 | 2574 | 1074 | 42 | 298 |
| PBANKA_0523900 | 81 | 92 | 14 | 17 | 183 | 126 | 1090 | 472 | 2290 | 1409 | 324 | 168 | 4925 | 1842 | 207 | 459 | 27839 | 6722 | 73999 | 64436 | 786 | 8806 |
| PBANKA_0524000 | 29 | 31 | 116 | 19 | 297 | 107 | 390 | 128 | 204 | 68 | 82 | 22 | 1522 | 915 | 777 | 3431 | 3853 | 667 | 991 | 1305 | 421 | 2615 |
| PBANKA_0524300 | 5 | 9 | 25 | 10 | 156 | 91 | 391 | 229 | 368 | 252 | 3024 | 2705 | 30341 | 14461 | 2496 | 13936 | 25966 | 2723 | 6363 | 4627 | 284 | 3053 |
| PBANKA_0524400 | 0 | 0 | 20 | 0 | 79 | 60 | 79 | 38 | 38 | 27 | 203 | 221 | 1548 | 856 | 291 | 1403 | 853 | 112 | 367 | 218 | 14 | 130 |
| PBANKA_0524600 | 0 | 0 | 16 | 5 | 61 | 35 | 47 | 25 | 129 | 70 | 22 | 15 | 362 | 174 | 196 | 720 | 544 | 217 | 993 | 461 | 73 | 487 |
| PBANKA_0524700 | 0 | 0 | 183 | 69 | 284 | 152 | 513 | 244 | 360 | 169 | 1986 | 1396 | 11049 | 16409 | 1693 | 14845 | 9478 | 989 | 3430 | 6956 | 503 | 3621 |
| PBANKA_0524800 | 0 | 0 | 7 | 6 | 54 | 34 | 88 | 78 | 125 | 167 | 937 | 989 | 3639 | 2535 | 2436 | 4561 | 3208 | 1262 | 1190 | 440 | 28 | 307 |
| PBANKA_0600091 | 0 | 0 | 6 | 0 | 0 | 0 | 6 | 6 | 0 | 11 | 58 | 48 | 297 | 115 | 59 | 246 | 94 | 17 | 21 | 17 | 0 | 24 |
| PBANKA_0600400 | 0 | 0 | 6 | 0 | 0 | 0 | 5 | 8 | 6 | 12 | 0 | 0 | 693 | 224 | 398 | 1042 | 382 | 65 | 634 | 395 | 17 | 198 |
| PBANKA_0600500 | 50 | 67 | 130 | 24 | 173 | 82 | 156 | 64 | 72 | 34 | 133 | 131 | 1240 | 885 | 257 | 1127 | 616 | 84 | 454 | 387 | 177 | 1428 |
| PBANKA_0600600 | 0 | 10 | 17 | 0 | 48 | 18 | 61 | 29 | 85 | 24 | 8 | 0 | 4777 | 1103 | 32 | 432 | 985 | 200 | 67677 | 55529 | 264 | 1492 |
| PBANKA_0600800 | 0 | 0 | 33 | 0 | 65 | 34 | 85 | 46 | 69 | 23 | 10 | 0 | 663 | 190 | 190 | 1345 | 1324 | 223 | 1147 | 1389 | 75 | 833 |
| PBANKA_0600900 | 21 | 21 | 10 | 0 | 45 | 29 | 61 | 35 | 27 | 23 | 0 | 7 | 348 | 74 | 91 | 323 | 810 | 227 | 2269 | 793 | 43 | 328 |
| PBANKA_0601000 | 13 | 31 | 56 | 8 | 143 | 63 | 181 | 39 | 55 | 25 | 6 | 0 | 820 | 178 | 182 | 1335 | 932 | 130 | 1763 | 1209 | 151 | 1005 |
| PBANKA_0601100 | 0 | 0 | 240 | 56 | 506 | 291 | 558 | 336 | 206 | 138 | 20 | 7 | 1025 | 170 | 862 | 3405 | 3063 | 400 | 6374 | 4247 | 197 | 1917 |
| PBANKA_0601200 | 0 | 0 | 44 | 9 | 82 | 41 | 82 | 30 | 84 | 23 | 18 | 9 | 283 | 438 | 79 | 240 | 860 | 214 | 2084 | 6597 | 76 | 458 |
| PBANKA_0601300 | 14 | 8 | 118 | 41 | 118 | 71 | 80 | 37 | 18 | 19 | 12 | 6 | 130 | 15 | 217 | 453 | 143 | 25 | 538 | 325 | 83 | 672 |
| PBANKA_0601500 | 6 | 8 | 77 | 15 | 150 | 49 | 201 | 63 | 115 | 43 | 21 | 7 | 703 | 313 | 253 | 1130 | 1047 | 312 | 2521 | 1131 | 209 | 1218 |
| PBANKA_0601600 | 398 | 567 | 275 | 68 | 1044 | 501 | 1433 | 579 | 805 | 445 | 194 | 70 | 2236 | 1678 | 3644 | 19144 | 21384 | 2673 | 1851 | 7336 | 2770 | 18586 |
| PBANKA_0601700 | 67 | 105 | 115 | 27 | 186 | 83 | 136 | 74 | 79 | 56 | 796 | 571 | 1500 | 1481 | 368 | 1462 | 188 | 79 | 467 | 320 | 356 | 1984 |
| PBANKA_0601800 | 0 | 0 | 14 | 0 | 29 | 8 | 32 | 7 | 0 | 5 | 0 | 0 | 62 | 10 | 57 | 98 | 43 | 18 | 108 | 58 | 35 | 204 |
| PBANKA_0601900 | 187 | 206 | 449 | 112 | 472 | 299 | 935 | 514 | 739 | 420 | 61 | 38 | 4252 | 867 | 896 | 3489 | 27822 | 5827 | 3746 | 1985 | 137 | 930 |
| PBANKA_0602000 | 278 | 448 | 116 | 22 | 512 | 215 | 582 | 253 | 240 | 113 | 59 | 25 | 2095 | 711 | 615 | 2014 | 5756 | 1039 | 6769 | 6416 | 307 | 2273 |
| PBANKA_0602100 | 0 | 0 | 41 | 12 | 77 | 31 | 117 | 34 | 50 | 13 | 20 | 13 | 470 | 161 | 213 | 1108 | 338 | 75 | 710 | 359 | 113 | 544 |
| PBANKA_0602200 | 0 | 0 | 25 | 9 | 44 | 13 | 38 | 26 | 15 | 6 | 0 | 0 | 249 | 118 | 130 | 283 | 140 | 26 | 214 | 107 | 38 | 209 |
| PBANKA_0602300 | 19 | 34 | 73 | 10 | 207 | 52 | 258 | 90 | 165 | 64 | 36 | 15 | 538 | 359 | 542 | 2711 | 2387 | 303 | 1422 | 1711 | 352 | 2003 |
| PBANKA_0602400 | 165 | 261 | 55 | 14 | 244 | 81 | 540 | 218 | 385 | 186 | 45 | 21 | 1037 | 265 | 510 | 2257 | 5441 | 933 | 3933 | 2598 | 452 | 3126 |
| PBANKA_0602500 | 24 | 14 | 101 | 16 | 69 | 37 | 57 | 22 | 38 | 10 | 163 | 109 | 997 | 729 | 115 | 550 | 93 | 25 | 816 | 424 | 392 | 2298 |
| PBANKA_0602600 | 0 | 5 | 46 | 6 | 112 | 53 | 93 | 34 | 38 | 17 | 28 | 25 | 165 | 109 | 98 | 333 | 37 | 16 | 188 | 76 | 245 | 1424 |
| PBANKA_0602700 | 0 | 0 | 782 | 236 | 1023 | 582 | 1291 | 596 | 372 | 232 | 64 | 29 | 5327 | 1285 | 2885 | 13165 | 6070 | 809 | 4025 | 2447 | 618 | 5600 |
| PBANKA_0602800 | 5 | 14 | 328 | 145 | 408 | 298 | 665 | 284 | 296 | 166 | 763 | 658 | 1073 | 542 | 864 | 1268 | 1096 | 470 | 6397 | 2767 | 305 | 1796 |
| PBANKA_0603000 | 0 | 0 | 57 | 18 | 188 | 74 | 167 | 70 | 94 | 34 | 15 | 12 | 518 | 280 | 458 | 2720 | 816 | 137 | 5391 | 3919 | 611 | 3206 |
| PBANKA_0603100 | 0 | 0 | 23 | 6 | 43 | 19 | 31 | 13 | 14 | 8 | 0 | 0 | 67 | 30 | 87 | 242 | 81 | 25 | 531 | 165 | 116 | 646 |
| PBANKA_0603200 | 40 | 30 | 1610 | 499 | 1027 | 666 | 941 | 520 | 324 | 276 | 3384 | 3399 | 5255 | 1669 | 2383 | 7232 | 2033 | 320 | 857 | 355 | 33 | 404 |
| PBANKA_0603300 | 0 | 0 | 37 | 12 | 105 | 81 | 101 | 61 | 42 | 13 | 8 | 0 | 467 | 220 | 17 | 40 | 187 | 108 | 10451 | 5483 | 164 | 1069 |
| PBANKA_0603400 | 0 | 0 | 49 | 10 | 89 | 53 | 145 | 45 | 45 | 21 | 49 | 19 | 1028 | 481 | 353 | 1295 | 1022 | 136 | 966 | 508 | 1659 | 8652 |
| PBANKA_0603500 | 0 | 0 | 37 | 6 | 96 | 43 | 91 | 35 | 38 | 25 | 20 | 18 | 334 | 214 | 102 | 511 | 274 | 32 | 113 | 75 | 1074 | 6866 |
| PBANKA_0603600 | 0 | 0 | 17 | 8 | 42 | 36 | 28 | 16 | 18 | 10 | 6 | 0 | 73 | 25 | 225 | 438 | 91 | 14 | 159 | 60 | 38 | 193 |
| PBANKA_0603700 | 77 | 84 | 39 | 12 | 144 | 37 | 271 | 113 | 203 | 85 | 28 | 6 | 1149 | 258 | 306 | 887 | 3689 | 858 | 6592 | 2305 | 241 | 1663 |
| PBANKA_0603800 | 0 | 0 | 130 | 20 | 100 | 51 | 92 | 39 | 33 | 14 | 0 | 12 | 1147 | 563 | 480 | 2230 | 654 | 82 | 73 | 72 | 26 | 133 |

| | | | | | | | | | | | | | | | | | | | | | | |
|---|---|---|---|---|---|---|---|---|---|---|---|---|---|---|---|---|---|---|---|---|---|---|
| PBANKA_0604000 | 0 | 0 | 57 | 0 | 153 | 65 | 148 | 68 | 44 | 33 | 0 | 0 | 227 | 76 | 232 | 723 | 853 | 163 | 188 | 132 | 358 | 2070 |
| PBANKA_0604100 | 24 | 21 | 60 | 14 | 325 | 154 | 603 | 217 | 459 | 166 | 292 | 143 | 1264 | 423 | 696 | 4228 | 3447 | 430 | 1620 | 1417 | 268 | 2080 |
| PBANKA_0604200 | 36 | 76 | 107 | 22 | 432 | 170 | 920 | 331 | 1143 | 563 | 3079 | 1563 | 4322 | 2563 | 1387 | 8544 | 7084 | 1030 | 4674 | 4524 | 768 | 6543 |
| PBANKA_0604300 | 7 | 0 | 528 | 147 | 653 | 373 | 976 | 612 | 401 | 228 | 83 | 28 | 2267 | 599 | 3039 | 8884 | 7422 | 1378 | 8141 | 3829 | 587 | 4185 |
| PBANKA_0604400 | 0 | 0 | 10 | 0 | 50 | 19 | 66 | 24 | 29 | 15 | 8 | 0 | 189 | 45 | 126 | 413 | 277 | 51 | 2351 | 893 | 86 | 707 |
| PBANKA_0604500 | 660 | 1093 | 917 | 172 | 1956 | 862 | 2940 | 972 | 2219 | 893 | 7827 | 4239 | 10033 | 21632 | 4789 | 28274 | 31233 | 5092 | 1868 | 5913 | 4125 | 28740 |
| PBANKA_0604600 | 80 | 89 | 121 | 24 | 398 | 166 | 559 | 184 | 361 | 144 | 89 | 22 | 1555 | 1267 | 746 | 4437 | 5901 | 655 | 4715 | 7681 | 749 | 5581 |
| PBANKA_0604700 | 0 | 0 | 79 | 14 | 131 | 73 | 177 | 76 | 55 | 32 | 0 | 0 | 290 | 74 | 353 | 1414 | 1240 | 142 | 671 | 468 | 131 | 1118 |
| PBANKA_0604800 | 86 | 103 | 918 | 280 | 1062 | 461 | 870 | 302 | 287 | 148 | 1626 | 927 | 7909 | 8320 | 3209 | 17981 | 3346 | 387 | 1231 | 2320 | 149 | 1225 |
| PBANKA_0604900 | 1392 | 2157 | 45 | 7 | 370 | 170 | 488 | 168 | 905 | 437 | 195 | 93 | 438 | 1406 | 105 | 615 | 27042 | 4348 | 445 | 2420 | 11062 | 69717 |
| PBANKA_0605000 | 34 | 42 | 6 | 0 | 45 | 14 | 20 | 20 | 29 | 14 | 0 | 0 | 98 | 23 | 31 | 83 | 392 | 102 | 467 | 128 | 636 | 2681 |
| PBANKA_0605200 | 99 | 115 | 251 | 56 | 500 | 250 | 554 | 172 | 293 | 138 | 1199 | 762 | 5634 | 3485 | 1617 | 8610 | 4582 | 682 | 1076 | 1575 | 137 | 1047 |
| PBANKA_0605300 | 93 | 85 | 257 | 43 | 373 | 171 | 326 | 91 | 151 | 75 | 197 | 102 | 2444 | 2714 | 1532 | 8400 | 2135 | 383 | 1448 | 3000 | 260 | 2272 |
| PBANKA_0605400 | 5 | 0 | 31 | 5 | 99 | 36 | 89 | 42 | 29 | 12 | 0 | 16 | 765 | 317 | 425 | 1607 | 478 | 74 | 508 | 276 | 20 | 167 |
| PBANKA_0605600 | 157 | 288 | 230 | 52 | 1297 | 466 | 2685 | 1037 | 1517 | 600 | 224 | 98 | 3048 | 4905 | 2001 | 15332 | 43568 | 5106 | 3366 | 38350 | 2380 | 11996 |
| PBANKA_0605700 | 39 | 45 | 235 | 43 | 281 | 116 | 247 | 70 | 226 | 133 | 2157 | 1390 | 5997 | 6101 | 1432 | 12748 | 4419 | 622 | 2692 | 4793 | 503 | 3934 |
| PBANKA_0605900 | 1753 | 2267 | 11 | 0 | 46 | 30 | 39 | 17 | 36 | 30 | 20 | 10 | 178 | 94 | 68 | 189 | 206 | 47 | 1269 | 1013 | 54 | 341 |
| PBANKA_0606000 | 0 | 0 | 37 | 6 | 94 | 31 | 97 | 41 | 29 | 11 | 6 | 0 | 153 | 29 | 112 | 432 | 283 | 60 | 227 | 106 | 0 | 67 |
| PBANKA_0606100 | 0 | 9 | 70 | 10 | 142 | 62 | 115 | 41 | 53 | 21 | 67 | 21 | 558 | 442 | 236 | 1054 | 470 | 77 | 246 | 272 | 14 | 118 |
| PBANKA_0606300 | 44 | 67 | 25 | 6 | 106 | 61 | 199 | 95 | 124 | 72 | 53 | 33 | 690 | 600 | 153 | 1565 | 1840 | 159 | 3120 | 7947 | 119 | 594 |
| PBANKA_0606400 | 23 | 23 | 46 | 7 | 121 | 66 | 123 | 53 | 61 | 25 | 15 | 8 | 345 | 83 | 85 | 610 | 306 | 33 | 1332 | 743 | 43 | 293 |
| PBANKA_0606500 | 0 | 0 | 17 | 0 | 64 | 32 | 47 | 28 | 19 | 13 | 0 | 0 | 58 | 13 | 58 | 139 | 161 | 26 | 215 | 111 | 11 | 146 |
| PBANKA_0606600 | 258 | 419 | 616 | 147 | 732 | 301 | 661 | 194 | 355 | 169 | 1826 | 1195 | 2369 | 6012 | 1070 | 5771 | 1201 | 155 | 801 | 2492 | 259 | 1795 |
| PBANKA_0606700 | 34 | 64 | 9 | 0 | 166 | 73 | 195 | 74 | 219 | 95 | 30 | 9 | 111 | 138 | 128 | 295 | 310 | 76 | 623 | 929 | 41 | 315 |
| PBANKA_0606800 | 0 | 0 | 173 | 21 | 132 | 86 | 159 | 62 | 38 | 29 | 0 | 6 | 838 | 353 | 330 | 1264 | 460 | 68 | 469 | 360 | 15 | 162 |
| PBANKA_0606900 | 0 | 0 | 15 | 0 | 591 | 284 | 640 | 219 | 227 | 98 | 19 | 20 | 266 | 135 | 470 | 2085 | 883 | 124 | 690 | 970 | 37 | 200 |
| PBANKA_0607000 | 0 | 0 | 108 | 36 | 482 | 197 | 463 | 137 | 187 | 67 | 29 | 20 | 176 | 133 | 79 | 273 | 362 | 105 | 2205 | 3849 | 216 | 1747 |
| PBANKA_0607100 | 0 | 0 | 40 | 12 | 108 | 74 | 92 | 39 | 35 | 20 | 0 | 0 | 58 | 6 | 111 | 234 | 87 | 28 | 250 | 86 | 38 | 254 |
| PBANKA_0607200 | 0 | 0 | 42 | 7 | 92 | 26 | 59 | 22 | 36 | 10 | 8 | 0 | 305 | 118 | 157 | 575 | 112 | 26 | 767 | 192 | 20 | 123 |
| PBANKA_0607300 | 255 | 391 | 68 | 20 | 702 | 212 | 730 | 197 | 414 | 135 | 1171 | 626 | 2074 | 1465 | 669 | 3508 | 1632 | 211 | 2947 | 2208 | 311 | 2398 |
| PBANKA_0607400 | 149 | 222 | 311 | 108 | 1833 | 835 | 2648 | 922 | 1255 | 509 | 236 | 143 | 3443 | 1183 | 1265 | 5727 | 17905 | 2751 | 8250 | 6525 | 380 | 2744 |
| PBANKA_0607500 | 0 | 0 | 31 | 7 | 61 | 23 | 26 | 15 | 17 | 13 | 0 | 0 | 234 | 42 | 237 | 851 | 93 | 35 | 199 | 57 | 11 | 52 |
| PBANKA_0607600 | 0 | 0 | 68 | 22 | 168 | 82 | 184 | 107 | 92 | 54 | 15 | 0 | 304 | 125 | 347 | 783 | 286 | 92 | 1602 | 449 | 29 | 124 |
| PBANKA_0607700 | 215 | 280 | 726 | 148 | 1646 | 570 | 3082 | 1273 | 2078 | 826 | 387 | 190 | 9748 | 5349 | 2991 | 19706 | 67537 | 8312 | 11692 | 21786 | 1410 | 8170 |
| PBANKA_0607800 | 30 | 38 | 23 | 9 | 179 | 88 | 540 | 181 | 547 | 269 | 145 | 66 | 965 | 237 | 136 | 477 | 3254 | 891 | 2801 | 1814 | 300 | 2358 |
| PBANKA_0607900 | 102 | 124 | 7 | 0 | 25 | 15 | 26 | 7 | 46 | 20 | 0 | 0 | 128 | 49 | 86 | 345 | 175 | 55 | 1391 | 753 | 39 | 283 |
| PBANKA_0608000 | 199 | 140 | 15 | 5 | 68 | 37 | 55 | 42 | 36 | 21 | 0 | 0 | 88 | 18 | 30 | 75 | 106 | 48 | 1833 | 865 | 1683 | 10929 |
| PBANKA_0608100 | 224 | 267 | 40 | 7 | 121 | 58 | 263 | 93 | 133 | 78 | 10 | 9 | 502 | 77 | 144 | 640 | 1574 | 289 | 391 | 267 | 47 | 264 |
| PBANKA_0608200 | 137 | 172 | 72 | 16 | 156 | 78 | 259 | 112 | 151 | 75 | 11 | 5 | 462 | 59 | 238 | 1000 | 1612 | 397 | 1276 | 599 | 151 | 1070 |
| PBANKA_0608300 | 6 | 0 | 63 | 12 | 195 | 76 | 225 | 90 | 168 | 110 | 155 | 59 | 822 | 404 | 320 | 1401 | 841 | 228 | 1842 | 870 | 106 | 485 |
| PBANKA_0608400 | 0 | 0 | 88 | 8 | 181 | 44 | 141 | 63 | 50 | 19 | 35 | 14 | 1284 | 590 | 406 | 1532 | 633 | 89 | 1349 | 556 | 54 | 441 |
| PBANKA_0608500 | 46 | 55 | 19 | 0 | 27 | 17 | 57 | 16 | 58 | 26 | 0 | 7 | 540 | 159 | 70 | 391 | 1114 | 306 | 2421 | 1474 | 630 | 4191 |
| PBANKA_0608600 | 16 | 11 | 12 | 0 | 18 | 12 | 20 | 0 | 10 | 5 | 5 | 16 | 3870 | 2350 | 152 | 628 | 1402 | 608 | 1E+05 | 1E+05 | 8967 | 76119 |
| PBANKA_0608700 | 7 | 14 | 46 | 7 | 61 | 40 | 61 | 21 | 21 | 7 | 14 | 8 | 422 | 175 | 163 | 746 | 112 | 34 | 381 | 322 | 74 | 576 |
| PBANKA_0608900 | 23 | 14 | 67 | 11 | 172 | 108 | 265 | 89 | 108 | 62 | 37 | 5 | 234 | 84 | 588 | 2441 | 880 | 123 | 786 | 468 | 49 | 406 |
| PBANKA_0609000 | 0 | 0 | 16 | 5 | 89 | 32 | 121 | 39 | 22 | 17 | 10 | 12 | 98 | 38 | 183 | 1015 | 293 | 40 | 537 | 266 | 17 | 113 |
| PBANKA_0609100 | 155 | 277 | 130 | 28 | 328 | 108 | 300 | 115 | 476 | 239 | 2780 | 1857 | 3273 | 3228 | 544 | 3845 | 1875 | 363 | 1325 | 1853 | 1309 | 9366 |
| PBANKA_0609200 | 24 | 36 | 24 | 0 | 82 | 45 | 133 | 58 | 52 | 27 | 9 | 0 | 359 | 102 | 128 | 566 | 665 | 116 | 1176 | 616 | 175 | 1326 |
| PBANKA_0609300 | 41 | 64 | 43 | 0 | 118 | 80 | 298 | 93 | 109 | 34 | 12 | 5 | 387 | 71 | 171 | 610 | 746 | 205 | 738 | 277 | 94 | 487 |
| PBANKA_0609500 | 996 | 1340 | 6 | 0 | 38 | 27 | 49 | 26 | 157 | 75 | 16 | 0 | 2155 | 2902 | 141 | 460 | 2069 | 741 | 33103 | 1E+05 | 7124 | 48771 |
| PBANKA_0609600 | 12 | 19 | 5 | 0 | 13 | 8 | 13 | 0 | 6 | 0 | 0 | 7 | 19 | 0 | 18 | 18 | 23 | 11 | 60 | 33 | 4672 | 32530 |
| PBANKA_0609700 | 34 | 56 | 125 | 29 | 365 | 132 | 468 | 167 | 178 | 106 | 191 | 105 | 2354 | 1471 | 892 | 4887 | 2807 | 379 | 964 | 1752 | 46 | 339 |
| PBANKA_0609800 | 0 | 0 | 29 | 5 | 179 | 76 | 253 | 92 | 101 | 38 | 28 | 0 | 288 | 96 | 182 | 609 | 851 | 150 | 873 | 575 | 23 | 172 |
| PBANKA_0609900 | 123 | 144 | 442 | 163 | 553 | 404 | 722 | 461 | 265 | 180 | 552 | 562 | 2001 | 661 | 889 | 2717 | 2160 | 407 | 1906 | 876 | 184 | 1618 |
| PBANKA_0610000 | 0 | 0 | 43 | 13 | 171 | 68 | 236 | 97 | 136 | 61 | 40 | 18 | 1548 | 730 | 404 | 1892 | 1306 | 257 | 1474 | 905 | 109 | 729 |
| PBANKA_0610100 | 77 | 82 | 309 | 66 | 1042 | 337 | 1281 | 458 | 766 | 315 | 1008 | 530 | 3819 | 9547 | 2961 | 17556 | 16337 | 2186 | 1488 | 7406 | 1265 | 7000 |
| PBANKA_0610200 | 63 | 70 | 166 | 64 | 598 | 457 | 1731 | 1091 | 934 | 660 | 108 | 33 | 1764 | 375 | 508 | 1105 | 8607 | 1378 | 7328 | 3477 | 379 | 4120 |
| PBANKA_0610300 | 249 | 302 | 338 | 83 | 944 | 493 | 1707 | 789 | 1117 | 846 | 146 | 113 | 2065 | 359 | 1689 | 3249 | 7211 | 2665 | 2757 | 1084 | 2003 | 16587 |
| PBANKA_0610400 | 109 | 145 | 439 | 143 | 754 | 428 | 598 | 200 | 2592 | 1592 | 1955 | 1424 | 11754 | 10661 | 3784 | 20094 | 3265 | 511 | 2545 | 3772 | 817 | 8219 |
| PBANKA_0610500 | 8 | 6 | 85 | 11 | 125 | 81 | 275 | 141 | 282 | 161 | 697 | 571 | 930 | 256 | 394 | 956 | 1121 | 420 | 841 | 274 | 69 | 544 |
| PBANKA_0610600 | 5 | 0 | 93 | 18 | 158 | 53 | 227 | 77 | 155 | 85 | 37 | 15 | 1104 | 420 | 352 | 1505 | 1656 | 301 | 1992 | 1171 | 91 | 685 |
| PBANKA_0610800 | 36 | 64 | 210 | 47 | 476 | 160 | 501 | 177 | 209 | 93 | 39 | 18 | 4769 | 1858 | 1098 | 5813 | 3142 | 481 | 1603 | 1287 | 1276 | 8871 |
| PBANKA_0610900 | 132 | 204 | 1036 | 267 | 762 | 348 | 546 | 233 | 270 | 128 | 1631 | 1104 | 12361 | 7490 | 4210 | 23276 | 3875 | 499 | 2279 | 2395 | 451 | 3701 |
| PBANKA_0611000 | 29 | 35 | 35 | 0 | 83 | 37 | 131 | 60 | 68 | 33 | 17 | 22 | 389 | 84 | 195 | 1061 | 628 | 77 | 317 | 212 | 186 | 1435 |
| PBANKA_0611100 | 7 | 18 | 46 | 11 | 229 | 86 | 353 | 147 | 229 | 75 | 59 | 23 | 631 | 233 | 816 | 3428 | 2368 | 336 | 760 | 882 | 378 | 2170 |
| PBANKA_0611200 | 0 | 0 | 61 | 15 | 82 | 32 | 74 | 26 | 30 | 6 | 11 | 0 | 81 | 27 | 234 | 676 | 124 | 38 | 220 | 125 | 708 | 3020 |
| PBANKA_0611300 | 0 | 0 | 62 | 15 | 74 | 39 | 75 | 29 | 82 | 57 | 15 | 0 | 104 | 35 | 149 | 494 | 331 | 146 | 1550 | 932 | 428 | 2237 |
| PBANKA_0611400 | 0 | 0 | 93 | 20 | 168 | 72 | 195 | 60 | 262 | 134 | 54 | 20 | 244 | 71 | 332 | 1634 | 1395 | 244 | 505 | 554 | 410 | 3350 |
| PBANKA_0611500 | 0 | 0 | 5 | 0 | 12 | 6 | 66 | 17 | 230 | 187 | 55 | 19 | 244 | 49 | 8 | 32 | 1174 | 617 | 698 | 293 | 92 | 599 |
| PBANKA_0611600 | 122 | 158 | 489 | 212 | 1378 | 758 | 2404 | 1071 | 1350 | 628 | 154 | 59 | 6636 | 2547 | 1080 | 5710 | 23419 | 3499 | 74944 | 82542 | 5716 | 49398 |
| PBANKA_0611700 | 0 | 0 | 9 | 0 | 120 | 48 | 242 | 99 | 77 | 43 | 16 | 5 | 207 | 30 | 40 | 61 | 612 | 111 | 540 | 182 | 57 | 483 |
| PBANKA_0611800 | 339 | 516 | 188 | 49 | 452 | 188 | 684 | 292 | 242 | 131 | 73 | 62 | 2031 | 1096 | 559 | 2569 | 3257 | 505 | 1788 | 1105 | 919 | 6503 |
| PBANKA_0611900 | 0 | 0 | 57 | 0 | 149 | 47 | 169 | 57 | 57 | 35 | 19 | 6 | 170 | 60 | 141 | 499 | 512 | 83 | 1334 | 652 | 84 | 543 |
| PBANKA_0612000 | 0 | 0 | 53 | 11 | 76 | 38 | 60 | 31 | 30 | 10 | 0 | 0 | 41 | 12 | 133 | 612 | 143 | 20 | 131 | 89 | 10 | 65 |
| PBANKA_0612100 | 6 | 7 | 57 | 8 | 103 | 42 | 236 | 86 | 106 | 62 | 18 | 8 | 259 | 116 | 489 | 2575 | 721 | 128 | 1544 | 731 | 150 | 971 |
| PBANKA_0612200 | 190 | 304 | 177 | 39 | 613 | 326 | 1248 | 493 | 944 | 520 | 126 | 62 | 1105 | 638 | 385 | 1626 | 12491 | 2486 | 2736 | 2972 | 1354 | 8800 |
| PBANKA_0612300 | 0 | 7 | 127 | 19 | 247 | 65 | 218 | 79 | 79 | 34 | 14 | 0 | 830 | 668 | 687 | 3557 | 2065 | 273 | 1807 | 1911 | 93 | 527 |
| PBANKA_0612400 | 0 | 0 | 18 | 0 | 39 | 16 | 61 | 29 | 57 | 39 | 22 | 0 | 5712 | 4879 | 165 | 690 | 3833 | 1441 | 1E+05 | 2E+05 | 2429 | 16291 |
| PBANKA_0612500 | 9 | 9 | 31 | 5 | 97 | 40 | 114 | 28 | 37 | 10 | 0 | 0 | 114 | 17 | 160 | 435 | 242 | 46 | 215 | 115 | 63 | 299 |
| PBANKA_0612600 | 13 | 24 | 54 | 7 | 76 | 37 | 133 | 40 | 139 | 61 | 52 | 10 | 1576 | 629 | 386 | 2051 | 1224 | 266 | 2642 | 1125 | 88 | 500 |
| PBANKA_0612700 | 42 | 55 | 16 | 0 | 98 | 50 | 173 | 86 | 111 | 52 | 50 | 38 | 335 | 87 | 91 | 420 | 903 | 191 | 1392 | 1166 | 233 | 1182 |
| PBANKA_0612800 | 9 | 0 | 55 | 7 | 165 | 68 | 211 | 86 | 144 | 68 | 16 | 9 | 1081 | 530 | 327 | 1600 | 1722 | 266 | 1127 | 1096 | 212 | 1656 |
| PBANKA_0613000 | 8 | 10 | 47 | 0 | 175 | 63 | 300 | 99 | 141 | 46 | 23 | 0 | 372 | 158 | 192 | 1049 | 3222 | 336 | 709 | 400 | 476 | 2487 |
| PBANKA_0613100 | 77 | 91 | 133 | 14 | 852 | 283 | 2849 | 876 | 1863 | 647 | 266 | 92 | 895 | 427 | 1286 | 5221 | 6472 | 1006 | 3061 | 2641 | 585 | 3351 |
| PBANKA_0613200 | 38 | 48 | 178 | 38 | 788 | 305 | 1336 | 627 | 728 | 346 | 108 | 36 | 1036 | 561 | 676 | 2125 | 10339 | 1348 | 2469 | 3046 | 724 | 4654 |
| PBANKA_0613300 | 0 | 13 | 20 | 6 | 151 | 69 | 189 | 105 | 94 | 78 | 24 | 18 | 1270 | 298 | 577 | 2399 | 5806 | 964 | 1460 | 1179 | 205 | 1315 |
| PBANKA_0613400 | 0 | 0 | 104 | 31 | 182 | 115 | 160 | 101 | 60 | 50 | 26 | 12 | 604 | 90 | 349 | 1808 | 308 | 43 | 139 | 76 | 183 | 1245 |
| PBANKA_0613500 | 0 | 0 | 26 | 12 | 91 | 38 | 104 | 21 | 41 | 25 | 15 | 5 | 594 | 219 | 243 | 927 | 508 | 74 | 947 | 410 | 355 | 1866 |
| PBANKA_0613600 | 0 | 0 | 30 | 0 | 134 | 63 | 155 | 59 | 83 | 27 | 8 | 15 | 325 | 133 | 127 | 493 | 395 | 62 | 1289 | 388 | 152 | 749 |
| PBANKA_0613700 | 0 | 0 | 248 | 88 | 502 | 235 | 468 | 238 | 388 | 183 | 4168 | 2843 | 6674 | 10360 | 2569 | 18338 | 6658 | 635 | 5151 | 15037 | 308 | 2148 |

| | | | | | | | | | | | | | | | | | | | | | | |
|---|---|---|---|---|---|---|---|---|---|---|---|---|---|---|---|---|---|---|---|---|---|---|
| PBANKA_0613800 | 174 | 286 | 235 | 51 | 428 | 141 | 282 | 118 | 437 | 274 | 2168 | 1360 | 3549 | 4616 | 1140 | 7959 | 2302 | 370 | 590 | 1132 | 139 | 584 |
| PBANKA_0613900 | 86 | 110 | 141 | 44 | 198 | 102 | 204 | 57 | 251 | 170 | 1236 | 838 | 1530 | 1140 | 241 | 898 | 336 | 106 | 328 | 186 | 146 | 609 |
| PBANKA_0614000 | 5 | 9 | 1040 | 276 | 2413 | 1318 | 3585 | 1708 | 956 | 607 | 76 | 52 | 7863 | 2301 | 3272 | 13573 | 19624 | 2632 | 2057 | 1127 | 90 | 757 |
| PBANKA_0614100 | 7 | 0 | 79 | 25 | 95 | 75 | 157 | 70 | 64 | 24 | 6 | 0 | 313 | 52 | 345 | 807 | 632 | 214 | 280 | 149 | 42 | 232 |
| PBANKA_0614200 | 10 | 12 | 96 | 25 | 192 | 128 | 217 | 62 | 100 | 45 | 14 | 6 | 166 | 24 | 265 | 895 | 337 | 48 | 384 | 171 | 54 | 261 |
| PBANKA_0614300 | 27 | 45 | 90 | 18 | 165 | 126 | 309 | 144 | 138 | 48 | 15 | 12 | 401 | 116 | 181 | 510 | 1533 | 378 | 206 | 135 | 26 | 117 |
| PBANKA_0614400 | 41 | 44 | 111 | 27 | 264 | 113 | 314 | 139 | 128 | 43 | 24 | 6 | 601 | 384 | 592 | 2782 | 532 | 90 | 664 | 763 | 222 | 1262 |
| PBANKA_0614500 | 45 | 49 | 423 | 145 | 377 | 288 | 414 | 183 | 135 | 87 | 277 | 304 | 2635 | 1533 | 1191 | 3128 | 629 | 133 | 543 | 340 | 109 | 754 |
| PBANKA_0614600 | 16 | 27 | 17 | 0 | 100 | 37 | 105 | 39 | 54 | 22 | 14 | 15 | 686 | 160 | 262 | 1520 | 440 | 57 | 381 | 209 | 38 | 326 |
| PBANKA_0614700 | 8 | 19 | 36 | 5 | 56 | 39 | 91 | 45 | 27 | 7 | 6 | 0 | 407 | 67 | 167 | 768 | 422 | 71 | 1763 | 713 | 16 | 141 |
| PBANKA_0614800 | 17 | 15 | 58 | 15 | 198 | 39 | 275 | 99 | 118 | 39 | 25 | 8 | 1552 | 761 | 602 | 2795 | 1808 | 369 | 3312 | 2425 | 169 | 1340 |
| PBANKA_0614900 | 159 | 214 | 77 | 19 | 265 | 88 | 387 | 133 | 115 | 70 | 20 | 12 | 1023 | 242 | 210 | 990 | 3188 | 430 | 1471 | 1007 | 130 | 818 |
| PBANKA_0615000 | 73 | 141 | 77 | 11 | 222 | 105 | 240 | 116 | 86 | 50 | 32 | 19 | 948 | 423 | 246 | 1238 | 881 | 102 | 627 | 569 | 161 | 1186 |
| PBANKA_0615100 | 20 | 37 | 191 | 41 | 247 | 77 | 202 | 62 | 134 | 80 | 925 | 543 | 1776 | 1341 | 391 | 2134 | 744 | 198 | 2204 | 675 | 211 | 1357 |
| PBANKA_0615200 | 46 | 103 | 10 | 0 | 160 | 112 | 595 | 293 | 328 | 181 | 43 | 22 | 2057 | 444 | 110 | 265 | 8626 | 1398 | 7041 | 4237 | 413 | 3486 |
| PBANKA_0615300 | 5 | 6 | 61 | 19 | 122 | 63 | 233 | 130 | 252 | 122 | 23 | 31 | 2047 | 1472 | 158 | 849 | 4674 | 1151 | 28589 | 38778 | 443 | 3018 |
| PBANKA_0615400 | 0 | 0 | 221 | 53 | 244 | 146 | 250 | 86 | 74 | 65 | 41 | 34 | 3743 | 2015 | 1439 | 6454 | 2434 | 357 | 475 | 573 | 36 | 324 |
| PBANKA_0615500 | 94 | 130 | 5 | 5 | 33 | 11 | 39 | 25 | 99 | 50 | 18 | 11 | 224 | 231 | 70 | 239 | 492 | 103 | 2354 | 5788 | 4461 | 30534 |
| PBANKA_0615600 | 0 | 0 | 85 | 20 | 109 | 84 | 98 | 64 | 23 | 13 | 5 | 0 | 418 | 129 | 213 | 880 | 236 | 23 | 383 | 350 | 207 | 1798 |
| PBANKA_0615700 | 337 | 446 | 56 | 16 | 131 | 50 | 140 | 73 | 148 | 37 | 30 | 14 | 592 | 847 | 322 | 1669 | 1976 | 254 | 1292 | 6663 | 490 | 3118 |
| PBANKA_0615800 | 197 | 299 | 104 | 28 | 712 | 282 | 983 | 347 | 462 | 159 | 66 | 39 | 655 | 676 | 1153 | 6035 | 9088 | 928 | 1788 | 6560 | 46 | 300 |
| PBANKA_0615900 | 35 | 23 | 5 | 0 | 26 | 15 | 25 | 5 | 21 | 8 | 0 | 6 | 13 | 0 | 21 | 26 | 21 | 5 | 89 | 42 | 0 | 9 |
| PBANKA_0616000 | 0 | 5 | 53 | 17 | 55 | 35 | 52 | 10 | 8 | 10 | 0 | 0 | 29 | 7 | 100 | 322 | 105 | 22 | 288 | 158 | 7 | 42 |
| PBANKA_0616100 | 0 | 6 | 157 | 34 | 192 | 111 | 158 | 80 | 50 | 29 | 0 | 9 | 548 | 128 | 420 | 973 | 361 | 82 | 494 | 243 | 39 | 328 |
| PBANKA_0616200 | 358 | 574 | 19 | 0 | 159 | 73 | 222 | 83 | 130 | 57 | 33 | 7 | 158 | 115 | 83 | 360 | 1659 | 212 | 291 | 631 | 5030 | 35621 |
| PBANKA_0616300 | 30 | 36 | 62 | 7 | 124 | 71 | 134 | 57 | 45 | 36 | 7 | 0 | 375 | 57 | 334 | 1219 | 722 | 132 | 233 | 129 | 220 | 1589 |
| PBANKA_0616400 | 0 | 0 | 14 | 0 | 16 | 13 | 24 | 10 | 7 | 6 | 0 | 0 | 119 | 16 | 37 | 164 | 81 | 39 | 1086 | 456 | 1727 | 14487 |
| PBANKA_0616500 | 0 | 0 | 70 | 12 | 128 | 76 | 150 | 76 | 45 | 22 | 9 | 8 | 342 | 63 | 241 | 692 | 395 | 63 | 927 | 584 | 2398 | 16617 |
| PBANKA_0616600 | 0 | 5 | 99 | 17 | 117 | 74 | 179 | 121 | 80 | 50 | 10 | 0 | 237 | 30 | 245 | 292 | 376 | 169 | 922 | 257 | 417 | 2144 |
| PBANKA_0616700 | 0 | 0 | 8 | 0 | 29 | 18 | 58 | 20 | 20 | 16 | 0 | 0 | 236 | 99 | 76 | 222 | 301 | 121 | 3868 | 1981 | 454 | 2984 |
| PBANKA_0616800 | 9 | 14 | 72 | 19 | 367 | 140 | 414 | 169 | 163 | 58 | 59 | 29 | 2092 | 1186 | 815 | 4883 | 2162 | 253 | 494 | 626 | 877 | 4591 |
| PBANKA_0616900 | 41 | 36 | 84 | 18 | 243 | 87 | 251 | 93 | 120 | 74 | 27 | 18 | 1386 | 740 | 474 | 2203 | 1536 | 252 | 3215 | 2222 | 225 | 1618 |
| PBANKA_0617100 | 29 | 30 | 1550 | 365 | 860 | 519 | 831 | 474 | 353 | 246 | 3057 | 2545 | 14919 | 4755 | 2694 | 10603 | 3212 | 524 | 3661 | 2159 | 221 | 1674 |
| PBANKA_0617200 | 48 | 66 | 1213 | 393 | 1084 | 633 | 1029 | 569 | 311 | 239 | 1196 | 1070 | 8662 | 2917 | 2050 | 4630 | 1536 | 455 | 1555 | 549 | 174 | 1893 |
| PBANKA_0617300 | 0 | 9 | 43 | 13 | 126 | 66 | 116 | 29 | 55 | 31 | 17 | 0 | 920 | 291 | 544 | 1953 | 2438 | 382 | 381 | 341 | 27 | 255 |
| PBANKA_0617400 | 184 | 239 | 220 | 56 | 913 | 467 | 916 | 295 | 354 | 134 | 74 | 44 | 1325 | 937 | 1741 | 7453 | 6059 | 726 | 234 | 451 | 45 | 280 |
| PBANKA_0617500 | 20 | 30 | 76 | 27 | 178 | 74 | 137 | 68 | 38 | 28 | 32 | 15 | 431 | 192 | 195 | 739 | 181 | 35 | 220 | 112 | 711 | 4704 |
| PBANKA_0617600 | 30 | 43 | 82 | 22 | 387 | 138 | 441 | 190 | 246 | 103 | 907 | 715 | 850 | 479 | 301 | 1285 | 1084 | 195 | 1057 | 652 | 759 | 3599 |
| PBANKA_0617700 | 21 | 24 | 31 | 12 | 63 | 32 | 116 | 62 | 110 | 61 | 191 | 208 | 180 | 94 | 83 | 230 | 222 | 60 | 236 | 105 | 149 | 1184 |
| PBANKA_0617900 | 0 | 0 | 94 | 37 | 194 | 107 | 225 | 81 | 143 | 71 | 31 | 10 | 1211 | 273 | 522 | 2614 | 597 | 149 | 1172 | 976 | 1649 | 10929 |
| PBANKA_0618000 | 0 | 0 | 19 | 5 | 55 | 27 | 68 | 23 | 14 | 9 | 0 | 0 | 64 | 6 | 51 | 92 | 50 | 23 | 92 | 43 | 0 | 11 |
| PBANKA_0618100 | 0 | 0 | 76 | 31 | 316 | 142 | 385 | 204 | 148 | 108 | 25 | 25 | 912 | 261 | 281 | 925 | 688 | 150 | 582 | 270 | 171 | 1215 |
| PBANKA_0618200 | 8 | 15 | 179 | 35 | 652 | 248 | 1103 | 362 | 630 | 228 | 786 | 487 | 8168 | 8638 | 3273 | 19210 | 14492 | 1879 | 3032 | 7133 | 1038 | 5802 |
| PBANKA_0618300 | 22 | 29 | 61 | 35 | 41 | 45 | 49 | 20 | 15 | 0 | 23 | 8 | 322 | 143 | 114 | 541 | 56 | 13 | 146 | 54 | 22 | 122 |
| PBANKA_0618400 | 19 | 21 | 111 | 54 | 191 | 137 | 240 | 160 | 110 | 87 | 41 | 17 | 602 | 229 | 360 | 1012 | 995 | 231 | 786 | 570 | 399 | 3371 |
| PBANKA_0618500 | 8 | 15 | 102 | 30 | 396 | 132 | 469 | 140 | 181 | 78 | 71 | 33 | 1131 | 849 | 625 | 3212 | 1605 | 253 | 3693 | 2838 | 1088 | 7117 |
| PBANKA_0618600 | 33 | 64 | 730 | 132 | 1156 | 433 | 1683 | 585 | 596 | 216 | 72 | 27 | 5311 | 1729 | 2022 | 7492 | 7614 | 1142 | 1182 | 585 | 150 | 1285 |
| PBANKA_0618700 | 8 | 14 | 26 | 7 | 122 | 60 | 169 | 41 | 51 | 35 | 46 | 12 | 358 | 197 | 261 | 1099 | 959 | 115 | 538 | 399 | 1019 | 6823 |
| PBANKA_0618800 | 37 | 79 | 111 | 21 | 201 | 69 | 228 | 101 | 143 | 64 | 495 | 311 | 963 | 1010 | 176 | 859 | 1052 | 188 | 234 | 245 | 1199 | 7748 |
| PBANKA_0618900 | 0 | 0 | 130 | 37 | 139 | 99 | 139 | 102 | 41 | 22 | 0 | 0 | 566 | 144 | 513 | 1188 | 762 | 155 | 119 | 64 | 11 | 79 |
| PBANKA_0619100 | 7 | 9 | 2325 | 858 | 1536 | 945 | 1506 | 854 | 477 | 346 | 2795 | 2380 | 10936 | 4275 | 3096 | 8691 | 2356 | 459 | 2631 | 1280 | 362 | 3041 |
| PBANKA_0619300 | 0 | 0 | 139 | 39 | 244 | 130 | 215 | 167 | 100 | 46 | 12 | 5 | 795 | 219 | 543 | 1693 | 650 | 145 | 378 | 264 | 104 | 655 |
| PBANKA_0619400 | 61 | 96 | 44 | 16 | 70 | 38 | 120 | 53 | 80 | 71 | 43 | 39 | 327 | 39 | 161 | 373 | 287 | 228 | 2103 | 446 | 197 | 1050 |
| PBANKA_0619500 | 31 | 36 | 11 | 0 | 39 | 18 | 47 | 24 | 27 | 11 | 7 | 0 | 420 | 135 | 110 | 406 | 429 | 124 | 1749 | 1240 | 416 | 2478 |
| PBANKA_0619800 | 89 | 118 | 289 | 78 | 503 | 144 | 419 | 141 | 155 | 68 | 512 | 276 | 3581 | 3966 | 888 | 4620 | 1783 | 291 | 1077 | 1939 | 376 | 2354 |
| PBANKA_0619900 | 36 | 51 | 316 | 101 | 368 | 221 | 452 | 154 | 122 | 81 | 294 | 235 | 3134 | 1123 | 1174 | 4479 | 1158 | 176 | 1295 | 615 | 109 | 864 |
| PBANKA_0620000 | 5 | 9 | 36 | 13 | 100 | 49 | 70 | 39 | 67 | 30 | 325 | 225 | 907 | 533 | 288 | 1436 | 831 | 126 | 389 | 270 | 206 | 1297 |
| PBANKA_0620100 | 227 | 393 | 137 | 35 | 251 | 86 | 261 | 110 | 273 | 115 | 1092 | 622 | 3390 | 3138 | 585 | 2695 | 2766 | 672 | 1668 | 2087 | 1090 | 6887 |
| PBANKA_0620200 | 37 | 86 | 64 | 21 | 130 | 96 | 271 | 97 | 195 | 134 | 179 | 126 | 261 | 41 | 105 | 189 | 364 | 329 | 1184 | 290 | 141 | 965 |
| PBANKA_0620300 | 0 | 0 | 7 | 0 | 40 | 34 | 50 | 36 | 89 | 45 | 22 | 16 | 1760 | 747 | 69 | 188 | 559 | 262 | 48504 | 30690 | 232 | 2559 |
| PBANKA_0620400 | 0 | 0 | 6 | 7 | 18 | 20 | 98 | 35 | 67 | 51 | 6 | 13 | 430 | 189 | 52 | 210 | 621 | 234 | 7325 | 3417 | 357 | 2675 |
| PBANKA_0620500 | 0 | 0 | 95 | 24 | 220 | 110 | 318 | 111 | 77 | 56 | 5 | 0 | 503 | 77 | 274 | 976 | 855 | 136 | 238 | 102 | 943 | 6823 |
| PBANKA_0620700 | 39 | 33 | 11 | 0 | 15 | 15 | 13 | 12 | 26 | 14 | 0 | 0 | 103 | 27 | 0 | 17 | 129 | 42 | 1947 | 1023 | 382 | 3163 |
| PBANKA_0620800 | 618 | 956 | 22 | 0 | 64 | 31 | 79 | 38 | 74 | 31 | 46 | 27 | 186 | 161 | 75 | 153 | 468 | 150 | 606 | 1306 | 1005 | 7724 |
| PBANKA_0620900 | 151 | 276 | 164 | 42 | 287 | 173 | 452 | 217 | 356 | 172 | 1546 | 916 | 3070 | 3069 | 904 | 5876 | 5047 | 721 | 1024 | 1260 | 2843 | 20142 |
| PBANKA_0621000 | 118 | 114 | 26 | 11 | 173 | 92 | 183 | 101 | 92 | 53 | 56 | 10 | 277 | 55 | 103 | 397 | 375 | 90 | 2424 | 724 | 294 | 2270 |
| PBANKA_0621100 | 8 | 6 | 69 | 17 | 137 | 89 | 114 | 35 | 26 | 16 | 0 | 0 | 208 | 29 | 155 | 502 | 83 | 27 | 386 | 87 | 37 | 184 |
| PBANKA_0621200 | 19 | 25 | 126 | 26 | 138 | 50 | 109 | 32 | 56 | 25 | 61 | 28 | 1019 | 660 | 237 | 1270 | 222 | 44 | 729 | 474 | 59 | 381 |
| PBANKA_0621300 | 278 | 411 | 263 | 71 | 817 | 314 | 1349 | 474 | 1077 | 390 | 210 | 85 | 1883 | 1816 | 1005 | 6479 | 21172 | 2782 | 3358 | 11981 | 315 | 1962 |
| PBANKA_0621400 | 15 | 24 | 189 | 45 | 585 | 255 | 688 | 283 | 420 | 200 | 1854 | 1313 | 3965 | 3227 | 1263 | 5942 | 4398 | 745 | 1325 | 1440 | 278 | 2456 |
| PBANKA_0621500 | 27 | 57 | 70 | 12 | 259 | 89 | 407 | 134 | 272 | 115 | 46 | 13 | 779 | 381 | 735 | 3382 | 2375 | 377 | 2013 | 1717 | 1123 | 6542 |
| PBANKA_0621600 | 69 | 90 | 98 | 27 | 237 | 92 | 269 | 85 | 149 | 84 | 140 | 55 | 2140 | 970 | 604 | 5385 | 4403 | 447 | 2305 | 2891 | 516 | 5157 |
| PBANKA_0621700 | 0 | 0 | 43 | 8 | 128 | 43 | 110 | 30 | 42 | 29 | 9 | 0 | 444 | 128 | 332 | 2246 | 337 | 50 | 922 | 655 | 73 | 523 |
| PBANKA_0621800 | 5 | 6 | 19 | 6 | 75 | 31 | 68 | 16 | 86 | 42 | 15 | 0 | 181 | 66 | 71 | 330 | 109 | 40 | 2511 | 816 | 120 | 805 |
| PBANKA_0621900 | 60 | 106 | 52 | 16 | 72 | 41 | 32 | 43 | 18 | 6 | 35 | 22 | 563 | 224 | 111 | 728 | 188 | 15 | 764 | 497 | 241 | 1468 |
| PBANKA_0622100 | 74 | 105 | 80 | 29 | 514 | 220 | 969 | 338 | 715 | 296 | 109 | 47 | 1083 | 597 | 251 | 599 | 8637 | 1601 | 3075 | 2472 | 660 | 4480 |
| PBANKA_0622200 | 35 | 25 | 351 | 83 | 1112 | 428 | 1303 | 451 | 555 | 280 | 408 | 236 | 2818 | 3742 | 3216 | 11912 | 7417 | 1120 | 911 | 2338 | 299 | 1778 |
| PBANKA_0622300 | 139 | 205 | 60 | 17 | 188 | 80 | 322 | 133 | 131 | 48 | 27 | 13 | 118 | 20 | 175 | 627 | 206 | 42 | 597 | 351 | 2233 | 14855 |
| PBANKA_0622400 | 70 | 124 | 66 | 19 | 150 | 93 | 250 | 78 | 132 | 91 | 48 | 12 | 1773 | 2744 | 352 | 2286 | 2610 | 528 | 20649 | 1E+05 | 7044 | 48718 |
| PBANKA_0622500 | 135 | 224 | 28 | 7 | 169 | 84 | 400 | 125 | 199 | 99 | 59 | 34 | 526 | 309 | 170 | 437 | 4314 | 787 | 659 | 768 | 532 | 3656 |
| PBANKA_0622600 | 59 | 83 | 61 | 8 | 163 | 35 | 272 | 77 | 244 | 92 | 225 | 138 | 1604 | 1269 | 306 | 2025 | 2179 | 395 | 1713 | 1403 | 565 | 2887 |
| PBANKA_0622700 | 11 | 15 | 69 | 16 | 119 | 89 | 191 | 138 | 55 | 29 | 13 | 6 | 1279 | 283 | 583 | 1242 | 1170 | 342 | 264 | 172 | 18 | 58 |
| PBANKA_0622800 | 380 | 561 | 140 | 41 | 263 | 103 | 261 | 116 | 115 | 48 | 439 | 275 | 2292 | 1781 | 500 | 3376 | 2585 | 360 | 1996 | 1650 | 396 | 2848 |
| PBANKA_0622921 | 1782 | 2089 | 797 | 219 | 1447 | 326 | 1551 | 390 | 1067 | 270 | 1544 | 372 | 2E+05 | 5631 | 31094 | 23042 | 15530 | 5879 | 60055 | 9438 | 62 | 1349 |
| PBANKA_0622961 | 57604 | 70432 | 5031 | 1743 | 7693 | 2125 | 9772 | 2658 | 6023 | 1876 | 8329 | 2018 | 16125 | 2189 | 13175 | 6806 | 7211 | 2918 | 27471 | 6627 | 61 | 1675 |
| PBANKA_0623000 | 562 | 651 | 377 | 147 | 292 | 213 | 290 | 130 | 1014 | 807 | 5948 | 4808 | 2745 | 2042 | 390 | 1268 | 1059 | 559 | 581 | 288 | 8 | 89 |
| PBANKA_0623100 | 28 | 30 | 245 | 88 | 640 | 391 | 1800 | 954 | 1992 | 1337 | 10715 | 9913 | 19415 | 49579 | 4857 | 41696 | 85146 | 6799 | 3708 | 12333 | 577 | 4484 |
| PBANKA_0623200 | 5 | 0 | 19 | 10 | 27 | 18 | 68 | 19 | 247 | 235 | 4262 | 3489 | 7316 | 4717 | 1004 | 5578 | 1684 | 226 | 2056 | 954 | 22 | 298 |
| PBANKA_0623300 | 0 | 0 | 92 | 52 | 137 | 93 | 123 | 68 | 281 | 238 | 4882 | 4468 | 11839 | 7766 | 1206 | 4999 | 2291 | 369 | 1399 | 1058 | 24 | 264 |
| PBANKA_0623400 | 0 | 0 | 6 | 0 | 9 | 19 | 39 | 12 | 181 | 168 | 126 | 126 | 540 | 383 | 55 | 199 | 1482 | 359 | 213 | 158 | 8 | 90 |

| | | | | | | | | | | | | | | | | | | | | | | |
|---|---|---|---|---|---|---|---|---|---|---|---|---|---|---|---|---|---|---|---|---|---|---|
| PBANKA_0623500 | 0 | 0 | 6 | 0 | 48 | 48 | 62 | 27 | 170 | 178 | 1627 | 1407 | 1904 | 855 | 588 | 1542 | 675 | 371 | 622 | 259 | 36 | 422 |
| PBANKA_0700521 | 0 | 0 | 6 | 5 | 10 | 10 | 8 | 0 | 0 | 9 | 56 | 38 | 200 | 79 | 95 | 141 | 123 | 35 | 12 | 0 | 0 | 0 |
| PBANKA_0700700 | 0 | 0 | 43 | 14 | 68 | 35 | 92 | 40 | 97 | 49 | 1860 | 1451 | 3455 | 2127 | 465 | 2131 | 653 | 134 | 654 | 424 | 24 | 201 |
| PBANKA_0700721 | 193 | 681 | 8604 | 843 | 18107 | 1105 | 22097 | 4454 | 15856 | 2689 | 25408 | 3428 | 4E+05 | 27400 | 37733 | 70823 | 46364 | 6934 | 1E+05 | 31066 | 631 | 6024 |
| PBANKA_0700741 | 0 | 0 | 776 | 361 | 1336 | 949 | 1346 | 566 | 481 | 249 | 207 | 137 | 10898 | 994 | 16936 | 6737 | 3245 | 1250 | 1E+05 | 4668 | 0 | 20 |
| PBANKA_0700761 | 3798 | 6750 | 26059 | 8208 | 71560 | 30987 | 94968 | 27470 | 54473 | 22569 | 81576 | 25159 | 3E+05 | 46188 | 43275 | 1E+05 | 1E+05 | 9591 | 3E+05 | 1E+05 | 551 | 9232 |
| PBANKA_0700800 | 0 | 0 | 10 | 0 | 63 | 21 | 88 | 33 | 42 | 23 | 0 | 0 | 410 | 124 | 79 | 442 | 3804 | 377 | 1039 | 599 | 19 | 302 |
| PBANKA_0700900 | 0 | 0 | 5 | 0 | 34 | 18 | 96 | 21 | 169 | 81 | 560 | 364 | 390 | 536 | 135 | 768 | 470 | 148 | 167 | 179 | 0 | 41 |
| PBANKA_0701000 | 0 | 0 | 12 | 0 | 56 | 22 | 31 | 34 | 307 | 193 | 3379 | 2041 | 4793 | 4209 | 442 | 2939 | 1241 | 199 | 243 | 315 | 14 | 126 |
| PBANKA_0701200 | 53 | 99 | 9 | 5 | 103 | 44 | 193 | 105 | 490 | 168 | 87 | 40 | 376 | 998 | 81 | 290 | 10415 | 1958 | 384 | 3340 | 96 | 482 |
| PBANKA_0701300 | 13 | 16 | 7 | 0 | 61 | 32 | 55 | 25 | 65 | 32 | 20 | 0 | 70 | 75 | 13 | 17 | 893 | 274 | 79 | 304 | 39 | 251 |
| PBANKA_0701400 | 13 | 20 | 28 | 0 | 78 | 28 | 61 | 23 | 16 | 11 | 0 | 0 | 402 | 56 | 147 | 708 | 319 | 51 | 306 | 149 | 47 | 314 |
| PBANKA_0701500 | 0 | 0 | 147 | 34 | 157 | 79 | 122 | 49 | 67 | 8 | 5 | 0 | 1021 | 289 | 827 | 3395 | 388 | 68 | 406 | 306 | 110 | 967 |
| PBANKA_0701600 | 7 | 0 | 51 | 9 | 101 | 75 | 135 | 59 | 48 | 22 | 19 | 25 | 1835 | 1020 | 540 | 3324 | 349 | 41 | 570 | 582 | 63 | 340 |
| PBANKA_0701700 | 158 | 202 | 122 | 26 | 440 | 146 | 429 | 178 | 191 | 78 | 74 | 42 | 1179 | 904 | 622 | 3103 | 1843 | 230 | 1205 | 1817 | 228 | 1642 |
| PBANKA_0701800 | 95 | 118 | 52 | 12 | 143 | 66 | 329 | 184 | 166 | 116 | 13 | 6 | 275 | 81 | 126 | 302 | 1989 | 369 | 347 | 275 | 126 | 784 |
| PBANKA_0702100 | 154 | 165 | 5 | 0 | 33 | 10 | 37 | 12 | 85 | 70 | 28 | 12 | 188 | 37 | 23 | 76 | 531 | 289 | 1453 | 599 | 43 | 344 |
| PBANKA_0702200 | 0 | 5 | 22 | 0 | 31 | 15 | 41 | 21 | 37 | 10 | 8 | 7 | 336 | 44 | 208 | 669 | 635 | 142 | 867 | 451 | 81 | 442 |
| PBANKA_0702300 | 0 | 0 | 96 | 19 | 191 | 108 | 173 | 103 | 114 | 46 | 21 | 0 | 303 | 70 | 403 | 1546 | 526 | 121 | 1369 | 643 | 192 | 1370 |
| PBANKA_0702400 | 86 | 103 | 102 | 28 | 334 | 150 | 626 | 344 | 497 | 261 | 93 | 34 | 1310 | 662 | 493 | 2474 | 8236 | 1455 | 2740 | 2712 | 346 | 2433 |
| PBANKA_0702500 | 111 | 154 | 166 | 45 | 313 | 124 | 401 | 105 | 193 | 75 | 55 | 32 | 1170 | 1618 | 458 | 2552 | 1435 | 431 | 1124 | 2088 | 1311 | 7573 |
| PBANKA_0702600 | 103 | 124 | 23 | 0 | 95 | 41 | 161 | 53 | 94 | 33 | 14 | 0 | 398 | 153 | 113 | 588 | 289 | 59 | 582 | 301 | 297 | 1654 |
| PBANKA_0702700 | 41 | 27 | 13 | 5 | 290 | 185 | 293 | 165 | 85 | 75 | 10 | 0 | 216 | 44 | 62 | 185 | 147 | 87 | 2840 | 807 | 136 | 730 |
| PBANKA_0702800 | 9 | 14 | 1344 | 409 | 2194 | 1326 | 5888 | 2941 | 2390 | 1328 | 256 | 217 | 11446 | 3158 | 6034 | 17636 | 33146 | 7082 | 9929 | 5218 | 624 | 7903 |
| PBANKA_0702900 | 5 | 0 | 147 | 31 | 1286 | 388 | 3646 | 1301 | 2083 | 824 | 3857 | 2128 | 2756 | 6028 | 1159 | 6943 | 21115 | 2432 | 1841 | 4073 | 174 | 1359 |
| PBANKA_0703000 | 0 | 0 | 9 | 0 | 45 | 16 | 67 | 40 | 67 | 32 | 30 | 17 | 50 | 48 | 12 | 64 | 241 | 54 | 314 | 354 | 53 | 384 |
| PBANKA_0703100 | 0 | 5 | 132 | 29 | 265 | 118 | 285 | 108 | 151 | 76 | 92 | 29 | 1292 | 2183 | 974 | 5186 | 2227 | 335 | 454 | 1948 | 44 | 349 |
| PBANKA_0703200 | 0 | 0 | 97 | 21 | 463 | 192 | 831 | 345 | 279 | 124 | 53 | 18 | 1516 | 418 | 547 | 2599 | 2239 | 353 | 486 | 394 | 762 | 5339 |
| PBANKA_0703300 | 15 | 15 | 20 | 0 | 111 | 43 | 152 | 71 | 47 | 32 | 5 | 0 | 709 | 273 | 97 | 361 | 255 | 143 | 14374 | 4804 | 350 | 2377 |
| PBANKA_0703400 | 95 | 127 | 115 | 24 | 280 | 147 | 377 | 169 | 146 | 82 | 97 | 44 | 2882 | 1673 | 520 | 3711 | 2135 | 255 | 1061 | 906 | 658 | 3135 |
| PBANKA_0703500 | 14 | 15 | 8 | 0 | 39 | 20 | 37 | 21 | 59 | 32 | 35 | 0 | 2388 | 1573 | 155 | 584 | 1474 | 707 | 26820 | 34966 | 332 | 1412 |
| PBANKA_0703600 | 10 | 5 | 101 | 28 | 250 | 70 | 444 | 141 | 259 | 101 | 363 | 229 | 2086 | 1919 | 695 | 3494 | 3843 | 563 | 1879 | 2639 | 449 | 2911 |
| PBANKA_0703700 | 27 | 50 | 199 | 41 | 325 | 133 | 359 | 134 | 260 | 116 | 1694 | 1054 | 3758 | 6487 | 750 | 4696 | 3575 | 573 | 1691 | 6607 | 807 | 4402 |
| PBANKA_0703800 | 28 | 51 | 18 | 0 | 102 | 23 | 284 | 100 | 190 | 102 | 22 | 6 | 1203 | 430 | 125 | 872 | 2837 | 447 | 2265 | 1506 | 135 | 733 |
| PBANKA_0703900 | 58 | 49 | 2657 | 805 | 2051 | 1416 | 1710 | 882 | 549 | 342 | 2448 | 2092 | 21622 | 8204 | 5017 | 18696 | 3132 | 489 | 3184 | 1634 | 192 | 2352 |
| PBANKA_0704000 | 0 | 0 | 36 | 0 | 51 | 40 | 50 | 24 | 28 | 13 | 5 | 5 | 467 | 189 | 156 | 384 | 290 | 61 | 213 | 122 | 65 | 370 |
| PBANKA_0704100 | 6 | 5 | 152 | 52 | 513 | 194 | 837 | 306 | 530 | 296 | 77 | 32 | 2153 | 639 | 1464 | 8455 | 5640 | 804 | 2591 | 3198 | 342 | 2683 |
| PBANKA_0704200 | 0 | 0 | 15 | 0 | 195 | 96 | 272 | 159 | 137 | 55 | 23 | 22 | 466 | 82 | 261 | 519 | 730 | 251 | 1023 | 401 | 64 | 373 |
| PBANKA_0704300 | 0 | 0 | 39 | 5 | 228 | 70 | 413 | 144 | 217 | 95 | 11 | 10 | 548 | 102 | 224 | 931 | 2367 | 342 | 1243 | 797 | 48 | 333 |
| PBANKA_0704400 | 11 | 11 | 23 | 0 | 112 | 28 | 147 | 64 | 97 | 42 | 29 | 7 | 532 | 161 | 99 | 429 | 1558 | 353 | 6026 | 4475 | 77 | 591 |
| PBANKA_0704500 | 474 | 746 | 2666 | 620 | 4746 | 1942 | 5640 | 1856 | 2205 | 805 | 2806 | 1354 | 6424 | 28418 | 9978 | 49587 | 16401 | 1672 | 1070 | 12244 | 584 | 5403 |
| PBANKA_0704600 | 19 | 21 | 106 | 20 | 190 | 80 | 163 | 66 | 83 | 45 | 157 | 85 | 2343 | 2263 | 658 | 4925 | 2229 | 235 | 648 | 1847 | 25 | 119 |
| PBANKA_0705000 | 10 | 8 | 21 | 5 | 43 | 17 | 28 | 10 | 19 | 13 | 37 | 14 | 124 | 76 | 68 | 193 | 82 | 32 | 194 | 85 | 17 | 149 |
| PBANKA_0705100 | 7 | 10 | 66 | 9 | 120 | 52 | 128 | 38 | 102 | 49 | 827 | 400 | 3826 | 3064 | 530 | 3479 | 1202 | 223 | 1474 | 1353 | 247 | 1657 |
| PBANKA_0705200 | 13 | 14 | 103 | 15 | 171 | 111 | 304 | 167 | 88 | 64 | 9 | 0 | 305 | 52 | 179 | 631 | 657 | 117 | 322 | 156 | 36 | 419 |
| PBANKA_0705300 | 9 | 11 | 24 | 6 | 97 | 33 | 142 | 59 | 40 | 17 | 0 | 0 | 136 | 26 | 88 | 236 | 370 | 105 | 200 | 102 | 10 | 103 |
| PBANKA_0705400 | 0 | 0 | 5 | 0 | 29 | 12 | 77 | 28 | 29 | 20 | 0 | 0 | 126 | 24 | 62 | 184 | 492 | 99 | 215 | 134 | 11 | 70 |
| PBANKA_0705600 | 61 | 97 | 99 | 29 | 426 | 163 | 660 | 218 | 393 | 145 | 69 | 30 | 1261 | 2014 | 350 | 2131 | 10743 | 1384 | 15688 | 48509 | 164 | 1007 |
| PBANKA_0705700 | 45 | 65 | 151 | 26 | 304 | 123 | 339 | 121 | 200 | 100 | 517 | 377 | 3427 | 2393 | 700 | 3615 | 2239 | 437 | 726 | 592 | 119 | 657 |
| PBANKA_0705800 | 67 | 106 | 221 | 50 | 653 | 315 | 800 | 330 | 399 | 228 | 472 | 260 | 4253 | 2592 | 1332 | 8637 | 7558 | 863 | 2554 | 3218 | 116 | 778 |
| PBANKA_0705900 | 142 | 200 | 56 | 17 | 345 | 163 | 540 | 186 | 154 | 110 | 25 | 10 | 619 | 103 | 280 | 1298 | 1612 | 264 | 2069 | 959 | 133 | 840 |
| PBANKA_0706000 | 85 | 115 | 77 | 10 | 77 | 53 | 131 | 31 | 37 | 18 | 45 | 36 | 1070 | 540 | 167 | 763 | 699 | 136 | 590 | 335 | 64 | 430 |
| PBANKA_0706100 | 0 | 0 | 39 | 7 | 52 | 40 | 50 | 18 | 5 | 6 | 6 | 0 | 196 | 56 | 116 | 782 | 86 | 15 | 252 | 141 | 6 | 106 |
| PBANKA_0706200 | 0 | 0 | 143 | 19 | 186 | 121 | 177 | 64 | 68 | 15 | 0 | 0 | 900 | 163 | 803 | 2371 | 410 | 79 | 374 | 216 | 0 | 46 |
| PBANKA_0706300 | 6 | 9 | 59 | 10 | 173 | 59 | 126 | 49 | 52 | 24 | 6 | 7 | 514 | 152 | 567 | 3856 | 345 | 42 | 635 | 896 | 61 | 526 |
| PBANKA_0706400 | 0 | 0 | 93 | 16 | 291 | 122 | 277 | 106 | 108 | 38 | 27 | 15 | 622 | 283 | 773 | 2525 | 1788 | 295 | 3059 | 2779 | 192 | 1715 |
| PBANKA_0706500 | 0 | 0 | 32 | 5 | 37 | 17 | 31 | 14 | 22 | 13 | 0 | 0 | 106 | 22 | 127 | 625 | 80 | 10 | 146 | 54 | 1395 | 9180 |
| PBANKA_0706600 | 0 | 6 | 42 | 15 | 107 | 52 | 105 | 43 | 83 | 28 | 27 | 0 | 41 | 18 | 64 | 138 | 92 | 48 | 114 | 243 | 11936 | 86887 |
| PBANKA_0706700 | 96 | 133 | 113 | 31 | 100 | 83 | 145 | 72 | 100 | 69 | 209 | 204 | 2312 | 1074 | 234 | 652 | 1682 | 470 | 14374 | 7773 | 593 | 5593 |
| PBANKA_0706800 | 0 | 6 | 112 | 20 | 261 | 122 | 459 | 169 | 185 | 74 | 9 | 7 | 891 | 297 | 774 | 3243 | 4975 | 612 | 1716 | 1344 | 227 | 2087 |
| PBANKA_0706900 | 0 | 0 | 34 | 6 | 43 | 34 | 51 | 34 | 13 | 12 | 0 | 0 | 136 | 20 | 139 | 134 | 75 | 47 | 160 | 36 | 0 | 35 |
| PBANKA_0707300 | 96 | 127 | 447 | 119 | 1254 | 507 | 1781 | 809 | 1024 | 470 | 3355 | 2148 | 6378 | 7203 | 2305 | 15608 | 17758 | 2274 | 2495 | 3555 | 698 | 3995 |
| PBANKA_0707400 | 32 | 46 | 312 | 127 | 446 | 332 | 580 | 295 | 386 | 260 | 3045 | 2689 | 1528 | 1625 | 883 | 2727 | 1594 | 284 | 1957 | 1350 | 133 | 1412 |
| PBANKA_0707500 | 0 | 0 | 77 | 31 | 110 | 77 | 90 | 36 | 27 | 15 | 0 | 0 | 995 | 281 | 336 | 1323 | 236 | 38 | 1322 | 560 | 117 | 903 |
| PBANKA_0707600 | 0 | 0 | 31 | 11 | 257 | 108 | 351 | 128 | 222 | 68 | 34 | 5 | 755 | 339 | 407 | 1660 | 2108 | 265 | 964 | 1046 | 537 | 2922 |
| PBANKA_0707700 | 5 | 9 | 28 | 7 | 241 | 71 | 298 | 104 | 3838 | 2404 | 1678 | 798 | 1985 | 2219 | 113 | 428 | 14398 | 4638 | 1216 | 2155 | 382 | 2991 |
| PBANKA_0707800 | 16 | 16 | 117 | 27 | 254 | 179 | 319 | 113 | 130 | 70 | 181 | 122 | 801 | 298 | 316 | 1276 | 903 | 137 | 337 | 240 | 50 | 412 |
| PBANKA_0708000 | 0 | 7 | 202 | 47 | 706 | 465 | 1170 | 471 | 664 | 319 | 135 | 61 | 3866 | 1286 | 1486 | 5291 | 5566 | 1048 | 983 | 538 | 130 | 1181 |
| PBANKA_0708100 | 8 | 0 | 6 | 0 | 6 | 6 | 14 | 9 | 28 | 13 | 0 | 0 | 131 | 41 | 11 | 39 | 151 | 81 | 726 | 275 | 70 | 348 |
| PBANKA_0708200 | 0 | 0 | 24 | 0 | 54 | 14 | 43 | 10 | 34 | 15 | 38 | 26 | 682 | 284 | 75 | 413 | 403 | 148 | 3592 | 2951 | 117 | 725 |
| PBANKA_0708400 | 0 | 0 | 47 | 7 | 113 | 45 | 97 | 36 | 26 | 10 | 7 | 5 | 287 | 58 | 205 | 492 | 252 | 48 | 353 | 103 | 10 | 76 |
| PBANKA_0708500 | 11 | 11 | 50 | 9 | 102 | 37 | 85 | 40 | 39 | 20 | 11 | 12 | 484 | 156 | 229 | 908 | 558 | 168 | 3253 | 849 | 429 | 2630 |
| PBANKA_0708600 | 0 | 0 | 37 | 10 | 102 | 59 | 74 | 43 | 25 | 10 | 6 | 0 | 226 | 70 | 113 | 441 | 154 | 25 | 657 | 219 | 314 | 2375 |
| PBANKA_0708700 | 0 | 0 | 27 | 9 | 111 | 51 | 106 | 25 | 25 | 14 | 7 | 0 | 410 | 79 | 235 | 771 | 246 | 58 | 418 | 86 | 55 | 341 |
| PBANKA_0708800 | 0 | 0 | 117 | 23 | 218 | 90 | 376 | 142 | 175 | 90 | 42 | 14 | 1553 | 617 | 602 | 3763 | 2515 | 286 | 1309 | 1153 | 436 | 3120 |
| PBANKA_0708900 | 0 | 0 | 9 | 16 | 47 | 23 | 77 | 51 | 21 | 17 | 0 | 0 | 127 | 14 | 83 | 185 | 139 | 55 | 312 | 118 | 67 | 462 |
| PBANKA_0709000 | 0 | 0 | 95 | 26 | 146 | 98 | 257 | 135 | 93 | 71 | 76 | 63 | 268 | 135 | 112 | 256 | 543 | 197 | 1425 | 361 | 92 | 622 |
| PBANKA_0709100 | 13 | 10 | 506 | 147 | 376 | 198 | 365 | 173 | 105 | 91 | 506 | 394 | 4839 | 1839 | 1045 | 3138 | 957 | 251 | 1033 | 395 | 162 | 703 |
| PBANKA_0709200 | 0 | 0 | 101 | 15 | 484 | 135 | 402 | 141 | 173 | 51 | 59 | 25 | 837 | 406 | 520 | 2214 | 806 | 152 | 672 | 410 | 754 | 5123 |
| PBANKA_0709300 | 0 | 0 | 48 | 11 | 64 | 32 | 59 | 24 | 15 | 6 | 10 | 9 | 476 | 251 | 153 | 544 | 81 | 24 | 127 | 50 | 109 | 717 |
| PBANKA_0709400 | 0 | 0 | 63 | 11 | 86 | 45 | 179 | 52 | 191 | 157 | 842 | 482 | 2771 | 1060 | 518 | 1574 | 3794 | 1478 | 951 | 436 | 110 | 666 |
| PBANKA_0709500 | 33 | 64 | 256 | 44 | 459 | 233 | 419 | 194 | 348 | 181 | 258 | 173 | 4148 | 2103 | 1005 | 4153 | 2073 | 549 | 5113 | 3082 | 685 | 4655 |
| PBANKA_0709600 | 46 | 69 | 110 | 26 | 127 | 56 | 209 | 84 | 206 | 91 | 109 | 59 | 4451 | 1712 | 419 | 2373 | 2990 | 583 | 10236 | 8304 | 305 | 2366 |
| PBANKA_0709700 | 76 | 155 | 48 | 11 | 189 | 93 | 454 | 169 | 293 | 169 | 60 | 12 | 619 | 189 | 284 | 1882 | 3723 | 567 | 1859 | 1609 | 923 | 5607 |
| PBANKA_0709800 | 8 | 12 | 215 | 42 | 287 | 152 | 660 | 258 | 349 | 147 | 23 | 8 | 810 | 166 | 171 | 684 | 5234 | 922 | 612 | 306 | 245 | 1635 |
| PBANKA_0709900 | 0 | 0 | 61 | 16 | 219 | 89 | 287 | 110 | 171 | 80 | 45 | 10 | 990 | 244 | 627 | 3079 | 2164 | 284 | 961 | 766 | 70 | 305 |
| PBANKA_0710000 | 12 | 12 | 68 | 12 | 193 | 82 | 285 | 59 | 107 | 52 | 18 | 13 | 465 | 135 | 312 | 1563 | 1464 | 211 | 823 | 480 | 115 | 547 |
| PBANKA_0710100 | 59 | 72 | 416 | 137 | 1215 | 713 | 1711 | 655 | 621 | 278 | 72 | 55 | 702 | 309 | 1432 | 6945 | 5109 | 643 | 3418 | 4966 | 1441 | 13927 |
| PBANKA_0710200 | 22 | 32 | 35 | 10 | 159 | 60 | 259 | 114 | 103 | 60 | 33 | 12 | 322 | 57 | 103 | 364 | 916 | 207 | 2272 | 977 | 66 | 403 |
| PBANKA_0710300 | 321 | 471 | 70 | 9 | 103 | 68 | 261 | 95 | 142 | 74 | 62 | 53 | 1496 | 646 | 229 | 1373 | 2597 | 532 | 1907 | 823 | 530 | 3787 |

| | | | | | | | | | | | | | | | | | | | | | | |
|---|---|---|---|---|---|---|---|---|---|---|---|---|---|---|---|---|---|---|---|---|---|---|
| PBANKA_0710400 | 6 | 5 | 31 | 6 | 306 | 178 | 645 | 349 | 343 | 189 | 35 | 9 | 128 | 39 | 195 | 777 | 362 | 58 | 396 | 164 | 75 | 616 |
| PBANKA_0710500 | 72 | 60 | 64 | 16 | 233 | 53 | 403 | 122 | 249 | 82 | 302 | 162 | 1799 | 697 | 448 | 2394 | 2101 | 344 | 2471 | 1279 | 177 | 1255 |
| PBANKA_0710600 | 0 | 0 | 33 | 14 | 81 | 38 | 90 | 28 | 51 | 12 | 30 | 6 | 518 | 179 | 261 | 816 | 289 | 84 | 1075 | 586 | 144 | 993 |
| PBANKA_0710700 | 9 | 11 | 50 | 5 | 94 | 24 | 82 | 28 | 38 | 14 | 0 | 0 | 662 | 244 | 236 | 1074 | 175 | 50 | 1523 | 682 | 599 | 3490 |
| PBANKA_0710800 | 6 | 11 | 303 | 80 | 983 | 375 | 1237 | 427 | 598 | 287 | 1445 | 971 | 3011 | 2116 | 2983 | 12925 | 7546 | 1000 | 3038 | 3316 | 1014 | 6937 |
| PBANKA_0710900 | 0 | 0 | 52 | 21 | 64 | 68 | 92 | 75 | 48 | 22 | 0 | 6 | 226 | 46 | 182 | 324 | 297 | 99 | 238 | 140 | 137 | 496 |
| PBANKA_0711000 | 5 | 11 | 13 | 0 | 98 | 24 | 91 | 36 | 54 | 23 | 14 | 6 | 463 | 150 | 89 | 409 | 552 | 134 | 2738 | 2272 | 731 | 4557 |
| PBANKA_0711100 | 0 | 0 | 6 | 0 | 135 | 57 | 270 | 136 | 110 | 56 | 30 | 0 | 366 | 68 | 214 | 788 | 1438 | 250 | 589 | 215 | 847 | 4817 |
| PBANKA_0711200 | 1784 | 1681 | 126 | 43 | 313 | 181 | 489 | 213 | 413 | 311 | 63 | 52 | 2115 | 1368 | 1218 | 8244 | 5780 | 1207 | 1618 | 3173 | 704 | 4138 |
| PBANKA_0711300 | 6 | 0 | 11 | 0 | 51 | 23 | 44 | 23 | 27 | 10 | 5 | 0 | 110 | 36 | 23 | 22 | 87 | 31 | 1678 | 872 | 224 | 1212 |
| PBANKA_0711400 | 0 | 0 | 6 | 0 | 24 | 14 | 54 | 25 | 73 | 26 | 0 | 0 | 27 | 10 | 12 | 18 | 93 | 26 | 205 | 162 | 7858 | 57720 |
| PBANKA_0711500 | 0 | 5 | 18 | 0 | 52 | 37 | 83 | 20 | 25 | 14 | 0 | 0 | 275 | 62 | 82 | 498 | 227 | 39 | 1550 | 391 | 342 | 1799 |
| PBANKA_0711600 | 13 | 14 | 6 | 0 | 32 | 13 | 27 | 17 | 21 | 7 | 22 | 13 | 244 | 106 | 40 | 142 | 34 | 7 | 158 | 147 | 6312 | 43554 |
| PBANKA_0711700 | 16 | 22 | 33 | 8 | 50 | 29 | 43 | 26 | 12 | 7 | 49 | 53 | 1011 | 668 | 163 | 765 | 182 | 61 | 822 | 661 | 204 | 1478 |
| PBANKA_0711800 | 386 | 469 | 266 | 57 | 808 | 329 | 813 | 291 | 425 | 170 | 649 | 256 | 8348 | 6742 | 2632 | 16651 | 7148 | 933 | 2039 | 3030 | 709 | 4231 |
| PBANKA_0711900 | 15842 | 19583 | 20877 | 6944 | 79048 | 45274 | 51571 | 18095 | 19735 | 10049 | 26003 | 16019 | 1E+05 | 46300 | 37466 | 1E+05 | 45651 | 4945 | 35692 | 29134 | 10118 | 1E+05 |
| PBANKA_0712000 | 12 | 19 | 126 | 20 | 144 | 68 | 162 | 49 | 135 | 48 | 998 | 842 | 2925 | 1677 | 291 | 1819 | 864 | 134 | 3934 | 2276 | 1440 | 9763 |
| PBANKA_0712100 | 105 | 162 | 173 | 38 | 694 | 170 | 1466 | 488 | 933 | 386 | 149 | 57 | 2224 | 883 | 795 | 3954 | 13711 | 1650 | 7085 | 7830 | 274 | 1877 |
| PBANKA_0712200 | 987 | 1133 | 615 | 205 | 374 | 262 | 424 | 152 | 149 | 81 | 52 | 10 | 1398 | 790 | 214 | 717 | 661 | 268 | 10705 | 6690 | 1913 | 16692 |
| PBANKA_0712300 | 60 | 61 | 111 | 39 | 244 | 109 | 351 | 139 | 394 | 238 | 111 | 68 | 2144 | 1071 | 502 | 2368 | 1960 | 456 | 2480 | 1394 | 313 | 2285 |
| PBANKA_0712400 | 89 | 156 | 46 | 15 | 54 | 32 | 49 | 33 | 87 | 59 | 316 | 254 | 639 | 451 | 101 | 407 | 181 | 61 | 691 | 374 | 308 | 2228 |
| PBANKA_0712600 | 32 | 42 | 1720 | 618 | 1810 | 1170 | 3323 | 2134 | 1486 | 1112 | 89 | 97 | 6180 | 1422 | 6646 | 17690 | 16778 | 3107 | 9344 | 4515 | 942 | 10941 |
| PBANKA_0712700 | 198 | 371 | 6 | 0 | 69 | 44 | 95 | 19 | 161 | 88 | 49 | 19 | 124 | 78 | 20 | 52 | 348 | 149 | 713 | 238 | 317 | 2124 |
| PBANKA_0712800 | 0 | 6 | 110 | 27 | 125 | 66 | 84 | 31 | 107 | 41 | 1128 | 693 | 3536 | 3903 | 549 | 5396 | 925 | 64 | 610 | 1184 | 41 | 403 |
| PBANKA_0712900 | 127 | 118 | 274 | 98 | 96 | 75 | 82 | 43 | 116 | 115 | 1202 | 1272 | 6964 | 2365 | 520 | 2004 | 1160 | 772 | 84715 | 39043 | 900 | 6467 |
| PBANKA_0713000 | 43 | 46 | 20 | 5 | 55 | 47 | 98 | 44 | 79 | 41 | 11 | 0 | 84 | 16 | 96 | 221 | 355 | 135 | 320 | 102 | 13 | 101 |
| PBANKA_0713100 | 30 | 61 | 31 | 13 | 124 | 74 | 781 | 335 | 2360 | 1508 | 504 | 226 | 2243 | 1030 | 89 | 162 | 21461 | 8174 | 851 | 876 | 294 | 1442 |
| PBANKA_0713200 | 406 | 596 | 29 | 12 | 80 | 41 | 180 | 67 | 713 | 295 | 232 | 79 | 801 | 420 | 41 | 106 | 6854 | 1714 | 4382 | 5015 | 6176 | 43952 |
| PBANKA_0713300 | 0 | 0 | 311 | 86 | 437 | 315 | 450 | 262 | 111 | 101 | 18 | 5 | 301 | 87 | 983 | 1741 | 714 | 139 | 426 | 201 | 99 | 646 |
| PBANKA_0713400 | 0 | 0 | 22 | 0 | 123 | 70 | 113 | 20 | 42 | 17 | 8 | 0 | 95 | 14 | 157 | 524 | 205 | 27 | 749 | 156 | 41 | 216 |
| PBANKA_0713500 | 0 | 0 | 158 | 30 | 236 | 82 | 177 | 72 | 101 | 48 | 46 | 35 | 740 | 329 | 821 | 4117 | 1653 | 261 | 1666 | 3051 | 549 | 3354 |
| PBANKA_0713600 | 31 | 50 | 15 | 0 | 53 | 21 | 89 | 20 | 39 | 25 | 272 | 174 | 945 | 548 | 120 | 544 | 396 | 56 | 1335 | 780 | 127 | 792 |
| PBANKA_0713700 | 7 | 0 | 131 | 22 | 238 | 79 | 220 | 73 | 119 | 49 | 31 | 10 | 1164 | 217 | 553 | 2538 | 1385 | 248 | 1832 | 1841 | 509 | 2823 |
| PBANKA_0713800 | 0 | 0 | 33 | 0 | 60 | 38 | 70 | 23 | 42 | 22 | 0 | 0 | 110 | 40 | 78 | 439 | 278 | 47 | 426 | 415 | 107 | 742 |
| PBANKA_0713900 | 0 | 0 | 14 | 10 | 117 | 54 | 154 | 73 | 60 | 38 | 12 | 6 | 186 | 34 | 44 | 103 | 1120 | 140 | 527 | 251 | 49 | 358 |
| PBANKA_0714000 | 0 | 0 | 10 | 0 | 30 | 23 | 46 | 21 | 15 | 26 | 10 | 7 | 2061 | 847 | 85 | 199 | 566 | 459 | 51850 | 31100 | 558 | 3813 |
| PBANKA_0714100 | 0 | 0 | 23 | 6 | 66 | 33 | 41 | 32 | 21 | 9 | 8 | 5 | 220 | 81 | 124 | 349 | 142 | 37 | 597 | 209 | 14 | 84 |
| PBANKA_0714200 | 12 | 29 | 604 | 228 | 1189 | 725 | 1003 | 334 | 268 | 152 | 80 | 16 | 2268 | 1595 | 2068 | 9890 | 3342 | 397 | 1687 | 2993 | 183 | 1419 |
| PBANKA_0714400 | 523 | 872 | 230 | 41 | 317 | 182 | 344 | 157 | 101 | 47 | 16 | 8 | 1228 | 192 | 418 | 1439 | 854 | 165 | 2594 | 1429 | 64 | 364 |
| PBANKA_0714500 | 114 | 160 | 7 | 0 | 42 | 17 | 33 | 17 | 38 | 18 | 9 | 0 | 165 | 66 | 8 | 19 | 126 | 30 | 1345 | 819 | 477 | 2790 |
| PBANKA_0714600 | 0 | 7 | 12 | 0 | 49 | 15 | 86 | 35 | 39 | 11 | 15 | 9 | 264 | 82 | 63 | 200 | 975 | 161 | 2122 | 1668 | 66 | 414 |
| PBANKA_0714700 | 0 | 0 | 13 | 5 | 65 | 38 | 46 | 26 | 24 | 12 | 138 | 132 | 144 | 49 | 129 | 253 | 91 | 18 | 34 | 15 | 6 | 53 |
| PBANKA_0714800 | 5 | 0 | 175 | 41 | 175 | 108 | 103 | 68 | 73 | 44 | 170 | 112 | 873 | 582 | 248 | 1105 | 247 | 48 | 100 | 97 | 77 | 625 |
| PBANKA_0714900 | 0 | 0 | 72 | 35 | 406 | 228 | 271 | 109 | 82 | 43 | 8 | 11 | 95 | 50 | 25 | 85 | 35 | 0 | 51 | 41 | 18 | 111 |
| PBANKA_0715000 | 15 | 32 | 164 | 48 | 385 | 169 | 550 | 227 | 428 | 197 | 249 | 150 | 3045 | 3328 | 1065 | 6325 | 19906 | 3138 | 2110 | 7604 | 2703 | 13209 |
| PBANKA_0715100 | 0 | 0 | 80 | 7 | 119 | 49 | 121 | 48 | 37 | 27 | 6 | 0 | 392 | 91 | 241 | 1303 | 1032 | 183 | 254 | 143 | 19 | 138 |
| PBANKA_0715200 | 13 | 5 | 452 | 172 | 465 | 234 | 339 | 171 | 132 | 123 | 511 | 550 | 3042 | 1308 | 1188 | 3604 | 1024 | 232 | 1111 | 594 | 167 | 877 |
| PBANKA_0715300 | 0 | 0 | 6 | 0 | 144 | 101 | 525 | 195 | 240 | 119 | 20 | 18 | 1092 | 228 | 138 | 236 | 4590 | 809 | 1035 | 451 | 203 | 1229 |
| PBANKA_0715400 | 18 | 14 | 72 | 9 | 147 | 64 | 189 | 73 | 68 | 47 | 12 | 6 | 310 | 143 | 219 | 801 | 1987 | 360 | 1215 | 686 | 103 | 754 |
| PBANKA_0715500 | 0 | 5 | 103 | 22 | 399 | 184 | 569 | 219 | 274 | 146 | 51 | 23 | 659 | 153 | 578 | 1641 | 3476 | 850 | 1773 | 1020 | 125 | 1079 |
| PBANKA_0715600 | 13 | 6 | 251 | 85 | 519 | 398 | 878 | 481 | 333 | 189 | 29 | 14 | 3038 | 627 | 2024 | 6728 | 5399 | 1035 | 1246 | 748 | 677 | 5486 |
| PBANKA_0715700 | 35 | 44 | 102 | 34 | 178 | 95 | 215 | 110 | 125 | 77 | 342 | 243 | 1013 | 324 | 220 | 770 | 3243 | 623 | 942 | 581 | 128 | 1246 |
| PBANKA_0715800 | 27 | 44 | 62 | 12 | 142 | 55 | 213 | 70 | 54 | 41 | 23 | 22 | 475 | 169 | 261 | 1123 | 1752 | 314 | 860 | 386 | 226 | 1599 |
| PBANKA_0715900 | 0 | 0 | 139 | 21 | 282 | 175 | 440 | 222 | 239 | 116 | 35 | 9 | 829 | 491 | 803 | 3478 | 2963 | 467 | 648 | 1161 | 270 | 2185 |
| PBANKA_0716000 | 8 | 19 | 567 | 106 | 857 | 341 | 999 | 362 | 414 | 220 | 109 | 32 | 774 | 837 | 1437 | 7388 | 7742 | 1000 | 2083 | 6438 | 399 | 2603 |
| PBANKA_0716100 | 9 | 8 | 53 | 14 | 142 | 37 | 104 | 49 | 71 | 26 | 8 | 10 | 622 | 267 | 339 | 1930 | 281 | 52 | 777 | 908 | 9 | 78 |
| PBANKA_0716300 | 37 | 31 | 18 | 0 | 30 | 8 | 36 | 15 | 18 | 0 | 0 | 0 | 114 | 38 | 26 | 95 | 192 | 30 | 200 | 55 | 187 | 1083 |
| PBANKA_0716500 | 0 | 0 | 38 | 20 | 99 | 73 | 104 | 59 | 39 | 18 | 0 | 7 | 474 | 98 | 257 | 822 | 601 | 207 | 3780 | 2036 | 118 | 996 |
| PBANKA_0716600 | 210 | 253 | 44 | 0 | 111 | 72 | 137 | 56 | 86 | 46 | 24 | 8 | 455 | 116 | 162 | 608 | 1609 | 368 | 2705 | 1670 | 85 | 865 |
| PBANKA_0716700 | 115 | 191 | 138 | 30 | 176 | 51 | 179 | 48 | 106 | 31 | 26 | 9 | 738 | 233 | 632 | 2669 | 2537 | 541 | 3407 | 2401 | 396 | 2642 |
| PBANKA_0716900 | 84 | 120 | 12 | 0 | 46 | 30 | 51 | 21 | 631 | 676 | 260 | 145 | 286 | 139 | 21 | 57 | 660 | 413 | 741 | 392 | 453 | 2258 |
| PBANKA_0717000 | 7 | 10 | 150 | 22 | 218 | 108 | 210 | 79 | 104 | 67 | 39 | 20 | 1405 | 561 | 568 | 2425 | 904 | 194 | 394 | 281 | 21 | 172 |
| PBANKA_0717100 | 47 | 59 | 44 | 6 | 58 | 14 | 61 | 15 | 54 | 24 | 6 | 0 | 234 | 72 | 77 | 303 | 312 | 96 | 328 | 175 | 25 | 172 |
| PBANKA_0717200 | 64 | 34 | 46 | 13 | 331 | 118 | 516 | 199 | 289 | 125 | 57 | 21 | 1545 | 990 | 711 | 4220 | 7593 | 1114 | 3199 | 6445 | 128 | 724 |
| PBANKA_0717300 | 46 | 51 | 384 | 82 | 1172 | 422 | 1232 | 413 | 551 | 195 | 465 | 216 | 4794 | 4836 | 1428 | 9098 | 5553 | 612 | 2826 | 6091 | 244 | 1541 |
| PBANKA_0717400 | 16 | 20 | 42 | 6 | 156 | 54 | 161 | 69 | 92 | 37 | 19 | 6 | 905 | 212 | 241 | 945 | 930 | 192 | 577 | 255 | 101 | 745 |
| PBANKA_0717500 | 7 | 0 | 110 | 21 | 105 | 48 | 139 | 55 | 63 | 41 | 24 | 24 | 1198 | 536 | 244 | 886 | 648 | 133 | 439 | 253 | 97 | 804 |
| PBANKA_0717600 | 0 | 0 | 39 | 9 | 40 | 12 | 41 | 14 | 18 | 10 | 0 | 0 | 306 | 79 | 139 | 606 | 221 | 40 | 1890 | 989 | 580 | 3307 |
| PBANKA_0717700 | 5 | 9 | 41 | 18 | 125 | 72 | 169 | 75 | 59 | 29 | 20 | 8 | 620 | 330 | 278 | 853 | 789 | 159 | 924 | 618 | 283 | 2322 |
| PBANKA_0717800 | 76 | 175 | 1604 | 429 | 942 | 653 | 922 | 453 | 395 | 268 | 2028 | 1629 | 17942 | 6239 | 3498 | 16198 | 5443 | 652 | 2950 | 1746 | 348 | 3113 |
| PBANKA_0717900 | 80 | 133 | 73 | 21 | 205 | 83 | 345 | 110 | 173 | 58 | 58 | 52 | 3577 | 1475 | 375 | 2168 | 2639 | 452 | 1264 | 593 | 434 | 3141 |
| PBANKA_0718000 | 0 | 0 | 30 | 0 | 63 | 18 | 68 | 34 | 55 | 39 | 7 | 5 | 145 | 52 | 28 | 106 | 97 | 18 | 75 | 53 | 168 | 1146 |
| PBANKA_0718100 | 0 | 8 | 138 | 17 | 250 | 99 | 260 | 128 | 196 | 84 | 59 | 16 | 441 | 306 | 298 | 1382 | 528 | 80 | 289 | 235 | 735 | 4558 |
| PBANKA_0718200 | 0 | 0 | 12 | 0 | 64 | 25 | 65 | 17 | 23 | 20 | 31 | 33 | 630 | 419 | 183 | 929 | 746 | 122 | 176 | 207 | 34 | 124 |
| PBANKA_0718300 | 18 | 18 | 80 | 23 | 387 | 150 | 473 | 160 | 217 | 82 | 41 | 18 | 121 | 56 | 269 | 1462 | 574 | 71 | 155 | 203 | 4512 | 33999 |
| PBANKA_0718400 | 49 | 84 | 207 | 68 | 1065 | 468 | 1708 | 740 | 1129 | 571 | 164 | 96 | 644 | 412 | 1485 | 7687 | 6153 | 671 | 1119 | 4148 | 1017 | 6128 |
| PBANKA_0718600 | 0 | 0 | 37 | 9 | 66 | 21 | 46 | 20 | 30 | 5 | 0 | 0 | 524 | 124 | 202 | 709 | 174 | 43 | 409 | 229 | 58 | 446 |
| PBANKA_0718700 | 0 | 8 | 31 | 0 | 54 | 32 | 63 | 22 | 21 | 13 | 0 | 0 | 85 | 18 | 50 | 185 | 147 | 40 | 277 | 124 | 169 | 1218 |
| PBANKA_0718800 | 44 | 42 | 262 | 76 | 451 | 327 | 719 | 353 | 668 | 494 | 145 | 64 | 1725 | 624 | 404 | 1574 | 5674 | 1150 | 1889 | 1054 | 190 | 1716 |
| PBANKA_0718900 | 87 | 66 | 116 | 37 | 334 | 128 | 309 | 93 | 141 | 76 | 30 | 15 | 1065 | 1212 | 643 | 3189 | 771 | 132 | 1447 | 1845 | 2064 | 11265 |
| PBANKA_0719300 | 240 | 180 | 160 | 37 | 302 | 151 | 417 | 139 | 148 | 58 | 36 | 8 | 538 | 96 | 601 | 2176 | 2486 | 329 | 1732 | 844 | 90 | 952 |
| PBANKA_0719400 | 89 | 88 | 112 | 20 | 220 | 91 | 213 | 68 | 127 | 55 | 93 | 39 | 2075 | 1141 | 642 | 3119 | 2099 | 427 | 1330 | 1219 | 469 | 3741 |
| PBANKA_0719500 | 291 | 329 | 478 | 97 | 1192 | 341 | 1980 | 651 | 2112 | 881 | 1197 | 585 | 2718 | 5598 | 2147 | 13067 | 20078 | 3395 | 3078 | 16093 | 2837 | 15123 |
| PBANKA_0719600 | 9 | 11 | 124 | 20 | 180 | 135 | 135 | 54 | 50 | 25 | 0 | 0 | 645 | 190 | 363 | 1216 | 161 | 47 | 298 | 125 | 204 | 893 |
| PBANKA_0719900 | 23 | 29 | 31 | 0 | 117 | 38 | 113 | 54 | 62 | 34 | 12 | 0 | 205 | 100 | 124 | 740 | 1048 | 167 | 976 | 984 | 394 | 2313 |
| PBANKA_0720100 | 0 | 0 | 8 | 0 | 33 | 5 | 45 | 19 | 73 | 40 | 28 | 12 | 454 | 484 | 59 | 251 | 2412 | 377 | 2075 | 9250 | 1385 | 8617 |
| PBANKA_0720200 | 71 | 89 | 60 | 26 | 273 | 79 | 302 | 80 | 172 | 55 | 199 | 103 | 2296 | 973 | 838 | 4873 | 2181 | 408 | 3462 | 2829 | 323 | 1968 |
| PBANKA_0720300 | 16 | 21 | 207 | 27 | 253 | 138 | 242 | 87 | 84 | 50 | 294 | 178 | 2814 | 1219 | 774 | 3406 | 1070 | 146 | 827 | 518 | 137 | 1021 |
| PBANKA_0720400 | 762 | 1032 | 224 | 57 | 1273 | 601 | 1395 | 670 | 1361 | 828 | 5829 | 3845 | 5913 | 6525 | 1597 | 10565 | 9661 | 1661 | 8113 | 12041 | 4630 | 32296 |
| PBANKA_0720500 | 0 | 0 | 71 | 15 | 327 | 139 | 381 | 179 | 98 | 39 | 15 | 10 | 115 | 9 | 134 | 274 | 81 | 19 | 301 | 157 | 8 | 36 |

| | | | | | | | | | | | | | | | | | | | | | | |
|---|---|---|---|---|---|---|---|---|---|---|---|---|---|---|---|---|---|---|---|---|---|---|
| PBANKA_0720600 | 0 | 0 | 67 | 21 | 129 | 91 | 139 | 75 | 90 | 70 | 17 | 18 | 272 | 56 | 248 | 579 | 294 | 61 | 144 | 81 | 26 | 119 |
| PBANKA_0720700 | 39 | 75 | 7 | 0 | 130 | 39 | 159 | 44 | 475 | 179 | 194 | 50 | 202 | 271 | 73 | 241 | 2500 | 771 | 397 | 790 | 117 | 667 |
| PBANKA_0720800 | 44 | 60 | 151 | 20 | 304 | 85 | 169 | 47 | 261 | 148 | 2425 | 1510 | 1687 | 1272 | 320 | 1804 | 617 | 124 | 823 | 764 | 169 | 1156 |
| PBANKA_0721200 | 0 | 9 | 21 | 0 | 68 | 16 | 42 | 7 | 13 | 0 | 0 | 0 | 189 | 78 | 51 | 256 | 88 | 23 | 361 | 158 | 24 | 106 |
| PBANKA_0721300 | 31 | 34 | 31 | 0 | 37 | 16 | 63 | 27 | 24 | 14 | 31 | 25 | 652 | 258 | 139 | 913 | 829 | 166 | 597 | 404 | 59 | 277 |
| PBANKA_0721400 | 17 | 16 | 21 | 0 | 136 | 60 | 132 | 60 | 44 | 25 | 9 | 0 | 486 | 124 | 260 | 730 | 394 | 57 | 404 | 130 | 24 | 110 |
| PBANKA_0721500 | 6 | 16 | 154 | 36 | 354 | 112 | 558 | 149 | 224 | 87 | 64 | 46 | 1557 | 1541 | 909 | 4592 | 3167 | 386 | 3941 | 3335 | 177 | 1025 |
| PBANKA_0721600 | 6 | 8 | 16 | 0 | 102 | 39 | 158 | 77 | 61 | 38 | 10 | 0 | 513 | 123 | 216 | 934 | 1041 | 125 | 923 | 287 | 38 | 149 |
| PBANKA_0721700 | 65 | 130 | 688 | 199 | 1072 | 598 | 1066 | 452 | 348 | 251 | 63 | 40 | 6122 | 3280 | 3556 | 22266 | 8793 | 1054 | 1929 | 3206 | 349 | 2759 |
| PBANKA_0721900 | 12 | 26 | 60 | 12 | 97 | 58 | 113 | 47 | 25 | 20 | 184 | 119 | 1782 | 948 | 289 | 1544 | 760 | 84 | 510 | 343 | 81 | 547 |
| PBANKA_0722000 | 188 | 240 | 173 | 34 | 465 | 134 | 897 | 290 | 793 | 337 | 208 | 76 | 1766 | 1946 | 1201 | 8009 | 15655 | 1835 | 1661 | 8563 | 645 | 3812 |
| PBANKA_0722600 | 0 | 0 | 6 | 0 | 17 | 10 | 18 | 0 | 20 | 8 | 9 | 5 | 5729 | 93 | 130 | 88 | 38 | 49 | 971 | 82 | 63 | 443 |
| PBANKA_0722801 | 0 | 0 | 25 | 0 | 58 | 37 | 66 | 25 | 17 | 11 | 20 | 12 | 2519 | 194 | 176 | 584 | 198 | 24 | 146 | 79 | 9 | 77 |
| PBANKA_0800400 | 0 | 0 | 28 | 5 | 63 | 34 | 101 | 52 | 77 | 43 | 629 | 497 | 13198 | 3652 | 780 | 4249 | 4117 | 560 | 1805 | 661 | 162 | 929 |
| PBANKA_0800600 | 0 | 0 | 27 | 0 | 69 | 34 | 65 | 26 | 13 | 8 | 0 | 0 | 188 | 54 | 141 | 502 | 434 | 62 | 298 | 170 | 11 | 58 |
| PBANKA_0800700 | 0 | 0 | 40 | 11 | 167 | 65 | 255 | 93 | 133 | 54 | 19 | 5 | 371 | 381 | 467 | 2136 | 1875 | 239 | 456 | 909 | 19 | 145 |
| PBANKA_0800800 | 40 | 66 | 26 | 0 | 52 | 27 | 87 | 57 | 38 | 40 | 11 | 0 | 265 | 94 | 88 | 195 | 503 | 143 | 369 | 204 | 47 | 289 |
| PBANKA_0800900 | 5 | 14 | 76 | 26 | 427 | 158 | 483 | 205 | 176 | 100 | 21 | 6 | 901 | 447 | 982 | 5386 | 2687 | 278 | 850 | 1061 | 59 | 233 |
| PBANKA_0801000 | 21 | 24 | 7 | 0 | 25 | 24 | 23 | 14 | 18 | 39 | 10 | 14 | 302 | 152 | 94 | 533 | 215 | 41 | 236 | 171 | 24 | 96 |
| PBANKA_0801100 | 0 | 0 | 38 | 0 | 87 | 29 | 93 | 28 | 52 | 33 | 11 | 13 | 392 | 152 | 142 | 854 | 707 | 162 | 2610 | 2401 | 275 | 1719 |
| PBANKA_0801200 | 269 | 407 | 514 | 145 | 865 | 486 | 924 | 301 | 380 | 158 | 735 | 509 | 7506 | 9627 | 2747 | 11717 | 3160 | 598 | 3159 | 4738 | 851 | 7790 |
| PBANKA_0801300 | 0 | 0 | 9 | 0 | 90 | 57 | 138 | 104 | 103 | 38 | 17 | 6 | 319 | 42 | 27 | 26 | 982 | 482 | 204 | 80 | 6 | 36 |
| PBANKA_0801400 | 0 | 0 | 6 | 0 | 17 | 13 | 33 | 16 | 13 | 9 | 84 | 88 | 1963 | 1228 | 398 | 1874 | 514 | 73 | 665 | 338 | 0 | 50 |
| PBANKA_0801600 | 154 | 207 | 500 | 98 | 803 | 272 | 723 | 256 | 452 | 189 | 1239 | 716 | 1771 | 3901 | 1526 | 8070 | 3982 | 422 | 1046 | 3180 | 152 | 932 |
| PBANKA_0801700 | 6 | 16 | 110 | 36 | 393 | 149 | 420 | 156 | 209 | 71 | 48 | 17 | 656 | 740 | 694 | 3486 | 1148 | 165 | 936 | 2027 | 80 | 672 |
| PBANKA_0801800 | 0 | 5 | 6 | 0 | 60 | 43 | 864 | 322 | 964 | 463 | 115 | 46 | 969 | 1865 | 56 | 174 | 22818 | 3605 | 8928 | 38883 | 137 | 1163 |
| PBANKA_0801900 | 20 | 38 | 214 | 53 | 455 | 184 | 430 | 141 | 182 | 82 | 114 | 60 | 2082 | 1490 | 732 | 4765 | 1353 | 126 | 365 | 761 | 89 | 592 |
| PBANKA_0802000 | 12 | 12 | 230 | 43 | 1200 | 551 | 2741 | 1081 | 2074 | 954 | 346 | 144 | 1746 | 2426 | 1017 | 6526 | 71055 | 7978 | 210 | 1589 | 64 | 410 |
| PBANKA_0802100 | 0 | 13 | 40 | 14 | 155 | 90 | 319 | 139 | 203 | 51 | 40 | 25 | 1028 | 367 | 381 | 1420 | 2487 | 449 | 2111 | 1397 | 379 | 3139 |
| PBANKA_0802200 | 0 | 0 | 15 | 6 | 63 | 22 | 705 | 237 | 722 | 291 | 63 | 29 | 1098 | 524 | 177 | 672 | 12375 | 1812 | 803 | 1154 | 23 | 152 |
| PBANKA_0802300 | 141 | 221 | 798 | 152 | 2035 | 694 | 2920 | 1091 | 1325 | 560 | 1082 | 705 | 9682 | 23631 | 10728 | 62680 | 35229 | 3581 | 4823 | 32145 | 2275 | 14385 |
| PBANKA_0802400 | 33 | 31 | 31 | 8 | 103 | 43 | 237 | 77 | 141 | 52 | 32 | 0 | 620 | 248 | 182 | 721 | 1897 | 320 | 1621 | 707 | 64 | 371 |
| PBANKA_0802500 | 35 | 30 | 17 | 0 | 147 | 37 | 183 | 71 | 351 | 168 | 65 | 30 | 419 | 233 | 103 | 265 | 2505 | 573 | 1458 | 1870 | 100 | 581 |
| PBANKA_0802600 | 0 | 0 | 33 | 8 | 132 | 54 | 149 | 52 | 54 | 28 | 7 | 0 | 335 | 115 | 220 | 1054 | 454 | 78 | 640 | 378 | 30 | 230 |
| PBANKA_0802700 | 0 | 0 | 83 | 21 | 312 | 103 | 390 | 103 | 237 | 115 | 79 | 36 | 2004 | 1493 | 898 | 5189 | 4055 | 663 | 1450 | 2141 | 73 | 479 |
| PBANKA_0802800 | 48 | 81 | 32 | 16 | 190 | 70 | 173 | 76 | 175 | 87 | 55 | 24 | 301 | 104 | 95 | 421 | 2454 | 543 | 1214 | 707 | 225 | 1550 |
| PBANKA_0802900 | 130 | 190 | 137 | 20 | 164 | 51 | 119 | 37 | 81 | 24 | 34 | 11 | 1430 | 794 | 444 | 2225 | 678 | 94 | 1416 | 839 | 99 | 522 |
| PBANKA_0803000 | 14 | 23 | 36 | 7 | 225 | 85 | 384 | 191 | 193 | 103 | 46 | 9 | 733 | 280 | 155 | 622 | 6695 | 977 | 1151 | 2050 | 21 | 233 |
| PBANKA_0803100 | 64 | 114 | 385 | 162 | 1248 | 611 | 1897 | 788 | 936 | 394 | 133 | 51 | 1796 | 570 | 1590 | 7323 | 10518 | 1191 | 4059 | 4190 | 136 | 1496 |
| PBANKA_0803200 | 520 | 857 | 284 | 84 | 1117 | 379 | 1838 | 698 | 1123 | 500 | 169 | 59 | 1899 | 1635 | 1922 | 11358 | 17025 | 1867 | 1896 | 5969 | 532 | 3509 |
| PBANKA_0803300 | 5 | 11 | 140 | 34 | 251 | 96 | 207 | 65 | 56 | 46 | 26 | 7 | 869 | 781 | 415 | 2167 | 352 | 64 | 581 | 775 | 76 | 526 |
| PBANKA_0803400 | 0 | 0 | 349 | 143 | 246 | 174 | 125 | 133 | 75 | 36 | 136 | 315 | 2139 | 750 | 445 | 1222 | 480 | 75 | 702 | 314 | 13 | 96 |
| PBANKA_0803500 | 9 | 0 | 252 | 135 | 685 | 306 | 799 | 518 | 330 | 196 | 18 | 35 | 826 | 141 | 1123 | 2182 | 1888 | 517 | 1158 | 428 | 127 | 1055 |
| PBANKA_0803600 | 22 | 30 | 324 | 109 | 618 | 259 | 690 | 245 | 226 | 105 | 364 | 332 | 2868 | 1795 | 2379 | 8762 | 2976 | 381 | 710 | 925 | 285 | 2505 |
| PBANKA_0803700 | 8 | 10 | 197 | 52 | 383 | 155 | 397 | 157 | 153 | 74 | 26 | 13 | 899 | 395 | 814 | 4011 | 5922 | 760 | 1336 | 2130 | 81 | 591 |
| PBANKA_0803800 | 0 | 0 | 18 | 8 | 21 | 11 | 15 | 14 | 6 | 0 | 0 | 0 | 17 | 0 | 10 | 90 | 37 | 13 | 110 | 60 | 0 | 0 |
| PBANKA_0804000 | 19 | 18 | 706 | 226 | 486 | 285 | 550 | 289 | 168 | 126 | 728 | 637 | 6429 | 2985 | 1395 | 4564 | 1726 | 281 | 984 | 508 | 46 | 430 |
| PBANKA_0804100 | 5 | 8 | 916 | 204 | 1234 | 491 | 1090 | 377 | 443 | 204 | 3474 | 1751 | 28150 | 29374 | 6263 | 54250 | 14134 | 1339 | 3842 | 8687 | 234 | 1612 |
| PBANKA_0804200 | 0 | 0 | 58 | 10 | 82 | 32 | 62 | 19 | 17 | 7 | 0 | 0 | 284 | 78 | 373 | 1219 | 163 | 25 | 401 | 180 | 8 | 95 |
| PBANKA_0804400 | 11 | 26 | 79 | 16 | 525 | 237 | 586 | 210 | 455 | 191 | 100 | 64 | 763 | 405 | 996 | 5716 | 4180 | 457 | 2255 | 2248 | 57 | 559 |
| PBANKA_0804500 | 6 | 7 | 16 | 12 | 52 | 35 | 823 | 317 | 4426 | 3346 | 742 | 547 | 2247 | 1579 | 42 | 168 | 34833 | 8323 | 543 | 1012 | 87 | 875 |
| PBANKA_0804600 | 51 | 51 | 201 | 58 | 267 | 122 | 132 | 51 | 66 | 25 | 58 | 36 | 3519 | 2632 | 985 | 5015 | 547 | 112 | 382 | 379 | 111 | 746 |
| PBANKA_0804700 | 0 | 0 | 85 | 9 | 520 | 128 | 493 | 159 | 226 | 87 | 208 | 123 | 2024 | 1551 | 982 | 5228 | 1739 | 279 | 1589 | 2205 | 70 | 714 |
| PBANKA_0804800 | 22 | 34 | 59 | 10 | 102 | 53 | 102 | 44 | 43 | 16 | 174 | 107 | 725 | 331 | 217 | 882 | 520 | 97 | 165 | 152 | 37 | 352 |
| PBANKA_0804900 | 14 | 6 | 241 | 58 | 564 | 347 | 700 | 278 | 358 | 201 | 102 | 62 | 2434 | 1627 | 668 | 4037 | 3094 | 420 | 929 | 973 | 124 | 1234 |
| PBANKA_0805000 | 12 | 25 | 157 | 61 | 519 | 218 | 541 | 220 | 138 | 94 | 13 | 8 | 844 | 320 | 569 | 2432 | 1128 | 175 | 706 | 457 | 96 | 774 |
| PBANKA_0805100 | 81 | 113 | 163 | 60 | 263 | 177 | 304 | 132 | 212 | 116 | 825 | 644 | 2794 | 1575 | 559 | 2796 | 1056 | 193 | 667 | 467 | 408 | 2935 |
| PBANKA_0805200 | 26 | 48 | 42 | 13 | 98 | 71 | 122 | 31 | 54 | 17 | 6 | 5 | 160 | 41 | 363 | 1657 | 501 | 78 | 726 | 496 | 43 | 354 |
| PBANKA_0805300 | 15 | 13 | 28 | 8 | 39 | 26 | 73 | 47 | 43 | 16 | 80 | 68 | 263 | 71 | 154 | 297 | 244 | 92 | 336 | 98 | 95 | 640 |
| PBANKA_0805400 | 7 | 16 | 33 | 13 | 57 | 36 | 50 | 28 | 42 | 22 | 141 | 117 | 387 | 95 | 163 | 534 | 342 | 71 | 262 | 109 | 49 | 240 |
| PBANKA_0805500 | 0 | 6 | 202 | 33 | 322 | 199 | 307 | 116 | 142 | 58 | 19 | 0 | 938 | 379 | 553 | 3033 | 800 | 110 | 1807 | 1678 | 8 | 117 |
| PBANKA_0805600 | 0 | 6 | 102 | 20 | 175 | 103 | 154 | 67 | 43 | 37 | 9 | 5 | 650 | 218 | 322 | 1042 | 239 | 56 | 718 | 249 | 0 | 38 |
| PBANKA_0805700 | 1617 | 2400 | 13989 | 4663 | 14638 | 9227 | 12515 | 5912 | 4013 | 2142 | 3774 | 2887 | 54016 | 33392 | 34098 | 2E+05 | 25571 | 2600 | 17285 | 17314 | 1742 | 17148 |
| PBANKA_0805800 | 0 | 0 | 29 | 16 | 121 | 98 | 130 | 41 | 67 | 40 | 7 | 6 | 782 | 364 | 312 | 1215 | 876 | 105 | 115 | 110 | 13 | 123 |
| PBANKA_0805900 | 0 | 10 | 171 | 55 | 680 | 378 | 2400 | 1132 | 1028 | 625 | 139 | 47 | 1194 | 641 | 796 | 4898 | 7493 | 684 | 3681 | 4830 | 40 | 417 |
| PBANKA_0806000 | 0 | 0 | 59 | 0 | 69 | 35 | 50 | 28 | 70 | 32 | 393 | 325 | 505 | 303 | 142 | 479 | 101 | 35 | 568 | 179 | 35 | 247 |
| PBANKA_0806100 | 20 | 28 | 123 | 28 | 288 | 133 | 274 | 135 | 142 | 79 | 520 | 398 | 2136 | 1470 | 668 | 3496 | 746 | 129 | 472 | 391 | 77 | 444 |
| PBANKA_0806200 | 0 | 0 | 54 | 27 | 117 | 90 | 162 | 104 | 47 | 40 | 0 | 0 | 103 | 15 | 259 | 820 | 339 | 46 | 448 | 217 | 6 | 39 |
| PBANKA_0806300 | 8 | 5 | 13 | 0 | 216 | 125 | 234 | 85 | 97 | 51 | 6 | 12 | 233 | 161 | 680 | 2027 | 1413 | 196 | 1698 | 2586 | 142 | 1094 |
| PBANKA_0806400 | 0 | 10 | 119 | 34 | 198 | 87 | 241 | 91 | 87 | 55 | 20 | 16 | 1411 | 797 | 526 | 1957 | 1552 | 254 | 1038 | 708 | 37 | 232 |
| PBANKA_0806500 | 56 | 96 | 48 | 12 | 167 | 91 | 145 | 73 | 74 | 18 | 19 | 0 | 349 | 68 | 411 | 1511 | 226 | 50 | 869 | 380 | 88 | 669 |
| PBANKA_0806600 | 51 | 60 | 92 | 25 | 276 | 140 | 229 | 87 | 85 | 49 | 37 | 6 | 555 | 225 | 465 | 2297 | 300 | 69 | 624 | 509 | 65 | 460 |
| PBANKA_0806700 | 591 | 990 | 256 | 62 | 1255 | 471 | 1948 | 742 | 1285 | 580 | 163 | 62 | 613 | 960 | 2023 | 7228 | 21128 | 2505 | 1191 | 7315 | 275 | 1858 |
| PBANKA_0806900 | 8 | 9 | 39 | 6 | 338 | 112 | 250 | 96 | 58 | 21 | 19 | 14 | 631 | 265 | 565 | 2135 | 634 | 89 | 456 | 448 | 7 | 78 |
| PBANKA_0807000 | 47 | 69 | 560 | 170 | 1526 | 690 | 1893 | 712 | 846 | 369 | 918 | 690 | 7538 | 6536 | 5012 | 25168 | 12263 | 1371 | 6290 | 9685 | 907 | 8197 |
| PBANKA_0807100 | 0 | 0 | 192 | 55 | 219 | 111 | 178 | 82 | 52 | 25 | 92 | 45 | 1448 | 735 | 317 | 1209 | 262 | 41 | 312 | 287 | 17 | 189 |
| PBANKA_0807300 | 33 | 50 | 121 | 53 | 259 | 119 | 245 | 119 | 54 | 36 | 7 | 0 | 358 | 105 | 549 | 2249 | 685 | 85 | 175 | 132 | 47 | 273 |
| PBANKA_0807400 | 20 | 23 | 136 | 27 | 173 | 71 | 182 | 63 | 48 | 18 | 5 | 9 | 244 | 69 | 249 | 905 | 275 | 49 | 69 | 70 | 11 | 56 |
| PBANKA_0807500 | 0 | 0 | 202 | 40 | 615 | 192 | 567 | 177 | 221 | 113 | 39 | 28 | 1634 | 1482 | 2645 | 11729 | 4376 | 673 | 2715 | 6690 | 408 | 3050 |
| PBANKA_0807600 | 102 | 137 | 1133 | 331 | 582 | 489 | 645 | 385 | 234 | 224 | 1210 | 920 | 8025 | 2669 | 1880 | 7433 | 2585 | 443 | 1963 | 1045 | 468 | 3052 |
| PBANKA_0807700 | 146 | 167 | 91 | 25 | 162 | 63 | 332 | 139 | 263 | 150 | 93 | 36 | 1664 | 996 | 397 | 1700 | 5669 | 776 | 2909 | 2457 | 969 | 6218 |
| PBANKA_0807800 | 0 | 0 | 35 | 0 | 99 | 72 | 252 | 76 | 89 | 48 | 9 | 6 | 235 | 60 | 55 | 167 | 1167 | 228 | 741 | 403 | 29 | 199 |
| PBANKA_0807900 | 148 | 229 | 574 | 153 | 1587 | 579 | 2348 | 859 | 2043 | 923 | 1097 | 661 | 6385 | 16279 | 4156 | 27070 | 53783 | 6213 | 4335 | 22094 | 3331 | 20610 |
| PBANKA_0808000 | 22 | 31 | 386 | 111 | 906 | 414 | 1356 | 495 | 659 | 327 | 99 | 52 | 1727 | 648 | 415 | 1156 | 19986 | 2486 | 2391 | 2460 | 2147 | 17125 |
| PBANKA_0808100 | 850 | 1242 | 22 | 0 | 31 | 9 | 18 | 12 | 17 | 0 | 13 | 0 | 69 | 24 | 68 | 280 | 113 | 23 | 120 | 126 | 140 | 877 |
| PBANKA_0808200 | 72 | 54 | 193 | 42 | 358 | 192 | 558 | 250 | 151 | 109 | 22 | 12 | 1132 | 123 | 1194 | 3695 | 1774 | 296 | 1121 | 523 | 149 | 973 |
| PBANKA_0808300 | 7 | 12 | 70 | 13 | 170 | 46 | 162 | 66 | 119 | 69 | 26 | 15 | 2010 | 665 | 322 | 1543 | 892 | 268 | 1421 | 926 | 257 | 1746 |
| PBANKA_0808380 | 0 | 0 | 6 | 0 | 24 | 6 | 33 | 9 | 43 | 14 | 12 | 0 | 37 | 8 | 22 | 91 | 51 | 26 | 183 | 51 | 7 | 41 |
| PBANKA_0808420 | 12 | 10 | 39 | 6 | 95 | 44 | 133 | 35 | 130 | 60 | 28 | 9 | 369 | 97 | 209 | 895 | 431 | 132 | 902 | 447 | 244 | 1080 |
| PBANKA_0808500 | 0 | 0 | 33 | 8 | 37 | 23 | 28 | 9 | 16 | 10 | 0 | 0 | 120 | 41 | 99 | 168 | 142 | 26 | 77 | 25 | 12 | 62 |

| | | | | | | | | | | | | | | | | | | | | | | |
|---|---|---|---|---|---|---|---|---|---|---|---|---|---|---|---|---|---|---|---|---|---|---|
| PBANKA_0808600 | 7 | 16 | 92 | 18 | 297 | 118 | 340 | 108 | 141 | 58 | 36 | 8 | 603 | 152 | 630 | 2441 | 664 | 145 | 1124 | 800 | 1621 | 9000 |
| PBANKA_0808700 | 47 | 99 | 66 | 9 | 264 | 84 | 255 | 76 | 207 | 96 | 1842 | 1366 | 1861 | 1245 | 898 | 5759 | 1399 | 236 | 1439 | 1794 | 309 | 2141 |
| PBANKA_0808800 | 33 | 27 | 63 | 14 | 94 | 32 | 46 | 32 | 30 | 9 | 9 | 0 | 442 | 202 | 271 | 1165 | 370 | 55 | 417 | 204 | 36 | 208 |
| PBANKA_0808900 | 99 | 150 | 357 | 75 | 935 | 362 | 1407 | 509 | 524 | 247 | 135 | 56 | 3481 | 2954 | 1601 | 9390 | 5047 | 540 | 2199 | 2578 | 684 | 4436 |
| PBANKA_0809000 | 31 | 38 | 153 | 60 | 453 | 234 | 559 | 212 | 185 | 86 | 32 | 12 | 581 | 197 | 791 | 3872 | 1459 | 171 | 1686 | 2223 | 3143 | 18488 |
| PBANKA_0809100 | 0 | 0 | 78 | 16 | 196 | 79 | 286 | 130 | 97 | 52 | 42 | 9 | 1967 | 1071 | 461 | 2221 | 1929 | 246 | 361 | 258 | 57 | 330 |
| PBANKA_0809200 | 0 | 0 | 30 | 10 | 95 | 37 | 112 | 54 | 51 | 12 | 6 | 5 | 423 | 122 | 197 | 678 | 850 | 197 | 377 | 164 | 176 | 1262 |
| PBANKA_0809300 | 18 | 18 | 114 | 24 | 269 | 127 | 414 | 161 | 150 | 91 | 25 | 9 | 223 | 21 | 161 | 812 | 645 | 89 | 130 | 92 | 436 | 3501 |
| PBANKA_0809400 | 0 | 9 | 31 | 6 | 64 | 43 | 792 | 448 | 668 | 445 | 51 | 35 | 612 | 145 | 87 | 315 | 4131 | 901 | 656 | 581 | 253 | 1725 |
| PBANKA_0809500 | 0 | 0 | 20 | 0 | 178 | 57 | 334 | 121 | 320 | 148 | 39 | 15 | 115 | 23 | 17 | 90 | 313 | 43 | 185 | 75 | 103 | 584 |
| PBANKA_0809600 | 18 | 46 | 102 | 19 | 365 | 104 | 808 | 266 | 1089 | 451 | 155 | 58 | 874 | 334 | 234 | 784 | 8068 | 1151 | 896 | 1066 | 382 | 2339 |
| PBANKA_0809700 | 0 | 5 | 66 | 11 | 235 | 110 | 349 | 147 | 83 | 57 | 20 | 11 | 301 | 62 | 494 | 1231 | 626 | 194 | 1519 | 605 | 134 | 922 |
| PBANKA_0809800 | 0 | 0 | 58 | 14 | 59 | 24 | 92 | 34 | 31 | 18 | 51 | 23 | 705 | 387 | 102 | 457 | 349 | 53 | 896 | 264 | 84 | 535 |
| PBANKA_0809900 | 0 | 0 | 71 | 18 | 159 | 93 | 179 | 73 | 60 | 33 | 13 | 11 | 589 | 194 | 237 | 681 | 585 | 145 | 737 | 371 | 92 | 531 |
| PBANKA_0810000 | 13 | 29 | 51 | 11 | 109 | 28 | 116 | 48 | 60 | 20 | 49 | 34 | 970 | 455 | 164 | 1060 | 596 | 114 | 1576 | 1001 | 93 | 686 |
| PBANKA_0810100 | 18 | 17 | 22 | 9 | 82 | 39 | 78 | 37 | 72 | 25 | 28 | 0 | 545 | 173 | 109 | 858 | 155 | 41 | 805 | 319 | 31 | 153 |
| PBANKA_0810200 | 0 | 10 | 22 | 10 | 82 | 33 | 87 | 32 | 68 | 28 | 29 | 0 | 719 | 635 | 306 | 1352 | 455 | 65 | 985 | 1216 | 7580 | 53210 |
| PBANKA_0810500 | 9 | 7 | 119 | 36 | 129 | 129 | 113 | 93 | 48 | 39 | 345 | 386 | 879 | 614 | 182 | 1205 | 537 | 47 | 1114 | 624 | 82 | 575 |
| PBANKA_0810600 | 30 | 48 | 497 | 113 | 1391 | 576 | 1674 | 582 | 759 | 333 | 3142 | 1764 | 3965 | 3227 | 1353 | 6198 | 5041 | 640 | 1413 | 2189 | 198 | 1935 |
| PBANKA_0810800 | 66 | 89 | 5 | 0 | 32 | 30 | 208 | 74 | 144 | 63 | 0 | 0 | 341 | 139 | 29 | 53 | 1642 | 301 | 1100 | 570 | 66 | 468 |
| PBANKA_0810900 | 98 | 192 | 28 | 12 | 84 | 38 | 159 | 61 | 189 | 113 | 78 | 62 | 1533 | 642 | 391 | 2230 | 1195 | 289 | 3936 | 3256 | 671 | 5547 |
| PBANKA_0811000 | 6 | 0 | 70 | 16 | 149 | 94 | 145 | 74 | 42 | 22 | 54 | 30 | 733 | 352 | 313 | 921 | 461 | 67 | 272 | 165 | 8 | 91 |
| PBANKA_0811100 | 20 | 30 | 149 | 46 | 391 | 184 | 355 | 98 | 139 | 51 | 565 | 436 | 1312 | 987 | 601 | 2537 | 996 | 136 | 499 | 493 | 58 | 731 |
| PBANKA_0811200 | 0 | 0 | 5 | 0 | 17 | 8 | 21 | 17 | 53 | 19 | 6 | 10 | 277 | 83 | 214 | 925 | 223 | 53 | 243 | 432 | 11 | 103 |
| PBANKA_0811300 | 18 | 58 | 273 | 77 | 1306 | 741 | 1881 | 788 | 912 | 494 | 423 | 276 | 4378 | 1363 | 2397 | 8961 | 6771 | 1242 | 4521 | 2485 | 153 | 1227 |
| PBANKA_0811400 | 161 | 217 | 293 | 65 | 1402 | 585 | 4623 | 1901 | 4009 | 2237 | 5683 | 3245 | 2274 | 7300 | 1702 | 10893 | 13462 | 1427 | 1312 | 4971 | 423 | 3208 |
| PBANKA_0811500 | 0 | 0 | 17 | 6 | 77 | 36 | 74 | 29 | 20 | 12 | 0 | 0 | 687 | 233 | 200 | 846 | 204 | 57 | 901 | 520 | 57 | 421 |
| PBANKA_0811600 | 0 | 5 | 76 | 5 | 154 | 82 | 484 | 155 | 244 | 101 | 20 | 10 | 627 | 260 | 551 | 2898 | 837 | 110 | 563 | 365 | 14 | 128 |
| PBANKA_0811700 | 20 | 56 | 23 | 11 | 35 | 46 | 76 | 42 | 72 | 46 | 7 | 5 | 217 | 21 | 127 | 533 | 481 | 163 | 477 | 194 | 42 | 315 |
| PBANKA_0811900 | 11 | 17 | 64 | 20 | 213 | 105 | 187 | 63 | 115 | 51 | 89 | 57 | 388 | 234 | 255 | 699 | 219 | 63 | 473 | 250 | 82 | 700 |
| PBANKA_0812000 | 70 | 94 | 51 | 7 | 280 | 171 | 477 | 206 | 208 | 89 | 22 | 17 | 484 | 125 | 332 | 695 | 1926 | 290 | 1694 | 764 | 41 | 339 |
| PBANKA_0812100 | 30 | 59 | 12 | 0 | 97 | 53 | 178 | 82 | 89 | 62 | 10 | 0 | 241 | 47 | 43 | 114 | 663 | 175 | 885 | 311 | 18 | 106 |
| PBANKA_0812200 | 9 | 19 | 25 | 0 | 181 | 99 | 425 | 178 | 1142 | 861 | 232 | 165 | 553 | 304 | 63 | 147 | 1840 | 950 | 2911 | 3248 | 178 | 967 |
| PBANKA_0812300 | 95 | 112 | 48 | 19 | 253 | 139 | 551 | 224 | 218 | 115 | 33 | 21 | 277 | 44 | 68 | 82 | 1056 | 221 | 1005 | 518 | 119 | 863 |
| PBANKA_0812400 | 0 | 0 | 72 | 5 | 182 | 135 | 268 | 101 | 142 | 41 | 41 | 14 | 489 | 246 | 482 | 1656 | 793 | 117 | 184 | 203 | 17 | 168 |
| PBANKA_0812500 | 0 | 0 | 94 | 15 | 154 | 52 | 88 | 24 | 41 | 24 | 14 | 6 | 680 | 301 | 280 | 1890 | 287 | 31 | 396 | 299 | 12 | 52 |
| PBANKA_0812700 | 10 | 5 | 103 | 18 | 321 | 121 | 688 | 279 | 363 | 136 | 26 | 13 | 1725 | 400 | 667 | 2820 | 7846 | 1091 | 1813 | 1186 | 69 | 496 |
| PBANKA_0812800 | 11 | 9 | 169 | 37 | 319 | 131 | 334 | 136 | 135 | 61 | 22 | 0 | 1268 | 285 | 986 | 3944 | 1284 | 194 | 4920 | 3328 | 171 | 1247 |
| PBANKA_0812900 | 6 | 0 | 51 | 13 | 118 | 49 | 101 | 29 | 46 | 15 | 5 | 6 | 919 | 360 | 221 | 859 | 373 | 124 | 22578 | 12897 | 78 | 787 |
| PBANKA_0813000 | 3596 | 4726 | 377 | 114 | 892 | 546 | 1067 | 511 | 699 | 397 | 779 | 644 | 4549 | 1625 | 2143 | 7778 | 3674 | 897 | 3542 | 1592 | 641 | 5221 |
| PBANKA_0813100 | 240 | 295 | 170 | 42 | 519 | 206 | 991 | 385 | 402 | 147 | 69 | 35 | 610 | 272 | 298 | 1009 | 4850 | 536 | 494 | 562 | 479 | 3014 |
| PBANKA_0813200 | 0 | 0 | 43 | 6 | 115 | 55 | 127 | 42 | 58 | 26 | 13 | 0 | 663 | 274 | 188 | 1097 | 251 | 51 | 469 | 260 | 145 | 1160 |
| PBANKA_0813300 | 8 | 18 | 9 | 0 | 53 | 10 | 165 | 71 | 103 | 55 | 17 | 0 | 838 | 286 | 225 | 752 | 1411 | 404 | 17061 | 12567 | 152 | 1537 |
| PBANKA_0813400 | 38 | 33 | 80 | 17 | 260 | 152 | 237 | 80 | 122 | 54 | 21 | 20 | 619 | 148 | 677 | 2212 | 2292 | 383 | 1218 | 627 | 80 | 538 |
| PBANKA_0813500 | 54 | 64 | 158 | 48 | 345 | 164 | 382 | 145 | 97 | 69 | 39 | 29 | 1817 | 621 | 394 | 1835 | 1511 | 247 | 1012 | 672 | 110 | 685 |
| PBANKA_0813700 | 0 | 0 | 19 | 0 | 23 | 7 | 33 | 8 | 18 | 5 | 6 | 0 | 95 | 43 | 66 | 178 | 123 | 46 | 92 | 35 | 10 | 37 |
| PBANKA_0813800 | 7 | 0 | 78 | 15 | 136 | 52 | 99 | 49 | 76 | 20 | 109 | 73 | 1008 | 629 | 222 | 1438 | 375 | 40 | 1035 | 1133 | 164 | 1451 |
| PBANKA_0813900 | 0 | 0 | 151 | 38 | 325 | 153 | 443 | 183 | 173 | 79 | 20 | 9 | 888 | 369 | 776 | 2565 | 1260 | 233 | 489 | 288 | 170 | 1289 |
| PBANKA_0814000 | 0 | 5 | 76 | 24 | 125 | 71 | 99 | 65 | 63 | 47 | 837 | 759 | 1033 | 606 | 157 | 934 | 353 | 60 | 532 | 300 | 115 | 747 |
| PBANKA_0814100 | 0 | 0 | 51 | 15 | 155 | 64 | 101 | 32 | 55 | 19 | 16 | 0 | 642 | 175 | 399 | 2044 | 347 | 54 | 1090 | 453 | 38 | 319 |
| PBANKA_0814200 | 0 | 0 | 1003 | 203 | 788 | 371 | 684 | 278 | 215 | 90 | 61 | 21 | 12922 | 3097 | 5654 | 24636 | 8590 | 919 | 3333 | 2479 | 396 | 2375 |
| PBANKA_0814300 | 0 | 0 | 114 | 25 | 187 | 85 | 175 | 43 | 39 | 28 | 23 | 0 | 1173 | 487 | 509 | 2721 | 457 | 70 | 560 | 334 | 489 | 3263 |
| PBANKA_0814400 | 8 | 7 | 94 | 28 | 179 | 89 | 240 | 108 | 154 | 87 | 33 | 15 | 280 | 37 | 213 | 647 | 459 | 199 | 331 | 184 | 176 | 1166 |
| PBANKA_0814500 | 0 | 0 | 29 | 6 | 101 | 53 | 145 | 31 | 55 | 17 | 7 | 0 | 496 | 124 | 148 | 575 | 730 | 175 | 5855 | 2658 | 124 | 787 |
| PBANKA_0814600 | 11 | 0 | 38 | 7 | 161 | 63 | 160 | 51 | 50 | 34 | 26 | 0 | 378 | 119 | 214 | 913 | 761 | 118 | 805 | 312 | 8 | 60 |
| PBANKA_0814700 | 12 | 23 | 126 | 23 | 319 | 147 | 336 | 126 | 131 | 52 | 30 | 15 | 888 | 210 | 422 | 2144 | 959 | 124 | 308 | 236 | 33 | 222 |
| PBANKA_0814800 | 170 | 145 | 15 | 6 | 26 | 24 | 39 | 25 | 185 | 177 | 124 | 57 | 184 | 62 | 51 | 108 | 236 | 281 | 202 | 95 | 154 | 1055 |
| PBANKA_0814900 | 0 | 7 | 46 | 10 | 172 | 77 | 218 | 70 | 119 | 42 | 24 | 7 | 541 | 170 | 359 | 2032 | 1429 | 168 | 837 | 302 | 184 | 970 |
| PBANKA_0815000 | 21 | 26 | 33 | 8 | 141 | 36 | 107 | 44 | 110 | 39 | 0 | 0 | 550 | 243 | 209 | 1102 | 831 | 125 | 467 | 305 | 565 | 2880 |
| PBANKA_0815100 | 167 | 268 | 11 | 0 | 83 | 20 | 87 | 37 | 742 | 355 | 232 | 116 | 838 | 705 | 72 | 337 | 3445 | 1156 | 8002 | 12907 | 363 | 1682 |
| PBANKA_0815200 | 8 | 13 | 27 | 15 | 225 | 134 | 450 | 189 | 209 | 136 | 26 | 27 | 2697 | 925 | 621 | 2790 | 3750 | 659 | 1864 | 1045 | 285 | 2403 |
| PBANKA_0815300 | 0 | 0 | 147 | 39 | 185 | 90 | 210 | 54 | 119 | 62 | 250 | 164 | 2319 | 1668 | 470 | 2263 | 893 | 149 | 1275 | 1159 | 788 | 4979 |
| PBANKA_0815400 | 0 | 0 | 6 | 0 | 10 | 6 | 59 | 47 | 339 | 227 | 53 | 57 | 871 | 345 | 25 | 83 | 1515 | 1037 | 24623 | 9021 | 370 | 2394 |
| PBANKA_0815500 | 17 | 30 | 21 | 0 | 128 | 44 | 137 | 48 | 245 | 143 | 103 | 47 | 518 | 402 | 239 | 808 | 1722 | 344 | 3490 | 8190 | 285 | 1424 |
| PBANKA_0815600 | 38 | 40 | 24 | 6 | 40 | 15 | 54 | 22 | 57 | 45 | 13 | 10 | 334 | 65 | 99 | 522 | 362 | 76 | 564 | 309 | 48 | 294 |
| PBANKA_0815700 | 917 | 1072 | 909 | 226 | 1170 | 645 | 1046 | 395 | 514 | 266 | 71 | 34 | 1447 | 438 | 265 | 1085 | 3599 | 869 | 19461 | 16048 | 775 | 5634 |
| PBANKA_0815800 | 55 | 42 | 50 | 9 | 68 | 32 | 115 | 39 | 58 | 38 | 8 | 10 | 86 | 35 | 25 | 49 | 469 | 128 | 837 | 357 | 82 | 729 |
| PBANKA_0815900 | 134 | 207 | 55 | 14 | 253 | 86 | 442 | 157 | 236 | 96 | 29 | 17 | 1145 | 368 | 271 | 1159 | 3496 | 695 | 8671 | 7490 | 163 | 1460 |
| PBANKA_0816000 | 0 | 0 | 106 | 24 | 336 | 116 | 500 | 214 | 310 | 115 | 34 | 23 | 480 | 165 | 230 | 626 | 1908 | 505 | 1613 | 727 | 103 | 661 |
| PBANKA_0816100 | 7 | 5 | 109 | 44 | 198 | 114 | 199 | 122 | 72 | 54 | 44 | 28 | 406 | 137 | 231 | 400 | 339 | 96 | 437 | 127 | 28 | 168 |
| PBANKA_0816200 | 181 | 243 | 10 | 0 | 59 | 16 | 46 | 22 | 27 | 9 | 6 | 0 | 463 | 111 | 163 | 452 | 624 | 115 | 2716 | 1643 | 357 | 2026 |
| PBANKA_0816300 | 41 | 63 | 51 | 11 | 164 | 78 | 183 | 47 | 47 | 22 | 25 | 9 | 807 | 404 | 249 | 936 | 367 | 113 | 5104 | 3082 | 140 | 740 |
| PBANKA_0816400 | 75 | 118 | 652 | 148 | 879 | 446 | 960 | 300 | 394 | 125 | 2374 | 1148 | 10576 | 6074 | 3294 | 16639 | 2756 | 374 | 2934 | 2502 | 2419 | 19464 |
| PBANKA_0816500 | 0 | 0 | 23 | 0 | 46 | 21 | 44 | 20 | 49 | 18 | 8 | 15 | 420 | 177 | 81 | 650 | 119 | 27 | 704 | 384 | 1404 | 8170 |
| PBANKA_0816600 | 0 | 0 | 73 | 11 | 120 | 55 | 103 | 49 | 103 | 49 | 248 | 149 | 2307 | 1733 | 334 | 2636 | 1439 | 216 | 1768 | 1787 | 75 | 268 |
| PBANKA_0816700 | 6 | 9 | 17 | 8 | 206 | 103 | 283 | 150 | 137 | 122 | 5 | 16 | 258 | 80 | 128 | 350 | 1216 | 294 | 2084 | 1124 | 141 | 1015 |
| PBANKA_0816800 | 76 | 101 | 5 | 0 | 20 | 10 | 42 | 19 | 28 | 12 | 17 | 5 | 371 | 93 | 32 | 123 | 269 | 92 | 10400 | 6150 | 278 | 1999 |
| PBANKA_0816900 | 0 | 0 | 49 | 9 | 138 | 79 | 299 | 137 | 104 | 69 | 10 | 12 | 1506 | 352 | 237 | 1138 | 2415 | 368 | 2998 | 2022 | 286 | 1881 |
| PBANKA_0817100 | 20 | 22 | 11 | 0 | 36 | 20 | 72 | 42 | 48 | 29 | 0 | 0 | 227 | 65 | 95 | 221 | 379 | 123 | 2011 | 903 | 196 | 1496 |
| PBANKA_0817200 | 10 | 7 | 175 | 47 | 270 | 141 | 256 | 84 | 129 | 68 | 65 | 36 | 2241 | 1909 | 901 | 5126 | 780 | 155 | 1049 | 591 | 363 | 2133 |
| PBANKA_0817300 | 87 | 80 | 8 | 7 | 72 | 45 | 108 | 43 | 218 | 129 | 22 | 46 | 37 | 26 | 26 | 35 | 257 | 101 | 317 | 245 | 47 | 344 |
| PBANKA_0817400 | 334 | 490 | 165 | 32 | 456 | 187 | 559 | 185 | 543 | 284 | 1590 | 996 | 3370 | 3020 | 1587 | 8466 | 3448 | 611 | 2240 | 2170 | 128 | 636 |
| PBANKA_0817500 | 0 | 0 | 50 | 13 | 127 | 49 | 122 | 37 | 15 | 11 | 13 | 0 | 142 | 23 | 104 | 399 | 208 | 34 | 45 | 16 | 0 | 23 |
| PBANKA_0817600 | 0 | 0 | 60 | 14 | 72 | 44 | 73 | 40 | 45 | 28 | 22 | 17 | 423 | 263 | 105 | 436 | 132 | 28 | 95 | 78 | 21 | 112 |
| PBANKA_0817700 | 443 | 743 | 38 | 8 | 158 | 46 | 164 | 65 | 196 | 114 | 1479 | 1189 | 4934 | 3136 | 746 | 4636 | 2856 | 463 | 2237 | 2363 | 277 | 2738 |
| PBANKA_0817800 | 6 | 0 | 9 | 0 | 33 | 11 | 44 | 13 | 13 | 9 | 0 | 0 | 264 | 74 | 87 | 399 | 219 | 39 | 131 | 49 | 5 | 24 |
| PBANKA_0817900 | 6 | 0 | 60 | 20 | 247 | 130 | 501 | 222 | 490 | 305 | 1746 | 1182 | 711 | 465 | 123 | 384 | 2007 | 356 | 167 | 167 | 239 | 1751 |
| PBANKA_0818000 | 9 | 34 | 83 | 15 | 499 | 147 | 853 | 274 | 401 | 157 | 56 | 16 | 687 | 515 | 730 | 2267 | 8286 | 987 | 1660 | 5479 | 123 | 821 |
| PBANKA_0818100 | 14 | 31 | 70 | 17 | 109 | 34 | 104 | 49 | 125 | 46 | 14 | 8 | 196 | 65 | 211 | 1247 | 1046 | 183 | 1029 | 703 | 61 | 387 |
| PBANKA_0818200 | 9 | 6 | 9 | 0 | 105 | 28 | 91 | 34 | 95 | 46 | 19 | 0 | 51 | 12 | 5 | 32 | 141 | 35 | 1077 | 273 | 91 | 730 |

| | | | | | | | | | | | | | | | | | | | | | | |
|---|---|---|---|---|---|---|---|---|---|---|---|---|---|---|---|---|---|---|---|---|---|---|
| PBANKA_0818300 | 0 | 0 | 6 | 0 | 20 | 10 | 50 | 30 | 135 | 101 | 19 | 8 | 92 | 17 | 0 | 0 | 949 | 346 | 125 | 71 | 19 | 140 |
| PBANKA_0818500 | 49 | 108 | 71 | 14 | 554 | 266 | 983 | 360 | 764 | 380 | 79 | 32 | 507 | 199 | 314 | 604 | 3186 | 615 | 1982 | 1801 | 74 | 499 |
| PBANKA_0818600 | 0 | 0 | 144 | 14 | 260 | 154 | 279 | 97 | 114 | 59 | 514 | 294 | 2086 | 1561 | 773 | 3124 | 1183 | 155 | 329 | 317 | 5 | 138 |
| PBANKA_0818700 | 0 | 0 | 13 | 6 | 53 | 30 | 62 | 22 | 15 | 12 | 61 | 40 | 176 | 71 | 100 | 271 | 117 | 20 | 147 | 36 | 9 | 79 |
| PBANKA_0818800 | 0 | 0 | 9 | 0 | 27 | 12 | 48 | 14 | 26 | 12 | 7 | 0 | 226 | 56 | 92 | 427 | 299 | 41 | 472 | 210 | 8 | 55 |
| PBANKA_0818900 | 24 | 35 | 2902 | 941 | 5110 | 3417 | 9303 | 4143 | 3001 | 1690 | 1274 | 1083 | 28262 | 9127 | 9363 | 40852 | 46893 | 6405 | 11349 | 5747 | 2008 | 22189 |
| PBANKA_0819100 | 0 | 0 | 63 | 14 | 115 | 71 | 92 | 37 | 56 | 45 | 76 | 55 | 485 | 246 | 455 | 1016 | 163 | 99 | 282 | 202 | 63 | 439 |
| PBANKA_0819200 | 68 | 67 | 128 | 25 | 249 | 128 | 395 | 180 | 300 | 142 | 46 | 61 | 1452 | 362 | 628 | 2544 | 2010 | 470 | 1928 | 749 | 164 | 1350 |
| PBANKA_0819300 | 42 | 48 | 440 | 118 | 453 | 246 | 485 | 208 | 141 | 84 | 529 | 478 | 3813 | 1587 | 1350 | 4862 | 935 | 205 | 1512 | 843 | 303 | 2270 |
| PBANKA_0819400 | 103 | 131 | 20 | 0 | 96 | 34 | 88 | 42 | 60 | 29 | 12 | 0 | 504 | 232 | 117 | 431 | 229 | 69 | 2834 | 2502 | 127 | 1064 |
| PBANKA_0819450 | 299 | 346 | 386 | 100 | 665 | 408 | 760 | 361 | 417 | 217 | 75 | 45 | 500 | 112 | 770 | 2441 | 2945 | 565 | 2873 | 1044 | 776 | 4918 |
| PBANKA_0819500 | 148 | 233 | 18 | 0 | 148 | 63 | 344 | 118 | 180 | 78 | 20 | 8 | 367 | 98 | 112 | 437 | 1850 | 325 | 674 | 649 | 229 | 1600 |
| PBANKA_0819600 | 0 | 0 | 40 | 9 | 96 | 78 | 82 | 33 | 296 | 238 | 92 | 44 | 154 | 134 | 29 | 53 | 843 | 506 | 980 | 355 | 1264 | 9450 |
| PBANKA_0819700 | 0 | 0 | 223 | 33 | 159 | 76 | 120 | 43 | 87 | 32 | 522 | 391 | 1704 | 998 | 385 | 1624 | 532 | 87 | 1406 | 880 | 467 | 3458 |
| PBANKA_0819800 | 162 | 189 | 118 | 26 | 399 | 196 | 341 | 115 | 221 | 131 | 486 | 286 | 1738 | 1135 | 1062 | 4554 | 2090 | 393 | 2692 | 2629 | 847 | 6482 |
| PBANKA_0819900 | 80 | 80 | 1148 | 336 | 1001 | 734 | 1019 | 576 | 407 | 274 | 2722 | 2533 | 6813 | 2703 | 2816 | 8148 | 4438 | 912 | 5016 | 2606 | 116 | 953 |
| PBANKA_0820000 | 65 | 96 | 224 | 62 | 497 | 298 | 936 | 465 | 330 | 225 | 268 | 141 | 2144 | 919 | 961 | 2458 | 2635 | 488 | 983 | 485 | 130 | 962 |
| PBANKA_0820200 | 583 | 729 | 5 | 5 | 63 | 40 | 79 | 24 | 43 | 23 | 9 | 7 | 17 | 0 | 10 | 12 | 68 | 23 | 136 | 85 | 10214 | 89421 |
| PBANKA_0820300 | 19 | 18 | 199 | 48 | 376 | 222 | 333 | 134 | 92 | 45 | 30 | 8 | 270 | 60 | 621 | 2188 | 1208 | 194 | 929 | 496 | 32 | 275 |
| PBANKA_0820400 | 539 | 782 | 14 | 0 | 127 | 51 | 105 | 37 | 49 | 21 | 11 | 0 | 27 | 9 | 20 | 112 | 38 | 8 | 150 | 96 | 8 | 87 |
| PBANKA_0820500 | 0 | 5 | 21 | 0 | 78 | 20 | 53 | 17 | 27 | 10 | 10 | 0 | 160 | 87 | 94 | 532 | 102 | 16 | 160 | 63 | 5 | 29 |
| PBANKA_0820600 | 92 | 125 | 23 | 6 | 80 | 37 | 64 | 29 | 30 | 10 | 0 | 0 | 166 | 62 | 149 | 483 | 77 | 26 | 95 | 63 | 227 | 1659 |
| PBANKA_0820700 | 83 | 180 | 15 | 0 | 87 | 36 | 360 | 145 | 386 | 201 | 106 | 54 | 868 | 488 | 383 | 1981 | 5001 | 796 | 2172 | 2388 | 467 | 3619 |
| PBANKA_0820800 | 75 | 89 | 111 | 35 | 1063 | 393 | 2308 | 885 | 1613 | 710 | 614 | 342 | 2312 | 2168 | 240 | 1013 | 42753 | 6109 | 1766 | 4506 | 537 | 3283 |
| PBANKA_0820900 | 85 | 82 | 75 | 17 | 479 | 174 | 586 | 152 | 221 | 54 | 33 | 15 | 47 | 24 | 20 | 18 | 49 | 8 | 42 | 31 | 24 | 162 |
| PBANKA_0821000 | 0 | 0 | 23 | 0 | 48 | 21 | 42 | 16 | 8 | 9 | 5 | 0 | 343 | 134 | 77 | 367 | 66 | 14 | 295 | 150 | 534 | 2283 |
| PBANKA_0821100 | 0 | 0 | 63 | 22 | 281 | 130 | 511 | 219 | 182 | 80 | 37 | 8 | 565 | 215 | 695 | 3005 | 3194 | 435 | 836 | 2024 | 212 | 1479 |
| PBANKA_0821300 | 7 | 0 | 41 | 5 | 99 | 28 | 286 | 88 | 108 | 67 | 23 | 7 | 207 | 73 | 150 | 803 | 515 | 63 | 1781 | 1045 | 328 | 1581 |
| PBANKA_0821500 | 36 | 16 | 84 | 17 | 228 | 64 | 349 | 116 | 114 | 37 | 29 | 12 | 513 | 136 | 327 | 3185 | 1670 | 180 | 1167 | 928 | 452 | 2644 |
| PBANKA_0821600 | 70 | 85 | 47 | 0 | 134 | 39 | 284 | 103 | 153 | 60 | 43 | 19 | 654 | 248 | 258 | 967 | 1890 | 313 | 2165 | 1253 | 126 | 605 |
| PBANKA_0821700 | 0 | 0 | 250 | 92 | 276 | 195 | 315 | 104 | 68 | 36 | 101 | 67 | 3614 | 1833 | 1661 | 6321 | 1534 | 163 | 1447 | 1029 | 123 | 1128 |
| PBANKA_0821800 | 20 | 22 | 100 | 19 | 219 | 82 | 224 | 57 | 66 | 27 | 37 | 29 | 938 | 351 | 412 | 1762 | 498 | 107 | 796 | 513 | 228 | 1783 |
| PBANKA_0821900 | 75 | 113 | 119 | 47 | 167 | 121 | 185 | 87 | 57 | 36 | 7 | 8 | 465 | 127 | 192 | 333 | 104 | 39 | 511 | 142 | 104 | 644 |
| PBANKA_0822000 | 0 | 5 | 25 | 5 | 131 | 44 | 235 | 113 | 234 | 132 | 39 | 18 | 137 | 21 | 108 | 351 | 597 | 181 | 429 | 218 | 38 | 263 |
| PBANKA_0822100 | 32 | 69 | 158 | 26 | 275 | 98 | 455 | 175 | 349 | 150 | 705 | 377 | 3108 | 2138 | 657 | 2695 | 2202 | 476 | 2620 | 1365 | 240 | 1501 |
| PBANKA_0822200 | 14 | 12 | 25 | 6 | 98 | 68 | 116 | 75 | 64 | 34 | 34 | 13 | 550 | 218 | 234 | 800 | 475 | 106 | 778 | 397 | 79 | 661 |
| PBANKA_0822300 | 0 | 0 | 31 | 0 | 96 | 47 | 161 | 52 | 49 | 32 | 11 | 0 | 86 | 20 | 155 | 485 | 515 | 88 | 138 | 90 | 54 | 421 |
| PBANKA_0822400 | 0 | 0 | 37 | 0 | 116 | 44 | 145 | 38 | 79 | 36 | 9 | 7 | 177 | 37 | 260 | 872 | 1083 | 192 | 139 | 104 | 120 | 783 |
| PBANKA_0822500 | 194 | 314 | 107 | 19 | 440 | 142 | 530 | 161 | 222 | 93 | 129 | 81 | 1736 | 1872 | 718 | 4185 | 2479 | 300 | 1133 | 2081 | 669 | 5113 |
| PBANKA_0822600 | 28 | 27 | 63 | 10 | 209 | 77 | 249 | 97 | 94 | 38 | 26 | 11 | 938 | 311 | 559 | 2848 | 2576 | 391 | 1123 | 730 | 171 | 1157 |
| PBANKA_0822700 | 0 | 0 | 72 | 11 | 115 | 47 | 95 | 31 | 59 | 52 | 427 | 330 | 883 | 643 | 224 | 777 | 231 | 69 | 1357 | 863 | 199 | 1185 |
| PBANKA_0822800 | 0 | 5 | 109 | 24 | 111 | 46 | 113 | 42 | 73 | 40 | 501 | 299 | 3879 | 3047 | 688 | 3459 | 1157 | 286 | 1652 | 1571 | 94 | 785 |
| PBANKA_0823000 | 192 | 309 | 299 | 86 | 937 | 351 | 1062 | 410 | 533 | 216 | 198 | 82 | 7328 | 6086 | 3654 | 20432 | 14462 | 1849 | 6617 | 9282 | 526 | 3802 |
| PBANKA_0823100 | 1177 | 1522 | 209 | 81 | 221 | 140 | 156 | 72 | 510 | 376 | 1598 | 1574 | 5183 | 2672 | 645 | 3329 | 1063 | 653 | 28168 | 16907 | 1609 | 12367 |
| PBANKA_0823300 | 124 | 170 | 48 | 8 | 120 | 44 | 115 | 38 | 67 | 17 | 31 | 10 | 840 | 364 | 552 | 2508 | 787 | 184 | 2555 | 2438 | 611 | 4307 |
| PBANKA_0823400 | 6 | 18 | 1846 | 446 | 1554 | 833 | 1457 | 484 | 439 | 195 | 176 | 66 | 17788 | 5953 | 6876 | 31378 | 6119 | 876 | 3764 | 2461 | 95 | 979 |
| PBANKA_0823500 | 66 | 134 | 58 | 12 | 192 | 115 | 198 | 89 | 86 | 58 | 245 | 150 | 1787 | 679 | 1049 | 4508 | 1077 | 230 | 1357 | 948 | 62 | 661 |
| PBANKA_0823600 | 0 | 0 | 32 | 7 | 53 | 14 | 65 | 28 | 19 | 11 | 56 | 24 | 1563 | 817 | 332 | 1874 | 708 | 135 | 1517 | 1286 | 109 | 734 |
| PBANKA_0823700 | 115 | 132 | 184 | 26 | 567 | 184 | 806 | 236 | 703 | 294 | 182 | 40 | 819 | 598 | 611 | 4123 | 5005 | 646 | 1506 | 3887 | 732 | 3699 |
| PBANKA_0823800 | 0 | 0 | 160 | 38 | 460 | 229 | 478 | 247 | 129 | 92 | 15 | 8 | 39 | 14 | 27 | 29 | 107 | 15 | 67 | 35 | 149 | 942 |
| PBANKA_0823900 | 8 | 9 | 77 | 38 | 181 | 95 | 145 | 74 | 54 | 24 | 169 | 151 | 788 | 615 | 427 | 1585 | 363 | 60 | 145 | 144 | 71 | 538 |
| PBANKA_0824000 | 0 | 0 | 9 | 0 | 25 | 13 | 21 | 6 | 11 | 8 | 0 | 0 | 71 | 25 | 5 | 30 | 30 | 15 | 588 | 305 | 47 | 266 |
| PBANKA_0824100 | 0 | 5 | 62 | 16 | 99 | 43 | 49 | 37 | 43 | 8 | 29 | 23 | 405 | 334 | 71 | 267 | 113 | 31 | 330 | 309 | 262 | 1524 |
| PBANKA_0824200 | 10 | 11 | 5 | 0 | 33 | 0 | 26 | 16 | 28 | 19 | 29 | 6 | 165 | 124 | 12 | 86 | 80 | 31 | 2238 | 2417 | 8172 | 60595 |
| PBANKA_0824300 | 330 | 557 | 153 | 40 | 461 | 163 | 642 | 231 | 236 | 146 | 324 | 173 | 2014 | 2576 | 549 | 3984 | 3778 | 404 | 1847 | 6266 | 490 | 2913 |
| PBANKA_0824500 | 17 | 21 | 13 | 0 | 64 | 24 | 68 | 31 | 23 | 11 | 5 | 0 | 64 | 10 | 31 | 56 | 219 | 45 | 269 | 116 | 837 | 5059 |
| PBANKA_0824700 | 0 | 5 | 143 | 48 | 320 | 140 | 550 | 286 | 640 | 274 | 134 | 67 | 892 | 180 | 894 | 3721 | 1970 | 295 | 2948 | 1478 | 664 | 5582 |
| PBANKA_0824800 | 61 | 78 | 178 | 50 | 218 | 162 | 243 | 112 | 170 | 123 | 470 | 524 | 1067 | 690 | 589 | 1181 | 413 | 129 | 1005 | 502 | 132 | 867 |
| PBANKA_0824900 | 82 | 100 | 59 | 8 | 182 | 55 | 228 | 57 | 559 | 268 | 230 | 125 | 1333 | 739 | 524 | 3237 | 1636 | 330 | 2629 | 2663 | 637 | 4473 |
| PBANKA_0825000 | 0 | 0 | 49 | 10 | 161 | 46 | 127 | 44 | 87 | 35 | 65 | 23 | 2427 | 2049 | 591 | 4159 | 768 | 145 | 597 | 531 | 1395 | 7494 |
| PBANKA_0825100 | 0 | 5 | 61 | 5 | 132 | 60 | 123 | 45 | 56 | 18 | 23 | 0 | 866 | 488 | 305 | 1484 | 281 | 46 | 516 | 274 | 499 | 3615 |
| PBANKA_0825200 | 0 | 0 | 5 | 0 | 25 | 12 | 60 | 18 | 30 | 17 | 0 | 8 | 410 | 207 | 93 | 573 | 406 | 81 | 1228 | 822 | 351 | 1961 |
| PBANKA_0825300 | 130 | 239 | 120 | 30 | 186 | 117 | 311 | 99 | 202 | 118 | 89 | 39 | 1414 | 967 | 345 | 2192 | 2139 | 317 | 1020 | 1135 | 1688 | 11181 |
| PBANKA_0825500 | 7 | 6 | 6 | 0 | 73 | 22 | 160 | 76 | 172 | 102 | 27 | 15 | 863 | 1052 | 40 | 179 | 1821 | 383 | 15658 | 40641 | 1540 | 9689 |
| PBANKA_0825600 | 17 | 24 | 68 | 14 | 216 | 80 | 200 | 102 | 86 | 58 | 142 | 74 | 1360 | 1047 | 414 | 1951 | 1137 | 193 | 1261 | 626 | 276 | 2117 |
| PBANKA_0825900 | 1185 | 1816 | 8 | 0 | 497 | 190 | 2907 | 1260 | 2195 | 1078 | 204 | 106 | 1081 | 457 | 13 | 71 | 330 | 140 | 22518 | 16171 | 2647 | 20118 |
| PBANKA_0826000 | 16 | 25 | 48 | 8 | 278 | 68 | 713 | 240 | 525 | 212 | 408 | 219 | 1898 | 1003 | 329 | 1578 | 2331 | 389 | 477 | 351 | 231 | 1155 |
| PBANKA_0826100 | 18 | 45 | 35 | 6 | 168 | 60 | 281 | 89 | 149 | 38 | 28 | 5 | 949 | 244 | 255 | 1476 | 1202 | 189 | 118 | 100 | 56 | 337 |
| PBANKA_0826200 | 17 | 36 | 31 | 13 | 276 | 154 | 402 | 161 | 225 | 75 | 42 | 7 | 452 | 121 | 345 | 1674 | 1108 | 188 | 2496 | 1523 | 253 | 2418 |
| PBANKA_0826300 | 0 | 0 | 39 | 15 | 64 | 48 | 76 | 45 | 13 | 8 | 0 | 0 | 35 | 17 | 126 | 206 | 107 | 31 | 187 | 75 | 62 | 252 |
| PBANKA_0826400 | 79 | 53 | 11 | 5 | 117 | 57 | 335 | 139 | 143 | 99 | 17 | 0 | 261 | 103 | 205 | 888 | 805 | 133 | 1502 | 612 | 89 | 605 |
| PBANKA_0826500 | 469 | 659 | 278 | 64 | 620 | 345 | 1446 | 533 | 772 | 370 | 200 | 126 | 2016 | 408 | 939 | 3264 | 4019 | 684 | 851 | 618 | 1686 | 14141 |
| PBANKA_0826600 | 15 | 15 | 19 | 0 | 135 | 27 | 188 | 61 | 143 | 45 | 28 | 17 | 211 | 148 | 74 | 246 | 476 | 124 | 1379 | 1536 | 230 | 1719 |
| PBANKA_0826700 | 8 | 6 | 11 | 0 | 79 | 37 | 172 | 66 | 202 | 108 | 2558 | 2373 | 26898 | 12831 | 3592 | 11388 | 4735 | 866 | 2403 | 1720 | 59 | 620 |
| PBANKA_0826800 | 155 | 201 | 42 | 7 | 189 | 87 | 674 | 267 | 586 | 265 | 62 | 35 | 1016 | 701 | 555 | 2387 | 638 | 172 | 3665 | 3143 | 823 | 4830 |
| PBANKA_0826900 | 77 | 119 | 394 | 97 | 2163 | 848 | 2442 | 764 | 1136 | 418 | 4478 | 2513 | 4676 | 6516 | 2705 | 13330 | 6811 | 860 | 2593 | 4255 | 467 | 3287 |
| PBANKA_0827000 | 0 | 11 | 10 | 0 | 27 | 9 | 37 | 16 | 46 | 15 | 15 | 7 | 49 | 14 | 37 | 61 | 65 | 47 | 90 | 28 | 7 | 36 |
| PBANKA_0827100 | 44 | 63 | 14 | 0 | 112 | 56 | 449 | 196 | 406 | 190 | 94 | 38 | 738 | 366 | 163 | 739 | 1149 | 292 | 4597 | 3812 | 166 | 1340 |
| PBANKA_0827200 | 5 | 9 | 91 | 18 | 804 | 349 | 1688 | 637 | 682 | 298 | 65 | 31 | 1843 | 217 | 920 | 3219 | 8526 | 1255 | 399 | 236 | 57 | 373 |
| PBANKA_0827300 | 0 | 0 | 59 | 23 | 182 | 89 | 209 | 85 | 154 | 78 | 39 | 40 | 322 | 373 | 134 | 604 | 570 | 102 | 615 | 2197 | 1108 | 6477 |
| PBANKA_0827500 | 0 | 0 | 108 | 28 | 433 | 250 | 756 | 320 | 309 | 161 | 764 | 720 | 3225 | 3098 | 1357 | 6063 | 4461 | 544 | 326 | 945 | 265 | 1676 |
| PBANKA_0827600 | 0 | 0 | 90 | 16 | 256 | 115 | 487 | 185 | 247 | 136 | 48 | 22 | 472 | 308 | 274 | 1389 | 3340 | 362 | 1149 | 4658 | 1442 | 10160 |
| PBANKA_0827700 | 0 | 0 | 42 | 10 | 94 | 63 | 157 | 74 | 54 | 26 | 8 | 0 | 245 | 36 | 293 | 821 | 658 | 153 | 284 | 215 | 143 | 629 |
| PBANKA_0827800 | 0 | 0 | 21 | 7 | 84 | 56 | 136 | 73 | 65 | 34 | 5 | 5 | 203 | 28 | 106 | 430 | 709 | 129 | 618 | 293 | 30 | 244 |
| PBANKA_0827900 | 0 | 0 | 36 | 18 | 76 | 56 | 134 | 62 | 49 | 25 | 8 | 0 | 175 | 34 | 197 | 714 | 626 | 101 | 676 | 272 | 119 | 774 |
| PBANKA_0828000 | 0 | 0 | 147 | 36 | 302 | 141 | 392 | 119 | 170 | 89 | 31 | 12 | 353 | 289 | 486 | 2330 | 863 | 224 | 4157 | 9435 | 2116 | 16179 |
| PBANKA_0828100 | 8 | 11 | 39 | 5 | 195 | 74 | 201 | 69 | 186 | 65 | 29 | 18 | 458 | 159 | 149 | 598 | 1799 | 581 | 3801 | 3778 | 326 | 2391 |
| PBANKA_0828200 | 23 | 40 | 34 | 0 | 97 | 30 | 73 | 26 | 62 | 30 | 5 | 8 | 517 | 155 | 30 | 116 | 633 | 220 | 4134 | 3783 | 196 | 1312 |
| PBANKA_0828400 | 71 | 104 | 102 | 19 | 347 | 163 | 702 | 288 | 325 | 145 | 68 | 36 | 1359 | 478 | 329 | 1223 | 6911 | 1281 | 4387 | 3210 | 190 | 1351 |
| PBANKA_0828500 | 10 | 8 | 56 | 15 | 152 | 70 | 154 | 72 | 72 | 29 | 15 | 15 | 293 | 77 | 202 | 770 | 686 | 200 | 2636 | 1330 | 83 | 596 |

| | | | | | | | | | | | | | | | | | | | | | | |
|---|---|---|---|---|---|---|---|---|---|---|---|---|---|---|---|---|---|---|---|---|---|---|
| PBANKA_0828600 | 0 | 0 | 79 | 25 | 250 | 130 | 356 | 159 | 145 | 111 | 12 | 9 | 539 | 96 | 464 | 2050 | 2262 | 354 | 1577 | 897 | 79 | 656 |
| PBANKA_0828700 | 18 | 23 | 113 | 22 | 750 | 394 | 1277 | 539 | 458 | 263 | 29 | 16 | 385 | 107 | 406 | 1660 | 2393 | 277 | 723 | 409 | 420 | 3109 |
| PBANKA_0828800 | 27 | 23 | 37 | 16 | 311 | 224 | 516 | 449 | 238 | 221 | 38 | 21 | 105 | 30 | 67 | 157 | 246 | 107 | 974 | 381 | 120 | 611 |
| PBANKA_0828900 | 0 | 0 | 7 | 0 | 145 | 81 | 116 | 48 | 24 | 14 | 20 | 0 | 428 | 195 | 61 | 96 | 343 | 65 | 2011 | 2080 | 30 | 241 |
| PBANKA_0829000 | 22 | 19 | 32 | 10 | 95 | 43 | 71 | 41 | 50 | 23 | 0 | 7 | 617 | 223 | 445 | 1825 | 390 | 74 | 1127 | 728 | 144 | 848 |
| PBANKA_0829100 | 100 | 170 | 112 | 20 | 177 | 127 | 240 | 94 | 83 | 47 | 17 | 5 | 487 | 171 | 295 | 1439 | 304 | 75 | 425 | 211 | 78 | 499 |
| PBANKA_0829200 | 15 | 28 | 15 | 0 | 52 | 23 | 162 | 62 | 112 | 56 | 31 | 10 | 546 | 160 | 129 | 561 | 1780 | 450 | 593 | 290 | 1077 | 7865 |
| PBANKA_0829300 | 25 | 33 | 19 | 0 | 23 | 26 | 45 | 18 | 23 | 16 | 19 | 9 | 103 | 49 | 38 | 168 | 235 | 67 | 184 | 125 | 318 | 1678 |
| PBANKA_0829400 | 0 | 0 | 5 | 0 | 8 | 0 | 19 | 9 | 10 | 0 | 0 | 0 | 221 | 44 | 54 | 173 | 108 | 27 | 597 | 148 | 53 | 368 |
| PBANKA_0829500 | 0 | 0 | 125 | 32 | 139 | 58 | 120 | 38 | 110 | 56 | 24 | 8 | 1786 | 782 | 578 | 3334 | 864 | 189 | 808 | 577 | 571 | 2659 |
| PBANKA_0829600 | 0 | 0 | 5 | 0 | 25 | 6 | 12 | 12 | 28 | 15 | 5 | 0 | 159 | 41 | 33 | 121 | 33 | 14 | 635 | 538 | 838 | 5057 |
| PBANKA_0829700 | 0 | 0 | 67 | 11 | 65 | 21 | 31 | 13 | 13 | 7 | 73 | 60 | 2057 | 1387 | 97 | 594 | 81 | 24 | 1140 | 644 | 34 | 184 |
| PBANKA_0829800 | 65 | 80 | 62 | 20 | 316 | 162 | 340 | 141 | 137 | 65 | 294 | 156 | 490 | 148 | 429 | 1789 | 1041 | 140 | 1746 | 722 | 624 | 4038 |
| PBANKA_0829900 | 0 | 0 | 18 | 0 | 53 | 20 | 53 | 18 | 18 | 11 | 0 | 0 | 243 | 28 | 84 | 462 | 268 | 39 | 971 | 262 | 184 | 887 |
| PBANKA_0830000 | 0 | 0 | 125 | 36 | 207 | 91 | 258 | 139 | 89 | 65 | 34 | 24 | 1804 | 749 | 494 | 1744 | 1236 | 210 | 1355 | 529 | 285 | 1962 |
| PBANKA_0830100 | 0 | 0 | 36 | 8 | 140 | 60 | 171 | 35 | 85 | 35 | 8 | 0 | 640 | 115 | 330 | 1817 | 923 | 209 | 1631 | 935 | 77 | 412 |
| PBANKA_0830200 | 24 | 29 | 91 | 19 | 2926 | 1816 | 9493 | 4888 | 3808 | 2581 | 224 | 123 | 9464 | 3631 | 8677 | 27408 | 83196 | 10217 | 3640 | 4200 | 413 | 3000 |
| PBANKA_0830450 | 5 | 7 | 60 | 16 | 183 | 98 | 244 | 109 | 85 | 47 | 12 | 0 | 212 | 36 | 282 | 1425 | 1017 | 155 | 1773 | 813 | 189 | 1276 |
| PBANKA_0830500 | 0 | 0 | 15 | 0 | 31 | 7 | 73 | 16 | 13 | 0 | 0 | 0 | 67 | 0 | 70 | 306 | 197 | 22 | 226 | 90 | 5 | 46 |
| PBANKA_0830600 | 6 | 0 | 26 | 12 | 88 | 40 | 121 | 79 | 61 | 50 | 14 | 0 | 204 | 22 | 136 | 343 | 603 | 142 | 441 | 164 | 37 | 304 |
| PBANKA_0830700 | 0 | 0 | 9 | 5 | 24 | 5 | 26 | 6 | 22 | 5 | 0 | 0 | 274 | 122 | 59 | 235 | 111 | 61 | 5626 | 3483 | 70 | 514 |
| PBANKA_0830800 | 0 | 0 | 137 | 29 | 291 | 111 | 363 | 117 | 165 | 51 | 30 | 0 | 1640 | 963 | 1247 | 5095 | 1621 | 222 | 1158 | 765 | 1148 | 6825 |
| PBANKA_0831000 | 57 | 59 | 40 | 25 | 3608 | 1507 | 27708 | 11809 | 47151 | 27098 | 7887 | 3266 | 20528 | 13495 | 153 | 250 | 3E+05 | 60092 | 3595 | 5565 | 213 | 2162 |
| PBANKA_0831100 | 130 | 201 | 118 | 29 | 203 | 67 | 251 | 80 | 126 | 48 | 86 | 27 | 1710 | 1061 | 467 | 3109 | 1897 | 438 | 2970 | 2657 | 560 | 3291 |
| PBANKA_0831200 | 68 | 131 | 28 | 9 | 155 | 58 | 467 | 146 | 637 | 324 | 111 | 64 | 1249 | 804 | 207 | 698 | 14489 | 2621 | 2483 | 4493 | 349 | 2338 |
| PBANKA_0831300 | 77 | 139 | 88 | 14 | 180 | 81 | 233 | 107 | 76 | 23 | 222 | 130 | 1001 | 790 | 352 | 1873 | 1427 | 204 | 545 | 582 | 289 | 1895 |
| PBANKA_0831400 | 8 | 13 | 32 | 0 | 104 | 39 | 116 | 27 | 39 | 13 | 30 | 26 | 314 | 127 | 226 | 787 | 299 | 54 | 571 | 359 | 55 | 368 |
| PBANKA_0831500 | 15 | 27 | 125 | 24 | 562 | 194 | 636 | 198 | 322 | 115 | 70 | 21 | 631 | 367 | 2157 | 10102 | 2763 | 386 | 1318 | 3589 | 154 | 956 |
| PBANKA_0831700 | 73 | 63 | 6 | 0 | 56 | 23 | 71 | 14 | 33 | 9 | 5 | 0 | 51 | 16 | 77 | 238 | 116 | 34 | 464 | 159 | 21 | 106 |
| PBANKA_0831800 | 59 | 67 | 42 | 10 | 155 | 63 | 195 | 61 | 72 | 44 | 174 | 115 | 653 | 317 | 317 | 1346 | 786 | 180 | 1688 | 762 | 170 | 1207 |
| PBANKA_0831900 | 132 | 147 | 177 | 51 | 258 | 86 | 194 | 72 | 121 | 69 | 787 | 393 | 2367 | 1758 | 458 | 2317 | 826 | 177 | 1018 | 763 | 1113 | 5868 |
| PBANKA_0832000 | 0 | 0 | 27 | 5 | 54 | 23 | 44 | 18 | 11 | 5 | 10 | 11 | 428 | 197 | 154 | 597 | 273 | 40 | 1906 | 830 | 108 | 670 |
| PBANKA_0832100 | 425 | 434 | 83 | 11 | 763 | 369 | 850 | 267 | 341 | 138 | 58 | 26 | 403 | 234 | 12 | 50 | 1095 | 159 | 225 | 199 | 132 | 1228 |
| PBANKA_0832200 | 0 | 0 | 102 | 17 | 285 | 145 | 287 | 125 | 87 | 40 | 29 | 7 | 2517 | 1247 | 1005 | 6813 | 3619 | 393 | 213 | 299 | 1105 | 6423 |
| PBANKA_0832300 | 0 | 0 | 10 | 0 | 25 | 10 | 24 | 15 | 52 | 42 | 6 | 12 | 236 | 55 | 75 | 245 | 242 | 92 | 665 | 219 | 75 | 369 |
| PBANKA_0832400 | 0 | 0 | 9 | 0 | 17 | 13 | 21 | 11 | 19 | 9 | 13 | 0 | 198 | 45 | 24 | 95 | 254 | 60 | 2049 | 837 | 674 | 3266 |
| PBANKA_0832500 | 0 | 0 | 133 | 24 | 208 | 123 | 250 | 70 | 75 | 26 | 10 | 5 | 309 | 150 | 198 | 557 | 970 | 140 | 2411 | 2970 | 512 | 3718 |
| PBANKA_0832600 | 0 | 0 | 157 | 38 | 247 | 125 | 297 | 144 | 93 | 56 | 7 | 6 | 891 | 137 | 776 | 2127 | 1169 | 201 | 420 | 179 | 53 | 414 |
| PBANKA_0832900 | 47 | 60 | 82 | 16 | 354 | 157 | 734 | 250 | 864 | 404 | 172 | 67 | 1397 | 714 | 569 | 1801 | 10388 | 2322 | 638 | 843 | 217 | 1261 |
| PBANKA_0833000 | 352 | 385 | 246 | 101 | 926 | 527 | 2001 | 762 | 694 | 456 | 140 | 82 | 2251 | 459 | 812 | 1693 | 10945 | 1930 | 2116 | 1171 | 578 | 3634 |
| PBANKA_0833100 | 10 | 14 | 469 | 121 | 2351 | 1281 | 1925 | 669 | 415 | 187 | 44 | 21 | 2060 | 446 | 1162 | 3877 | 755 | 189 | 1333 | 627 | 353 | 2856 |
| PBANKA_0833200 | 10 | 10 | 114 | 30 | 398 | 154 | 389 | 161 | 152 | 72 | 78 | 64 | 1240 | 358 | 412 | 2119 | 3206 | 417 | 1864 | 1351 | 629 | 3566 |
| PBANKA_0833300 | 12 | 19 | 5 | 0 | 167 | 58 | 392 | 153 | 273 | 108 | 43 | 21 | 559 | 146 | 22 | 103 | 2120 | 354 | 1638 | 1159 | 377 | 2324 |
| PBANKA_0833400 | 61 | 106 | 96 | 18 | 196 | 65 | 233 | 99 | 93 | 40 | 21 | 7 | 483 | 119 | 366 | 1039 | 873 | 275 | 383 | 201 | 114 | 612 |
| PBANKA_0833500 | 135 | 221 | 87 | 13 | 194 | 99 | 186 | 62 | 125 | 56 | 151 | 72 | 669 | 570 | 312 | 1607 | 686 | 138 | 494 | 347 | 116 | 575 |
| PBANKA_0833600 | 25 | 22 | 99 | 11 | 241 | 77 | 374 | 134 | 248 | 75 | 246 | 191 | 2162 | 963 | 719 | 3450 | 5536 | 776 | 1692 | 1082 | 67 | 490 |
| PBANKA_0833800 | 21 | 19 | 72 | 13 | 273 | 96 | 409 | 132 | 243 | 134 | 91 | 35 | 473 | 203 | 245 | 1180 | 1151 | 197 | 1032 | 691 | 1075 | 7249 |
| PBANKA_0833900 | 0 | 0 | 11 | 0 | 105 | 51 | 116 | 66 | 50 | 17 | 17 | 6 | 328 | 135 | 213 | 884 | 383 | 77 | 1011 | 1475 | 207 | 1363 |
| PBANKA_0834000 | 0 | 0 | 52 | 7 | 133 | 63 | 116 | 59 | 61 | 18 | 9 | 0 | 285 | 74 | 170 | 574 | 486 | 142 | 545 | 312 | 16 | 135 |
| PBANKA_0834100 | 13 | 15 | 40 | 0 | 185 | 62 | 216 | 71 | 143 | 52 | 36 | 9 | 201 | 105 | 204 | 609 | 880 | 131 | 1490 | 2250 | 289 | 2058 |
| PBANKA_0834200 | 11 | 14 | 46 | 15 | 117 | 43 | 90 | 40 | 65 | 30 | 38 | 16 | 77 | 25 | 156 | 694 | 221 | 38 | 579 | 421 | 273 | 1489 |
| PBANKA_0834300 | 21 | 35 | 84 | 25 | 311 | 120 | 478 | 178 | 224 | 86 | 36 | 21 | 761 | 275 | 693 | 3376 | 3466 | 393 | 1917 | 2169 | 434 | 2933 |
| PBANKA_0834400 | 76 | 146 | 245 | 62 | 564 | 346 | 753 | 326 | 247 | 152 | 44 | 17 | 2157 | 693 | 1215 | 4713 | 3879 | 657 | 3244 | 2446 | 912 | 8069 |
| PBANKA_0834500 | 12 | 16 | 21 | 11 | 64 | 30 | 95 | 51 | 28 | 12 | 0 | 0 | 86 | 12 | 37 | 112 | 133 | 39 | 590 | 267 | 125 | 747 |
| PBANKA_0834600 | 17 | 29 | 41 | 14 | 154 | 57 | 238 | 65 | 73 | 36 | 16 | 10 | 952 | 332 | 249 | 995 | 1090 | 242 | 2129 | 1914 | 668 | 4528 |
| PBANKA_0834700 | 15 | 30 | 88 | 19 | 209 | 98 | 240 | 97 | 117 | 52 | 43 | 18 | 1230 | 696 | 363 | 1411 | 1484 | 266 | 1066 | 651 | 240 | 1710 |
| PBANKA_0834800 | 157 | 207 | 67 | 17 | 111 | 44 | 146 | 50 | 64 | 23 | 14 | 6 | 502 | 201 | 181 | 611 | 372 | 67 | 1109 | 369 | 659 | 3804 |
| PBANKA_0834900 | 92 | 137 | 142 | 26 | 396 | 138 | 398 | 139 | 207 | 119 | 41 | 31 | 1413 | 694 | 842 | 3875 | 1639 | 275 | 1681 | 1482 | 1395 | 8358 |
| PBANKA_0835000 | 0 | 0 | 24 | 5 | 107 | 43 | 148 | 52 | 50 | 21 | 5 | 0 | 48 | 17 | 89 | 231 | 162 | 46 | 349 | 156 | 29 | 218 |
| PBANKA_0835100 | 5 | 0 | 64 | 7 | 109 | 72 | 142 | 76 | 44 | 31 | 0 | 0 | 128 | 8 | 162 | 504 | 579 | 117 | 311 | 138 | 95 | 637 |
| PBANKA_0835200 | 0 | 6 | 244 | 78 | 265 | 174 | 319 | 190 | 95 | 71 | 16 | 11 | 399 | 62 | 635 | 2472 | 2023 | 299 | 735 | 499 | 160 | 1389 |
| PBANKA_0835300 | 39 | 45 | 97 | 23 | 240 | 110 | 375 | 162 | 176 | 67 | 68 | 67 | 1263 | 546 | 575 | 2039 | 944 | 226 | 1771 | 1020 | 340 | 2686 |
| PBANKA_0835400 | 87 | 77 | 16 | 0 | 43 | 24 | 50 | 12 | 133 | 101 | 48 | 39 | 235 | 128 | 16 | 98 | 179 | 225 | 2846 | 967 | 69 | 416 |
| PBANKA_0835500 | 66 | 123 | 262 | 62 | 285 | 75 | 359 | 110 | 274 | 135 | 53 | 11 | 1861 | 504 | 386 | 1928 | 4127 | 902 | 7844 | 7021 | 223 | 1652 |
| PBANKA_0835700 | 85 | 121 | 7 | 0 | 29 | 17 | 83 | 24 | 62 | 28 | 20 | 15 | 443 | 325 | 111 | 434 | 641 | 144 | 312 | 174 | 525 | 2527 |
| PBANKA_0835800 | 10 | 13 | 63 | 25 | 146 | 68 | 139 | 75 | 52 | 27 | 34 | 28 | 546 | 242 | 307 | 843 | 608 | 146 | 306 | 121 | 296 | 1873 |
| PBANKA_0835900 | 9 | 8 | 34 | 10 | 54 | 34 | 75 | 41 | 31 | 13 | 25 | 19 | 166 | 52 | 109 | 330 | 212 | 50 | 82 | 33 | 126 | 639 |
| PBANKA_0836000 | 12 | 7 | 79 | 22 | 276 | 113 | 271 | 105 | 188 | 82 | 98 | 36 | 964 | 628 | 776 | 3125 | 1966 | 352 | 781 | 673 | 153 | 1224 |
| PBANKA_0836100 | 0 | 0 | 12 | 0 | 63 | 31 | 47 | 30 | 25 | 20 | 15 | 0 | 161 | 45 | 92 | 262 | 282 | 76 | 180 | 78 | 0 | 47 |
| PBANKA_0836300 | 6 | 16 | 33 | 9 | 405 | 195 | 1144 | 500 | 607 | 297 | 39 | 19 | 2706 | 706 | 2361 | 7610 | 11984 | 2237 | 3567 | 2129 | 99 | 777 |
| PBANKA_0836700 | 0 | 0 | 19 | 5 | 120 | 58 | 102 | 51 | 51 | 27 | 8 | 0 | 816 | 249 | 450 | 1202 | 1700 | 328 | 413 | 161 | 0 | 13 |
| PBANKA_0900900 | 6 | 16 | 33 | 12 | 165 | 76 | 179 | 54 | 487 | 301 | 151 | 73 | 160 | 427 | 135 | 541 | 1672 | 450 | 135 | 400 | 20 | 129 |
| PBANKA_0901000 | 0 | 12 | 18 | 0 | 82 | 33 | 56 | 10 | 59 | 50 | 47 | 28 | 1276 | 379 | 357 | 1193 | 409 | 154 | 1751 | 694 | 44 | 463 |
| PBANKA_0901100 | 246 | 385 | 156 | 45 | 570 | 242 | 800 | 312 | 389 | 151 | 72 | 26 | 639 | 295 | 589 | 2293 | 5552 | 642 | 594 | 833 | 145 | 900 |
| PBANKA_0901200 | 6 | 14 | 35 | 7 | 105 | 67 | 136 | 72 | 76 | 66 | 7 | 7 | 161 | 30 | 158 | 630 | 784 | 141 | 828 | 321 | 17 | 134 |
| PBANKA_0901300 | 1070 | 1493 | 118 | 29 | 461 | 225 | 364 | 181 | 181 | 97 | 42 | 11 | 594 | 330 | 483 | 3365 | 2134 | 184 | 1474 | 5195 | 144 | 801 |
| PBANKA_0901400 | 25 | 35 | 229 | 69 | 325 | 172 | 304 | 162 | 110 | 77 | 34 | 14 | 268 | 68 | 412 | 1219 | 1017 | 121 | 866 | 443 | 155 | 1136 |
| PBANKA_0901500 | 7 | 8 | 13 | 0 | 113 | 32 | 175 | 49 | 56 | 22 | 9 | 0 | 482 | 72 | 222 | 1159 | 960 | 125 | 520 | 326 | 12 | 69 |
| PBANKA_0901600 | 11 | 5 | 53 | 9 | 109 | 88 | 181 | 61 | 49 | 30 | 6 | 10 | 422 | 255 | 334 | 1191 | 422 | 105 | 383 | 212 | 17 | 155 |
| PBANKA_0901700 | 19 | 25 | 31 | 14 | 51 | 48 | 89 | 44 | 32 | 31 | 5 | 0 | 30 | 16 | 71 | 104 | 119 | 47 | 415 | 82 | 24 | 223 |
| PBANKA_0901800 | 19 | 49 | 348 | 55 | 1806 | 652 | 1863 | 630 | 712 | 328 | 139 | 44 | 455 | 313 | 1115 | 6331 | 7838 | 1246 | 304 | 1091 | 194 | 1231 |
| PBANKA_0901900 | 21 | 15 | 70 | 14 | 147 | 93 | 304 | 109 | 1022 | 448 | 157 | 99 | 1381 | 3253 | 473 | 2757 | 6454 | 1250 | 13661 | 1E+05 | 545 | 4268 |
| PBANKA_0902100 | 2743 | 3844 | 2237 | 561 | 3781 | 1723 | 2857 | 830 | 1026 | 411 | 278 | 144 | 323 | 445 | 121 | 599 | 304 | 37 | 407 | 432 | 68 | 488 |
| PBANKA_0902300 | 7 | 0 | 8 | 0 | 46 | 31 | 39 | 29 | 62 | 52 | 10 | 0 | 139 | 295 | 196 | 609 | 627 | 243 | 2110 | 5244 | 388 | 3080 |
| PBANKA_0902400 | 0 | 0 | 16 | 0 | 18 | 0 | 24 | 9 | 15 | 11 | 0 | 0 | 197 | 92 | 25 | 108 | 258 | 76 | 1586 | 1357 | 25 | 96 |
| PBANKA_0902500 | 184 | 317 | 124 | 48 | 233 | 99 | 254 | 81 | 200 | 68 | 24 | 12 | 985 | 798 | 447 | 3428 | 6557 | 698 | 2169 | 5710 | 818 | 4831 |
| PBANKA_0902700 | 31 | 53 | 76 | 17 | 71 | 30 | 91 | 39 | 27 | 8 | 11 | 0 | 63 | 38 | 0 | 11 | 174 | 35 | 606 | 621 | 36 | 273 |
| PBANKA_0902900 | 12 | 11 | 17 | 6 | 114 | 46 | 220 | 168 | 62 | 52 | 33 | 16 | 627 | 323 | 134 | 211 | 484 | 346 | 18508 | 5881 | 166 | 963 |
| PBANKA_0903000 | 35 | 43 | 201 | 61 | 290 | 120 | 245 | 80 | 122 | 69 | 1560 | 1034 | 2048 | 1683 | 511 | 2656 | 547 | 133 | 1991 | 1790 | 50 | 373 |
| PBANKA_0903100 | 93 | 152 | 6 | 0 | 28 | 14 | 35 | 8 | 16 | 7 | 5 | 0 | 181 | 50 | 16 | 71 | 102 | 32 | 1421 | 621 | 75 | 424 |

| | | | | | | | | | | | | | | | | | | | | | | |
|---|---|---|---|---|---|---|---|---|---|---|---|---|---|---|---|---|---|---|---|---|---|---|
| PBANKA_0903200 | 103 | 142 | 115 | 17 | 251 | 78 | 239 | 63 | 95 | 31 | 70 | 38 | 517 | 423 | 240 | 1098 | 265 | 54 | 1112 | 722 | 173 | 1097 |
| PBANKA_0903300 | 8 | 6 | 25 | 13 | 101 | 48 | 148 | 28 | 50 | 18 | 12 | 0 | 184 | 53 | 32 | 187 | 416 | 48 | 790 | 786 | 65 | 429 |
| PBANKA_0903400 | 8 | 17 | 51 | 6 | 192 | 73 | 255 | 104 | 294 | 123 | 46 | 20 | 2615 | 982 | 421 | 2138 | 2932 | 678 | 1462 | 1238 | 243 | 1734 |
| PBANKA_0903500 | 10 | 13 | 77 | 14 | 235 | 60 | 352 | 110 | 192 | 67 | 27 | 16 | 398 | 96 | 144 | 814 | 1717 | 243 | 714 | 409 | 49 | 289 |
| PBANKA_0903600 | 11 | 6 | 128 | 35 | 823 | 542 | 1724 | 611 | 532 | 334 | 78 | 37 | 2222 | 409 | 1756 | 7225 | 3869 | 590 | 2090 | 1394 | 97 | 778 |
| PBANKA_0903700 | 0 | 7 | 28 | 7 | 49 | 56 | 58 | 40 | 34 | 31 | 9 | 5 | 167 | 38 | 189 | 301 | 256 | 123 | 510 | 149 | 14 | 86 |
| PBANKA_0903800 | 0 | 6 | 30 | 0 | 107 | 45 | 473 | 201 | 1191 | 667 | 147 | 105 | 2635 | 2626 | 134 | 474 | 13883 | 3513 | 48174 | 76716 | 3761 | 25297 |
| PBANKA_0903900 | 14 | 16 | 79 | 14 | 196 | 104 | 409 | 142 | 240 | 96 | 27 | 12 | 1219 | 510 | 691 | 2323 | 3839 | 773 | 637 | 490 | 132 | 893 |
| PBANKA_0904100 | 119 | 110 | 133 | 29 | 211 | 139 | 415 | 199 | 214 | 174 | 81 | 39 | 1599 | 458 | 299 | 1028 | 3847 | 847 | 1035 | 489 | 304 | 2098 |
| PBANKA_0904200 | 8 | 17 | 55 | 6 | 160 | 61 | 137 | 47 | 49 | 19 | 20 | 17 | 898 | 629 | 116 | 556 | 511 | 207 | 13393 | 12050 | 135 | 905 |
| PBANKA_0904300 | 5 | 6 | 89 | 18 | 119 | 52 | 104 | 32 | 24 | 17 | 6 | 5 | 182 | 20 | 224 | 476 | 150 | 79 | 1783 | 562 | 19 | 141 |
| PBANKA_0904400 | 39 | 35 | 37 | 12 | 135 | 58 | 209 | 73 | 55 | 40 | 57 | 34 | 539 | 250 | 170 | 911 | 582 | 71 | 486 | 416 | 518 | 3399 |
| PBANKA_0904500 | 43 | 55 | 51 | 12 | 189 | 92 | 245 | 105 | 88 | 56 | 11 | 0 | 501 | 75 | 339 | 1451 | 977 | 133 | 734 | 464 | 1018 | 7204 |
| PBANKA_0904600 | 0 | 0 | 11 | 0 | 52 | 22 | 83 | 38 | 79 | 34 | 23 | 8 | 629 | 474 | 98 | 420 | 386 | 90 | 4904 | 5451 | 367 | 2346 |
| PBANKA_0904700 | 7 | 6 | 88 | 30 | 62 | 28 | 78 | 34 | 19 | 8 | 61 | 24 | 915 | 208 | 792 | 756 | 189 | 147 | 344 | 135 | 8 | 190 |
| PBANKA_0904800 | 0 | 0 | 21 | 0 | 15 | 6 | 20 | 8 | 14 | 7 | 0 | 0 | 1181 | 396 | 38 | 226 | 219 | 44 | 1614 | 731 | 13 | 138 |
| PBANKA_0904900 | 0 | 0 | 19 | 0 | 53 | 62 | 64 | 33 | 10 | 10 | 0 | 0 | 163 | 45 | 120 | 338 | 128 | 28 | 391 | 136 | 15 | 100 |
| PBANKA_0905000 | 32 | 21 | 399 | 94 | 620 | 375 | 484 | 217 | 113 | 125 | 545 | 349 | 1867 | 1040 | 1045 | 1953 | 538 | 178 | 483 | 262 | 64 | 407 |
| PBANKA_0905100 | 0 | 0 | 39 | 5 | 141 | 58 | 149 | 46 | 74 | 47 | 12 | 5 | 160 | 80 | 162 | 610 | 270 | 52 | 349 | 157 | 17 | 128 |
| PBANKA_0905200 | 10 | 19 | 87 | 17 | 657 | 302 | 812 | 282 | 1631 | 815 | 240 | 170 | 672 | 178 | 490 | 1981 | 1659 | 520 | 2345 | 1688 | 1164 | 7167 |
| PBANKA_0905600 | 36 | 51 | 230 | 83 | 262 | 159 | 293 | 164 | 155 | 139 | 503 | 463 | 3302 | 1195 | 1148 | 3724 | 884 | 186 | 3850 | 1797 | 372 | 2094 |
| PBANKA_0905700 | 7 | 8 | 64 | 29 | 110 | 66 | 113 | 25 | 34 | 26 | 136 | 94 | 826 | 391 | 287 | 922 | 224 | 51 | 664 | 385 | 90 | 687 |
| PBANKA_0905800 | 0 | 0 | 90 | 26 | 220 | 78 | 282 | 84 | 256 | 110 | 29 | 20 | 1735 | 528 | 415 | 1541 | 2506 | 487 | 1186 | 645 | 138 | 849 |
| PBANKA_0905900 | 209 | 269 | 123 | 20 | 373 | 149 | 331 | 81 | 553 | 282 | 162 | 56 | 1549 | 2238 | 132 | 461 | 2926 | 886 | 28208 | 75280 | 466 | 3329 |
| PBANKA_0906000 | 0 | 0 | 8 | 0 | 70 | 28 | 55 | 30 | 30 | 13 | 0 | 0 | 49 | 15 | 20 | 54 | 102 | 35 | 318 | 206 | 124 | 837 |
| PBANKA_0906200 | 6 | 5 | 134 | 26 | 281 | 105 | 364 | 134 | 116 | 60 | 10 | 0 | 1286 | 362 | 375 | 1895 | 1542 | 178 | 1402 | 799 | 368 | 2851 |
| PBANKA_0906300 | 0 | 0 | 63 | 11 | 133 | 91 | 121 | 54 | 58 | 19 | 17 | 0 | 790 | 366 | 213 | 846 | 585 | 108 | 359 | 265 | 536 | 3587 |
| PBANKA_0906400 | 19 | 20 | 1185 | 288 | 846 | 422 | 753 | 353 | 251 | 212 | 1702 | 1226 | 5116 | 1903 | 1287 | 2997 | 897 | 253 | 1284 | 518 | 81 | 633 |
| PBANKA_0906500 | 14 | 10 | 672 | 184 | 571 | 294 | 724 | 327 | 182 | 133 | 606 | 516 | 7710 | 3181 | 1240 | 4349 | 1960 | 459 | 1458 | 529 | 38 | 382 |
| PBANKA_0906600 | 13 | 19 | 74 | 23 | 132 | 66 | 126 | 55 | 101 | 37 | 126 | 103 | 2067 | 1145 | 339 | 1615 | 641 | 174 | 1588 | 1207 | 543 | 3364 |
| PBANKA_0906700 | 208 | 297 | 103 | 27 | 443 | 123 | 524 | 228 | 341 | 150 | 70 | 20 | 1256 | 659 | 727 | 3694 | 5099 | 772 | 3644 | 3744 | 643 | 3798 |
| PBANKA_0906900 | 220 | 419 | 319 | 77 | 1610 | 586 | 2140 | 714 | 891 | 348 | 1201 | 696 | 3479 | 6830 | 2797 | 15947 | 11774 | 1229 | 3047 | 8213 | 1459 | 9657 |
| PBANKA_0907000 | 6 | 5 | 36 | 8 | 40 | 20 | 53 | 5 | 16 | 8 | 0 | 0 | 326 | 67 | 142 | 731 | 141 | 62 | 2021 | 1917 | 1865 | 11984 |
| PBANKA_0907100 | 0 | 0 | 14 | 0 | 28 | 10 | 17 | 10 | 40 | 11 | 8 | 0 | 1139 | 678 | 34 | 108 | 477 | 298 | 35330 | 29011 | 1062 | 8633 |
| PBANKA_0907200 | 109 | 152 | 62 | 16 | 177 | 93 | 533 | 206 | 345 | 130 | 24 | 15 | 785 | 125 | 17 | 42 | 3157 | 760 | 1631 | 835 | 126 | 879 |
| PBANKA_0907300 | 0 | 0 | 60 | 7 | 141 | 74 | 314 | 91 | 74 | 36 | 8 | 9 | 954 | 244 | 421 | 2472 | 2794 | 319 | 939 | 908 | 54 | 331 |
| PBANKA_0907600 | 0 | 0 | 44 | 6 | 195 | 63 | 189 | 88 | 63 | 40 | 0 | 0 | 146 | 32 | 46 | 99 | 503 | 84 | 785 | 230 | 86 | 589 |
| PBANKA_0907700 | 0 | 0 | 18 | 6 | 199 | 66 | 551 | 215 | 277 | 106 | 40 | 15 | 965 | 469 | 92 | 308 | 7469 | 1022 | 1840 | 3230 | 298 | 1938 |
| PBANKA_0907800 | 22 | 35 | 54 | 18 | 187 | 104 | 236 | 107 | 94 | 77 | 17 | 7 | 135 | 24 | 105 | 289 | 455 | 97 | 583 | 242 | 46 | 438 |
| PBANKA_0907900 | 5 | 12 | 6 | 0 | 25 | 13 | 54 | 23 | 56 | 23 | 24 | 8 | 1770 | 4174 | 139 | 566 | 2431 | 659 | 21270 | 1E+05 | 506 | 4179 |
| PBANKA_0908000 | 30 | 55 | 58 | 12 | 215 | 69 | 279 | 88 | 128 | 56 | 26 | 9 | 228 | 282 | 978 | 6533 | 3092 | 335 | 507 | 1599 | 133 | 846 |
| PBANKA_0908100 | 49 | 76 | 99 | 25 | 446 | 172 | 525 | 209 | 220 | 93 | 33 | 16 | 299 | 194 | 1034 | 4629 | 1583 | 200 | 1450 | 2868 | 50 | 287 |
| PBANKA_0908200 | 91 | 162 | 85 | 20 | 460 | 188 | 1348 | 536 | 674 | 369 | 252 | 126 | 1385 | 779 | 336 | 1962 | 1146 | 157 | 1213 | 922 | 241 | 1504 |
| PBANKA_0908300 | 57 | 77 | 48 | 12 | 830 | 327 | 2409 | 872 | 1579 | 740 | 3910 | 2414 | 1518 | 3294 | 1123 | 6401 | 2508 | 342 | 1184 | 3109 | 413 | 3022 |
| PBANKA_0908500 | 0 | 0 | 13 | 0 | 36 | 11 | 35 | 8 | 49 | 10 | 17 | 0 | 254 | 219 | 33 | 88 | 456 | 146 | 1865 | 5279 | 116 | 979 |
| PBANKA_0908600 | 69 | 57 | 12 | 0 | 67 | 45 | 70 | 34 | 64 | 50 | 8 | 6 | 31 | 14 | 0 | 14 | 332 | 96 | 431 | 76 | 1649 | 13906 |
| PBANKA_0908700 | 0 | 0 | 39 | 18 | 96 | 28 | 86 | 33 | 48 | 33 | 0 | 5 | 256 | 80 | 323 | 1357 | 1439 | 239 | 2277 | 1202 | 340 | 2612 |
| PBANKA_0908800 | 5 | 0 | 99 | 34 | 272 | 121 | 474 | 224 | 222 | 131 | 57 | 19 | 1416 | 346 | 531 | 2629 | 5502 | 776 | 5325 | 3715 | 207 | 1475 |
| PBANKA_0908900 | 119 | 130 | 48 | 15 | 150 | 79 | 174 | 89 | 52 | 29 | 9 | 5 | 380 | 118 | 219 | 539 | 309 | 93 | 912 | 508 | 126 | 645 |
| PBANKA_0909000 | 0 | 0 | 9 | 0 | 27 | 24 | 34 | 16 | 23 | 11 | 0 | 0 | 98 | 25 | 25 | 145 | 105 | 28 | 388 | 129 | 28 | 412 |
| PBANKA_0909100 | 0 | 0 | 40 | 16 | 119 | 84 | 179 | 78 | 38 | 27 | 6 | 0 | 361 | 117 | 221 | 793 | 399 | 63 | 559 | 241 | 0 | 49 |
| PBANKA_0909200 | 100 | 138 | 46 | 13 | 350 | 101 | 329 | 126 | 158 | 69 | 44 | 6 | 506 | 265 | 647 | 2541 | 2133 | 297 | 1205 | 1531 | 206 | 1154 |
| PBANKA_0909300 | 11 | 29 | 31 | 0 | 209 | 86 | 366 | 149 | 128 | 63 | 21 | 6 | 165 | 59 | 172 | 491 | 732 | 164 | 510 | 454 | 50 | 291 |
| PBANKA_0909400 | 901 | 1312 | 7 | 0 | 24 | 13 | 48 | 9 | 31 | 24 | 13 | 5 | 173 | 34 | 11 | 29 | 202 | 52 | 2205 | 1248 | 3991 | 27260 |
| PBANKA_0909500 | 51 | 71 | 30 | 0 | 163 | 110 | 186 | 56 | 53 | 29 | 11 | 0 | 164 | 64 | 27 | 55 | 145 | 80 | 2739 | 1587 | 212 | 1340 |
| PBANKA_0909600 | 208 | 319 | 562 | 146 | 1120 | 398 | 1030 | 346 | 387 | 167 | 502 | 363 | 3247 | 7616 | 3088 | 15118 | 3900 | 552 | 2904 | 8500 | 323 | 2490 |
| PBANKA_0909700 | 0 | 0 | 19 | 0 | 61 | 29 | 48 | 16 | 18 | 18 | 0 | 0 | 377 | 177 | 236 | 512 | 144 | 30 | 280 | 95 | 16 | 113 |
| PBANKA_0909800 | 143 | 172 | 390 | 102 | 607 | 244 | 451 | 147 | 126 | 71 | 24 | 16 | 186 | 75 | 589 | 2932 | 460 | 44 | 320 | 719 | 6493 | 49260 |
| PBANKA_0909900 | 1119 | 1803 | 221 | 45 | 750 | 422 | 1181 | 379 | 481 | 216 | 123 | 49 | 1885 | 1537 | 895 | 4215 | 3396 | 474 | 1196 | 1409 | 130 | 948 |
| PBANKA_0910000 | 11 | 15 | 7 | 0 | 74 | 24 | 56 | 42 | 340 | 195 | 1213 | 563 | 1465 | 2832 | 992 | 5166 | 3904 | 1014 | 1955 | 6464 | 37 | 148 |
| PBANKA_0910100 | 0 | 0 | 60 | 21 | 141 | 74 | 175 | 69 | 57 | 32 | 9 | 15 | 1382 | 457 | 433 | 1871 | 1682 | 242 | 609 | 353 | 16 | 82 |
| PBANKA_0910200 | 110 | 154 | 398 | 105 | 736 | 392 | 742 | 362 | 246 | 153 | 44 | 16 | 2169 | 670 | 2258 | 13249 | 4958 | 480 | 2051 | 2462 | 571 | 4450 |
| PBANKA_0910300 | 0 | 0 | 13 | 5 | 395 | 203 | 1320 | 525 | 1083 | 533 | 146 | 50 | 1025 | 751 | 207 | 361 | 20026 | 2794 | 2075 | 6138 | 44 | 390 |
| PBANKA_0910400 | 0 | 0 | 14 | 0 | 22 | 12 | 35 | 9 | 18 | 7 | 0 | 0 | 54 | 11 | 34 | 89 | 54 | 14 | 45 | 60 | 0 | 0 |
| PBANKA_0910600 | 0 | 0 | 36 | 0 | 49 | 21 | 54 | 21 | 26 | 20 | 0 | 5 | 165 | 64 | 78 | 222 | 317 | 65 | 268 | 147 | 27 | 224 |
| PBANKA_0910700 | 68 | 67 | 7 | 0 | 84 | 42 | 371 | 132 | 976 | 420 | 198 | 75 | 391 | 723 | 66 | 228 | 17504 | 3086 | 621 | 2958 | 284 | 1882 |
| PBANKA_0910800 | 91 | 103 | 26 | 0 | 53 | 32 | 53 | 22 | 26 | 8 | 20 | 14 | 146 | 83 | 18 | 73 | 188 | 34 | 892 | 893 | 20 | 81 |
| PBANKA_0911000 | 0 | 13 | 5 | 0 | 103 | 47 | 155 | 71 | 85 | 40 | 21 | 7 | 144 | 63 | 0 | 15 | 138 | 36 | 1487 | 1016 | 12 | 58 |
| PBANKA_0911200 | 57 | 71 | 86 | 0 | 81 | 63 | 91 | 46 | 29 | 18 | 7 | 0 | 284 | 51 | 192 | 496 | 802 | 152 | 390 | 164 | 14 | 179 |
| PBANKA_0911300 | 23 | 29 | 290 | 66 | 768 | 400 | 665 | 201 | 147 | 64 | 22 | 8 | 762 | 269 | 447 | 1853 | 411 | 71 | 157 | 64 | 0 | 30 |
| PBANKA_0911400 | 0 | 0 | 176 | 40 | 267 | 103 | 224 | 63 | 88 | 59 | 183 | 125 | 2090 | 1155 | 775 | 3196 | 1193 | 188 | 890 | 818 | 74 | 616 |
| PBANKA_0911500 | 46 | 86 | 5 | 0 | 25 | 20 | 84 | 36 | 178 | 123 | 35 | 19 | 227 | 93 | 38 | 85 | 1559 | 473 | 1880 | 1690 | 99 | 837 |
| PBANKA_0911600 | 9 | 0 | 48 | 7 | 104 | 61 | 125 | 57 | 61 | 27 | 9 | 6 | 235 | 69 | 215 | 981 | 930 | 139 | 1113 | 647 | 47 | 340 |
| PBANKA_0911650 | 55 | 52 | 124 | 12 | 67 | 50 | 65 | 24 | 92 | 59 | 26 | 0 | 800 | 165 | 1535 | 593 | 1095 | 337 | 13911 | 1223 | 0 | 24 |
| PBANKA_0911700 | 416 | 549 | 19 | 8 | 66 | 47 | 87 | 40 | 833 | 559 | 415 | 187 | 1140 | 903 | 266 | 1110 | 3883 | 985 | 5790 | 7198 | 1157 | 7793 |
| PBANKA_0911800 | 38 | 83 | 44 | 7 | 131 | 63 | 208 | 77 | 113 | 37 | 19 | 14 | 1628 | 657 | 266 | 1359 | 1123 | 155 | 949 | 637 | 147 | 915 |
| PBANKA_0911900 | 0 | 0 | 80 | 12 | 69 | 48 | 72 | 31 | 102 | 77 | 42 | 21 | 618 | 166 | 480 | 1888 | 1231 | 395 | 243 | 198 | 38 | 155 |
| PBANKA_0912000 | 19 | 27 | 341 | 68 | 434 | 150 | 389 | 132 | 333 | 147 | 2805 | 1528 | 4272 | 6546 | 1124 | 8850 | 1735 | 203 | 1449 | 3312 | 543 | 3377 |
| PBANKA_0912100 | 13 | 13 | 300 | 107 | 505 | 299 | 879 | 383 | 408 | 292 | 41 | 48 | 2872 | 907 | 1166 | 3384 | 4668 | 1096 | 2764 | 1111 | 1795 | 11736 |
| PBANKA_0912200 | 0 | 0 | 142 | 33 | 281 | 88 | 202 | 53 | 86 | 46 | 77 | 42 | 1803 | 1191 | 896 | 3830 | 807 | 133 | 691 | 697 | 23 | 177 |
| PBANKA_0912300 | 14 | 14 | 704 | 209 | 798 | 327 | 757 | 305 | 269 | 162 | 623 | 435 | 4234 | 5584 | 3299 | 18392 | 5347 | 603 | 882 | 2684 | 737 | 5183 |
| PBANKA_0912400 | 9 | 14 | 56 | 7 | 106 | 48 | 106 | 42 | 297 | 163 | 47 | 26 | 1007 | 882 | 37 | 91 | 3176 | 806 | 1868 | 1224 | 45 | 419 |
| PBANKA_0912600 | 0 | 0 | 31 | 9 | 346 | 147 | 738 | 254 | 474 | 233 | 77 | 28 | 761 | 1270 | 644 | 2247 | 9817 | 1021 | 1501 | 6568 | 242 | 1821 |
| PBANKA_0912700 | 7 | 0 | 71 | 10 | 209 | 81 | 319 | 171 | 163 | 99 | 16 | 13 | 923 | 303 | 426 | 1330 | 1730 | 466 | 702 | 304 | 19 | 148 |
| PBANKA_0912800 | 151 | 230 | 7 | 0 | 19 | 13 | 25 | 21 | 13 | 6 | 36 | 9 | 128 | 53 | 46 | 284 | 88 | 25 | 175 | 98 | 50 | 326 |
| PBANKA_0912900 | 75 | 75 | 35 | 6 | 51 | 31 | 58 | 20 | 22 | 14 | 88 | 53 | 343 | 179 | 99 | 414 | 237 | 75 | 297 | 167 | 87 | 677 |
| PBANKA_0913000 | 85 | 92 | 108 | 29 | 314 | 132 | 356 | 112 | 174 | 87 | 119 | 71 | 444 | 255 | 441 | 1367 | 1116 | 183 | 1981 | 3601 | 89 | 591 |
| PBANKA_0913100 | 0 | 0 | 31 | 5 | 89 | 56 | 84 | 28 | 25 | 25 | 12 | 0 | 91 | 27 | 140 | 422 | 131 | 37 | 723 | 326 | 20 | 153 |
| PBANKA_0913200 | 14 | 23 | 48 | 8 | 148 | 73 | 329 | 97 | 113 | 61 | 10 | 13 | 994 | 272 | 373 | 1297 | 1305 | 284 | 1271 | 653 | 49 | 370 |
| PBANKA_0913300 | 6 | 16 | 76 | 18 | 270 | 153 | 421 | 131 | 158 | 75 | 33 | 24 | 312 | 105 | 283 | 1181 | 1158 | 152 | 325 | 314 | 38 | 292 |

| | | | | | | | | | | | | | | | | | | | | | | |
|---|---|---|---|---|---|---|---|---|---|---|---|---|---|---|---|---|---|---|---|---|---|---|
| PBANKA_0913400 | 0 | 0 | 51 | 14 | 203 | 155 | 326 | 133 | 1705 | 1199 | 8716 | 7778 | 5670 | 2871 | 2545 | 8600 | 3359 | 887 | 1062 | 715 | 108 | 1002 |
| PBANKA_0913500 | 0 | 0 | 13 | 9 | 76 | 70 | 81 | 36 | 49 | 9 | 0 | 0 | 65 | 6 | 114 | 300 | 200 | 44 | 9 | 11 | 0 | 7 |
| PBANKA_0913700 | 23 | 57 | 88 | 16 | 350 | 150 | 424 | 166 | 166 | 85 | 90 | 53 | 1132 | 507 | 536 | 2831 | 1981 | 224 | 568 | 444 | 44 | 466 |
| PBANKA_0913800 | 61 | 90 | 208 | 66 | 453 | 261 | 485 | 189 | 230 | 102 | 351 | 229 | 3588 | 2113 | 984 | 4257 | 1948 | 253 | 954 | 946 | 70 | 478 |
| PBANKA_0913900 | 40 | 65 | 84 | 14 | 352 | 123 | 486 | 170 | 353 | 144 | 70 | 18 | 1436 | 840 | 498 | 2754 | 10425 | 1745 | 1627 | 2824 | 81 | 610 |
| PBANKA_0914000 | 9 | 30 | 63 | 13 | 177 | 100 | 176 | 69 | 58 | 32 | 15 | 0 | 302 | 149 | 149 | 465 | 373 | 145 | 179 | 173 | 18 | 109 |
| PBANKA_0914100 | 12 | 24 | 40 | 9 | 196 | 78 | 589 | 244 | 485 | 220 | 86 | 26 | 544 | 604 | 366 | 1800 | 10315 | 1307 | 1052 | 4234 | 402 | 2816 |
| PBANKA_0914200 | 139 | 189 | 334 | 58 | 585 | 185 | 503 | 204 | 660 | 253 | 5325 | 2996 | 3580 | 5407 | 978 | 4614 | 1607 | 372 | 1098 | 1856 | 366 | 2079 |
| PBANKA_0914300 | 12 | 15 | 104 | 25 | 343 | 228 | 391 | 216 | 103 | 87 | 9 | 20 | 519 | 197 | 1084 | 3161 | 2573 | 439 | 629 | 377 | 44 | 450 |
| PBANKA_0914400 | 1215 | 1776 | 2162 | 734 | 1777 | 1092 | 1607 | 735 | 395 | 262 | 328 | 274 | 11009 | 6466 | 4132 | 19523 | 1985 | 278 | 2040 | 2226 | 934 | 8600 |
| PBANKA_0914500 | 31 | 30 | 92 | 13 | 307 | 124 | 830 | 291 | 403 | 191 | 94 | 64 | 783 | 514 | 456 | 2184 | 3822 | 532 | 741 | 928 | 329 | 2521 |
| PBANKA_0914600 | 5 | 6 | 161 | 22 | 246 | 96 | 450 | 173 | 247 | 100 | 40 | 0 | 1639 | 742 | 464 | 2580 | 3922 | 651 | 10060 | 9198 | 185 | 1726 |
| PBANKA_0914700 | 0 | 0 | 122 | 37 | 283 | 159 | 320 | 148 | 78 | 54 | 128 | 71 | 1642 | 1462 | 560 | 3264 | 1394 | 144 | 321 | 533 | 21 | 151 |
| PBANKA_0914800 | 15 | 12 | 52 | 10 | 50 | 28 | 33 | 21 | 37 | 5 | 16 | 0 | 844 | 457 | 205 | 1034 | 297 | 37 | 617 | 239 | 8 | 47 |
| PBANKA_0914900 | 0 | 0 | 22 | 0 | 58 | 26 | 51 | 22 | 36 | 9 | 6 | 0 | 160 | 30 | 169 | 747 | 150 | 28 | 768 | 504 | 291 | 1622 |
| PBANKA_0915200 | 23 | 59 | 277 | 47 | 863 | 406 | 1331 | 436 | 668 | 274 | 93 | 38 | 2565 | 1411 | 308 | 1016 | 29400 | 3245 | 132 | 386 | 31 | 215 |
| PBANKA_0915300 | 58 | 92 | 236 | 50 | 346 | 109 | 253 | 95 | 400 | 155 | 2126 | 1145 | 2500 | 3230 | 906 | 4888 | 1508 | 254 | 727 | 1497 | 458 | 2872 |
| PBANKA_0915500 | 29 | 29 | 23 | 5 | 93 | 29 | 116 | 37 | 66 | 17 | 11 | 0 | 262 | 118 | 114 | 637 | 312 | 41 | 366 | 191 | 22 | 126 |
| PBANKA_0915600 | 318 | 385 | 23 | 0 | 113 | 67 | 1865 | 913 | 3423 | 2000 | 266 | 162 | 1161 | 263 | 121 | 323 | 10259 | 3812 | 1503 | 661 | 207 | 1567 |
| PBANKA_0915700 | 0 | 0 | 68 | 15 | 232 | 155 | 337 | 132 | 176 | 84 | 34 | 10 | 399 | 190 | 186 | 758 | 644 | 105 | 259 | 134 | 42 | 452 |
| PBANKA_0915800 | 8 | 6 | 48 | 10 | 131 | 78 | 205 | 61 | 75 | 33 | 64 | 39 | 1054 | 425 | 265 | 1157 | 827 | 161 | 653 | 457 | 82 | 501 |
| PBANKA_0915900 | 41 | 46 | 14 | 0 | 65 | 16 | 66 | 19 | 62 | 19 | 8 | 0 | 164 | 126 | 47 | 324 | 271 | 86 | 757 | 486 | 245 | 1109 |
| PBANKA_0916000 | 32 | 52 | 65 | 16 | 248 | 93 | 250 | 116 | 77 | 51 | 134 | 120 | 1338 | 1230 | 564 | 2993 | 1268 | 165 | 379 | 363 | 3594 | 19993 |
| PBANKA_0916100 | 6 | 8 | 85 | 9 | 208 | 81 | 356 | 153 | 153 | 56 | 13 | 9 | 798 | 133 | 362 | 1224 | 2148 | 375 | 3700 | 1955 | 116 | 891 |
| PBANKA_0916200 | 45 | 50 | 661 | 211 | 889 | 439 | 885 | 385 | 251 | 183 | 48 | 51 | 3996 | 1226 | 3323 | 14440 | 3416 | 452 | 2105 | 1807 | 172 | 1679 |
| PBANKA_0916400 | 133 | 206 | 165 | 30 | 526 | 249 | 701 | 241 | 298 | 132 | 73 | 44 | 1026 | 615 | 652 | 3167 | 3245 | 435 | 1792 | 1655 | 3410 | 25306 |
| PBANKA_0916600 | 7 | 8 | 84 | 18 | 399 | 171 | 553 | 217 | 245 | 128 | 44 | 13 | 409 | 209 | 337 | 1300 | 3656 | 475 | 1389 | 1767 | 659 | 4546 |
| PBANKA_0916800 | 0 | 0 | 32 | 5 | 143 | 35 | 115 | 33 | 79 | 46 | 15 | 0 | 560 | 383 | 355 | 1710 | 427 | 67 | 602 | 431 | 132 | 709 |
| PBANKA_0916900 | 23 | 25 | 51 | 6 | 217 | 67 | 221 | 82 | 70 | 30 | 26 | 9 | 439 | 218 | 167 | 695 | 847 | 167 | 1732 | 1134 | 425 | 2632 |
| PBANKA_0917000 | 36 | 44 | 16 | 0 | 76 | 20 | 108 | 44 | 70 | 41 | 0 | 14 | 668 | 204 | 97 | 259 | 1966 | 421 | 2256 | 2214 | 704 | 4152 |
| PBANKA_0917100 | 8 | 28 | 24 | 0 | 49 | 18 | 114 | 35 | 151 | 63 | 39 | 0 | 330 | 99 | 26 | 121 | 2560 | 468 | 2866 | 2012 | 446 | 2849 |
| PBANKA_0917200 | 86 | 140 | 42 | 12 | 144 | 50 | 98 | 29 | 50 | 27 | 14 | 6 | 1007 | 183 | 260 | 885 | 291 | 79 | 3122 | 1774 | 291 | 1441 |
| PBANKA_0917300 | 49 | 69 | 97 | 19 | 145 | 57 | 125 | 45 | 88 | 50 | 85 | 32 | 730 | 678 | 222 | 1202 | 718 | 109 | 1014 | 753 | 856 | 5800 |
| PBANKA_0917400 | 9 | 10 | 20 | 5 | 75 | 37 | 98 | 39 | 46 | 34 | 8 | 0 | 175 | 54 | 30 | 101 | 250 | 99 | 2608 | 894 | 438 | 3078 |
| PBANKA_0917500 | 76 | 109 | 220 | 48 | 650 | 214 | 762 | 257 | 366 | 153 | 151 | 56 | 1858 | 1332 | 1820 | 9271 | 9955 | 1437 | 3816 | 13207 | 800 | 3325 |
| PBANKA_0917700 | 80 | 146 | 84 | 11 | 204 | 69 | 212 | 80 | 142 | 63 | 660 | 328 | 1744 | 1222 | 422 | 1564 | 532 | 176 | 476 | 503 | 235 | 1274 |
| PBANKA_0917800 | 11 | 29 | 41 | 8 | 167 | 63 | 206 | 81 | 144 | 52 | 45 | 19 | 1421 | 760 | 351 | 1611 | 1153 | 185 | 1012 | 673 | 1100 | 7511 |
| PBANKA_0917900 | 9 | 0 | 203 | 60 | 470 | 275 | 774 | 311 | 303 | 120 | 49 | 22 | 2932 | 833 | 1420 | 5829 | 4312 | 655 | 1788 | 1149 | 185 | 1798 |
| PBANKA_0918000 | 864 | 1201 | 3183 | 1036 | 2920 | 1760 | 2971 | 1364 | 967 | 548 | 4017 | 3481 | 19038 | 8841 | 4346 | 11177 | 3748 | 855 | 4305 | 2018 | 3110 | 29105 |
| PBANKA_0918100 | 9 | 20 | 379 | 63 | 241 | 79 | 240 | 75 | 68 | 26 | 339 | 180 | 4007 | 1773 | 725 | 2397 | 1056 | 259 | 690 | 310 | 141 | 1094 |
| PBANKA_0918200 | 0 | 0 | 27 | 10 | 61 | 31 | 45 | 14 | 23 | 12 | 7 | 0 | 128 | 42 | 248 | 899 | 210 | 44 | 281 | 127 | 19 | 102 |
| PBANKA_0918300 | 399 | 539 | 572 | 173 | 1145 | 668 | 1151 | 429 | 429 | 281 | 59 | 46 | 5979 | 2031 | 1097 | 6434 | 2736 | 513 | 2534 | 2011 | 2528 | 17741 |
| PBANKA_0918400 | 0 | 0 | 25 | 6 | 46 | 27 | 41 | 15 | 16 | 0 | 6 | 0 | 124 | 31 | 66 | 359 | 133 | 20 | 449 | 354 | 150 | 1050 |
| PBANKA_0918700 | 5 | 6 | 47 | 10 | 577 | 256 | 924 | 328 | 252 | 115 | 36 | 17 | 319 | 111 | 193 | 784 | 675 | 113 | 219 | 113 | 11 | 102 |
| PBANKA_0918800 | 137 | 196 | 244 | 85 | 583 | 289 | 741 | 268 | 301 | 129 | 840 | 466 | 3043 | 2184 | 563 | 2764 | 562 | 135 | 488 | 391 | 94 | 600 |
| PBANKA_0918900 | 186 | 236 | 23 | 0 | 136 | 45 | 361 | 110 | 304 | 127 | 68 | 9 | 376 | 438 | 112 | 357 | 5371 | 921 | 436 | 1788 | 85 | 496 |
| PBANKA_0919000 | 0 | 0 | 210 | 39 | 389 | 209 | 546 | 282 | 172 | 87 | 35 | 11 | 1681 | 319 | 1249 | 4476 | 2910 | 363 | 339 | 268 | 16 | 90 |
| PBANKA_0919100 | 29 | 37 | 226 | 57 | 590 | 303 | 1056 | 468 | 1469 | 994 | 6972 | 5099 | 10602 | 7358 | 2079 | 10224 | 14688 | 3052 | 8467 | 5606 | 184 | 1547 |
| PBANKA_0919200 | 0 | 0 | 33 | 9 | 148 | 79 | 223 | 117 | 117 | 111 | 684 | 572 | 1842 | 632 | 523 | 1872 | 1374 | 262 | 1040 | 457 | 15 | 128 |
| PBANKA_0919300 | 18 | 11 | 404 | 132 | 438 | 267 | 340 | 154 | 115 | 80 | 458 | 354 | 3248 | 1586 | 1115 | 5023 | 770 | 124 | 574 | 418 | 67 | 566 |
| PBANKA_0919600 | 29 | 39 | 112 | 18 | 298 | 117 | 177 | 77 | 86 | 46 | 23 | 5 | 620 | 455 | 2281 | 15391 | 3993 | 497 | 1606 | 4377 | 118 | 774 |
| PBANKA_0919700 | 0 | 5 | 79 | 21 | 193 | 58 | 160 | 72 | 84 | 37 | 43 | 19 | 938 | 560 | 466 | 2724 | 1032 | 149 | 750 | 934 | 38 | 323 |
| PBANKA_0919800 | 0 | 0 | 128 | 35 | 229 | 144 | 247 | 145 | 84 | 42 | 12 | 7 | 314 | 132 | 471 | 2103 | 874 | 140 | 1005 | 451 | 36 | 298 |
| PBANKA_0919900 | 0 | 5 | 6 | 0 | 47 | 45 | 57 | 21 | 59 | 16 | 11 | 6 | 266 | 151 | 97 | 245 | 630 | 222 | 3940 | 4252 | 1917 | 12990 |
| PBANKA_0920000 | 31 | 33 | 75 | 35 | 168 | 115 | 170 | 72 | 57 | 26 | 15 | 8 | 691 | 240 | 490 | 1721 | 456 | 76 | 821 | 490 | 120 | 963 |
| PBANKA_0920100 | 19 | 36 | 95 | 23 | 182 | 73 | 154 | 73 | 38 | 11 | 12 | 0 | 361 | 99 | 385 | 1325 | 331 | 69 | 688 | 336 | 103 | 710 |
| PBANKA_0920300 | 26 | 30 | 47 | 6 | 202 | 97 | 230 | 72 | 81 | 43 | 16 | 8 | 485 | 146 | 452 | 1453 | 854 | 171 | 1373 | 938 | 229 | 1196 |
| PBANKA_0920400 | 210 | 287 | 238 | 50 | 513 | 129 | 922 | 300 | 778 | 352 | 1263 | 599 | 2804 | 3682 | 1041 | 5626 | 8337 | 1303 | 2017 | 4572 | 1175 | 6361 |
| PBANKA_0920500 | 222 | 235 | 44 | 6 | 110 | 53 | 469 | 229 | 1024 | 598 | 271 | 134 | 1450 | 701 | 118 | 606 | 7759 | 2056 | 5284 | 4409 | 926 | 6762 |
| PBANKA_0920600 | 84 | 89 | 158 | 36 | 397 | 113 | 483 | 173 | 462 | 168 | 212 | 106 | 2762 | 2795 | 949 | 7053 | 4904 | 814 | 2929 | 4798 | 497 | 2104 |
| PBANKA_0920700 | 5 | 7 | 6 | 0 | 47 | 25 | 25 | 26 | 45 | 16 | 33 | 7 | 228 | 109 | 58 | 164 | 283 | 109 | 1385 | 656 | 16 | 121 |
| PBANKA_0920800 | 0 | 0 | 288 | 66 | 751 | 485 | 1044 | 400 | 366 | 197 | 42 | 25 | 916 | 160 | 1185 | 4017 | 1815 | 303 | 2906 | 1331 | 93 | 795 |
| PBANKA_0920900 | 0 | 11 | 170 | 21 | 776 | 323 | 1605 | 561 | 551 | 290 | 63 | 48 | 1631 | 501 | 631 | 1399 | 2226 | 861 | 2067 | 754 | 270 | 1419 |
| PBANKA_0921000 | 0 | 0 | 13 | 0 | 39 | 12 | 51 | 13 | 17 | 7 | 0 | 0 | 193 | 91 | 41 | 232 | 170 | 37 | 232 | 198 | 26 | 63 |
| PBANKA_0921100 | 0 | 0 | 60 | 11 | 66 | 20 | 53 | 19 | 36 | 15 | 6 | 10 | 766 | 315 | 226 | 766 | 269 | 53 | 1547 | 826 | 29 | 204 |
| PBANKA_0921300 | 9 | 0 | 63 | 11 | 195 | 97 | 190 | 78 | 70 | 39 | 6 | 0 | 204 | 45 | 478 | 1294 | 660 | 114 | 1115 | 297 | 47 | 272 |
| PBANKA_0921400 | 0 | 0 | 14 | 0 | 52 | 48 | 74 | 32 | 533 | 373 | 703 | 511 | 2188 | 1676 | 628 | 1572 | 883 | 561 | 266 | 194 | 91 | 715 |
| PBANKA_0921500 | 0 | 0 | 82 | 30 | 176 | 85 | 198 | 51 | 40 | 25 | 147 | 207 | 418 | 308 | 372 | 555 | 337 | 73 | 157 | 101 | 12 | 41 |
| PBANKA_0921600 | 0 | 6 | 95 | 17 | 182 | 69 | 242 | 92 | 191 | 101 | 61 | 14 | 829 | 309 | 384 | 2092 | 2629 | 446 | 2360 | 3998 | 1069 | 7707 |
| PBANKA_0921700 | 21 | 34 | 179 | 39 | 680 | 311 | 743 | 283 | 1014 | 831 | 364 | 179 | 1842 | 1263 | 237 | 1483 | 8095 | 1707 | 5240 | 11638 | 3409 | 21168 |
| PBANKA_0921800 | 0 | 0 | 16 | 0 | 120 | 57 | 263 | 121 | 226 | 94 | 682 | 400 | 3433 | 1687 | 509 | 2413 | 5124 | 774 | 1094 | 837 | 126 | 838 |
| PBANKA_0921900 | 8 | 12 | 218 | 46 | 481 | 133 | 859 | 280 | 547 | 210 | 1004 | 541 | 3068 | 2542 | 691 | 3132 | 913 | 163 | 328 | 455 | 1515 | 9983 |
| PBANKA_0922000 | 36 | 63 | 52 | 6 | 152 | 29 | 221 | 87 | 84 | 40 | 39 | 21 | 1750 | 1055 | 349 | 1899 | 1803 | 356 | 839 | 570 | 84 | 571 |
| PBANKA_0922100 | 64 | 66 | 1726 | 542 | 1217 | 923 | 1301 | 685 | 423 | 401 | 2463 | 1920 | 10266 | 4468 | 2141 | 7663 | 3059 | 434 | 2800 | 1221 | 134 | 1402 |
| PBANKA_0922200 | 11 | 15 | 77 | 16 | 410 | 185 | 373 | 127 | 138 | 66 | 84 | 18 | 665 | 378 | 375 | 1833 | 537 | 100 | 942 | 783 | 1355 | 8985 |
| PBANKA_0922300 | 203 | 283 | 908 | 213 | 1676 | 768 | 2130 | 989 | 953 | 500 | 165 | 90 | 2912 | 1900 | 1310 | 5115 | 2644 | 337 | 1333 | 1768 | 280 | 1804 |
| PBANKA_0922400 | 0 | 0 | 27 | 0 | 35 | 11 | 62 | 22 | 34 | 18 | 6 | 0 | 198 | 79 | 168 | 451 | 295 | 61 | 263 | 142 | 187 | 1164 |
| PBANKA_0922500 | 0 | 5 | 113 | 26 | 430 | 254 | 1408 | 820 | 1020 | 699 | 109 | 52 | 2369 | 466 | 843 | 2631 | 15115 | 3094 | 1365 | 818 | 3131 | 27761 |
| PBANKA_0922700 | 0 | 0 | 32 | 9 | 54 | 30 | 47 | 28 | 30 | 25 | 5 | 0 | 68 | 9 | 88 | 92 | 144 | 124 | 275 | 56 | 33 | 191 |
| PBANKA_0922800 | 0 | 0 | 74 | 21 | 141 | 71 | 144 | 64 | 42 | 30 | 20 | 12 | 846 | 433 | 349 | 1492 | 576 | 62 | 722 | 310 | 280 | 1967 |
| PBANKA_0922900 | 118 | 198 | 156 | 41 | 232 | 108 | 185 | 88 | 81 | 30 | 166 | 118 | 2371 | 1133 | 492 | 2789 | 626 | 145 | 2540 | 1975 | 1150 | 8545 |
| PBANKA_0923000 | 220 | 361 | 328 | 55 | 573 | 250 | 392 | 102 | 183 | 64 | 44 | 25 | 2752 | 2389 | 1242 | 7233 | 1075 | 200 | 2196 | 2972 | 1781 | 10968 |
| PBANKA_0923100 | 0 | 0 | 30 | 17 | 176 | 98 | 175 | 119 | 52 | 33 | 8 | 6 | 233 | 61 | 103 | 541 | 304 | 61 | 362 | 195 | 27 | 200 |
| PBANKA_0923200 | 0 | 0 | 108 | 23 | 222 | 81 | 314 | 132 | 99 | 46 | 10 | 5 | 391 | 86 | 295 | 1016 | 1265 | 196 | 1236 | 433 | 201 | 1399 |
| PBANKA_0923300 | 0 | 0 | 30 | 10 | 150 | 61 | 256 | 122 | 123 | 68 | 28 | 14 | 641 | 190 | 186 | 752 | 792 | 183 | 7503 | 3084 | 293 | 2015 |
| PBANKA_0923400 | 9 | 6 | 872 | 194 | 471 | 264 | 416 | 213 | 184 | 121 | 1205 | 1100 | 9685 | 2448 | 1699 | 4433 | 1428 | 387 | 1607 | 515 | 111 | 543 |
| PBANKA_0923500 | 26 | 38 | 152 | 32 | 147 | 72 | 119 | 79 | 136 | 75 | 1108 | 758 | 3284 | 1613 | 417 | 1334 | 643 | 152 | 1075 | 683 | 189 | 1095 |
| PBANKA_0923600 | 0 | 0 | 243 | 66 | 139 | 94 | 97 | 38 | 32 | 11 | 24 | 11 | 1745 | 759 | 577 | 2449 | 217 | 45 | 2769 | 1389 | 616 | 4089 |
| PBANKA_0923700 | 0 | 0 | 79 | 10 | 172 | 46 | 825 | 323 | 1091 | 585 | 91 | 59 | 914 | 439 | 283 | 1051 | 3125 | 606 | 728 | 535 | 186 | 1426 |
| PBANKA_0923800 | 5 | 5 | 222 | 49 | 685 | 282 | 1009 | 363 | 434 | 183 | 56 | 18 | 99 | 28 | 37 | 166 | 472 | 95 | 13 | 16 | 23 | 103 |

| | | | | | | | | | | | | | | | | | | | | | | |
|---|---|---|---|---|---|---|---|---|---|---|---|---|---|---|---|---|---|---|---|---|---|---|
| PBANKA_0923900 | 13 | 20 | 32 | 5 | 174 | 95 | 313 | 143 | 142 | 92 | 78 | 44 | 324 | 206 | 94 | 350 | 269 | 50 | 150 | 63 | 18 | 94 |
| PBANKA_0924000 | 33 | 36 | 50 | 16 | 128 | 31 | 400 | 267 | 386 | 290 | 725 | 382 | 1674 | 1581 | 336 | 1635 | 475 | 128 | 1905 | 1437 | 427 | 2780 |
| PBANKA_0924100 | 0 | 10 | 20 | 7 | 102 | 39 | 216 | 137 | 182 | 73 | 72 | 19 | 1118 | 401 | 115 | 553 | 3425 | 538 | 5841 | 3654 | 366 | 2495 |
| PBANKA_0924200 | 5 | 8 | 39 | 5 | 172 | 81 | 194 | 72 | 77 | 31 | 41 | 11 | 1189 | 580 | 570 | 3093 | 812 | 131 | 266 | 333 | 13 | 123 |
| PBANKA_0924300 | 232 | 384 | 93 | 29 | 212 | 125 | 562 | 223 | 378 | 192 | 61 | 21 | 1113 | 247 | 355 | 1053 | 2915 | 709 | 3052 | 1329 | 198 | 1572 |
| PBANKA_0924400 | 55 | 55 | 198 | 78 | 251 | 177 | 234 | 120 | 129 | 120 | 1602 | 1828 | 1155 | 570 | 352 | 767 | 263 | 57 | 231 | 77 | 119 | 819 |
| PBANKA_0924500 | 52 | 66 | 59 | 13 | 427 | 141 | 679 | 247 | 428 | 140 | 64 | 14 | 1341 | 597 | 267 | 1183 | 10916 | 1360 | 356 | 1087 | 2887 | 17538 |
| PBANKA_0924600 | 0 | 6 | 30 | 7 | 81 | 57 | 72 | 35 | 14 | 19 | 0 | 0 | 42 | 7 | 176 | 594 | 100 | 22 | 212 | 128 | 42 | 461 |
| PBANKA_0924700 | 7 | 14 | 83 | 36 | 201 | 130 | 237 | 64 | 80 | 45 | 31 | 17 | 634 | 357 | 381 | 2402 | 921 | 184 | 1841 | 1556 | 1401 | 10539 |
| PBANKA_0924800 | 57 | 95 | 252 | 64 | 541 | 335 | 763 | 286 | 326 | 225 | 706 | 388 | 5419 | 4088 | 1655 | 9782 | 7177 | 678 | 1533 | 1777 | 3437 | 24764 |
| PBANKA_0924900 | 9 | 11 | 869 | 225 | 1375 | 704 | 1085 | 362 | 358 | 169 | 1630 | 1073 | 4282 | 2623 | 2374 | 10822 | 1844 | 274 | 1577 | 1476 | 42 | 559 |
| PBANKA_0925000 | 64 | 100 | 65 | 9 | 154 | 99 | 413 | 167 | 274 | 133 | 210 | 145 | 2001 | 773 | 502 | 1940 | 3553 | 738 | 5209 | 3305 | 215 | 1550 |
| PBANKA_0925100 | 37 | 30 | 8 | 0 | 76 | 35 | 156 | 73 | 79 | 52 | 8 | 0 | 587 | 137 | 98 | 426 | 1265 | 196 | 4259 | 2158 | 43 | 373 |
| PBANKA_0925200 | 52 | 53 | 456 | 138 | 1116 | 431 | 964 | 332 | 401 | 174 | 1383 | 901 | 4283 | 8268 | 3190 | 20424 | 5214 | 560 | 1942 | 8028 | 810 | 7038 |
| PBANKA_0925300 | 11 | 21 | 163 | 27 | 417 | 176 | 435 | 134 | 269 | 132 | 2507 | 1146 | 4745 | 7345 | 2152 | 11420 | 4018 | 572 | 2417 | 5916 | 351 | 2287 |
| PBANKA_0925400 | 29 | 51 | 7 | 0 | 73 | 37 | 97 | 56 | 140 | 48 | 32 | 14 | 629 | 981 | 760 | 3736 | 3577 | 585 | 1979 | 17214 | 65 | 523 |
| PBANKA_0925500 | 370 | 704 | 159 | 30 | 523 | 218 | 741 | 281 | 275 | 141 | 41 | 30 | 440 | 276 | 789 | 3957 | 3720 | 363 | 1579 | 3398 | 3290 | 23495 |
| PBANKA_0925600 | 24 | 21 | 18 | 7 | 146 | 106 | 530 | 236 | 290 | 154 | 18 | 16 | 596 | 120 | 43 | 102 | 3251 | 634 | 485 | 264 | 264 | 1944 |
| PBANKA_0925700 | 33 | 47 | 16 | 0 | 128 | 77 | 187 | 58 | 74 | 38 | 9 | 0 | 219 | 93 | 84 | 173 | 963 | 403 | 1689 | 1114 | 60 | 418 |
| PBANKA_0925800 | 37 | 58 | 10 | 0 | 62 | 39 | 160 | 65 | 108 | 65 | 32 | 0 | 324 | 248 | 150 | 493 | 2491 | 330 | 973 | 1170 | 62 | 424 |
| PBANKA_0925900 | 19 | 19 | 156 | 39 | 194 | 101 | 307 | 128 | 127 | 67 | 8 | 0 | 1845 | 892 | 522 | 2855 | 2040 | 288 | 880 | 820 | 598 | 4462 |
| PBANKA_0926000 | 32 | 32 | 11 | 0 | 101 | 57 | 90 | 40 | 27 | 22 | 20 | 5 | 168 | 61 | 115 | 499 | 146 | 24 | 210 | 109 | 111 | 631 |
| PBANKA_0926100 | 42 | 73 | 128 | 26 | 256 | 144 | 250 | 88 | 78 | 32 | 47 | 11 | 570 | 317 | 177 | 803 | 157 | 26 | 286 | 187 | 82 | 543 |
| PBANKA_0926200 | 9 | 8 | 87 | 19 | 125 | 84 | 177 | 40 | 37 | 19 | 6 | 0 | 194 | 45 | 281 | 934 | 542 | 117 | 391 | 259 | 14 | 92 |
| PBANKA_0926300 | 0 | 0 | 48 | 9 | 111 | 45 | 84 | 37 | 23 | 13 | 7 | 0 | 807 | 294 | 242 | 1066 | 475 | 83 | 1417 | 991 | 158 | 1195 |
| PBANKA_0926400 | 0 | 0 | 26 | 0 | 51 | 17 | 50 | 23 | 35 | 29 | 0 | 0 | 164 | 69 | 138 | 505 | 664 | 85 | 443 | 453 | 0 | 47 |
| PBANKA_0926500 | 0 | 0 | 6 | 0 | 34 | 13 | 25 | 18 | 27 | 19 | 0 | 0 | 356 | 464 | 31 | 126 | 374 | 143 | 8628 | 24116 | 3725 | 25592 |
| PBANKA_0926600 | 0 | 0 | 163 | 30 | 238 | 90 | 263 | 69 | 136 | 49 | 27 | 15 | 1273 | 619 | 626 | 3085 | 1303 | 215 | 1381 | 2377 | 266 | 1811 |
| PBANKA_0926700 | 23 | 21 | 1007 | 489 | 1827 | 1396 | 2042 | 1352 | 3378 | 3768 | 9923 | 9074 | 6233 | 3684 | 3719 | 11324 | 6229 | 3725 | 1919 | 1131 | 58 | 610 |
| PBANKA_0926800 | 77 | 93 | 92 | 23 | 186 | 63 | 120 | 33 | 44 | 26 | 33 | 9 | 698 | 240 | 185 | 911 | 409 | 115 | 1488 | 1441 | 70 | 394 |
| PBANKA_0926900 | 19 | 20 | 31 | 11 | 167 | 72 | 396 | 166 | 389 | 253 | 103 | 46 | 593 | 163 | 121 | 434 | 710 | 234 | 4259 | 2571 | 176 | 1251 |
| PBANKA_0927000 | 0 | 0 | 12 | 0 | 205 | 99 | 939 | 394 | 390 | 217 | 49 | 9 | 880 | 317 | 85 | 335 | 5100 | 532 | 2063 | 2205 | 275 | 1848 |
| PBANKA_0927100 | 0 | 0 | 11 | 0 | 49 | 19 | 52 | 23 | 29 | 6 | 6 | 0 | 97 | 23 | 126 | 405 | 325 | 131 | 380 | 167 | 68 | 429 |
| PBANKA_0927200 | 20 | 34 | 188 | 50 | 445 | 188 | 642 | 215 | 436 | 235 | 1735 | 1175 | 2243 | 4907 | 1159 | 6377 | 5302 | 711 | 1386 | 1639 | 140 | 778 |
| PBANKA_0927400 | 6 | 0 | 14 | 0 | 57 | 29 | 42 | 7 | 15 | 5 | 0 | 6 | 249 | 64 | 87 | 331 | 492 | 129 | 1495 | 1060 | 58 | 482 |
| PBANKA_0927500 | 5 | 0 | 17 | 5 | 33 | 15 | 40 | 17 | 11 | 7 | 5 | 0 | 215 | 78 | 68 | 213 | 293 | 95 | 1088 | 621 | 32 | 307 |
| PBANKA_0927700 | 5 | 8 | 9 | 0 | 12 | 0 | 9 | 9 | 13 | 6 | 0 | 12 | 580 | 851 | 65 | 321 | 829 | 97 | 2316 | 11011 | 41 | 254 |
| PBANKA_0927800 | 9 | 12 | 114 | 22 | 259 | 92 | 213 | 68 | 56 | 27 | 199 | 135 | 1379 | 828 | 769 | 3154 | 1054 | 168 | 457 | 500 | 15 | 60 |
| PBANKA_0927900 | 0 | 0 | 110 | 17 | 128 | 71 | 132 | 52 | 83 | 54 | 14 | 17 | 833 | 294 | 376 | 1409 | 1050 | 277 | 1020 | 497 | 77 | 438 |
| PBANKA_0928000 | 173 | 269 | 24 | 9 | 26 | 14 | 42 | 16 | 19 | 10 | 15 | 15 | 67 | 52 | 25 | 108 | 71 | 38 | 191 | 168 | 176 | 923 |
| PBANKA_0928100 | 0 | 5 | 1950 | 688 | 1182 | 950 | 947 | 439 | 255 | 151 | 84 | 56 | 7715 | 2221 | 4436 | 14937 | 2598 | 305 | 1445 | 813 | 81 | 784 |
| PBANKA_0928200 | 35 | 89 | 406 | 129 | 467 | 162 | 384 | 114 | 194 | 109 | 1110 | 803 | 8336 | 10458 | 2489 | 16306 | 3034 | 388 | 3047 | 5384 | 222 | 1491 |
| PBANKA_0928300 | 0 | 7 | 60 | 11 | 187 | 72 | 226 | 101 | 122 | 66 | 31 | 11 | 846 | 620 | 552 | 2701 | 2561 | 325 | 1458 | 1878 | 34 | 292 |
| PBANKA_0928400 | 14 | 9 | 33 | 7 | 103 | 33 | 132 | 53 | 90 | 46 | 15 | 0 | 264 | 79 | 111 | 427 | 1443 | 237 | 552 | 474 | 170 | 1372 |
| PBANKA_0928500 | 441 | 758 | 119 | 45 | 342 | 156 | 228 | 70 | 104 | 41 | 36 | 18 | 1394 | 766 | 730 | 3305 | 557 | 102 | 1474 | 1389 | 334 | 2929 |
| PBANKA_0928600 | 0 | 0 | 35 | 11 | 155 | 62 | 134 | 61 | 73 | 29 | 7 | 0 | 418 | 138 | 237 | 876 | 398 | 59 | 256 | 162 | 125 | 924 |
| PBANKA_0928700 | 143 | 202 | 107 | 31 | 487 | 188 | 630 | 257 | 291 | 134 | 65 | 53 | 1235 | 804 | 725 | 3916 | 4163 | 464 | 1897 | 3001 | 429 | 2962 |
| PBANKA_0928800 | 48 | 72 | 89 | 31 | 137 | 147 | 179 | 95 | 76 | 69 | 169 | 190 | 1407 | 716 | 526 | 2873 | 1524 | 127 | 914 | 762 | 294 | 2841 |
| PBANKA_0928900 | 69 | 63 | 20 | 0 | 48 | 20 | 53 | 28 | 30 | 11 | 15 | 10 | 1088 | 321 | 177 | 726 | 624 | 210 | 755 | 386 | 141 | 869 |
| PBANKA_0929000 | 62 | 81 | 69 | 23 | 137 | 49 | 125 | 69 | 104 | 71 | 84 | 37 | 964 | 720 | 129 | 661 | 517 | 119 | 228 | 256 | 318 | 2028 |
| PBANKA_0929100 | 141 | 227 | 91 | 18 | 146 | 69 | 201 | 97 | 105 | 62 | 25 | 12 | 435 | 124 | 107 | 328 | 1150 | 298 | 2439 | 1108 | 498 | 3427 |
| PBANKA_0929200 | 286 | 355 | 76 | 15 | 73 | 53 | 69 | 29 | 48 | 26 | 9 | 0 | 112 | 25 | 12 | 31 | 125 | 84 | 2098 | 860 | 119 | 846 |
| PBANKA_0929300 | 49 | 32 | 184 | 48 | 212 | 137 | 244 | 123 | 164 | 124 | 40 | 17 | 242 | 102 | 134 | 263 | 365 | 134 | 4690 | 2162 | 556 | 3503 |
| PBANKA_0929400 | 9 | 16 | 6 | 0 | 73 | 43 | 226 | 112 | 533 | 340 | 83 | 46 | 679 | 349 | 13 | 58 | 3968 | 1174 | 2938 | 2460 | 2622 | 12636 |
| PBANKA_0929500 | 84 | 132 | 80 | 14 | 663 | 321 | 1142 | 463 | 385 | 203 | 49 | 24 | 520 | 181 | 240 | 871 | 1164 | 200 | 569 | 272 | 283 | 2027 |
| PBANKA_0929600 | 104 | 138 | 430 | 120 | 765 | 364 | 645 | 294 | 371 | 187 | 2217 | 1678 | 4853 | 3777 | 1073 | 5384 | 1423 | 204 | 1446 | 1499 | 573 | 4577 |
| PBANKA_0929700 | 43 | 53 | 44 | 7 | 113 | 53 | 183 | 47 | 117 | 52 | 22 | 10 | 301 | 94 | 38 | 190 | 306 | 63 | 3799 | 2802 | 170 | 1402 |
| PBANKA_0929800 | 13 | 9 | 199 | 63 | 476 | 199 | 521 | 164 | 165 | 92 | 87 | 44 | 2551 | 2169 | 1176 | 5484 | 2493 | 332 | 816 | 1116 | 229 | 2011 |
| PBANKA_0929900 | 258 | 357 | 757 | 222 | 472 | 308 | 373 | 121 | 168 | 70 | 60 | 24 | 2151 | 1174 | 1044 | 4581 | 474 | 97 | 802 | 685 | 294 | 2301 |
| PBANKA_0930000 | 0 | 0 | 71 | 21 | 251 | 148 | 395 | 153 | 136 | 73 | 19 | 9 | 528 | 100 | 368 | 934 | 649 | 206 | 490 | 141 | 21 | 115 |
| PBANKA_0930100 | 87 | 113 | 44 | 14 | 183 | 52 | 450 | 245 | 594 | 281 | 85 | 43 | 806 | 478 | 284 | 1392 | 6185 | 1207 | 2265 | 5929 | 1803 | 11075 |
| PBANKA_0930200 | 79 | 137 | 84 | 24 | 336 | 128 | 523 | 170 | 279 | 102 | 49 | 24 | 399 | 151 | 318 | 1427 | 3157 | 506 | 1608 | 1734 | 1780 | 13025 |
| PBANKA_0930300 | 371 | 428 | 1058 | 381 | 983 | 720 | 1551 | 915 | 684 | 414 | 1448 | 1328 | 7105 | 2443 | 2265 | 7119 | 5831 | 992 | 1384 | 696 | 221 | 1688 |
| PBANKA_0930400 | 0 | 0 | 29 | 8 | 80 | 33 | 62 | 40 | 30 | 11 | 5 | 0 | 84 | 32 | 45 | 140 | 90 | 32 | 332 | 170 | 17 | 99 |
| PBANKA_0930500 | 0 | 0 | 128 | 34 | 346 | 142 | 417 | 123 | 201 | 88 | 24 | 9 | 805 | 212 | 419 | 1744 | 1434 | 321 | 974 | 588 | 42 | 309 |
| PBANKA_0930600 | 0 | 0 | 61 | 11 | 153 | 50 | 248 | 69 | 391 | 226 | 67 | 37 | 823 | 234 | 386 | 2072 | 2321 | 683 | 305 | 320 | 52 | 473 |
| PBANKA_0930700 | 0 | 8 | 53 | 7 | 201 | 88 | 274 | 112 | 81 | 57 | 101 | 82 | 1728 | 411 | 582 | 1699 | 870 | 170 | 1506 | 571 | 50 | 375 |
| PBANKA_0930800 | 32 | 35 | 5 | 0 | 53 | 13 | 45 | 19 | 55 | 29 | 10 | 0 | 138 | 49 | 44 | 147 | 210 | 71 | 1361 | 695 | 86 | 416 |
| PBANKA_0930900 | 0 | 0 | 78 | 22 | 216 | 134 | 314 | 165 | 118 | 64 | 10 | 7 | 834 | 182 | 726 | 2404 | 2888 | 374 | 1802 | 1078 | 195 | 1225 |
| PBANKA_0931000 | 23 | 21 | 31 | 0 | 165 | 71 | 194 | 88 | 104 | 49 | 13 | 0 | 296 | 61 | 294 | 1624 | 1052 | 135 | 1227 | 655 | 99 | 614 |
| PBANKA_0931300 | 651 | 930 | 129 | 34 | 329 | 199 | 295 | 123 | 149 | 76 | 1399 | 1025 | 5764 | 2356 | 3494 | 15660 | 2207 | 365 | 2952 | 2707 | 3254 | 28438 |
| PBANKA_0931400 | 0 | 0 | 92 | 15 | 295 | 132 | 176 | 78 | 68 | 24 | 19 | 9 | 743 | 206 | 762 | 3196 | 413 | 77 | 123 | 83 | 12 | 62 |
| PBANKA_0931500 | 17 | 19 | 55 | 13 | 149 | 69 | 132 | 60 | 141 | 105 | 22 | 34 | 1316 | 342 | 423 | 1484 | 1802 | 581 | 13335 | 6007 | 104 | 791 |
| PBANKA_0931600 | 16 | 14 | 77 | 12 | 174 | 55 | 278 | 83 | 209 | 106 | 29 | 17 | 639 | 278 | 223 | 1135 | 1714 | 346 | 1715 | 1285 | 71 | 568 |
| PBANKA_0931700 | 0 | 0 | 69 | 18 | 158 | 98 | 164 | 128 | 59 | 66 | 7 | 0 | 194 | 32 | 248 | 533 | 243 | 95 | 449 | 179 | 9 | 110 |
| PBANKA_0931800 | 9 | 34 | 179 | 43 | 104 | 92 | 177 | 68 | 64 | 34 | 0 | 0 | 388 | 75 | 353 | 892 | 249 | 79 | 320 | 144 | 17 | 139 |
| PBANKA_0931900 | 401 | 511 | 124 | 27 | 338 | 115 | 632 | 205 | 1034 | 415 | 200 | 81 | 1442 | 1218 | 606 | 3235 | 8951 | 1623 | 1733 | 2940 | 941 | 5796 |
| PBANKA_0932100 | 0 | 0 | 28 | 5 | 133 | 83 | 303 | 117 | 117 | 72 | 10 | 0 | 318 | 87 | 349 | 930 | 793 | 196 | 1553 | 824 | 26 | 287 |
| PBANKA_0932200 | 0 | 0 | 247 | 94 | 2869 | 2389 | 5799 | 3458 | 1281 | 990 | 120 | 71 | 1400 | 183 | 3044 | 7284 | 4956 | 979 | 3854 | 1663 | 117 | 1251 |
| PBANKA_0932300 | 41 | 89 | 89 | 11 | 103 | 93 | 135 | 71 | 42 | 25 | 0 | 0 | 370 | 90 | 101 | 336 | 108 | 34 | 636 | 208 | 11 | 64 |
| PBANKA_0932400 | 0 | 0 | 57 | 18 | 262 | 176 | 270 | 153 | 132 | 71 | 1175 | 1056 | 7967 | 2797 | 5358 | 20113 | 2676 | 334 | 685 | 445 | 16 | 169 |
| PBANKA_0932500 | 0 | 0 | 26 | 7 | 229 | 116 | 549 | 196 | 961 | 521 | 75 | 64 | 3438 | 1918 | 434 | 1629 | 12922 | 2477 | 55907 | 77474 | 229 | 2124 |
| PBANKA_0932700 | 0 | 0 | 110 | 21 | 77 | 40 | 100 | 55 | 32 | 12 | 7 | 0 | 54 | 6 | 137 | 365 | 122 | 30 | 78 | 45 | 5 | 41 |
| PBANKA_0932800 | 20 | 34 | 18 | 0 | 363 | 186 | 593 | 204 | 403 | 198 | 83 | 21 | 392 | 303 | 192 | 675 | 6906 | 916 | 832 | 1988 | 29 | 204 |
| PBANKA_0932900 | 6 | 13 | 23 | 18 | 369 | 253 | 439 | 240 | 160 | 103 | 11 | 10 | 92 | 16 | 59 | 92 | 302 | 51 | 302 | 140 | 5 | 25 |
| PBANKA_0933000 | 178 | 225 | 172 | 75 | 555 | 251 | 994 | 389 | 617 | 310 | 58 | 43 | 2969 | 1080 | 1525 | 7207 | 7956 | 1155 | 5251 | 3862 | 346 | 2915 |
| PBANKA_0933200 | 100 | 190 | 22 | 0 | 99 | 49 | 117 | 35 | 47 | 36 | 9 | 0 | 283 | 72 | 226 | 953 | 547 | 86 | 310 | 214 | 267 | 1565 |
| PBANKA_0933300 | 64 | 168 | 177 | 36 | 303 | 145 | 380 | 128 | 105 | 48 | 17 | 11 | 1586 | 537 | 486 | 2461 | 1536 | 189 | 1226 | 1016 | 181 | 1336 |
| PBANKA_0933400 | 0 | 8 | 55 | 8 | 162 | 91 | 270 | 98 | 85 | 41 | 24 | 13 | 1259 | 484 | 629 | 1756 | 879 | 221 | 480 | 329 | 11 | 105 |
| PBANKA_0933500 | 580 | 779 | 68 | 25 | 162 | 109 | 286 | 121 | 295 | 236 | 68 | 20 | 325 | 83 | 183 | 340 | 1166 | 349 | 1033 | 443 | 618 | 4316 |

| | | | | | | | | | | | | | | | | | | | | | | |
|---|---|---|---|---|---|---|---|---|---|---|---|---|---|---|---|---|---|---|---|---|---|---|
| PBANKA_0933700 | 0 | 0 | 174 | 27 | 459 | 260 | 427 | 186 | 110 | 63 | 8 | 5 | 396 | 134 | 79 | 238 | 604 | 103 | 3602 | 2248 | 21 | 205 |
| PBANKA_0933800 | 0 | 0 | 81 | 17 | 186 | 64 | 129 | 37 | 46 | 15 | 0 | 0 | 262 | 62 | 390 | 1438 | 629 | 100 | 1276 | 540 | 15 | 101 |
| PBANKA_0933900 | 0 | 0 | 181 | 28 | 228 | 140 | 202 | 72 | 39 | 24 | 24 | 10 | 681 | 251 | 496 | 1987 | 423 | 57 | 741 | 331 | 28 | 328 |
| PBANKA_0934000 | 14 | 27 | 39 | 5 | 74 | 27 | 62 | 20 | 63 | 49 | 65 | 42 | 356 | 157 | 80 | 343 | 657 | 276 | 380 | 172 | 16 | 114 |
| PBANKA_0934100 | 7 | 7 | 23 | 13 | 187 | 77 | 198 | 75 | 108 | 36 | 14 | 11 | 327 | 171 | 136 | 771 | 315 | 73 | 463 | 265 | 9 | 65 |
| PBANKA_0934200 | 128 | 187 | 465 | 139 | 505 | 421 | 582 | 213 | 226 | 177 | 64 | 50 | 3149 | 1343 | 1169 | 4374 | 1963 | 477 | 2057 | 1253 | 81 | 600 |
| PBANKA_0934300 | 0 | 0 | 90 | 15 | 275 | 104 | 409 | 168 | 191 | 95 | 18 | 8 | 1079 | 149 | 333 | 1179 | 4636 | 955 | 2360 | 1136 | 189 | 1640 |
| PBANKA_0934400 | 36 | 39 | 9 | 0 | 48 | 28 | 72 | 26 | 40 | 25 | 11 | 20 | 159 | 40 | 101 | 295 | 198 | 99 | 800 | 280 | 56 | 353 |
| PBANKA_0934500 | 114 | 161 | 79 | 23 | 171 | 77 | 228 | 59 | 316 | 174 | 822 | 517 | 2837 | 1730 | 1038 | 4103 | 3350 | 791 | 3278 | 2240 | 291 | 1726 |
| PBANKA_0934600 | 0 | 0 | 7 | 0 | 70 | 29 | 116 | 52 | 205 | 90 | 36 | 0 | 130 | 109 | 67 | 111 | 296 | 54 | 610 | 1636 | 81 | 722 |
| PBANKA_0934700 | 0 | 0 | 6 | 0 | 68 | 27 | 121 | 46 | 83 | 50 | 5 | 0 | 566 | 135 | 45 | 122 | 326 | 169 | 3156 | 1549 | 65 | 337 |
| PBANKA_0935000 | 6 | 14 | 24 | 16 | 219 | 94 | 639 | 267 | 474 | 203 | 67 | 21 | 702 | 188 | 419 | 2332 | 3563 | 421 | 1220 | 899 | 28 | 167 |
| PBANKA_0935100 | 0 | 0 | 28 | 12 | 49 | 25 | 173 | 84 | 139 | 67 | 17 | 10 | 197 | 41 | 103 | 412 | 142 | 22 | 382 | 98 | 41 | 157 |
| PBANKA_0935200 | 30 | 49 | 35 | 7 | 133 | 45 | 156 | 72 | 57 | 30 | 11 | 0 | 122 | 29 | 143 | 491 | 303 | 62 | 495 | 177 | 24 | 209 |
| PBANKA_0935300 | 14 | 24 | 84 | 13 | 287 | 82 | 327 | 106 | 129 | 47 | 46 | 10 | 367 | 112 | 288 | 1608 | 520 | 80 | 1237 | 774 | 110 | 680 |
| PBANKA_0935400 | 7 | 12 | 17 | 5 | 54 | 32 | 58 | 12 | 13 | 9 | 0 | 5 | 117 | 35 | 115 | 427 | 78 | 24 | 169 | 78 | 8 | 111 |
| PBANKA_0935500 | 16 | 11 | 253 | 54 | 807 | 296 | 1514 | 592 | 1028 | 474 | 158 | 90 | 2245 | 2012 | 653 | 2665 | 27413 | 3564 | 2221 | 8758 | 1598 | 11680 |
| PBANKA_0935600 | 39 | 30 | 32 | 8 | 41 | 25 | 39 | 27 | 21 | 16 | 9 | 8 | 107 | 76 | 86 | 165 | 80 | 38 | 217 | 43 | 0 | 104 |
| PBANKA_0935800 | 13 | 36 | 75 | 18 | 129 | 37 | 199 | 74 | 63 | 31 | 10 | 0 | 516 | 141 | 288 | 1165 | 1588 | 272 | 3417 | 2064 | 60 | 411 |
| PBANKA_0935900 | 19 | 28 | 119 | 20 | 228 | 80 | 375 | 91 | 136 | 54 | 31 | 18 | 1243 | 534 | 388 | 2158 | 1383 | 165 | 599 | 530 | 85 | 408 |
| PBANKA_0936100 | 0 | 0 | 31 | 0 | 81 | 48 | 119 | 60 | 29 | 30 | 0 | 0 | 82 | 22 | 118 | 383 | 352 | 67 | 284 | 122 | 36 | 128 |
| PBANKA_0936200 | 0 | 0 | 19 | 0 | 38 | 22 | 75 | 37 | 12 | 10 | 0 | 0 | 225 | 41 | 128 | 769 | 645 | 104 | 535 | 302 | 12 | 67 |
| PBANKA_0936300 | 14 | 13 | 33 | 11 | 47 | 29 | 115 | 41 | 75 | 23 | 160 | 91 | 840 | 402 | 99 | 433 | 858 | 213 | 924 | 431 | 331 | 2146 |
| PBANKA_0936400 | 0 | 0 | 125 | 27 | 418 | 125 | 568 | 224 | 202 | 72 | 40 | 25 | 2947 | 702 | 1512 | 8689 | 5097 | 837 | 4888 | 3745 | 228 | 1358 |
| PBANKA_0936500 | 11 | 10 | 62 | 15 | 258 | 137 | 401 | 140 | 156 | 70 | 11 | 0 | 424 | 88 | 279 | 756 | 1339 | 236 | 1568 | 479 | 45 | 273 |
| PBANKA_0936600 | 20 | 42 | 71 | 19 | 117 | 64 | 158 | 49 | 147 | 127 | 324 | 217 | 810 | 522 | 157 | 532 | 277 | 94 | 892 | 501 | 1992 | 12333 |
| PBANKA_0936700 | 18 | 23 | 39 | 7 | 98 | 59 | 139 | 59 | 169 | 93 | 451 | 319 | 601 | 475 | 290 | 1565 | 1300 | 249 | 1521 | 1690 | 2391 | 14865 |
| PBANKA_0936900 | 0 | 0 | 37 | 5 | 333 | 167 | 828 | 425 | 283 | 178 | 30 | 20 | 725 | 164 | 550 | 1774 | 1977 | 342 | 1429 | 423 | 221 | 1545 |
| PBANKA_0937000 | 0 | 0 | 74 | 21 | 94 | 77 | 108 | 75 | 53 | 42 | 13 | 9 | 124 | 26 | 168 | 482 | 319 | 81 | 198 | 130 | 50 | 298 |
| PBANKA_0937100 | 9 | 10 | 62 | 10 | 198 | 89 | 364 | 145 | 198 | 99 | 40 | 19 | 1179 | 307 | 606 | 2211 | 4328 | 723 | 3444 | 2572 | 117 | 861 |
| PBANKA_0937200 | 169 | 304 | 574 | 109 | 990 | 426 | 1052 | 361 | 573 | 223 | 3122 | 1896 | 9960 | 22440 | 5072 | 34920 | 9860 | 778 | 896 | 5528 | 3870 | 24988 |
| PBANKA_0937300 | 21 | 26 | 83 | 11 | 189 | 71 | 336 | 125 | 163 | 77 | 208 | 121 | 822 | 402 | 223 | 1260 | 443 | 92 | 1915 | 808 | 155 | 889 |
| PBANKA_0937400 | 0 | 5 | 107 | 23 | 234 | 102 | 445 | 158 | 246 | 136 | 116 | 72 | 1205 | 1117 | 796 | 3682 | 2369 | 330 | 529 | 880 | 72 | 567 |
| PBANKA_0937500 | 95 | 161 | 45 | 11 | 155 | 57 | 642 | 249 | 393 | 156 | 38 | 32 | 1774 | 672 | 286 | 1530 | 7536 | 951 | 12749 | 23213 | 1387 | 10539 |
| PBANKA_0937600 | 7 | 0 | 8 | 0 | 10 | 9 | 21 | 12 | 6 | 9 | 0 | 0 | 83 | 24 | 61 | 195 | 50 | 15 | 704 | 354 | 6 | 34 |
| PBANKA_0937700 | 19 | 28 | 657 | 185 | 501 | 267 | 465 | 279 | 150 | 112 | 673 | 547 | 20727 | 8577 | 2781 | 13570 | 4164 | 508 | 3562 | 2120 | 81 | 736 |
| PBANKA_0937800 | 59 | 82 | 236 | 48 | 245 | 81 | 242 | 65 | 154 | 58 | 1625 | 1120 | 3296 | 2279 | 326 | 1705 | 680 | 135 | 1117 | 1181 | 776 | 4535 |
| PBANKA_0937900 | 0 | 0 | 6 | 0 | 13 | 5 | 44 | 30 | 103 | 45 | 12 | 6 | 783 | 424 | 53 | 175 | 1744 | 513 | 2574 | 1484 | 24 | 121 |
| PBANKA_0938000 | 0 | 0 | 185 | 38 | 206 | 77 | 228 | 101 | 79 | 43 | 13 | 11 | 2101 | 2126 | 330 | 1313 | 1421 | 496 | 34309 | 91680 | 290 | 2129 |
| PBANKA_0938100 | 0 | 0 | 50 | 12 | 26 | 24 | 38 | 9 | 21 | 11 | 5 | 15 | 472 | 226 | 113 | 456 | 93 | 23 | 777 | 215 | 30 | 165 |
| PBANKA_0938200 | 0 | 0 | 16 | 0 | 46 | 34 | 57 | 24 | 21 | 6 | 0 | 0 | 82 | 20 | 16 | 48 | 83 | 15 | 1203 | 579 | 35 | 390 |
| PBANKA_0938300 | 0 | 0 | 150 | 47 | 464 | 250 | 831 | 322 | 321 | 178 | 49 | 13 | 3023 | 1132 | 1623 | 6896 | 5964 | 916 | 2815 | 1287 | 162 | 1123 |
| PBANKA_0938400 | 0 | 0 | 1140 | 308 | 2089 | 1296 | 3201 | 1654 | 937 | 601 | 98 | 77 | 5270 | 803 | 6600 | 22626 | 19424 | 2780 | 3543 | 2000 | 361 | 3520 |
| PBANKA_0938500 | 0 | 0 | 257 | 84 | 578 | 315 | 768 | 388 | 323 | 164 | 60 | 30 | 698 | 117 | 450 | 1585 | 2565 | 504 | 3707 | 2182 | 86 | 861 |
| PBANKA_0938600 | 0 | 0 | 238 | 50 | 341 | 145 | 328 | 128 | 111 | 45 | 40 | 31 | 3648 | 1426 | 1325 | 6212 | 1070 | 153 | 1518 | 1008 | 36 | 352 |
| PBANKA_0938700 | 8 | 11 | 61 | 10 | 113 | 32 | 70 | 41 | 52 | 23 | 0 | 5 | 86 | 10 | 95 | 488 | 105 | 22 | 77 | 38 | 33 | 176 |
| PBANKA_0938800 | 6 | 6 | 8 | 0 | 20 | 10 | 61 | 15 | 174 | 73 | 60 | 24 | 450 | 301 | 41 | 140 | 1085 | 323 | 4465 | 5792 | 1934 | 12117 |
| PBANKA_0938900 | 45 | 43 | 54 | 26 | 147 | 81 | 119 | 75 | 180 | 110 | 365 | 335 | 811 | 806 | 527 | 2547 | 2033 | 376 | 2273 | 2850 | 838 | 5663 |
| PBANKA_0939000 | 248 | 363 | 24 | 0 | 107 | 76 | 116 | 63 | 125 | 66 | 35 | 29 | 933 | 675 | 81 | 582 | 1053 | 278 | 17507 | 24053 | 867 | 6607 |
| PBANKA_0939100 | 637 | 861 | 362 | 92 | 727 | 219 | 719 | 277 | 2483 | 1290 | 4720 | 3186 | 8023 | 12096 | 1284 | 9154 | 20620 | 4278 | 5191 | 10068 | 2784 | 23116 |
| PBANKA_0939200 | 0 | 15 | 151 | 29 | 245 | 78 | 180 | 59 | 128 | 86 | 1431 | 803 | 3337 | 1982 | 696 | 2929 | 530 | 107 | 774 | 555 | 293 | 1466 |
| PBANKA_0939300 | 15 | 20 | 10 | 0 | 37 | 17 | 40 | 20 | 31 | 10 | 19 | 11 | 1158 | 790 | 275 | 1304 | 716 | 146 | 3098 | 6306 | 27 | 142 |
| PBANKA_0939400 | 11 | 24 | 42 | 8 | 66 | 61 | 63 | 34 | 25 | 30 | 0 | 0 | 381 | 111 | 86 | 368 | 279 | 58 | 134 | 118 | 78 | 644 |
| PBANKA_0939500 | 29 | 56 | 157 | 49 | 281 | 306 | 443 | 204 | 150 | 90 | 15 | 25 | 570 | 232 | 296 | 736 | 2296 | 397 | 6741 | 4192 | 894 | 9477 |
| PBANKA_0939600 | 109 | 150 | 1339 | 362 | 2359 | 1057 | 2826 | 1220 | 1752 | 752 | 12441 | 6529 | 12513 | 34942 | 7727 | 49526 | 33561 | 3731 | 4812 | 17150 | 693 | 4386 |
| PBANKA_0939700 | 0 | 0 | 48 | 7 | 229 | 126 | 271 | 117 | 103 | 53 | 23 | 14 | 523 | 168 | 396 | 1200 | 969 | 179 | 479 | 383 | 597 | 4010 |
| PBANKA_0939800 | 0 | 7 | 85 | 16 | 303 | 173 | 282 | 111 | 136 | 54 | 12 | 5 | 352 | 74 | 348 | 1623 | 881 | 128 | 567 | 398 | 25 | 263 |
| PBANKA_0939900 | 0 | 7 | 39 | 10 | 82 | 51 | 88 | 40 | 23 | 19 | 5 | 0 | 735 | 228 | 287 | 1289 | 719 | 70 | 295 | 174 | 14 | 218 |
| PBANKA_0940000 | 7 | 14 | 10 | 0 | 39 | 9 | 77 | 26 | 22 | 12 | 0 | 0 | 112 | 20 | 37 | 106 | 175 | 63 | 782 | 300 | 45 | 276 |
| PBANKA_0940100 | 0 | 0 | 98 | 19 | 352 | 101 | 514 | 224 | 512 | 307 | 82 | 82 | 3052 | 1300 | 1147 | 5793 | 2404 | 463 | 2446 | 2054 | 101 | 613 |
| PBANKA_0940200 | 7 | 7 | 57 | 17 | 337 | 105 | 373 | 146 | 239 | 105 | 57 | 12 | 717 | 474 | 526 | 2957 | 1781 | 258 | 672 | 1278 | 49 | 339 |
| PBANKA_0940300 | 0 | 0 | 57 | 20 | 234 | 96 | 173 | 57 | 76 | 46 | 11 | 19 | 413 | 298 | 327 | 1392 | 403 | 120 | 2827 | 1905 | 56 | 440 |
| PBANKA_0940400 | 0 | 0 | 55 | 10 | 111 | 49 | 120 | 38 | 53 | 18 | 21 | 8 | 596 | 394 | 316 | 1830 | 251 | 37 | 941 | 461 | 92 | 465 |
| PBANKA_0940500 | 0 | 0 | 13 | 0 | 29 | 10 | 21 | 7 | 11 | 5 | 0 | 0 | 58 | 18 | 51 | 125 | 39 | 13 | 351 | 112 | 0 | 17 |
| PBANKA_0940600 | 0 | 0 | 109 | 23 | 272 | 138 | 298 | 109 | 122 | 46 | 19 | 9 | 556 | 114 | 341 | 1552 | 602 | 91 | 1281 | 759 | 18 | 149 |
| PBANKA_0940700 | 0 | 5 | 148 | 23 | 293 | 118 | 347 | 98 | 94 | 35 | 49 | 17 | 2453 | 1432 | 448 | 2003 | 1319 | 217 | 778 | 332 | 36 | 236 |
| PBANKA_0940800 | 15 | 11 | 125 | 37 | 273 | 138 | 368 | 156 | 115 | 78 | 32 | 8 | 320 | 132 | 209 | 553 | 657 | 164 | 416 | 311 | 43 | 321 |
| PBANKA_0940900 | 0 | 0 | 223 | 43 | 418 | 189 | 358 | 151 | 111 | 47 | 62 | 21 | 2003 | 980 | 991 | 4202 | 1875 | 223 | 977 | 745 | 201 | 1465 |
| PBANKA_0941100 | 255 | 290 | 243 | 69 | 334 | 231 | 395 | 184 | 127 | 57 | 83 | 97 | 2399 | 1237 | 701 | 2214 | 1505 | 331 | 1781 | 791 | 1364 | 9505 |
| PBANKA_0941200 | 73 | 88 | 113 | 43 | 466 | 254 | 441 | 161 | 229 | 124 | 70 | 28 | 624 | 247 | 506 | 1853 | 568 | 144 | 887 | 489 | 197 | 1798 |
| PBANKA_0941300 | 0 | 0 | 49 | 8 | 164 | 63 | 201 | 57 | 620 | 316 | 475 | 256 | 2051 | 1219 | 668 | 3052 | 1382 | 745 | 1784 | 835 | 27 | 278 |
| PBANKA_0941400 | 0 | 0 | 28 | 6 | 84 | 25 | 55 | 41 | 45 | 16 | 13 | 0 | 224 | 71 | 31 | 138 | 376 | 63 | 1132 | 542 | 12 | 87 |
| PBANKA_0941500 | 17 | 35 | 2095 | 550 | 1311 | 777 | 1434 | 630 | 539 | 366 | 1721 | 1385 | 14361 | 4959 | 3040 | 11163 | 4424 | 622 | 5590 | 2235 | 363 | 3091 |
| PBANKA_0941600 | 0 | 0 | 17 | 6 | 142 | 50 | 539 | 196 | 458 | 252 | 58 | 27 | 2063 | 703 | 381 | 1471 | 10944 | 2028 | 4073 | 1992 | 183 | 1697 |
| PBANKA_0941700 | 14 | 18 | 9 | 0 | 40 | 11 | 54 | 39 | 19 | 21 | 0 | 5 | 150 | 25 | 9 | 15 | 135 | 49 | 1278 | 539 | 127 | 749 |
| PBANKA_0941800 | 227 | 301 | 6406 | 2523 | 16856 | 13151 | 29820 | 19800 | 23103 | 21657 | 3518 | 2418 | 2091 | 896 | 530 | 1506 | 15270 | 4651 | 3730 | 4069 | 568 | 6551 |
| PBANKA_0941900 | 639 | 658 | 2130 | 793 | 6787 | 4319 | 14033 | 8079 | 11829 | 8305 | 2246 | 1438 | 7042 | 2114 | 705 | 3080 | 22377 | 4592 | 12844 | 7048 | 1022 | 7245 |
| PBANKA_0942000 | 23 | 32 | 27 | 0 | 120 | 26 | 159 | 74 | 324 | 158 | 71 | 28 | 557 | 488 | 162 | 549 | 4718 | 966 | 2727 | 3325 | 75 | 526 |
| PBANKA_0942100 | 0 | 0 | 56 | 15 | 54 | 52 | 75 | 14 | 16 | 11 | 0 | 14 | 892 | 300 | 266 | 988 | 200 | 18 | 155 | 70 | 141 | 634 |
| PBANKA_0942200 | 0 | 0 | 127 | 39 | 185 | 86 | 136 | 59 | 38 | 17 | 0 | 0 | 737 | 258 | 368 | 1492 | 226 | 24 | 114 | 58 | 48 | 354 |
| PBANKA_0942500 | 8 | 0 | 270 | 82 | 583 | 345 | 751 | 352 | 231 | 153 | 336 | 278 | 4736 | 1660 | 2626 | 8399 | 5510 | 1014 | 1893 | 860 | 137 | 1078 |
| PBANKA_0942600 | 96 | 59 | 122 | 31 | 317 | 119 | 348 | 110 | 138 | 55 | 25 | 18 | 1021 | 544 | 343 | 2324 | 789 | 133 | 1441 | 1186 | 25 | 180 |
| PBANKA_0942700 | 117 | 182 | 578 | 145 | 1528 | 599 | 1513 | 501 | 659 | 236 | 254 | 179 | 3395 | 3743 | 1239 | 7816 | 24375 | 2779 | 1980 | 5759 | 262 | 1942 |
| PBANKA_0942800 | 62 | 85 | 53 | 9 | 108 | 60 | 109 | 34 | 48 | 16 | 36 | 21 | 1027 | 502 | 219 | 1482 | 360 | 54 | 216 | 212 | 783 | 4391 |
| PBANKA_0942900 | 12 | 14 | 85 | 26 | 248 | 154 | 359 | 107 | 96 | 54 | 296 | 247 | 1854 | 1216 | 631 | 3246 | 834 | 148 | 1445 | 1019 | 71 | 646 |
| PBANKA_0943000 | 0 | 0 | 6 | 0 | 9 | 5 | 18 | 8 | 7 | 0 | 12 | 9 | 75 | 29 | 57 | 81 | 29 | 36 | 345 | 104 | 0 | 11 |
| PBANKA_0943100 | 122 | 249 | 743 | 143 | 1613 | 719 | 1793 | 672 | 713 | 292 | 86 | 50 | 3618 | 4579 | 5899 | 24353 | 10720 | 1552 | 1422 | 4760 | 372 | 2632 |
| PBANKA_0943200 | 12 | 20 | 147 | 32 | 208 | 112 | 315 | 102 | 71 | 44 | 18 | 6 | 1536 | 520 | 496 | 1454 | 656 | 123 | 886 | 303 | 89 | 592 |
| PBANKA_0943300 | 7 | 6 | 90 | 16 | 210 | 108 | 246 | 98 | 72 | 44 | 17 | 5 | 483 | 139 | 376 | 1423 | 304 | 38 | 696 | 412 | 19 | 232 |
| PBANKA_0943400 | 13 | 23 | 23 | 0 | 85 | 47 | 71 | 22 | 26 | 15 | 7 | 0 | 2652 | 1387 | 50 | 163 | 969 | 446 | 61098 | 46999 | 545 | 5083 |

| | | | | | | | | | | | | | | | | | | | | | | |
|---|---|---|---|---|---|---|---|---|---|---|---|---|---|---|---|---|---|---|---|---|---|---|
| PBANKA_0943500 | 27 | 39 | 161 | 51 | 397 | 163 | 273 | 93 | 112 | 47 | 711 | 341 | 1413 | 718 | 688 | 2764 | 1003 | 154 | 406 | 254 | 8 | 72 |
| PBANKA_0943600 | 33 | 35 | 1459 | 457 | 616 | 398 | 623 | 379 | 228 | 120 | 1000 | 975 | 10506 | 3371 | 2652 | 9044 | 2971 | 426 | 3868 | 2146 | 124 | 1389 |
| PBANKA_0943900 | 0 | 0 | 6 | 0 | 14 | 7 | 50 | 48 | 43 | 57 | 17 | 0 | 220 | 11 | 30 | 7 | 61 | 512 | 55 | 0 | 0 | 25 |
| PBANKA_1000011 | 11 | 22 | 6 | 5 | 22 | 18 | 51 | 16 | 87 | 79 | 24 | 15 | 189 | 707 | 135 | 687 | 3688 | 439 | 75 | 676 | 52 | 222 |
| PBANKA_1000700 | 107 | 143 | 167 | 37 | 302 | 194 | 350 | 134 | 122 | 65 | 31 | 17 | 978 | 620 | 572 | 2813 | 897 | 97 | 436 | 630 | 27 | 204 |
| PBANKA_1000800 | 9 | 6 | 60 | 10 | 138 | 37 | 141 | 30 | 47 | 19 | 10 | 5 | 1309 | 962 | 719 | 5984 | 1290 | 88 | 287 | 753 | 30 | 179 |
| PBANKA_1000900 | 0 | 0 | 9 | 0 | 32 | 13 | 49 | 22 | 22 | 13 | 0 | 0 | 61 | 25 | 43 | 90 | 70 | 19 | 81 | 28 | 23 | 50 |
| PBANKA_1001000 | 285 | 415 | 282 | 54 | 1151 | 534 | 1817 | 715 | 1540 | 646 | 578 | 214 | 1772 | 3680 | 1991 | 7754 | 17293 | 2985 | 765 | 2254 | 222 | 1510 |
| PBANKA_1001100 | 175 | 262 | 63 | 10 | 104 | 62 | 135 | 55 | 58 | 31 | 12 | 7 | 149 | 53 | 351 | 1211 | 733 | 121 | 719 | 561 | 122 | 813 |
| PBANKA_1001200 | 23 | 32 | 6 | 0 | 29 | 6 | 27 | 13 | 43 | 12 | 5 | 0 | 109 | 24 | 92 | 177 | 261 | 65 | 444 | 438 | 10 | 51 |
| PBANKA_1001300 | 0 | 6 | 10 | 0 | 146 | 64 | 1044 | 443 | 1007 | 485 | 122 | 45 | 702 | 137 | 197 | 563 | 4133 | 507 | 1247 | 750 | 8 | 40 |
| PBANKA_1001400 | 0 | 0 | 219 | 31 | 482 | 208 | 772 | 327 | 368 | 184 | 164 | 87 | 2694 | 1575 | 895 | 3659 | 2868 | 558 | 3592 | 4610 | 34 | 312 |
| PBANKA_1001500 | 22 | 39 | 41 | 11 | 129 | 56 | 269 | 113 | 441 | 278 | 135 | 71 | 850 | 540 | 107 | 633 | 4840 | 717 | 1339 | 1275 | 157 | 1381 |
| PBANKA_1001600 | 25 | 34 | 111 | 36 | 496 | 205 | 661 | 275 | 312 | 170 | 78 | 70 | 525 | 414 | 742 | 3105 | 3727 | 570 | 1610 | 2735 | 113 | 858 |
| PBANKA_1001700 | 38 | 54 | 6 | 0 | 20 | 17 | 34 | 29 | 66 | 94 | 13 | 7 | 29 | 11 | 6 | 6 | 284 | 137 | 45 | 46 | 26 | 155 |
| PBANKA_1001800 | 55 | 96 | 118 | 39 | 616 | 275 | 750 | 302 | 378 | 169 | 70 | 22 | 262 | 452 | 435 | 1853 | 2063 | 232 | 130 | 442 | 324 | 3004 |
| PBANKA_1001900 | 9 | 17 | 30 | 8 | 212 | 102 | 240 | 101 | 59 | 19 | 15 | 18 | 271 | 77 | 126 | 479 | 224 | 27 | 49 | 24 | 0 | 23 |
| PBANKA_1002000 | 63 | 124 | 385 | 119 | 979 | 600 | 921 | 396 | 412 | 250 | 170 | 106 | 5759 | 6666 | 2024 | 18488 | 7808 | 508 | 3378 | 7102 | 166 | 1177 |
| PBANKA_1002100 | 1344 | 1804 | 27 | 0 | 125 | 46 | 122 | 35 | 93 | 41 | 8 | 8 | 422 | 194 | 0 | 36 | 1034 | 162 | 8270 | 7707 | 186 | 1455 |
| PBANKA_1002300 | 23 | 30 | 851 | 181 | 4108 | 1583 | 3178 | 839 | 1209 | 389 | 231 | 89 | 62 | 52 | 13 | 61 | 827 | 222 | 311 | 655 | 109 | 992 |
| PBANKA_1002400 | 9 | 12 | 41 | 11 | 198 | 60 | 368 | 114 | 1967 | 1061 | 384 | 150 | 985 | 481 | 14 | 48 | 11602 | 2962 | 586 | 808 | 24 | 133 |
| PBANKA_1002700 | 107 | 149 | 109 | 23 | 118 | 67 | 96 | 33 | 83 | 49 | 1132 | 851 | 1323 | 717 | 153 | 881 | 470 | 76 | 258 | 225 | 127 | 825 |
| PBANKA_1002800 | 14 | 16 | 84 | 22 | 522 | 362 | 1058 | 401 | 497 | 306 | 108 | 43 | 2581 | 1667 | 886 | 4199 | 4338 | 599 | 1361 | 1690 | 129 | 1378 |
| PBANKA_1003000 | 0 | 0 | 2799 | 600 | 37579 | 18117 | 46140 | 15216 | 13728 | 6225 | 1603 | 613 | 280 | 115 | 251 | 1023 | 531 | 69 | 22 | 43 | 0 | 10 |
| PBANKA_1003100 | 28 | 64 | 1063 | 304 | 2118 | 1183 | 3013 | 1081 | 1777 | 810 | 1772 | 1289 | 8236 | 7510 | 4820 | 19066 | 11187 | 1710 | 2756 | 3537 | 242 | 2583 |
| PBANKA_1003200 | 9 | 13 | 65 | 26 | 94 | 51 | 100 | 61 | 34 | 34 | 462 | 361 | 360 | 203 | 111 | 314 | 224 | 47 | 309 | 162 | 63 | 483 |
| PBANKA_1003300 | 0 | 0 | 44 | 9 | 202 | 88 | 269 | 152 | 145 | 77 | 440 | 266 | 877 | 512 | 227 | 853 | 846 | 172 | 604 | 259 | 30 | 212 |
| PBANKA_1003400 | 312 | 557 | 97 | 20 | 1086 | 457 | 1003 | 335 | 396 | 184 | 90 | 45 | 651 | 594 | 207 | 611 | 3134 | 507 | 6048 | 16841 | 692 | 6091 |
| PBANKA_1003500 | 0 | 0 | 26 | 6 | 82 | 44 | 78 | 34 | 14 | 9 | 0 | 0 | 341 | 78 | 162 | 892 | 267 | 37 | 963 | 545 | 18 | 130 |
| PBANKA_1003700 | 153 | 201 | 86 | 27 | 236 | 84 | 378 | 155 | 432 | 184 | 1274 | 757 | 3688 | 2237 | 638 | 3186 | 5749 | 1308 | 3342 | 2159 | 566 | 3658 |
| PBANKA_1003800 | 0 | 9 | 255 | 97 | 1283 | 856 | 1691 | 609 | 727 | 381 | 148 | 92 | 4072 | 1141 | 1700 | 8412 | 1941 | 256 | 3117 | 1860 | 121 | 1348 |
| PBANKA_1003900 | 0 | 0 | 5 | 0 | 682 | 311 | 5700 | 2696 | 5531 | 3346 | 492 | 251 | 34 | 18 | 8 | 36 | 55 | 8 | 61 | 28 | 0 | 34 |
| PBANKA_1004000 | 0 | 0 | 103 | 55 | 202 | 176 | 195 | 94 | 53 | 31 | 40 | 0 | 1493 | 469 | 605 | 1396 | 216 | 29 | 388 | 194 | 14 | 114 |
| PBANKA_1004100 | 0 | 0 | 26 | 0 | 276 | 101 | 338 | 138 | 189 | 96 | 38 | 14 | 334 | 85 | 115 | 616 | 1219 | 212 | 973 | 1107 | 149 | 833 |
| PBANKA_1004200 | 0 | 0 | 26 | 6 | 54 | 27 | 31 | 26 | 20 | 13 | 0 | 0 | 193 | 28 | 141 | 409 | 80 | 22 | 177 | 85 | 22 | 161 |
| PBANKA_1004300 | 0 | 0 | 25 | 0 | 103 | 37 | 260 | 116 | 145 | 60 | 79 | 70 | 908 | 234 | 257 | 788 | 2855 | 496 | 1456 | 832 | 52 | 400 |
| PBANKA_1004400 | 25 | 32 | 96 | 22 | 253 | 210 | 326 | 143 | 98 | 49 | 33 | 9 | 218 | 41 | 347 | 657 | 454 | 158 | 657 | 443 | 36 | 286 |
| PBANKA_1004500 | 0 | 0 | 13 | 0 | 40 | 34 | 68 | 46 | 76 | 25 | 11 | 6 | 170 | 314 | 59 | 354 | 684 | 159 | 1143 | 5967 | 390 | 2566 |
| PBANKA_1004600 | 0 | 0 | 44 | 11 | 83 | 42 | 106 | 44 | 43 | 18 | 15 | 6 | 550 | 249 | 92 | 370 | 863 | 168 | 570 | 257 | 81 | 525 |
| PBANKA_1004700 | 0 | 0 | 58 | 7 | 232 | 110 | 285 | 77 | 88 | 46 | 16 | 0 | 270 | 131 | 131 | 460 | 556 | 146 | 291 | 181 | 23 | 151 |
| PBANKA_1004800 | 6 | 13 | 101 | 32 | 233 | 144 | 266 | 140 | 103 | 80 | 75 | 53 | 423 | 191 | 339 | 1087 | 714 | 157 | 535 | 223 | 50 | 230 |
| PBANKA_1004900 | 43 | 70 | 82 | 11 | 153 | 56 | 142 | 56 | 86 | 47 | 406 | 223 | 2012 | 1268 | 560 | 3527 | 1246 | 175 | 1241 | 1056 | 187 | 1119 |
| PBANKA_1005000 | 6 | 6 | 5 | 0 | 37 | 13 | 39 | 13 | 16 | 6 | 24 | 17 | 167 | 97 | 38 | 291 | 88 | 34 | 202 | 172 | 28 | 277 |
| PBANKA_1005100 | 21 | 19 | 34 | 9 | 67 | 32 | 84 | 19 | 42 | 22 | 390 | 255 | 1735 | 950 | 281 | 1339 | 317 | 53 | 613 | 327 | 82 | 520 |
| PBANKA_1005200 | 556 | 787 | 31 | 13 | 99 | 50 | 90 | 45 | 70 | 25 | 23 | 6 | 61 | 20 | 94 | 431 | 495 | 62 | 104 | 87 | 0 | 15 |
| PBANKA_1005300 | 16 | 27 | 135 | 28 | 573 | 255 | 1079 | 371 | 822 | 446 | 85 | 32 | 1945 | 599 | 1675 | 12751 | 5375 | 637 | 2750 | 3655 | 82 | 583 |
| PBANKA_1005400 | 0 | 18 | 74 | 32 | 295 | 97 | 664 | 214 | 982 | 499 | 139 | 55 | 821 | 550 | 743 | 2779 | 14250 | 2715 | 725 | 2228 | 274 | 2019 |
| PBANKA_1005500 | 14 | 14 | 187 | 43 | 410 | 214 | 460 | 169 | 172 | 113 | 55 | 42 | 446 | 103 | 451 | 1981 | 1570 | 195 | 243 | 394 | 242 | 1611 |
| PBANKA_1005700 | 51 | 76 | 72 | 22 | 613 | 203 | 630 | 181 | 304 | 171 | 495 | 300 | 1207 | 1795 | 540 | 3116 | 3172 | 425 | 686 | 1606 | 90 | 669 |
| PBANKA_1005800 | 0 | 6 | 6 | 0 | 71 | 25 | 75 | 24 | 59 | 28 | 20 | 15 | 492 | 542 | 107 | 841 | 866 | 135 | 1797 | 3184 | 61 | 329 |
| PBANKA_1005900 | 25 | 35 | 23 | 10 | 108 | 76 | 160 | 89 | 72 | 47 | 35 | 23 | 1478 | 796 | 322 | 1786 | 1484 | 188 | 323 | 228 | 19 | 101 |
| PBANKA_1006000 | 0 | 0 | 74 | 16 | 134 | 45 | 103 | 48 | 44 | 27 | 6 | 0 | 375 | 78 | 373 | 1697 | 246 | 36 | 43 | 80 | 0 | 19 |
| PBANKA_1006100 | 173 | 255 | 61 | 15 | 216 | 85 | 443 | 162 | 179 | 109 | 25 | 11 | 1296 | 366 | 275 | 1341 | 7055 | 850 | 4245 | 3204 | 383 | 1922 |
| PBANKA_1006200 | 2310 | 3226 | 14 | 8 | 46 | 28 | 22 | 26 | 42 | 11 | 41 | 0 | 119 | 38 | 11 | 11 | 56 | 14 | 1371 | 686 | 6072 | 54078 |
| PBANKA_1006300 | 4223 | 5957 | 8 | 0 | 88 | 55 | 95 | 21 | 42 | 24 | 14 | 0 | 76 | 13 | 5 | 10 | 70 | 13 | 752 | 418 | 108 | 1043 |
| PBANKA_1006500 | 0 | 0 | 58 | 5 | 180 | 68 | 104 | 64 | 62 | 26 | 9 | 6 | 311 | 160 | 181 | 658 | 151 | 32 | 144 | 60 | 448 | 2760 |
| PBANKA_1006600 | 5 | 0 | 44 | 12 | 167 | 100 | 233 | 71 | 106 | 56 | 17 | 6 | 537 | 380 | 517 | 1972 | 2127 | 300 | 659 | 825 | 28 | 222 |
| PBANKA_1006700 | 103 | 132 | 120 | 44 | 202 | 95 | 300 | 110 | 168 | 112 | 951 | 586 | 2001 | 1093 | 567 | 2528 | 1459 | 290 | 1002 | 526 | 297 | 1738 |
| PBANKA_1006800 | 103 | 131 | 84 | 24 | 101 | 59 | 110 | 65 | 126 | 92 | 979 | 743 | 1472 | 1255 | 314 | 1113 | 510 | 129 | 327 | 296 | 209 | 1417 |
| PBANKA_1006900 | 46 | 50 | 148 | 38 | 233 | 104 | 152 | 36 | 60 | 30 | 153 | 134 | 1981 | 1102 | 598 | 3000 | 368 | 57 | 495 | 622 | 29 | 170 |
| PBANKA_1007000 | 110 | 157 | 278 | 108 | 564 | 375 | 452 | 155 | 130 | 58 | 387 | 259 | 1990 | 1191 | 692 | 3050 | 347 | 57 | 441 | 284 | 13 | 177 |
| PBANKA_1007100 | 0 | 7 | 29 | 5 | 113 | 39 | 176 | 80 | 42 | 31 | 62 | 38 | 1324 | 597 | 399 | 1318 | 1221 | 233 | 293 | 190 | 15 | 117 |
| PBANKA_1007200 | 15 | 32 | 148 | 52 | 712 | 430 | 905 | 427 | 381 | 185 | 57 | 40 | 920 | 435 | 748 | 3513 | 3299 | 436 | 2287 | 2801 | 214 | 1867 |
| PBANKA_1007300 | 7 | 16 | 35 | 14 | 170 | 88 | 185 | 80 | 61 | 25 | 21 | 10 | 282 | 51 | 239 | 935 | 377 | 74 | 1266 | 477 | 32 | 299 |
| PBANKA_1007400 | 126 | 154 | 90 | 25 | 231 | 60 | 229 | 93 | 344 | 173 | 1248 | 828 | 4119 | 2500 | 706 | 4439 | 3296 | 659 | 1454 | 1122 | 418 | 3239 |
| PBANKA_1007500 | 44 | 46 | 37 | 5 | 78 | 32 | 78 | 29 | 107 | 63 | 325 | 180 | 733 | 393 | 317 | 856 | 624 | 273 | 1409 | 657 | 144 | 785 |
| PBANKA_1007600 | 72 | 106 | 23 | 0 | 100 | 63 | 97 | 45 | 54 | 31 | 0 | 13 | 2331 | 815 | 433 | 2550 | 788 | 127 | 2016 | 1255 | 78 | 498 |
| PBANKA_1007700 | 80 | 104 | 192 | 68 | 624 | 430 | 901 | 370 | 383 | 191 | 48 | 37 | 3014 | 1427 | 1166 | 5717 | 3758 | 439 | 3081 | 2185 | 129 | 1202 |
| PBANKA_1007800 | 0 | 0 | 79 | 16 | 217 | 92 | 227 | 105 | 74 | 39 | 16 | 6 | 522 | 264 | 258 | 1287 | 535 | 76 | 333 | 236 | 26 | 206 |
| PBANKA_1008000 | 226 | 262 | 123 | 35 | 203 | 148 | 180 | 87 | 87 | 45 | 134 | 120 | 871 | 445 | 444 | 1463 | 258 | 53 | 905 | 377 | 255 | 1525 |
| PBANKA_1008100 | 38 | 51 | 81 | 34 | 222 | 73 | 185 | 65 | 62 | 34 | 32 | 19 | 938 | 401 | 709 | 4026 | 1559 | 183 | 776 | 636 | 104 | 720 |
| PBANKA_1008200 | 0 | 0 | 121 | 21 | 376 | 220 | 543 | 234 | 230 | 111 | 16 | 15 | 814 | 155 | 796 | 2419 | 2552 | 350 | 450 | 284 | 6 | 48 |
| PBANKA_1008300 | 15 | 31 | 64 | 21 | 142 | 68 | 102 | 39 | 38 | 20 | 12 | 10 | 415 | 147 | 202 | 795 | 210 | 63 | 7180 | 4424 | 28 | 289 |
| PBANKA_1008400 | 0 | 0 | 97 | 13 | 220 | 86 | 200 | 66 | 68 | 33 | 18 | 5 | 398 | 109 | 528 | 2963 | 502 | 84 | 206 | 277 | 7 | 49 |
| PBANKA_1008500 | 16 | 10 | 166 | 57 | 189 | 112 | 228 | 164 | 4074 | 2842 | 13220 | 9826 | 22375 | 21958 | 4571 | 35790 | 15976 | 3094 | 3064 | 4573 | 420 | 4371 |
| PBANKA_1008600 | 62 | 76 | 8 | 0 | 64 | 24 | 292 | 113 | 2352 | 1095 | 743 | 307 | 701 | 1291 | 148 | 514 | 12322 | 2980 | 1024 | 1220 | 29 | 167 |
| PBANKA_1008700 | 168 | 233 | 51 | 8 | 243 | 76 | 387 | 146 | 253 | 81 | 46 | 19 | 1234 | 538 | 320 | 1653 | 7984 | 930 | 1771 | 3061 | 120 | 747 |
| PBANKA_1008800 | 29 | 35 | 390 | 85 | 336 | 301 | 735 | 372 | 298 | 171 | 57 | 33 | 5756 | 1594 | 1716 | 7980 | 3551 | 646 | 5793 | 3648 | 363 | 2779 |
| PBANKA_1008900 | 20 | 37 | 30 | 9 | 133 | 54 | 123 | 61 | 38 | 15 | 8 | 0 | 280 | 72 | 152 | 689 | 195 | 59 | 3058 | 1443 | 158 | 1227 |
| PBANKA_1009000 | 6 | 0 | 88 | 11 | 84 | 32 | 87 | 26 | 37 | 11 | 0 | 0 | 459 | 81 | 214 | 1477 | 149 | 28 | 1529 | 907 | 65 | 557 |
| PBANKA_1009100 | 0 | 0 | 249 | 66 | 746 | 447 | 772 | 294 | 285 | 178 | 36 | 19 | 4034 | 610 | 1423 | 6157 | 2518 | 568 | 6377 | 4013 | 442 | 4318 |
| PBANKA_1009200 | 0 | 5 | 67 | 20 | 489 | 186 | 534 | 148 | 1396 | 710 | 465 | 233 | 817 | 1262 | 86 | 209 | 9137 | 1920 | 12282 | 36945 | 123 | 883 |
| PBANKA_1009300 | 8 | 0 | 110 | 8 | 287 | 106 | 572 | 239 | 171 | 91 | 105 | 59 | 2730 | 1072 | 989 | 4514 | 3296 | 409 | 1113 | 953 | 87 | 656 |
| PBANKA_1009400 | 0 | 0 | 49 | 10 | 123 | 57 | 218 | 60 | 52 | 31 | 22 | 30 | 643 | 300 | 363 | 1346 | 826 | 127 | 304 | 246 | 34 | 326 |
| PBANKA_1009500 | 28 | 50 | 778 | 243 | 1493 | 761 | 1415 | 471 | 1114 | 648 | 199 | 74 | 3285 | 2408 | 1159 | 5322 | 8493 | 1834 | 12142 | 16058 | 2127 | 16822 |
| PBANKA_1009700 | 19 | 26 | 43 | 5 | 99 | 51 | 98 | 46 | 29 | 16 | 0 | 0 | 162 | 42 | 216 | 959 | 411 | 69 | 1398 | 453 | 11 | 74 |
| PBANKA_1009800 | 12 | 14 | 132 | 25 | 161 | 79 | 128 | 40 | 41 | 16 | 5 | 5 | 196 | 60 | 287 | 1208 | 284 | 61 | 1118 | 781 | 115 | 1162 |
| PBANKA_1010000 | 185 | 201 | 849 | 308 | 582 | 443 | 449 | 237 | 86 | 50 | 17 | 24 | 2842 | 683 | 1831 | 6061 | 533 | 88 | 610 | 318 | 49 | 444 |
| PBANKA_1010100 | 47 | 60 | 64 | 15 | 116 | 82 | 124 | 34 | 38 | 20 | 18 | 0 | 711 | 197 | 159 | 1047 | 168 | 34 | 254 | 194 | 31 | 323 |
| PBANKA_1010200 | 143 | 165 | 179 | 37 | 294 | 159 | 254 | 88 | 121 | 36 | 752 | 532 | 3566 | 2380 | 1463 | 8583 | 1145 | 129 | 1068 | 1326 | 445 | 3304 |

| | | | | | | | | | | | | | | | | | | | | | | |
|---|---|---|---|---|---|---|---|---|---|---|---|---|---|---|---|---|---|---|---|---|---|---|
| PBANKA_1010300 | 320 | 444 | 1459 | 353 | 2378 | 833 | 2106 | 718 | 1353 | 622 | 5315 | 3118 | 2591 | 11189 | 2390 | 14399 | 4150 | 461 | 1241 | 10730 | 1479 | 10505 |
| PBANKA_1010500 | 239 | 330 | 959 | 283 | 1148 | 546 | 1183 | 443 | 372 | 205 | 844 | 581 | 7956 | 5829 | 5041 | 21549 | 4796 | 862 | 1957 | 2924 | 348 | 2769 |
| PBANKA_1010600 | 19 | 27 | 341 | 96 | 645 | 479 | 1452 | 938 | 507 | 419 | 63 | 54 | 3243 | 692 | 1156 | 4135 | 6560 | 1048 | 2639 | 1271 | 99 | 848 |
| PBANKA_1010700 | 0 | 0 | 69 | 6 | 80 | 22 | 59 | 17 | 21 | 7 | 41 | 31 | 1023 | 714 | 133 | 714 | 169 | 39 | 722 | 441 | 16 | 78 |
| PBANKA_1010800 | 0 | 0 | 236 | 77 | 247 | 189 | 210 | 122 | 59 | 35 | 84 | 72 | 2051 | 897 | 962 | 2370 | 463 | 113 | 428 | 258 | 26 | 268 |
| PBANKA_1010900 | 39 | 76 | 26 | 0 | 134 | 53 | 268 | 77 | 150 | 50 | 42 | 10 | 308 | 131 | 267 | 749 | 1288 | 173 | 1963 | 2730 | 170 | 942 |
| PBANKA_1011100 | 35 | 65 | 76 | 24 | 630 | 245 | 1018 | 357 | 579 | 199 | 83 | 15 | 1127 | 487 | 525 | 1571 | 6049 | 782 | 4189 | 4535 | 198 | 1517 |
| PBANKA_1011200 | 50 | 56 | 76 | 16 | 221 | 87 | 288 | 98 | 293 | 113 | 91 | 41 | 541 | 255 | 397 | 2061 | 2557 | 472 | 1791 | 2009 | 357 | 2381 |
| PBANKA_1011300 | 0 | 0 | 66 | 11 | 110 | 61 | 121 | 42 | 31 | 13 | 0 | 0 | 232 | 81 | 119 | 394 | 153 | 69 | 201 | 96 | 16 | 175 |
| PBANKA_1011400 | 17 | 14 | 119 | 32 | 401 | 182 | 399 | 162 | 373 | 172 | 220 | 151 | 1323 | 650 | 1703 | 7200 | 9045 | 1451 | 6019 | 6681 | 61 | 550 |
| PBANKA_1011500 | 107 | 130 | 341 | 66 | 2407 | 762 | 4228 | 1382 | 3351 | 1266 | 1832 | 873 | 3232 | 5164 | 6620 | 29662 | 46267 | 7286 | 2260 | 9646 | 228 | 1164 |
| PBANKA_1011600 | 13 | 5 | 77 | 19 | 194 | 136 | 216 | 117 | 201 | 73 | 31 | 15 | 317 | 109 | 215 | 692 | 1210 | 216 | 2427 | 1271 | 856 | 6174 |
| PBANKA_1011700 | 0 | 0 | 10 | 0 | 84 | 33 | 68 | 37 | 63 | 44 | 14 | 0 | 396 | 200 | 208 | 829 | 782 | 180 | 1848 | 3131 | 24 | 122 |
| PBANKA_1011800 | 10 | 6 | 54 | 9 | 224 | 107 | 324 | 137 | 145 | 71 | 150 | 85 | 2149 | 835 | 494 | 2168 | 2068 | 374 | 1262 | 605 | 173 | 902 |
| PBANKA_1011900 | 0 | 0 | 17 | 0 | 38 | 21 | 29 | 7 | 10 | 8 | 20 | 16 | 235 | 66 | 119 | 139 | 78 | 50 | 90 | 53 | 15 | 79 |
| PBANKA_1012000 | 10 | 9 | 13 | 0 | 45 | 15 | 96 | 37 | 225 | 94 | 39 | 7 | 483 | 139 | 49 | 198 | 1539 | 370 | 555 | 285 | 26 | 219 |
| PBANKA_1012100 | 181 | 362 | 86 | 17 | 335 | 133 | 375 | 156 | 200 | 59 | 122 | 81 | 3058 | 1830 | 457 | 2551 | 2500 | 367 | 1183 | 806 | 724 | 5654 |
| PBANKA_1012200 | 12 | 23 | 65 | 28 | 103 | 70 | 113 | 45 | 39 | 25 | 6 | 5 | 110 | 44 | 203 | 582 | 307 | 42 | 1006 | 413 | 193 | 1148 |
| PBANKA_1012300 | 290 | 341 | 161 | 37 | 414 | 141 | 376 | 121 | 226 | 100 | 104 | 60 | 849 | 528 | 831 | 5018 | 3140 | 527 | 1051 | 2320 | 58 | 502 |
| PBANKA_1012400 | 12 | 26 | 37 | 6 | 109 | 45 | 83 | 32 | 117 | 71 | 486 | 527 | 1062 | 1009 | 321 | 1452 | 936 | 284 | 1398 | 1322 | 416 | 2539 |
| PBANKA_1012500 | 33 | 30 | 43 | 6 | 132 | 58 | 94 | 52 | 143 | 85 | 126 | 72 | 702 | 816 | 386 | 1672 | 768 | 153 | 2466 | 2441 | 2885 | 14718 |
| PBANKA_1012600 | 114 | 189 | 39 | 10 | 207 | 67 | 216 | 76 | 78 | 31 | 0 | 7 | 160 | 71 | 227 | 811 | 750 | 102 | 689 | 840 | 881 | 5231 |
| PBANKA_1012700 | 72 | 110 | 23 | 0 | 94 | 20 | 99 | 35 | 92 | 44 | 39 | 9 | 554 | 202 | 142 | 601 | 1049 | 185 | 3523 | 4197 | 585 | 3080 |
| PBANKA_1012800 | 24 | 30 | 292 | 63 | 390 | 184 | 431 | 211 | 143 | 69 | 47 | 21 | 1501 | 403 | 708 | 1609 | 683 | 184 | 676 | 227 | 99 | 523 |
| PBANKA_1012900 | 19 | 21 | 47 | 10 | 166 | 52 | 266 | 115 | 84 | 33 | 37 | 18 | 1152 | 396 | 369 | 1637 | 1681 | 228 | 381 | 249 | 91 | 511 |
| PBANKA_1013000 | 5 | 0 | 5 | 0 | 39 | 15 | 76 | 25 | 52 | 20 | 15 | 0 | 810 | 175 | 135 | 409 | 379 | 184 | 3280 | 1569 | 107 | 858 |
| PBANKA_1013100 | 0 | 8 | 1593 | 340 | 850 | 406 | 796 | 262 | 239 | 200 | 1053 | 642 | 7150 | 3039 | 1752 | 4900 | 982 | 268 | 1621 | 660 | 183 | 1168 |
| PBANKA_1013200 | 216 | 333 | 254 | 61 | 507 | 212 | 747 | 278 | 313 | 151 | 22 | 13 | 610 | 234 | 441 | 1606 | 2099 | 297 | 3233 | 2645 | 1367 | 10914 |
| PBANKA_1013300 | 103 | 175 | 32 | 5 | 198 | 65 | 210 | 106 | 113 | 59 | 29 | 11 | 307 | 148 | 187 | 752 | 999 | 145 | 1099 | 1101 | 1310 | 9179 |
| PBANKA_1013400 | 10 | 10 | 32 | 0 | 99 | 33 | 158 | 77 | 367 | 166 | 130 | 67 | 1067 | 367 | 132 | 503 | 6944 | 1456 | 985 | 1077 | 127 | 840 |
| PBANKA_1013500 | 38 | 50 | 167 | 62 | 372 | 144 | 384 | 133 | 288 | 148 | 650 | 354 | 4487 | 2759 | 969 | 6661 | 2635 | 743 | 1448 | 1535 | 265 | 1886 |
| PBANKA_1013700 | 6 | 0 | 88 | 20 | 168 | 81 | 144 | 80 | 86 | 42 | 10 | 5 | 333 | 46 | 222 | 682 | 279 | 53 | 864 | 313 | 53 | 344 |
| PBANKA_1013800 | 80 | 101 | 129 | 30 | 342 | 175 | 497 | 220 | 182 | 148 | 40 | 27 | 393 | 109 | 344 | 1149 | 1522 | 372 | 2880 | 1612 | 248 | 2240 |
| PBANKA_1014000 | 0 | 8 | 227 | 63 | 1343 | 870 | 1775 | 681 | 583 | 361 | 57 | 47 | 321 | 77 | 416 | 1706 | 1239 | 209 | 285 | 202 | 83 | 667 |
| PBANKA_1014100 | 0 | 0 | 27 | 6 | 144 | 55 | 170 | 66 | 59 | 34 | 7 | 0 | 238 | 63 | 157 | 545 | 641 | 103 | 849 | 650 | 529 | 3879 |
| PBANKA_1014300 | 0 | 0 | 101 | 35 | 248 | 103 | 256 | 88 | 82 | 42 | 23 | 7 | 1328 | 415 | 712 | 3320 | 1347 | 242 | 969 | 525 | 813 | 5560 |
| PBANKA_1014400 | 0 | 0 | 221 | 60 | 755 | 406 | 602 | 257 | 210 | 113 | 68 | 15 | 631 | 584 | 1560 | 8871 | 1832 | 248 | 612 | 2578 | 908 | 6562 |
| PBANKA_1014600 | 0 | 7 | 41 | 0 | 103 | 44 | 137 | 68 | 91 | 43 | 8 | 8 | 301 | 62 | 204 | 696 | 699 | 190 | 421 | 192 | 106 | 750 |
| PBANKA_1014700 | 0 | 8 | 24 | 6 | 124 | 62 | 214 | 81 | 98 | 49 | 0 | 0 | 215 | 24 | 101 | 246 | 788 | 187 | 197 | 91 | 211 | 999 |
| PBANKA_1014800 | 0 | 12 | 28 | 13 | 216 | 126 | 278 | 115 | 172 | 78 | 39 | 0 | 565 | 303 | 236 | 766 | 1790 | 324 | 4387 | 8808 | 1187 | 9473 |
| PBANKA_1014900 | 0 | 5 | 165 | 39 | 524 | 304 | 924 | 345 | 318 | 178 | 69 | 39 | 2474 | 1018 | 1421 | 6098 | 5111 | 688 | 2298 | 1779 | 797 | 5602 |
| PBANKA_1015000 | 0 | 0 | 57 | 9 | 187 | 87 | 162 | 66 | 84 | 39 | 7 | 0 | 639 | 177 | 471 | 1900 | 730 | 127 | 2298 | 1740 | 329 | 2430 |
| PBANKA_1015100 | 30 | 43 | 82 | 15 | 145 | 59 | 157 | 74 | 52 | 24 | 5 | 5 | 229 | 33 | 259 | 1053 | 385 | 70 | 716 | 494 | 160 | 957 |
| PBANKA_1015200 | 46 | 64 | 88 | 32 | 175 | 94 | 231 | 85 | 110 | 90 | 191 | 143 | 1256 | 829 | 284 | 1397 | 1113 | 161 | 517 | 335 | 3348 | 21895 |
| PBANKA_1015300 | 23 | 17 | 98 | 19 | 312 | 98 | 302 | 137 | 222 | 83 | 77 | 35 | 504 | 334 | 1034 | 4581 | 4572 | 596 | 1061 | 2315 | 701 | 3651 |
| PBANKA_1015400 | 31 | 50 | 16 | 15 | 118 | 57 | 132 | 53 | 160 | 84 | 57 | 31 | 352 | 158 | 65 | 188 | 388 | 399 | 5295 | 2468 | 404 | 3067 |
| PBANKA_1015500 | 18 | 17 | 44 | 13 | 166 | 82 | 132 | 43 | 110 | 32 | 32 | 6 | 357 | 139 | 153 | 578 | 1263 | 319 | 1517 | 1674 | 503 | 2705 |
| PBANKA_1015600 | 29 | 36 | 52 | 19 | 184 | 72 | 223 | 75 | 103 | 39 | 22 | 0 | 887 | 344 | 483 | 2755 | 2869 | 348 | 152 | 254 | 136 | 750 |
| PBANKA_1015700 | 23 | 31 | 56 | 11 | 46 | 19 | 44 | 13 | 26 | 10 | 5 | 7 | 474 | 146 | 76 | 374 | 211 | 19 | 204 | 129 | 66 | 480 |
| PBANKA_1015800 | 137 | 223 | 127 | 24 | 510 | 201 | 824 | 362 | 422 | 177 | 62 | 14 | 522 | 325 | 357 | 1621 | 4423 | 535 | 459 | 891 | 2596 | 15161 |
| PBANKA_1015900 | 42 | 63 | 24 | 9 | 155 | 60 | 269 | 118 | 151 | 80 | 14 | 6 | 270 | 95 | 173 | 711 | 1145 | 312 | 738 | 321 | 417 | 2851 |
| PBANKA_1016000 | 53 | 82 | 42 | 10 | 74 | 24 | 107 | 48 | 80 | 39 | 17 | 5 | 772 | 212 | 247 | 834 | 1624 | 602 | 3471 | 2263 | 127 | 852 |
| PBANKA_1016100 | 0 | 0 | 38 | 9 | 120 | 70 | 152 | 69 | 55 | 22 | 8 | 0 | 325 | 64 | 150 | 549 | 811 | 139 | 235 | 145 | 5 | 45 |
| PBANKA_1016200 | 118 | 158 | 96 | 18 | 211 | 85 | 262 | 91 | 127 | 96 | 28 | 13 | 581 | 383 | 438 | 1344 | 3560 | 719 | 1609 | 3084 | 868 | 5621 |
| PBANKA_1016300 | 126 | 155 | 452 | 117 | 537 | 247 | 531 | 222 | 280 | 164 | 205 | 129 | 3518 | 1937 | 1731 | 6324 | 2434 | 656 | 1257 | 912 | 195 | 1469 |
| PBANKA_1016400 | 72 | 68 | 80 | 15 | 169 | 51 | 254 | 89 | 213 | 118 | 44 | 12 | 894 | 356 | 415 | 1422 | 3719 | 820 | 1448 | 1307 | 216 | 1186 |
| PBANKA_1016500 | 32 | 37 | 80 | 12 | 136 | 44 | 215 | 54 | 181 | 76 | 133 | 96 | 1738 | 1121 | 426 | 2298 | 4519 | 739 | 1803 | 1541 | 209 | 1491 |
| PBANKA_1016600 | 0 | 0 | 29 | 5 | 58 | 25 | 70 | 35 | 21 | 5 | 9 | 0 | 258 | 53 | 340 | 802 | 648 | 160 | 757 | 298 | 40 | 215 |
| PBANKA_1016700 | 5 | 10 | 969 | 182 | 956 | 401 | 1403 | 578 | 446 | 226 | 50 | 20 | 10492 | 1797 | 5007 | 26359 | 27660 | 3152 | 13758 | 8261 | 204 | 1727 |
| PBANKA_1016800 | 0 | 0 | 55 | 13 | 101 | 58 | 99 | 55 | 32 | 12 | 15 | 7 | 523 | 307 | 217 | 1063 | 273 | 38 | 198 | 78 | 14 | 102 |
| PBANKA_1017000 | 0 | 5 | 6 | 0 | 10 | 0 | 22 | 12 | 14 | 8 | 9 | 0 | 18 | 0 | 8 | 50 | 55 | 17 | 44 | 58 | 1469 | 10595 |
| PBANKA_1017100 | 57 | 72 | 173 | 46 | 474 | 152 | 834 | 349 | 379 | 203 | 75 | 24 | 483 | 119 | 604 | 2073 | 4600 | 663 | 273 | 258 | 130 | 713 |
| PBANKA_1017200 | 138 | 209 | 62 | 20 | 158 | 106 | 329 | 159 | 197 | 86 | 20 | 9 | 880 | 272 | 339 | 1101 | 2882 | 621 | 943 | 499 | 134 | 786 |
| PBANKA_1017300 | 79 | 68 | 157 | 39 | 317 | 138 | 627 | 272 | 295 | 130 | 27 | 5 | 1916 | 945 | 488 | 1978 | 7556 | 1079 | 20057 | 18717 | 65 | 565 |
| PBANKA_1017500 | 2509 | 3457 | 27 | 8 | 256 | 112 | 249 | 91 | 151 | 51 | 35 | 8 | 157 | 69 | 105 | 764 | 629 | 114 | 608 | 1173 | 191 | 1015 |
| PBANKA_1017600 | 8 | 22 | 111 | 21 | 297 | 140 | 500 | 196 | 300 | 126 | 30 | 14 | 618 | 289 | 400 | 2745 | 3600 | 395 | 1193 | 2498 | 1256 | 8327 |
| PBANKA_1017700 | 0 | 0 | 77 | 21 | 142 | 105 | 250 | 168 | 96 | 80 | 11 | 6 | 532 | 65 | 382 | 929 | 1841 | 392 | 1095 | 530 | 124 | 831 |
| PBANKA_1017800 | 498 | 697 | 96 | 18 | 489 | 205 | 604 | 209 | 287 | 133 | 111 | 63 | 2224 | 1331 | 1225 | 6060 | 3651 | 436 | 2722 | 2474 | 114 | 860 |
| PBANKA_1017900 | 352 | 347 | 56 | 9 | 219 | 96 | 231 | 101 | 158 | 78 | 593 | 439 | 883 | 754 | 622 | 2192 | 429 | 115 | 983 | 960 | 1091 | 7459 |
| PBANKA_1018000 | 54 | 78 | 113 | 22 | 279 | 99 | 243 | 77 | 125 | 56 | 215 | 131 | 1378 | 1335 | 317 | 1703 | 596 | 124 | 1342 | 660 | 1491 | 8435 |
| PBANKA_1018100 | 29 | 24 | 38 | 16 | 145 | 78 | 144 | 58 | 54 | 31 | 142 | 72 | 1260 | 574 | 133 | 497 | 485 | 243 | 22511 | 15445 | 371 | 2994 |
| PBANKA_1018300 | 0 | 0 | 6 | 0 | 26 | 13 | 23 | 10 | 21 | 0 | 7 | 0 | 262 | 123 | 45 | 150 | 112 | 54 | 2645 | 1734 | 52 | 483 |
| PBANKA_1018400 | 18 | 40 | 221 | 40 | 193 | 104 | 133 | 48 | 45 | 27 | 13 | 17 | 3815 | 1560 | 711 | 3075 | 287 | 62 | 1539 | 908 | 212 | 1681 |
| PBANKA_1018500 | 49 | 34 | 728 | 159 | 421 | 231 | 332 | 154 | 357 | 226 | 457 | 284 | 7698 | 2184 | 1888 | 6530 | 1289 | 332 | 1205 | 559 | 400 | 2191 |
| PBANKA_1018600 | 93 | 110 | 2191 | 572 | 1488 | 850 | 1390 | 664 | 592 | 420 | 2728 | 2131 | 16685 | 7717 | 3083 | 13249 | 5514 | 851 | 2108 | 1600 | 360 | 3034 |
| PBANKA_1018700 | 0 | 0 | 75 | 33 | 83 | 77 | 111 | 58 | 38 | 17 | 218 | 179 | 1481 | 723 | 539 | 1712 | 308 | 72 | 632 | 276 | 45 | 353 |
| PBANKA_1018800 | 39 | 86 | 17 | 9 | 224 | 83 | 404 | 147 | 172 | 85 | 21 | 9 | 708 | 315 | 157 | 488 | 3927 | 386 | 2141 | 5472 | 875 | 6195 |
| PBANKA_1018900 | 0 | 0 | 18 | 0 | 57 | 15 | 106 | 35 | 77 | 39 | 23 | 5 | 824 | 268 | 117 | 569 | 1922 | 354 | 933 | 510 | 198 | 1269 |
| PBANKA_1019100 | 38 | 39 | 28 | 0 | 86 | 61 | 192 | 92 | 146 | 52 | 16 | 6 | 259 | 120 | 127 | 554 | 2110 | 354 | 798 | 764 | 910 | 6725 |
| PBANKA_1019200 | 0 | 0 | 57 | 14 | 120 | 56 | 197 | 77 | 107 | 58 | 17 | 0 | 458 | 111 | 304 | 1116 | 1948 | 419 | 994 | 818 | 586 | 3581 |
| PBANKA_1019300 | 0 | 0 | 17 | 5 | 72 | 50 | 104 | 45 | 37 | 23 | 12 | 0 | 232 | 106 | 197 | 677 | 324 | 42 | 144 | 64 | 15 | 129 |
| PBANKA_1019310 | 0 | 0 | 5 | 0 | 11 | 0 | 13 | 7 | 0 | 0 | 7 | 0 | 19 | 14 | 17 | 23 | 9 | 7 | 13 | 6 | 0 | 25 |
| PBANKA_1019330 | 0 | 0 | 5 | 0 | 12 | 0 | 28 | 0 | 8 | 0 | 13 | 5 | 60 | 7 | 63 | 28 | 19 | 10 | 403 | 22 | 13 | 127 |
| PBANKA_1019400 | 147 | 199 | 2699 | 604 | 1826 | 888 | 2048 | 727 | 627 | 334 | 2548 | 1927 | 25278 | 11871 | 4269 | 14955 | 6305 | 1283 | 3946 | 2372 | 526 | 4410 |
| PBANKA_1019500 | 198 | 224 | 2568 | 836 | 1597 | 1116 | 1597 | 823 | 555 | 398 | 6453 | 6234 | 15571 | 6786 | 3487 | 11142 | 3410 | 721 | 5072 | 2280 | 990 | 9116 |
| PBANKA_1019600 | 35 | 56 | 91 | 19 | 403 | 126 | 436 | 151 | 306 | 110 | 288 | 175 | 2539 | 1455 | 873 | 5037 | 1645 | 310 | 2167 | 1943 | 1711 | 10327 |
| PBANKA_1019700 | 220 | 371 | 602 | 137 | 929 | 376 | 1063 | 368 | 570 | 249 | 5140 | 3027 | 6147 | 7354 | 2952 | 18522 | 5990 | 630 | 1749 | 4066 | 1092 | 7936 |
| PBANKA_1019800 | 23 | 17 | 22 | 5 | 116 | 42 | 94 | 29 | 37 | 32 | 19 | 0 | 302 | 178 | 54 | 283 | 240 | 121 | 6102 | 4021 | 119 | 579 |
| PBANKA_1019900 | 23 | 20 | 14 | 0 | 77 | 31 | 75 | 26 | 104 | 57 | 54 | 8 | 211 | 182 | 19 | 119 | 3256 | 676 | 319 | 743 | 203 | 1336 |
| PBANKA_1020100 | 1335 | 1922 | 90 | 30 | 498 | 224 | 1000 | 509 | 735 | 364 | 73 | 36 | 1553 | 386 | 351 | 777 | 8306 | 1923 | 1616 | 1002 | 1801 | 12186 |

| | | | | | | | | | | | | | | | | | | | | | | |
|---|---|---|---|---|---|---|---|---|---|---|---|---|---|---|---|---|---|---|---|---|---|---|
| PBANKA_1020200 | 190 | 341 | 107 | 21 | 239 | 126 | 190 | 70 | 75 | 27 | 33 | 0 | 958 | 338 | 392 | 2381 | 1330 | 195 | 626 | 477 | 75 | 464 |
| PBANKA_1020400 | 13 | 28 | 84 | 11 | 128 | 54 | 104 | 53 | 116 | 51 | 1045 | 683 | 1203 | 1331 | 249 | 1331 | 833 | 164 | 530 | 586 | 231 | 1631 |
| PBANKA_1020600 | 61 | 85 | 40 | 13 | 120 | 74 | 99 | 45 | 40 | 22 | 0 | 0 | 420 | 106 | 334 | 1579 | 387 | 54 | 316 | 250 | 2770 | 17989 |
| PBANKA_1020700 | 231 | 319 | 202 | 49 | 373 | 106 | 339 | 124 | 187 | 82 | 64 | 25 | 5622 | 3174 | 1416 | 8326 | 2275 | 292 | 1591 | 2186 | 1178 | 6300 |
| PBANKA_1020800 | 26 | 54 | 20 | 0 | 109 | 37 | 173 | 57 | 96 | 49 | 27 | 12 | 418 | 162 | 89 | 362 | 1573 | 201 | 4265 | 4556 | 1391 | 7757 |
| PBANKA_1020900 | 39 | 45 | 93 | 8 | 183 | 95 | 299 | 135 | 329 | 128 | 62 | 26 | 1864 | 604 | 289 | 1340 | 3370 | 789 | 1869 | 934 | 317 | 1983 |
| PBANKA_1021000 | 10 | 17 | 140 | 19 | 176 | 58 | 160 | 78 | 107 | 38 | 186 | 73 | 2315 | 2283 | 641 | 3474 | 810 | 130 | 1282 | 1392 | 1259 | 7005 |
| PBANKA_1021100 | 12 | 8 | 24 | 7 | 49 | 15 | 42 | 18 | 37 | 22 | 320 | 186 | 693 | 425 | 94 | 488 | 383 | 103 | 3588 | 1425 | 940 | 5313 |
| PBANKA_1021300 | 0 | 0 | 62 | 14 | 96 | 43 | 88 | 29 | 111 | 36 | 12 | 0 | 882 | 140 | 262 | 1269 | 722 | 154 | 542 | 516 | 94 | 600 |
| PBANKA_1021400 | 0 | 0 | 15 | 8 | 74 | 37 | 119 | 63 | 51 | 24 | 5 | 0 | 409 | 57 | 359 | 466 | 868 | 366 | 233 | 86 | 22 | 109 |
| PBANKA_1021500 | 0 | 0 | 36 | 8 | 124 | 47 | 207 | 101 | 84 | 47 | 12 | 0 | 775 | 95 | 557 | 1303 | 2016 | 604 | 517 | 183 | 81 | 401 |
| PBANKA_1021600 | 17 | 11 | 62 | 14 | 229 | 65 | 263 | 116 | 87 | 32 | 15 | 0 | 421 | 121 | 86 | 331 | 765 | 91 | 344 | 256 | 280 | 1573 |
| PBANKA_1021700 | 24 | 40 | 113 | 17 | 245 | 81 | 231 | 65 | 82 | 24 | 17 | 9 | 2071 | 1336 | 529 | 2081 | 590 | 123 | 2061 | 1243 | 308 | 1804 |
| PBANKA_1021800 | 8 | 17 | 43 | 9 | 62 | 29 | 76 | 28 | 11 | 8 | 0 | 0 | 142 | 92 | 37 | 213 | 44 | 7 | 84 | 58 | 20 | 143 |
| PBANKA_1021900 | 17 | 42 | 54 | 11 | 83 | 26 | 52 | 14 | 30 | 0 | 7 | 0 | 784 | 294 | 137 | 683 | 202 | 32 | 347 | 200 | 96 | 463 |
| PBANKA_1022000 | 50 | 70 | 1009 | 328 | 692 | 452 | 788 | 371 | 234 | 224 | 1374 | 1138 | 8977 | 2175 | 1863 | 5929 | 2047 | 382 | 2382 | 864 | 109 | 733 |
| PBANKA_1022200 | 0 | 0 | 34 | 8 | 107 | 57 | 220 | 110 | 78 | 77 | 87 | 49 | 1125 | 528 | 253 | 956 | 787 | 196 | 1424 | 503 | 16 | 93 |
| PBANKA_1022300 | 0 | 0 | 23 | 0 | 204 | 81 | 342 | 128 | 181 | 109 | 39 | 17 | 456 | 84 | 86 | 170 | 1631 | 413 | 2719 | 1045 | 28 | 247 |
| PBANKA_1022500 | 2541 | 3137 | 313 | 90 | 569 | 252 | 470 | 176 | 312 | 145 | 229 | 151 | 1297 | 907 | 228 | 1177 | 655 | 225 | 1028 | 902 | 1035 | 7589 |
| PBANKA_1022600 | 45 | 38 | 27 | 7 | 115 | 47 | 202 | 49 | 207 | 99 | 87 | 36 | 363 | 211 | 69 | 298 | 413 | 132 | 1544 | 896 | 592 | 3757 |
| PBANKA_1022700 | 59 | 72 | 91 | 21 | 281 | 94 | 477 | 96 | 214 | 81 | 19 | 9 | 854 | 342 | 500 | 2000 | 1500 | 273 | 2643 | 2009 | 290 | 1849 |
| PBANKA_1022800 | 86 | 75 | 294 | 97 | 422 | 193 | 436 | 130 | 155 | 73 | 40 | 15 | 2465 | 1205 | 1129 | 5007 | 1631 | 270 | 1039 | 928 | 317 | 2268 |
| PBANKA_1022900 | 28 | 41 | 67 | 21 | 136 | 51 | 227 | 71 | 137 | 85 | 56 | 10 | 1151 | 391 | 185 | 930 | 1578 | 278 | 1140 | 746 | 165 | 861 |
| PBANKA_1023000 | 99 | 145 | 45 | 10 | 391 | 136 | 869 | 275 | 504 | 247 | 76 | 19 | 1001 | 212 | 652 | 2238 | 6230 | 1114 | 2123 | 1404 | 400 | 2561 |
| PBANKA_1023100 | 0 | 0 | 10 | 0 | 47 | 14 | 62 | 29 | 30 | 8 | 9 | 0 | 110 | 70 | 79 | 282 | 245 | 60 | 603 | 1903 | 6531 | 37418 |
| PBANKA_1023200 | 0 | 0 | 142 | 42 | 328 | 167 | 388 | 123 | 144 | 73 | 144 | 106 | 2424 | 800 | 574 | 2261 | 1017 | 202 | 1582 | 805 | 74 | 513 |
| PBANKA_1023300 | 11 | 12 | 12 | 0 | 95 | 30 | 153 | 66 | 71 | 36 | 23 | 10 | 884 | 354 | 300 | 1742 | 904 | 120 | 722 | 659 | 125 | 900 |
| PBANKA_1023400 | 70 | 90 | 229 | 58 | 240 | 174 | 264 | 143 | 107 | 57 | 36 | 30 | 2853 | 919 | 670 | 3330 | 1655 | 205 | 1146 | 623 | 802 | 6154 |
| PBANKA_1023500 | 0 | 0 | 366 | 90 | 522 | 233 | 526 | 221 | 173 | 101 | 557 | 428 | 8629 | 3937 | 2497 | 10796 | 2274 | 333 | 1415 | 1329 | 105 | 733 |
| PBANKA_1023600 | 22 | 24 | 104 | 25 | 443 | 160 | 625 | 277 | 282 | 160 | 168 | 151 | 1766 | 754 | 639 | 3584 | 3019 | 509 | 3412 | 4629 | 194 | 1097 |
| PBANKA_1023700 | 253 | 239 | 24 | 5 | 54 | 29 | 93 | 37 | 25 | 27 | 8 | 0 | 341 | 84 | 77 | 326 | 1034 | 167 | 1108 | 496 | 60 | 366 |
| PBANKA_1023800 | 119 | 154 | 154 | 21 | 528 | 197 | 995 | 366 | 553 | 244 | 780 | 451 | 3987 | 5100 | 1247 | 6910 | 7685 | 1067 | 1608 | 2848 | 214 | 1261 |
| PBANKA_1023900 | 0 | 0 | 22 | 0 | 82 | 44 | 125 | 51 | 24 | 19 | 11 | 0 | 285 | 87 | 233 | 904 | 193 | 52 | 1218 | 478 | 87 | 711 |
| PBANKA_1024000 | 0 | 0 | 152 | 27 | 526 | 184 | 882 | 317 | 252 | 112 | 19 | 18 | 1260 | 599 | 515 | 2242 | 1627 | 211 | 1125 | 831 | 55 | 371 |
| PBANKA_1024100 | 9 | 9 | 140 | 41 | 403 | 135 | 341 | 130 | 383 | 179 | 1987 | 1172 | 1994 | 2919 | 580 | 3247 | 1080 | 310 | 244 | 946 | 160 | 1102 |
| PBANKA_1024200 | 61 | 124 | 122 | 16 | 442 | 151 | 726 | 225 | 396 | 124 | 72 | 16 | 1105 | 514 | 243 | 1239 | 9126 | 1135 | 469 | 978 | 66 | 349 |
| PBANKA_1024300 | 71 | 119 | 136 | 26 | 123 | 46 | 126 | 38 | 178 | 85 | 2197 | 1522 | 1450 | 2696 | 481 | 3359 | 932 | 121 | 657 | 2895 | 494 | 4084 |
| PBANKA_1024311 | 13 | 12 | 29 | 15 | 58 | 11 | 49 | 17 | 42 | 7 | 40 | 6 | 143 | 249 | 2018 | 511 | 398 | 221 | 149 | 313 | 0 | 29 |
| PBANKA_1024351 | 5 | 0 | 9 | 0 | 5 | 0 | 20 | 0 | 13 | 0 | 17 | 11 | 21 | 0 | 1362 | 11 | 50 | 74 | 57 | 7 | 0 | 5 |
| PBANKA_1024400 | 25 | 34 | 83 | 22 | 278 | 165 | 328 | 147 | 135 | 69 | 17 | 13 | 124 | 26 | 87 | 293 | 329 | 57 | 1084 | 613 | 4657 | 35076 |
| PBANKA_1024500 | 0 | 5 | 61 | 26 | 219 | 102 | 281 | 111 | 78 | 39 | 12 | 5 | 124 | 23 | 100 | 342 | 301 | 48 | 26 | 36 | 16 | 159 |
| PBANKA_1024600 | 24 | 13 | 4283 | 961 | 12625 | 5467 | 21738 | 7884 | 12267 | 5260 | 1295 | 524 | 36 | 21 | 69 | 191 | 90 | 16 | 121 | 167 | 22 | 187 |
| PBANKA_1024700 | 158 | 206 | 76 | 23 | 260 | 230 | 335 | 206 | 110 | 54 | 8 | 16 | 1414 | 304 | 358 | 1719 | 1032 | 196 | 2580 | 1041 | 200 | 1179 |
| PBANKA_1024800 | 0 | 0 | 90 | 28 | 251 | 155 | 230 | 210 | 115 | 73 | 0 | 0 | 264 | 59 | 344 | 627 | 710 | 227 | 1152 | 434 | 151 | 735 |
| PBANKA_1024900 | 64 | 35 | 248 | 86 | 846 | 482 | 1346 | 621 | 509 | 354 | 70 | 34 | 1284 | 421 | 1288 | 4194 | 9682 | 1099 | 3487 | 3104 | 67 | 594 |
| PBANKA_1025000 | 7 | 17 | 78 | 16 | 244 | 157 | 404 | 197 | 201 | 87 | 28 | 8 | 930 | 169 | 308 | 1163 | 2663 | 409 | 1328 | 783 | 134 | 904 |
| PBANKA_1025100 | 425 | 566 | 424 | 141 | 1857 | 989 | 3480 | 1291 | 1393 | 669 | 322 | 193 | 1810 | 5157 | 3405 | 17436 | 14211 | 1397 | 1158 | 5490 | 835 | 6240 |
| PBANKA_1025200 | 19 | 30 | 222 | 60 | 277 | 226 | 351 | 143 | 110 | 64 | 706 | 688 | 2202 | 1336 | 550 | 2841 | 849 | 99 | 368 | 376 | 80 | 710 |
| PBANKA_1025300 | 276 | 504 | 7 | 0 | 40 | 21 | 265 | 126 | 782 | 433 | 295 | 116 | 165 | 316 | 44 | 69 | 2879 | 689 | 382 | 3207 | 157 | 696 |
| PBANKA_1025400 | 15 | 15 | 18 | 6 | 155 | 59 | 322 | 147 | 228 | 128 | 34 | 13 | 462 | 182 | 135 | 383 | 2960 | 542 | 1377 | 1301 | 28 | 217 |
| PBANKA_1025500 | 63 | 90 | 765 | 215 | 1749 | 836 | 2907 | 1092 | 1848 | 852 | 1132 | 618 | 5595 | 13257 | 5182 | 27289 | 27314 | 3315 | 2591 | 9284 | 500 | 3843 |
| PBANKA_1025600 | 0 | 0 | 27 | 5 | 76 | 32 | 80 | 19 | 11 | 16 | 0 | 0 | 115 | 26 | 168 | 508 | 219 | 50 | 292 | 162 | 23 | 138 |
| PBANKA_1025800 | 8 | 0 | 106 | 40 | 144 | 134 | 116 | 54 | 29 | 25 | 0 | 5 | 466 | 176 | 302 | 759 | 170 | 24 | 241 | 134 | 68 | 444 |
| PBANKA_1025900 | 0 | 0 | 30 | 6 | 114 | 62 | 145 | 61 | 34 | 21 | 11 | 0 | 345 | 201 | 282 | 1116 | 1022 | 226 | 838 | 924 | 37 | 250 |
| PBANKA_1026200 | 174 | 232 | 275 | 74 | 1958 | 1236 | 2174 | 713 | 879 | 504 | 2589 | 1726 | 3718 | 2220 | 1059 | 4070 | 544 | 142 | 1104 | 818 | 619 | 5200 |
| PBANKA_1026300 | 10 | 0 | 19 | 0 | 393 | 116 | 459 | 192 | 190 | 72 | 52 | 8 | 250 | 141 | 101 | 254 | 298 | 127 | 1810 | 2965 | 341 | 2114 |
| PBANKA_1026400 | 5 | 5 | 70 | 13 | 151 | 48 | 167 | 52 | 73 | 39 | 10 | 5 | 500 | 131 | 377 | 1859 | 705 | 118 | 607 | 375 | 24 | 201 |
| PBANKA_1026500 | 19 | 30 | 18 | 0 | 146 | 61 | 314 | 103 | 265 | 99 | 49 | 13 | 396 | 257 | 167 | 611 | 3249 | 499 | 1720 | 4130 | 874 | 6675 |
| PBANKA_1026600 | 59 | 137 | 66 | 30 | 426 | 155 | 464 | 195 | 188 | 99 | 25 | 32 | 1537 | 814 | 641 | 3463 | 2834 | 439 | 1435 | 1630 | 69 | 594 |
| PBANKA_1026700 | 24 | 36 | 6 | 0 | 17 | 15 | 54 | 35 | 55 | 34 | 9 | 5 | 189 | 26 | 30 | 160 | 190 | 26 | 1147 | 393 | 45 | 231 |
| PBANKA_1026800 | 139 | 130 | 34 | 6 | 94 | 72 | 182 | 96 | 54 | 46 | 19 | 7 | 428 | 104 | 126 | 471 | 581 | 185 | 2499 | 1460 | 81 | 623 |
| PBANKA_1026900 | 65 | 98 | 256 | 58 | 214 | 103 | 269 | 113 | 394 | 185 | 1041 | 720 | 2147 | 1621 | 327 | 1435 | 1636 | 335 | 1038 | 703 | 337 | 2159 |
| PBANKA_1027000 | 44 | 42 | 61 | 34 | 235 | 137 | 371 | 196 | 529 | 264 | 284 | 191 | 1190 | 421 | 475 | 2324 | 3060 | 587 | 19472 | 10728 | 400 | 3801 |
| PBANKA_1027100 | 0 | 0 | 148 | 23 | 347 | 114 | 602 | 232 | 277 | 119 | 106 | 47 | 629 | 333 | 424 | 1861 | 3250 | 385 | 1476 | 1548 | 109 | 786 |
| PBANKA_1027200 | 75 | 62 | 31 | 9 | 125 | 35 | 117 | 45 | 93 | 51 | 7 | 0 | 287 | 83 | 287 | 1018 | 853 | 178 | 1477 | 821 | 148 | 898 |
| PBANKA_1027300 | 9 | 14 | 273 | 58 | 504 | 197 | 785 | 255 | 392 | 121 | 40 | 14 | 1411 | 1114 | 2176 | 13244 | 13690 | 1434 | 4883 | 5308 | 409 | 2357 |
| PBANKA_1027400 | 0 | 0 | 40 | 0 | 82 | 23 | 47 | 31 | 24 | 13 | 9 | 0 | 66 | 25 | 58 | 215 | 40 | 20 | 123 | 89 | 18 | 161 |
| PBANKA_1027500 | 35 | 18 | 68 | 14 | 49 | 38 | 56 | 34 | 38 | 21 | 175 | 177 | 106 | 78 | 94 | 94 | 22 | 15 | 78 | 30 | 12 | 28 |
| PBANKA_1027600 | 9 | 6 | 5 | 0 | 24 | 5 | 16 | 9 | 73 | 46 | 36 | 11 | 108 | 28 | 12 | 24 | 329 | 120 | 942 | 352 | 1125 | 8079 |
| PBANKA_1027700 | 26 | 44 | 5 | 0 | 49 | 20 | 69 | 31 | 78 | 25 | 23 | 7 | 368 | 73 | 26 | 63 | 1051 | 219 | 3717 | 2070 | 501 | 3159 |
| PBANKA_1027800 | 27 | 46 | 135 | 23 | 158 | 60 | 183 | 50 | 135 | 59 | 79 | 8 | 1049 | 511 | 450 | 2304 | 1204 | 223 | 883 | 844 | 477 | 2526 |
| PBANKA_1027900 | 6 | 0 | 105 | 29 | 155 | 97 | 198 | 103 | 84 | 50 | 63 | 58 | 890 | 397 | 446 | 1454 | 578 | 89 | 396 | 315 | 501 | 3093 |
| PBANKA_1028000 | 59 | 118 | 119 | 20 | 181 | 53 | 210 | 56 | 265 | 117 | 120 | 57 | 2402 | 2012 | 1250 | 7252 | 6847 | 1415 | 3531 | 4824 | 307 | 1329 |
| PBANKA_1028100 | 148 | 158 | 57 | 0 | 106 | 45 | 175 | 79 | 163 | 60 | 37 | 25 | 1213 | 668 | 286 | 1713 | 1720 | 342 | 1187 | 778 | 260 | 1638 |
| PBANKA_1028200 | 786 | 1074 | 31 | 0 | 233 | 91 | 375 | 169 | 489 | 264 | 134 | 62 | 2126 | 1782 | 627 | 3371 | 11618 | 2258 | 14726 | 37445 | 2021 | 11464 |
| PBANKA_1028300 | 138 | 138 | 153 | 45 | 361 | 151 | 413 | 226 | 213 | 110 | 26 | 21 | 1427 | 493 | 697 | 2687 | 2560 | 385 | 2035 | 1162 | 226 | 1873 |
| PBANKA_1028400 | 105 | 98 | 2086 | 630 | 1463 | 966 | 1601 | 819 | 470 | 292 | 1966 | 1615 | 14485 | 3968 | 3926 | 9876 | 3742 | 814 | 4086 | 1743 | 716 | 6740 |
| PBANKA_1028500 | 306 | 436 | 141 | 25 | 239 | 116 | 274 | 77 | 245 | 112 | 1112 | 591 | 2362 | 1267 | 421 | 2497 | 1683 | 297 | 1713 | 983 | 644 | 4434 |
| PBANKA_1028600 | 25 | 16 | 12 | 0 | 26 | 16 | 125 | 63 | 96 | 44 | 13 | 0 | 875 | 307 | 171 | 1090 | 1234 | 219 | 2019 | 1496 | 689 | 4176 |
| PBANKA_1028700 | 0 | 0 | 20 | 0 | 115 | 37 | 242 | 140 | 114 | 72 | 18 | 7 | 572 | 200 | 214 | 525 | 736 | 147 | 4269 | 2270 | 851 | 2756 |
| PBANKA_1028800 | 13 | 18 | 47 | 11 | 212 | 109 | 270 | 111 | 98 | 57 | 16 | 0 | 728 | 274 | 406 | 1358 | 1865 | 387 | 642 | 352 | 95 | 584 |
| PBANKA_1028900 | 6 | 0 | 70 | 23 | 152 | 45 | 90 | 45 | 113 | 81 | 161 | 74 | 839 | 386 | 147 | 634 | 229 | 90 | 400 | 212 | 667 | 3852 |
| PBANKA_1029000 | 17 | 18 | 13 | 0 | 234 | 96 | 344 | 109 | 465 | 276 | 525 | 304 | 830 | 855 | 225 | 984 | 3364 | 1404 | 409 | 3100 | 118 | 730 |
| PBANKA_1029100 | 10 | 14 | 9 | 0 | 40 | 0 | 30 | 11 | 30 | 9 | 22 | 9 | 77 | 23 | 22 | 114 | 158 | 56 | 75 | 73 | 23 | 112 |
| PBANKA_1029200 | 30 | 43 | 22 | 5 | 113 | 47 | 217 | 81 | 58 | 30 | 10 | 0 | 270 | 38 | 98 | 307 | 556 | 113 | 236 | 163 | 133 | 709 |
| PBANKA_1029300 | 7 | 9 | 78 | 14 | 218 | 70 | 292 | 107 | 96 | 47 | 23 | 6 | 1058 | 382 | 973 | 5581 | 2934 | 377 | 683 | 1430 | 165 | 650 |
| PBANKA_1029400 | 0 | 0 | 5 | 0 | 28 | 7 | 27 | 9 | 65 | 37 | 19 | 15 | 3975 | 996 | 56 | 324 | 3749 | 1162 | 21714 | 20642 | 392 | 1772 |
| PBANKA_1029700 | 71 | 82 | 65 | 18 | 97 | 68 | 155 | 56 | 44 | 17 | 16 | 15 | 1171 | 429 | 308 | 1435 | 303 | 64 | 2384 | 859 | 118 | 775 |
| PBANKA_1029800 | 20 | 22 | 70 | 20 | 141 | 109 | 175 | 87 | 28 | 28 | 6 | 6 | 504 | 192 | 398 | 1319 | 418 | 68 | 610 | 414 | 42 | 329 |
| PBANKA_1029900 | 0 | 0 | 128 | 32 | 368 | 145 | 231 | 76 | 90 | 26 | 101 | 54 | 1177 | 681 | 999 | 4179 | 948 | 108 | 276 | 168 | 0 | 26 |

| | | | | | | | | | | | | | | | | | | | | | |
|---|---|---|---|---|---|---|---|---|---|---|---|---|---|---|---|---|---|---|---|---|---|
| PBANKA_1030000 | 0 | 0 | 136 | 49 | 203 | 97 | 136 | 58 | 52 | 29 | 336 | 252 | 1147 | 701 | 397 | 1673 | 237 | 35 | 87 | 81 | 0 | 7 |
| PBANKA_1030100 | 0 | 0 | 196 | 68 | 146 | 95 | 95 | 48 | 78 | 33 | 10 | 5 | 396 | 81 | 459 | 1929 | 704 | 174 | 4123 | 1891 | 17 | 189 |
| PBANKA_1030200 | 18 | 39 | 35 | 11 | 231 | 116 | 325 | 117 | 189 | 62 | 230 | 89 | 1154 | 1865 | 781 | 4221 | 2519 | 352 | 692 | 1318 | 276 | 2174 |
| PBANKA_1030300 | 35 | 51 | 776 | 260 | 672 | 366 | 712 | 318 | 151 | 144 | 107 | 94 | 2982 | 824 | 1736 | 5241 | 1798 | 379 | 1071 | 341 | 49 | 488 |
| PBANKA_1030500 | 10 | 11 | 149 | 17 | 454 | 146 | 456 | 123 | 136 | 83 | 63 | 15 | 330 | 295 | 696 | 2637 | 2108 | 229 | 275 | 271 | 0 | 38 |
| PBANKA_1030600 | 0 | 0 | 7 | 0 | 26 | 13 | 63 | 31 | 78 | 52 | 913 | 728 | 5612 | 2745 | 1862 | 5283 | 7523 | 1324 | 5465 | 1733 | 68 | 719 |
| PBANKA_1030700 | 0 | 0 | 41 | 9 | 101 | 33 | 84 | 33 | 48 | 23 | 498 | 343 | 623 | 558 | 151 | 783 | 1160 | 141 | 260 | 188 | 23 | 107 |
| PBANKA_1030800 | 0 | 0 | 12 | 6 | 29 | 17 | 39 | 18 | 41 | 21 | 121 | 95 | 384 | 267 | 114 | 508 | 50 | 13 | 94 | 69 | 0 | 30 |
| PBANKA_1030900 | 0 | 0 | 40 | 10 | 68 | 31 | 64 | 21 | 27 | 12 | 6 | 11 | 443 | 233 | 144 | 716 | 405 | 52 | 132 | 111 | 0 | 31 |
| PBANKA_1031000 | 10 | 15 | 78 | 18 | 260 | 86 | 140 | 53 | 105 | 52 | 857 | 589 | 621 | 704 | 279 | 1950 | 427 | 43 | 593 | 772 | 17 | 80 |
| PBANKA_1031100 | 119 | 202 | 146 | 37 | 743 | 391 | 512 | 158 | 232 | 110 | 361 | 240 | 268 | 227 | 997 | 4936 | 1603 | 191 | 306 | 758 | 13 | 89 |
| PBANKA_1031200 | 0 | 0 | 68 | 17 | 194 | 86 | 358 | 138 | 187 | 68 | 33 | 13 | 1108 | 475 | 299 | 1750 | 2380 | 431 | 1381 | 832 | 127 | 812 |
| PBANKA_1031300 | 7 | 11 | 50 | 8 | 80 | 40 | 172 | 61 | 140 | 69 | 46 | 23 | 1947 | 813 | 344 | 1856 | 3845 | 884 | 2681 | 1602 | 56 | 422 |
| PBANKA_1031400 | 5 | 9 | 40 | 6 | 151 | 91 | 231 | 114 | 91 | 36 | 16 | 8 | 388 | 128 | 351 | 990 | 614 | 167 | 562 | 316 | 11 | 233 |
| PBANKA_1031600 | 144 | 173 | 87 | 32 | 187 | 81 | 201 | 92 | 212 | 115 | 69 | 32 | 743 | 364 | 332 | 1750 | 1519 | 337 | 1094 | 1617 | 587 | 3812 |
| PBANKA_1031800 | 0 | 0 | 40 | 12 | 172 | 122 | 260 | 148 | 115 | 68 | 15 | 11 | 1118 | 194 | 314 | 1558 | 2623 | 377 | 571 | 457 | 101 | 563 |
| PBANKA_1031900 | 27 | 19 | 480 | 172 | 566 | 373 | 541 | 243 | 209 | 151 | 308 | 211 | 7569 | 3956 | 2117 | 8324 | 2408 | 401 | 1533 | 1137 | 60 | 631 |
| PBANKA_1032000 | 0 | 5 | 33 | 22 | 97 | 53 | 103 | 46 | 22 | 20 | 0 | 0 | 262 | 86 | 122 | 467 | 158 | 26 | 165 | 99 | 0 | 41 |
| PBANKA_1032100 | 0 | 0 | 10 | 6 | 120 | 54 | 1340 | 507 | 7353 | 4503 | 896 | 645 | 3173 | 1431 | 80 | 252 | 43506 | 16006 | 912 | 900 | 64 | 639 |
| PBANKA_1032200 | 0 | 5 | 546 | 103 | 992 | 432 | 951 | 317 | 573 | 230 | 222 | 118 | 3189 | 6637 | 4501 | 22670 | 4737 | 538 | 805 | 5579 | 65 | 614 |
| PBANKA_1032300 | 5 | 9 | 213 | 63 | 230 | 101 | 117 | 43 | 109 | 45 | 970 | 629 | 3631 | 2028 | 647 | 4121 | 395 | 71 | 2878 | 2287 | 130 | 1245 |
| PBANKA_1032500 | 9 | 8 | 66 | 16 | 181 | 78 | 182 | 84 | 81 | 30 | 24 | 14 | 1016 | 365 | 332 | 1500 | 969 | 146 | 1433 | 660 | 46 | 283 |
| PBANKA_1032600 | 149 | 160 | 113 | 42 | 352 | 212 | 438 | 203 | 214 | 107 | 42 | 28 | 2886 | 825 | 727 | 4882 | 4937 | 584 | 1853 | 1132 | 197 | 1718 |
| PBANKA_1032700 | 48 | 44 | 474 | 126 | 556 | 293 | 592 | 198 | 108 | 69 | 279 | 215 | 4047 | 2275 | 1615 | 8125 | 1633 | 229 | 2276 | 1810 | 642 | 5794 |
| PBANKA_1032800 | 0 | 11 | 31 | 6 | 114 | 57 | 149 | 61 | 58 | 25 | 13 | 0 | 462 | 215 | 237 | 839 | 542 | 92 | 1405 | 779 | 74 | 621 |
| PBANKA_1032900 | 0 | 0 | 73 | 17 | 547 | 239 | 725 | 289 | 501 | 297 | 515 | 210 | 1956 | 956 | 486 | 1696 | 3941 | 835 | 667 | 422 | 76 | 556 |
| PBANKA_1033000 | 113 | 155 | 184 | 34 | 253 | 100 | 328 | 132 | 109 | 75 | 26 | 14 | 248 | 117 | 465 | 2219 | 736 | 127 | 1099 | 946 | 286 | 1905 |
| PBANKA_1033100 | 54 | 67 | 59 | 16 | 257 | 126 | 680 | 320 | 1507 | 850 | 194 | 128 | 584 | 169 | 121 | 235 | 4327 | 1466 | 1486 | 752 | 72 | 726 |
| PBANKA_1033200 | 5 | 9 | 55 | 22 | 294 | 172 | 480 | 217 | 165 | 110 | 27 | 15 | 423 | 115 | 435 | 1463 | 2913 | 390 | 455 | 397 | 12 | 164 |
| PBANKA_1033300 | 0 | 0 | 114 | 34 | 243 | 136 | 266 | 167 | 203 | 135 | 11 | 6 | 379 | 66 | 108 | 320 | 2152 | 388 | 929 | 678 | 64 | 665 |
| PBANKA_1033400 | 32 | 62 | 90 | 32 | 426 | 146 | 509 | 172 | 205 | 80 | 27 | 21 | 2175 | 1195 | 1453 | 8326 | 7850 | 797 | 655 | 1883 | 129 | 804 |
| PBANKA_1033500 | 0 | 0 | 56 | 6 | 89 | 37 | 89 | 35 | 40 | 11 | 80 | 31 | 1201 | 849 | 134 | 747 | 351 | 58 | 193 | 162 | 24 | 129 |
| PBANKA_1033600 | 0 | 0 | 70 | 14 | 135 | 83 | 80 | 44 | 24 | 18 | 415 | 347 | 516 | 306 | 107 | 433 | 97 | 19 | 632 | 678 | 5 | 57 |
| PBANKA_1033800 | 94 | 141 | 713 | 173 | 2050 | 829 | 2474 | 872 | 928 | 374 | 207 | 88 | 1758 | 2425 | 5602 | 26356 | 18180 | 1767 | 1362 | 9080 | 327 | 2580 |
| PBANKA_1033900 | 367 | 437 | 2460 | 866 | 1416 | 1147 | 1350 | 872 | 534 | 418 | 2427 | 2240 | 28264 | 11611 | 4811 | 24469 | 7378 | 1050 | 4899 | 2976 | 402 | 3223 |
| PBANKA_1034000 | 18 | 25 | 73 | 8 | 125 | 47 | 104 | 42 | 35 | 17 | 130 | 80 | 1790 | 990 | 209 | 1089 | 518 | 101 | 2681 | 1666 | 249 | 1550 |
| PBANKA_1034100 | 12 | 0 | 93 | 29 | 319 | 114 | 401 | 157 | 192 | 89 | 21 | 6 | 1715 | 1117 | 696 | 4435 | 3148 | 406 | 4580 | 3831 | 79 | 676 |
| PBANKA_1034300 | 13 | 13 | 103 | 21 | 414 | 175 | 384 | 130 | 359 | 182 | 4173 | 2270 | 1773 | 2799 | 793 | 5059 | 3062 | 385 | 3051 | 5435 | 17 | 148 |
| PBANKA_1034400 | 0 | 0 | 140 | 39 | 1396 | 761 | 2160 | 1055 | 742 | 474 | 4810 | 4005 | 27343 | 8551 | 9645 | 30680 | 8049 | 1690 | 4141 | 2356 | 171 | 2105 |
| PBANKA_1034600 | 33 | 45 | 97 | 17 | 146 | 53 | 161 | 83 | 144 | 88 | 34 | 17 | 609 | 207 | 160 | 273 | 1101 | 323 | 2357 | 1072 | 97 | 689 |
| PBANKA_1034700 | 57 | 68 | 94 | 12 | 80 | 48 | 138 | 45 | 46 | 48 | 20 | 11 | 942 | 311 | 174 | 646 | 426 | 128 | 2488 | 1159 | 158 | 791 |
| PBANKA_1034800 | 39 | 51 | 72 | 5 | 117 | 30 | 97 | 49 | 66 | 16 | 5 | 0 | 666 | 147 | 142 | 666 | 898 | 153 | 2370 | 1108 | 111 | 758 |
| PBANKA_1034900 | 98 | 100 | 238 | 42 | 556 | 293 | 575 | 235 | 573 | 285 | 709 | 502 | 7251 | 3971 | 1497 | 8943 | 4874 | 715 | 2132 | 1956 | 353 | 2380 |
| PBANKA_1035000 | 64 | 84 | 21 | 0 | 50 | 14 | 81 | 23 | 110 | 62 | 64 | 38 | 1217 | 426 | 162 | 987 | 1042 | 220 | 1367 | 778 | 198 | 1311 |
| PBANKA_1035100 | 23 | 24 | 237 | 91 | 213 | 93 | 177 | 55 | 91 | 57 | 1846 | 775 | 12493 | 3241 | 937 | 6585 | 1550 | 229 | 1551 | 801 | 330 | 2535 |
| PBANKA_1035300 | 0 | 0 | 87 | 16 | 241 | 160 | 258 | 99 | 69 | 54 | 29 | 5 | 632 | 381 | 421 | 1756 | 462 | 64 | 652 | 546 | 103 | 786 |
| PBANKA_1035400 | 0 | 0 | 24 | 12 | 141 | 96 | 858 | 470 | 1661 | 1138 | 228 | 170 | 3315 | 736 | 132 | 517 | 12822 | 4085 | 56696 | 35472 | 881 | 7645 |
| PBANKA_1035500 | 0 | 0 | 49 | 19 | 204 | 128 | 214 | 97 | 79 | 38 | 15 | 0 | 133 | 45 | 308 | 932 | 500 | 132 | 515 | 217 | 131 | 1068 |
| PBANKA_1035600 | 0 | 6 | 288 | 41 | 333 | 107 | 225 | 77 | 95 | 29 | 106 | 44 | 1220 | 1331 | 973 | 4615 | 1307 | 197 | 611 | 861 | 23 | 147 |
| PBANKA_1035700 | 0 | 0 | 9 | 0 | 49 | 23 | 44 | 22 | 43 | 11 | 15 | 0 | 652 | 282 | 123 | 773 | 689 | 129 | 1476 | 3229 | 20 | 153 |
| PBANKA_1035800 | 5 | 15 | 42 | 9 | 210 | 72 | 565 | 213 | 433 | 183 | 48 | 18 | 389 | 121 | 347 | 1393 | 1436 | 273 | 1332 | 1117 | 344 | 2139 |
| PBANKA_1035900 | 83 | 138 | 565 | 86 | 604 | 246 | 586 | 174 | 305 | 172 | 2400 | 1610 | 2740 | 1531 | 437 | 1822 | 1394 | 255 | 5137 | 3184 | 225 | 1869 |
| PBANKA_1036000 | 29 | 38 | 652 | 118 | 1710 | 588 | 1933 | 712 | 979 | 433 | 4571 | 2625 | 7116 | 13669 | 6688 | 30300 | 16525 | 2140 | 2394 | 7878 | 203 | 1527 |
| PBANKA_1036100 | 14 | 17 | 38 | 0 | 138 | 50 | 178 | 63 | 75 | 37 | 49 | 46 | 410 | 348 | 178 | 984 | 624 | 96 | 525 | 550 | 67 | 413 |
| PBANKA_1036200 | 0 | 0 | 30 | 17 | 67 | 38 | 93 | 51 | 24 | 27 | 0 | 0 | 222 | 64 | 171 | 338 | 128 | 54 | 630 | 236 | 81 | 548 |
| PBANKA_1036300 | 14 | 15 | 91 | 25 | 132 | 54 | 108 | 41 | 200 | 128 | 1670 | 849 | 4708 | 4953 | 661 | 4062 | 1102 | 308 | 1873 | 3334 | 776 | 3710 |
| PBANKA_1036400 | 71 | 122 | 169 | 45 | 413 | 164 | 330 | 124 | 226 | 99 | 1483 | 1028 | 3397 | 3440 | 465 | 2858 | 804 | 129 | 1136 | 1676 | 2238 | 15642 |
| PBANKA_1036500 | 23 | 15 | 29 | 7 | 95 | 12 | 135 | 26 | 223 | 97 | 120 | 80 | 1057 | 455 | 256 | 1394 | 1409 | 286 | 2877 | 1470 | 147 | 934 |
| PBANKA_1036600 | 42 | 51 | 83 | 22 | 387 | 222 | 813 | 411 | 277 | 194 | 35 | 7 | 1759 | 215 | 1090 | 3380 | 8544 | 1489 | 3126 | 1835 | 99 | 994 |
| PBANKA_1036700 | 0 | 0 | 88 | 23 | 112 | 48 | 43 | 13 | 23 | 0 | 0 | 0 | 374 | 58 | 232 | 960 | 164 | 40 | 581 | 343 | 214 | 1423 |
| PBANKA_1036800 | 15 | 16 | 46 | 11 | 94 | 23 | 120 | 32 | 59 | 21 | 5 | 0 | 359 | 75 | 413 | 2112 | 615 | 103 | 503 | 907 | 152 | 971 |
| PBANKA_1036900 | 147 | 216 | 96 | 20 | 286 | 122 | 481 | 148 | 210 | 118 | 47 | 16 | 1194 | 726 | 672 | 3120 | 2640 | 437 | 876 | 872 | 540 | 3520 |
| PBANKA_1037000 | 136 | 184 | 44 | 13 | 177 | 108 | 411 | 143 | 102 | 88 | 20 | 12 | 222 | 147 | 126 | 562 | 639 | 89 | 174 | 171 | 94 | 606 |
| PBANKA_1037100 | 71 | 108 | 53 | 20 | 193 | 88 | 260 | 85 | 130 | 49 | 53 | 38 | 2634 | 1643 | 536 | 3360 | 1417 | 149 | 1752 | 1208 | 229 | 1108 |
| PBANKA_1037200 | 24 | 19 | 26 | 6 | 107 | 46 | 190 | 60 | 80 | 46 | 167 | 77 | 848 | 429 | 288 | 965 | 748 | 129 | 947 | 689 | 772 | 5336 |
| PBANKA_1037300 | 901 | 1331 | 43 | 14 | 84 | 38 | 141 | 71 | 105 | 77 | 641 | 541 | 3569 | 2150 | 550 | 2392 | 1931 | 378 | 2699 | 1604 | 274 | 2049 |
| PBANKA_1037400 | 1314 | 1982 | 18 | 5 | 51 | 33 | 52 | 28 | 264 | 156 | 593 | 497 | 593 | 408 | 24 | 109 | 548 | 343 | 614 | 288 | 188 | 1195 |
| PBANKA_1037500 | 10 | 22 | 25 | 11 | 314 | 144 | 347 | 135 | 129 | 67 | 50 | 21 | 351 | 160 | 75 | 247 | 671 | 146 | 1807 | 1440 | 23 | 182 |
| PBANKA_1037600 | 21 | 32 | 149 | 20 | 413 | 153 | 536 | 223 | 174 | 87 | 616 | 305 | 3248 | 2054 | 463 | 3003 | 592 | 115 | 894 | 888 | 130 | 892 |
| PBANKA_1037700 | 0 | 0 | 22 | 0 | 48 | 47 | 48 | 18 | 18 | 7 | 0 | 5 | 117 | 20 | 106 | 279 | 122 | 26 | 123 | 62 | 69 | 378 |
| PBANKA_1038000 | 141 | 154 | 227 | 59 | 316 | 181 | 482 | 186 | 271 | 180 | 45 | 19 | 415 | 113 | 309 | 816 | 1383 | 438 | 210 | 87 | 38 | 376 |
| PBANKA_1038100 | 0 | 0 | 70 | 7 | 190 | 74 | 132 | 47 | 37 | 9 | 0 | 0 | 328 | 53 | 322 | 1277 | 259 | 36 | 106 | 42 | 0 | 10 |
| PBANKA_1038200 | 0 | 0 | 7 | 0 | 10 | 7 | 25 | 10 | 35 | 15 | 0 | 0 | 287 | 207 | 126 | 377 | 558 | 128 | 1740 | 4954 | 23 | 184 |
| PBANKA_1038300 | 0 | 0 | 57 | 7 | 188 | 99 | 341 | 154 | 114 | 38 | 28 | 13 | 1645 | 670 | 578 | 2063 | 2535 | 421 | 1452 | 879 | 148 | 1256 |
| PBANKA_1038400 | 0 | 0 | 19 | 7 | 48 | 25 | 89 | 28 | 26 | 18 | 0 | 0 | 301 | 52 | 91 | 273 | 430 | 100 | 3739 | 1496 | 42 | 380 |
| PBANKA_1038600 | 0 | 0 | 6 | 5 | 34 | 14 | 50 | 22 | 16 | 5 | 0 | 0 | 220 | 53 | 19 | 87 | 185 | 54 | 1244 | 907 | 15 | 87 |
| PBANKA_1038900 | 57 | 109 | 169 | 41 | 856 | 311 | 1334 | 546 | 733 | 301 | 107 | 52 | 1166 | 760 | 1055 | 5825 | 13437 | 1343 | 1967 | 7035 | 555 | 4420 |
| PBANKA_1039000 | 6 | 5 | 22 | 0 | 37 | 14 | 86 | 42 | 88 | 37 | 5 | 0 | 368 | 108 | 46 | 226 | 195 | 52 | 3514 | 1920 | 24 | 206 |
| PBANKA_1039100 | 13 | 23 | 26 | 0 | 97 | 34 | 137 | 54 | 58 | 35 | 9 | 5 | 632 | 237 | 223 | 1084 | 686 | 154 | 1316 | 696 | 71 | 450 |
| PBANKA_1039300 | 16 | 25 | 93 | 18 | 159 | 101 | 194 | 97 | 129 | 87 | 24 | 8 | 1346 | 499 | 558 | 2426 | 2445 | 491 | 778 | 683 | 11 | 102 |
| PBANKA_1039400 | 25 | 32 | 1344 | 468 | 720 | 385 | 897 | 393 | 379 | 182 | 1569 | 1378 | 10451 | 3170 | 2507 | 5822 | 2152 | 700 | 1720 | 673 | 216 | 1264 |
| PBANKA_1039500 | 15 | 31 | 115 | 25 | 589 | 256 | 1904 | 949 | 1095 | 706 | 170 | 61 | 3222 | 2222 | 658 | 4308 | 4250 | 481 | 746 | 1158 | 51 | 389 |
| PBANKA_1039600 | 0 | 0 | 166 | 32 | 387 | 203 | 652 | 277 | 281 | 122 | 15 | 22 | 2719 | 932 | 1172 | 4804 | 3834 | 426 | 1052 | 672 | 78 | 612 |
| PBANKA_1039700 | 33 | 29 | 129 | 30 | 289 | 122 | 506 | 207 | 285 | 204 | 51 | 18 | 609 | 157 | 311 | 972 | 1771 | 627 | 1178 | 560 | 107 | 849 |
| PBANKA_1039800 | 0 | 0 | 67 | 15 | 157 | 57 | 132 | 65 | 75 | 38 | 5 | 5 | 387 | 116 | 269 | 1189 | 737 | 102 | 44 | 18 | 0 | 5 |
| PBANKA_1039900 | 0 | 0 | 30 | 8 | 177 | 96 | 277 | 110 | 69 | 36 | 0 | 14 | 287 | 58 | 364 | 1411 | 831 | 100 | 36 | 28 | 0 | 12 |
| PBANKA_1040000 | 0 | 0 | 19 | 0 | 61 | 36 | 74 | 52 | 75 | 51 | 22 | 6 | 948 | 317 | 180 | 681 | 875 | 233 | 2581 | 1803 | 14 | 66 |
| PBANKA_1040100 | 9 | 12 | 29 | 11 | 75 | 58 | 123 | 55 | 97 | 57 | 9 | 13 | 1420 | 542 | 332 | 1615 | 1695 | 355 | 37371 | 28569 | 49 | 349 |
| PBANKA_1040521 | 0 | 0 | 9 | 0 | 19 | 10 | 18 | 0 | 11 | 0 | 0 | 0 | 268 | 0 | 17 | 0 | 0 | 7 | 38 | 7 | 8 | 64 |
| PBANKA_1100441 | 0 | 0 | 5 | 0 | 26 | 30 | 36 | 14 | 18 | 0 | 7 | 0 | 999 | 0 | 63 | 11 | 16 | 18 | 89 | 25 | 0 | 30 |

| | | | | | | | | | | | | | | | | | | | | | | |
|---|---|---|---|---|---|---|---|---|---|---|---|---|---|---|---|---|---|---|---|---|---|---|
| PBANKA_1100860 | 0 | 0 | 11 | 0 | 39 | 23 | 46 | 14 | 77 | 88 | 871 | 659 | 1190 | 451 | 262 | 962 | 313 | 82 | 2969 | 1347 | 93 | 555 |
| PBANKA_1101000 | 0 | 0 | 18 | 0 | 546 | 334 | 500 | 247 | 154 | 65 | 8 | 19 | 159 | 52 | 304 | 867 | 113 | 23 | 75 | 47 | 0 | 19 |
| PBANKA_1101100 | 0 | 0 | 58 | 16 | 87 | 54 | 59 | 33 | 690 | 535 | 7793 | 5626 | 5380 | 5672 | 421 | 3037 | 1922 | 584 | 544 | 756 | 88 | 545 |
| PBANKA_1101200 | 0 | 0 | 22 | 5 | 31 | 33 | 34 | 10 | 709 | 382 | 4330 | 2881 | 11522 | 8339 | 472 | 2658 | 2029 | 1206 | 33162 | 23598 | 538 | 3683 |
| PBANKA_1101300 | 0 | 0 | 13 | 0 | 49 | 23 | 65 | 27 | 586 | 343 | 7646 | 6040 | 5668 | 6425 | 1906 | 7676 | 4798 | 923 | 1211 | 1275 | 9 | 128 |
| PBANKA_1101600 | 0 | 5 | 66 | 21 | 522 | 321 | 915 | 330 | 291 | 141 | 45 | 16 | 1564 | 561 | 637 | 1867 | 8710 | 952 | 4558 | 5700 | 169 | 1391 |
| PBANKA_1101700 | 0 | 0 | 56 | 11 | 156 | 55 | 150 | 56 | 55 | 22 | 0 | 9 | 517 | 151 | 186 | 842 | 359 | 68 | 639 | 325 | 14 | 73 |
| PBANKA_1101800 | 0 | 0 | 49 | 6 | 110 | 52 | 174 | 51 | 46 | 24 | 9 | 0 | 1219 | 580 | 435 | 2041 | 494 | 85 | 485 | 303 | 15 | 100 |
| PBANKA_1101900 | 0 | 0 | 11 | 0 | 72 | 35 | 68 | 14 | 25 | 13 | 23 | 22 | 549 | 241 | 248 | 1206 | 509 | 86 | 773 | 498 | 17 | 114 |
| PBANKA_1102000 | 0 | 0 | 47 | 10 | 117 | 55 | 134 | 21 | 57 | 23 | 42 | 26 | 866 | 761 | 236 | 1016 | 298 | 70 | 743 | 488 | 45 | 232 |
| PBANKA_1102200 | 0 | 0 | 29 | 10 | 176 | 91 | 269 | 144 | 966 | 714 | 6154 | 5223 | 33000 | 12986 | 6211 | 29151 | 10555 | 2255 | 1899 | 1834 | 43 | 363 |
| PBANKA_1102300 | 0 | 0 | 18 | 0 | 56 | 20 | 86 | 28 | 43 | 18 | 18 | 0 | 814 | 440 | 156 | 724 | 888 | 175 | 2872 | 4799 | 51 | 267 |
| PBANKA_1102400 | 0 | 0 | 42 | 0 | 125 | 50 | 114 | 32 | 39 | 13 | 9 | 0 | 317 | 87 | 142 | 629 | 227 | 30 | 446 | 246 | 18 | 89 |
| PBANKA_1102500 | 0 | 0 | 52 | 5 | 172 | 70 | 174 | 47 | 43 | 12 | 0 | 0 | 298 | 195 | 222 | 1003 | 344 | 55 | 181 | 141 | 62 | 399 |
| PBANKA_1102600 | 0 | 5 | 13 | 0 | 42 | 22 | 24 | 19 | 8 | 9 | 0 | 7 | 231 | 80 | 67 | 93 | 52 | 17 | 198 | 99 | 0 | 32 |
| PBANKA_1102700 | 13 | 14 | 77 | 18 | 277 | 75 | 227 | 89 | 77 | 60 | 26 | 6 | 767 | 276 | 365 | 1216 | 867 | 179 | 613 | 307 | 28 | 187 |
| PBANKA_1102800 | 0 | 0 | 33 | 9 | 87 | 40 | 136 | 41 | 47 | 17 | 6 | 0 | 389 | 91 | 252 | 1129 | 496 | 136 | 1312 | 726 | 88 | 480 |
| PBANKA_1102900 | 70 | 138 | 36 | 0 | 263 | 97 | 369 | 133 | 230 | 92 | 47 | 12 | 438 | 190 | 80 | 263 | 2444 | 316 | 3083 | 2297 | 1416 | 8191 |
| PBANKA_1103000 | 5 | 0 | 345 | 111 | 367 | 199 | 314 | 131 | 261 | 162 | 1586 | 1353 | 6910 | 3860 | 1355 | 8032 | 2095 | 247 | 3101 | 1989 | 70 | 450 |
| PBANKA_1103100 | 41 | 35 | 374 | 153 | 381 | 275 | 678 | 432 | 198 | 187 | 29 | 6 | 691 | 172 | 971 | 1755 | 2934 | 751 | 1269 | 501 | 379 | 3306 |
| PBANKA_1103200 | 0 | 5 | 49 | 10 | 113 | 38 | 116 | 54 | 71 | 40 | 29 | 7 | 1295 | 696 | 315 | 2144 | 888 | 119 | 806 | 895 | 836 | 5227 |
| PBANKA_1103300 | 0 | 0 | 15 | 0 | 69 | 28 | 49 | 26 | 257 | 208 | 56 | 43 | 311 | 262 | 122 | 549 | 741 | 286 | 209 | 480 | 2160 | 12444 |
| PBANKA_1103400 | 49 | 41 | 1312 | 351 | 801 | 604 | 1167 | 602 | 347 | 350 | 2169 | 1674 | 7293 | 1844 | 1833 | 4804 | 1738 | 391 | 2729 | 918 | 83 | 519 |
| PBANKA_1103500 | 0 | 0 | 157 | 25 | 369 | 123 | 398 | 161 | 126 | 65 | 17 | 19 | 2212 | 849 | 659 | 2839 | 2540 | 362 | 377 | 175 | 0 | 32 |
| PBANKA_1103600 | 0 | 0 | 185 | 24 | 416 | 137 | 434 | 121 | 230 | 76 | 96 | 72 | 1718 | 692 | 835 | 3648 | 1995 | 368 | 565 | 299 | 44 | 284 |
| PBANKA_1103700 | 0 | 0 | 15 | 0 | 17 | 6 | 44 | 8 | 95 | 62 | 30 | 9 | 130 | 22 | 37 | 101 | 325 | 253 | 1116 | 300 | 185 | 761 |
| PBANKA_1103800 | 398 | 502 | 48 | 10 | 153 | 61 | 166 | 68 | 570 | 294 | 135 | 41 | 782 | 256 | 291 | 945 | 3445 | 1242 | 2914 | 1218 | 456 | 2304 |
| PBANKA_1103900 | 0 | 7 | 7 | 0 | 20 | 7 | 18 | 8 | 22 | 0 | 0 | 0 | 241 | 71 | 158 | 930 | 394 | 97 | 1352 | 774 | 1023 | 6569 |
| PBANKA_1104000 | 0 | 6 | 115 | 44 | 131 | 76 | 92 | 25 | 43 | 23 | 9 | 5 | 357 | 90 | 388 | 1767 | 544 | 49 | 517 | 313 | 389 | 2663 |
| PBANKA_1104200 | 15 | 33 | 81 | 29 | 356 | 262 | 478 | 262 | 139 | 84 | 26 | 9 | 243 | 56 | 630 | 1585 | 963 | 163 | 1048 | 524 | 1308 | 10948 |
| PBANKA_1104300 | 6 | 13 | 92 | 16 | 245 | 127 | 377 | 182 | 158 | 76 | 111 | 68 | 2289 | 1206 | 465 | 2436 | 1279 | 191 | 428 | 394 | 101 | 724 |
| PBANKA_1104400 | 96 | 151 | 185 | 21 | 632 | 190 | 1363 | 457 | 1145 | 420 | 236 | 63 | 2112 | 2319 | 1223 | 4743 | 19720 | 3027 | 2406 | 13761 | 1362 | 8030 |
| PBANKA_1104500 | 63 | 83 | 5 | 0 | 59 | 47 | 186 | 118 | 128 | 79 | 13 | 0 | 302 | 46 | 29 | 37 | 477 | 230 | 732 | 272 | 85 | 381 |
| PBANKA_1104600 | 47 | 68 | 155 | 33 | 297 | 100 | 347 | 120 | 257 | 140 | 1495 | 842 | 1204 | 1464 | 546 | 3292 | 1729 | 225 | 784 | 1475 | 690 | 4599 |
| PBANKA_1104700 | 72 | 97 | 66 | 10 | 340 | 124 | 343 | 127 | 209 | 82 | 32 | 0 | 1169 | 1010 | 1043 | 4615 | 1364 | 188 | 968 | 1704 | 151 | 1208 |
| PBANKA_1104800 | 0 | 9 | 27 | 10 | 216 | 116 | 614 | 273 | 338 | 147 | 42 | 11 | 739 | 330 | 414 | 1948 | 1000 | 198 | 709 | 458 | 546 | 3506 |
| PBANKA_1104900 | 5 | 5 | 32 | 6 | 92 | 30 | 176 | 75 | 108 | 46 | 368 | 216 | 1330 | 498 | 207 | 795 | 1954 | 414 | 2444 | 686 | 371 | 2070 |
| PBANKA_1105000 | 0 | 0 | 223 | 39 | 322 | 105 | 355 | 129 | 112 | 48 | 8 | 0 | 1107 | 510 | 1457 | 7406 | 3864 | 380 | 130 | 264 | 5 | 20 |
| PBANKA_1105100 | 26 | 44 | 61 | 15 | 389 | 163 | 814 | 314 | 424 | 157 | 61 | 23 | 1032 | 413 | 411 | 2037 | 6723 | 876 | 3588 | 4013 | 116 | 885 |
| PBANKA_1105200 | 5 | 7 | 19 | 7 | 57 | 23 | 145 | 39 | 57 | 9 | 10 | 0 | 643 | 149 | 170 | 727 | 617 | 96 | 2940 | 1348 | 113 | 808 |
| PBANKA_1105300 | 79 | 117 | 22 | 0 | 120 | 45 | 184 | 63 | 102 | 32 | 19 | 0 | 461 | 118 | 56 | 253 | 224 | 47 | 1953 | 1136 | 178 | 1115 |
| PBANKA_1105400 | 20 | 18 | 90 | 38 | 190 | 165 | 280 | 184 | 109 | 95 | 48 | 51 | 782 | 409 | 261 | 901 | 856 | 123 | 470 | 272 | 28 | 210 |
| PBANKA_1105500 | 128 | 127 | 39 | 20 | 90 | 38 | 111 | 79 | 56 | 36 | 12 | 7 | 212 | 50 | 63 | 259 | 183 | 20 | 750 | 378 | 129 | 882 |
| PBANKA_1105600 | 99 | 102 | 29 | 0 | 82 | 19 | 111 | 43 | 60 | 19 | 15 | 0 | 402 | 111 | 86 | 275 | 359 | 94 | 1293 | 405 | 105 | 664 |
| PBANKA_1105700 | 32 | 27 | 259 | 73 | 215 | 90 | 159 | 75 | 138 | 57 | 790 | 514 | 3006 | 1659 | 349 | 1474 | 378 | 63 | 1637 | 750 | 102 | 402 |
| PBANKA_1105800 | 20 | 47 | 78 | 19 | 115 | 79 | 114 | 35 | 89 | 68 | 710 | 535 | 647 | 287 | 221 | 893 | 220 | 61 | 859 | 502 | 480 | 3107 |
| PBANKA_1105900 | 58 | 58 | 73 | 5 | 104 | 37 | 96 | 39 | 89 | 40 | 20 | 10 | 461 | 166 | 122 | 747 | 361 | 51 | 1155 | 939 | 656 | 3538 |
| PBANKA_1106000 | 0 | 9 | 10 | 0 | 71 | 59 | 57 | 24 | 15 | 8 | 0 | 0 | 292 | 122 | 0 | 8 | 93 | 63 | 6066 | 2511 | 59 | 483 |
| PBANKA_1106100 | 185 | 240 | 170 | 55 | 1080 | 466 | 6847 | 2938 | 4957 | 2335 | 503 | 275 | 496 | 727 | 604 | 2493 | 3429 | 502 | 1382 | 8414 | 166 | 1322 |
| PBANKA_1106200 | 0 | 0 | 29 | 6 | 100 | 41 | 106 | 17 | 29 | 14 | 0 | 0 | 39 | 12 | 70 | 176 | 131 | 23 | 369 | 153 | 37 | 176 |
| PBANKA_1106300 | 0 | 0 | 8 | 0 | 57 | 22 | 107 | 34 | 33 | 18 | 7 | 0 | 185 | 45 | 144 | 531 | 648 | 110 | 308 | 188 | 62 | 538 |
| PBANKA_1106400 | 7 | 16 | 120 | 28 | 251 | 98 | 406 | 121 | 241 | 139 | 500 | 259 | 2846 | 2221 | 872 | 4868 | 950 | 208 | 707 | 633 | 162 | 1141 |
| PBANKA_1106500 | 224 | 267 | 15 | 0 | 216 | 96 | 735 | 374 | 367 | 219 | 96 | 31 | 3432 | 1014 | 284 | 858 | 9477 | 1418 | 46027 | 31875 | 675 | 6554 |
| PBANKA_1106600 | 0 | 0 | 22 | 9 | 19 | 11 | 26 | 0 | 0 | 0 | 0 | 0 | 65 | 5 | 58 | 278 | 32 | 8 | 28 | 6 | 0 | 11 |
| PBANKA_1106700 | 247 | 334 | 4327 | 1414 | 3462 | 1937 | 3137 | 1355 | 1216 | 693 | 7626 | 4913 | 38931 | 21893 | 6645 | 30929 | 9522 | 1176 | 3605 | 3941 | 703 | 5447 |
| PBANKA_1106800 | 20 | 34 | 55 | 5 | 103 | 39 | 164 | 68 | 661 | 314 | 101 | 49 | 1242 | 434 | 134 | 737 | 805 | 319 | 5249 | 2117 | 248 | 1726 |
| PBANKA_1107100 | 0 | 0 | 19 | 9 | 109 | 51 | 223 | 81 | 587 | 402 | 146 | 84 | 701 | 244 | 165 | 672 | 1991 | 683 | 6325 | 3909 | 469 | 3752 |
| PBANKA_1107200 | 82 | 97 | 113 | 32 | 293 | 130 | 441 | 192 | 274 | 192 | 115 | 110 | 1358 | 609 | 584 | 2220 | 2621 | 632 | 1666 | 882 | 181 | 1603 |
| PBANKA_1107300 | 0 | 0 | 63 | 11 | 200 | 86 | 194 | 98 | 78 | 31 | 30 | 7 | 771 | 359 | 513 | 2137 | 938 | 155 | 730 | 621 | 137 | 1027 |
| PBANKA_1107400 | 76 | 91 | 112 | 25 | 260 | 149 | 418 | 140 | 309 | 134 | 727 | 548 | 3625 | 2138 | 1003 | 4476 | 4210 | 812 | 1900 | 1527 | 674 | 5051 |
| PBANKA_1107500 | 0 | 0 | 50 | 9 | 31 | 14 | 70 | 61 | 25 | 17 | 0 | 6 | 307 | 47 | 99 | 283 | 279 | 103 | 354 | 141 | 86 | 392 |
| PBANKA_1107600 | 2228 | 3266 | 86 | 26 | 326 | 198 | 1031 | 508 | 694 | 419 | 44 | 25 | 959 | 153 | 26 | 35 | 6440 | 1341 | 1228 | 684 | 3631 | 29820 |
| PBANKA_1107700 | 1097 | 1569 | 84 | 18 | 514 | 260 | 1265 | 653 | 1066 | 538 | 133 | 68 | 828 | 393 | 113 | 227 | 9040 | 1646 | 1026 | 1988 | 5127 | 33956 |
| PBANKA_1107800 | 50 | 65 | 84 | 26 | 173 | 90 | 222 | 78 | 124 | 108 | 48 | 40 | 725 | 163 | 450 | 1603 | 2091 | 621 | 5087 | 2738 | 339 | 3161 |
| PBANKA_1107900 | 7 | 12 | 233 | 69 | 564 | 346 | 613 | 227 | 239 | 126 | 681 | 453 | 1011 | 301 | 755 | 2660 | 1615 | 353 | 4285 | 2298 | 142 | 1220 |
| PBANKA_1108000 | 0 | 0 | 6 | 0 | 118 | 57 | 92 | 54 | 106 | 59 | 26 | 12 | 415 | 220 | 37 | 129 | 578 | 149 | 2781 | 2197 | 54 | 338 |
| PBANKA_1108100 | 32 | 30 | 55 | 9 | 214 | 98 | 495 | 177 | 566 | 242 | 98 | 29 | 1058 | 986 | 446 | 2721 | 9023 | 1521 | 1048 | 2300 | 505 | 3497 |
| PBANKA_1108200 | 6 | 5 | 50 | 13 | 126 | 74 | 233 | 104 | 191 | 152 | 37 | 22 | 861 | 522 | 150 | 917 | 3466 | 420 | 395 | 404 | 157 | 1437 |
| PBANKA_1108300 | 300 | 550 | 317 | 59 | 1047 | 391 | 1147 | 345 | 506 | 161 | 369 | 152 | 5042 | 7260 | 2234 | 15832 | 13761 | 1491 | 3026 | 10418 | 6078 | 45794 |
| PBANKA_1108400 | 0 | 0 | 53 | 18 | 147 | 66 | 191 | 75 | 97 | 59 | 22 | 9 | 519 | 76 | 343 | 1767 | 1013 | 133 | 456 | 382 | 114 | 1032 |
| PBANKA_1108500 | 8 | 10 | 8 | 10 | 148 | 71 | 602 | 200 | 3143 | 1431 | 1033 | 431 | 1334 | 3446 | 898 | 4273 | 33882 | 5685 | 7391 | 49502 | 1781 | 11940 |
| PBANKA_1108600 | 198 | 245 | 196 | 57 | 437 | 200 | 520 | 229 | 316 | 161 | 57 | 28 | 613 | 250 | 392 | 1374 | 2260 | 651 | 875 | 1820 | 131 | 935 |
| PBANKA_1108700 | 22 | 32 | 146 | 46 | 668 | 238 | 1225 | 529 | 703 | 272 | 96 | 32 | 2498 | 2608 | 782 | 3108 | 21561 | 2515 | 3442 | 45957 | 955 | 5602 |
| PBANKA_1108900 | 9 | 20 | 45 | 9 | 54 | 28 | 60 | 27 | 18 | 20 | 0 | 0 | 818 | 177 | 253 | 1149 | 730 | 140 | 1010 | 506 | 47 | 264 |
| PBANKA_1109000 | 101 | 174 | 1034 | 252 | 2138 | 921 | 1826 | 581 | 653 | 285 | 1124 | 686 | 5295 | 7598 | 4592 | 22082 | 5129 | 606 | 2145 | 5500 | 460 | 3577 |
| PBANKA_1109200 | 48 | 77 | 105 | 23 | 237 | 153 | 192 | 72 | 76 | 34 | 15 | 0 | 529 | 106 | 371 | 1359 | 279 | 51 | 500 | 214 | 9 | 68 |
| PBANKA_1109300 | 0 | 0 | 36 | 11 | 71 | 38 | 53 | 13 | 12 | 10 | 7 | 0 | 987 | 451 | 46 | 301 | 346 | 164 | 21526 | 12648 | 355 | 2504 |
| PBANKA_1109400 | 0 | 0 | 229 | 66 | 498 | 202 | 664 | 240 | 242 | 114 | 182 | 91 | 1599 | 1786 | 1073 | 5065 | 3097 | 381 | 1802 | 4600 | 122 | 1170 |
| PBANKA_1109500 | 0 | 0 | 12 | 0 | 54 | 9 | 52 | 17 | 22 | 12 | 8 | 0 | 157 | 25 | 142 | 573 | 338 | 68 | 274 | 105 | 19 | 189 |
| PBANKA_1109700 | 130 | 234 | 43 | 11 | 658 | 276 | 2093 | 865 | 999 | 457 | 93 | 23 | 1159 | 946 | 1103 | 4541 | 15068 | 1588 | 1758 | 2685 | 361 | 2638 |
| PBANKA_1109800 | 0 | 0 | 75 | 19 | 112 | 50 | 233 | 99 | 121 | 52 | 103 | 41 | 949 | 539 | 457 | 2217 | 971 | 145 | 569 | 419 | 56 | 328 |
| PBANKA_1110000 | 0 | 0 | 38 | 6 | 47 | 25 | 20 | 8 | 16 | 0 | 24 | 24 | 450 | 248 | 76 | 477 | 132 | 13 | 361 | 252 | 1628 | 7923 |
| PBANKA_1110100 | 111 | 180 | 97 | 14 | 310 | 125 | 561 | 231 | 315 | 145 | 55 | 13 | 1498 | 815 | 381 | 2393 | 7394 | 988 | 2203 | 3672 | 71 | 421 |
| PBANKA_1110200 | 6 | 11 | 5 | 0 | 12 | 11 | 17 | 11 | 19 | 17 | 5 | 0 | 907 | 375 | 64 | 270 | 232 | 147 | 24237 | 15820 | 446 | 3783 |
| PBANKA_1110300 | 0 | 0 | 6 | 0 | 19 | 5 | 35 | 16 | 24 | 0 | 9 | 15 | 364 | 106 | 39 | 129 | 294 | 86 | 2468 | 1483 | 11 | 150 |
| PBANKA_1110350 | 11 | 11 | 18 | 0 | 34 | 19 | 34 | 13 | 20 | 6 | 14 | 11 | 95 | 42 | 125 | 74 | 62 | 20 | 238 | 50 | 0 | 35 |
| PBANKA_1110500 | 14 | 8 | 656 | 183 | 598 | 354 | 988 | 488 | 559 | 321 | 195 | 162 | 3351 | 1898 | 1665 | 6095 | 2684 | 368 | 1557 | 1005 | 115 | 1241 |
| PBANKA_1110600 | 0 | 0 | 5 | 0 | 11 | 6 | 40 | 12 | 37 | 11 | 0 | 0 | 282 | 66 | 35 | 95 | 63 | 49 | 6509 | 2934 | 68 | 488 |
| PBANKA_1110700 | 62 | 59 | 284 | 44 | 1883 | 771 | 3779 | 1552 | 2661 | 1317 | 297 | 123 | 750 | 262 | 417 | 955 | 4336 | 811 | 6989 | 4943 | 329 | 2721 |
| PBANKA_1110800 | 156 | 168 | 33 | 0 | 212 | 95 | 387 | 153 | 162 | 81 | 46 | 19 | 773 | 370 | 354 | 1894 | 1924 | 229 | 510 | 628 | 330 | 1865 |

| | | | | | | | | | | | | | | | | | | | | | | |
|---|---|---|---|---|---|---|---|---|---|---|---|---|---|---|---|---|---|---|---|---|---|---|
| PBANKA_1110900 | 1131 | 1503 | 8 | 0 | 47 | 21 | 73 | 35 | 71 | 28 | 8 | 10 | 104 | 52 | 27 | 117 | 304 | 43 | 490 | 662 | 140 | 817 |
| PBANKA_1111000 | 718 | 1263 | 1426 | 342 | 3563 | 1350 | 5365 | 1899 | 3296 | 1372 | 4393 | 2478 | 6718 | 43098 | 7984 | 44028 | 56041 | 6576 | 771 | 12278 | 1198 | 6605 |
| PBANKA_1111200 | 27 | 36 | 7 | 0 | 37 | 8 | 54 | 16 | 121 | 59 | 7 | 7 | 111 | 42 | 63 | 206 | 619 | 208 | 366 | 120 | 42 | 213 |
| PBANKA_1111300 | 0 | 0 | 46 | 11 | 122 | 57 | 140 | 46 | 61 | 34 | 8 | 0 | 145 | 17 | 137 | 250 | 155 | 25 | 505 | 163 | 62 | 370 |
| PBANKA_1111400 | 0 | 8 | 250 | 58 | 1493 | 525 | 1817 | 704 | 777 | 345 | 84 | 37 | 2533 | 778 | 1389 | 6592 | 3291 | 449 | 1831 | 1119 | 156 | 1457 |
| PBANKA_1111500 | 0 | 0 | 19 | 0 | 171 | 54 | 213 | 68 | 142 | 64 | 30 | 18 | 323 | 98 | 86 | 663 | 621 | 115 | 914 | 661 | 35 | 253 |
| PBANKA_1111600 | 0 | 0 | 104 | 19 | 65 | 43 | 81 | 26 | 42 | 22 | 14 | 0 | 256 | 76 | 179 | 384 | 227 | 59 | 630 | 207 | 55 | 147 |
| PBANKA_1111700 | 0 | 0 | 39 | 6 | 39 | 24 | 33 | 11 | 17 | 0 | 0 | 0 | 293 | 72 | 198 | 499 | 147 | 44 | 711 | 295 | 62 | 385 |
| PBANKA_1111800 | 0 | 0 | 56 | 10 | 146 | 60 | 156 | 69 | 39 | 19 | 5 | 5 | 654 | 164 | 404 | 1617 | 525 | 68 | 109 | 55 | 299 | 1823 |
| PBANKA_1111900 | 31 | 56 | 63 | 14 | 131 | 49 | 153 | 65 | 42 | 17 | 18 | 17 | 423 | 173 | 196 | 1120 | 643 | 102 | 975 | 674 | 272 | 1992 |
| PBANKA_1112000 | 279 | 453 | 5 | 0 | 15 | 0 | 11 | 0 | 10 | 8 | 0 | 0 | 34 | 9 | 14 | 88 | 56 | 12 | 159 | 92 | 1304 | 9340 |
| PBANKA_1112100 | 9 | 7 | 55 | 9 | 122 | 48 | 143 | 37 | 133 | 53 | 9 | 0 | 136 | 53 | 167 | 742 | 834 | 211 | 302 | 131 | 72 | 284 |
| PBANKA_1112200 | 0 | 0 | 133 | 32 | 382 | 126 | 315 | 99 | 138 | 60 | 20 | 14 | 512 | 133 | 634 | 3182 | 895 | 170 | 1665 | 1038 | 193 | 1162 |
| PBANKA_1112300 | 0 | 0 | 130 | 19 | 260 | 178 | 404 | 231 | 154 | 99 | 18 | 17 | 2364 | 1048 | 710 | 2671 | 2957 | 495 | 1136 | 788 | 164 | 937 |
| PBANKA_1112400 | 5 | 6 | 144 | 32 | 132 | 46 | 126 | 47 | 28 | 17 | 31 | 14 | 1981 | 987 | 466 | 2076 | 463 | 71 | 593 | 581 | 53 | 309 |
| PBANKA_1112500 | 20 | 27 | 107 | 29 | 333 | 148 | 301 | 101 | 135 | 74 | 31 | 6 | 1959 | 588 | 791 | 3831 | 1387 | 261 | 3809 | 2466 | 466 | 2574 |
| PBANKA_1112600 | 0 | 0 | 33 | 5 | 46 | 23 | 64 | 16 | 18 | 11 | 0 | 0 | 763 | 200 | 305 | 1452 | 220 | 20 | 675 | 259 | 158 | 1051 |
| PBANKA_1112800 | 0 | 0 | 11 | 0 | 76 | 25 | 253 | 110 | 326 | 165 | 46 | 22 | 664 | 153 | 54 | 321 | 2476 | 667 | 2542 | 1560 | 145 | 1018 |
| PBANKA_1112900 | 39 | 60 | 91 | 34 | 454 | 116 | 791 | 253 | 603 | 230 | 112 | 45 | 1696 | 720 | 754 | 3920 | 4853 | 826 | 2542 | 2493 | 443 | 1973 |
| PBANKA_1113000 | 8 | 7 | 543 | 145 | 1212 | 804 | 1579 | 935 | 384 | 267 | 90 | 55 | 15776 | 2770 | 6001 | 29539 | 1490 | 609 | 3437 | 1640 | 395 | 4209 |
| PBANKA_1113100 | 34 | 37 | 24 | 14 | 93 | 56 | 71 | 31 | 18 | 25 | 6 | 0 | 307 | 76 | 100 | 312 | 40 | 15 | 335 | 134 | 219 | 1341 |
| PBANKA_1113200 | 24 | 48 | 19 | 0 | 49 | 12 | 51 | 19 | 25 | 15 | 25 | 8 | 416 | 117 | 99 | 267 | 242 | 99 | 734 | 265 | 106 | 648 |
| PBANKA_1113300 | 225 | 395 | 207 | 49 | 665 | 369 | 1142 | 376 | 513 | 273 | 67 | 40 | 840 | 257 | 196 | 654 | 4518 | 431 | 3174 | 2207 | 493 | 3551 |
| PBANKA_1113400 | 241 | 298 | 9 | 0 | 177 | 137 | 407 | 158 | 247 | 110 | 42 | 22 | 660 | 329 | 65 | 213 | 1791 | 760 | 11725 | 9013 | 2116 | 17344 |
| PBANKA_1113500 | 0 | 0 | 140 | 35 | 238 | 129 | 340 | 208 | 123 | 60 | 13 | 0 | 980 | 141 | 652 | 2263 | 1548 | 287 | 3805 | 2680 | 207 | 1650 |
| PBANKA_1113600 | 0 | 9 | 41 | 0 | 96 | 22 | 82 | 21 | 19 | 16 | 0 | 0 | 317 | 23 | 146 | 642 | 179 | 52 | 468 | 180 | 182 | 1217 |
| PBANKA_1113800 | 89 | 117 | 59 | 22 | 256 | 93 | 394 | 149 | 208 | 68 | 19 | 12 | 332 | 209 | 167 | 881 | 1919 | 233 | 1452 | 1587 | 1097 | 7190 |
| PBANKA_1113900 | 87 | 154 | 98 | 32 | 661 | 245 | 726 | 253 | 305 | 130 | 154 | 87 | 1147 | 589 | 445 | 2407 | 1468 | 240 | 793 | 703 | 838 | 3880 |
| PBANKA_1114000 | 31 | 22 | 100 | 12 | 363 | 158 | 476 | 190 | 343 | 143 | 51 | 13 | 707 | 225 | 331 | 1490 | 1957 | 334 | 2446 | 1661 | 147 | 933 |
| PBANKA_1114200 | 10 | 30 | 47 | 10 | 89 | 65 | 61 | 36 | 46 | 30 | 11 | 9 | 691 | 175 | 111 | 347 | 281 | 151 | 10860 | 4419 | 176 | 1207 |
| PBANKA_1114300 | 13 | 34 | 12 | 0 | 128 | 60 | 214 | 96 | 164 | 88 | 13 | 7 | 623 | 329 | 280 | 1050 | 1807 | 349 | 2199 | 3680 | 94 | 520 |
| PBANKA_1114400 | 426 | 715 | 77 | 17 | 222 | 110 | 296 | 112 | 164 | 79 | 216 | 176 | 1683 | 1163 | 281 | 1760 | 1273 | 205 | 1252 | 897 | 579 | 4038 |
| PBANKA_1114500 | 127 | 123 | 18 | 5 | 53 | 23 | 104 | 47 | 53 | 30 | 41 | 20 | 568 | 221 | 109 | 425 | 626 | 127 | 854 | 308 | 283 | 1608 |
| PBANKA_1114600 | 6 | 5 | 79 | 27 | 179 | 107 | 318 | 141 | 157 | 75 | 432 | 231 | 2281 | 917 | 441 | 2437 | 1772 | 270 | 7547 | 5256 | 412 | 3436 |
| PBANKA_1114700 | 18 | 13 | 30 | 11 | 99 | 40 | 140 | 43 | 61 | 25 | 18 | 0 | 350 | 76 | 172 | 632 | 635 | 105 | 869 | 431 | 108 | 677 |
| PBANKA_1114800 | 7 | 0 | 21 | 5 | 41 | 17 | 24 | 14 | 17 | 7 | 9 | 0 | 87 | 20 | 36 | 144 | 150 | 42 | 110 | 64 | 11 | 45 |
| PBANKA_1114900 | 59 | 108 | 139 | 27 | 547 | 174 | 504 | 166 | 268 | 131 | 73 | 24 | 1147 | 654 | 1197 | 4966 | 2618 | 441 | 2220 | 3877 | 279 | 2058 |
| PBANKA_1115000 | 0 | 0 | 5 | 0 | 17 | 7 | 39 | 14 | 21 | 18 | 7 | 0 | 2079 | 1601 | 70 | 197 | 1147 | 429 | 50395 | 68078 | 1860 | 12005 |
| PBANKA_1115100 | 0 | 0 | 10 | 0 | 100 | 37 | 243 | 66 | 212 | 109 | 42 | 10 | 644 | 144 | 252 | 910 | 2094 | 659 | 3237 | 1287 | 79 | 727 |
| PBANKA_1115200 | 6 | 8 | 13 | 0 | 241 | 80 | 253 | 145 | 526 | 233 | 78 | 49 | 953 | 693 | 202 | 979 | 2905 | 859 | 11201 | 12536 | 989 | 7872 |
| PBANKA_1115300 | 31 | 34 | 9 | 0 | 150 | 107 | 1632 | 814 | 3265 | 2092 | 329 | 206 | 7674 | 2022 | 113 | 296 | 32843 | 8698 | 2E+05 | 93476 | 698 | 8606 |
| PBANKA_1115400 | 12 | 5 | 66 | 22 | 107 | 52 | 101 | 42 | 23 | 23 | 6 | 0 | 285 | 39 | 237 | 453 | 100 | 35 | 510 | 215 | 18 | 96 |
| PBANKA_1115500 | 14 | 26 | 94 | 16 | 213 | 89 | 401 | 139 | 230 | 110 | 86 | 52 | 1240 | 346 | 553 | 2034 | 1507 | 370 | 1083 | 565 | 194 | 1445 |
| PBANKA_1115600 | 8 | 8 | 18 | 0 | 44 | 13 | 48 | 14 | 27 | 14 | 15 | 9 | 429 | 143 | 125 | 839 | 410 | 113 | 1541 | 998 | 115 | 690 |
| PBANKA_1115700 | 14 | 6 | 133 | 34 | 252 | 111 | 740 | 296 | 545 | 284 | 66 | 26 | 1512 | 642 | 302 | 1402 | 9741 | 1481 | 4765 | 4925 | 2056 | 14025 |
| PBANKA_1115800 | 22 | 25 | 62 | 13 | 104 | 49 | 166 | 57 | 103 | 46 | 8 | 0 | 845 | 160 | 202 | 755 | 963 | 369 | 708 | 307 | 233 | 1328 |
| PBANKA_1115900 | 71 | 101 | 58 | 17 | 252 | 67 | 303 | 93 | 252 | 115 | 53 | 15 | 618 | 249 | 294 | 1302 | 2364 | 521 | 1004 | 539 | 98 | 459 |
| PBANKA_1116100 | 66 | 89 | 99 | 33 | 573 | 239 | 739 | 303 | 227 | 131 | 35 | 16 | 526 | 148 | 598 | 2670 | 1254 | 201 | 1393 | 1207 | 640 | 4588 |
| PBANKA_1116200 | 0 | 0 | 12 | 7 | 156 | 70 | 339 | 159 | 137 | 47 | 9 | 0 | 429 | 93 | 173 | 674 | 1252 | 259 | 2323 | 823 | 274 | 1573 |
| PBANKA_1116300 | 0 | 8 | 109 | 25 | 351 | 150 | 557 | 230 | 276 | 133 | 25 | 20 | 749 | 199 | 276 | 1141 | 2519 | 407 | 899 | 664 | 106 | 851 |
| PBANKA_1116400 | 45 | 40 | 176 | 36 | 398 | 143 | 395 | 142 | 194 | 70 | 521 | 295 | 2209 | 3931 | 1129 | 7384 | 2651 | 344 | 842 | 1754 | 2068 | 12503 |
| PBANKA_1116500 | 0 | 0 | 199 | 71 | 282 | 174 | 160 | 86 | 68 | 43 | 19 | 10 | 2274 | 918 | 731 | 2773 | 762 | 163 | 1825 | 944 | 158 | 1499 |
| PBANKA_1116700 | 17 | 7 | 46 | 7 | 136 | 39 | 281 | 123 | 242 | 100 | 272 | 116 | 2756 | 2055 | 426 | 2107 | 3638 | 625 | 1532 | 1285 | 296 | 2277 |
| PBANKA_1116800 | 39 | 58 | 1143 | 187 | 1249 | 546 | 1498 | 724 | 513 | 266 | 808 | 552 | 44266 | 25723 | 5994 | 40284 | 14116 | 1706 | 5285 | 7667 | 3743 | 17433 |
| PBANKA_1116900 | 468 | 565 | 20 | 5 | 36 | 27 | 129 | 54 | 72 | 69 | 13 | 9 | 103 | 13 | 57 | 127 | 463 | 262 | 453 | 146 | 295 | 2171 |
| PBANKA_1117000 | 733 | 809 | 1804 | 403 | 5430 | 1873 | 11309 | 3425 | 8000 | 3813 | 1352 | 448 | 3404 | 791 | 733 | 1671 | 19734 | 8100 | 6835 | 2485 | 342 | 2601 |
| PBANKA_1117100 | 1326 | 1530 | 871 | 279 | 1456 | 1002 | 3018 | 1324 | 2428 | 1747 | 1753 | 1370 | 2737 | 1198 | 1299 | 3881 | 6042 | 2555 | 3472 | 1773 | 946 | 7249 |
| PBANKA_1117300 | 0 | 0 | 66 | 21 | 206 | 70 | 207 | 99 | 115 | 36 | 22 | 0 | 891 | 271 | 354 | 1328 | 811 | 128 | 575 | 473 | 2898 | 17729 |
| PBANKA_1117400 | 0 | 0 | 145 | 19 | 116 | 50 | 88 | 28 | 30 | 10 | 11 | 0 | 5778 | 1712 | 775 | 4582 | 664 | 85 | 1044 | 601 | 292 | 1398 |
| PBANKA_1117500 | 61 | 82 | 1313 | 434 | 842 | 551 | 955 | 519 | 362 | 266 | 1184 | 982 | 8651 | 2968 | 2364 | 6232 | 2873 | 651 | 4670 | 2089 | 302 | 2221 |
| PBANKA_1117600 | 63 | 69 | 37 | 9 | 191 | 68 | 156 | 40 | 78 | 17 | 30 | 6 | 355 | 89 | 50 | 125 | 45 | 16 | 881 | 368 | 605 | 3637 |
| PBANKA_1117700 | 345 | 458 | 219 | 52 | 539 | 287 | 1028 | 504 | 314 | 256 | 31 | 13 | 1125 | 162 | 755 | 1932 | 5080 | 831 | 2337 | 1130 | 371 | 3223 |
| PBANKA_1117800 | 0 | 0 | 26 | 12 | 134 | 95 | 139 | 46 | 56 | 30 | 10 | 0 | 855 | 186 | 418 | 1020 | 194 | 62 | 364 | 121 | 9 | 74 |
| PBANKA_1117900 | 0 | 5 | 19 | 0 | 68 | 32 | 74 | 27 | 24 | 9 | 6 | 0 | 447 | 88 | 89 | 535 | 304 | 43 | 696 | 276 | 66 | 416 |
| PBANKA_1118000 | 59 | 75 | 39 | 9 | 248 | 85 | 271 | 98 | 219 | 112 | 40 | 0 | 979 | 351 | 281 | 1428 | 1061 | 184 | 4365 | 1818 | 596 | 2859 |
| PBANKA_1118100 | 171 | 217 | 36 | 8 | 185 | 74 | 359 | 138 | 245 | 84 | 49 | 7 | 613 | 181 | 230 | 1092 | 1986 | 343 | 2012 | 1351 | 74 | 494 |
| PBANKA_1118200 | 168 | 221 | 77 | 10 | 167 | 58 | 235 | 100 | 209 | 116 | 37 | 6 | 1339 | 713 | 410 | 1885 | 1867 | 469 | 2108 | 1797 | 873 | 4719 |
| PBANKA_1118300 | 31 | 26 | 36 | 8 | 65 | 31 | 70 | 24 | 52 | 23 | 16 | 0 | 516 | 100 | 43 | 267 | 532 | 133 | 4098 | 1278 | 182 | 1149 |
| PBANKA_1118400 | 41 | 52 | 94 | 11 | 320 | 113 | 467 | 128 | 205 | 87 | 41 | 28 | 1715 | 1143 | 500 | 3213 | 3265 | 396 | 1124 | 925 | 991 | 4454 |
| PBANKA_1118500 | 91 | 169 | 120 | 53 | 553 | 291 | 759 | 298 | 306 | 143 | 51 | 16 | 478 | 313 | 544 | 2357 | 3866 | 508 | 1196 | 3394 | 9942 | 68296 |
| PBANKA_1118600 | 10 | 17 | 678 | 194 | 1423 | 977 | 2289 | 1023 | 1068 | 522 | 106 | 35 | 3845 | 562 | 3768 | 11481 | 8315 | 1273 | 4958 | 2634 | 144 | 1630 |
| PBANKA_1118700 | 8 | 29 | 85 | 12 | 427 | 175 | 775 | 304 | 363 | 190 | 275 | 180 | 3054 | 1332 | 1320 | 6038 | 6048 | 885 | 2277 | 934 | 1331 | 7556 |
| PBANKA_1118800 | 32 | 44 | 72 | 17 | 192 | 78 | 269 | 99 | 166 | 67 | 46 | 18 | 785 | 455 | 291 | 1614 | 1154 | 176 | 269 | 181 | 486 | 3013 |
| PBANKA_1118900 | 57 | 78 | 29 | 0 | 117 | 27 | 163 | 48 | 124 | 47 | 32 | 5 | 723 | 235 | 138 | 543 | 3052 | 553 | 1107 | 848 | 233 | 1265 |
| PBANKA_1119000 | 58 | 55 | 64 | 16 | 283 | 94 | 441 | 140 | 281 | 111 | 103 | 59 | 1052 | 684 | 395 | 2493 | 2359 | 355 | 1577 | 1352 | 91 | 521 |
| PBANKA_1119100 | 0 | 0 | 30 | 9 | 71 | 32 | 95 | 45 | 53 | 34 | 74 | 33 | 598 | 251 | 169 | 611 | 849 | 263 | 500 | 240 | 370 | 1656 |
| PBANKA_1119200 | 0 | 0 | 24 | 16 | 27 | 9 | 16 | 14 | 8 | 0 | 6 | 0 | 102 | 39 | 10 | 30 | 64 | 29 | 1455 | 948 | 19225 | 1E+05 |
| PBANKA_1119300 | 5 | 8 | 65 | 22 | 125 | 99 | 203 | 146 | 115 | 67 | 16 | 0 | 248 | 37 | 125 | 403 | 751 | 240 | 583 | 278 | 158 | 953 |
| PBANKA_1119400 | 28 | 21 | 13 | 0 | 17 | 17 | 23 | 8 | 11 | 6 | 8 | 9 | 230 | 55 | 55 | 212 | 23 | 18 | 127 | 76 | 106 | 666 |
| PBANKA_1119500 | 13 | 7 | 7 | 6 | 29 | 22 | 62 | 10 | 266 | 120 | 77 | 32 | 268 | 85 | 74 | 389 | 1104 | 343 | 317 | 346 | 107 | 653 |
| PBANKA_1119700 | 21 | 20 | 131 | 24 | 289 | 154 | 348 | 159 | 174 | 89 | 67 | 17 | 2166 | 1149 | 874 | 4736 | 2596 | 340 | 2926 | 2071 | 281 | 1930 |
| PBANKA_1119900 | 71 | 68 | 48 | 0 | 205 | 80 | 336 | 138 | 142 | 80 | 26 | 12 | 252 | 38 | 38 | 81 | 966 | 229 | 1127 | 426 | 119 | 885 |
| PBANKA_1120000 | 24 | 16 | 69 | 5 | 160 | 51 | 172 | 49 | 70 | 37 | 25 | 7 | 2272 | 1285 | 329 | 1850 | 1065 | 175 | 956 | 895 | 348 | 2097 |
| PBANKA_1120100 | 0 | 0 | 5 | 0 | 88 | 31 | 855 | 388 | 909 | 469 | 73 | 37 | 1491 | 453 | 132 | 307 | 11536 | 2097 | 3721 | 3587 | 503 | 3988 |
| PBANKA_1120200 | 0 | 0 | 178 | 33 | 84 | 90 | 60 | 26 | 31 | 11 | 101 | 69 | 544 | 264 | 160 | 747 | 76 | 18 | 872 | 513 | 40 | 408 |
| PBANKA_1120300 | 12 | 15 | 27 | 10 | 42 | 6 | 23 | 0 | 16 | 14 | 173 | 108 | 1501 | 1153 | 190 | 1222 | 161 | 44 | 819 | 687 | 97 | 603 |
| PBANKA_1120500 | 0 | 0 | 29 | 0 | 38 | 26 | 67 | 15 | 15 | 5 | 0 | 0 | 235 | 33 | 150 | 652 | 200 | 29 | 110 | 55 | 0 | 18 |
| PBANKA_1120600 | 14 | 15 | 223 | 63 | 416 | 136 | 411 | 150 | 185 | 60 | 16 | 14 | 4233 | 2398 | 759 | 5035 | 1171 | 163 | 1746 | 1312 | 117 | 704 |
| PBANKA_1120700 | 113 | 177 | 1052 | 366 | 807 | 567 | 513 | 333 | 171 | 80 | 635 | 674 | 7746 | 2428 | 1692 | 5826 | 709 | 151 | 1448 | 589 | 111 | 708 |
| PBANKA_1120800 | 0 | 0 | 124 | 28 | 273 | 98 | 238 | 73 | 67 | 28 | 124 | 97 | 2224 | 1254 | 797 | 3640 | 966 | 158 | 1102 | 851 | 62 | 441 |

| | | | | | | | | | | | | | | | | | | | | | | |
|---|---|---|---|---|---|---|---|---|---|---|---|---|---|---|---|---|---|---|---|---|---|---|
| PBANKA_1120900 | 0 | 0 | 6 | 0 | 20 | 14 | 24 | 11 | 6 | 6 | 0 | 5 | 60 | 31 | 28 | 91 | 47 | 30 | 815 | 322 | 15 | 136 |
| PBANKA_1121000 | 0 | 0 | 14 | 0 | 153 | 28 | 349 | 156 | 190 | 84 | 31 | 0 | 203 | 289 | 82 | 191 | 668 | 161 | 2667 | 8711 | 3456 | 22189 |
| PBANKA_1121100 | 0 | 8 | 111 | 28 | 187 | 55 | 206 | 71 | 103 | 25 | 192 | 116 | 2997 | 2239 | 562 | 3232 | 2081 | 234 | 500 | 434 | 117 | 844 |
| PBANKA_1121200 | 0 | 5 | 72 | 14 | 232 | 97 | 212 | 84 | 98 | 35 | 15 | 0 | 1099 | 477 | 613 | 2942 | 873 | 138 | 1177 | 812 | 95 | 673 |
| PBANKA_1121300 | 0 | 0 | 34 | 5 | 60 | 27 | 56 | 21 | 16 | 0 | 0 | 0 | 96 | 16 | 113 | 370 | 87 | 11 | 639 | 203 | 29 | 226 |
| PBANKA_1121400 | 0 | 0 | 73 | 22 | 94 | 39 | 90 | 37 | 37 | 20 | 15 | 9 | 509 | 118 | 218 | 798 | 379 | 84 | 1014 | 622 | 458 | 3755 |
| PBANKA_1121500 | 0 | 0 | 97 | 20 | 204 | 108 | 348 | 199 | 108 | 94 | 21 | 10 | 228 | 19 | 255 | 450 | 460 | 152 | 819 | 299 | 211 | 1379 |
| PBANKA_1121600 | 11 | 0 | 65 | 9 | 138 | 44 | 226 | 70 | 75 | 32 | 21 | 6 | 1088 | 323 | 365 | 1605 | 1110 | 216 | 1735 | 795 | 230 | 1412 |
| PBANKA_1121700 | 51 | 94 | 508 | 123 | 1163 | 450 | 1247 | 486 | 436 | 195 | 1255 | 816 | 5396 | 5654 | 4318 | 23656 | 8421 | 956 | 2989 | 6586 | 446 | 2950 |
| PBANKA_1121800 | 8 | 14 | 135 | 39 | 541 | 183 | 455 | 162 | 252 | 86 | 156 | 87 | 2239 | 1667 | 615 | 2966 | 1450 | 250 | 460 | 1357 | 152 | 1070 |
| PBANKA_1121900 | 13 | 8 | 203 | 61 | 454 | 207 | 427 | 160 | 93 | 61 | 10 | 0 | 833 | 97 | 969 | 5049 | 1468 | 149 | 803 | 610 | 232 | 1401 |
| PBANKA_1122000 | 61 | 56 | 207 | 61 | 439 | 215 | 682 | 340 | 378 | 201 | 553 | 449 | 5193 | 2975 | 1071 | 5675 | 4295 | 678 | 3293 | 2168 | 394 | 2586 |
| PBANKA_1122100 | 61 | 61 | 78 | 9 | 115 | 31 | 138 | 39 | 76 | 22 | 39 | 27 | 715 | 410 | 121 | 643 | 732 | 132 | 1317 | 1171 | 489 | 3223 |
| PBANKA_1122200 | 0 | 5 | 79 | 18 | 141 | 99 | 237 | 119 | 110 | 40 | 8 | 13 | 242 | 66 | 86 | 284 | 854 | 153 | 50 | 48 | 170 | 1234 |
| PBANKA_1122300 | 5 | 0 | 90 | 13 | 338 | 101 | 422 | 150 | 177 | 88 | 26 | 11 | 124 | 49 | 168 | 632 | 492 | 106 | 1295 | 1241 | 382 | 3069 |
| PBANKA_1122400 | 15 | 0 | 20 | 0 | 81 | 58 | 130 | 60 | 40 | 21 | 6 | 9 | 238 | 48 | 108 | 461 | 209 | 55 | 606 | 261 | 118 | 559 |
| PBANKA_1122500 | 53 | 40 | 80 | 11 | 376 | 148 | 424 | 137 | 285 | 141 | 1067 | 486 | 1068 | 742 | 402 | 1795 | 711 | 174 | 1035 | 700 | 492 | 3800 |
| PBANKA_1122600 | 12 | 11 | 95 | 34 | 235 | 116 | 255 | 96 | 124 | 74 | 89 | 50 | 1290 | 400 | 471 | 2260 | 1618 | 404 | 565 | 426 | 716 | 5231 |
| PBANKA_1122700 | 35 | 41 | 200 | 46 | 438 | 146 | 502 | 171 | 2314 | 1084 | 18158 | 9354 | 4330 | 9547 | 1885 | 12824 | 7461 | 1369 | 1695 | 4247 | 3660 | 25130 |
| PBANKA_1122800 | 210 | 243 | 26 | 8 | 121 | 58 | 176 | 62 | 77 | 37 | 60 | 35 | 314 | 79 | 334 | 926 | 769 | 134 | 1332 | 617 | 172 | 1069 |
| PBANKA_1122900 | 309 | 276 | 573 | 197 | 525 | 359 | 598 | 318 | 224 | 118 | 181 | 189 | 8397 | 4375 | 2813 | 12512 | 7782 | 833 | 1692 | 890 | 418 | 3814 |
| PBANKA_1123000 | 0 | 0 | 54 | 16 | 79 | 39 | 67 | 23 | 23 | 9 | 10 | 0 | 214 | 71 | 207 | 604 | 262 | 58 | 428 | 261 | 20 | 117 |
| PBANKA_1123100 | 0 | 0 | 24 | 0 | 35 | 25 | 92 | 35 | 47 | 20 | 12 | 5 | 344 | 100 | 101 | 357 | 661 | 163 | 1309 | 772 | 315 | 2077 |
| PBANKA_1123300 | 104 | 135 | 232 | 48 | 275 | 157 | 356 | 141 | 112 | 49 | 15 | 14 | 1535 | 834 | 797 | 3770 | 2249 | 280 | 3615 | 2774 | 2895 | 16781 |
| PBANKA_1123400 | 13 | 14 | 6 | 0 | 33 | 17 | 66 | 24 | 19 | 10 | 0 | 0 | 47 | 24 | 39 | 133 | 113 | 27 | 286 | 85 | 202 | 1168 |
| PBANKA_1123500 | 644 | 865 | 1201 | 311 | 4073 | 1744 | 6882 | 2540 | 3604 | 1636 | 4836 | 2851 | 8695 | 16082 | 7495 | 37796 | 68933 | 7612 | 5164 | 15612 | 2461 | 17769 |
| PBANKA_1123600 | 0 | 0 | 64 | 12 | 127 | 89 | 115 | 42 | 34 | 20 | 0 | 0 | 368 | 60 | 201 | 662 | 357 | 77 | 137 | 54 | 8 | 94 |
| PBANKA_1123800 | 148 | 225 | 10 | 0 | 41 | 12 | 36 | 6 | 23 | 6 | 5 | 5 | 445 | 89 | 10 | 33 | 234 | 106 | 2991 | 1624 | 48 | 220 |
| PBANKA_1123900 | 11 | 7 | 16 | 0 | 62 | 29 | 114 | 56 | 86 | 67 | 42 | 30 | 596 | 148 | 443 | 1725 | 1241 | 378 | 893 | 500 | 237 | 1598 |
| PBANKA_1124000 | 29 | 45 | 14 | 5 | 74 | 30 | 89 | 33 | 40 | 21 | 27 | 19 | 705 | 243 | 80 | 301 | 262 | 96 | 18924 | 15256 | 267 | 1887 |
| PBANKA_1124100 | 63 | 86 | 77 | 10 | 123 | 45 | 130 | 44 | 99 | 54 | 457 | 298 | 1379 | 861 | 179 | 1101 | 582 | 125 | 1473 | 654 | 406 | 2544 |
| PBANKA_1124200 | 0 | 0 | 90 | 21 | 354 | 142 | 778 | 314 | 443 | 183 | 69 | 31 | 780 | 363 | 299 | 1684 | 4434 | 656 | 469 | 448 | 55 | 381 |
| PBANKA_1124300 | 0 | 0 | 27 | 14 | 183 | 79 | 211 | 82 | 592 | 443 | 154 | 72 | 749 | 281 | 294 | 1251 | 1704 | 581 | 365 | 326 | 232 | 1516 |
| PBANKA_1124400 | 0 | 0 | 30 | 0 | 128 | 59 | 162 | 51 | 124 | 68 | 28 | 10 | 188 | 59 | 152 | 736 | 387 | 126 | 1171 | 705 | 75 | 482 |
| PBANKA_1124500 | 15 | 12 | 31 | 5 | 142 | 79 | 221 | 91 | 178 | 66 | 35 | 34 | 1428 | 518 | 377 | 1664 | 2757 | 538 | 2803 | 1768 | 120 | 961 |
| PBANKA_1124600 | 0 | 0 | 31 | 13 | 137 | 63 | 159 | 70 | 61 | 25 | 11 | 0 | 143 | 40 | 117 | 462 | 631 | 113 | 424 | 288 | 155 | 1103 |
| PBANKA_1124800 | 6 | 0 | 75 | 23 | 200 | 100 | 190 | 61 | 80 | 22 | 8 | 20 | 1257 | 710 | 613 | 2379 | 675 | 92 | 796 | 739 | 72 | 489 |
| PBANKA_1124900 | 0 | 0 | 113 | 30 | 295 | 181 | 301 | 165 | 75 | 57 | 0 | 0 | 299 | 94 | 363 | 1071 | 352 | 87 | 983 | 556 | 40 | 366 |
| PBANKA_1125000 | 0 | 0 | 78 | 14 | 166 | 62 | 174 | 52 | 48 | 18 | 6 | 5 | 882 | 359 | 318 | 1652 | 371 | 61 | 649 | 275 | 22 | 111 |
| PBANKA_1125100 | 146 | 186 | 260 | 102 | 1349 | 1086 | 2539 | 1216 | 755 | 443 | 46 | 41 | 20 | 15 | 16 | 16 | 70 | 20 | 83 | 52 | 185 | 1628 |
| PBANKA_1125200 | 104 | 118 | 22 | 12 | 198 | 97 | 378 | 173 | 264 | 127 | 38 | 14 | 456 | 144 | 17 | 57 | 2588 | 296 | 1679 | 1471 | 365 | 2095 |
| PBANKA_1125400 | 5 | 0 | 31 | 0 | 38 | 19 | 47 | 20 | 6 | 6 | 0 | 0 | 105 | 37 | 72 | 220 | 134 | 45 | 293 | 229 | 8 | 54 |
| PBANKA_1125500 | 110 | 89 | 86 | 28 | 211 | 166 | 318 | 157 | 69 | 58 | 19 | 16 | 846 | 341 | 521 | 2570 | 1422 | 137 | 900 | 671 | 21 | 232 |
| PBANKA_1125600 | 1268 | 1640 | 1995 | 717 | 1670 | 1386 | 2283 | 1052 | 714 | 437 | 2229 | 2297 | 9846 | 6132 | 7137 | 20828 | 4246 | 592 | 5305 | 3508 | 434 | 4821 |
| PBANKA_1125700 | 0 | 0 | 62 | 9 | 238 | 114 | 260 | 108 | 60 | 32 | 11 | 10 | 443 | 130 | 573 | 2422 | 1632 | 194 | 498 | 258 | 0 | 54 |
| PBANKA_1125900 | 380 | 546 | 377 | 125 | 1054 | 506 | 1105 | 402 | 1104 | 638 | 3966 | 2386 | 3232 | 3459 | 1136 | 5517 | 5907 | 1081 | 19020 | 24270 | 255 | 1856 |
| PBANKA_1126000 | 239 | 358 | 124 | 45 | 369 | 167 | 466 | 267 | 174 | 92 | 184 | 127 | 1783 | 843 | 734 | 3007 | 2324 | 350 | 2446 | 1676 | 324 | 2794 |
| PBANKA_1126100 | 0 | 0 | 34 | 12 | 53 | 22 | 55 | 31 | 13 | 12 | 0 | 0 | 149 | 25 | 115 | 387 | 125 | 22 | 303 | 126 | 17 | 94 |
| PBANKA_1126200 | 0 | 0 | 226 | 85 | 263 | 185 | 333 | 212 | 105 | 73 | 12 | 10 | 781 | 160 | 1334 | 2734 | 626 | 133 | 1204 | 607 | 111 | 799 |
| PBANKA_1126300 | 0 | 0 | 20 | 0 | 72 | 46 | 103 | 54 | 40 | 22 | 0 | 0 | 350 | 122 | 272 | 994 | 634 | 87 | 3478 | 2002 | 105 | 955 |
| PBANKA_1126400 | 238 | 294 | 242 | 84 | 469 | 288 | 552 | 262 | 169 | 121 | 517 | 555 | 4719 | 2108 | 1547 | 6431 | 2494 | 317 | 2382 | 1469 | 105 | 1034 |
| PBANKA_1126500 | 15 | 12 | 236 | 59 | 588 | 323 | 1059 | 468 | 433 | 222 | 343 | 190 | 5743 | 2747 | 1919 | 10166 | 7470 | 761 | 971 | 776 | 119 | 882 |
| PBANKA_1126600 | 0 | 0 | 85 | 25 | 211 | 103 | 271 | 116 | 70 | 69 | 13 | 16 | 997 | 342 | 355 | 1407 | 1004 | 147 | 589 | 340 | 106 | 981 |
| PBANKA_1126700 | 0 | 6 | 33 | 7 | 99 | 32 | 45 | 22 | 8 | 9 | 0 | 0 | 64 | 36 | 124 | 300 | 36 | 15 | 73 | 47 | 5 | 29 |
| PBANKA_1126800 | 823 | 1420 | 37 | 10 | 445 | 195 | 1298 | 421 | 4788 | 2081 | 546 | 333 | 730 | 2445 | 196 | 445 | 24030 | 4263 | 4354 | 57232 | 965 | 9123 |
| PBANKA_1126900 | 73 | 122 | 432 | 93 | 976 | 491 | 1323 | 498 | 554 | 256 | 635 | 304 | 4201 | 8365 | 2877 | 14557 | 8808 | 964 | 960 | 3330 | 204 | 1553 |
| PBANKA_1127000 | 7 | 8 | 635 | 150 | 902 | 648 | 2298 | 920 | 3096 | 1976 | 13666 | 10278 | 8589 | 4399 | 2852 | 6155 | 8418 | 2953 | 1732 | 1058 | 287 | 1242 |
| PBANKA_1127200 | 0 | 5 | 21 | 0 | 100 | 45 | 85 | 44 | 42 | 13 | 5 | 0 | 39 | 19 | 29 | 43 | 31 | 12 | 246 | 116 | 0 | 0 |
| PBANKA_1127300 | 0 | 0 | 36 | 0 | 91 | 43 | 148 | 50 | 49 | 23 | 10 | 7 | 686 | 239 | 325 | 1661 | 954 | 188 | 365 | 237 | 0 | 53 |
| PBANKA_1127400 | 0 | 0 | 112 | 14 | 195 | 97 | 184 | 47 | 57 | 33 | 38 | 18 | 1296 | 700 | 279 | 1165 | 371 | 85 | 608 | 217 | 0 | 28 |
| PBANKA_1127500 | 0 | 0 | 50 | 6 | 116 | 43 | 126 | 33 | 40 | 11 | 0 | 0 | 840 | 374 | 494 | 2722 | 541 | 119 | 975 | 862 | 27 | 251 |
| PBANKA_1127600 | 0 | 0 | 22 | 8 | 120 | 43 | 100 | 42 | 28 | 14 | 7 | 0 | 231 | 99 | 138 | 537 | 158 | 15 | 52 | 35 | 0 | 37 |
| PBANKA_1127700 | 0 | 0 | 94 | 27 | 171 | 130 | 371 | 150 | 125 | 60 | 121 | 101 | 4318 | 2157 | 1559 | 5597 | 1548 | 212 | 379 | 314 | 9 | 57 |
| PBANKA_1127800 | 0 | 0 | 145 | 38 | 446 | 268 | 654 | 302 | 236 | 147 | 11 | 12 | 1569 | 491 | 1036 | 5090 | 3096 | 373 | 2790 | 1765 | 92 | 961 |
| PBANKA_1127900 | 18 | 22 | 97 | 24 | 279 | 139 | 367 | 117 | 163 | 88 | 138 | 91 | 1226 | 1281 | 412 | 2191 | 428 | 72 | 245 | 171 | 12 | 68 |
| PBANKA_1128100 | 5806 | 7562 | 1356 | 470 | 2239 | 1400 | 2270 | 1064 | 1328 | 967 | 246 | 149 | 359 | 155 | 210 | 843 | 3071 | 487 | 2317 | 3452 | 481 | 3581 |
| PBANKA_1128200 | 7 | 0 | 80 | 24 | 78 | 29 | 71 | 27 | 45 | 25 | 539 | 506 | 3495 | 2783 | 363 | 2676 | 274 | 69 | 903 | 706 | 16 | 171 |
| PBANKA_1128300 | 22 | 37 | 23 | 0 | 151 | 84 | 113 | 49 | 39 | 27 | 79 | 74 | 1509 | 588 | 803 | 4287 | 498 | 60 | 320 | 244 | 30 | 167 |
| PBANKA_1128400 | 14 | 14 | 21 | 10 | 43 | 24 | 36 | 22 | 22 | 15 | 39 | 14 | 469 | 271 | 88 | 341 | 46 | 14 | 69 | 45 | 0 | 23 |
| PBANKA_1128500 | 647 | 913 | 715 | 136 | 4669 | 1456 | 11636 | 4213 | 4695 | 1904 | 1410 | 803 | 2592 | 10992 | 4492 | 23842 | 30666 | 2962 | 661 | 8471 | 448 | 3107 |
| PBANKA_1128600 | 153 | 239 | 95 | 23 | 332 | 134 | 482 | 162 | 167 | 95 | 41 | 15 | 596 | 279 | 445 | 2202 | 1369 | 203 | 1385 | 1551 | 235 | 1763 |
| PBANKA_1128700 | 98 | 131 | 34 | 6 | 78 | 35 | 183 | 54 | 49 | 31 | 18 | 0 | 779 | 353 | 282 | 1172 | 1365 | 242 | 1136 | 530 | 114 | 986 |
| PBANKA_1128900 | 165 | 251 | 12 | 0 | 67 | 39 | 100 | 17 | 311 | 206 | 76 | 47 | 224 | 99 | 84 | 217 | 578 | 173 | 1641 | 1234 | 2763 | 16093 |
| PBANKA_1129000 | 31 | 31 | 5 | 0 | 24 | 10 | 9 | 0 | 35 | 42 | 5 | 9 | 252 | 91 | 8 | 21 | 101 | 63 | 6225 | 2509 | 214 | 1284 |
| PBANKA_1129100 | 6 | 10 | 208 | 61 | 745 | 311 | 914 | 307 | 354 | 192 | 25 | 20 | 666 | 363 | 1076 | 4136 | 5235 | 580 | 3348 | 4553 | 26 | 228 |
| PBANKA_1129200 | 26 | 48 | 5 | 0 | 18 | 0 | 65 | 31 | 66 | 41 | 8 | 8 | 142 | 69 | 23 | 82 | 1367 | 368 | 852 | 347 | 62 | 517 |
| PBANKA_1129300 | 14 | 0 | 39 | 11 | 62 | 24 | 54 | 21 | 51 | 19 | 381 | 262 | 981 | 605 | 111 | 663 | 156 | 22 | 516 | 185 | 64 | 384 |
| PBANKA_1129400 | 243 | 357 | 103 | 18 | 333 | 121 | 376 | 146 | 158 | 74 | 20 | 9 | 651 | 204 | 929 | 3793 | 3610 | 606 | 1501 | 1309 | 137 | 1144 |
| PBANKA_1129500 | 7 | 16 | 35 | 0 | 131 | 44 | 104 | 29 | 88 | 29 | 33 | 28 | 982 | 331 | 218 | 1320 | 230 | 50 | 968 | 403 | 519 | 2556 |
| PBANKA_1129700 | 125 | 143 | 59 | 15 | 221 | 81 | 292 | 81 | 255 | 117 | 1608 | 946 | 738 | 599 | 191 | 1058 | 735 | 117 | 480 | 335 | 478 | 3289 |
| PBANKA_1129800 | 53 | 49 | 8 | 0 | 62 | 18 | 137 | 51 | 209 | 76 | 107 | 28 | 599 | 308 | 55 | 100 | 2935 | 594 | 11567 | 5808 | 1030 | 8583 |
| PBANKA_1129900 | 7 | 6 | 63 | 11 | 425 | 131 | 460 | 157 | 239 | 98 | 95 | 63 | 1503 | 1487 | 1589 | 7968 | 4636 | 604 | 2038 | 3488 | 118 | 478 |
| PBANKA_1130000 | 0 | 0 | 117 | 13 | 413 | 155 | 460 | 98 | 148 | 75 | 57 | 40 | 1465 | 1402 | 793 | 4523 | 1871 | 250 | 668 | 1097 | 212 | 1451 |
| PBANKA_1130100 | 0 | 0 | 50 | 14 | 173 | 64 | 193 | 70 | 48 | 23 | 6 | 0 | 426 | 135 | 301 | 1018 | 311 | 43 | 1047 | 613 | 588 | 5382 |
| PBANKA_1130200 | 139 | 180 | 186 | 53 | 384 | 198 | 478 | 223 | 227 | 150 | 71 | 14 | 983 | 152 | 387 | 682 | 1428 | 653 | 888 | 312 | 101 | 684 |
| PBANKA_1130300 | 10 | 17 | 94 | 32 | 71 | 77 | 70 | 24 | 38 | 13 | 0 | 0 | 830 | 244 | 205 | 879 | 193 | 46 | 327 | 148 | 166 | 1217 |
| PBANKA_1130400 | 0 | 0 | 201 | 79 | 375 | 256 | 459 | 250 | 160 | 79 | 30 | 0 | 1567 | 304 | 1709 | 6105 | 2747 | 376 | 1501 | 750 | 218 | 1813 |
| PBANKA_1130500 | 0 | 7 | 273 | 93 | 593 | 393 | 652 | 347 | 202 | 124 | 30 | 15 | 979 | 181 | 1086 | 3766 | 1325 | 242 | 806 | 449 | 184 | 1429 |
| PBANKA_1130600 | 51 | 60 | 18 | 0 | 84 | 28 | 45 | 17 | 40 | 17 | 12 | 0 | 104 | 35 | 69 | 184 | 80 | 27 | 1257 | 522 | 53 | 256 |
| PBANKA_1130700 | 33 | 53 | 113 | 22 | 159 | 81 | 142 | 76 | 63 | 31 | 8 | 10 | 273 | 74 | 312 | 1578 | 475 | 71 | 711 | 438 | 116 | 785 |

| | | | | | | | | | | | | | | | | | | | | | | |
|---|---|---|---|---|---|---|---|---|---|---|---|---|---|---|---|---|---|---|---|---|---|---|
| PBANKA_1130800 | 32 | 36 | 90 | 7 | 199 | 126 | 439 | 158 | 233 | 139 | 57 | 28 | 1286 | 686 | 509 | 2649 | 4205 | 706 | 796 | 1015 | 2136 | 15236 |
| PBANKA_1130900 | 68 | 88 | 389 | 71 | 679 | 242 | 798 | 260 | 600 | 286 | 5638 | 3246 | 6254 | 9990 | 1511 | 8599 | 5060 | 869 | 5428 | 15086 | 1307 | 8701 |
| PBANKA_1131000 | 145 | 163 | 32 | 26 | 148 | 134 | 241 | 110 | 81 | 70 | 25 | 9 | 525 | 174 | 152 | 611 | 602 | 175 | 557 | 196 | 166 | 926 |
| PBANKA_1131100 | 14 | 6 | 58 | 23 | 130 | 59 | 192 | 55 | 72 | 23 | 20 | 10 | 1545 | 674 | 352 | 2125 | 754 | 97 | 1374 | 733 | 700 | 4784 |
| PBANKA_1131200 | 5 | 5 | 148 | 30 | 170 | 89 | 185 | 84 | 122 | 54 | 261 | 189 | 945 | 462 | 245 | 943 | 371 | 180 | 392 | 204 | 115 | 640 |
| PBANKA_1131300 | 87 | 79 | 11 | 0 | 24 | 31 | 21 | 11 | 24 | 16 | 0 | 0 | 29 | 11 | 43 | 184 | 48 | 7 | 403 | 194 | 994 | 6149 |
| PBANKA_1131400 | 113 | 183 | 26 | 11 | 164 | 85 | 222 | 79 | 79 | 48 | 25 | 5 | 558 | 297 | 28 | 85 | 817 | 398 | 12551 | 6443 | 1444 | 9463 |
| PBANKA_1131500 | 67 | 108 | 50 | 13 | 220 | 93 | 461 | 191 | 196 | 99 | 31 | 9 | 752 | 280 | 394 | 1906 | 2766 | 406 | 3202 | 5040 | 912 | 6537 |
| PBANKA_1131600 | 22 | 28 | 222 | 61 | 833 | 477 | 1278 | 476 | 632 | 318 | 95 | 27 | 1131 | 420 | 1060 | 3440 | 8155 | 948 | 3034 | 2962 | 384 | 3673 |
| PBANKA_1131700 | 66 | 107 | 66 | 9 | 135 | 73 | 162 | 65 | 79 | 43 | 12 | 8 | 604 | 317 | 268 | 1213 | 673 | 139 | 1251 | 649 | 40 | 265 |
| PBANKA_1131800 | 19 | 19 | 22 | 0 | 45 | 21 | 56 | 19 | 31 | 17 | 56 | 34 | 408 | 222 | 124 | 499 | 401 | 92 | 492 | 216 | 78 | 417 |
| PBANKA_1131900 | 20 | 19 | 154 | 31 | 251 | 84 | 209 | 76 | 108 | 71 | 263 | 140 | 5162 | 2612 | 872 | 4998 | 1611 | 247 | 2455 | 1991 | 1887 | 10981 |
| PBANKA_1132000 | 30 | 10 | 64 | 8 | 252 | 113 | 296 | 91 | 194 | 106 | 353 | 160 | 1323 | 933 | 529 | 3741 | 1711 | 257 | 1395 | 1942 | 201 | 1472 |
| PBANKA_1132100 | 29 | 27 | 49 | 14 | 80 | 74 | 76 | 42 | 24 | 18 | 39 | 16 | 147 | 149 | 54 | 121 | 117 | 29 | 168 | 76 | 8 | 98 |
| PBANKA_1132200 | 0 | 0 | 60 | 12 | 121 | 82 | 169 | 72 | 64 | 30 | 29 | 11 | 1238 | 521 | 239 | 1186 | 772 | 133 | 363 | 267 | 175 | 1257 |
| PBANKA_1132400 | 6 | 6 | 67 | 15 | 271 | 134 | 360 | 124 | 167 | 62 | 37 | 22 | 452 | 264 | 402 | 1502 | 1662 | 245 | 331 | 580 | 15 | 199 |
| PBANKA_1132500 | 27 | 30 | 109 | 38 | 90 | 49 | 73 | 28 | 17 | 9 | 10 | 0 | 966 | 185 | 199 | 597 | 83 | 38 | 255 | 117 | 49 | 342 |
| PBANKA_1132600 | 28 | 28 | 147 | 22 | 178 | 164 | 232 | 105 | 86 | 46 | 12 | 8 | 447 | 110 | 311 | 996 | 738 | 181 | 709 | 381 | 56 | 435 |
| PBANKA_1132700 | 17 | 33 | 87 | 6 | 301 | 133 | 306 | 87 | 133 | 57 | 22 | 0 | 355 | 102 | 301 | 1021 | 1327 | 188 | 1338 | 1348 | 51 | 405 |
| PBANKA_1133000 | 0 | 0 | 11 | 8 | 13 | 12 | 31 | 5 | 7 | 8 | 0 | 0 | 273 | 76 | 113 | 247 | 80 | 20 | 228 | 88 | 7 | 30 |
| PBANKA_1133100 | 0 | 0 | 6 | 0 | 28 | 16 | 178 | 89 | 1709 | 821 | 417 | 230 | 521 | 1970 | 121 | 543 | 12177 | 2384 | 4362 | 48489 | 352 | 2868 |
| PBANKA_1133200 | 0 | 5 | 12 | 5 | 110 | 79 | 98 | 48 | 34 | 30 | 5 | 0 | 379 | 101 | 373 | 947 | 275 | 64 | 1334 | 511 | 721 | 4603 |
| PBANKA_1133300 | 67 | 22 | 11316 | 4649 | 8128 | 6457 | 7737 | 3471 | 2250 | 1327 | 3672 | 2640 | 36496 | 10497 | 14200 | 50075 | 16219 | 2377 | 7060 | 3169 | 529 | 6327 |
| PBANKA_1133400 | 0 | 5 | 711 | 275 | 509 | 456 | 433 | 251 | 132 | 98 | 233 | 179 | 10132 | 2180 | 12611 | 9664 | 3277 | 1784 | 2713 | 731 | 36 | 335 |
| PBANKA_1133500 | 5 | 13 | 75 | 26 | 126 | 47 | 88 | 47 | 28 | 17 | 5 | 0 | 1210 | 434 | 376 | 1913 | 1448 | 175 | 666 | 445 | 112 | 773 |
| PBANKA_1133700 | 7 | 12 | 360 | 91 | 858 | 326 | 988 | 398 | 418 | 205 | 85 | 22 | 2363 | 2048 | 1070 | 5833 | 7021 | 785 | 1012 | 1880 | 136 | 1020 |
| PBANKA_1133800 | 238 | 471 | 28 | 9 | 200 | 101 | 489 | 200 | 232 | 115 | 29 | 10 | 452 | 163 | 138 | 559 | 1755 | 269 | 1466 | 1241 | 337 | 2858 |
| PBANKA_1133900 | 5 | 0 | 25 | 0 | 27 | 16 | 26 | 20 | 6 | 0 | 0 | 0 | 91 | 9 | 80 | 302 | 74 | 17 | 50 | 24 | 0 | 29 |
| PBANKA_1134000 | 9 | 17 | 216 | 58 | 369 | 124 | 331 | 118 | 153 | 57 | 733 | 537 | 3646 | 3279 | 852 | 4702 | 1783 | 203 | 431 | 792 | 69 | 388 |
| PBANKA_1134100 | 50 | 141 | 872 | 222 | 1034 | 852 | 1308 | 556 | 353 | 194 | 133 | 102 | 6488 | 1943 | 3683 | 15809 | 3521 | 390 | 1874 | 1162 | 126 | 1185 |
| PBANKA_1134200 | 0 | 0 | 36 | 8 | 77 | 88 | 65 | 40 | 30 | 23 | 10 | 0 | 192 | 25 | 425 | 1690 | 230 | 31 | 34 | 23 | 0 | 9 |
| PBANKA_1134300 | 37 | 68 | 106 | 17 | 241 | 104 | 468 | 160 | 180 | 83 | 78 | 25 | 705 | 466 | 402 | 2005 | 3337 | 348 | 589 | 1422 | 70 | 361 |
| PBANKA_1134400 | 79 | 124 | 256 | 55 | 375 | 97 | 261 | 74 | 241 | 94 | 2693 | 1498 | 7643 | 9579 | 1343 | 12060 | 2173 | 301 | 1765 | 5024 | 608 | 3549 |
| PBANKA_1134500 | 9 | 0 | 72 | 13 | 249 | 70 | 193 | 69 | 113 | 44 | 164 | 57 | 856 | 651 | 582 | 3172 | 1070 | 162 | 1095 | 802 | 74 | 599 |
| PBANKA_1134600 | 9 | 16 | 26 | 10 | 62 | 25 | 105 | 33 | 39 | 17 | 0 | 0 | 103 | 17 | 72 | 206 | 523 | 114 | 258 | 77 | 16 | 159 |
| PBANKA_1134700 | 35 | 26 | 55 | 9 | 90 | 33 | 208 | 101 | 202 | 77 | 22 | 8 | 335 | 67 | 132 | 309 | 1554 | 464 | 770 | 267 | 20 | 205 |
| PBANKA_1134800 | 21 | 38 | 121 | 34 | 107 | 70 | 103 | 32 | 64 | 29 | 388 | 310 | 1264 | 856 | 222 | 1276 | 222 | 47 | 413 | 392 | 30 | 231 |
| PBANKA_1134900 | 35 | 67 | 7 | 0 | 60 | 33 | 61 | 38 | 104 | 68 | 15 | 13 | 4645 | 4591 | 126 | 543 | 2402 | 895 | 84424 | 2E+05 | 2866 | 20811 |
| PBANKA_1135000 | 0 | 0 | 100 | 18 | 153 | 84 | 230 | 108 | 80 | 26 | 75 | 33 | 2577 | 1596 | 742 | 2707 | 1469 | 252 | 728 | 401 | 78 | 637 |
| PBANKA_1135100 | 46 | 70 | 1374 | 421 | 1032 | 623 | 1161 | 571 | 365 | 237 | 2598 | 2227 | 6458 | 2926 | 1693 | 4641 | 1848 | 446 | 1548 | 544 | 90 | 693 |
| PBANKA_1135200 | 0 | 0 | 108 | 57 | 177 | 129 | 240 | 153 | 72 | 49 | 89 | 44 | 1243 | 519 | 290 | 1117 | 377 | 57 | 933 | 376 | 48 | 535 |
| PBANKA_1135300 | 251 | 281 | 516 | 148 | 748 | 279 | 491 | 179 | 507 | 200 | 2263 | 1421 | 3257 | 5908 | 1317 | 5771 | 1370 | 179 | 789 | 1641 | 263 | 1477 |
| PBANKA_1135400 | 0 | 0 | 64 | 20 | 123 | 60 | 95 | 28 | 29 | 19 | 0 | 11 | 1169 | 453 | 529 | 2504 | 295 | 66 | 1022 | 603 | 26 | 211 |
| PBANKA_1135500 | 7 | 7 | 39 | 16 | 137 | 90 | 294 | 156 | 165 | 123 | 35 | 20 | 531 | 155 | 266 | 689 | 1144 | 352 | 1453 | 663 | 55 | 469 |
| PBANKA_1135600 | 13 | 15 | 201 | 35 | 376 | 155 | 392 | 129 | 154 | 62 | 38 | 11 | 1304 | 767 | 969 | 4368 | 1480 | 240 | 340 | 257 | 27 | 166 |
| PBANKA_1135700 | 216 | 332 | 888 | 279 | 1702 | 873 | 1798 | 714 | 766 | 446 | 7820 | 5159 | 9219 | 7909 | 2531 | 9124 | 4720 | 749 | 3207 | 1762 | 440 | 3671 |
| PBANKA_1135800 | 52 | 41 | 20 | 0 | 266 | 124 | 227 | 81 | 59 | 19 | 7 | 0 | 125 | 44 | 54 | 225 | 105 | 33 | 1185 | 1306 | 68 | 423 |
| PBANKA_1135900 | 23 | 28 | 7 | 0 | 98 | 36 | 222 | 79 | 244 | 99 | 44 | 22 | 270 | 267 | 90 | 412 | 5602 | 687 | 761 | 1674 | 36 | 289 |
| PBANKA_1136100 | 10 | 11 | 102 | 31 | 224 | 123 | 316 | 123 | 219 | 129 | 67 | 57 | 1864 | 1668 | 986 | 5031 | 5327 | 772 | 1481 | 4335 | 59 | 506 |
| PBANKA_1136200 | 23 | 34 | 175 | 46 | 248 | 204 | 308 | 157 | 95 | 60 | 12 | 5 | 502 | 141 | 744 | 2820 | 2726 | 473 | 774 | 653 | 25 | 215 |
| PBANKA_1136300 | 0 | 0 | 71 | 22 | 122 | 64 | 168 | 100 | 136 | 87 | 447 | 420 | 1102 | 641 | 189 | 472 | 1402 | 327 | 1078 | 606 | 17 | 116 |
| PBANKA_1136400 | 0 | 0 | 80 | 25 | 155 | 109 | 173 | 120 | 62 | 46 | 6 | 6 | 364 | 98 | 299 | 756 | 701 | 162 | 624 | 224 | 48 | 346 |
| PBANKA_1136500 | 0 | 0 | 20 | 7 | 93 | 38 | 105 | 34 | 51 | 16 | 0 | 0 | 170 | 61 | 127 | 378 | 321 | 58 | 271 | 133 | 13 | 93 |
| PBANKA_1136600 | 0 | 0 | 67 | 8 | 142 | 53 | 107 | 40 | 48 | 20 | 28 | 21 | 396 | 198 | 189 | 679 | 243 | 53 | 520 | 267 | 28 | 187 |
| PBANKA_1136700 | 15 | 25 | 12 | 0 | 93 | 64 | 134 | 86 | 169 | 80 | 39 | 17 | 387 | 354 | 131 | 375 | 1160 | 273 | 4990 | 8611 | 156 | 1361 |
| PBANKA_1136900 | 82 | 132 | 60 | 14 | 188 | 73 | 338 | 142 | 257 | 126 | 656 | 407 | 2798 | 1596 | 496 | 2504 | 5570 | 754 | 2533 | 1742 | 34 | 351 |
| PBANKA_1137000 | 630 | 1004 | 1188 | 351 | 1537 | 767 | 1497 | 593 | 515 | 281 | 376 | 248 | 5001 | 3152 | 4065 | 17052 | 6629 | 930 | 2454 | 4067 | 631 | 5346 |
| PBANKA_1137100 | 0 | 0 | 74 | 12 | 192 | 102 | 133 | 56 | 66 | 26 | 336 | 170 | 1366 | 764 | 406 | 1873 | 301 | 27 | 257 | 166 | 0 | 29 |
| PBANKA_1137200 | 17 | 43 | 232 | 80 | 850 | 580 | 1205 | 591 | 731 | 455 | 783 | 632 | 4767 | 2008 | 1653 | 6339 | 6309 | 1341 | 2071 | 1254 | 98 | 1107 |
| PBANKA_1137300 | 441 | 650 | 187 | 50 | 641 | 373 | 973 | 436 | 634 | 297 | 147 | 101 | 2780 | 3846 | 1677 | 8942 | 11443 | 2202 | 2721 | 7204 | 316 | 2333 |
| PBANKA_1137400 | 0 | 0 | 34 | 6 | 272 | 196 | 252 | 98 | 39 | 35 | 10 | 0 | 375 | 111 | 19 | 67 | 226 | 92 | 6529 | 2043 | 58 | 413 |
| PBANKA_1137500 | 0 | 0 | 7 | 0 | 40 | 15 | 24 | 8 | 20 | 0 | 7 | 0 | 71 | 17 | 0 | 16 | 68 | 20 | 1054 | 302 | 0 | 36 |
| PBANKA_1137600 | 0 | 0 | 17 | 7 | 83 | 27 | 46 | 33 | 31 | 22 | 0 | 0 | 87 | 14 | 51 | 205 | 235 | 53 | 320 | 125 | 42 | 449 |
| PBANKA_1137700 | 6 | 0 | 85 | 43 | 351 | 264 | 590 | 339 | 381 | 311 | 36 | 31 | 515 | 137 | 301 | 911 | 1987 | 514 | 3671 | 1768 | 165 | 1519 |
| PBANKA_1137800 | 0 | 0 | 78 | 15 | 932 | 677 | 1916 | 882 | 987 | 439 | 78 | 38 | 1570 | 761 | 5059 | 13887 | 24347 | 2496 | 997 | 1494 | 43 | 398 |
| PBANKA_1137900 | 0 | 6 | 327 | 93 | 657 | 353 | 1028 | 542 | 328 | 203 | 32 | 20 | 994 | 143 | 1654 | 4035 | 3812 | 793 | 4108 | 1331 | 28 | 342 |
| PBANKA_1138000 | 22 | 28 | 87 | 32 | 481 | 391 | 865 | 621 | 501 | 396 | 94 | 60 | 273 | 63 | 246 | 902 | 1937 | 340 | 2019 | 943 | 87 | 1018 |
| PBANKA_1138100 | 457 | 600 | 16 | 0 | 57 | 37 | 60 | 25 | 28 | 24 | 0 | 0 | 70 | 24 | 32 | 94 | 28 | 20 | 930 | 372 | 1678 | 10466 |
| PBANKA_1138200 | 0 | 0 | 166 | 56 | 209 | 138 | 227 | 158 | 129 | 66 | 62 | 59 | 3171 | 1536 | 718 | 3589 | 1359 | 256 | 19130 | 15631 | 641 | 7121 |
| PBANKA_1138300 | 5 | 0 | 73 | 14 | 114 | 57 | 202 | 96 | 70 | 39 | 14 | 11 | 1594 | 504 | 310 | 1212 | 1496 | 249 | 1205 | 759 | 95 | 791 |
| PBANKA_1138400 | 30 | 44 | 129 | 40 | 335 | 144 | 470 | 191 | 395 | 141 | 299 | 130 | 3306 | 4372 | 1199 | 7242 | 10226 | 1536 | 2915 | 6410 | 504 | 3193 |
| PBANKA_1138500 | 80 | 122 | 29 | 13 | 104 | 40 | 64 | 29 | 46 | 15 | 10 | 0 | 71 | 57 | 174 | 815 | 190 | 27 | 342 | 193 | 117 | 839 |
| PBANKA_1138600 | 8 | 7 | 5 | 0 | 17 | 16 | 25 | 8 | 15 | 6 | 0 | 0 | 1145 | 698 | 88 | 450 | 450 | 188 | 27968 | 21214 | 26 | 245 |
| PBANKA_1138700 | 208 | 332 | 350 | 93 | 671 | 290 | 784 | 246 | 220 | 93 | 64 | 29 | 673 | 679 | 1779 | 11120 | 3158 | 288 | 396 | 620 | 39 | 289 |
| PBANKA_1138800 | 15 | 27 | 11 | 0 | 60 | 33 | 146 | 42 | 294 | 158 | 36 | 13 | 340 | 176 | 62 | 357 | 4102 | 740 | 1458 | 2319 | 432 | 3308 |
| PBANKA_1139000 | 18 | 36 | 46 | 14 | 181 | 63 | 119 | 37 | 66 | 16 | 11 | 8 | 905 | 489 | 467 | 2860 | 811 | 106 | 1166 | 754 | 10 | 46 |
| PBANKA_1139100 | 0 | 0 | 88 | 22 | 217 | 124 | 257 | 97 | 91 | 40 | 23 | 8 | 1437 | 752 | 453 | 2726 | 1241 | 138 | 648 | 1085 | 19 | 131 |
| PBANKA_1139200 | 5 | 9 | 70 | 13 | 104 | 53 | 91 | 33 | 37 | 14 | 6 | 0 | 296 | 128 | 307 | 1085 | 750 | 142 | 460 | 272 | 10 | 60 |
| PBANKA_1139300 | 0 | 15 | 45 | 5 | 104 | 22 | 88 | 30 | 22 | 6 | 0 | 0 | 184 | 31 | 178 | 563 | 308 | 41 | 291 | 132 | 6 | 52 |
| PBANKA_1139400 | 0 | 0 | 53 | 11 | 99 | 53 | 188 | 61 | 38 | 23 | 5 | 5 | 844 | 221 | 356 | 1738 | 592 | 76 | 589 | 389 | 5 | 61 |
| PBANKA_1139500 | 0 | 0 | 28 | 11 | 76 | 35 | 132 | 49 | 38 | 10 | 9 | 0 | 1050 | 264 | 270 | 1314 | 548 | 90 | 1661 | 1024 | 29 | 155 |
| PBANKA_1139600 | 0 | 0 | 13 | 0 | 59 | 30 | 375 | 135 | 124 | 68 | 0 | 6 | 72 | 7 | 88 | 220 | 109 | 25 | 71 | 24 | 0 | 12 |
| PBANKA_1139700 | 14 | 13 | 62 | 18 | 248 | 93 | 667 | 299 | 226 | 121 | 26 | 12 | 798 | 176 | 441 | 1663 | 1158 | 195 | 486 | 464 | 20 | 161 |
| PBANKA_1139900 | 62 | 57 | 130 | 21 | 351 | 186 | 635 | 293 | 376 | 168 | 352 | 273 | 2767 | 980 | 886 | 3603 | 3581 | 507 | 957 | 545 | 62 | 552 |
| PBANKA_1140000 | 17 | 18 | 49 | 8 | 151 | 61 | 196 | 87 | 122 | 64 | 46 | 32 | 546 | 213 | 191 | 976 | 1795 | 326 | 742 | 522 | 51 | 437 |
| PBANKA_1140100 | 1130 | 1279 | 183 | 44 | 821 | 352 | 1181 | 384 | 645 | 284 | 284 | 136 | 2414 | 1780 | 751 | 2961 | 5940 | 932 | 1356 | 1885 | 637 | 5202 |
| PBANKA_1140200 | 156 | 208 | 35 | 12 | 185 | 81 | 373 | 122 | 179 | 84 | 150 | 128 | 1383 | 1009 | 331 | 1894 | 2846 | 374 | 1344 | 1164 | 97 | 978 |
| PBANKA_1140300 | 228 | 318 | 270 | 62 | 720 | 214 | 851 | 272 | 416 | 127 | 705 | 324 | 3278 | 7589 | 2039 | 11592 | 6602 | 835 | 2116 | 6136 | 247 | 1691 |
| PBANKA_1140400 | 0 | 0 | 35 | 8 | 83 | 41 | 132 | 39 | 46 | 14 | 5 | 11 | 696 | 308 | 528 | 2017 | 754 | 127 | 340 | 305 | 36 | 291 |
| PBANKA_1140500 | 0 | 0 | 78 | 18 | 287 | 121 | 353 | 119 | 129 | 51 | 20 | 19 | 645 | 377 | 557 | 2799 | 1879 | 210 | 622 | 958 | 83 | 515 |

| | | | | | | | | | | | | | | | | | | | | | | |
|---|---|---|---|---|---|---|---|---|---|---|---|---|---|---|---|---|---|---|---|---|---|---|
| PBANKA_1140600 | 112 | 190 | 126 | 38 | 168 | 128 | 165 | 74 | 83 | 34 | 297 | 289 | 1160 | 566 | 455 | 867 | 317 | 98 | 780 | 288 | 157 | 898 |
| PBANKA_1140700 | 0 | 11 | 90 | 23 | 220 | 105 | 683 | 270 | 387 | 224 | 43 | 25 | 99 | 25 | 73 | 166 | 118 | 23 | 86 | 51 | 0 | 27 |
| PBANKA_1140800 | 23 | 33 | 79 | 13 | 312 | 139 | 1177 | 529 | 644 | 337 | 62 | 42 | 1358 | 483 | 473 | 2432 | 1128 | 189 | 276 | 217 | 31 | 116 |
| PBANKA_1140900 | 0 | 0 | 15 | 24 | 62 | 54 | 151 | 78 | 81 | 35 | 0 | 0 | 158 | 55 | 176 | 338 | 144 | 62 | 979 | 454 | 76 | 473 |
| PBANKA_1141000 | 0 | 0 | 40 | 6 | 45 | 28 | 45 | 25 | 13 | 8 | 0 | 0 | 94 | 14 | 52 | 111 | 83 | 32 | 187 | 55 | 0 | 42 |
| PBANKA_1141100 | 18 | 15 | 41 | 9 | 66 | 28 | 175 | 75 | 169 | 80 | 1798 | 1005 | 978 | 637 | 140 | 678 | 465 | 98 | 558 | 410 | 95 | 650 |
| PBANKA_1141200 | 0 | 0 | 12 | 0 | 51 | 21 | 37 | 11 | 21 | 10 | 8 | 0 | 980 | 475 | 134 | 886 | 171 | 32 | 342 | 151 | 16 | 107 |
| PBANKA_1141300 | 0 | 0 | 67 | 11 | 127 | 52 | 146 | 51 | 34 | 21 | 31 | 35 | 464 | 212 | 300 | 1220 | 395 | 50 | 111 | 125 | 21 | 170 |
| PBANKA_1141400 | 36 | 53 | 10 | 0 | 47 | 15 | 70 | 36 | 76 | 34 | 15 | 7 | 435 | 159 | 71 | 244 | 339 | 175 | 3061 | 2068 | 435 | 2525 |
| PBANKA_1141500 | 266 | 334 | 46 | 17 | 202 | 78 | 389 | 205 | 727 | 466 | 87 | 50 | 1990 | 419 | 545 | 1923 | 4871 | 1584 | 3285 | 1622 | 764 | 5441 |
| PBANKA_1141600 | 0 | 0 | 11 | 0 | 70 | 27 | 113 | 38 | 178 | 126 | 68 | 34 | 813 | 268 | 196 | 1006 | 761 | 216 | 982 | 739 | 1077 | 6722 |
| PBANKA_1141700 | 578 | 589 | 1123 | 275 | 827 | 564 | 947 | 455 | 678 | 522 | 1762 | 1255 | 6653 | 2683 | 2259 | 5029 | 3262 | 1329 | 3347 | 1283 | 383 | 2931 |
| PBANKA_1141800 | 6 | 8 | 32 | 11 | 82 | 54 | 66 | 25 | 29 | 18 | 12 | 8 | 531 | 139 | 71 | 345 | 70 | 25 | 105 | 36 | 81 | 430 |
| PBANKA_1142000 | 0 | 0 | 35 | 14 | 39 | 18 | 48 | 18 | 22 | 6 | 0 | 0 | 92 | 56 | 65 | 232 | 63 | 24 | 863 | 376 | 35 | 168 |
| PBANKA_1142100 | 115 | 201 | 317 | 44 | 726 | 256 | 909 | 354 | 581 | 208 | 554 | 301 | 2722 | 2581 | 1128 | 4690 | 5036 | 798 | 1378 | 2393 | 189 | 1387 |
| PBANKA_1142300 | 8 | 9 | 25 | 9 | 179 | 60 | 228 | 91 | 164 | 67 | 708 | 352 | 3291 | 2749 | 990 | 4403 | 3991 | 690 | 1010 | 1133 | 183 | 1230 |
| PBANKA_1142400 | 0 | 0 | 37 | 9 | 115 | 71 | 304 | 124 | 120 | 71 | 23 | 5 | 594 | 82 | 182 | 523 | 1952 | 410 | 2524 | 1137 | 102 | 816 |
| PBANKA_1142500 | 124 | 160 | 120 | 23 | 260 | 114 | 294 | 109 | 165 | 54 | 47 | 7 | 1912 | 1053 | 345 | 1558 | 1579 | 255 | 1511 | 885 | 297 | 2301 |
| PBANKA_1142600 | 5 | 0 | 16 | 11 | 56 | 33 | 120 | 45 | 33 | 18 | 7 | 0 | 419 | 139 | 121 | 645 | 339 | 62 | 1364 | 781 | 23 | 122 |
| PBANKA_1142700 | 0 | 0 | 89 | 36 | 104 | 66 | 316 | 109 | 118 | 80 | 116 | 78 | 1208 | 812 | 344 | 1112 | 2013 | 505 | 1984 | 1151 | 112 | 612 |
| PBANKA_1142800 | 148 | 208 | 487 | 148 | 865 | 414 | 900 | 347 | 313 | 170 | 375 | 226 | 7156 | 4424 | 2555 | 16165 | 5502 | 702 | 4095 | 3478 | 285 | 1780 |
| PBANKA_1142900 | 7 | 16 | 161 | 38 | 327 | 212 | 379 | 232 | 135 | 125 | 36 | 26 | 513 | 161 | 290 | 637 | 475 | 121 | 918 | 409 | 48 | 321 |
| PBANKA_1143000 | 0 | 5 | 99 | 18 | 202 | 78 | 183 | 79 | 82 | 28 | 20 | 20 | 1272 | 839 | 577 | 3554 | 1570 | 115 | 705 | 964 | 42 | 484 |
| PBANKA_1143100 | 0 | 0 | 8 | 0 | 14 | 6 | 10 | 7 | 15 | 0 | 0 | 0 | 197 | 77 | 26 | 77 | 95 | 45 | 1374 | 705 | 16 | 98 |
| PBANKA_1143300 | 146 | 228 | 165 | 40 | 286 | 110 | 309 | 87 | 220 | 91 | 1026 | 654 | 1741 | 1548 | 354 | 1829 | 1047 | 203 | 1289 | 841 | 593 | 3594 |
| PBANKA_1143400 | 18 | 26 | 45 | 19 | 156 | 66 | 181 | 60 | 81 | 31 | 12 | 10 | 485 | 230 | 21 | 83 | 533 | 131 | 10246 | 6440 | 1272 | 6518 |
| PBANKA_1143500 | 21 | 32 | 128 | 39 | 128 | 68 | 100 | 31 | 64 | 26 | 698 | 530 | 901 | 617 | 141 | 534 | 244 | 53 | 459 | 288 | 44 | 355 |
| PBANKA_1143600 | 302 | 504 | 766 | 206 | 1086 | 472 | 1265 | 407 | 508 | 243 | 680 | 491 | 2363 | 2381 | 1931 | 9524 | 3072 | 358 | 479 | 1155 | 307 | 2839 |
| PBANKA_1143700 | 0 | 6 | 9 | 0 | 13 | 11 | 38 | 15 | 28 | 10 | 9 | 0 | 136 | 175 | 130 | 522 | 132 | 35 | 731 | 3452 | 37662 | 3E+05 |
| PBANKA_1143800 | 8 | 18 | 32 | 0 | 41 | 19 | 101 | 35 | 42 | 26 | 11 | 0 | 902 | 295 | 130 | 304 | 1059 | 333 | 11210 | 8861 | 3057 | 23989 |
| PBANKA_1143900 | 0 | 0 | 9 | 0 | 22 | 8 | 47 | 18 | 30 | 16 | 7 | 0 | 94 | 28 | 24 | 74 | 440 | 88 | 230 | 118 | 30 | 191 |
| PBANKA_1144000 | 0 | 0 | 220 | 70 | 512 | 347 | 715 | 426 | 313 | 195 | 29 | 13 | 1470 | 173 | 1049 | 4152 | 3263 | 519 | 1988 | 996 | 31 | 304 |
| PBANKA_1144100 | 173 | 279 | 160 | 46 | 236 | 103 | 255 | 109 | 133 | 69 | 45 | 10 | 2199 | 866 | 651 | 3131 | 2741 | 530 | 563 | 417 | 66 | 523 |
| PBANKA_1144200 | 51 | 54 | 52 | 10 | 192 | 68 | 299 | 128 | 124 | 72 | 53 | 7 | 217 | 48 | 138 | 423 | 514 | 255 | 658 | 285 | 28 | 202 |
| PBANKA_1144300 | 21 | 15 | 65 | 20 | 160 | 108 | 332 | 119 | 241 | 145 | 28 | 20 | 574 | 197 | 126 | 607 | 2243 | 510 | 609 | 402 | 39 | 326 |
| PBANKA_1144400 | 5 | 7 | 81 | 21 | 179 | 74 | 214 | 77 | 77 | 33 | 15 | 0 | 1022 | 498 | 438 | 2245 | 2590 | 444 | 692 | 696 | 38 | 277 |
| PBANKA_1144500 | 0 | 0 | 41 | 10 | 43 | 19 | 37 | 14 | 10 | 9 | 0 | 0 | 76 | 23 | 98 | 338 | 72 | 25 | 204 | 96 | 12 | 95 |
| PBANKA_1144600 | 28 | 25 | 37 | 12 | 230 | 81 | 229 | 93 | 58 | 36 | 6 | 5 | 575 | 226 | 313 | 1683 | 282 | 66 | 816 | 430 | 41 | 280 |
| PBANKA_1144700 | 15 | 23 | 93 | 17 | 351 | 190 | 816 | 435 | 340 | 152 | 47 | 14 | 455 | 230 | 962 | 4209 | 2129 | 253 | 639 | 1032 | 70 | 526 |
| PBANKA_1144800 | 229 | 362 | 123 | 17 | 283 | 159 | 657 | 273 | 316 | 226 | 57 | 10 | 488 | 137 | 321 | 978 | 2296 | 463 | 2114 | 865 | 379 | 3235 |
| PBANKA_1144900 | 10 | 15 | 36 | 6 | 64 | 38 | 86 | 33 | 46 | 13 | 16 | 0 | 205 | 48 | 247 | 852 | 771 | 136 | 407 | 322 | 14 | 138 |
| PBANKA_1145000 | 6 | 0 | 8 | 0 | 34 | 10 | 28 | 11 | 14 | 7 | 10 | 9 | 112 | 81 | 67 | 396 | 209 | 36 | 600 | 483 | 13 | 140 |
| PBANKA_1145100 | 0 | 0 | 86 | 19 | 360 | 120 | 524 | 196 | 225 | 111 | 340 | 275 | 798 | 966 | 534 | 3606 | 1887 | 173 | 838 | 1421 | 81 | 578 |
| PBANKA_1145200 | 0 | 0 | 9 | 0 | 15 | 8 | 8 | 13 | 13 | 5 | 0 | 0 | 667 | 484 | 55 | 293 | 179 | 120 | 15972 | 12159 | 65 | 600 |
| PBANKA_1145300 | 0 | 0 | 84 | 9 | 185 | 85 | 212 | 65 | 73 | 20 | 36 | 25 | 421 | 348 | 501 | 2408 | 742 | 90 | 720 | 1361 | 38 | 247 |
| PBANKA_1145400 | 0 | 0 | 32 | 15 | 90 | 73 | 102 | 58 | 1008 | 856 | 10380 | 7832 | 5729 | 7001 | 490 | 1961 | 3812 | 1528 | 478 | 765 | 19 | 116 |
| PBANKA_1145500 | 7 | 0 | 269 | 44 | 484 | 214 | 573 | 236 | 207 | 148 | 786 | 553 | 2968 | 3091 | 1100 | 7513 | 6552 | 581 | 531 | 1118 | 27 | 203 |
| PBANKA_1145600 | 347 | 576 | 191 | 54 | 495 | 210 | 455 | 219 | 153 | 101 | 27 | 11 | 787 | 370 | 1008 | 4692 | 2967 | 371 | 118 | 191 | 7 | 38 |
| PBANKA_1145700 | 0 | 0 | 22 | 5 | 52 | 37 | 66 | 41 | 33 | 6 | 230 | 233 | 818 | 421 | 163 | 618 | 283 | 88 | 619 | 374 | 12 | 122 |
| PBANKA_1145800 | 0 | 0 | 5 | 0 | 12 | 17 | 60 | 21 | 181 | 113 | 7673 | 6693 | 26306 | 18489 | 6716 | 22330 | 7204 | 1027 | 3147 | 2931 | 73 | 716 |
| PBANKA_1145900 | 0 | 0 | 14 | 10 | 53 | 30 | 60 | 31 | 171 | 97 | 12800 | 7435 | 26966 | 21657 | 5291 | 24358 | 4480 | 555 | 1092 | 1145 | 17 | 263 |
| PBANKA_1146000 | 0 | 0 | 7 | 0 | 71 | 16 | 67 | 27 | 23 | 20 | 43 | 28 | 4428 | 4306 | 1447 | 7400 | 3855 | 442 | 460 | 415 | 0 | 36 |
| PBANKA_1200091 | 0 | 0 | 5 | 0 | 0 | 0 | 0 | 0 | 0 | 0 | 0 | 0 | 58 | 68 | 20 | 38 | 37 | 6 | 83 | 20 | 6 | 15 |
| PBANKA_1200096 | 0 | 0 | 16 | 0 | 37 | 35 | 33 | 23 | 21 | 12 | 5 | 0 | 510 | 7 | 34 | 30 | 7 | 18 | 56 | 11 | 26 | 134 |
| PBANKA_1200600 | 63 | 59 | 11 | 12 | 42 | 26 | 105 | 49 | 310 | 240 | 8046 | 8762 | 1650 | 5653 | 360 | 2528 | 14173 | 1426 | 358 | 1281 | 150 | 1462 |
| PBANKA_1200700 | 39 | 76 | 34 | 8 | 42 | 16 | 38 | 27 | 35 | 13 | 6 | 0 | 87 | 83 | 47 | 105 | 152 | 28 | 275 | 143 | 1402 | 10878 |
| PBANKA_1200800 | 55 | 71 | 365 | 116 | 503 | 246 | 624 | 390 | 336 | 282 | 2088 | 1312 | 2332 | 1060 | 1155 | 2095 | 1580 | 324 | 3775 | 1948 | 243 | 1851 |
| PBANKA_1200900 | 81 | 125 | 30 | 13 | 65 | 40 | 58 | 40 | 24 | 10 | 5 | 7 | 319 | 186 | 216 | 579 | 306 | 115 | 219 | 145 | 44 | 287 |
| PBANKA_1201000 | 116 | 165 | 107 | 22 | 168 | 90 | 180 | 67 | 67 | 39 | 16 | 6 | 364 | 89 | 457 | 2188 | 646 | 105 | 165 | 194 | 17 | 68 |
| PBANKA_1201100 | 160 | 310 | 50 | 12 | 245 | 148 | 438 | 237 | 165 | 75 | 52 | 18 | 193 | 110 | 215 | 706 | 1822 | 262 | 421 | 689 | 46 | 539 |
| PBANKA_1201200 | 6 | 15 | 83 | 23 | 332 | 132 | 349 | 98 | 178 | 68 | 36 | 7 | 617 | 298 | 772 | 3228 | 3078 | 448 | 2297 | 3347 | 63 | 559 |
| PBANKA_1201300 | 61 | 70 | 65 | 26 | 59 | 31 | 69 | 33 | 33 | 21 | 0 | 0 | 456 | 154 | 71 | 283 | 457 | 161 | 1063 | 388 | 40 | 245 |
| PBANKA_1201500 | 0 | 0 | 17 | 0 | 86 | 31 | 85 | 32 | 19 | 14 | 0 | 0 | 80 | 28 | 87 | 183 | 261 | 84 | 377 | 134 | 10 | 84 |
| PBANKA_1201600 | 0 | 0 | 81 | 8 | 130 | 70 | 114 | 60 | 38 | 22 | 0 | 0 | 226 | 33 | 272 | 1013 | 350 | 57 | 786 | 476 | 80 | 618 |
| PBANKA_1201700 | 13 | 22 | 32 | 6 | 48 | 29 | 46 | 12 | 23 | 9 | 86 | 79 | 415 | 187 | 122 | 423 | 137 | 26 | 754 | 433 | 79 | 521 |
| PBANKA_1201800 | 15 | 16 | 11 | 13 | 51 | 30 | 139 | 55 | 1163 | 501 | 278 | 125 | 2229 | 2260 | 260 | 821 | 4587 | 1151 | 19914 | 35004 | 103 | 775 |
| PBANKA_1201900 | 40 | 46 | 933 | 325 | 606 | 452 | 500 | 302 | 123 | 106 | 815 | 738 | 4725 | 1772 | 1340 | 4059 | 1190 | 227 | 1679 | 678 | 100 | 892 |
| PBANKA_1202000 | 0 | 10 | 29 | 8 | 48 | 40 | 89 | 45 | 1510 | 1243 | 420 | 301 | 2303 | 1183 | 223 | 992 | 2921 | 1698 | 28504 | 20449 | 17381 | 1E+05 |
| PBANKA_1202100 | 0 | 18 | 210 | 43 | 547 | 268 | 458 | 181 | 153 | 78 | 30 | 7 | 512 | 183 | 2003 | 9475 | 1370 | 199 | 1938 | 2622 | 501 | 3728 |
| PBANKA_1202200 | 0 | 0 | 288 | 78 | 695 | 302 | 671 | 218 | 219 | 84 | 379 | 227 | 3663 | 3390 | 2323 | 13861 | 2890 | 300 | 2840 | 4850 | 102 | 712 |
| PBANKA_1202300 | 0 | 0 | 7 | 0 | 24 | 13 | 32 | 13 | 21 | 0 | 11 | 0 | 605 | 150 | 91 | 382 | 141 | 73 | 2082 | 845 | 9 | 60 |
| PBANKA_1202400 | 477 | 599 | 2512 | 800 | 1424 | 1018 | 1657 | 906 | 754 | 572 | 5200 | 4554 | 16324 | 6736 | 3373 | 13252 | 4459 | 875 | 3092 | 1750 | 648 | 5366 |
| PBANKA_1202500 | 182 | 253 | 198 | 41 | 588 | 240 | 1522 | 621 | 977 | 369 | 179 | 66 | 2295 | 2673 | 1684 | 8930 | 6075 | 535 | 1213 | 2290 | 331 | 2118 |
| PBANKA_1202600 | 523 | 805 | 333 | 94 | 820 | 465 | 1150 | 486 | 895 | 539 | 2303 | 2023 | 5792 | 4828 | 1602 | 9913 | 7007 | 1195 | 40015 | 62553 | 832 | 6369 |
| PBANKA_1202700 | 99 | 146 | 375 | 113 | 981 | 384 | 913 | 311 | 363 | 164 | 169 | 101 | 5046 | 5510 | 2287 | 18093 | 7305 | 577 | 10418 | 31580 | 1787 | 13543 |
| PBANKA_1202800 | 25 | 28 | 75 | 5 | 159 | 49 | 164 | 58 | 102 | 48 | 43 | 14 | 637 | 177 | 154 | 592 | 1279 | 308 | 2256 | 1175 | 61 | 473 |
| PBANKA_1203000 | 0 | 0 | 49 | 8 | 130 | 101 | 151 | 52 | 57 | 24 | 14 | 0 | 105 | 41 | 113 | 318 | 223 | 29 | 62 | 23 | 5 | 63 |
| PBANKA_1203100 | 0 | 0 | 44 | 0 | 71 | 30 | 80 | 31 | 28 | 14 | 5 | 0 | 101 | 75 | 74 | 221 | 147 | 19 | 43 | 26 | 14 | 54 |
| PBANKA_1203200 | 0 | 0 | 25 | 0 | 61 | 25 | 39 | 10 | 12 | 9 | 10 | 0 | 74 | 30 | 110 | 435 | 106 | 22 | 149 | 68 | 31 | 155 |
| PBANKA_1203300 | 5 | 0 | 32 | 7 | 85 | 45 | 111 | 29 | 50 | 22 | 39 | 29 | 633 | 369 | 207 | 987 | 316 | 54 | 417 | 201 | 97 | 623 |
| PBANKA_1203400 | 0 | 0 | 126 | 22 | 207 | 50 | 175 | 63 | 94 | 42 | 364 | 201 | 3920 | 3491 | 745 | 4355 | 1555 | 240 | 1365 | 1305 | 384 | 2078 |
| PBANKA_1203700 | 16 | 20 | 85 | 35 | 598 | 288 | 605 | 210 | 356 | 180 | 76 | 46 | 243 | 301 | 770 | 3511 | 1389 | 212 | 2667 | 6347 | 144 | 784 |
| PBANKA_1203800 | 72 | 81 | 63 | 11 | 118 | 38 | 144 | 44 | 226 | 100 | 261 | 104 | 2385 | 1588 | 332 | 1773 | 951 | 194 | 6996 | 6390 | 1046 | 4788 |
| PBANKA_1203900 | 6 | 6 | 26 | 0 | 94 | 28 | 133 | 38 | 222 | 115 | 46 | 17 | 425 | 79 | 190 | 1064 | 342 | 153 | 1820 | 755 | 346 | 2202 |
| PBANKA_1204100 | 46 | 34 | 98 | 23 | 250 | 114 | 214 | 95 | 111 | 55 | 47 | 11 | 769 | 366 | 632 | 2367 | 2218 | 377 | 3048 | 3557 | 602 | 4971 |
| PBANKA_1204300 | 0 | 0 | 109 | 38 | 141 | 115 | 162 | 70 | 27 | 17 | 0 | 0 | 292 | 93 | 447 | 2289 | 673 | 74 | 1486 | 890 | 63 | 372 |
| PBANKA_1204400 | 7 | 0 | 837 | 324 | 514 | 316 | 427 | 270 | 150 | 100 | 523 | 380 | 3519 | 1056 | 1769 | 2846 | 771 | 230 | 967 | 328 | 81 | 673 |
| PBANKA_1204600 | 0 | 8 | 20 | 7 | 88 | 24 | 116 | 41 | 62 | 23 | 27 | 0 | 185 | 81 | 129 | 621 | 174 | 31 | 1302 | 767 | 979 | 6238 |
| PBANKA_1204700 | 0 | 0 | 17 | 0 | 92 | 41 | 101 | 30 | 67 | 25 | 0 | 0 | 104 | 25 | 157 | 586 | 115 | 20 | 256 | 146 | 205 | 1130 |
| PBANKA_1204800 | 0 | 0 | 21 | 0 | 57 | 27 | 55 | 12 | 20 | 11 | 0 | 0 | 1046 | 292 | 392 | 1788 | 625 | 98 | 1239 | 616 | 138 | 904 |

| | | | | | | | | | | | | | | | | | | | | | | |
|---|---|---|---|---|---|---|---|---|---|---|---|---|---|---|---|---|---|---|---|---|---|---|
| PBANKA_1204900 | 12 | 15 | 153 | 35 | 159 | 89 | 227 | 71 | 62 | 34 | 25 | 9 | 1896 | 589 | 409 | 1623 | 550 | 115 | 1970 | 609 | 62 | 353 |
| PBANKA_1205000 | 5 | 0 | 662 | 171 | 438 | 196 | 501 | 153 | 125 | 96 | 197 | 201 | 9211 | 4567 | 1499 | 8384 | 3139 | 351 | 1827 | 1699 | 50 | 355 |
| PBANKA_1205100 | 0 | 0 | 16 | 0 | 36 | 15 | 64 | 58 | 118 | 79 | 103 | 95 | 1273 | 478 | 265 | 606 | 701 | 487 | 8198 | 2910 | 252 | 1463 |
| PBANKA_1205200 | 0 | 5 | 80 | 9 | 195 | 124 | 215 | 124 | 97 | 65 | 9 | 5 | 721 | 188 | 226 | 637 | 1101 | 368 | 9301 | 4684 | 96 | 624 |
| PBANKA_1205400 | 874 | 1219 | 46 | 14 | 396 | 189 | 539 | 286 | 305 | 177 | 70 | 34 | 600 | 144 | 287 | 899 | 2749 | 395 | 2237 | 1378 | 964 | 7021 |
| PBANKA_1205600 | 13 | 16 | 28 | 0 | 94 | 35 | 95 | 41 | 72 | 37 | 10 | 6 | 505 | 186 | 231 | 1208 | 1877 | 325 | 408 | 269 | 21 | 84 |
| PBANKA_1205700 | 0 | 0 | 8 | 0 | 18 | 16 | 11 | 8 | 8 | 0 | 0 | 0 | 221 | 85 | 50 | 298 | 305 | 49 | 257 | 176 | 10 | 60 |
| PBANKA_1205800 | 0 | 0 | 41 | 16 | 78 | 58 | 165 | 50 | 55 | 32 | 6 | 5 | 515 | 62 | 329 | 1218 | 1105 | 296 | 1131 | 406 | 41 | 421 |
| PBANKA_1205900 | 396 | 489 | 327 | 72 | 1099 | 506 | 1622 | 646 | 1000 | 486 | 1911 | 1084 | 2813 | 2446 | 1125 | 5966 | 9900 | 1332 | 1067 | 1496 | 428 | 3366 |
| PBANKA_1206000 | 74 | 69 | 71 | 19 | 237 | 75 | 352 | 145 | 273 | 97 | 69 | 26 | 1897 | 1322 | 202 | 1337 | 5398 | 1042 | 15224 | 26511 | 746 | 4377 |
| PBANKA_1206100 | 8 | 10 | 404 | 99 | 459 | 225 | 470 | 165 | 128 | 89 | 344 | 272 | 3533 | 2260 | 1167 | 5168 | 1172 | 192 | 1499 | 922 | 113 | 797 |
| PBANKA_1206200 | 11 | 8 | 138 | 32 | 419 | 216 | 1132 | 448 | 710 | 326 | 170 | 53 | 4807 | 4991 | 3309 | 14482 | 14037 | 2157 | 2133 | 4338 | 69 | 416 |
| PBANKA_1206300 | 593 | 817 | 874 | 194 | 2063 | 850 | 3339 | 1174 | 1875 | 813 | 5090 | 2534 | 6690 | 15397 | 6495 | 30212 | 39588 | 4923 | 1396 | 8098 | 921 | 7020 |
| PBANKA_1206400 | 5 | 0 | 47 | 11 | 234 | 108 | 320 | 134 | 119 | 56 | 14 | 7 | 380 | 106 | 540 | 1765 | 1474 | 283 | 723 | 439 | 71 | 492 |
| PBANKA_1206500 | 0 | 0 | 9 | 0 | 56 | 28 | 98 | 41 | 47 | 43 | 0 | 0 | 151 | 40 | 126 | 370 | 548 | 151 | 238 | 151 | 7 | 61 |
| PBANKA_1206600 | 5 | 0 | 488 | 137 | 968 | 533 | 1252 | 556 | 481 | 250 | 58 | 41 | 4398 | 731 | 2783 | 10114 | 7518 | 1193 | 4182 | 2355 | 68 | 525 |
| PBANKA_1206700 | 35 | 39 | 49 | 9 | 234 | 113 | 270 | 105 | 211 | 110 | 45 | 15 | 591 | 190 | 269 | 1879 | 2159 | 395 | 1420 | 944 | 189 | 1192 |
| PBANKA_1206900 | 17 | 14 | 585 | 228 | 2111 | 1980 | 3350 | 1843 | 1403 | 1028 | 139 | 91 | 3381 | 802 | 4063 | 9861 | 14307 | 3203 | 21148 | 8482 | 365 | 5287 |
| PBANKA_1207000 | 213 | 283 | 302 | 117 | 254 | 126 | 111 | 58 | 262 | 170 | 2780 | 2370 | 2986 | 2006 | 603 | 2521 | 621 | 181 | 1035 | 583 | 377 | 2706 |
| PBANKA_1207100 | 54 | 85 | 39 | 6 | 82 | 46 | 114 | 66 | 620 | 497 | 4934 | 4556 | 893 | 549 | 336 | 946 | 499 | 240 | 1319 | 1075 | 93 | 684 |
| PBANKA_1207200 | 0 | 0 | 228 | 57 | 296 | 186 | 513 | 264 | 173 | 110 | 272 | 248 | 1992 | 1009 | 749 | 2918 | 2722 | 388 | 1184 | 758 | 17 | 123 |
| PBANKA_1207300 | 0 | 0 | 146 | 28 | 160 | 107 | 182 | 56 | 79 | 24 | 95 | 46 | 914 | 472 | 198 | 751 | 336 | 51 | 263 | 153 | 8 | 101 |
| PBANKA_1207500 | 0 | 0 | 42 | 7 | 58 | 26 | 44 | 6 | 18 | 8 | 15 | 7 | 614 | 261 | 439 | 1818 | 177 | 52 | 1117 | 854 | 7 | 83 |
| PBANKA_1207600 | 0 | 0 | 102 | 29 | 202 | 76 | 170 | 64 | 61 | 21 | 13 | 10 | 853 | 511 | 378 | 2031 | 409 | 40 | 269 | 150 | 33 | 219 |
| PBANKA_1207700 | 0 | 6 | 152 | 47 | 365 | 264 | 346 | 134 | 124 | 63 | 278 | 271 | 1945 | 1051 | 633 | 2454 | 877 | 190 | 386 | 189 | 31 | 220 |
| PBANKA_1207800 | 0 | 0 | 15 | 7 | 63 | 27 | 90 | 55 | 158 | 81 | 32 | 19 | 163 | 70 | 134 | 439 | 416 | 117 | 534 | 369 | 9 | 82 |
| PBANKA_1207900 | 0 | 0 | 5 | 0 | 14 | 5 | 31 | 21 | 611 | 588 | 2513 | 1830 | 743 | 565 | 82 | 351 | 1920 | 861 | 522 | 392 | 54 | 472 |
| PBANKA_1208100 | 71 | 76 | 40 | 11 | 162 | 44 | 148 | 45 | 50 | 18 | 8 | 0 | 372 | 175 | 480 | 1934 | 394 | 82 | 462 | 318 | 176 | 1017 |
| PBANKA_1208200 | 210 | 282 | 212 | 44 | 823 | 328 | 1444 | 472 | 568 | 241 | 115 | 46 | 997 | 1897 | 1200 | 4765 | 16835 | 2154 | 2189 | 16951 | 629 | 3716 |
| PBANKA_1208300 | 15 | 28 | 55 | 0 | 119 | 57 | 142 | 39 | 56 | 23 | 12 | 8 | 227 | 89 | 148 | 586 | 243 | 64 | 1197 | 548 | 21 | 153 |
| PBANKA_1208400 | 105 | 128 | 87 | 27 | 120 | 61 | 170 | 64 | 56 | 34 | 6 | 13 | 319 | 46 | 177 | 775 | 532 | 115 | 831 | 411 | 26 | 185 |
| PBANKA_1208500 | 14 | 17 | 22 | 0 | 95 | 32 | 124 | 46 | 55 | 21 | 0 | 0 | 309 | 57 | 156 | 835 | 684 | 78 | 540 | 195 | 8 | 34 |
| PBANKA_1208600 | 6 | 0 | 170 | 32 | 210 | 73 | 227 | 75 | 127 | 58 | 739 | 382 | 1553 | 2666 | 504 | 2739 | 508 | 74 | 1285 | 1906 | 78 | 632 |
| PBANKA_1208700 | 0 | 0 | 60 | 17 | 87 | 33 | 116 | 36 | 84 | 67 | 1370 | 1075 | 1024 | 904 | 144 | 468 | 451 | 103 | 871 | 347 | 77 | 467 |
| PBANKA_1208800 | 28 | 47 | 12 | 0 | 57 | 14 | 102 | 23 | 28 | 23 | 303 | 146 | 538 | 175 | 59 | 146 | 668 | 81 | 2129 | 3225 | 317 | 1548 |
| PBANKA_1209000 | 0 | 0 | 179 | 50 | 153 | 107 | 120 | 70 | 46 | 19 | 79 | 47 | 3049 | 1382 | 782 | 3450 | 1066 | 128 | 1083 | 685 | 156 | 1738 |
| PBANKA_1209100 | 0 | 0 | 62 | 9 | 166 | 77 | 237 | 91 | 75 | 51 | 0 | 0 | 761 | 122 | 351 | 1056 | 440 | 126 | 706 | 335 | 45 | 326 |
| PBANKA_1209200 | 0 | 0 | 17 | 0 | 33 | 12 | 36 | 9 | 5 | 0 | 0 | 0 | 96 | 16 | 87 | 410 | 123 | 21 | 354 | 154 | 25 | 165 |
| PBANKA_1209300 | 0 | 0 | 26 | 0 | 30 | 25 | 71 | 40 | 32 | 17 | 0 | 0 | 132 | 33 | 185 | 651 | 234 | 52 | 311 | 135 | 11 | 84 |
| PBANKA_1209500 | 0 | 5 | 45 | 7 | 109 | 35 | 83 | 29 | 55 | 25 | 0 | 0 | 471 | 180 | 207 | 1110 | 628 | 90 | 520 | 397 | 111 | 928 |
| PBANKA_1209600 | 0 | 0 | 18 | 0 | 55 | 30 | 63 | 17 | 39 | 24 | 0 | 5 | 194 | 57 | 49 | 108 | 123 | 28 | 176 | 100 | 11 | 84 |
| PBANKA_1209700 | 0 | 0 | 24 | 0 | 83 | 32 | 178 | 43 | 83 | 41 | 9 | 5 | 347 | 65 | 62 | 226 | 1354 | 193 | 1392 | 501 | 966 | 5889 |
| PBANKA_1209800 | 6 | 0 | 337 | 84 | 619 | 366 | 957 | 449 | 281 | 171 | 34 | 9 | 2361 | 506 | 1796 | 7271 | 3911 | 507 | 1641 | 918 | 63 | 489 |
| PBANKA_1209900 | 147 | 173 | 8 | 0 | 41 | 29 | 113 | 56 | 46 | 27 | 0 | 0 | 452 | 217 | 32 | 129 | 588 | 190 | 4166 | 2269 | 3896 | 22701 |
| PBANKA_1210000 | 37 | 38 | 43 | 10 | 196 | 69 | 222 | 74 | 103 | 25 | 5 | 5 | 603 | 176 | 201 | 1108 | 502 | 88 | 1239 | 288 | 599 | 3039 |
| PBANKA_1210100 | 26 | 43 | 140 | 33 | 176 | 103 | 217 | 87 | 70 | 53 | 19 | 13 | 1331 | 711 | 567 | 2246 | 1364 | 179 | 411 | 247 | 223 | 1686 |
| PBANKA_1210200 | 11 | 22 | 1321 | 339 | 1364 | 626 | 1115 | 430 | 490 | 209 | 4223 | 2580 | 33392 | 46140 | 7906 | 67122 | 14175 | 1978 | 53896 | 1E+05 | 1174 | 7468 |
| PBANKA_1210300 | 26 | 32 | 86 | 24 | 223 | 99 | 278 | 127 | 81 | 59 | 6 | 0 | 694 | 106 | 438 | 1365 | 1162 | 269 | 2009 | 615 | 106 | 742 |
| PBANKA_1210400 | 88 | 125 | 23 | 9 | 56 | 24 | 54 | 26 | 104 | 40 | 5 | 0 | 584 | 150 | 242 | 986 | 2856 | 614 | 1652 | 914 | 8 | 73 |
| PBANKA_1210500 | 0 | 0 | 57 | 14 | 78 | 36 | 113 | 26 | 227 | 119 | 44 | 14 | 721 | 201 | 306 | 1517 | 2512 | 609 | 233 | 259 | 97 | 785 |
| PBANKA_1210700 | 0 | 0 | 153 | 40 | 346 | 273 | 566 | 356 | 436 | 369 | 48 | 25 | 763 | 177 | 538 | 1459 | 2283 | 652 | 2136 | 1144 | 123 | 1350 |
| PBANKA_1210800 | 0 | 0 | 269 | 76 | 290 | 210 | 321 | 152 | 112 | 62 | 63 | 30 | 21831 | 1078 | 6970 | 41976 | 1534 | 591 | 2214 | 1477 | 20 | 227 |
| PBANKA_1210900 | 0 | 11 | 66 | 14 | 244 | 219 | 281 | 120 | 122 | 79 | 122 | 143 | 3027 | 1140 | 937 | 4312 | 2074 | 356 | 5307 | 3742 | 1026 | 9964 |
| PBANKA_1211000 | 0 | 5 | 105 | 35 | 162 | 83 | 176 | 72 | 80 | 33 | 58 | 52 | 2468 | 1285 | 541 | 2222 | 533 | 115 | 1313 | 709 | 109 | 880 |
| PBANKA_1211100 | 0 | 0 | 93 | 22 | 443 | 194 | 565 | 169 | 226 | 109 | 52 | 20 | 1605 | 1112 | 459 | 2356 | 2077 | 290 | 762 | 771 | 77 | 694 |
| PBANKA_1211200 | 0 | 0 | 115 | 22 | 187 | 77 | 140 | 62 | 69 | 36 | 10 | 0 | 909 | 317 | 569 | 2752 | 431 | 76 | 712 | 329 | 11 | 55 |
| PBANKA_1211300 | 0 | 0 | 44 | 14 | 188 | 83 | 241 | 113 | 100 | 51 | 147 | 107 | 1488 | 523 | 533 | 2380 | 1339 | 259 | 985 | 493 | 196 | 1695 |
| PBANKA_1211400 | 0 | 0 | 20 | 7 | 113 | 68 | 101 | 50 | 59 | 26 | 7 | 0 | 275 | 69 | 149 | 488 | 320 | 65 | 1158 | 520 | 55 | 387 |
| PBANKA_1211500 | 42 | 52 | 227 | 53 | 346 | 132 | 264 | 88 | 166 | 95 | 1111 | 610 | 1669 | 1643 | 506 | 3108 | 689 | 105 | 533 | 611 | 218 | 1547 |
| PBANKA_1211600 | 27 | 38 | 80 | 14 | 306 | 169 | 498 | 136 | 547 | 295 | 71 | 54 | 1009 | 514 | 419 | 2051 | 5340 | 989 | 1243 | 1038 | 74 | 582 |
| PBANKA_1211700 | 5 | 0 | 112 | 34 | 287 | 156 | 518 | 198 | 221 | 110 | 19 | 14 | 760 | 165 | 402 | 1257 | 3064 | 439 | 902 | 512 | 58 | 493 |
| PBANKA_1211800 | 38 | 67 | 21 | 0 | 185 | 51 | 1511 | 754 | 1087 | 553 | 121 | 56 | 266 | 184 | 49 | 299 | 215 | 31 | 483 | 479 | 5 | 23 |
| PBANKA_1211900 | 59 | 87 | 214 | 62 | 630 | 275 | 777 | 301 | 669 | 360 | 429 | 229 | 3733 | 3464 | 1145 | 5380 | 8073 | 1647 | 4323 | 4554 | 643 | 4805 |
| PBANKA_1212000 | 159 | 213 | 263 | 67 | 1677 | 637 | 2447 | 905 | 979 | 455 | 186 | 65 | 771 | 604 | 339 | 1212 | 11191 | 1100 | 1420 | 5409 | 264 | 1994 |
| PBANKA_1212100 | 0 | 0 | 42 | 20 | 50 | 25 | 64 | 29 | 58 | 51 | 7 | 0 | 99 | 15 | 127 | 202 | 288 | 145 | 28 | 18 | 0 | 41 |
| PBANKA_1212300 | 0 | 6 | 119 | 17 | 310 | 149 | 409 | 175 | 138 | 73 | 18 | 24 | 1271 | 325 | 676 | 3596 | 1783 | 214 | 1153 | 986 | 179 | 1244 |
| PBANKA_1212700 | 561 | 734 | 272 | 94 | 1795 | 723 | 1504 | 524 | 739 | 351 | 213 | 93 | 1141 | 1324 | 1594 | 8759 | 7700 | 1091 | 5512 | 12398 | 255 | 2189 |
| PBANKA_1212800 | 7 | 9 | 107 | 18 | 218 | 88 | 298 | 153 | 108 | 69 | 18 | 20 | 251 | 71 | 229 | 964 | 753 | 148 | 66 | 62 | 45 | 335 |
| PBANKA_1212900 | 25 | 28 | 47 | 11 | 90 | 43 | 171 | 55 | 102 | 43 | 565 | 305 | 662 | 655 | 203 | 774 | 852 | 103 | 548 | 892 | 673 | 5470 |
| PBANKA_1213000 | 428 | 641 | 549 | 133 | 1373 | 576 | 2995 | 1099 | 2204 | 968 | 3551 | 1798 | 7088 | 10198 | 4376 | 19656 | 29587 | 4585 | 3657 | 6413 | 869 | 4598 |
| PBANKA_1213100 | 113 | 151 | 160 | 50 | 209 | 167 | 252 | 85 | 54 | 34 | 12 | 14 | 252 | 51 | 369 | 1394 | 359 | 57 | 1335 | 800 | 341 | 3002 |
| PBANKA_1213400 | 0 | 0 | 13 | 0 | 44 | 20 | 49 | 20 | 30 | 12 | 6 | 0 | 144 | 32 | 98 | 405 | 565 | 116 | 404 | 253 | 42 | 295 |
| PBANKA_1213500 | 42 | 92 | 22 | 5 | 77 | 35 | 113 | 50 | 38 | 35 | 27 | 23 | 594 | 227 | 207 | 809 | 830 | 127 | 764 | 465 | 24 | 142 |
| PBANKA_1213600 | 0 | 0 | 472 | 180 | 677 | 469 | 571 | 267 | 162 | 66 | 58 | 68 | 1971 | 870 | 770 | 3317 | 712 | 146 | 2998 | 1763 | 576 | 4196 |
| PBANKA_1213700 | 0 | 0 | 226 | 67 | 274 | 157 | 347 | 162 | 165 | 86 | 153 | 88 | 1231 | 630 | 363 | 1507 | 292 | 58 | 1513 | 1138 | 1612 | 12692 |
| PBANKA_1213800 | 19 | 28 | 32 | 11 | 366 | 132 | 519 | 211 | 176 | 86 | 20 | 11 | 178 | 56 | 172 | 584 | 739 | 127 | 693 | 462 | 135 | 1043 |
| PBANKA_1213900 | 10 | 17 | 136 | 46 | 263 | 167 | 329 | 137 | 88 | 61 | 13 | 8 | 976 | 453 | 555 | 2670 | 1176 | 189 | 485 | 360 | 40 | 226 |
| PBANKA_1214000 | 0 | 0 | 1484 | 566 | 696 | 533 | 405 | 176 | 158 | 76 | 19 | 36 | 3275 | 530 | 2921 | 13545 | 865 | 162 | 1287 | 1035 | 166 | 1684 |
| PBANKA_1214200 | 17 | 10 | 164 | 44 | 348 | 133 | 682 | 277 | 240 | 153 | 31 | 8 | 1700 | 294 | 906 | 2130 | 5541 | 1046 | 4555 | 1817 | 256 | 1704 |
| PBANKA_1214300 | 390 | 495 | 4678 | 1486 | 2699 | 1941 | 2235 | 1072 | 728 | 465 | 287 | 224 | 19470 | 5286 | 13409 | 46425 | 6598 | 1047 | 5455 | 2922 | 1621 | 13962 |
| PBANKA_1214400 | 25 | 33 | 5 | 0 | 45 | 17 | 64 | 28 | 150 | 85 | 42 | 12 | 572 | 331 | 83 | 304 | 584 | 213 | 9448 | 9677 | 178 | 1069 |
| PBANKA_1214600 | 20 | 24 | 27 | 12 | 209 | 40 | 269 | 71 | 112 | 43 | 10 | 11 | 447 | 103 | 309 | 1399 | 1086 | 203 | 1712 | 864 | 299 | 1857 |
| PBANKA_1214700 | 352 | 518 | 193 | 44 | 787 | 274 | 1244 | 440 | 516 | 234 | 48 | 24 | 1533 | 581 | 1484 | 6283 | 7050 | 1035 | 9036 | 7385 | 457 | 3997 |
| PBANKA_1214800 | 133 | 227 | 246 | 49 | 434 | 130 | 638 | 236 | 316 | 113 | 495 | 298 | 6949 | 4576 | 1827 | 12213 | 8657 | 1127 | 5034 | 11047 | 1190 | 6017 |
| PBANKA_1214900 | 161 | 220 | 126 | 13 | 568 | 174 | 657 | 243 | 234 | 111 | 465 | 238 | 2032 | 1427 | 1102 | 5336 | 2693 | 380 | 1646 | 1604 | 305 | 2264 |
| PBANKA_1215000 | 0 | 7 | 62 | 19 | 127 | 63 | 163 | 66 | 117 | 77 | 47 | 14 | 1807 | 659 | 297 | 1760 | 2095 | 388 | 195 | 293 | 458 | 2765 |
| PBANKA_1215100 | 0 | 0 | 23 | 8 | 162 | 85 | 264 | 89 | 622 | 376 | 131 | 64 | 397 | 136 | 263 | 1134 | 1982 | 590 | 261 | 236 | 264 | 1752 |
| PBANKA_1215200 | 0 | 0 | 48 | 5 | 26 | 8 | 20 | 7 | 7 | 0 | 0 | 0 | 145 | 22 | 192 | 1162 | 69 | 31 | 405 | 166 | 5 | 20 |
| PBANKA_1215300 | 0 | 0 | 42 | 0 | 59 | 17 | 86 | 36 | 45 | 30 | 5 | 6 | 367 | 109 | 117 | 548 | 621 | 117 | 1220 | 358 | 75 | 457 |
| PBANKA_1215400 | 64 | 85 | 135 | 21 | 381 | 118 | 843 | 274 | 714 | 314 | 389 | 200 | 2141 | 1550 | 923 | 5472 | 6164 | 868 | 4122 | 7441 | 486 | 3161 |

| | | | | | | | | | | | | | | | | | | | | | | |
|---|---|---|---|---|---|---|---|---|---|---|---|---|---|---|---|---|---|---|---|---|---|---|
| PBANKA_1215500 | 18 | 8 | 23 | 9 | 26 | 10 | 53 | 10 | 16 | 7 | 8 | 9 | 317 | 119 | 236 | 301 | 298 | 256 | 368 | 106 | 38 | 79 |
| PBANKA_1215600 | 5 | 5 | 1688 | 431 | 1077 | 538 | 1154 | 461 | 401 | 249 | 1452 | 1100 | 15288 | 6534 | 2786 | 12306 | 5113 | 794 | 2797 | 1331 | 113 | 1082 |
| PBANKA_1215800 | 88 | 122 | 115 | 22 | 599 | 180 | 1005 | 305 | 507 | 243 | 370 | 198 | 1688 | 1317 | 858 | 4255 | 3259 | 413 | 1846 | 2681 | 278 | 1636 |
| PBANKA_1215900 | 23 | 22 | 12 | 0 | 39 | 12 | 54 | 17 | 31 | 12 | 21 | 0 | 501 | 298 | 111 | 617 | 1102 | 132 | 1375 | 1684 | 63 | 320 |
| PBANKA_1216000 | 89 | 102 | 28 | 8 | 90 | 21 | 67 | 24 | 36 | 6 | 0 | 7 | 389 | 164 | 99 | 570 | 238 | 46 | 676 | 378 | 319 | 1607 |
| PBANKA_1216100 | 0 | 0 | 73 | 17 | 178 | 71 | 147 | 36 | 43 | 20 | 116 | 65 | 803 | 542 | 227 | 1296 | 212 | 37 | 550 | 243 | 9 | 71 |
| PBANKA_1216200 | 0 | 0 | 30 | 0 | 351 | 188 | 513 | 182 | 191 | 77 | 12 | 9 | 1009 | 293 | 340 | 1774 | 1773 | 265 | 533 | 281 | 41 | 296 |
| PBANKA_1216300 | 0 | 0 | 6 | 0 | 158 | 64 | 324 | 92 | 180 | 71 | 31 | 0 | 446 | 208 | 28 | 129 | 1498 | 213 | 1271 | 3046 | 324 | 1619 |
| PBANKA_1216400 | 0 | 0 | 79 | 34 | 147 | 75 | 172 | 65 | 64 | 41 | 71 | 34 | 694 | 473 | 378 | 1525 | 554 | 55 | 109 | 103 | 10 | 73 |
| PBANKA_1216500 | 0 | 11 | 1548 | 568 | 1107 | 875 | 1066 | 713 | 342 | 246 | 319 | 266 | 10047 | 3028 | 7472 | 18520 | 3845 | 774 | 1839 | 771 | 144 | 1397 |
| PBANKA_1216600 | 67 | 63 | 219 | 51 | 705 | 237 | 929 | 288 | 609 | 186 | 674 | 379 | 2251 | 1654 | 701 | 3006 | 4533 | 771 | 1041 | 1315 | 672 | 4063 |
| PBANKA_1216700 | 411 | 672 | 101 | 32 | 568 | 228 | 1727 | 644 | 1441 | 609 | 332 | 159 | 1957 | 4298 | 1294 | 7385 | 5108 | 974 | 1013 | 4298 | 564 | 3983 |
| PBANKA_1216800 | 517 | 781 | 144 | 29 | 723 | 247 | 1814 | 831 | 1306 | 623 | 138 | 55 | 1202 | 1985 | 1162 | 6939 | 16624 | 1776 | 1130 | 13080 | 418 | 2389 |
| PBANKA_1216900 | 0 | 14 | 54 | 13 | 231 | 93 | 303 | 108 | 148 | 78 | 709 | 401 | 1911 | 1333 | 625 | 2988 | 2463 | 353 | 698 | 713 | 33 | 392 |
| PBANKA_1217000 | 0 | 0 | 141 | 27 | 218 | 175 | 382 | 125 | 122 | 51 | 36 | 38 | 930 | 535 | 649 | 2819 | 1178 | 135 | 332 | 409 | 71 | 373 |
| PBANKA_1217200 | 0 | 0 | 14 | 5 | 169 | 83 | 356 | 143 | 443 | 222 | 95 | 41 | 182 | 110 | 114 | 443 | 2527 | 389 | 503 | 887 | 40 | 294 |
| PBANKA_1217300 | 29 | 24 | 50 | 6 | 95 | 49 | 180 | 78 | 98 | 44 | 13 | 8 | 181 | 27 | 91 | 294 | 619 | 175 | 586 | 336 | 85 | 643 |
| PBANKA_1217400 | 98 | 141 | 94 | 23 | 1437 | 619 | 1925 | 630 | 858 | 320 | 162 | 66 | 580 | 348 | 178 | 503 | 5582 | 634 | 225 | 366 | 149 | 1053 |
| PBANKA_1217500 | 0 | 0 | 167 | 26 | 246 | 84 | 173 | 60 | 74 | 32 | 34 | 17 | 2529 | 1817 | 899 | 5316 | 918 | 202 | 592 | 835 | 47 | 367 |
| PBANKA_1217600 | 189 | 252 | 617 | 272 | 1080 | 936 | 1721 | 1140 | 2224 | 1739 | 440 | 344 | 3006 | 1160 | 813 | 2153 | 17818 | 7148 | 1549 | 809 | 526 | 4089 |
| PBANKA_1217700 | 9 | 13 | 168 | 44 | 459 | 305 | 619 | 334 | 240 | 127 | 29 | 34 | 1489 | 465 | 1131 | 2902 | 2613 | 814 | 6900 | 3474 | 351 | 2971 |
| PBANKA_1217800 | 0 | 0 | 76 | 11 | 115 | 78 | 150 | 64 | 70 | 20 | 0 | 0 | 273 | 134 | 257 | 352 | 306 | 96 | 376 | 296 | 32 | 235 |
| PBANKA_1217900 | 0 | 0 | 19 | 0 | 57 | 22 | 65 | 31 | 61 | 36 | 0 | 0 | 112 | 26 | 45 | 113 | 290 | 63 | 798 | 318 | 13 | 107 |
| PBANKA_1218000 | 0 | 0 | 38 | 7 | 29 | 24 | 22 | 0 | 179 | 118 | 2223 | 1510 | 1939 | 1570 | 128 | 712 | 702 | 250 | 17271 | 9973 | 24 | 188 |
| PBANKA_1218100 | 105 | 164 | 324 | 76 | 385 | 239 | 296 | 114 | 194 | 96 | 137 | 41 | 516 | 463 | 668 | 2915 | 807 | 106 | 329 | 1045 | 79962 | 6E+05 |
| PBANKA_1218200 | 0 | 0 | 844 | 208 | 1103 | 647 | 1045 | 381 | 302 | 148 | 33 | 19 | 4424 | 1996 | 3818 | 16940 | 3304 | 411 | 2763 | 6398 | 152 | 1240 |
| PBANKA_1218300 | 0 | 0 | 5 | 0 | 71 | 21 | 50 | 33 | 19 | 11 | 7 | 6 | 1251 | 599 | 35 | 188 | 643 | 253 | 22339 | 25549 | 534 | 3996 |
| PBANKA_1218400 | 63 | 94 | 34 | 0 | 107 | 57 | 217 | 92 | 112 | 90 | 17 | 10 | 577 | 146 | 131 | 353 | 827 | 164 | 783 | 478 | 220 | 969 |
| PBANKA_1218500 | 8 | 13 | 186 | 43 | 224 | 135 | 281 | 154 | 114 | 77 | 14 | 9 | 1054 | 322 | 683 | 1641 | 784 | 278 | 4276 | 1860 | 232 | 1052 |
| PBANKA_1218600 | 6 | 0 | 48 | 11 | 113 | 57 | 145 | 46 | 49 | 30 | 11 | 0 | 315 | 114 | 146 | 508 | 681 | 140 | 1015 | 554 | 68 | 788 |
| PBANKA_1218800 | 7 | 13 | 147 | 35 | 253 | 101 | 321 | 128 | 305 | 124 | 627 | 380 | 3438 | 4246 | 756 | 4207 | 1949 | 271 | 1275 | 1620 | 107 | 642 |
| PBANKA_1218900 | 0 | 0 | 277 | 67 | 271 | 146 | 281 | 147 | 79 | 68 | 16 | 12 | 1769 | 482 | 1205 | 2506 | 1116 | 264 | 1104 | 434 | 51 | 268 |
| PBANKA_1219000 | 5 | 8 | 267 | 46 | 618 | 223 | 608 | 169 | 286 | 131 | 250 | 143 | 9381 | 5469 | 3041 | 17651 | 5554 | 649 | 1341 | 2222 | 209 | 1325 |
| PBANKA_1219100 | 0 | 0 | 7 | 0 | 22 | 7 | 16 | 0 | 20 | 7 | 0 | 0 | 625 | 296 | 29 | 102 | 190 | 150 | 18930 | 6382 | 999 | 7313 |
| PBANKA_1219200 | 25 | 40 | 133 | 49 | 349 | 305 | 559 | 330 | 302 | 191 | 38 | 10 | 689 | 241 | 414 | 1403 | 3055 | 497 | 4773 | 2860 | 181 | 2097 |
| PBANKA_1219300 | 40 | 59 | 2577 | 644 | 3359 | 1649 | 3462 | 1334 | 1059 | 514 | 279 | 158 | 15932 | 12239 | 8617 | 46558 | 13352 | 1332 | 1939 | 4114 | 458 | 4296 |
| PBANKA_1219400 | 0 | 0 | 44 | 13 | 229 | 116 | 210 | 94 | 78 | 45 | 8 | 5 | 374 | 105 | 199 | 622 | 147 | 31 | 297 | 115 | 13 | 89 |
| PBANKA_1219500 | 0 | 0 | 41 | 7 | 116 | 61 | 177 | 79 | 97 | 48 | 132 | 89 | 2478 | 1062 | 956 | 2823 | 1003 | 202 | 532 | 267 | 24 | 182 |
| PBANKA_1219600 | 0 | 0 | 41 | 14 | 124 | 58 | 170 | 52 | 34 | 15 | 16 | 6 | 328 | 188 | 400 | 1465 | 410 | 77 | 885 | 532 | 51 | 322 |
| PBANKA_1219700 | 6 | 18 | 5 | 0 | 62 | 32 | 145 | 38 | 61 | 32 | 10 | 8 | 369 | 151 | 16 | 108 | 430 | 70 | 831 | 298 | 118 | 764 |
| PBANKA_1219800 | 0 | 0 | 122 | 22 | 788 | 314 | 1286 | 434 | 416 | 180 | 52 | 31 | 716 | 492 | 1496 | 6424 | 2779 | 279 | 335 | 575 | 17 | 182 |
| PBANKA_1219900 | 17 | 19 | 97 | 20 | 239 | 92 | 422 | 161 | 172 | 85 | 24 | 8 | 521 | 253 | 412 | 1974 | 2777 | 348 | 2839 | 3017 | 136 | 723 |
| PBANKA_1220000 | 8 | 7 | 34 | 14 | 131 | 68 | 139 | 56 | 67 | 32 | 0 | 0 | 100 | 13 | 148 | 339 | 259 | 62 | 542 | 168 | 33 | 274 |
| PBANKA_1220100 | 8 | 7 | 131 | 17 | 389 | 122 | 402 | 137 | 189 | 69 | 114 | 58 | 2622 | 1254 | 1117 | 6397 | 1707 | 234 | 1499 | 1557 | 60 | 437 |
| PBANKA_1220300 | 190 | 236 | 25 | 0 | 41 | 29 | 122 | 45 | 53 | 31 | 103 | 57 | 1472 | 801 | 376 | 2046 | 1942 | 252 | 835 | 787 | 389 | 2648 |
| PBANKA_1220400 | 41 | 27 | 10 | 0 | 17 | 7 | 22 | 13 | 0 | 10 | 11 | 99 | 27 | 56 | 97 | 76 | 30 | 147 | 28 | 29 | 249 | |
| PBANKA_1220500 | 361 | 429 | 158 | 33 | 675 | 266 | 996 | 329 | 997 | 411 | 156 | 52 | 942 | 661 | 387 | 1047 | 18493 | 3295 | 1341 | 2993 | 162 | 1034 |
| PBANKA_1220600 | 235 | 315 | 40 | 10 | 388 | 92 | 879 | 298 | 666 | 244 | 181 | 74 | 569 | 580 | 176 | 405 | 9686 | 1674 | 1541 | 4355 | 841 | 4706 |
| PBANKA_1220700 | 6 | 6 | 43 | 11 | 131 | 51 | 173 | 57 | 116 | 45 | 42 | 8 | 216 | 79 | 213 | 894 | 1539 | 267 | 620 | 385 | 54 | 416 |
| PBANKA_1220800 | 2023 | 3098 | 310 | 100 | 1073 | 484 | 1736 | 781 | 998 | 512 | 184 | 103 | 1137 | 1495 | 523 | 3267 | 23963 | 2245 | 300 | 2416 | 2891 | 16007 |
| PBANKA_1220900 | 0 | 0 | 91 | 19 | 148 | 60 | 98 | 68 | 43 | 21 | 12 | 0 | 551 | 108 | 177 | 607 | 107 | 36 | 679 | 320 | 7 | 41 |
| PBANKA_1221000 | 0 | 0 | 141 | 30 | 309 | 120 | 257 | 90 | 90 | 46 | 20 | 7 | 1994 | 763 | 435 | 2378 | 327 | 59 | 510 | 189 | 12 | 88 |
| PBANKA_1221100 | 0 | 0 | 43 | 8 | 73 | 43 | 72 | 27 | 17 | 10 | 8 | 0 | 257 | 88 | 160 | 420 | 128 | 49 | 234 | 69 | 14 | 153 |
| PBANKA_1221200 | 143 | 149 | 966 | 272 | 697 | 376 | 761 | 375 | 252 | 213 | 1110 | 810 | 16973 | 6441 | 3312 | 12275 | 4716 | 970 | 2499 | 1701 | 330 | 2312 |
| PBANKA_1221300 | 25 | 54 | 55 | 11 | 82 | 31 | 86 | 49 | 95 | 61 | 249 | 158 | 1507 | 1051 | 258 | 1683 | 1440 | 322 | 584 | 603 | 537 | 2645 |
| PBANKA_1221400 | 0 | 0 | 54 | 15 | 120 | 39 | 181 | 93 | 60 | 50 | 14 | 8 | 925 | 170 | 176 | 952 | 1232 | 154 | 413 | 330 | 339 | 1997 |
| PBANKA_1221500 | 13 | 16 | 49 | 6 | 136 | 63 | 165 | 64 | 78 | 26 | 19 | 7 | 466 | 177 | 266 | 940 | 752 | 144 | 844 | 744 | 1972 | 13138 |
| PBANKA_1221600 | 132 | 194 | 403 | 89 | 682 | 301 | 850 | 356 | 448 | 217 | 706 | 431 | 2664 | 1650 | 1757 | 8891 | 4265 | 720 | 1816 | 3228 | 928 | 7116 |
| PBANKA_1221700 | 12 | 10 | 17 | 0 | 77 | 52 | 84 | 32 | 50 | 44 | 263 | 228 | 683 | 229 | 112 | 467 | 155 | 34 | 178 | 66 | 104 | 1108 |
| PBANKA_1221800 | 28 | 18 | 78 | 10 | 300 | 123 | 266 | 123 | 84 | 34 | 43 | 56 | 961 | 467 | 386 | 1781 | 541 | 88 | 1037 | 728 | 75 | 656 |
| PBANKA_1221900 | 0 | 0 | 9 | 0 | 25 | 12 | 29 | 18 | 18 | 13 | 7 | 6 | 184 | 66 | 37 | 110 | 62 | 19 | 309 | 124 | 67 | 539 |
| PBANKA_1222000 | 0 | 0 | 141 | 26 | 277 | 101 | 218 | 89 | 101 | 41 | 37 | 13 | 2672 | 1914 | 929 | 5976 | 1530 | 205 | 813 | 1325 | 35 | 248 |
| PBANKA_1222100 | 0 | 0 | 79 | 31 | 166 | 89 | 242 | 94 | 293 | 183 | 43 | 17 | 874 | 190 | 283 | 1039 | 1666 | 451 | 1541 | 1159 | 193 | 1499 |
| PBANKA_1222200 | 10 | 13 | 29 | 7 | 49 | 24 | 51 | 33 | 25 | 9 | 6 | 0 | 90 | 17 | 96 | 285 | 188 | 42 | 192 | 113 | 92 | 810 |
| PBANKA_1222300 | 75 | 127 | 69 | 15 | 146 | 53 | 503 | 169 | 438 | 185 | 92 | 53 | 1690 | 534 | 458 | 1606 | 3197 | 769 | 1373 | 677 | 1204 | 6495 |
| PBANKA_1222400 | 60 | 56 | 105 | 32 | 256 | 140 | 270 | 128 | 102 | 65 | 24 | 13 | 1578 | 282 | 663 | 3554 | 2347 | 291 | 3817 | 1795 | 210 | 1542 |
| PBANKA_1222500 | 754 | 1122 | 5 | 0 | 34 | 14 | 56 | 17 | 424 | 272 | 147 | 70 | 508 | 244 | 37 | 140 | 1006 | 498 | 2607 | 3015 | 5433 | 43738 |
| PBANKA_1222600 | 6 | 8 | 28 | 12 | 139 | 71 | 244 | 81 | 103 | 47 | 37 | 10 | 1124 | 492 | 424 | 2082 | 1140 | 148 | 2787 | 1612 | 733 | 3791 |
| PBANKA_1222700 | 65 | 104 | 84 | 16 | 216 | 85 | 279 | 97 | 226 | 98 | 811 | 471 | 468 | 765 | 244 | 1267 | 379 | 100 | 824 | 736 | 1968 | 12007 |
| PBANKA_1222800 | 119 | 98 | 438 | 135 | 656 | 381 | 741 | 271 | 215 | 126 | 70 | 45 | 2394 | 911 | 825 | 2880 | 2135 | 279 | 1146 | 816 | 1362 | 10945 |
| PBANKA_1222900 | 63 | 94 | 330 | 66 | 724 | 330 | 1043 | 443 | 424 | 211 | 72 | 22 | 2108 | 650 | 857 | 4200 | 4515 | 669 | 3012 | 2070 | 1588 | 10072 |
| PBANKA_1223000 | 58 | 135 | 237 | 98 | 607 | 205 | 786 | 325 | 370 | 191 | 142 | 88 | 2204 | 2534 | 1185 | 8144 | 9191 | 664 | 559 | 1406 | 239 | 2060 |
| PBANKA_1223100 | 0 | 0 | 273 | 75 | 534 | 324 | 632 | 352 | 198 | 120 | 29 | 9 | 2024 | 336 | 1524 | 5795 | 2324 | 315 | 1515 | 617 | 165 | 1382 |
| PBANKA_1223200 | 43 | 46 | 25 | 6 | 81 | 36 | 95 | 24 | 15 | 19 | 6 | 0 | 85 | 19 | 54 | 155 | 73 | 25 | 595 | 315 | 109 | 813 |
| PBANKA_1223300 | 0 | 0 | 48 | 8 | 218 | 123 | 472 | 242 | 192 | 103 | 636 | 465 | 2353 | 647 | 546 | 2236 | 945 | 202 | 1171 | 581 | 97 | 880 |
| PBANKA_1223400 | 11 | 10 | 32 | 10 | 80 | 54 | 168 | 53 | 68 | 46 | 14 | 0 | 721 | 126 | 263 | 875 | 1437 | 334 | 3124 | 1005 | 125 | 738 |
| PBANKA_1223500 | 19 | 14 | 88 | 23 | 281 | 91 | 355 | 110 | 438 | 248 | 407 | 278 | 2199 | 1452 | 972 | 5101 | 4236 | 733 | 1158 | 1474 | 380 | 1973 |
| PBANKA_1223600 | 0 | 0 | 59 | 12 | 141 | 111 | 176 | 137 | 57 | 59 | 9 | 0 | 554 | 177 | 316 | 793 | 381 | 125 | 267 | 119 | 33 | 226 |
| PBANKA_1223700 | 0 | 0 | 6 | 0 | 42 | 17 | 52 | 15 | 38 | 16 | 0 | 0 | 249 | 110 | 116 | 618 | 271 | 36 | 380 | 185 | 54 | 427 |
| PBANKA_1223800 | 632 | 918 | 327 | 112 | 1084 | 490 | 1613 | 596 | 1251 | 707 | 334 | 174 | 3421 | 1985 | 1809 | 9211 | 11147 | 1788 | 3463 | 4802 | 5539 | 46606 |
| PBANKA_1223900 | 7 | 7 | 33 | 0 | 76 | 23 | 147 | 46 | 60 | 19 | 8 | 0 | 372 | 96 | 226 | 944 | 1334 | 196 | 315 | 288 | 242 | 1191 |
| PBANKA_1224000 | 120 | 165 | 61 | 9 | 192 | 74 | 201 | 64 | 172 | 94 | 53 | 14 | 807 | 281 | 637 | 2931 | 1154 | 179 | 1301 | 1006 | 287 | 2158 |
| PBANKA_1224100 | 0 | 0 | 7 | 0 | 56 | 28 | 115 | 43 | 185 | 128 | 24 | 9 | 250 | 133 | 205 | 910 | 972 | 262 | 563 | 494 | 339 | 2213 |
| PBANKA_1224200 | 183 | 183 | 107 | 28 | 156 | 78 | 349 | 117 | 810 | 408 | 3397 | 1712 | 12763 | 13353 | 1117 | 6041 | 3020 | 784 | 1761 | 1872 | 955 | 8573 |
| PBANKA_1224300 | 0 | 0 | 12 | 0 | 28 | 12 | 49 | 22 | 14 | 9 | 0 | 0 | 55 | 13 | 58 | 166 | 112 | 22 | 84 | 26 | 24 | 88 |
| PBANKA_1224400 | 0 | 0 | 39 | 11 | 413 | 198 | 982 | 406 | 479 | 210 | 89 | 17 | 442 | 196 | 752 | 3177 | 2841 | 335 | 1382 | 1668 | 124 | 1093 |
| PBANKA_1224500 | 12 | 6 | 25 | 0 | 149 | 58 | 284 | 117 | 125 | 63 | 20 | 7 | 437 | 162 | 149 | 470 | 1259 | 269 | 4608 | 2618 | 133 | 1010 |
| PBANKA_1224600 | 0 | 0 | 39 | 10 | 152 | 60 | 188 | 53 | 92 | 37 | 18 | 0 | 734 | 307 | 305 | 1631 | 1138 | 177 | 798 | 411 | 2223 | 12345 |
| PBANKA_1224700 | 5 | 7 | 93 | 27 | 253 | 116 | 330 | 107 | 131 | 48 | 31 | 0 | 1483 | 799 | 725 | 3874 | 1670 | 244 | 2642 | 1384 | 274 | 1829 |
| PBANKA_1224800 | 0 | 0 | 14 | 5 | 47 | 27 | 74 | 23 | 29 | 10 | 7 | 0 | 346 | 86 | 128 | 781 | 594 | 77 | 853 | 348 | 41 | 283 |

| | | | | | | | | | | | | | | | | | | | | | | |
|---|---|---|---|---|---|---|---|---|---|---|---|---|---|---|---|---|---|---|---|---|---|---|
| PBANKA_1224900 | 5 | 7 | 6 | 0 | 139 | 44 | 960 | 392 | 630 | 256 | 65 | 34 | 989 | 285 | 36 | 88 | 3200 | 673 | 4218 | 4205 | 549 | 4147 |
| PBANKA_1225000 | 173 | 237 | 39 | 7 | 358 | 146 | 1334 | 505 | 1032 | 488 | 75 | 52 | 2264 | 970 | 474 | 1269 | 27084 | 3975 | 2339 | 3750 | 818 | 6191 |
| PBANKA_1225100 | 70 | 89 | 33 | 17 | 111 | 65 | 224 | 92 | 147 | 47 | 46 | 11 | 597 | 358 | 139 | 654 | 3408 | 535 | 2255 | 2475 | 30 | 151 |
| PBANKA_1225200 | 0 | 0 | 28 | 10 | 20 | 6 | 6 | 0 | 7 | 0 | 0 | 0 | 40 | 5 | 186 | 460 | 88 | 19 | 24 | 11 | 0 | 8 |
| PBANKA_1225700 | 143 | 202 | 126 | 35 | 231 | 142 | 409 | 191 | 235 | 142 | 33 | 16 | 1187 | 595 | 358 | 1569 | 3691 | 535 | 1448 | 1027 | 2104 | 14690 |
| PBANKA_1225800 | 5 | 0 | 127 | 31 | 585 | 333 | 870 | 397 | 346 | 197 | 53 | 36 | 983 | 382 | 328 | 1059 | 1343 | 355 | 938 | 473 | 137 | 1030 |
| PBANKA_1225900 | 205 | 304 | 59 | 9 | 610 | 186 | 571 | 254 | 403 | 164 | 135 | 39 | 2015 | 4274 | 181 | 765 | 6641 | 1519 | 22522 | 1E+05 | 743 | 3891 |
| PBANKA_1226000 | 0 | 0 | 97 | 19 | 102 | 42 | 99 | 29 | 115 | 52 | 217 | 146 | 2207 | 1679 | 530 | 2678 | 1207 | 277 | 624 | 832 | 74 | 674 |
| PBANKA_1226100 | 507 | 715 | 689 | 173 | 1784 | 636 | 2368 | 764 | 1215 | 558 | 348 | 127 | 3449 | 7182 | 4979 | 24690 | 17254 | 2024 | 1879 | 11573 | 4047 | 22968 |
| PBANKA_1226200 | 31 | 41 | 152 | 27 | 430 | 197 | 484 | 179 | 150 | 63 | 23 | 9 | 570 | 156 | 576 | 2515 | 1639 | 227 | 431 | 315 | 603 | 3726 |
| PBANKA_1226300 | 62 | 70 | 45 | 0 | 123 | 35 | 158 | 34 | 109 | 47 | 675 | 286 | 1187 | 885 | 235 | 1205 | 768 | 96 | 608 | 541 | 621 | 3986 |
| PBANKA_1226400 | 47 | 55 | 260 | 56 | 476 | 176 | 472 | 171 | 187 | 110 | 690 | 473 | 10528 | 5910 | 1913 | 9890 | 5032 | 653 | 2142 | 1793 | 624 | 4245 |
| PBANKA_1226500 | 0 | 0 | 226 | 48 | 286 | 203 | 424 | 207 | 172 | 79 | 42 | 13 | 1324 | 291 | 1116 | 3155 | 2211 | 249 | 440 | 272 | 584 | 5145 |
| PBANKA_1226600 | 0 | 0 | 115 | 38 | 276 | 146 | 286 | 101 | 154 | 60 | 27 | 5 | 617 | 174 | 554 | 2090 | 1064 | 157 | 1280 | 614 | 24 | 373 |
| PBANKA_1226700 | 64 | 108 | 90 | 14 | 149 | 49 | 135 | 34 | 279 | 113 | 152 | 61 | 1876 | 1201 | 665 | 3615 | 1155 | 467 | 3793 | 3127 | 198 | 1007 |
| PBANKA_1226800 | 0 | 0 | 86 | 38 | 345 | 181 | 474 | 198 | 218 | 131 | 34 | 7 | 584 | 397 | 632 | 2950 | 6204 | 632 | 1598 | 4666 | 4517 | 31396 |
| PBANKA_1226900 | 23 | 15 | 81 | 30 | 269 | 154 | 312 | 149 | 273 | 207 | 44 | 19 | 391 | 74 | 174 | 646 | 1881 | 496 | 623 | 343 | 159 | 1116 |
| PBANKA_1227000 | 54 | 133 | 46 | 10 | 239 | 80 | 306 | 101 | 444 | 210 | 85 | 28 | 742 | 332 | 100 | 430 | 8881 | 1976 | 2109 | 3775 | 229 | 1588 |
| PBANKA_1227100 | 58 | 67 | 184 | 45 | 214 | 81 | 159 | 44 | 205 | 81 | 2497 | 1385 | 5461 | 6448 | 1054 | 7414 | 3329 | 599 | 1367 | 4011 | 1094 | 5608 |
| PBANKA_1227200 | 337 | 521 | 64 | 9 | 235 | 107 | 357 | 146 | 183 | 85 | 58 | 19 | 1774 | 1171 | 427 | 2366 | 2467 | 375 | 1750 | 2246 | 971 | 5798 |
| PBANKA_1227300 | 76 | 84 | 35 | 8 | 61 | 33 | 73 | 32 | 22 | 12 | 6 | 0 | 96 | 61 | 50 | 132 | 123 | 46 | 273 | 87 | 61 | 484 |
| PBANKA_1227400 | 138 | 167 | 46 | 16 | 162 | 68 | 260 | 119 | 107 | 67 | 5 | 7 | 1096 | 161 | 572 | 2716 | 3395 | 573 | 4046 | 2082 | 353 | 3066 |
| PBANKA_1227600 | 187 | 323 | 164 | 42 | 355 | 105 | 357 | 117 | 356 | 137 | 132 | 40 | 449 | 321 | 129 | 538 | 5374 | 1177 | 1261 | 1590 | 296 | 1995 |
| PBANKA_1227700 | 5 | 0 | 49 | 5 | 189 | 72 | 226 | 89 | 243 | 91 | 63 | 20 | 1080 | 606 | 446 | 2423 | 2593 | 537 | 1170 | 1538 | 60 | 373 |
| PBANKA_1227800 | 198 | 307 | 327 | 69 | 562 | 159 | 650 | 205 | 570 | 278 | 1349 | 774 | 5150 | 5951 | 1842 | 10136 | 3915 | 891 | 976 | 2618 | 669 | 3030 |
| PBANKA_1227900 | 33 | 43 | 179 | 51 | 252 | 128 | 252 | 58 | 282 | 118 | 1649 | 1070 | 1883 | 2380 | 525 | 3606 | 807 | 89 | 336 | 726 | 339 | 2352 |
| PBANKA_1228000 | 16 | 16 | 198 | 46 | 466 | 287 | 657 | 306 | 345 | 223 | 37 | 23 | 902 | 361 | 317 | 1100 | 298 | 62 | 664 | 353 | 252 | 1915 |
| PBANKA_1228100 | 43 | 106 | 152 | 33 | 1448 | 534 | 2996 | 1048 | 2635 | 1166 | 435 | 175 | 2582 | 2786 | 1038 | 5304 | 51412 | 5696 | 4372 | 14852 | 2747 | 18835 |
| PBANKA_1228200 | 11 | 20 | 79 | 8 | 266 | 159 | 237 | 92 | 128 | 61 | 10 | 10 | 1422 | 366 | 1305 | 5224 | 2829 | 474 | 3907 | 3258 | 406 | 3731 |
| PBANKA_1228300 | 2552 | 3669 | 4647 | 1154 | 8324 | 3056 | 5640 | 1577 | 2329 | 1081 | 573 | 241 | 740 | 2372 | 921 | 5855 | 2714 | 696 | 3164 | 37911 | 18178 | 2E+05 |
| PBANKA_1228400 | 124 | 105 | 140 | 31 | 1348 | 410 | 1256 | 498 | 551 | 192 | 93 | 49 | 5201 | 2751 | 1580 | 10872 | 5016 | 559 | 1234 | 1395 | 109 | 477 |
| PBANKA_1228500 | 21 | 46 | 98 | 19 | 325 | 95 | 427 | 169 | 224 | 103 | 30 | 30 | 1170 | 500 | 1139 | 5604 | 3848 | 540 | 1726 | 6305 | 145 | 1085 |
| PBANKA_1228600 | 0 | 0 | 94 | 25 | 196 | 72 | 231 | 88 | 67 | 33 | 0 | 8 | 192 | 28 | 303 | 584 | 545 | 120 | 365 | 170 | 33 | 311 |
| PBANKA_1228700 | 31 | 72 | 84 | 21 | 121 | 59 | 108 | 41 | 143 | 73 | 797 | 548 | 1131 | 1018 | 243 | 1235 | 399 | 164 | 726 | 738 | 378 | 2780 |
| PBANKA_1229000 | 12 | 22 | 59 | 24 | 152 | 118 | 137 | 75 | 685 | 423 | 20206 | 14691 | 8611 | 10890 | 2832 | 10935 | 2137 | 296 | 575 | 763 | 10 | 206 |
| PBANKA_1229100 | 24 | 32 | 47 | 6 | 133 | 55 | 179 | 61 | 42 | 19 | 22 | 10 | 1275 | 547 | 310 | 1560 | 850 | 151 | 775 | 498 | 215 | 1359 |
| PBANKA_1229200 | 6 | 15 | 1856 | 466 | 1245 | 599 | 1313 | 518 | 509 | 261 | 2355 | 1883 | 26484 | 9779 | 3644 | 14131 | 4127 | 675 | 3429 | 1781 | 98 | 739 |
| PBANKA_1229300 | 8 | 12 | 43 | 8 | 47 | 10 | 50 | 22 | 78 | 28 | 30 | 11 | 488 | 106 | 135 | 821 | 757 | 156 | 1311 | 1097 | 121 | 843 |
| PBANKA_1229500 | 8 | 11 | 46 | 13 | 125 | 44 | 133 | 40 | 81 | 31 | 25 | 11 | 599 | 483 | 173 | 977 | 568 | 117 | 1175 | 981 | 265 | 2086 |
| PBANKA_1229600 | 23 | 35 | 41 | 7 | 151 | 51 | 273 | 79 | 259 | 95 | 38 | 14 | 1798 | 987 | 392 | 2132 | 4337 | 838 | 2088 | 1794 | 413 | 2857 |
| PBANKA_1229700 | 5 | 14 | 27 | 6 | 37 | 13 | 40 | 20 | 19 | 10 | 53 | 43 | 783 | 441 | 129 | 981 | 399 | 61 | 358 | 298 | 127 | 666 |
| PBANKA_1229800 | 14 | 22 | 138 | 43 | 458 | 218 | 541 | 239 | 127 | 90 | 32 | 0 | 168 | 82 | 31 | 99 | 316 | 66 | 1722 | 1088 | 161 | 1162 |
| PBANKA_1229900 | 36 | 59 | 12 | 0 | 95 | 32 | 84 | 12 | 19 | 14 | 0 | 0 | 533 | 321 | 31 | 59 | 283 | 169 | 14105 | 9047 | 548 | 3293 |
| PBANKA_1230000 | 0 | 5 | 14 | 0 | 79 | 28 | 132 | 49 | 58 | 28 | 0 | 5 | 202 | 87 | 193 | 510 | 562 | 99 | 612 | 324 | 30 | 153 |
| PBANKA_1230100 | 736 | 786 | 466 | 107 | 1136 | 436 | 1972 | 709 | 1123 | 444 | 262 | 92 | 5082 | 9657 | 2692 | 13990 | 45044 | 5082 | 2762 | 17495 | 1688 | 10773 |
| PBANKA_1230300 | 20 | 32 | 198 | 67 | 468 | 200 | 594 | 245 | 325 | 130 | 121 | 48 | 1062 | 957 | 948 | 4932 | 1819 | 220 | 1249 | 1779 | 60 | 385 |
| PBANKA_1230500 | 9 | 15 | 106 | 38 | 142 | 122 | 193 | 122 | 84 | 59 | 548 | 309 | 788 | 339 | 254 | 822 | 234 | 46 | 740 | 263 | 51 | 335 |
| PBANKA_1230600 | 0 | 0 | 21 | 0 | 111 | 29 | 118 | 51 | 77 | 27 | 94 | 54 | 579 | 221 | 193 | 955 | 548 | 46 | 1203 | 560 | 502 | 2576 |
| PBANKA_1230700 | 0 | 0 | 50 | 9 | 225 | 85 | 269 | 106 | 154 | 58 | 28 | 9 | 365 | 225 | 536 | 2159 | 2349 | 232 | 174 | 535 | 350 | 2419 |
| PBANKA_1230800 | 0 | 0 | 97 | 22 | 91 | 61 | 82 | 41 | 54 | 30 | 8 | 5 | 397 | 86 | 176 | 673 | 203 | 25 | 330 | 126 | 280 | 1661 |
| PBANKA_1230900 | 0 | 0 | 111 | 25 | 114 | 52 | 34 | 26 | 48 | 19 | 6 | 7 | 1434 | 488 | 369 | 1774 | 151 | 33 | 230 | 174 | 180 | 1226 |
| PBANKA_1231000 | 67 | 93 | 1777 | 720 | 1571 | 1071 | 1140 | 677 | 479 | 250 | 2089 | 1680 | 5424 | 1958 | 1960 | 4485 | 1136 | 219 | 1094 | 642 | 73 | 454 |
| PBANKA_1231100 | 0 | 0 | 42 | 16 | 122 | 40 | 86 | 37 | 36 | 22 | 58 | 24 | 3827 | 1682 | 267 | 1727 | 539 | 105 | 1157 | 818 | 55 | 462 |
| PBANKA_1231200 | 0 | 0 | 68 | 12 | 204 | 58 | 176 | 47 | 52 | 24 | 19 | 7 | 947 | 227 | 393 | 1895 | 857 | 146 | 1844 | 801 | 491 | 2917 |
| PBANKA_1231300 | 0 | 0 | 26 | 13 | 145 | 104 | 272 | 92 | 81 | 44 | 13 | 0 | 1849 | 539 | 71 | 268 | 932 | 463 | 47662 | 22160 | 361 | 3035 |
| PBANKA_1231400 | 18 | 30 | 123 | 23 | 504 | 139 | 469 | 163 | 275 | 133 | 56 | 30 | 2800 | 2182 | 559 | 3260 | 5976 | 978 | 2749 | 3808 | 379 | 2544 |
| PBANKA_1231500 | 0 | 0 | 121 | 24 | 574 | 311 | 871 | 314 | 375 | 160 | 54 | 30 | 2892 | 798 | 1472 | 7056 | 3999 | 544 | 5050 | 6663 | 416 | 2826 |
| PBANKA_1231600 | 6 | 14 | 86 | 15 | 1812 | 539 | 3785 | 1302 | 2188 | 892 | 8784 | 5011 | 6571 | 15600 | 2531 | 13837 | 14008 | 1942 | 3102 | 13815 | 427 | 3154 |
| PBANKA_1231700 | 115 | 169 | 3554 | 1170 | 2430 | 1453 | 2578 | 1279 | 1174 | 774 | 4411 | 3276 | 23884 | 9060 | 5085 | 20925 | 5632 | 904 | 3743 | 1941 | 364 | 2842 |
| PBANKA_1231800 | 64 | 110 | 1321 | 397 | 854 | 561 | 921 | 530 | 347 | 273 | 1064 | 991 | 11027 | 3969 | 3190 | 7104 | 2958 | 813 | 2344 | 829 | 145 | 1217 |
| PBANKA_1231900 | 0 | 0 | 51 | 0 | 50 | 15 | 24 | 15 | 15 | 5 | 7 | 0 | 161 | 58 | 66 | 366 | 56 | 13 | 426 | 159 | 66 | 331 |
| PBANKA_1232100 | 0 | 0 | 378 | 149 | 406 | 326 | 388 | 252 | 396 | 377 | 2711 | 2797 | 6248 | 3390 | 2238 | 8890 | 3618 | 520 | 1223 | 990 | 79 | 983 |
| PBANKA_1232200 | 0 | 0 | 373 | 126 | 1156 | 475 | 1376 | 518 | 541 | 181 | 156 | 73 | 1998 | 1247 | 2842 | 16307 | 15057 | 1592 | 3746 | 8305 | 84 | 702 |
| PBANKA_1232400 | 21 | 35 | 7 | 0 | 20 | 5 | 24 | 5 | 23 | 0 | 8 | 8 | 150 | 67 | 26 | 83 | 98 | 49 | 902 | 364 | 70 | 561 |
| PBANKA_1232500 | 101 | 165 | 1150 | 333 | 1659 | 807 | 1479 | 557 | 390 | 198 | 833 | 651 | 6310 | 5566 | 3354 | 14987 | 2995 | 473 | 1996 | 2789 | 639 | 5792 |
| PBANKA_1232700 | 14 | 23 | 16 | 5 | 140 | 70 | 267 | 96 | 120 | 83 | 11 | 8 | 360 | 103 | 43 | 163 | 2242 | 414 | 442 | 295 | 191 | 1566 |
| PBANKA_1232800 | 6 | 15 | 47 | 20 | 156 | 97 | 229 | 119 | 129 | 75 | 6 | 6 | 479 | 111 | 88 | 309 | 757 | 162 | 444 | 283 | 514 | 3298 |
| PBANKA_1232900 | 67 | 76 | 164 | 20 | 361 | 146 | 317 | 110 | 166 | 94 | 36 | 21 | 1036 | 861 | 935 | 4581 | 2808 | 434 | 1409 | 2643 | 822 | 5418 |
| PBANKA_1233000 | 16 | 22 | 24 | 0 | 130 | 74 | 305 | 143 | 153 | 94 | 11 | 6 | 452 | 75 | 64 | 163 | 1137 | 278 | 1534 | 574 | 21 | 161 |
| PBANKA_1233100 | 0 | 0 | 231 | 60 | 440 | 272 | 602 | 270 | 205 | 136 | 15 | 16 | 1247 | 300 | 1620 | 3823 | 2184 | 371 | 1007 | 538 | 124 | 880 |
| PBANKA_1233200 | 0 | 0 | 5 | 0 | 0 | 10 | 15 | 0 | 13 | 9 | 0 | 0 | 148 | 52 | 14 | 28 | 121 | 48 | 4105 | 1459 | 1448 | 8192 |
| PBANKA_1233300 | 155 | 176 | 230 | 53 | 505 | 294 | 633 | 257 | 264 | 98 | 200 | 168 | 3744 | 3694 | 1468 | 9171 | 6189 | 611 | 1589 | 2163 | 1545 | 12066 |
| PBANKA_1233400 | 335 | 496 | 82 | 9 | 1098 | 381 | 993 | 346 | 358 | 174 | 83 | 31 | 1225 | 841 | 726 | 3732 | 1168 | 283 | 2928 | 4440 | 352 | 3050 |
| PBANKA_1233800 | 31 | 49 | 68 | 16 | 272 | 157 | 217 | 106 | 55 | 56 | 14 | 7 | 243 | 39 | 366 | 1403 | 233 | 44 | 529 | 349 | 163 | 1186 |
| PBANKA_1233900 | 0 | 0 | 58 | 20 | 96 | 47 | 93 | 44 | 23 | 20 | 0 | 0 | 224 | 54 | 368 | 1984 | 565 | 112 | 977 | 833 | 146 | 1261 |
| PBANKA_1234000 | 320 | 328 | 242 | 96 | 335 | 251 | 587 | 423 | 420 | 411 | 79 | 57 | 2371 | 734 | 1030 | 1876 | 2398 | 1379 | 1365 | 716 | 880 | 5304 |
| PBANKA_1234100 | 17 | 11 | 41 | 11 | 125 | 51 | 181 | 76 | 112 | 56 | 23 | 13 | 916 | 315 | 338 | 1295 | 1674 | 259 | 2270 | 1373 | 701 | 5031 |
| PBANKA_1234200 | 16 | 23 | 1543 | 471 | 1080 | 658 | 1381 | 615 | 438 | 332 | 2070 | 1750 | 21146 | 5728 | 3118 | 9421 | 4563 | 845 | 3420 | 1543 | 331 | 2297 |
| PBANKA_1234300 | 0 | 7 | 204 | 50 | 597 | 219 | 957 | 308 | 551 | 170 | 293 | 140 | 2907 | 2217 | 860 | 4328 | 8860 | 1327 | 1848 | 2367 | 127 | 1069 |
| PBANKA_1234400 | 0 | 0 | 90 | 7 | 152 | 74 | 267 | 117 | 71 | 49 | 45 | 0 | 834 | 215 | 487 | 2600 | 1371 | 137 | 1230 | 890 | 217 | 1462 |
| PBANKA_1234500 | 127 | 117 | 119 | 44 | 176 | 115 | 490 | 255 | 214 | 155 | 402 | 319 | 2366 | 904 | 558 | 1888 | 1882 | 581 | 1766 | 823 | 418 | 2545 |
| PBANKA_1234600 | 244 | 407 | 93 | 22 | 328 | 124 | 343 | 112 | 176 | 57 | 69 | 45 | 4238 | 3780 | 1322 | 7438 | 2553 | 246 | 630 | 1045 | 3544 | 24440 |
| PBANKA_1234700 | 0 | 7 | 47 | 6 | 148 | 32 | 231 | 58 | 130 | 20 | 49 | 5 | 384 | 108 | 101 | 634 | 473 | 73 | 438 | 258 | 465 | 3380 |
| PBANKA_1234800 | 19 | 13 | 1352 | 514 | 867 | 530 | 770 | 469 | 299 | 231 | 1828 | 1708 | 13042 | 3930 | 2491 | 8054 | 2560 | 571 | 3658 | 1379 | 147 | 1005 |
| PBANKA_1234900 | 12 | 10 | 109 | 13 | 270 | 92 | 358 | 138 | 471 | 166 | 125 | 27 | 1404 | 1014 | 517 | 3281 | 1496 | 175 | 2596 | 2036 | 196 | 1093 |
| PBANKA_1235000 | 0 | 0 | 41 | 0 | 94 | 49 | 89 | 33 | 51 | 13 | 31 | 9 | 706 | 499 | 279 | 1493 | 522 | 82 | 761 | 730 | 389 | 2518 |
| PBANKA_1235100 | 0 | 0 | 42 | 19 | 125 | 55 | 105 | 56 | 26 | 15 | 10 | 11 | 774 | 210 | 355 | 1065 | 515 | 84 | 368 | 165 | 73 | 377 |
| PBANKA_1235200 | 38 | 46 | 47 | 12 | 232 | 82 | 364 | 144 | 135 | 56 | 7 | 9 | 371 | 95 | 287 | 925 | 1006 | 192 | 852 | 302 | 159 | 1006 |
| PBANKA_1235300 | 0 | 6 | 29 | 6 | 66 | 39 | 123 | 49 | 22 | 23 | 0 | 5 | 644 | 255 | 296 | 1452 | 1080 | 191 | 844 | 358 | 10 | 68 |
| PBANKA_1235400 | 48 | 61 | 91 | 7 | 158 | 48 | 151 | 37 | 71 | 37 | 49 | 7 | 1228 | 563 | 345 | 1609 | 735 | 112 | 1572 | 754 | 42 | 227 |

| | | | | | | | | | | | | | | | | | | | | | | |
|---|---|---|---|---|---|---|---|---|---|---|---|---|---|---|---|---|---|---|---|---|---|---|
| PBANKA_1235500 | 72 | 97 | 171 | 46 | 224 | 67 | 224 | 73 | 160 | 49 | 76 | 25 | 915 | 595 | 660 | 3579 | 892 | 113 | 333 | 639 | 54 | 414 |
| PBANKA_1235600 | 26 | 28 | 748 | 218 | 802 | 565 | 666 | 282 | 286 | 147 | 90 | 51 | 13784 | 4710 | 8146 | 44035 | 5231 | 712 | 1026 | 1119 | 99 | 806 |
| PBANKA_1235700 | 0 | 0 | 11 | 0 | 40 | 24 | 53 | 18 | 39 | 15 | 5 | 0 | 144 | 36 | 153 | 536 | 463 | 81 | 872 | 488 | 60 | 389 |
| PBANKA_1235800 | 7 | 14 | 31 | 10 | 77 | 28 | 57 | 20 | 22 | 14 | 13 | 0 | 98 | 27 | 111 | 496 | 118 | 48 | 604 | 209 | 63 | 315 |
| PBANKA_1235900 | 68 | 102 | 111 | 12 | 230 | 77 | 285 | 86 | 186 | 69 | 41 | 15 | 1554 | 833 | 658 | 3500 | 2636 | 418 | 1350 | 1690 | 377 | 2239 |
| PBANKA_1236000 | 6 | 7 | 75 | 19 | 204 | 96 | 189 | 68 | 72 | 29 | 16 | 0 | 701 | 251 | 308 | 1381 | 550 | 59 | 139 | 64 | 410 | 3010 |
| PBANKA_1236100 | 9 | 17 | 41 | 10 | 158 | 60 | 163 | 80 | 64 | 8 | 21 | 17 | 387 | 145 | 138 | 732 | 366 | 45 | 98 | 66 | 506 | 2974 |
| PBANKA_1236300 | 85 | 110 | 113 | 19 | 384 | 173 | 460 | 129 | 130 | 68 | 34 | 6 | 430 | 333 | 475 | 2442 | 824 | 122 | 529 | 703 | 144 | 880 |
| PBANKA_1236400 | 7 | 21 | 93 | 15 | 1685 | 720 | 2013 | 691 | 549 | 219 | 49 | 29 | 479 | 143 | 481 | 2379 | 1454 | 297 | 1853 | 911 | 277 | 1508 |
| PBANKA_1236600 | 55 | 50 | 10 | 0 | 21 | 8 | 38 | 24 | 55 | 37 | 18 | 13 | 534 | 735 | 155 | 948 | 3208 | 415 | 1750 | 10194 | 104 | 492 |
| PBANKA_1236700 | 89 | 56 | 20 | 0 | 46 | 23 | 55 | 27 | 18 | 7 | 7 | 0 | 126 | 36 | 76 | 160 | 76 | 54 | 219 | 114 | 36 | 201 |
| PBANKA_1236800 | 5 | 0 | 6 | 0 | 109 | 49 | 222 | 45 | 836 | 516 | 265 | 104 | 399 | 268 | 69 | 229 | 2407 | 738 | 2039 | 1080 | 283 | 1534 |
| PBANKA_1236900 | 8 | 5 | 32 | 0 | 60 | 38 | 85 | 34 | 119 | 122 | 146 | 166 | 387 | 104 | 167 | 380 | 381 | 398 | 682 | 261 | 114 | 590 |
| PBANKA_1237000 | 166 | 217 | 87 | 50 | 214 | 125 | 145 | 74 | 228 | 155 | 450 | 349 | 1153 | 614 | 205 | 774 | 494 | 184 | 1416 | 511 | 179 | 998 |
| PBANKA_1237100 | 40 | 83 | 38 | 8 | 2069 | 680 | 4955 | 1846 | 3453 | 1428 | 361 | 205 | 1668 | 3923 | 970 | 4092 | 61531 | 8124 | 1234 | 21393 | 375 | 2948 |
| PBANKA_1237200 | 0 | 0 | 34 | 20 | 71 | 50 | 104 | 51 | 18 | 19 | 5 | 0 | 287 | 56 | 206 | 502 | 408 | 83 | 1045 | 658 | 39 | 354 |
| PBANKA_1237400 | 0 | 0 | 33 | 5 | 55 | 24 | 48 | 17 | 13 | 5 | 0 | 0 | 68 | 5 | 74 | 322 | 75 | 33 | 87 | 20 | 0 | 26 |
| PBANKA_1237500 | 0 | 0 | 94 | 15 | 117 | 83 | 82 | 27 | 21 | 15 | 32 | 21 | 475 | 209 | 183 | 770 | 113 | 20 | 228 | 139 | 13 | 72 |
| PBANKA_1237600 | 0 | 0 | 60 | 12 | 291 | 148 | 285 | 98 | 151 | 48 | 31 | 19 | 959 | 699 | 315 | 1635 | 1668 | 302 | 7957 | 24890 | 3551 | 28404 |
| PBANKA_1237800 | 0 | 0 | 5 | 0 | 39 | 37 | 65 | 53 | 323 | 275 | 2230 | 2492 | 10479 | 9079 | 6925 | 32148 | 5424 | 923 | 14951 | 22227 | 328 | 3692 |
| PBANKA_1237900 | 0 | 0 | 334 | 71 | 636 | 426 | 1106 | 481 | 422 | 239 | 79 | 21 | 1583 | 298 | 992 | 3491 | 3328 | 592 | 1421 | 920 | 54 | 503 |
| PBANKA_1238000 | 0 | 0 | 53 | 11 | 92 | 52 | 124 | 38 | 56 | 33 | 11 | 8 | 343 | 57 | 406 | 1318 | 833 | 128 | 941 | 377 | 17 | 144 |
| PBANKA_1238100 | 67 | 93 | 37 | 11 | 123 | 75 | 179 | 112 | 108 | 79 | 17 | 7 | 346 | 133 | 118 | 537 | 1470 | 220 | 2229 | 1271 | 86 | 662 |
| PBANKA_1238200 | 544 | 787 | 252 | 67 | 645 | 402 | 650 | 219 | 414 | 277 | 733 | 590 | 3214 | 1040 | 1040 | 4875 | 1998 | 376 | 8702 | 4375 | 612 | 4938 |
| PBANKA_1238400 | 0 | 0 | 40 | 5 | 117 | 84 | 206 | 108 | 58 | 58 | 6 | 7 | 248 | 54 | 223 | 706 | 834 | 148 | 1303 | 426 | 30 | 256 |
| PBANKA_1238500 | 0 | 0 | 40 | 14 | 102 | 25 | 106 | 31 | 52 | 23 | 21 | 0 | 852 | 504 | 234 | 1052 | 366 | 81 | 4775 | 2232 | 74 | 473 |
| PBANKA_1238600 | 15 | 23 | 78 | 22 | 158 | 110 | 222 | 100 | 96 | 47 | 117 | 152 | 2828 | 1196 | 621 | 3308 | 1490 | 189 | 1750 | 1142 | 185 | 1617 |
| PBANKA_1238700 | 7 | 6 | 146 | 39 | 259 | 239 | 378 | 159 | 89 | 84 | 40 | 26 | 840 | 385 | 340 | 910 | 337 | 69 | 694 | 360 | 83 | 842 |
| PBANKA_1238800 | 43 | 72 | 1180 | 355 | 1304 | 889 | 1855 | 752 | 685 | 328 | 2491 | 2080 | 28698 | 16008 | 7555 | 35226 | 10585 | 1241 | 8040 | 6428 | 1165 | 10463 |
| PBANKA_1238900 | 198 | 300 | 97 | 21 | 255 | 85 | 233 | 74 | 852 | 484 | 317 | 140 | 1920 | 992 | 556 | 2596 | 4289 | 1248 | 1673 | 2523 | 902 | 5547 |
| PBANKA_1239000 | 31 | 48 | 32 | 8 | 138 | 76 | 153 | 63 | 85 | 65 | 7 | 7 | 1575 | 381 | 50 | 50 | 707 | 559 | 48998 | 17936 | 1072 | 10408 |
| PBANKA_1239100 | 0 | 0 | 28 | 0 | 111 | 33 | 91 | 33 | 33 | 19 | 62 | 45 | 1962 | 941 | 450 | 2101 | 365 | 76 | 679 | 390 | 21 | 73 |
| PBANKA_1239200 | 89 | 102 | 111 | 30 | 230 | 91 | 181 | 67 | 52 | 20 | 47 | 41 | 843 | 466 | 253 | 1230 | 455 | 66 | 280 | 123 | 82 | 610 |
| PBANKA_1239400 | 0 | 0 | 17 | 6 | 36 | 15 | 61 | 31 | 26 | 11 | 0 | 0 | 369 | 99 | 171 | 858 | 769 | 116 | 2670 | 1581 | 110 | 702 |
| PBANKA_1239500 | 35 | 40 | 92 | 29 | 106 | 78 | 119 | 90 | 28 | 21 | 8 | 9 | 1070 | 418 | 276 | 1115 | 610 | 94 | 377 | 231 | 231 | 1723 |
| PBANKA_1239600 | 21 | 27 | 71 | 7 | 217 | 57 | 220 | 56 | 111 | 41 | 16 | 6 | 1074 | 330 | 289 | 1208 | 1014 | 216 | 1202 | 525 | 78 | 473 |
| PBANKA_1239700 | 40 | 73 | 89 | 41 | 257 | 152 | 204 | 72 | 69 | 57 | 369 | 222 | 1184 | 897 | 450 | 2221 | 402 | 56 | 327 | 345 | 24 | 207 |
| PBANKA_1239800 | 144 | 213 | 173 | 59 | 505 | 322 | 565 | 230 | 339 | 135 | 806 | 606 | 9983 | 7866 | 1277 | 7908 | 4296 | 538 | 1709 | 2010 | 313 | 2719 |
| PBANKA_1239900 | 15 | 18 | 302 | 87 | 573 | 350 | 1311 | 537 | 1228 | 590 | 162 | 127 | 3926 | 1089 | 2035 | 8456 | 11218 | 1941 | 4102 | 2699 | 703 | 5320 |
| PBANKA_1240000 | 151 | 235 | 219 | 50 | 601 | 258 | 706 | 280 | 345 | 151 | 42 | 28 | 2094 | 1027 | 1637 | 6886 | 18838 | 2529 | 1575 | 2807 | 150 | 1197 |
| PBANKA_1240100 | 117 | 194 | 17 | 0 | 79 | 41 | 108 | 55 | 201 | 113 | 72 | 35 | 793 | 463 | 36 | 133 | 1050 | 389 | 17654 | 11944 | 1401 | 8002 |
| PBANKA_1240200 | 9 | 9 | 83 | 17 | 408 | 110 | 538 | 160 | 315 | 133 | 55 | 11 | 802 | 251 | 428 | 2119 | 2965 | 582 | 1508 | 837 | 93 | 409 |
| PBANKA_1240300 | 6 | 8 | 77 | 12 | 167 | 65 | 180 | 57 | 92 | 38 | 124 | 62 | 926 | 578 | 382 | 1749 | 393 | 58 | 278 | 161 | 17 | 127 |
| PBANKA_1240400 | 0 | 0 | 14 | 0 | 109 | 43 | 99 | 30 | 31 | 14 | 17 | 8 | 186 | 52 | 126 | 500 | 178 | 31 | 374 | 299 | 75 | 482 |
| PBANKA_1240500 | 0 | 0 | 27 | 5 | 52 | 33 | 41 | 27 | 7 | 15 | 0 | 0 | 145 | 35 | 75 | 332 | 149 | 62 | 125 | 66 | 19 | 109 |
| PBANKA_1240600 | 122 | 160 | 34 | 18 | 143 | 62 | 144 | 61 | 1985 | 1779 | 501 | 289 | 1082 | 457 | 26 | 87 | 2968 | 2270 | 7018 | 4352 | 7624 | 57423 |
| PBANKA_1240800 | 0 | 0 | 34 | 8 | 51 | 16 | 54 | 13 | 19 | 0 | 5 | 8 | 400 | 222 | 157 | 710 | 82 | 23 | 620 | 274 | 52 | 335 |
| PBANKA_1240900 | 0 | 0 | 27 | 7 | 41 | 11 | 25 | 18 | 22 | 11 | 14 | 5 | 218 | 86 | 61 | 369 | 31 | 17 | 282 | 108 | 37 | 258 |
| PBANKA_1241000 | 5 | 14 | 59 | 13 | 119 | 99 | 134 | 68 | 56 | 16 | 30 | 7 | 513 | 153 | 255 | 1382 | 383 | 66 | 2196 | 1042 | 1007 | 6920 |
| PBANKA_1241100 | 0 | 0 | 68 | 10 | 167 | 71 | 316 | 154 | 170 | 75 | 30 | 12 | 313 | 82 | 115 | 258 | 1097 | 230 | 3069 | 1854 | 64 | 579 |
| PBANKA_1241200 | 0 | 0 | 53 | 0 | 129 | 65 | 239 | 71 | 76 | 52 | 0 | 7 | 121 | 31 | 97 | 386 | 882 | 130 | 306 | 289 | 43 | 296 |
| PBANKA_1241300 | 10 | 26 | 187 | 35 | 659 | 186 | 810 | 252 | 706 | 253 | 146 | 52 | 2936 | 1073 | 1223 | 7223 | 9963 | 1475 | 3199 | 4313 | 1013 | 5645 |
| PBANKA_1241400 | 0 | 0 | 12 | 0 | 6 | 0 | 0 | 0 | 0 | 0 | 0 | 0 | 10 | 0 | 22 | 89 | 11 | 5 | 214 | 152 | 886 | 5170 |
| PBANKA_1241500 | 47 | 57 | 185 | 55 | 421 | 221 | 546 | 252 | 336 | 131 | 98 | 42 | 725 | 595 | 652 | 3897 | 3312 | 397 | 231 | 491 | 1884 | 12784 |
| PBANKA_1241600 | 0 | 0 | 24 | 7 | 73 | 27 | 101 | 23 | 38 | 12 | 5 | 0 | 345 | 111 | 106 | 420 | 326 | 66 | 190 | 88 | 8 | 77 |
| PBANKA_1241700 | 58 | 71 | 103 | 19 | 230 | 91 | 233 | 138 | 100 | 42 | 15 | 9 | 654 | 193 | 217 | 709 | 347 | 93 | 907 | 307 | 44 | 285 |
| PBANKA_1241800 | 393 | 515 | 333 | 122 | 1681 | 956 | 4222 | 1736 | 2663 | 1302 | 253 | 149 | 3299 | 926 | 621 | 2127 | 21016 | 2706 | 8153 | 4489 | 81 | 940 |
| PBANKA_1241900 | 0 | 0 | 210 | 58 | 417 | 204 | 445 | 184 | 123 | 79 | 25 | 18 | 704 | 198 | 731 | 1420 | 686 | 196 | 523 | 260 | 26 | 127 |
| PBANKA_1242000 | 5 | 10 | 201 | 40 | 364 | 198 | 606 | 277 | 249 | 128 | 25 | 9 | 1010 | 220 | 761 | 2435 | 2053 | 416 | 1213 | 1079 | 124 | 1169 |
| PBANKA_1242100 | 25 | 25 | 38 | 5 | 160 | 111 | 185 | 64 | 49 | 34 | 57 | 20 | 514 | 217 | 227 | 820 | 264 | 94 | 3520 | 1826 | 409 | 3093 |
| PBANKA_1242200 | 0 | 0 | 12 | 0 | 42 | 13 | 69 | 20 | 42 | 25 | 222 | 136 | 1614 | 628 | 421 | 1366 | 379 | 101 | 584 | 207 | 13 | 93 |
| PBANKA_1242300 | 55 | 80 | 1061 | 262 | 1249 | 506 | 1014 | 323 | 320 | 134 | 404 | 236 | 15613 | 7341 | 5167 | 24028 | 5133 | 703 | 5935 | 4042 | 398 | 2667 |
| PBANKA_1242400 | 23 | 30 | 126 | 26 | 723 | 362 | 828 | 358 | 278 | 167 | 53 | 26 | 556 | 181 | 176 | 590 | 2022 | 320 | 2769 | 2236 | 178 | 1519 |
| PBANKA_1242500 | 5 | 0 | 84 | 18 | 313 | 167 | 432 | 154 | 175 | 64 | 233 | 151 | 2278 | 1452 | 1000 | 6524 | 3633 | 386 | 1567 | 2167 | 170 | 1271 |
| PBANKA_1242600 | 26 | 28 | 141 | 40 | 254 | 170 | 377 | 199 | 114 | 74 | 19 | 6 | 1063 | 184 | 770 | 1888 | 1033 | 234 | 992 | 412 | 265 | 1916 |
| PBANKA_1242700 | 0 | 0 | 103 | 30 | 337 | 142 | 560 | 215 | 264 | 139 | 67 | 52 | 2913 | 1165 | 971 | 3483 | 3672 | 580 | 1376 | 791 | 758 | 4880 |
| PBANKA_1242800 | 39 | 72 | 530 | 112 | 683 | 349 | 617 | 184 | 214 | 103 | 866 | 491 | 4487 | 2475 | 1142 | 4212 | 1128 | 208 | 935 | 627 | 178 | 1225 |
| PBANKA_1242900 | 77 | 100 | 15 | 0 | 43 | 17 | 80 | 26 | 109 | 72 | 61 | 27 | 308 | 143 | 54 | 186 | 641 | 207 | 360 | 192 | 1044 | 5871 |
| PBANKA_1243100 | 9 | 7 | 9 | 6 | 55 | 23 | 112 | 50 | 81 | 41 | 41 | 12 | 168 | 48 | 28 | 59 | 265 | 82 | 2682 | 999 | 186 | 1157 |
| PBANKA_1243200 | 32 | 37 | 37 | 9 | 54 | 31 | 66 | 39 | 42 | 37 | 238 | 161 | 733 | 298 | 139 | 518 | 404 | 120 | 241 | 105 | 298 | 1742 |
| PBANKA_1243300 | 8 | 10 | 23 | 8 | 82 | 47 | 59 | 22 | 44 | 16 | 42 | 46 | 319 | 214 | 209 | 649 | 130 | 17 | 155 | 139 | 135 | 1343 |
| PBANKA_1243400 | 38 | 69 | 62 | 10 | 100 | 44 | 114 | 25 | 86 | 39 | 491 | 335 | 1615 | 1377 | 319 | 1507 | 498 | 95 | 949 | 680 | 414 | 2635 |
| PBANKA_1243600 | 52 | 52 | 62 | 13 | 144 | 133 | 265 | 114 | 112 | 49 | 26 | 32 | 1388 | 456 | 481 | 2303 | 1398 | 186 | 711 | 587 | 444 | 3787 |
| PBANKA_1243700 | 0 | 10 | 53 | 11 | 103 | 59 | 97 | 29 | 26 | 12 | 8 | 0 | 274 | 62 | 242 | 665 | 158 | 22 | 119 | 36 | 111 | 580 |
| PBANKA_1243800 | 19 | 17 | 136 | 40 | 187 | 105 | 226 | 104 | 84 | 40 | 16 | 13 | 537 | 182 | 306 | 1236 | 200 | 36 | 256 | 109 | 271 | 1763 |
| PBANKA_1243900 | 76 | 154 | 340 | 100 | 837 | 268 | 1963 | 660 | 1578 | 691 | 176 | 93 | 2466 | 1270 | 754 | 4338 | 27573 | 3572 | 5144 | 6708 | 125 | 1307 |
| PBANKA_1244000 | 0 | 0 | 87 | 22 | 70 | 48 | 88 | 59 | 34 | 18 | 10 | 0 | 375 | 115 | 248 | 502 | 200 | 66 | 214 | 107 | 157 | 892 |
| PBANKA_1244100 | 0 | 0 | 35 | 8 | 85 | 31 | 39 | 23 | 13 | 9 | 11 | 0 | 1007 | 466 | 199 | 859 | 195 | 14 | 473 | 227 | 66 | 427 |
| PBANKA_1244200 | 22 | 28 | 201 | 43 | 490 | 139 | 792 | 296 | 392 | 195 | 77 | 26 | 577 | 785 | 725 | 4024 | 7963 | 889 | 1176 | 4534 | 142 | 1040 |
| PBANKA_1244300 | 217 | 228 | 78 | 17 | 162 | 106 | 240 | 103 | 127 | 100 | 101 | 79 | 475 | 210 | 340 | 767 | 998 | 270 | 1557 | 575 | 354 | 2708 |
| PBANKA_1244400 | 7 | 8 | 153 | 59 | 124 | 110 | 115 | 62 | 49 | 39 | 10 | 0 | 797 | 253 | 561 | 2065 | 456 | 82 | 771 | 501 | 21 | 219 |
| PBANKA_1244500 | 6 | 8 | 10 | 0 | 30 | 10 | 41 | 15 | 10 | 5 | 5 | 0 | 1120 | 358 | 233 | 1998 | 867 | 101 | 501 | 399 | 25 | 162 |
| PBANKA_1244600 | 0 | 5 | 108 | 29 | 483 | 272 | 648 | 245 | 177 | 81 | 24 | 6 | 636 | 108 | 559 | 2312 | 2005 | 273 | 764 | 378 | 44 | 357 |
| PBANKA_1244700 | 0 | 0 | 31 | 5 | 118 | 46 | 242 | 64 | 248 | 110 | 23 | 6 | 556 | 252 | 206 | 883 | 4637 | 798 | 1304 | 1774 | 59 | 403 |
| PBANKA_1244800 | 0 | 0 | 85 | 23 | 217 | 124 | 298 | 76 | 131 | 47 | 15 | 0 | 401 | 191 | 343 | 1494 | 444 | 99 | 332 | 389 | 34 | 245 |
| PBANKA_1244900 | 0 | 0 | 6 | 0 | 45 | 9 | 27 | 11 | 23 | 7 | 10 | 0 | 127 | 37 | 28 | 178 | 27 | 18 | 1104 | 892 | 23 | 138 |
| PBANKA_1245000 | 21 | 23 | 75 | 6 | 129 | 55 | 110 | 44 | 30 | 15 | 68 | 39 | 696 | 587 | 230 | 1291 | 380 | 65 | 182 | 211 | 37 | 207 |
| PBANKA_1245100 | 5 | 0 | 153 | 38 | 259 | 83 | 183 | 68 | 74 | 44 | 118 | 85 | 1342 | 1427 | 570 | 3324 | 852 | 123 | 1574 | 2263 | 86 | 707 |
| PBANKA_1245150 | 0 | 0 | 9 | 0 | 44 | 10 | 43 | 13 | 14 | 0 | 6 | 0 | 120 | 40 | 77 | 218 | 81 | 25 | 850 | 283 | 20 | 136 |
| PBANKA_1245200 | 0 | 0 | 127 | 34 | 171 | 144 | 234 | 97 | 44 | 43 | 23 | 13 | 441 | 236 | 198 | 370 | 193 | 111 | 8200 | 2329 | 214 | 1424 |

| | | | | | | | | | | | | | | | | | | | | | | |
|---|---|---|---|---|---|---|---|---|---|---|---|---|---|---|---|---|---|---|---|---|---|---|
| PBANKA_1245300 | 53 | 100 | 49 | 11 | 177 | 105 | 420 | 140 | 564 | 311 | 275 | 229 | 1073 | 981 | 482 | 2002 | 7175 | 1498 | 1103 | 2784 | 530 | 3649 |
| PBANKA_1245400 | 33 | 72 | 99 | 20 | 244 | 80 | 285 | 104 | 116 | 57 | 38 | 17 | 848 | 393 | 527 | 2472 | 2297 | 379 | 895 | 856 | 164 | 1122 |
| PBANKA_1245500 | 0 | 0 | 6 | 0 | 44 | 26 | 260 | 81 | 475 | 187 | 87 | 33 | 315 | 455 | 57 | 195 | 6354 | 1129 | 817 | 3356 | 154 | 1080 |
| PBANKA_1245600 | 0 | 0 | 20 | 0 | 52 | 20 | 57 | 17 | 18 | 6 | 0 | 6 | 498 | 161 | 167 | 610 | 208 | 40 | 194 | 86 | 0 | 39 |
| PBANKA_1245700 | 17 | 13 | 41 | 8 | 59 | 27 | 87 | 51 | 42 | 27 | 17 | 10 | 695 | 342 | 145 | 824 | 342 | 37 | 373 | 322 | 26 | 150 |
| PBANKA_1245800 | 373 | 482 | 191 | 28 | 674 | 193 | 709 | 206 | 481 | 217 | 1585 | 679 | 2963 | 2449 | 1463 | 8352 | 1573 | 337 | 1114 | 1151 | 103 | 527 |
| PBANKA_1245821 | 11 | 6 | 101 | 11 | 382 | 39 | 411 | 38 | 281 | 39 | 396 | 42 | 19644 | 1328 | 3040 | 6832 | 5029 | 1087 | 9101 | 3009 | 30 | 215 |
| PBANKA_1245841 | 0 | 0 | 38 | 21 | 61 | 39 | 51 | 20 | 21 | 8 | 0 | 8 | 161 | 20 | 169 | 108 | 74 | 20 | 1289 | 100 | 0 | 0 |
| PBANKA_1245861 | 1092 | 1236 | 4698 | 1685 | 12226 | 3875 | 15113 | 3785 | 8810 | 3239 | 14448 | 3655 | 43411 | 6046 | 18860 | 18547 | 31930 | 3532 | 72573 | 30359 | 67 | 1119 |
| PBANKA_1246600 | 0 | 0 | 102 | 37 | 260 | 145 | 192 | 79 | 85 | 55 | 43 | 7 | 3125 | 16 | 101 | 66 | 11 | 19 | 226 | 49 | 17 | 78 |
| PBANKA_1300096 | 0 | 0 | 12 | 0 | 58 | 40 | 42 | 7 | 18 | 5 | 0 | 0 | 594 | 28 | 55 | 26 | 17 | 17 | 94 | 48 | 14 | 64 |
| PBANKA_1300600 | 0 | 9 | 47 | 9 | 158 | 81 | 202 | 92 | 99 | 61 | 50 | 35 | 910 | 688 | 283 | 1291 | 1728 | 304 | 595 | 546 | 64 | 497 |
| PBANKA_1300700 | 89 | 96 | 28 | 0 | 290 | 151 | 285 | 94 | 92 | 43 | 22 | 10 | 839 | 833 | 88 | 354 | 2603 | 940 | 20342 | 31378 | 2436 | 30917 |
| PBANKA_1300800 | 207 | 336 | 38 | 13 | 565 | 334 | 728 | 288 | 376 | 210 | 54 | 39 | 184 | 194 | 457 | 1015 | 3063 | 600 | 562 | 2102 | 172 | 1467 |
| PBANKA_1300900 | 0 | 0 | 54 | 18 | 67 | 43 | 87 | 26 | 13 | 13 | 0 | 5 | 123 | 23 | 160 | 413 | 127 | 38 | 795 | 356 | 16 | 106 |
| PBANKA_1301000 | 26 | 58 | 86 | 20 | 261 | 131 | 315 | 132 | 120 | 47 | 28 | 12 | 1911 | 1281 | 826 | 4358 | 3187 | 363 | 636 | 817 | 20 | 100 |
| PBANKA_1301100 | 26 | 62 | 103 | 11 | 222 | 106 | 198 | 74 | 71 | 35 | 14 | 0 | 589 | 264 | 468 | 2707 | 1107 | 132 | 231 | 361 | 0 | 47 |
| PBANKA_1301200 | 16 | 16 | 28 | 7 | 50 | 29 | 64 | 14 | 20 | 0 | 10 | 0 | 67 | 18 | 33 | 212 | 116 | 14 | 15 | 17 | 0 | 0 |
| PBANKA_1301300 | 0 | 0 | 58 | 25 | 191 | 90 | 190 | 71 | 63 | 41 | 0 | 11 | 661 | 246 | 474 | 2352 | 2098 | 299 | 4652 | 4141 | 179 | 1654 |
| PBANKA_1301400 | 0 | 0 | 221 | 66 | 343 | 244 | 362 | 228 | 82 | 85 | 14 | 13 | 1426 | 324 | 1412 | 4298 | 1609 | 290 | 1476 | 563 | 30 | 272 |
| PBANKA_1301500 | 0 | 0 | 48 | 16 | 229 | 96 | 295 | 114 | 84 | 32 | 21 | 11 | 265 | 121 | 576 | 2226 | 526 | 172 | 614 | 428 | 0 | 29 |
| PBANKA_1301600 | 0 | 6 | 38 | 14 | 79 | 63 | 122 | 59 | 228 | 110 | 312 | 244 | 1627 | 897 | 470 | 1808 | 2013 | 489 | 2211 | 684 | 28 | 144 |
| PBANKA_1301700 | 0 | 9 | 131 | 37 | 257 | 108 | 296 | 101 | 206 | 104 | 173 | 138 | 2771 | 2076 | 1127 | 6194 | 4519 | 773 | 795 | 1137 | 45 | 328 |
| PBANKA_1301900 | 121 | 167 | 55 | 28 | 976 | 401 | 3142 | 1322 | 3289 | 1911 | 450 | 223 | 499 | 153 | 209 | 1053 | 4232 | 776 | 915 | 818 | 94 | 579 |
| PBANKA_1302000 | 32 | 29 | 47 | 11 | 426 | 175 | 1594 | 549 | 1525 | 573 | 200 | 76 | 194 | 113 | 205 | 399 | 411 | 126 | 1193 | 2503 | 17 | 93 |
| PBANKA_1302100 | 0 | 0 | 40 | 5 | 64 | 34 | 61 | 33 | 14 | 13 | 0 | 0 | 97 | 17 | 109 | 513 | 155 | 16 | 63 | 34 | 0 | 0 |
| PBANKA_1302200 | 0 | 0 | 161 | 26 | 331 | 271 | 626 | 348 | 273 | 189 | 49 | 23 | 2147 | 617 | 1051 | 4128 | 4732 | 670 | 1037 | 992 | 28 | 260 |
| PBANKA_1302300 | 0 | 0 | 52 | 10 | 939 | 408 | 1692 | 563 | 1320 | 687 | 216 | 123 | 862 | 263 | 441 | 1862 | 4051 | 1024 | 1372 | 1405 | 15 | 94 |
| PBANKA_1302400 | 225 | 329 | 108 | 30 | 526 | 284 | 505 | 238 | 171 | 112 | 572 | 466 | 1881 | 1717 | 910 | 4789 | 1584 | 215 | 695 | 1151 | 71 | 587 |
| PBANKA_1302500 | 8 | 16 | 54 | 18 | 120 | 79 | 170 | 71 | 65 | 37 | 208 | 152 | 707 | 310 | 231 | 965 | 508 | 82 | 372 | 228 | 18 | 121 |
| PBANKA_1302600 | 0 | 11 | 136 | 52 | 342 | 254 | 515 | 245 | 145 | 89 | 21 | 24 | 458 | 103 | 415 | 1101 | 1006 | 169 | 1091 | 432 | 29 | 147 |
| PBANKA_1302700 | 0 | 0 | 33 | 7 | 134 | 65 | 218 | 101 | 111 | 81 | 7 | 0 | 1498 | 84 | 696 | 2300 | 953 | 253 | 4114 | 1042 | 57 | 486 |
| PBANKA_1302800 | 0 | 10 | 1161 | 388 | 840 | 632 | 1025 | 571 | 244 | 179 | 21 | 13 | 1139 | 132 | 2379 | 6250 | 2440 | 464 | 2475 | 1725 | 164 | 1686 |
| PBANKA_1302900 | 0 | 0 | 78 | 14 | 171 | 105 | 409 | 189 | 121 | 60 | 13 | 0 | 460 | 92 | 557 | 1737 | 2283 | 393 | 283 | 172 | 17 | 210 |
| PBANKA_1303000 | 0 | 0 | 152 | 31 | 324 | 174 | 313 | 72 | 106 | 31 | 7 | 6 | 1387 | 757 | 797 | 3283 | 470 | 77 | 160 | 253 | 22 | 171 |
| PBANKA_1303100 | 0 | 0 | 5 | 0 | 51 | 19 | 49 | 21 | 35 | 18 | 0 | 0 | 183 | 115 | 138 | 267 | 127 | 35 | 506 | 647 | 9 | 84 |
| PBANKA_1303200 | 58 | 80 | 473 | 107 | 862 | 383 | 1258 | 474 | 534 | 278 | 1021 | 662 | 4574 | 6660 | 2320 | 9869 | 12099 | 1535 | 1264 | 2268 | 243 | 1865 |
| PBANKA_1303300 | 11 | 34 | 284 | 67 | 764 | 539 | 1404 | 602 | 942 | 571 | 95 | 45 | 1282 | 329 | 888 | 3546 | 8352 | 1556 | 3180 | 2076 | 145 | 1295 |
| PBANKA_1303400 | 24 | 28 | 62 | 20 | 183 | 129 | 302 | 130 | 383 | 258 | 74 | 24 | 414 | 179 | 190 | 981 | 4755 | 831 | 1232 | 976 | 11 | 111 |
| PBANKA_1303500 | 39 | 81 | 135 | 51 | 459 | 329 | 2023 | 1189 | 1024 | 826 | 110 | 51 | 572 | 155 | 531 | 1415 | 1920 | 552 | 2823 | 1093 | 71 | 836 |
| PBANKA_1303600 | 11 | 28 | 120 | 26 | 421 | 201 | 453 | 224 | 138 | 73 | 30 | 13 | 216 | 33 | 144 | 702 | 361 | 44 | 315 | 144 | 24 | 109 |
| PBANKA_1303700 | 97 | 124 | 114 | 35 | 276 | 153 | 543 | 221 | 185 | 139 | 65 | 64 | 1467 | 462 | 412 | 1474 | 1769 | 343 | 1128 | 542 | 173 | 1160 |
| PBANKA_1303800 | 12 | 0 | 927 | 336 | 1167 | 778 | 1096 | 524 | 302 | 187 | 59 | 41 | 4392 | 802 | 3169 | 9034 | 1688 | 285 | 2593 | 1161 | 172 | 1404 |
| PBANKA_1303900 | 13 | 10 | 95 | 30 | 316 | 118 | 592 | 226 | 374 | 192 | 47 | 47 | 907 | 1171 | 95 | 362 | 7842 | 1109 | 9053 | 26968 | 1003 | 6657 |
| PBANKA_1304000 | 85 | 142 | 92 | 27 | 228 | 114 | 425 | 165 | 296 | 165 | 43 | 30 | 658 | 222 | 25 | 114 | 7630 | 1179 | 2093 | 1442 | 1045 | 8582 |
| PBANKA_1304200 | 253 | 343 | 185 | 46 | 171 | 117 | 168 | 71 | 84 | 41 | 49 | 24 | 436 | 219 | 402 | 1711 | 362 | 106 | 448 | 523 | 25 | 264 |
| PBANKA_1304100 | 332 | 435 | 79 | 18 | 115 | 44 | 118 | 51 | 69 | 44 | 8 | 0 | 207 | 91 | 111 | 613 | 246 | 66 | 343 | 199 | 42 | 275 |
| PBANKA_1304300 | 0 | 0 | 34 | 16 | 165 | 86 | 407 | 162 | 221 | 125 | 42 | 6 | 547 | 301 | 43 | 128 | 5742 | 712 | 1285 | 3073 | 84 | 700 |
| PBANKA_1304400 | 0 | 0 | 49 | 5 | 176 | 78 | 144 | 38 | 77 | 32 | 14 | 8 | 271 | 76 | 126 | 447 | 320 | 80 | 2383 | 1178 | 124 | 958 |
| PBANKA_1304500 | 13 | 41 | 12 | 0 | 51 | 15 | 64 | 19 | 26 | 21 | 24 | 0 | 370 | 145 | 16 | 56 | 255 | 91 | 2444 | 1767 | 110 | 738 |
| PBANKA_1304600 | 20 | 31 | 97 | 20 | 215 | 106 | 317 | 98 | 102 | 53 | 22 | 0 | 385 | 194 | 254 | 1096 | 751 | 91 | 302 | 236 | 10 | 115 |
| PBANKA_1304700 | 13 | 0 | 60 | 14 | 164 | 79 | 246 | 102 | 130 | 94 | 19 | 6 | 413 | 88 | 183 | 746 | 935 | 199 | 180 | 122 | 0 | 32 |
| PBANKA_1304800 | 0 | 0 | 41 | 13 | 107 | 60 | 214 | 112 | 115 | 85 | 13 | 10 | 189 | 32 | 51 | 175 | 630 | 211 | 108 | 61 | 0 | 8 |
| PBANKA_1304900 | 10 | 0 | 103 | 27 | 132 | 45 | 348 | 145 | 352 | 171 | 77 | 25 | 874 | 508 | 208 | 1215 | 462 | 165 | 200 | 145 | 7 | 52 |
| PBANKA_1305000 | 34 | 55 | 690 | 233 | 710 | 363 | 800 | 400 | 446 | 329 | 3247 | 2595 | 34050 | 17487 | 4712 | 37553 | 24767 | 2100 | 9550 | 8241 | 755 | 7084 |
| PBANKA_1305100 | 0 | 0 | 1161 | 367 | 812 | 506 | 914 | 469 | 242 | 198 | 1651 | 1570 | 11526 | 4852 | 2565 | 9428 | 2780 | 409 | 2316 | 1366 | 212 | 1853 |
| PBANKA_1305200 | 11 | 16 | 16 | 0 | 221 | 105 | 914 | 376 | 1802 | 918 | 300 | 110 | 3027 | 5080 | 525 | 665 | 17081 | 2869 | 42016 | 92020 | 768 | 5731 |
| PBANKA_1305300 | 0 | 0 | 151 | 38 | 307 | 180 | 325 | 203 | 121 | 59 | 11 | 18 | 1312 | 333 | 1166 | 4082 | 3040 | 453 | 1944 | 1159 | 13 | 127 |
| PBANKA_1305400 | 0 | 7 | 42 | 8 | 206 | 82 | 363 | 129 | 134 | 76 | 14 | 6 | 290 | 58 | 125 | 380 | 554 | 105 | 829 | 455 | 20 | 182 |
| PBANKA_1305500 | 328 | 495 | 17 | 0 | 50 | 29 | 177 | 48 | 352 | 211 | 25 | 25 | 585 | 195 | 57 | 205 | 2968 | 670 | 4673 | 3344 | 667 | 6318 |
| PBANKA_1305600 | 55 | 78 | 170 | 35 | 258 | 146 | 296 | 104 | 111 | 57 | 21 | 11 | 168 | 23 | 236 | 780 | 315 | 74 | 386 | 220 | 36 | 219 |
| PBANKA_1305700 | 47 | 47 | 48 | 14 | 66 | 38 | 105 | 34 | 33 | 15 | 8 | 0 | 608 | 213 | 203 | 676 | 485 | 115 | 4163 | 1899 | 475 | 3245 |
| PBANKA_1305800 | 11 | 8 | 6 | 0 | 24 | 15 | 38 | 9 | 8 | 7 | 0 | 0 | 91 | 14 | 35 | 168 | 107 | 17 | 147 | 53 | 11 | 102 |
| PBANKA_1305900 | 0 | 0 | 45 | 9 | 122 | 80 | 137 | 69 | 37 | 21 | 0 | 0 | 436 | 38 | 227 | 852 | 451 | 68 | 1562 | 766 | 19 | 196 |
| PBANKA_1306000 | 0 | 7 | 107 | 26 | 148 | 99 | 228 | 102 | 57 | 20 | 15 | 0 | 374 | 71 | 225 | 530 | 343 | 121 | 1738 | 563 | 27 | 195 |
| PBANKA_1306100 | 0 | 0 | 61 | 14 | 132 | 40 | 65 | 25 | 42 | 13 | 9 | 0 | 306 | 150 | 278 | 1498 | 175 | 33 | 145 | 416 | 106 | 760 |
| PBANKA_1306200 | 0 | 0 | 353 | 109 | 274 | 174 | 244 | 95 | 65 | 32 | 38 | 24 | 2622 | 1041 | 1195 | 5719 | 882 | 104 | 1341 | 1121 | 29 | 216 |
| PBANKA_1306300 | 0 | 0 | 163 | 29 | 549 | 243 | 803 | 282 | 572 | 265 | 105 | 31 | 1299 | 2147 | 2058 | 10649 | 8753 | 1348 | 583 | 3194 | 411 | 3475 |
| PBANKA_1306400 | 0 | 0 | 51 | 29 | 158 | 59 | 123 | 44 | 44 | 24 | 7 | 0 | 404 | 78 | 368 | 1641 | 419 | 48 | 466 | 129 | 8 | 45 |
| PBANKA_1306500 | 74 | 80 | 28 | 7 | 37 | 16 | 19 | 18 | 33 | 20 | 17 | 5 | 7 | 0 | 25 | 182 | 47 | 17 | 65 | 62 | 0 | 20 |
| PBANKA_1306600 | 81 | 162 | 63 | 10 | 818 | 292 | 2271 | 899 | 1486 | 675 | 259 | 148 | 513 | 627 | 276 | 987 | 11454 | 1337 | 443 | 2780 | 196 | 1364 |
| PBANKA_1306700 | 0 | 0 | 5 | 5 | 23 | 15 | 20 | 11 | 20 | 9 | 0 | 0 | 35 | 10 | 45 | 208 | 58 | 15 | 90 | 40 | 5 | 20 |
| PBANKA_1306800 | 94 | 174 | 521 | 109 | 1267 | 511 | 1503 | 491 | 607 | 240 | 96 | 60 | 1670 | 2051 | 3247 | 15613 | 9355 | 948 | 735 | 2486 | 141 | 820 |
| PBANKA_1306900 | 11 | 26 | 64 | 16 | 537 | 354 | 1205 | 502 | 518 | 261 | 71 | 37 | 334 | 276 | 236 | 705 | 3448 | 576 | 1983 | 3612 | 404 | 3517 |
| PBANKA_1307000 | 153 | 221 | 146 | 36 | 458 | 168 | 653 | 252 | 490 | 245 | 73 | 33 | 1884 | 1252 | 594 | 3693 | 14354 | 1785 | 5283 | 11188 | 212 | 1934 |
| PBANKA_1307100 | 12 | 17 | 82 | 20 | 334 | 169 | 346 | 113 | 114 | 37 | 18 | 9 | 336 | 78 | 127 | 588 | 433 | 71 | 1051 | 591 | 54 | 512 |
| PBANKA_1307200 | 0 | 0 | 57 | 19 | 257 | 71 | 278 | 107 | 74 | 38 | 41 | 24 | 267 | 149 | 190 | 413 | 256 | 41 | 125 | 49 | 15 | 58 |
| PBANKA_1307300 | 0 | 0 | 198 | 42 | 843 | 374 | 871 | 267 | 234 | 99 | 46 | 8 | 623 | 199 | 1660 | 6170 | 3389 | 325 | 168 | 167 | 0 | 22 |
| PBANKA_1307400 | 0 | 0 | 71 | 16 | 190 | 103 | 170 | 57 | 74 | 29 | 7 | 7 | 797 | 175 | 434 | 2025 | 448 | 92 | 445 | 582 | 0 | 35 |
| PBANKA_1307500 | 6 | 14 | 106 | 25 | 401 | 144 | 508 | 147 | 198 | 74 | 47 | 0 | 322 | 154 | 741 | 3004 | 1639 | 212 | 934 | 1649 | 48 | 431 |
| PBANKA_1307600 | 211 | 233 | 146 | 26 | 283 | 129 | 424 | 179 | 245 | 85 | 20 | 12 | 678 | 111 | 538 | 1781 | 1465 | 249 | 911 | 529 | 67 | 630 |
| PBANKA_1307700 | 76 | 82 | 48 | 11 | 196 | 77 | 305 | 121 | 114 | 58 | 40 | 13 | 1434 | 935 | 534 | 3518 | 3949 | 448 | 2154 | 1848 | 68 | 434 |
| PBANKA_1307800 | 104 | 126 | 254 | 75 | 552 | 408 | 878 | 343 | 284 | 173 | 79 | 56 | 1809 | 1259 | 1086 | 4825 | 3043 | 412 | 665 | 685 | 119 | 1076 |
| PBANKA_1307900 | 0 | 0 | 6 | 0 | 6 | 0 | 11 | 0 | 0 | 0 | 0 | 0 | 56 | 34 | 46 | 89 | 61 | 47 | 265 | 159 | 6 | 18 |
| PBANKA_1308000 | 12 | 31 | 48 | 17 | 117 | 64 | 173 | 63 | 276 | 110 | 52 | 23 | 538 | 1269 | 398 | 1386 | 6478 | 1135 | 1348 | 18978 | 165 | 1015 |
| PBANKA_1308200 | 0 | 13 | 88 | 34 | 222 | 154 | 371 | 188 | 96 | 63 | 13 | 14 | 1362 | 223 | 628 | 1400 | 2692 | 673 | 1413 | 596 | 69 | 674 |
| PBANKA_1308300 | 18 | 10 | 26 | 0 | 64 | 14 | 69 | 17 | 20 | 15 | 12 | 8 | 301 | 180 | 115 | 472 | 106 | 32 | 281 | 160 | 44 | 283 |
| PBANKA_1308400 | 339 | 395 | 107 | 27 | 418 | 119 | 446 | 149 | 295 | 137 | 39 | 17 | 411 | 277 | 502 | 2075 | 3616 | 550 | 829 | 1834 | 52 | 434 |
| PBANKA_1308500 | 75 | 66 | 29 | 13 | 148 | 72 | 106 | 68 | 50 | 24 | 0 | 6 | 161 | 45 | 79 | 184 | 161 | 58 | 239 | 62 | 15 | 115 |
| PBANKA_1308600 | 1003 | 1026 | 1713 | 605 | 2340 | 1833 | 2045 | 1043 | 599 | 360 | 234 | 213 | 25259 | 8188 | 6489 | 21937 | 4418 | 845 | 45146 | 10748 | 600 | 5524 |

| | | | | | | | | | | | | | | | | | | | | | | |
|---|---|---|---|---|---|---|---|---|---|---|---|---|---|---|---|---|---|---|---|---|---|---|
| PBANKA_1308700 | 97 | 142 | 74 | 22 | 103 | 72 | 98 | 34 | 51 | 25 | 5 | 0 | 120 | 21 | 126 | 262 | 134 | 41 | 601 | 185 | 13 | 117 |
| PBANKA_1308800 | 55 | 87 | 30 | 0 | 69 | 42 | 111 | 38 | 41 | 29 | 11 | 0 | 250 | 70 | 123 | 420 | 596 | 84 | 1066 | 665 | 30 | 248 |
| PBANKA_1308900 | 93 | 138 | 145 | 40 | 422 | 184 | 377 | 82 | 136 | 51 | 125 | 105 | 2228 | 1027 | 929 | 4380 | 712 | 107 | 941 | 461 | 96 | 571 |
| PBANKA_1309000 | 24 | 36 | 78 | 31 | 225 | 79 | 195 | 86 | 132 | 63 | 532 | 301 | 2022 | 1966 | 391 | 2072 | 740 | 145 | 428 | 580 | 105 | 687 |
| PBANKA_1309100 | 0 | 0 | 42 | 18 | 145 | 59 | 137 | 71 | 53 | 25 | 10 | 5 | 479 | 150 | 252 | 884 | 322 | 75 | 180 | 101 | 7 | 60 |
| PBANKA_1309200 | 86 | 113 | 199 | 34 | 291 | 116 | 278 | 113 | 446 | 250 | 893 | 579 | 3271 | 3549 | 365 | 2353 | 3913 | 664 | 896 | 1316 | 298 | 1980 |
| PBANKA_1309300 | 10 | 17 | 10 | 0 | 40 | 11 | 48 | 18 | 56 | 23 | 27 | 17 | 336 | 294 | 91 | 305 | 251 | 108 | 303 | 171 | 52 | 407 |
| PBANKA_1309400 | 300 | 303 | 81 | 15 | 346 | 150 | 797 | 347 | 634 | 335 | 403 | 227 | 2708 | 1124 | 365 | 1662 | 2426 | 678 | 3625 | 1368 | 510 | 2634 |
| PBANKA_1309500 | 153 | 252 | 465 | 93 | 526 | 251 | 607 | 188 | 194 | 93 | 1180 | 801 | 4638 | 2704 | 1522 | 8083 | 2355 | 311 | 638 | 745 | 140 | 842 |
| PBANKA_1309600 | 7 | 6 | 8 | 0 | 39 | 8 | 25 | 8 | 27 | 11 | 0 | 5 | 159 | 110 | 32 | 89 | 90 | 16 | 348 | 398 | 94 | 831 |
| PBANKA_1309700 | 240 | 315 | 201 | 61 | 212 | 124 | 301 | 123 | 159 | 73 | 850 | 752 | 4749 | 1953 | 777 | 2744 | 1522 | 333 | 2673 | 1122 | 2764 | 23533 |
| PBANKA_1309750 | 0 | 0 | 27 | 0 | 20 | 7 | 10 | 0 | 23 | 11 | 11 | 8 | 6 | 26 | 202 | 38 | 49 | 28 | 42 | 42 | 0 | 0 |
| PBANKA_1309800 | 6 | 0 | 14 | 0 | 26 | 6 | 30 | 10 | 63 | 41 | 26 | 6 | 130 | 67 | 46 | 181 | 259 | 78 | 889 | 1226 | 165 | 1068 |
| PBANKA_1309900 | 48 | 49 | 620 | 202 | 894 | 539 | 786 | 308 | 201 | 130 | 69 | 30 | 6644 | 1228 | 3580 | 14297 | 2580 | 402 | 2369 | 1256 | 128 | 992 |
| PBANKA_1310000 | 222 | 239 | 25 | 0 | 96 | 28 | 84 | 28 | 186 | 74 | 75 | 50 | 693 | 486 | 263 | 2092 | 646 | 171 | 2447 | 2630 | 452 | 2810 |
| PBANKA_1310100 | 0 | 0 | 45 | 14 | 281 | 204 | 374 | 140 | 108 | 48 | 13 | 13 | 68 | 22 | 35 | 217 | 346 | 46 | 109 | 78 | 30 | 260 |
| PBANKA_1310200 | 426 | 588 | 362 | 85 | 1046 | 324 | 1439 | 476 | 897 | 336 | 792 | 558 | 2611 | 2204 | 1463 | 5915 | 15470 | 2277 | 2903 | 5004 | 431 | 2832 |
| PBANKA_1310400 | 0 | 0 | 77 | 19 | 148 | 83 | 474 | 234 | 612 | 409 | 62 | 40 | 775 | 144 | 224 | 638 | 3093 | 1288 | 1511 | 586 | 36 | 369 |
| PBANKA_1310500 | 136 | 163 | 35 | 17 | 107 | 44 | 381 | 196 | 352 | 150 | 20 | 9 | 964 | 260 | 121 | 491 | 2878 | 661 | 8586 | 4165 | 210 | 1464 |
| PBANKA_1310600 | 11 | 12 | 66 | 12 | 202 | 104 | 197 | 111 | 92 | 49 | 17 | 0 | 185 | 32 | 175 | 569 | 401 | 108 | 1180 | 475 | 20 | 185 |
| PBANKA_1310700 | 1844 | 2600 | 22 | 8 | 189 | 76 | 267 | 139 | 108 | 52 | 5 | 9 | 623 | 211 | 90 | 340 | 640 | 157 | 6682 | 4494 | 951 | 8225 |
| PBANKA_1310800 | 183 | 251 | 2374 | 801 | 1646 | 1450 | 1977 | 1046 | 666 | 431 | 2816 | 2500 | 17507 | 5862 | 4299 | 14890 | 3957 | 642 | 3020 | 1337 | 622 | 5315 |
| PBANKA_1310900 | 53 | 67 | 64 | 9 | 87 | 33 | 59 | 30 | 33 | 29 | 46 | 78 | 794 | 304 | 165 | 593 | 355 | 82 | 484 | 274 | 47 | 294 |
| PBANKA_1311000 | 70 | 102 | 96 | 16 | 186 | 87 | 202 | 84 | 84 | 47 | 120 | 78 | 1779 | 1018 | 518 | 2374 | 1193 | 215 | 1216 | 1026 | 325 | 2004 |
| PBANKA_1311100 | 0 | 0 | 32 | 12 | 55 | 32 | 68 | 22 | 14 | 9 | 0 | 0 | 97 | 36 | 192 | 622 | 197 | 37 | 229 | 51 | 13 | 128 |
| PBANKA_1311200 | 0 | 0 | 78 | 26 | 132 | 80 | 113 | 60 | 35 | 16 | 8 | 0 | 517 | 230 | 146 | 449 | 82 | 25 | 583 | 258 | 143 | 627 |
| PBANKA_1311500 | 0 | 0 | 28 | 5 | 234 | 104 | 358 | 147 | 125 | 83 | 10 | 5 | 426 | 87 | 256 | 1290 | 1672 | 210 | 640 | 525 | 119 | 789 |
| PBANKA_1311600 | 6 | 6 | 15 | 5 | 217 | 82 | 343 | 128 | 247 | 116 | 503 | 299 | 2050 | 2015 | 374 | 2794 | 3112 | 567 | 861 | 4395 | 159 | 842 |
| PBANKA_1311700 | 6 | 0 | 9 | 0 | 58 | 30 | 91 | 35 | 215 | 116 | 19 | 8 | 616 | 244 | 408 | 2010 | 2347 | 389 | 662 | 461 | 130 | 1015 |
| PBANKA_1311800 | 47 | 79 | 190 | 47 | 247 | 116 | 244 | 77 | 125 | 91 | 1355 | 789 | 2017 | 2188 | 421 | 2533 | 1193 | 131 | 700 | 982 | 654 | 4400 |
| PBANKA_1312100 | 16 | 18 | 47 | 10 | 256 | 73 | 637 | 208 | 366 | 108 | 133 | 39 | 1344 | 720 | 526 | 2806 | 4872 | 619 | 2650 | 2369 | 156 | 1201 |
| PBANKA_1312200 | 0 | 8 | 192 | 41 | 393 | 243 | 352 | 155 | 104 | 40 | 16 | 14 | 2755 | 1042 | 1242 | 5802 | 1024 | 151 | 3177 | 3316 | 79 | 694 |
| PBANKA_1312300 | 74 | 111 | 16 | 7 | 79 | 40 | 112 | 46 | 85 | 36 | 34 | 16 | 427 | 219 | 120 | 434 | 1111 | 203 | 744 | 602 | 1136 | 7329 |
| PBANKA_1312400 | 130 | 169 | 56 | 16 | 135 | 62 | 189 | 87 | 84 | 59 | 65 | 40 | 676 | 369 | 159 | 612 | 753 | 151 | 342 | 207 | 428 | 2258 |
| PBANKA_1312500 | 41 | 41 | 101 | 17 | 177 | 66 | 203 | 71 | 66 | 25 | 22 | 0 | 393 | 145 | 320 | 982 | 482 | 134 | 1272 | 499 | 228 | 1241 |
| PBANKA_1312600 | 0 | 0 | 41 | 8 | 65 | 27 | 54 | 19 | 12 | 11 | 0 | 0 | 46 | 11 | 82 | 211 | 97 | 27 | 297 | 97 | 6 | 58 |
| PBANKA_1312800 | 102 | 104 | 68 | 9 | 495 | 267 | 594 | 248 | 670 | 419 | 182 | 114 | 696 | 357 | 259 | 817 | 3347 | 639 | 4674 | 4129 | 3619 | 27268 |
| PBANKA_1313000 | 29 | 34 | 74 | 22 | 221 | 110 | 217 | 77 | 150 | 99 | 1484 | 1073 | 1008 | 615 | 216 | 1069 | 476 | 154 | 1118 | 642 | 248 | 1946 |
| PBANKA_1313100 | 7 | 9 | 62 | 20 | 376 | 173 | 558 | 188 | 231 | 107 | 33 | 13 | 964 | 636 | 117 | 371 | 5860 | 763 | 6482 | 14696 | 3939 | 24702 |
| PBANKA_1313200 | 10 | 17 | 42 | 0 | 72 | 45 | 94 | 35 | 36 | 25 | 13 | 0 | 1330 | 749 | 205 | 1454 | 741 | 141 | 29794 | 31604 | 882 | 6643 |
| PBANKA_1313400 | 56 | 46 | 96 | 13 | 143 | 50 | 120 | 45 | 172 | 103 | 1032 | 642 | 2288 | 1879 | 237 | 1804 | 1217 | 188 | 1930 | 1467 | 480 | 2822 |
| PBANKA_1313500 | 17 | 33 | 22 | 0 | 38 | 51 | 91 | 37 | 79 | 73 | 201 | 132 | 212 | 74 | 113 | 296 | 403 | 79 | 416 | 169 | 93 | 691 |
| PBANKA_1313600 | 0 | 0 | 15 | 5 | 61 | 29 | 140 | 63 | 65 | 28 | 5 | 0 | 363 | 65 | 123 | 551 | 563 | 139 | 1629 | 594 | 49 | 295 |
| PBANKA_1313700 | 0 | 0 | 23 | 0 | 71 | 36 | 159 | 86 | 233 | 147 | 70 | 24 | 1764 | 587 | 415 | 1222 | 1536 | 524 | 1672 | 800 | 32 | 245 |
| PBANKA_1313800 | 79 | 99 | 86 | 20 | 179 | 73 | 268 | 97 | 101 | 45 | 74 | 53 | 1153 | 609 | 384 | 2050 | 1527 | 180 | 1449 | 1203 | 101 | 943 |
| PBANKA_1313900 | 0 | 0 | 10 | 0 | 21 | 6 | 30 | 11 | 19 | 7 | 0 | 0 | 220 | 80 | 84 | 439 | 283 | 45 | 234 | 217 | 6 | 36 |
| PBANKA_1314000 | 0 | 0 | 61 | 5 | 129 | 66 | 176 | 71 | 81 | 30 | 5 | 0 | 774 | 271 | 347 | 1582 | 966 | 158 | 940 | 620 | 14 | 113 |
| PBANKA_1314100 | 12 | 8 | 87 | 20 | 429 | 121 | 333 | 124 | 180 | 72 | 547 | 319 | 2662 | 2391 | 1272 | 6288 | 1803 | 270 | 1599 | 2517 | 61 | 404 |
| PBANKA_1314200 | 105 | 78 | 176 | 42 | 372 | 146 | 347 | 134 | 201 | 76 | 35 | 19 | 591 | 302 | 693 | 3367 | 2541 | 297 | 1129 | 1011 | 49 | 293 |
| PBANKA_1314300 | 205 | 326 | 100 | 23 | 251 | 129 | 373 | 153 | 130 | 89 | 53 | 37 | 674 | 389 | 235 | 908 | 786 | 131 | 3986 | 2743 | 204 | 1639 |
| PBANKA_1314400 | 227 | 277 | 123 | 38 | 349 | 221 | 539 | 230 | 330 | 194 | 722 | 542 | 3868 | 5624 | 1272 | 9646 | 8983 | 811 | 1444 | 5654 | 305 | 2273 |
| PBANKA_1314500 | 0 | 0 | 92 | 27 | 75 | 48 | 49 | 10 | 12 | 5 | 0 | 0 | 1359 | 283 | 766 | 2752 | 450 | 83 | 429 | 241 | 11 | 139 |
| PBANKA_1314600 | 5 | 0 | 59 | 7 | 49 | 46 | 40 | 6 | 14 | 6 | 0 | 0 | 472 | 154 | 210 | 902 | 123 | 23 | 115 | 84 | 6 | 60 |
| PBANKA_1314700 | 62 | 101 | 164 | 31 | 195 | 73 | 175 | 55 | 110 | 44 | 21 | 6 | 1464 | 968 | 512 | 2538 | 1433 | 215 | 915 | 1458 | 149 | 965 |
| PBANKA_1314800 | 69 | 84 | 3986 | 1394 | 3799 | 3116 | 3639 | 1792 | 1077 | 675 | 922 | 681 | 21781 | 11806 | 12317 | 46561 | 10528 | 1201 | 4278 | 3353 | 1986 | 21547 |
| PBANKA_1314900 | 392 | 468 | 75 | 18 | 364 | 157 | 321 | 101 | 805 | 421 | 4866 | 3262 | 2921 | 3113 | 842 | 4750 | 2362 | 630 | 10182 | 15447 | 113 | 1114 |
| PBANKA_1315000 | 39 | 11 | 29 | 15 | 180 | 135 | 199 | 92 | 73 | 73 | 13 | 7 | 216 | 63 | 73 | 200 | 842 | 186 | 741 | 431 | 11 | 87 |
| PBANKA_1315100 | 6 | 5 | 82 | 29 | 252 | 125 | 366 | 113 | 180 | 54 | 32 | 28 | 561 | 360 | 637 | 2669 | 1715 | 238 | 604 | 791 | 77 | 427 |
| PBANKA_1315200 | 0 | 5 | 22 | 0 | 54 | 19 | 86 | 26 | 22 | 17 | 48 | 20 | 306 | 97 | 125 | 467 | 382 | 82 | 185 | 97 | 0 | 29 |
| PBANKA_1315300 | 0 | 0 | 5 | 0 | 12 | 5 | 15 | 10 | 7 | 0 | 0 | 0 | 1889 | 1269 | 71 | 242 | 1199 | 561 | 51258 | 66722 | 1540 | 15102 |
| PBANKA_1315400 | 87 | 100 | 60 | 6 | 163 | 98 | 425 | 192 | 233 | 185 | 21 | 9 | 438 | 68 | 162 | 558 | 1274 | 369 | 840 | 413 | 83 | 429 |
| PBANKA_1315500 | 172 | 282 | 362 | 126 | 981 | 517 | 1298 | 451 | 611 | 356 | 149 | 102 | 5166 | 1900 | 1446 | 6396 | 11252 | 1461 | 2958 | 2485 | 228 | 2176 |
| PBANKA_1315600 | 249 | 310 | 133 | 32 | 274 | 159 | 261 | 147 | 181 | 81 | 1145 | 1024 | 2025 | 2514 | 412 | 2549 | 639 | 105 | 484 | 772 | 289 | 2261 |
| PBANKA_1315700 | 22 | 9 | 45 | 15 | 291 | 191 | 1247 | 552 | 5338 | 3328 | 1082 | 849 | 2780 | 2009 | 429 | 1658 | 43956 | 10842 | 2612 | 2676 | 42 | 392 |
| PBANKA_1315800 | 0 | 0 | 26 | 0 | 41 | 16 | 28 | 13 | 16 | 6 | 61 | 40 | 333 | 128 | 54 | 207 | 124 | 26 | 183 | 74 | 0 | 17 |
| PBANKA_1315900 | 22 | 22 | 82 | 11 | 103 | 68 | 119 | 59 | 86 | 22 | 72 | 43 | 582 | 274 | 280 | 1224 | 634 | 95 | 887 | 669 | 46 | 333 |
| PBANKA_1316000 | 21 | 25 | 37 | 7 | 119 | 31 | 174 | 67 | 77 | 33 | 25 | 20 | 579 | 200 | 219 | 746 | 1009 | 220 | 405 | 305 | 135 | 802 |
| PBANKA_1316100 | 82 | 149 | 140 | 23 | 189 | 118 | 365 | 134 | 187 | 110 | 389 | 279 | 2663 | 2306 | 536 | 3014 | 3010 | 448 | 9715 | 11795 | 417 | 3038 |
| PBANKA_1316200 | 0 | 0 | 12 | 0 | 63 | 35 | 150 | 70 | 109 | 89 | 14 | 0 | 194 | 44 | 102 | 426 | 423 | 89 | 369 | 158 | 18 | 193 |
| PBANKA_1316300 | 18 | 36 | 245 | 52 | 868 | 262 | 1473 | 541 | 1024 | 428 | 153 | 64 | 1890 | 1279 | 1849 | 8300 | 6105 | 939 | 2240 | 3850 | 73 | 578 |
| PBANKA_1316400 | 0 | 0 | 244 | 68 | 241 | 253 | 302 | 128 | 85 | 51 | 41 | 16 | 4216 | 1266 | 1131 | 5584 | 2013 | 281 | 1684 | 799 | 23 | 192 |
| PBANKA_1316500 | 0 | 6 | 7 | 0 | 22 | 8 | 31 | 17 | 25 | 9 | 0 | 0 | 246 | 42 | 41 | 132 | 100 | 52 | 2196 | 689 | 8 | 93 |
| PBANKA_1316600 | 11 | 35 | 33 | 10 | 107 | 55 | 94 | 46 | 61 | 22 | 13 | 0 | 299 | 48 | 155 | 601 | 454 | 111 | 928 | 343 | 14 | 91 |
| PBANKA_1316700 | 0 | 0 | 107 | 44 | 223 | 106 | 176 | 68 | 49 | 21 | 13 | 23 | 1674 | 964 | 371 | 1893 | 1046 | 129 | 85 | 46 | 0 | 26 |
| PBANKA_1316900 | 23 | 56 | 26 | 10 | 176 | 65 | 449 | 221 | 292 | 151 | 47 | 25 | 576 | 273 | 202 | 684 | 4233 | 707 | 901 | 1154 | 39 | 283 |
| PBANKA_1317000 | 0 | 0 | 7 | 0 | 17 | 10 | 31 | 13 | 15 | 7 | 5 | 0 | 30 | 0 | 16 | 54 | 52 | 11 | 108 | 36 | 0 | 7 |
| PBANKA_1317100 | 40 | 62 | 114 | 41 | 460 | 207 | 560 | 214 | 355 | 167 | 44 | 17 | 837 | 324 | 589 | 2666 | 2547 | 446 | 1409 | 1446 | 48 | 426 |
| PBANKA_1317200 | 25 | 59 | 14 | 12 | 72 | 72 | 59 | 33 | 36 | 10 | 5 | 0 | 1779 | 1348 | 106 | 322 | 1386 | 558 | 37377 | 52852 | 617 | 4671 |
| PBANKA_1317300 | 0 | 5 | 191 | 57 | 414 | 231 | 406 | 148 | 174 | 57 | 40 | 0 | 1970 | 1591 | 850 | 3603 | 617 | 105 | 1309 | 658 | 129 | 962 |
| PBANKA_1317400 | 13 | 18 | 2704 | 781 | 2350 | 1179 | 1797 | 640 | 389 | 228 | 475 | 286 | 38232 | 8871 | 7308 | 52871 | 6870 | 577 | 6012 | 4346 | 511 | 4891 |
| PBANKA_1317500 | 135 | 184 | 166 | 30 | 606 | 407 | 757 | 255 | 310 | 183 | 344 | 202 | 1929 | 1082 | 1215 | 4716 | 2884 | 420 | 2431 | 1838 | 349 | 2570 |
| PBANKA_1317600 | 181 | 198 | 117 | 31 | 290 | 176 | 334 | 104 | 290 | 137 | 2279 | 1219 | 971 | 815 | 274 | 1434 | 1181 | 198 | 792 | 869 | 859 | 6975 |
| PBANKA_1317700 | 0 | 6 | 93 | 27 | 264 | 127 | 318 | 139 | 129 | 74 | 150 | 68 | 876 | 546 | 600 | 2997 | 1700 | 233 | 231 | 366 | 19 | 238 |
| PBANKA_1317800 | 0 | 0 | 15 | 0 | 10 | 12 | 36 | 12 | 0 | 5 | 0 | 0 | 127 | 39 | 28 | 71 | 71 | 16 | 277 | 114 | 13 | 73 |
| PBANKA_1317900 | 38 | 89 | 261 | 50 | 384 | 160 | 382 | 121 | 151 | 55 | 1046 | 556 | 1972 | 2108 | 978 | 4603 | 1010 | 139 | 623 | 1026 | 351 | 2817 |
| PBANKA_1318100 | 61 | 88 | 486 | 135 | 1839 | 1026 | 2112 | 776 | 768 | 433 | 2580 | 1251 | 18495 | 10819 | 6997 | 28287 | 6520 | 1059 | 3036 | 2145 | 77 | 848 |
| PBANKA_1318200 | 638 | 679 | 41 | 6 | 58 | 50 | 68 | 38 | 28 | 37 | 78 | 68 | 443 | 91 | 151 | 469 | 445 | 78 | 2352 | 801 | 338 | 2171 |
| PBANKA_1318300 | 0 | 0 | 61 | 7 | 135 | 75 | 87 | 30 | 20 | 7 | 6 | 0 | 788 | 312 | 182 | 905 | 305 | 33 | 225 | 180 | 6 | 33 |
| PBANKA_1318400 | 0 | 0 | 374 | 140 | 903 | 530 | 740 | 259 | 214 | 116 | 41 | 38 | 2869 | 837 | 1714 | 6754 | 1602 | 178 | 498 | 222 | 9 | 84 |
| PBANKA_1318900 | 0 | 0 | 57 | 12 | 160 | 95 | 168 | 61 | 53 | 43 | 0 | 5 | 240 | 44 | 200 | 491 | 393 | 85 | 1077 | 389 | 15 | 196 |

| | | | | | | | | | | | | | | | | | | | | | | |
|---|---|---|---|---|---|---|---|---|---|---|---|---|---|---|---|---|---|---|---|---|---|---|
| PBANKA_1319000 | 15 | 7 | 9 | 0 | 116 | 88 | 112 | 43 | 1735 | 1071 | 876 | 342 | 1669 | 938 | 60 | 157 | 3185 | 1878 | 18146 | 12531 | 146 | 1203 |
| PBANKA_1319100 | 20 | 12 | 25 | 12 | 64 | 40 | 67 | 22 | 29 | 11 | 0 | 5 | 185 | 44 | 257 | 794 | 333 | 68 | 2941 | 1523 | 33 | 212 |
| PBANKA_1319200 | 21 | 28 | 91 | 34 | 459 | 264 | 677 | 288 | 266 | 112 | 18 | 15 | 1878 | 521 | 1194 | 4198 | 3753 | 552 | 1961 | 1347 | 699 | 5740 |
| PBANKA_1319300 | 26 | 40 | 35 | 0 | 508 | 228 | 1115 | 382 | 731 | 309 | 104 | 37 | 857 | 234 | 122 | 243 | 6355 | 945 | 459 | 628 | 12 | 113 |
| PBANKA_1319400 | 50 | 134 | 71 | 19 | 219 | 108 | 223 | 79 | 305 | 171 | 43 | 28 | 183 | 91 | 252 | 930 | 883 | 195 | 1113 | 899 | 337 | 2617 |
| PBANKA_1319500 | 0 | 0 | 11 | 0 | 29 | 11 | 23 | 11 | 32 | 9 | 0 | 0 | 3355 | 2181 | 78 | 321 | 2024 | 779 | 1E+05 | 1E+05 | 1430 | 12253 |
| PBANKA_1319600 | 0 | 0 | 33 | 8 | 250 | 88 | 205 | 81 | 79 | 50 | 16 | 8 | 73 | 9 | 145 | 391 | 327 | 90 | 116 | 47 | 0 | 42 |
| PBANKA_1319700 | 19 | 31 | 11 | 0 | 221 | 121 | 205 | 78 | 89 | 53 | 33 | 0 | 180 | 51 | 44 | 148 | 387 | 78 | 2234 | 1475 | 196 | 1612 |
| PBANKA_1319800 | 15 | 14 | 57 | 15 | 129 | 43 | 92 | 14 | 23 | 11 | 15 | 12 | 398 | 103 | 238 | 889 | 78 | 9 | 384 | 152 | 13 | 62 |
| PBANKA_1319900 | 5 | 0 | 60 | 16 | 127 | 49 | 58 | 17 | 38 | 15 | 0 | 0 | 791 | 254 | 221 | 1592 | 106 | 19 | 170 | 126 | 0 | 15 |
| PBANKA_1320000 | 55 | 65 | 18 | 0 | 141 | 48 | 128 | 64 | 135 | 53 | 21 | 6 | 308 | 76 | 93 | 448 | 1553 | 338 | 136 | 111 | 53 | 309 |
| PBANKA_1320200 | 428 | 634 | 374 | 84 | 1297 | 488 | 1633 | 565 | 705 | 266 | 117 | 47 | 1552 | 1071 | 1816 | 7859 | 14832 | 1567 | 6550 | 18864 | 285 | 2383 |
| PBANKA_1320300 | 0 | 0 | 102 | 21 | 172 | 70 | 123 | 53 | 57 | 20 | 8 | 0 | 404 | 58 | 236 | 726 | 220 | 48 | 76 | 40 | 0 | 23 |
| PBANKA_1320400 | 25 | 35 | 127 | 28 | 538 | 255 | 696 | 266 | 352 | 149 | 26 | 16 | 2150 | 710 | 780 | 3407 | 4471 | 690 | 1929 | 1321 | 90 | 950 |
| PBANKA_1320500 | 24 | 16 | 387 | 106 | 642 | 522 | 544 | 251 | 1393 | 1060 | 6844 | 5801 | 16515 | 5734 | 3319 | 16002 | 4743 | 1140 | 2845 | 1837 | 119 | 1046 |
| PBANKA_1320600 | 0 | 0 | 24 | 8 | 29 | 28 | 30 | 12 | 5 | 0 | 84 | 67 | 203 | 100 | 44 | 94 | 35 | 19 | 123 | 37 | 0 | 49 |
| PBANKA_1320700 | 0 | 0 | 220 | 88 | 477 | 332 | 1472 | 681 | 840 | 554 | 37 | 36 | 3616 | 687 | 926 | 2458 | 8858 | 2646 | 5063 | 2384 | 454 | 4076 |
| PBANKA_1320800 | 19 | 21 | 6 | 0 | 68 | 48 | 191 | 81 | 78 | 34 | 27 | 11 | 2392 | 1630 | 126 | 368 | 2356 | 706 | 57964 | 70812 | 2208 | 21060 |
| PBANKA_1320900 | 75 | 83 | 489 | 214 | 958 | 542 | 825 | 636 | 253 | 208 | 21 | 23 | 655 | 106 | 1887 | 3124 | 1567 | 562 | 1190 | 526 | 42 | 335 |
| PBANKA_1321000 | 0 | 0 | 184 | 53 | 658 | 253 | 654 | 262 | 172 | 78 | 807 | 561 | 3467 | 2133 | 882 | 4353 | 638 | 114 | 2199 | 1720 | 339 | 2641 |
| PBANKA_1321100 | 13 | 10 | 153 | 31 | 740 | 277 | 770 | 305 | 279 | 127 | 176 | 109 | 1104 | 985 | 1321 | 6450 | 2100 | 257 | 1383 | 1741 | 51 | 336 |
| PBANKA_1321200 | 5 | 11 | 35 | 15 | 262 | 162 | 396 | 169 | 228 | 84 | 37 | 12 | 544 | 173 | 301 | 1417 | 1226 | 220 | 1528 | 894 | 57 | 453 |
| PBANKA_1321300 | 26 | 52 | 107 | 23 | 207 | 67 | 211 | 78 | 112 | 41 | 26 | 8 | 1674 | 624 | 310 | 1226 | 1381 | 269 | 1140 | 645 | 67 | 469 |
| PBANKA_1321400 | 120 | 186 | 370 | 80 | 416 | 136 | 366 | 135 | 364 | 157 | 3438 | 1956 | 4780 | 8465 | 1122 | 7364 | 1749 | 341 | 872 | 1893 | 655 | 4628 |
| PBANKA_1321600 | 0 | 0 | 90 | 28 | 394 | 159 | 668 | 254 | 407 | 187 | 157 | 53 | 980 | 1356 | 1764 | 7728 | 9086 | 1139 | 974 | 5578 | 191 | 1385 |
| PBANKA_1321700 | 36 | 70 | 113 | 64 | 398 | 256 | 465 | 167 | 207 | 127 | 250 | 227 | 6271 | 2661 | 2179 | 9525 | 2082 | 338 | 1634 | 1083 | 396 | 2958 |
| PBANKA_1321800 | 0 | 0 | 14 | 0 | 13 | 10 | 36 | 18 | 13 | 0 | 0 | 0 | 43 | 6 | 56 | 121 | 45 | 22 | 102 | 26 | 0 | 16 |
| PBANKA_1321900 | 0 | 5 | 22 | 0 | 44 | 17 | 70 | 20 | 25 | 6 | 0 | 0 | 64 | 10 | 113 | 306 | 164 | 83 | 172 | 91 | 8 | 59 |
| PBANKA_1322000 | 147 | 239 | 52 | 15 | 250 | 118 | 531 | 143 | 359 | 149 | 27 | 13 | 730 | 371 | 947 | 4680 | 5640 | 907 | 1732 | 2728 | 242 | 1631 |
| PBANKA_1322100 | 0 | 5 | 56 | 8 | 185 | 103 | 271 | 125 | 109 | 53 | 22 | 5 | 166 | 61 | 82 | 271 | 1008 | 159 | 236 | 118 | 20 | 164 |
| PBANKA_1322200 | 0 | 0 | 30 | 11 | 76 | 22 | 191 | 80 | 149 | 83 | 46 | 11 | 620 | 525 | 221 | 1003 | 6153 | 712 | 890 | 3712 | 24 | 193 |
| PBANKA_1322300 | 8 | 12 | 70 | 10 | 211 | 99 | 278 | 99 | 162 | 76 | 66 | 32 | 756 | 612 | 487 | 2341 | 2082 | 347 | 811 | 1038 | 113 | 681 |
| PBANKA_1322400 | 6 | 26 | 93 | 13 | 184 | 96 | 215 | 95 | 84 | 34 | 17 | 0 | 585 | 173 | 340 | 1588 | 2023 | 253 | 2614 | 1875 | 40 | 261 |
| PBANKA_1322500 | 42 | 60 | 57 | 14 | 205 | 69 | 177 | 64 | 100 | 46 | 163 | 94 | 747 | 495 | 254 | 1543 | 480 | 66 | 378 | 411 | 166 | 1228 |
| PBANKA_1322600 | 47 | 75 | 29 | 13 | 125 | 79 | 172 | 99 | 65 | 63 | 9 | 0 | 174 | 45 | 96 | 186 | 581 | 152 | 266 | 162 | 39 | 295 |
| PBANKA_1322700 | 20 | 37 | 148 | 33 | 199 | 125 | 228 | 59 | 74 | 36 | 499 | 410 | 2919 | 2299 | 1081 | 6237 | 916 | 128 | 852 | 1151 | 693 | 5063 |
| PBANKA_1322800 | 13 | 24 | 94 | 20 | 178 | 97 | 145 | 45 | 38 | 19 | 8 | 7 | 889 | 467 | 586 | 2893 | 342 | 74 | 949 | 1227 | 224 | 1880 |
| PBANKA_1322900 | 0 | 0 | 52 | 15 | 135 | 47 | 157 | 44 | 72 | 22 | 20 | 0 | 178 | 61 | 46 | 190 | 828 | 188 | 852 | 662 | 28 | 137 |
| PBANKA_1323000 | 0 | 9 | 31 | 0 | 93 | 33 | 151 | 44 | 61 | 45 | 328 | 213 | 741 | 466 | 218 | 774 | 490 | 83 | 224 | 122 | 30 | 209 |
| PBANKA_1323100 | 15 | 10 | 68 | 15 | 93 | 52 | 147 | 38 | 47 | 11 | 32 | 18 | 988 | 707 | 180 | 825 | 413 | 52 | 613 | 315 | 54 | 382 |
| PBANKA_1323200 | 66 | 79 | 90 | 10 | 213 | 76 | 366 | 121 | 133 | 49 | 15 | 7 | 461 | 137 | 392 | 1897 | 1633 | 253 | 577 | 606 | 121 | 873 |
| PBANKA_1323300 | 0 | 0 | 22 | 12 | 51 | 47 | 53 | 19 | 17 | 14 | 0 | 5 | 103 | 32 | 152 | 399 | 276 | 88 | 730 | 351 | 36 | 253 |
| PBANKA_1323400 | 400 | 480 | 143 | 51 | 738 | 531 | 1210 | 657 | 665 | 403 | 230 | 175 | 1498 | 432 | 629 | 1292 | 2165 | 756 | 2638 | 1180 | 429 | 3165 |
| PBANKA_1323700 | 0 | 0 | 21 | 14 | 90 | 50 | 92 | 34 | 72 | 38 | 507 | 426 | 808 | 299 | 182 | 819 | 274 | 80 | 459 | 169 | 53 | 264 |
| PBANKA_1323900 | 0 | 0 | 29 | 12 | 80 | 49 | 86 | 23 | 15 | 17 | 6 | 0 | 478 | 146 | 223 | 680 | 226 | 82 | 4555 | 2808 | 26 | 204 |
| PBANKA_1324000 | 0 | 0 | 164 | 32 | 167 | 68 | 404 | 169 | 94 | 92 | 387 | 128 | 704 | 252 | 339 | 428 | 192 | 219 | 312 | 149 | 53 | 153 |
| PBANKA_1324100 | 0 | 8 | 94 | 30 | 160 | 165 | 206 | 109 | 32 | 26 | 6 | 0 | 394 | 107 | 204 | 480 | 205 | 31 | 463 | 370 | 27 | 255 |
| PBANKA_1324200 | 7 | 13 | 71 | 16 | 690 | 282 | 781 | 231 | 337 | 156 | 78 | 35 | 170 | 79 | 163 | 448 | 959 | 142 | 440 | 586 | 160 | 1360 |
| PBANKA_1324300 | 26 | 46 | 31 | 7 | 30 | 16 | 56 | 37 | 166 | 113 | 37 | 12 | 103 | 32 | 29 | 59 | 607 | 256 | 227 | 119 | 355 | 2705 |
| PBANKA_1324400 | 94 | 57 | 694 | 204 | 528 | 310 | 637 | 302 | 236 | 148 | 1214 | 1070 | 11514 | 3178 | 2119 | 6949 | 1912 | 419 | 2829 | 1121 | 284 | 1511 |
| PBANKA_1324500 | 0 | 0 | 30 | 0 | 110 | 53 | 127 | 65 | 50 | 34 | 11 | 15 | 202 | 57 | 156 | 627 | 295 | 41 | 66 | 33 | 12 | 46 |
| PBANKA_1324600 | 0 | 0 | 62 | 6 | 122 | 71 | 149 | 66 | 40 | 19 | 7 | 0 | 154 | 21 | 228 | 557 | 316 | 90 | 76 | 31 | 0 | 25 |
| PBANKA_1324700 | 0 | 0 | 18 | 0 | 56 | 26 | 40 | 13 | 35 | 16 | 0 | 8 | 27 | 8 | 16 | 83 | 36 | 14 | 677 | 261 | 30 | 217 |
| PBANKA_1324800 | 0 | 0 | 39 | 15 | 146 | 74 | 242 | 104 | 82 | 58 | 14 | 0 | 128 | 51 | 19 | 41 | 350 | 82 | 1498 | 1181 | 31 | 208 |
| PBANKA_1324900 | 8 | 26 | 54 | 11 | 68 | 30 | 47 | 23 | 31 | 29 | 436 | 359 | 787 | 325 | 139 | 433 | 192 | 57 | 1010 | 382 | 116 | 800 |
| PBANKA_1325000 | 106 | 99 | 367 | 106 | 241 | 136 | 238 | 136 | 153 | 120 | 685 | 581 | 6697 | 1494 | 1284 | 4296 | 2479 | 527 | 4369 | 2005 | 111 | 870 |
| PBANKA_1325300 | 5 | 14 | 128 | 33 | 285 | 170 | 370 | 148 | 148 | 69 | 155 | 134 | 2392 | 1385 | 624 | 3326 | 1565 | 159 | 1673 | 1295 | 143 | 1294 |
| PBANKA_1325500 | 17 | 50 | 37 | 9 | 46 | 24 | 46 | 18 | 45 | 17 | 11 | 7 | 582 | 484 | 186 | 1034 | 512 | 128 | 2480 | 4610 | 395 | 2817 |
| PBANKA_1325600 | 0 | 0 | 448 | 111 | 977 | 451 | 734 | 256 | 217 | 100 | 82 | 48 | 3776 | 2634 | 2054 | 8967 | 1895 | 185 | 184 | 235 | 32 | 285 |
| PBANKA_1325800 | 0 | 18 | 30 | 13 | 187 | 115 | 248 | 88 | 98 | 35 | 16 | 22 | 337 | 165 | 288 | 929 | 659 | 126 | 1127 | 548 | 67 | 555 |
| PBANKA_1325900 | 0 | 0 | 31 | 11 | 70 | 41 | 85 | 30 | 31 | 12 | 27 | 11 | 76 | 34 | 66 | 213 | 121 | 18 | 228 | 107 | 21 | 136 |
| PBANKA_1326000 | 385 | 679 | 285 | 81 | 442 | 162 | 930 | 439 | 395 | 178 | 98 | 48 | 3785 | 2902 | 967 | 5271 | 11056 | 1080 | 1735 | 2632 | 1039 | 7955 |
| PBANKA_1326200 | 0 | 0 | 8 | 0 | 29 | 9 | 12 | 12 | 16 | 0 | 0 | 0 | 233 | 52 | 55 | 96 | 203 | 60 | 2078 | 857 | 11 | 86 |
| PBANKA_1326300 | 0 | 0 | 286 | 109 | 773 | 473 | 1134 | 534 | 601 | 359 | 66 | 15 | 1296 | 257 | 720 | 1913 | 3436 | 901 | 1877 | 958 | 133 | 1335 |
| PBANKA_1326400 | 13 | 0 | 7694 | 3700 | 5412 | 4916 | 5032 | 2890 | 1413 | 1062 | 280 | 192 | 26773 | 5697 | 14968 | 34000 | 5445 | 1236 | 6570 | 1833 | 222 | 2599 |
| PBANKA_1326500 | 0 | 0 | 56 | 9 | 121 | 64 | 117 | 40 | 52 | 24 | 42 | 21 | 849 | 342 | 439 | 1786 | 551 | 77 | 461 | 284 | 34 | 184 |
| PBANKA_1326600 | 19 | 21 | 41 | 22 | 114 | 58 | 137 | 56 | 92 | 39 | 632 | 530 | 1554 | 715 | 166 | 948 | 1178 | 173 | 5433 | 2732 | 66 | 547 |
| PBANKA_1326700 | 14 | 16 | 272 | 34 | 1137 | 298 | 1430 | 461 | 653 | 240 | 84 | 45 | 483 | 444 | 2332 | 8612 | 3739 | 432 | 757 | 2390 | 52 | 273 |
| PBANKA_1326800 | 5 | 6 | 117 | 35 | 285 | 197 | 634 | 303 | 250 | 146 | 18 | 6 | 262 | 79 | 349 | 1111 | 1455 | 211 | 968 | 341 | 6 | 62 |
| PBANKA_1326900 | 20 | 23 | 31 | 5 | 145 | 70 | 1431 | 687 | 1017 | 619 | 62 | 39 | 316 | 79 | 253 | 713 | 1124 | 242 | 1225 | 629 | 0 | 40 |
| PBANKA_1327000 | 120 | 179 | 128 | 33 | 312 | 113 | 403 | 167 | 143 | 86 | 217 | 174 | 1758 | 1041 | 335 | 1783 | 513 | 87 | 914 | 604 | 76 | 523 |
| PBANKA_1327021 | 11 | 40 | 68 | 22 | 165 | 140 | 419 | 218 | 428 | 395 | 965 | 940 | 3579 | 2180 | 2053 | 6380 | 6057 | 1350 | 688 | 536 | 26 | 176 |
| PBANKA_1327041 | 0 | 7 | 8 | 7 | 12 | 6 | 19 | 7 | 8 | 0 | 0 | 0 | 22 | 7 | 16 | 39 | 33 | 29 | 80 | 9 | 5 | 8 |
| PBANKA_1327061 | 22 | 48 | 23 | 0 | 118 | 33 | 96 | 60 | 38 | 19 | 0 | 0 | 152 | 31 | 167 | 543 | 555 | 64 | 394 | 192 | 41 | 284 |
| PBANKA_1327080 | 10 | 0 | 30 | 9 | 40 | 16 | 68 | 34 | 10 | 9 | 0 | 0 | 57 | 9 | 106 | 202 | 193 | 41 | 303 | 143 | 16 | 67 |
| PBANKA_1327200 | 7 | 12 | 28 | 10 | 61 | 38 | 74 | 35 | 16 | 9 | 0 | 7 | 111 | 20 | 174 | 461 | 98 | 26 | 113 | 75 | 7 | 27 |
| PBANKA_1327251 | 0 | 11 | 5 | 8 | 80 | 42 | 113 | 55 | 218 | 198 | 881 | 696 | 1645 | 814 | 421 | 959 | 809 | 383 | 504 | 220 | 19 | 129 |
| PBANKA_1327300 | 0 | 0 | 102 | 29 | 218 | 210 | 339 | 218 | 618 | 644 | 2220 | 1752 | 2929 | 2298 | 1337 | 3849 | 2438 | 791 | 588 | 295 | 0 | 78 |
| PBANKA_1327400 | 11 | 29 | 196 | 40 | 418 | 237 | 481 | 192 | 207 | 110 | 62 | 17 | 1989 | 2509 | 1025 | 7267 | 4197 | 354 | 96 | 390 | 66 | 358 |
| PBANKA_1327500 | 15 | 5 | 62 | 21 | 162 | 85 | 243 | 96 | 129 | 43 | 14 | 14 | 420 | 273 | 204 | 737 | 690 | 115 | 99 | 52 | 30 | 174 |
| PBANKA_1327600 | 156 | 213 | 52 | 5 | 256 | 105 | 214 | 77 | 82 | 34 | 143 | 120 | 443 | 258 | 100 | 414 | 311 | 56 | 494 | 222 | 30 | 98 |
| PBANKA_1327700 | 12 | 13 | 95 | 15 | 97 | 35 | 102 | 37 | 41 | 16 | 223 | 148 | 583 | 189 | 71 | 197 | 165 | 55 | 1044 | 791 | 28 | 254 |
| PBANKA_1327800 | 5 | 0 | 111 | 38 | 71 | 63 | 55 | 42 | 27 | 28 | 298 | 246 | 624 | 314 | 92 | 563 | 197 | 29 | 416 | 242 | 81 | 499 |
| PBANKA_1327900 | 52 | 66 | 352 | 92 | 1334 | 413 | 1458 | 423 | 654 | 262 | 689 | 306 | 1970 | 3636 | 2887 | 15243 | 7461 | 840 | 1242 | 5090 | 222 | 1279 |
| PBANKA_1328000 | 5108 | 7602 | 186 | 38 | 535 | 238 | 425 | 129 | 359 | 178 | 167 | 99 | 1577 | 1234 | 557 | 3296 | 1232 | 339 | 865 | 1077 | 32 | 172 |
| PBANKA_1328100 | 132 | 136 | 41 | 18 | 181 | 93 | 325 | 134 | 96 | 85 | 18 | 5 | 758 | 249 | 399 | 1556 | 1724 | 304 | 1902 | 964 | 1132 | 8860 |
| PBANKA_1328300 | 48 | 95 | 247 | 61 | 351 | 211 | 379 | 115 | 127 | 74 | 143 | 104 | 2011 | 1350 | 596 | 2674 | 827 | 131 | 924 | 1020 | 100 | 802 |
| PBANKA_1328400 | 39 | 57 | 156 | 42 | 422 | 236 | 641 | 274 | 321 | 228 | 51 | 11 | 460 | 148 | 605 | 1740 | 2195 | 539 | 2924 | 1065 | 322 | 3052 |
| PBANKA_1328500 | 7 | 9 | 22 | 5 | 68 | 26 | 111 | 58 | 50 | 15 | 17 | 6 | 550 | 194 | 245 | 1314 | 695 | 113 | 280 | 211 | 33 | 284 |
| PBANKA_1328600 | 0 | 0 | 56 | 6 | 68 | 50 | 118 | 70 | 32 | 24 | 0 | 16 | 217 | 70 | 181 | 751 | 552 | 61 | 24 | 20 | 0 | 8 |

| | | | | | | | | | | | | | | | | | | | | | | |
|---|---|---|---|---|---|---|---|---|---|---|---|---|---|---|---|---|---|---|---|---|---|---|
| PBANKA_1328700 | 0 | 0 | 47 | 9 | 116 | 60 | 172 | 81 | 78 | 21 | 26 | 6 | 900 | 320 | 483 | 2136 | 675 | 113 | 612 | 221 | 6 | 50 |
| PBANKA_1328800 | 329 | 318 | 71 | 27 | 103 | 70 | 161 | 71 | 123 | 104 | 241 | 180 | 458 | 268 | 150 | 432 | 563 | 253 | 1513 | 704 | 178 | 879 |
| PBANKA_1329200 | 59 | 45 | 11 | 0 | 626 | 200 | 655 | 169 | 414 | 178 | 95 | 30 | 888 | 1178 | 527 | 2066 | 4461 | 693 | 3461 | 20758 | 143 | 987 |
| PBANKA_1329300 | 134 | 228 | 2242 | 584 | 1842 | 1311 | 2035 | 1030 | 543 | 399 | 2856 | 2488 | 15511 | 5931 | 3627 | 14778 | 4479 | 561 | 2241 | 1628 | 329 | 3318 |
| PBANKA_1329500 | 482 | 727 | 111 | 21 | 395 | 282 | 821 | 310 | 461 | 211 | 39 | 31 | 616 | 238 | 396 | 910 | 4296 | 643 | 3524 | 2541 | 168 | 1644 |
| PBANKA_1329600 | 19 | 26 | 32 | 0 | 106 | 59 | 165 | 60 | 351 | 216 | 3313 | 2574 | 2065 | 2025 | 542 | 2668 | 1976 | 393 | 2899 | 3978 | 51 | 432 |
| PBANKA_1329700 | 67 | 65 | 547 | 153 | 1522 | 809 | 2056 | 785 | 776 | 361 | 151 | 48 | 4712 | 1893 | 4043 | 18498 | 12545 | 1508 | 3956 | 3330 | 374 | 3112 |
| PBANKA_1329800 | 1604 | 1705 | 345 | 126 | 498 | 263 | 552 | 191 | 853 | 452 | 2573 | 1890 | 3209 | 2910 | 340 | 2353 | 4071 | 598 | 1878 | 2095 | 253 | 2185 |
| PBANKA_1330000 | 0 | 0 | 5 | 0 | 13 | 6 | 11 | 0 | 13 | 0 | 0 | 0 | 495 | 187 | 12 | 98 | 158 | 59 | 3143 | 1391 | 31 | 160 |
| PBANKA_1330100 | 108 | 122 | 110 | 33 | 68 | 56 | 83 | 39 | 34 | 26 | 371 | 314 | 436 | 216 | 76 | 294 | 73 | 15 | 467 | 212 | 500 | 2897 |
| PBANKA_1330200 | 64 | 130 | 99 | 16 | 179 | 70 | 223 | 58 | 153 | 72 | 192 | 103 | 2253 | 1659 | 367 | 2088 | 1178 | 349 | 1564 | 1633 | 855 | 4899 |
| PBANKA_1330300 | 20 | 43 | 137 | 8 | 466 | 129 | 567 | 187 | 261 | 125 | 64 | 26 | 960 | 593 | 486 | 2558 | 4379 | 558 | 515 | 1254 | 100 | 668 |
| PBANKA_1330400 | 0 | 5 | 32 | 0 | 42 | 26 | 55 | 20 | 17 | 8 | 6 | 0 | 603 | 333 | 228 | 899 | 260 | 52 | 66 | 37 | 318 | 2359 |
| PBANKA_1330500 | 0 | 11 | 12 | 0 | 40 | 12 | 49 | 18 | 12 | 9 | 0 | 11 | 238 | 184 | 176 | 785 | 216 | 30 | 40 | 27 | 494 | 4238 |
| PBANKA_1330600 | 0 | 5 | 182 | 28 | 397 | 155 | 336 | 141 | 155 | 48 | 7 | 10 | 475 | 105 | 598 | 2366 | 1324 | 257 | 176 | 101 | 285 | 1892 |
| PBANKA_1330700 | 6 | 6 | 80 | 19 | 210 | 79 | 197 | 84 | 74 | 22 | 10 | 0 | 1142 | 392 | 456 | 2084 | 1260 | 208 | 9597 | 6594 | 1518 | 10314 |
| PBANKA_1330800 | 99 | 166 | 119 | 17 | 252 | 126 | 264 | 138 | 90 | 46 | 31 | 14 | 1374 | 829 | 406 | 2566 | 1513 | 202 | 1105 | 978 | 744 | 4789 |
| PBANKA_1330900 | 5 | 0 | 7 | 0 | 35 | 0 | 63 | 21 | 37 | 24 | 15 | 0 | 1121 | 346 | 59 | 321 | 713 | 197 | 12695 | 8219 | 196 | 981 |
| PBANKA_1331000 | 0 | 0 | 44 | 13 | 34 | 12 | 38 | 16 | 12 | 8 | 0 | 0 | 127 | 25 | 93 | 215 | 78 | 18 | 182 | 83 | 25 | 129 |
| PBANKA_1331100 | 47 | 51 | 27 | 12 | 134 | 74 | 243 | 138 | 96 | 78 | 14 | 9 | 216 | 32 | 103 | 307 | 605 | 172 | 496 | 129 | 33 | 226 |
| PBANKA_1331200 | 306 | 461 | 74 | 16 | 338 | 142 | 992 | 379 | 716 | 297 | 101 | 33 | 1491 | 748 | 749 | 2822 | 11501 | 1892 | 2792 | 3844 | 1418 | 10565 |
| PBANKA_1331300 | 14 | 6 | 15 | 0 | 138 | 37 | 150 | 54 | 89 | 49 | 33 | 18 | 485 | 491 | 85 | 455 | 1505 | 304 | 2237 | 10466 | 141 | 854 |
| PBANKA_1331400 | 3547 | 6034 | 1233 | 330 | 2666 | 1209 | 3704 | 1515 | 2192 | 1119 | 841 | 590 | 12180 | 22666 | 7910 | 48893 | 65359 | 6944 | 3002 | 14387 | 838 | 6446 |
| PBANKA_1331500 | 5 | 10 | 13 | 0 | 24 | 0 | 29 | 10 | 14 | 7 | 0 | 0 | 420 | 269 | 72 | 486 | 249 | 53 | 359 | 197 | 26 | 160 |
| PBANKA_1331800 | 30 | 44 | 71 | 15 | 138 | 72 | 234 | 107 | 93 | 57 | 9 | 7 | 622 | 52 | 350 | 1044 | 1297 | 326 | 903 | 428 | 80 | 559 |
| PBANKA_1331900 | 28 | 25 | 1621 | 500 | 1139 | 846 | 747 | 354 | 371 | 238 | 4025 | 3267 | 18429 | 9343 | 3405 | 19090 | 2595 | 306 | 2052 | 1905 | 302 | 2838 |
| PBANKA_1332000 | 0 | 0 | 357 | 71 | 722 | 348 | 783 | 309 | 390 | 199 | 597 | 325 | 5415 | 6280 | 2105 | 12868 | 5919 | 648 | 1026 | 1767 | 98 | 932 |
| PBANKA_1332100 | 0 | 5 | 165 | 35 | 402 | 109 | 383 | 109 | 145 | 53 | 150 | 88 | 3603 | 2711 | 1211 | 7699 | 4418 | 489 | 1160 | 1784 | 364 | 2439 |
| PBANKA_1332200 | 0 | 0 | 28 | 5 | 58 | 20 | 87 | 30 | 36 | 19 | 8 | 0 | 499 | 232 | 80 | 536 | 475 | 93 | 335 | 237 | 8 | 100 |
| PBANKA_1332400 | 0 | 0 | 14 | 0 | 165 | 80 | 407 | 237 | 198 | 112 | 16 | 10 | 546 | 113 | 275 | 1076 | 2285 | 406 | 1148 | 794 | 64 | 586 |
| PBANKA_1332500 | 10 | 7 | 71 | 14 | 82 | 41 | 69 | 20 | 62 | 27 | 520 | 486 | 1598 | 781 | 296 | 1410 | 343 | 82 | 708 | 421 | 135 | 904 |
| PBANKA_1332600 | 0 | 0 | 125 | 19 | 263 | 115 | 182 | 88 | 146 | 66 | 57 | 34 | 1938 | 855 | 1137 | 5347 | 1415 | 197 | 693 | 796 | 19 | 131 |
| PBANKA_1332700 | 5 | 0 | 7 | 11 | 207 | 143 | 243 | 108 | 1972 | 1212 | 770 | 644 | 1981 | 1836 | 74 | 412 | 4248 | 2404 | 739 | 959 | 79 | 607 |
| PBANKA_1332800 | 269 | 372 | 344 | 75 | 6625 | 3216 | 6864 | 2203 | 2313 | 966 | 334 | 200 | 215 | 370 | 294 | 1210 | 1444 | 309 | 2602 | 11309 | 456 | 4176 |
| PBANKA_1332900 | 0 | 0 | 35 | 8 | 106 | 22 | 97 | 34 | 40 | 18 | 6 | 0 | 134 | 40 | 225 | 995 | 664 | 125 | 264 | 100 | 14 | 78 |
| PBANKA_1333000 | 16 | 13 | 25 | 0 | 90 | 43 | 123 | 38 | 31 | 18 | 0 | 6 | 139 | 51 | 169 | 622 | 368 | 90 | 1259 | 441 | 17 | 125 |
| PBANKA_1333200 | 9 | 12 | 15 | 9 | 54 | 22 | 60 | 24 | 19 | 9 | 0 | 0 | 52 | 0 | 38 | 127 | 76 | 14 | 241 | 108 | 9 | 28 |
| PBANKA_1333300 | 49 | 46 | 64 | 13 | 391 | 123 | 396 | 128 | 192 | 81 | 90 | 65 | 633 | 301 | 677 | 2294 | 1365 | 235 | 947 | 1346 | 110 | 791 |
| PBANKA_1333400 | 26 | 40 | 27 | 0 | 91 | 24 | 70 | 17 | 64 | 32 | 175 | 107 | 520 | 322 | 107 | 509 | 309 | 93 | 304 | 256 | 83 | 576 |
| PBANKA_1333500 | 50 | 65 | 45 | 7 | 74 | 26 | 76 | 41 | 47 | 19 | 66 | 45 | 571 | 242 | 190 | 693 | 578 | 215 | 344 | 165 | 98 | 628 |
| PBANKA_1333600 | 84 | 118 | 47 | 24 | 92 | 49 | 147 | 75 | 30 | 30 | 8 | 0 | 117 | 29 | 108 | 276 | 336 | 104 | 385 | 145 | 27 | 210 |
| PBANKA_1333800 | 47 | 95 | 181 | 28 | 246 | 119 | 209 | 102 | 83 | 64 | 61 | 25 | 2534 | 1387 | 548 | 2256 | 753 | 188 | 810 | 483 | 124 | 757 |
| PBANKA_1333900 | 8 | 17 | 74 | 20 | 196 | 67 | 173 | 70 | 113 | 30 | 22 | 15 | 1374 | 785 | 397 | 1897 | 910 | 178 | 780 | 664 | 87 | 665 |
| PBANKA_1334000 | 0 | 0 | 135 | 26 | 342 | 99 | 502 | 156 | 257 | 62 | 61 | 11 | 1275 | 620 | 1065 | 5181 | 6037 | 769 | 515 | 1016 | 19 | 158 |
| PBANKA_1334100 | 0 | 0 | 297 | 57 | 408 | 234 | 697 | 322 | 217 | 142 | 28 | 11 | 1231 | 191 | 995 | 3406 | 2924 | 463 | 848 | 318 | 12 | 135 |
| PBANKA_1334200 | 17 | 25 | 138 | 42 | 172 | 85 | 135 | 59 | 116 | 65 | 968 | 594 | 1861 | 1456 | 392 | 2243 | 309 | 69 | 389 | 471 | 555 | 4324 |
| PBANKA_1334300 | 17 | 10 | 631 | 179 | 837 | 439 | 672 | 288 | 1585 | 1228 | 9661 | 7266 | 11552 | 5067 | 2709 | 11591 | 2431 | 866 | 741 | 598 | 49 | 462 |
| PBANKA_1334400 | 77 | 68 | 63 | 16 | 115 | 85 | 144 | 61 | 111 | 37 | 42 | 41 | 1444 | 413 | 582 | 2182 | 1270 | 321 | 2400 | 1242 | 73 | 653 |
| PBANKA_1334500 | 0 | 0 | 19 | 7 | 80 | 27 | 74 | 29 | 23 | 12 | 11 | 7 | 294 | 57 | 128 | 668 | 199 | 35 | 425 | 231 | 12 | 69 |
| PBANKA_1334600 | 0 | 0 | 60 | 17 | 184 | 110 | 242 | 88 | 519 | 327 | 573 | 392 | 2369 | 1179 | 250 | 1135 | 1486 | 510 | 1203 | 534 | 60 | 545 |
| PBANKA_1334800 | 0 | 0 | 7 | 5 | 25 | 11 | 27 | 13 | 34 | 12 | 9 | 0 | 1420 | 1588 | 53 | 296 | 1200 | 491 | 32475 | 47408 | 2053 | 15388 |
| PBANKA_1335100 | 97 | 133 | 93 | 22 | 355 | 159 | 968 | 361 | 608 | 249 | 250 | 137 | 877 | 1234 | 477 | 2412 | 2272 | 342 | 1357 | 3476 | 291 | 1990 |
| PBANKA_1335200 | 58 | 79 | 40 | 12 | 94 | 37 | 131 | 64 | 52 | 32 | 16 | 18 | 886 | 429 | 217 | 846 | 515 | 130 | 1173 | 478 | 73 | 438 |
| PBANKA_1335300 | 41 | 65 | 52 | 14 | 129 | 85 | 239 | 100 | 107 | 54 | 32 | 12 | 1067 | 379 | 290 | 1199 | 1199 | 244 | 1046 | 462 | 95 | 888 |
| PBANKA_1335400 | 133 | 177 | 101 | 27 | 335 | 120 | 558 | 179 | 334 | 97 | 204 | 133 | 3583 | 3191 | 1035 | 5354 | 6126 | 753 | 2166 | 4100 | 781 | 4920 |
| PBANKA_1335500 | 0 | 5 | 24 | 7 | 125 | 101 | 153 | 57 | 138 | 109 | 50 | 49 | 1375 | 511 | 521 | 1631 | 755 | 210 | 564 | 334 | 95 | 502 |
| PBANKA_1335600 | 10 | 12 | 17 | 5 | 59 | 17 | 54 | 21 | 49 | 22 | 9 | 0 | 119 | 53 | 211 | 861 | 143 | 24 | 1374 | 634 | 37 | 245 |
| PBANKA_1335700 | 0 | 0 | 6 | 0 | 15 | 0 | 45 | 10 | 12 | 13 | 0 | 0 | 157 | 52 | 15 | 35 | 141 | 57 | 1793 | 796 | 135 | 956 |
| PBANKA_1335900 | 13 | 29 | 114 | 20 | 253 | 144 | 349 | 118 | 127 | 61 | 12 | 5 | 971 | 181 | 839 | 3499 | 3553 | 584 | 796 | 405 | 6 | 75 |
| PBANKA_1336000 | 0 | 0 | 90 | 23 | 124 | 71 | 101 | 76 | 26 | 10 | 6 | 0 | 368 | 41 | 351 | 1634 | 415 | 51 | 507 | 131 | 38 | 183 |
| PBANKA_1336100 | 0 | 0 | 42 | 0 | 132 | 74 | 116 | 60 | 46 | 16 | 0 | 5 | 216 | 34 | 313 | 953 | 160 | 29 | 434 | 127 | 6 | 41 |
| PBANKA_1336300 | 18 | 20 | 39 | 5 | 178 | 75 | 317 | 153 | 142 | 43 | 19 | 0 | 527 | 136 | 438 | 1404 | 1758 | 291 | 1842 | 899 | 142 | 970 |
| PBANKA_1336400 | 5 | 12 | 81 | 12 | 271 | 138 | 734 | 298 | 364 | 168 | 61 | 27 | 2048 | 286 | 538 | 1896 | 4019 | 654 | 3671 | 2277 | 104 | 540 |
| PBANKA_1336500 | 0 | 0 | 7 | 0 | 43 | 22 | 80 | 34 | 75 | 18 | 10 | 0 | 225 | 88 | 171 | 686 | 659 | 94 | 1080 | 966 | 15 | 90 |
| PBANKA_1336700 | 64 | 110 | 84 | 27 | 170 | 88 | 238 | 61 | 134 | 65 | 21 | 28 | 1121 | 462 | 260 | 1197 | 525 | 127 | 1626 | 769 | 227 | 1760 |
| PBANKA_1336800 | 0 | 0 | 13 | 5 | 23 | 19 | 23 | 9 | 15 | 11 | 0 | 0 | 28 | 5 | 44 | 128 | 21 | 9 | 221 | 85 | 519 | 3530 |
| PBANKA_1336900 | 130 | 186 | 43 | 19 | 276 | 125 | 561 | 232 | 243 | 131 | 36 | 11 | 882 | 350 | 17 | 89 | 3263 | 496 | 5311 | 3186 | 229 | 1549 |
| PBANKA_1337000 | 359 | 521 | 147 | 45 | 392 | 126 | 428 | 130 | 307 | 168 | 1372 | 896 | 1556 | 1287 | 296 | 1641 | 1005 | 227 | 2089 | 1380 | 773 | 3942 |
| PBANKA_1337100 | 5 | 15 | 432 | 101 | 881 | 414 | 951 | 425 | 339 | 172 | 318 | 206 | 7529 | 8807 | 2941 | 23298 | 10249 | 658 | 1589 | 6988 | 229 | 2185 |
| PBANKA_1337200 | 22 | 28 | 43 | 9 | 110 | 52 | 209 | 105 | 254 | 167 | 85 | 58 | 2244 | 756 | 358 | 1766 | 3184 | 910 | 1622 | 814 | 67 | 549 |
| PBANKA_1337300 | 0 | 0 | 8 | 0 | 77 | 38 | 91 | 42 | 45 | 24 | 17 | 5 | 666 | 151 | 294 | 1056 | 632 | 229 | 1287 | 529 | 151 | 1451 |
| PBANKA_1337400 | 0 | 0 | 86 | 14 | 204 | 82 | 219 | 54 | 72 | 23 | 21 | 0 | 1324 | 460 | 540 | 2116 | 728 | 176 | 1744 | 994 | 156 | 1015 |
| PBANKA_1337500 | 174 | 298 | 197 | 49 | 615 | 238 | 1035 | 389 | 577 | 219 | 424 | 227 | 1095 | 867 | 538 | 1707 | 10558 | 1555 | 3965 | 7616 | 791 | 4019 |
| PBANKA_1337600 | 319 | 467 | 48 | 14 | 179 | 73 | 270 | 118 | 167 | 63 | 31 | 0 | 721 | 269 | 233 | 1138 | 3916 | 510 | 3600 | 2992 | 731 | 4938 |
| PBANKA_1337800 | 19 | 32 | 99 | 24 | 361 | 178 | 442 | 181 | 188 | 115 | 25 | 17 | 1099 | 291 | 551 | 2443 | 4157 | 606 | 1648 | 1121 | 319 | 2332 |
| PBANKA_1337900 | 0 | 0 | 71 | 12 | 135 | 51 | 104 | 42 | 48 | 13 | 0 | 0 | 93 | 35 | 48 | 208 | 93 | 35 | 836 | 696 | 277 | 2222 |
| PBANKA_1338000 | 0 | 0 | 125 | 19 | 96 | 50 | 70 | 35 | 21 | 12 | 0 | 0 | 104 | 18 | 215 | 422 | 163 | 57 | 987 | 326 | 54 | 343 |
| PBANKA_1338200 | 10 | 0 | 83 | 24 | 265 | 200 | 463 | 306 | 167 | 106 | 12 | 9 | 92 | 27 | 22 | 23 | 140 | 57 | 1985 | 643 | 186 | 998 |
| PBANKA_1338300 | 0 | 13 | 1046 | 301 | 839 | 427 | 953 | 468 | 332 | 192 | 1700 | 1436 | 6498 | 2789 | 2079 | 5057 | 1854 | 398 | 986 | 465 | 290 | 2345 |
| PBANKA_1338400 | 0 | 0 | 100 | 32 | 156 | 134 | 249 | 160 | 86 | 59 | 28 | 14 | 1204 | 334 | 264 | 766 | 646 | 142 | 1327 | 633 | 76 | 608 |
| PBANKA_1338500 | 0 | 0 | 27 | 0 | 76 | 17 | 106 | 35 | 27 | 12 | 13 | 6 | 679 | 205 | 129 | 998 | 706 | 103 | 1605 | 1566 | 130 | 830 |
| PBANKA_1338600 | 79 | 118 | 1153 | 280 | 637 | 443 | 881 | 504 | 399 | 229 | 1640 | 1410 | 25151 | 10664 | 3307 | 14655 | 7663 | 1356 | 3174 | 2250 | 185 | 1033 |
| PBANKA_1338700 | 0 | 0 | 111 | 39 | 329 | 264 | 503 | 211 | 361 | 166 | 55 | 19 | 1325 | 390 | 289 | 930 | 6949 | 1494 | 5304 | 3985 | 366 | 2861 |
| PBANKA_1338800 | 0 | 5 | 76 | 21 | 267 | 103 | 264 | 104 | 94 | 31 | 6 | 6 | 897 | 204 | 423 | 1648 | 1526 | 224 | 430 | 354 | 37 | 243 |
| PBANKA_1338900 | 0 | 0 | 9 | 0 | 136 | 115 | 1082 | 472 | 2370 | 1430 | 365 | 217 | 2791 | 900 | 59 | 144 | 15658 | 4846 | 46814 | 26865 | 1305 | 15328 |
| PBANKA_1339000 | 176 | 226 | 49 | 10 | 374 | 151 | 437 | 152 | 198 | 120 | 403 | 332 | 1111 | 687 | 364 | 1802 | 1477 | 215 | 636 | 682 | 327 | 2166 |
| PBANKA_1339100 | 24 | 31 | 52 | 13 | 107 | 41 | 103 | 51 | 68 | 26 | 213 | 113 | 1808 | 834 | 269 | 1669 | 800 | 148 | 490 | 343 | 179 | 1075 |
| PBANKA_1339200 | 18 | 7 | 120 | 16 | 176 | 55 | 163 | 41 | 127 | 22 | 247 | 126 | 793 | 640 | 259 | 1435 | 321 | 61 | 630 | 448 | 80 | 497 |
| PBANKA_1339300 | 0 | 0 | 62 | 26 | 278 | 132 | 402 | 155 | 99 | 61 | 10 | 10 | 806 | 186 | 60 | 114 | 2797 | 369 | 1946 | 954 | 247 | 1942 |
| PBANKA_1339400 | 71 | 83 | 153 | 40 | 365 | 128 | 275 | 127 | 235 | 132 | 49 | 38 | 3680 | 2106 | 1294 | 8100 | 5142 | 850 | 2757 | 5796 | 338 | 1925 |

| | | | | | | | | | | | | | | | | | | | | | | |
|---|---|---|---|---|---|---|---|---|---|---|---|---|---|---|---|---|---|---|---|---|---|---|
| PBANKA_1339500 | 129 | 225 | 234 | 37 | 578 | 146 | 504 | 118 | 335 | 137 | 185 | 73 | 925 | 1605 | 1016 | 5508 | 4388 | 712 | 1401 | 5529 | 902 | 5537 |
| PBANKA_1339600 | 95 | 99 | 97 | 29 | 431 | 218 | 369 | 149 | 134 | 67 | 65 | 36 | 511 | 199 | 176 | 658 | 357 | 97 | 2593 | 1016 | 839 | 5911 |
| PBANKA_1339700 | 0 | 6 | 35 | 9 | 123 | 48 | 141 | 36 | 71 | 30 | 14 | 5 | 550 | 271 | 524 | 2718 | 2016 | 241 | 429 | 516 | 43 | 152 |
| PBANKA_1339800 | 0 | 0 | 62 | 6 | 177 | 75 | 205 | 68 | 148 | 52 | 16 | 7 | 438 | 112 | 193 | 832 | 631 | 156 | 1429 | 838 | 56 | 371 |
| PBANKA_1339900 | 0 | 14 | 57 | 28 | 139 | 91 | 176 | 87 | 170 | 125 | 27 | 16 | 1497 | 406 | 373 | 1391 | 642 | 154 | 479 | 339 | 166 | 1379 |
| PBANKA_1340000 | 0 | 0 | 202 | 40 | 186 | 116 | 185 | 70 | 30 | 28 | 0 | 0 | 512 | 57 | 627 | 2241 | 284 | 47 | 36 | 19 | 124 | 991 |
| PBANKA_1340100 | 159 | 241 | 4817 | 1433 | 4122 | 2833 | 3444 | 1391 | 1023 | 593 | 611 | 303 | 21038 | 6322 | 8760 | 27241 | 3321 | 636 | 2884 | 1391 | 425 | 3838 |
| PBANKA_1340200 | 0 | 0 | 16 | 6 | 35 | 20 | 62 | 14 | 14 | 9 | 6 | 0 | 287 | 60 | 161 | 596 | 179 | 30 | 1023 | 458 | 136 | 1027 |
| PBANKA_1340300 | 0 | 0 | 230 | 69 | 198 | 125 | 142 | 59 | 76 | 30 | 940 | 793 | 4107 | 1752 | 1109 | 4395 | 357 | 68 | 484 | 294 | 7 | 63 |
| PBANKA_1340500 | 112 | 199 | 322 | 100 | 772 | 357 | 1325 | 560 | 1367 | 639 | 287 | 180 | 513 | 829 | 359 | 958 | 14752 | 1221 | 558 | 1044 | 145 | 930 |
| PBANKA_1340600 | 64 | 98 | 95 | 14 | 157 | 90 | 216 | 75 | 93 | 39 | 9 | 7 | 493 | 108 | 209 | 950 | 420 | 83 | 3075 | 2859 | 100 | 560 |
| PBANKA_1340700 | 12 | 17 | 30 | 0 | 79 | 34 | 164 | 73 | 115 | 74 | 15 | 6 | 42 | 14 | 19 | 52 | 279 | 62 | 357 | 174 | 50 | 395 |
| PBANKA_1340800 | 14 | 21 | 23 | 6 | 121 | 87 | 148 | 75 | 56 | 40 | 8 | 0 | 31 | 10 | 39 | 90 | 110 | 33 | 128 | 58 | 13 | 83 |
| PBANKA_1340900 | 294 | 399 | 443 | 118 | 1519 | 486 | 1290 | 404 | 456 | 153 | 81 | 27 | 130 | 86 | 112 | 544 | 491 | 86 | 420 | 1088 | 96 | 483 |
| PBANKA_1341000 | 12 | 13 | 14 | 0 | 297 | 99 | 749 | 304 | 490 | 210 | 411 | 220 | 806 | 1196 | 306 | 1190 | 12276 | 1254 | 1004 | 6662 | 200 | 1319 |
| PBANKA_1341100 | 12 | 23 | 64 | 25 | 185 | 110 | 480 | 195 | 354 | 236 | 83 | 24 | 569 | 114 | 369 | 1252 | 2305 | 827 | 2217 | 1169 | 121 | 934 |
| PBANKA_1341200 | 7 | 7 | 76 | 19 | 253 | 97 | 349 | 149 | 147 | 82 | 20 | 0 | 381 | 96 | 406 | 1493 | 1189 | 159 | 717 | 495 | 223 | 1312 |
| PBANKA_1341300 | 18 | 29 | 17 | 6 | 46 | 14 | 117 | 35 | 81 | 13 | 5 | 0 | 252 | 59 | 87 | 256 | 903 | 187 | 694 | 625 | 339 | 2141 |
| PBANKA_1341500 | 0 | 7 | 329 | 92 | 554 | 294 | 718 | 258 | 313 | 110 | 111 | 87 | 3652 | 2282 | 1241 | 6830 | 2669 | 282 | 852 | 693 | 68 | 628 |
| PBANKA_1341600 | 0 | 0 | 133 | 25 | 220 | 104 | 254 | 103 | 127 | 37 | 15 | 20 | 916 | 328 | 337 | 1252 | 446 | 80 | 310 | 209 | 46 | 294 |
| PBANKA_1341700 | 0 | 0 | 24 | 0 | 106 | 36 | 100 | 26 | 74 | 25 | 20 | 9 | 281 | 159 | 49 | 194 | 461 | 120 | 840 | 2172 | 970 | 6849 |
| PBANKA_1341800 | 16 | 37 | 332 | 78 | 1219 | 358 | 1552 | 548 | 781 | 356 | 247 | 68 | 1353 | 1669 | 2678 | 10663 | 7763 | 1027 | 2839 | 6496 | 456 | 2604 |
| PBANKA_1341900 | 52 | 85 | 23 | 0 | 84 | 28 | 189 | 72 | 155 | 58 | 28 | 7 | 490 | 258 | 146 | 759 | 1035 | 156 | 794 | 603 | 672 | 4217 |
| PBANKA_1342000 | 151 | 227 | 62 | 11 | 225 | 55 | 318 | 107 | 143 | 78 | 24 | 7 | 715 | 194 | 194 | 913 | 1810 | 325 | 1256 | 790 | 3819 | 19443 |
| PBANKA_1342100 | 47 | 26 | 19 | 0 | 139 | 45 | 104 | 34 | 55 | 21 | 26 | 10 | 494 | 243 | 226 | 1008 | 445 | 67 | 739 | 436 | 155 | 793 |
| PBANKA_1342200 | 14 | 16 | 34 | 10 | 68 | 38 | 61 | 31 | 57 | 29 | 25 | 7 | 545 | 237 | 151 | 740 | 144 | 27 | 631 | 282 | 364 | 2523 |
| PBANKA_1342400 | 0 | 5 | 52 | 8 | 88 | 62 | 79 | 37 | 63 | 59 | 270 | 165 | 1019 | 472 | 162 | 733 | 478 | 102 | 701 | 580 | 17 | 178 |
| PBANKA_1342500 | 1367 | 2244 | 49 | 7 | 179 | 82 | 217 | 91 | 1502 | 917 | 764 | 331 | 2518 | 3887 | 142 | 627 | 9612 | 2482 | 32535 | 1E+05 | 17845 | 1E+05 |
| PBANKA_1342600 | 147 | 110 | 80 | 8 | 243 | 89 | 316 | 101 | 153 | 84 | 48 | 20 | 315 | 138 | 289 | 1021 | 1575 | 411 | 460 | 406 | 35 | 191 |
| PBANKA_1342700 | 103 | 171 | 9 | 0 | 50 | 19 | 125 | 30 | 181 | 71 | 35 | 16 | 304 | 155 | 69 | 315 | 3316 | 677 | 374 | 316 | 24 | 242 |
| PBANKA_1342800 | 0 | 5 | 26 | 11 | 67 | 29 | 80 | 25 | 147 | 116 | 419 | 267 | 523 | 370 | 128 | 420 | 375 | 111 | 299 | 283 | 42 | 236 |
| PBANKA_1342900 | 0 | 0 | 24 | 9 | 56 | 21 | 84 | 25 | 28 | 17 | 27 | 19 | 157 | 106 | 54 | 343 | 445 | 60 | 391 | 465 | 39 | 230 |
| PBANKA_1343000 | 0 | 0 | 306 | 96 | 362 | 245 | 288 | 112 | 153 | 126 | 2624 | 2280 | 2379 | 1495 | 938 | 4311 | 430 | 73 | 1570 | 1078 | 60 | 499 |
| PBANKA_1343200 | 29 | 48 | 87 | 10 | 466 | 159 | 1633 | 588 | 1233 | 562 | 147 | 51 | 1711 | 1216 | 568 | 3081 | 13067 | 1755 | 9931 | 18566 | 3644 | 30398 |
| PBANKA_1343300 | 6 | 14 | 487 | 174 | 1397 | 951 | 1961 | 997 | 627 | 365 | 89 | 29 | 2455 | 424 | 2270 | 10829 | 5646 | 583 | 1334 | 851 | 106 | 938 |
| PBANKA_1343400 | 291 | 582 | 215 | 64 | 462 | 202 | 1185 | 424 | 1689 | 882 | 258 | 118 | 435 | 502 | 248 | 809 | 8056 | 1215 | 1924 | 4621 | 5692 | 43321 |
| PBANKA_1343500 | 20 | 28 | 42 | 12 | 328 | 105 | 422 | 159 | 611 | 377 | 116 | 55 | 873 | 366 | 9 | 53 | 7762 | 1714 | 1926 | 962 | 284 | 2028 |
| PBANKA_1343600 | 74 | 123 | 72 | 10 | 133 | 35 | 111 | 40 | 45 | 15 | 9 | 5 | 413 | 172 | 211 | 1251 | 230 | 37 | 1014 | 817 | 244 | 1394 |
| PBANKA_1343700 | 6 | 0 | 7 | 0 | 84 | 38 | 85 | 53 | 26 | 17 | 7 | 0 | 317 | 93 | 17 | 151 | 256 | 85 | 2695 | 1390 | 260 | 1643 |
| PBANKA_1343750 | 48 | 47 | 137 | 23 | 342 | 117 | 552 | 191 | 404 | 123 | 399 | 79 | 207 | 112 | 297 | 307 | 287 | 87 | 1072 | 224 | 31 | 233 |
| PBANKA_1343800 | 39 | 39 | 30 | 11 | 164 | 64 | 177 | 84 | 120 | 72 | 49 | 33 | 389 | 267 | 190 | 431 | 355 | 110 | 1322 | 549 | 1164 | 8356 |
| PBANKA_1343900 | 10 | 6 | 32 | 8 | 92 | 20 | 90 | 29 | 43 | 16 | 11 | 19 | 536 | 526 | 306 | 1798 | 427 | 117 | 736 | 647 | 257 | 1598 |
| PBANKA_1344000 | 19 | 35 | 170 | 42 | 333 | 162 | 416 | 142 | 292 | 136 | 201 | 111 | 1588 | 1974 | 1141 | 5629 | 1572 | 258 | 878 | 1511 | 517 | 3771 |
| PBANKA_1344100 | 562 | 852 | 60 | 12 | 550 | 262 | 921 | 322 | 437 | 193 | 69 | 19 | 560 | 360 | 595 | 2351 | 6264 | 792 | 1818 | 4464 | 638 | 5066 |
| PBANKA_1344300 | 38 | 58 | 104 | 42 | 444 | 237 | 1415 | 495 | 1545 | 728 | 224 | 74 | 1285 | 241 | 471 | 1325 | 4119 | 645 | 3422 | 1543 | 247 | 1720 |
| PBANKA_1344400 | 71 | 120 | 23 | 0 | 203 | 155 | 993 | 454 | 1954 | 1232 | 301 | 142 | 614 | 188 | 6 | 14 | 5490 | 1430 | 772 | 404 | 599 | 4547 |
| PBANKA_1344500 | 8 | 5 | 299 | 66 | 1875 | 804 | 2663 | 1022 | 2229 | 1047 | 23916 | 11934 | 6109 | 10416 | 2231 | 11588 | 7280 | 1099 | 1256 | 2583 | 278 | 2542 |
| PBANKA_1344600 | 202 | 257 | 173 | 29 | 391 | 139 | 547 | 172 | 378 | 146 | 442 | 193 | 1749 | 724 | 378 | 2445 | 3743 | 643 | 2334 | 1504 | 290 | 2442 |
| PBANKA_1344800 | 233 | 192 | 44 | 28 | 115 | 44 | 109 | 21 | 137 | 65 | 77 | 40 | 784 | 386 | 201 | 1296 | 387 | 103 | 756 | 413 | 98 | 424 |
| PBANKA_1344900 | 41 | 49 | 52 | 6 | 86 | 31 | 146 | 57 | 122 | 49 | 102 | 41 | 964 | 568 | 112 | 500 | 716 | 135 | 738 | 494 | 378 | 2086 |
| PBANKA_1345000 | 16 | 17 | 70 | 14 | 133 | 64 | 195 | 75 | 101 | 37 | 13 | 7 | 60 | 13 | 199 | 323 | 142 | 42 | 197 | 98 | 412 | 1539 |
| PBANKA_1345100 | 33 | 59 | 91 | 0 | 117 | 39 | 160 | 55 | 116 | 41 | 19 | 9 | 396 | 86 | 203 | 954 | 686 | 103 | 351 | 210 | 181 | 1139 |
| PBANKA_1345200 | 13 | 17 | 25 | 0 | 123 | 45 | 539 | 249 | 317 | 152 | 48 | 19 | 291 | 52 | 126 | 600 | 417 | 87 | 220 | 133 | 98 | 540 |
| PBANKA_1345300 | 57 | 72 | 80 | 22 | 198 | 92 | 506 | 254 | 302 | 193 | 40 | 25 | 1162 | 283 | 326 | 1223 | 1507 | 386 | 1464 | 605 | 277 | 2079 |
| PBANKA_1345400 | 52 | 44 | 63 | 24 | 214 | 151 | 648 | 280 | 525 | 275 | 57 | 24 | 1722 | 219 | 595 | 1772 | 4933 | 1450 | 1125 | 577 | 916 | 6333 |
| PBANKA_1345500 | 119 | 183 | 93 | 19 | 340 | 151 | 396 | 163 | 227 | 123 | 43 | 13 | 625 | 233 | 433 | 2079 | 2132 | 373 | 5864 | 8679 | 1821 | 14292 |
| PBANKA_1345600 | 0 | 10 | 204 | 56 | 445 | 186 | 349 | 163 | 108 | 57 | 31 | 8 | 593 | 140 | 798 | 3817 | 882 | 133 | 1881 | 969 | 537 | 4742 |
| PBANKA_1345700 | 0 | 0 | 34 | 8 | 125 | 76 | 176 | 72 | 63 | 28 | 14 | 14 | 245 | 74 | 258 | 692 | 640 | 132 | 1699 | 670 | 177 | 1070 |
| PBANKA_1345800 | 82 | 92 | 527 | 146 | 202 | 107 | 147 | 53 | 249 | 147 | 2522 | 1615 | 6874 | 5363 | 1027 | 6359 | 1160 | 260 | 1580 | 1604 | 1800 | 12234 |
| PBANKA_1345900 | 57 | 108 | 7 | 0 | 13 | 8 | 41 | 6 | 332 | 159 | 86 | 42 | 245 | 156 | 33 | 82 | 1978 | 540 | 740 | 592 | 1286 | 7788 |
| PBANKA_1346000 | 0 | 0 | 31 | 0 | 151 | 85 | 128 | 53 | 87 | 29 | 28 | 7 | 412 | 200 | 352 | 1842 | 632 | 75 | 636 | 561 | 210 | 1484 |
| PBANKA_1346100 | 7 | 10 | 82 | 17 | 218 | 80 | 154 | 57 | 85 | 35 | 523 | 298 | 1901 | 940 | 517 | 2383 | 524 | 108 | 691 | 537 | 77 | 704 |
| PBANKA_1346200 | 9 | 8 | 86 | 19 | 242 | 171 | 344 | 166 | 101 | 70 | 12 | 9 | 187 | 58 | 181 | 700 | 479 | 98 | 750 | 311 | 155 | 1088 |
| PBANKA_1346300 | 114 | 220 | 64 | 11 | 279 | 140 | 286 | 105 | 120 | 55 | 20 | 9 | 656 | 241 | 60 | 417 | 457 | 133 | 18562 | 15995 | 172 | 1401 |
| PBANKA_1346400 | 19 | 38 | 30 | 0 | 91 | 39 | 184 | 56 | 494 | 295 | 176 | 77 | 713 | 318 | 88 | 252 | 2555 | 649 | 3041 | 1938 | 232 | 1208 |
| PBANKA_1346500 | 29 | 26 | 52 | 17 | 193 | 129 | 370 | 179 | 185 | 114 | 16 | 15 | 134 | 31 | 177 | 333 | 576 | 184 | 518 | 199 | 139 | 960 |
| PBANKA_1346600 | 10 | 0 | 305 | 79 | 729 | 306 | 811 | 264 | 252 | 127 | 75 | 53 | 2291 | 1123 | 1391 | 6376 | 1953 | 291 | 1341 | 1796 | 557 | 4644 |
| PBANKA_1346700 | 47 | 36 | 1093 | 459 | 743 | 520 | 890 | 532 | 305 | 302 | 1256 | 1129 | 9313 | 1985 | 2185 | 5107 | 1656 | 506 | 2382 | 625 | 529 | 4052 |
| PBANKA_1346800 | 261 | 377 | 103 | 37 | 212 | 149 | 267 | 131 | 102 | 47 | 11 | 17 | 2251 | 634 | 405 | 1674 | 1259 | 229 | 1447 | 885 | 434 | 3154 |
| PBANKA_1346900 | 28 | 41 | 40 | 11 | 147 | 28 | 145 | 34 | 72 | 29 | 9 | 11 | 84 | 26 | 33 | 101 | 73 | 15 | 1153 | 504 | 162 | 1112 |
| PBANKA_1347000 | 585 | 1034 | 6 | 0 | 39 | 30 | 149 | 38 | 2186 | 1352 | 724 | 401 | 1665 | 1075 | 59 | 197 | 8413 | 1965 | 10921 | 14330 | 4655 | 32385 |
| PBANKA_1347100 | 0 | 0 | 41 | 11 | 63 | 22 | 38 | 20 | 16 | 6 | 6 | 0 | 176 | 63 | 95 | 338 | 102 | 21 | 23 | 16 | 5 | 19 |
| PBANKA_1347200 | 93 | 124 | 97 | 31 | 238 | 102 | 260 | 129 | 161 | 63 | 121 | 65 | 871 | 625 | 297 | 1257 | 1631 | 292 | 1038 | 1060 | 5220 | 35575 |
| PBANKA_1347300 | 9 | 7 | 63 | 17 | 128 | 47 | 129 | 37 | 69 | 27 | 19 | 7 | 484 | 212 | 186 | 778 | 603 | 114 | 342 | 197 | 518 | 3189 |
| PBANKA_1347400 | 65 | 125 | 117 | 23 | 292 | 121 | 387 | 169 | 185 | 90 | 51 | 19 | 1354 | 443 | 490 | 2597 | 2857 | 311 | 739 | 683 | 537 | 4221 |
| PBANKA_1347500 | 9 | 16 | 30 | 9 | 58 | 20 | 64 | 32 | 43 | 22 | 108 | 68 | 563 | 253 | 246 | 1072 | 362 | 108 | 533 | 440 | 635 | 4140 |
| PBANKA_1347600 | 0 | 0 | 162 | 39 | 224 | 113 | 228 | 75 | 82 | 36 | 312 | 220 | 1986 | 617 | 385 | 2205 | 414 | 75 | 369 | 221 | 316 | 2383 |
| PBANKA_1347700 | 0 | 0 | 220 | 62 | 326 | 177 | 269 | 90 | 112 | 82 | 375 | 256 | 3048 | 2002 | 1176 | 5930 | 1545 | 229 | 1938 | 1908 | 852 | 5557 |
| PBANKA_1347800 | 0 | 0 | 189 | 68 | 212 | 135 | 234 | 136 | 61 | 50 | 131 | 142 | 1233 | 390 | 414 | 1483 | 507 | 96 | 1591 | 950 | 827 | 5215 |
| PBANKA_1347900 | 0 | 0 | 160 | 39 | 322 | 127 | 534 | 257 | 269 | 148 | 34 | 12 | 1037 | 546 | 908 | 3485 | 8159 | 1213 | 2292 | 995 | 711 | 4871 |
| PBANKA_1348000 | 0 | 0 | 87 | 28 | 189 | 104 | 195 | 101 | 61 | 41 | 6 | 0 | 651 | 145 | 240 | 678 | 372 | 90 | 338 | 180 | 66 | 474 |
| PBANKA_1348100 | 0 | 0 | 131 | 30 | 240 | 123 | 225 | 94 | 89 | 39 | 11 | 7 | 847 | 164 | 265 | 836 | 749 | 119 | 475 | 326 | 101 | 743 |
| PBANKA_1348200 | 33 | 39 | 92 | 15 | 132 | 41 | 101 | 27 | 69 | 19 | 22 | 5 | 1271 | 634 | 406 | 2421 | 417 | 46 | 1153 | 1378 | 991 | 5356 |
| PBANKA_1348300 | 177 | 253 | 283 | 80 | 461 | 267 | 471 | 159 | 309 | 149 | 1746 | 1142 | 1648 | 1554 | 587 | 2658 | 681 | 144 | 732 | 743 | 1132 | 7097 |
| PBANKA_1348400 | 24 | 50 | 98 | 38 | 188 | 111 | 240 | 125 | 163 | 89 | 36 | 20 | 397 | 120 | 132 | 449 | 1034 | 191 | 1279 | 491 | 204 | 2145 |
| PBANKA_1348500 | 6 | 16 | 166 | 25 | 171 | 66 | 150 | 68 | 65 | 33 | 25 | 8 | 786 | 335 | 365 | 1933 | 194 | 47 | 694 | 352 | 669 | 4415 |
| PBANKA_1348600 | 13 | 11 | 81 | 14 | 333 | 76 | 560 | 158 | 387 | 186 | 81 | 25 | 844 | 391 | 615 | 3086 | 4964 | 721 | 1163 | 1256 | 525 | 3266 |
| PBANKA_1348700 | 0 | 0 | 19 | 9 | 73 | 35 | 133 | 53 | 95 | 47 | 17 | 0 | 470 | 193 | 107 | 301 | 2338 | 504 | 1800 | 1987 | 430 | 2766 |
| PBANKA_1348800 | 9 | 11 | 80 | 7 | 217 | 76 | 341 | 110 | 183 | 58 | 38 | 17 | 860 | 393 | 191 | 967 | 2603 | 413 | 3535 | 3123 | 95 | 768 |
| PBANKA_1348900 | 0 | 0 | 375 | 60 | 329 | 149 | 307 | 98 | 119 | 40 | 78 | 47 | 6190 | 3328 | 2128 | 12507 | 2593 | 376 | 1866 | 2435 | 59 | 425 |

| | | | | | | | | | | | | | | | | | | | | | | |
|---|---|---|---|---|---|---|---|---|---|---|---|---|---|---|---|---|---|---|---|---|---|---|
| PBANKA_1349200 | 9 | 8 | 127 | 35 | 135 | 86 | 245 | 128 | 215 | 146 | 2354 | 1943 | 10722 | 5771 | 2386 | 9158 | 11256 | 1567 | 5598 | 3627 | 133 | 1002 |
| PBANKA_1349300 | 13175 | 19013 | 2001 | 561 | 384 | 212 | 229 | 98 | 75 | 46 | 25 | 20 | 288 | 69 | 25 | 113 | 128 | 37 | 3992 | 1999 | 73 | 691 |
| PBANKA_1349400 | 22 | 24 | 14 | 7 | 14 | 12 | 24 | 0 | 0 | 5 | 12 | 0 | 208 | 45 | 7 | 24 | 115 | 40 | 2515 | 798 | 7 | 63 |
| PBANKA_1349500 | 137 | 167 | 59 | 16 | 118 | 53 | 123 | 59 | 106 | 47 | 53 | 38 | 740 | 340 | 183 | 709 | 587 | 173 | 610 | 231 | 162 | 1086 |
| PBANKA_1349600 | 18 | 18 | 85 | 15 | 270 | 140 | 323 | 133 | 127 | 51 | 23 | 13 | 685 | 610 | 661 | 3949 | 2031 | 220 | 475 | 808 | 57 | 406 |
| PBANKA_1349700 | 0 | 0 | 82 | 27 | 140 | 89 | 248 | 112 | 97 | 57 | 17 | 12 | 1008 | 458 | 535 | 2683 | 1949 | 303 | 807 | 900 | 113 | 708 |
| PBANKA_1349800 | 25670 | 35972 | 54 | 25 | 305 | 214 | 409 | 167 | 194 | 104 | 37 | 35 | 303 | 80 | 96 | 388 | 756 | 137 | 393 | 248 | 312 | 2424 |
| PBANKA_1349900 | 128 | 159 | 113 | 26 | 317 | 155 | 551 | 244 | 233 | 101 | 39 | 31 | 2886 | 1075 | 522 | 2302 | 2726 | 455 | 811 | 424 | 218 | 1156 |
| PBANKA_1350000 | 0 | 5 | 41 | 16 | 165 | 112 | 186 | 83 | 49 | 40 | 0 | 0 | 248 | 49 | 157 | 924 | 542 | 83 | 656 | 496 | 66 | 419 |
| PBANKA_1350100 | 0 | 0 | 60 | 9 | 125 | 63 | 150 | 54 | 82 | 39 | 29 | 17 | 756 | 481 | 322 | 2237 | 1404 | 216 | 340 | 283 | 70 | 569 |
| PBANKA_1350200 | 6 | 0 | 13 | 0 | 26 | 8 | 43 | 11 | 21 | 7 | 5 | 0 | 554 | 234 | 157 | 753 | 499 | 126 | 3104 | 1371 | 29 | 131 |
| PBANKA_1350300 | 409 | 569 | 209 | 45 | 418 | 198 | 357 | 218 | 176 | 87 | 31 | 10 | 541 | 112 | 365 | 1072 | 717 | 201 | 1361 | 772 | 194 | 1136 |
| PBANKA_1350400 | 22 | 42 | 35 | 12 | 96 | 43 | 88 | 67 | 52 | 21 | 5 | 0 | 36 | 16 | 57 | 139 | 140 | 24 | 250 | 98 | 8 | 88 |
| PBANKA_1350500 | 241 | 223 | 39 | 0 | 44 | 36 | 77 | 39 | 66 | 61 | 327 | 302 | 1083 | 600 | 349 | 858 | 1178 | 250 | 1576 | 736 | 48 | 520 |
| PBANKA_1350600 | 211 | 225 | 221 | 43 | 253 | 101 | 224 | 90 | 73 | 32 | 77 | 55 | 4401 | 2957 | 1359 | 9325 | 1495 | 191 | 2201 | 2042 | 421 | 2285 |
| PBANKA_1350700 | 33 | 50 | 136 | 49 | 395 | 203 | 420 | 155 | 100 | 54 | 24 | 8 | 2192 | 984 | 613 | 2474 | 1425 | 253 | 592 | 411 | 82 | 689 |
| PBANKA_1350800 | 0 | 7 | 52 | 10 | 188 | 87 | 193 | 57 | 57 | 26 | 16 | 7 | 508 | 135 | 457 | 1806 | 695 | 91 | 260 | 165 | 352 | 2160 |
| PBANKA_1350900 | 0 | 11 | 33 | 8 | 95 | 58 | 117 | 74 | 38 | 25 | 0 | 0 | 316 | 41 | 168 | 703 | 364 | 68 | 136 | 75 | 64 | 361 |
| PBANKA_1351000 | 128 | 155 | 147 | 32 | 354 | 136 | 337 | 121 | 277 | 164 | 107 | 51 | 1637 | 876 | 862 | 3517 | 1559 | 322 | 1023 | 1140 | 765 | 4512 |
| PBANKA_1351100 | 5 | 0 | 115 | 18 | 227 | 80 | 275 | 90 | 137 | 51 | 35 | 10 | 1660 | 733 | 322 | 1621 | 2006 | 270 | 1853 | 1755 | 351 | 2331 |
| PBANKA_1351200 | 116 | 145 | 91 | 10 | 239 | 78 | 243 | 78 | 84 | 44 | 10 | 8 | 2328 | 1199 | 790 | 3706 | 1483 | 229 | 665 | 494 | 95 | 550 |
| PBANKA_1351300 | 71 | 108 | 314 | 90 | 613 | 256 | 1277 | 504 | 1139 | 579 | 224 | 80 | 1469 | 755 | 938 | 4748 | 11821 | 1441 | 6690 | 9435 | 586 | 4335 |
| PBANKA_1351400 | 8 | 12 | 14 | 0 | 69 | 44 | 160 | 55 | 761 | 397 | 124 | 59 | 279 | 191 | 8 | 41 | 2857 | 813 | 771 | 1168 | 185 | 1089 |
| PBANKA_1351600 | 0 | 0 | 31 | 12 | 59 | 40 | 88 | 14 | 28 | 10 | 13 | 5 | 127 | 60 | 113 | 496 | 244 | 47 | 71 | 41 | 68 | 503 |
| PBANKA_1351700 | 60 | 84 | 82 | 29 | 332 | 172 | 554 | 361 | 365 | 185 | 46 | 40 | 1101 | 200 | 429 | 1331 | 3043 | 792 | 4280 | 2087 | 906 | 4078 |
| PBANKA_1351800 | 49 | 107 | 231 | 66 | 557 | 323 | 746 | 369 | 247 | 180 | 48 | 18 | 1458 | 231 | 1276 | 5035 | 3358 | 483 | 1061 | 816 | 113 | 864 |
| PBANKA_1351900 | 17 | 18 | 779 | 178 | 489 | 192 | 484 | 140 | 170 | 90 | 604 | 335 | 10874 | 4010 | 2191 | 7662 | 2092 | 319 | 1967 | 1004 | 307 | 2371 |
| PBANKA_1352000 | 6 | 5 | 1907 | 466 | 1266 | 713 | 1293 | 516 | 320 | 164 | 187 | 139 | 8454 | 3469 | 4321 | 13714 | 3647 | 725 | 2306 | 1299 | 182 | 1729 |
| PBANKA_1352100 | 48 | 89 | 31 | 0 | 81 | 30 | 70 | 25 | 47 | 16 | 12 | 0 | 1044 | 364 | 505 | 1604 | 613 | 105 | 14129 | 7034 | 378 | 2178 |
| PBANKA_1352200 | 19 | 42 | 68 | 23 | 136 | 40 | 129 | 39 | 58 | 32 | 15 | 0 | 399 | 77 | 311 | 1041 | 336 | 88 | 883 | 407 | 344 | 2085 |
| PBANKA_1352300 | 0 | 16 | 229 | 33 | 432 | 239 | 635 | 223 | 225 | 98 | 44 | 13 | 802 | 251 | 860 | 4923 | 3212 | 408 | 1528 | 1855 | 510 | 3380 |
| PBANKA_1352400 | 10 | 6 | 38 | 9 | 200 | 103 | 386 | 166 | 219 | 105 | 24 | 14 | 940 | 329 | 113 | 443 | 6466 | 897 | 2095 | 1932 | 184 | 1680 |
| PBANKA_1352500 | 246 | 266 | 20 | 0 | 98 | 60 | 163 | 57 | 56 | 16 | 14 | 5 | 1048 | 456 | 37 | 73 | 505 | 319 | 24130 | 15034 | 2282 | 13974 |
| PBANKA_1352600 | 22 | 17 | 65 | 17 | 303 | 131 | 757 | 308 | 317 | 223 | 36 | 25 | 458 | 147 | 612 | 2402 | 1311 | 223 | 3613 | 1890 | 1572 | 10221 |
| PBANKA_1352700 | 135 | 186 | 121 | 34 | 309 | 153 | 301 | 85 | 142 | 85 | 793 | 582 | 3814 | 2433 | 619 | 3982 | 1411 | 194 | 1653 | 978 | 750 | 5151 |
| PBANKA_1352800 | 74 | 69 | 55 | 16 | 230 | 100 | 370 | 196 | 193 | 183 | 12 | 20 | 593 | 155 | 297 | 1430 | 2734 | 495 | 1103 | 809 | 472 | 2544 |
| PBANKA_1352900 | 19 | 22 | 29 | 11 | 78 | 40 | 74 | 46 | 11 | 16 | 7 | 7 | 168 | 31 | 154 | 397 | 120 | 38 | 326 | 126 | 23 | 187 |
| PBANKA_1353000 | 115 | 125 | 63 | 12 | 155 | 69 | 158 | 64 | 66 | 30 | 14 | 7 | 391 | 103 | 219 | 993 | 349 | 81 | 681 | 356 | 104 | 570 |
| PBANKA_1353100 | 18 | 23 | 90 | 18 | 322 | 116 | 1248 | 501 | 1226 | 521 | 215 | 111 | 2094 | 2554 | 317 | 2279 | 27181 | 3581 | 2203 | 18272 | 9579 | 58942 |
| PBANKA_1353200 | 5 | 13 | 21 | 5 | 45 | 11 | 42 | 19 | 17 | 11 | 0 | 0 | 141 | 12 | 61 | 276 | 226 | 37 | 378 | 206 | 26 | 295 |
| PBANKA_1353300 | 11 | 19 | 68 | 16 | 237 | 123 | 288 | 111 | 117 | 62 | 13 | 0 | 406 | 85 | 264 | 1297 | 1131 | 157 | 1785 | 1012 | 91 | 767 |
| PBANKA_1353400 | 12 | 28 | 16 | 6 | 58 | 29 | 63 | 28 | 47 | 33 | 14 | 6 | 119 | 88 | 78 | 461 | 203 | 47 | 362 | 1199 | 13182 | 1E+05 |
| PBANKA_1353500 | 0 | 9 | 13 | 10 | 79 | 39 | 89 | 29 | 36 | 9 | 20 | 12 | 313 | 122 | 175 | 705 | 288 | 40 | 129 | 121 | 609 | 4216 |
| PBANKA_1353700 | 0 | 8 | 231 | 38 | 327 | 91 | 205 | 81 | 185 | 85 | 1573 | 939 | 6261 | 5550 | 884 | 5387 | 1073 | 176 | 893 | 2093 | 103 | 685 |
| PBANKA_1353800 | 87 | 151 | 9 | 0 | 14 | 15 | 34 | 15 | 30 | 25 | 0 | 0 | 1911 | 978 | 40 | 177 | 545 | 368 | 56496 | 35291 | 6247 | 62151 |
| PBANKA_1354000 | 0 | 0 | 120 | 32 | 182 | 104 | 197 | 66 | 347 | 195 | 252 | 132 | 1178 | 464 | 312 | 1427 | 599 | 251 | 266 | 141 | 52 | 350 |
| PBANKA_1354100 | 0 | 0 | 59 | 7 | 177 | 70 | 351 | 197 | 267 | 127 | 24 | 13 | 926 | 223 | 128 | 477 | 3683 | 883 | 2555 | 1871 | 325 | 2111 |
| PBANKA_1354200 | 0 | 0 | 5 | 0 | 54 | 23 | 63 | 27 | 23 | 21 | 0 | 0 | 49 | 19 | 16 | 20 | 142 | 29 | 658 | 216 | 34 | 262 |
| PBANKA_1354300 | 373 | 531 | 184 | 69 | 442 | 322 | 518 | 179 | 419 | 319 | 1504 | 1237 | 10358 | 4200 | 2058 | 8266 | 2308 | 512 | 7634 | 3821 | 1103 | 10384 |
| PBANKA_1354380 | 25 | 13 | 26 | 24 | 47 | 25 | 61 | 31 | 62 | 26 | 282 | 187 | 174 | 82 | 1110 | 109 | 114 | 139 | 5194 | 90 | 0 | 7 |
| PBANKA_1354400 | 8 | 15 | 3743 | 955 | 2129 | 1221 | 2498 | 1212 | 651 | 443 | 3161 | 2691 | 19191 | 8711 | 3801 | 10524 | 3451 | 623 | 3997 | 1892 | 653 | 4680 |
| PBANKA_1354500 | 18 | 20 | 1910 | 667 | 1591 | 1097 | 1633 | 998 | 547 | 414 | 4548 | 4088 | 7549 | 4495 | 2276 | 6431 | 1951 | 439 | 1822 | 875 | 271 | 2386 |
| PBANKA_1354700 | 5 | 12 | 26 | 8 | 237 | 89 | 341 | 139 | 161 | 52 | 24 | 7 | 326 | 62 | 285 | 1327 | 1514 | 199 | 983 | 710 | 63 | 414 |
| PBANKA_1355000 | 199 | 230 | 50 | 31 | 181 | 113 | 344 | 148 | 150 | 90 | 32 | 31 | 1096 | 380 | 402 | 1408 | 1144 | 207 | 811 | 346 | 57 | 482 |
| PBANKA_1355100 | 35 | 33 | 2953 | 744 | 1673 | 962 | 1598 | 793 | 503 | 319 | 3343 | 2681 | 34723 | 14716 | 5780 | 25182 | 6278 | 923 | 5917 | 3864 | 207 | 1666 |
| PBANKA_1355200 | 642 | 1034 | 195 | 59 | 1103 | 447 | 1422 | 502 | 781 | 327 | 110 | 58 | 1275 | 506 | 879 | 3404 | 4474 | 763 | 2763 | 2253 | 245 | 2334 |
| PBANKA_1355300 | 126 | 139 | 37 | 7 | 152 | 59 | 141 | 60 | 85 | 40 | 15 | 0 | 244 | 77 | 149 | 660 | 366 | 66 | 959 | 476 | 95 | 549 |
| PBANKA_1355400 | 5 | 6 | 25 | 15 | 105 | 43 | 73 | 58 | 30 | 5 | 20 | 0 | 162 | 14 | 91 | 398 | 173 | 19 | 291 | 139 | 54 | 447 |
| PBANKA_1355500 | 71 | 70 | 235 | 89 | 480 | 301 | 493 | 221 | 139 | 81 | 49 | 25 | 2309 | 1213 | 1263 | 6169 | 1264 | 180 | 2057 | 1731 | 291 | 2666 |
| PBANKA_1355600 | 2761 | 3616 | 22 | 0 | 29 | 14 | 27 | 11 | 13 | 5 | 0 | 0 | 28 | 8 | 21 | 52 | 40 | 17 | 274 | 134 | 36 | 235 |
| PBANKA_1355700 | 2256 | 3032 | 79 | 20 | 189 | 143 | 280 | 116 | 3948 | 2826 | 992 | 641 | 3489 | 2105 | 165 | 835 | 6971 | 3759 | 44028 | 21150 | 16746 | 1E+05 |
| PBANKA_1355800 | 38 | 23 | 54 | 12 | 69 | 43 | 57 | 60 | 53 | 20 | 75 | 41 | 130 | 74 | 98 | 185 | 58 | 17 | 60 | 46 | 14 | 142 |
| PBANKA_1355900 | 35 | 35 | 282 | 85 | 706 | 569 | 753 | 281 | 194 | 102 | 43 | 32 | 2723 | 941 | 978 | 4220 | 1400 | 387 | 45988 | 34718 | 1336 | 13862 |
| PBANKA_1356000 | 640 | 894 | 293 | 72 | 956 | 402 | 1239 | 420 | 761 | 353 | 2006 | 1156 | 970 | 3427 | 1191 | 5248 | 2369 | 312 | 299 | 1864 | 1354 | 8817 |
| PBANKA_1356100 | 11 | 13 | 66 | 11 | 110 | 58 | 263 | 107 | 449 | 248 | 442 | 272 | 1050 | 731 | 43 | 368 | 5539 | 1247 | 2961 | 2946 | 76 | 671 |
| PBANKA_1356200 | 49 | 29 | 136 | 24 | 279 | 120 | 322 | 115 | 167 | 84 | 658 | 442 | 1095 | 432 | 156 | 783 | 519 | 133 | 603 | 272 | 9 | 83 |
| PBANKA_1356300 | 326 | 577 | 175 | 53 | 561 | 317 | 961 | 410 | 467 | 215 | 56 | 40 | 1244 | 424 | 522 | 1965 | 7215 | 1111 | 3862 | 3049 | 105 | 906 |
| PBANKA_1356400 | 10 | 19 | 118 | 25 | 254 | 139 | 346 | 141 | 165 | 86 | 38 | 10 | 713 | 221 | 344 | 1673 | 1819 | 234 | 5059 | 5006 | 57 | 448 |
| PBANKA_1356500 | 251 | 334 | 138 | 25 | 229 | 121 | 291 | 165 | 144 | 70 | 12 | 5 | 511 | 114 | 367 | 1484 | 1967 | 408 | 757 | 556 | 14 | 105 |
| PBANKA_1356700 | 0 | 0 | 127 | 16 | 423 | 191 | 602 | 248 | 325 | 214 | 2399 | 2055 | 3795 | 2711 | 915 | 4310 | 4573 | 708 | 1032 | 1480 | 24 | 268 |
| PBANKA_1356800 | 224 | 367 | 79 | 9 | 738 | 282 | 2819 | 1191 | 4604 | 1991 | 618 | 226 | 1249 | 2690 | 1213 | 4814 | 58810 | 10785 | 960 | 11960 | 458 | 3629 |
| PBANKA_1356900 | 9 | 0 | 75 | 9 | 210 | 97 | 238 | 102 | 142 | 44 | 55 | 33 | 863 | 316 | 310 | 1664 | 1610 | 230 | 867 | 511 | 21 | 154 |
| PBANKA_1357000 | 7 | 14 | 72 | 12 | 141 | 82 | 118 | 56 | 49 | 22 | 14 | 5 | 464 | 165 | 369 | 2287 | 829 | 64 | 89 | 99 | 0 | 11 |
| PBANKA_1357100 | 11 | 29 | 13 | 0 | 23 | 16 | 23 | 5 | 30 | 20 | 7 | 0 | 620 | 110 | 87 | 389 | 314 | 87 | 657 | 483 | 15 | 101 |
| PBANKA_1357200 | 5 | 5 | 335 | 121 | 1015 | 565 | 1715 | 684 | 564 | 295 | 432 | 384 | 4520 | 6243 | 3599 | 22150 | 13550 | 1392 | 901 | 3421 | 296 | 2671 |
| PBANKA_1357300 | 0 | 0 | 11 | 0 | 129 | 106 | 422 | 212 | 965 | 658 | 112 | 123 | 454 | 290 | 85 | 324 | 4756 | 623 | 1210 | 1250 | 232 | 2037 |
| PBANKA_1357400 | 0 | 0 | 57 | 12 | 43 | 31 | 66 | 30 | 18 | 11 | 0 | 9 | 170 | 83 | 154 | 354 | 66 | 24 | 63 | 29 | 0 | 5 |
| PBANKA_1357500 | 0 | 0 | 221 | 44 | 520 | 323 | 459 | 177 | 110 | 52 | 39 | 15 | 467 | 199 | 554 | 2780 | 938 | 119 | 187 | 190 | 0 | 11 |
| PBANKA_1357600 | 46 | 54 | 128 | 28 | 440 | 158 | 531 | 201 | 221 | 86 | 33 | 17 | 676 | 311 | 520 | 2387 | 1858 | 226 | 566 | 450 | 80 | 606 |
| PBANKA_1357700 | 85 | 150 | 147 | 53 | 171 | 115 | 296 | 207 | 105 | 78 | 8 | 7 | 558 | 131 | 231 | 836 | 1882 | 272 | 4072 | 2484 | 94 | 763 |
| PBANKA_1357800 | 88 | 142 | 150 | 23 | 308 | 127 | 301 | 87 | 176 | 88 | 650 | 365 | 4479 | 4414 | 934 | 5962 | 1858 | 278 | 5378 | 6302 | 155 | 1506 |
| PBANKA_1357900 | 0 | 0 | 20 | 0 | 70 | 31 | 244 | 89 | 572 | 381 | 124 | 77 | 398 | 85 | 92 | 152 | 177 | 128 | 3101 | 2150 | 111 | 914 |
| PBANKA_1358000 | 0 | 0 | 99 | 26 | 202 | 104 | 168 | 69 | 232 | 131 | 387 | 280 | 1044 | 582 | 302 | 1061 | 604 | 280 | 995 | 428 | 56 | 409 |
| PBANKA_1358100 | 6 | 11 | 152 | 34 | 322 | 161 | 311 | 110 | 100 | 45 | 28 | 9 | 985 | 349 | 577 | 2023 | 396 | 57 | 244 | 149 | 33 | 201 |
| PBANKA_1358200 | 18 | 22 | 27 | 6 | 89 | 36 | 195 | 108 | 196 | 139 | 28 | 16 | 249 | 61 | 133 | 343 | 987 | 398 | 799 | 233 | 41 | 330 |
| PBANKA_1358300 | 147 | 197 | 226 | 55 | 904 | 351 | 1493 | 642 | 877 | 414 | 144 | 54 | 2353 | 1541 | 1332 | 7446 | 20761 | 2261 | 3089 | 10103 | 269 | 1886 |
| PBANKA_1358400 | 0 | 0 | 78 | 17 | 168 | 102 | 297 | 120 | 117 | 68 | 9 | 0 | 417 | 87 | 180 | 987 | 1252 | 150 | 410 | 149 | 14 | 55 |
| PBANKA_1358600 | 0 | 0 | 168 | 44 | 551 | 336 | 702 | 325 | 172 | 119 | 26 | 22 | 415 | 129 | 488 | 1689 | 1498 | 211 | 1637 | 927 | 191 | 1756 |
| PBANKA_1358700 | 0 | 0 | 183 | 57 | 455 | 229 | 548 | 206 | 206 | 89 | 33 | 0 | 1978 | 627 | 756 | 3600 | 2981 | 419 | 1563 | 1319 | 259 | 1888 |
| PBANKA_1358800 | 15 | 24 | 47 | 6 | 129 | 63 | 214 | 86 | 88 | 53 | 0 | 12 | 263 | 75 | 92 | 404 | 942 | 133 | 744 | 328 | 64 | 387 |

| | | | | | | | | | | | | | | | | | | | | | | |
|---|---|---|---|---|---|---|---|---|---|---|---|---|---|---|---|---|---|---|---|---|---|---|
| PBANKA_1358900 | 0 | 0 | 7 | 0 | 119 | 53 | 149 | 71 | 63 | 29 | 9 | 11 | 347 | 164 | 110 | 338 | 668 | 146 | 2197 | 1638 | 262 | 2309 |
| PBANKA_1359000 | 63 | 74 | 751 | 218 | 1086 | 712 | 1935 | 878 | 779 | 476 | 215 | 104 | 7046 | 2444 | 2404 | 8188 | 18911 | 2944 | 16248 | 9727 | 1038 | 11945 |
| PBANKA_1359100 | 0 | 0 | 39 | 5 | 103 | 41 | 101 | 40 | 26 | 13 | 15 | 17 | 315 | 124 | 135 | 907 | 102 | 24 | 549 | 341 | 25 | 272 |
| PBANKA_1359200 | 55 | 107 | 603 | 199 | 377 | 273 | 390 | 143 | 178 | 93 | 1748 | 1281 | 7569 | 3040 | 1287 | 6276 | 779 | 107 | 2023 | 1197 | 85 | 798 |
| PBANKA_1359300 | 175 | 255 | 880 | 216 | 2279 | 1023 | 2412 | 824 | 770 | 385 | 451 | 226 | 1734 | 3974 | 3841 | 19072 | 4205 | 430 | 350 | 4134 | 236 | 2146 |
| PBANKA_1359400 | 53 | 82 | 164 | 35 | 471 | 258 | 358 | 168 | 126 | 58 | 383 | 310 | 1993 | 1355 | 710 | 3464 | 960 | 264 | 1445 | 1057 | 119 | 978 |
| PBANKA_1359500 | 0 | 0 | 81 | 26 | 188 | 93 | 203 | 79 | 60 | 22 | 16 | 6 | 293 | 94 | 135 | 705 | 469 | 106 | 2170 | 1315 | 201 | 1579 |
| PBANKA_1359600 | 0 | 0 | 15 | 5 | 92 | 42 | 112 | 55 | 37 | 22 | 0 | 0 | 370 | 98 | 49 | 114 | 333 | 112 | 4360 | 3201 | 91 | 1090 |
| PBANKA_1359700 | 0 | 0 | 8 | 0 | 40 | 27 | 43 | 9 | 28 | 6 | 5 | 0 | 164 | 76 | 15 | 13 | 348 | 256 | 3619 | 1815 | 63 | 652 |
| PBANKA_1359800 | 0 | 0 | 10 | 0 | 22 | 10 | 77 | 38 | 38 | 18 | 0 | 0 | 98 | 23 | 19 | 49 | 347 | 43 | 857 | 290 | 30 | 183 |
| PBANKA_1359900 | 0 | 0 | 38 | 0 | 198 | 116 | 429 | 214 | 142 | 87 | 9 | 7 | 114 | 20 | 226 | 726 | 245 | 42 | 257 | 83 | 0 | 17 |
| PBANKA_1360000 | 12 | 14 | 8 | 0 | 195 | 86 | 686 | 324 | 223 | 138 | 38 | 12 | 387 | 124 | 111 | 244 | 1871 | 226 | 2595 | 1493 | 11 | 213 |
| PBANKA_1360100 | 5 | 0 | 81 | 24 | 345 | 185 | 341 | 170 | 215 | 172 | 1533 | 1523 | 6979 | 1780 | 3119 | 10732 | 1315 | 320 | 454 | 327 | 12 | 207 |
| PBANKA_1360200 | 30 | 35 | 28 | 8 | 103 | 34 | 82 | 51 | 37 | 27 | 0 | 6 | 898 | 258 | 161 | 560 | 561 | 133 | 3323 | 1615 | 164 | 1165 |
| PBANKA_1360300 | 0 | 0 | 840 | 248 | 1103 | 753 | 1052 | 470 | 251 | 199 | 123 | 177 | 9740 | 4540 | 3119 | 11635 | 2069 | 373 | 3496 | 2287 | 161 | 1659 |
| PBANKA_1360500 | 21 | 27 | 139 | 39 | 274 | 164 | 357 | 109 | 316 | 203 | 47 | 56 | 5123 | 1671 | 838 | 3576 | 6924 | 1497 | 1461 | 904 | 16 | 167 |
| PBANKA_1360700 | 33 | 51 | 264 | 45 | 462 | 205 | 475 | 162 | 174 | 66 | 28 | 26 | 1337 | 1007 | 1126 | 5008 | 1583 | 226 | 630 | 989 | 41 | 302 |
| PBANKA_1360800 | 0 | 0 | 10 | 0 | 101 | 26 | 109 | 40 | 46 | 28 | 0 | 0 | 94 | 55 | 16 | 94 | 622 | 110 | 307 | 1119 | 34 | 263 |
| PBANKA_1361000 | 63 | 73 | 39 | 6 | 50 | 16 | 53 | 24 | 86 | 51 | 88 | 32 | 627 | 216 | 100 | 318 | 855 | 306 | 1327 | 873 | 402 | 2101 |
| PBANKA_1361100 | 0 | 9 | 129 | 38 | 286 | 165 | 256 | 94 | 123 | 65 | 102 | 67 | 2008 | 1588 | 783 | 3393 | 1545 | 231 | 846 | 1151 | 30 | 332 |
| PBANKA_1361200 | 205 | 290 | 196 | 39 | 868 | 287 | 1004 | 368 | 488 | 221 | 921 | 421 | 1717 | 3428 | 1199 | 4911 | 9098 | 1829 | 15776 | 57451 | 593 | 3666 |
| PBANKA_1361300 | 6 | 10 | 20 | 0 | 84 | 29 | 89 | 43 | 32 | 13 | 13 | 7 | 983 | 299 | 344 | 1695 | 533 | 160 | 4743 | 2791 | 257 | 1812 |
| PBANKA_1361500 | 0 | 6 | 21 | 0 | 66 | 40 | 62 | 16 | 33 | 6 | 0 | 0 | 246 | 67 | 67 | 230 | 286 | 71 | 2264 | 1163 | 37 | 182 |
| PBANKA_1361600 | 0 | 8 | 5 | 0 | 20 | 5 | 11 | 7 | 33 | 17 | 10 | 5 | 611 | 258 | 37 | 81 | 703 | 183 | 4716 | 6988 | 81 | 404 |
| PBANKA_1361700 | 0 | 0 | 16 | 0 | 40 | 8 | 27 | 14 | 10 | 0 | 0 | 0 | 143 | 81 | 78 | 323 | 197 | 34 | 104 | 58 | 19 | 143 |
| PBANKA_1361800 | 0 | 0 | 13 | 0 | 100 | 68 | 133 | 69 | 42 | 29 | 0 | 0 | 107 | 39 | 18 | 16 | 122 | 73 | 1725 | 868 | 98 | 611 |
| PBANKA_1361900 | 70 | 114 | 58 | 13 | 116 | 60 | 137 | 76 | 54 | 41 | 9 | 8 | 353 | 97 | 189 | 635 | 640 | 134 | 112 | 61 | 13 | 91 |
| PBANKA_1362000 | 5 | 7 | 359 | 62 | 321 | 188 | 334 | 122 | 116 | 96 | 113 | 80 | 7144 | 2749 | 2895 | 12508 | 2282 | 352 | 3585 | 2501 | 6339 | 51982 |
| PBANKA_1362100 | 10 | 0 | 62 | 12 | 304 | 113 | 601 | 191 | 337 | 103 | 64 | 18 | 832 | 416 | 180 | 478 | 5062 | 593 | 2477 | 3837 | 275 | 1696 |
| PBANKA_1362200 | 14 | 23 | 20 | 8 | 80 | 37 | 92 | 32 | 27 | 22 | 6 | 10 | 174 | 78 | 96 | 296 | 150 | 61 | 443 | 170 | 40 | 223 |
| PBANKA_1362300 | 70 | 105 | 247 | 41 | 1066 | 330 | 1982 | 666 | 1166 | 446 | 175 | 57 | 2868 | 2155 | 2547 | 12570 | 32108 | 3804 | 1466 | 4572 | 659 | 3555 |
| PBANKA_1362400 | 14 | 8 | 58 | 16 | 187 | 86 | 386 | 152 | 163 | 86 | 24 | 7 | 512 | 91 | 199 | 538 | 2237 | 346 | 604 | 328 | 67 | 546 |
| PBANKA_1362500 | 141 | 207 | 17 | 0 | 115 | 76 | 176 | 50 | 60 | 33 | 22 | 5 | 253 | 88 | 45 | 220 | 422 | 131 | 531 | 306 | 486 | 3079 |
| PBANKA_1362600 | 54 | 52 | 148 | 28 | 197 | 71 | 143 | 64 | 69 | 39 | 286 | 179 | 1473 | 826 | 233 | 1428 | 261 | 40 | 403 | 362 | 145 | 685 |
| PBANKA_1362700 | 9 | 6 | 39 | 0 | 87 | 23 | 63 | 16 | 31 | 17 | 53 | 16 | 406 | 99 | 128 | 356 | 90 | 41 | 1660 | 360 | 772 | 3942 |
| PBANKA_1362900 | 174 | 249 | 313 | 89 | 594 | 318 | 845 | 449 | 317 | 141 | 43 | 14 | 2618 | 636 | 1006 | 4370 | 4104 | 637 | 780 | 569 | 247 | 2102 |
| PBANKA_1363000 | 14 | 12 | 56 | 15 | 125 | 83 | 123 | 57 | 27 | 27 | 152 | 106 | 299 | 133 | 123 | 355 | 167 | 54 | 361 | 129 | 88 | 569 |
| PBANKA_1363100 | 49 | 60 | 95 | 10 | 135 | 46 | 131 | 46 | 86 | 49 | 661 | 391 | 1954 | 1415 | 244 | 1578 | 922 | 179 | 1260 | 861 | 415 | 1461 |
| PBANKA_1363200 | 59 | 64 | 19 | 0 | 164 | 41 | 171 | 49 | 90 | 50 | 20 | 6 | 344 | 112 | 160 | 835 | 706 | 140 | 3259 | 1111 | 194 | 1013 |
| PBANKA_1363300 | 0 | 0 | 43 | 8 | 128 | 65 | 238 | 97 | 704 | 405 | 161 | 114 | 998 | 614 | 149 | 759 | 5241 | 1343 | 512 | 400 | 89 | 871 |
| PBANKA_1363400 | 0 | 0 | 7 | 0 | 47 | 20 | 65 | 38 | 158 | 91 | 14 | 7 | 159 | 32 | 77 | 208 | 731 | 295 | 334 | 250 | 36 | 292 |
| PBANKA_1363500 | 0 | 0 | 186 | 33 | 221 | 116 | 240 | 95 | 104 | 32 | 17 | 5 | 998 | 268 | 380 | 2287 | 1072 | 128 | 2092 | 1002 | 110 | 734 |
| PBANKA_1363700 | 238 | 388 | 254 | 61 | 837 | 305 | 691 | 188 | 182 | 87 | 41 | 14 | 62 | 32 | 21 | 76 | 188 | 25 | 855 | 715 | 1447 | 10629 |
| PBANKA_1363800 | 32 | 30 | 474 | 105 | 1884 | 740 | 2355 | 627 | 689 | 296 | 133 | 60 | 2110 | 1743 | 1001 | 6134 | 1899 | 204 | 1587 | 2192 | 363 | 2224 |
| PBANKA_1363900 | 0 | 0 | 51 | 16 | 104 | 84 | 123 | 82 | 38 | 26 | 0 | 10 | 237 | 70 | 177 | 410 | 190 | 44 | 383 | 113 | 27 | 190 |
| PBANKA_1364000 | 158 | 261 | 143 | 25 | 391 | 142 | 580 | 203 | 255 | 102 | 41 | 10 | 2012 | 1570 | 1277 | 7368 | 6203 | 638 | 807 | 1886 | 1464 | 9688 |
| PBANKA_1364100 | 7 | 0 | 20 | 11 | 40 | 23 | 50 | 19 | 23 | 19 | 186 | 151 | 1255 | 333 | 240 | 1210 | 383 | 98 | 1580 | 627 | 416 | 3055 |
| PBANKA_1364200 | 158 | 226 | 2036 | 604 | 1379 | 846 | 1262 | 814 | 525 | 314 | 2784 | 2238 | 13864 | 4906 | 2651 | 7714 | 2466 | 522 | 3254 | 1095 | 397 | 3630 |
| PBANKA_1364250 | 0 | 0 | 5 | 0 | 0 | 0 | 0 | 0 | 0 | 0 | 0 | 0 | 398 | 70 | 35 | 94 | 71 | 8 | 1052 | 84 | 5 | 15 |
| PBANKA_1364300 | 0 | 14 | 105 | 42 | 565 | 545 | 1031 | 595 | 270 | 182 | 15 | 10 | 1824 | 517 | 1691 | 5232 | 4772 | 825 | 16201 | 9203 | 529 | 5308 |
| PBANKA_1364500 | 0 | 0 | 51 | 14 | 75 | 58 | 59 | 26 | 107 | 83 | 114 | 75 | 1305 | 326 | 314 | 1145 | 230 | 154 | 314 | 121 | 15 | 109 |
| PBANKA_1364600 | 10 | 14 | 42 | 9 | 112 | 62 | 183 | 84 | 48 | 28 | 36 | 18 | 1382 | 637 | 381 | 2325 | 1707 | 205 | 851 | 680 | 131 | 807 |
| PBANKA_1364700 | 53 | 60 | 234 | 41 | 633 | 208 | 590 | 240 | 360 | 200 | 249 | 117 | 4343 | 4803 | 1644 | 12103 | 7172 | 1028 | 9354 | 40965 | 1477 | 7508 |
| PBANKA_1364800 | 203 | 251 | 71 | 17 | 191 | 92 | 177 | 73 | 142 | 86 | 92 | 88 | 924 | 574 | 215 | 953 | 899 | 247 | 698 | 501 | 604 | 3644 |
| PBANKA_1364900 | 99 | 114 | 117 | 32 | 90 | 53 | 83 | 23 | 52 | 39 | 509 | 410 | 1509 | 894 | 198 | 901 | 194 | 32 | 1171 | 691 | 445 | 2366 |
| PBANKA_1365000 | 0 | 6 | 38 | 9 | 90 | 36 | 79 | 22 | 29 | 11 | 17 | 5 | 832 | 283 | 131 | 627 | 391 | 72 | 1190 | 413 | 25 | 184 |
| PBANKA_1365100 | 44 | 64 | 97 | 15 | 225 | 75 | 338 | 136 | 162 | 70 | 47 | 25 | 1603 | 1059 | 716 | 4165 | 6880 | 639 | 2345 | 2608 | 62 | 388 |
| PBANKA_1365200 | 0 | 0 | 362 | 82 | 700 | 433 | 1390 | 798 | 665 | 639 | 69 | 32 | 3213 | 407 | 1550 | 2858 | 11642 | 3894 | 1627 | 566 | 101 | 1033 |
| PBANKA_1365300 | 53 | 86 | 244 | 51 | 564 | 242 | 882 | 308 | 415 | 176 | 169 | 95 | 2455 | 3054 | 1609 | 8387 | 6557 | 718 | 1067 | 2085 | 208 | 1508 |
| PBANKA_1365400 | 15 | 20 | 105 | 17 | 308 | 130 | 349 | 153 | 132 | 69 | 15 | 8 | 723 | 265 | 868 | 4347 | 2341 | 278 | 1031 | 1076 | 47 | 272 |
| PBANKA_1365500 | 0 | 0 | 157 | 61 | 217 | 126 | 201 | 119 | 169 | 107 | 4217 | 3832 | 58231 | 28103 | 6334 | 19130 | 5897 | 1048 | 2916 | 1895 | 93 | 1357 |
| PBANKA_1365600 | 0 | 0 | 11 | 0 | 14 | 16 | 33 | 18 | 27 | 27 | 532 | 439 | 469 | 220 | 208 | 515 | 431 | 139 | 261 | 139 | 18 | 178 |
| PBANKA_1365680 | 0 | 0 | 5 | 0 | 0 | 0 | 0 | 0 | 0 | 0 | 15 | 21 | 189 | 37 | 89 | 133 | 70 | 22 | 24 | 16 | 0 | 14 |
| PBANKA_1400091 | 5 | 7 | 21 | 0 | 145 | 47 | 120 | 62 | 608 | 374 | 164 | 81 | 211 | 500 | 172 | 518 | 2010 | 555 | 158 | 511 | 78 | 273 |
| PBANKA_1400600 | 6 | 8 | 129 | 46 | 3921 | 2368 | 23833 | 10657 | 16327 | 8783 | 1293 | 626 | 26377 | 6178 | 11385 | 38565 | 2E+05 | 19339 | 2385 | 2343 | 179 | 2470 |
| PBANKA_1400800 | 7485 | 8318 | 125 | 24 | 219 | 104 | 162 | 72 | 141 | 91 | 113 | 38 | 737 | 380 | 63 | 331 | 733 | 205 | 601 | 252 | 43 | 229 |
| PBANKA_1401000 | 230 | 315 | 118 | 35 | 439 | 178 | 642 | 247 | 284 | 116 | 47 | 18 | 726 | 350 | 1605 | 7528 | 2959 | 376 | 1555 | 3068 | 30 | 171 |
| PBANKA_1401100 | 35 | 46 | 104 | 31 | 202 | 118 | 230 | 84 | 106 | 54 | 370 | 273 | 1161 | 688 | 385 | 1620 | 743 | 101 | 156 | 180 | 68 | 402 |
| PBANKA_1401200 | 63 | 89 | 218 | 52 | 324 | 147 | 297 | 104 | 205 | 105 | 2173 | 1272 | 2457 | 3197 | 576 | 2710 | 978 | 166 | 253 | 509 | 82 | 513 |
| PBANKA_1401300 | 0 | 5 | 1536 | 421 | 998 | 583 | 1071 | 570 | 312 | 207 | 1205 | 1000 | 18258 | 7900 | 4500 | 14277 | 4310 | 626 | 2642 | 1574 | 55 | 453 |
| PBANKA_1401400 | 5 | 6 | 77 | 29 | 108 | 75 | 107 | 26 | 65 | 22 | 27 | 0 | 1284 | 624 | 533 | 2001 | 467 | 94 | 450 | 264 | 6 | 61 |
| PBANKA_1401500 | 96 | 111 | 91 | 30 | 125 | 81 | 204 | 133 | 69 | 54 | 9 | 5 | 423 | 60 | 244 | 749 | 794 | 195 | 494 | 267 | 12 | 82 |
| PBANKA_1401600 | 140 | 174 | 227 | 57 | 250 | 153 | 409 | 237 | 181 | 110 | 16 | 9 | 751 | 106 | 583 | 1742 | 1747 | 323 | 1060 | 545 | 25 | 254 |
| PBANKA_1401700 | 0 | 0 | 44 | 13 | 78 | 72 | 88 | 39 | 24 | 13 | 5 | 0 | 274 | 69 | 135 | 480 | 115 | 18 | 323 | 131 | 49 | 306 |
| PBANKA_1401800 | 0 | 0 | 46 | 14 | 124 | 42 | 162 | 51 | 38 | 18 | 5 | 0 | 610 | 264 | 298 | 1350 | 369 | 82 | 235 | 224 | 27 | 159 |
| PBANKA_1401900 | 17 | 16 | 112 | 31 | 257 | 134 | 264 | 93 | 127 | 67 | 821 | 693 | 1050 | 831 | 188 | 1256 | 534 | 76 | 277 | 229 | 31 | 227 |
| PBANKA_1402000 | 33 | 64 | 159 | 35 | 838 | 458 | 891 | 377 | 606 | 279 | 68 | 41 | 2144 | 1072 | 999 | 5376 | 9026 | 1353 | 938 | 1583 | 530 | 4211 |
| PBANKA_1402100 | 14 | 26 | 11 | 0 | 60 | 35 | 78 | 36 | 30 | 13 | 0 | 0 | 123 | 16 | 36 | 127 | 124 | 62 | 136 | 63 | 19 | 126 |
| PBANKA_1402200 | 66 | 169 | 52 | 12 | 402 | 144 | 475 | 232 | 152 | 106 | 23 | 7 | 1133 | 202 | 508 | 2049 | 980 | 219 | 2366 | 1382 | 145 | 979 |
| PBANKA_1402300 | 63 | 95 | 497 | 164 | 1280 | 447 | 2651 | 964 | 1918 | 854 | 215 | 75 | 284 | 451 | 1312 | 5489 | 6445 | 664 | 611 | 5726 | 62 | 354 |
| PBANKA_1402400 | 5 | 0 | 9 | 5 | 19 | 39 | 75 | 46 | 76 | 74 | 15 | 13 | 160 | 30 | 32 | 47 | 105 | 40 | 1106 | 311 | 5 | 39 |
| PBANKA_1402500 | 94 | 164 | 91 | 34 | 206 | 105 | 427 | 215 | 288 | 185 | 26 | 25 | 784 | 409 | 542 | 1860 | 5073 | 872 | 4577 | 8014 | 74 | 642 |
| PBANKA_1402600 | 142 | 195 | 172 | 56 | 634 | 265 | 842 | 333 | 373 | 211 | 60 | 27 | 1330 | 291 | 650 | 2511 | 5334 | 861 | 6450 | 4442 | 135 | 1426 |
| PBANKA_1402700 | 0 | 0 | 55 | 16 | 87 | 79 | 221 | 90 | 49 | 37 | 9 | 5 | 568 | 84 | 252 | 1512 | 1361 | 208 | 288 | 203 | 12 | 163 |
| PBANKA_1402800 | 0 | 0 | 66 | 23 | 87 | 55 | 81 | 35 | 35 | 18 | 0 | 6 | 202 | 20 | 143 | 501 | 170 | 32 | 213 | 121 | 11 | 80 |
| PBANKA_1402900 | 0 | 0 | 42 | 15 | 155 | 67 | 178 | 51 | 63 | 24 | 15 | 0 | 390 | 262 | 344 | 1627 | 326 | 42 | 832 | 984 | 5 | 70 |
| PBANKA_1403000 | 0 | 0 | 57 | 22 | 168 | 102 | 145 | 78 | 31 | 20 | 9 | 5 | 167 | 10 | 273 | 850 | 83 | 13 | 117 | 62 | 0 | 8 |
| PBANKA_1403100 | 15 | 15 | 115 | 33 | 466 | 252 | 479 | 201 | 162 | 65 | 23 | 17 | 180 | 32 | 479 | 2097 | 630 | 86 | 106 | 80 | 11 | 86 |
| PBANKA_1403200 | 0 | 0 | 35 | 6 | 98 | 81 | 137 | 65 | 42 | 35 | 0 | 5 | 243 | 84 | 128 | 408 | 451 | 93 | 178 | 70 | 0 | 28 |
| PBANKA_1403300 | 370 | 583 | 36 | 7 | 208 | 78 | 200 | 62 | 98 | 52 | 44 | 7 | 304 | 330 | 51 | 151 | 1182 | 212 | 1551 | 5964 | 175 | 1159 |

| | | | | | | | | | | | | | | | | | | | | | | |
|---|---|---|---|---|---|---|---|---|---|---|---|---|---|---|---|---|---|---|---|---|---|---|
| PBANKA_1403400 | 5 | 17 | 63 | 13 | 93 | 44 | 62 | 38 | 13 | 7 | 19 | 13 | 201 | 91 | 107 | 224 | 36 | 10 | 266 | 84 | 5 | 64 |
| PBANKA_1403500 | 181 | 357 | 166 | 46 | 472 | 223 | 759 | 275 | 318 | 140 | 51 | 19 | 663 | 372 | 520 | 2125 | 7211 | 1039 | 589 | 1273 | 111 | 971 |
| PBANKA_1403600 | 0 | 0 | 42 | 8 | 121 | 25 | 112 | 43 | 51 | 17 | 10 | 0 | 318 | 73 | 497 | 2110 | 1058 | 184 | 462 | 449 | 18 | 137 |
| PBANKA_1403700 | 0 | 0 | 83 | 15 | 121 | 62 | 94 | 37 | 38 | 19 | 8 | 0 | 303 | 64 | 192 | 978 | 289 | 56 | 2703 | 1875 | 42 | 201 |
| PBANKA_1403800 | 8 | 19 | 497 | 128 | 876 | 371 | 948 | 328 | 317 | 119 | 313 | 198 | 3276 | 6797 | 2828 | 16102 | 4282 | 343 | 908 | 4385 | 124 | 841 |
| PBANKA_1403900 | 0 | 6 | 36 | 0 | 105 | 50 | 128 | 44 | 74 | 37 | 15 | 7 | 879 | 245 | 318 | 1658 | 606 | 86 | 513 | 261 | 25 | 126 |
| PBANKA_1404000 | 0 | 7 | 37 | 8 | 120 | 70 | 166 | 67 | 69 | 28 | 0 | 0 | 307 | 142 | 346 | 1309 | 301 | 52 | 316 | 253 | 9 | 56 |
| PBANKA_1404100 | 7 | 7 | 89 | 27 | 259 | 127 | 327 | 144 | 112 | 74 | 14 | 10 | 539 | 53 | 351 | 1357 | 999 | 221 | 645 | 304 | 42 | 275 |
| PBANKA_1404200 | 0 | 0 | 30 | 10 | 52 | 42 | 63 | 37 | 38 | 18 | 0 | 0 | 133 | 57 | 41 | 135 | 107 | 26 | 169 | 66 | 6 | 70 |
| PBANKA_1404400 | 36 | 43 | 69 | 23 | 124 | 106 | 150 | 50 | 83 | 35 | 83 | 66 | 1403 | 667 | 187 | 1270 | 800 | 119 | 1163 | 865 | 934 | 6526 |
| PBANKA_1404500 | 0 | 0 | 5 | 0 | 44 | 21 | 117 | 58 | 189 | 87 | 16 | 15 | 271 | 168 | 25 | 57 | 2712 | 570 | 286 | 150 | 21 | 122 |
| PBANKA_1404600 | 0 | 0 | 28 | 9 | 60 | 17 | 38 | 17 | 20 | 0 | 0 | 0 | 104 | 11 | 57 | 279 | 67 | 8 | 204 | 69 | 12 | 72 |
| PBANKA_1404700 | 88 | 157 | 114 | 33 | 436 | 205 | 798 | 326 | 262 | 155 | 16 | 0 | 965 | 235 | 437 | 1795 | 5153 | 585 | 641 | 477 | 260 | 2484 |
| PBANKA_1404800 | 0 | 0 | 98 | 14 | 147 | 60 | 139 | 51 | 45 | 12 | 5 | 0 | 526 | 97 | 683 | 3235 | 498 | 88 | 586 | 261 | 19 | 121 |
| PBANKA_1404900 | 0 | 0 | 211 | 46 | 395 | 231 | 696 | 292 | 326 | 162 | 19 | 23 | 3341 | 541 | 1247 | 7779 | 8136 | 892 | 1687 | 1456 | 51 | 409 |
| PBANKA_1405100 | 84 | 94 | 76 | 23 | 175 | 97 | 262 | 103 | 101 | 52 | 16 | 8 | 996 | 273 | 533 | 1981 | 1411 | 243 | 1856 | 785 | 79 | 612 |
| PBANKA_1405200 | 174 | 242 | 64 | 27 | 162 | 87 | 263 | 114 | 129 | 59 | 207 | 144 | 2052 | 697 | 519 | 3325 | 3293 | 487 | 2088 | 1276 | 118 | 1074 |
| PBANKA_1405300 | 27 | 46 | 47 | 6 | 80 | 28 | 96 | 43 | 50 | 16 | 90 | 43 | 1196 | 816 | 306 | 1836 | 945 | 149 | 399 | 509 | 86 | 745 |
| PBANKA_1405400 | 59 | 109 | 281 | 53 | 440 | 309 | 518 | 176 | 188 | 87 | 293 | 217 | 1412 | 1005 | 542 | 2251 | 788 | 141 | 454 | 305 | 34 | 244 |
| PBANKA_1405500 | 0 | 0 | 91 | 26 | 159 | 94 | 195 | 69 | 53 | 22 | 150 | 117 | 830 | 322 | 356 | 1223 | 337 | 73 | 303 | 165 | 33 | 320 |
| PBANKA_1405600 | 6 | 9 | 90 | 19 | 95 | 39 | 208 | 98 | 221 | 104 | 426 | 290 | 2159 | 1786 | 229 | 996 | 663 | 154 | 938 | 1014 | 107 | 738 |
| PBANKA_1405700 | 0 | 0 | 69 | 26 | 137 | 72 | 134 | 55 | 46 | 33 | 11 | 10 | 1304 | 811 | 344 | 2261 | 732 | 71 | 283 | 310 | 12 | 97 |
| PBANKA_1405800 | 0 | 0 | 173 | 31 | 318 | 161 | 334 | 121 | 100 | 33 | 29 | 11 | 1436 | 929 | 598 | 2675 | 926 | 92 | 358 | 393 | 31 | 267 |
| PBANKA_1405900 | 13 | 6 | 17 | 0 | 32 | 8 | 23 | 6 | 34 | 11 | 10 | 7 | 160 | 60 | 127 | 607 | 289 | 84 | 998 | 696 | 73 | 582 |
| PBANKA_1406000 | 61 | 65 | 7 | 0 | 14 | 6 | 14 | 0 | 35 | 13 | 10 | 0 | 68 | 16 | 12 | 27 | 207 | 70 | 146 | 98 | 81 | 368 |
| PBANKA_1406100 | 30 | 59 | 59 | 19 | 389 | 177 | 748 | 230 | 524 | 276 | 126 | 42 | 543 | 350 | 449 | 2208 | 2542 | 355 | 1143 | 1423 | 430 | 3917 |
| PBANKA_1406200 | 184 | 281 | 156 | 26 | 410 | 170 | 732 | 287 | 552 | 275 | 431 | 332 | 2527 | 3076 | 970 | 5703 | 7603 | 1191 | 1535 | 2916 | 1333 | 8936 |
| PBANKA_1406300 | 9 | 7 | 92 | 18 | 344 | 137 | 441 | 162 | 190 | 95 | 39 | 10 | 1111 | 346 | 511 | 2824 | 2669 | 505 | 844 | 715 | 82 | 655 |
| PBANKA_1406400 | 0 | 0 | 8 | 0 | 70 | 25 | 310 | 129 | 518 | 267 | 56 | 39 | 622 | 260 | 14 | 24 | 6206 | 1623 | 3362 | 3050 | 188 | 1432 |
| PBANKA_1406600 | 0 | 0 | 31 | 0 | 124 | 47 | 404 | 163 | 212 | 120 | 35 | 11 | 277 | 71 | 159 | 423 | 903 | 195 | 1405 | 512 | 21 | 251 |
| PBANKA_1406700 | 91 | 111 | 1140 | 315 | 1527 | 727 | 1358 | 407 | 575 | 242 | 4302 | 2344 | 7765 | 11066 | 4570 | 25136 | 3563 | 325 | 1151 | 2489 | 386 | 3718 |
| PBANKA_1406800 | 18 | 31 | 504 | 192 | 210 | 169 | 300 | 154 | 73 | 70 | 601 | 695 | 973 | 971 | 554 | 948 | 279 | 89 | 295 | 148 | 20 | 111 |
| PBANKA_1406900 | 22 | 54 | 525 | 117 | 1469 | 633 | 1629 | 613 | 722 | 319 | 108 | 61 | 1743 | 1796 | 2991 | 14779 | 13439 | 1035 | 1120 | 9696 | 290 | 2022 |
| PBANKA_1407000 | 0 | 0 | 26 | 0 | 46 | 24 | 79 | 43 | 50 | 26 | 0 | 0 | 165 | 62 | 91 | 396 | 655 | 115 | 1278 | 790 | 155 | 1140 |
| PBANKA_1407100 | 0 | 5 | 88 | 19 | 179 | 93 | 228 | 104 | 77 | 56 | 0 | 0 | 582 | 211 | 257 | 1430 | 913 | 102 | 541 | 475 | 36 | 220 |
| PBANKA_1407200 | 0 | 0 | 35 | 0 | 127 | 61 | 215 | 82 | 94 | 46 | 13 | 8 | 283 | 57 | 168 | 569 | 661 | 122 | 385 | 234 | 9 | 119 |
| PBANKA_1407300 | 0 | 0 | 64 | 26 | 198 | 106 | 212 | 103 | 126 | 84 | 24 | 12 | 475 | 131 | 186 | 711 | 2186 | 390 | 513 | 296 | 28 | 219 |
| PBANKA_1407400 | 96 | 157 | 243 | 71 | 669 | 332 | 822 | 319 | 451 | 255 | 2476 | 1576 | 4108 | 5414 | 1918 | 14476 | 6943 | 798 | 3885 | 9877 | 773 | 5891 |
| PBANKA_1407500 | 0 | 0 | 51 | 10 | 87 | 45 | 73 | 43 | 40 | 9 | 11 | 8 | 506 | 303 | 162 | 569 | 131 | 40 | 579 | 211 | 27 | 84 |
| PBANKA_1407600 | 12 | 11 | 1489 | 438 | 1091 | 809 | 1192 | 779 | 385 | 344 | 1953 | 1807 | 14520 | 6370 | 3884 | 12291 | 4263 | 1012 | 3556 | 1850 | 179 | 1716 |
| PBANKA_1407700 | 13 | 20 | 31 | 8 | 112 | 48 | 156 | 46 | 250 | 141 | 71 | 38 | 2904 | 1737 | 553 | 2551 | 1949 | 416 | 2507 | 1877 | 376 | 3344 |
| PBANKA_1407800 | 15 | 27 | 114 | 33 | 344 | 119 | 295 | 104 | 151 | 61 | 749 | 442 | 2277 | 1582 | 598 | 3344 | 1990 | 277 | 2143 | 2724 | 34 | 240 |
| PBANKA_1407900 | 0 | 6 | 80 | 23 | 230 | 116 | 223 | 98 | 153 | 69 | 520 | 333 | 3397 | 3519 | 667 | 3987 | 2313 | 538 | 35989 | 55657 | 56 | 565 |
| PBANKA_1408000 | 0 | 0 | 61 | 15 | 76 | 29 | 55 | 20 | 45 | 17 | 57 | 25 | 1947 | 947 | 206 | 1057 | 742 | 174 | 403 | 439 | 31 | 214 |
| PBANKA_1408100 | 0 | 0 | 61 | 17 | 81 | 30 | 65 | 26 | 18 | 23 | 19 | 10 | 582 | 236 | 241 | 642 | 227 | 66 | 78 | 43 | 7 | 77 |
| PBANKA_1408200 | 10 | 21 | 71 | 23 | 185 | 90 | 192 | 58 | 92 | 29 | 41 | 40 | 1114 | 1045 | 791 | 3428 | 968 | 130 | 644 | 863 | 34 | 241 |
| PBANKA_1408300 | 0 | 0 | 104 | 29 | 93 | 75 | 90 | 33 | 20 | 17 | 8 | 0 | 846 | 179 | 421 | 1574 | 237 | 44 | 150 | 68 | 0 | 25 |
| PBANKA_1408400 | 0 | 0 | 6 | 0 | 32 | 29 | 35 | 17 | 11 | 8 | 0 | 0 | 70 | 18 | 65 | 159 | 37 | 10 | 78 | 21 | 0 | 7 |
| PBANKA_1408500 | 0 | 0 | 56 | 13 | 126 | 53 | 227 | 74 | 50 | 38 | 9 | 0 | 231 | 64 | 223 | 787 | 903 | 167 | 911 | 462 | 47 | 335 |
| PBANKA_1408600 | 0 | 0 | 59 | 13 | 162 | 103 | 160 | 74 | 72 | 40 | 12 | 0 | 206 | 71 | 257 | 941 | 372 | 72 | 495 | 339 | 34 | 285 |
| PBANKA_1408800 | 13 | 20 | 9 | 0 | 53 | 24 | 71 | 29 | 35 | 15 | 142 | 102 | 265 | 102 | 58 | 268 | 104 | 23 | 523 | 126 | 71 | 388 |
| PBANKA_1408900 | 0 | 0 | 59 | 26 | 321 | 220 | 447 | 192 | 168 | 90 | 101 | 45 | 2063 | 755 | 1053 | 4096 | 1956 | 374 | 661 | 494 | 13 | 125 |
| PBANKA_1409000 | 0 | 0 | 38 | 7 | 151 | 72 | 206 | 80 | 81 | 33 | 8 | 8 | 569 | 150 | 297 | 805 | 514 | 144 | 915 | 482 | 13 | 97 |
| PBANKA_1409100 | 0 | 0 | 9 | 0 | 94 | 74 | 248 | 145 | 186 | 133 | 33 | 5 | 294 | 111 | 163 | 521 | 706 | 118 | 77 | 64 | 13 | 137 |
| PBANKA_1409200 | 419 | 679 | 11 | 8 | 66 | 40 | 127 | 57 | 657 | 619 | 206 | 134 | 518 | 244 | 38 | 63 | 2149 | 1457 | 1645 | 732 | 14682 | 1E+05 |
| PBANKA_1409300 | 7 | 6 | 134 | 36 | 255 | 166 | 242 | 68 | 125 | 71 | 941 | 595 | 1257 | 1302 | 544 | 3048 | 449 | 58 | 347 | 497 | 290 | 1963 |
| PBANKA_1409400 | 0 | 5 | 102 | 24 | 320 | 180 | 324 | 143 | 86 | 53 | 232 | 192 | 323 | 152 | 312 | 902 | 198 | 28 | 164 | 97 | 22 | 222 |
| PBANKA_1409500 | 22 | 36 | 42 | 5 | 66 | 21 | 53 | 25 | 25 | 13 | 0 | 0 | 290 | 62 | 86 | 209 | 54 | 16 | 319 | 91 | 13 | 182 |
| PBANKA_1409600 | 5 | 7 | 12 | 0 | 99 | 30 | 66 | 35 | 53 | 21 | 22 | 0 | 1711 | 1195 | 75 | 447 | 2539 | 486 | 5302 | 21871 | 86 | 632 |
| PBANKA_1409700 | 13 | 16 | 51 | 16 | 204 | 82 | 359 | 130 | 189 | 92 | 28 | 8 | 704 | 364 | 386 | 2626 | 880 | 92 | 388 | 617 | 22 | 140 |
| PBANKA_1409800 | 9 | 16 | 153 | 34 | 312 | 216 | 492 | 279 | 204 | 152 | 29 | 16 | 213 | 68 | 270 | 772 | 889 | 160 | 380 | 186 | 45 | 531 |
| PBANKA_1409900 | 0 | 0 | 82 | 24 | 265 | 102 | 570 | 242 | 327 | 157 | 17 | 6 | 975 | 212 | 609 | 2043 | 2057 | 434 | 757 | 501 | 118 | 959 |
| PBANKA_1410000 | 353 | 427 | 363 | 124 | 1310 | 710 | 1899 | 970 | 756 | 452 | 87 | 55 | 2897 | 855 | 2040 | 8913 | 6770 | 774 | 2979 | 1863 | 87 | 995 |
| PBANKA_1410100 | 238 | 372 | 101 | 23 | 208 | 80 | 327 | 141 | 561 | 373 | 1890 | 1447 | 3800 | 2033 | 945 | 4403 | 2767 | 928 | 21452 | 16203 | 1315 | 9419 |
| PBANKA_1410200 | 51 | 96 | 80 | 17 | 89 | 59 | 84 | 33 | 60 | 49 | 18 | 16 | 279 | 164 | 230 | 465 | 231 | 79 | 257 | 146 | 47 | 308 |
| PBANKA_1410300 | 52 | 60 | 1772 | 536 | 1808 | 1167 | 1820 | 663 | 760 | 456 | 484 | 367 | 13852 | 9246 | 7175 | 27542 | 5927 | 1088 | 2474 | 3869 | 635 | 6797 |
| PBANKA_1410400 | 7 | 13 | 143 | 43 | 682 | 508 | 877 | 317 | 309 | 129 | 68 | 27 | 1165 | 407 | 1260 | 5490 | 2761 | 381 | 2360 | 1596 | 178 | 1943 |
| PBANKA_1410500 | 5 | 0 | 54 | 11 | 335 | 154 | 322 | 117 | 79 | 41 | 11 | 10 | 117 | 16 | 37 | 43 | 339 | 134 | 93 | 35 | 11 | 136 |
| PBANKA_1410600 | 220 | 267 | 17 | 10 | 173 | 66 | 531 | 204 | 302 | 161 | 39 | 25 | 409 | 231 | 85 | 345 | 5690 | 719 | 334 | 1221 | 177 | 1026 |
| PBANKA_1410700 | 13 | 21 | 65 | 12 | 145 | 63 | 182 | 89 | 87 | 51 | 0 | 10 | 312 | 46 | 130 | 509 | 620 | 116 | 156 | 103 | 11 | 67 |
| PBANKA_1410800 | 0 | 0 | 46 | 5 | 83 | 48 | 87 | 33 | 27 | 15 | 8 | 0 | 167 | 38 | 270 | 1290 | 254 | 44 | 109 | 56 | 0 | 14 |
| PBANKA_1410900 | 0 | 0 | 102 | 40 | 153 | 88 | 106 | 45 | 21 | 17 | 0 | 0 | 675 | 244 | 408 | 1975 | 220 | 25 | 35 | 80 | 7 | 44 |
| PBANKA_1411000 | 1057 | 1092 | 6 | 0 | 11 | 14 | 286 | 164 | 239 | 163 | 22 | 8 | 10 | 5 | 0 | 11 | 57 | 14 | 124 | 42 | 10 | 66 |
| PBANKA_1411100 | 332 | 391 | 146 | 48 | 386 | 284 | 1292 | 714 | 589 | 368 | 41 | 28 | 409 | 54 | 514 | 1259 | 1508 | 308 | 1442 | 671 | 214 | 2066 |
| PBANKA_1411200 | 175 | 253 | 226 | 52 | 596 | 222 | 1618 | 658 | 1558 | 802 | 311 | 152 | 1496 | 2629 | 2193 | 11084 | 14932 | 2673 | 2628 | 17083 | 329 | 2410 |
| PBANKA_1411300 | 359 | 533 | 740 | 137 | 1182 | 535 | 1678 | 658 | 747 | 344 | 781 | 569 | 10539 | 11682 | 4739 | 29435 | 24032 | 2489 | 2767 | 6268 | 671 | 4374 |
| PBANKA_1411400 | 0 | 5 | 28 | 8 | 62 | 24 | 53 | 33 | 17 | 14 | 0 | 0 | 104 | 13 | 127 | 148 | 164 | 64 | 617 | 209 | 35 | 225 |
| PBANKA_1411500 | 76 | 108 | 112 | 33 | 323 | 149 | 273 | 108 | 173 | 91 | 63 | 15 | 41 | 38 | 41 | 77 | 237 | 35 | 203 | 363 | 697 | 4517 |
| PBANKA_1411600 | 0 | 0 | 43 | 7 | 167 | 70 | 162 | 64 | 46 | 20 | 0 | 0 | 133 | 35 | 133 | 406 | 275 | 55 | 349 | 190 | 14 | 75 |
| PBANKA_1411700 | 0 | 0 | 29 | 0 | 24 | 31 | 54 | 19 | 10 | 7 | 0 | 0 | 74 | 39 | 58 | 230 | 178 | 43 | 273 | 113 | 7 | 77 |
| PBANKA_1411800 | 0 | 0 | 253 | 33 | 372 | 139 | 318 | 115 | 102 | 57 | 23 | 20 | 2064 | 627 | 784 | 4699 | 890 | 137 | 421 | 396 | 30 | 246 |
| PBANKA_1411900 | 0 | 0 | 8 | 5 | 93 | 34 | 245 | 79 | 333 | 150 | 32 | 11 | 543 | 227 | 24 | 72 | 6652 | 1605 | 368 | 250 | 19 | 155 |
| PBANKA_1412100 | 0 | 0 | 23 | 7 | 50 | 26 | 25 | 14 | 9 | 6 | 148 | 131 | 680 | 317 | 353 | 1121 | 187 | 25 | 101 | 59 | 5 | 22 |
| PBANKA_1412200 | 106 | 139 | 110 | 24 | 443 | 124 | 500 | 175 | 204 | 79 | 103 | 58 | 955 | 1222 | 1111 | 4429 | 3075 | 415 | 818 | 2490 | 67 | 413 |
| PBANKA_1412300 | 126 | 181 | 63 | 16 | 237 | 112 | 324 | 128 | 121 | 49 | 40 | 18 | 1230 | 587 | 227 | 1334 | 1489 | 223 | 317 | 237 | 147 | 1069 |
| PBANKA_1412400 | 44 | 64 | 71 | 25 | 365 | 148 | 500 | 166 | 154 | 88 | 31 | 10 | 913 | 568 | 325 | 1630 | 1234 | 167 | 1207 | 982 | 130 | 887 |
| PBANKA_1412500 | 0 | 0 | 16 | 0 | 49 | 21 | 40 | 17 | 20 | 9 | 13 | 0 | 436 | 181 | 213 | 832 | 292 | 38 | 42 | 26 | 0 | 18 |
| PBANKA_1412600 | 38 | 31 | 167 | 23 | 514 | 147 | 497 | 149 | 266 | 88 | 80 | 28 | 841 | 462 | 2210 | 9118 | 4736 | 676 | 808 | 2039 | 45 | 208 |
| PBANKA_1412800 | 0 | 0 | 43 | 10 | 123 | 71 | 130 | 44 | 25 | 10 | 7 | 0 | 300 | 45 | 173 | 956 | 344 | 52 | 300 | 217 | 15 | 73 |
| PBANKA_1412900 | 26 | 37 | 31 | 5 | 40 | 23 | 68 | 30 | 17 | 5 | 0 | 0 | 124 | 17 | 136 | 362 | 437 | 94 | 194 | 107 | 29 | 154 |

| | | | | | | | | | | | | | | | | | | | | | | |
|---|---|---|---|---|---|---|---|---|---|---|---|---|---|---|---|---|---|---|---|---|---|---|
| PBANKA_1413000 | 13 | 19 | 35 | 8 | 78 | 27 | 80 | 58 | 31 | 33 | 0 | 5 | 558 | 204 | 154 | 416 | 416 | 99 | 1219 | 606 | 89 | 513 |
| PBANKA_1413100 | 18 | 38 | 84 | 18 | 115 | 59 | 114 | 59 | 37 | 17 | 15 | 0 | 1319 | 324 | 226 | 884 | 305 | 64 | 512 | 192 | 88 | 523 |
| PBANKA_1413200 | 0 | 0 | 79 | 7 | 116 | 47 | 130 | 40 | 48 | 36 | 14 | 10 | 2006 | 1600 | 830 | 6493 | 2113 | 307 | 1264 | 1526 | 25 | 203 |
| PBANKA_1413400 | 226 | 169 | 187 | 59 | 390 | 268 | 406 | 174 | 120 | 82 | 20 | 0 | 575 | 93 | 713 | 1896 | 635 | 139 | 1591 | 542 | 165 | 1331 |
| PBANKA_1413600 | 0 | 0 | 8 | 0 | 37 | 0 | 49 | 10 | 115 | 68 | 22 | 6 | 87 | 17 | 10 | 35 | 166 | 32 | 766 | 323 | 6 | 38 |
| PBANKA_1413700 | 0 | 0 | 61 | 0 | 263 | 132 | 288 | 97 | 137 | 61 | 30 | 7 | 279 | 110 | 116 | 589 | 551 | 120 | 895 | 504 | 57 | 382 |
| PBANKA_1413800 | 0 | 0 | 16 | 0 | 127 | 46 | 460 | 141 | 574 | 251 | 60 | 28 | 377 | 116 | 18 | 55 | 2865 | 654 | 348 | 286 | 68 | 563 |
| PBANKA_1413900 | 142 | 235 | 130 | 34 | 295 | 153 | 323 | 101 | 358 | 158 | 996 | 697 | 2267 | 1235 | 733 | 3580 | 1504 | 303 | 1925 | 924 | 349 | 2053 |
| PBANKA_1414000 | 7 | 11 | 34 | 17 | 114 | 55 | 107 | 58 | 115 | 91 | 85 | 61 | 2189 | 1003 | 385 | 1169 | 685 | 161 | 634 | 305 | 256 | 1718 |
| PBANKA_1414100 | 114 | 151 | 140 | 36 | 185 | 135 | 324 | 192 | 347 | 224 | 139 | 93 | 414 | 153 | 265 | 482 | 700 | 351 | 2382 | 1433 | 152 | 971 |
| PBANKA_1414200 | 48 | 68 | 123 | 34 | 282 | 137 | 360 | 125 | 262 | 138 | 36 | 18 | 702 | 217 | 347 | 1439 | 1596 | 326 | 893 | 473 | 107 | 774 |
| PBANKA_1414300 | 0 | 0 | 10 | 0 | 81 | 33 | 112 | 30 | 60 | 24 | 20 | 21 | 1054 | 439 | 183 | 983 | 617 | 134 | 667 | 444 | 44 | 237 |
| PBANKA_1414400 | 0 | 0 | 122 | 38 | 222 | 119 | 352 | 136 | 120 | 56 | 267 | 170 | 2882 | 1699 | 810 | 3412 | 1484 | 235 | 1285 | 887 | 43 | 337 |
| PBANKA_1414600 | 0 | 12 | 129 | 30 | 481 | 171 | 467 | 128 | 128 | 65 | 44 | 9 | 509 | 175 | 735 | 3173 | 658 | 155 | 3036 | 2948 | 85 | 708 |
| PBANKA_1414700 | 0 | 0 | 31 | 5 | 144 | 39 | 170 | 48 | 45 | 28 | 7 | 0 | 154 | 24 | 205 | 575 | 544 | 117 | 699 | 255 | 64 | 413 |
| PBANKA_1414900 | 9 | 12 | 44 | 10 | 156 | 114 | 261 | 109 | 137 | 90 | 20 | 7 | 261 | 74 | 63 | 98 | 726 | 327 | 1643 | 807 | 56 | 444 |
| PBANKA_1415000 | 64 | 97 | 136 | 38 | 340 | 159 | 457 | 184 | 226 | 124 | 123 | 83 | 2002 | 1554 | 718 | 3447 | 3336 | 539 | 845 | 1027 | 670 | 5102 |
| PBANKA_1415100 | 0 | 0 | 137 | 39 | 388 | 139 | 340 | 145 | 266 | 163 | 279 | 161 | 2957 | 1327 | 802 | 4082 | 2714 | 529 | 2591 | 1780 | 86 | 644 |
| PBANKA_1415200 | 19 | 36 | 24 | 5 | 75 | 17 | 126 | 51 | 69 | 49 | 14 | 9 | 838 | 355 | 103 | 328 | 655 | 290 | 22230 | 14253 | 113 | 912 |
| PBANKA_1415300 | 0 | 0 | 46 | 13 | 157 | 76 | 236 | 90 | 83 | 54 | 11 | 0 | 512 | 100 | 162 | 667 | 1068 | 130 | 1568 | 988 | 46 | 391 |
| PBANKA_1415400 | 157 | 274 | 148 | 28 | 453 | 149 | 654 | 257 | 303 | 141 | 49 | 15 | 2652 | 1063 | 626 | 3381 | 4953 | 762 | 25512 | 21657 | 301 | 2292 |
| PBANKA_1415500 | 31 | 32 | 68 | 23 | 209 | 116 | 336 | 147 | 117 | 42 | 21 | 9 | 1538 | 600 | 657 | 3508 | 2147 | 265 | 306 | 277 | 93 | 783 |
| PBANKA_1415600 | 20 | 44 | 335 | 79 | 1124 | 714 | 2299 | 942 | 1289 | 666 | 172 | 66 | 1601 | 428 | 806 | 2509 | 8211 | 1076 | 6238 | 3762 | 131 | 1508 |
| PBANKA_1415700 | 50 | 83 | 107 | 45 | 1317 | 527 | 3631 | 1343 | 1820 | 935 | 276 | 114 | 1057 | 1077 | 1634 | 8074 | 7948 | 945 | 1873 | 4939 | 756 | 6865 |
| PBANKA_1415800 | 11 | 37 | 139 | 19 | 259 | 120 | 221 | 108 | 54 | 37 | 25 | 13 | 1369 | 524 | 391 | 1647 | 378 | 42 | 89 | 94 | 0 | 37 |
| PBANKA_1415900 | 9 | 19 | 129 | 33 | 197 | 114 | 173 | 70 | 37 | 16 | 14 | 6 | 2046 | 701 | 431 | 2520 | 324 | 37 | 74 | 49 | 0 | 32 |
| PBANKA_1416000 | 0 | 0 | 16 | 0 | 49 | 25 | 69 | 32 | 25 | 11 | 0 | 0 | 119 | 18 | 50 | 146 | 206 | 47 | 401 | 163 | 12 | 121 |
| PBANKA_1416100 | 0 | 0 | 62 | 11 | 341 | 123 | 527 | 254 | 205 | 119 | 23 | 11 | 688 | 186 | 327 | 1567 | 2637 | 362 | 353 | 459 | 73 | 505 |
| PBANKA_1416200 | 0 | 5 | 15 | 6 | 39 | 9 | 14 | 9 | 18 | 15 | 41 | 27 | 215 | 55 | 48 | 252 | 50 | 12 | 456 | 164 | 34 | 230 |
| PBANKA_1416300 | 92 | 133 | 1475 | 346 | 1164 | 585 | 1083 | 532 | 337 | 202 | 1849 | 1586 | 7961 | 3499 | 1641 | 5816 | 2008 | 383 | 2107 | 905 | 432 | 3574 |
| PBANKA_1416400 | 101 | 153 | 96 | 23 | 219 | 76 | 134 | 41 | 107 | 58 | 1233 | 673 | 2088 | 1723 | 399 | 2477 | 673 | 96 | 1206 | 1188 | 193 | 1535 |
| PBANKA_1416600 | 18 | 38 | 88 | 16 | 186 | 92 | 153 | 67 | 53 | 25 | 14 | 0 | 277 | 56 | 243 | 1098 | 373 | 37 | 149 | 133 | 24 | 273 |
| PBANKA_1416700 | 41 | 38 | 63 | 19 | 200 | 126 | 221 | 105 | 60 | 28 | 12 | 0 | 49 | 6 | 77 | 218 | 169 | 28 | 38 | 19 | 19 | 161 |
| PBANKA_1416800 | 294 | 408 | 40 | 7 | 155 | 64 | 259 | 105 | 151 | 56 | 21 | 16 | 177 | 66 | 22 | 29 | 1421 | 226 | 107 | 310 | 2664 | 18810 |
| PBANKA_1416900 | 66 | 80 | 61 | 19 | 309 | 121 | 528 | 214 | 244 | 99 | 55 | 11 | 788 | 885 | 44 | 143 | 9185 | 1124 | 2049 | 11570 | 169 | 1135 |
| PBANKA_1417100 | 0 | 8 | 80 | 28 | 237 | 104 | 243 | 109 | 105 | 49 | 11 | 6 | 281 | 80 | 482 | 1714 | 796 | 166 | 849 | 556 | 20 | 79 |
| PBANKA_1417300 | 27 | 27 | 369 | 136 | 675 | 477 | 771 | 214 | 650 | 390 | 594 | 381 | 4823 | 2589 | 1791 | 7631 | 6331 | 1113 | 3940 | 2858 | 541 | 5223 |
| PBANKA_1417400 | 0 | 0 | 7 | 0 | 53 | 19 | 58 | 26 | 33 | 18 | 6 | 9 | 382 | 227 | 55 | 241 | 151 | 59 | 9647 | 6194 | 2390 | 19742 |
| PBANKA_1417500 | 0 | 0 | 58 | 15 | 137 | 66 | 139 | 43 | 70 | 27 | 10 | 0 | 601 | 242 | 1160 | 6589 | 1083 | 163 | 1151 | 636 | 29 | 348 |
| PBANKA_1417600 | 15 | 14 | 39 | 7 | 136 | 70 | 170 | 74 | 298 | 158 | 292 | 127 | 2174 | 1035 | 471 | 2191 | 3350 | 763 | 1236 | 726 | 134 | 810 |
| PBANKA_1417800 | 40 | 80 | 230 | 53 | 288 | 88 | 237 | 86 | 108 | 37 | 508 | 350 | 3114 | 3125 | 721 | 4735 | 536 | 74 | 1061 | 1381 | 265 | 1995 |
| PBANKA_1417900 | 132 | 210 | 161 | 34 | 814 | 347 | 1503 | 627 | 727 | 342 | 91 | 57 | 2967 | 1594 | 1493 | 7062 | 20899 | 2690 | 4487 | 11274 | 96 | 819 |
| PBANKA_1418100 | 6 | 16 | 33 | 12 | 349 | 133 | 269 | 111 | 177 | 94 | 29 | 7 | 259 | 319 | 122 | 329 | 808 | 236 | 2003 | 6832 | 190 | 1454 |
| PBANKA_1418200 | 18 | 46 | 45 | 5 | 237 | 98 | 387 | 156 | 149 | 72 | 26 | 17 | 533 | 159 | 261 | 1203 | 1615 | 217 | 814 | 913 | 78 | 452 |
| PBANKA_1418300 | 73 | 115 | 213 | 55 | 592 | 334 | 840 | 380 | 538 | 348 | 555 | 455 | 6323 | 3766 | 2293 | 9923 | 5389 | 808 | 1341 | 1238 | 383 | 3197 |
| PBANKA_1418400 | 0 | 0 | 53 | 11 | 124 | 114 | 201 | 149 | 78 | 76 | 15 | 8 | 278 | 45 | 244 | 768 | 623 | 100 | 935 | 471 | 31 | 175 |
| PBANKA_1418500 | 0 | 0 | 16 | 17 | 134 | 65 | 111 | 58 | 64 | 15 | 7 | 0 | 256 | 54 | 188 | 655 | 554 | 88 | 885 | 605 | 25 | 247 |
| PBANKA_1418600 | 129 | 120 | 14 | 8 | 52 | 17 | 74 | 40 | 35 | 21 | 7 | 0 | 163 | 72 | 120 | 417 | 195 | 34 | 535 | 369 | 5 | 27 |
| PBANKA_1418700 | 6 | 0 | 36 | 14 | 99 | 44 | 68 | 40 | 22 | 7 | 6 | 0 | 276 | 121 | 217 | 917 | 361 | 62 | 129 | 85 | 0 | 12 |
| PBANKA_1418800 | 7 | 14 | 147 | 38 | 266 | 155 | 296 | 156 | 112 | 51 | 21 | 16 | 899 | 264 | 319 | 1148 | 718 | 154 | 818 | 724 | 40 | 267 |
| PBANKA_1418900 | 8 | 5 | 109 | 41 | 159 | 114 | 307 | 160 | 125 | 82 | 18 | 11 | 1730 | 516 | 513 | 1628 | 1598 | 405 | 7968 | 3359 | 80 | 713 |
| PBANKA_1419000 | 0 | 0 | 30 | 11 | 202 | 66 | 536 | 176 | 488 | 211 | 47 | 24 | 735 | 406 | 188 | 600 | 9719 | 1505 | 1264 | 1757 | 130 | 1028 |
| PBANKA_1419100 | 0 | 7 | 517 | 111 | 724 | 383 | 569 | 259 | 150 | 107 | 23 | 41 | 1866 | 524 | 1377 | 5305 | 1416 | 284 | 4946 | 2705 | 105 | 1074 |
| PBANKA_1419200 | 6 | 10 | 109 | 26 | 149 | 114 | 170 | 80 | 40 | 25 | 20 | 18 | 387 | 170 | 189 | 856 | 341 | 49 | 633 | 499 | 69 | 584 |
| PBANKA_1419500 | 0 | 0 | 26 | 8 | 35 | 28 | 16 | 14 | 8 | 0 | 0 | 7 | 203 | 37 | 47 | 306 | 150 | 20 | 48 | 45 | 0 | 0 |
| PBANKA_1419600 | 58 | 44 | 162 | 32 | 363 | 142 | 600 | 229 | 324 | 169 | 127 | 71 | 1036 | 1186 | 373 | 2119 | 1698 | 207 | 802 | 606 | 24 | 118 |
| PBANKA_1419700 | 42 | 47 | 23 | 17 | 91 | 104 | 331 | 229 | 175 | 121 | 88 | 83 | 635 | 200 | 152 | 522 | 766 | 202 | 305 | 211 | 21 | 166 |
| PBANKA_1419800 | 0 | 0 | 8 | 0 | 118 | 29 | 429 | 139 | 633 | 250 | 59 | 26 | 1249 | 369 | 86 | 299 | 9810 | 1957 | 311 | 439 | 33 | 160 |
| PBANKA_1420000 | 32 | 36 | 39 | 7 | 105 | 64 | 210 | 106 | 94 | 40 | 14 | 11 | 1597 | 330 | 559 | 1469 | 1458 | 406 | 798 | 316 | 48 | 253 |
| PBANKA_1420100 | 0 | 0 | 6 | 0 | 20 | 5 | 52 | 21 | 25 | 19 | 0 | 0 | 395 | 227 | 31 | 116 | 401 | 64 | 1574 | 1236 | 16 | 111 |
| PBANKA_1420200 | 0 | 0 | 113 | 23 | 209 | 99 | 211 | 90 | 90 | 30 | 19 | 10 | 156 | 66 | 501 | 2039 | 507 | 63 | 206 | 263 | 13 | 102 |
| PBANKA_1420300 | 190 | 263 | 51 | 14 | 143 | 89 | 233 | 128 | 88 | 52 | 20 | 17 | 1013 | 328 | 215 | 801 | 788 | 186 | 1042 | 511 | 135 | 631 |
| PBANKA_1420400 | 5 | 11 | 8 | 0 | 30 | 13 | 55 | 20 | 25 | 12 | 15 | 6 | 300 | 171 | 331 | 1435 | 702 | 105 | 475 | 392 | 9 | 72 |
| PBANKA_1420500 | 30 | 67 | 859 | 236 | 1255 | 545 | 1197 | 459 | 814 | 338 | 8644 | 5936 | 5464 | 19433 | 3782 | 22997 | 7455 | 768 | 1533 | 9502 | 645 | 5198 |
| PBANKA_1420600 | 477 | 539 | 542 | 184 | 599 | 425 | 1318 | 746 | 1732 | 1537 | 284 | 175 | 1953 | 684 | 517 | 1258 | 9528 | 3550 | 1263 | 662 | 551 | 4496 |
| PBANKA_1420800 | 57 | 101 | 221 | 61 | 421 | 279 | 875 | 326 | 485 | 206 | 880 | 569 | 2439 | 2232 | 515 | 2778 | 3736 | 524 | 12833 | 15566 | 353 | 2903 |
| PBANKA_1420900 | 36 | 48 | 53 | 20 | 368 | 171 | 670 | 272 | 241 | 135 | 539 | 335 | 1847 | 1068 | 476 | 2359 | 1866 | 341 | 1178 | 1030 | 188 | 1252 |
| PBANKA_1421100 | 0 | 0 | 141 | 42 | 260 | 126 | 289 | 113 | 86 | 26 | 57 | 25 | 1964 | 1209 | 730 | 4168 | 1404 | 131 | 1836 | 951 | 17 | 139 |
| PBANKA_1421200 | 42 | 47 | 44 | 24 | 134 | 87 | 184 | 86 | 85 | 46 | 10 | 8 | 1112 | 613 | 326 | 1579 | 1660 | 209 | 504 | 707 | 174 | 1238 |
| PBANKA_1421300 | 32 | 29 | 52 | 13 | 105 | 52 | 68 | 31 | 12 | 0 | 9 | 10 | 1110 | 267 | 440 | 1869 | 283 | 53 | 698 | 421 | 48 | 342 |
| PBANKA_1421500 | 0 | 0 | 14 | 0 | 137 | 50 | 141 | 62 | 84 | 40 | 9 | 0 | 1549 | 2028 | 65 | 259 | 1845 | 550 | 16923 | 69207 | 230 | 1268 |
| PBANKA_1421600 | 0 | 0 | 10 | 0 | 131 | 67 | 189 | 71 | 72 | 30 | 17 | 0 | 74 | 21 | 11 | 16 | 88 | 21 | 1284 | 737 | 94 | 783 |
| PBANKA_1421800 | 0 | 0 | 34 | 13 | 69 | 39 | 161 | 79 | 51 | 20 | 5 | 0 | 83 | 16 | 122 | 164 | 160 | 40 | 250 | 132 | 37 | 227 |
| PBANKA_1421900 | 0 | 0 | 140 | 39 | 311 | 148 | 347 | 149 | 181 | 63 | 20 | 8 | 907 | 251 | 551 | 2299 | 1069 | 155 | 676 | 608 | 28 | 182 |
| PBANKA_1422000 | 0 | 0 | 6 | 6 | 41 | 33 | 46 | 24 | 9 | 0 | 0 | 0 | 41 | 0 | 43 | 168 | 48 | 12 | 241 | 114 | 13 | 75 |
| PBANKA_1422100 | 5 | 5 | 22 | 14 | 211 | 113 | 250 | 66 | 81 | 34 | 8 | 0 | 475 | 68 | 377 | 2131 | 623 | 73 | 567 | 359 | 33 | 234 |
| PBANKA_1422200 | 30 | 37 | 49 | 14 | 135 | 82 | 120 | 56 | 66 | 30 | 5 | 0 | 838 | 107 | 658 | 2989 | 577 | 98 | 875 | 474 | 90 | 731 |
| PBANKA_1422300 | 11 | 13 | 51 | 11 | 140 | 47 | 158 | 34 | 54 | 32 | 12 | 8 | 304 | 83 | 219 | 948 | 445 | 90 | 831 | 585 | 312 | 1709 |
| PBANKA_1422400 | 49 | 75 | 131 | 33 | 288 | 159 | 319 | 116 | 74 | 77 | 109 | 103 | 3728 | 1619 | 1387 | 5867 | 3012 | 358 | 1696 | 1468 | 199 | 1658 |
| PBANKA_1422500 | 5 | 11 | 40 | 6 | 103 | 27 | 87 | 29 | 42 | 13 | 9 | 5 | 703 | 154 | 312 | 1963 | 233 | 47 | 1695 | 954 | 17 | 106 |
| PBANKA_1422600 | 0 | 0 | 158 | 38 | 225 | 116 | 196 | 63 | 64 | 18 | 9 | 7 | 446 | 125 | 451 | 1557 | 221 | 34 | 176 | 81 | 12 | 57 |
| PBANKA_1422700 | 246 | 285 | 276 | 89 | 545 | 332 | 590 | 285 | 176 | 94 | 40 | 19 | 776 | 278 | 748 | 1752 | 305 | 115 | 1635 | 457 | 211 | 1371 |
| PBANKA_1422800 | 266 | 348 | 6 | 0 | 13 | 12 | 23 | 7 | 28 | 13 | 9 | 0 | 127 | 50 | 10 | 29 | 69 | 34 | 1351 | 1152 | 2755 | 16001 |
| PBANKA_1423000 | 76 | 109 | 19 | 0 | 141 | 53 | 159 | 60 | 457 | 286 | 189 | 81 | 669 | 385 | 25 | 178 | 1986 | 861 | 1133 | 1067 | 1793 | 10932 |
| PBANKA_1423100 | 380 | 696 | 32 | 6 | 216 | 108 | 406 | 169 | 424 | 233 | 70 | 33 | 423 | 115 | 70 | 246 | 1844 | 342 | 1617 | 1147 | 1229 | 7018 |
| PBANKA_1423200 | 45 | 73 | 127 | 28 | 253 | 75 | 210 | 65 | 172 | 85 | 883 | 567 | 1287 | 1435 | 304 | 1606 | 335 | 83 | 652 | 599 | 209 | 1474 |
| PBANKA_1423300 | 84 | 99 | 3447 | 1294 | 2766 | 1985 | 3051 | 1386 | 979 | 570 | 5722 | 4073 | 57623 | 15394 | 13232 | 70804 | 23587 | 2227 | 10724 | 6445 | 845 | 9376 |
| PBANKA_1423400 | 6 | 11 | 22 | 0 | 90 | 65 | 145 | 69 | 47 | 34 | 9 | 18 | 208 | 68 | 87 | 298 | 150 | 62 | 185 | 49 | 15 | 71 |
| PBANKA_1423500 | 9 | 16 | 2104 | 617 | 1047 | 641 | 978 | 484 | 391 | 229 | 2102 | 1597 | 27706 | 9815 | 3880 | 17272 | 4376 | 705 | 2907 | 1394 | 302 | 2444 |
| PBANKA_1423600 | 11 | 7 | 1529 | 390 | 880 | 612 | 933 | 493 | 259 | 172 | 1394 | 1122 | 7757 | 2913 | 1574 | 5359 | 1519 | 321 | 1623 | 567 | 227 | 1160 |

| | | | | | | | | | | | | | | | | | | | | | | |
|---|---|---|---|---|---|---|---|---|---|---|---|---|---|---|---|---|---|---|---|---|---|---|
| PBANKA_1423700 | 0 | 0 | 6 | 0 | 89 | 37 | 102 | 38 | 37 | 22 | 0 | 0 | 527 | 246 | 35 | 80 | 244 | 79 | 18744 | 10915 | 1001 | 7034 |
| PBANKA_1423800 | 0 | 0 | 129 | 23 | 354 | 115 | 408 | 139 | 204 | 66 | 61 | 13 | 815 | 234 | 606 | 2449 | 2104 | 320 | 271 | 458 | 140 | 834 |
| PBANKA_1423900 | 0 | 0 | 187 | 36 | 137 | 53 | 144 | 37 | 37 | 19 | 19 | 13 | 2568 | 794 | 601 | 3106 | 946 | 123 | 1120 | 770 | 50 | 297 |
| PBANKA_1424000 | 155 | 280 | 26 | 0 | 88 | 29 | 131 | 65 | 75 | 38 | 15 | 0 | 767 | 158 | 476 | 3161 | 1483 | 182 | 584 | 325 | 38 | 278 |
| PBANKA_1424100 | 0 | 0 | 37 | 8 | 67 | 26 | 49 | 16 | 21 | 8 | 0 | 0 | 503 | 91 | 302 | 991 | 367 | 62 | 41 | 25 | 0 | 12 |
| PBANKA_1424200 | 0 | 5 | 575 | 105 | 781 | 324 | 783 | 309 | 279 | 120 | 91 | 51 | 7708 | 5884 | 2290 | 15309 | 2425 | 272 | 2701 | 2232 | 225 | 1398 |
| PBANKA_1424300 | 205 | 233 | 2405 | 690 | 2439 | 1304 | 2232 | 941 | 1458 | 976 | 15428 | 9809 | 13352 | 9151 | 4591 | 24485 | 7192 | 756 | 4321 | 5127 | 6525 | 46458 |
| PBANKA_1424400 | 0 | 0 | 58 | 19 | 150 | 109 | 211 | 97 | 51 | 45 | 0 | 0 | 263 | 85 | 365 | 1451 | 767 | 104 | 615 | 539 | 106 | 667 |
| PBANKA_1424500 | 32 | 42 | 8 | 0 | 27 | 15 | 75 | 24 | 161 | 115 | 27 | 5 | 546 | 282 | 49 | 136 | 2172 | 452 | 7269 | 6397 | 713 | 6167 |
| PBANKA_1424600 | 0 | 12 | 40 | 9 | 109 | 48 | 169 | 61 | 299 | 152 | 1550 | 997 | 3245 | 1876 | 429 | 2748 | 2236 | 397 | 1598 | 1028 | 270 | 1964 |
| PBANKA_1424700 | 0 | 0 | 19 | 0 | 32 | 19 | 40 | 35 | 13 | 5 | 0 | 0 | 229 | 102 | 92 | 287 | 162 | 29 | 102 | 52 | 0 | 40 |
| PBANKA_1424800 | 0 | 0 | 36 | 7 | 74 | 29 | 64 | 17 | 21 | 11 | 173 | 96 | 391 | 185 | 160 | 615 | 138 | 28 | 357 | 173 | 20 | 125 |
| PBANKA_1424900 | 0 | 0 | 26 | 5 | 81 | 57 | 97 | 49 | 27 | 13 | 248 | 149 | 556 | 215 | 191 | 493 | 138 | 41 | 694 | 270 | 18 | 164 |
| PBANKA_1425000 | 0 | 0 | 290 | 77 | 730 | 281 | 878 | 328 | 351 | 157 | 2665 | 1765 | 4176 | 7536 | 2330 | 18722 | 5463 | 375 | 272 | 1835 | 49 | 298 |
| PBANKA_1425100 | 37 | 55 | 359 | 116 | 1215 | 748 | 1878 | 710 | 621 | 334 | 279 | 158 | 4570 | 1680 | 1544 | 6420 | 5910 | 841 | 2496 | 1362 | 490 | 5153 |
| PBANKA_1425300 | 63 | 81 | 48 | 8 | 61 | 31 | 73 | 36 | 24 | 9 | 0 | 0 | 25 | 7 | 157 | 672 | 94 | 10 | 32 | 16 | 0 | 8 |
| PBANKA_1425350 | 24 | 38 | 74 | 13 | 163 | 55 | 131 | 41 | 49 | 17 | 14 | 5 | 194 | 104 | 286 | 1075 | 297 | 53 | 474 | 360 | 52 | 357 |
| PBANKA_1425400 | 5 | 9 | 265 | 41 | 975 | 362 | 770 | 301 | 293 | 124 | 72 | 38 | 1522 | 1863 | 1617 | 6427 | 2821 | 356 | 605 | 1590 | 35 | 240 |
| PBANKA_1425500 | 0 | 0 | 159 | 46 | 530 | 293 | 368 | 151 | 98 | 40 | 14 | 0 | 854 | 372 | 606 | 2715 | 412 | 94 | 643 | 586 | 36 | 295 |
| PBANKA_1425600 | 0 | 0 | 73 | 18 | 216 | 73 | 238 | 92 | 93 | 32 | 11 | 0 | 467 | 205 | 355 | 2208 | 1989 | 186 | 553 | 660 | 15 | 62 |
| PBANKA_1425700 | 0 | 0 | 32 | 5 | 93 | 30 | 121 | 46 | 44 | 23 | 0 | 0 | 213 | 55 | 121 | 524 | 492 | 82 | 735 | 255 | 0 | 16 |
| PBANKA_1425800 | 0 | 0 | 8 | 0 | 32 | 12 | 68 | 32 | 176 | 95 | 51 | 20 | 345 | 92 | 29 | 75 | 1058 | 267 | 3140 | 1930 | 13 | 62 |
| PBANKA_1425900 | 0 | 0 | 186 | 64 | 552 | 304 | 636 | 215 | 1570 | 846 | 196 | 111 | 1111 | 599 | 149 | 350 | 21690 | 5232 | 1840 | 1465 | 25 | 245 |
| PBANKA_1426000 | 0 | 0 | 106 | 23 | 246 | 114 | 244 | 94 | 294 | 110 | 90 | 44 | 792 | 872 | 407 | 1963 | 2600 | 827 | 371 | 263 | 5 | 64 |
| PBANKA_1426100 | 7 | 0 | 107 | 29 | 336 | 172 | 331 | 145 | 116 | 46 | 81 | 37 | 1093 | 599 | 922 | 4228 | 2096 | 216 | 606 | 755 | 6 | 52 |
| PBANKA_1426200 | 680 | 873 | 235 | 72 | 1009 | 441 | 1869 | 675 | 1111 | 550 | 4337 | 2571 | 6207 | 8569 | 2680 | 17655 | 19886 | 1900 | 3630 | 4591 | 995 | 7054 |
| PBANKA_1426300 | 109 | 194 | 199 | 50 | 633 | 274 | 784 | 346 | 268 | 159 | 51 | 38 | 3276 | 1713 | 1640 | 6947 | 3315 | 466 | 1360 | 1312 | 457 | 3555 |
| PBANKA_1426400 | 86 | 127 | 96 | 26 | 383 | 153 | 461 | 147 | 209 | 94 | 41 | 15 | 2456 | 706 | 922 | 3441 | 2600 | 592 | 1205 | 677 | 205 | 1521 |
| PBANKA_1426500 | 356 | 478 | 38 | 11 | 119 | 54 | 267 | 77 | 353 | 151 | 41 | 17 | 529 | 194 | 124 | 370 | 3187 | 805 | 2621 | 2385 | 324 | 2372 |
| PBANKA_1426600 | 113 | 193 | 24 | 0 | 45 | 23 | 92 | 22 | 164 | 116 | 27 | 14 | 244 | 91 | 109 | 192 | 1622 | 497 | 441 | 297 | 26 | 170 |
| PBANKA_1426700 | 58 | 60 | 89 | 26 | 61 | 25 | 78 | 43 | 53 | 23 | 21 | 6 | 939 | 169 | 989 | 3928 | 492 | 146 | 556 | 448 | 23 | 201 |
| PBANKA_1426800 | 0 | 0 | 23 | 0 | 41 | 33 | 27 | 16 | 16 | 10 | 0 | 0 | 39 | 0 | 28 | 116 | 6 | 0 | 118 | 47 | 0 | 21 |
| PBANKA_1426900 | 118 | 85 | 406 | 118 | 690 | 400 | 700 | 237 | 268 | 140 | 1048 | 682 | 6388 | 4546 | 2469 | 15644 | 3468 | 405 | 9124 | 9404 | 845 | 6997 |
| PBANKA_1427000 | 195 | 258 | 168 | 56 | 352 | 228 | 285 | 151 | 64 | 41 | 8 | 11 | 574 | 180 | 389 | 2035 | 295 | 36 | 883 | 594 | 73 | 543 |
| PBANKA_1427100 | 58 | 52 | 99 | 36 | 204 | 101 | 177 | 72 | 66 | 31 | 15 | 11 | 2000 | 785 | 514 | 2255 | 1418 | 374 | 22386 | 12210 | 336 | 2082 |
| PBANKA_1427200 | 0 | 0 | 15 | 5 | 116 | 76 | 254 | 119 | 120 | 79 | 10 | 0 | 1231 | 319 | 945 | 3490 | 2946 | 542 | 5774 | 4309 | 122 | 1017 |
| PBANKA_1427300 | 0 | 0 | 113 | 34 | 313 | 131 | 244 | 96 | 101 | 25 | 26 | 22 | 873 | 505 | 563 | 2259 | 953 | 151 | 938 | 989 | 62 | 522 |
| PBANKA_1427400 | 0 | 0 | 34 | 6 | 80 | 41 | 99 | 41 | 30 | 19 | 0 | 0 | 307 | 80 | 132 | 385 | 228 | 62 | 311 | 137 | 22 | 153 |
| PBANKA_1427500 | 0 | 0 | 101 | 28 | 135 | 95 | 235 | 152 | 126 | 98 | 12 | 17 | 151 | 69 | 288 | 744 | 259 | 81 | 510 | 192 | 47 | 328 |
| PBANKA_1427800 | 8 | 33 | 296 | 64 | 1044 | 466 | 1106 | 396 | 454 | 153 | 69 | 25 | 281 | 349 | 2367 | 9409 | 3919 | 426 | 830 | 3827 | 49 | 327 |
| PBANKA_1427900 | 5 | 5 | 8 | 5 | 41 | 29 | 60 | 18 | 17 | 8 | 56 | 35 | 349 | 139 | 96 | 373 | 336 | 58 | 65 | 57 | 54 | 416 |
| PBANKA_1428000 | 79 | 105 | 255 | 69 | 410 | 216 | 324 | 123 | 117 | 67 | 24 | 15 | 547 | 178 | 585 | 2952 | 1224 | 176 | 872 | 1389 | 300 | 2404 |
| PBANKA_1428100 | 17 | 21 | 72 | 8 | 185 | 88 | 311 | 131 | 154 | 77 | 81 | 59 | 1538 | 1008 | 562 | 2586 | 5160 | 851 | 2640 | 3031 | 153 | 1131 |
| PBANKA_1428200 | 6 | 8 | 16 | 0 | 70 | 32 | 79 | 20 | 70 | 41 | 24 | 15 | 331 | 156 | 205 | 887 | 293 | 65 | 979 | 527 | 48 | 369 |
| PBANKA_1428300 | 0 | 0 | 14 | 0 | 74 | 39 | 115 | 51 | 51 | 21 | 67 | 33 | 433 | 463 | 160 | 599 | 510 | 105 | 1435 | 1527 | 12 | 120 |
| PBANKA_1428400 | 15 | 14 | 59 | 14 | 99 | 39 | 133 | 71 | 62 | 21 | 23 | 23 | 463 | 194 | 156 | 695 | 605 | 103 | 660 | 231 | 37 | 221 |
| PBANKA_1428500 | 19 | 19 | 39 | 13 | 165 | 157 | 259 | 134 | 126 | 50 | 22 | 10 | 1247 | 341 | 449 | 1834 | 2219 | 323 | 604 | 350 | 40 | 305 |
| PBANKA_1428600 | 0 | 0 | 48 | 0 | 59 | 22 | 68 | 31 | 22 | 0 | 0 | 0 | 356 | 150 | 195 | 592 | 355 | 79 | 492 | 275 | 7 | 73 |
| PBANKA_1428700 | 219 | 390 | 2157 | 492 | 2883 | 1154 | 2455 | 846 | 771 | 330 | 3578 | 2097 | 19468 | 38471 | 10976 | 77398 | 14688 | 1235 | 4960 | 18478 | 448 | 3740 |
| PBANKA_1428800 | 224 | 350 | 162 | 35 | 228 | 105 | 231 | 97 | 72 | 43 | 13 | 5 | 1020 | 411 | 345 | 1400 | 1543 | 254 | 1765 | 1306 | 104 | 830 |
| PBANKA_1428900 | 247 | 335 | 246 | 57 | 427 | 182 | 557 | 214 | 593 | 275 | 4487 | 2739 | 3082 | 5252 | 795 | 5011 | 4591 | 599 | 1499 | 2609 | 335 | 2212 |
| PBANKA_1428980 | 0 | 0 | 6 | 0 | 0 | 0 | 6 | 0 | 0 | 0 | 9 | 0 | 137 | 54 | 988 | 112 | 134 | 109 | 983 | 98 | 8 | 36 |
| PBANKA_1429000 | 0 | 0 | 38 | 9 | 94 | 49 | 66 | 27 | 15 | 8 | 18 | 13 | 292 | 101 | 212 | 710 | 158 | 38 | 455 | 160 | 12 | 56 |
| PBANKA_1429300 | 77 | 96 | 10 | 0 | 46 | 23 | 103 | 45 | 67 | 23 | 11 | 6 | 118 | 27 | 26 | 46 | 220 | 65 | 2038 | 897 | 268 | 1729 |
| PBANKA_1429500 | 128 | 128 | 109 | 27 | 157 | 96 | 152 | 104 | 43 | 26 | 54 | 50 | 524 | 219 | 217 | 509 | 288 | 68 | 472 | 211 | 99 | 502 |
| PBANKA_1429600 | 7 | 12 | 630 | 189 | 471 | 280 | 371 | 147 | 319 | 135 | 92 | 76 | 4540 | 2556 | 1676 | 7422 | 3319 | 662 | 351 | 344 | 0 | 49 |
| PBANKA_1429700 | 8 | 5 | 304 | 71 | 638 | 347 | 511 | 171 | 160 | 47 | 92 | 32 | 4785 | 2566 | 2405 | 14068 | 2168 | 233 | 3783 | 3771 | 134 | 1014 |
| PBANKA_1429800 | 0 | 0 | 36 | 7 | 76 | 44 | 69 | 38 | 19 | 11 | 0 | 0 | 112 | 17 | 145 | 634 | 175 | 21 | 200 | 141 | 21 | 183 |
| PBANKA_1429900 | 28 | 39 | 59 | 8 | 240 | 89 | 255 | 97 | 146 | 63 | 21 | 5 | 550 | 111 | 548 | 2200 | 952 | 189 | 1110 | 838 | 122 | 865 |
| PBANKA_1430000 | 92 | 152 | 61 | 9 | 91 | 41 | 88 | 18 | 24 | 7 | 7 | 0 | 70 | 12 | 69 | 267 | 90 | 30 | 101 | 77 | 13 | 95 |
| PBANKA_1430100 | 0 | 5 | 57 | 13 | 209 | 117 | 341 | 109 | 87 | 46 | 13 | 0 | 1276 | 360 | 571 | 2674 | 1499 | 249 | 1721 | 1039 | 188 | 1359 |
| PBANKA_1430200 | 0 | 0 | 15 | 5 | 25 | 39 | 86 | 29 | 17 | 7 | 0 | 0 | 281 | 77 | 126 | 582 | 377 | 60 | 385 | 242 | 54 | 425 |
| PBANKA_1430400 | 137 | 156 | 101 | 46 | 168 | 150 | 187 | 78 | 62 | 20 | 5 | 9 | 579 | 238 | 250 | 928 | 350 | 49 | 369 | 195 | 26 | 169 |
| PBANKA_1430500 | 31 | 25 | 60 | 13 | 70 | 42 | 69 | 23 | 20 | 12 | 0 | 0 | 246 | 73 | 130 | 425 | 123 | 23 | 167 | 83 | 24 | 151 |
| PBANKA_1430600 | 5 | 0 | 6 | 0 | 10 | 13 | 41 | 18 | 21 | 9 | 0 | 0 | 146 | 56 | 21 | 51 | 115 | 132 | 1962 | 605 | 278 | 1611 |
| PBANKA_1430700 | 0 | 0 | 58 | 27 | 103 | 99 | 221 | 111 | 345 | 243 | 44 | 35 | 1405 | 357 | 354 | 1151 | 3444 | 1340 | 1703 | 774 | 47 | 515 |
| PBANKA_1430800 | 0 | 0 | 100 | 30 | 312 | 217 | 547 | 245 | 123 | 101 | 21 | 0 | 164 | 40 | 364 | 1050 | 691 | 116 | 854 | 377 | 26 | 364 |
| PBANKA_1430900 | 0 | 0 | 161 | 28 | 180 | 82 | 184 | 47 | 66 | 23 | 28 | 7 | 421 | 103 | 381 | 1581 | 647 | 144 | 3285 | 2080 | 230 | 1808 |
| PBANKA_1431000 | 0 | 0 | 81 | 15 | 127 | 65 | 101 | 38 | 35 | 30 | 8 | 0 | 147 | 17 | 325 | 1759 | 366 | 29 | 66 | 91 | 6 | 40 |
| PBANKA_1431050 | 21 | 28 | 396 | 187 | 178 | 95 | 174 | 97 | 45 | 32 | 25 | 31 | 591 | 187 | 588 | 1681 | 161 | 50 | 904 | 236 | 64 | 416 |
| PBANKA_1431100 | 176 | 265 | 46 | 11 | 289 | 108 | 465 | 189 | 255 | 104 | 196 | 92 | 1464 | 678 | 559 | 3170 | 3694 | 525 | 3793 | 3217 | 460 | 3422 |
| PBANKA_1431300 | 17 | 21 | 5 | 0 | 25 | 12 | 15 | 6 | 9 | 0 | 5 | 0 | 146 | 24 | 34 | 210 | 70 | 29 | 1365 | 485 | 16 | 113 |
| PBANKA_1431600 | 29 | 42 | 125 | 22 | 174 | 95 | 265 | 104 | 117 | 55 | 14 | 0 | 911 | 278 | 341 | 1367 | 2332 | 413 | 915 | 590 | 52 | 429 |
| PBANKA_1431700 | 0 | 6 | 74 | 20 | 190 | 79 | 171 | 59 | 56 | 31 | 31 | 24 | 962 | 496 | 510 | 2273 | 1111 | 147 | 251 | 186 | 15 | 99 |
| PBANKA_1431800 | 0 | 0 | 34 | 0 | 84 | 38 | 87 | 24 | 16 | 10 | 0 | 0 | 204 | 45 | 323 | 1267 | 412 | 79 | 345 | 197 | 17 | 122 |
| PBANKA_1431900 | 20 | 8 | 131 | 36 | 356 | 204 | 493 | 193 | 134 | 78 | 40 | 22 | 1169 | 585 | 579 | 1675 | 1063 | 192 | 372 | 186 | 15 | 130 |
| PBANKA_1432000 | 116 | 193 | 91 | 22 | 150 | 48 | 166 | 54 | 81 | 25 | 25 | 15 | 1655 | 706 | 272 | 1573 | 974 | 135 | 1021 | 650 | 14 | 120 |
| PBANKA_1432100 | 45 | 46 | 63 | 14 | 178 | 71 | 158 | 87 | 82 | 34 | 26 | 12 | 992 | 290 | 407 | 2581 | 1152 | 144 | 1059 | 626 | 28 | 159 |
| PBANKA_1432300 | 10830 | 11090 | 7 | 0 | 0 | 0 | 0 | 0 | 0 | 5 | 0 | 5 | 91 | 17 | 5 | 6 | 23 | 8 | 1658 | 638 | 9591 | 93707 |
| PBANKA_1432400 | 61 | 45 | 12 | 0 | 75 | 33 | 79 | 40 | 22 | 12 | 11 | 6 | 630 | 176 | 53 | 157 | 679 | 262 | 8623 | 4267 | 243 | 1940 |
| PBANKA_1432500 | 49 | 58 | 285 | 85 | 284 | 124 | 233 | 62 | 224 | 77 | 2335 | 1429 | 9724 | 8422 | 2208 | 13948 | 6525 | 796 | 1903 | 3145 | 165 | 1114 |
| PBANKA_1432600 | 8 | 17 | 24 | 5 | 49 | 17 | 31 | 13 | 19 | 7 | 0 | 0 | 109 | 32 | 126 | 499 | 225 | 36 | 1122 | 355 | 27 | 190 |
| PBANKA_1432700 | 83 | 102 | 7 | 0 | 82 | 49 | 88 | 27 | 36 | 22 | 11 | 8 | 60 | 42 | 9 | 40 | 88 | 51 | 637 | 280 | 121 | 732 |
| PBANKA_1432800 | 46 | 86 | 157 | 60 | 268 | 97 | 254 | 91 | 263 | 105 | 1323 | 728 | 2572 | 3529 | 452 | 2995 | 1241 | 165 | 667 | 1078 | 743 | 4749 |
| PBANKA_1432900 | 52 | 95 | 106 | 24 | 358 | 178 | 311 | 99 | 99 | 59 | 62 | 19 | 900 | 313 | 774 | 4004 | 939 | 112 | 2231 | 2466 | 326 | 2372 |
| PBANKA_1433000 | 88 | 166 | 166 | 16 | 292 | 201 | 302 | 105 | 545 | 286 | 144 | 50 | 968 | 345 | 504 | 2260 | 1826 | 453 | 4404 | 2963 | 1115 | 7611 |
| PBANKA_1433100 | 29 | 54 | 26 | 16 | 128 | 66 | 154 | 70 | 187 | 89 | 952 | 603 | 2348 | 807 | 626 | 3033 | 1921 | 310 | 952 | 569 | 141 | 1212 |
| PBANKA_1433200 | 11 | 16 | 27 | 0 | 37 | 6 | 42 | 22 | 20 | 12 | 48 | 24 | 455 | 205 | 91 | 269 | 206 | 59 | 959 | 404 | 38 | 254 |
| PBANKA_1433300 | 101 | 139 | 135 | 27 | 258 | 103 | 258 | 73 | 107 | 57 | 486 | 343 | 2008 | 1407 | 460 | 1975 | 1128 | 183 | 1034 | 835 | 291 | 1952 |
| PBANKA_1433400 | 58 | 63 | 54 | 22 | 106 | 42 | 100 | 48 | 30 | 20 | 181 | 169 | 201 | 169 | 86 | 214 | 137 | 21 | 107 | 108 | 20 | 148 |

| | | | | | | | | | | | | | | | | | | | | | | |
|---|---|---|---|---|---|---|---|---|---|---|---|---|---|---|---|---|---|---|---|---|---|---|
| PBANKA_1433500 | 34 | 47 | 42 | 14 | 244 | 107 | 283 | 155 | 122 | 56 | 53 | 28 | 591 | 171 | 415 | 1214 | 968 | 149 | 838 | 469 | 133 | 908 |
| PBANKA_1433600 | 310 | 458 | 9 | 0 | 19 | 0 | 31 | 11 | 72 | 57 | 33 | 10 | 133 | 79 | 21 | 75 | 150 | 81 | 291 | 118 | 384 | 2223 |
| PBANKA_1433700 | 200 | 275 | 7 | 0 | 78 | 43 | 146 | 77 | 82 | 47 | 9 | 6 | 285 | 133 | 23 | 32 | 1251 | 172 | 1724 | 1669 | 896 | 5649 |
| PBANKA_1433800 | 33 | 74 | 370 | 85 | 950 | 332 | 1243 | 464 | 474 | 211 | 312 | 198 | 2281 | 2929 | 2162 | 10835 | 7128 | 713 | 987 | 2263 | 77 | 491 |
| PBANKA_1433900 | 35 | 25 | 26 | 0 | 57 | 35 | 115 | 42 | 95 | 16 | 167 | 142 | 740 | 510 | 162 | 653 | 917 | 152 | 845 | 668 | 86 | 552 |
| PBANKA_1434000 | 124 | 182 | 23 | 5 | 66 | 44 | 147 | 53 | 61 | 20 | 8 | 10 | 370 | 95 | 248 | 871 | 1004 | 212 | 454 | 325 | 62 | 399 |
| PBANKA_1434100 | 27 | 38 | 64 | 25 | 273 | 187 | 322 | 183 | 297 | 140 | 1895 | 1547 | 1682 | 1320 | 873 | 4029 | 2211 | 344 | 1218 | 1165 | 56 | 596 |
| PBANKA_1434200 | 584 | 939 | 432 | 89 | 467 | 326 | 626 | 252 | 167 | 93 | 41 | 23 | 2014 | 886 | 826 | 2865 | 1839 | 300 | 608 | 471 | 55 | 495 |
| PBANKA_1434500 | 0 | 0 | 20 | 14 | 103 | 42 | 90 | 23 | 63 | 30 | 56 | 12 | 850 | 522 | 243 | 1183 | 651 | 92 | 760 | 526 | 27 | 190 |
| PBANKA_1434600 | 72 | 108 | 142 | 38 | 351 | 140 | 619 | 213 | 900 | 407 | 143 | 38 | 881 | 1378 | 215 | 653 | 10894 | 1847 | 10031 | 43309 | 88 | 498 |
| PBANKA_1434700 | 21 | 23 | 351 | 73 | 936 | 418 | 1548 | 571 | 837 | 351 | 103 | 68 | 1927 | 1496 | 2689 | 11818 | 15670 | 1920 | 1600 | 3088 | 150 | 1484 |
| PBANKA_1434800 | 0 | 0 | 78 | 23 | 371 | 224 | 691 | 270 | 827 | 382 | 100 | 56 | 2596 | 1405 | 1798 | 10399 | 8622 | 1238 | 6147 | 8030 | 79 | 561 |
| PBANKA_1435100 | 61 | 73 | 29 | 6 | 59 | 45 | 127 | 60 | 60 | 40 | 8 | 0 | 272 | 29 | 205 | 717 | 530 | 89 | 611 | 353 | 45 | 297 |
| PBANKA_1435300 | 143 | 229 | 93 | 15 | 390 | 160 | 485 | 220 | 245 | 130 | 144 | 84 | 1718 | 939 | 624 | 3240 | 2927 | 501 | 1229 | 1494 | 1215 | 9088 |
| PBANKA_1435500 | 54 | 109 | 17 | 0 | 390 | 174 | 792 | 332 | 407 | 164 | 55 | 24 | 672 | 390 | 470 | 1776 | 7915 | 943 | 1045 | 2029 | 44 | 245 |
| PBANKA_1435600 | 13 | 20 | 26 | 0 | 92 | 27 | 93 | 35 | 63 | 45 | 27 | 5 | 1304 | 564 | 306 | 1363 | 1130 | 227 | 573 | 526 | 102 | 663 |
| PBANKA_1435700 | 0 | 0 | 97 | 9 | 151 | 88 | 179 | 75 | 58 | 42 | 98 | 99 | 2374 | 903 | 495 | 2814 | 1124 | 197 | 609 | 703 | 132 | 874 |
| PBANKA_1435800 | 6 | 0 | 18 | 0 | 158 | 102 | 255 | 130 | 106 | 57 | 31 | 16 | 361 | 88 | 225 | 687 | 950 | 235 | 505 | 318 | 0 | 50 |
| PBANKA_1435900 | 698 | 1013 | 5 | 0 | 321 | 119 | 342 | 120 | 101 | 37 | 32 | 9 | 23 | 26 | 7 | 14 | 31 | 13 | 192 | 119 | 92 | 604 |
| PBANKA_1436100 | 0 | 5 | 34 | 0 | 66 | 46 | 371 | 205 | 507 | 263 | 47 | 26 | 545 | 251 | 190 | 1153 | 5468 | 579 | 403 | 427 | 110 | 740 |
| PBANKA_1436200 | 0 | 8 | 217 | 57 | 807 | 350 | 1641 | 646 | 1206 | 560 | 130 | 68 | 1412 | 1884 | 825 | 3520 | 13354 | 1817 | 2561 | 4832 | 152 | 1310 |
| PBANKA_1436400 | 59 | 66 | 47 | 13 | 70 | 36 | 73 | 62 | 29 | 18 | 49 | 38 | 219 | 171 | 47 | 223 | 88 | 21 | 548 | 152 | 15 | 207 |
| PBANKA_1436500 | 10 | 13 | 14 | 6 | 46 | 36 | 189 | 90 | 757 | 419 | 156 | 92 | 1085 | 590 | 44 | 119 | 8190 | 2118 | 7253 | 12456 | 196 | 1572 |
| PBANKA_1436700 | 0 | 0 | 77 | 16 | 148 | 76 | 131 | 57 | 42 | 15 | 13 | 7 | 364 | 139 | 360 | 1667 | 448 | 76 | 772 | 378 | 18 | 122 |
| PBANKA_1436800 | 8 | 20 | 40 | 5 | 231 | 117 | 375 | 135 | 119 | 79 | 21 | 10 | 401 | 88 | 345 | 1228 | 1481 | 174 | 763 | 444 | 13 | 139 |
| PBANKA_1436900 | 0 | 0 | 48 | 9 | 182 | 71 | 147 | 57 | 59 | 25 | 0 | 6 | 662 | 509 | 312 | 1499 | 539 | 120 | 311 | 391 | 7 | 52 |
| PBANKA_1437000 | 0 | 5 | 168 | 36 | 524 | 298 | 1013 | 418 | 425 | 215 | 66 | 46 | 1209 | 270 | 857 | 4243 | 4281 | 617 | 709 | 989 | 31 | 253 |
| PBANKA_1437100 | 0 | 0 | 17 | 0 | 94 | 53 | 195 | 83 | 642 | 429 | 1682 | 1293 | 4701 | 4117 | 511 | 3286 | 3961 | 1006 | 1261 | 1810 | 24 | 200 |
| PBANKA_1437200 | 0 | 0 | 17 | 0 | 104 | 67 | 169 | 96 | 71 | 45 | 13 | 13 | 762 | 229 | 795 | 2620 | 909 | 165 | 1672 | 1071 | 28 | 235 |
| PBANKA_1437300 | 172 | 244 | 1799 | 469 | 2910 | 1530 | 4095 | 1832 | 1739 | 991 | 287 | 125 | 8000 | 2648 | 10173 | 37239 | 37821 | 5227 | 5344 | 6041 | 609 | 6157 |
| PBANKA_1437400 | 32 | 53 | 106 | 38 | 261 | 138 | 455 | 195 | 270 | 154 | 56 | 37 | 904 | 326 | 168 | 484 | 387 | 150 | 533 | 288 | 16 | 194 |
| PBANKA_1437800 | 162 | 274 | 91 | 10 | 413 | 104 | 718 | 154 | 934 | 302 | 441 | 109 | 1488 | 932 | 479 | 2142 | 1915 | 460 | 2027 | 1296 | 1034 | 5997 |
| PBANKA_1437900 | 20 | 20 | 20 | 0 | 59 | 25 | 72 | 30 | 105 | 76 | 29 | 11 | 164 | 53 | 68 | 108 | 326 | 132 | 279 | 87 | 121 | 750 |
| PBANKA_1438000 | 5 | 14 | 119 | 44 | 656 | 337 | 1666 | 824 | 840 | 544 | 71 | 42 | 2258 | 602 | 346 | 1239 | 12405 | 1912 | 4016 | 2896 | 324 | 2829 |
| PBANKA_1438100 | 7 | 5 | 51 | 12 | 153 | 54 | 165 | 88 | 94 | 52 | 80 | 23 | 2434 | 1127 | 432 | 2081 | 1944 | 363 | 1767 | 1691 | 196 | 1742 |
| PBANKA_1438200 | 197 | 310 | 71 | 14 | 389 | 140 | 700 | 271 | 454 | 159 | 107 | 38 | 1324 | 629 | 885 | 5285 | 9733 | 1458 | 1072 | 2516 | 168 | 908 |
| PBANKA_1438300 | 248 | 298 | 287 | 90 | 1022 | 336 | 1523 | 512 | 1023 | 396 | 214 | 64 | 7974 | 5793 | 3918 | 20959 | 25443 | 4178 | 3008 | 7070 | 358 | 2532 |
| PBANKA_1438400 | 73 | 109 | 443 | 93 | 1018 | 316 | 1561 | 501 | 1085 | 391 | 764 | 298 | 2389 | 3640 | 3148 | 13704 | 15764 | 2116 | 3146 | 13229 | 527 | 3319 |
| PBANKA_1438500 | 203 | 256 | 97 | 20 | 262 | 125 | 373 | 140 | 147 | 78 | 32 | 0 | 510 | 205 | 889 | 3962 | 2211 | 337 | 1023 | 763 | 601 | 3461 |
| PBANKA_1438600 | 0 | 0 | 53 | 14 | 201 | 142 | 204 | 136 | 61 | 22 | 10 | 8 | 423 | 123 | 717 | 2907 | 1221 | 163 | 363 | 246 | 28 | 327 |
| PBANKA_1438700 | 0 | 0 | 42 | 8 | 67 | 42 | 71 | 15 | 24 | 11 | 0 | 5 | 527 | 241 | 150 | 687 | 44 | 15 | 179 | 95 | 34 | 255 |
| PBANKA_1438800 | 0 | 0 | 36 | 0 | 58 | 35 | 52 | 39 | 46 | 15 | 35 | 26 | 399 | 231 | 208 | 885 | 218 | 66 | 428 | 317 | 13 | 95 |
| PBANKA_1438900 | 0 | 0 | 27 | 5 | 81 | 30 | 69 | 29 | 26 | 10 | 5 | 11 | 425 | 170 | 220 | 1037 | 213 | 36 | 754 | 338 | 45 | 307 |
| PBANKA_1439100 | 420 | 680 | 78 | 13 | 158 | 66 | 310 | 112 | 219 | 104 | 40 | 16 | 2254 | 1209 | 185 | 634 | 3567 | 1130 | 56669 | 44769 | 560 | 3519 |
| PBANKA_1439200 | 689 | 1050 | 1938 | 578 | 2971 | 1821 | 3380 | 1359 | 3238 | 1805 | 45238 | 27512 | 78435 | 51835 | 19308 | 91070 | 32986 | 8104 | 25326 | 36963 | 2688 | 22845 |
| PBANKA_1439300 | 87 | 83 | 84 | 21 | 295 | 86 | 276 | 112 | 179 | 60 | 358 | 184 | 2153 | 993 | 349 | 1740 | 1302 | 246 | 6228 | 3628 | 334 | 2057 |
| PBANKA_1439400 | 118 | 211 | 178 | 43 | 855 | 509 | 1377 | 723 | 630 | 384 | 51 | 36 | 980 | 182 | 212 | 874 | 4405 | 853 | 2834 | 1428 | 201 | 1508 |
| PBANKA_1439500 | 0 | 0 | 20 | 0 | 61 | 22 | 54 | 32 | 16 | 9 | 0 | 0 | 200 | 38 | 123 | 482 | 286 | 40 | 312 | 167 | 17 | 124 |
| PBANKA_1439600 | 0 | 5 | 48 | 11 | 281 | 140 | 547 | 214 | 310 | 152 | 63 | 19 | 489 | 286 | 317 | 1434 | 5657 | 611 | 717 | 1270 | 223 | 1698 |
| PBANKA_1439700 | 0 | 0 | 68 | 9 | 111 | 41 | 157 | 47 | 57 | 20 | 17 | 0 | 115 | 29 | 162 | 541 | 632 | 104 | 782 | 427 | 261 | 1456 |
| PBANKA_1439800 | 5 | 0 | 20 | 7 | 65 | 46 | 105 | 77 | 49 | 32 | 203 | 123 | 375 | 143 | 119 | 360 | 269 | 80 | 360 | 207 | 16 | 146 |
| PBANKA_1439900 | 127 | 132 | 116 | 14 | 242 | 78 | 246 | 53 | 171 | 30 | 75 | 9 | 1884 | 1258 | 434 | 2668 | 1452 | 244 | 2039 | 1117 | 201 | 865 |
| PBANKA_1440000 | 75 | 109 | 77 | 19 | 249 | 84 | 344 | 130 | 237 | 102 | 85 | 31 | 998 | 409 | 562 | 2181 | 2016 | 378 | 1243 | 840 | 77 | 525 |
| PBANKA_1440100 | 25 | 51 | 84 | 38 | 276 | 150 | 435 | 161 | 203 | 123 | 1395 | 1114 | 2242 | 1280 | 685 | 3150 | 2280 | 377 | 1339 | 919 | 50 | 472 |
| PBANKA_1440200 | 17 | 37 | 60 | 12 | 90 | 45 | 95 | 30 | 60 | 38 | 491 | 430 | 704 | 543 | 146 | 632 | 356 | 66 | 412 | 279 | 31 | 128 |
| PBANKA_1440300 | 27 | 21 | 126 | 33 | 98 | 52 | 68 | 39 | 50 | 20 | 418 | 240 | 1826 | 1167 | 249 | 1487 | 168 | 34 | 310 | 239 | 55 | 346 |
| PBANKA_1440400 | 26 | 29 | 51 | 16 | 63 | 14 | 63 | 28 | 42 | 21 | 158 | 86 | 789 | 539 | 231 | 1110 | 142 | 56 | 235 | 126 | 156 | 951 |
| PBANKA_1440500 | 17 | 24 | 114 | 33 | 366 | 137 | 395 | 178 | 155 | 77 | 33 | 20 | 2922 | 2129 | 1105 | 6172 | 2279 | 415 | 548 | 682 | 374 | 1876 |
| PBANKA_1440600 | 16 | 36 | 121 | 18 | 345 | 134 | 401 | 101 | 171 | 57 | 110 | 48 | 2757 | 2541 | 843 | 6131 | 2909 | 328 | 1150 | 2427 | 645 | 4289 |
| PBANKA_1440700 | 13 | 36 | 198 | 68 | 683 | 436 | 938 | 353 | 289 | 101 | 63 | 21 | 2017 | 787 | 1574 | 6508 | 4288 | 542 | 2299 | 1737 | 265 | 1948 |
| PBANKA_1440800 | 6 | 5 | 193 | 28 | 465 | 148 | 407 | 112 | 116 | 35 | 20 | 10 | 507 | 202 | 355 | 1979 | 587 | 84 | 720 | 434 | 710 | 4562 |
| PBANKA_1440900 | 0 | 0 | 55 | 7 | 99 | 43 | 124 | 35 | 43 | 20 | 0 | 0 | 285 | 131 | 278 | 1072 | 414 | 62 | 393 | 247 | 16 | 82 |
| PBANKA_1441000 | 6 | 8 | 9 | 0 | 88 | 41 | 77 | 24 | 57 | 11 | 10 | 0 | 37 | 22 | 7 | 18 | 16 | 6 | 637 | 199 | 0 | 9 |
| PBANKA_1441100 | 0 | 0 | 428 | 74 | 852 | 449 | 1380 | 672 | 497 | 322 | 59 | 41 | 1916 | 325 | 2390 | 7820 | 6322 | 1157 | 3698 | 1276 | 70 | 539 |
| PBANKA_1441200 | 0 | 0 | 48 | 8 | 139 | 58 | 260 | 114 | 268 | 136 | 23 | 16 | 832 | 130 | 281 | 1073 | 2487 | 593 | 1374 | 822 | 120 | 917 |
| PBANKA_1441300 | 0 | 0 | 81 | 13 | 110 | 48 | 185 | 58 | 269 | 143 | 46 | 15 | 561 | 281 | 347 | 1515 | 2745 | 588 | 1042 | 1685 | 158 | 1209 |
| PBANKA_1441400 | 43 | 92 | 122 | 37 | 298 | 200 | 456 | 194 | 143 | 117 | 17 | 7 | 421 | 128 | 401 | 911 | 2467 | 448 | 1479 | 708 | 62 | 541 |
| PBANKA_1441500 | 81 | 82 | 176 | 50 | 248 | 113 | 297 | 72 | 161 | 84 | 1137 | 751 | 1643 | 1314 | 383 | 1893 | 833 | 117 | 647 | 543 | 445 | 2556 |
| PBANKA_1441600 | 0 | 0 | 61 | 16 | 180 | 112 | 411 | 182 | 237 | 128 | 27 | 11 | 513 | 103 | 149 | 511 | 1078 | 226 | 960 | 483 | 209 | 1404 |
| PBANKA_1441700 | 0 | 0 | 157 | 36 | 345 | 218 | 368 | 146 | 118 | 78 | 1382 | 1144 | 3513 | 1992 | 813 | 2574 | 1541 | 275 | 625 | 272 | 7 | 66 |
| PBANKA_1441900 | 0 | 0 | 255 | 71 | 179 | 88 | 107 | 42 | 57 | 15 | 78 | 58 | 1532 | 1323 | 547 | 3684 | 286 | 68 | 1173 | 1698 | 471 | 3298 |
| PBANKA_1442000 | 83 | 134 | 78 | 20 | 313 | 148 | 331 | 86 | 319 | 131 | 68 | 39 | 358 | 193 | 263 | 1356 | 5231 | 777 | 395 | 621 | 39 | 238 |
| PBANKA_1442100 | 35 | 42 | 34 | 8 | 189 | 52 | 337 | 155 | 171 | 64 | 28 | 7 | 544 | 297 | 196 | 814 | 3849 | 519 | 501 | 839 | 44 | 349 |
| PBANKA_1442200 | 1109 | 1489 | 1321 | 382 | 4979 | 2162 | 6829 | 2820 | 2942 | 1376 | 466 | 176 | 2691 | 4266 | 9050 | 40052 | 54012 | 6829 | 8515 | 24739 | 6149 | 59973 |
| PBANKA_1442300 | 85 | 112 | 131 | 25 | 356 | 123 | 525 | 236 | 334 | 173 | 62 | 19 | 947 | 403 | 178 | 638 | 7331 | 1064 | 2201 | 2006 | 1290 | 9472 |
| PBANKA_1442400 | 0 | 0 | 71 | 10 | 90 | 36 | 79 | 37 | 237 | 111 | 850 | 652 | 555 | 953 | 148 | 888 | 2511 | 615 | 258 | 346 | 12 | 88 |
| PBANKA_1442500 | 55 | 87 | 180 | 47 | 340 | 110 | 277 | 103 | 192 | 91 | 1073 | 591 | 2407 | 2660 | 523 | 4250 | 777 | 115 | 494 | 998 | 198 | 1506 |
| PBANKA_1442600 | 10 | 10 | 87 | 21 | 156 | 87 | 156 | 42 | 54 | 21 | 73 | 43 | 1003 | 553 | 293 | 1910 | 496 | 82 | 4609 | 4031 | 530 | 3566 |
| PBANKA_1442700 | 0 | 0 | 13 | 0 | 32 | 15 | 25 | 10 | 5 | 0 | 0 | 0 | 157 | 31 | 90 | 297 | 97 | 16 | 83 | 89 | 6 | 14 |
| PBANKA_1442800 | 0 | 0 | 43 | 13 | 100 | 34 | 63 | 35 | 15 | 13 | 0 | 0 | 200 | 38 | 128 | 395 | 201 | 37 | 264 | 282 | 0 | 34 |
| PBANKA_1442900 | 9 | 0 | 29 | 9 | 124 | 50 | 82 | 51 | 51 | 9 | 16 | 0 | 476 | 83 | 164 | 753 | 256 | 61 | 266 | 151 | 12 | 68 |
| PBANKA_1443000 | 328 | 499 | 110 | 32 | 564 | 306 | 990 | 447 | 456 | 304 | 100 | 30 | 1704 | 598 | 375 | 1930 | 8710 | 775 | 5177 | 4464 | 4973 | 35367 |
| PBANKA_1443100 | 5 | 19 | 168 | 33 | 259 | 121 | 320 | 106 | 100 | 60 | 159 | 120 | 2509 | 1914 | 1170 | 5805 | 2208 | 246 | 342 | 397 | 16 | 95 |
| PBANKA_1443200 | 0 | 0 | 166 | 56 | 250 | 135 | 266 | 119 | 65 | 39 | 9 | 9 | 1197 | 204 | 685 | 3515 | 1117 | 144 | 894 | 567 | 23 | 113 |
| PBANKA_1443300 | 0 | 0 | 23 | 5 | 42 | 34 | 173 | 84 | 207 | 122 | 26 | 15 | 10261 | 2657 | 73 | 134 | 94813 | 14485 | 485 | 674 | 75 | 742 |
| PBANKA_1443400 | 6 | 13 | 222 | 61 | 690 | 321 | 720 | 297 | 250 | 110 | 60 | 24 | 1172 | 770 | 849 | 3572 | 4222 | 483 | 99 | 140 | 1704 | 11032 |
| PBANKA_1443500 | 95 | 172 | 279 | 71 | 1241 | 484 | 1945 | 706 | 940 | 429 | 212 | 75 | 2812 | 5277 | 3652 | 22223 | 29645 | 3075 | 674 | 4952 | 323 | 1771 |
| PBANKA_1443600 | 57 | 50 | 51 | 7 | 130 | 61 | 107 | 42 | 50 | 19 | 6 | 7 | 1147 | 313 | 313 | 1679 | 505 | 72 | 706 | 532 | 103 | 495 |
| PBANKA_1443700 | 315 | 370 | 190 | 39 | 476 | 260 | 514 | 215 | 171 | 65 | 41 | 9 | 1519 | 503 | 543 | 2351 | 937 | 192 | 3922 | 2758 | 1340 | 6158 |
| PBANKA_1443800 | 346 | 566 | 1185 | 259 | 1260 | 431 | 986 | 322 | 713 | 295 | 2088 | 966 | 4769 | 6100 | 795 | 4872 | 4120 | 593 | 1855 | 2336 | 931 | 6079 |

| | | | | | | | | | | | | | | | | | | | | | | |
|---|---|---|---|---|---|---|---|---|---|---|---|---|---|---|---|---|---|---|---|---|---|---|
| PBANKA_1443900 | 131 | 123 | 43 | 13 | 170 | 58 | 594 | 266 | 682 | 324 | 298 | 119 | 1666 | 777 | 131 | 679 | 9926 | 2339 | 5836 | 4083 | 330 | 2066 |
| PBANKA_1444000 | 0 | 5 | 256 | 81 | 586 | 347 | 640 | 311 | 128 | 84 | 23 | 9 | 1104 | 216 | 1607 | 2619 | 664 | 227 | 318 | 102 | 19 | 105 |
| PBANKA_1444100 | 29 | 60 | 621 | 199 | 852 | 490 | 831 | 353 | 239 | 109 | 24 | 15 | 4277 | 1543 | 3452 | 16180 | 2708 | 327 | 692 | 629 | 158 | 1474 |
| PBANKA_1444200 | 0 | 6 | 267 | 54 | 384 | 164 | 471 | 160 | 164 | 68 | 40 | 6 | 1574 | 1362 | 740 | 4029 | 3853 | 434 | 1901 | 6451 | 832 | 4170 |
| PBANKA_1444300 | 0 | 0 | 17 | 5 | 53 | 38 | 103 | 37 | 25 | 14 | 0 | 0 | 60 | 21 | 37 | 59 | 91 | 51 | 227 | 113 | 11 | 69 |
| PBANKA_1444400 | 0 | 11 | 44 | 6 | 102 | 34 | 165 | 45 | 90 | 35 | 22 | 0 | 260 | 51 | 119 | 512 | 594 | 223 | 1284 | 768 | 98 | 610 |
| PBANKA_1444500 | 11 | 12 | 16 | 0 | 79 | 41 | 210 | 109 | 491 | 241 | 123 | 44 | 328 | 270 | 166 | 511 | 5207 | 1118 | 1128 | 2658 | 1122 | 7680 |
| PBANKA_1444600 | 43 | 80 | 60 | 0 | 166 | 98 | 194 | 69 | 60 | 32 | 0 | 0 | 323 | 53 | 300 | 911 | 408 | 68 | 1779 | 812 | 260 | 1279 |
| PBANKA_1444700 | 12 | 8 | 21 | 0 | 274 | 96 | 381 | 136 | 155 | 83 | 21 | 12 | 819 | 302 | 1087 | 5470 | 2829 | 430 | 723 | 1283 | 1450 | 9010 |
| PBANKA_1444900 | 0 | 0 | 140 | 46 | 340 | 152 | 290 | 92 | 132 | 60 | 20 | 9 | 863 | 195 | 773 | 3183 | 861 | 156 | 1445 | 958 | 76 | 665 |
| PBANKA_1445100 | 12 | 7 | 208 | 130 | 533 | 341 | 355 | 190 | 105 | 83 | 29 | 24 | 760 | 178 | 534 | 1418 | 302 | 81 | 481 | 231 | 156 | 815 |
| PBANKA_1445200 | 20 | 32 | 50 | 0 | 104 | 55 | 113 | 32 | 61 | 28 | 295 | 281 | 650 | 388 | 157 | 808 | 125 | 23 | 164 | 149 | 137 | 1065 |
| PBANKA_1445300 | 51 | 72 | 56 | 8 | 154 | 81 | 211 | 61 | 49 | 19 | 33 | 38 | 715 | 328 | 359 | 1353 | 377 | 98 | 769 | 524 | 105 | 982 |
| PBANKA_1445400 | 45 | 62 | 364 | 138 | 1028 | 613 | 2079 | 898 | 843 | 607 | 2095 | 1722 | 4350 | 2300 | 1481 | 5989 | 4694 | 775 | 2270 | 1220 | 131 | 1254 |
| PBANKA_1445500 | 5 | 5 | 190 | 34 | 404 | 166 | 376 | 130 | 164 | 50 | 771 | 467 | 2563 | 3400 | 957 | 6969 | 3753 | 291 | 465 | 1359 | 46 | 340 |
| PBANKA_1445600 | 11 | 6 | 162 | 34 | 209 | 108 | 167 | 49 | 137 | 63 | 336 | 249 | 935 | 1130 | 365 | 1440 | 378 | 113 | 739 | 351 | 52 | 299 |
| PBANKA_1445700 | 80 | 145 | 95 | 19 | 135 | 76 | 102 | 47 | 62 | 27 | 20 | 5 | 601 | 363 | 457 | 3163 | 468 | 159 | 5709 | 7796 | 322 | 2406 |
| PBANKA_1445800 | 0 | 0 | 76 | 29 | 97 | 73 | 106 | 50 | 288 | 357 | 62 | 59 | 440 | 141 | 294 | 1029 | 604 | 274 | 817 | 366 | 39 | 260 |
| PBANKA_1445900 | 49 | 88 | 76 | 16 | 135 | 69 | 157 | 57 | 347 | 270 | 146 | 56 | 904 | 214 | 510 | 2286 | 1083 | 405 | 5120 | 3058 | 2056 | 13770 |
| PBANKA_1446100 | 0 | 8 | 67 | 22 | 363 | 184 | 487 | 133 | 535 | 306 | 183 | 100 | 675 | 500 | 533 | 2229 | 4644 | 913 | 1970 | 3904 | 184 | 1471 |
| PBANKA_1446200 | 0 | 0 | 26 | 0 | 50 | 26 | 29 | 13 | 19 | 13 | 82 | 84 | 128 | 43 | 60 | 366 | 119 | 23 | 697 | 355 | 14 | 78 |
| PBANKA_1446300 | 69 | 95 | 48 | 11 | 139 | 51 | 200 | 73 | 87 | 58 | 50 | 9 | 1552 | 1021 | 604 | 3572 | 2800 | 379 | 1144 | 1473 | 122 | 873 |
| PBANKA_1446400 | 0 | 0 | 9 | 0 | 38 | 13 | 32 | 22 | 16 | 12 | 0 | 0 | 82 | 20 | 50 | 91 | 15 | 7 | 80 | 33 | 11 | 52 |
| PBANKA_1446500 | 82 | 82 | 326 | 86 | 2174 | 839 | 2737 | 925 | 1457 | 517 | 230 | 108 | 5581 | 1833 | 6333 | 40207 | 9855 | 930 | 2702 | 4093 | 156 | 1282 |
| PBANKA_1446600 | 97 | 82 | 69 | 6 | 153 | 49 | 148 | 40 | 195 | 70 | 62 | 22 | 975 | 486 | 406 | 1906 | 788 | 207 | 1494 | 657 | 148 | 913 |
| PBANKA_1446700 | 6 | 0 | 61 | 12 | 140 | 49 | 173 | 93 | 68 | 29 | 9 | 0 | 942 | 276 | 313 | 1270 | 1038 | 180 | 975 | 694 | 1060 | 6771 |
| PBANKA_1446800 | 0 | 7 | 619 | 189 | 900 | 477 | 776 | 271 | 209 | 91 | 40 | 14 | 5087 | 1729 | 4121 | 21379 | 5010 | 488 | 950 | 964 | 33 | 383 |
| PBANKA_1446900 | 24 | 16 | 210 | 39 | 845 | 404 | 838 | 295 | 291 | 110 | 47 | 26 | 890 | 167 | 811 | 2910 | 2420 | 368 | 1891 | 793 | 44 | 399 |
| PBANKA_1447000 | 15 | 25 | 70 | 22 | 231 | 141 | 229 | 89 | 59 | 41 | 6 | 6 | 154 | 26 | 160 | 479 | 441 | 143 | 1006 | 462 | 34 | 185 |
| PBANKA_1447100 | 200 | 238 | 136 | 28 | 211 | 93 | 276 | 94 | 179 | 73 | 220 | 125 | 1099 | 745 | 249 | 1488 | 1330 | 243 | 2107 | 1446 | 235 | 1463 |
| PBANKA_1447200 | 25 | 51 | 6 | 0 | 92 | 37 | 377 | 138 | 373 | 141 | 53 | 22 | 623 | 178 | 67 | 218 | 4194 | 834 | 2666 | 1632 | 308 | 2048 |
| PBANKA_1447300 | 27 | 37 | 10 | 0 | 38 | 19 | 57 | 27 | 32 | 15 | 6 | 5 | 130 | 42 | 52 | 200 | 218 | 56 | 720 | 311 | 141 | 849 |
| PBANKA_1447400 | 0 | 0 | 26 | 6 | 103 | 51 | 251 | 84 | 89 | 48 | 7 | 5 | 229 | 35 | 104 | 283 | 1184 | 122 | 524 | 293 | 5 | 91 |
| PBANKA_1447500 | 0 | 0 | 6 | 0 | 13 | 5 | 27 | 9 | 11 | 10 | 0 | 0 | 47 | 10 | 59 | 214 | 71 | 21 | 151 | 96 | 0 | 17 |
| PBANKA_1447600 | 0 | 0 | 44 | 16 | 106 | 46 | 130 | 54 | 36 | 20 | 12 | 0 | 896 | 431 | 195 | 738 | 247 | 58 | 547 | 361 | 195 | 1145 |
| PBANKA_1447700 | 5 | 20 | 175 | 55 | 404 | 251 | 382 | 153 | 144 | 63 | 32 | 19 | 587 | 236 | 584 | 2161 | 840 | 133 | 615 | 450 | 615 | 3989 |
| PBANKA_1447800 | 12 | 18 | 18 | 0 | 54 | 23 | 61 | 18 | 41 | 18 | 25 | 19 | 735 | 430 | 278 | 1674 | 869 | 180 | 622 | 455 | 353 | 1569 |
| PBANKA_1447900 | 0 | 0 | 16 | 0 | 59 | 28 | 101 | 30 | 143 | 80 | 49 | 30 | 673 | 599 | 490 | 3658 | 2803 | 554 | 4785 | 13693 | 195 | 1342 |
| PBANKA_1448000 | 0 | 0 | 341 | 95 | 449 | 197 | 482 | 163 | 158 | 66 | 37 | 10 | 1072 | 994 | 2160 | 13812 | 1521 | 263 | 3827 | 13773 | 2953 | 18169 |
| PBANKA_1448100 | 0 | 0 | 11 | 0 | 86 | 44 | 112 | 42 | 52 | 30 | 84 | 35 | 1285 | 407 | 526 | 2176 | 433 | 89 | 449 | 283 | 126 | 622 |
| PBANKA_1448200 | 365 | 583 | 867 | 225 | 2451 | 903 | 2427 | 797 | 1071 | 416 | 250 | 99 | 2622 | 3205 | 6453 | 30730 | 11354 | 1468 | 1997 | 9335 | 1498 | 8983 |
| PBANKA_1448300 | 0 | 0 | 34 | 9 | 80 | 28 | 124 | 54 | 54 | 14 | 23 | 6 | 370 | 166 | 231 | 708 | 706 | 189 | 808 | 238 | 30 | 240 |
| PBANKA_1448400 | 0 | 0 | 90 | 36 | 202 | 90 | 221 | 76 | 140 | 81 | 94 | 35 | 1193 | 450 | 493 | 2115 | 1821 | 343 | 1540 | 820 | 51 | 400 |
| PBANKA_1448500 | 16 | 23 | 101 | 19 | 173 | 63 | 341 | 127 | 397 | 151 | 330 | 130 | 4580 | 2827 | 613 | 3540 | 11384 | 2272 | 4999 | 4149 | 366 | 2413 |
| PBANKA_1448600 | 8 | 6 | 44 | 8 | 96 | 27 | 91 | 52 | 52 | 21 | 21 | 10 | 267 | 73 | 115 | 435 | 525 | 117 | 601 | 238 | 169 | 994 |
| PBANKA_1448700 | 244 | 379 | 365 | 60 | 1451 | 439 | 1824 | 596 | 1018 | 392 | 1148 | 626 | 4943 | 11438 | 5224 | 29979 | 24067 | 2933 | 1770 | 9497 | 1365 | 7474 |
| PBANKA_1448800 | 5 | 5 | 160 | 50 | 556 | 178 | 739 | 268 | 286 | 149 | 57 | 19 | 392 | 380 | 747 | 2414 | 5142 | 647 | 840 | 1860 | 1462 | 9668 |
| PBANKA_1448900 | 0 | 0 | 59 | 6 | 294 | 105 | 336 | 154 | 133 | 70 | 15 | 10 | 528 | 156 | 587 | 1900 | 1741 | 321 | 2754 | 1626 | 279 | 2303 |
| PBANKA_1449100 | 9 | 17 | 18 | 7 | 78 | 38 | 77 | 17 | 25 | 12 | 5 | 0 | 257 | 73 | 167 | 860 | 319 | 50 | 936 | 485 | 88 | 470 |
| PBANKA_1449200 | 142 | 205 | 119 | 38 | 213 | 120 | 360 | 109 | 154 | 59 | 8 | 0 | 594 | 193 | 430 | 1726 | 2916 | 360 | 1544 | 809 | 533 | 2856 |
| PBANKA_1449300 | 123 | 134 | 6 | 7 | 244 | 88 | 244 | 81 | 87 | 41 | 10 | 0 | 692 | 350 | 30 | 82 | 362 | 166 | 20720 | 16553 | 191 | 1464 |
| PBANKA_1449400 | 37 | 29 | 138 | 42 | 592 | 315 | 606 | 236 | 264 | 132 | 1078 | 721 | 3703 | 1845 | 1259 | 5892 | 2195 | 265 | 2313 | 1751 | 319 | 2822 |
| PBANKA_1449500 | 1464 | 1684 | 937 | 461 | 1373 | 1151 | 2214 | 1652 | 1534 | 1226 | 209 | 125 | 1089 | 174 | 941 | 2847 | 3636 | 839 | 3421 | 1461 | 571 | 5739 |
| PBANKA_1449700 | 182 | 139 | 21 | 0 | 71 | 156 | 465 | 240 | 387 | 191 | 64 | 38 | 144 | 55 | 90 | 203 | 286 | 155 | 876 | 503 | 259 | 1684 |
| PBANKA_1449800 | 41 | 27 | 19 | 9 | 296 | 122 | 801 | 368 | 817 | 311 | 77 | 43 | 307 | 91 | 41 | 152 | 905 | 224 | 3373 | 1255 | 160 | 1375 |
| PBANKA_1449900 | 143 | 237 | 288 | 58 | 647 | 219 | 992 | 340 | 692 | 258 | 461 | 228 | 4506 | 6222 | 1715 | 12344 | 11073 | 1236 | 3591 | 6769 | 1673 | 10802 |
| PBANKA_1450000 | 0 | 0 | 34 | 6 | 151 | 60 | 175 | 64 | 117 | 42 | 41 | 33 | 286 | 241 | 167 | 715 | 416 | 86 | 875 | 692 | 339 | 2495 |
| PBANKA_1450100 | 0 | 0 | 63 | 18 | 230 | 128 | 275 | 99 | 123 | 47 | 347 | 257 | 894 | 512 | 316 | 1442 | 524 | 82 | 925 | 442 | 319 | 2548 |
| PBANKA_1450200 | 0 | 0 | 66 | 26 | 520 | 323 | 1890 | 926 | 1205 | 740 | 104 | 58 | 2113 | 487 | 1152 | 2167 | 9623 | 1657 | 3848 | 1958 | 99 | 921 |
| PBANKA_1450300 | 0 | 0 | 594 | 181 | 1170 | 875 | 1850 | 776 | 526 | 332 | 47 | 29 | 815 | 239 | 1644 | 6681 | 4283 | 645 | 2481 | 1882 | 774 | 8787 |
| PBANKA_1450400 | 13 | 24 | 10 | 0 | 31 | 30 | 131 | 36 | 101 | 35 | 17 | 0 | 418 | 191 | 30 | 65 | 789 | 169 | 4089 | 4148 | 84 | 616 |
| PBANKA_1450500 | 9 | 8 | 103 | 27 | 218 | 126 | 374 | 182 | 203 | 105 | 39 | 17 | 1593 | 620 | 397 | 2258 | 2406 | 331 | 1467 | 1042 | 208 | 1397 |
| PBANKA_1450600 | 16 | 24 | 35 | 0 | 70 | 37 | 129 | 50 | 48 | 29 | 5 | 0 | 703 | 250 | 196 | 935 | 951 | 179 | 882 | 417 | 95 | 591 |
| PBANKA_1450700 | 22 | 51 | 208 | 54 | 505 | 262 | 553 | 238 | 214 | 95 | 43 | 44 | 2347 | 1127 | 1186 | 5011 | 2240 | 347 | 1881 | 1123 | 100 | 823 |
| PBANKA_1450800 | 189 | 214 | 18 | 7 | 95 | 51 | 89 | 42 | 66 | 35 | 11 | 5 | 158 | 76 | 51 | 284 | 136 | 43 | 1355 | 2744 | 5216 | 35076 |
| PBANKA_1450900 | 68 | 82 | 54 | 16 | 217 | 95 | 250 | 83 | 82 | 35 | 0 | 0 | 417 | 84 | 268 | 1167 | 395 | 53 | 676 | 322 | 479 | 2352 |
| PBANKA_1451000 | 20 | 33 | 18 | 6 | 163 | 67 | 279 | 143 | 117 | 50 | 19 | 7 | 651 | 218 | 185 | 883 | 1589 | 208 | 1240 | 536 | 43 | 371 |
| PBANKA_1451200 | 28 | 42 | 8 | 0 | 24 | 22 | 21 | 20 | 25 | 14 | 9 | 18 | 62 | 28 | 9 | 17 | 23 | 12 | 689 | 203 | 2086 | 18522 |
| PBANKA_1451300 | 0 | 0 | 129 | 28 | 213 | 61 | 134 | 43 | 34 | 9 | 0 | 15 | 734 | 214 | 386 | 1447 | 471 | 62 | 451 | 194 | 340 | 1938 |
| PBANKA_1451400 | 0 | 0 | 7 | 0 | 97 | 32 | 83 | 37 | 40 | 27 | 7 | 0 | 261 | 79 | 152 | 714 | 449 | 132 | 546 | 231 | 27 | 115 |
| PBANKA_1451600 | 0 | 0 | 98 | 26 | 386 | 116 | 537 | 215 | 238 | 117 | 54 | 29 | 762 | 194 | 627 | 2834 | 2485 | 414 | 2013 | 2577 | 621 | 4421 |
| PBANKA_1451700 | 102 | 89 | 142 | 18 | 285 | 79 | 248 | 45 | 214 | 94 | 72 | 36 | 1453 | 1761 | 761 | 3595 | 1428 | 349 | 1762 | 1759 | 189 | 1374 |
| PBANKA_1451800 | 318 | 437 | 247 | 49 | 461 | 134 | 516 | 158 | 254 | 93 | 881 | 369 | 2812 | 2825 | 577 | 3446 | 2767 | 424 | 2608 | 4007 | 276 | 2267 |
| PBANKA_1451900 | 106 | 66 | 13 | 0 | 108 | 50 | 127 | 47 | 40 | 27 | 17 | 0 | 173 | 74 | 36 | 187 | 119 | 16 | 1322 | 407 | 7 | 73 |
| PBANKA_1452000 | 669 | 876 | 109 | 21 | 221 | 105 | 270 | 118 | 110 | 62 | 24 | 10 | 918 | 162 | 186 | 685 | 1822 | 436 | 1770 | 768 | 114 | 795 |
| PBANKA_1452100 | 103 | 123 | 112 | 30 | 522 | 186 | 682 | 224 | 381 | 159 | 321 | 253 | 2075 | 1735 | 864 | 4088 | 7014 | 1069 | 2022 | 3078 | 1077 | 8123 |
| PBANKA_1452200 | 17 | 11 | 60 | 18 | 137 | 55 | 127 | 53 | 72 | 32 | 163 | 80 | 3254 | 1910 | 464 | 2544 | 842 | 137 | 1061 | 1302 | 376 | 2760 |
| PBANKA_1452300 | 78 | 74 | 243 | 59 | 388 | 267 | 488 | 230 | 204 | 149 | 1217 | 1060 | 3691 | 1015 | 1715 | 4083 | 1072 | 301 | 1353 | 537 | 238 | 1238 |
| PBANKA_1452400 | 7 | 0 | 18 | 7 | 29 | 25 | 37 | 21 | 11 | 5 | 10 | 11 | 133 | 52 | 66 | 252 | 125 | 41 | 66 | 55 | 9 | 79 |
| PBANKA_1452500 | 208 | 270 | 316 | 92 | 721 | 322 | 656 | 327 | 421 | 210 | 976 | 793 | 7309 | 5483 | 3293 | 17125 | 6673 | 978 | 3848 | 4233 | 869 | 8190 |
| PBANKA_1452600 | 78 | 115 | 333 | 74 | 358 | 150 | 263 | 118 | 92 | 58 | 33 | 28 | 2309 | 1115 | 1115 | 3072 | 670 | 132 | 787 | 471 | 29 | 193 |
| PBANKA_1452700 | 17 | 16 | 39 | 7 | 107 | 58 | 91 | 40 | 158 | 142 | 65 | 25 | 110 | 89 | 31 | 75 | 204 | 85 | 775 | 681 | 41 | 320 |
| PBANKA_1452800 | 361 | 544 | 161 | 33 | 393 | 136 | 434 | 131 | 256 | 117 | 426 | 289 | 2685 | 2932 | 1042 | 5839 | 6066 | 710 | 3057 | 9209 | 1004 | 5446 |
| PBANKA_1452900 | 50 | 77 | 8 | 0 | 11 | 9 | 32 | 14 | 32 | 11 | 0 | 0 | 142 | 40 | 8 | 17 | 338 | 93 | 970 | 264 | 122 | 747 |
| PBANKA_1453000 | 162 | 196 | 70 | 14 | 227 | 83 | 515 | 206 | 492 | 234 | 86 | 36 | 1346 | 833 | 40 | 173 | 10068 | 1595 | 10250 | 17314 | 200 | 1592 |
| PBANKA_1453100 | 0 | 5 | 35 | 6 | 105 | 42 | 138 | 39 | 661 | 334 | 153 | 88 | 2033 | 661 | 371 | 2086 | 1369 | 528 | 15921 | 14684 | 69 | 623 |
| PBANKA_1453200 | 0 | 0 | 148 | 43 | 388 | 189 | 670 | 326 | 979 | 673 | 246 | 151 | 2771 | 612 | 1322 | 6966 | 2329 | 463 | 3910 | 1891 | 337 | 2369 |
| PBANKA_1453300 | 24 | 30 | 271 | 59 | 644 | 349 | 893 | 430 | 599 | 302 | 76 | 42 | 4298 | 1685 | 1666 | 7470 | 8624 | 1654 | 19392 | 16780 | 291 | 2362 |
| PBANKA_1453400 | 235 | 442 | 32 | 0 | 79 | 60 | 94 | 39 | 586 | 479 | 205 | 102 | 594 | 314 | 115 | 542 | 1653 | 601 | 2722 | 2659 | 2524 | 18730 |
| PBANKA_1453500 | 73 | 136 | 95 | 14 | 295 | 100 | 270 | 84 | 307 | 176 | 61 | 42 | 151 | 65 | 285 | 1028 | 612 | 169 | 1318 | 1010 | 323 | 2675 |

| | | | | | | | | | | | | | | | | | | | | | | |
|---|---|---|---|---|---|---|---|---|---|---|---|---|---|---|---|---|---|---|---|---|---|---|
| PBANKA_1453600 | 12 | 11 | 81 | 25 | 172 | 109 | 161 | 63 | 55 | 24 | 23 | 0 | 178 | 25 | 219 | 650 | 285 | 90 | 313 | 150 | 20 | 160 |
| PBANKA_1453700 | 5 | 0 | 92 | 23 | 1424 | 518 | 2410 | 871 | 989 | 421 | 137 | 41 | 3530 | 2191 | 911 | 3179 | 42131 | 3650 | 2476 | 5040 | 115 | 965 |
| PBANKA_1453800 | 0 | 0 | 10 | 0 | 30 | 15 | 25 | 16 | 14 | 9 | 11 | 7 | 327 | 107 | 131 | 472 | 130 | 47 | 643 | 245 | 11 | 100 |
| PBANKA_1454000 | 0 | 0 | 64 | 22 | 353 | 147 | 573 | 216 | 202 | 104 | 14 | 13 | 1584 | 628 | 666 | 2959 | 5375 | 783 | 687 | 1070 | 55 | 403 |
| PBANKA_1454100 | 166 | 236 | 310 | 126 | 1345 | 817 | 2129 | 1004 | 936 | 527 | 705 | 498 | 3676 | 2041 | 1214 | 5260 | 5876 | 799 | 550 | 371 | 44 | 381 |
| PBANKA_1454200 | 7 | 16 | 1023 | 215 | 2251 | 823 | 4123 | 1650 | 2714 | 1104 | 380 | 156 | 1423 | 4824 | 1691 | 6722 | 72084 | 8875 | 664 | 16956 | 502 | 3144 |
| PBANKA_1454300 | 28 | 60 | 28 | 5 | 297 | 112 | 275 | 104 | 103 | 41 | 19 | 9 | 1078 | 488 | 650 | 2632 | 1084 | 159 | 491 | 390 | 63 | 536 |
| PBANKA_1454400 | 0 | 0 | 12 | 0 | 101 | 21 | 107 | 21 | 35 | 23 | 6 | 0 | 95 | 43 | 156 | 643 | 213 | 59 | 351 | 371 | 23 | 231 |
| PBANKA_1454500 | 9 | 0 | 21 | 13 | 184 | 94 | 294 | 100 | 117 | 40 | 71 | 40 | 632 | 218 | 276 | 1141 | 647 | 140 | 756 | 530 | 45 | 467 |
| PBANKA_1454600 | 15 | 35 | 210 | 55 | 639 | 247 | 903 | 282 | 541 | 246 | 77 | 37 | 1896 | 1571 | 983 | 5478 | 15264 | 1324 | 2712 | 5300 | 85 | 793 |
| PBANKA_1454700 | 0 | 0 | 68 | 18 | 106 | 63 | 97 | 31 | 33 | 11 | 10 | 5 | 624 | 167 | 279 | 1472 | 405 | 50 | 212 | 144 | 0 | 49 |
| PBANKA_1454800 | 5 | 5 | 137 | 33 | 250 | 95 | 228 | 93 | 150 | 70 | 32 | 0 | 521 | 693 | 381 | 1146 | 1900 | 582 | 6474 | 18830 | 378 | 3331 |
| PBANKA_1454900 | 1055 | 1757 | 81 | 19 | 478 | 181 | 751 | 256 | 297 | 110 | 67 | 36 | 482 | 170 | 115 | 715 | 1735 | 192 | 1220 | 583 | 87 | 581 |
| PBANKA_1455000 | 378 | 422 | 125 | 35 | 395 | 222 | 408 | 135 | 117 | 67 | 27 | 15 | 265 | 68 | 65 | 237 | 201 | 80 | 2282 | 758 | 30 | 143 |
| PBANKA_1455100 | 0 | 0 | 294 | 102 | 713 | 475 | 871 | 452 | 254 | 152 | 18 | 8 | 410 | 90 | 647 | 2498 | 1542 | 258 | 2162 | 1025 | 16 | 177 |
| PBANKA_1455200 | 93 | 158 | 267 | 56 | 408 | 249 | 375 | 132 | 323 | 202 | 2090 | 1162 | 2029 | 2207 | 462 | 2461 | 1278 | 238 | 407 | 513 | 275 | 2300 |
| PBANKA_1455300 | 55 | 45 | 175 | 35 | 371 | 222 | 503 | 210 | 205 | 153 | 615 | 458 | 1885 | 1236 | 703 | 3941 | 2644 | 381 | 903 | 719 | 301 | 2301 |
| PBANKA_1455400 | 876 | 1367 | 85 | 16 | 168 | 90 | 222 | 72 | 218 | 118 | 49 | 29 | 1435 | 698 | 293 | 1084 | 3061 | 796 | 1356 | 842 | 382 | 2563 |
| PBANKA_1455500 | 121 | 133 | 34 | 12 | 60 | 46 | 89 | 49 | 54 | 32 | 10 | 0 | 346 | 61 | 102 | 387 | 366 | 76 | 358 | 243 | 23 | 110 |
| PBANKA_1455600 | 627 | 872 | 252 | 68 | 514 | 269 | 722 | 323 | 426 | 221 | 82 | 41 | 3765 | 1630 | 1083 | 6145 | 4648 | 783 | 3895 | 3647 | 337 | 2212 |
| PBANKA_1455700 | 44 | 24 | 19 | 0 | 6 | 0 | 11 | 0 | 0 | 0 | 0 | 0 | 313 | 61 | 147 | 531 | 76 | 17 | 620 | 224 | 13 | 108 |
| PBANKA_1455800 | 0 | 0 | 5 | 0 | 0 | 0 | 8 | 6 | 6 | 0 | 0 | 0 | 331 | 98 | 355 | 2131 | 494 | 82 | 2299 | 2013 | 17 | 145 |
| PBANKA_1455900 | 0 | 0 | 29 | 7 | 149 | 61 | 146 | 56 | 72 | 32 | 10 | 0 | 329 | 80 | 239 | 1066 | 700 | 115 | 973 | 672 | 27 | 298 |
| PBANKA_1456000 | 59 | 55 | 96 | 16 | 144 | 74 | 160 | 71 | 43 | 30 | 13 | 5 | 576 | 199 | 274 | 1145 | 341 | 36 | 305 | 164 | 27 | 163 |
| PBANKA_1456100 | 50 | 68 | 2361 | 698 | 1594 | 933 | 1638 | 758 | 460 | 334 | 2089 | 1880 | 19303 | 8437 | 2777 | 13704 | 5780 | 682 | 2875 | 1697 | 52 | 489 |
| PBANKA_1456200 | 0 | 7 | 164 | 29 | 204 | 134 | 352 | 175 | 99 | 87 | 25 | 10 | 2065 | 600 | 963 | 3482 | 1269 | 186 | 1495 | 696 | 193 | 1997 |
| PBANKA_1456300 | 7 | 0 | 72 | 23 | 101 | 39 | 79 | 54 | 14 | 13 | 9 | 0 | 523 | 116 | 149 | 454 | 159 | 32 | 662 | 194 | 23 | 179 |
| PBANKA_1456400 | 0 | 0 | 41 | 7 | 135 | 41 | 225 | 83 | 55 | 29 | 18 | 0 | 803 | 510 | 423 | 1729 | 1378 | 199 | 478 | 301 | 27 | 256 |
| PBANKA_1456500 | 6 | 0 | 23 | 0 | 94 | 51 | 111 | 50 | 76 | 71 | 44 | 34 | 377 | 123 | 118 | 408 | 284 | 140 | 335 | 188 | 67 | 411 |
| PBANKA_1456600 | 0 | 0 | 14 | 0 | 24 | 15 | 35 | 14 | 17 | 7 | 6 | 0 | 43 | 10 | 27 | 55 | 10 | 9 | 23 | 10 | 0 | 25 |
| PBANKA_1456800 | 382 | 457 | 91 | 12 | 260 | 161 | 416 | 234 | 256 | 141 | 28 | 8 | 571 | 117 | 60 | 194 | 1581 | 449 | 1132 | 537 | 616 | 3348 |
| PBANKA_1456900 | 1194 | 1313 | 545 | 172 | 671 | 403 | 1307 | 533 | 1097 | 673 | 1516 | 860 | 2480 | 623 | 820 | 1937 | 3256 | 1683 | 3666 | 1298 | 594 | 5737 |
| PBANKA_1457100 | 0 | 0 | 75 | 22 | 202 | 128 | 189 | 86 | 103 | 72 | 22 | 0 | 588 | 168 | 266 | 1005 | 789 | 276 | 344 | 244 | 24 | 176 |
| PBANKA_1457300 | 190 | 253 | 178 | 49 | 453 | 248 | 364 | 111 | 134 | 43 | 80 | 25 | 1216 | 673 | 753 | 4344 | 720 | 100 | 578 | 598 | 80 | 590 |
| PBANKA_1457400 | 63 | 80 | 335 | 74 | 442 | 232 | 339 | 102 | 110 | 60 | 384 | 243 | 2653 | 2888 | 1277 | 6217 | 518 | 61 | 799 | 1149 | 20 | 185 |
| PBANKA_1457500 | 213 | 335 | 102 | 27 | 674 | 288 | 738 | 233 | 250 | 99 | 122 | 52 | 1088 | 720 | 501 | 2799 | 786 | 75 | 852 | 904 | 236 | 1896 |
| PBANKA_1457550 | 101 | 62 | 538 | 114 | 255 | 193 | 659 | 312 | 138 | 113 | 443 | 391 | 219 | 111 | 667 | 103 | 103 | 78 | 342 | 42 | 6 | 108 |
| PBANKA_1457600 | 10 | 0 | 214 | 61 | 463 | 210 | 496 | 179 | 210 | 86 | 295 | 127 | 2395 | 2992 | 1476 | 7711 | 3250 | 479 | 460 | 1102 | 65 | 361 |
| PBANKA_1457700 | 11 | 5 | 6 | 0 | 0 | 0 | 20 | 5 | 13 | 13 | 70 | 18 | 87 | 91 | 63 | 142 | 81 | 42 | 188 | 526 | 18028 | 1E+05 |
| PBANKA_1457800 | 210 | 344 | 51 | 12 | 99 | 82 | 290 | 124 | 172 | 92 | 14 | 19 | 674 | 286 | 380 | 1535 | 2747 | 458 | 1907 | 1095 | 102 | 935 |
| PBANKA_1457900 | 68 | 122 | 143 | 24 | 303 | 115 | 259 | 118 | 155 | 89 | 40 | 25 | 1756 | 536 | 572 | 3041 | 4385 | 813 | 2320 | 1746 | 33 | 178 |
| PBANKA_1458000 | 90 | 144 | 145 | 23 | 146 | 81 | 181 | 83 | 62 | 29 | 102 | 55 | 5218 | 2076 | 1132 | 5208 | 1497 | 227 | 972 | 788 | 98 | 537 |
| PBANKA_1458100 | 339 | 475 | 138 | 20 | 348 | 124 | 324 | 126 | 116 | 54 | 17 | 0 | 913 | 272 | 480 | 2459 | 554 | 92 | 1135 | 459 | 56 | 331 |
| PBANKA_1458200 | 47 | 52 | 30 | 6 | 102 | 57 | 156 | 53 | 63 | 35 | 11 | 9 | 303 | 123 | 115 | 588 | 647 | 134 | 1923 | 878 | 106 | 846 |
| PBANKA_1458300 | 153 | 272 | 73 | 22 | 383 | 153 | 821 | 297 | 387 | 192 | 57 | 21 | 1059 | 518 | 527 | 1467 | 8567 | 1098 | 4155 | 5536 | 791 | 5824 |
| PBANKA_1458400 | 0 | 0 | 75 | 17 | 140 | 51 | 123 | 48 | 70 | 27 | 15 | 0 | 384 | 109 | 298 | 1481 | 514 | 94 | 1677 | 684 | 66 | 438 |
| PBANKA_1458500 | 5 | 0 | 7 | 5 | 34 | 14 | 28 | 16 | 12 | 0 | 0 | 0 | 36 | 7 | 50 | 192 | 40 | 7 | 214 | 71 | 20 | 133 |
| PBANKA_1458600 | 28 | 41 | 69 | 15 | 146 | 54 | 87 | 22 | 33 | 17 | 5 | 0 | 148 | 35 | 217 | 992 | 195 | 24 | 736 | 692 | 95 | 626 |
| PBANKA_1458700 | 0 | 0 | 44 | 11 | 147 | 74 | 165 | 68 | 68 | 41 | 32 | 7 | 801 | 249 | 382 | 1926 | 1461 | 220 | 846 | 602 | 213 | 1162 |
| PBANKA_1458800 | 0 | 0 | 29 | 0 | 51 | 19 | 102 | 37 | 63 | 42 | 13 | 0 | 753 | 719 | 265 | 921 | 1214 | 273 | 3651 | 11900 | 206 | 1288 |
| PBANKA_1458900 | 0 | 0 | 38 | 14 | 142 | 62 | 160 | 45 | 108 | 61 | 113 | 77 | 1878 | 1130 | 482 | 2229 | 2344 | 378 | 3419 | 2749 | 100 | 825 |
| PBANKA_1459000 | 5 | 5 | 44 | 8 | 36 | 20 | 17 | 8 | 21 | 5 | 181 | 73 | 676 | 427 | 160 | 646 | 51 | 6 | 395 | 153 | 118 | 751 |
| PBANKA_1459100 | 0 | 7 | 13 | 0 | 72 | 29 | 64 | 26 | 30 | 13 | 5 | 0 | 121 | 30 | 15 | 41 | 163 | 29 | 1159 | 689 | 303 | 1600 |
| PBANKA_1459200 | 10 | 11 | 68 | 9 | 194 | 67 | 182 | 33 | 41 | 20 | 12 | 0 | 42 | 35 | 49 | 163 | 82 | 28 | 159 | 200 | 240 | 1346 |
| PBANKA_1459300 | 276 | 417 | 1271 | 408 | 3241 | 2367 | 8831 | 3887 | 6502 | 3893 | 1386 | 690 | 7906 | 2173 | 6373 | 18154 | 60532 | 11432 | 9123 | 4983 | 1611 | 19659 |
| PBANKA_1459400 | 11 | 13 | 112 | 28 | 224 | 97 | 244 | 61 | 340 | 205 | 151 | 76 | 1183 | 859 | 473 | 2629 | 1291 | 376 | 3575 | 2151 | 73 | 583 |
| PBANKA_1459500 | 55 | 51 | 30 | 10 | 62 | 35 | 61 | 23 | 1197 | 1004 | 262 | 159 | 3011 | 1583 | 72 | 203 | 3268 | 2043 | 69707 | 51170 | 9262 | 80468 |
| PBANKA_1459600 | 23 | 50 | 18 | 8 | 57 | 28 | 66 | 19 | 16 | 10 | 14 | 7 | 431 | 211 | 141 | 589 | 365 | 81 | 286 | 175 | 66 | 559 |
| PBANKA_1459700 | 14 | 14 | 108 | 23 | 176 | 100 | 260 | 104 | 93 | 73 | 105 | 87 | 1657 | 874 | 495 | 3122 | 3002 | 383 | 885 | 948 | 98 | 662 |
| PBANKA_1459800 | 0 | 0 | 105 | 35 | 301 | 126 | 273 | 118 | 77 | 47 | 20 | 16 | 819 | 421 | 472 | 2084 | 602 | 89 | 737 | 517 | 487 | 3185 |
| PBANKA_1459900 | 54 | 75 | 124 | 38 | 257 | 138 | 455 | 169 | 187 | 90 | 9 | 10 | 370 | 69 | 238 | 605 | 1013 | 289 | 663 | 301 | 55 | 349 |
| PBANKA_1460100 | 14 | 15 | 25 | 0 | 90 | 37 | 81 | 29 | 40 | 10 | 10 | 0 | 122 | 45 | 59 | 172 | 327 | 55 | 143 | 88 | 415 | 1825 |
| PBANKA_1460200 | 81 | 142 | 59 | 9 | 126 | 56 | 176 | 68 | 82 | 42 | 37 | 14 | 1121 | 505 | 184 | 932 | 903 | 197 | 1015 | 554 | 183 | 869 |
| PBANKA_1460300 | 0 | 0 | 11 | 0 | 108 | 57 | 191 | 82 | 78 | 35 | 8 | 9 | 123 | 49 | 102 | 440 | 754 | 117 | 537 | 340 | 64 | 296 |
| PBANKA_1460400 | 111 | 97 | 338 | 100 | 379 | 177 | 259 | 137 | 99 | 60 | 18 | 10 | 2110 | 345 | 1688 | 8483 | 1955 | 194 | 1140 | 820 | 129 | 874 |
| PBANKA_1460500 | 173 | 277 | 24 | 0 | 49 | 25 | 104 | 36 | 208 | 86 | 28 | 10 | 463 | 149 | 51 | 270 | 3326 | 673 | 2898 | 1949 | 376 | 3162 |
| PBANKA_1460600 | 58 | 99 | 24 | 0 | 87 | 24 | 111 | 26 | 190 | 131 | 180 | 98 | 155 | 47 | 28 | 135 | 836 | 362 | 841 | 661 | 332 | 1899 |
| PBANKA_1460700 | 0 | 0 | 10 | 0 | 44 | 34 | 39 | 24 | 34 | 16 | 29 | 11 | 543 | 267 | 36 | 116 | 248 | 93 | 10522 | 9109 | 118 | 814 |
| PBANKA_1460800 | 0 | 0 | 55 | 15 | 68 | 26 | 55 | 13 | 37 | 15 | 176 | 99 | 963 | 573 | 205 | 920 | 383 | 78 | 463 | 352 | 31 | 182 |
| PBANKA_1460900 | 0 | 0 | 6 | 0 | 18 | 10 | 50 | 24 | 12 | 5 | 0 | 9 | 146 | 62 | 23 | 122 | 392 | 48 | 342 | 278 | 24 | 97 |
| PBANKA_1461000 | 0 | 0 | 52 | 9 | 44 | 18 | 39 | 15 | 26 | 12 | 75 | 31 | 787 | 624 | 137 | 791 | 280 | 49 | 428 | 527 | 2394 | 15798 |
| PBANKA_1461100 | 292 | 386 | 78 | 32 | 175 | 92 | 475 | 238 | 315 | 194 | 1176 | 1033 | 1249 | 570 | 334 | 1282 | 1493 | 265 | 1527 | 884 | 797 | 5824 |
| PBANKA_1461200 | 7 | 8 | 56 | 25 | 202 | 130 | 164 | 78 | 30 | 29 | 0 | 6 | 130 | 23 | 98 | 331 | 69 | 34 | 1073 | 474 | 107 | 937 |
| PBANKA_1461300 | 31 | 62 | 20 | 0 | 41 | 17 | 21 | 10 | 20 | 9 | 6 | 0 | 1893 | 826 | 57 | 211 | 410 | 276 | 57164 | 32844 | 5327 | 45307 |
| PBANKA_1461400 | 186 | 267 | 296 | 75 | 542 | 268 | 1102 | 539 | 1143 | 768 | 459 | 308 | 5289 | 1621 | 1612 | 5912 | 7670 | 2039 | 4190 | 1860 | 1029 | 8398 |
| PBANKA_1461500 | 0 | 0 | 33 | 5 | 162 | 101 | 295 | 140 | 133 | 58 | 14 | 11 | 342 | 91 | 498 | 1872 | 1401 | 166 | 1650 | 770 | 105 | 753 |
| PBANKA_1461600 | 271 | 355 | 551 | 113 | 1213 | 343 | 1502 | 481 | 881 | 379 | 502 | 325 | 4744 | 7951 | 1876 | 12849 | 38160 | 5507 | 1310 | 7362 | 755 | 4130 |
| PBANKA_1461700 | 8 | 0 | 27 | 0 | 36 | 15 | 29 | 13 | 21 | 5 | 0 | 0 | 132 | 40 | 91 | 448 | 79 | 20 | 490 | 288 | 32 | 191 |
| PBANKA_1461800 | 6 | 21 | 259 | 75 | 895 | 528 | 1327 | 619 | 502 | 319 | 97 | 12 | 1480 | 397 | 1363 | 5849 | 5609 | 715 | 1447 | 1503 | 893 | 8453 |
| PBANKA_1461900 | 0 | 6 | 7 | 0 | 57 | 18 | 63 | 17 | 28 | 5 | 8 | 0 | 248 | 175 | 138 | 562 | 116 | 18 | 109 | 71 | 39 | 231 |
| PBANKA_1462000 | 64 | 88 | 236 | 72 | 265 | 106 | 181 | 78 | 78 | 44 | 522 | 400 | 2774 | 1646 | 458 | 2503 | 311 | 58 | 1014 | 887 | 164 | 1057 |
| PBANKA_1462200 | 135 | 185 | 86 | 15 | 252 | 116 | 245 | 102 | 81 | 39 | 159 | 98 | 3218 | 1046 | 1001 | 6348 | 1225 | 143 | 1983 | 1838 | 603 | 4363 |
| PBANKA_1462300 | 15 | 11 | 6 | 0 | 0 | 0 | 15 | 0 | 9 | 5 | 0 | 0 | 66 | 18 | 72 | 245 | 117 | 22 | 183 | 177 | 131 | 787 |
| PBANKA_1462400 | 0 | 0 | 21 | 8 | 77 | 28 | 55 | 24 | 20 | 6 | 5 | 0 | 99 | 14 | 188 | 618 | 163 | 33 | 54 | 23 | 46 | 317 |
| PBANKA_1462500 | 0 | 0 | 8 | 0 | 55 | 18 | 50 | 16 | 21 | 5 | 0 | 0 | 27 | 16 | 36 | 97 | 63 | 20 | 71 | 70 | 30 | 271 |
| PBANKA_1462600 | 8 | 5 | 133 | 20 | 1741 | 664 | 2854 | 1131 | 1374 | 598 | 185 | 55 | 21 | 6 | 20 | 78 | 57 | 15 | 85 | 110 | 35 | 329 |
| PBANKA_1462700 | 175 | 291 | 597 | 92 | 1237 | 452 | 1277 | 456 | 605 | 215 | 363 | 106 | 3104 | 3020 | 2139 | 8062 | 6427 | 948 | 1380 | 2281 | 830 | 5962 |
| PBANKA_1462800 | 23 | 23 | 68 | 13 | 166 | 63 | 177 | 55 | 95 | 36 | 194 | 153 | 948 | 519 | 319 | 1536 | 607 | 115 | 512 | 509 | 399 | 3335 |
| PBANKA_1462900 | 16 | 18 | 82 | 24 | 130 | 52 | 124 | 45 | 76 | 51 | 752 | 502 | 953 | 592 | 262 | 1216 | 439 | 86 | 814 | 703 | 589 | 4310 |
| PBANKA_1463000 | 5 | 0 | 14 | 0 | 37 | 18 | 35 | 28 | 52 | 34 | 24 | 0 | 647 | 1481 | 610 | 2729 | 6400 | 2301 | 6895 | 28645 | 7124 | 52904 |

| | | | | | | | | | | | | | | | | | | | | | | |
|---|---|---|---|---|---|---|---|---|---|---|---|---|---|---|---|---|---|---|---|---|---|---|
| PBANKA_1463100 | 0 | 7 | 57 | 9 | 111 | 51 | 212 | 92 | 222 | 97 | 47 | 27 | 1250 | 567 | 295 | 1963 | 3669 | 724 | 2794 | 2561 | 762 | 5452 |
| PBANKA_1463200 | 0 | 0 | 39 | 12 | 156 | 68 | 195 | 83 | 74 | 41 | 17 | 5 | 298 | 87 | 320 | 1098 | 1441 | 302 | 823 | 732 | 248 | 2051 |
| PBANKA_1463400 | 0 | 0 | 30 | 10 | 81 | 45 | 74 | 35 | 20 | 15 | 12 | 9 | 144 | 45 | 123 | 287 | 127 | 50 | 134 | 84 | 11 | 75 |
| PBANKA_1463500 | 24 | 32 | 94 | 14 | 256 | 114 | 336 | 100 | 101 | 49 | 144 | 79 | 1709 | 626 | 503 | 2486 | 1247 | 174 | 830 | 658 | 369 | 2558 |
| PBANKA_1463600 | 5 | 0 | 35 | 8 | 83 | 39 | 62 | 34 | 17 | 5 | 0 | 0 | 119 | 32 | 166 | 609 | 216 | 43 | 199 | 85 | 8 | 23 |
| PBANKA_1463700 | 135 | 237 | 432 | 94 | 529 | 209 | 394 | 137 | 164 | 90 | 34 | 15 | 1322 | 807 | 2597 | 14613 | 2407 | 270 | 3407 | 9106 | 1169 | 7410 |
| PBANKA_1463800 | 0 | 0 | 56 | 13 | 213 | 75 | 486 | 179 | 223 | 103 | 597 | 338 | 1063 | 945 | 224 | 1189 | 1049 | 184 | 159 | 148 | 56 | 344 |
| PBANKA_1464000 | 0 | 0 | 20 | 0 | 29 | 12 | 69 | 21 | 28 | 27 | 229 | 224 | 856 | 317 | 215 | 768 | 528 | 199 | 507 | 198 | 41 | 290 |
| PBANKA_1464200 | 13 | 12 | 82 | 24 | 196 | 138 | 258 | 84 | 76 | 28 | 27 | 8 | 148 | 23 | 166 | 431 | 415 | 65 | 494 | 181 | 16 | 143 |
| PBANKA_1464400 | 15 | 15 | 42 | 5 | 146 | 66 | 153 | 61 | 73 | 32 | 31 | 17 | 363 | 229 | 143 | 647 | 399 | 80 | 386 | 157 | 51 | 286 |
| PBANKA_1464500 | 51 | 76 | 252 | 51 | 296 | 155 | 340 | 103 | 476 | 247 | 2653 | 1629 | 3013 | 3076 | 567 | 3218 | 2830 | 737 | 1363 | 1734 | 214 | 1401 |
| PBANKA_1464600 | 0 | 0 | 37 | 9 | 115 | 36 | 117 | 54 | 70 | 43 | 944 | 538 | 1771 | 1781 | 402 | 3299 | 1129 | 145 | 868 | 1256 | 53 | 284 |
| PBANKA_1464700 | 0 | 0 | 65 | 9 | 129 | 54 | 148 | 79 | 37 | 27 | 18 | 7 | 1353 | 727 | 333 | 1303 | 1393 | 283 | 375 | 165 | 6 | 60 |
| PBANKA_1464800 | 138 | 256 | 19 | 5 | 101 | 43 | 149 | 75 | 137 | 51 | 351 | 254 | 1289 | 993 | 134 | 942 | 1158 | 167 | 1151 | 767 | 163 | 1105 |
| PBANKA_1465051 | 7606 | 10920 | 561 | 111 | 988 | 398 | 869 | 293 | 297 | 140 | 88 | 32 | 13 | 7 | 7 | 0 | 0 | 0 | 118 | 69 | 6 | 31 |
| PBANKA_1465100 | 5 | 0 | 116 | 48 | 141 | 74 | 76 | 28 | 24 | 12 | 163 | 142 | 1189 | 67 | 278 | 77 | 27 | 66 | 144 | 27 | 0 | 52 |
| PBANKA_API00011 | 76 | 89 | 14 | 10 | 60 | 28 | 186 | 71 | 183 | 63 | 296 | 134 | 248 | 40 | 56 | 100 | 442 | 44 | 346 | 214 | 0 | 38 |
| PBANKA_API00051 | 19 | 17 | 13 | 0 | 43 | 7 | 92 | 11 | 93 | 11 | 89 | 29 | 238 | 7 | 47 | 42 | 248 | 39 | 380 | 141 | 0 | 0 |
| PBANKA_API00095 | 16 | 21 | 5 | 0 | 23 | 7 | 52 | 9 | 50 | 18 | 39 | 19 | 186 | 22 | 46 | 35 | 193 | 31 | 331 | 85 | 0 | 12 |
| PBANKA_API00170 | 0 | 0 | 5 | 0 | 16 | 0 | 41 | 6 | 31 | 0 | 8 | 0 | 8 | 0 | 0 | 0 | 12 | 0 | 7 | 0 | 0 | 0 |
| PBANKA_API00280 | 0 | 0 | 8 | 0 | 16 | 5 | 33 | 8 | 38 | 11 | 14 | 9 | 49 | 11 | 17 | 30 | 57 | 0 | 30 | 12 | 0 | 0 |
| PBANKA_MIT00600 | 0 | 0 | 43 | 6 | 42 | 5 | 40 | 10 | 34 | 9 | 20 | 0 | 664 | 37 | 793 | 742 | 457 | 246 | 4607 | 263 | 0 | 14 |
| PBANKA_MIT00700 | 7 | 0 | 16 | 5 | 43 | 24 | 38 | 28 | 29 | 24 | 0 | 0 | 150 | 83 | 572 | 2049 | 796 | 147 | 1814 | 258 | 0 | 8 |
| PBANKA_MIT00800 | 0 | 0 | 36 | 0 | 49 | 8 | 41 | 8 | 30 | 5 | 16 | 5 | 38 | 7 | 656 | 162 | 92 | 151 | 237 | 61 | 0 | 10 |
| PBANKA_MIT01000 | 0 | 0 | 84 | 0 | 131 | 14 | 93 | 21 | 71 | 12 | 35 | 5 | 130 | 99 | 2874 | 746 | 590 | 295 | 3552 | 917 | 0 | 8 |
| PBANKA_MIT01100 | 0 | 0 | 101 | 15 | 424 | 161 | 375 | 147 | 372 | 198 | 47 | 21 | 425 | 98 | 281 | 1558 | 1180 | 273 | 1524 | 705 | 7 | 45 |
| PBANKA_MIT01200 | 0 | 0 | 36 | 0 | 82 | 0 | 69 | 7 | 57 | 5 | 67 | 10 | 22 | 17 | 216 | 133 | 66 | 37 | 118 | 33 | 0 | 0 |
| PBANKA_MIT01400 | 0 | 0 | 19 | 0 | 27 | 0 | 20 | 0 | 14 | 0 | 10 | 0 | 0 | 0 | 94 | 29 | 20 | 28 | 54 | 52 | 0 | 0 |
| PBANKA_MIT01600 | 0 | 0 | 5 | 0 | 6 | 0 | 0 | 0 | 0 | 0 | 0 | 0 | 77 | 12 | 157 | 120 | 83 | 104 | 1119 | 51 | 0 | 0 |
| PBANKA_MIT01700 | 7 | 7 | 88 | 0 | 141 | 15 | 156 | 19 | 133 | 11 | 78 | 14 | 433 | 315 | 1926 | 1546 | 1414 | 589 | 5563 | 1657 | 40 | 266 |
| PBANKA_MIT01800 | 12 | 17 | 456 | 46 | 1281 | 380 | 1459 | 341 | 794 | 266 | 189 | 87 | 433 | 124 | 335 | 2644 | 2241 | 337 | 1774 | 1120 | 57 | 629 |
| PBANKA_MIT01900 | 11 | 10 | 279 | 68 | 972 | 411 | 1137 | 519 | 719 | 340 | 139 | 66 | 699 | 324 | 476 | 6143 | 10791 | 1037 | 4355 | 2735 | 44 | 401 |
| PBANKA_MIT02200 | 0 | 0 | 38 | 0 | 50 | 7 | 62 | 12 | 43 | 0 | 25 | 6 | 47 | 49 | 422 | 151 | 177 | 93 | 349 | 143 | 0 | 0 |
| PBANKA_MIT02700 | 33 | 23 | 223 | 30 | 402 | 170 | 322 | 182 | 250 | 104 | 109 | 85 | 264 | 205 | 597 | 1969 | 1463 | 338 | 2262 | 830 | 0 | 22 |
| PBANKA_MIT02800 | 0 | 0 | 26 | 0 | 50 | 7 | 34 | 5 | 20 | 0 | 10 | 0 | 109 | 55 | 690 | 155 | 166 | 84 | 8517 | 213 | 8 | 48 |
| PBANKA_MIT03100 | 0 | 0 | 12 | 0 | 15 | 0 | 5 | 0 | 0 | 0 | 0 | 0 | 10 | 6 | 341 | 113 | 43 | 57 | 420 | 24 | 5 | 26 |
| PBANKA_MIT03200 | 0 | 0 | 5 | 0 | 0 | 0 | 6 | 0 | 6 | 0 | 0 | 0 | 5 | 0 | 69 | 10 | 0 | 15 | 85 | 45 | 6 | 25 |
| PBANKA_MIT03500 | 1379 | 1372 | 3132 | 779 | 8677 | 4872 | 10698 | 4482 | 6985 | 3248 | 8509 | 4662 | 1044 | 838 | 1391 | 4648 | 3729 | 639 | 1934 | 2252 | 15 | 105 |
| PBANKA_MIT03600 | 9 | 11 | 46 | 8 | 67 | 21 | 50 | 16 | 39 | 7 | 31 | 22 | 130 | 18 | 474 | 198 | 160 | 99 | 2616 | 95 | 0 | 6 |
| PBANKA_MIT03700 | 0 | 0 | 10 | 0 | 20 | 0 | 5 | 0 | 16 | 0 | 0 | 0 | 78 | 120 | 977 | 271 | 320 | 416 | 6605 | 152 | 0 | 0 |
| PBANKA_0100041 | 112 | 46 | 0 | 7 | 30 | 6 | 93 | 37 | 108 | 57 | 6 | 18 | 61 | 233 | 174 | 1040 | 997 | 86 | 261 | 846 | 423 | 3940 |
| PBANKA_0300600 | 5 | 6 | 0 | 7 | 27 | 18 | 67 | 46 | 718 | 484 | 17626 | 10996 | 11370 | 10659 | 2582 | 10932 | 3561 | 696 | 2720 | 1981 | 179 | 1375 |
| PBANKA_0316600 | 8 | 0 | 0 | 6 | 268 | 142 | 1183 | 524 | 1327 | 833 | 179 | 96 | 692 | 1059 | 412 | 1184 | 21371 | 2787 | 133 | 573 | 47 | 193 |
| PBANKA_0418500 | 0 | 0 | 0 | 5 | 24 | 19 | 40 | 14 | 1058 | 872 | 8649 | 6035 | 901 | 1127 | 95 | 184 | 1477 | 1205 | 138 | 66 | 0 | 33 |
| PBANKA_0519100 | 9 | 10 | 0 | 5 | 153 | 57 | 3693 | 1892 | 7325 | 5060 | 1359 | 842 | 6273 | 2507 | 15 | 64 | 69980 | 8211 | 650 | 1249 | 9 | 123 |
| PBANKA_0523700 | 323 | 393 | 0 | 6 | 49 | 16 | 552 | 203 | 2436 | 1321 | 283 | 153 | 5682 | 2106 | 48 | 47 | 43244 | 13003 | 2421 | 2317 | 2217 | 12674 |
| PBANKA_0918600 | 21 | 42 | 0 | 8 | 205 | 65 | 286 | 114 | 168 | 60 | 22 | 14 | 777 | 302 | 89 | 372 | 1431 | 251 | 7698 | 7767 | 118 | 868 |
| PBANKA_1116000 | 3239 | 4707 | 0 | 5 | 90 | 24 | 77 | 34 | 83 | 44 | 181 | 119 | 240 | 223 | 105 | 293 | 187 | 46 | 515 | 553 | 183 | 1318 |
| PBANKA_1201400 | 0 | 0 | 0 | 5 | 157 | 67 | 247 | 77 | 410 | 244 | 54 | 59 | 855 | 402 | 241 | 1304 | 2714 | 542 | 1129 | 974 | 42 | 259 |
| PBANKA_1419900 | 8 | 7 | 0 | 5 | 77 | 43 | 187 | 71 | 66 | 34 | 17 | 0 | 87 | 27 | 10 | 41 | 566 | 83 | 281 | 200 | 48 | 443 |
| PBANKA_0006300 | 0 | 0 | 0 | 0 | 5 | 0 | 0 | 0 | 6 | 0 | 0 | 14 | 21 | 6 | 0 | 0 | 0 | 0 | 0 | 0 | 0 | 0 |
| PBANKA_0100100 | 0 | 0 | 0 | 0 | 10 | 6 | 11 | 6 | 0 | 0 | 0 | 0 | 48 | 0 | 56 | 186 | 8 | 6 | 86 | 46 | 0 | 8 |
| PBANKA_0100300 | 0 | 0 | 0 | 0 | 6 | 0 | 25 | 17 | 0 | 5 | 128 | 133 | 479 | 256 | 73 | 236 | 89 | 44 | 144 | 63 | 0 | 11 |
| PBANKA_0100400 | 0 | 0 | 0 | 0 | 32 | 14 | 58 | 34 | 1084 | 742 | 347 | 184 | 645 | 459 | 95 | 177 | 2297 | 1781 | 239 | 336 | 0 | 21 |
| PBANKA_0100500 | 0 | 0 | 0 | 0 | 19 | 9 | 73 | 28 | 131 | 109 | 1242 | 1132 | 1807 | 832 | 467 | 1705 | 1068 | 244 | 860 | 487 | 7 | 39 |
| PBANKA_0100600 | 0 | 0 | 0 | 0 | 28 | 19 | 206 | 131 | 275 | 162 | 283 | 213 | 2639 | 580 | 407 | 1142 | 8052 | 2638 | 777 | 350 | 0 | 24 |
| PBANKA_0102700 | 0 | 0 | 0 | 0 | 14 | 8 | 31 | 11 | 8 | 10 | 0 | 0 | 349 | 124 | 40 | 288 | 234 | 81 | 2438 | 2314 | 30 | 198 |
| PBANKA_0102900 | 0 | 0 | 0 | 0 | 53 | 16 | 998 | 414 | 1217 | 600 | 119 | 91 | 1166 | 904 | 39 | 157 | 29040 | 3830 | 5433 | 6975 | 181 | 1614 |
| PBANKA_0104300 | 0 | 0 | 0 | 0 | 32 | 16 | 65 | 17 | 20 | 10 | 11 | 0 | 156 | 68 | 150 | 646 | 465 | 62 | 409 | 151 | 20 | 121 |
| PBANKA_0104400 | 0 | 0 | 0 | 0 | 35 | 14 | 44 | 17 | 70 | 38 | 22 | 7 | 510 | 249 | 35 | 154 | 1625 | 285 | 4884 | 3743 | 39 | 254 |
| PBANKA_0104500 | 10 | 12 | 0 | 0 | 27 | 7 | 24 | 11 | 20 | 7 | 0 | 0 | 543 | 210 | 9 | 51 | 228 | 123 | 16261 | 11482 | 2036 | 14393 |
| PBANKA_0108100 | 159 | 198 | 0 | 0 | 60 | 16 | 270 | 107 | 816 | 483 | 155 | 64 | 260 | 76 | 48 | 191 | 452 | 109 | 1399 | 955 | 132 | 1039 |
| PBANKA_0108300 | 0 | 0 | 0 | 0 | 28 | 22 | 270 | 167 | 361 | 213 | 34 | 17 | 570 | 113 | 14 | 47 | 3278 | 863 | 7988 | 3649 | 250 | 2092 |
| PBANKA_0109900 | 5 | 0 | 0 | 0 | 43 | 29 | 394 | 186 | 515 | 262 | 107 | 65 | 2409 | 1059 | 58 | 230 | 7747 | 1715 | 28886 | 27734 | 149 | 1025 |
| PBANKA_0110300 | 0 | 0 | 0 | 0 | 64 | 16 | 220 | 91 | 38 | 44 | 6 | 7 | 66 | 0 | 0 | 0 | 139 | 57 | 10 | 0 | 0 | 5 |
| PBANKA_0112641 | 0 | 0 | 0 | 0 | 16 | 7 | 20 | 13 | 15 | 15 | 82 | 50 | 976 | 686 | 159 | 494 | 278 | 47 | 408 | 285 | 17 | 125 |
| PBANKA_0112661 | 0 | 0 | 0 | 0 | 5 | 0 | 0 | 0 | 90 | 81 | 603 | 482 | 286 | 289 | 53 | 88 | 256 | 193 | 67 | 74 | 0 | 15 |
| PBANKA_0112721 | 0 | 0 | 0 | 0 | 9 | 0 | 0 | 0 | 51 | 44 | 926 | 510 | 753 | 588 | 304 | 727 | 94 | 82 | 619 | 533 | 16 | 69 |
| PBANKA_0200500 | 5 | 0 | 0 | 0 | 12 | 7 | 16 | 12 | 18 | 0 | 0 | 0 | 70 | 165 | 30 | 166 | 94 | 25 | 32 | 14 | 0 | 6 |
| PBANKA_0200650 | 0 | 0 | 0 | 0 | 14 | 0 | 18 | 9 | 0 | 8 | 0 | 0 | 29 | 26 | 27 | 52 | 31 | 16 | 9 | 0 | 0 | 0 |
| PBANKA_0201000 | 0 | 0 | 0 | 0 | 9 | 0 | 7 | 0 | 0 | 0 | 0 | 0 | 1605 | 397 | 280 | 1129 | 217 | 32 | 126 | 58 | 12 | 59 |
| PBANKA_0201250 | 0 | 0 | 0 | 0 | 57 | 47 | 93 | 65 | 143 | 108 | 171 | 130 | 456 | 178 | 183 | 467 | 454 | 256 | 65 | 54 | 6 | 37 |
| PBANKA_0201300 | 0 | 0 | 0 | 0 | 83 | 36 | 47 | 24 | 235 | 190 | 556 | 604 | 571 | 349 | 152 | 678 | 941 | 378 | 210 | 161 | 19 | 202 |
| PBANKA_0206700 | 0 | 0 | 0 | 0 | 18 | 0 | 34 | 20 | 20 | 11 | 16 | 0 | 297 | 127 | 42 | 256 | 225 | 47 | 896 | 822 | 8 | 94 |
| PBANKA_0207100 | 9 | 16 | 0 | 0 | 69 | 18 | 95 | 50 | 61 | 36 | 5 | 0 | 200 | 53 | 17 | 46 | 706 | 205 | 1465 | 561 | 14 | 109 |
| PBANKA_0207900 | 0 | 0 | 0 | 0 | 36 | 16 | 96 | 27 | 87 | 78 | 14 | 5 | 139 | 15 | 6 | 18 | 867 | 415 | 299 | 143 | 70 | 396 |
| PBANKA_0208900 | 0 | 0 | 0 | 0 | 662 | 269 | 3561 | 1616 | 3482 | 1931 | 343 | 159 | 843 | 411 | 17 | 58 | 3333 | 1822 | 495 | 538 | 653 | 3560 |
| PBANKA_0209000 | 433 | 699 | 0 | 0 | 48 | 29 | 277 | 131 | 386 | 148 | 41 | 16 | 471 | 142 | 15 | 23 | 3876 | 759 | 468 | 675 | 238 | 1553 |
| PBANKA_0210300 | 0 | 0 | 0 | 0 | 60 | 33 | 96 | 30 | 46 | 15 | 7 | 0 | 327 | 55 | 46 | 210 | 1004 | 135 | 1395 | 1048 | 194 | 1192 |
| PBANKA_0212600 | 0 | 7 | 0 | 0 | 13 | 0 | 20 | 8 | 18 | 6 | 0 | 0 | 60 | 0 | 7 | 24 | 20 | 10 | 806 | 240 | 13 | 126 |
| PBANKA_0213900 | 0 | 0 | 0 | 0 | 12 | 0 | 8 | 7 | 16 | 15 | 0 | 0 | 112 | 10 | 5 | 7 | 137 | 81 | 855 | 181 | 6 | 85 |
| PBANKA_0214000 | 0 | 0 | 0 | 0 | 331 | 151 | 737 | 304 | 475 | 184 | 175 | 60 | 167 | 351 | 79 | 203 | 1394 | 320 | 2252 | 8387 | 1044 | 8765 |
| PBANKA_0214550 | 0 | 0 | 0 | 0 | 8 | 5 | 17 | 16 | 5 | 5 | 0 | 0 | 649 | 205 | 485 | 1710 | 674 | 111 | 237 | 98 | 0 | 11 |
| PBANKA_0214600 | 0 | 0 | 0 | 0 | 123 | 82 | 215 | 104 | 895 | 529 | 8110 | 5390 | 4055 | 2682 | 796 | 3485 | 1403 | 510 | 448 | 352 | 12 | 93 |
| PBANKA_0215800 | 0 | 0 | 0 | 0 | 22 | 13 | 32 | 17 | 12 | 8 | 6 | 0 | 1067 | 273 | 254 | 1572 | 538 | 81 | 44 | 36 | 0 | 7 |
| PBANKA_0216000 | 0 | 0 | 0 | 0 | 62 | 12 | 112 | 56 | 62 | 38 | 10 | 0 | 252 | 9 | 67 | 0 | 15 | 223 | 132 | 8 | 30 | 195 |
| PBANKA_0216721 | 0 | 0 | 0 | 0 | 9 | 0 | 8 | 0 | 0 | 0 | 12 | 6 | 181 | 57 | 220 | 191 | 51 | 20 | 185 | 58 | 0 | 17 |
| PBANKA_0216741 | 0 | 0 | 0 | 0 | 5 | 0 | 10 | 0 | 6 | 5 | 89 | 66 | 422 | 288 | 83 | 214 | 114 | 33 | 137 | 51 | 0 | 0 |
| PBANKA_0216761 | 0 | 0 | 0 | 0 | 5 | 6 | 0 | 0 | 184 | 132 | 74 | 46 | 354 | 281 | 35 | 64 | 2133 | 940 | 131 | 153 | 0 | 22 |
| PBANKA_0216781 | 0 | 5 | 0 | 0 | 65 | 53 | 298 | 150 | 226 | 110 | 45 | 16 | 276 | 81 | 12 | 20 | 1669 | 510 | 101 | 85 | 0 | 26 |

| | | | | | | | | | | | | | | | | | | | | | | |
|---|---|---|---|---|---|---|---|---|---|---|---|---|---|---|---|---|---|---|---|---|---|---|
| PBANKA_0216801 | 6 | 0 | 0 | 0 | 9 | 0 | 5 | 5 | 13 | 8 | 0 | 0 | 380 | 357 | 75 | 394 | 1015 | 472 | 76 | 161 | 28 | 227 |
| PBANKA_0300100 | 0 | 0 | 0 | 0 | 89 | 72 | 262 | 92 | 74 | 53 | 37 | 13 | 61 | 23 | 28 | 20 | 90 | 94 | 74 | 25 | 10 | 47 |
| PBANKA_0302200 | 0 | 0 | 0 | 0 | 5 | 0 | 18 | 11 | 10 | 0 | 0 | 0 | 21 | 11 | 21 | 73 | 54 | 19 | 233 | 115 | 131 | 702 |
| PBANKA_0310700 | 0 | 6 | 0 | 0 | 46 | 16 | 31 | 13 | 20 | 12 | 14 | 0 | 433 | 260 | 28 | 107 | 243 | 117 | 10374 | 10195 | 183 | 1184 |
| PBANKA_0311300 | 15 | 12 | 0 | 0 | 54 | 29 | 33 | 21 | 34 | 7 | 38 | 25 | 203 | 108 | 21 | 126 | 69 | 16 | 191 | 92 | 384 | 1449 |
| PBANKA_0312000 | 8 | 11 | 0 | 0 | 10 | 7 | 15 | 7 | 0 | 8 | 0 | 0 | 468 | 174 | 6 | 29 | 147 | 57 | 10953 | 6039 | 3069 | 21428 |
| PBANKA_0312600 | 6 | 0 | 0 | 0 | 44 | 11 | 117 | 52 | 100 | 66 | 11 | 14 | 85 | 11 | 16 | 27 | 457 | 152 | 238 | 109 | 14 | 122 |
| PBANKA_0313500 | 0 | 5 | 0 | 0 | 31 | 17 | 95 | 49 | 128 | 50 | 17 | 12 | 460 | 103 | 11 | 13 | 2453 | 499 | 849 | 368 | 556 | 3456 |
| PBANKA_0313600 | 0 | 0 | 0 | 0 | 10 | 0 | 51 | 35 | 66 | 28 | 6 | 5 | 182 | 90 | 0 | 6 | 876 | 144 | 1042 | 865 | 216 | 1456 |
| PBANKA_0316300 | 0 | 0 | 0 | 0 | 12 | 5 | 37 | 15 | 92 | 62 | 1368 | 1190 | 1930 | 1357 | 206 | 599 | 2327 | 576 | 661 | 432 | 100 | 620 |
| PBANKA_0316800 | 0 | 0 | 0 | 0 | 53 | 50 | 65 | 31 | 42 | 33 | 21 | 16 | 4110 | 2422 | 1612 | 6006 | 1298 | 168 | 264 | 267 | 33 | 287 |
| PBANKA_0316961 | 17 | 12 | 0 | 0 | 15 | 16 | 78 | 48 | 84 | 26 | 9 | 0 | 67 | 15 | 7 | 12 | 308 | 94 | 9 | 5 | 0 | 0 |
| PBANKA_0400500 | 0 | 0 | 0 | 0 | 160 | 95 | 1492 | 708 | 1284 | 696 | 176 | 83 | 58 | 24 | 22 | 57 | 123 | 27 | 809 | 325 | 11 | 74 |
| PBANKA_0400600 | 0 | 0 | 0 | 0 | 28 | 24 | 67 | 18 | 30 | 24 | 23 | 10 | 45 | 46 | 10 | 39 | 8 | 5 | 439 | 192 | 0 | 6 |
| PBANKA_0400700 | 0 | 0 | 0 | 0 | 6 | 0 | 8 | 0 | 5 | 5 | 0 | 0 | 9 | 0 | 10 | 8 | 0 | 25 | 27 | 13 | 0 | 8 |
| PBANKA_0400800 | 0 | 0 | 0 | 0 | 35 | 21 | 36 | 12 | 6 | 5 | 0 | 0 | 14 | 0 | 20 | 25 | 13 | 0 | 20 | 0 | 0 | 0 |
| PBANKA_0402000 | 0 | 0 | 0 | 0 | 54 | 34 | 178 | 60 | 81 | 35 | 13 | 0 | 831 | 191 | 120 | 594 | 1208 | 229 | 1967 | 876 | 56 | 347 |
| PBANKA_0402500 | 0 | 8 | 0 | 0 | 30 | 17 | 50 | 21 | 61 | 46 | 7 | 7 | 592 | 126 | 16 | 14 | 470 | 250 | 13450 | 4406 | 109 | 808 |
| PBANKA_0402700 | 0 | 8 | 0 | 0 | 17 | 11 | 57 | 15 | 567 | 544 | 169 | 84 | 3705 | 2173 | 48 | 190 | 1867 | 969 | 1E+05 | 82615 | 2295 | 20675 |
| PBANKA_0406400 | 0 | 0 | 0 | 0 | 52 | 20 | 547 | 217 | 766 | 392 | 76 | 58 | 349 | 310 | 59 | 146 | 8523 | 1395 | 381 | 1026 | 929 | 5639 |
| PBANKA_0407300 | 6 | 9 | 0 | 0 | 7 | 0 | 14 | 6 | 9 | 7 | 0 | 0 | 90 | 25 | 5 | 6 | 86 | 48 | 1649 | 707 | 47 | 293 |
| PBANKA_0408200 | 0 | 0 | 0 | 0 | 13 | 0 | 10 | 5 | 13 | 6 | 9 | 0 | 208 | 99 | 18 | 72 | 103 | 23 | 2009 | 2009 | 15830 | 1E+05 |
| PBANKA_0409700 | 0 | 6 | 0 | 0 | 14 | 5 | 25 | 21 | 63 | 35 | 13 | 10 | 342 | 128 | 19 | 52 | 167 | 81 | 6349 | 3761 | 586 | 4165 |
| PBANKA_0410650 | 0 | 0 | 0 | 0 | 13 | 0 | 13 | 11 | 16 | 8 | 6 | 8 | 358 | 268 | 25 | 154 | 82 | 18 | 172 | 187 | 3613 | 29742 |
| PBANKA_0413400 | 0 | 0 | 0 | 0 | 71 | 48 | 51 | 17 | 34 | 6 | 5 | 0 | 119 | 50 | 59 | 203 | 53 | 17 | 1683 | 757 | 33 | 270 |
| PBANKA_0415800 | 0 | 0 | 0 | 0 | 29 | 8 | 47 | 15 | 28 | 18 | 7 | 0 | 216 | 57 | 18 | 48 | 302 | 57 | 1963 | 1462 | 127 | 825 |
| PBANKA_0417100 | 60 | 94 | 0 | 0 | 30 | 17 | 24 | 17 | 40 | 22 | 11 | 7 | 203 | 121 | 71 | 172 | 149 | 49 | 1017 | 1513 | 67 | 419 |
| PBANKA_0417200 | 0 | 0 | 0 | 0 | 13 | 11 | 0 | 10 | 19 | 6 | 0 | 0 | 2141 | 750 | 33 | 81 | 612 | 399 | 56751 | 29808 | 668 | 6558 |
| PBANKA_0417600 | 0 | 0 | 0 | 0 | 85 | 45 | 95 | 31 | 50 | 23 | 13 | 0 | 854 | 621 | 45 | 100 | 630 | 246 | 22514 | 22448 | 103 | 1009 |
| PBANKA_0500781 | 0 | 0 | 0 | 0 | 5 | 0 | 0 | 6 | 10 | 9 | 0 | 0 | 1116 | 572 | 413 | 1454 | 7687 | 922 | 244 | 270 | 5 | 33 |
| PBANKA_0501100 | 0 | 0 | 0 | 0 | 46 | 26 | 64 | 33 | 25 | 26 | 34 | 22 | 1649 | 547 | 1415 | 5338 | 1413 | 341 | 2624 | 1157 | 42 | 473 |
| PBANKA_0501400 | 0 | 0 | 0 | 0 | 13 | 5 | 257 | 72 | 1213 | 805 | 167 | 63 | 1044 | 302 | 20 | 111 | 10986 | 3380 | 102 | 213 | 18 | 147 |
| PBANKA_0503400 | 0 | 0 | 0 | 0 | 5 | 0 | 16 | 0 | 39 | 23 | 6 | 9 | 66 | 25 | 9 | 25 | 249 | 81 | 418 | 182 | 23 | 115 |
| PBANKA_0504400 | 0 | 0 | 0 | 0 | 8 | 0 | 10 | 6 | 10 | 0 | 0 | 0 | 2393 | 2347 | 54 | 194 | 1647 | 733 | 34605 | 77410 | 2532 | 14539 |
| PBANKA_0507900 | 0 | 0 | 0 | 0 | 7 | 0 | 0 | 0 | 0 | 0 | 0 | 0 | 104 | 37 | 7 | 22 | 169 | 21 | 1320 | 640 | 7 | 39 |
| PBANKA_0508000 | 0 | 0 | 0 | 0 | 26 | 10 | 32 | 13 | 39 | 11 | 9 | 0 | 11 | 9 | 31 | 41 | 51 | 21 | 17 | 13 | 5 | 7 |
| PBANKA_0512000 | 0 | 0 | 0 | 0 | 10 | 7 | 16 | 0 | 7 | 0 | 0 | 0 | 438 | 163 | 14 | 41 | 137 | 92 | 6911 | 3316 | 164 | 970 |
| PBANKA_0513000 | 250 | 363 | 0 | 0 | 59 | 30 | 98 | 50 | 211 | 118 | 33 | 28 | 97 | 50 | 7 | 24 | 801 | 263 | 720 | 477 | 3939 | 28953 |
| PBANKA_0513700 | 1242 | 1607 | 0 | 0 | 7 | 0 | 31 | 11 | 73 | 49 | 19 | 0 | 53 | 14 | 33 | 70 | 173 | 37 | 108 | 91 | 551 | 3657 |
| PBANKA_0514900 | 8 | 8 | 0 | 0 | 6 | 0 | 40 | 28 | 27 | 33 | 6 | 10 | 12415 | 5298 | 237 | 472 | 2870 | 2307 | 4E+05 | 2E+05 | 36073 | 4E+05 |
| PBANKA_0515600 | 0 | 0 | 0 | 0 | 82 | 51 | 153 | 75 | 67 | 40 | 7 | 8 | 134 | 40 | 43 | 51 | 563 | 123 | 55 | 21 | 7 | 64 |
| PBANKA_0517600 | 0 | 0 | 0 | 0 | 5 | 8 | 9 | 5 | 6 | 6 | 0 | 0 | 1693 | 525 | 33 | 105 | 419 | 219 | 43650 | 23179 | 370 | 2780 |
| PBANKA_0518100 | 7 | 11 | 0 | 0 | 69 | 28 | 84 | 29 | 47 | 37 | 0 | 7 | 38 | 26 | 32 | 105 | 98 | 13 | 233 | 321 | 4313 | 26107 |
| PBANKA_0519200 | 6 | 0 | 0 | 0 | 78 | 75 | 452 | 211 | 3067 | 2518 | 623 | 354 | 1776 | 1378 | 11 | 29 | 13985 | 2837 | 516 | 815 | 59 | 662 |
| PBANKA_0519400 | 7 | 0 | 0 | 0 | 505 | 328 | 4259 | 2430 | 4997 | 3106 | 481 | 263 | 7372 | 2664 | 23 | 75 | 78234 | 7927 | 1509 | 2209 | 87 | 910 |
| PBANKA_0521200 | 0 | 0 | 0 | 0 | 33 | 11 | 38 | 12 | 23 | 6 | 7 | 0 | 173 | 34 | 20 | 62 | 116 | 74 | 1689 | 740 | 41 | 383 |
| PBANKA_0522100 | 48 | 46 | 0 | 0 | 20 | 10 | 24 | 17 | 32 | 23 | 20 | 22 | 267 | 255 | 44 | 88 | 573 | 241 | 1609 | 3807 | 203 | 1271 |
| PBANKA_0524100 | 0 | 0 | 0 | 0 | 17 | 5 | 22 | 19 | 102 | 115 | 641 | 477 | 1789 | 914 | 218 | 966 | 227 | 80 | 251 | 155 | 14 | 76 |
| PBANKA_0524200 | 8 | 0 | 0 | 0 | 74 | 49 | 209 | 106 | 524 | 427 | 3820 | 2540 | 4877 | 2728 | 2300 | 5254 | 4709 | 2063 | 3528 | 1329 | 94 | 794 |
| PBANKA_0524821 | 0 | 0 | 0 | 0 | 26 | 20 | 33 | 9 | 14 | 10 | 56 | 56 | 249 | 203 | 67 | 70 | 44 | 36 | 40 | 6 | 0 | 0 |
| PBANKA_0600041 | 0 | 0 | 0 | 0 | 12 | 0 | 8 | 0 | 30 | 9 | 213 | 158 | 321 | 221 | 63 | 227 | 80 | 87 | 81 | 72 | 0 | 18 |
| PBANKA_0600051 | 0 | 0 | 0 | 0 | 6 | 0 | 6 | 0 | 0 | 0 | 59 | 61 | 468 | 120 | 140 | 614 | 111 | 27 | 1497 | 1139 | 12 | 79 |
| PBANKA_0600351 | 0 | 0 | 0 | 0 | 27 | 12 | 27 | 0 | 136 | 49 | 47 | 15 | 193 | 1504 | 105 | 745 | 6820 | 739 | 132 | 816 | 30 | 213 |
| PBANKA_0600700 | 11 | 11 | 0 | 0 | 7 | 7 | 7 | 0 | 15 | 0 | 0 | 0 | 157 | 54 | 13 | 65 | 105 | 30 | 1916 | 1033 | 63 | 470 |
| PBANKA_0603650 | 0 | 0 | 0 | 0 | 6 | 5 | 10 | 0 | 106 | 117 | 27 | 11 | 141 | 66 | 0 | 18 | 418 | 260 | 516 | 287 | 12 | 74 |
| PBANKA_0603900 | 0 | 0 | 0 | 0 | 5 | 0 | 30 | 12 | 87 | 64 | 18 | 9 | 291 | 68 | 5 | 14 | 2197 | 575 | 160 | 177 | 2997 | 17308 |
| PBANKA_0605100 | 61 | 77 | 0 | 0 | 44 | 22 | 11 | 12 | 36 | 40 | 10 | 6 | 510 | 517 | 32 | 140 | 984 | 235 | 2911 | 13738 | 294 | 2054 |
| PBANKA_0605500 | 490 | 930 | 0 | 0 | 7 | 0 | 22 | 18 | 25 | 5 | 10 | 0 | 274 | 138 | 37 | 178 | 578 | 138 | 1524 | 1948 | 79 | 547 |
| PBANKA_0606200 | 0 | 0 | 0 | 0 | 11 | 6 | 7 | 0 | 6 | 5 | 0 | 0 | 1219 | 519 | 76 | 193 | 411 | 346 | 25627 | 12599 | 205 | 1405 |
| PBANKA_0608800 | 64 | 89 | 0 | 0 | 27 | 6 | 95 | 35 | 75 | 20 | 14 | 0 | 248 | 54 | 22 | 88 | 672 | 201 | 489 | 642 | 97 | 589 |
| PBANKA_0610700 | 0 | 0 | 0 | 0 | 8 | 5 | 19 | 9 | 12 | 0 | 6 | 0 | 94 | 21 | 0 | 17 | 107 | 28 | 881 | 293 | 94 | 595 |
| PBANKA_0612900 | 0 | 0 | 0 | 0 | 6 | 0 | 9 | 0 | 12 | 0 | 0 | 0 | 109 | 45 | 7 | 28 | 85 | 35 | 2766 | 853 | 67 | 404 |
| PBANKA_0617000 | 0 | 0 | 0 | 0 | 9 | 5 | 14 | 15 | 27 | 15 | 14 | 9 | 297 | 120 | 26 | 66 | 258 | 150 | 2767 | 2040 | 462 | 3690 |
| PBANKA_0617800 | 0 | 0 | 0 | 0 | 6 | 0 | 17 | 0 | 9 | 5 | 0 | 0 | 43 | 22 | 6 | 17 | 29 | 11 | 390 | 232 | 509 | 3296 |
| PBANKA_0619000 | 0 | 0 | 0 | 0 | 47 | 15 | 177 | 69 | 176 | 104 | 12 | 9 | 1446 | 399 | 106 | 287 | 5363 | 896 | 2051 | 1638 | 100 | 622 |
| PBANKA_0619200 | 0 | 5 | 0 | 0 | 8 | 11 | 69 | 18 | 57 | 32 | 14 | 5 | 1483 | 504 | 63 | 153 | 1295 | 552 | 12993 | 6952 | 55992 | 5E+05 |
| PBANKA_0619600 | 38 | 43 | 0 | 0 | 37 | 19 | 54 | 22 | 44 | 17 | 14 | 9 | 311 | 178 | 107 | 479 | 439 | 190 | 314 | 309 | 1541 | 9838 |
| PBANKA_0620600 | 0 | 0 | 0 | 0 | 13 | 12 | 16 | 20 | 18 | 15 | 14 | 0 | 170 | 52 | 15 | 46 | 231 | 92 | 2587 | 1529 | 339 | 2932 |
| PBANKA_0622000 | 6 | 21 | 0 | 0 | 29 | 0 | 40 | 15 | 25 | 14 | 0 | 5 | 1283 | 276 | 18 | 64 | 494 | 229 | 13348 | 9439 | 131 | 749 |
| PBANKA_0623150 | 0 | 0 | 0 | 0 | 5 | 0 | 18 | 10 | 40 | 24 | 783 | 538 | 528 | 319 | 88 | 290 | 123 | 45 | 1503 | 578 | 29 | 113 |
| PBANKA_0623450 | 9 | 12 | 0 | 0 | 11 | 0 | 81 | 45 | 81 | 49 | 6 | 0 | 954 | 252 | 11 | 17 | 3309 | 1511 | 4723 | 2122 | 107 | 587 |
| PBANKA_0700061 | 0 | 0 | 0 | 0 | 6 | 0 | 8 | 0 | 7 | 0 | 181 | 106 | 81 | 34 | 17 | 66 | 12 | 6 | 407 | 166 | 0 | 6 |
| PBANKA_0700100 | 0 | 0 | 0 | 0 | 5 | 7 | 25 | 7 | 19 | 9 | 7 | 0 | 189 | 98 | 51 | 207 | 54 | 12 | 89 | 73 | 0 | 23 |
| PBANKA_0700561 | 0 | 0 | 0 | 0 | 6 | 0 | 0 | 0 | 0 | 0 | 41 | 8 | 771 | 1077 | 142 | 659 | 262 | 23 | 240 | 287 | 0 | 42 |
| PBANKA_0700600 | 0 | 0 | 0 | 0 | 9 | 0 | 13 | 0 | 15 | 24 | 914 | 720 | 3911 | 3158 | 427 | 2138 | 318 | 64 | 435 | 321 | 17 | 125 |
| PBANKA_0701100 | 0 | 0 | 0 | 0 | 22 | 12 | 163 | 93 | 95 | 58 | 395 | 299 | 5367 | 1744 | 2422 | 7986 | 4918 | 955 | 1439 | 640 | 34 | 278 |
| PBANKA_0701900 | 4841 | 7583 | 0 | 0 | 70 | 41 | 106 | 27 | 290 | 224 | 109 | 55 | 286 | 169 | 23 | 55 | 874 | 457 | 338 | 432 | 10956 | 80513 |
| PBANKA_0702000 | 96 | 165 | 0 | 0 | 40 | 16 | 67 | 30 | 379 | 215 | 159 | 84 | 1491 | 1272 | 106 | 460 | 4276 | 1028 | 5726 | 25399 | 868 | 3402 |
| PBANKA_0704700 | 0 | 0 | 0 | 0 | 21 | 17 | 22 | 14 | 20 | 11 | 0 | 0 | 420 | 259 | 25 | 45 | 572 | 278 | 6381 | 4941 | 185 | 1568 |
| PBANKA_0704800 | 0 | 0 | 0 | 0 | 5 | 8 | 12 | 9 | 12 | 6 | 0 | 0 | 3452 | 1542 | 33 | 133 | 905 | 535 | 86610 | 78888 | 883 | 8496 |
| PBANKA_0704900 | 0 | 0 | 0 | 0 | 19 | 0 | 5 | 8 | 9 | 0 | 0 | 0 | 1830 | 700 | 16 | 54 | 483 | 312 | 55952 | 29923 | 415 | 3510 |
| PBANKA_0707000 | 0 | 0 | 0 | 0 | 87 | 21 | 98 | 26 | 27 | 10 | 8 | 0 | 48 | 21 | 10 | 22 | 122 | 28 | 1094 | 309 | 53 | 296 |
| PBANKA_0707100 | 0 | 0 | 0 | 0 | 9 | 7 | 0 | 0 | 9 | 0 | 0 | 0 | 2686 | 1553 | 57 | 164 | 1097 | 749 | 78784 | 53901 | 17674 | 2E+05 |
| PBANKA_0707200 | 10 | 0 | 0 | 0 | 18 | 9 | 23 | 5 | 10 | 6 | 10 | 0 | 325 | 138 | 25 | 80 | 119 | 51 | 2534 | 1016 | 45 | 368 |
| PBANKA_0707900 | 0 | 0 | 0 | 0 | 34 | 7 | 33 | 22 | 50 | 31 | 23 | 0 | 92 | 49 | 40 | 81 | 244 | 55 | 384 | 296 | 133 | 1330 |
| PBANKA_0712500 | 0 | 0 | 0 | 0 | 14 | 0 | 16 | 14 | 33 | 9 | 7 | 0 | 99 | 29 | 50 | 122 | 91 | 41 | 1558 | 510 | 88 | 713 |
| PBANKA_0714300 | 292 | 369 | 0 | 0 | 10 | 11 | 8 | 0 | 8 | 8 | 0 | 0 | 145 | 55 | 11 | 32 | 42 | 23 | 2936 | 2109 | 4312 | 40619 |
| PBANKA_0716200 | 0 | 0 | 0 | 0 | 69 | 41 | 125 | 62 | 39 | 28 | 0 | 6 | 85 | 17 | 25 | 35 | 332 | 95 | 576 | 346 | 19 | 157 |
| PBANKA_0716400 | 961 | 1343 | 0 | 0 | 52 | 19 | 41 | 17 | 55 | 11 | 8 | 0 | 160 | 48 | 8 | 47 | 140 | 52 | 781 | 286 | 700 | 4120 |
| PBANKA_0718500 | 0 | 0 | 0 | 0 | 79 | 25 | 87 | 32 | 25 | 12 | 10 | 5 | 325 | 107 | 100 | 529 | 310 | 70 | 2035 | 983 | 13 | 108 |

| | | | | | | | | | | | | | | | | | | | | | | |
|---|---|---|---|---|---|---|---|---|---|---|---|---|---|---|---|---|---|---|---|---|---|---|
| PBANKA_0719100 | 0 | 0 | 0 | 0 | 5 | 0 | 10 | 7 | 13 | 6 | 0 | 0 | 179 | 50 | 19 | 65 | 399 | 129 | 3660 | 1720 | 520 | 3383 |
| PBANKA_0719200 | 3345 | 4918 | 0 | 0 | 39 | 29 | 49 | 15 | 36 | 18 | 10 | 7 | 72 | 24 | 47 | 210 | 113 | 36 | 572 | 301 | 296 | 1946 |
| PBANKA_0719700 | 0 | 14 | 0 | 0 | 10 | 10 | 30 | 14 | 36 | 8 | 17 | 14 | 367 | 435 | 23 | 90 | 1389 | 220 | 1878 | 7477 | 1065 | 7369 |
| PBANKA_0719800 | 10 | 7 | 0 | 0 | 10 | 7 | 15 | 0 | 15 | 0 | 0 | 0 | 288 | 110 | 14 | 55 | 126 | 75 | 4783 | 2276 | 744 | 3833 |
| PBANKA_0720000 | 5 | 0 | 0 | 0 | 20 | 8 | 52 | 18 | 85 | 33 | 33 | 16 | 654 | 214 | 109 | 1143 | 778 | 172 | 4572 | 3452 | 79 | 549 |
| PBANKA_0720900 | 11 | 14 | 0 | 0 | 27 | 21 | 120 | 38 | 562 | 423 | 88 | 47 | 1696 | 468 | 32 | 91 | 395 | 232 | 31526 | 12403 | 273 | 1700 |
| PBANKA_0721000 | 426 | 618 | 0 | 0 | 25 | 9 | 40 | 17 | 69 | 45 | 7 | 0 | 144 | 45 | 15 | 37 | 339 | 136 | 1337 | 943 | 1378 | 8532 |
| PBANKA_0721100 | 0 | 0 | 0 | 0 | 18 | 9 | 14 | 10 | 19 | 18 | 121 | 69 | 813 | 280 | 90 | 422 | 328 | 86 | 4775 | 2694 | 253 | 1001 |
| PBANKA_0721800 | 13 | 17 | 0 | 0 | 22 | 6 | 63 | 37 | 137 | 83 | 9 | 14 | 306 | 85 | 11 | 70 | 2133 | 534 | 2513 | 740 | 27 | 243 |
| PBANKA_0722921 | 0 | 0 | 0 | 0 | 65 | 39 | 62 | 36 | 40 | 16 | 14 | 0 | 292 | 28 | 68 | 141 | 137 | 211 | 120 | 25 | 12 | 33 |
| PBANKA_0800300 | 0 | 0 | 0 | 0 | 22 | 14 | 30 | 18 | 33 | 42 | 737 | 542 | 1957 | 1810 | 188 | 885 | 237 | 48 | 1203 | 1034 | 163 | 1695 |
| PBANKA_0800500 | 0 | 0 | 0 | 0 | 16 | 10 | 56 | 5 | 24 | 11 | 86 | 68 | 1236 | 888 | 113 | 853 | 168 | 35 | 527 | 333 | 3E+05 | 2E+06 |
| PBANKA_0801500 | 0 | 0 | 0 | 0 | 86 | 25 | 347 | 123 | 126 | 46 | 11 | 13 | 367 | 220 | 75 | 287 | 958 | 136 | 1557 | 2938 | 32 | 97 |
| PBANKA_0803900 | 0 | 0 | 0 | 0 | 8 | 7 | 17 | 6 | 110 | 93 | 17 | 24 | 160 | 73 | 14 | 47 | 988 | 468 | 71 | 68 | 0 | 37 |
| PBANKA_0804300 | 0 | 0 | 0 | 0 | 7 | 0 | 13 | 0 | 0 | 0 | 0 | 0 | 243 | 72 | 44 | 162 | 217 | 70 | 2521 | 1683 | 22 | 170 |
| PBANKA_0805750 | 0 | 0 | 0 | 0 | 8 | 0 | 6 | 0 | 0 | 0 | 5 | 5 | 42 | 23 | 80 | 56 | 36 | 19 | 6 | 28 | 0 | 20 |
| PBANKA_0806800 | 0 | 0 | 0 | 0 | 19 | 9 | 29 | 10 | 12 | 7 | 0 | 6 | 486 | 258 | 10 | 55 | 356 | 65 | 2750 | 5060 | 51 | 371 |
| PBANKA_0807200 | 0 | 0 | 0 | 0 | 11 | 0 | 31 | 19 | 23 | 0 | 0 | 7 | 1127 | 382 | 28 | 119 | 617 | 175 | 28980 | 16474 | 1529 | 9535 |
| PBANKA_0810300 | 0 | 0 | 0 | 0 | 9 | 0 | 26 | 18 | 23 | 13 | 7 | 5 | 121 | 22 | 27 | 78 | 100 | 34 | 1356 | 732 | 288 | 1916 |
| PBANKA_0810400 | 0 | 0 | 0 | 0 | 10 | 0 | 20 | 11 | 27 | 6 | 0 | 0 | 74 | 23 | 18 | 77 | 64 | 19 | 795 | 598 | 223 | 1678 |
| PBANKA_0810700 | 0 | 0 | 0 | 0 | 60 | 37 | 67 | 44 | 33 | 16 | 0 | 0 | 1694 | 787 | 23 | 112 | 670 | 334 | 41350 | 23391 | 2307 | 24723 |
| PBANKA_0811800 | 0 | 0 | 0 | 0 | 6 | 0 | 25 | 8 | 16 | 0 | 0 | 0 | 128 | 43 | 8 | 24 | 145 | 63 | 1792 | 1268 | 29 | 230 |
| PBANKA_0812600 | 0 | 0 | 0 | 0 | 11 | 6 | 27 | 6 | 11 | 10 | 0 | 0 | 1019 | 358 | 19 | 46 | 425 | 179 | 25744 | 10542 | 26 | 196 |
| PBANKA_0813600 | 0 | 0 | 0 | 0 | 28 | 22 | 53 | 28 | 56 | 27 | 12 | 0 | 357 | 180 | 78 | 346 | 179 | 53 | 1714 | 1810 | 48 | 201 |
| PBANKA_0817000 | 6 | 11 | 0 | 0 | 34 | 15 | 155 | 77 | 387 | 173 | 45 | 24 | 1043 | 381 | 55 | 210 | 5548 | 1243 | 6378 | 5088 | 145 | 1172 |
| PBANKA_0818400 | 8 | 19 | 0 | 0 | 23 | 6 | 89 | 40 | 133 | 69 | 31 | 9 | 34 | 60 | 7 | 6 | 594 | 120 | 318 | 699 | 3205 | 21915 |
| PBANKA_0819000 | 0 | 7 | 0 | 0 | 137 | 68 | 1033 | 506 | 1245 | 718 | 155 | 76 | 4729 | 1540 | 78 | 190 | 23479 | 4715 | 76154 | 43759 | 1649 | 16981 |
| PBANKA_0820100 | 0 | 0 | 0 | 0 | 48 | 24 | 62 | 21 | 169 | 176 | 47 | 39 | 113 | 86 | 114 | 265 | 430 | 245 | 1073 | 548 | 17 | 126 |
| PBANKA_0821200 | 0 | 0 | 0 | 0 | 32 | 14 | 56 | 19 | 15 | 10 | 0 | 0 | 586 | 241 | 29 | 122 | 226 | 91 | 15938 | 10770 | 134 | 774 |
| PBANKA_0821400 | 12 | 47 | 0 | 0 | 16 | 0 | 22 | 15 | 9 | 0 | 0 | 0 | 331 | 58 | 9 | 7 | 83 | 72 | 9990 | 3230 | 402 | 3269 |
| PBANKA_0822900 | 16 | 13 | 0 | 0 | 21 | 21 | 68 | 32 | 51 | 26 | 34 | 13 | 619 | 284 | 56 | 268 | 643 | 143 | 3420 | 3750 | 927 | 5524 |
| PBANKA_0823200 | 115 | 172 | 0 | 0 | 15 | 5 | 16 | 0 | 9 | 6 | 0 | 0 | 839 | 366 | 28 | 88 | 256 | 124 | 18490 | 13988 | 463 | 3296 |
| PBANKA_0824400 | 0 | 0 | 0 | 0 | 44 | 15 | 656 | 293 | 758 | 389 | 65 | 51 | 1240 | 383 | 13 | 48 | 3597 | 751 | 13179 | 10262 | 233 | 2675 |
| PBANKA_0824600 | 17 | 8 | 0 | 0 | 27 | 20 | 45 | 11 | 12 | 0 | 0 | 0 | 98 | 15 | 34 | 136 | 114 | 39 | 330 | 151 | 0 | 8 |
| PBANKA_0825400 | 15 | 14 | 0 | 0 | 59 | 21 | 424 | 204 | 404 | 196 | 32 | 13 | 829 | 247 | 38 | 145 | 5981 | 1171 | 3708 | 2677 | 695 | 4035 |
| PBANKA_0825700 | 0 | 0 | 0 | 0 | 11 | 0 | 67 | 57 | 60 | 33 | 0 | 0 | 476 | 118 | 0 | 9 | 304 | 93 | 4613 | 2406 | 123 | 267 |
| PBANKA_0825800 | 5 | 0 | 0 | 0 | 36 | 16 | 205 | 75 | 134 | 56 | 17 | 5 | 239 | 43 | 5 | 15 | 176 | 51 | 1208 | 640 | 23 | 127 |
| PBANKA_0827400 | 0 | 0 | 0 | 0 | 53 | 30 | 44 | 20 | 55 | 37 | 13 | 7 | 521 | 356 | 33 | 132 | 392 | 178 | 10545 | 16172 | 1314 | 10707 |
| PBANKA_0828300 | 0 | 0 | 0 | 0 | 26 | 10 | 24 | 17 | 11 | 0 | 0 | 0 | 41 | 10 | 0 | 0 | 44 | 24 | 401 | 112 | 69 | 154 |
| PBANKA_0830300 | 20 | 16 | 0 | 0 | 5 | 0 | 8 | 0 | 11 | 6 | 0 | 0 | 153 | 21 | 27 | 35 | 150 | 129 | 1661 | 900 | 19 | 121 |
| PBANKA_0830400 | 0 | 5 | 0 | 0 | 25 | 0 | 37 | 31 | 19 | 24 | 8 | 0 | 1380 | 619 | 41 | 79 | 441 | 281 | 42688 | 19947 | 3974 | 34495 |
| PBANKA_0830900 | 24 | 66 | 0 | 0 | 58 | 32 | 175 | 54 | 380 | 171 | 63 | 30 | 420 | 114 | 9 | 31 | 360 | 128 | 4734 | 1839 | 98 | 617 |
| PBANKA_0832700 | 0 | 7 | 0 | 0 | 13 | 0 | 27 | 6 | 27 | 12 | 0 | 0 | 90 | 10 | 0 | 11 | 132 | 49 | 897 | 272 | 200 | 948 |
| PBANKA_0832800 | 6 | 11 | 0 | 0 | 8 | 0 | 22 | 13 | 443 | 436 | 177 | 116 | 74 | 38 | 25 | 45 | 277 | 148 | 52 | 62 | 889 | 4113 |
| PBANKA_0833700 | 0 | 0 | 0 | 0 | 22 | 20 | 46 | 21 | 33 | 17 | 6 | 0 | 205 | 39 | 31 | 131 | 125 | 48 | 953 | 323 | 120 | 761 |
| PBANKA_0835600 | 0 | 0 | 0 | 0 | 23 | 16 | 125 | 69 | 275 | 195 | 23 | 14 | 573 | 114 | 51 | 86 | 3927 | 1260 | 1606 | 572 | 10 | 73 |
| PBANKA_0836200 | 0 | 0 | 0 | 0 | 17 | 6 | 16 | 7 | 651 | 470 | 5015 | 4448 | 2559 | 1655 | 875 | 3732 | 664 | 301 | 302 | 236 | 6 | 80 |
| PBANKA_0836500 | 0 | 0 | 0 | 0 | 7 | 0 | 0 | 5 | 8 | 0 | 81 | 57 | 260 | 176 | 71 | 201 | 77 | 10 | 448 | 210 | 5 | 13 |
| PBANKA_0836800 | 0 | 0 | 0 | 0 | 30 | 16 | 27 | 12 | 15 | 5 | 34 | 20 | 2202 | 1785 | 1334 | 5250 | 963 | 93 | 722 | 893 | 7 | 152 |
| PBANKA_0902200 | 0 | 0 | 0 | 0 | 52 | 23 | 142 | 82 | 58 | 29 | 0 | 0 | 194 | 18 | 104 | 235 | 405 | 165 | 783 | 144 | 20 | 154 |
| PBANKA_0902600 | 0 | 0 | 0 | 0 | 12 | 0 | 23 | 5 | 11 | 6 | 0 | 0 | 88 | 13 | 0 | 0 | 58 | 17 | 1145 | 363 | 0 | 8 |
| PBANKA_0904000 | 197 | 323 | 0 | 0 | 66 | 20 | 56 | 30 | 98 | 108 | 41 | 32 | 343 | 112 | 21 | 145 | 444 | 205 | 2930 | 954 | 189 | 1492 |
| PBANKA_0905300 | 0 | 0 | 0 | 0 | 45 | 20 | 72 | 30 | 38 | 27 | 10 | 5 | 188 | 47 | 9 | 19 | 148 | 70 | 2477 | 1138 | 360 | 1806 |
| PBANKA_0905500 | 72 | 119 | 0 | 0 | 12 | 0 | 20 | 11 | 59 | 42 | 12 | 5 | 187 | 77 | 19 | 62 | 384 | 102 | 1332 | 1047 | 247 | 1520 |
| PBANKA_0906100 | 54 | 103 | 0 | 0 | 5 | 0 | 17 | 10 | 55 | 27 | 11 | 0 | 99 | 65 | 7 | 30 | 654 | 139 | 458 | 291 | 218 | 1408 |
| PBANKA_0906800 | 5 | 0 | 0 | 0 | 19 | 9 | 16 | 6 | 13 | 0 | 5 | 0 | 37 | 13 | 0 | 32 | 17 | 5 | 394 | 122 | 306 | 1814 |
| PBANKA_0907400 | 0 | 0 | 0 | 0 | 31 | 13 | 33 | 13 | 12 | 13 | 0 | 0 | 157 | 32 | 57 | 146 | 329 | 84 | 994 | 464 | 31 | 230 |
| PBANKA_0908400 | 54 | 105 | 0 | 0 | 43 | 14 | 23 | 15 | 44 | 25 | 35 | 8 | 82 | 265 | 84 | 194 | 294 | 59 | 572 | 1650 | 1028 | 6091 |
| PBANKA_0910500 | 0 | 0 | 0 | 0 | 6 | 0 | 0 | 0 | 6 | 0 | 0 | 0 | 0 | 0 | 0 | 0 | 6 | 0 | 22 | 11 | 0 | 0 |
| PBANKA_0910900 | 5882 | 8159 | 0 | 0 | 30 | 15 | 25 | 14 | 18 | 12 | 15 | 0 | 87 | 10 | 0 | 0 | 91 | 31 | 932 | 551 | 43 | 276 |
| PBANKA_0911100 | 9 | 8 | 0 | 0 | 50 | 18 | 58 | 32 | 63 | 34 | 8 | 8 | 145 | 46 | 5 | 26 | 108 | 37 | 1353 | 748 | 10 | 38 |
| PBANKA_0912500 | 28 | 31 | 0 | 0 | 23 | 9 | 46 | 38 | 184 | 98 | 30 | 16 | 824 | 518 | 31 | 75 | 886 | 356 | 19477 | 16194 | 278 | 2361 |
| PBANKA_0913600 | 0 | 0 | 0 | 0 | 12 | 7 | 17 | 8 | 9 | 5 | 0 | 0 | 177 | 68 | 15 | 61 | 126 | 37 | 1489 | 1108 | 7 | 25 |
| PBANKA_0915000 | 2505 | 3744 | 0 | 0 | 20 | 12 | 44 | 12 | 1288 | 1059 | 333 | 256 | 879 | 355 | 9 | 0 | 1457 | 1280 | 529 | 523 | 58 | 365 |
| PBANKA_0915100 | 0 | 0 | 0 | 0 | 28 | 14 | 80 | 33 | 83 | 44 | 10 | 7 | 235 | 54 | 0 | 0 | 1557 | 462 | 98 | 62 | 39 | 246 |
| PBANKA_0916500 | 0 | 0 | 0 | 0 | 9 | 5 | 25 | 7 | 28 | 7 | 6 | 7 | 58 | 47 | 8 | 14 | 259 | 42 | 121 | 101 | 196 | 1512 |
| PBANKA_0916700 | 0 | 9 | 0 | 0 | 10 | 10 | 52 | 37 | 73 | 45 | 5 | 0 | 1629 | 734 | 27 | 78 | 997 | 285 | 49355 | 33977 | 2593 | 22413 |
| PBANKA_0918500 | 0 | 0 | 0 | 0 | 36 | 21 | 62 | 19 | 19 | 9 | 0 | 0 | 211 | 45 | 68 | 375 | 311 | 48 | 1272 | 798 | 74 | 461 |
| PBANKA_0919400 | 0 | 0 | 0 | 0 | 64 | 24 | 167 | 66 | 385 | 190 | 59 | 34 | 659 | 203 | 89 | 261 | 4703 | 1308 | 3802 | 2323 | 138 | 964 |
| PBANKA_0919500 | 0 | 0 | 0 | 0 | 30 | 15 | 43 | 9 | 18 | 12 | 0 | 0 | 132 | 40 | 63 | 230 | 545 | 101 | 821 | 399 | 76 | 582 |
| PBANKA_0921200 | 433 | 340 | 0 | 0 | 9 | 7 | 15 | 13 | 20 | 7 | 10 | 0 | 8 | 5 | 6 | 0 | 8 | 0 | 80 | 36 | 1355 | 8386 |
| PBANKA_0922600 | 5 | 5 | 0 | 0 | 32 | 8 | 191 | 70 | 930 | 546 | 191 | 109 | 271 | 88 | 23 | 75 | 1770 | 600 | 555 | 297 | 185 | 1353 |
| PBANKA_0927300 | 0 | 0 | 0 | 0 | 8 | 0 | 40 | 23 | 158 | 123 | 21 | 12 | 186 | 47 | 19 | 27 | 937 | 374 | 2626 | 1132 | 102 | 652 |
| PBANKA_0927600 | 0 | 5 | 0 | 0 | 34 | 13 | 57 | 13 | 27 | 21 | 14 | 10 | 3065 | 2164 | 141 | 518 | 2128 | 845 | 85647 | 98529 | 675 | 4594 |
| PBANKA_0931100 | 821 | 813 | 0 | 0 | 119 | 49 | 366 | 260 | 286 | 278 | 28 | 46 | 314 | 88 | 9 | 19 | 1313 | 498 | 1700 | 740 | 654 | 2961 |
| PBANKA_0931200 | 14 | 26 | 0 | 0 | 36 | 16 | 88 | 28 | 2091 | 1483 | 5301 | 3479 | 9509 | 5797 | 2373 | 11517 | 6365 | 3020 | 2598 | 2265 | 54 | 512 |
| PBANKA_0932000 | 0 | 0 | 0 | 0 | 17 | 17 | 1064 | 448 | 3116 | 2028 | 287 | 215 | 1652 | 681 | 49 | 98 | 25255 | 7064 | 1191 | 1268 | 128 | 1352 |
| PBANKA_0932600 | 0 | 6 | 0 | 0 | 13 | 8 | 53 | 13 | 91 | 50 | 5 | 6 | 470 | 226 | 16 | 62 | 316 | 174 | 8488 | 5155 | 936 | 5308 |
| PBANKA_0933100 | 0 | 6 | 0 | 0 | 9 | 0 | 12 | 10 | 9 | 0 | 0 | 0 | 80 | 16 | 16 | 70 | 65 | 16 | 400 | 148 | 15 | 103 |
| PBANKA_0933600 | 23 | 13 | 0 | 0 | 5 | 0 | 0 | 0 | 6 | 0 | 0 | 0 | 77 | 9 | 0 | 7 | 150 | 81 | 820 | 266 | 9 | 87 |
| PBANKA_0934800 | 0 | 0 | 0 | 0 | 19 | 0 | 81 | 31 | 77 | 33 | 6 | 0 | 74 | 13 | 9 | 44 | 148 | 37 | 618 | 224 | 0 | 40 |
| PBANKA_0934900 | 0 | 0 | 0 | 0 | 11 | 0 | 38 | 17 | 27 | 16 | 0 | 8 | 71 | 13 | 23 | 115 | 142 | 30 | 742 | 370 | 0 | 29 |
| PBANKA_0935700 | 0 | 0 | 0 | 0 | 10 | 0 | 18 | 11 | 24 | 12 | 0 | 0 | 148 | 15 | 6 | 31 | 576 | 139 | 1368 | 377 | 11 | 56 |
| PBANKA_0936000 | 469 | 622 | 0 | 0 | 25 | 5 | 65 | 25 | 58 | 28 | 6 | 0 | 170 | 71 | 51 | 155 | 228 | 101 | 2654 | 2020 | 452 | 3201 |
| PBANKA_0936800 | 0 | 0 | 0 | 0 | 43 | 18 | 94 | 26 | 32 | 13 | 17 | 6 | 332 | 79 | 26 | 98 | 210 | 44 | 1379 | 501 | 16 | 100 |
| PBANKA_0942300 | 0 | 0 | 0 | 0 | 9 | 6 | 31 | 17 | 27 | 15 | 0 | 0 | 212 | 56 | 19 | 41 | 288 | 75 | 2063 | 850 | 32 | 209 |
| PBANKA_0944041 | 0 | 0 | 0 | 0 | 8 | 0 | 7 | 0 | 53 | 37 | 11 | 0 | 48 | 232 | 52 | 101 | 1044 | 379 | 38 | 147 | 0 | 51 |
| PBANKA_1000021 | 0 | 0 | 0 | 0 | 5 | 0 | 16 | 13 | 8 | 5 | 0 | 0 | 0 | 0 | 0 | 0 | 16 | 6 | 0 | 0 | 0 | 0 |
| PBANKA_1000200 | 0 | 0 | 0 | 0 | 5 | 0 | 0 | 0 | 5 | 0 | 10 | 12 | 142 | 202 | 48 | 91 | 39 | 22 | 25 | 9 | 0 | 6 |
| PBANKA_1000300 | 0 | 0 | 0 | 0 | 55 | 46 | 74 | 41 | 29 | 26 | 103 | 106 | 1175 | 583 | 821 | 2929 | 946 | 208 | 1067 | 429 | 28 | 241 |

| | | | | | | | | | | | | | | | | | | | | | | |
|---|---|---|---|---|---|---|---|---|---|---|---|---|---|---|---|---|---|---|---|---|---|---|
| PBANKA_1000500 | 0 | 0 | 0 | 0 | 53 | 34 | 79 | 38 | 74 | 36 | 1233 | 1024 | 2252 | 1538 | 605 | 2450 | 932 | 148 | 738 | 481 | 19 | 171 |
| PBANKA_1000600 | 0 | 0 | 0 | 0 | 13 | 19 | 12 | 13 | 370 | 316 | 6675 | 7397 | 4267 | 2531 | 2179 | 6999 | 1089 | 228 | 1586 | 751 | 30 | 416 |
| PBANKA_1002200 | 858 | 1065 | 0 | 0 | 10 | 7 | 13 | 6 | 19 | 13 | 6 | 0 | 0 | 0 | 0 | 0 | 7 | 0 | 15 | 10 | 60 | 434 |
| PBANKA_1002500 | 201 | 263 | 0 | 0 | 16 | 7 | 27 | 18 | 116 | 59 | 39 | 8 | 200 | 111 | 8 | 21 | 431 | 125 | 2495 | 2453 | 68 | 443 |
| PBANKA_1002600 | 19 | 20 | 0 | 0 | 54 | 25 | 113 | 50 | 1408 | 1212 | 465 | 215 | 707 | 384 | 17 | 25 | 3863 | 1904 | 650 | 567 | 34 | 348 |
| PBANKA_1002900 | 0 | 0 | 0 | 0 | 86 | 52 | 95 | 48 | 125 | 77 | 31 | 20 | 147 | 61 | 34 | 138 | 793 | 194 | 112 | 129 | 7 | 24 |
| PBANKA_1003600 | 8 | 8 | 0 | 0 | 20 | 17 | 153 | 71 | 478 | 349 | 68 | 46 | 963 | 619 | 106 | 473 | 12279 | 2280 | 943 | 950 | 161 | 1163 |
| PBANKA_1005600 | 0 | 0 | 0 | 0 | 8 | 0 | 9 | 0 | 99 | 80 | 242 | 225 | 385 | 137 | 39 | 161 | 225 | 134 | 3397 | 1305 | 0 | 18 |
| PBANKA_1006400 | 6 | 0 | 0 | 0 | 5 | 5 | 0 | 0 | 6 | 0 | 0 | 0 | 23 | 0 | 7 | 11 | 0 | 0 | 83 | 38 | 4400 | 35556 |
| PBANKA_1007900 | 0 | 0 | 0 | 0 | 16 | 0 | 16 | 6 | 5 | 5 | 5 | 0 | 776 | 309 | 55 | 241 | 224 | 121 | 16679 | 10645 | 1024 | 7646 |
| PBANKA_1009600 | 12 | 15 | 0 | 0 | 28 | 15 | 42 | 16 | 13 | 9 | 18 | 0 | 1479 | 640 | 47 | 157 | 496 | 194 | 23351 | 20798 | 679 | 4637 |
| PBANKA_1009900 | 0 | 0 | 0 | 0 | 40 | 28 | 44 | 13 | 31 | 9 | 6 | 0 | 538 | 319 | 49 | 119 | 196 | 134 | 18994 | 8948 | 1590 | 11817 |
| PBANKA_1010400 | 446 | 618 | 0 | 0 | 7 | 6 | 40 | 22 | 325 | 251 | 80 | 33 | 272 | 115 | 30 | 38 | 2060 | 646 | 290 | 177 | 238 | 1598 |
| PBANKA_1013600 | 0 | 0 | 0 | 0 | 73 | 34 | 91 | 34 | 48 | 34 | 8 | 0 | 216 | 45 | 137 | 659 | 337 | 73 | 1369 | 463 | 24 | 169 |
| PBANKA_1013900 | 0 | 0 | 0 | 0 | 16 | 0 | 76 | 20 | 102 | 44 | 19 | 9 | 459 | 103 | 19 | 53 | 592 | 192 | 4534 | 1552 | 207 | 1523 |
| PBANKA_1014200 | 0 | 0 | 0 | 0 | 10 | 0 | 10 | 0 | 5 | 0 | 0 | 5 | 79 | 38 | 42 | 119 | 135 | 33 | 447 | 382 | 61 | 325 |
| PBANKA_1014500 | 0 | 0 | 0 | 0 | 46 | 24 | 159 | 69 | 575 | 320 | 143 | 44 | 531 | 271 | 23 | 37 | 5110 | 1938 | 1654 | 1271 | 161 | 1062 |
| PBANKA_1016900 | 0 | 0 | 0 | 0 | 36 | 14 | 63 | 32 | 24 | 19 | 0 | 0 | 69 | 8 | 30 | 43 | 322 | 169 | 138 | 58 | 33 | 270 |
| PBANKA_1017400 | 0 | 0 | 0 | 0 | 20 | 5 | 58 | 24 | 51 | 29 | 6 | 0 | 405 | 64 | 58 | 274 | 1194 | 283 | 648 | 216 | 11 | 97 |
| PBANKA_1019320 | 9 | 0 | 0 | 0 | 59 | 9 | 93 | 16 | 35 | 12 | 46 | 20 | 98 | 41 | 18 | 40 | 39 | 6 | 66 | 9 | 86 | 312 |
| PBANKA_1020000 | 36 | 39 | 0 | 0 | 23 | 12 | 49 | 18 | 79 | 43 | 15 | 0 | 252 | 195 | 49 | 133 | 1011 | 330 | 2277 | 4394 | 3923 | 24546 |
| PBANKA_1020300 | 29 | 30 | 0 | 0 | 22 | 16 | 13 | 13 | 65 | 33 | 13 | 7 | 740 | 396 | 31 | 107 | 556 | 168 | 17301 | 14516 | 626 | 5138 |
| PBANKA_1020500 | 84 | 105 | 0 | 0 | 60 | 22 | 52 | 25 | 22 | 15 | 7 | 0 | 100 | 35 | 16 | 96 | 282 | 58 | 678 | 969 | 2435 | 14973 |
| PBANKA_1021200 | 0 | 7 | 0 | 0 | 50 | 16 | 80 | 25 | 198 | 115 | 43 | 14 | 285 | 100 | 18 | 44 | 1561 | 403 | 1913 | 1070 | 345 | 2384 |
| PBANKA_1022100 | 0 | 0 | 0 | 0 | 74 | 34 | 200 | 55 | 116 | 50 | 42 | 41 | 1243 | 628 | 385 | 1465 | 1822 | 270 | 694 | 427 | 23 | 174 |
| PBANKA_1022400 | 0 | 0 | 0 | 0 | 8 | 0 | 99 | 55 | 69 | 42 | 13 | 5 | 89 | 19 | 6 | 15 | 66 | 31 | 854 | 211 | 0 | 50 |
| PBANKA_1024331 | 0 | 0 | 0 | 0 | 16 | 0 | 6 | 0 | 16 | 0 | 26 | 0 | 11 | 9 | 764 | 12 | 6 | 27 | 59 | 0 | 0 | 0 |
| PBANKA_1025700 | 30 | 49 | 0 | 0 | 6 | 0 | 7 | 0 | 22 | 20 | 22 | 8 | 43 | 20 | 14 | 52 | 48 | 15 | 316 | 292 | 2747 | 20410 |
| PBANKA_1026000 | 0 | 0 | 0 | 0 | 20 | 6 | 25 | 8 | 0 | 6 | 0 | 0 | 188 | 68 | 31 | 154 | 162 | 57 | 3001 | 1825 | 24 | 157 |
| PBANKA_1026100 | 8 | 9 | 0 | 0 | 47 | 26 | 28 | 18 | 70 | 28 | 10 | 8 | 49 | 130 | 38 | 76 | 219 | 76 | 450 | 2473 | 63 | 334 |
| PBANKA_1029500 | 0 | 0 | 0 | 0 | 8 | 5 | 9 | 0 | 22 | 14 | 16 | 0 | 215 | 55 | 8 | 22 | 81 | 34 | 1886 | 905 | 38 | 291 |
| PBANKA_1029600 | 0 | 0 | 0 | 0 | 7 | 0 | 5 | 0 | 0 | 8 | 0 | 0 | 835 | 238 | 7 | 51 | 455 | 118 | 8815 | 5102 | 168 | 975 |
| PBANKA_1030400 | 0 | 0 | 0 | 0 | 5 | 0 | 14 | 0 | 15 | 12 | 13 | 0 | 524 | 248 | 67 | 246 | 656 | 154 | 3364 | 4563 | 7 | 85 |
| PBANKA_1031500 | 50 | 77 | 0 | 0 | 14 | 14 | 23 | 28 | 38 | 25 | 10 | 9 | 117 | 76 | 38 | 33 | 268 | 76 | 1270 | 2299 | 59 | 678 |
| PBANKA_1031700 | 0 | 0 | 0 | 0 | 6 | 0 | 30 | 7 | 104 | 75 | 5 | 10 | 91 | 13 | 0 | 15 | 531 | 162 | 250 | 109 | 0 | 35 |
| PBANKA_1032400 | 0 | 0 | 0 | 0 | 14 | 12 | 35 | 11 | 25 | 16 | 9 | 5 | 172 | 46 | 16 | 66 | 182 | 81 | 1807 | 767 | 133 | 956 |
| PBANKA_1033700 | 5 | 6 | 0 | 0 | 24 | 10 | 24 | 17 | 24 | 8 | 0 | 0 | 320 | 64 | 16 | 99 | 256 | 72 | 2439 | 1394 | 11 | 116 |
| PBANKA_1034500 | 165 | 244 | 0 | 0 | 12 | 0 | 24 | 15 | 23 | 10 | 9 | 0 | 34 | 20 | 11 | 21 | 41 | 11 | 71 | 37 | 4288 | 35379 |
| PBANKA_1035200 | 0 | 0 | 0 | 0 | 15 | 5 | 10 | 11 | 37 | 14 | 18 | 0 | 400 | 302 | 40 | 104 | 940 | 414 | 6058 | 10136 | 1611 | 17162 |
| PBANKA_1037800 | 9 | 16 | 0 | 0 | 16 | 8 | 25 | 19 | 18 | 10 | 21 | 10 | 71 | 36 | 0 | 11 | 17 | 12 | 1872 | 975 | 42757 | 3E+05 |
| PBANKA_1038500 | 0 | 0 | 0 | 0 | 23 | 14 | 54 | 18 | 16 | 14 | 0 | 0 | 203 | 70 | 58 | 195 | 273 | 98 | 1341 | 491 | 27 | 169 |
| PBANKA_1038700 | 0 | 0 | 0 | 0 | 8 | 0 | 19 | 16 | 14 | 5 | 0 | 6 | 159 | 79 | 5 | 33 | 177 | 43 | 1227 | 1924 | 14 | 90 |
| PBANKA_1038800 | 0 | 0 | 0 | 0 | 12 | 9 | 12 | 8 | 0 | 7 | 0 | 0 | 319 | 135 | 10 | 25 | 710 | 275 | 7565 | 5089 | 186 | 2485 |
| PBANKA_1039200 | 22 | 43 | 0 | 0 | 46 | 18 | 137 | 76 | 188 | 128 | 36 | 5 | 411 | 104 | 80 | 449 | 3092 | 521 | 411 | 337 | 0 | 38 |
| PBANKA_1040300 | 0 | 0 | 0 | 0 | 5 | 0 | 0 | 0 | 0 | 5 | 0 | 0 | 75 | 0 | 18 | 12 | 6 | 0 | 30 | 17 | 0 | 18 |
| PBANKA_1040561 | 0 | 0 | 0 | 0 | 6 | 0 | 0 | 0 | 0 | 0 | 14 | 13 | 1262 | 783 | 80 | 383 | 58 | 27 | 162 | 135 | 0 | 16 |
| PBANKA_1100700 | 0 | 0 | 0 | 0 | 6 | 0 | 0 | 0 | 9 | 6 | 36 | 19 | 12 | 0 | 6 | 0 | 0 | 11 | 18 | 7 | 0 | 66 |
| PBANKA_1100900 | 0 | 0 | 0 | 0 | 115 | 85 | 142 | 52 | 176 | 183 | 166 | 193 | 1821 | 1309 | 368 | 1363 | 1250 | 327 | 573 | 426 | 18 | 183 |
| PBANKA_1101400 | 0 | 0 | 0 | 0 | 7 | 8 | 606 | 224 | 3870 | 2535 | 375 | 230 | 1967 | 688 | 55 | 90 | 28146 | 9722 | 3329 | 2593 | 52 | 580 |
| PBANKA_1101500 | 0 | 0 | 0 | 0 | 34 | 12 | 46 | 12 | 13 | 9 | 7 | 0 | 115 | 24 | 24 | 47 | 213 | 53 | 1579 | 1130 | 0 | 41 |
| PBANKA_1102100 | 8 | 11 | 0 | 0 | 18 | 5 | 119 | 32 | 502 | 286 | 126 | 38 | 434 | 226 | 34 | 138 | 291 | 126 | 9738 | 6769 | 83 | 420 |
| PBANKA_1104100 | 0 | 0 | 0 | 0 | 38 | 20 | 22 | 6 | 23 | 5 | 6 | 11 | 517 | 270 | 17 | 44 | 109 | 78 | 17976 | 8206 | 89 | 718 |
| PBANKA_1106900 | 0 | 0 | 0 | 0 | 8 | 6 | 7 | 10 | 14 | 0 | 8 | 5 | 34 | 7 | 12 | 16 | 23 | 6 | 147 | 95 | 8580 | 56151 |
| PBANKA_1107000 | 0 | 0 | 0 | 0 | 12 | 5 | 11 | 0 | 10 | 0 | 0 | 0 | 73 | 5 | 0 | 17 | 80 | 32 | 1118 | 388 | 10 | 74 |
| PBANKA_1108800 | 5 | 0 | 0 | 0 | 29 | 19 | 55 | 22 | 16 | 12 | 7 | 0 | 159 | 44 | 47 | 126 | 346 | 93 | 950 | 584 | 73 | 488 |
| PBANKA_1109100 | 16 | 7 | 0 | 0 | 80 | 76 | 164 | 79 | 56 | 22 | 6 | 8 | 128 | 50 | 62 | 169 | 508 | 92 | 65 | 49 | 410 | 3425 |
| PBANKA_1109600 | 9 | 11 | 0 | 0 | 14 | 12 | 15 | 8 | 6 | 11 | 0 | 0 | 583 | 144 | 51 | 113 | 502 | 179 | 4146 | 3801 | 220 | 1708 |
| PBANKA_1109900 | 66 | 63 | 0 | 0 | 13 | 0 | 23 | 10 | 9 | 14 | 6 | 0 | 174 | 104 | 19 | 84 | 143 | 72 | 3541 | 1863 | 1303 | 10510 |
| PBANKA_1112700 | 0 | 0 | 0 | 0 | 14 | 7 | 24 | 9 | 20 | 10 | 8 | 0 | 120 | 57 | 19 | 98 | 131 | 80 | 2338 | 1147 | 232 | 1450 |
| PBANKA_1113700 | 39 | 63 | 0 | 0 | 16 | 15 | 32 | 14 | 28 | 14 | 0 | 5 | 243 | 49 | 68 | 192 | 207 | 89 | 2657 | 1277 | 1241 | 8178 |
| PBANKA_1114100 | 0 | 0 | 0 | 0 | 7 | 0 | 10 | 8 | 7 | 7 | 0 | 0 | 440 | 178 | 9 | 20 | 140 | 84 | 11764 | 4055 | 87 | 573 |
| PBANKA_1117200 | 0 | 0 | 0 | 0 | 41 | 17 | 161 | 50 | 428 | 185 | 67 | 32 | 656 | 290 | 124 | 588 | 3581 | 903 | 651 | 373 | 123 | 695 |
| PBANKA_1119600 | 0 | 0 | 0 | 0 | 9 | 9 | 74 | 20 | 550 | 421 | 222 | 87 | 223 | 107 | 12 | 119 | 1843 | 392 | 292 | 468 | 213 | 1461 |
| PBANKA_1119800 | 6 | 5 | 0 | 0 | 14 | 0 | 21 | 9 | 11 | 6 | 0 | 0 | 230 | 36 | 18 | 21 | 144 | 45 | 1857 | 603 | 43 | 265 |
| PBANKA_1120400 | 21 | 63 | 0 | 0 | 47 | 30 | 53 | 38 | 94 | 40 | 35 | 7 | 2731 | 1781 | 49 | 222 | 1433 | 686 | 66422 | 71784 | 2082 | 19296 |
| PBANKA_1123700 | 71 | 84 | 0 | 0 | 16 | 6 | 36 | 8 | 18 | 0 | 0 | 0 | 92 | 20 | 25 | 78 | 85 | 24 | 1387 | 420 | 22 | 237 |
| PBANKA_1124700 | 0 | 0 | 0 | 0 | 9 | 7 | 12 | 0 | 13 | 5 | 6 | 0 | 79 | 37 | 28 | 46 | 61 | 26 | 982 | 810 | 3092 | 24516 |
| PBANKA_1125300 | 5 | 0 | 0 | 0 | 91 | 32 | 184 | 68 | 62 | 30 | 48 | 12 | 354 | 407 | 78 | 537 | 864 | 151 | 1086 | 6279 | 31 | 209 |
| PBANKA_1125800 | 0 | 0 | 0 | 0 | 8 | 0 | 9 | 0 | 9 | 0 | 7 | 0 | 114 | 15 | 6 | 14 | 44 | 20 | 1479 | 443 | 9 | 42 |
| PBANKA_1127100 | 0 | 0 | 0 | 0 | 13 | 0 | 8 | 0 | 8 | 0 | 0 | 0 | 42 | 26 | 9 | 35 | 69 | 30 | 355 | 188 | 21 | 179 |
| PBANKA_1128000 | 8 | 0 | 0 | 0 | 11 | 10 | 13 | 9 | 12 | 7 | 0 | 0 | 0 | 0 | 0 | 0 | 0 | 0 | 0 | 0 | 0 | 0 |
| PBANKA_1128800 | 5 | 0 | 0 | 0 | 10 | 5 | 26 | 8 | 29 | 24 | 0 | 0 | 394 | 136 | 39 | 73 | 263 | 138 | 7013 | 4125 | 403 | 3154 |
| PBANKA_1129600 | 0 | 0 | 0 | 0 | 17 | 0 | 11 | 0 | 32 | 12 | 5 | 0 | 1989 | 527 | 36 | 230 | 1346 | 416 | 7066 | 6003 | 135 | 649 |
| PBANKA_1132300 | 0 | 0 | 0 | 0 | 46 | 9 | 37 | 16 | 79 | 27 | 23 | 10 | 1147 | 1779 | 144 | 550 | 1723 | 460 | 16504 | 49204 | 339 | 1711 |
| PBANKA_1132900 | 64 | 108 | 0 | 0 | 95 | 41 | 438 | 186 | 558 | 266 | 109 | 44 | 441 | 147 | 163 | 899 | 2884 | 638 | 825 | 514 | 38 | 255 |
| PBANKA_1133600 | 0 | 7 | 0 | 0 | 7 | 6 | 24 | 5 | 7 | 0 | 0 | 0 | 71 | 44 | 19 | 69 | 68 | 16 | 844 | 287 | 7 | 41 |
| PBANKA_1136000 | 0 | 5 | 0 | 0 | 30 | 13 | 44 | 23 | 78 | 49 | 5 | 11 | 65 | 21 | 0 | 6 | 488 | 124 | 213 | 180 | 18 | 132 |
| PBANKA_1138900 | 44 | 51 | 0 | 0 | 18 | 17 | 24 | 6 | 59 | 28 | 17 | 7 | 997 | 971 | 33 | 118 | 526 | 214 | 21993 | 41696 | 61 | 351 |
| PBANKA_1139800 | 0 | 0 | 0 | 0 | 18 | 14 | 41 | 18 | 18 | 20 | 0 | 0 | 234 | 52 | 10 | 19 | 192 | 46 | 2206 | 1322 | 106 | 919 |
| PBANKA_1141900 | 9 | 26 | 0 | 0 | 10 | 0 | 11 | 0 | 34 | 14 | 8 | 0 | 149 | 85 | 11 | 77 | 436 | 82 | 419 | 297 | 616 | 5720 |
| PBANKA_1143200 | 0 | 0 | 0 | 0 | 30 | 6 | 19 | 5 | 12 | 7 | 0 | 0 | 66 | 30 | 61 | 215 | 46 | 13 | 566 | 262 | 7 | 109 |
| PBANKA_1146241 | 0 | 0 | 0 | 0 | 24 | 14 | 38 | 8 | 12 | 0 | 11 | 7 | 254 | 156 | 49 | 282 | 38 | 0 | 40 | 32 | 0 | 0 |
| PBANKA_1146261 | 0 | 0 | 0 | 0 | 42 | 25 | 144 | 67 | 90 | 50 | 21 | 5 | 21 | 8 | 25 | 38 | 39 | 6 | 114 | 51 | 5 | 35 |
| PBANKA_1146600 | 0 | 0 | 0 | 0 | 15 | 5 | 11 | 0 | 9 | 8 | 0 | 0 | 141 | 6 | 34 | 20 | 0 | 9 | 78 | 30 | 0 | 31 |
| PBANKA_1202900 | 14 | 6 | 0 | 0 | 22 | 10 | 17 | 20 | 32 | 24 | 14 | 0 | 72 | 39 | 13 | 49 | 174 | 28 | 164 | 228 | 17 | 147 |
| PBANKA_1204000 | 0 | 10 | 0 | 0 | 20 | 5 | 28 | 10 | 57 | 48 | 8 | 12 | 43 | 16 | 5 | 37 | 294 | 151 | 106 | 55 | 20 | 116 |
| PBANKA_1204200 | 479 | 701 | 0 | 0 | 18 | 15 | 21 | 5 | 20 | 11 | 7 | 0 | 3384 | 1021 | 86 | 357 | 1094 | 339 | 51879 | 37801 | 2817 | 16015 |
| PBANKA_1205300 | 0 | 7 | 0 | 0 | 17 | 7 | 16 | 0 | 10 | 0 | 0 | 0 | 148 | 46 | 27 | 64 | 136 | 44 | 1847 | 944 | 60 | 481 |
| PBANKA_1205500 | 0 | 0 | 0 | 0 | 7 | 0 | 14 | 8 | 164 | 93 | 30 | 19 | 117 | 65 | 20 | 56 | 880 | 237 | 496 | 448 | 38 | 311 |
| PBANKA_1206800 | 239 | 336 | 0 | 0 | 13 | 9 | 6 | 7 | 11 | 0 | 7 | 0 | 59 | 12 | 7 | 26 | 124 | 28 | 286 | 233 | 0 | 33 |

| | | | | | | | | | | | | | | | | | | | | | | |
|---|---|---|---|---|---|---|---|---|---|---|---|---|---|---|---|---|---|---|---|---|---|---|
| PBANKA_1207400 | 5 | 0 | 0 | 0 | 8 | 0 | 0 | 0 | 8 | 0 | 0 | 0 | 31 | 15 | 9 | 23 | 40 | 7 | 819 | 485 | 15 | 158 |
| PBANKA_1208000 | 5 | 0 | 0 | 0 | 12 | 0 | 23 | 8 | 60 | 25 | 14 | 0 | 149 | 58 | 24 | 61 | 524 | 175 | 1677 | 1447 | 295 | 2771 |
| PBANKA_1209400 | 0 | 0 | 0 | 0 | 10 | 8 | 23 | 11 | 11 | 8 | 0 | 0 | 257 | 86 | 7 | 7 | 148 | 107 | 4872 | 1832 | 1096 | 7543 |
| PBANKA_1210600 | 0 | 0 | 0 | 0 | 8 | 6 | 128 | 55 | 628 | 408 | 81 | 59 | 499 | 207 | 22 | 85 | 4000 | 1617 | 533 | 393 | 51 | 452 |
| PBANKA_1212200 | 0 | 0 | 0 | 0 | 8 | 0 | 124 | 30 | 723 | 437 | 167 | 89 | 1059 | 498 | 22 | 85 | 10238 | 2744 | 437 | 743 | 16 | 98 |
| PBANKA_1212400 | 0 | 0 | 0 | 0 | 37 | 6 | 62 | 19 | 208 | 165 | 54 | 13 | 83 | 13 | 0 | 7 | 193 | 106 | 235 | 120 | 21 | 205 |
| PBANKA_1212500 | 0 | 10 | 0 | 0 | 6 | 11 | 104 | 22 | 1356 | 989 | 265 | 151 | 248 | 230 | 31 | 101 | 2287 | 613 | 699 | 1318 | 163 | 1202 |
| PBANKA_1212600 | 0 | 0 | 0 | 0 | 28 | 16 | 23 | 13 | 18 | 10 | 0 | 5 | 192 | 64 | 17 | 45 | 163 | 47 | 1604 | 871 | 9 | 82 |
| PBANKA_1213200 | 0 | 0 | 0 | 0 | 9 | 0 | 12 | 6 | 19 | 5 | 0 | 0 | 232 | 140 | 39 | 143 | 458 | 69 | 1771 | 1848 | 16 | 108 |
| PBANKA_1213300 | 0 | 0 | 0 | 0 | 28 | 27 | 282 | 95 | 931 | 384 | 105 | 70 | 710 | 1175 | 25 | 76 | 18367 | 3190 | 6837 | 22967 | 154 | 1292 |
| PBANKA_1214100 | 0 | 0 | 0 | 0 | 22 | 13 | 22 | 14 | 18 | 9 | 0 | 0 | 101 | 23 | 73 | 375 | 239 | 39 | 1015 | 439 | 20 | 127 |
| PBANKA_1215700 | 7 | 8 | 0 | 0 | 37 | 22 | 52 | 13 | 30 | 11 | 0 | 0 | 193 | 40 | 32 | 115 | 163 | 68 | 1907 | 617 | 64 | 373 |
| PBANKA_1217100 | 0 | 0 | 0 | 0 | 38 | 6 | 261 | 97 | 691 | 400 | 85 | 52 | 307 | 100 | 16 | 23 | 2627 | 755 | 508 | 238 | 26 | 190 |
| PBANKA_1218700 | 13 | 11 | 0 | 0 | 36 | 12 | 69 | 30 | 34 | 23 | 21 | 5 | 165 | 111 | 27 | 173 | 186 | 60 | 1015 | 884 | 107 | 660 |
| PBANKA_1220200 | 7 | 0 | 0 | 0 | 13 | 15 | 34 | 25 | 370 | 316 | 3264 | 2817 | 6980 | 6383 | 1323 | 13413 | 1482 | 256 | 514 | 775 | 9 | 109 |
| PBANKA_1225300 | 16 | 15 | 0 | 0 | 7 | 0 | 26 | 0 | 16 | 6 | 8 | 0 | 284 | 344 | 20 | 86 | 439 | 123 | 4969 | 11244 | 46 | 321 |
| PBANKA_1225400 | 752 | 813 | 0 | 0 | 17 | 8 | 17 | 13 | 280 | 293 | 1468 | 1329 | 107 | 91 | 6 | 5 | 316 | 175 | 103 | 50 | 5241 | 32586 |
| PBANKA_1225500 | 21 | 0 | 0 | 0 | 7 | 0 | 7 | 7 | 7 | 0 | 11 | 9 | 1099 | 630 | 28 | 58 | 490 | 305 | 25927 | 30218 | 815 | 7807 |
| PBANKA_1225600 | 0 | 0 | 0 | 0 | 109 | 56 | 139 | 70 | 34 | 27 | 0 | 5 | 216 | 28 | 131 | 343 | 161 | 37 | 1166 | 516 | 72 | 553 |
| PBANKA_1227500 | 0 | 0 | 0 | 0 | 35 | 21 | 85 | 40 | 29 | 24 | 0 | 0 | 70 | 29 | 6 | 7 | 145 | 40 | 1251 | 475 | 83 | 515 |
| PBANKA_1228800 | 0 | 0 | 0 | 0 | 24 | 17 | 34 | 14 | 252 | 223 | 47 | 28 | 247 | 148 | 39 | 120 | 1830 | 682 | 146 | 194 | 5 | 24 |
| PBANKA_1228900 | 0 | 0 | 0 | 0 | 13 | 13 | 13 | 8 | 7 | 5 | 16 | 0 | 333 | 115 | 13 | 33 | 72 | 50 | 8541 | 5273 | 20137 | 2E+05 |
| PBANKA_1230200 | 14 | 18 | 0 | 0 | 85 | 62 | 331 | 153 | 380 | 154 | 61 | 23 | 891 | 231 | 69 | 115 | 5045 | 486 | 2499 | 2131 | 135 | 1163 |
| PBANKA_1232000 | 7 | 0 | 0 | 0 | 15 | 10 | 29 | 16 | 21 | 14 | 44 | 16 | 22 | 44 | 15 | 65 | 188 | 27 | 138 | 314 | 1209 | 9075 |
| PBANKA_1232300 | 5 | 0 | 0 | 0 | 19 | 7 | 17 | 7 | 10 | 7 | 7 | 7 | 218 | 188 | 36 | 113 | 170 | 90 | 2374 | 2018 | 366 | 3045 |
| PBANKA_1232600 | 0 | 0 | 0 | 0 | 108 | 38 | 180 | 61 | 97 | 46 | 11 | 9 | 228 | 95 | 10 | 34 | 123 | 53 | 5454 | 3949 | 95 | 768 |
| PBANKA_1233500 | 1051 | 1598 | 0 | 0 | 56 | 20 | 33 | 29 | 94 | 39 | 23 | 5 | 326 | 282 | 29 | 91 | 319 | 97 | 3293 | 6810 | 1758 | 11311 |
| PBANKA_1233600 | 6 | 0 | 0 | 0 | 7 | 0 | 5 | 0 | 7 | 0 | 0 | 0 | 760 | 258 | 19 | 35 | 181 | 118 | 21329 | 12160 | 111 | 947 |
| PBANKA_1233700 | 869 | 1189 | 0 | 0 | 10 | 6 | 6 | 0 | 10 | 6 | 0 | 0 | 61 | 24 | 8 | 20 | 26 | 26 | 633 | 400 | 1822 | 17104 |
| PBANKA_1236200 | 23 | 31 | 0 | 0 | 8 | 0 | 0 | 0 | 0 | 0 | 0 | 0 | 36 | 12 | 0 | 10 | 32 | 8 | 398 | 247 | 5 | 43 |
| PBANKA_1237300 | 0 | 0 | 0 | 0 | 97 | 45 | 202 | 83 | 175 | 111 | 24 | 19 | 475 | 118 | 35 | 111 | 3557 | 881 | 1185 | 593 | 206 | 1759 |
| PBANKA_1238300 | 0 | 0 | 0 | 0 | 38 | 30 | 42 | 29 | 20 | 16 | 0 | 0 | 32 | 0 | 18 | 55 | 75 | 11 | 435 | 145 | 19 | 77 |
| PBANKA_1239300 | 0 | 14 | 0 | 0 | 23 | 6 | 30 | 13 | 6 | 0 | 6 | 0 | 168 | 44 | 35 | 184 | 69 | 18 | 229 | 72 | 22 | 93 |
| PBANKA_1240700 | 0 | 0 | 0 | 0 | 20 | 0 | 27 | 7 | 24 | 5 | 0 | 0 | 89 | 30 | 13 | 53 | 46 | 29 | 2199 | 987 | 59 | 470 |
| PBANKA_1243000 | 23 | 37 | 0 | 0 | 24 | 10 | 58 | 20 | 167 | 79 | 32 | 8 | 301 | 122 | 6 | 0 | 1780 | 615 | 2177 | 1150 | 825 | 4033 |
| PBANKA_1243500 | 0 | 0 | 0 | 0 | 17 | 10 | 30 | 20 | 0 | 6 | 0 | 0 | 375 | 140 | 21 | 79 | 193 | 85 | 5689 | 3144 | 282 | 1322 |
| PBANKA_1245900 | 0 | 0 | 0 | 0 | 7 | 9 | 34 | 19 | 94 | 43 | 624 | 407 | 3017 | 1818 | 772 | 4207 | 891 | 220 | 1731 | 791 | 73 | 449 |
| PBANKA_1246141 | 0 | 0 | 0 | 0 | 6 | 0 | 8 | 0 | 0 | 11 | 155 | 29 | 212 | 101 | 60 | 165 | 173 | 20 | 6 | 0 | 0 | 0 |
| PBANKA_1246161 | 0 | 0 | 0 | 0 | 10 | 14 | 15 | 0 | 179 | 168 | 3189 | 2433 | 2984 | 2028 | 717 | 3270 | 571 | 143 | 2452 | 1525 | 71 | 483 |
| PBANKA_1246500 | 0 | 0 | 0 | 0 | 45 | 12 | 105 | 35 | 58 | 31 | 9 | 0 | 104 | 34 | 38 | 63 | 26 | 6 | 76 | 37 | 5 | 22 |
| PBANKA_1246700 | 0 | 0 | 0 | 0 | 5 | 0 | 0 | 0 | 9 | 0 | 0 | 0 | 195 | 79 | 24 | 113 | 32 | 0 | 407 | 164 | 0 | 12 |
| PBANKA_1300031 | 0 | 0 | 0 | 0 | 12 | 0 | 8 | 0 | 17 | 8 | 201 | 133 | 118 | 30 | 19 | 30 | 11 | 8 | 44 | 9 | 0 | 33 |
| PBANKA_1300500 | 0 | 0 | 0 | 0 | 9 | 0 | 19 | 5 | 5 | 0 | 0 | 0 | 214 | 67 | 13 | 67 | 137 | 54 | 1632 | 861 | 5 | 50 |
| PBANKA_1301800 | 14 | 21 | 0 | 0 | 70 | 44 | 181 | 54 | 254 | 71 | 29 | 12 | 382 | 356 | 153 | 624 | 1404 | 293 | 2081 | 6725 | 133 | 776 |
| PBANKA_1308100 | 0 | 0 | 0 | 0 | 9 | 0 | 13 | 0 | 0 | 0 | 5 | 0 | 270 | 17 | 6 | 14 | 121 | 31 | 2880 | 650 | 10 | 63 |
| PBANKA_1311300 | 0 | 8 | 0 | 0 | 25 | 16 | 47 | 18 | 13 | 9 | 0 | 6 | 57 | 24 | 6 | 38 | 24 | 13 | 1364 | 590 | 893 | 7558 |
| PBANKA_1311900 | 0 | 0 | 0 | 0 | 30 | 6 | 22 | 8 | 14 | 0 | 10 | 6 | 164 | 77 | 51 | 225 | 115 | 17 | 189 | 89 | 48 | 193 |
| PBANKA_1312000 | 7 | 0 | 0 | 0 | 31 | 15 | 55 | 26 | 34 | 20 | 5 | 0 | 214 | 57 | 18 | 93 | 261 | 80 | 1983 | 1312 | 471 | 2340 |
| PBANKA_1312700 | 6297 | 9321 | 0 | 0 | 8 | 0 | 16 | 13 | 9 | 5 | 0 | 0 | 345 | 146 | 14 | 36 | 626 | 404 | 4098 | 2042 | 4574 | 39542 |
| PBANKA_1312900 | 0 | 0 | 0 | 0 | 61 | 12 | 176 | 64 | 928 | 555 | 113 | 72 | 927 | 409 | 6 | 34 | 5923 | 3183 | 7668 | 3248 | 37 | 352 |
| PBANKA_1313300 | 0 | 0 | 0 | 0 | 26 | 8 | 87 | 24 | 107 | 61 | 17 | 7 | 304 | 158 | 27 | 120 | 1897 | 431 | 2030 | 1150 | 59 | 411 |
| PBANKA_1316800 | 0 | 8 | 0 | 0 | 65 | 51 | 67 | 25 | 37 | 26 | 0 | 0 | 194 | 49 | 46 | 113 | 456 | 86 | 1576 | 867 | 23 | 307 |
| PBANKA_1318000 | 7 | 7 | 0 | 0 | 45 | 18 | 47 | 13 | 33 | 10 | 10 | 10 | 312 | 191 | 34 | 140 | 220 | 57 | 373 | 240 | 23 | 176 |
| PBANKA_1318500 | 0 | 0 | 0 | 0 | 6 | 5 | 0 | 8 | 19 | 9 | 0 | 0 | 917 | 487 | 19 | 49 | 187 | 148 | 27660 | 11123 | 40 | 314 |
| PBANKA_1318600 | 17 | 21 | 0 | 0 | 90 | 50 | 71 | 53 | 68 | 85 | 7 | 7 | 1365 | 706 | 37 | 62 | 360 | 388 | 49311 | 19671 | 1921 | 21591 |
| PBANKA_1318700 | 407 | 513 | 0 | 0 | 18 | 0 | 26 | 18 | 25 | 15 | 0 | 0 | 78 | 15 | 0 | 5 | 61 | 27 | 945 | 464 | 63 | 489 |
| PBANKA_1318800 | 0 | 0 | 0 | 0 | 62 | 44 | 51 | 43 | 16 | 15 | 0 | 0 | 142 | 54 | 10 | 14 | 85 | 47 | 3829 | 1881 | 33 | 307 |
| PBANKA_1320100 | 543 | 809 | 0 | 0 | 59 | 15 | 131 | 32 | 80 | 30 | 17 | 7 | 360 | 154 | 8 | 51 | 365 | 90 | 4679 | 4168 | 47 | 370 |
| PBANKA_1321500 | 0 | 0 | 0 | 0 | 21 | 0 | 63 | 26 | 34 | 13 | 0 | 0 | 242 | 61 | 61 | 386 | 427 | 76 | 2407 | 1713 | 45 | 305 |
| PBANKA_1323500 | 0 | 0 | 0 | 0 | 10 | 0 | 8 | 9 | 30 | 19 | 10 | 0 | 56 | 31 | 9 | 24 | 431 | 151 | 196 | 154 | 10 | 102 |
| PBANKA_1323600 | 0 | 0 | 0 | 0 | 35 | 20 | 25 | 9 | 30 | 30 | 7 | 0 | 86 | 22 | 22 | 36 | 224 | 77 | 467 | 276 | 84 | 712 |
| PBANKA_1323800 | 0 | 7 | 0 | 0 | 43 | 11 | 44 | 13 | 6 | 9 | 6 | 8 | 654 | 305 | 73 | 185 | 230 | 105 | 18941 | 15057 | 51 | 453 |
| PBANKA_1325100 | 60 | 109 | 0 | 0 | 11 | 9 | 20 | 8 | 100 | 46 | 15 | 5 | 213 | 81 | 19 | 50 | 309 | 103 | 2159 | 1648 | 88 | 621 |
| PBANKA_1325200 | 0 | 0 | 0 | 0 | 15 | 6 | 6 | 0 | 0 | 0 | 0 | 0 | 193 | 51 | 17 | 11 | 52 | 44 | 1696 | 1057 | 96 | 678 |
| PBANKA_1325700 | 0 | 0 | 0 | 0 | 5 | 0 | 9 | 0 | 0 | 0 | 0 | 0 | 40 | 19 | 18 | 39 | 27 | 19 | 686 | 224 | 6 | 51 |
| PBANKA_1326100 | 7 | 11 | 0 | 0 | 17 | 9 | 17 | 9 | 5 | 0 | 0 | 0 | 701 | 181 | 53 | 148 | 492 | 107 | 3721 | 1754 | 48 | 366 |
| PBANKA_1327100 | 0 | 0 | 0 | 0 | 17 | 19 | 136 | 62 | 655 | 378 | 119 | 72 | 1061 | 352 | 70 | 158 | 7196 | 2291 | 807 | 461 | 13 | 150 |
| PBANKA_1328200 | 0 | 0 | 0 | 0 | 15 | 0 | 9 | 8 | 6 | 0 | 0 | 0 | 63 | 15 | 21 | 93 | 109 | 23 | 489 | 149 | 7 | 49 |
| PBANKA_1328900 | 0 | 5 | 0 | 0 | 44 | 24 | 58 | 33 | 83 | 34 | 21 | 0 | 167 | 99 | 75 | 231 | 311 | 68 | 1709 | 1363 | 11 | 91 |
| PBANKA_1329000 | 0 | 6 | 0 | 0 | 6 | 0 | 12 | 0 | 13 | 7 | 6 | 0 | 330 | 77 | 60 | 254 | 233 | 68 | 2339 | 1586 | 5 | 53 |
| PBANKA_1329100 | 6 | 5 | 0 | 0 | 77 | 53 | 96 | 18 | 60 | 51 | 13 | 5 | 741 | 306 | 70 | 198 | 372 | 150 | 20107 | 12649 | 37 | 247 |
| PBANKA_1329400 | 0 | 0 | 0 | 0 | 14 | 13 | 21 | 6 | 6 | 0 | 0 | 6 | 552 | 232 | 15 | 37 | 129 | 84 | 10122 | 3940 | 133 | 1143 |
| PBANKA_1329900 | 0 | 0 | 0 | 0 | 20 | 10 | 21 | 7 | 0 | 0 | 0 | 0 | 182 | 88 | 5 | 17 | 79 | 46 | 4280 | 1212 | 280 | 1134 |
| PBANKA_1331600 | 7 | 12 | 0 | 0 | 19 | 6 | 8 | 6 | 11 | 0 | 0 | 0 | 99 | 34 | 13 | 41 | 97 | 47 | 930 | 297 | 6 | 65 |
| PBANKA_1331700 | 301 | 398 | 0 | 0 | 11 | 0 | 15 | 17 | 470 | 367 | 209 | 131 | 251 | 177 | 29 | 64 | 2143 | 475 | 984 | 683 | 81 | 936 |
| PBANKA_1332300 | 0 | 0 | 0 | 0 | 27 | 17 | 51 | 19 | 44 | 0 | 12 | 0 | 276 | 131 | 65 | 220 | 344 | 54 | 431 | 285 | 53 | 369 |
| PBANKA_1333100 | 0 | 0 | 0 | 0 | 23 | 0 | 22 | 9 | 5 | 0 | 0 | 0 | 585 | 201 | 60 | 274 | 368 | 71 | 3744 | 4062 | 34 | 177 |
| PBANKA_1333700 | 0 | 6 | 0 | 0 | 6 | 0 | 22 | 7 | 25 | 8 | 16 | 0 | 1663 | 765 | 67 | 197 | 664 | 350 | 35331 | 26062 | 1070 | 7270 |
| PBANKA_1334900 | 0 | 0 | 0 | 0 | 17 | 13 | 21 | 7 | 13 | 7 | 6 | 0 | 1179 | 1445 | 145 | 494 | 1144 | 420 | 26128 | 48078 | 2399 | 18253 |
| PBANKA_1335800 | 15 | 22 | 0 | 0 | 35 | 19 | 63 | 28 | 27 | 19 | 0 | 0 | 211 | 34 | 91 | 391 | 852 | 168 | 917 | 418 | 14 | 146 |
| PBANKA_1336600 | 113 | 179 | 0 | 0 | 12 | 6 | 42 | 6 | 318 | 256 | 64 | 30 | 428 | 183 | 62 | 121 | 685 | 273 | 2751 | 2921 | 6806 | 51912 |
| PBANKA_1337700 | 190 | 228 | 0 | 0 | 24 | 13 | 39 | 18 | 27 | 9 | 6 | 0 | 148 | 37 | 50 | 295 | 348 | 54 | 1471 | 1223 | 128 | 602 |
| PBANKA_1338100 | 0 | 0 | 0 | 0 | 24 | 12 | 93 | 49 | 64 | 34 | 8 | 0 | 668 | 240 | 12 | 50 | 156 | 119 | 15644 | 5449 | 290 | 1401 |
| PBANKA_1340400 | 0 | 0 | 0 | 0 | 31 | 9 | 16 | 5 | 15 | 5 | 8 | 0 | 309 | 90 | 41 | 106 | 581 | 207 | 2683 | 904 | 15 | 181 |
| PBANKA_1341400 | 0 | 0 | 0 | 0 | 14 | 5 | 29 | 12 | 18 | 0 | 0 | 0 | 170 | 38 | 24 | 47 | 114 | 62 | 1321 | 489 | 41 | 262 |
| PBANKA_1344200 | 0 | 7 | 0 | 0 | 17 | 0 | 55 | 23 | 56 | 14 | 11 | 0 | 383 | 83 | 18 | 99 | 361 | 78 | 3023 | 1586 | 89 | 738 |
| PBANKA_1349000 | 0 | 0 | 0 | 0 | 426 | 235 | 7769 | 4336 | 7276 | 4714 | 886 | 460 | 7312 | 1923 | 49 | 154 | 58971 | 9451 | 585 | 575 | 6 | 109 |
| PBANKA_1349100 | 6 | 0 | 0 | 0 | 11 | 5 | 798 | 342 | 3534 | 2570 | 580 | 351 | 3391 | 1239 | 108 | 283 | 42003 | 11038 | 713 | 898 | 20 | 221 |
| PBANKA_1351500 | 0 | 0 | 0 | 0 | 12 | 6 | 46 | 16 | 277 | 121 | 605 | 341 | 397 | 181 | 15 | 26 | 1073 | 538 | 356 | 337 | 143 | 852 |
| PBANKA_1353600 | 0 | 0 | 0 | 0 | 14 | 5 | 15 | 0 | 6 | 0 | 6 | 0 | 24 | 15 | 16 | 54 | 22 | 0 | 7 | 14 | 0 | 5 |

| | | | | | | | | | | | | | | | | | | | | | | |
|---|---|---|---|---|---|---|---|---|---|---|---|---|---|---|---|---|---|---|---|---|---|---|
| PBANKA_1353900 | 0 | 5 | 0 | 0 | 19 | 6 | 19 | 7 | 93 | 32 | 26 | 0 | 333 | 288 | 39 | 164 | 777 | 228 | 2837 | 4442 | 133 | 1021 |
| PBANKA_1354600 | 0 | 0 | 0 | 0 | 15 | 14 | 27 | 10 | 35 | 19 | 19 | 7 | 69 | 40 | 35 | 112 | 13 | 7 | 93 | 72 | 1301 | 11444 |
| PBANKA_1354800 | 0 | 0 | 0 | 0 | 13 | 0 | 21 | 14 | 18 | 8 | 5 | 0 | 126 | 34 | 20 | 77 | 197 | 90 | 1066 | 427 | 41 | 177 |
| PBANKA_1354900 | 0 | 0 | 0 | 0 | 49 | 17 | 51 | 26 | 31 | 19 | 7 | 0 | 1698 | 1220 | 82 | 168 | 558 | 337 | 45722 | 38537 | 3355 | 32216 |
| PBANKA_1356600 | 0 | 0 | 0 | 0 | 40 | 11 | 27 | 15 | 13 | 10 | 11 | 0 | 107 | 30 | 139 | 516 | 196 | 42 | 168 | 51 | 0 | 7 |
| PBANKA_1358500 | 28 | 36 | 0 | 0 | 32 | 6 | 34 | 19 | 312 | 231 | 66 | 42 | 137 | 94 | 6 | 13 | 1130 | 411 | 382 | 225 | 639 | 3662 |
| PBANKA_1360400 | 0 | 0 | 0 | 0 | 30 | 6 | 17 | 10 | 6 | 0 | 0 | 0 | 455 | 170 | 38 | 218 | 216 | 78 | 8380 | 3900 | 150 | 1235 |
| PBANKA_1360600 | 8 | 5 | 0 | 0 | 17 | 8 | 40 | 22 | 16 | 0 | 0 | 0 | 112 | 44 | 106 | 286 | 145 | 41 | 763 | 238 | 14 | 123 |
| PBANKA_1360900 | 0 | 0 | 0 | 0 | 30 | 16 | 104 | 76 | 36 | 47 | 0 | 0 | 155 | 26 | 5 | 0 | 649 | 92 | 404 | 310 | 7 | 72 |
| PBANKA_1361400 | 73 | 93 | 0 | 0 | 24 | 12 | 79 | 29 | 184 | 86 | 16 | 0 | 291 | 89 | 12 | 35 | 2162 | 658 | 1102 | 335 | 92 | 586 |
| PBANKA_1362800 | 0 | 0 | 0 | 0 | 22 | 9 | 20 | 5 | 9 | 5 | 0 | 0 | 82 | 34 | 6 | 37 | 65 | 27 | 1159 | 641 | 37 | 173 |
| PBANKA_1363600 | 18 | 22 | 0 | 0 | 7 | 0 | 18 | 0 | 6 | 0 | 7 | 0 | 1282 | 549 | 28 | 52 | 298 | 194 | 41212 | 21794 | 585 | 3662 |
| PBANKA_1364400 | 19 | 40 | 0 | 0 | 52 | 45 | 109 | 42 | 1042 | 629 | 351 | 176 | 776 | 844 | 150 | 195 | 5278 | 1418 | 6722 | 13596 | 17175 | 1E+05 |
| PBANKA_1365721 | 30 | 22 | 0 | 0 | 17 | 10 | 19 | 19 | 20 | 11 | 10 | 0 | 41 | 66 | 102 | 310 | 343 | 60 | 95 | 338 | 77 | 580 |
| PBANKA_1400081 | 0 | 0 | 0 | 0 | 7 | 0 | 7 | 0 | 6 | 7 | 6 | 0 | 197 | 83 | 35 | 179 | 49 | 10 | 204 | 118 | 0 | 18 |
| PBANKA_1400096 | 0 | 6 | 0 | 0 | 6 | 0 | 83 | 30 | 31 | 26 | 6 | 0 | 33 | 11 | 0 | 0 | 104 | 25 | 8 | 0 | 0 | 0 |
| PBANKA_1400400 | 0 | 0 | 0 | 0 | 5 | 0 | 23 | 0 | 8 | 0 | 0 | 5 | 4672 | 77 | 164 | 42 | 33 | 90 | 544 | 31 | 43 | 293 |
| PBANKA_1400700 | 0 | 0 | 0 | 0 | 104 | 69 | 1077 | 708 | 1122 | 830 | 3965 | 3456 | 13525 | 8106 | 3423 | 8656 | 13682 | 3899 | 3831 | 1593 | 320 | 3058 |
| PBANKA_1404300 | 0 | 0 | 0 | 0 | 5 | 0 | 9 | 0 | 6 | 0 | 0 | 0 | 22 | 13 | 14 | 34 | 40 | 13 | 245 | 154 | 0 | 48 |
| PBANKA_1405000 | 0 | 8 | 0 | 0 | 68 | 38 | 99 | 39 | 98 | 46 | 20 | 0 | 586 | 241 | 46 | 160 | 834 | 175 | 5132 | 5483 | 180 | 1168 |
| PBANKA_1406500 | 0 | 0 | 0 | 0 | 21 | 10 | 107 | 50 | 450 | 282 | 54 | 30 | 351 | 175 | 30 | 81 | 2992 | 805 | 1729 | 1118 | 91 | 825 |
| PBANKA_1407650 | 0 | 0 | 0 | 0 | 8 | 0 | 9 | 0 | 5 | 0 | 34 | 17 | 104 | 92 | 32 | 29 | 26 | 7 | 15 | 28 | 0 | 0 |
| PBANKA_1408700 | 12 | 24 | 0 | 0 | 8 | 0 | 34 | 18 | 138 | 87 | 7 | 12 | 78 | 13 | 0 | 10 | 424 | 348 | 124 | 42 | 5 | 21 |
| PBANKA_1410950 | 10 | 13 | 0 | 0 | 7 | 0 | 7 | 0 | 10 | 9 | 11 | 0 | 9 | 15 | 0 | 10 | 36 | 30 | 217 | 134 | 5253 | 40349 |
| PBANKA_1412000 | 0 | 0 | 0 | 0 | 16 | 0 | 84 | 29 | 326 | 147 | 71 | 25 | 250 | 208 | 30 | 77 | 3635 | 788 | 1192 | 1794 | 325 | 2171 |
| PBANKA_1413300 | 8 | 0 | 0 | 0 | 26 | 12 | 37 | 15 | 22 | 5 | 0 | 0 | 97 | 19 | 31 | 114 | 145 | 43 | 649 | 305 | 10 | 59 |
| PBANKA_1413500 | 19 | 14 | 0 | 0 | 33 | 21 | 29 | 10 | 112 | 77 | 46 | 21 | 313 | 126 | 154 | 483 | 176 | 81 | 571 | 256 | 73 | 340 |
| PBANKA_1414500 | 6 | 10 | 0 | 0 | 15 | 5 | 17 | 6 | 0 | 0 | 0 | 0 | 749 | 353 | 26 | 36 | 195 | 139 | 17156 | 9272 | 214 | 1175 |
| PBANKA_1416500 | 0 | 0 | 0 | 0 | 10 | 9 | 37 | 18 | 151 | 79 | 15 | 5 | 111 | 19 | 0 | 37 | 738 | 281 | 193 | 142 | 0 | 65 |
| PBANKA_1417000 | 0 | 0 | 0 | 0 | 10 | 6 | 15 | 10 | 16 | 5 | 0 | 0 | 40 | 0 | 0 | 6 | 65 | 11 | 333 | 290 | 0 | 51 |
| PBANKA_1417200 | 42 | 79 | 0 | 0 | 16 | 5 | 40 | 19 | 629 | 380 | 224 | 85 | 1267 | 955 | 51 | 119 | 2640 | 987 | 23926 | 32859 | 743 | 5940 |
| PBANKA_1417700 | 0 | 0 | 0 | 0 | 7 | 5 | 0 | 6 | 7 | 6 | 0 | 0 | 108 | 86 | 102 | 411 | 133 | 31 | 262 | 150 | 5 | 21 |
| PBANKA_1418000 | 0 | 0 | 0 | 0 | 9 | 0 | 13 | 0 | 0 | 0 | 0 | 0 | 38 | 0 | 36 | 109 | 22 | 13 | 203 | 71 | 14 | 69 |
| PBANKA_1419400 | 0 | 0 | 0 | 0 | 9 | 11 | 17 | 0 | 37 | 0 | 17 | 0 | 45 | 23 | 27 | 49 | 338 | 62 | 419 | 192 | 7 | 32 |
| PBANKA_1420700 | 0 | 0 | 0 | 0 | 44 | 25 | 135 | 37 | 97 | 35 | 35 | 10 | 628 | 443 | 23 | 229 | 565 | 117 | 3189 | 5772 | 42 | 359 |
| PBANKA_1421000 | 0 | 0 | 0 | 0 | 17 | 8 | 29 | 16 | 8 | 0 | 0 | 0 | 528 | 145 | 39 | 294 | 368 | 74 | 3745 | 2488 | 10 | 77 |
| PBANKA_1421400 | 0 | 0 | 0 | 0 | 9 | 0 | 6 | 0 | 0 | 0 | 0 | 0 | 94 | 12 | 23 | 63 | 54 | 42 | 675 | 453 | 0 | 50 |
| PBANKA_1422900 | 1015 | 1324 | 0 | 0 | 10 | 0 | 6 | 9 | 18 | 10 | 17 | 0 | 1040 | 318 | 40 | 76 | 249 | 164 | 28841 | 13974 | 9655 | 84591 |
| PBANKA_1425200 | 1078 | 1796 | 0 | 0 | 41 | 12 | 78 | 39 | 82 | 42 | 9 | 5 | 95 | 30 | 29 | 103 | 432 | 70 | 560 | 441 | 173 | 1559 |
| PBANKA_1427600 | 0 | 0 | 0 | 0 | 32 | 0 | 248 | 136 | 263 | 126 | 34 | 17 | 217 | 88 | 38 | 155 | 779 | 177 | 1830 | 1636 | 45 | 279 |
| PBANKA_1429400 | 102 | 115 | 0 | 0 | 41 | 21 | 122 | 54 | 62 | 43 | 6 | 5 | 171 | 29 | 62 | 70 | 581 | 176 | 1137 | 317 | 39 | 295 |
| PBANKA_1430300 | 0 | 0 | 0 | 0 | 10 | 9 | 15 | 9 | 5 | 0 | 0 | 0 | 494 | 228 | 14 | 25 | 115 | 73 | 17188 | 7660 | 128 | 1040 |
| PBANKA_1434300 | 5 | 12 | 0 | 0 | 18 | 6 | 18 | 9 | 20 | 5 | 0 | 0 | 90 | 44 | 39 | 104 | 101 | 30 | 740 | 667 | 69 | 690 |
| PBANKA_1434400 | 7 | 8 | 0 | 0 | 14 | 6 | 11 | 0 | 11 | 12 | 0 | 6 | 167 | 36 | 30 | 107 | 187 | 103 | 2940 | 1329 | 120 | 912 |
| PBANKA_1434900 | 26 | 42 | 0 | 0 | 9 | 0 | 9 | 6 | 5 | 0 | 0 | 0 | 36 | 21 | 0 | 10 | 53 | 24 | 676 | 530 | 79 | 582 |
| PBANKA_1435200 | 8 | 5 | 0 | 0 | 86 | 25 | 118 | 56 | 46 | 22 | 14 | 0 | 122 | 63 | 105 | 237 | 199 | 99 | 3129 | 1761 | 2487 | 19006 |
| PBANKA_1435400 | 5 | 0 | 0 | 0 | 150 | 63 | 279 | 89 | 168 | 50 | 22 | 10 | 245 | 123 | 463 | 1725 | 1427 | 236 | 1090 | 1593 | 34 | 189 |
| PBANKA_1436600 | 0 | 0 | 0 | 0 | 5 | 5 | 8 | 9 | 21 | 8 | 15 | 0 | 3308 | 1479 | 55 | 165 | 933 | 617 | 89470 | 60526 | 176 | 2036 |
| PBANKA_1437500 | 0 | 0 | 0 | 0 | 150 | 86 | 300 | 105 | 142 | 96 | 59 | 14 | 2129 | 3345 | 309 | 1371 | 637 | 91 | 1064 | 2854 | 31 | 279 |
| PBANKA_1437600 | 93 | 145 | 0 | 0 | 12 | 11 | 130 | 67 | 1115 | 1011 | 310 | 167 | 1946 | 1024 | 40 | 81 | 3174 | 1374 | 58075 | 37990 | 2952 | 31449 |
| PBANKA_1437700 | 0 | 0 | 0 | 0 | 28 | 6 | 32 | 10 | 102 | 71 | 38 | 14 | 1087 | 482 | 38 | 241 | 369 | 236 | 26808 | 13685 | 441 | 2659 |
| PBANKA_1439000 | 32 | 32 | 0 | 0 | 26 | 25 | 75 | 32 | 98 | 61 | 9 | 8 | 128 | 22 | 9 | 20 | 757 | 306 | 371 | 119 | 230 | 1431 |
| PBANKA_1441800 | 0 | 0 | 0 | 0 | 17 | 7 | 80 | 48 | 69 | 34 | 10 | 0 | 189 | 58 | 7 | 18 | 976 | 213 | 933 | 289 | 44 | 285 |
| PBANKA_1445000 | 78 | 89 | 0 | 0 | 18 | 8 | 7 | 0 | 35 | 23 | 0 | 0 | 341 | 139 | 24 | 85 | 136 | 62 | 7670 | 5794 | 1429 | 11650 |
| PBANKA_1446000 | 21 | 16 | 0 | 0 | 8 | 6 | 9 | 0 | 15 | 0 | 6 | 0 | 105 | 35 | 28 | 200 | 144 | 56 | 1590 | 1497 | 11 | 60 |
| PBANKA_1449000 | 6 | 10 | 0 | 0 | 19 | 8 | 24 | 13 | 33 | 11 | 17 | 0 | 855 | 196 | 54 | 183 | 948 | 436 | 11098 | 3711 | 119 | 1190 |
| PBANKA_1449600 | 44 | 44 | 0 | 0 | 12 | 11 | 28 | 11 | 20 | 0 | 0 | 0 | 62 | 5 | 21 | 28 | 29 | 22 | 498 | 251 | 76 | 547 |
| PBANKA_1451100 | 0 | 5 | 0 | 0 | 12 | 0 | 0 | 9 | 14 | 0 | 0 | 0 | 1018 | 357 | 16 | 61 | 300 | 189 | 27259 | 21184 | 2664 | 19548 |
| PBANKA_1451500 | 0 | 0 | 0 | 0 | 130 | 73 | 383 | 177 | 353 | 189 | 44 | 12 | 314 | 73 | 12 | 26 | 1534 | 451 | 1790 | 720 | 158 | 1402 |
| PBANKA_1453900 | 0 | 0 | 0 | 0 | 8 | 0 | 19 | 7 | 15 | 5 | 0 | 0 | 152 | 70 | 0 | 28 | 80 | 53 | 3828 | 1941 | 54 | 433 |
| PBANKA_1456700 | 117 | 217 | 0 | 0 | 63 | 21 | 79 | 16 | 27 | 14 | 0 | 0 | 7 | 0 | 0 | 0 | 0 | 0 | 44 | 21 | 336 | 1174 |
| PBANKA_1457000 | 7 | 5 | 0 | 0 | 41 | 32 | 152 | 51 | 317 | 202 | 81 | 48 | 570 | 213 | 58 | 195 | 2594 | 593 | 699 | 501 | 61 | 615 |
| PBANKA_1457200 | 20 | 35 | 0 | 0 | 200 | 73 | 834 | 355 | 638 | 329 | 71 | 45 | 1512 | 1201 | 200 | 1052 | 19308 | 2672 | 2485 | 9118 | 165 | 1300 |
| PBANKA_1460000 | 253 | 399 | 0 | 0 | 8 | 5 | 34 | 31 | 25 | 9 | 9 | 6 | 72 | 18 | 15 | 37 | 218 | 68 | 245 | 140 | 2642 | 14210 |
| PBANKA_1462100 | 0 | 0 | 0 | 0 | 25 | 14 | 46 | 16 | 18 | 6 | 0 | 0 | 209 | 57 | 72 | 232 | 272 | 32 | 955 | 252 | 88 | 390 |
| PBANKA_1463300 | 6 | 8 | 0 | 0 | 17 | 9 | 44 | 16 | 71 | 38 | 65 | 28 | 8820 | 3891 | 116 | 662 | 2332 | 920 | 1E+05 | 1E+05 | 318 | 1112 |
| PBANKA_1464900 | 0 | 0 | 0 | 0 | 224 | 87 | 1466 | 606 | 1688 | 853 | 216 | 135 | 963 | 745 | 62 | 180 | 17138 | 3842 | 2564 | 3111 | 352 | 3426 |
| PBANKA_1465000 | 7 | 8 | 0 | 0 | 13 | 0 | 21 | 0 | 346 | 210 | 2399 | 2064 | 381 | 1199 | 18 | 40 | 690 | 182 | 281 | 338 | 52 | 610 |
| PBANKA_1465600 | 0 | 0 | 0 | 0 | 7 | 14 | 21 | 12 | 26 | 21 | 144 | 101 | 77 | 81 | 19 | 117 | 45 | 0 | 114 | 125 | 0 | 5 |
| PBANKA_1465700 | 0 | 0 | 0 | 0 | 8 | 0 | 19 | 0 | 5 | 0 | 0 | 0 | 0 | 0 | 0 | 0 | 0 | 0 | 0 | 0 | 13 | 110 |
| PBANKA_1466000 | 0 | 0 | 0 | 0 | 7 | 0 | 7 | 0 | 10 | 0 | 78 | 64 | 177 | 113 | 49 | 273 | 123 | 7 | 74 | 37 | 0 | 0 |
| PBANKA_API00055 | 0 | 8 | 0 | 0 | 12 | 5 | 30 | 0 | 16 | 0 | 40 | 0 | 65 | 6 | 30 | 14 | 68 | 23 | 133 | 24 | 0 | 0 |
| PBANKA_API00120 | 0 | 0 | 0 | 0 | 11 | 0 | 18 | 0 | 14 | 6 | 0 | 0 | 8 | 0 | 0 | 9 | 12 | 0 | 5 | 0 | 0 | 0 |
| PBANKA_API00150 | 0 | 0 | 0 | 0 | 16 | 5 | 50 | 8 | 36 | 9 | 5 | 0 | 22 | 5 | 0 | 8 | 19 | 0 | 6 | 0 | 0 | 0 |
| PBANKA_API00160 | 0 | 0 | 0 | 0 | 5 | 0 | 7 | 8 | 18 | 0 | 0 | 0 | 0 | 0 | 0 | 0 | 0 | 0 | 6 | 0 | 0 | 0 |
| PBANKA_API00206 | 0 | 0 | 0 | 0 | 9 | 0 | 20 | 0 | 21 | 0 | 6 | 0 | 0 | 0 | 0 | 0 | 17 | 0 | 8 | 0 | 0 | 0 |
| PBANKA_API00300 | 0 | 0 | 0 | 0 | 8 | 0 | 7 | 8 | 16 | 0 | 0 | 5 | 0 | 0 | 0 | 0 | 9 | 0 | 0 | 0 | 0 | 0 |
| PBANKA_API00304 | 0 | 0 | 0 | 0 | 5 | 0 | 16 | 5 | 12 | 0 | 8 | 0 | 0 | 0 | 0 | 0 | 0 | 0 | 0 | 0 | 0 | 0 |
| PBANKA_API00320 | 0 | 0 | 0 | 0 | 5 | 0 | 13 | 0 | 5 | 0 | 0 | 0 | 0 | 0 | 0 | 0 | 0 | 0 | 0 | 0 | 0 | 0 |
| PBANKA_API00330 | 0 | 0 | 0 | 0 | 14 | 0 | 23 | 0 | 17 | 0 | 0 | 0 | 0 | 0 | 0 | 0 | 9 | 0 | 0 | 0 | 0 | 0 |
| PBANKA_API00340 | 0 | 0 | 0 | 0 | 5 | 0 | 11 | 0 | 7 | 0 | 0 | 0 | 0 | 0 | 0 | 0 | 0 | 0 | 0 | 0 | 0 | 0 |
| PBANKA_API00380 | 0 | 0 | 0 | 0 | 13 | 0 | 20 | 5 | 11 | 0 | 8 | 0 | 5 | 0 | 0 | 7 | 9 | 0 | 6 | 7 | 0 | 0 |
| PBANKA_API00390 | 0 | 0 | 0 | 0 | 6 | 0 | 8 | 0 | 11 | 0 | 0 | 0 | 0 | 0 | 0 | 0 | 5 | 0 | 0 | 0 | 0 | 0 |
| PBANKA_API00410 | 0 | 0 | 0 | 0 | 13 | 8 | 25 | 6 | 23 | 5 | 0 | 0 | 14 | 0 | 0 | 8 | 10 | 0 | 9 | 0 | 0 | 0 |
| PBANKA_MIT02100 | 0 | 0 | 0 | 0 | 9 | 0 | 0 | 0 | 0 | 0 | 0 | 0 | 0 | 5 | 124 | 26 | 12 | 30 | 54 | 7 | 0 | 6 |
| PBANKA_0100021 | 0 | 0 | 0 | 0 | 0 | 9 | 0 | 0 | 45 | 7 | 890 | 662 | 758 | 713 | 106 | 374 | 209 | 76 | 200 | 103 | 0 | 69 |
| PBANKA_0112681 | 0 | 0 | 0 | 0 | 0 | 14 | 0 | 0 | 7 | 0 | 0 | 0 | 38 | 6 | 9 | 0 | 0 | 6 | 49 | 22 | 0 | 0 |
| PBANKA_0201200 | 0 | 0 | 0 | 0 | 0 | 9 | 14 | 0 | 6 | 0 | 5 | 6 | 14 | 8 | 19 | 16 | 18 | 0 | 34 | 27 | 0 | 9 |
| PBANKA_0314800 | 0 | 0 | 0 | 0 | 0 | 6 | 35 | 13 | 28 | 16 | 0 | 0 | 317 | 109 | 28 | 66 | 383 | 131 | 6043 | 3096 | 29 | 193 |
| PBANKA_0404300 | 0 | 0 | 0 | 0 | 0 | 7 | 13 | 8 | 21 | 15 | 12 | 0 | 361 | 227 | 82 | 344 | 695 | 157 | 1588 | 2488 | 168 | 1154 |

| | | | | | | | | | | | | | | | | | | | | | | |
|---|---|---|---|---|---|---|---|---|---|---|---|---|---|---|---|---|---|---|---|---|---|---|
| PBANKA_0800100 | 0 | 0 | 0 | 0 | 0 | 6 | 0 | 0 | 5 | 0 | 0 | 12 | 292 | 154 | 70 | 178 | 40 | 14 | 829 | 263 | 0 | 57 |
| PBANKA_0942400 | 0 | 0 | 0 | 0 | 0 | 6 | 60 | 18 | 44 | 19 | 11 | 0 | 270 | 56 | 25 | 93 | 280 | 72 | 2433 | 1135 | 31 | 197 |
| PBANKA_1203600 | 0 | 0 | 0 | 0 | 0 | 5 | 0 | 0 | 9 | 0 | 0 | 0 | 22 | 0 | 0 | 5 | 0 | 0 | 293 | 52 | 32 | 209 |
| PBANKA_1325400 | 0 | 10 | 0 | 0 | 0 | 6 | 5 | 0 | 0 | 0 | 0 | 0 | 164 | 56 | 7 | 22 | 69 | 55 | 3013 | 1062 | 7 | 67 |
| PBANKA_1342300 | 0 | 0 | 0 | 0 | 0 | 5 | 10 | 6 | 0 | 0 | 0 | 0 | 598 | 249 | 12 | 51 | 266 | 148 | 14814 | 7171 | 312 | 1515 |
| PBANKA_1414800 | 14 | 0 | 0 | 0 | 0 | 7 | 9 | 11 | 12 | 0 | 0 | 0 | 259 | 54 | 5 | 8 | 43 | 35 | 3392 | 1279 | 157 | 445 |
| PBANKA_1444800 | 0 | 0 | 0 | 0 | 0 | 6 | 0 | 0 | 12 | 9 | 24 | 18 | 1716 | 1080 | 68 | 119 | 602 | 333 | 53196 | 53959 | 285 | 2862 |
| PBANKA_0001201 | 0 | 0 | 0 | 0 | 0 | 0 | 10 | 8 | 7 | 0 | 6 | 8 | 88 | 38 | 36 | 61 | 9 | 0 | 91 | 27 | 0 | 0 |
| PBANKA_0103200 | 0 | 0 | 0 | 0 | 0 | 0 | 9 | 0 | 7 | 6 | 0 | 0 | 432 | 164 | 8 | 45 | 104 | 92 | 12526 | 5449 | 220 | 1279 |
| PBANKA_0104200 | 0 | 0 | 0 | 0 | 0 | 0 | 6 | 0 | 6 | 0 | 0 | 0 | 99 | 31 | 29 | 173 | 160 | 52 | 789 | 235 | 248 | 1073 |
| PBANKA_0112621 | 0 | 0 | 0 | 0 | 0 | 0 | 10 | 0 | 9 | 0 | 0 | 0 | 77 | 92 | 20 | 97 | 22 | 8 | 49 | 52 | 0 | 0 |
| PBANKA_0201700 | 0 | 0 | 0 | 0 | 0 | 0 | 6 | 0 | 0 | 0 | 7 | 0 | 25 | 8 | 7 | 49 | 8 | 0 | 454 | 169 | 1054 | 7199 |
| PBANKA_0300300 | 0 | 0 | 0 | 0 | 0 | 0 | 6 | 0 | 7 | 0 | 0 | 0 | 421 | 160 | 66 | 289 | 67 | 12 | 370 | 125 | 0 | 0 |
| PBANKA_0300500 | 0 | 0 | 0 | 0 | 0 | 0 | 8 | 0 | 0 | 0 | 0 | 0 | 205 | 178 | 40 | 263 | 20 | 6 | 61 | 30 | 0 | 0 |
| PBANKA_0301400 | 0 | 0 | 0 | 0 | 0 | 0 | 19 | 10 | 35 | 22 | 0 | 0 | 183 | 55 | 461 | 2094 | 429 | 74 | 1505 | 784 | 79 | 444 |
| PBANKA_0301800 | 0 | 17 | 0 | 0 | 0 | 0 | 51 | 32 | 114 | 68 | 15 | 0 | 187 | 69 | 19 | 88 | 322 | 85 | 2207 | 992 | 140 | 878 |
| PBANKA_0302900 | 0 | 0 | 0 | 0 | 0 | 0 | 6 | 0 | 5 | 0 | 0 | 0 | 175 | 18 | 0 | 0 | 38 | 25 | 1468 | 380 | 7 | 41 |
| PBANKA_0303000 | 0 | 0 | 0 | 0 | 0 | 0 | 15 | 7 | 42 | 11 | 0 | 0 | 124 | 29 | 13 | 23 | 113 | 33 | 1200 | 510 | 57 | 494 |
| PBANKA_0311500 | 0 | 0 | 0 | 0 | 0 | 0 | 106 | 57 | 260 | 186 | 84 | 53 | 237 | 64 | 0 | 16 | 507 | 231 | 1196 | 512 | 2075 | 14237 |
| PBANKA_0317021 | 0 | 0 | 0 | 0 | 0 | 0 | 6 | 0 | 6 | 6 | 8 | 7 | 95 | 34 | 29 | 53 | 8 | 0 | 21 | 19 | 0 | 7 |
| PBANKA_0317161 | 0 | 0 | 0 | 0 | 0 | 0 | 7 | 0 | 12 | 0 | 70 | 53 | 103 | 74 | 20 | 34 | 20 | 19 | 261 | 68 | 0 | 37 |
| PBANKA_0400100 | 0 | 0 | 0 | 0 | 0 | 0 | 5 | 8 | 12 | 8 | 45 | 65 | 252 | 224 | 53 | 208 | 48 | 16 | 7 | 14 | 0 | 11 |
| PBANKA_0400200 | 0 | 0 | 0 | 0 | 0 | 0 | 12 | 5 | 17 | 0 | 0 | 0 | 0 | 0 | 5 | 38 | 0 | 0 | 14 | 6 | 0 | 0 |
| PBANKA_0500100 | 0 | 0 | 0 | 0 | 0 | 0 | 21 | 5 | 16 | 7 | 0 | 0 | 17 | 0 | 0 | 0 | 83 | 24 | 0 | 0 | 0 | 0 |
| PBANKA_0501000 | 0 | 0 | 0 | 0 | 0 | 0 | 7 | 0 | 115 | 110 | 14 | 9 | 72 | 81 | 125 | 221 | 268 | 74 | 21 | 74 | 0 | 25 |
| PBANKA_0522300 | 0 | 0 | 0 | 0 | 0 | 0 | 5 | 0 | 9 | 5 | 0 | 0 | 107 | 79 | 0 | 6 | 154 | 90 | 4483 | 1282 | 98 | 677 |
| PBANKA_0523800 | 0 | 0 | 0 | 0 | 0 | 0 | 62 | 29 | 430 | 298 | 50 | 34 | 282 | 166 | 0 | 9 | 4067 | 1018 | 336 | 159 | 35 | 257 |
| PBANKA_0524500 | 0 | 0 | 0 | 0 | 0 | 0 | 5 | 0 | 121 | 130 | 66 | 47 | 136 | 89 | 5 | 35 | 302 | 149 | 29 | 38 | 6 | 50 |
| PBANKA_0524841 | 0 | 0 | 0 | 0 | 0 | 0 | 5 | 0 | 35 | 12 | 0 | 0 | 15 | 26 | 69 | 59 | 138 | 75 | 12 | 30 | 0 | 7 |
| PBANKA_0600021 | 0 | 0 | 0 | 0 | 0 | 0 | 10 | 0 | 40 | 19 | 421 | 436 | 271 | 111 | 85 | 109 | 25 | 39 | 77 | 37 | 0 | 5 |
| PBANKA_0600031 | 0 | 0 | 0 | 0 | 0 | 0 | 8 | 0 | 53 | 30 | 1344 | 1034 | 39 | 57 | 8 | 30 | 11 | 0 | 99 | 153 | 0 | 14 |
| PBANKA_0600071 | 8 | 7 | 0 | 0 | 0 | 0 | 19 | 0 | 6 | 0 | 0 | 0 | 10 | 35 | 23 | 131 | 150 | 12 | 24 | 137 | 15 | 170 |
| PBANKA_0600251 | 0 | 0 | 0 | 0 | 0 | 0 | 6 | 6 | 8 | 6 | 5 | 0 | 39 | 27 | 17 | 45 | 0 | 5 | 355 | 117 | 0 | 0 |
| PBANKA_0600300 | 0 | 0 | 0 | 0 | 0 | 0 | 8 | 0 | 11 | 0 | 9 | 0 | 155 | 61 | 79 | 255 | 73 | 15 | 254 | 119 | 0 | 9 |
| PBANKA_0605800 | 24 | 18 | 0 | 0 | 0 | 0 | 8 | 0 | 0 | 0 | 0 | 0 | 267 | 141 | 10 | 34 | 56 | 38 | 6028 | 2862 | 200 | 1395 |
| PBANKA_0619700 | 5 | 7 | 0 | 0 | 0 | 0 | 182 | 52 | 1020 | 651 | 120 | 81 | 811 | 430 | 42 | 207 | 9028 | 2440 | 342 | 286 | 51 | 370 |
| PBANKA_0622900 | 0 | 0 | 0 | 0 | 0 | 0 | 19 | 19 | 269 | 317 | 51 | 40 | 59 | 45 | 0 | 0 | 241 | 182 | 105 | 59 | 9 | 57 |
| PBANKA_0623700 | 0 | 0 | 0 | 0 | 0 | 0 | 9 | 8 | 28 | 0 | 0 | 0 | 39 | 0 | 18 | 33 | 137 | 32 | 10 | 26 | 0 | 76 |
| PBANKA_0700051 | 0 | 0 | 0 | 0 | 0 | 0 | 25 | 0 | 15 | 14 | 24 | 24 | 218 | 94 | 80 | 204 | 41 | 0 | 228 | 118 | 0 | 7 |
| PBANKA_0705500 | 0 | 0 | 0 | 0 | 0 | 0 | 14 | 0 | 0 | 0 | 0 | 0 | 79 | 22 | 0 | 12 | 58 | 17 | 968 | 314 | 0 | 45 |
| PBANKA_0800021 | 0 | 0 | 0 | 0 | 0 | 0 | 9 | 0 | 15 | 6 | 0 | 0 | 16 | 37 | 20 | 30 | 184 | 86 | 13 | 30 | 0 | 5 |
| PBANKA_0800041 | 0 | 0 | 0 | 0 | 0 | 0 | 53 | 29 | 21 | 17 | 0 | 0 | 49 | 7 | 0 | 0 | 191 | 93 | 8 | 5 | 0 | 0 |
| PBANKA_0831600 | 0 | 0 | 0 | 0 | 0 | 0 | 6 | 0 | 0 | 6 | 0 | 0 | 140 | 96 | 11 | 28 | 106 | 63 | 2684 | 2194 | 15 | 98 |
| PBANKA_0836550 | 0 | 0 | 0 | 0 | 0 | 0 | 11 | 0 | 0 | 0 | 0 | 0 | 27 | 10 | 26 | 87 | 54 | 5 | 13 | 11 | 0 | 9 |
| PBANKA_0902800 | 0 | 0 | 0 | 0 | 0 | 0 | 11 | 0 | 11 | 11 | 0 | 0 | 59 | 24 | 43 | 166 | 114 | 33 | 87 | 38 | 9 | 37 |
| PBANKA_0905400 | 0 | 0 | 0 | 0 | 0 | 0 | 5 | 0 | 11 | 10 | 0 | 0 | 103 | 34 | 0 | 14 | 37 | 14 | 1315 | 598 | 21 | 106 |
| PBANKA_0907500 | 0 | 0 | 0 | 0 | 0 | 0 | 6 | 0 | 5 | 0 | 0 | 0 | 185 | 59 | 6 | 34 | 69 | 32 | 1260 | 612 | 43 | 267 |
| PBANKA_0915400 | 8 | 0 | 0 | 0 | 0 | 0 | 7 | 0 | 22 | 11 | 0 | 0 | 50 | 8 | 0 | 0 | 108 | 80 | 1049 | 570 | 60 | 425 |
| PBANKA_0916300 | 6 | 0 | 0 | 0 | 0 | 0 | 8 | 0 | 6 | 9 | 7 | 0 | 16 | 5 | 17 | 78 | 35 | 25 | 16 | 39 | 170 | 1279 |
| PBANKA_1000100 | 0 | 0 | 0 | 0 | 0 | 0 | 7 | 0 | 41 | 22 | 0 | 0 | 24 | 58 | 61 | 85 | 336 | 82 | 18 | 77 | 0 | 15 |
| PBANKA_1019360 | 0 | 0 | 0 | 0 | 0 | 0 | 5 | 0 | 5 | 0 | 6 | 0 | 125 | 11 | 60 | 27 | 19 | 11 | 598 | 12 | 14 | 76 |
| PBANKA_1019520 | 0 | 0 | 0 | 0 | 0 | 0 | 5 | 0 | 0 | 0 | 25 | 31 | 27 | 32 | 14 | 23 | 23 | 0 | 41 | 20 | 0 | 12 |
| PBANKA_1034200 | 11 | 7 | 0 | 0 | 0 | 0 | 8 | 0 | 8 | 0 | 0 | 0 | 0 | 0 | 0 | 14 | 8 | 0 | 121 | 37 | 34 | 118 |
| PBANKA_1037900 | 0 | 0 | 0 | 0 | 0 | 0 | 9 | 0 | 6 | 0 | 0 | 0 | 152 | 36 | 0 | 14 | 110 | 35 | 2199 | 637 | 90 | 485 |
| PBANKA_1100500 | 0 | 0 | 0 | 0 | 0 | 0 | 6 | 0 | 0 | 6 | 0 | 0 | 0 | 0 | 0 | 0 | 0 | 0 | 14 | 0 | 0 | 6 |
| PBANKA_1123200 | 0 | 0 | 0 | 0 | 0 | 0 | 13 | 9 | 14 | 16 | 0 | 0 | 391 | 155 | 42 | 163 | 346 | 104 | 5285 | 2515 | 130 | 792 |
| PBANKA_1146400 | 0 | 0 | 0 | 0 | 0 | 0 | 6 | 0 | 6 | 5 | 0 | 0 | 0 | 0 | 0 | 6 | 0 | 0 | 31 | 0 | 0 | 0 |
| PBANKA_1208900 | 0 | 0 | 0 | 0 | 0 | 0 | 6 | 0 | 6 | 7 | 0 | 0 | 138 | 29 | 6 | 29 | 92 | 37 | 1188 | 379 | 207 | 1076 |
| PBANKA_1229400 | 0 | 0 | 0 | 0 | 0 | 0 | 34 | 10 | 217 | 75 | 26 | 8 | 390 | 148 | 9 | 24 | 1450 | 453 | 2463 | 1174 | 82 | 512 |
| PBANKA_1300100 | 0 | 0 | 0 | 0 | 0 | 0 | 5 | 0 | 0 | 0 | 0 | 6 | 71 | 32 | 10 | 48 | 12 | 0 | 421 | 158 | 0 | 0 |
| PBANKA_1334700 | 0 | 0 | 0 | 0 | 0 | 0 | 7 | 9 | 19 | 9 | 0 | 0 | 646 | 316 | 17 | 75 | 459 | 163 | 18628 | 15092 | 814 | 6551 |
| PBANKA_1335000 | 0 | 0 | 0 | 0 | 0 | 0 | 7 | 0 | 18 | 7 | 0 | 0 | 614 | 206 | 32 | 49 | 191 | 133 | 17100 | 8893 | 128 | 1242 |
| PBANKA_1344700 | 321 | 512 | 0 | 0 | 0 | 0 | 21 | 0 | 148 | 85 | 28 | 15 | 189 | 77 | 12 | 14 | 619 | 156 | 1244 | 835 | 420 | 3853 |
| PBANKA_1365650 | 0 | 0 | 0 | 0 | 0 | 0 | 5 | 0 | 0 | 0 | 7 | 0 | 41 | 28 | 54 | 179 | 31 | 5 | 14 | 10 | 0 | 0 |
| PBANKA_1400071 | 0 | 0 | 0 | 0 | 0 | 0 | 6 | 0 | 0 | 0 | 0 | 6 | 284 | 112 | 58 | 363 | 121 | 14 | 195 | 191 | 0 | 10 |
| PBANKA_1400500 | 0 | 0 | 0 | 0 | 0 | 0 | 13 | 0 | 13 | 6 | 0 | 0 | 11 | 7 | 15 | 0 | 7 | 0 | 0 | 0 | 0 | 0 |
| PBANKA_1419300 | 0 | 0 | 0 | 0 | 0 | 0 | 5 | 0 | 6 | 0 | 0 | 0 | 1626 | 914 | 47 | 131 | 725 | 400 | 54198 | 43244 | 1978 | 12887 |
| PBANKA_1429200 | 1238 | 1603 | 0 | 0 | 0 | 0 | 11 | 11 | 18 | 7 | 0 | 6 | 361 | 185 | 11 | 25 | 191 | 127 | 8294 | 5930 | 1938 | 14825 |
| PBANKA_1432200 | 7 | 10 | 0 | 0 | 0 | 0 | 10 | 0 | 0 | 0 | 0 | 0 | 1267 | 436 | 23 | 53 | 2309 | 909 | 19546 | 10215 | 1918 | 16751 |
| PBANKA_1436300 | 0 | 0 | 0 | 0 | 0 | 0 | 6 | 0 | 0 | 0 | 0 | 0 | 343 | 174 | 22 | 77 | 186 | 75 | 7171 | 3332 | 27 | 197 |
| PBANKA_1463900 | 0 | 0 | 0 | 0 | 0 | 0 | 10 | 8 | 6 | 0 | 0 | 0 | 80 | 29 | 0 | 0 | 78 | 36 | 1453 | 473 | 214 | 1540 |
| PBANKA_1464100 | 0 | 0 | 0 | 0 | 0 | 0 | 8 | 0 | 11 | 0 | 6 | 0 | 75 | 62 | 15 | 41 | 55 | 15 | 82 | 23 | 12 | 84 |
| PBANKA_API00360 | 0 | 0 | 0 | 0 | 0 | 0 | 8 | 0 | 10 | 0 | 0 | 0 | 0 | 0 | 0 | 0 | 0 | 0 | 0 | 0 | 0 | 0 |
| PBANKA_API00430 | 0 | 0 | 0 | 0 | 0 | 0 | 11 | 0 | 5 | 0 | 0 | 0 | 0 | 0 | 0 | 0 | 0 | 0 | 0 | 0 | 0 | 0 |
| PBANKA_0700011 | 0 | 0 | 0 | 0 | 0 | 0 | 0 | 10 | 26 | 0 | 0 | 0 | 6 | 0 | 19 | 96 | 0 | 0 | 45 | 22 | 0 | 0 |
| PBANKA_1311400 | 171 | 174 | 0 | 0 | 0 | 0 | 0 | 6 | 7 | 0 | 0 | 0 | 105 | 42 | 7 | 0 | 41 | 28 | 2306 | 548 | 104 | 424 |
| PBANKA_1336200 | 5 | 0 | 0 | 0 | 0 | 0 | 0 | 5 | 0 | 0 | 0 | 0 | 159 | 55 | 9 | 40 | 53 | 62 | 5082 | 1349 | 13 | 153 |
| PBANKA_1431500 | 0 | 0 | 0 | 0 | 0 | 0 | 0 | 5 | 5 | 0 | 0 | 0 | 924 | 231 | 13 | 92 | 589 | 111 | 7962 | 5109 | 19 | 115 |
| PBANKA_1465400 | 0 | 0 | 0 | 0 | 0 | 0 | 0 | 6 | 0 | 0 | 7 | 0 | 77 | 39 | 32 | 71 | 7 | 0 | 32 | 22 | 0 | 0 |
| PBANKA_0001101 | 0 | 0 | 0 | 0 | 0 | 0 | 0 | 0 | 7 | 0 | 19 | 42 | 50 | 20 | 14 | 18 | 0 | 5 | 80 | 17 | 0 | 0 |
| PBANKA_0001601 | 0 | 0 | 0 | 0 | 0 | 0 | 0 | 0 | 10 | 13 | 293 | 303 | 22 | 0 | 12 | 6 | 0 | 0 | 10 | 0 | 0 | 0 |
| PBANKA_0007501 | 0 | 0 | 0 | 0 | 0 | 0 | 0 | 0 | 5 | 12 | 257 | 189 | 13 | 8 | 0 | 5 | 0 | 0 | 0 | 0 | 0 | 0 |
| PBANKA_0007801 | 0 | 0 | 0 | 0 | 0 | 0 | 0 | 0 | 7 | 8 | 167 | 121 | 62 | 12 | 7 | 5 | 0 | 0 | 11 | 0 | 0 | 0 |
| PBANKA_0008101 | 0 | 0 | 0 | 0 | 0 | 0 | 0 | 0 | 40 | 22 | 447 | 561 | 64 | 56 | 24 | 43 | 7 | 7 | 15 | 11 | 0 | 7 |
| PBANKA_0109800 | 25 | 40 | 0 | 0 | 0 | 0 | 0 | 0 | 8 | 0 | 0 | 0 | 542 | 256 | 11 | 33 | 256 | 125 | 13726 | 10812 | 111 | 951 |
| PBANKA_0216041 | 0 | 0 | 0 | 0 | 0 | 0 | 0 | 0 | 9 | 11 | 123 | 86 | 501 | 180 | 65 | 80 | 66 | 0 | 219 | 133 | 0 | 26 |
| PBANKA_0316400 | 0 | 0 | 0 | 0 | 0 | 0 | 0 | 0 | 6 | 0 | 9 | 0 | 210 | 85 | 305 | 1265 | 525 | 102 | 1118 | 711 | 6 | 70 |
| PBANKA_0316500 | 0 | 0 | 0 | 0 | 0 | 0 | 0 | 0 | 27 | 33 | 1438 | 1269 | 428 | 660 | 32 | 52 | 34 | 23 | 123 | 42 | 0 | 16 |
| PBANKA_0316900 | 0 | 0 | 0 | 0 | 0 | 0 | 0 | 0 | 5 | 0 | 13 | 9 | 26 | 35 | 11 | 34 | 0 | 0 | 58 | 31 | 0 | 14 |
| PBANKA_0316921 | 0 | 0 | 0 | 0 | 0 | 0 | 0 | 0 | 103 | 28 | 436 | 350 | 1264 | 881 | 150 | 955 | 383 | 83 | 37 | 36 | 0 | 0 |

| | | | | | | | | | | | | | | | | | | | | | | |
|---|---|---|---|---|---|---|---|---|---|---|---|---|---|---|---|---|---|---|---|---|---|---|
| PBANKA_0317121 | 0 | 0 | 0 | 0 | 0 | 0 | 0 | 0 | 5 | 6 | 18 | 22 | 210 | 58 | 84 | 132 | 71 | 43 | 17 | 18 | 0 | 8 |
| PBANKA_0317181 | 0 | 0 | 0 | 0 | 0 | 0 | 0 | 0 | 11 | 0 | 0 | 0 | 47 | 18 | 23 | 36 | 16 | 19 | 631 | 353 | 0 | 17 |
| PBANKA_0500600 | 0 | 0 | 0 | 0 | 0 | 0 | 0 | 0 | 29 | 28 | 223 | 232 | 295 | 238 | 51 | 201 | 129 | 29 | 14 | 9 | 0 | 5 |
| PBANKA_0500700 | 0 | 0 | 0 | 0 | 0 | 0 | 0 | 0 | 137 | 95 | 732 | 643 | 1401 | 1047 | 378 | 1277 | 316 | 103 | 78 | 72 | 0 | 25 |
| PBANKA_0500741 | 0 | 0 | 0 | 0 | 0 | 0 | 0 | 0 | 13 | 27 | 17 | 48 | 1820 | 683 | 194 | 697 | 897 | 329 | 464 | 275 | 0 | 47 |
| PBANKA_0504300 | 0 | 0 | 0 | 0 | 0 | 0 | 0 | 0 | 8 | 5 | 0 | 0 | 164 | 32 | 0 | 15 | 95 | 30 | 2205 | 712 | 130 | 957 |
| PBANKA_0516000 | 0 | 0 | 0 | 0 | 0 | 0 | 0 | 0 | 11 | 0 | 0 | 0 | 195 | 78 | 13 | 13 | 68 | 42 | 3693 | 1937 | 58 | 336 |
| PBANKA_0522200 | 129 | 171 | 0 | 0 | 0 | 0 | 0 | 0 | 7 | 0 | 0 | 0 | 273 | 113 | 0 | 13 | 89 | 49 | 7426 | 5075 | 15 | 112 |
| PBANKA_0600011 | 0 | 0 | 0 | 0 | 0 | 0 | 0 | 0 | 11 | 0 | 0 | 10 | 20 | 0 | 0 | 10 | 0 | 0 | 7 | 7 | 0 | 0 |
| PBANKA_0600061 | 0 | 0 | 0 | 0 | 0 | 0 | 0 | 0 | 6 | 6 | 177 | 101 | 440 | 347 | 98 | 393 | 189 | 61 | 146 | 80 | 0 | 22 |
| PBANKA_0623600 | 0 | 0 | 0 | 0 | 0 | 0 | 0 | 0 | 6 | 0 | 0 | 6 | 109 | 36 | 27 | 63 | 23 | 0 | 1063 | 442 | 0 | 17 |
| PBANKA_0700021 | 0 | 0 | 0 | 0 | 0 | 0 | 0 | 0 | 63 | 0 | 0 | 0 | 0 | 0 | 7 | 34 | 0 | 0 | 34 | 0 | 0 | 0 |
| PBANKA_0700081 | 0 | 0 | 0 | 0 | 0 | 0 | 0 | 0 | 17 | 7 | 195 | 252 | 269 | 61 | 49 | 130 | 17 | 6 | 269 | 131 | 0 | 27 |
| PBANKA_0716800 | 6 | 5 | 0 | 0 | 0 | 0 | 0 | 0 | 8 | 0 | 0 | 0 | 7 | 0 | 0 | 8 | 10 | 7 | 18 | 6 | 1328 | 5848 |
| PBANKA_0836600 | 0 | 0 | 0 | 0 | 0 | 0 | 0 | 0 | 64 | 52 | 100 | 68 | 397 | 218 | 71 | 258 | 212 | 149 | 64 | 32 | 0 | 0 |
| PBANKA_0836961 | 0 | 0 | 0 | 0 | 0 | 0 | 0 | 0 | 5 | 0 | 0 | 0 | 37 | 7 | 15 | 33 | 8 | 0 | 45 | 13 | 0 | 13 |
| PBANKA_0836981 | 0 | 0 | 0 | 0 | 0 | 0 | 0 | 0 | 13 | 0 | 45 | 51 | 60 | 16 | 8 | 16 | 0 | 0 | 35 | 12 | 0 | 0 |
| PBANKA_0837021 | 0 | 0 | 0 | 0 | 0 | 0 | 0 | 0 | 5 | 0 | 0 | 0 | 24 | 12 | 0 | 10 | 8 | 0 | 10 | 0 | 0 | 0 |
| PBANKA_0902000 | 0 | 0 | 0 | 0 | 0 | 0 | 0 | 0 | 8 | 0 | 0 | 0 | 45 | 10 | 0 | 0 | 36 | 28 | 261 | 89 | 5 | 27 |
| PBANKA_0944121 | 0 | 0 | 0 | 0 | 0 | 0 | 0 | 0 | 6 | 0 | 0 | 0 | 0 | 0 | 0 | 0 | 0 | 0 | 0 | 0 | 0 | 0 |
| PBANKA_1000041 | 0 | 0 | 0 | 0 | 0 | 0 | 0 | 0 | 5 | 10 | 44 | 40 | 489 | 265 | 153 | 216 | 124 | 76 | 25 | 16 | 0 | 0 |
| PBANKA_1000051 | 0 | 0 | 0 | 0 | 0 | 0 | 0 | 0 | 38 | 31 | 136 | 134 | 299 | 223 | 88 | 334 | 80 | 24 | 14 | 5 | 0 | 7 |
| PBANKA_1000071 | 0 | 0 | 0 | 0 | 0 | 0 | 0 | 0 | 6 | 0 | 9 | 0 | 468 | 109 | 95 | 69 | 224 | 174 | 184 | 62 | 0 | 11 |
| PBANKA_1040151 | 0 | 0 | 0 | 0 | 0 | 0 | 0 | 0 | 10 | 0 | 0 | 0 | 0 | 0 | 0 | 0 | 0 | 0 | 48 | 6 | 0 | 11 |
| PBANKA_1040581 | 0 | 0 | 0 | 0 | 0 | 0 | 0 | 0 | 11 | 0 | 268 | 184 | 1512 | 1130 | 163 | 1215 | 79 | 9 | 158 | 160 | 0 | 0 |
| PBANKA_1100011 | 0 | 0 | 0 | 0 | 0 | 0 | 0 | 0 | 7 | 0 | 0 | 0 | 0 | 0 | 14 | 13 | 0 | 0 | 6 | 0 | 0 | 0 |
| PBANKA_1100461 | 0 | 0 | 0 | 0 | 0 | 0 | 0 | 0 | 13 | 0 | 12 | 0 | 13 | 5 | 14 | 15 | 0 | 0 | 23 | 21 | 0 | 7 |
| PBANKA_1146200 | 0 | 0 | 0 | 0 | 0 | 0 | 0 | 0 | 10 | 0 | 0 | 0 | 92 | 197 | 10 | 49 | 321 | 33 | 173 | 89 | 0 | 0 |
| PBANKA_1146761 | 0 | 0 | 0 | 0 | 0 | 0 | 0 | 0 | 16 | 5 | 0 | 0 | 28 | 14 | 15 | 14 | 122 | 64 | 26 | 9 | 0 | 15 |
| PBANKA_1200041 | 0 | 0 | 0 | 0 | 0 | 0 | 0 | 0 | 8 | 10 | 405 | 353 | 31 | 18 | 9 | 0 | 0 | 6 | 25 | 0 | 0 | 0 |
| PBANKA_1203500 | 0 | 0 | 0 | 0 | 0 | 0 | 0 | 0 | 8 | 0 | 0 | 0 | 95 | 25 | 6 | 35 | 58 | 14 | 1047 | 403 | 23 | 116 |
| PBANKA_1246100 | 0 | 0 | 0 | 0 | 0 | 0 | 0 | 0 | 22 | 11 | 123 | 121 | 92 | 63 | 26 | 27 | 59 | 91 | 25 | 11 | 0 | 0 |
| PBANKA_1246400 | 0 | 0 | 0 | 0 | 0 | 0 | 0 | 0 | 5 | 0 | 10 | 12 | 296 | 180 | 110 | 296 | 65 | 18 | 867 | 281 | 0 | 22 |
| PBANKA_1365550 | 0 | 0 | 0 | 0 | 0 | 0 | 0 | 0 | 5 | 0 | 6 | 0 | 235 | 157 | 85 | 501 | 93 | 18 | 114 | 96 | 0 | 0 |
| PBANKA_1400031 | 0 | 0 | 0 | 0 | 0 | 0 | 0 | 0 | 15 | 0 | 76 | 58 | 150 | 71 | 45 | 51 | 20 | 20 | 136 | 70 | 0 | 0 |
| PBANKA_1400051 | 0 | 0 | 0 | 0 | 0 | 0 | 0 | 0 | 6 | 0 | 161 | 88 | 579 | 322 | 132 | 278 | 17 | 15 | 57 | 47 | 0 | 0 |
| PBANKA_1400200 | 0 | 0 | 0 | 0 | 0 | 0 | 0 | 0 | 7 | 0 | 8 | 0 | 397 | 264 | 44 | 289 | 60 | 13 | 43 | 51 | 0 | 14 |
| PBANKA_1429100 | 18 | 26 | 0 | 0 | 0 | 0 | 0 | 0 | 13 | 15 | 0 | 0 | 185 | 44 | 6 | 9 | 59 | 32 | 2409 | 618 | 46 | 413 |
| PBANKA_1431400 | 0 | 0 | 0 | 0 | 0 | 0 | 0 | 0 | 7 | 7 | 8 | 0 | 221 | 166 | 22 | 80 | 290 | 60 | 1736 | 4489 | 33 | 242 |
| PBANKA_1465500 | 0 | 0 | 0 | 0 | 0 | 0 | 0 | 0 | 6 | 0 | 12 | 6 | 94 | 35 | 16 | 107 | 28 | 0 | 103 | 37 | 0 | 6 |
| PBANKA_1466121 | 0 | 0 | 0 | 0 | 0 | 0 | 0 | 0 | 6 | 0 | 34 | 29 | 165 | 81 | 50 | 134 | 20 | 7 | 96 | 46 | 0 | 12 |
| PBANKA_API00550 | 0 | 0 | 0 | 0 | 0 | 0 | 0 | 0 | 5 | 0 | 0 | 0 | 0 | 0 | 0 | 0 | 0 | 0 | 0 | 0 | 0 | 0 |
| PBANKA_MIT03300 | 0 | 0 | 0 | 0 | 0 | 0 | 0 | 0 | 5 | 0 | 0 | 0 | 0 | 0 | 48 | 0 | 6 | 0 | 0 | 53 | 0 | 0 |
| PBANKA_0837001 | 0 | 0 | 0 | 0 | 0 | 0 | 0 | 0 | 0 | 8 | 128 | 137 | 122 | 31 | 26 | 45 | 11 | 0 | 62 | 48 | 0 | 28 |
| PBANKA_0837121 | 0 | 0 | 0 | 0 | 0 | 0 | 0 | 0 | 0 | 5 | 7 | 9 | 78 | 17 | 0 | 58 | 25 | 0 | 88 | 24 | 0 | 13 |
| PBANKA_0837141 | 0 | 0 | 0 | 0 | 0 | 0 | 0 | 0 | 0 | 5 | 167 | 94 | 221 | 95 | 25 | 209 | 49 | 5 | 100 | 55 | 0 | 13 |
| PBANKA_1146781 | 0 | 0 | 0 | 0 | 0 | 0 | 0 | 0 | 0 | 5 | 0 | 0 | 261 | 106 | 84 | 63 | 138 | 130 | 129 | 44 | 0 | 0 |
| PBANKA_1214500 | 0 | 0 | 0 | 0 | 0 | 0 | 0 | 0 | 0 | 5 | 5 | 0 | 48 | 10 | 17 | 32 | 28 | 16 | 568 | 189 | 0 | 14 |
| PBANKA_1427700 | 0 | 0 | 0 | 0 | 0 | 0 | 0 | 0 | 0 | 5 | 0 | 0 | 99 | 30 | 46 | 124 | 168 | 55 | 778 | 504 | 7 | 34 |
| PBANKA_0000101 | 0 | 0 | 0 | 0 | 0 | 0 | 0 | 0 | 0 | 0 | 18 | 0 | 87 | 50 | 16 | 44 | 5 | 0 | 10 | 15 | 0 | 0 |
| PBANKA_0000301 | 0 | 0 | 0 | 0 | 0 | 0 | 0 | 0 | 0 | 0 | 12 | 8 | 13 | 5 | 0 | 0 | 0 | 0 | 63 | 26 | 0 | 0 |
| PBANKA_0000600 | 0 | 0 | 0 | 0 | 0 | 0 | 0 | 0 | 0 | 0 | 5 | 0 | 31 | 11 | 6 | 7 | 0 | 0 | 15 | 5 | 0 | 0 |
| PBANKA_0000701 | 0 | 0 | 0 | 0 | 0 | 0 | 0 | 0 | 0 | 0 | 6 | 5 | 12 | 0 | 0 | 0 | 0 | 0 | 145 | 35 | 0 | 0 |
| PBANKA_0001001 | 0 | 0 | 0 | 0 | 0 | 0 | 0 | 0 | 0 | 0 | 34 | 44 | 146 | 91 | 33 | 171 | 58 | 5 | 857 | 500 | 0 | 21 |
| PBANKA_0001301 | 0 | 0 | 0 | 0 | 0 | 0 | 0 | 0 | 0 | 0 | 27 | 28 | 0 | 0 | 0 | 0 | 0 | 0 | 0 | 0 | 0 | 0 |
| PBANKA_0001401 | 0 | 0 | 0 | 0 | 0 | 0 | 0 | 0 | 0 | 0 | 9 | 0 | 12 | 0 | 0 | 7 | 0 | 0 | 0 | 0 | 0 | 0 |
| PBANKA_0007601 | 0 | 0 | 0 | 0 | 0 | 0 | 0 | 0 | 0 | 0 | 6 | 0 | 0 | 0 | 0 | 0 | 0 | 0 | 5 | 6 | 0 | 0 |
| PBANKA_0008001 | 0 | 0 | 0 | 0 | 0 | 0 | 0 | 0 | 0 | 0 | 7 | 0 | 0 | 0 | 0 | 5 | 0 | 0 | 11 | 5 | 0 | 0 |
| PBANKA_0008201 | 0 | 0 | 0 | 0 | 0 | 0 | 0 | 0 | 0 | 0 | 10 | 0 | 17 | 0 | 0 | 9 | 0 | 0 | 5 | 0 | 0 | 0 |
| PBANKA_0216021 | 0 | 0 | 0 | 0 | 0 | 0 | 0 | 0 | 0 | 0 | 55 | 52 | 45 | 0 | 0 | 0 | 0 | 0 | 204 | 188 | 0 | 0 |
| PBANKA_0300200 | 0 | 0 | 0 | 0 | 0 | 0 | 0 | 0 | 0 | 0 | 5 | 10 | 51 | 42 | 14 | 63 | 8 | 0 | 16 | 10 | 0 | 0 |
| PBANKA_0500950 | 0 | 0 | 0 | 0 | 0 | 0 | 0 | 0 | 0 | 0 | 12 | 17 | 1265 | 539 | 332 | 1505 | 384 | 49 | 691 | 246 | 0 | 13 |
| PBANKA_0500971 | 0 | 0 | 0 | 0 | 0 | 0 | 0 | 0 | 0 | 0 | 37 | 27 | 231 | 125 | 79 | 292 | 56 | 9 | 232 | 42 | 0 | 7 |
| PBANKA_0600081 | 0 | 0 | 0 | 0 | 0 | 0 | 0 | 0 | 0 | 0 | 16 | 15 | 84 | 28 | 14 | 34 | 11 | 0 | 172 | 95 | 5 | 10 |
| PBANKA_0600086 | 0 | 0 | 0 | 0 | 0 | 0 | 0 | 0 | 0 | 0 | 21 | 7 | 72 | 28 | 13 | 90 | 11 | 0 | 359 | 179 | 0 | 26 |
| PBANKA_0600100 | 0 | 0 | 0 | 0 | 0 | 0 | 0 | 0 | 0 | 0 | 6 | 37 | 93 | 56 | 20 | 85 | 16 | 0 | 156 | 75 | 0 | 13 |
| PBANKA_0600200 | 0 | 0 | 0 | 0 | 0 | 0 | 0 | 0 | 0 | 0 | 37 | 13 | 55 | 20 | 18 | 43 | 10 | 0 | 231 | 66 | 0 | 0 |
| PBANKA_0623651 | 0 | 0 | 0 | 0 | 0 | 0 | 0 | 0 | 0 | 0 | 34 | 42 | 83 | 35 | 70 | 98 | 28 | 11 | 144 | 71 | 0 | 0 |
| PBANKA_0623751 | 0 | 0 | 0 | 0 | 0 | 0 | 0 | 0 | 0 | 0 | 10 | 7 | 0 | 0 | 0 | 0 | 0 | 0 | 0 | 0 | 0 | 0 |
| PBANKA_0700031 | 0 | 0 | 0 | 0 | 0 | 0 | 0 | 0 | 0 | 0 | 14 | 12 | 14 | 0 | 0 | 15 | 0 | 0 | 16 | 0 | 0 | 0 |
| PBANKA_0700041 | 0 | 0 | 0 | 0 | 0 | 0 | 0 | 0 | 0 | 0 | 42 | 60 | 6 | 0 | 0 | 0 | 0 | 0 | 10 | 0 | 0 | 0 |
| PBANKA_0700071 | 0 | 0 | 0 | 0 | 0 | 0 | 0 | 0 | 0 | 0 | 43 | 68 | 115 | 56 | 23 | 77 | 16 | 0 | 421 | 130 | 0 | 21 |
| PBANKA_0700200 | 0 | 0 | 0 | 0 | 0 | 0 | 0 | 0 | 0 | 0 | 9 | 16 | 115 | 31 | 25 | 118 | 32 | 11 | 442 | 248 | 0 | 18 |
| PBANKA_0700300 | 0 | 0 | 0 | 0 | 0 | 0 | 0 | 0 | 0 | 0 | 10 | 0 | 40 | 5 | 12 | 27 | 13 | 0 | 266 | 116 | 0 | 10 |
| PBANKA_0700400 | 0 | 0 | 0 | 0 | 0 | 0 | 0 | 0 | 0 | 0 | 13 | 6 | 87 | 23 | 21 | 37 | 0 | 12 | 18 | 15 | 0 | 0 |
| PBANKA_0700500 | 0 | 0 | 0 | 0 | 0 | 0 | 0 | 0 | 0 | 0 | 19 | 21 | 61 | 58 | 0 | 28 | 0 | 0 | 44 | 17 | 0 | 10 |
| PBANKA_0722941 | 0 | 0 | 0 | 0 | 0 | 0 | 0 | 0 | 0 | 0 | 8 | 0 | 1483 | 409 | 344 | 1256 | 513 | 107 | 192 | 104 | 0 | 15 |
| PBANKA_0800200 | 0 | 0 | 0 | 0 | 0 | 0 | 0 | 0 | 0 | 0 | 5 | 0 | 81 | 30 | 23 | 94 | 11 | 9 | 400 | 146 | 0 | 27 |
| PBANKA_0836941 | 0 | 0 | 0 | 0 | 0 | 0 | 0 | 0 | 0 | 0 | 16 | 8 | 14 | 0 | 16 | 0 | 0 | 6 | 19 | 10 | 0 | 12 |
| PBANKA_0837101 | 0 | 0 | 0 | 0 | 0 | 0 | 0 | 0 | 0 | 0 | 21 | 15 | 66 | 17 | 26 | 21 | 6 | 0 | 55 | 21 | 0 | 22 |
| PBANKA_0917980 | 5 | 8 | 0 | 0 | 0 | 0 | 0 | 0 | 0 | 0 | 7 | 0 | 212 | 31 | 95 | 39 | 58 | 14 | 2961 | 74 | 104 | 737 |
| PBANKA_0937860 | 0 | 0 | 0 | 0 | 0 | 0 | 0 | 0 | 0 | 0 | 6 | 0 | 115 | 21 | 46 | 21 | 17 | 10 | 542 | 12 | 0 | 0 |
| PBANKA_0943541 | 0 | 0 | 0 | 0 | 0 | 0 | 0 | 0 | 0 | 0 | 5 | 0 | 0 | 0 | 0 | 0 | 0 | 0 | 0 | 0 | 0 | 0 |
| PBANKA_0943800 | 0 | 0 | 0 | 0 | 0 | 0 | 0 | 0 | 0 | 0 | 8 | 0 | 42 | 13 | 13 | 16 | 0 | 6 | 10 | 0 | 0 | 0 |
| PBANKA_0944000 | 0 | 0 | 0 | 0 | 0 | 0 | 0 | 0 | 0 | 0 | 8 | 0 | 186 | 42 | 36 | 88 | 20 | 12 | 70 | 30 | 0 | 0 |
| PBANKA_0944061 | 0 | 0 | 0 | 0 | 0 | 0 | 0 | 0 | 0 | 0 | 26 | 33 | 70 | 40 | 19 | 69 | 15 | 0 | 29 | 16 | 0 | 5 |
| PBANKA_0944081 | 0 | 0 | 0 | 0 | 0 | 0 | 0 | 0 | 0 | 0 | 6 | 6 | 25 | 8 | 13 | 39 | 13 | 0 | 77 | 47 | 0 | 0 |
| PBANKA_0944101 | 0 | 0 | 0 | 0 | 0 | 0 | 0 | 0 | 0 | 0 | 63 | 122 | 149 | 75 | 39 | 90 | 27 | 10 | 36 | 25 | 0 | 0 |
| PBANKA_1000091 | 0 | 0 | 0 | 0 | 0 | 0 | 0 | 0 | 0 | 0 | 10 | 9 | 115 | 47 | 79 | 127 | 31 | 0 | 86 | 45 | 0 | 6 |
| PBANKA_1100351 | 0 | 0 | 0 | 0 | 0 | 0 | 0 | 0 | 0 | 0 | 23 | 26 | 9 | 7 | 5 | 8 | 0 | 0 | 125 | 32 | 0 | 10 |

| | | | | | | | | | | | | | | | | | | | | | | |
|---|---|---|---|---|---|---|---|---|---|---|---|---|---|---|---|---|---|---|---|---|---|---|
| PBANKA_1100400 | 0 | 0 | 0 | 0 | 0 | 0 | 0 | 0 | 0 | 0 | 8 | 0 | 17 | 0 | 6 | 0 | 0 | 0 | 50 | 12 | 0 | 0 |
| PBANKA_1100781 | 0 | 0 | 0 | 0 | 0 | 0 | 0 | 0 | 0 | 0 | 5 | 0 | 30 | 9 | 6 | 24 | 7 | 0 | 38 | 21 | 0 | 6 |
| PBANKA_1100850 | 0 | 0 | 0 | 0 | 0 | 0 | 0 | 0 | 0 | 0 | 7 | 0 | 36 | 16 | 6 | 33 | 61 | 10 | 10 | 8 | 0 | 0 |
| PBANKA_1125650 | 0 | 0 | 0 | 0 | 0 | 0 | 0 | 0 | 0 | 0 | 7 | 0 | 18 | 11 | 27 | 15 | 7 | 0 | 8 | 6 | 0 | 0 |
| PBANKA_1146700 | 0 | 0 | 0 | 0 | 0 | 0 | 0 | 0 | 0 | 0 | 36 | 24 | 97 | 15 | 27 | 77 | 20 | 0 | 227 | 84 | 0 | 44 |
| PBANKA_1146721 | 0 | 0 | 0 | 0 | 0 | 0 | 0 | 0 | 0 | 0 | 65 | 57 | 228 | 114 | 24 | 242 | 64 | 5 | 159 | 66 | 11 | 48 |
| PBANKA_1146741 | 0 | 0 | 0 | 0 | 0 | 0 | 0 | 0 | 0 | 0 | 10 | 9 | 179 | 57 | 110 | 204 | 28 | 8 | 90 | 34 | 0 | 0 |
| PBANKA_1200011 | 0 | 0 | 0 | 0 | 0 | 0 | 0 | 0 | 0 | 0 | 9 | 6 | 0 | 0 | 0 | 0 | 0 | 0 | 0 | 0 | 0 | 0 |
| PBANKA_1200021 | 0 | 0 | 0 | 0 | 0 | 0 | 0 | 0 | 0 | 0 | 7 | 6 | 0 | 0 | 0 | 0 | 0 | 0 | 27 | 14 | 0 | 0 |
| PBANKA_1200031 | 0 | 0 | 0 | 0 | 0 | 0 | 0 | 0 | 0 | 0 | 47 | 49 | 17 | 0 | 0 | 0 | 0 | 0 | 11 | 0 | 0 | 0 |
| PBANKA_1200071 | 0 | 0 | 0 | 0 | 0 | 0 | 0 | 0 | 0 | 0 | 14 | 6 | 75 | 34 | 38 | 88 | 20 | 5 | 64 | 11 | 0 | 0 |
| PBANKA_1200400 | 0 | 0 | 0 | 0 | 0 | 0 | 0 | 0 | 0 | 0 | 8 | 0 | 390 | 179 | 103 | 271 | 69 | 13 | 1174 | 404 | 9 | 69 |
| PBANKA_1245920 | 0 | 0 | 0 | 0 | 0 | 0 | 0 | 0 | 0 | 0 | 21 | 12 | 1331 | 594 | 74 | 387 | 287 | 54 | 438 | 184 | 7 | 59 |
| PBANKA_1246800 | 0 | 0 | 0 | 0 | 0 | 0 | 0 | 0 | 0 | 0 | 28 | 28 | 19 | 11 | 6 | 10 | 0 | 0 | 367 | 95 | 0 | 8 |
| PBANKA_1246900 | 0 | 0 | 0 | 0 | 0 | 0 | 0 | 0 | 0 | 0 | 8 | 0 | 41 | 25 | 29 | 75 | 20 | 0 | 67 | 16 | 0 | 0 |
| PBANKA_1246941 | 0 | 0 | 0 | 0 | 0 | 0 | 0 | 0 | 0 | 0 | 7 | 0 | 0 | 0 | 0 | 0 | 0 | 0 | 0 | 0 | 0 | 0 |
| PBANKA_1246961 | 0 | 0 | 0 | 0 | 0 | 0 | 0 | 0 | 0 | 0 | 73 | 53 | 7 | 8 | 7 | 0 | 0 | 0 | 0 | 0 | 0 | 0 |
| PBANKA_1300011 | 0 | 0 | 0 | 0 | 0 | 0 | 0 | 0 | 0 | 0 | 77 | 54 | 22 | 7 | 12 | 5 | 0 | 0 | 14 | 8 | 0 | 0 |
| PBANKA_1300021 | 0 | 0 | 0 | 0 | 0 | 0 | 0 | 0 | 0 | 0 | 6 | 0 | 7 | 6 | 0 | 0 | 0 | 0 | 21 | 7 | 0 | 0 |
| PBANKA_1300041 | 0 | 0 | 0 | 0 | 0 | 0 | 0 | 0 | 0 | 0 | 42 | 33 | 6 | 0 | 0 | 0 | 0 | 0 | 0 | 0 | 0 | 0 |
| PBANKA_1300051 | 0 | 0 | 0 | 0 | 0 | 0 | 0 | 0 | 0 | 0 | 10 | 5 | 0 | 0 | 0 | 0 | 0 | 0 | 0 | 0 | 0 | 0 |
| PBANKA_1300081 | 0 | 0 | 0 | 0 | 0 | 0 | 0 | 0 | 0 | 0 | 13 | 8 | 32 | 6 | 7 | 56 | 6 | 0 | 63 | 38 | 0 | 0 |
| PBANKA_1300091 | 0 | 0 | 0 | 0 | 0 | 0 | 0 | 0 | 0 | 0 | 9 | 7 | 87 | 37 | 32 | 85 | 24 | 0 | 103 | 32 | 0 | 0 |
| PBANKA_1300200 | 0 | 0 | 0 | 0 | 0 | 0 | 0 | 0 | 0 | 0 | 5 | 0 | 2146 | 1333 | 218 | 2495 | 375 | 46 | 717 | 580 | 0 | 27 |
| PBANKA_1400041 | 0 | 0 | 0 | 0 | 0 | 0 | 0 | 0 | 0 | 0 | 14 | 8 | 65 | 20 | 10 | 18 | 0 | 0 | 698 | 646 | 0 | 13 |
| PBANKA_1465300 | 0 | 0 | 0 | 0 | 0 | 0 | 0 | 0 | 0 | 0 | 7 | 0 | 179 | 107 | 59 | 181 | 34 | 0 | 36 | 18 | 0 | 0 |
| PBANKA_1465921 | 0 | 0 | 0 | 0 | 0 | 0 | 0 | 0 | 0 | 0 | 6 | 11 | 28 | 17 | 7 | 37 | 6 | 0 | 127 | 74 | 0 | 10 |
| PBANKA_1466181 | 0 | 0 | 0 | 0 | 0 | 0 | 0 | 0 | 0 | 0 | 5 | 0 | 16 | 18 | 29 | 46 | 10 | 6 | 30 | 20 | 0 | 0 |
| PBANKA_1466221 | 0 | 0 | 0 | 0 | 0 | 0 | 0 | 0 | 0 | 0 | 8 | 5 | 0 | 0 | 0 | 0 | 0 | 0 | 0 | 0 | 0 | 0 |
| PBANKA_1466241 | 0 | 0 | 0 | 0 | 0 | 0 | 0 | 0 | 0 | 0 | 30 | 28 | 47 | 13 | 0 | 0 | 0 | 0 | 0 | 0 | 0 | 0 |
| PBANKA_0722961 | 0 | 0 | 0 | 0 | 0 | 0 | 0 | 0 | 0 | 0 | 0 | 21 | 0 | 8 | 0 | 0 | 0 | 0 | 6 | 0 | 0 | 0 |
| PBANKA_0837201 | 0 | 0 | 0 | 0 | 0 | 0 | 0 | 0 | 0 | 0 | 0 | 10 | 104 | 133 | 23 | 108 | 57 | 12 | 69 | 50 | 0 | 0 |
| PBANKA_1040541 | 0 | 0 | 0 | 0 | 0 | 0 | 0 | 0 | 0 | 0 | 0 | 6 | 104 | 32 | 16 | 17 | 9 | 0 | 578 | 138 | 0 | 21 |
| PBANKA_1200051 | 0 | 0 | 0 | 0 | 0 | 0 | 0 | 0 | 0 | 0 | 0 | 8 | 0 | 0 | 0 | 0 | 0 | 0 | 0 | 0 | 0 | 0 |
| PBANKA_1200081 | 0 | 0 | 0 | 0 | 0 | 0 | 0 | 0 | 0 | 0 | 0 | 20 | 33 | 19 | 25 | 18 | 8 | 0 | 369 | 74 | 0 | 8 |
| PBANKA_1246981 | 0 | 0 | 0 | 0 | 0 | 0 | 0 | 0 | 0 | 0 | 0 | 5 | 0 | 0 | 0 | 0 | 0 | 0 | 0 | 0 | 0 | 0 |
| PBANKA_1400011 | 0 | 0 | 0 | 0 | 0 | 0 | 0 | 0 | 0 | 0 | 0 | 7 | 0 | 0 | 0 | 0 | 0 | 0 | 93 | 46 | 0 | 0 |
| PBANKA_1400061 | 0 | 0 | 0 | 0 | 0 | 0 | 0 | 0 | 0 | 0 | 0 | 11 | 529 | 261 | 66 | 121 | 16 | 17 | 107 | 43 | 0 | 0 |
| PBANKA_1465800 | 0 | 0 | 0 | 0 | 0 | 0 | 0 | 0 | 0 | 0 | 0 | 12 | 5 | 0 | 0 | 5 | 0 | 0 | 14 | 0 | 0 | 0 |
| PBANKA_0007901 | 0 | 0 | 0 | 0 | 0 | 0 | 0 | 0 | 0 | 0 | 0 | 0 | 0 | 0 | 0 | 0 | 0 | 0 | 0 | 0 | 0 | 0 |
| PBANKA_0216061 | 0 | 0 | 0 | 0 | 0 | 0 | 0 | 0 | 0 | 0 | 0 | 0 | 368 | 33 | 38 | 13 | 0 | 0 | 26 | 9 | 0 | 6 |
| PBANKA_0300400 | 0 | 0 | 0 | 0 | 0 | 0 | 0 | 0 | 0 | 0 | 0 | 0 | 20 | 24 | 0 | 20 | 0 | 0 | 26 | 20 | 0 | 0 |
| PBANKA_0316650 | 0 | 0 | 0 | 0 | 0 | 0 | 0 | 0 | 0 | 0 | 0 | 0 | 0 | 0 | 0 | 0 | 17 | 0 | 8 | 5 | 0 | 0 |
| PBANKA_0836900 | 0 | 0 | 0 | 0 | 0 | 0 | 0 | 0 | 0 | 0 | 0 | 0 | 59 | 37 | 19 | 58 | 15 | 0 | 562 | 171 | 0 | 6 |
| PBANKA_0837081 | 0 | 0 | 0 | 0 | 0 | 0 | 0 | 0 | 0 | 0 | 0 | 0 | 35 | 10 | 0 | 0 | 0 | 0 | 13 | 6 | 0 | 0 |
| PBANKA_0837161 | 0 | 0 | 0 | 0 | 0 | 0 | 0 | 0 | 0 | 0 | 0 | 0 | 39 | 5 | 11 | 8 | 0 | 0 | 451 | 130 | 0 | 0 |
| PBANKA_1200061 | 0 | 0 | 0 | 0 | 0 | 0 | 0 | 0 | 0 | 0 | 0 | 0 | 8 | 0 | 0 | 0 | 55 | 6 | 0 | 5 | 0 | 8 |
| PBANKA_1365700 | 0 | 0 | 0 | 0 | 0 | 0 | 0 | 0 | 0 | 0 | 0 | 0 | 184 | 72 | 47 | 186 | 48 | 11 | 561 | 391 | 0 | 28 |
| PBANKA_0316941 | 0 | 0 | 0 | 0 | 0 | 0 | 0 | 0 | 0 | 0 | 0 | 0 | 462 | 302 | 42 | 419 | 16 | 8 | 59 | 9 | 0 | 0 |
| PBANKA_0722100 | 0 | 0 | 0 | 0 | 0 | 0 | 0 | 0 | 0 | 0 | 0 | 0 | 37 | 0 | 6 | 0 | 0 | 0 | 12 | 6 | 0 | 5 |
| PBANKA_0900600 | 0 | 0 | 0 | 0 | 0 | 0 | 0 | 0 | 0 | 0 | 0 | 0 | 0 | 23 | 30 | 5 | 0 | 0 | 181 | 108 | 0 | 0 |
| PBANKA_1403250 | 0 | 0 | 0 | 0 | 0 | 0 | 0 | 0 | 0 | 0 | 0 | 0 | 21 | 9 | 30 | 15 | 19 | 0 | 26 | 21 | 0 | 0 |
| PBANKA_1465821 | 0 | 0 | 0 | 0 | 0 | 0 | 0 | 0 | 0 | 0 | 0 | 0 | 0 | 0 | 0 | 0 | 12 | 0 | 0 | 0 | 0 | 0 |
| PBANKA_0105100 | 0 | 0 | 0 | 0 | 0 | 0 | 0 | 0 | 0 | 0 | 0 | 0 | 362 | 168 | 10 | 27 | 123 | 53 | 7741 | 4272 | 66 | 392 |
| PBANKA_0110800 | 0 | 0 | 0 | 0 | 0 | 0 | 0 | 0 | 0 | 0 | 0 | 0 | 618 | 134 | 8 | 96 | 161 | 94 | 14818 | 6240 | 266 | 1606 |
| PBANKA_0602900 | 0 | 0 | 0 | 0 | 0 | 0 | 0 | 0 | 0 | 0 | 0 | 0 | 152 | 19 | 7 | 14 | 40 | 22 | 1387 | 353 | 6 | 66 |
| PBANKA_0916650 | 487 | 573 | 0 | 0 | 0 | 0 | 0 | 0 | 0 | 0 | 0 | 0 | 33 | 12 | 0 | 6 | 25 | 0 | 834 | 512 | 353 | 2037 |
| PBANKA_0917990 | 0 | 0 | 0 | 0 | 0 | 0 | 0 | 0 | 0 | 0 | 0 | 0 | 83 | 6 | 38 | 13 | 6 | 6 | 939 | 11 | 12 | 53 |
| PBANKA_0941000 | 0 | 7 | 0 | 0 | 0 | 0 | 0 | 0 | 0 | 0 | 0 | 0 | 39 | 17 | 16 | 18 | 21 | 5 | 101 | 48 | 312 | 2209 |
| PBANKA_1018200 | 0 | 7 | 0 | 0 | 0 | 0 | 0 | 0 | 0 | 0 | 0 | 0 | 144 | 42 | 5 | 27 | 44 | 22 | 1053 | 582 | 20 | 181 |
| PBANKA_1019000 | 0 | 0 | 0 | 0 | 0 | 0 | 0 | 0 | 0 | 0 | 0 | 0 | 86 | 20 | 6 | 10 | 60 | 18 | 842 | 269 | 379 | 1776 |
| PBANKA_1040200 | 0 | 0 | 0 | 0 | 0 | 0 | 0 | 0 | 0 | 0 | 0 | 0 | 78 | 37 | 15 | 84 | 19 | 0 | 60 | 21 | 6 | 24 |
| PBANKA_1132800 | 0 | 0 | 0 | 0 | 0 | 0 | 0 | 0 | 0 | 0 | 0 | 0 | 84 | 25 | 9 | 24 | 68 | 22 | 1335 | 620 | 132 | 1026 |
| PBANKA_1203450 | 0 | 0 | 0 | 0 | 0 | 0 | 0 | 0 | 0 | 0 | 0 | 0 | 9 | 0 | 0 | 0 | 0 | 9 | 85 | 26 | 24 | 228 |
| PBANKA_1204500 | 0 | 0 | 0 | 0 | 0 | 0 | 0 | 0 | 0 | 0 | 0 | 0 | 324 | 109 | 7 | 8 | 70 | 52 | 6399 | 2798 | 141 | 573 |
| PBANKA_1400900 | 0 | 0 | 0 | 0 | 0 | 0 | 0 | 0 | 0 | 0 | 0 | 0 | 103 | 17 | 5 | 6 | 52 | 47 | 1326 | 424 | 85 | 654 |
| PBANKA_1421700 | 0 | 0 | 0 | 0 | 0 | 0 | 0 | 0 | 0 | 0 | 0 | 0 | 1073 | 305 | 20 | 30 | 130 | 128 | 36040 | 10980 | 41 | 235 |
| PBANKA_0100081 | 0 | 0 | 0 | 0 | 0 | 0 | 0 | 0 | 0 | 0 | 0 | 0 | 40 | 0 | 5 | 21 | 0 | 0 | 60 | 16 | 0 | 6 |
| PBANKA_0315420 | 0 | 0 | 0 | 0 | 0 | 0 | 0 | 0 | 0 | 0 | 0 | 0 | 14 | 27 | 10 | 18 | 27 | 9 | 30 | 0 | 0 | 8 |
| PBANKA_0316861 | 0 | 0 | 0 | 0 | 0 | 0 | 0 | 0 | 0 | 0 | 0 | 0 | 132 | 69 | 41 | 101 | 20 | 0 | 255 | 103 | 0 | 30 |
| PBANKA_0317001 | 0 | 0 | 0 | 0 | 0 | 0 | 0 | 0 | 0 | 0 | 0 | 0 | 64 | 18 | 9 | 20 | 0 | 0 | 23 | 16 | 0 | 7 |
| PBANKA_0405310 | 0 | 0 | 0 | 0 | 0 | 0 | 0 | 0 | 0 | 0 | 0 | 0 | 21 | 15 | 27 | 29 | 17 | 0 | 6 | 11 | 0 | 28 |
| PBANKA_0609250 | 0 | 0 | 0 | 0 | 0 | 0 | 0 | 0 | 0 | 0 | 0 | 0 | 9 | 0 | 9 | 17 | 38 | 0 | 120 | 11 | 0 | 11 |
| PBANKA_0700541 | 0 | 0 | 0 | 0 | 0 | 0 | 0 | 0 | 0 | 0 | 0 | 0 | 223 | 102 | 51 | 157 | 48 | 7 | 179 | 106 | 0 | 36 |
| PBANKA_0703910 | 0 | 0 | 0 | 0 | 0 | 0 | 0 | 0 | 0 | 0 | 0 | 0 | 60 | 11 | 72 | 9 | 15 | 0 | 346 | 10 | 0 | 5 |
| PBANKA_0722900 | 0 | 0 | 0 | 0 | 0 | 0 | 0 | 0 | 0 | 0 | 0 | 0 | 200 | 34 | 148 | 113 | 26 | 7 | 76 | 6 | 0 | 11 |
| PBANKA_0836400 | 0 | 0 | 0 | 0 | 0 | 0 | 0 | 0 | 0 | 0 | 0 | 0 | 195 | 39 | 528 | 1810 | 390 | 47 | 156 | 27 | 0 | 9 |
| PBANKA_0943700 | 0 | 0 | 0 | 0 | 0 | 0 | 0 | 0 | 0 | 0 | 0 | 0 | 46 | 29 | 7 | 8 | 0 | 0 | 687 | 477 | 0 | 52 |
| PBANKA_1000081 | 0 | 0 | 0 | 0 | 0 | 0 | 0 | 0 | 0 | 0 | 0 | 0 | 382 | 153 | 248 | 460 | 2218 | 334 | 111 | 79 | 0 | 5 |
| PBANKA_1000400 | 0 | 0 | 0 | 0 | 0 | 0 | 0 | 0 | 0 | 0 | 0 | 0 | 121 | 58 | 70 | 219 | 15 | 6 | 127 | 46 | 0 | 7 |
| PBANKA_1019380 | 0 | 0 | 0 | 0 | 0 | 0 | 0 | 0 | 0 | 0 | 0 | 0 | 24 | 46 | 23 | 16 | 6 | 7 | 10 | 0 | 0 | 17 |
| PBANKA_1040400 | 0 | 0 | 0 | 0 | 0 | 0 | 0 | 0 | 0 | 0 | 0 | 0 | 80 | 39 | 16 | 97 | 18 | 0 | 75 | 55 | 0 | 6 |
| PBANKA_1040621 | 0 | 0 | 0 | 0 | 0 | 0 | 0 | 0 | 0 | 0 | 0 | 0 | 25 | 9 | 11 | 7 | 0 | 0 | 275 | 129 | 0 | 9 |
| PBANKA_1226470 | 0 | 0 | 0 | 0 | 0 | 0 | 0 | 0 | 0 | 0 | 0 | 0 | 19 | 7 | 24 | 10 | 0 | 10 | 136 | 7 | 0 | 6 |
| PBANKA_1246200 | 0 | 0 | 0 | 0 | 0 | 0 | 0 | 0 | 0 | 0 | 0 | 0 | 270 | 79 | 49 | 67 | 69 | 32 | 127 | 30 | 0 | 9 |
| PBANKA_1246300 | 0 | 0 | 0 | 0 | 0 | 0 | 0 | 0 | 0 | 0 | 0 | 0 | 618 | 197 | 71 | 527 | 90 | 12 | 608 | 266 | 0 | 5 |
| PBANKA_1343100 | 0 | 0 | 0 | 0 | 0 | 0 | 0 | 0 | 0 | 0 | 0 | 0 | 25 | 8 | 0 | 6 | 29 | 16 | 299 | 87 | 0 | 15 |
| PBANKA_1346750 | 0 | 0 | 0 | 0 | 0 | 0 | 0 | 0 | 0 | 0 | 0 | 0 | 41 | 10 | 71 | 19 | 12 | 10 | 137 | 13 | 0 | 6 |
| PBANKA_1354550 | 0 | 0 | 0 | 0 | 0 | 0 | 0 | 0 | 0 | 0 | 0 | 0 | 31 | 0 | 51 | 0 | 0 | 7 | 36 | 0 | 0 | 11 |
| PBANKA_1400100 | 0 | 0 | 0 | 0 | 0 | 0 | 0 | 0 | 0 | 0 | 0 | 0 | 186 | 59 | 60 | 201 | 40 | 12 | 621 | 345 | 0 | 28 |

| | | | | | | | | | | | | | | | | | | | | | | |
|---|---|---|---|---|---|---|---|---|---|---|---|---|---|---|---|---|---|---|---|---|---|---|
| PBANKA_1400300 | 0 | 0 | 0 | 0 | 0 | 0 | 0 | 0 | 0 | 0 | 0 | 0 | 75 | 55 | 33 | 86 | 26 | 7 | 84 | 58 | 0 | 14 |
| PBANKA_1431200 | 0 | 0 | 0 | 0 | 0 | 0 | 0 | 0 | 0 | 0 | 0 | 0 | 16 | 6 | 0 | 7 | 5 | 12 | 136 | 74 | 0 | 27 |
| PBANKA_1435000 | 0 | 5 | 0 | 0 | 0 | 0 | 0 | 0 | 0 | 0 | 0 | 0 | 0 | 0 | 0 | 0 | 5 | 0 | 27 | 7 | 0 | 12 |
| PBANKA_1465200 | 0 | 0 | 0 | 0 | 0 | 0 | 0 | 0 | 0 | 0 | 0 | 0 | 143 | 45 | 25 | 38 | 10 | 0 | 407 | 38 | 0 | 5 |
| PBANKA_1466141 | 0 | 0 | 0 | 0 | 0 | 0 | 0 | 0 | 0 | 0 | 0 | 0 | 48 | 76 | 16 | 60 | 25 | 9 | 55 | 35 | 0 | 10 |
| PBANKA_0000401 | 0 | 0 | 0 | 0 | 0 | 0 | 0 | 0 | 0 | 0 | 0 | 0 | 6 | 0 | 0 | 0 | 0 | 0 | 0 | 0 | 0 | 0 |
| PBANKA_0000801 | 0 | 0 | 0 | 0 | 0 | 0 | 0 | 0 | 0 | 0 | 0 | 0 | 26 | 0 | 8 | 10 | 0 | 0 | 157 | 18 | 0 | 0 |
| PBANKA_0000901 | 0 | 0 | 0 | 0 | 0 | 0 | 0 | 0 | 0 | 0 | 0 | 0 | 91 | 119 | 35 | 66 | 32 | 10 | 65 | 43 | 0 | 0 |
| PBANKA_0200751 | 0 | 0 | 0 | 0 | 0 | 0 | 0 | 0 | 0 | 0 | 0 | 0 | 13 | 18 | 8 | 20 | 5 | 0 | 12 | 5 | 0 | 0 |
| PBANKA_0201100 | 0 | 0 | 0 | 0 | 0 | 0 | 0 | 0 | 0 | 0 | 0 | 0 | 8 | 7 | 5 | 9 | 0 | 0 | 44 | 43 | 0 | 0 |
| PBANKA_0201360 | 0 | 0 | 0 | 0 | 0 | 0 | 0 | 0 | 0 | 0 | 0 | 0 | 13 | 15 | 20 | 17 | 10 | 5 | 28 | 10 | 0 | 0 |
| PBANKA_0201400 | 0 | 0 | 0 | 0 | 0 | 0 | 0 | 0 | 0 | 0 | 0 | 0 | 75 | 57 | 66 | 211 | 73 | 22 | 53 | 22 | 0 | 0 |
| PBANKA_0201450 | 0 | 0 | 0 | 0 | 0 | 0 | 0 | 0 | 0 | 0 | 0 | 0 | 33 | 7 | 54 | 226 | 78 | 14 | 56 | 17 | 0 | 0 |
| PBANKA_0215700 | 0 | 0 | 0 | 0 | 0 | 0 | 0 | 0 | 0 | 0 | 0 | 0 | 52 | 42 | 19 | 37 | 0 | 0 | 64 | 18 | 0 | 0 |
| PBANKA_0215900 | 0 | 0 | 0 | 0 | 0 | 0 | 0 | 0 | 0 | 0 | 0 | 0 | 18 | 0 | 0 | 0 | 6 | 0 | 9 | 0 | 0 | 0 |
| PBANKA_0216701 | 0 | 0 | 0 | 0 | 0 | 0 | 0 | 0 | 0 | 0 | 0 | 0 | 62 | 12 | 30 | 161 | 15 | 0 | 72 | 32 | 0 | 0 |
| PBANKA_0315430 | 0 | 0 | 0 | 0 | 0 | 0 | 0 | 0 | 0 | 0 | 0 | 0 | 13 | 0 | 14 | 5 | 5 | 7 | 42 | 0 | 0 | 0 |
| PBANKA_0317041 | 0 | 0 | 0 | 0 | 0 | 0 | 0 | 0 | 0 | 0 | 0 | 0 | 89 | 43 | 11 | 73 | 12 | 0 | 33 | 23 | 0 | 0 |
| PBANKA_0317081 | 0 | 0 | 0 | 0 | 0 | 0 | 0 | 0 | 0 | 0 | 0 | 0 | 29 | 0 | 52 | 60 | 40 | 17 | 9 | 0 | 0 | 0 |
| PBANKA_0317101 | 0 | 0 | 0 | 0 | 0 | 0 | 0 | 0 | 0 | 0 | 0 | 0 | 16 | 0 | 10 | 15 | 0 | 0 | 19 | 10 | 0 | 0 |
| PBANKA_0500200 | 0 | 0 | 0 | 0 | 0 | 0 | 0 | 0 | 0 | 0 | 0 | 0 | 183 | 199 | 20 | 134 | 21 | 0 | 43 | 6 | 0 | 0 |
| PBANKA_0500721 | 0 | 0 | 0 | 0 | 0 | 0 | 0 | 0 | 0 | 0 | 0 | 0 | 466 | 252 | 215 | 970 | 162 | 26 | 75 | 65 | 0 | 0 |
| PBANKA_0722200 | 0 | 0 | 0 | 0 | 0 | 0 | 0 | 0 | 0 | 0 | 0 | 0 | 30 | 5 | 14 | 10 | 0 | 0 | 6 | 0 | 0 | 0 |
| PBANKA_0722700 | 0 | 0 | 0 | 0 | 0 | 0 | 0 | 0 | 0 | 0 | 0 | 0 | 7 | 0 | 6 | 0 | 0 | 0 | 6 | 0 | 0 | 0 |
| PBANKA_0800061 | 0 | 0 | 0 | 0 | 0 | 0 | 0 | 0 | 0 | 0 | 0 | 0 | 22 | 0 | 12 | 18 | 0 | 0 | 50 | 16 | 0 | 0 |
| PBANKA_0836921 | 0 | 0 | 0 | 0 | 0 | 0 | 0 | 0 | 0 | 0 | 0 | 0 | 5 | 5 | 0 | 0 | 0 | 0 | 0 | 7 | 0 | 0 |
| PBANKA_0837181 | 0 | 0 | 0 | 0 | 0 | 0 | 0 | 0 | 0 | 0 | 0 | 0 | 21 | 0 | 8 | 7 | 0 | 0 | 151 | 11 | 0 | 0 |
| PBANKA_1000031 | 0 | 0 | 0 | 0 | 0 | 0 | 0 | 0 | 0 | 0 | 0 | 0 | 520 | 148 | 61 | 300 | 24 | 0 | 48 | 30 | 0 | 0 |
| PBANKA_1000061 | 0 | 0 | 0 | 0 | 0 | 0 | 0 | 0 | 0 | 0 | 0 | 0 | 99 | 38 | 159 | 126 | 34 | 14 | 16 | 5 | 0 | 0 |
| PBANKA_1000451 | 0 | 0 | 0 | 0 | 0 | 0 | 0 | 0 | 0 | 0 | 0 | 0 | 28 | 8 | 11 | 23 | 0 | 0 | 33 | 6 | 0 | 0 |
| PBANKA_1019540 | 0 | 0 | 0 | 0 | 0 | 0 | 0 | 0 | 0 | 0 | 0 | 0 | 33 | 6 | 93 | 16 | 11 | 10 | 54 | 12 | 0 | 0 |
| PBANKA_1019560 | 0 | 0 | 0 | 0 | 0 | 0 | 0 | 0 | 0 | 0 | 0 | 0 | 11 | 0 | 25 | 11 | 0 | 0 | 0 | 0 | 0 | 0 |
| PBANKA_1040500 | 0 | 0 | 0 | 0 | 0 | 0 | 0 | 0 | 0 | 0 | 0 | 0 | 43 | 0 | 0 | 0 | 0 | 0 | 45 | 10 | 0 | 0 |
| PBANKA_1100600 | 0 | 0 | 0 | 0 | 0 | 0 | 0 | 0 | 0 | 0 | 0 | 0 | 0 | 7 | 0 | 0 | 0 | 0 | 0 | 0 | 0 | 0 |
| PBANKA_1100750 | 0 | 0 | 0 | 0 | 0 | 0 | 0 | 0 | 0 | 0 | 0 | 0 | 47 | 47 | 41 | 74 | 40 | 5 | 7 | 5 | 0 | 0 |
| PBANKA_1100821 | 0 | 0 | 0 | 0 | 0 | 0 | 0 | 0 | 0 | 0 | 0 | 0 | 5 | 0 | 0 | 0 | 0 | 0 | 0 | 0 | 0 | 0 |
| PBANKA_1146100 | 0 | 0 | 0 | 0 | 0 | 0 | 0 | 0 | 0 | 0 | 0 | 0 | 262 | 183 | 195 | 347 | 196 | 57 | 67 | 45 | 0 | 0 |
| PBANKA_1200300 | 0 | 0 | 0 | 0 | 0 | 0 | 0 | 0 | 0 | 0 | 0 | 0 | 8 | 0 | 8 | 12 | 0 | 0 | 26 | 16 | 0 | 0 |
| PBANKA_1246121 | 0 | 0 | 0 | 0 | 0 | 0 | 0 | 0 | 0 | 0 | 0 | 0 | 0 | 0 | 5 | 0 | 0 | 0 | 17 | 0 | 0 | 0 |
| PBANKA_1300061 | 0 | 0 | 0 | 0 | 0 | 0 | 0 | 0 | 0 | 0 | 0 | 0 | 5 | 0 | 0 | 0 | 13 | 6 | 0 | 5 | 0 | 0 |
| PBANKA_1300071 | 0 | 0 | 0 | 0 | 0 | 0 | 0 | 0 | 0 | 0 | 0 | 0 | 16 | 9 | 11 | 25 | 0 | 0 | 10 | 5 | 0 | 0 |
| PBANKA_1400021 | 0 | 0 | 0 | 0 | 0 | 0 | 0 | 0 | 0 | 0 | 0 | 0 | 43 | 0 | 12 | 10 | 0 | 0 | 0 | 0 | 0 | 0 |
| PBANKA_1465841 | 0 | 0 | 0 | 0 | 0 | 0 | 0 | 0 | 0 | 0 | 0 | 0 | 24 | 0 | 13 | 0 | 0 | 0 | 257 | 83 | 0 | 0 |
| PBANKA_1465861 | 0 | 0 | 0 | 0 | 0 | 0 | 0 | 0 | 0 | 0 | 0 | 0 | 31 | 16 | 18 | 0 | 0 | 0 | 35 | 0 | 0 | 0 |
| PBANKA_1465881 | 0 | 0 | 0 | 0 | 0 | 0 | 0 | 0 | 0 | 0 | 0 | 0 | 0 | 0 | 0 | 5 | 0 | 0 | 0 | 0 | 0 | 0 |
| PBANKA_1466100 | 0 | 0 | 0 | 0 | 0 | 0 | 0 | 0 | 0 | 0 | 0 | 0 | 30 | 13 | 0 | 14 | 7 | 0 | 171 | 77 | 0 | 0 |
| PBANKA_1466161 | 0 | 0 | 0 | 0 | 0 | 0 | 0 | 0 | 0 | 0 | 0 | 0 | 5 | 0 | 5 | 10 | 0 | 0 | 92 | 11 | 0 | 0 |
| PBANKA_MIT02300 | 0 | 0 | 0 | 0 | 0 | 0 | 0 | 0 | 0 | 0 | 0 | 0 | 5 | 0 | 66 | 16 | 10 | 6 | 18 | 0 | 0 | 0 |
| PBANKA_0007701 | 0 | 0 | 0 | 0 | 0 | 0 | 0 | 0 | 0 | 0 | 0 | 0 | 11 | 0 | 0 | 11 | 36 | 11 | 0 | 0 | 0 | 0 |
| PBANKA_1200500 | 0 | 0 | 0 | 0 | 0 | 0 | 0 | 0 | 0 | 0 | 0 | 0 | 7 | 0 | 0 | 8 | 0 | 0 | 13 | 6 | 0 | 0 |
| PBANKA_API00290 | 0 | 0 | 0 | 0 | 0 | 0 | 0 | 0 | 0 | 0 | 0 | 0 | 5 | 0 | 0 | 0 | 5 | 0 | 0 | 0 | 0 | 0 |
| PBANKA_MIT02400 | 0 | 0 | 0 | 0 | 0 | 0 | 0 | 0 | 0 | 0 | 0 | 0 | 5 | 29 | 231 | 93 | 97 | 108 | 25 | 93 | 0 | 0 |
| PBANKA_MIT02900 | 0 | 0 | 0 | 0 | 0 | 0 | 0 | 0 | 0 | 0 | 0 | 0 | 26 | 5 | 500 | 42 | 37 | 66 | 166 | 28 | 0 | 0 |
| PBANKA_0200670 | 0 | 0 | 0 | 0 | 0 | 0 | 0 | 0 | 0 | 0 | 0 | 0 | 5 | 10 | 0 | 9 | 0 | 0 | 9 | 0 | 0 | 0 |
| PBANKA_0406650 | 0 | 0 | 0 | 0 | 0 | 0 | 0 | 0 | 0 | 0 | 0 | 0 | 8 | 0 | 92 | 12 | 15 | 10 | 24 | 10 | 0 | 0 |
| PBANKA_0407550 | 0 | 0 | 0 | 0 | 0 | 0 | 0 | 0 | 0 | 0 | 0 | 0 | 18 | 0 | 51 | 11 | 0 | 0 | 244 | 10 | 0 | 0 |
| PBANKA_0937750 | 0 | 0 | 0 | 0 | 0 | 0 | 0 | 0 | 0 | 0 | 0 | 0 | 22 | 0 | 60 | 6 | 6 | 0 | 118 | 0 | 0 | 0 |
| PBANKA_0938050 | 0 | 0 | 0 | 0 | 0 | 0 | 0 | 0 | 0 | 0 | 0 | 0 | 37 | 6 | 44 | 6 | 6 | 0 | 126 | 7 | 0 | 0 |
| PBANKA_0944021 | 0 | 0 | 0 | 0 | 0 | 0 | 0 | 0 | 0 | 0 | 0 | 0 | 11 | 0 | 0 | 0 | 23 | 10 | 0 | 0 | 0 | 0 |
| PBANKA_1000651 | 0 | 0 | 0 | 0 | 0 | 0 | 0 | 0 | 0 | 0 | 0 | 0 | 5 | 0 | 0 | 0 | 0 | 0 | 0 | 0 | 0 | 0 |
| PBANKA_1019370 | 0 | 0 | 0 | 0 | 0 | 0 | 0 | 0 | 0 | 0 | 0 | 0 | 37 | 0 | 32 | 0 | 0 | 0 | 9 | 0 | 0 | 0 |
| PBANKA_1019580 | 0 | 0 | 0 | 0 | 0 | 0 | 0 | 0 | 0 | 0 | 0 | 0 | 22 | 7 | 60 | 25 | 7 | 5 | 48 | 0 | 0 | 0 |
| PBANKA_1039450 | 0 | 0 | 0 | 0 | 0 | 0 | 0 | 0 | 0 | 0 | 0 | 0 | 11 | 0 | 23 | 5 | 0 | 0 | 10 | 0 | 0 | 0 |
| PBANKA_0937840 | 0 | 0 | 0 | 0 | 0 | 0 | 0 | 0 | 0 | 0 | 0 | 0 | 0 | 5 | 7 | 0 | 0 | 0 | 5 | 0 | 0 | 0 |
| PBANKA_1145741 | 0 | 0 | 0 | 0 | 0 | 0 | 0 | 0 | 0 | 0 | 0 | 0 | 0 | 6 | 0 | 0 | 0 | 0 | 0 | 0 | 0 | 0 |
| PBANKA_0612841 | 0 | 0 | 0 | 0 | 0 | 0 | 0 | 0 | 0 | 0 | 0 | 0 | 0 | 0 | 6 | 0 | 0 | 0 | 0 | 0 | 0 | 0 |
| PBANKA_1345850 | 0 | 0 | 0 | 0 | 0 | 0 | 0 | 0 | 0 | 0 | 0 | 0 | 0 | 0 | 44 | 0 | 0 | 0 | 6 | 0 | 0 | 0 |
| PBANKA_1437250 | 0 | 0 | 0 | 0 | 0 | 0 | 0 | 0 | 0 | 0 | 0 | 0 | 0 | 0 | 12 | 13 | 7 | 0 | 43 | 7 | 0 | 0 |
| PBANKA_MIT00200 | 0 | 0 | 0 | 0 | 0 | 0 | 0 | 0 | 0 | 0 | 0 | 0 | 0 | 0 | 5 | 0 | 0 | 0 | 0 | 0 | 0 | 0 |
| PBANKA_MIT00300 | 0 | 0 | 0 | 0 | 0 | 0 | 0 | 0 | 0 | 0 | 0 | 0 | 0 | 0 | 24 | 0 | 0 | 31 | 6 | 0 | 0 | 0 |
| PBANKA_0216081 | 0 | 0 | 0 | 0 | 0 | 0 | 0 | 0 | 0 | 0 | 0 | 0 | 0 | 0 | 0 | 6 | 0 | 0 | 8 | 0 | 0 | 0 |
| PBANKA_0836850 | 0 | 0 | 0 | 0 | 0 | 0 | 0 | 0 | 0 | 0 | 0 | 0 | 0 | 0 | 0 | 7 | 0 | 0 | 9 | 0 | 0 | 0 |
| PBANKA_1100031 | 0 | 0 | 0 | 0 | 0 | 0 | 0 | 0 | 0 | 0 | 0 | 0 | 0 | 0 | 0 | 5 | 0 | 0 | 0 | 0 | 0 | 0 |
| PBANKA_1246921 | 0 | 0 | 0 | 0 | 0 | 0 | 0 | 0 | 0 | 0 | 0 | 0 | 0 | 0 | 0 | 0 | 15 | 8 | 0 | 0 | 0 | 0 |
| PBANKA_1421261 | 0 | 0 | 0 | 0 | 0 | 0 | 0 | 0 | 0 | 0 | 0 | 0 | 0 | 0 | 0 | 0 | 5 | 0 | 0 | 0 | 0 | 0 |
| PBANKA_1466201 | 0 | 0 | 0 | 0 | 0 | 0 | 0 | 0 | 0 | 0 | 0 | 0 | 0 | 0 | 0 | 0 | 13 | 0 | 0 | 0 | 0 | 0 |
| PBANKA_0600096 | 0 | 0 | 0 | 0 | 0 | 0 | 0 | 0 | 0 | 0 | 0 | 0 | 0 | 0 | 0 | 0 | 0 | 0 | 5 | 7 | 0 | 0 |
| PBANKA_0837041 | 0 | 0 | 0 | 0 | 0 | 0 | 0 | 0 | 0 | 0 | 0 | 0 | 0 | 0 | 0 | 0 | 0 | 0 | 17 | 14 | 0 | 0 |
| PBANKA_0837061 | 0 | 0 | 0 | 0 | 0 | 0 | 0 | 0 | 0 | 0 | 0 | 0 | 0 | 0 | 0 | 0 | 0 | 0 | 5 | 6 | 0 | 0 |
| PBANKA_1302241 | 0 | 0 | 0 | 0 | 0 | 0 | 0 | 0 | 0 | 0 | 0 | 0 | 0 | 0 | 0 | 0 | 0 | 0 | 6 | 0 | 0 | 0 |
| PBANKA_1354351 | 0 | 0 | 0 | 0 | 0 | 0 | 0 | 0 | 0 | 0 | 0 | 0 | 0 | 0 | 0 | 0 | 0 | 0 | 5 | 0 | 0 | 0 |
| PBANKA_1400086 | 0 | 0 | 0 | 0 | 0 | 0 | 0 | 0 | 0 | 0 | 0 | 0 | 0 | 0 | 0 | 0 | 0 | 0 | 8 | 0 | 0 | 0 |
| PBANKA_1351351 | 0 | 0 | 0 | 0 | 0 | 0 | 0 | 0 | 0 | 0 | 0 | 0 | 0 | 0 | 0 | 0 | 0 | 0 | 0 | 6 | 0 | 0 |
