## Supplementary material for "Transcriptome analysis of *Plasmodium berghei* during exo-erythrocytic development": Table S3

BP: Biological Process  CC: Cellular Component  MF: Molecular Function

| | BP | Process Description | Annotated | Observed | pValue |
|---|---|---|---|---|---|
| 1 | GO:0010467 | gene expression | 462 | 126 | 8.8e-23 |
| | GO:0042254 | ribosome biogenesis | 63 | 36 | 1.2e-17 |
| | GO:0097659 | nucleic acid-templated transcription | 100 | 23 | 0.0002 |
| | GO:0008295 | spermidine biosynthetic process | 2 | 2 | 0.0131 |
| | GO:0000290 | deadenylation-dependent decapping of nuclear-transcribed mRNA | 3 | 2 | 0.0363 |

| | CC | Process Description | Annotated | Observed | pValue |
|---|---|---|---|---|---|
| | GO:0030529 | intracellular ribonucleoprotein complex | 220 | 88 | < 1e-30 |
| | GO:0043232 | intracellular non-membrane-bounded organelle | 264 | 77 | 3.5e-19 |
| | GO:0005852 | eukaryotic translation initiation factor 3 complex | 10 | 7 | 2.5e-05 |
| | GO:0005666 | DNA-directed RNA polymerase III complex | 8 | 4 | 0.009 |
| | GO:0044228 | host cell surface | 2 | 2 | 0.014 |
| | GO:0005665 | DNA-directed RNA polymerase II, core complex | 7 | 3 | 0.040 |

| | MF | Process Description | Annotated | Observed | pValue |
|---|---|---|---|---|---|
| | GO:0003735 | structural constituent of ribosome | 129 | 49 | 9.4e-15 |
| | GO:0003676 | nucleic acid binding | 481 | 112 | 3.6e-12 |
| | GO:0003724 | RNA helicase activity | 10 | 7 | 3.5e-05 |
| | GO:0003899 | DNA-directed 5'-3' RNA polymerase activity | 31 | 11 | 0.00072 |
| | GO:0003743 | translation initiation factor activity | 27 | 10 | 0.00086 |
| | GO:0003682 | chromatin binding | 9 | 5 | 0.00229 |
| | GO:0031072 | heat shock protein binding | 4 | 3 | 0.00677 |
| | GO:0000179 | rRNA (adenine-N6,N6-)-dimethyltransferase activity | 2 | 2 | 0.01520 |
| | GO:0017025 | TBP-class protein binding | 2 | 2 | 0.01520 |
| | GO:0036094 | small molecule binding | 554 | 73 | 0.03825 |
| | GO:0031369 | translation initiation factor binding | 3 | 2 | 0.04187 |

| | BP | Process Description | Annotated | Observed | pValue |
|---|---|---|---|---|---|
| 2 | GO:0006397 | mRNA processing | 57 | 22 | 1.5e-06 |
| | GO:0006412 | translation | 211 | 45 | 0.00013 |
| | GO:0032784 | regulation of DNA-templated transcription, elongation | 3 | 3 | 0.00194 |
| | GO:0032259 | methylation | 25 | 9 | 0.00777 |
| | GO:0042255 | ribosome assembly | 8 | 4 | 0.01114 |
| | GO:0043484 | regulation of RNA splicing | 2 | 2 | 0.01563 |
| | GO:0042026 | protein refolding | 3 | 2 | 0.04302 |
| | GO:0044053 | translocation of peptides or proteins into host cell cytoplasm | 3 | 2 | 0.04302 |

| | CC | Process Description | Annotated | Observed | pValue |
|---|---|---|---|---|---|
| | GO:0097619 | PTEX complex | 6 | 6 | 4.0e-06 |
| | GO:0033646 | host intracellular part | 85 | 28 | 7.4e-05 |
| | GO:0030529 | intracellular ribonucleoprotein complex | 220 | 49 | 0.00014 |
| | GO:0005850 | eukaryotic translation initiation factor 2 complex | 3 | 3 | 0.00206 |
| | GO:0020020 | food vacuole | 15 | 5 | 0.03302 |
| | GO:0005685 | U1 snRNP | 3 | 2 | 0.04462 |

| MF | Process Description | Annotated | Observed | pValue |
|---|---|---|---|---|
| GO:0003723 | RNA binding | 183 | 43 | 6.6e-06 |
| GO:0003735 | structural constituent of ribosome | 129 | 25 | 0.0091 |
| GO:0004812 | aminoacyl-tRNA ligase activity | 37 | 10 | 0.0098 |
| GO:0016763 | transferase activity, transferring pentosyl groups | 5 | 3 | 0.0144 |
| GO:0004749 | ribose phosphate diphosphokinase activity | 2 | 2 | 0.0145 |
| GO:0004526 | ribonuclease P activity | 2 | 2 | 0.0145 |
| GO:0017150 | tRNA dihydrouridine synthase activity | 2 | 2 | 0.0145 |
| GO:0003924 | GTPase activity | 52 | 12 | 0.0179 |
| GO:0002161 | aminoacyl-tRNA editing activity | 6 | 3 | 0.0263 |

3

| BP | Process Description | Annotated | Observed | pValue |
|---|---|---|---|---|
| GO:0032958 | inositol phosphate biosynthetic process | 1 | 1 | 0.0019 |
| GO:0006891 | intra-Golgi vesicle-mediated transport | 2 | 1 | 0.0037 |
| GO:0048870 | cell motility | 16 | 1 | 0.0295 |

| CC | Process Description | Annotated | Observed | pValue |
|---|---|---|---|---|
| GO:0030430 | host cell cytoplasm | 85 | 6 | 8e-06 |
| GO:0017119 | Golgi transport complex | 2 | 1 | 0.014 |
| GO:0031514 | motile cilium | 2 | 1 | 0.014 |

| MF | Process Description | Annotated | Observed | pValue |
|---|---|---|---|---|
| GO:0008440 | inositol-1,4,5-trisphosphate 3-kinase activity | 3 | 1 | 0.0056 |
| GO:0004252 | serine-type endopeptidase activity | 15 | 1 | 0.0276 |
| GO:0008026 | ATP-dependent helicase activity | 25 | 1 | 0.0457 |

4

| BP | Process Description | Annotated | Observed | pValue |
|---|---|---|---|---|
| GO:0016567 | protein ubiquitination | 6 | 1 | 0.018 |
| GO:0009405 | pathogenesis | 8 | 1 | 0.025 |

| CC | Process Description | Annotated | Observed | pValue |
|---|---|---|---|---|
| GO:0005680 | anaphase-promoting complex | 2 | 1 | 0.0055 |
| GO:0030131 | clathrin adaptor complex | 7 | 1 | 0.0191 |
| GO:0034399 | nuclear periphery | 12 | 1 | 0.0326 |

| MF | Process Description | Annotated | Observed | pValue |
|---|---|---|---|---|
| GO:0005515 | protein binding | 478 | 7 | 0.0016 |
| GO:0016307 | phosphatidylinositol phosphate kinase activity | 2 | 1 | 0.0093 |
| GO:0046872 | metal ion binding | 243 | 5 | 0.0170 |

5

| BP | Process Description | Annotated | Observed | pValue |
|---|---|---|---|---|
| GO:0051187 | cofactor catabolic process | 3 | 3 | 0.0083 |
| GO:0006470 | protein dephosphorylation | 10 | 4 | 0.0092 |
| GO:0034982 | mitochondrial protein processing | 3 | 2 | 0.0236 |
| GO:0015991 | ATP hydrolysis coupled proton transport | 14 | 4 | 0.0329 |
| GO:0042168 | heme metabolic process | 5 | 3 | 0.0439 |
| GO:0071897 | DNA biosynthetic process | 4 | 2 | 0.0444 |
| GO:0007034 | vacuolar transport | 4 | 2 | 0.0444 |

| CC | Process Description | Annotated | Observed | pValue |
|---|---|---|---|---|
| GO:0016272 | prefoldin complex | 9 | 4 | 0.0058 |
| GO:0005955 | calcineurin complex | 2 | 2 | 0.0083 |
| GO:0043625 | delta DNA polymerase complex | 2 | 2 | 0.0083 |
| GO:0016021 | integral component of membrane | 126 | 20 | 0.0127 |
| GO:0033176 | proton-transporting V-type ATPase complex | 10 | 4 | 0.0408 |
| GO:0020020 | food vacuole | 15 | 4 | 0.0410 |
| GO:0005657 | replication fork | 8 | 4 | 0.0429 |

| MF | Process Description | Annotated | Observed | pValue |
|---|---|---|---|---|
| GO:0009055 | electron carrier activity | 17 | 6 | 0.0019 |
| GO:0008408 | 3'-5' exonuclease activity | 10 | 4 | 0.0070 |
| GO:0004392 | heme oxygenase (decyclizing) activity | 2 | 2 | 0.0072 |
| GO:0015035 | protein disulfide oxidoreductase activity | 6 | 3 | 0.0100 |
| GO:0008374 | O-acyltransferase activity | 3 | 2 | 0.0203 |
| GO:0005381 | iron ion transmembrane transporter activity | 3 | 2 | 0.0203 |
| GO:0051537 | 2 iron, 2 sulfur cluster binding | 8 | 3 | 0.0245 |

6

| BP | Process Description | Annotated | Observed | pValue |
|---|---|---|---|---|
| GO:0006928 | movement of cell or subcellular component | 34 | 17 | 1.1e-07 |
| GO:0044409 | entry into host | 33 | 16 | 4.6e-07 |
| GO:0006468 | protein phosphorylation | 82 | 24 | 3.7e-05 |
| GO:0035891 | exit from host cell | 8 | 4 | 0.012 |
| GO:0006281 | DNA repair | 38 | 10 | 0.017 |
| GO:0016226 | iron-sulfur cluster assembly | 11 | 4 | 0.041 |
| GO:0006744 | ubiquinone biosynthetic process | 3 | 2 | 0.045 |

| CC | Process Description | Annotated | Observed | pValue |
|---|---|---|---|---|
| GO:0020009 | microneme | 25 | 19 | 5.4e-13 |
| GO:0009986 | cell surface | 32 | 18 | 4.7e-09 |
| GO:0070258 | inner membrane complex | 32 | 13 | 7.3e-05 |
| GO:0005875 | microtubule associated complex | 17 | 8 | 0.0006 |
| GO:0046658 | anchored component of plasma membrane | 3 | 3 | 0.0022 |
| GO:0045177 | apical part of cell | 68 | 31 | 0.0030 |
| GO:0015629 | actin cytoskeleton | 13 | 6 | 0.0035 |
| GO:0005886 | plasma membrane | 28 | 12 | 0.0039 |
| GO:0032299 | ribonuclease H2 complex | 2 | 2 | 0.0168 |
| GO:0005664 | nuclear origin of replication recognition complex | 2 | 2 | 0.0168 |

| MF | Process Description | Annotated | Observed | pValue |
|---|---|---|---|---|
| GO:0003774 | motor activity | 23 | 13 | 7.9e-07 |
| GO:0004672 | protein kinase activity | 81 | 24 | 8.6e-05 |
| GO:0008017 | microtubule binding | 14 | 6 | 0.0055 |
| GO:0043142 | single-stranded DNA-dependent ATPase activity | 2 | 2 | 0.0168 |
| GO:0005509 | calcium ion binding | 32 | 9 | 0.0171 |
| GO:0005515 | protein binding | 478 | 81 | 0.0174 |
| GO:0004252 | serine-type endopeptidase activity | 15 | 5 | 0.0356 |
| GO:0005524 | ATP binding | 335 | 54 | 0.0417 |

| | | | | | |
|---|---|---|---|---|---|
| | GO:0016798 | hydrolase activity, acting on glycosyl bonds | 6 | 4 | 0.0457 |
| | GO:0043138 | 3'-5' DNA helicase activity | 3 | 2 | 0.0462 |
| | GO:0004427 | inorganic diphosphatase activity | 3 | 2 | 0.0462 |

| 7 | BP | Process Description | Annotated | Observed | pValue |
|---|---|---|---|---|---|
| | GO:0006863 | purine nucleobase transport | 1 | 1 | 0.0037 |
| | GO:0015860 | purine nucleoside transmembrane transport | 1 | 1 | 0.0037 |
| | GO:0045454 | cell redox homeostasis | 28 | 2 | 0.0042 |
| | **CC** | **Process Description** | **Annotated** | **Observed** | **pValue** |
| | GO:0009376 | HslUV protease complex | 2 | 1 | 0.0082 |
| | GO:0005829 | cytosol | 78 | 3 | 0.0243 |
| | **MF** | **Process Description** | **Annotated** | **Observed** | **pValue** |
| | GO:0005345 | purine nucleobase transmembrane transporter activity | 1 | 1 | 0.0028 |
| | GO:0015211 | purine nucleoside transmembrane transporter activity | 1 | 1 | 0.0028 |
| | GO:0016668 | oxidoreductase activity, acting on a sulfur group of donors, NAD(P) as acceptor | 4 | 1 | 0.0111 |
| | GO:0015035 | protein disulfide oxidoreductase activity | 6 | 1 | 0.0166 |
| | GO:0004177 | aminopeptidase activity | 7 | 1 | 0.0194 |
| | GO:0050660 | flavin adenine dinucleotide binding | 12 | 1 | 0.0330 |
| | GO:0004298 | threonine-type endopeptidase activity | 15 | 1 | 0.0411 |
| | GO:0009055 | electron carrier activity | 17 | 1 | 0.0465 |

| 8 | BP | Process Description | Annotated | Observed | pValue |
|---|---|---|---|---|---|
| | **CC** | **Process Description** | **Annotated** | **Observed** | **pValue** |
| | GO:0005739 | mitochondrion | 105 | 2 | 0.028 |
| | **MF** | **Process Description** | **Annotated** | **Observed** | **pValue** |
| | GO:0004768 | stearoyl-CoA 9-desaturase activity | 1 | 1 | 0.0023 |
| | GO:0004075 | biotin carboxylase activity | 1 | 1 | 0.0023 |
| | GO:0004129 | cytochrome-c oxidase activity | 6 | 1 | 0.0139 |

| 9 | BP | Process Description | Annotated | Observed | pValue |
|---|---|---|---|---|---|
| | GO:0034645 | cellular macromolecule biosynthetic process | 377 | 12 | 2.4e-07 |
| | GO:0010467 | gene expression | 462 | 12 | 2.7e-06 |
| | **CC** | **Process Description** | **Annotated** | **Observed** | **pValue** |
| | GO:0005840 | ribosome | 132 | 9 | 7.1e-08 |
| | **MF** | **Process Description** | **Annotated** | **Observed** | **pValue** |
| | GO:0003735 | structural constituent of ribosome | 129 | 9 | 4.4e-09 |
| | GO:0003899 | DNA-directed 5'-3' RNA polymerase activity | 31 | 3 | 0.0007 |
| | GO:0019843 | rRNA binding | 10 | 2 | 0.0015 |

| 10 | BP | Process Description | Annotated | Observed | pValue |
|---|---|---|---|---|---|
| | GO:0006259 | DNA metabolic process | 86 | 6 | 5.6e-05 |
| | GO:0009186 | deoxyribonucleoside diphosphate metabolic process | 3 | 1 | 0.020 |

| | | | | |
|---|---|---|---|---|
| GO:0009263 | deoxyribonucleotide biosynthetic process | 5 | 1 | 0.034 |
| GO:0006298 | mismatch repair | 6 | 1 | 0.040 |
| GO:0007067 | mitotic nuclear division | 7 | 1 | 0.047 |

| CC | Process Description | Annotated | Observed | pValue |
|---|---|---|---|---|
| GO:0000808 | origin recognition complex | 4 | 2 | 0.00025 |
| GO:0005658 | alpha DNA polymerase:primase complex | 2 | 1 | 0.01369 |
| GO:0005971 | ribonucleoside-diphosphate reductase complex | 3 | 1 | 0.02048 |

| MF | Process Description | Annotated | Observed | pValue |
|---|---|---|---|---|
| GO:0003677 | DNA binding | 162 | 5 | 0.011 |
| GO:0003917 | DNA topoisomerase type I activity | 2 | 1 | 0.013 |
| GO:0004748 | ribonucleoside-diphosphate reductase activity, thioredoxin disulfide as acceptor | 3 | 1 | 0.019 |
| GO:0003918 | DNA topoisomerase type II (ATP-hydrolyzing) activity | 4 | 1 | 0.026 |
| GO:0030983 | mismatched DNA binding | 6 | 1 | 0.038 |

11

| BP | Process Description | Annotated | Observed | pValue |
|---|---|---|---|---|
| GO:0006606 | protein import into nucleus | 3 | 2 | 0.0062 |
| GO:0006633 | fatty acid biosynthetic process | 9 | 3 | 0.0066 |
| GO:0006457 | protein folding | 53 | 7 | 0.0098 |
| GO:0006270 | DNA replication initiation | 4 | 2 | 0.0121 |
| GO:0006950 | response to stress | 48 | 7 | 0.0222 |
| GO:0022904 | respiratory electron transport chain | 5 | 2 | 0.0460 |
| GO:0000289 | nuclear-transcribed mRNA poly(A) tail shortening | 1 | 1 | 0.0465 |
| GO:0006433 | prolyl-tRNA aminoacylation | 1 | 1 | 0.0465 |
| GO:0019510 | S-adenosylhomocysteine catabolic process | 1 | 1 | 0.0465 |
| GO:0000077 | DNA damage checkpoint | 1 | 1 | 0.0465 |
| GO:0042787 | protein ubiquitination involved in ubiquitin-dependent protein catabolic process | 1 | 1 | 0.0465 |
| GO:0051382 | kinetochore assembly | 1 | 1 | 0.0465 |
| GO:0034477 | U6 snRNA 3'-end processing | 1 | 1 | 0.0465 |
| GO:0030433 | ubiquitin-dependent ERAD pathway | 1 | 1 | 0.0465 |
| GO:0006561 | proline biosynthetic process | 1 | 1 | 0.0465 |
| GO:0010564 | regulation of cell cycle process | 1 | 1 | 0.0465 |

| CC | Process Description | Annotated | Observed | pValue |
|---|---|---|---|---|
| GO:0005832 | chaperonin-containing T-complex | 8 | 4 | 0.00024 |
| GO:0000408 | EKC/KEOPS complex | 3 | 2 | 0.00590 |
| GO:0042555 | MCM complex | 4 | 2 | 0.01145 |
| GO:0030896 | checkpoint clamp complex | 1 | 1 | 0.04533 |
| GO:0030014 | CCR4-NOT complex | 1 | 1 | 0.04533 |

| MF | Process Description | Annotated | Observed | pValue |
|---|---|---|---|---|
| GO:0051082 | unfolded protein binding | 28 | 6 | 0.0015 |
| GO:0000175 | 3'-5'-exoribonuclease activity | 3 | 2 | 0.0063 |
| GO:0004222 | metalloendopeptidase activity | 10 | 3 | 0.0094 |
| GO:0004045 | aminoacyl-tRNA hydrolase activity | 4 | 2 | 0.0123 |
| GO:0003678 | DNA helicase activity | 17 | 3 | 0.0422 |

| | | | | |
|---|---|---|---|---|
| GO:0004325 | ferrochelatase activity | 1 | 1 | 0.0469 |
| GO:0035064 | methylated histone binding | 1 | 1 | 0.0469 |
| GO:0004013 | adenosylhomocysteinase activity | 1 | 1 | 0.0469 |
| GO:0004425 | indole-3-glycerol-phosphate synthase activity | 1 | 1 | 0.0469 |
| GO:0004735 | pyrroline-5-carboxylate reductase activity | 1 | 1 | 0.0469 |
| GO:0004827 | proline-tRNA ligase activity | 1 | 1 | 0.0469 |
| GO:0016992 | lipoate synthase activity | 1 | 1 | 0.0469 |
| GO:0046429 | 4-hydroxy-3-methylbut-2-en-1-yl diphosphate synthase activity | 1 | 1 | 0.0469 |
| GO:0008897 | holo-[acyl-carrier-protein] synthase activity | 1 | 1 | 0.0469 |
| GO:0004148 | dihydrolipoyl dehydrogenase activity | 1 | 1 | 0.0469 |

12

| BP | Process Description | Annotated | Observed | pValue |
|---|---|---|---|---|
| GO:0006511 | ubiquitin-dependent protein catabolic process | 38 | 17 | 1.8e-06 |
| GO:0051603 | proteolysis involved in cellular protein catabolic process | 45 | 22 | 0.0013 |
| GO:0006122 | mitochondrial electron transport, ubiquinol to cytochrome c | 4 | 3 | 0.0087 |
| GO:0045184 | establishment of protein localization | 93 | 22 | 0.0127 |
| GO:0006913 | nucleocytoplasmic transport | 20 | 7 | 0.0147 |
| GO:0032446 | protein modification by small protein conjugation | 12 | 5 | 0.0150 |
| GO:0007264 | small GTPase mediated signal transduction | 26 | 8 | 0.0168 |
| GO:0006875 | cellular metal ion homeostasis | 3 | 3 | 0.0179 |
| GO:0015991 | ATP hydrolysis coupled proton transport | 14 | 5 | 0.0302 |
| GO:0006099 | tricarboxylic acid cycle | 10 | 4 | 0.0347 |
| GO:0070682 | proteasome regulatory particle assembly | 6 | 3 | 0.0353 |
| GO:0034613 | cellular protein localization | 82 | 18 | 0.0378 |
| GO:0030163 | protein catabolic process | 52 | 25 | 0.0454 |

| CC | Process Description | Annotated | Observed | pValue |
|---|---|---|---|---|
| GO:0000502 | proteasome complex | 31 | 20 | 9e-12 |
| GO:0012505 | endomembrane system | 94 | 23 | 0.00068 |
| GO:0005750 | mitochondrial respiratory chain complex III | 4 | 3 | 0.00710 |
| GO:0045259 | proton-transporting ATP synthase complex | 6 | 3 | 0.02927 |
| GO:0033178 | proton-transporting two-sector ATPase complex, catalytic domain | 11 | 4 | 0.03905 |
| GO:0005853 | eukaryotic translation elongation factor 1 complex | 3 | 2 | 0.04325 |
| GO:0030532 | small nuclear ribonucleoprotein complex | 12 | 4 | 0.04631 |

| MF | Process Description | Annotated | Observed | pValue |
|---|---|---|---|---|
| GO:0004298 | threonine-type endopeptidase activity | 15 | 11 | 1.3e-07 |
| GO:0008565 | protein transporter activity | 15 | 7 | 0.0015 |
| GO:0048037 | cofactor binding | 41 | 11 | 0.0126 |
| GO:0005092 | GDP-dissociation inhibitor activity | 2 | 2 | 0.0168 |
| GO:0015078 | hydrogen ion transmembrane transporter activity | 29 | 8 | 0.0270 |
| GO:0004843 | thiol-dependent ubiquitin-specific protease activity | 6 | 3 | 0.0322 |
| GO:0000287 | magnesium ion binding | 15 | 5 | 0.0356 |
| GO:0043168 | anion binding | 455 | 71 | 0.0452 |
| GO:0008242 | omega peptidase activity | 3 | 2 | 0.0462 |

13

| BP | Process Description | Annotated | Observed | pValue |
|---|---|---|---|---|
| GO:0006465 | signal peptide processing | 5 | 4 | 0.0007 |
| GO:0006825 | copper ion transport | 5 | 3 | 0.0117 |
| GO:0045039 | protein import into mitochondrial inner membrane | 2 | 2 | 0.0125 |
| GO:0006415 | translational termination | 7 | 3 | 0.0346 |

| CC | Process Description | Annotated | Observed | pValue |
|---|---|---|---|---|
| GO:0005787 | signal peptidase complex | 3 | 3 | 0.0012 |
| GO:0020011 | apicoplast | 94 | 20 | 0.0015 |
| GO:0005758 | mitochondrial intermembrane space | 4 | 3 | 0.0045 |
| GO:0005739 | mitochondrion | 105 | 22 | 0.0049 |
| GO:0016272 | prefoldin complex | 9 | 4 | 0.0106 |
| GO:0005672 | transcription factor TFIIA complex | 2 | 2 | 0.0116 |
| GO:0000932 | P-body | 3 | 2 | 0.0322 |
| GO:0005732 | small nucleolar ribonucleoprotein complex | 14 | 4 | 0.0550 |

| MF | Process Description | Annotated | Observed | pValue |
|---|---|---|---|---|
| GO:0043022 | ribosome binding | 8 | 4 | 0.0046 |
| GO:0005375 | copper ion transmembrane transporter activity | 2 | 2 | 0.0096 |
| GO:0003747 | translation release factor activity | 6 | 3 | 0.0148 |
| GO:0004312 | fatty acid synthase activity | 3 | 2 | 0.0268 |

14

| BP | Process Description | Annotated | Observed | pValue |
|---|---|---|---|---|
| GO:0006497 | protein lipidation | 21 | 8 | 0.0011 |
| GO:0006468 | protein phosphorylation | 82 | 19 | 0.0019 |
| GO:0007017 | microtubule-based process | 35 | 11 | 0.0021 |
| GO:0006928 | movement of cell or subcellular component | 34 | 9 | 0.0093 |
| GO:0016197 | endosomal transport | 2 | 2 | 0.0123 |
| GO:0015693 | magnesium ion transport | 2 | 2 | 0.0123 |
| GO:0051301 | cell division | 7 | 3 | 0.0336 |
| GO:0006388 | tRNA splicing, via endonucleolytic cleavage and ligation | 3 | 2 | 0.0341 |
| GO:0019673 | GDP-mannose metabolic process | 3 | 2 | 0.0341 |
| GO:0006836 | neurotransmitter transport | 3 | 2 | 0.0341 |
| GO:0035891 | exit from host cell | 8 | 3 | 0.0495 |

| CC | Process Description | Annotated | Observed | pValue |
|---|---|---|---|---|
| GO:0020039 | pellicle | 33 | 18 | 3.3e-09 |
| GO:0020008 | rhoptry | 26 | 14 | 2.6e-07 |
| GO:0044430 | cytoskeletal part | 50 | 19 | 0.00020 |
| GO:0015630 | microtubule cytoskeleton | 38 | 15 | 0.00039 |
| GO:0015629 | actin cytoskeleton | 13 | 6 | 0.00252 |
| GO:0044310 | osmiophilic body | 7 | 4 | 0.00562 |
| GO:0044312 | crystalloid | 7 | 4 | 0.00562 |
| GO:0020003 | symbiont-containing vacuole | 44 | 10 | 0.03450 |
| GO:0044164 | host cell cytosol | 16 | 5 | 0.03651 |

| MF | Process Description | Annotated | Observed | pValue |
|---|---|---|---|---|
| GO:0008017 | microtubule binding | 14 | 6 | 0.0030 |
| GO:0004672 | protein kinase activity | 81 | 18 | 0.0039 |

| | | | | |
|---|---|---|---|---|
| GO:0016409 | palmitoyltransferase activity | 11 | 5 | 0.0051 |
| GO:0008234 | cysteine-type peptidase activity | 30 | 8 | 0.0071 |
| GO:0005509 | calcium ion binding | 32 | 9 | 0.0081 |
| GO:0003774 | motor activity | 23 | 7 | 0.0121 |
| GO:0070569 | uridylyltransferase activity | 2 | 2 | 0.0133 |
| GO:0071949 | FAD binding | 2 | 2 | 0.0133 |
| GO:0015095 | magnesium ion transmembrane transporter activity | 2 | 2 | 0.0133 |
| GO:0000213 | tRNA-intron endonuclease activity | 2 | 2 | 0.0133 |
| GO:0030246 | carbohydrate binding | 6 | 3 | 0.0234 |
| GO:0005328 | neurotransmitter:sodium symporter activity | 3 | 2 | 0.0369 |
| GO:0046789 | host cell surface receptor binding | 3 | 2 | 0.0369 |
| GO:0016538 | cyclin-dependent protein serine/threonine kinase regulator activity | 3 | 2 | 0.0369 |
| GO:0046812 | host cell surface binding | 7 | 3 | 0.3863 |
