## Supplementary material for "Transcriptome analysis of *Plasmodium berghei* during exo-erythrocytic development": Table S4

Table S4: Gene IDs (PBANKA)

**Community 1**

_0001301
_0001401
_0001601
_0007501
_0007801
_0008201
_0100061
_0100900
_0105300
_0107700
_0108700
_0110000
_0111200
_0200400
_0202900
_0204100
_0205000
_0205200
_0213000
_0213100
_0213400
_0213500
_0301500
_0302300
_0304300
_0305300
_0307000
_0308500
_0308600
_0309100
_0311200
_0311800
_0312400
_0314500
_0315100
_0315400
_0315600
_0316100
_0401100
_0401900
_0403000
_0404000
_0407700
_0410600
_0412200
_0412300
_0413600
_0414600
_0414700
_0415600
_0416200
_0417500
_0502700
_0502800
_0502900
_0504900
_0505700
_0506200
_0506300
_0508100
_0510900
_0511200
_0512400
_0512500
_0513200
_0515900
_0516900
_0517400
_0517500
_0518700
_0521261
_0521300
_0521400
_0522800
_0523100
_0600011
_0600031
_0600500
_0601700
_0602500
_0603200
_0604500
_0604800
_0605200
_0606600
_0609100
_0609700
_0609900
_0610900
_0613800
_0613900
_0614500
_0615100
_0617100
_0617200
_0617600
_0617700
_0618800
_0619800
_0619900
_0620000
_0620100
_0620900
_0622600
_0622800
_0623000
_0623751
_0700031
_0700041
_0700761
_0703700
_0703900
_0704500
_0704600
_0705000
_0705700
_0705800
_0707400
_0707800
_0709100
_0709500
_0710500
_0711800
_0711900
_0712000
_0712400
_0717800
_0720300
_0720400
_0720800
_0721900
_0722961
_0801200
_0801600
_0804000
_0804800
_0805100
_0805300
_0805400
_0806100
_0807600
_0808700
_0811100
_0816400
_0817400
_0817700
_0819300
_0819800
_0819900
_0822100
_0823900
_0824300
_0824800
_0825600
_0826900
_0831300
_0831800
_0831900
_0836941
_0836981
_0903000
_0904700
_0905000
_0905600
_0906400
_0906600
_0908300
_0909600
_0912900
_0913800
_0914200
_0915300
_0916000
_0917700
_0918000
_0918100
_0918800
_0919300
_0920400
_0922100
_0923500
_0924400
_0924800
_0927800
_0928800
_0929600
_0930300
_0931300
_0934500
_0936300
_0936600
_0936700
_0937200
_0937800
_0938900
_0939100
_0943500
_0943541
_0943600
_1002700
_1003200
_1003700
_1004900
_1005100
_1005700
_1006700
_1006800
_1006900
_1007000
_1007400
_1007500
_1010200
_1010300
_1010500
_1012400
_1015200
_1017900
_1018000
_1018600
_1019330
_1019400
_1019500
_1019600
_1019700
_1020400
_1021000
_1021100
_1022000
_1023800
_1024300
_1025200
_1026900
_1027500
_1028400
_1028500
_1031000
_1033900
_1035100
_1035900
_1036300
_1036400
_1037200
_1037300
_1037400
_1037600
_1039400
_1100700
_1103400
_1104600
_1105700
_1105800
_1106700
_1107400
_1109000
_1111000
_1114600
_1116400
_1117100
_1117500
_1119000
_1120300
_1120700
_1122500
_1122700
_1123500
_1124100
_1125600
_1126000
_1126400
_1126900
_1127900
_1128400
_1129700
_1130900
_1131900
_1134000
_1134400
_1134800
_1135100
_1135300
_1135700
_1136900
_1139900
_1140200
_1140600
_1141100
_1141700
_1142100
_1142800
_1143300
_1143500
_1143600
_1200011
_1200031
_1200041
_1200051
_1200600
_1200800
_1201700
_1201900
_1202400
_1206300
_1207000
_1207100
_1211500
_1212900
_1213000
_1215800
_1221200
_1221300
_1221600
_1222700
_1224200
_1227100
_1227800
_1227900
_1228700
_1229700
_1230500
_1231000
_1231700
_1231800
_1232500
_1234500
_1237000
_1238200
_1239700
_1239800
_1240300
_1242800
_1243200
_1243400
_1245000
_1245800
_1245861
_1246941
_1246961
_1246981
_1300011
_1300041
_1300051
_1302400
_1302500
_1309000
_1309200
_1309500
_1309700
_1310800
_1310900
_1311000
_1311800
_1313000
_1313400
_1314400
_1315600
_1317600
_1317900
_1318100
_1321400
_1322500
_1322700
_1324400
_1324900
_1327000
_1327021
_1327700
_1328300
_1328800
_1329300
_1329800
_1330100
_1331900
_1332500
_1333500
_1334200
_1337000
_1338600
_1339100
_1339200
_1343750
_1344600
_1345800
_1346700
_1347500
_1348300
_1350500
_1352700
_1354300
_1354380
_1354500
_1355800
_1356000
_1356200
_1359200
_1359400
_1362600
_1363000
_1363100
_1364100
_1364200
_1364900
_1401100
_1401200
_1401900
_1405200
_1405300
_1405400
_1406700
_1406800
_1407400
_1413900
_1416300
_1416400
_1417800
_1420900
_1423200
_1424300
_1426200
_1426900
_1428700
_1428900
_1432800
_1433100
_1433300
_1433400
_1434100
_1439200
_1440100
_1440200
_1440300
_1440400
_1441500
_1442500
_1443100
_1443800
_1445600
_1447100
_1451800
_1452200
_1452300
_1452500
_1452800
_1455200
_1455300
_1456100
_1456900
_1457400
_1457550
_1459000
_1461100
_1462000
_1462200
_1462800
_1462900
_1463500
_1464500
_1464800
_1465000
_1465100
_1465800
_1466221
_1466241
_API00011
_API00051
_API00095
_MIT02700
_MIT03500
_MIT03600

Table S4: Gene IDs (PBANKA)

| | | | | | | | | |
|---|---|---|---|---|---|---|---|---|
| **Community 2** | _0316300 | _0600100 | _0811300 | _0943800 | _1107900 | _1216400 | _1326600 | _1417300 |
| _0000101 | _0316500 | _0600200 | _0812500 | _0944061 | _1109800 | _1216500 | _1327251 | _1417600 |
| _0000301 | _0316800 | _0602800 | _0813800 | _0944081 | _1110500 | _1216900 | _1327300 | _1418300 |
| _0000600 | _0316821 | _0604200 | _0814000 | _0944101 | _1112400 | _1217500 | _1327800 | _1420500 |
| _0000701 | _0316841 | _0605700 | _0815300 | _1000041 | _1116700 | _1218000 | _1327900 | _1420800 |
| _0001001 | _0316900 | _0606100 | _0816600 | _1000051 | _1116800 | _1218800 | _1328700 | _1421100 |
| _0001101 | _0316921 | _0608300 | _0817600 | _1000091 | _1120200 | _1219000 | _1329600 | _1423300 |
| _0007601 | _0317021 | _0610100 | _0817900 | _1000200 | _1120800 | _1219500 | _1332000 | _1423500 |
| _0008001 | _0317061 | _0610500 | _0818600 | _1000300 | _1121100 | _1220100 | _1332100 | _1423600 |
| _0008101 | _0317121 | _0613700 | _0818700 | _1000500 | _1121700 | _1220200 | _1332600 | _1424600 |
| _0100021 | _0317161 | _0615400 | _0819100 | _1000600 | _1121800 | _1221000 | _1334300 | _1424800 |
| _0100200 | _0400100 | _0618200 | _0819700 | _1003100 | _1122000 | _1223300 | _1334600 | _1424900 |
| _0100300 | _0401000 | _0619100 | _0821700 | _1003300 | _1125650 | _1223500 | _1335500 | _1425000 |
| _0100500 | _0401700 | _0621400 | _0822700 | _1005600 | _1126500 | _1226000 | _1337100 | _1426000 |
| _0100600 | _0402800 | _0622200 | _0822800 | _1007100 | _1127000 | _1226400 | _1338300 | _1426100 |
| _0100700 | _0405300 | _0623100 | _0823400 | _1008500 | _1127400 | _1229000 | _1340300 | _1429000 |
| _0102200 | _0405400 | _0623150 | _0823600 | _1009300 | _1127700 | _1229200 | _1341500 | _1429700 |
| _0103300 | _0405500 | _0623200 | _0826700 | _1010700 | _1128200 | _1231100 | _1342400 | _1431700 |
| _0104800 | _0407100 | _0623300 | _0827500 | _1010800 | _1129300 | _1231600 | _1342800 | _1432500 |
| _0104900 | _0407600 | _0623400 | _0829700 | _1011800 | _1131200 | _1232100 | _1343000 | _1434500 |
| _0105200 | _0409100 | _0623500 | _0836000 | _1011900 | _1132000 | _1234200 | _1343900 | _1435700 |
| _0105600 | _0409500 | _0623651 | _0836200 | _1013100 | _1133300 | _1234300 | _1344500 | _1437100 |
| _0108200 | _0411100 | _0700051 | _0836500 | _1013500 | _1133400 | _1234800 | _1346100 | _1438100 |
| _0109200 | _0411200 | _0700061 | _0836600 | _1016500 | _1134500 | _1237500 | _1347600 | _1438800 |
| _0110400 | _0412100 | _0700071 | _0836800 | _1016800 | _1135000 | _1237800 | _1347700 | _1439800 |
| _0112600 | _0412800 | _0700081 | _0837001 | _1018700 | _1135200 | _1238600 | _1348900 | _1440600 |
| _0112641 | _0418000 | _0700200 | _0837101 | _1019520 | _1136300 | _1238800 | _1349200 | _1441700 |
| _0112661 | _0418500 | _0700300 | _0837121 | _1022200 | _1137100 | _1239100 | _1351900 | _1442400 |
| _0112701 | _0500600 | _0700400 | _0837141 | _1023200 | _1137200 | _1242200 | _1352000 | _1445400 |
| _0112721 | _0500700 | _0700500 | _0905700 | _1023500 | _1138400 | _1242300 | _1353700 | _1445500 |
| _0201200 | _0500741 | _0700521 | _0906500 | _1024100 | _1141300 | _1242500 | _1354000 | _1446200 |
| _0201250 | _0500971 | _0700600 | _0910000 | _1025500 | _1142300 | _1245100 | _1354400 | _1446800 |
| _0201300 | _0501000 | _0700700 | _0911400 | _1029900 | _1142700 | _1245900 | _1355100 | _1448100 |
| _0201500 | _0501051 | _0700721 | _0912000 | _1030000 | _1145100 | _1246100 | _1356700 | _1448500 |
| _0201600 | _0503600 | _0700741 | _0912200 | _1030600 | _1145400 | _1246141 | _1356900 | _1449400 |
| _0203200 | _0504200 | _0700900 | _0912300 | _1030700 | _1145500 | _1246161 | _1357200 | _1450100 |
| _0203300 | _0505900 | _0701000 | _0913400 | _1030800 | _1145700 | _1246800 | _1357800 | _1452400 |
| _0203800 | _0506400 | _0701100 | _0914700 | _1030900 | _1145800 | _1300021 | _1358000 | _1454500 |
| _0203900 | _0509000 | _0701600 | _0915800 | _1031900 | _1145900 | _1300031 | _1360100 | _1454700 |
| _0204900 | _0510200 | _0702900 | _0919100 | _1032200 | _1146000 | _1300081 | _1361100 | _1457600 |
| _0205100 | _0511900 | _0703100 | _0919200 | _1032300 | _1146241 | _1300091 | _1363300 | _1458900 |
| _0205500 | _0514000 | _0703600 | _0921400 | _1032900 | _1146700 | _1300100 | _1364500 | _1459700 |
| _0206600 | _0514100 | _0705100 | _0921500 | _1033500 | _1146721 | _1300600 | _1365500 | _1460800 |
| _0206900 | _0515800 | _0707300 | _0921800 | _1033600 | _1146741 | _1301600 | _1365600 | _1463800 |
| _0208700 | _0517000 | _0709400 | _0923400 | _1034300 | _1200021 | _1301700 | _1365680 | _1464000 |
| _0208800 | _0517100 | _0710800 | _0924900 | _1034400 | _1200071 | _1303200 | _1400011 | _1464600 |
| _0210200 | _0517200 | _0712800 | _0925200 | _1034900 | _1200081 | _1305000 | _1400031 | _1464700 |
| _0212000 | _0517300 | _0714700 | _0925300 | _1035600 | _1202200 | _1305100 | _1400041 | _1465500 |
| _0212100 | _0518800 | _0714800 | _0926700 | _1036000 | _1203300 | _1306200 | _1400051 | _1465600 |
| _0214600 | _0519900 | _0715200 | _0927200 | _1040541 | _1203400 | _1307200 | _1400700 | _1465921 |
| _0214700 | _0520200 | _0717300 | _0928100 | _1040581 | _1204400 | _1311600 | _1401300 | _1466000 |
| _0214800 | _0520600 | _0718200 | _0928200 | _1100351 | _1205000 | _1314100 | _1403800 | _1466121 |
| _0214900 | _0520900 | _0722801 | _0929800 | _1100400 | _1206100 | _1315200 | _1405500 | _MIT01200 |
| _0216021 | _0524100 | _0800300 | _0930700 | _1100461 | _1207200 | _1315800 | _1405600 | |
| _0216041 | _0524200 | _0800400 | _0931200 | _1100781 | _1207300 | _1316700 | _1405700 | |
| _0216741 | _0524300 | _0801400 | _0931400 | _1100860 | _1207500 | _1317400 | _1407600 | |
| _0216761 | _0524400 | _0803400 | _0932400 | _1100900 | _1207700 | _1317700 | _1407650 | |
| _0300200 | _0524500 | _0803600 | _0933400 | _1101100 | _1207900 | _1318400 | _1407800 | |
| _0300600 | _0524700 | _0804100 | _0937400 | _1101200 | _1208600 | _1320500 | _1408000 | |
| _0302000 | _0524800 | _0804700 | _0937700 | _1101300 | _1208700 | _1320600 | _1408100 | |
| _0307800 | _0524821 | _0804900 | _0938600 | _1102000 | _1209000 | _1321000 | _1408900 | |
| _0310800 | _0600021 | _0806000 | _0939200 | _1102200 | _1210200 | _1321100 | _1409300 | |
| _0311600 | _0600041 | _0807000 | _0939600 | _1103000 | _1210800 | _1323000 | _1409400 | |
| _0314300 | _0600051 | _0807100 | _0940700 | _1103500 | _1211000 | _1323700 | _1410300 | |
| _0314600 | _0600061 | _0809800 | _0941300 | _1103600 | _1211300 | _1324000 | _1412100 | |
| _0315200 | _0600081 | _0810500 | _0941500 | _1104300 | _1211900 | _1325300 | _1414400 | |
| _0315300 | _0600086 | _0810600 | _0942500 | _1104900 | _1215600 | _1325600 | _1415100 | |
| _0316200 | _0600091 | _0811000 | _0942900 | _1106400 | _1216100 | _1326500 | _1416200 | |

Table S4: Gene IDs (PBANKA)

| | | |
|---|---|---|
| **Community 3** | _0900600 | _1403250 |
| _0000801 | _0902800 | _1428100 |
| _0000901 | _0937750 | _1428980 |
| _0100081 | _0937840 | _1437250 |
| _0112621 | _0937860 | _1446300 |
| _0200670 | _0938050 | _1455800 |
| _0200751 | _0943700 | _1465200 |
| _0201000 | _0944000 | _1465300 |
| _0201100 | _1000031 | _1465400 |
| _0201360 | _1000061 | _1465841 |
| _0201400 | _1000071 | _1465861 |
| _0201450 | _1000081 | _1465881 |
| _0215700 | _1000400 | _1466100 |
| _0215900 | _1000451 | _1466141 |
| _0216061 | _1019370 | _1466161 |
| _0216081 | _1019380 | _1466181 |
| _0216701 | _1019540 | _MIT00200 |
| _0216721 | _1019560 | _MIT00300 |
| _0216801 | _1019580 | _MIT01600 |
| _0300300 | _1031300 | _MIT02100 |
| _0300400 | _1039450 | _MIT02300 |
| _0300500 | _1040200 | _MIT02400 |
| _0301400 | _1040300 | _MIT02900 |
| _0315420 | _1040400 | _MIT03200 |
| _0315430 | _1040500 | _MIT03700 |
| _0316400 | _1040561 | |
| _0316861 | _1040621 | |
| _0316941 | _1100031 | |
| _0317001 | _1100750 | |
| _0317041 | _1100850 | |
| _0317081 | _1136100 | |
| _0317101 | _1146100 | |
| _0405310 | _1146200 | |
| _0406650 | _1146781 | |
| _0407550 | _1200091 | |
| _0500200 | _1200300 | |
| _0500721 | _1200400 | |
| _0500781 | _1200500 | |
| _0500950 | _1206200 | |
| _0513500 | _1226470 | |
| _0516100 | _1229600 | |
| _0600300 | _1245821 | |
| _0600400 | _1245920 | |
| _0612841 | _1246121 | |
| _0623600 | _1246200 | |
| _0700541 | _1246300 | |
| _0700561 | _1246400 | |
| _0703910 | _1246700 | |
| _0722100 | _1246900 | |
| _0722200 | _1300071 | |
| _0722700 | _1300200 | |
| _0722900 | _1328500 | |
| _0722941 | _1339400 | |
| _0800061 | _1345850 | |
| _0800100 | _1346750 | |
| _0800200 | _1354550 | |
| _0805750 | _1364250 | |
| _0823000 | _1364600 | |
| _0836400 | _1365550 | |
| _0836550 | _1365650 | |
| _0836850 | _1365700 | |
| _0836900 | _1400021 | |
| _0836921 | _1400061 | |
| _0836961 | _1400071 | |
| _0837081 | _1400081 | |
| _0837161 | _1400100 | |
| _0837181 | _1400200 | |
| _0837201 | _1400300 | |

Table S4: Gene IDs (PBANKA)

**Community 4**

_0100800
_0200600
_0211200
_0304100
_0317141
_0408300
_0509200
_0514300
_0706700
_0711700
_0712900
_0713600
_0810000
_0831400
_1005000
_1030200
_1034000
_1036100
_1131800
_1202600
_1203800
_1208800
_1221700
_1226300
_1309300
_1313500
_1316100
_1333400
_1361200
_1408800
_1410100
_1413500
_1433200
_1433900
_1439300
_1445200

Table S4: Gene IDs (PBANKA)

| | | | | | |
|---|---|---|---|---|---|
| **Community 5** | _0507600 | _0908900 | _1112100 | _1303700 | _1401600 |
| _0101300 | _0508400 | _0909200 | _1113100 | _1304100 | _1402500 |
| _0101700 | _0508500 | _0909900 | _1114700 | _1304200 | _1402600 |
| _0102000 | _0508700 | _0910200 | _1114900 | _1304600 | _1403500 |
| _0103500 | _0511500 | _0914000 | _1115800 | _1305600 | _1404700 |
| _0103800 | _0511700 | _0915500 | _1115900 | _1305700 | _1405100 |
| _0104000 | _0516300 | _0917200 | _1117700 | _1306500 | _1409500 |
| _0105000 | _0518600 | _0917400 | _1118000 | _1307000 | _1410000 |
| _0106000 | _0601000 | _0918300 | _1118100 | _1307600 | _1410700 |
| _0106800 | _0602300 | _0920000 | _1118300 | _1307800 | _1411100 |
| _0107400 | _0608100 | _0920100 | _1123300 | _1308400 | _1412400 |
| _0108800 | _0608200 | _0920300 | _1125100 | _1308500 | _1412900 |
| _0203000 | _0612500 | _0924300 | _1125500 | _1308700 | _1413000 |
| _0203400 | _0614400 | _0925900 | _1128100 | _1310600 | _1413100 |
| _0203500 | _0614700 | _0928400 | _1128600 | _1312500 | _1413400 |
| _0203650 | _0614900 | _0931800 | _1128700 | _1314200 | _1414200 |
| _0204700 | _0615000 | _0932300 | _1129400 | _1314700 | _1416600 |
| _0206200 | _0615800 | _0933300 | _1130200 | _1315400 | _1416700 |
| _0207500 | _0616300 | _0934200 | _1130300 | _1316600 | _1418600 |
| _0208100 | _0617400 | _0935200 | _1130700 | _1319100 | _1420300 |
| _0208200 | _0621100 | _0935400 | _1131000 | _1320200 | _1422200 |
| _0208300 | _0621500 | _0935600 | _1131700 | _1320900 | _1422700 |
| _0209200 | _0622961 | _0935800 | _1132500 | _1322000 | _1425300 |
| _0211100 | _0701700 | _0936500 | _1132600 | _1322400 | _1425350 |
| _0212500 | _0701800 | _0938700 | _1132700 | _1322600 | _1426400 |
| _0213700 | _0705200 | _0939400 | _1134600 | _1323200 | _1427000 |
| _0214400 | _0705300 | _0939500 | _1138500 | _1327200 | _1428000 |
| _0302600 | _0705900 | _0942600 | _1138700 | _1328000 | _1428800 |
| _0302700 | _0710700 | _1000700 | _1139000 | _1329500 | _1429900 |
| _0306000 | _0711200 | _1001100 | _1144200 | _1331100 | _1430000 |
| _0306500 | _0713000 | _1005200 | _1144600 | _1331800 | _1430400 |
| _0307900 | _0714400 | _1007700 | _1144900 | _1333200 | _1430500 |
| _0308300 | _0716700 | _1008900 | _1145600 | _1333600 | _1431600 |
| _0308700 | _0717100 | _1009700 | _1201000 | _1335300 | _1432100 |
| _0309600 | _0719300 | _1009800 | _1201100 | _1335600 | _1432600 |
| _0310100 | _0719600 | _1010000 | _1201300 | _1336300 | _1434000 |
| _0313900 | _0803200 | _1010100 | _1202700 | _1337600 | _1434200 |
| _0315500 | _0806500 | _1012200 | _1208100 | _1340600 | _1435100 |
| _0316000 | _0806600 | _1012300 | _1208400 | _1340800 | _1438200 |
| _0403100 | _0806700 | _1012600 | _1210000 | _1340900 | _1438500 |
| _0403900 | _0807300 | _1015100 | _1210300 | _1341300 | _1439400 |
| _0404700 | _0807400 | _1015500 | _1212000 | _1342600 | _1441400 |
| _0406300 | _0808100 | _1015600 | _1212700 | _1346500 | _1442000 |
| _0407000 | _0808200 | _1017200 | _1213100 | _1350300 | _1445700 |
| _0407200 | _0808800 | _1020200 | _1213800 | _1350400 | _1449100 |
| _0409600 | _0811700 | _1020600 | _1214600 | _1351800 | _1449200 |
| _0411300 | _0812000 | _1021600 | _1214700 | _1352200 | _1449500 |
| _0411600 | _0813100 | _1021800 | _1216800 | _1352900 | _1450900 |
| _0412400 | _0815000 | _1022700 | _1220000 | _1353000 | _1452000 |
| _0412500 | _0820300 | _1022800 | _1220500 | _1353200 | _1452600 |
| _0412600 | _0820600 | _1023700 | _1220800 | _1353300 | _1453600 |
| _0412700 | _0820900 | _1024700 | _1222200 | _1355000 | _1455000 |
| _0413700 | _0821900 | _1027200 | _1222400 | _1355200 | _1455500 |
| _0414000 | _0829000 | _1028300 | _1222900 | _1355300 | _1455600 |
| _0415200 | _0829100 | _1028800 | _1223400 | _1355400 | _1456000 |
| _0415300 | _0832100 | _1029200 | _1227300 | _1355500 | _1457300 |
| _0416300 | _0833000 | _1029800 | _1227400 | _1355600 | _1457800 |
| _0416400 | _0834300 | _1032600 | _1232800 | _1356300 | _1458100 |
| _0416600 | _0834400 | _1033000 | _1232900 | _1356500 | _1458500 |
| _0417400 | _0834500 | _1033100 | _1234000 | _1357600 | _1458600 |
| _0501200 | _0901100 | _1034700 | _1235200 | _1358200 | _1459900 |
| _0501300 | _0901300 | _1034800 | _1235800 | _1358800 | _1460400 |
| _0501900 | _0901700 | _1036800 | _1236300 | _1360200 | _1461200 |
| _0502300 | _0902100 | _1037000 | _1239600 | _1361900 | _1465051 |
| _0503700 | _0902500 | _1038000 | _1241700 | _1362200 | |
| _0504000 | _0904500 | _1108600 | _1300800 | _1400800 | |
| _0506500 | _0907000 | _1108900 | _1301100 | _1401000 | |
| _0506700 | _0907800 | _1109200 | _1301200 | _1401500 | |

Table S4: Gene IDs (PBANKA)

| | | | | | | | | |
|---|---|---|---|---|---|---|---|---|
| **Community 6** | _0312600 | _0600096 | _0713200 | _0830900 | _0928900 | _1023300 | _1122800 | _1225700 |
| _0100041 | _0313000 | _0600700 | _0714300 | _0831100 | _0929100 | _1024400 | _1123400 | _1225900 |
| _0101400 | _0313100 | _0600900 | _0714500 | _0831200 | _0929300 | _1025300 | _1123700 | _1226100 |
| _0101800 | _0313400 | _0602000 | _0716300 | _0831700 | _0929400 | _1025700 | _1123800 | _1227000 |
| _0103900 | _0314000 | _0602400 | _0716400 | _0832800 | _0929700 | _1026100 | _1124000 | _1227200 |
| _0104500 | _0314100 | _0603700 | _0716600 | _0833300 | _0930100 | _1026500 | _1125200 | _1228300 |
| _0106100 | _0314200 | _0604600 | _0716800 | _0833500 | _0930200 | _1026700 | _1126800 | _1229900 |
| _0106300 | _0315700 | _0604900 | _0716900 | _0834100 | _0930800 | _1026800 | _1128900 | _1230200 |
| _0107800 | _0401400 | _0605000 | _0717400 | _0834800 | _0931000 | _1027600 | _1129000 | _1232400 |
| _0107900 | _0403200 | _0605100 | _0718900 | _0834900 | _0931100 | _1027700 | _1129200 | _1232700 |
| _0108000 | _0403300 | _0605500 | _0719200 | _0835400 | _0931900 | _1028100 | _1129500 | _1233000 |
| _0108100 | _0404500 | _0605800 | _0719500 | _0835700 | _0932800 | _1028200 | _1129800 | _1233400 |
| _0109800 | _0405700 | _0605900 | _0719800 | _0837041 | _0933100 | _1028600 | _1130600 | _1233500 |
| _0110100 | _0406200 | _0606300 | _0719900 | _0837061 | _0933200 | _1029000 | _1130800 | _1233700 |
| _0111000 | _0406900 | _0606700 | _0720700 | _0901500 | _0933500 | _1029100 | _1131300 | _1234600 |
| _0111900 | _0407300 | _0607800 | _0720900 | _0902900 | _0933600 | _1031500 | _1131400 | _1235900 |
| _0112100 | _0407400 | _0607900 | _0721000 | _0903100 | _0934400 | _1031600 | _1131500 | _1236100 |
| _0112300 | _0408900 | _0608000 | _0721300 | _0904000 | _0936000 | _1033700 | _1132900 | _1236200 |
| _0112400 | _0409000 | _0608500 | _0721400 | _0904200 | _0937500 | _1034200 | _1133600 | _1236600 |
| _0201051 | _0412900 | _0608800 | _0721600 | _0904400 | _0939000 | _1034500 | _1133800 | _1236700 |
| _0202400 | _0413000 | _0609200 | _0721800 | _0905500 | _0939300 | _1035000 | _1134900 | _1238100 |
| _0202700 | _0413500 | _0609300 | _0800800 | _0905900 | _0940000 | _1036900 | _1135800 | _1238900 |
| _0203750 | _0414100 | _0609500 | _0802500 | _0906100 | _0941100 | _1037800 | _1135900 | _1239000 |
| _0204500 | _0414300 | _0609600 | _0802800 | _0906700 | _0941700 | _1039000 | _1136000 | _1240100 |
| _0204600 | _0414900 | _0610300 | _0807700 | _0907200 | _0942000 | _1039100 | _1136700 | _1240600 |
| _0204800 | _0415700 | _0611000 | _0810800 | _0908400 | _0942800 | _1039200 | _1138100 | _1242900 |
| _0205800 | _0416100 | _0611600 | _0810900 | _0908600 | _0943400 | _1040100 | _1138600 | _1243000 |
| _0207100 | _0417100 | _0611800 | _0812100 | _0909300 | _1001200 | _1102100 | _1138800 | _1243600 |
| _0207300 | _0417800 | _0612200 | _0812200 | _0909400 | _1001700 | _1102900 | _1138900 | _1244300 |
| _0208400 | _0501600 | _0612700 | _0812300 | _0909500 | _1002100 | _1103800 | _1140100 | _1245300 |
| _0209000 | _0501700 | _0613000 | _0813000 | _0909800 | _1002200 | _1104500 | _1141400 | _1300700 |
| _0209100 | _0501800 | _0615200 | _0813300 | _0910700 | _1002500 | _1104700 | _1141500 | _1301800 |
| _0209600 | _0502000 | _0615500 | _0814800 | _0910800 | _1002600 | _1105300 | _1141900 | _1302241 |
| _0209700 | _0502500 | _0615700 | _0815100 | _0910900 | _1003400 | _1105600 | _1142500 | _1304000 |
| _0209800 | _0502650 | _0616200 | _0815500 | _0911000 | _1003600 | _1105900 | _1143400 | _1304500 |
| _0210100 | _0503500 | _0616800 | _0815800 | _0911100 | _1006100 | _1106500 | _1143800 | _1305500 |
| _0210400 | _0503800 | _0616900 | _0815900 | _0911200 | _1006200 | _1107600 | _1144800 | _1305800 |
| _0211300 | _0503900 | _0617500 | _0816200 | _0911500 | _1006300 | _1107700 | _1200700 | _1308800 |
| _0212600 | _0506000 | _0619500 | _0816800 | _0911700 | _1006400 | _1109600 | _1202900 | _1309400 |
| _0212800 | _0506900 | _0619600 | _0817000 | _0912500 | _1007600 | _1109900 | _1204000 | _1309600 |
| _0300700 | _0507200 | _0620700 | _0817100 | _0912800 | _1008600 | _1110200 | _1204200 | _1310000 |
| _0300800 | _0508300 | _0620800 | _0817300 | _0915000 | _1009600 | _1110800 | _1205400 | _1310500 |
| _0301300 | _0509500 | _0621000 | _0818200 | _0915600 | _1010400 | _1110900 | _1206800 | _1310700 |
| _0301600 | _0509600 | _0621600 | _0818400 | _0915900 | _1010900 | _1111200 | _1207400 | _1311400 |
| _0301900 | _0510100 | _0622000 | _0819400 | _0916400 | _1012000 | _1111900 | _1208200 | _1312300 |
| _0302500 | _0510700 | _0622100 | _0819500 | _0916650 | _1012100 | _1112000 | _1208500 | _1312400 |
| _0302800 | _0510800 | _0622300 | _0820200 | _0917000 | _1012500 | _1113200 | _1209900 | _1312700 |
| _0303800 | _0512700 | _0622400 | _0820400 | _0917100 | _1012700 | _1113300 | _1214400 | _1312800 |
| _0303900 | _0512800 | _0622500 | _0820700 | _0917300 | _1013200 | _1113400 | _1215700 | _1313200 |
| _0304500 | _0513000 | _0623450 | _0821400 | _0917980 | _1013300 | _1113700 | _1215900 | _1317200 |
| _0304600 | _0513700 | _0701200 | _0821600 | _0918600 | _1013400 | _1113800 | _1216000 | _1318000 |
| _0304700 | _0513800 | _0701300 | _0822500 | _0918900 | _1015800 | _1114200 | _1218100 | _1318200 |
| _0305500 | _0513900 | _0701400 | _0822900 | _0920500 | _1015900 | _1114300 | _1218400 | _1318600 |
| _0305700 | _0514200 | _0701900 | _0823200 | _0920700 | _1016000 | _1114400 | _1218700 | _1318700 |
| _0306100 | _0514800 | _0702000 | _0823300 | _0921200 | _1016200 | _1114500 | _1219700 | _1319000 |
| _0306200 | _0515000 | _0702100 | _0824200 | _0922600 | _1017500 | _1115300 | _1220300 | _1319300 |
| _0306400 | _0515200 | _0702400 | _0824500 | _0923000 | _1018100 | _1116000 | _1220600 | _1319700 |
| _0306600 | _0515300 | _0702500 | _0825300 | _0924500 | _1018800 | _1116900 | _1221500 | _1320100 |
| _0307400 | _0515350 | _0702600 | _0825400 | _0925000 | _1019100 | _1117600 | _1222500 | _1323400 |
| _0307500 | _0515400 | _0702700 | _0825800 | _0925100 | _1019800 | _1118200 | _1222800 | _1324300 |
| _0307700 | _0517700 | _0703300 | _0825900 | _0925400 | _1019900 | _1118400 | _1223200 | _1325100 |
| _0308000 | _0518100 | _0703400 | _0826400 | _0925500 | _1020000 | _1118500 | _1223800 | _1325500 |
| _0309500 | _0521700 | _0703800 | _0826600 | _0925600 | _1020100 | _1118900 | _1223900 | _1326100 |
| _0309700 | _0522100 | _0704400 | _0827100 | _0925700 | _1020300 | _1119400 | _1224500 | _1327061 |
| _0310400 | _0522200 | _0709700 | _0828200 | _0925800 | _1020500 | _1119500 | _1224900 | _1328100 |
| _0310600 | _0523700 | _0710300 | _0829200 | _0928000 | _1020800 | _1119800 | _1225000 | _1329100 |
| _0311400 | _0523900 | _0711600 | _0829300 | _0928500 | _1022500 | _1119900 | _1225300 | _1329200 |
| _0312000 | _0600071 | _0712700 | _0830300 | _0928700 | _1022600 | _1120400 | _1225400 | _1330400 |

Table S4: Gene IDs (PBANKA)

| | | |
|---|---|---|
| _1330500 | _1402200 | _1449600 |
| _1330800 | _1403300 | _1449700 |
| _1331200 | _1404400 | _1449800 |
| _1331300 | _1405900 | _1450400 |
| _1331600 | _1408700 | _1450600 |
| _1331700 | _1409200 | _1450800 |
| _1333000 | _1410600 | _1451000 |
| _1335800 | _1410950 | _1451200 |
| _1336600 | _1411000 | _1452900 |
| _1336900 | _1411500 | _1453000 |
| _1337700 | _1413300 | _1453400 |
| _1339600 | _1414500 | _1454300 |
| _1340700 | _1414800 | _1454900 |
| _1341000 | _1415200 | _1455400 |
| _1341900 | _1415400 | _1456700 |
| _1342000 | _1416800 | _1456800 |
| _1342200 | _1416900 | _1457000 |
| _1342500 | _1417200 | _1457200 |
| _1342700 | _1418200 | _1457700 |
| _1343400 | _1419900 | _1458200 |
| _1343600 | _1422800 | _1458300 |
| _1343800 | _1422900 | _1460000 |
| _1344100 | _1423000 | _1460100 |
| _1344400 | _1423100 | _1460200 |
| _1344700 | _1424000 | _1460500 |
| _1345100 | _1424500 | _1460600 |
| _1345400 | _1425200 | _1461300 |
| _1345500 | _1426500 | _1461400 |
| _1345900 | _1426600 | _1462300 |
| _1346300 | _1427100 | |
| _1346400 | _1429100 | |
| _1346800 | _1429200 | |
| _1346900 | _1429300 | |
| _1347000 | _1429400 | |
| _1347200 | _1431100 | |
| _1347400 | _1431300 | |
| _1348200 | _1432300 | |
| _1349500 | _1432400 | |
| _1349800 | _1432700 | |
| _1351351 | _1433600 | |
| _1351400 | _1433700 | |
| _1351700 | _1434300 | |
| _1352100 | _1434400 | |
| _1352500 | _1434900 | |
| _1352800 | _1435000 | |
| _1353400 | _1435200 | |
| _1353800 | _1435300 | |
| _1354351 | _1435500 | |
| _1355700 | _1435600 | |
| _1356800 | _1435900 | |
| _1357100 | _1436500 | |
| _1358500 | _1437600 | |
| _1360000 | _1437800 | |
| _1360600 | _1437900 | |
| _1361000 | _1439000 | |
| _1361400 | _1439100 | |
| _1362500 | _1442300 | |
| _1362700 | _1443000 | |
| _1363200 | _1443700 | |
| _1363600 | _1443900 | |
| _1363700 | _1444500 | |
| _1364000 | _1444600 | |
| _1364400 | _1445000 | |
| _1364800 | _1446000 | |
| _1365721 | _1447200 | |
| _1400086 | _1447300 | |
| _1400091 | _1449000 | |
| _1402100 | _1449300 | |

Table S4: Gene IDs (PBANKA)

Community 7

_0105500
_0209900
_0305800
_0315410
_0714900
_0833100
_1238700
_1246600
_1314800
_1363800
_1445100

Table S4: Gene IDs (PBANKA)

**Community 8**

_0303400
_0828800
_1000021
_1106100
_1110700
_1138000
_1301900
_1302000
_1303500
_1306600
_1332800
_1344300

Table S4: Gene IDs (PBANKA)

**Community 9**

_0400500
_0518900
_0519500
_0827000
_1003900
_1146261
_1423400
_API00120
_API00150
_API00160
_API00170
_API00206
_API00300
_API00304
_API00320
_API00330
_API00340
_API00360
_API00380
_API00390
_API00410
_API00430

Table S4: Gene IDs (PBANKA)

**Community 10**

_0213800
_0312500
_0315800
_0402300
_0613200
_0621300
_0712100
_0803000
_0808000
_0816700
_0828400
_0902600
_0913900
_1011100
_1025400
_1036600
_1038900
_1105100
_1110100
_1114000
_1219200
_1225100
_1243900
_1303400
_1306900
_1316900
_1358300
_1417000
_1417900
_1418100
_1442100

Table S4: Gene IDs (PBANKA)

**Community 11**

_0101000
_0101900
_0107300
_0108900
_0206300
_0207800
_0210900
_0213200
_0213300
_0214300
_0305600
_0406000
_0412000
_0418100
_0418400
_0507000
_0512900
_0514600
_0514700
_0516700
_0520500
_0521600
_0614000
_0618600
_0622700
_0709800
_0717000
_0721700
_0801900
_0803100
_0803800
_0804400
_0805600
_0805700
_0814700
_0816100
_0817500
_0820000
_0901200
_0901600
_0901800
_0903500
_0911300
_0913300
_0913500
_0914300
_0916200
_0918700
_0934100
_0935000
_0935900
_0940800
_0943000
_0943100
_1001600
_1001900
_1003000
_1004800
_1005300
_1006000
_1010600
_1011400
_1012800
_1024900
_1030300
_1030500
_1031400
_1033200
_1033800
_1038100
_1039300
_1039700
_1039800
_1039900
_1101000
_1102700
_1106600
_1125700
_1127200
_1127600
_1132400
_1134100
_1134200
_1135500
_1135600
_1139600
_1139700
_1140700
_1142900
_1144700
_1203000
_1211600
_1212100
_1213900
_1219300
_1220700
_1225200
_1230300
_1235600
_1240200
_1244200
_1245841
_1301000
_1301500
_1302100
_1303300
_1304700
_1304800
_1304900
_1307300
_1307400
_1309900
_1316300
_1317000
_1317100
_1319600
_1319900
_1320300
_1321200
_1324600
_1326700
_1326800
_1327400
_1328600
_1330300
_1335900
_1340100
_1343300
_1347100
_1349600
_1354700
_1357000
_1357400
_1357500
_1358100
_1359900
_1360500
_1360700
_1362400
_1401400
_1402300
_1403000
_1403200
_1404100
_1408300
_1409700
_1409800
_1410800
_1415600
_1415800
_1415900
_1418700
_1418800
_1419500
_1424100
_1425400
_1427800
_1429600
_1431900
_1433800
_1434700
_1435800
_1436800
_1437300
_1438400
_1444000
_1444100
_1446900
_1450700
_1454100
_1456600
_MIT00800
_MIT01000
_MIT01800
_MIT01900
_MIT02200

Table S4: Gene IDs (PBANKA)

| | | | | | | | |
|---|---|---|---|---|---|---|---|
| **Community 12** | _0501500 | _0715500 | _0921600 | _1108200 | _1222000 | _1341600 | _1446100 |
| _0102500 | _0502100 | _0715900 | _0922500 | _1108400 | _1222100 | _1341800 | _1447400 |
| _0102800 | _0502200 | _0717500 | _0922800 | _1109400 | _1223100 | _1342900 | _1448000 |
| _0103100 | _0504600 | _0718000 | _0923100 | _1110000 | _1224400 | _1345700 | _1448300 |
| _0104600 | _0504700 | _0718100 | _0923200 | _1111400 | _1225800 | _1346600 | _1448400 |
| _0104700 | _0505100 | _0720500 | _0923300 | _1111800 | _1226500 | _1347800 | _1448900 |
| _0105400 | _0505200 | _0720600 | _0923600 | _1112200 | _1226600 | _1347900 | _1450000 |
| _0106500 | _0505600 | _0721500 | _0923700 | _1112300 | _1226800 | _1348100 | _1450200 |
| _0106700 | _0507100 | _0800700 | _0926600 | _1113000 | _1227700 | _1349700 | _1450300 |
| _0106950 | _0507400 | _0802100 | _0927900 | _1115500 | _1230700 | _1350100 | _1451600 |
| _0107100 | _0510500 | _0802200 | _0928300 | _1116300 | _1230800 | _1350800 | _1453200 |
| _0108400 | _0511100 | _0802700 | _0930000 | _1116500 | _1230900 | _1351100 | _1453700 |
| _0109000 | _0511800 | _0803500 | _0930500 | _1119100 | _1231200 | _1351600 | _1454000 |
| _0109300 | _0514500 | _0805800 | _0930600 | _1121400 | _1231500 | _1352300 | _1454200 |
| _0110900 | _0516200 | _0805900 | _0930900 | _1121500 | _1232200 | _1354100 | _1455100 |
| _0204200 | _0516600 | _0806400 | _0932200 | _1121600 | _1233100 | _1358600 | _1456200 |
| _0204400 | _0517800 | _0807500 | _0933900 | _1122300 | _1234700 | _1359100 | _1458700 |
| _0205300 | _0518000 | _0808300 | _0934300 | _1124200 | _1235100 | _1359500 | _1459800 |
| _0205400 | _0518200 | _0809100 | _0935100 | _1124300 | _1237600 | _1360300 | _1461000 |
| _0206500 | _0518300 | _0809200 | _0935500 | _1124800 | _1237900 | _1362000 | _1461500 |
| _0206800 | _0519800 | _0809700 | _0936400 | _1125000 | _1238000 | _1362100 | _1463100 |
| _0207000 | _0520000 | _0809900 | _0936900 | _1126200 | _1238400 | _1363500 | _1463200 |
| _0208500 | _0520700 | _0811600 | _0937000 | _1126600 | _1239900 | _1365000 | _1463400 |
| _0208600 | _0520800 | _0812400 | _0938100 | _1127800 | _1240800 | _1365200 | _MIT01100 |
| _0210000 | _0521000 | _0813900 | _0938300 | _1130000 | _1240900 | _1402700 | |
| _0210700 | _0522500 | _0814200 | _0938400 | _1130500 | _1241100 | _1404900 | |
| _0211500 | _0522700 | _0814900 | _0938500 | _1132200 | _1241300 | _1405800 | |
| _0211600 | _0524600 | _0816000 | _0939700 | _1133700 | _1241900 | _1407300 | |
| _0211900 | _0601100 | _0816900 | _0939800 | _1136400 | _1242700 | _1407500 | |
| _0212400 | _0601200 | _0817200 | _0940100 | _1136600 | _1244600 | _1409000 | |
| _0212900 | _0602100 | _0818900 | _0940200 | _1137700 | _1244700 | _1409900 | |
| _0301100 | _0602700 | _0821100 | _0940300 | _1137800 | _1245200 | _1410400 | |
| _0303100 | _0603000 | _0821300 | _0940400 | _1137900 | _1301400 | _1410500 | |
| _0303200 | _0603400 | _0822000 | _0940600 | _1138200 | _1302200 | _1411800 | |
| _0303600 | _0603500 | _0824100 | _0940900 | _1138300 | _1302300 | _1413200 | |
| _0305200 | _0604300 | _0824700 | _0941600 | _1139100 | _1302600 | _1414600 | |
| _0306300 | _0607000 | _0825000 | _1001400 | _1139400 | _1303000 | _1416100 | |
| _0306900 | _0608400 | _0827300 | _1003800 | _1140400 | _1303800 | _1418400 | |
| _0307100 | _0609000 | _0827600 | _1004600 | _1140500 | _1304300 | _1419000 | |
| _0308100 | _0610000 | _0827800 | _1005400 | _1142400 | _1304400 | _1419100 | |
| _0308900 | _0610600 | _0828000 | _1006500 | _1143000 | _1305300 | _1420200 | |
| _0309800 | _0611400 | _0828600 | _1006600 | _1144000 | _1305400 | _1421900 | |
| _0312200 | _0612800 | _0829500 | _1007800 | _1145300 | _1306300 | _1422600 | |
| _0312800 | _0613400 | _0830000 | _1008200 | _1205800 | _1308200 | _1423800 | |
| _0313800 | _0613500 | _0832000 | _1008400 | _1206400 | _1309100 | _1423900 | |
| _0314400 | _0616500 | _0832200 | _1009100 | _1206600 | _1310100 | _1424200 | |
| _0314700 | _0617900 | _0832500 | _1009400 | _1207600 | _1310400 | _1425100 | |
| _0314900 | _0618100 | _0832600 | _1014000 | _1207800 | _1311500 | _1425900 | |
| _0402900 | _0619300 | _0833200 | _1014300 | _1209800 | _1315100 | _1427300 | |
| _0404200 | _0702800 | _0903600 | _1014400 | _1210500 | _1316400 | _1427500 | |
| _0404400 | _0703200 | _0903700 | _1014900 | _1210700 | _1320700 | _1430700 | |
| _0404900 | _0704100 | _0905100 | _1017700 | _1210900 | _1321600 | _1430900 | |
| _0405200 | _0704300 | _0905800 | _1024000 | _1211100 | _1322100 | _1434800 | |
| _0406500 | _0706400 | _0907300 | _1027100 | _1211700 | _1322200 | _1436200 | |
| _0406800 | _0707600 | _0907700 | _1027900 | _1212300 | _1322300 | _1436700 | |
| _0407500 | _0708000 | _0908800 | _1028900 | _1213600 | _1325800 | _1437000 | |
| _0407800 | _0708400 | _0910100 | _1030100 | _1213700 | _1325900 | _1438000 | |
| _0407900 | _0708800 | _0911900 | _1031200 | _1214000 | _1326300 | _1438600 | |
| _0409200 | _0709000 | _0912600 | _1031800 | _1215000 | _1326400 | _1438900 | |
| _0409400 | _0709200 | _0912700 | _1033300 | _1215100 | _1334000 | _1439600 | |
| _0410300 | _0709300 | _0915700 | _1035300 | _1217000 | _1334100 | _1441100 | |
| _0410400 | _0709900 | _0917900 | _1038300 | _1217200 | _1334500 | _1441200 | |
| _0410500 | _0710600 | _0919000 | _1039600 | _1218200 | _1338400 | _1441300 | |
| _0413800 | _0713300 | _0919700 | _1101600 | _1218900 | _1338700 | _1441600 | |
| _0416000 | _0713500 | _0919800 | _1103200 | _1219400 | _1339300 | _1441900 | |
| _0416700 | _0713700 | _0920800 | _1104800 | _1219600 | _1339700 | _1443200 | |
| _0417300 | _0713900 | _0920900 | _1107100 | _1219800 | _1339800 | _1444900 | |
| _0418200 | _0714100 | _0921100 | _1107300 | _1221400 | _1339900 | _1445800 | |

Table S4: Gene IDs (PBANKA)

| | | | | | | | |
|---|---|---|---|---|---|---|---|
| **Community 13** | _0414500 | _0708900 | _0930400 | _1115400 | _1234400 | _1358700 | _1461800 |
| _0103000 | _0415000 | _0710000 | _0931700 | _1116100 | _1235300 | _1359000 | _1462400 |
| _0103400 | _0415500 | _0710100 | _0932100 | _1116200 | _1236000 | _1359600 | _1463600 |
| _0105700 | _0416800 | _0710900 | _0932700 | _1117300 | _1237200 | _1363900 | _MIT00600 |
| _0105800 | _0416900 | _0712600 | _0933700 | _1117400 | _1237400 | _1364300 | _MIT00700 |
| _0105900 | _0417000 | _0714200 | _0933800 | _1117800 | _1238500 | _1365400 | |
| _0106600 | _0504500 | _0715100 | _0938000 | _1118600 | _1239400 | _1401700 | |
| _0107000 | _0505000 | _0715600 | _0939900 | _1119300 | _1240500 | _1401800 | |
| _0107500 | _0507300 | _0716000 | _0941400 | _1119700 | _1241600 | _1402800 | |
| _0108500 | _0507500 | _0716500 | _0942100 | _1120600 | _1242000 | _1402900 | |
| _0109500 | _0508600 | _0717600 | _0942200 | _1121200 | _1242600 | _1403600 | |
| _0109600 | _0508800 | _0717700 | _0943200 | _1121300 | _1243700 | _1403700 | |
| _0110200 | _0508900 | _0718600 | _1000800 | _1121900 | _1244000 | _1404000 | |
| _0111700 | _0509400 | _0800900 | _1003500 | _1123000 | _1244100 | _1404200 | |
| _0200700 | _0509800 | _0801700 | _1004000 | _1123600 | _1244400 | _1404600 | |
| _0201800 | _0510300 | _0802600 | _1004200 | _1124600 | _1244800 | _1404800 | |
| _0201900 | _0510400 | _0803300 | _1004700 | _1124900 | _1300900 | _1406300 | |
| _0202000 | _0516800 | _0803700 | _1007200 | _1126100 | _1301300 | _1406900 | |
| _0202200 | _0518400 | _0804200 | _1008800 | _1126700 | _1302700 | _1407100 | |
| _0202300 | _0519600 | _0805000 | _1009000 | _1127500 | _1302800 | _1408500 | |
| _0203600 | _0519700 | _0805500 | _1009500 | _1129100 | _1302900 | _1408600 | |
| _0206000 | _0521100 | _0806200 | _1011300 | _1130100 | _1305900 | _1410900 | |
| _0206400 | _0521241 | _0808500 | _1013700 | _1130400 | _1306000 | _1411400 | |
| _0211400 | _0521800 | _0811500 | _1014100 | _1131600 | _1306100 | _1411600 | |
| _0301200 | _0521900 | _0812800 | _1014700 | _1133000 | _1306400 | _1411700 | |
| _0302400 | _0522400 | _0812900 | _1015000 | _1133200 | _1306700 | _1412800 | |
| _0305900 | _0523300 | _0813200 | _1016100 | _1133500 | _1307500 | _1414700 | |
| _0307300 | _0523500 | _0814100 | _1016600 | _1135400 | _1311100 | _1415300 | |
| _0308200 | _0523600 | _0814300 | _1016700 | _1136200 | _1311200 | _1417100 | |
| _0308400 | _0601500 | _0814500 | _1019200 | _1136500 | _1312200 | _1417500 | |
| _0309000 | _0602200 | _0814600 | _1019300 | _1137400 | _1312600 | _1418500 | |
| _0310200 | _0603100 | _0823800 | _1021300 | _1137600 | _1313600 | _1418900 | |
| _0310300 | _0603300 | _0825100 | _1021400 | _1139200 | _1314000 | _1421800 | |
| _0315000 | _0603600 | _0826300 | _1021500 | _1139300 | _1314500 | _1422000 | |
| _0315900 | _0603800 | _0827700 | _1024800 | _1139500 | _1314600 | _1422100 | |
| _0400900 | _0604700 | _0827900 | _1025600 | _1140900 | _1317300 | _1424400 | |
| _0401200 | _0605400 | _0830100 | _1025800 | _1141000 | _1318300 | _1425500 | |
| _0401300 | _0606000 | _0830450 | _1025900 | _1142000 | _1318900 | _1425600 | |
| _0401500 | _0606800 | _0830600 | _1026400 | _1142600 | _1322900 | _1425700 | |
| _0401600 | _0607100 | _0830700 | _1027300 | _1144400 | _1323300 | _1427400 | |
| _0401800 | _0607200 | _0830800 | _1029300 | _1144500 | _1323900 | _1429800 | |
| _0402200 | _0607500 | _0834000 | _1032000 | _1201600 | _1324100 | _1430100 | |
| _0403500 | _0607600 | _0834700 | _1032500 | _1202100 | _1324800 | _1430200 | |
| _0405000 | _0609800 | _0835000 | _1032800 | _1204300 | _1327080 | _1430800 | |
| _0405100 | _0611200 | _0835100 | _1034100 | _1204900 | _1329700 | _1431000 | |
| _0405800 | _0611300 | _0835200 | _1035500 | _1205200 | _1330600 | _1436900 | |
| _0405900 | _0612000 | _0836700 | _1036200 | _1206900 | _1330700 | _1438700 | |
| _0406100 | _0612300 | _0903300 | _1036700 | _1209100 | _1331000 | _1439700 | |
| _0406700 | _0614100 | _0906200 | _1038400 | _1209500 | _1332200 | _1440700 | |
| _0408000 | _0615600 | _0906300 | _1038600 | _1211200 | _1332900 | _1440800 | |
| _0408100 | _0616000 | _0907600 | _1101700 | _1211400 | _1333900 | _1440900 | |
| _0409300 | _0616100 | _0908700 | _1101800 | _1214200 | _1336000 | _1442800 | |
| _0409800 | _0616600 | _0909100 | _1102500 | _1215200 | _1336800 | _1442900 | |
| _0409900 | _0617300 | _0911600 | _1102800 | _1217700 | _1337400 | _1444200 | |
| _0410000 | _0618000 | _0912100 | _1104000 | _1217800 | _1337900 | _1444300 | |
| _0410100 | _0618900 | _0913100 | _1104200 | _1218500 | _1338000 | _1444400 | |
| _0410700 | _0620500 | _0914600 | _1105000 | _1218600 | _1338200 | _1446700 | |
| _0410800 | _0621700 | _0916100 | _1105200 | _1219900 | _1338800 | _1447600 | |
| _0410900 | _0701500 | _0916800 | _1106200 | _1220900 | _1340000 | _1447700 | |
| _0411000 | _0702300 | _0918200 | _1107500 | _1221100 | _1340200 | _1448800 | |
| _0411400 | _0706100 | _0918400 | _1109300 | _1223600 | _1341200 | _1450500 | |
| _0411500 | _0706200 | _0921300 | _1111300 | _1224600 | _1345600 | _1451300 | |
| _0411700 | _0706500 | _0922700 | _1111600 | _1224700 | _1346200 | _1454800 | |
| _0411900 | _0706800 | _0924600 | _1111700 | _1224800 | _1348000 | _1455900 | |
| _0411950 | _0706900 | _0926200 | _1112500 | _1228600 | _1348700 | _1456300 | |
| _0413300 | _0707500 | _0926300 | _1112600 | _1231300 | _1350000 | _1456400 | |
| _0413900 | _0708600 | _0927500 | _1113500 | _1231900 | _1350900 | _1457100 | |
| _0414200 | _0708700 | _0928600 | _1114800 | _1233900 | _1358400 | _1458400 | |

Table S4: Gene IDs (PBANKA)

| | | | | | | | | |
|---|---|---|---|---|---|---|---|---|
| **Community 14** | _0310700 | _0515700 | _0704900 | _0818000 | _0914100 | _1013900 | _1111500 | _1205700 |
| _0100400 | _0311500 | _0516000 | _0705400 | _0818300 | _0914900 | _1014200 | _1112700 | _1206000 |
| _0101500 | _0311700 | _0516400 | _0705500 | _0818800 | _0915100 | _1014500 | _1112800 | _1206500 |
| _0102400 | _0312100 | _0517600 | _0706600 | _0819000 | _0915400 | _1014600 | _1112900 | _1208000 |
| _0102600 | _0312700 | _0517900 | _0707000 | _0820100 | _0916500 | _1014800 | _1113600 | _1208900 |
| _0102700 | _0313200 | _0518500 | _0707100 | _0820500 | _0916600 | _1016900 | _1114100 | _1209200 |
| _0102900 | _0313300 | _0519000 | _0707200 | _0820800 | _0916700 | _1017000 | _1115000 | _1209300 |
| _0103200 | _0313500 | _0519100 | _0707700 | _0821000 | _0916900 | _1017400 | _1115100 | _1209400 |
| _0103700 | _0313600 | _0519200 | _0707900 | _0821200 | _0917800 | _1017600 | _1115200 | _1209600 |
| _0104200 | _0313700 | _0519300 | _0708100 | _0821500 | _0917990 | _1018200 | _1115600 | _1209700 |
| _0104300 | _0314800 | _0519400 | _0708200 | _0822200 | _0918500 | _1018300 | _1115700 | _1210600 |
| _0104400 | _0316600 | _0520100 | _0708500 | _0822300 | _0919400 | _1018900 | _1117200 | _1212200 |
| _0105100 | _0316700 | _0520400 | _0709600 | _0822400 | _0919500 | _1019000 | _1117900 | _1212400 |
| _0106200 | _0316981 | _0521200 | _0711000 | _0824000 | _0919900 | _1019360 | _1118700 | _1212500 |
| _0106900 | _0317181 | _0521500 | _0711100 | _0824400 | _0921000 | _1020900 | _1119200 | _1212600 |
| _0107200 | _0402000 | _0522300 | _0711300 | _0825200 | _0922200 | _1021200 | _1119600 | _1213200 |
| _0107600 | _0402100 | _0523000 | _0711400 | _0825500 | _0922400 | _1022100 | _1120100 | _1213300 |
| _0108300 | _0402400 | _0523400 | _0711500 | _0825700 | _0924100 | _1022300 | _1120500 | _1213400 |
| _0109400 | _0402500 | _0523800 | _0712500 | _0827200 | _0924200 | _1022400 | _1120900 | _1214100 |
| _0109900 | _0402700 | _0600600 | _0713100 | _0827400 | _0924700 | _1023100 | _1121000 | _1214500 |
| _0110300 | _0403600 | _0600800 | _0713400 | _0828100 | _0926400 | _1023600 | _1122400 | _1215300 |
| _0110600 | _0403700 | _0601800 | _0713800 | _0828300 | _0926500 | _1023900 | _1123100 | _1215400 |
| _0110700 | _0404100 | _0602900 | _0714000 | _0828900 | _0927000 | _1025000 | _1123200 | _1216200 |
| _0110800 | _0404300 | _0603650 | _0714600 | _0829400 | _0927100 | _1026000 | _1124400 | _1216300 |
| _0111500 | _0405600 | _0603900 | _0715300 | _0829600 | _0927300 | _1026300 | _1124500 | _1217100 |
| _0111600 | _0406400 | _0604000 | _0716200 | _0829900 | _0927400 | _1027400 | _1124700 | _1217900 |
| _0112000 | _0406600 | _0604400 | _0718500 | _0830200 | _0927600 | _1028700 | _1125300 | _1218300 |
| _0112200 | _0408200 | _0605600 | _0718700 | _0830400 | _0927700 | _1029400 | _1125400 | _1219100 |
| _0201340 | _0408500 | _0606200 | _0719100 | _0830500 | _0931500 | _1029500 | _1125800 | _1221900 |
| _0201700 | _0409700 | _0606500 | _0719400 | _0831000 | _0932000 | _1029600 | _1126300 | _1222600 |
| _0202100 | _0410200 | _0606900 | _0719700 | _0831600 | _0932500 | _1030400 | _1127100 | _1223700 |
| _0204300 | _0410650 | _0608600 | _0720000 | _0832300 | _0932600 | _1031700 | _1127300 | _1224100 |
| _0206700 | _0411800 | _0609250 | _0720100 | _0832400 | _0934600 | _1032100 | _1128800 | _1224300 |
| _0207400 | _0413400 | _0610700 | _0721100 | _0832700 | _0934700 | _1032400 | _1129600 | _1225500 |
| _0207900 | _0415100 | _0611100 | _0722921 | _0833700 | _0934800 | _1035200 | _1129900 | _1225600 |
| _0208000 | _0415400 | _0611500 | _0800500 | _0833800 | _0934900 | _1035400 | _1131100 | _1227500 |
| _0208900 | _0415800 | _0611700 | _0800600 | _0833900 | _0935700 | _1035700 | _1132300 | _1228100 |
| _0209300 | _0415900 | _0611900 | _0801100 | _0834600 | _0936100 | _1036500 | _1132800 | _1228200 |
| _0210300 | _0417200 | _0612400 | _0801300 | _0835600 | _0936200 | _1037700 | _1133100 | _1228800 |
| _0211800 | _0417600 | _0612600 | _0801500 | _0836100 | _0936800 | _1037900 | _1139800 | _1228900 |
| _0212300 | _0418300 | _0612900 | _0801800 | _0836300 | _0937100 | _1038200 | _1141200 | _1229400 |
| _0213900 | _0501100 | _0613300 | _0802000 | _0901000 | _0937600 | _1038500 | _1141600 | _1229500 |
| _0214000 | _0501400 | _0613600 | _0803900 | _0901900 | _0937900 | _1038700 | _1143100 | _1230000 |
| _0214500 | _0503300 | _0614800 | _0804300 | _0902000 | _0938200 | _1038800 | _1143200 | _1230600 |
| _0214550 | _0503400 | _0615300 | _0804500 | _0902200 | _0938800 | _1040000 | _1143700 | _1231400 |
| _0214951 | _0504100 | _0616400 | _0806300 | _0902300 | _0940500 | _1101400 | _1143900 | _1232000 |
| _0215800 | _0504300 | _0616700 | _0806800 | _0902400 | _0941000 | _1101500 | _1145000 | _1232300 |
| _0216000 | _0504400 | _0617000 | _0807200 | _0903400 | _0942300 | _1101900 | _1145200 | _1232600 |
| _0216781 | _0504800 | _0617800 | _0807800 | _0903800 | _0942400 | _1102300 | _1201200 | _1233200 |
| _0300100 | _0505400 | _0618400 | _0808380 | _0904600 | _0943900 | _1102400 | _1201400 | _1233600 |
| _0300900 | _0505500 | _0618500 | _0808600 | _0904800 | _1000900 | _1103300 | _1201500 | _1234100 |
| _0301000 | _0506600 | _0618700 | _0809000 | _0904900 | _1001300 | _1103700 | _1201800 | _1234900 |
| _0301700 | _0507700 | _0619000 | _0809400 | _0905300 | _1001500 | _1103900 | _1202000 | _1235000 |
| _0301800 | _0507800 | _0619200 | _0809500 | _0905400 | _1002400 | _1104100 | _1202300 | _1235700 |
| _0302100 | _0507900 | _0619700 | _0809600 | _0906000 | _1002800 | _1104400 | _1203100 | _1236800 |
| _0302200 | _0508000 | _0620300 | _0810200 | _0906800 | _1002900 | _1106000 | _1203200 | _1236900 |
| _0302900 | _0509300 | _0620400 | _0810300 | _0907100 | _1004100 | _1106300 | _1203450 | _1237100 |
| _0303000 | _0509700 | _0620600 | _0810400 | _0907400 | _1004300 | _1106900 | _1203500 | _1237300 |
| _0303300 | _0509900 | _0622900 | _0810700 | _0907500 | _1004500 | _1107000 | _1203600 | _1238300 |
| _0303500 | _0510000 | _0700100 | _0811200 | _0907900 | _1005800 | _1108000 | _1203900 | _1240400 |
| _0304800 | _0510600 | _0700800 | _0811800 | _0908500 | _1007900 | _1108100 | _1204100 | _1240700 |
| _0304900 | _0512000 | _0702200 | _0812600 | _0909000 | _1009200 | _1108300 | _1204500 | _1241000 |
| _0305000 | _0512300 | _0703000 | _0812700 | _0909700 | _1009900 | _1108500 | _1204600 | _1241200 |
| _0305100 | _0513100 | _0703500 | _0813600 | _0910300 | _1011500 | _1108700 | _1204700 | _1241400 |
| _0306700 | _0514400 | _0704000 | _0813700 | _0910400 | _1011600 | _1108800 | _1204800 | _1243500 |
| _0307600 | _0514900 | _0704200 | _0815200 | _0910600 | _1011700 | _1109500 | _1205100 | _1244900 |
| _0309900 | _0515100 | _0704700 | _0815400 | _0913200 | _1013000 | _1110300 | _1205300 | _1245150 |
| _0310500 | _0515600 | _0704800 | _0816500 | _0913600 | _1013600 | _1110600 | _1205500 | _1245500 |

Table S4: Gene IDs (PBANKA)

| | | | |
|---|---|---|---|
| _1245600 | _1338900 | _1417700 | _1464100 |
| _1300096 | _1340400 | _1418000 | _1464900 |
| _1300500 | _1341400 | _1419300 | _MIT02800 |
| _1303100 | _1341700 | _1419400 | |
| _1303900 | _1342300 | _1419800 | |
| _1305200 | _1343100 | _1420100 | |
| _1307900 | _1343200 | _1420700 | |
| _1308100 | _1343700 | _1421000 | |
| _1309800 | _1344200 | _1421400 | |
| _1311300 | _1346000 | _1421500 | |
| _1311700 | _1347300 | _1421600 | |
| _1312000 | _1348600 | _1421700 | |
| _1312100 | _1348800 | _1422300 | |
| _1312900 | _1349000 | _1423700 | |
| _1313100 | _1349100 | _1424700 | |
| _1313300 | _1350200 | _1425800 | |
| _1313700 | _1351500 | _1427200 | |
| _1313900 | _1352400 | _1427600 | |
| _1315300 | _1353100 | _1427700 | |
| _1315700 | _1353900 | _1428300 | |
| _1316200 | _1354200 | _1428600 | |
| _1316500 | _1354600 | _1430300 | |
| _1316800 | _1354800 | _1430600 | |
| _1317800 | _1354900 | _1431200 | |
| _1318500 | _1356600 | _1431400 | |
| _1318800 | _1357300 | _1431500 | |
| _1319200 | _1357900 | _1431800 | |
| _1319500 | _1358900 | _1432200 | |
| _1320400 | _1359700 | _1435400 | |
| _1320800 | _1359800 | _1436100 | |
| _1321500 | _1360400 | _1436300 | |
| _1321800 | _1360800 | _1436600 | |
| _1321900 | _1360900 | _1437200 | |
| _1323500 | _1361300 | _1437500 | |
| _1323600 | _1361500 | _1437700 | |
| _1323800 | _1361600 | _1439500 | |
| _1324500 | _1361700 | _1441800 | |
| _1324700 | _1361800 | _1442700 | |
| _1325200 | _1362800 | _1444800 | |
| _1325400 | _1363400 | _1446400 | |
| _1325700 | _1364700 | _1447500 | |
| _1326200 | _1400400 | _1447800 | |
| _1327100 | _1400600 | _1447900 | |
| _1328200 | _1400900 | _1448600 | |
| _1328900 | _1403900 | _1451100 | |
| _1329000 | _1404300 | _1451400 | |
| _1329400 | _1404500 | _1451500 | |
| _1329900 | _1405000 | _1453100 | |
| _1330000 | _1406400 | _1453300 | |
| _1330900 | _1406500 | _1453800 | |
| _1332300 | _1406600 | _1453900 | |
| _1332400 | _1407000 | _1454400 | |
| _1332700 | _1407200 | _1454600 | |
| _1333100 | _1407700 | _1456500 | |
| _1333700 | _1407900 | _1458800 | |
| _1334700 | _1408400 | _1459100 | |
| _1334800 | _1409100 | _1459400 | |
| _1334900 | _1409600 | _1459500 | |
| _1335000 | _1411900 | _1460300 | |
| _1335700 | _1412000 | _1460700 | |
| _1336100 | _1412500 | _1460900 | |
| _1336200 | _1413600 | _1461700 | |
| _1336400 | _1413700 | _1461900 | |
| _1336500 | _1413800 | _1462100 | |
| _1337300 | _1414300 | _1462500 | |
| _1337800 | _1416000 | _1463000 | |
| _1338100 | _1416500 | _1463300 | |
| _1338500 | _1417400 | _1463900 | |

Table S4: Gene IDs (PBANKA)

**Community mixed**

_0000401
_0007701
_0101100
_0101200
_0101600
_0104100
_0106400
_0111300
_0111400
_0112500
_0203100
_0205900
_0206100
_0207700
_0210500
_0214100
_0214200
_0304200
_0305400
_0306800
_0309200
_0309400
_0310000
_0311000
_0311100
_0311900
_0312300
_0316650
_0402600
_0404600
_0414400
_0414800
_0417900
_0500100
_0502400
_0505300
_0511300
_0513400
_0513600
_0520300
_0521221
_0524841
_0605300
_0607300
_0608700
_0610200
_0613100
_0614200
_0621900
_0622921
_0700021
_0705600
_0710400
_0712200
_0715000
_0718300
_0718800
_0720200
_0800021
_0800041
_0802400
_0802900
_0804600
_0807900
_0808420
_0809300
_0814400
_0815700
_0818500
_0819600
_0821800
_0823500
_0833400
_0901400
_0902700
_0903200
_0903900
_0911650
_0914400
_0921700
_0922900
_0923800
_0929200
_0929900
_0932900
_0933000
_0941200
_0941800
_0941900
_0944021
_0944041
_0944121
_1000100
_1000651
_1002000
_1002300
_1004400
_1005500
_1008000
_1008100
_1008700
_1013800
_1015400
_1017300
_1017800
_1021700
_1023000
_1023400
_1024500
_1024600
_1032700
_1100011
_1100500
_1100600
_1100821
_1109700
_1117000
_1118800
_1122600
_1125900
_1137300
_1140300
_1140800
_1144300
_1145741
_1146400
_1146761
_1200061
_1202800
_1206700
_1210400
_1212800
_1213500
_1214800
_1214900
_1217300
_1217600
_1222300
_1236400
_1239200
_1239500
_1241500
_1241800
_1242400
_1243100
_1246921
_1300061
_1303600
_1307700
_1308300
_1314300
_1314900
_1315000
_1317500
_1319400
_1322800
_1326900
_1333800
_1335200
_1335400
_1336700
_1340500
_1341100
_1342100
_1345000
_1348400
_1349300
_1349900
_1351300
_1352600
_1353500
_1362300
_1362900
_1365100
_1408200
_1412300
_1414900
_1415700
_1420000
_1421261
_1422400
_1433000
_1434600
_1438300
_1439900
_1444700
_1445900
_1447000
_1449900
_1453500
_1457900
_1462600
_1464200
_1465700
_1465821
_1466201
_API00550

Table S4: Gene IDs (PBANKA)

| | | | | | | |
|---|---|---|---|---|---|---|
| **Not in GCN** | _0512200 | _0824900 | _1018400 | _1216700 | _1345200 | _1448700 |
| _0001201 | _0512600 | _0826000 | _1018500 | _1217400 | _1345300 | _1451700 |
| _0006300 | _0513300 | _0826100 | _1019310 | _1220400 | _1348500 | _1451900 |
| _0100100 | _0515500 | _0826200 | _1019320 | _1221800 | _1349400 | _1452100 |
| _0102100 | _0516500 | _0826500 | _1020700 | _1223000 | _1350600 | _1452700 |
| _0102300 | _0522000 | _0826800 | _1021900 | _1224000 | _1350700 | _1455700 |
| _0108600 | _0522600 | _0828500 | _1022900 | _1226200 | _1351000 | _1457500 |
| _0109100 | _0522900 | _0828700 | _1024200 | _1226700 | _1351200 | _1458000 |
| _0109700 | _0523200 | _0829800 | _1024311 | _1226900 | _1353600 | _1459200 |
| _0110500 | _0524000 | _0831500 | _1024331 | _1227600 | _1355900 | _1459300 |
| _0111100 | _0600251 | _0832900 | _1024351 | _1228000 | _1356100 | _1459600 |
| _0111800 | _0600351 | _0833600 | _1025100 | _1228400 | _1356400 | _1461600 |
| _0112681 | _0601300 | _0834200 | _1026200 | _1228500 | _1357700 | _1462700 |
| _0200500 | _0601600 | _0835300 | _1026600 | _1229100 | _1359300 | _1463700 |
| _0200650 | _0601900 | _0835500 | _1027000 | _1229300 | _1365300 | _1464400 |
| _0202500 | _0602600 | _0835800 | _1027800 | _1229800 | _1400096 | _API00055 |
| _0202600 | _0604100 | _0835900 | _1028000 | _1230100 | _1400500 | _API00280 |
| _0202800 | _0606400 | _0837021 | _1029700 | _1233300 | _1402000 | _API00290 |
| _0204000 | _0607400 | _0900900 | _1031100 | _1233800 | _1402400 | _MIT01400 |
| _0205600 | _0607700 | _0904100 | _1033400 | _1235400 | _1403100 | _MIT01700 |
| _0205700 | _0608900 | _0904300 | _1034600 | _1235500 | _1403400 | _MIT03100 |
| _0207200 | _0610400 | _0905200 | _1035800 | _1239300 | _1406000 | _MIT03300 |
| _0207600 | _0610800 | _0906900 | _1037100 | _1240000 | _1406100 | |
| _0209400 | _0612100 | _0908000 | _1037500 | _1242100 | _1406200 | |
| _0209500 | _0614300 | _0908100 | _1039500 | _1243300 | _1410200 | |
| _0210600 | _0614600 | _0908200 | _1040151 | _1243800 | _1411200 | |
| _0210800 | _0615900 | _0910500 | _1040521 | _1244500 | _1411300 | |
| _0211000 | _0618300 | _0911800 | _1100441 | _1245400 | _1412200 | |
| _0211700 | _0619400 | _0912400 | _1102600 | _1245700 | _1412600 | |
| _0212200 | _0620200 | _0913000 | _1103100 | _1246500 | _1414000 | |
| _0212700 | _0621200 | _0913700 | _1105400 | _1306800 | _1414100 | |
| _0213600 | _0621800 | _0914500 | _1105500 | _1307100 | _1415000 | |
| _0303700 | _0623700 | _0914800 | _1106800 | _1308000 | _1415500 | |
| _0304000 | _0700011 | _0915200 | _1107200 | _1308600 | _1419200 | |
| _0304400 | _0706000 | _0916300 | _1107800 | _1308900 | _1419600 | |
| _0307200 | _0706300 | _0917500 | _1109100 | _1309750 | _1419700 | |
| _0308800 | _0710200 | _0919600 | _1110350 | _1310200 | _1420400 | |
| _0310900 | _0712300 | _0920600 | _1113900 | _1311900 | _1420600 | |
| _0311300 | _0715400 | _0921900 | _1120000 | _1313800 | _1421200 | |
| _0312900 | _0715700 | _0922000 | _1122100 | _1315500 | _1421300 | |
| _0316961 | _0715800 | _0922300 | _1122200 | _1315900 | _1422500 | |
| _0400200 | _0716100 | _0923900 | _1122900 | _1316000 | _1426300 | |
| _0400600 | _0717200 | _0924000 | _1123900 | _1319800 | _1426700 | |
| _0400700 | _0717900 | _0926000 | _1128000 | _1320000 | _1426800 | |
| _0400800 | _0718400 | _0926100 | _1128300 | _1321300 | _1427900 | |
| _0403400 | _0721200 | _0926800 | _1128500 | _1321700 | _1428200 | |
| _0403800 | _0722000 | _0926900 | _1132100 | _1323100 | _1428400 | |
| _0404800 | _0722600 | _0929000 | _1133900 | _1324200 | _1428500 | |
| _0408400 | _0801000 | _0929500 | _1134300 | _1325000 | _1429500 | |
| _0408600 | _0802300 | _0931600 | _1134700 | _1326000 | _1431050 | |
| _0408700 | _0805200 | _0934000 | _1137000 | _1327041 | _1432000 | |
| _0408800 | _0806900 | _0935300 | _1137500 | _1327500 | _1432900 | |
| _0413200 | _0808900 | _0937300 | _1140000 | _1327600 | _1433500 | |
| _0416500 | _0810100 | _0942700 | _1141800 | _1328400 | _1436400 | |
| _0417700 | _0811400 | _0943300 | _1144100 | _1330200 | _1437400 | |
| _0502600 | _0811900 | _1000011 | _1146600 | _1331400 | _1440000 | |
| _0503000 | _0813400 | _1001000 | _1200096 | _1331500 | _1440500 | |
| _0503100 | _0813500 | _1001800 | _1200900 | _1333300 | _1441000 | |
| _0503200 | _0815600 | _1005900 | _1202500 | _1334400 | _1442200 | |
| _0505800 | _0816300 | _1007300 | _1203700 | _1335100 | _1442600 | |
| _0506100 | _0817800 | _1008300 | _1205600 | _1337200 | _1443300 | |
| _0506800 | _0818100 | _1011200 | _1205900 | _1337500 | _1443400 | |
| _0508200 | _0819200 | _1012900 | _1208300 | _1339000 | _1443500 | |
| _0509100 | _0819450 | _1015300 | _1210100 | _1339500 | _1443600 | |
| _0511000 | _0822600 | _1015700 | _1211800 | _1343500 | _1445300 | |
| _0511400 | _0823100 | _1016300 | _1214300 | _1344000 | _1446500 | |
| _0511600 | _0823700 | _1016400 | _1215500 | _1344800 | _1446600 | |
| _0512100 | _0824600 | _1017100 | _1216600 | _1344900 | _1448200 | |
