## Supplementary material for "Transcriptome analysis of *Plasmodium berghei* during exo-erythrocytic development": Table S5

**Preferentially expressed genes in developing erythrocytic stage**
(significantly **down-regulated** in
developing exo-erythrocytic compared to erythrocytic stage)

| GeneID | logBaseMean | logFC | pVal | adjP |
|---|---|---|---|---|
| PBANKA_1300200 | *6.2* | *-9.79* | 5.16e-08 | 2.37E-07 |
| PBANKA_0500950 | *5.72* | *-9.17* | 6.21e-29 | 1.90E-27 |
| PBANKA_0722941 | *5.46* | *-9.11* | 4.09e-11 | 2.73E-10 |
| PBANKA_0500721 | *4.4* | *-8.21* | 1.37e-21 | 2.48E-20 |
| PBANKA_1245920 | *5.38* | *-8.19* | 1.83e-24 | 4.02E-23 |
| PBANKA_1040581 | *7.03* | *-8.02* | 3.06e-32 | 1.13E-30 |
| PBANKA_0836400 | *4.83* | *-7.87* | 1.04e-03 | 2.40E-03 |
| PBANKA_1000081 | *5.35* | *-7.82* | 1.17e-21 | 2.15E-20 |
| PBANKA_0700561 | *5.6* | *-7.77* | 1.29e-22 | 2.56E-21 |
| PBANKA_1246300 | *4.49* | *-7.66* | 6.42e-17 | 7.38E-16 |
| PBANKA_1040561 | *5.26* | *-7.4* | 4.26e-09 | 2.25E-08 |
| PBANKA_1200400 | *4.81* | *-7.33* | 3.01e-17 | 3.59E-16 |
| PBANKA_0500741 | *6.31* | *-7.17* | 1.68e-07 | 7.18E-07 |
| PBANKA_0500700 | *8.49* | *-7.12* | 3.74e-26 | 9.37E-25 |
| PBANKA_0201000 | *5.24* | *-6.97* | 1.60e-23 | 3.31E-22 |
| PBANKA_0500971 | *4.58* | *-6.92* | 1.56e-14 | 1.46E-13 |
| PBANKA_1145800 | *11.94* | *-6.87* | 6.97e-114 | 1.98E-111 |
| PBANKA_0316400 | *5.43* | *-6.85* | 2.21e-09 | 1.21E-08 |
| PBANKA_0316921 | *7.78* | *-6.84* | 3.06e-25 | 7.11E-24 |
| PBANKA_1400051 | *5.98* | *-6.84* | 1.16e-19 | 1.79E-18 |
| PBANKA_0722900 | *3.01* | *-6.64* | 6.59e-04 | 1.59E-03 |
| PBANKA_1146741 | *3.59* | *-6.62* | 9.67e-09 | 4.89E-08 |
| PBANKA_1421700 | *8.84* | *-6.61* | 1.03e-06 | 3.99E-06 |
| PBANKA_0600061 | *6.24* | *-6.46* | 3.20e-20 | 5.16E-19 |
| PBANKA_0700541 | *3.29* | *-6.46* | 6.13e-05 | 1.77E-04 |
| PBANKA_1146721 | *5.16* | *-6.45* | 5.22e-15 | 5.05E-14 |
| PBANKA_0700600 | *8.8* | *-6.42* | 6.19e-43 | 4.11E-41 |
| PBANKA_1246400 | *4.75* | *-6.41* | 7.98e-13 | 6.42E-12 |
| PBANKA_1365550 | *3.87* | *-6.39* | 2.53e-07 | 1.06E-06 |
| PBANKA_1000061 | *2.85* | *-6.38* | 4.67e-03 | 9.38E-03 |
| PBANKA_1000041 | *5.18* | *-6.32* | 3.32e-15 | 3.29E-14 |
| PBANKA_1365700 | *3.97* | *-6.29* | 1.42e-08 | 7.03E-08 |
| PBANKA_0917980 | *5.61* | *-6.29* | 9.60e-06 | 3.21E-05 |
| PBANKA_1000400 | *2.88* | *-6.27* | 8.81e-04 | 2.06E-03 |
| PBANKA_1246200 | *3.16* | *-6.25* | 2.64e-04 | 6.85E-04 |
| PBANKA_0110800 | *7.89* | *-6.24* | 1.71e-09 | 9.56E-09 |
| PBANKA_1400200 | *3.91* | *-6.22* | 7.05e-07 | 2.79E-06 |
| PBANKA_1428980 | *5.38* | *-6.18* | 6.14e-05 | 1.78E-04 |
| PBANKA_0001001 | *5.19* | *-6.13* | 2.60e-13 | 2.19E-12 |
| PBANKA_1000091 | *3.43* | *-6.11* | 2.93e-05 | 9.02E-05 |
| PBANKA_1000071 | *4.34* | *-6.09* | 4.58e-04 | 1.13E-03 |
| PBANKA_1400071 | *3.92* | *-6.05* | 2.43e-07 | 1.01E-06 |
| PBANKA_1419300 | *10.21* | *-6.02* | 6.61e-21 | 1.12E-19 |
| PBANKA_0836800 | *7.17* | *-5.98* | 2.01e-39 | 1.06E-37 |
| PBANKA_0707100 | *11.8* | *-5.95* | 3.97e-19 | 5.73E-18 |
| PBANKA_0216061 | *2.85* | *-5.95* | 4.13e-03 | 8.40E-03 |
| PBANKA_0944101 | *5.71* | *-5.92* | 1.22e-12 | 9.55E-12 |
| PBANKA_1432200 | *9.21* | *-5.91* | 9.30e-09 | 4.71E-08 |
| PBANKA_0837201 | *3.22* | *-5.91* | 1.03e-03 | 2.39E-03 |
| PBANKA_1220200 | *10.74* | *-5.9* | 1.17e-32 | 4.54E-31 |
| PBANKA_0500781 | *6.94* | *-5.88* | 2.49e-32 | 9.31E-31 |
| PBANKA_1145900 | *12.18* | *-5.87* | 7.79e-117 | 2.49E-114 |
| PBANKA_0944000 | *2.93* | *-5.82* | 2.41e-03 | 5.19E-03 |
| PBANKA_0300500 | *3.29* | *-5.79* | 2.91e-04 | 7.45E-04 |
| PBANKA_0600051 | *6.03* | *-5.78* | 5.01e-20 | 7.91E-19 |
| PBANKA_0623651 | *4.69* | *-5.76* | 1.64e-10 | 1.03E-09 |
| PBANKA_0112721 | *8.36* | *-5.67* | 8.02e-16 | 8.40E-15 |
| PBANKA_0514900 | *13.53* | *-5.64* | 7.61e-22 | 1.42E-20 |
| PBANKA_1237800 | *11.1* | *-5.62* | 8.95e-58 | 1.17E-55 |
| PBANKA_1444800 | *10.13* | *-5.6* | 2.75e-18 | 3.62E-17 |
| PBANKA_1400400 | *5.98* | *-5.59* | 2.73e-04 | 7.04E-04 |
| PBANKA_1365680 | *4.03* | *-5.56* | 6.65e-09 | 3.41E-08 |
| PBANKA_0105100 | *7.01* | *-5.55* | 1.59e-12 | 1.22E-11 |
| PBANKA_1364250 | *4.21* | *-5.55* | 2.16e-07 | 9.07E-07 |
| PBANKA_1000600 | *11.69* | *-5.5* | 1.90e-27 | 5.19E-26 |
| PBANKA_0216041 | *5.86* | *-5.47* | 1.81e-07 | 7.70E-07 |
| PBANKA_0700200 | *4.03* | *-5.46* | 4.31e-07 | 1.75E-06 |
| PBANKA_0700071 | *5.14* | *-5.45* | 2.51e-09 | 1.36E-08 |
| PBANKA_1000051 | *6.25* | *-5.42* | 3.52e-15 | 3.48E-14 |

**Preferentially expressed genes in developing exo-erythrocytic stage**
(significantly **up-regulated** in
developing exo-erythrocytic compared to erythrocytic stage)

| GeneID | logBaseMean | logFC | pVal | adjP |
|---|---|---|---|---|
| PBANKA_1024600 | *12.26* | *9.71* | 9.99E-500 | 0.00E+00 |
| PBANKA_1465051 | *12.02* | *8.74* | 5.91E-99 | 1.37E-96 |
| PBANKA_1125100 | *9.22* | *8.18* | 1.76e-153 | 1.13E-150 |
| PBANKA_1003000 | *12.98* | *7.83* | 0.00e+00 | 0.00E+00 |
| PBANKA_1002300 | *9.73* | *7.55* | 5.66e-184 | 4.82E-181 |
| PBANKA_1462600 | *9.1* | *7.52* | 4.57e-190 | 4.67E-187 |
| PBANKA_0518900 | *12.82* | *7.3* | 5.45e-222 | 9.30E-219 |
| PBANKA_0501200 | *15.87* | *7.3* | 2.93e-147 | 1.66E-144 |
| PBANKA_1002200 | *8.71* | *7.2* | 6.41e-11 | 4.19E-10 |
| PBANKA_1003900 | *10.51* | *6.95* | 2.30e-109 | 5.89E-107 |
| PBANKA_0941800 | *13.63* | *6.49* | 9.05e-120 | 3.31E-117 |
| PBANKA_1456700 | *6.79* | *6.33* | 2.58e-12 | 1.95E-11 |
| PBANKA_0505000 | *8.01* | *5.95* | 1.69e-146 | 8.64E-144 |
| PBANKA_0818300 | *5.92* | *5.59* | 5.47e-12 | 3.99E-11 |
| PBANKA_0110300 | *5.15* | *5.58* | 2.65e-05 | 8.22E-05 |
| PBANKA_0823800 | *7.21* | *5.53* | 3.29e-65 | 5.42E-63 |
| PBANKA_1411000 | *9.04* | *5.52* | 7.38e-36 | 3.28E-34 |
| PBANKA_1363700 | *8.93* | *5.34* | 2.74e-194 | 3.51E-191 |
| PBANKA_0820900 | *7.24* | *5.23* | 9.92e-62 | 1.49E-59 |
| PBANKA_0923800 | *7.89* | *5.01* | 1.15e-153 | 8.43E-151 |
| PBANKA_0308200 | *6.06* | *5* | 7.32e-40 | 3.94E-38 |
| PBANKA_1338200 | *7.33* | *4.9* | 2.70e-34 | 1.12E-32 |
| PBANKA_1128100 | *11.95* | *4.86* | 1.32e-140 | 5.63E-138 |
| PBANKA_0902100 | *11.28* | *4.81* | 8.67e-117 | 2.61E-114 |
| PBANKA_1344400 | *9.53* | *4.72* | 8.04e-53 | 8.57E-51 |
| PBANKA_0412400 | *9.15* | *4.69* | 6.04e-72 | 1.14E-69 |
| PBANKA_0501600 | *9.03* | *4.68* | 2.70e-129 | 1.06E-126 |
| PBANKA_0519300 | *9.76* | *4.61* | 4.84e-54 | 5.26E-52 |
| PBANKA_0209100 | *9.61* | *4.6* | 4.70e-34 | 1.94E-32 |
| PBANKA_1332800 | *10.76* | *4.56* | 1.99e-45 | 1.54E-43 |
| PBANKA_1435900 | *8.76* | *4.51* | 2.44e-40 | 1.37E-38 |
| PBANKA_0519500 | *11.91* | *4.46* | 5.81e-37 | 2.75E-35 |
| PBANKA_0908600 | *8.09* | *4.37* | 1.56e-22 | 3.08E-21 |
| PBANKA_MIT03500 | *12.59* | *4.31* | 6.69e-144 | 3.11E-141 |
| PBANKA_1340900 | *9.02* | *4.31* | 2.29e-73 | 4.69E-71 |
| PBANKA_1411500 | *7.88* | *4.29* | 2.48e-64 | 3.97E-62 |
| PBANKA_0820200 | *10.64* | *4.28* | 3.35e-25 | 7.75E-24 |
| PBANKA_1400096 | *4.04* | *4.25* | 2.46e-04 | 6.41E-04 |
| PBANKA_0902700 | *6.08* | *4.19* | 9.01e-39 | 4.66E-37 |
| PBANKA_1349300 | *12.82* | *4.18* | 2.08e-58 | 2.88E-56 |
| PBANKA_0832100 | *8.67* | *4.14* | 6.64e-117 | 2.27E-114 |
| PBANKA_0941900 | *12.63* | *3.98* | 2.51e-72 | 4.94E-70 |
| PBANKA_1229800 | *7.54* | *3.98* | 9.13e-72 | 1.67E-69 |
| PBANKA_1140700 | *7.32* | *3.98* | 3.63e-49 | 3.44E-47 |
| PBANKA_1117000 | *11.91* | *3.97* | 3.07e-37 | 1.47E-35 |
| PBANKA_0305600 | *6.81* | *3.96* | 5.46e-40 | 3.00E-38 |
| PBANKA_1107600 | *10.92* | *3.96* | 9.36e-38 | 4.52E-36 |
| PBANKA_0915100 | *5.9* | *3.96* | 3.47e-06 | 1.24E-05 |
| PBANKA_0932900 | *6.92* | *3.95* | 5.83e-29 | 1.80E-27 |
| PBANKA_0828800 | *7.58* | *3.94* | 6.31e-32 | 2.31E-30 |
| PBANKA_0910900 | *11.53* | *3.93* | 9.33e-06 | 3.13E-05 |
| PBANKA_0714900 | *6.6* | *3.86* | 2.25e-41 | 1.37E-39 |
| PBANKA_0501800 | *8.92* | *3.86* | 1.26e-30 | 4.28E-29 |
| PBANKA_0828300 | *4.06* | *3.86* | 6.24e-04 | 1.51E-03 |
| PBANKA_0400500 | *8.56* | *3.78* | 2.18e-18 | 2.90E-17 |
| PBANKA_1001700 | *5.88* | *3.67* | 1.42e-14 | 1.33E-13 |
| PBANKA_1410500 | *6.39* | *3.66* | 1.85e-18 | 2.49E-17 |
| PBANKA_0403200 | *13.4* | *3.65* | 1.52e-44 | 1.14E-42 |
| PBANKA_1340700 | *6.01* | *3.63* | 1.70e-31 | 6.09E-30 |
| PBANKA_1310100 | *6.51* | *3.57* | 1.03e-46 | 8.82E-45 |
| PBANKA_1110700 | *10.15* | *3.56* | 5.28e-67 | 9.01E-65 |
| PBANKA_1301900 | *10.17* | *3.53* | 3.61e-90 | 8.03E-88 |
| PBANKA_0809500 | *6.77* | *3.48* | 4.84e-27 | 1.30E-25 |
| PBANKA_1421600 | *6.13* | *3.48* | 6.20e-19 | 8.77E-18 |
| PBANKA_0825900 | *10.93* | *3.47* | 8.20e-41 | 4.93E-39 |
| PBANKA_1324800 | *6.48* | *3.46* | 4.81e-40 | 2.67E-38 |
| PBANKA_0402600 | *5.93* | *3.45* | 4.68e-16 | 4.96E-15 |
| PBANKA_1006300 | *11.08* | *3.44* | 4.66e-18 | 5.95E-17 |
| PBANKA_0818200 | *5.97* | *3.43* | 6.12e-14 | 5.47E-13 |

| | | | | | | | | | |
|---|---|---|---|---|---|---|---|---|---|
| PBANKA_1200071 | 3.05 | -5.38 | 4.22e-04 | 1.05E-03 | PBANKA_1340800 | 5.75 | 3.42 | 6.46e-25 | 1.46E-23 |
| PBANKA_1300091 | 3.01 | -5.38 | 2.18e-03 | 4.74E-03 | PBANKA_0817300 | 6.98 | 3.4 | 4.32e-20 | 6.84E-19 |
| PBANKA_0937860 | 3.31 | -5.37 | 2.90e-03 | 6.11E-03 | PBANKA_0522400 | 7.22 | 3.35 | 2.27e-07 | 9.51E-07 |
| PBANKA_1245900 | 8.49 | -5.36 | 1.49e-36 | 6.86E-35 | PBANKA_0101100 | 6.54 | 3.34 | 3.45e-28 | 1.01E-26 |
| PBANKA_0600100 | 3.99 | -5.36 | 1.51e-05 | 4.81E-05 | PBANKA_0831000 | 14 | 3.33 | 1.42e-32 | 5.48E-31 |
| PBANKA_0837141 | 5.96 | -5.35 | 3.34e-13 | 2.78E-12 | PBANKA_1416700 | 6.34 | 3.32 | 1.15e-26 | 3.03E-25 |
| PBANKA_0701100 | 8.71 | -5.33 | 9.34e-36 | 4.11E-34 | PBANKA_1449500 | 10.86 | 3.31 | 1.57e-24 | 3.48E-23 |
| PBANKA_0300300 | 4.05 | -5.32 | 3.37e-06 | 1.21E-05 | PBANKA_1449800 | 8.25 | 3.3 | 2.64e-36 | 1.19E-34 |
| PBANKA_1436300 | 6.81 | -5.28 | 1.22e-13 | 1.05E-12 | PBANKA_1306500 | 5.88 | 3.3 | 1.74e-13 | 1.48E-12 |
| PBANKA_1466121 | 4.42 | -5.27 | 4.30e-10 | 2.58E-09 | PBANKA_0911000 | 6.12 | 3.28 | 1.09e-16 | 1.23E-15 |
| PBANKA_0316800 | 7.56 | -5.26 | 1.18e-33 | 4.81E-32 | PBANKA_1014000 | 8.77 | 3.27 | 3.52e-58 | 4.74E-56 |
| PBANKA_1246161 | 10.34 | -5.25 | 2.01e-28 | 5.93E-27 | PBANKA_1416800 | 9.08 | 3.27 | 4.45e-27 | 1.20E-25 |
| PBANKA_0317121 | 4.23 | -5.22 | 2.47e-09 | 1.34E-08 | PBANKA_0507600 | 5.95 | 3.26 | 6.20e-16 | 6.52E-15 |
| PBANKA_0100021 | 8.43 | -5.22 | 1.30e-07 | 5.63E-07 | PBANKA_0907200 | 8.02 | 3.25 | 4.06e-31 | 1.41E-29 |
| PBANKA_0800200 | 3.23 | -5.18 | 1.31e-03 | 2.96E-03 | PBANKA_1343500 | 8.67 | 3.24 | 1.02e-54 | 1.13E-52 |
| PBANKA_1455800 | 6.19 | -5.16 | 3.53e-24 | 7.59E-23 | PBANKA_1228300 | 12.63 | 3.23 | 1.32e-12 | 1.03E-11 |
| PBANKA_0112701 | 6.41 | -5.15 | 2.99e-26 | 7.62E-25 | PBANKA_0929200 | 7.75 | 3.22 | 1.34e-19 | 2.03E-18 |
| PBANKA_0836600 | 6.15 | -5.15 | 8.27e-13 | 6.62E-12 | PBANKA_1302000 | 8.9 | 3.21 | 8.24e-35 | 3.51E-33 |
| PBANKA_0826700 | 10.81 | -5.13 | 3.77e-109 | 9.18E-107 | PBANKA_0306000 | 8.99 | 3.19 | 1.85e-10 | 1.15E-09 |
| PBANKA_0500600 | 6.81 | -5.13 | 8.36e-14 | 7.37E-13 | PBANKA_1002400 | 9.5 | 3.17 | 3.47e-47 | 3.06E-45 |
| PBANKA_1146700 | 4.28 | -5.12 | 1.92e-06 | 7.21E-06 | PBANKA_0622900 | 6.91 | 3.17 | 1.03e-03 | 2.38E-03 |
| PBANKA_0931200 | 11.5 | -5.1 | 1.25e-31 | 4.50E-30 | PBANKA_0511000 | 7.56 | 3.16 | 1.10e-43 | 7.69E-42 |
| PBANKA_0316500 | 9.14 | -5.09 | 1.12e-08 | 5.59E-08 | PBANKA_0501700 | 7.01 | 3.16 | 3.00e-25 | 7.01E-24 |
| PBANKA_0504400 | 10.5 | -5.04 | 7.25e-20 | 1.13E-18 | PBANKA_0812300 | 7.66 | 3.16 | 1.75e-24 | 3.85E-23 |
| PBANKA_0944061 | 4.04 | -5.04 | 2.36e-07 | 9.87E-07 | PBANKA_0105700 | 6.14 | 3.16 | 3.83e-15 | 3.77E-14 |
| PBANKA_0600400 | 5.47 | -4.98 | 5.07e-18 | 6.43E-17 | PBANKA_1212400 | 6.37 | 3.15 | 4.36e-07 | 1.77E-06 |
| PBANKA_0216721 | 4.02 | -4.98 | 1.99e-06 | 7.45E-06 | PBANKA_1441000 | 5.08 | 3.14 | 1.47e-08 | 7.24E-08 |
| PBANKA_1146000 | 7.96 | -4.96 | 3.59e-48 | 3.23E-46 | PBANKA_0210100 | 6.31 | 3.09 | 4.22e-25 | 9.62E-24 |
| PBANKA_1204500 | 6.73 | -4.96 | 4.56e-05 | 1.36E-04 | PBANKA_1024400 | 9.39 | 3.06 | 1.51e-43 | 1.05E-41 |
| PBANKA_0109800 | 8.11 | -4.94 | 9.87e-10 | 5.65E-09 | PBANKA_1138000 | 8.47 | 3.06 | 2.67e-27 | 7.24E-26 |
| PBANKA_0600086 | 3.82 | -4.91 | 6.14e-06 | 2.12E-05 | PBANKA_1106100 | 10.72 | 3.01 | 1.38e-18 | 1.89E-17 |
| PBANKA_0836900 | 3.31 | -4.9 | 3.67e-03 | 7.58E-03 | PBANKA_1125200 | 7.95 | 3 | 2.54e-43 | 1.71E-41 |
| PBANKA_0917990 | 3.64 | -4.86 | 3.10e-03 | 6.50E-03 | PBANKA_0208900 | 10.15 | 3 | 1.67e-11 | 1.17E-10 |
| PBANKA_0700081 | 6.75 | -4.85 | 3.80e-11 | 2.56E-10 | PBANKA_1324200 | 7.88 | 2.99 | 1.25e-56 | 1.56E-54 |
| PBANKA_0300200 | 2.81 | -4.81 | 2.24e-03 | 4.86E-03 | PBANKA_1217400 | 9.06 | 2.97 | 1.61e-49 | 1.55E-47 |
| PBANKA_1431400 | 6.29 | -4.76 | 2.62e-11 | 1.81E-10 | PBANKA_0208400 | 8.15 | 2.97 | 6.25e-33 | 2.48E-31 |
| PBANKA_1000200 | 3.76 | -4.76 | 1.10e-05 | 3.64E-05 | PBANKA_0820400 | 8.26 | 2.95 | 1.35e-30 | 4.59E-29 |
| PBANKA_1040541 | 3.61 | -4.72 | 1.30e-03 | 2.95E-03 | PBANKA_1455000 | 8.31 | 2.94 | 5.55e-27 | 1.48E-25 |
| PBANKA_1431500 | 7.2 | -4.71 | 9.53e-04 | 2.22E-03 | PBANKA_0720500 | 6.49 | 2.9 | 2.10e-16 | 2.31E-15 |
| PBANKA_0300600 | 12.56 | -4.7 | 6.28e-44 | 4.53E-42 | PBANKA_0607000 | 7.78 | 2.89 | 1.12e-38 | 5.71E-37 |
| PBANKA_0836200 | 11.12 | -4.69 | 2.26e-24 | 4.90E-23 | PBANKA_0711400 | 9.87 | 2.89 | 8.31e-15 | 7.91E-14 |
| PBANKA_0214550 | 5.43 | -4.69 | 2.68e-18 | 3.53E-17 | PBANKA_1107700 | 10.68 | 2.88 | 5.24e-38 | 2.55E-36 |
| PBANKA_1030600 | 9.41 | -4.68 | 1.72e-51 | 1.76E-49 | PBANKA_0305100 | 13.08 | 2.88 | 4.56e-21 | 7.90E-20 |
| PBANKA_0837101 | 3.81 | -4.67 | 4.16e-05 | 1.25E-04 | PBANKA_1413800 | 7.6 | 2.87 | 3.28e-35 | 1.41E-33 |
| PBANKA_0600081 | 3.65 | -4.65 | 7.96e-06 | 2.70E-05 | PBANKA_0108800 | 9.39 | 2.87 | 6.09e-18 | 7.62E-17 |
| PBANKA_1400031 | 5.21 | -4.62 | 1.12e-07 | 4.90E-07 | PBANKA_1304000 | 8.76 | 2.86 | 1.92e-63 | 2.97E-61 |
| PBANKA_1427700 | 4.34 | -4.62 | 9.59e-07 | 3.72E-06 | PBANKA_1136000 | 5.27 | 2.86 | 9.04e-07 | 3.52E-06 |
| PBANKA_0501100 | 7.37 | -4.61 | 2.77e-25 | 6.49E-24 | PBANKA_0206500 | 8.02 | 2.85 | 3.39e-25 | 7.77E-24 |
| PBANKA_1463300 | 11.43 | -4.57 | 3.75e-24 | 8.02E-23 | PBANKA_0209300 | 8.83 | 2.84 | 2.10e-20 | 3.47E-19 |
| PBANKA_0301400 | 6.03 | -4.56 | 2.03e-15 | 2.05E-14 | PBANKA_1119900 | 7.21 | 2.82 | 4.72e-15 | 4.58E-14 |
| PBANKA_0600200 | 3.81 | -4.55 | 2.31e-05 | 7.19E-05 | PBANKA_1419900 | 5.79 | 2.8 | 1.77e-11 | 1.24E-10 |
| PBANKA_1018200 | 4.54 | -4.54 | 1.51e-06 | 5.77E-06 | PBANKA_1211800 | 8.51 | 2.79 | 1.11e-40 | 6.50E-39 |
| PBANKA_0605800 | 6.93 | -4.44 | 1.77e-09 | 9.85E-09 | PBANKA_1033100 | 9.13 | 2.79 | 2.63e-28 | 7.72E-27 |
| PBANKA_1146200 | 3.55 | -4.44 | 1.28e-03 | 2.91E-03 | PBANKA_0815800 | 6.43 | 2.79 | 3.34e-20 | 5.36E-19 |
| PBANKA_1365500 | 11.6 | -4.43 | 1.15e-67 | 2.02E-65 | PBANKA_0706600 | 10.48 | 2.79 | 1.25e-12 | 9.76E-12 |
| PBANKA_0600300 | 3.65 | -4.39 | 2.63e-04 | 6.83E-04 | PBANKA_1351400 | 8.18 | 2.77 | 5.41e-28 | 1.55E-26 |
| PBANKA_0606200 | 8.71 | -4.34 | 6.26e-09 | 3.23E-08 | PBANKA_0309500 | 8.38 | 2.77 | 1.45e-16 | 1.62E-15 |
| PBANKA_1334900 | 10.06 | -4.33 | 3.51e-19 | 5.09E-18 | PBANKA_0506900 | 8.49 | 2.76 | 3.90e-26 | 9.74E-25 |
| PBANKA_0602900 | 4.37 | -4.33 | 4.76e-05 | 1.41E-04 | PBANKA_0207600 | 9.86 | 2.75 | 8.89e-46 | 7.00E-44 |
| PBANKA_1436600 | 10.57 | -4.31 | 1.90e-15 | 1.92E-14 | PBANKA_0402200 | 5.54 | 2.75 | 7.51e-14 | 6.65E-13 |
| PBANKA_1132800 | 5.23 | -4.31 | 2.55e-07 | 1.06E-06 | PBANKA_0208300 | 9.17 | 2.73 | 1.92e-31 | 6.82E-30 |
| PBANKA_1465500 | 3.42 | -4.31 | 3.00e-04 | 7.68E-04 | PBANKA_1350400 | 5.75 | 2.73 | 2.46e-13 | 2.08E-12 |
| PBANKA_0804300 | 5.72 | -4.16 | 8.49e-10 | 4.88E-09 | PBANKA_MIT01800 | 8.9 | 2.72 | 2.09e-25 | 4.98E-24 |
| PBANKA_0400100 | 5.22 | -4.16 | 9.27e-10 | 5.32E-09 | PBANKA_1212000 | 9.65 | 2.72 | 1.86e-21 | 3.34E-20 |
| PBANKA_0608600 | 11.75 | -4.15 | 3.33e-23 | 6.81E-22 | PBANKA_1439400 | 9.05 | 2.71 | 4.85e-31 | 1.68E-29 |
| PBANKA_0216741 | 5.65 | -4.14 | 1.52e-13 | 1.30E-12 | PBANKA_0819600 | 8.35 | 2.71 | 9.53e-16 | 9.90E-15 |
| PBANKA_1400700 | 11.29 | -4.12 | 4.44e-19 | 6.38E-18 | PBANKA_1361800 | 6.1 | 2.71 | 8.48e-10 | 4.88E-09 |
| PBANKA_1335000 | 8.13 | -4.12 | 4.96e-08 | 2.28E-07 | PBANKA_0511100 | 7.5 | 2.69 | 6.75e-24 | 1.42E-22 |
| PBANKA_1000500 | 9.17 | -4.11 | 2.33e-24 | 5.02E-23 | PBANKA_0104100 | 7.57 | 2.69 | 3.90e-23 | 7.92E-22 |
| PBANKA_1102200 | 11.83 | -4.09 | 1.16e-88 | 2.48E-86 | PBANKA_1346900 | 6.51 | 2.69 | 1.25e-19 | 1.91E-18 |
| PBANKA_1204200 | 10.51 | -4.09 | 2.46e-19 | 3.64E-18 | PBANKA_1459200 | 6.28 | 2.69 | 5.69e-18 | 7.20E-17 |
| PBANKA_0801400 | 6.86 | -4.08 | 2.24e-31 | 7.89E-30 | PBANKA_1336900 | 8.4 | 2.68 | 4.11e-49 | 3.75E-47 |
| PBANKA_0941000 | 5.34 | -4.08 | 3.08e-06 | 1.12E-05 | PBANKA_1343400 | 10.61 | 2.68 | 6.32e-27 | 1.67E-25 |
| PBANKA_0800300 | 8.55 | -4.07 | 5.22e-20 | 8.16E-19 | PBANKA_1303500 | 9.16 | 2.68 | 1.18e-21 | 2.15E-20 |

| | | | | |
|---|---|---|---|---|
| PBANKA_0524200 | 10.77 | -4.07 | 3.04e-17 | 3.62E-16 |
| PBANKA_0704800 | 10.79 | -4.07 | 9.31e-16 | 9.70E-15 |
| PBANKA_1101300 | 11.72 | -4.05 | 5.33e-50 | 5.34E-48 |
| PBANKA_0509000 | 10.35 | -4.04 | 3.70e-112 | 9.95E-110 |
| PBANKA_1246700 | 3.63 | -4.04 | 9.58e-04 | 2.23E-03 |
| PBANKA_0215800 | 5.38 | -4.03 | 1.51e-12 | 1.17E-11 |
| PBANKA_1333700 | 9.56 | -4.02 | 4.48e-15 | 4.36E-14 |
| PBANKA_1455700 | 5.25 | -4.01 | 4.27e-09 | 2.26E-08 |
| PBANKA_1363600 | 9.43 | -4.01 | 9.03e-05 | 2.54E-04 |
| PBANKA_0837001 | 5.92 | -4.01 | 1.83e-03 | 4.04E-03 |
| PBANKA_0927600 | 10.94 | -4 | 2.95e-19 | 4.33E-18 |
| PBANKA_0516000 | 6.03 | -3.97 | 1.48e-03 | 3.32E-03 |
| PBANKA_0700500 | 3.69 | -3.97 | 3.06e-03 | 6.43E-03 |
| PBANKA_0800500 | 15.03 | -3.9 | 3.28e-15 | 3.25E-14 |
| PBANKA_0316300 | 9.32 | -3.89 | 1.12e-19 | 1.73E-18 |
| PBANKA_0836500 | 5.46 | -3.89 | 1.09e-09 | 6.23E-09 |
| PBANKA_0216801 | 5.63 | -3.87 | 5.70e-09 | 2.96E-08 |
| PBANKA_1225500 | 9.44 | -3.87 | 1.19e-05 | 3.92E-05 |
| PBANKA_0112641 | 6.14 | -3.85 | 5.69e-19 | 8.09E-18 |
| PBANKA_0600021 | 7.53 | -3.85 | 5.89e-06 | 2.04E-05 |
| PBANKA_1000300 | 7.17 | -3.84 | 9.27e-18 | 1.14E-16 |
| PBANKA_1342300 | 7.98 | -3.83 | 3.04e-06 | 1.10E-05 |
| PBANKA_1019000 | 5.62 | -3.82 | 1.21e-05 | 3.96E-05 |
| PBANKA_1465921 | 3.18 | -3.82 | 2.48e-03 | 5.31E-03 |
| PBANKA_1129600 | 7.61 | -3.81 | 1.86e-08 | 9.02E-08 |
| PBANKA_0201500 | 9.92 | -3.8 | 1.56e-61 | 2.28E-59 |
| PBANKA_1414500 | 8.2 | -3.8 | 1.42e-07 | 6.13E-07 |
| PBANKA_0524300 | 11.19 | -3.76 | 2.43e-55 | 2.83E-53 |
| PBANKA_0100500 | 9.26 | -3.76 | 9.65e-19 | 1.35E-17 |
| PBANKA_0622921 | 12.39 | -3.74 | 3.43e-10 | 2.08E-09 |
| PBANKA_0104200 | 5.29 | -3.73 | 3.65e-07 | 1.50E-06 |
| PBANKA_0112661 | 8.01 | -3.73 | 1.07e-06 | 4.14E-06 |
| PBANKA_0700051 | 4.61 | -3.73 | 9.96e-06 | 3.32E-05 |
| PBANKA_0524100 | 8.33 | -3.7 | 1.80e-16 | 1.98E-15 |
| PBANKA_1123200 | 6.71 | -3.7 | 3.81e-10 | 2.30E-09 |
| PBANKA_0100300 | 6.18 | -3.7 | 7.06e-10 | 4.11E-09 |
| PBANKA_1315300 | 10.46 | -3.69 | 5.19e-22 | 9.79E-21 |
| PBANKA_1233600 | 8.44 | -3.69 | 3.41e-03 | 7.08E-03 |
| PBANKA_0800400 | 9.13 | -3.68 | 5.66e-46 | 4.52E-44 |
| PBANKA_1334700 | 8.79 | -3.68 | 1.07e-10 | 6.86E-10 |
| PBANKA_0623200 | 10.89 | -3.67 | 3.58e-36 | 1.61E-34 |
| PBANKA_1030400 | 6.72 | -3.66 | 4.68e-13 | 3.83E-12 |
| PBANKA_0722600 | 6.35 | -3.63 | 1.89e-04 | 5.00E-04 |
| PBANKA_0301000 | 9.96 | -3.62 | 1.36e-18 | 1.87E-17 |
| PBANKA_0517600 | 9.46 | -3.62 | 2.43e-07 | 1.01E-06 |
| PBANKA_1246100 | 5.98 | -3.62 | 9.39e-06 | 3.15E-05 |
| PBANKA_1007900 | 8.64 | -3.59 | 2.14e-08 | 1.03E-07 |
| PBANKA_1466000 | 5.35 | -3.5 | 1.53e-08 | 7.54E-08 |
| PBANKA_1132300 | 9.59 | -3.49 | 4.24e-12 | 3.13E-11 |
| PBANKA_0600351 | 7.37 | -3.46 | 2.73e-05 | 8.45E-05 |
| PBANKA_0103200 | 7.68 | -3.45 | 4.53e-03 | 9.11E-03 |
| PBANKA_1333100 | 6.62 | -3.44 | 1.13e-11 | 8.08E-11 |
| PBANKA_0704900 | 9.81 | -3.42 | 2.24e-03 | 4.85E-03 |
| PBANKA_1101200 | 11.3 | -3.41 | 8.36e-30 | 2.74E-28 |
| PBANKA_1210800 | 9.56 | -3.4 | 3.06e-18 | 3.96E-17 |
| PBANKA_0507900 | 4.64 | -3.4 | 3.31e-05 | 1.01E-04 |
| PBANKA_1019520 | 3.96 | -3.4 | 1.12e-03 | 2.57E-03 |
| PBANKA_0524800 | 9.46 | -3.37 | 6.13e-18 | 7.65E-17 |
| PBANKA_0600091 | 5.13 | -3.36 | 1.18e-06 | 4.55E-06 |
| PBANKA_1203500 | 4.28 | -3.35 | 2.21e-04 | 5.82E-04 |
| PBANKA_0522200 | 7.57 | -3.34 | 1.02e-04 | 2.84E-04 |
| PBANKA_0623150 | 8.27 | -3.3 | 6.78e-12 | 4.91E-11 |
| PBANKA_0417200 | 9.9 | -3.3 | 3.91e-04 | 9.81E-04 |
| PBANKA_1300011 | 4.76 | -3.27 | 9.28e-04 | 2.16E-03 |
| PBANKA_0008101 | 7.83 | -3.26 | 1.16e-05 | 3.82E-05 |
| PBANKA_0910000 | 9.44 | -3.25 | 5.91e-18 | 7.43E-17 |
| PBANKA_1329000 | 5.76 | -3.25 | 2.50e-08 | 1.19E-07 |
| PBANKA_0701000 | 10.33 | -3.24 | 7.52e-31 | 2.58E-29 |
| PBANKA_0600041 | 6.51 | -3.24 | 1.30e-07 | 5.61E-07 |
| PBANKA_0501000 | 5.84 | -3.21 | 5.50e-06 | 1.91E-05 |
| PBANKA_1244500 | 5.95 | -3.2 | 1.01e-22 | 2.03E-21 |
| PBANKA_0907900 | 10.88 | -3.2 | 4.34e-12 | 3.21E-11 |
| PBANKA_0932400 | 9.84 | -3.19 | 4.15e-47 | 3.59E-45 |
| PBANKA_0404300 | 6.47 | -3.19 | 1.86e-10 | 1.16E-09 |

| | | | | |
|---|---|---|---|---|
| PBANKA_0507100 | 6.58 | 2.66 | 1.65e-10 | 1.03E-09 |
| PBANKA_1127200 | 4.84 | 2.65 | 1.23e-07 | 5.33E-07 |
| PBANKA_0933700 | 7.55 | 2.64 | 2.11e-40 | 1.20E-38 |
| PBANKA_1452700 | 6.92 | 2.64 | 1.56e-21 | 2.82E-20 |
| PBANKA_0406900 | 9.02 | 2.64 | 1.59e-18 | 2.16E-17 |
| PBANKA_0203650 | 6.78 | 2.64 | 9.58e-13 | 7.61E-12 |
| PBANKA_0519000 | 11.38 | 2.64 | 2.73e-09 | 1.48E-08 |
| PBANKA_1425900 | 9.69 | 2.62 | 3.44e-46 | 2.79E-44 |
| PBANKA_1411100 | 9 | 2.62 | 1.88e-19 | 2.83E-18 |
| PBANKA_1449700 | 8 | 2.62 | 4.43e-12 | 3.26E-11 |
| PBANKA_1346500 | 7.18 | 2.61 | 1.07e-17 | 1.30E-16 |
| PBANKA_0411000 | 5.58 | 2.61 | 7.43e-11 | 4.83E-10 |
| PBANKA_0901700 | 5.7 | 2.6 | 4.38e-09 | 2.31E-08 |
| PBANKA_0710400 | 7.41 | 2.59 | 1.44e-28 | 4.32E-27 |
| PBANKA_1303600 | 7.15 | 2.59 | 2.81e-21 | 4.95E-20 |
| PBANKA_0815700 | 10.41 | 2.58 | 3.17e-57 | 4.05E-55 |
| PBANKA_0417800 | 10.31 | 2.58 | 2.56e-19 | 3.77E-18 |
| PBANKA_0907600 | 6.17 | 2.58 | 5.07e-14 | 4.55E-13 |
| PBANKA_1456800 | 8.65 | 2.57 | 3.59e-22 | 6.86E-21 |
| PBANKA_0311900 | 7.86 | 2.56 | 1.01e-28 | 3.04E-27 |
| PBANKA_1024500 | 6.27 | 2.55 | 2.05e-21 | 3.66E-20 |
| PBANKA_0907800 | 6.7 | 2.54 | 1.55e-18 | 2.12E-17 |
| PBANKA_1113300 | 9.1 | 2.53 | 2.32e-39 | 1.21E-37 |
| PBANKA_0929300 | 8.12 | 2.53 | 3.19e-21 | 5.60E-20 |
| PBANKA_1240600 | 10.95 | 2.51 | 4.05e-15 | 3.97E-14 |
| PBANKA_1409800 | 7.5 | 2.5 | 1.29e-25 | 3.12E-24 |
| PBANKA_1340500 | 9.59 | 2.5 | 1.40e-19 | 2.13E-18 |
| PBANKA_1304800 | 6.29 | 2.5 | 1.59e-13 | 1.36E-12 |
| PBANKA_1137400 | 7.25 | 2.5 | 2.42e-13 | 2.05E-12 |
| PBANKA_1402300 | 9.83 | 2.5 | 1.44e-10 | 9.08E-10 |
| PBANKA_1324700 | 4.89 | 2.48 | 3.91e-06 | 1.39E-05 |
| PBANKA_1122300 | 7.48 | 2.47 | 3.60e-42 | 2.27E-40 |
| PBANKA_1349800 | 13.67 | 2.46 | 1.13e-24 | 2.53E-23 |
| PBANKA_1033300 | 7.32 | 2.46 | 7.85e-20 | 1.22E-18 |
| PBANKA_0417700 | 10.19 | 2.46 | 1.51e-18 | 2.06E-17 |
| PBANKA_1414900 | 6.96 | 2.46 | 6.54e-15 | 6.28E-14 |
| PBANKA_0512900 | 7.73 | 2.46 | 1.08e-12 | 8.52E-12 |
| PBANKA_0404700 | 8.16 | 2.46 | 6.21e-10 | 3.65E-09 |
| PBANKA_1106000 | 6.8 | 2.46 | 1.13e-06 | 4.33E-06 |
| PBANKA_1324300 | 7.13 | 2.45 | 1.27e-15 | 1.31E-14 |
| PBANKA_1242400 | 8.28 | 2.44 | 2.58e-55 | 2.93E-53 |
| PBANKA_0909500 | 7.06 | 2.43 | 1.53e-11 | 1.08E-10 |
| PBANKA_0620200 | 7.72 | 2.43 | 1.80e-09 | 1.00E-08 |
| PBANKA_1006200 | 10.95 | 2.43 | 4.17e-07 | 1.70E-06 |
| PBANKA_0103000 | 5 | 2.43 | 4.25e-07 | 1.73E-06 |
| PBANKA_1241800 | 10.58 | 2.41 | 1.15e-28 | 3.46E-27 |
| PBANKA_0306700 | 8.64 | 2.41 | 4.83e-24 | 1.03E-22 |
| PBANKA_1319600 | 6.05 | 2.41 | 2.74e-10 | 1.67E-09 |
| PBANKA_1420600 | 10.34 | 2.4 | 2.39e-19 | 3.54E-18 |
| PBANKA_1345000 | 6.72 | 2.39 | 2.31e-14 | 2.11E-13 |
| PBANKA_0931100 | 9.19 | 2.39 | 2.59e-06 | 9.49E-06 |
| PBANKA_0519100 | 11.51 | 2.37 | 7.69e-12 | 5.56E-11 |
| PBANKA_1343750 | 8 | 2.37 | 1.85e-10 | 1.15E-09 |
| PBANKA_1457550 | 8.93 | 2.37 | 3.60e-04 | 9.09E-04 |
| PBANKA_0615900 | 4.75 | 2.37 | 4.64e-04 | 1.15E-03 |
| PBANKA_0921200 | 8.33 | 2.37 | 4.42e-03 | 8.91E-03 |
| PBANKA_0828700 | 8.45 | 2.36 | 1.18e-36 | 5.48E-35 |
| PBANKA_1002100 | 9.79 | 2.36 | 3.24e-16 | 3.50E-15 |
| PBANKA_1432700 | 6.48 | 2.36 | 2.77e-11 | 1.90E-10 |
| PBANKA_1306900 | 8.68 | 2.35 | 1.56e-29 | 5.08E-28 |
| PBANKA_0925600 | 7.66 | 2.35 | 2.42e-17 | 2.90E-16 |
| PBANKA_0204600 | 10.11 | 2.35 | 1.53e-09 | 8.56E-09 |
| PBANKA_1360900 | 5.23 | 2.34 | 6.98e-04 | 1.67E-03 |
| PBANKA_0809300 | 7.47 | 2.33 | 4.02e-17 | 4.70E-16 |
| PBANKA_0702700 | 7.15 | 2.33 | 3.41e-09 | 1.82E-08 |
| PBANKA_0213800 | 7.79 | 2.32 | 2.61e-34 | 1.10E-32 |
| PBANKA_0932200 | 10.32 | 2.32 | 5.47e-12 | 3.99E-11 |
| PBANKA_0510500 | 7.42 | 2.31 | 2.56e-33 | 1.03E-31 |
| PBANKA_1406400 | 8.28 | 2.3 | 1.65e-19 | 2.49E-18 |
| PBANKA_0510700 | 8.01 | 2.3 | 9.31e-09 | 4.71E-08 |
| PBANKA_1236400 | 8.77 | 2.29 | 6.83e-29 | 2.08E-27 |
| PBANKA_0404400 | 7.79 | 2.28 | 4.73e-11 | 3.13E-10 |
| PBANKA_1146261 | 5.19 | 2.28 | 1.13e-05 | 3.74E-05 |
| PBANKA_1423100 | 9.11 | 2.27 | 1.53e-32 | 5.83E-31 |

| | | | | |
|---|---|---|---|---|
| PBANKA_1326100 | 6.35 | -3.19 | 1.98e-10 | 1.23E-09 |
| PBANKA_0316200 | 15.57 | -3.19 | 8.00e-07 | 3.15E-06 |
| PBANKA_0721100 | 7.35 | -3.18 | 8.85e-13 | 7.07E-12 |
| PBANKA_0515000 | 12.94 | -3.17 | 7.17e-10 | 4.18E-09 |
| PBANKA_1029600 | 7.42 | -3.17 | 1.96e-03 | 4.30E-03 |
| PBANKA_1463000 | 10.43 | -3.16 | 1.21e-05 | 3.96E-05 |
| PBANKA_0214600 | 11.55 | -3.12 | 9.85e-14 | 8.66E-13 |
| PBANKA_0317161 | 5.15 | -3.12 | 4.07e-04 | 1.02E-03 |
| PBANKA_1422900 | 11.07 | -3.11 | 1.37e-07 | 5.91E-07 |
| PBANKA_0831600 | 5.82 | -3.1 | 1.62e-05 | 5.14E-05 |
| PBANKA_0907500 | 4.82 | -3.1 | 7.03e-05 | 2.02E-04 |
| PBANKA_1417700 | 4.15 | -3.09 | 9.48e-06 | 3.17E-05 |
| PBANKA_0200700 | 4.68 | -3.08 | 1.42e-05 | 4.55E-05 |
| PBANKA_1245821 | 9.24 | -3.06 | 1.64e-06 | 6.21E-06 |
| PBANKA_0105600 | 9.67 | -3.05 | 1.57e-16 | 1.75E-15 |
| PBANKA_1400081 | 3.6 | -3.04 | 2.33e-03 | 5.03E-03 |
| PBANKA_1229000 | 12.87 | -3.03 | 7.57e-25 | 1.71E-23 |
| PBANKA_0927700 | 7.47 | -3.03 | 4.14e-05 | 1.24E-04 |
| PBANKA_1246800 | 4.21 | -3.03 | 2.81e-03 | 5.95E-03 |
| PBANKA_0612400 | 11.74 | -3.01 | 1.15e-11 | 8.18E-11 |
| PBANKA_0001201 | 3.38 | -2.98 | 4.08e-03 | 8.31E-03 |
| PBANKA_1120300 | 7.2 | -2.96 | 6.51e-24 | 1.38E-22 |
| PBANKA_0807200 | 9.28 | -2.96 | 4.03e-03 | 8.22E-03 |
| PBANKA_1117400 | 7.68 | -2.95 | 7.69e-30 | 2.54E-28 |
| PBANKA_0823200 | 8.79 | -2.95 | 8.14e-10 | 4.70E-09 |
| PBANKA_1008500 | 12.82 | -2.94 | 1.11e-27 | 3.08E-26 |
| PBANKA_1319500 | 11.16 | -2.92 | 2.48e-20 | 4.07E-19 |
| PBANKA_1437500 | 7.72 | -2.92 | 6.70e-10 | 3.92E-09 |
| PBANKA_0216761 | 7.06 | -2.91 | 2.73e-06 | 9.97E-06 |
| PBANKA_0517200 | 7.22 | -2.88 | 7.29e-40 | 3.94E-38 |
| PBANKA_1210200 | 12.34 | -2.88 | 4.80e-28 | 1.39E-26 |
| PBANKA_0100700 | 12.72 | -2.85 | 1.10e-13 | 9.57E-13 |
| PBANKA_1138600 | 9 | -2.84 | 3.08e-19 | 4.47E-18 |
| PBANKA_1360400 | 7.23 | -2.84 | 2.09e-08 | 1.01E-07 |
| PBANKA_0939300 | 7.36 | -2.82 | 5.94e-26 | 1.47E-24 |
| PBANKA_1009600 | 9.09 | -2.82 | 2.76e-10 | 1.69E-09 |
| PBANKA_1421000 | 6.29 | -2.82 | 5.79e-09 | 3.00E-08 |
| PBANKA_1354900 | 10.36 | -2.82 | 1.57e-08 | 7.72E-08 |
| PBANKA_1000100 | 4.27 | -2.79 | 2.04e-03 | 4.45E-03 |
| PBANKA_1349200 | 10.55 | -2.78 | 3.77e-49 | 3.51E-47 |
| PBANKA_0100600 | 8.5 | -2.78 | 4.92e-09 | 2.58E-08 |
| PBANKA_1022100 | 6.97 | -2.76 | 1.73e-11 | 1.21E-10 |
| PBANKA_0720000 | 7.21 | -2.76 | 3.22e-10 | 1.96E-09 |
| PBANKA_1318500 | 8.61 | -2.76 | 2.61e-04 | 6.78E-04 |
| PBANKA_1429100 | 5.69 | -2.76 | 5.33e-04 | 1.30E-03 |
| PBANKA_1432500 | 10.41 | -2.75 | 1.89e-38 | 9.49E-37 |
| PBANKA_0409500 | 6.23 | -2.74 | 2.79e-10 | 1.70E-09 |
| PBANKA_1325200 | 5.45 | -2.74 | 5.03e-04 | 1.24E-03 |
| PBANKA_0804100 | 11.41 | -2.73 | 2.06e-24 | 4.49E-23 |
| PBANKA_1115000 | 10.47 | -2.71 | 3.00e-16 | 3.26E-15 |
| PBANKA_0836981 | 4.68 | -2.7 | 4.15e-03 | 8.42E-03 |
| PBANKA_1242200 | 7.15 | -2.68 | 6.02e-21 | 1.03E-19 |
| PBANKA_1407650 | 3.94 | -2.67 | 2.50e-03 | 5.36E-03 |
| PBANKA_1429200 | 9.94 | -2.65 | 6.18e-04 | 1.50E-03 |
| PBANKA_1225300 | 7.58 | -2.64 | 8.50e-05 | 2.40E-04 |
| PBANKA_0510200 | 9.06 | -2.63 | 1.19e-40 | 6.90E-39 |
| PBANKA_1100900 | 7.84 | -2.63 | 2.26e-09 | 1.24E-08 |
| PBANKA_0214700 | 10.5 | -2.62 | 1.21e-58 | 1.72E-56 |
| PBANKA_1029400 | 9.18 | -2.61 | 1.07e-16 | 1.21E-15 |
| PBANKA_0201600 | 12.21 | -2.61 | 2.17e-14 | 2.00E-13 |
| PBANKA_0504300 | 5.61 | -2.61 | 1.57e-03 | 3.49E-03 |
| PBANKA_0703500 | 9.53 | -2.59 | 1.67e-12 | 1.29E-11 |
| PBANKA_1323800 | 8.48 | -2.58 | 4.80e-08 | 2.22E-07 |
| PBANKA_1128200 | 8.6 | -2.57 | 6.82e-28 | 1.92E-26 |
| PBANKA_1134900 | 11.63 | -2.57 | 2.59e-14 | 2.36E-13 |
| PBANKA_0405300 | 9.02 | -2.56 | 2.68e-11 | 1.85E-10 |
| PBANKA_0712800 | 9.2 | -2.55 | 7.04e-18 | 8.70E-17 |
| PBANKA_1014200 | 4.48 | -2.55 | 3.89e-03 | 7.96E-03 |
| PBANKA_0524700 | 10.63 | -2.54 | 4.67e-18 | 5.96E-17 |
| PBANKA_1134400 | 10.5 | -2.54 | 1.81e-17 | 2.18E-16 |
| PBANKA_0823600 | 6.98 | -2.52 | 2.30e-43 | 1.57E-41 |
| PBANKA_1413200 | 7.54 | -2.52 | 8.06e-24 | 1.69E-22 |
| PBANKA_0410650 | 8.86 | -2.52 | 4.04e-08 | 1.89E-07 |
| PBANKA_0925400 | 8.51 | -2.52 | 1.12e-06 | 4.30E-06 |

| | | | | |
|---|---|---|---|---|
| PBANKA_1326900 | 8.45 | 2.27 | 2.87e-18 | 3.74E-17 |
| PBANKA_1217600 | 10.73 | 2.27 | 5.48e-15 | 5.29E-14 |
| PBANKA_1430800 | 7.24 | 2.27 | 1.33e-12 | 1.04E-11 |
| PBANKA_1322100 | 6.43 | 2.26 | 3.75e-20 | 5.99E-19 |
| PBANKA_0812200 | 9.17 | 2.26 | 1.82e-14 | 1.69E-13 |
| PBANKA_1305600 | 7.19 | 2.24 | 5.53e-19 | 7.88E-18 |
| PBANKA_1306600 | 9.55 | 2.24 | 9.18e-11 | 5.92E-10 |
| PBANKA_0818500 | 8.65 | 2.23 | 1.67e-29 | 5.42E-28 |
| PBANKA_0405000 | 5.55 | 2.23 | 2.20e-08 | 1.05E-07 |
| PBANKA_1455100 | 8.31 | 2.22 | 1.95e-20 | 3.23E-19 |
| PBANKA_0408100 | 6.57 | 2.22 | 5.10e-20 | 8.00E-19 |
| PBANKA_1139600 | 5.79 | 2.22 | 8.95e-06 | 3.01E-05 |
| PBANKA_0622300 | 8.68 | 2.21 | 4.36e-22 | 8.29E-21 |
| PBANKA_1319300 | 8.41 | 2.2 | 5.48e-16 | 5.78E-15 |
| PBANKA_0929500 | 8.3 | 2.19 | 1.51e-35 | 6.52E-34 |
| PBANKA_1461200 | 6.52 | 2.19 | 6.04e-09 | 3.13E-08 |
| PBANKA_0822000 | 6.71 | 2.18 | 1.31e-13 | 1.13E-12 |
| PBANKA_0915600 | 10.19 | 2.18 | 7.53e-13 | 6.07E-12 |
| PBANKA_1315000 | 6.65 | 2.18 | 4.46e-10 | 2.67E-09 |
| PBANKA_0607100 | 5.43 | 2.18 | 1.06e-07 | 4.68E-07 |
| PBANKA_1117100 | 11.37 | 2.17 | 2.53e-18 | 3.37E-17 |
| PBANKA_0518200 | 7.77 | 2.17 | 4.68e-17 | 5.43E-16 |
| PBANKA_1451500 | 7.44 | 2.17 | 4.74e-06 | 1.67E-05 |
| PBANKA_0918700 | 7.48 | 2.15 | 1.80e-33 | 7.30E-32 |
| PBANKA_1213800 | 7.24 | 2.15 | 1.79e-32 | 6.79E-31 |
| PBANKA_1326800 | 7.53 | 2.15 | 3.75e-18 | 4.83E-17 |
| PBANKA_0620700 | 6.7 | 2.15 | 6.65e-07 | 2.64E-06 |
| PBANKA_1354200 | 4.84 | 2.15 | 2.29e-04 | 6.02E-04 |
| PBANKA_0718300 | 9.39 | 2.14 | 2.26e-34 | 9.54E-33 |
| PBANKA_1137700 | 8.28 | 2.14 | 6.53e-14 | 5.82E-13 |
| PBANKA_1121500 | 7.11 | 2.14 | 1.52e-08 | 7.48E-08 |
| PBANKA_1337900 | 6.45 | 2.13 | 2.77e-18 | 3.62E-17 |
| PBANKA_0316821 | 5.69 | 2.13 | 3.87e-13 | 3.19E-12 |
| PBANKA_1464200 | 6.51 | 2.13 | 1.93e-11 | 1.35E-10 |
| PBANKA_1444300 | 4.77 | 2.13 | 1.23e-05 | 4.01E-05 |
| PBANKA_0207500 | 8.77 | 2.12 | 6.86e-44 | 4.87E-42 |
| PBANKA_1038000 | 8.06 | 2.12 | 1.81e-18 | 2.45E-17 |
| PBANKA_0303800 | 7.54 | 2.12 | 2.11e-12 | 1.62E-11 |
| PBANKA_1454900 | 9.62 | 2.11 | 1.79e-22 | 3.50E-21 |
| PBANKA_1304300 | 7.78 | 2.11 | 1.88e-14 | 1.74E-13 |
| PBANKA_0413800 | 6.02 | 2.1 | 2.15e-22 | 4.17E-21 |
| PBANKA_1415600 | 9.63 | 2.1 | 1.01e-21 | 1.86E-20 |
| PBANKA_0819450 | 9.23 | 2.1 | 3.84e-18 | 4.93E-17 |
| PBANKA_0610200 | 9.51 | 2.1 | 3.78e-13 | 3.12E-12 |
| PBANKA_1009200 | 10.17 | 2.1 | 3.97e-11 | 2.66E-10 |
| PBANKA_0834500 | 5.76 | 2.1 | 3.41e-09 | 1.82E-08 |
| PBANKA_0932700 | 5.54 | 2.1 | 4.81e-09 | 2.52E-08 |
| PBANKA_1106200 | 5.14 | 2.1 | 1.89e-08 | 9.16E-08 |
| PBANKA_1226900 | 7.69 | 2.09 | 2.80e-16 | 3.05E-15 |
| PBANKA_0933500 | 9.07 | 2.09 | 1.20e-13 | 1.04E-12 |
| PBANKA_0608000 | 8.27 | 2.09 | 1.98e-08 | 9.56E-08 |
| PBANKA_0312500 | 8.19 | 2.08 | 1.67e-23 | 3.44E-22 |
| PBANKA_1122200 | 6.66 | 2.08 | 6.44e-18 | 8.02E-17 |
| PBANKA_1460600 | 7.92 | 2.08 | 6.78e-13 | 5.50E-12 |
| PBANKA_0835000 | 5.42 | 2.08 | 3.01e-11 | 2.06E-10 |
| PBANKA_0713900 | 6.02 | 2.08 | 2.45e-09 | 1.34E-08 |
| PBANKA_0701300 | 5.79 | 2.08 | 3.17e-05 | 9.68E-05 |
| PBANKA_0818400 | 8.72 | 2.08 | 8.06e-05 | 2.29E-04 |
| PBANKA_1413600 | 5.56 | 2.08 | 8.38e-05 | 2.37E-04 |
| PBANKA_1346200 | 7.01 | 2.07 | 6.77e-17 | 7.75E-16 |
| PBANKA_0513000 | 9.4 | 2.07 | 1.76e-06 | 6.63E-06 |
| PBANKA_0906000 | 5.18 | 2.07 | 1.98e-06 | 7.41E-06 |
| PBANKA_1339300 | 7.49 | 2.06 | 1.03e-13 | 9.01E-13 |
| PBANKA_1447000 | 6.66 | 2.06 | 5.40e-12 | 3.95E-11 |
| PBANKA_0703000 | 5.68 | 2.06 | 8.82e-12 | 6.34E-11 |
| PBANKA_0402400 | 7.25 | 2.06 | 6.50e-11 | 4.24E-10 |
| PBANKA_1359900 | 6.49 | 2.06 | 2.05e-07 | 8.65E-07 |
| PBANKA_1333200 | 4.74 | 2.06 | 6.02e-04 | 1.46E-03 |
| PBANKA_1116900 | 8.27 | 2.05 | 5.58e-08 | 2.55E-07 |
| PBANKA_1318700 | 7.81 | 2.05 | 8.79e-04 | 2.06E-03 |
| PBANKA_1408700 | 5.89 | 2.05 | 2.53e-03 | 5.41E-03 |
| PBANKA_0513900 | 10.63 | 2.04 | 3.09e-32 | 1.14E-30 |
| PBANKA_1451200 | 8.27 | 2.04 | 1.99e-06 | 7.45E-06 |
| PBANKA_0300900 | 4.8 | 2.03 | 1.01e-05 | 3.36E-05 |

| | | | | |
|---|---|---|---|---|
| PBANKA_0619200 | *12.85* | *-2.5* | 3.68e-07 | 1.51E-06 |
| PBANKA_0623300 | *11.25* | *-2.49* | 5.26e-22 | 9.89E-21 |
| PBANKA_1213200 | *5.73* | *-2.49* | 7.45e-08 | 3.37E-07 |
| PBANKA_1120400 | *10.75* | *-2.49* | 1.26e-07 | 5.45E-07 |
| PBANKA_0409400 | *10.11* | *-2.48* | 2.97e-09 | 1.60E-08 |
| PBANKA_1109600 | *7.04* | *-2.47* | 1.03e-06 | 3.99E-06 |
| PBANKA_1305000 | *11.52* | *-2.46* | 8.49e-26 | 2.07E-24 |
| PBANKA_1227100 | *10.35* | *-2.46* | 2.11e-18 | 2.82E-17 |
| PBANKA_1449000 | *7.65* | *-2.46* | 1.95e-06 | 7.31E-06 |
| PBANKA_1138900 | *9.46* | *-2.44* | 2.03e-07 | 8.60E-07 |
| PBANKA_0623100 | *12.82* | *-2.43* | 5.63e-10 | 3.33E-09 |
| PBANKA_0830400 | *10.15* | *-2.43* | 2.16e-03 | 4.69E-03 |
| PBANKA_0931300 | *10.68* | *-2.42* | 2.14e-46 | 1.77E-44 |
| PBANKA_0817700 | *10.13* | *-2.42* | 2.09e-38 | 1.04E-36 |
| PBANKA_0928200 | *10.13* | *-2.41* | 1.83e-21 | 3.30E-20 |
| PBANKA_0517000 | *12.6* | *-2.41* | 9.71e-16 | 1.01E-14 |
| PBANKA_0822800 | *8.57* | *-2.4* | 4.62e-52 | 4.82E-50 |
| PBANKA_1239100 | *6.81* | *-2.4* | 2.73e-20 | 4.47E-19 |
| PBANKA_1200600 | *11.99* | *-2.4* | 1.13e-07 | 4.95E-07 |
| PBANKA_0705100 | *8.86* | *-2.39* | 1.67e-46 | 1.40E-44 |
| PBANKA_1204800 | *6.39* | *-2.39* | 1.70e-15 | 1.73E-14 |
| PBANKA_1035100 | *9.76* | *-2.38* | 3.36e-25 | 7.75E-24 |
| PBANKA_1116800 | *11.28* | *-2.36* | 8.00e-32 | 2.90E-30 |
| PBANKA_0605700 | *10.3* | *-2.36* | 6.14e-23 | 1.24E-21 |
| PBANKA_0623500 | *9.58* | *-2.36* | 3.59e-10 | 2.17E-09 |
| PBANKA_1005600 | *7.51* | *-2.36* | 4.90e-04 | 1.21E-03 |
| PBANKA_1146241 | *4.14* | *-2.36* | 9.52e-04 | 2.22E-03 |
| PBANKA_1458000 | *8.25* | *-2.35* | 4.22e-56 | 5.14E-54 |
| PBANKA_1224200 | *10.85* | *-2.35* | 9.39e-33 | 3.67E-31 |
| PBANKA_0201340 | *6.38* | *-2.35* | 5.75e-09 | 2.98E-08 |
| PBANKA_0609500 | *11.49* | *-2.35* | 6.57e-08 | 2.98E-07 |
| PBANKA_0916650 | *8.1* | *-2.35* | 4.49e-03 | 9.03E-03 |
| PBANKA_1142300 | *8.91* | *-2.34* | 1.40e-42 | 9.21E-41 |
| PBANKA_1314500 | *6.5* | *-2.34* | 2.16e-14 | 1.99E-13 |
| PBANKA_0937200 | *11.42* | *-2.34* | 2.51e-09 | 1.36E-08 |
| PBANKA_0102700 | *6.08* | *-2.33* | 1.35e-06 | 5.18E-06 |
| PBANKA_0513600 | *9.09* | *-2.32* | 3.33e-17 | 3.94E-16 |
| PBANKA_1036300 | *9.78* | *-2.31* | 2.07e-22 | 4.04E-21 |
| PBANKA_0605500 | *8.47* | *-2.31* | 3.74e-07 | 1.53E-06 |
| PBANKA_0402700 | *11.21* | *-2.31* | 8.16e-07 | 3.21E-06 |
| PBANKA_1145200 | *8.24* | *-2.31* | 9.31e-06 | 3.13E-05 |
| PBANKA_0702000 | *9.43* | *-2.3* | 1.40e-06 | 5.38E-06 |
| PBANKA_1462300 | *4.95* | *-2.3* | 7.00e-05 | 2.01E-04 |
| PBANKA_1208900 | *5.35* | *-2.3* | 1.68e-03 | 3.72E-03 |
| PBANKA_1350600 | *8.83* | *-2.29* | 1.44e-42 | 9.35E-41 |
| PBANKA_0700700 | *9.66* | *-2.29* | 1.09e-40 | 6.49E-39 |
| PBANKA_0825000 | *8.24* | *-2.29* | 2.65e-29 | 8.42E-28 |
| PBANKA_0509200 | *10.01* | *-2.28* | 2.61e-25 | 6.15E-24 |
| PBANKA_1412100 | *6.61* | *-2.28* | 2.92e-18 | 3.79E-17 |
| PBANKA_1000800 | *7.06* | *-2.28* | 2.24e-12 | 1.71E-11 |
| PBANKA_0812600 | *8.52* | *-2.28* | 4.74e-06 | 1.67E-05 |
| PBANKA_1457700 | *11.07* | *-2.28* | 5.76e-05 | 1.68E-04 |
| PBANKA_0512000 | *7* | *-2.27* | 7.17e-05 | 2.05E-04 |
| PBANKA_1128300 | *7.2* | *-2.26* | 2.35e-22 | 4.53E-21 |
| PBANKA_1007600 | *7.41* | *-2.24* | 8.06e-13 | 6.47E-12 |
| PBANKA_1141200 | *5.48* | *-2.24* | 1.61e-11 | 1.14E-10 |
| PBANKA_0925300 | *10.3* | *-2.23* | 4.49e-16 | 4.78E-15 |
| PBANKA_1348900 | *8.75* | *-2.22* | 1.06e-41 | 6.64E-40 |
| PBANKA_0311600 | *9.53* | *-2.21* | 1.33e-39 | 7.10E-38 |
| PBANKA_1035200 | *8.74* | *-2.21* | 1.49e-05 | 4.76E-05 |
| PBANKA_1437100 | *9.99* | *-2.2* | 8.75e-19 | 1.23E-17 |
| PBANKA_1437700 | *8.95* | *-2.2* | 4.18e-06 | 1.48E-05 |
| PBANKA_1203400 | *8.58* | *-2.19* | 2.54e-31 | 8.89E-30 |
| PBANKA_1334800 | *10.09* | *-2.19* | 2.13e-10 | 1.31E-09 |
| PBANKA_1231100 | *7.28* | *-2.18* | 2.08e-30 | 7.01E-29 |
| PBANKA_1110200 | *8.87* | *-2.18* | 1.27e-09 | 7.15E-09 |
| PBANKA_1236600 | *7.88* | *-2.18* | 8.65e-08 | 3.86E-07 |
| PBANKA_0937700 | *10.03* | *-2.17* | 1.08e-27 | 3.01E-26 |
| PBANKA_1034400 | *11.61* | *-2.17* | 3.23e-20 | 5.20E-19 |
| PBANKA_0619700 | *8.72* | *-2.17* | 5.65e-05 | 1.64E-04 |
| PBANKA_1219000 | *9.39* | *-2.14* | 5.35e-29 | 1.66E-27 |
| PBANKA_1235600 | *9.99* | *-2.13* | 2.40e-36 | 1.10E-34 |
| PBANKA_0409000 | *7.14* | *-2.13* | 5.15e-16 | 5.44E-15 |
| PBANKA_0316841 | *7.36* | *-2.13* | 1.66e-11 | 1.16E-10 |

| | | | | |
|---|---|---|---|---|
| PBANKA_0300100 | *5.8* | *2.02* | 8.65e-04 | 2.03E-03 |
| PBANKA_0206100 | *7.01* | *2.01* | 5.21e-21 | 9.00E-20 |
| PBANKA_0413000 | *7.6* | *2.01* | 4.52e-19 | 6.47E-18 |
| PBANKA_0709800 | *8.09* | *2.01* | 8.30e-19 | 1.17E-17 |
| PBANKA_0938500 | *8.45* | *2.01* | 7.02e-17 | 8.00E-16 |
| PBANKA_1119300 | *6.83* | *2.01* | 6.60e-09 | 3.39E-08 |
| PBANKA_1003400 | *9.63* | *2* | 3.67e-15 | 3.61E-14 |
| PBANKA_1227500 | *5.45* | *2* | 2.67e-04 | 6.91E-04 |

| | | | | |
|---|---|---|---|---|
| PBANKA_1220300 | *8.07* | *-2.12* | 2.12e-32 | 7.99E-31 |
| PBANKA_1464600 | *8.8* | *-2.12* | 1.27e-19 | 1.93E-18 |
| PBANKA_0613700 | *11.13* | *-2.12* | 3.34e-11 | 2.26E-10 |
| PBANKA_1448100 | *6.82* | *-2.11* | 4.36e-19 | 6.29E-18 |
| PBANKA_1360100 | *10.01* | *-2.1* | 1.03e-14 | 9.77E-14 |
| PBANKA_0700100 | *4.14* | *-2.1* | 7.84e-04 | 1.85E-03 |
| PBANKA_1300500 | *5.04* | *-2.1* | 8.15e-04 | 1.92E-03 |
| PBANKA_1362000 | *10.3* | *-2.09* | 3.13e-33 | 1.25E-31 |
| PBANKA_1234600 | *9.71* | *-2.09* | 1.83e-20 | 3.05E-19 |
| PBANKA_1232300 | *6.75* | *-2.09* | 1.75e-05 | 5.54E-05 |
| PBANKA_1037300 | *9.9* | *-2.07* | 1.86e-29 | 6.00E-28 |
| PBANKA_0206700 | *5.31* | *-2.07* | 1.99e-04 | 5.27E-04 |
| PBANKA_1446000 | *5.65* | *-2.07* | 2.63e-04 | 6.83E-04 |
| PBANKA_0403800 | *9.35* | *-2.06* | 1.66e-44 | 1.23E-42 |
| PBANKA_0829700 | *6.85* | *-2.06* | 6.31e-17 | 7.27E-16 |
| PBANKA_1103900 | *7.28* | *-2.06* | 3.56e-09 | 1.90E-08 |
| PBANKA_1350200 | *6.28* | *-2.06* | 7.24e-07 | 2.86E-06 |
| PBANKA_1226400 | *9.68* | *-2.05* | 1.06e-55 | 1.26E-53 |
| PBANKA_1101100 | *11.62* | *-2.05* | 6.88e-15 | 6.58E-14 |
| PBANKA_0403400 | *6.06* | *-2.05* | 3.47e-10 | 2.10E-09 |
| PBANKA_1207500 | *6.14* | *-2.02* | 2.28e-12 | 1.73E-11 |
| PBANKA_0804600 | *8.04* | *-2.01* | 4.52e-44 | 3.30E-42 |
| PBANKA_0921400 | *9.05* | *-2.01* | 9.97e-10 | 5.70E-09 |
| PBANKA_1428700 | *11.92* | *-2.01* | 6.17e-09 | 3.19E-08 |
| PBANKA_1133400 | *9.83* | *-2.01* | 3.90e-06 | 1.39E-05 |
| PBANKA_1447800 | *6.81* | *-2* | 3.23e-16 | 3.50E-15 |
| PBANKA_1024311 | *6.61* | *-2* | 3.99e-03 | 8.14E-03 |
