## Supplementary material for "Transcriptome analysis of *Plasmodium berghei* during exo-erythrocytic development": Table S6

**Preferentially express genes in eryhtrocytic schizonts (22h)**
(significantly **down-regulated** in
Detached Cells compared to erythrocytic Schizonts)

| GeneID | logBaseMean | logFC | pVal | adjP |
|---|---|---|---|---|
| PBANKA_0500781 | 7.13 | -10.24 | 8.79e-12 | 5.31E-11 |
| PBANKA_1432200 | 9.41 | -9.45 | 5.19e-10 | 2.67E-09 |
| PBANKA_1319500 | 11.38 | -9.24 | 7.05e-10 | 3.59E-09 |
| PBANKA_0700800 | 6.53 | -9.12 | 1.83e-09 | 8.82E-09 |
| PBANKA_0504400 | 10.71 | -9.09 | 6.37e-09 | 2.92E-08 |
| PBANKA_0707100 | 12.01 | -8.97 | 9.35e-09 | 4.22E-08 |
| PBANKA_1315300 | 10.67 | -8.68 | 1.13e-08 | 5.03E-08 |
| PBANKA_1000081 | 5.56 | -8.59 | 3.63e-08 | 1.52E-07 |
| PBANKA_0704800 | 11 | -8.53 | 2.73e-08 | 1.17E-07 |
| PBANKA_0216801 | 5.71 | -8.44 | 1.67e-07 | 6.50E-07 |
| PBANKA_0810800 | 7.03 | -8.29 | 2.89e-08 | 1.23E-07 |
| PBANKA_1419300 | 10.42 | -8.13 | 1.08e-07 | 4.31E-07 |
| PBANKA_1312700 | 12.13 | -8.09 | 1.88e-07 | 7.25E-07 |
| PBANKA_0417200 | 10.11 | -8.08 | 1.45e-05 | 4.17E-05 |
| PBANKA_1353800 | 11.04 | -7.94 | 2.41e-07 | 9.14E-07 |
| PBANKA_0810700 | 10.17 | -7.9 | 1.80e-07 | 6.98E-07 |
| PBANKA_0606200 | 8.92 | -7.79 | 5.30e-07 | 1.92E-06 |
| PBANKA_1038800 | 7.64 | -7.73 | 5.12e-07 | 1.85E-06 |
| PBANKA_0704900 | 10.02 | -7.72 | 5.89e-05 | 1.51E-04 |
| PBANKA_0704700 | 7.52 | -7.64 | 4.80e-07 | 1.74E-06 |
| PBANKA_0604700 | 6.84 | -7.59 | 3.27e-07 | 1.22E-06 |
| PBANKA_1024800 | 7.26 | -7.54 | 1.08e-06 | 3.74E-06 |
| PBANKA_1014700 | 6.51 | -7.44 | 1.75e-06 | 5.84E-06 |
| PBANKA_1335800 | 5.86 | -7.4 | 1.36e-06 | 4.65E-06 |
| PBANKA_0604000 | 6.77 | -7.38 | 1.19e-06 | 4.08E-06 |
| PBANKA_1443300 | 10.54 | -7.28 | 5.28e-49 | 1.91E-47 |
| PBANKA_1209900 | 9.35 | -7.27 | 9.49e-07 | 3.32E-06 |
| PBANKA_0517600 | 9.67 | -7.27 | 2.14e-06 | 7.08E-06 |
| PBANKA_0618900 | 6.75 | -7.26 | 1.61e-06 | 5.40E-06 |
| PBANKA_MIT00700 | 6.33 | -7.26 | 6.24e-06 | 1.92E-05 |
| PBANKA_0815000 | 7.25 | -7.18 | 1.12e-06 | 3.88E-06 |
| PBANKA_1108900 | 6.42 | -7.16 | 2.21e-06 | 7.26E-06 |
| PBANKA_1407100 | 6.84 | -7.15 | 1.33e-06 | 4.53E-06 |
| PBANKA_0203600 | 8.3 | -7.15 | 2.13e-06 | 7.04E-06 |
| PBANKA_1109600 | 7.22 | -7.14 | 2.86e-06 | 9.25E-06 |
| PBANKA_1138600 | 9.21 | -7.13 | 1.82e-06 | 6.03E-06 |
| PBANKA_0507200 | 7.46 | -7.1 | 3.46e-06 | 1.11E-05 |
| PBANKA_1239400 | 6.65 | -7.07 | 2.09e-06 | 6.90E-06 |
| PBANKA_0812600 | 8.74 | -7.06 | 3.42e-06 | 1.10E-05 |
| PBANKA_1206500 | 5.48 | -7.03 | 4.40e-06 | 1.39E-05 |
| PBANKA_0110700 | 6.2 | -7.02 | 5.59e-06 | 1.73E-05 |
| PBANKA_1451100 | 9.84 | -7.01 | 3.73e-03 | 6.64E-03 |
| PBANKA_1424400 | 6.82 | -7 | 2.40e-06 | 7.88E-06 |
| PBANKA_1334700 | 8.99 | -7 | 2.55e-06 | 8.32E-06 |
| PBANKA_0935700 | 5.23 | -7 | 2.56e-05 | 7.01E-05 |
| PBANKA_1201300 | 6.75 | -6.99 | 3.67e-06 | 1.17E-05 |
| PBANKA_0315700 | 6.58 | -6.98 | 8.59e-06 | 2.59E-05 |
| PBANKA_0902200 | 5.71 | -6.97 | 2.33e-05 | 6.42E-05 |
| PBANKA_1407000 | 6.33 | -6.94 | 3.39e-06 | 1.09E-05 |
| PBANKA_0214550 | 5.6 | -6.93 | 8.86e-06 | 2.66E-05 |
| PBANKA_0202300 | 6.99 | -6.92 | 4.87e-06 | 1.53E-05 |
| PBANKA_1038200 | 6.76 | -6.91 | 5.95e-06 | 1.84E-05 |
| PBANKA_1016900 | 4.92 | -6.9 | 3.03e-05 | 8.21E-05 |
| PBANKA_1360800 | 5.78 | -6.87 | 1.04e-05 | 3.09E-05 |
| PBANKA_0936200 | 5.45 | -6.86 | 1.12e-05 | 3.30E-05 |
| PBANKA_0835100 | 6.24 | -6.86 | 1.67e-05 | 4.75E-05 |
| PBANKA_1229900 | 8.56 | -6.85 | 8.36e-06 | 2.52E-05 |
| PBANKA_1431500 | 7.4 | -6.84 | 2.58e-04 | 5.86E-04 |
| PBANKA_1127500 | 6.69 | -6.82 | 4.10e-06 | 1.30E-05 |
| PBANKA_1233900 | 6.76 | -6.81 | 4.79e-06 | 1.50E-05 |
| PBANKA_0933800 | 6.56 | -6.81 | 6.35e-06 | 1.95E-05 |
| PBANKA_1204800 | 6.56 | -6.79 | 7.19e-06 | 2.19E-05 |
| PBANKA_0926500 | 9.79 | -6.78 | 1.13e-05 | 3.33E-05 |
| PBANKA_1360900 | 5.37 | -6.78 | 5.80e-05 | 1.49E-04 |
| PBANKA_1134600 | 5.61 | -6.77 | 1.29e-05 | 3.74E-05 |
| PBANKA_1209500 | 6.37 | -6.75 | 4.95e-06 | 1.55E-05 |
| PBANKA_1209100 | 6.67 | -6.75 | 1.02e-05 | 3.02E-05 |
| PBANKA_0926400 | 5.47 | -6.75 | 2.10e-05 | 5.83E-05 |
| PBANKA_1126300 | 6.96 | -6.73 | 6.61e-06 | 2.02E-05 |

**Preferentially expressed genes in Detached cells**
(significantly **up-regulated** in
Detached Cells compared to erythrocytic Schizonts)

| feature | logBaseMean | logFC | pVal | adjP |
|---|---|---|---|---|
| PBANKA_0007501 | 6.57 | 13.85 | 6.99e-19 | 6.50E-18 |
| PBANKA_0007801 | 5.97 | 13.22 | 2.95e-04 | 6.65E-04 |
| PBANKA_0600031 | 8.98 | 12.17 | 6.15e-31 | 9.95E-30 |
| PBANKA_1300011 | 4.84 | 12.07 | 4.68e-11 | 2.65E-10 |
| PBANKA_0700041 | 4.61 | 11.87 | 1.06e-04 | 2.59E-04 |
| PBANKA_1465051 | 12.21 | 11.81 | 1.55e-14 | 1.14E-13 |
| PBANKA_0836981 | 4.62 | 11.74 | 4.51e-04 | 9.76E-04 |
| PBANKA_1246800 | 4.22 | 10.92 | 8.26e-09 | 3.75E-08 |
| PBANKA_1100351 | 3.8 | 10.76 | 3.79e-07 | 1.40E-06 |
| PBANKA_0008101 | 7.88 | 9.85 | 1.19e-43 | 3.44E-42 |
| PBANKA_0316900 | 3.04 | 9.5 | 2.08e-04 | 4.82E-04 |
| PBANKA_1400041 | 4.43 | 9.45 | 9.83e-08 | 3.94E-07 |
| PBANKA_0837001 | 6 | 9.38 | 1.08e-04 | 2.63E-04 |
| PBANKA_0316500 | 9.23 | 8.98 | 3.72e-15 | 2.85E-14 |
| PBANKA_0700081 | 6.83 | 8.27 | 6.98e-30 | 1.08E-28 |
| PBANKA_0316200 | 15.58 | 8.24 | 1.07e-14 | 7.98E-14 |
| PBANKA_1024600 | 12.33 | 8.06 | 7.21e-110 | 2.63E-107 |
| PBANKA_1229000 | 12.92 | 8.05 | 2.47e-62 | 1.50E-60 |
| PBANKA_0700061 | 6.08 | 7.7 | 1.45e-25 | 1.86E-24 |
| PBANKA_0700071 | 5.17 | 7.67 | 1.90e-04 | 4.44E-04 |
| PBANKA_1003900 | 10.46 | 7.62 | 2.81e-68 | 2.18E-66 |
| PBANKA_0304300 | 9.05 | 7.57 | 2.37e-129 | 1.73E-126 |
| PBANKA_0924400 | 9.92 | 7.52 | 4.77e-53 | 2.03E-51 |
| PBANKA_1000600 | 11.72 | 7.52 | 6.05e-42 | 1.59E-40 |
| PBANKA_1300031 | 6.23 | 7.47 | 1.85e-07 | 7.13E-07 |
| PBANKA_1343000 | 10.38 | 7.44 | 7.49e-112 | 3.19E-109 |
| PBANKA_1207100 | 11.15 | 7.28 | 4.61e-61 | 2.62E-59 |
| PBANKA_0837101 | 3.48 | 7.12 | 4.09e-04 | 8.93E-04 |
| PBANKA_1330100 | 8.35 | 7.04 | 5.33e-72 | 5.34E-70 |
| PBANKA_1465600 | 6.02 | 6.98 | 2.41e-20 | 2.43E-19 |
| PBANKA_0836200 | 11.12 | 6.91 | 9.39e-54 | 4.14E-52 |
| PBANKA_1246161 | 10.41 | 6.9 | 1.70e-54 | 7.85E-53 |
| PBANKA_1033600 | 7.85 | 6.81 | 6.74e-91 | 1.04E-88 |
| PBANKA_1027500 | 7 | 6.8 | 2.03e-21 | 2.16E-20 |
| PBANKA_0600021 | 7.66 | 6.77 | 2.56e-08 | 1.10E-07 |
| PBANKA_0300600 | 12.65 | 6.75 | 3.57e-38 | 8.00E-37 |
| PBANKA_0305300 | 9.16 | 6.74 | 1.20e-193 | 6.13E-190 |
| PBANKA_0623150 | 8.32 | 6.71 | 2.38e-49 | 9.01E-48 |
| PBANKA_0600200 | 3.9 | 6.69 | 1.19e-05 | 3.47E-05 |
| PBANKA_0944061 | 4.1 | 6.63 | 2.26e-07 | 8.60E-07 |
| PBANKA_0301500 | 9.63 | 6.61 | 3.30e-49 | 1.22E-47 |
| PBANKA_0100900 | 11.27 | 6.59 | 1.71e-74 | 1.82E-72 |
| PBANKA_1207000 | 10.58 | 6.56 | 1.57e-79 | 1.95E-77 |
| PBANKA_1040581 | 7.17 | 6.56 | 3.33e-46 | 1.09E-44 |
| PBANKA_1026200 | 10.76 | 6.54 | 1.20e-132 | 1.02E-129 |
| PBANKA_0206900 | 6.58 | 6.52 | 1.54e-51 | 6.24E-50 |
| PBANKA_0214600 | 11.58 | 6.5 | 2.42e-74 | 2.52E-72 |
| PBANKA_0837141 | 5.93 | 6.48 | 2.77e-20 | 2.77E-19 |
| PBANKA_0107700 | 9.61 | 6.46 | 2.56e-55 | 1.23E-53 |
| PBANKA_0216041 | 5.92 | 6.45 | 1.53e-04 | 3.63E-04 |
| PBANKA_0720800 | 9.98 | 6.41 | 1.60e-102 | 3.90E-100 |
| PBANKA_0600100 | 4.17 | 6.41 | 5.17e-05 | 1.34E-04 |
| PBANKA_0205000 | 11.83 | 6.38 | 4.72e-57 | 2.42E-55 |
| PBANKA_0518900 | 12.74 | 6.35 | 1.93e-66 | 1.39E-64 |
| PBANKA_0944101 | 5.66 | 6.3 | 1.39e-16 | 1.15E-15 |
| PBANKA_1141100 | 9.38 | 6.28 | 1.52e-191 | 3.90E-188 |
| PBANKA_0623200 | 10.93 | 6.27 | 5.16e-41 | 1.33E-39 |
| PBANKA_1465000 | 10.04 | 6.27 | 2.58e-10 | 1.36E-09 |
| PBANKA_1400051 | 6.09 | 6.26 | 2.37e-18 | 2.13E-17 |
| PBANKA_1465921 | 3.05 | 6.26 | 4.49e-03 | 7.88E-03 |
| PBANKA_0623000 | 11.55 | 6.23 | 2.11e-37 | 4.52E-36 |
| PBANKA_0213500 | 9.61 | 6.22 | 2.23e-104 | 6.71E-102 |
| PBANKA_1101100 | 11.62 | 6.22 | 2.91e-47 | 9.92E-46 |
| PBANKA_1145900 | 12.3 | 6.21 | 8.17e-56 | 4.02E-54 |
| PBANKA_1459000 | 6.77 | 6.21 | 2.64e-42 | 7.15E-41 |
| PBANKA_1146700 | 4.27 | 6.21 | 4.29e-05 | 1.13E-04 |
| PBANKA_0800300 | 8.62 | 6.2 | 6.47e-57 | 3.28E-55 |
| PBANKA_0112721 | 8.41 | 6.2 | 2.56e-26 | 3.41E-25 |
| PBANKA_0806000 | 7.7 | 6.19 | 4.34e-28 | 6.28E-27 |

| | | | | |
|---|---|---|---|---|
| PBANKA_1208500 | 5.93 | -6.73 | 1.64e-05 | 4.67E-05 |
| PBANKA_1029600 | 7.6 | -6.72 | 8.76e-04 | 1.79E-03 |
| PBANKA_0719100 | 7.11 | -6.71 | 9.29e-06 | 2.77E-05 |
| PBANKA_0919500 | 5.48 | -6.71 | 9.82e-06 | 2.92E-05 |
| PBANKA_0314800 | 6.96 | -6.71 | 1.19e-05 | 3.49E-05 |
| PBANKA_1204300 | 7.03 | -6.7 | 7.31e-06 | 2.22E-05 |
| PBANKA_1342300 | 8.19 | -6.69 | 9.43e-05 | 2.33E-04 |
| PBANKA_1326100 | 6.56 | -6.68 | 1.02e-05 | 3.02E-05 |
| PBANKA_0817100 | 6.66 | -6.64 | 1.27e-05 | 3.69E-05 |
| PBANKA_0705400 | 4.91 | -6.62 | 1.91e-05 | 5.35E-05 |
| PBANKA_0523600 | 6.75 | -6.61 | 1.30e-05 | 3.78E-05 |
| PBANKA_1219100 | 8.75 | -6.6 | 1.12e-04 | 2.73E-04 |
| PBANKA_1128800 | 7.65 | -6.59 | 1.75e-05 | 4.94E-05 |
| PBANKA_1318500 | 8.82 | -6.59 | 3.48e-04 | 7.71E-04 |
| PBANKA_1350000 | 6.53 | -6.57 | 1.34e-05 | 3.86E-05 |
| PBANKA_1038400 | 6.6 | -6.54 | 1.95e-05 | 5.44E-05 |
| PBANKA_0907600 | 6.28 | -6.52 | 2.37e-05 | 6.53E-05 |
| PBANKA_0616700 | 7.19 | -6.51 | 1.59e-05 | 4.54E-05 |
| PBANKA_1412900 | 5.71 | -6.5 | 2.34e-05 | 6.46E-05 |
| PBANKA_1414500 | 8.41 | -6.5 | 2.64e-05 | 7.24E-05 |
| PBANKA_0605000 | 6.8 | -6.5 | 2.76e-05 | 7.53E-05 |
| PBANKA_1455800 | 6.4 | -6.49 | 1.54e-05 | 4.41E-05 |
| PBANKA_0705300 | 5.63 | -6.49 | 2.22e-05 | 6.16E-05 |
| PBANKA_1425700 | 5.74 | -6.49 | 2.64e-05 | 7.22E-05 |
| PBANKA_0521241 | 7.49 | -6.49 | 2.26e-04 | 5.18E-04 |
| PBANKA_1359600 | 7.12 | -6.48 | 1.73e-05 | 4.91E-05 |
| PBANKA_0510000 | 5.21 | -6.48 | 4.94e-05 | 1.28E-04 |
| PBANKA_0109800 | 8.32 | -6.47 | 1.76e-05 | 4.99E-05 |
| PBANKA_1316800 | 5.96 | -6.47 | 2.75e-05 | 7.49E-05 |
| PBANKA_0823200 | 9 | -6.46 | 1.79e-05 | 5.06E-05 |
| PBANKA_1103900 | 7.47 | -6.46 | 2.26e-05 | 6.25E-05 |
| PBANKA_1335000 | 8.33 | -6.46 | 1.35e-04 | 3.23E-04 |
| PBANKA_0403100 | 7.39 | -6.45 | 2.31e-05 | 6.37E-05 |
| PBANKA_0521900 | 5.69 | -6.45 | 5.70e-05 | 1.47E-04 |
| PBANKA_1123200 | 6.91 | -6.44 | 2.01e-05 | 5.61E-05 |
| PBANKA_1327061 | 6.09 | -6.44 | 3.14e-05 | 8.52E-05 |
| PBANKA_0110600 | 4.49 | -6.44 | 1.97e-04 | 4.58E-04 |
| PBANKA_1314500 | 6.68 | -6.43 | 2.26e-05 | 6.26E-05 |
| PBANKA_0104500 | 9.22 | -6.4 | 2.11e-05 | 5.88E-05 |
| PBANKA_0830300 | 5.67 | -6.36 | 1.51e-04 | 3.58E-04 |
| PBANKA_1321500 | 6.21 | -6.33 | 3.46e-05 | 9.30E-05 |
| PBANKA_1452900 | 6.24 | -6.33 | 4.32e-05 | 1.14E-04 |
| PBANKA_1431800 | 5.66 | -6.33 | 4.46e-05 | 1.17E-04 |
| PBANKA_0706200 | 6.89 | -6.33 | 4.96e-05 | 1.29E-04 |
| PBANKA_1430600 | 6.32 | -6.33 | 1.11e-04 | 2.70E-04 |
| PBANKA_1421700 | 9.05 | -6.33 | 1.61e-03 | 3.11E-03 |
| PBANKA_1213200 | 5.87 | -6.32 | 2.28e-05 | 6.30E-05 |
| PBANKA_1233600 | 8.65 | -6.32 | 1.75e-03 | 3.36E-03 |
| PBANKA_0301400 | 6.18 | -6.31 | 2.59e-05 | 7.09E-05 |
| PBANKA_0417000 | 6.22 | -6.31 | 3.16e-05 | 8.55E-05 |
| PBANKA_1217800 | 6.22 | -6.3 | 3.91e-05 | 1.04E-04 |
| PBANKA_1145200 | 8.44 | -6.3 | 4.29e-05 | 1.13E-04 |
| PBANKA_1305900 | 6.45 | -6.3 | 5.50e-05 | 1.42E-04 |
| PBANKA_1124900 | 7.13 | -6.29 | 2.52e-05 | 6.92E-05 |
| PBANKA_0825700 | 6.86 | -6.28 | 8.03e-05 | 2.01E-04 |
| PBANKA_0623450 | 7.71 | -6.27 | 4.63e-14 | 3.29E-13 |
| PBANKA_0519700 | 6.45 | -6.27 | 4.52e-05 | 1.18E-04 |
| PBANKA_1038500 | 5.46 | -6.26 | 5.23e-05 | 1.36E-04 |
| PBANKA_0508700 | 6.23 | -6.25 | 4.37e-05 | 1.15E-04 |
| PBANKA_0510300 | 8.17 | -6.24 | 4.04e-05 | 1.07E-04 |
| PBANKA_1009700 | 6.29 | -6.23 | 3.97e-05 | 1.05E-04 |
| PBANKA_0907400 | 5.25 | -6.23 | 6.59e-05 | 1.67E-04 |
| PBANKA_1428600 | 5.53 | -6.23 | 1.52e-04 | 3.62E-04 |
| PBANKA_1000100 | 4.07 | -6.23 | 1.53e-03 | 2.97E-03 |
| PBANKA_1444600 | 7.24 | -6.22 | 6.46e-05 | 1.64E-04 |
| PBANKA_1123600 | 5.94 | -6.21 | 5.78e-05 | 1.48E-04 |
| PBANKA_0800600 | 5.21 | -6.21 | 6.07e-05 | 1.55E-04 |
| PBANKA_1421000 | 6.5 | -6.2 | 4.05e-05 | 1.07E-04 |
| PBANKA_1440900 | 5.97 | -6.18 | 3.22e-05 | 8.70E-05 |
| PBANKA_0103400 | 6.68 | -6.18 | 3.98e-05 | 1.05E-04 |
| PBANKA_1420100 | 5.86 | -6.17 | 3.17e-05 | 8.57E-05 |
| PBANKA_1333100 | 6.8 | -6.17 | 4.44e-05 | 1.16E-04 |
| PBANKA_0600400 | 5.64 | -6.14 | 1.06e-04 | 2.61E-04 |
| PBANKA_1350900 | 5.97 | -6.13 | 8.43e-05 | 2.10E-04 |

| | | | | |
|---|---|---|---|---|
| PBANKA_1344500 | 12.99 | 6.18 | 3.55e-40 | 8.69E-39 |
| PBANKA_1364900 | 8.67 | 6.16 | 1.13e-87 | 1.52E-85 |
| PBANKA_0512400 | 9.55 | 6.16 | 1.92e-70 | 1.75E-68 |
| PBANKA_0700600 | 8.9 | 6.15 | 6.81e-47 | 2.29E-45 |
| PBANKA_0100021 | 8.49 | 6.14 | 1.01e-04 | 2.48E-04 |
| PBANKA_1208700 | 9.26 | 6.13 | 7.91e-55 | 3.75E-53 |
| PBANKA_1340300 | 9.22 | 6.12 | 8.62e-88 | 1.19E-85 |
| PBANKA_0700700 | 9.72 | 6.11 | 2.33e-113 | 1.08E-110 |
| PBANKA_0814000 | 8.81 | 6.11 | 1.20e-68 | 9.93E-67 |
| PBANKA_0512500 | 9.45 | 6.1 | 2.77e-71 | 2.57E-69 |
| PBANKA_1445200 | 7.72 | 6.1 | 5.17e-51 | 2.07E-49 |
| PBANKA_1143500 | 8.53 | 6.07 | 2.76e-100 | 6.14E-98 |
| PBANKA_1334300 | 12.12 | 6.07 | 4.36e-61 | 2.53E-59 |
| PBANKA_0600081 | 3.69 | 6.07 | 3.41e-05 | 9.19E-05 |
| PBANKA_1019520 | 3.99 | 6.07 | 5.26e-05 | 1.36E-04 |
| PBANKA_1105800 | 8.73 | 6.05 | 8.71e-72 | 8.40E-70 |
| PBANKA_1024300 | 10.08 | 6.05 | 2.26e-26 | 3.02E-25 |
| PBANKA_0613900 | 9.34 | 6.02 | 7.67e-66 | 5.45E-64 |
| PBANKA_0601700 | 8.96 | 6.02 | 6.60e-57 | 3.31E-55 |
| PBANKA_1129300 | 7.67 | 6.02 | 3.90e-38 | 8.67E-37 |
| PBANKA_0519500 | 11.86 | 6.02 | 1.18e-29 | 1.81E-28 |
| PBANKA_1225400 | 10.73 | 6.02 | 3.35e-25 | 4.24E-24 |
| PBANKA_1334200 | 9.12 | 6.01 | 5.84e-110 | 2.30E-107 |
| PBANKA_0701000 | 10.36 | 6.01 | 3.10e-47 | 1.05E-45 |
| PBANKA_0623300 | 11.27 | 6.01 | 1.00e-46 | 3.33E-45 |
| PBANKA_1002700 | 9.19 | 5.99 | 8.32e-142 | 1.06E-138 |
| PBANKA_1359200 | 10.18 | 5.98 | 1.11e-74 | 1.20E-72 |
| PBANKA_1424300 | 12.95 | 5.96 | 1.65e-88 | 2.41E-86 |
| PBANKA_1003000 | 13.1 | 5.96 | 5.01e-68 | 3.82E-66 |
| PBANKA_0914200 | 11.11 | 5.96 | 5.64e-46 | 1.84E-44 |
| PBANKA_1220200 | 10.72 | 5.95 | 1.67e-26 | 2.25E-25 |
| PBANKA_0100700 | 12.74 | 5.94 | 2.74e-23 | 3.22E-22 |
| PBANKA_1031000 | 8.7 | 5.92 | 6.25e-54 | 2.80E-52 |
| PBANKA_0913400 | 12.01 | 5.88 | 3.34e-57 | 1.74E-55 |
| PBANKA_1354500 | 11.6 | 5.87 | 6.83e-46 | 2.18E-44 |
| PBANKA_1227900 | 9.71 | 5.87 | 7.86e-41 | 2.00E-39 |
| PBANKA_1100860 | 8.77 | 5.86 | 2.98e-43 | 8.43E-42 |
| PBANKA_0903000 | 9.62 | 5.85 | 3.44e-108 | 1.17E-105 |
| PBANKA_1032300 | 9.26 | 5.84 | 1.53e-116 | 8.67E-114 |
| PBANKA_1313000 | 9.47 | 5.81 | 6.49e-78 | 7.72E-76 |
| PBANKA_1348300 | 10.11 | 5.79 | 1.06e-99 | 2.25E-97 |
| PBANKA_1409300 | 9.02 | 5.79 | 4.60e-63 | 3.05E-61 |
| PBANKA_0937800 | 9.83 | 5.78 | 4.94e-102 | 1.15E-99 |
| PBANKA_0939200 | 9.41 | 5.78 | 2.68e-93 | 4.56E-91 |
| PBANKA_0603200 | 11.2 | 5.75 | 1.04e-34 | 2.01E-33 |
| PBANKA_1030800 | 6.28 | 5.75 | 9.86e-33 | 1.73E-31 |
| PBANKA_0100061 | 7.85 | 5.74 | 7.93e-46 | 2.52E-44 |
| PBANKA_0707400 | 10.68 | 5.74 | 4.73e-45 | 1.45E-43 |
| PBANKA_0600086 | 3.83 | 5.74 | 2.87e-04 | 6.48E-04 |
| PBANKA_0213400 | 10.2 | 5.73 | 5.99e-62 | 3.52E-60 |
| PBANKA_0111200 | 10.77 | 5.72 | 6.57e-104 | 1.87E-101 |
| PBANKA_1315600 | 9.73 | 5.72 | 8.80e-48 | 3.10E-46 |
| PBANKA_1406800 | 9.13 | 5.71 | 4.49e-28 | 6.49E-27 |
| PBANKA_0311200 | 9.68 | 5.7 | 1.79e-144 | 3.05E-141 |
| PBANKA_1440300 | 8.03 | 5.7 | 1.02e-95 | 1.86E-93 |
| PBANKA_1423200 | 8.97 | 5.7 | 1.97e-68 | 1.60E-66 |
| PBANKA_1019500 | 12.16 | 5.68 | 6.42e-45 | 1.94E-43 |
| PBANKA_0517000 | 12.61 | 5.68 | 1.38e-31 | 2.29E-30 |
| PBANKA_1129700 | 9.57 | 5.67 | 5.07e-106 | 1.62E-103 |
| PBANKA_1218000 | 10.19 | 5.67 | 7.40e-36 | 1.50E-34 |
| PBANKA_0418500 | 11.68 | 5.67 | 1.00e-06 | 3.49E-06 |
| PBANKA_1105700 | 8.98 | 5.65 | 1.02e-65 | 7.14E-64 |
| PBANKA_1401200 | 9.94 | 5.65 | 1.05e-50 | 4.11E-49 |
| PBANKA_1003200 | 7.96 | 5.64 | 5.98e-51 | 2.37E-49 |
| PBANKA_MIT03500 | 12.68 | 5.63 | 2.98e-96 | 5.64E-94 |
| PBANKA_1331900 | 11.47 | 5.63 | 1.11e-62 | 7.17E-61 |
| PBANKA_1122700 | 12.72 | 5.63 | 9.50e-27 | 1.29E-25 |
| PBANKA_0524100 | 8.28 | 5.61 | 7.14e-39 | 1.66E-37 |
| PBANKA_1230500 | 8.21 | 5.6 | 1.62e-41 | 4.21E-40 |
| PBANKA_1128200 | 8.63 | 5.59 | 2.71e-49 | 1.02E-47 |
| PBANKA_0500600 | 6.87 | 5.59 | 8.39e-33 | 1.49E-31 |
| PBANKA_1036400 | 10.15 | 5.55 | 2.41e-63 | 1.64E-61 |
| PBANKA_0414700 | 10.74 | 5.55 | 4.08e-53 | 1.75E-51 |
| PBANKA_1462600 | 9.15 | 5.55 | 1.54e-48 | 5.50E-47 |

| | | | | | | | | | |
|---|---|---|---|---|---|---|---|---|---|
| PBANKA_1209400 | 7.84 | -6.12 | 9.54e-05 | 2.36E-04 | PBANKA_1231000 | 10.88 | 5.55 | 3.11e-44 | 9.25E-43 |
| PBANKA_0936100 | 5.53 | -6.1 | 1.15e-04 | 2.78E-04 | PBANKA_1324900 | 7.99 | 5.55 | 6.22e-38 | 1.37E-36 |
| PBANKA_1201500 | 5.1 | -6.1 | 1.21e-04 | 2.92E-04 | PBANKA_0500700 | 8.53 | 5.54 | 1.44e-37 | 3.10E-36 |
| PBANKA_1424100 | 5.47 | -6.08 | 8.60e-05 | 2.14E-04 | PBANKA_1401900 | 8.86 | 5.53 | 1.48e-77 | 1.72E-75 |
| PBANKA_0821200 | 8.38 | -6.07 | 4.52e-05 | 1.18E-04 | PBANKA_1457550 | 8.85 | 5.52 | 8.77e-07 | 3.08E-06 |
| PBANKA_1430200 | 5.54 | -6.07 | 7.73e-05 | 1.94E-04 | PBANKA_1345800 | 10.62 | 5.49 | 3.87e-95 | 6.82E-93 |
| PBANKA_0800041 | 3.86 | -6.07 | 1.48e-03 | 2.89E-03 | PBANKA_0317021 | 3.11 | 5.47 | 1.59e-03 | 3.08E-03 |
| PBANKA_0942300 | 5.71 | -6.05 | 8.76e-05 | 2.18E-04 | PBANKA_1327800 | 7.66 | 5.46 | 1.69e-36 | 3.51E-35 |
| PBANKA_0316700 | 5.48 | -6.05 | 2.47e-04 | 5.63E-04 | PBANKA_0929600 | 10.48 | 5.45 | 5.49e-103 | 1.40E-100 |
| PBANKA_1020600 | 8.82 | -6.04 | 4.26e-05 | 1.12E-04 | PBANKA_1135300 | 10.47 | 5.45 | 6.31e-33 | 1.13E-31 |
| PBANKA_1450900 | 7.5 | -6.04 | 5.53e-05 | 1.43E-04 | PBANKA_1416400 | 9.3 | 5.42 | 4.28e-93 | 7.06E-91 |
| PBANKA_0110800 | 8.1 | -6.02 | 7.75e-04 | 1.60E-03 | PBANKA_1134800 | 8.08 | 5.42 | 1.28e-76 | 1.45E-74 |
| PBANKA_0931800 | 6.83 | -6.01 | 9.31e-05 | 2.30E-04 | PBANKA_0200400 | 9.88 | 5.42 | 1.60e-34 | 3.07E-33 |
| PBANKA_1361500 | 6.13 | -6 | 1.13e-04 | 2.75E-04 | PBANKA_1221700 | 7.43 | 5.41 | 2.10e-30 | 3.31E-29 |
| PBANKA_0902400 | 5.7 | -6 | 1.23e-04 | 2.97E-04 | PBANKA_0714700 | 6.41 | 5.41 | 7.01e-27 | 9.55E-26 |
| PBANKA_0102700 | 6.23 | -5.99 | 7.38e-05 | 1.86E-04 | PBANKA_1145400 | 12.06 | 5.41 | 1.88e-25 | 2.39E-24 |
| PBANKA_1423700 | 8.93 | -5.98 | 5.80e-05 | 1.49E-04 | PBANKA_1462000 | 8.84 | 5.4 | 1.41e-113 | 7.20E-111 |
| PBANKA_1201600 | 6.39 | -5.98 | 9.05e-05 | 2.24E-04 | PBANKA_1317600 | 10.08 | 5.39 | 1.52e-97 | 3.11E-95 |
| PBANKA_1243500 | 7.17 | -5.96 | 1.07e-04 | 2.61E-04 | PBANKA_0912000 | 10.45 | 5.38 | 1.95e-30 | 3.07E-29 |
| PBANKA_1102500 | 6.13 | -5.95 | 6.20e-05 | 1.58E-04 | PBANKA_1135700 | 11.92 | 5.37 | 1.93e-69 | 1.70E-67 |
| PBANKA_1033700 | 5.99 | -5.95 | 1.08e-04 | 2.63E-04 | PBANKA_0606600 | 10.37 | 5.37 | 1.20e-22 | 1.36E-21 |
| PBANKA_0201051 | 5.65 | -5.95 | 2.13e-04 | 4.91E-04 | PBANKA_1006800 | 9.14 | 5.36 | 4.61e-67 | 3.42E-65 |
| PBANKA_0522300 | 6.39 | -5.94 | 1.99e-04 | 4.62E-04 | PBANKA_1329600 | 10.56 | 5.36 | 6.47e-43 | 1.78E-41 |
| PBANKA_0512000 | 7.19 | -5.94 | 6.99e-04 | 1.45E-03 | PBANKA_0502700 | 8.33 | 5.36 | 1.03e-18 | 9.42E-18 |
| PBANKA_1136500 | 5.46 | -5.93 | 8.69e-05 | 2.16E-04 | PBANKA_1321400 | 10.75 | 5.35 | 3.93e-31 | 6.43E-30 |
| PBANKA_1360400 | 7.43 | -5.92 | 9.41e-05 | 2.33E-04 | PBANKA_1035900 | 10.35 | 5.34 | 5.21e-88 | 7.40E-86 |
| PBANKA_1217900 | 5.41 | -5.92 | 1.55e-04 | 3.66E-04 | PBANKA_1433400 | 7.29 | 5.34 | 1.10e-42 | 2.99E-41 |
| PBANKA_0314000 | 5.62 | -5.91 | 1.89e-04 | 4.42E-04 | PBANKA_1101300 | 11.69 | 5.33 | 2.59e-39 | 6.13E-38 |
| PBANKA_1321900 | 4.8 | -5.88 | 4.13e-04 | 8.99E-04 | PBANKA_0908300 | 10.96 | 5.33 | 1.26e-25 | 1.61E-24 |
| PBANKA_0107200 | 7.12 | -5.86 | 1.03e-04 | 2.54E-04 | PBANKA_0001201 | 3.28 | 5.33 | 3.39e-03 | 6.12E-03 |
| PBANKA_0103200 | 7.87 | -5.86 | 6.08e-04 | 1.28E-03 | PBANKA_1314900 | 11.29 | 5.3 | 1.45e-47 | 5.09E-46 |
| PBANKA_0701400 | 5.69 | -5.84 | 2.00e-04 | 4.64E-04 | PBANKA_0712800 | 9.24 | 5.3 | 4.89e-27 | 6.74E-26 |
| PBANKA_1220000 | 6.08 | -5.83 | 1.81e-04 | 4.25E-04 | PBANKA_1000051 | 6.29 | 5.3 | 2.77e-23 | 3.24E-22 |
| PBANKA_0804300 | 5.92 | -5.83 | 1.82e-04 | 4.27E-04 | PBANKA_1462900 | 8.81 | 5.29 | 2.57e-136 | 2.63E-133 |
| PBANKA_0933600 | 4.86 | -5.83 | 1.11e-03 | 2.22E-03 | PBANKA_1323700 | 8.07 | 5.27 | 3.10e-33 | 5.68E-32 |
| PBANKA_0939400 | 6.01 | -5.82 | 1.20e-04 | 2.89E-04 | PBANKA_1030000 | 7.92 | 5.26 | 6.26e-81 | 8.21E-79 |
| PBANKA_0806200 | 6.3 | -5.82 | 1.74e-04 | 4.10E-04 | PBANKA_1222700 | 9.38 | 5.26 | 3.70e-43 | 1.03E-41 |
| PBANKA_1114100 | 7.62 | -5.82 | 1.93e-04 | 4.50E-04 | PBANKA_1332500 | 8.38 | 5.25 | 3.65e-63 | 2.46E-61 |
| PBANKA_0708100 | 5.12 | -5.82 | 3.92e-04 | 8.58E-04 | PBANKA_1440200 | 8.14 | 5.24 | 7.63e-58 | 4.06E-56 |
| PBANKA_1404000 | 6.2 | -5.81 | 8.10e-05 | 2.03E-04 | PBANKA_1000500 | 9.23 | 5.23 | 7.59e-72 | 7.47E-70 |
| PBANKA_1436300 | 7.02 | -5.81 | 1.35e-04 | 3.24E-04 | PBANKA_0311800 | 12.5 | 5.22 | 2.83e-55 | 1.35E-53 |
| PBANKA_1359800 | 5.2 | -5.8 | 2.20e-04 | 5.08E-04 | PBANKA_1211500 | 9.23 | 5.21 | 9.61e-75 | 1.07E-72 |
| PBANKA_0213900 | 4.53 | -5.8 | 1.77e-03 | 3.39E-03 | PBANKA_1146721 | 5.22 | 5.21 | 7.78e-13 | 5.16E-12 |
| PBANKA_0307900 | 6.52 | -5.78 | 1.23e-04 | 2.95E-04 | PBANKA_0822700 | 8.12 | 5.2 | 4.07e-49 | 1.49E-47 |
| PBANKA_0918500 | 5.72 | -5.78 | 1.39e-04 | 3.31E-04 | PBANKA_0906400 | 10.26 | 5.19 | 2.24e-36 | 4.59E-35 |
| PBANKA_1411600 | 5.85 | -5.77 | 1.24e-04 | 2.98E-04 | PBANKA_0923500 | 9.23 | 5.17 | 3.35e-72 | 3.43E-70 |
| PBANKA_1205700 | 4.48 | -5.77 | 3.09e-04 | 6.94E-04 | PBANKA_1441500 | 9.36 | 5.16 | 2.60e-80 | 3.33E-78 |
| PBANKA_0109400 | 5.18 | -5.75 | 4.47e-04 | 9.69E-04 | PBANKA_1420500 | 12.1 | 5.15 | 2.75e-13 | 1.87E-12 |
| PBANKA_1427400 | 5.56 | -5.74 | 1.66e-04 | 3.92E-04 | PBANKA_1010300 | 11.73 | 5.13 | 6.07e-12 | 3.71E-11 |
| PBANKA_0915400 | 5.18 | -5.71 | 1.30e-03 | 2.56E-03 | PBANKA_0602500 | 7.56 | 5.12 | 1.66e-39 | 3.99E-38 |
| PBANKA_1340000 | 7.05 | -5.69 | 2.13e-04 | 4.91E-04 | PBANKA_1455200 | 9.99 | 5.11 | 1.58e-58 | 8.50E-57 |
| PBANKA_0713800 | 5.74 | -5.68 | 1.77e-04 | 4.17E-04 | PBANKA_1409400 | 7.68 | 5.1 | 2.64e-50 | 1.02E-48 |
| PBANKA_0203400 | 7.02 | -5.68 | 2.78e-04 | 6.28E-04 | PBANKA_0808700 | 9.9 | 5.09 | 7.02e-59 | 3.86E-57 |
| PBANKA_1331500 | 5.21 | -5.68 | 3.28e-04 | 7.32E-04 | PBANKA_1353700 | 9.78 | 5.09 | 2.93e-34 | 5.59E-33 |
| PBANKA_0719800 | 7.47 | -5.68 | 1.08e-03 | 2.17E-03 | PBANKA_1034300 | 10.68 | 5.09 | 5.24e-32 | 8.92E-31 |
| PBANKA_1139300 | 5.47 | -5.67 | 4.43e-04 | 9.61E-04 | PBANKA_1406700 | 11.32 | 5.09 | 1.46e-28 | 2.15E-27 |
| PBANKA_0106800 | 5.01 | -5.67 | 9.75e-04 | 1.97E-03 | PBANKA_0609100 | 10.58 | 5.08 | 2.12e-50 | 8.27E-49 |
| PBANKA_1313900 | 4.65 | -5.66 | 2.33e-04 | 5.33E-04 | PBANKA_0712000 | 9.63 | 5.07 | 8.08e-59 | 4.40E-57 |
| PBANKA_1215700 | 5.83 | -5.66 | 2.94e-04 | 6.61E-04 | PBANKA_1320500 | 11.85 | 5.07 | 1.02e-38 | 2.35E-37 |
| PBANKA_1326200 | 5.6 | -5.66 | 5.70e-04 | 1.20E-03 | PBANKA_0712400 | 8.17 | 5.05 | 5.49e-42 | 1.46E-40 |
| PBANKA_1137600 | 5.33 | -5.64 | 3.16e-04 | 7.08E-04 | PBANKA_1208600 | 8.79 | 5.05 | 1.13e-31 | 1.90E-30 |
| PBANKA_1011300 | 5.73 | -5.63 | 2.34e-04 | 5.35E-04 | PBANKA_1466121 | 4.44 | 5.05 | 1.70e-07 | 6.61E-07 |
| PBANKA_0523500 | 5.39 | -5.63 | 2.81e-04 | 6.35E-04 | PBANKA_1145800 | 12.01 | 5.04 | 1.41e-38 | 3.23E-37 |
| PBANKA_1209300 | 5.22 | -5.63 | 3.62e-04 | 8.00E-04 | PBANKA_1135100 | 10.88 | 5.04 | 3.57e-35 | 7.03E-34 |
| PBANKA_0940000 | 5.36 | -5.62 | 3.73e-04 | 8.21E-04 | PBANKA_1320600 | 5.74 | 5.04 | 1.09e-10 | 5.97E-10 |
| PBANKA_1361800 | 6.24 | -5.62 | 4.21e-04 | 9.17E-04 | PBANKA_0623500 | 9.6 | 5.03 | 2.63e-19 | 2.51E-18 |
| PBANKA_1411400 | 5.43 | -5.61 | 5.99e-04 | 1.26E-03 | PBANKA_0506300 | 7.64 | 5.01 | 1.60e-51 | 6.45E-50 |
| PBANKA_0811500 | 6.07 | -5.59 | 1.83e-04 | 4.28E-04 | PBANKA_1442500 | 9.3 | 5.01 | 7.51e-37 | 1.58E-35 |
| PBANKA_1430300 | 8.24 | -5.59 | 3.05e-04 | 6.86E-04 | PBANKA_0510900 | 11.33 | 4.98 | 3.95e-34 | 7.48E-33 |
| PBANKA_1439500 | 5.08 | -5.59 | 3.82e-04 | 8.39E-04 | PBANKA_0700051 | 4.7 | 4.98 | 3.91e-05 | 1.04E-04 |
| PBANKA_0310100 | 6.5 | -5.58 | 3.15e-04 | 7.06E-04 | PBANKA_1024100 | 9.85 | 4.97 | 7.19e-22 | 7.74E-21 |
| PBANKA_1210400 | 7.44 | -5.57 | 1.13e-13 | 7.94E-13 | PBANKA_0619100 | 11.3 | 4.96 | 1.11e-35 | 2.23E-34 |
| PBANKA_0612500 | 5.65 | -5.56 | 3.24e-04 | 7.23E-04 | PBANKA_1424900 | 7.1 | 4.96 | 8.60e-29 | 1.29E-27 |
| PBANKA_1025600 | 5.29 | -5.55 | 3.40e-04 | 7.55E-04 | PBANKA_1202400 | 11.97 | 4.95 | 3.72e-43 | 1.03E-41 |

| | | | | |
|---|---|---|---|---|
| PBANKA_0811800 | 5.56 | -5.54 | 5.28e-04 | 1.12E-03 |
| PBANKA_1240500 | 4.96 | -5.53 | 5.22e-04 | 1.11E-03 |
| PBANKA_1140900 | 6.31 | -5.52 | 5.44e-04 | 1.15E-03 |
| PBANKA_0829900 | 5.84 | -5.52 | 6.00e-04 | 1.26E-03 |
| PBANKA_0821400 | 7.8 | -5.52 | 5.46e-03 | 9.44E-03 |
| PBANKA_0400900 | 6.2 | -5.51 | 2.21e-04 | 5.09E-04 |
| PBANKA_1330000 | 6.11 | -5.51 | 5.29e-04 | 1.13E-03 |
| PBANKA_1003500 | 5.84 | -5.5 | 2.76e-04 | 6.24E-04 |
| PBANKA_1038600 | 5.48 | -5.5 | 3.38e-04 | 7.51E-04 |
| PBANKA_1223700 | 5.32 | -5.49 | 2.76e-04 | 6.24E-04 |
| PBANKA_1338000 | 6.19 | -5.49 | 4.45e-04 | 9.63E-04 |
| PBANKA_1427700 | 4.53 | -5.49 | 1.67e-03 | 3.21E-03 |
| PBANKA_0907000 | 8.28 | -5.48 | 3.19e-04 | 7.13E-04 |
| PBANKA_1026000 | 6.09 | -5.48 | 3.67e-04 | 8.10E-04 |
| PBANKA_1113600 | 6.03 | -5.47 | 1.21e-03 | 2.39E-03 |
| PBANKA_0607900 | 6.7 | -5.46 | 2.52e-04 | 5.75E-04 |
| PBANKA_1416000 | 4.9 | -5.46 | 6.74e-04 | 1.41E-03 |
| PBANKA_0904800 | 5.89 | -5.45 | 3.84e-04 | 8.43E-04 |
| PBANKA_1038100 | 5.96 | -5.45 | 5.20e-04 | 1.11E-03 |
| PBANKA_1402100 | 5.23 | -5.45 | 6.35e-04 | 1.33E-03 |
| PBANKA_1445000 | 8.67 | -5.43 | 2.74e-04 | 6.19E-04 |
| PBANKA_1214100 | 5.08 | -5.43 | 4.54e-04 | 9.80E-04 |
| PBANKA_1341400 | 5.18 | -5.43 | 1.19e-03 | 2.37E-03 |
| PBANKA_0312000 | 9.24 | -5.42 | 3.62e-04 | 8.00E-04 |
| PBANKA_1463600 | 5.31 | -5.42 | 4.81e-04 | 1.03E-03 |
| PBANKA_1462100 | 5.31 | -5.42 | 5.78e-04 | 1.22E-03 |
| PBANKA_1335700 | 5.88 | -5.42 | 6.64e-04 | 1.39E-03 |
| PBANKA_0707500 | 6.84 | -5.41 | 3.27e-04 | 7.30E-04 |
| PBANKA_0104200 | 5.49 | -5.4 | 8.52e-04 | 1.75E-03 |
| PBANKA_0717600 | 7.03 | -5.39 | 3.12e-04 | 6.99E-04 |
| PBANKA_0710700 | 7.23 | -5.39 | 3.34e-04 | 7.43E-04 |
| PBANKA_1130300 | 6.7 | -5.39 | 3.67e-04 | 8.09E-04 |
| PBANKA_1139800 | 6.13 | -5.39 | 4.61e-04 | 9.95E-04 |
| PBANKA_0711500 | 6.47 | -5.39 | 6.10e-04 | 1.28E-03 |
| PBANKA_0913500 | 5.12 | -5.39 | 1.04e-03 | 2.10E-03 |
| PBANKA_0708900 | 5.48 | -5.39 | 1.18e-03 | 2.36E-03 |
| PBANKA_0817800 | 4.61 | -5.37 | 9.11e-04 | 1.85E-03 |
| PBANKA_0830700 | 7 | -5.36 | 4.50e-04 | 9.73E-04 |
| PBANKA_1106000 | 6.99 | -5.36 | 6.88e-04 | 1.43E-03 |
| PBANKA_1363900 | 6.02 | -5.35 | 5.53e-04 | 1.17E-03 |
| PBANKA_1353200 | 5.19 | -5.35 | 7.46e-04 | 1.55E-03 |
| PBANKA_1300500 | 5.24 | -5.35 | 8.04e-04 | 1.66E-03 |
| PBANKA_1432600 | 5.62 | -5.34 | 4.86e-04 | 1.04E-03 |
| PBANKA_0409900 | 6.24 | -5.34 | 7.81e-04 | 1.61E-03 |
| PBANKA_1036200 | 5.97 | -5.32 | 7.25e-04 | 1.50E-03 |
| PBANKA_1327080 | 5.13 | -5.31 | 1.09e-03 | 2.18E-03 |
| PBANKA_1411700 | 4.66 | -5.3 | 1.15e-03 | 2.29E-03 |
| PBANKA_0719600 | 6.89 | -5.29 | 4.69e-04 | 1.01E-03 |
| PBANKA_0509800 | 5.31 | -5.28 | 5.81e-04 | 1.23E-03 |
| PBANKA_0718600 | 5.74 | -5.27 | 5.65e-04 | 1.20E-03 |
| PBANKA_1442800 | 5.45 | -5.27 | 7.87e-04 | 1.63E-03 |
| PBANKA_0416900 | 5.37 | -5.27 | 1.46e-03 | 2.86E-03 |
| PBANKA_0105100 | 7.22 | -5.25 | 5.37e-04 | 1.14E-03 |
| PBANKA_1311100 | 5.28 | -5.25 | 7.72e-04 | 1.60E-03 |
| PBANKA_0832700 | 5.53 | -5.25 | 2.37e-03 | 4.43E-03 |
| PBANKA_0201000 | 5.45 | -5.23 | 1.03e-03 | 2.07E-03 |
| PBANKA_0111500 | 4.32 | -5.23 | 4.85e-03 | 8.48E-03 |
| PBANKA_1463000 | 10.63 | -5.22 | 3.16e-05 | 8.56E-05 |
| PBANKA_0103000 | 5.01 | -5.21 | 1.47e-03 | 2.86E-03 |
| PBANKA_1119800 | 5.5 | -5.21 | 1.59e-03 | 3.08E-03 |
| PBANKA_1111700 | 5.58 | -5.2 | 1.03e-03 | 2.07E-03 |
| PBANKA_0510600 | 5.98 | -5.19 | 6.87e-04 | 1.43E-03 |
| PBANKA_1361700 | 4.42 | -5.19 | 1.49e-03 | 2.90E-03 |
| PBANKA_1036700 | 6.49 | -5.18 | 8.80e-04 | 1.80E-03 |
| PBANKA_1316500 | 5.64 | -5.18 | 1.39e-03 | 2.73E-03 |
| PBANKA_1125400 | 4.79 | -5.17 | 1.40e-03 | 2.75E-03 |
| PBANKA_1413300 | 4.75 | -5.17 | 1.56e-03 | 3.02E-03 |
| PBANKA_1205300 | 5.63 | -5.16 | 1.58e-03 | 3.07E-03 |
| PBANKA_0921000 | 4.67 | -5.15 | 1.15e-03 | 2.29E-03 |
| PBANKA_0942200 | 6.68 | -5.14 | 5.43e-04 | 1.15E-03 |
| PBANKA_1360600 | 4.97 | -5.13 | 1.51e-03 | 2.94E-03 |
| PBANKA_1317200 | 10.24 | -5.12 | 1.62e-09 | 7.87E-09 |
| PBANKA_1410900 | 6.53 | -5.12 | 7.03e-04 | 1.46E-03 |
| PBANKA_1444300 | 4.94 | -5.12 | 1.30e-03 | 2.56E-03 |

| | | | | |
|---|---|---|---|---|
| PBANKA_0506200 | 11.37 | 4.95 | 5.22e-40 | 1.27E-38 |
| PBANKA_1321000 | 9.36 | 4.94 | 3.31e-117 | 2.12E-114 |
| PBANKA_1311800 | 9.56 | 4.94 | 2.57e-45 | 7.91E-44 |
| PBANKA_0915300 | 10.03 | 4.94 | 4.20e-35 | 8.17E-34 |
| PBANKA_1408800 | 6.6 | 4.93 | 3.14e-22 | 3.44E-21 |
| PBANKA_0400600 | 4.82 | 4.93 | 5.72e-09 | 2.64E-08 |
| PBANKA_0214800 | 10.11 | 4.92 | 6.11e-42 | 1.60E-40 |
| PBANKA_1134400 | 10.57 | 4.91 | 7.71e-23 | 8.80E-22 |
| PBANKA_1360100 | 10.03 | 4.9 | 1.95e-27 | 2.75E-26 |
| PBANKA_1400031 | 5.21 | 4.9 | 4.46e-03 | 7.83E-03 |
| PBANKA_1005100 | 7.89 | 4.89 | 1.12e-62 | 7.17E-61 |
| PBANKA_1025200 | 9.08 | 4.89 | 4.84e-50 | 1.86E-48 |
| PBANKA_1228700 | 8.88 | 4.89 | 3.27e-43 | 9.18E-42 |
| PBANKA_0100500 | 9.28 | 4.87 | 4.74e-44 | 1.40E-42 |
| PBANKA_1137100 | 7.66 | 4.87 | 3.54e-42 | 9.49E-41 |
| PBANKA_0836500 | 5.57 | 4.87 | 4.40e-09 | 2.05E-08 |
| PBANKA_0317161 | 5.14 | 4.86 | 3.85e-05 | 1.02E-04 |
| PBANKA_1337000 | 9.96 | 4.83 | 1.63e-57 | 8.57E-56 |
| PBANKA_1120200 | 7.03 | 4.83 | 1.83e-29 | 2.77E-28 |
| PBANKA_1103400 | 10.66 | 4.83 | 1.03e-22 | 1.17E-21 |
| PBANKA_1017900 | 9.38 | 4.82 | 4.11e-59 | 2.29E-57 |
| PBANKA_1200800 | 10.18 | 4.82 | 1.50e-33 | 2.78E-32 |
| PBANKA_1020400 | 8.99 | 4.81 | 1.50e-49 | 5.72E-48 |
| PBANKA_0918800 | 9.38 | 4.8 | 7.06e-91 | 1.06E-88 |
| PBANKA_0810500 | 8.22 | 4.8 | 8.80e-24 | 1.05E-22 |
| PBANKA_1127000 | 12.62 | 4.79 | 5.26e-29 | 7.92E-28 |
| PBANKA_1007000 | 8.87 | 4.78 | 1.30e-54 | 6.08E-53 |
| PBANKA_0623651 | 4.62 | 4.78 | 9.55e-07 | 3.34E-06 |
| PBANKA_0316921 | 7.78 | 4.77 | 8.22e-39 | 1.90E-37 |
| PBANKA_1364200 | 11.25 | 4.77 | 1.11e-28 | 1.65E-27 |
| PBANKA_0824800 | 8.81 | 4.77 | 3.71e-28 | 5.39E-27 |
| PBANKA_0617100 | 11.12 | 4.75 | 2.97e-43 | 8.43E-42 |
| PBANKA_0001001 | 5.28 | 4.75 | 7.15e-10 | 3.64E-09 |
| PBANKA_1362600 | 7.99 | 4.73 | 5.69e-65 | 3.93E-63 |
| PBANKA_0505700 | 10.1 | 4.72 | 3.94e-103 | 1.06E-100 |
| PBANKA_0400100 | 5.29 | 4.72 | 1.11e-09 | 5.49E-09 |
| PBANKA_0302300 | 7.78 | 4.71 | 7.75e-57 | 3.85E-55 |
| PBANKA_1439200 | 14.34 | 4.7 | 2.86e-53 | 1.25E-51 |
| PBANKA_0417500 | 10.82 | 4.7 | 1.74e-45 | 5.40E-44 |
| PBANKA_0918000 | 12.18 | 4.7 | 3.60e-35 | 7.06E-34 |
| PBANKA_0400500 | 8.51 | 4.7 | 1.25e-33 | 2.33E-32 |
| PBANKA_1432800 | 9.63 | 4.7 | 3.92e-31 | 6.42E-30 |
| PBANKA_0819700 | 8.67 | 4.69 | 2.43e-67 | 1.82E-65 |
| PBANKA_1120700 | 10.01 | 4.69 | 1.99e-27 | 2.79E-26 |
| PBANKA_1232100 | 10.79 | 4.69 | 2.72e-26 | 3.61E-25 |
| PBANKA_1417800 | 8.97 | 4.68 | 7.22e-41 | 1.85E-39 |
| PBANKA_0924000 | 9 | 4.68 | 5.87e-39 | 1.37E-37 |
| PBANKA_1424800 | 6.61 | 4.68 | 2.11e-38 | 4.80E-37 |
| PBANKA_1428900 | 11.03 | 4.67 | 6.44e-28 | 9.17E-27 |
| PBANKA_0316841 | 7.44 | 4.65 | 8.79e-22 | 9.44E-21 |
| PBANKA_1122500 | 9.15 | 4.64 | 1.23e-66 | 9.00E-65 |
| PBANKA_1103000 | 10.11 | 4.64 | 5.89e-48 | 2.09E-46 |
| PBANKA_1356200 | 8.57 | 4.64 | 1.47e-39 | 3.55E-38 |
| PBANKA_1354400 | 11.67 | 4.64 | 2.48e-38 | 5.58E-37 |
| PBANKA_0205100 | 8.09 | 4.64 | 9.84e-34 | 1.85E-32 |
| PBANKA_1130900 | 11.47 | 4.64 | 2.77e-22 | 3.04E-21 |
| PBANKA_0317061 | 6.53 | 4.64 | 2.54e-14 | 1.83E-13 |
| PBANKA_1466000 | 5.35 | 4.63 | 1.26e-08 | 5.59E-08 |
| PBANKA_1412100 | 6.72 | 4.62 | 5.67e-32 | 9.63E-31 |
| PBANKA_1452300 | 9.76 | 4.62 | 1.51e-24 | 1.85E-23 |
| PBANKA_1434100 | 9.96 | 4.61 | 5.77e-46 | 1.87E-44 |
| PBANKA_1416300 | 10.7 | 4.61 | 1.42e-43 | 4.09E-42 |
| PBANKA_1317900 | 9.35 | 4.61 | 2.38e-38 | 5.39E-37 |
| PBANKA_1035100 | 9.89 | 4.61 | 3.73e-34 | 7.10E-33 |
| PBANKA_1036300 | 9.82 | 4.6 | 1.03e-32 | 1.79E-31 |
| PBANKA_0100300 | 6.29 | 4.6 | 1.57e-10 | 8.50E-10 |
| PBANKA_1441700 | 9.6 | 4.59 | 3.32e-40 | 8.16E-39 |
| PBANKA_1019320 | 5.31 | 4.59 | 5.65e-07 | 2.03E-06 |
| PBANKA_0516900 | 11.8 | 4.58 | 1.02e-62 | 6.72E-61 |
| PBANKA_1019700 | 11.42 | 4.58 | 3.03e-26 | 4.00E-25 |
| PBANKA_1365600 | 7.95 | 4.57 | 1.47e-22 | 1.65E-21 |
| PBANKA_0112661 | 8.05 | 4.57 | 1.65e-06 | 5.52E-06 |
| PBANKA_1200600 | 12.01 | 4.55 | 5.09e-10 | 2.62E-09 |
| PBANKA_0921900 | 9.68 | 4.54 | 4.76e-52 | 1.96E-50 |

| | | | | |
|---|---|---|---|---|
| PBANKA_1453900 | 6.32 | -5.12 | 1.44e-03 | 2.82E-03 |
| PBANKA_0718700 | 5.67 | -5.11 | 1.23e-03 | 2.45E-03 |
| PBANKA_1227500 | 5.59 | -5.1 | 1.02e-03 | 2.06E-03 |
| PBANKA_1120500 | 4.8 | -5.1 | 2.61e-03 | 4.82E-03 |
| PBANKA_0608000 | 8.46 | -5.09 | 8.80e-04 | 1.80E-03 |
| PBANKA_0716300 | 5.81 | -5.09 | 1.46e-03 | 2.85E-03 |
| PBANKA_0620700 | 6.88 | -5.07 | 1.06e-03 | 2.13E-03 |
| PBANKA_1331600 | 4.76 | -5.07 | 2.11e-03 | 3.97E-03 |
| PBANKA_0515100 | 4.84 | -5.04 | 2.18e-03 | 4.09E-03 |
| PBANKA_0522200 | 7.78 | -5.03 | 1.01e-03 | 2.03E-03 |
| PBANKA_0407300 | 5.41 | -5.03 | 1.75e-03 | 3.36E-03 |
| PBANKA_0834500 | 5.92 | -5.02 | 1.14e-03 | 2.28E-03 |
| PBANKA_0514900 | 13.74 | -5.01 | 1.07e-10 | 5.89E-10 |
| PBANKA_0315500 | 6.17 | -5.01 | 8.71e-04 | 1.78E-03 |
| PBANKA_1112600 | 6.24 | -5.01 | 1.05e-03 | 2.11E-03 |
| PBANKA_1143100 | 5.18 | -5.01 | 1.99e-03 | 3.76E-03 |
| PBANKA_1318800 | 6.46 | -4.99 | 1.49e-03 | 2.90E-03 |
| PBANKA_0507700 | 4.87 | -4.99 | 2.91e-03 | 5.31E-03 |
| PBANKA_0706900 | 4.9 | -4.98 | 4.74e-03 | 8.28E-03 |
| PBANKA_0824600 | 4.59 | -4.96 | 3.75e-03 | 6.67E-03 |
| PBANKA_0515000 | 13.15 | -4.95 | 5.30e-08 | 2.18E-07 |
| PBANKA_0913600 | 5.31 | -4.94 | 1.21e-03 | 2.40E-03 |
| PBANKA_1329900 | 6.61 | -4.94 | 1.67e-03 | 3.22E-03 |
| PBANKA_1424700 | 4.59 | -4.94 | 3.13e-03 | 5.68E-03 |
| PBANKA_1110600 | 6.99 | -4.93 | 1.65e-03 | 3.18E-03 |
| PBANKA_0412700 | 5.75 | -4.92 | 1.82e-03 | 3.47E-03 |
| PBANKA_1129600 | 7.8 | -4.91 | 1.05e-05 | 3.12E-05 |
| PBANKA_0517900 | 4.38 | -4.91 | 2.90e-03 | 5.30E-03 |
| PBANKA_1303100 | 5.27 | -4.9 | 1.40e-03 | 2.76E-03 |
| PBANKA_0201900 | 5.05 | -4.9 | 1.59e-03 | 3.09E-03 |
| PBANKA_0214951 | 4 | -4.9 | 5.72e-03 | 9.87E-03 |
| PBANKA_0909700 | 5.26 | -4.87 | 2.01e-03 | 3.80E-03 |
| PBANKA_0500721 | 4.6 | -4.86 | 2.19e-03 | 4.12E-03 |
| PBANKA_0606500 | 4.7 | -4.86 | 2.48e-03 | 4.60E-03 |
| PBANKA_1105000 | 7.94 | -4.85 | 1.79e-13 | 1.24E-12 |
| PBANKA_0804200 | 5.74 | -4.85 | 1.47e-03 | 2.87E-03 |
| PBANKA_1009000 | 6.49 | -4.85 | 1.66e-03 | 3.20E-03 |
| PBANKA_1417700 | 4.17 | -4.85 | 5.36e-03 | 9.27E-03 |
| PBANKA_1032000 | 5.67 | -4.84 | 2.05e-03 | 3.87E-03 |
| PBANKA_1354200 | 5 | -4.84 | 2.37e-03 | 4.43E-03 |
| PBANKA_1429800 | 5.35 | -4.81 | 1.77e-03 | 3.39E-03 |
| PBANKA_0303000 | 5.22 | -4.79 | 2.66e-03 | 4.90E-03 |
| PBANKA_1208900 | 5.56 | -4.79 | 3.15e-03 | 5.71E-03 |
| PBANKA_1325200 | 5.65 | -4.79 | 4.66e-03 | 8.15E-03 |
| PBANKA_0932300 | 6.6 | -4.78 | 1.98e-03 | 3.75E-03 |
| PBANKA_0616400 | 8.11 | -4.77 | 2.26e-03 | 4.23E-03 |
| PBANKA_0507900 | 4.77 | -4.77 | 3.23e-03 | 5.85E-03 |
| PBANKA_0906000 | 5.34 | -4.76 | 1.87e-03 | 3.57E-03 |
| PBANKA_0808500 | 4.68 | -4.76 | 2.66e-03 | 4.91E-03 |
| PBANKA_0602200 | 5.14 | -4.75 | 2.04e-03 | 3.86E-03 |
| PBANKA_0607500 | 5.19 | -4.74 | 2.95e-03 | 5.37E-03 |
| PBANKA_0904900 | 5.11 | -4.74 | 3.32e-03 | 6.00E-03 |
| PBANKA_0826300 | 5.44 | -4.7 | 2.72e-03 | 5.00E-03 |
| PBANKA_0612900 | 5.74 | -4.7 | 4.35e-03 | 7.66E-03 |
| PBANKA_0417400 | 8.01 | -4.69 | 7.47e-15 | 5.59E-14 |
| PBANKA_1463900 | 5.77 | -4.69 | 3.61e-03 | 6.46E-03 |
| PBANKA_1337900 | 6.61 | -4.67 | 1.79e-03 | 3.43E-03 |
| PBANKA_0600700 | 5.76 | -4.66 | 2.84e-03 | 5.21E-03 |
| PBANKA_0310200 | 4.86 | -4.65 | 5.18e-03 | 8.99E-03 |
| PBANKA_1434300 | 5.36 | -4.62 | 2.43e-03 | 4.52E-03 |
| PBANKA_1106200 | 5.26 | -4.62 | 2.94e-03 | 5.37E-03 |
| PBANKA_0612000 | 5.28 | -4.61 | 2.16e-03 | 4.06E-03 |
| PBANKA_1123400 | 5.65 | -4.61 | 2.58e-03 | 4.77E-03 |
| PBANKA_0909000 | 4.73 | -4.61 | 4.35e-03 | 7.66E-03 |
| PBANKA_0504300 | 5.78 | -4.61 | 4.60e-03 | 8.06E-03 |
| PBANKA_1016700 | 10.48 | -4.6 | 3.66e-24 | 4.44E-23 |
| PBANKA_0605800 | 7.14 | -4.58 | 3.18e-03 | 5.77E-03 |
| PBANKA_1314600 | 5.47 | -4.57 | 3.23e-03 | 5.85E-03 |
| PBANKA_1430500 | 5.86 | -4.56 | 2.47e-03 | 4.58E-03 |
| PBANKA_1302100 | 4.92 | -4.56 | 4.00e-03 | 7.09E-03 |
| PBANKA_1449200 | 8.34 | -4.55 | 1.88e-13 | 1.30E-12 |
| PBANKA_1404200 | 5.1 | -4.55 | 3.24e-03 | 5.86E-03 |
| PBANKA_1126100 | 5.19 | -4.55 | 3.67e-03 | 6.54E-03 |
| PBANKA_1021400 | 6.17 | -4.54 | 3.60e-06 | 1.15E-05 |

| | | | | |
|---|---|---|---|---|
| PBANKA_0922100 | 11.02 | 4.54 | 5.66e-28 | 8.11E-27 |
| PBANKA_1101200 | 11.35 | 4.54 | 2.28e-21 | 2.41E-20 |
| PBANKA_1327700 | 7.3 | 4.53 | 3.66e-33 | 6.66E-32 |
| PBANKA_1243400 | 8.51 | 4.52 | 7.31e-54 | 3.25E-52 |
| PBANKA_0703900 | 11.44 | 4.52 | 7.08e-31 | 1.14E-29 |
| PBANKA_0514000 | 7.33 | 4.52 | 8.15e-19 | 7.54E-18 |
| PBANKA_1239700 | 8.18 | 4.51 | 1.21e-45 | 3.79E-44 |
| PBANKA_0613800 | 10.27 | 4.51 | 2.90e-26 | 3.84E-25 |
| PBANKA_1465100 | 7.21 | 4.51 | 2.72e-03 | 5.01E-03 |
| PBANKA_0924900 | 10.29 | 4.5 | 4.36e-70 | 3.91E-68 |
| PBANKA_1338300 | 10.44 | 4.5 | 1.24e-29 | 1.89E-28 |
| PBANKA_0617700 | 7.43 | 4.48 | 6.14e-28 | 8.77E-27 |
| PBANKA_1423600 | 10.4 | 4.48 | 4.71e-27 | 6.51E-26 |
| PBANKA_0517500 | 10.08 | 4.47 | 1.29e-45 | 4.02E-44 |
| PBANKA_0902100 | 11.45 | 4.47 | 7.43e-33 | 1.32E-31 |
| PBANKA_0615100 | 9.04 | 4.47 | 9.28e-30 | 1.43E-28 |
| PBANKA_1106700 | 12.53 | 4.46 | 4.86e-69 | 4.14E-67 |
| PBANKA_1461100 | 9.88 | 4.46 | 9.35e-49 | 3.37E-47 |
| PBANKA_1355800 | 6.34 | 4.45 | 2.28e-16 | 1.84E-15 |
| PBANKA_0617600 | 9.26 | 4.44 | 3.18e-59 | 1.79E-57 |
| PBANKA_1143300 | 9.46 | 4.44 | 3.46e-57 | 1.79E-55 |
| PBANKA_0521261 | 16.05 | 4.44 | 1.41e-10 | 7.69E-10 |
| PBANKA_0112500 | 9.27 | 4.43 | 2.81e-62 | 1.69E-60 |
| PBANKA_1104600 | 9.65 | 4.43 | 1.11e-31 | 1.87E-30 |
| PBANKA_1407650 | 4.04 | 4.43 | 8.27e-04 | 1.70E-03 |
| PBANKA_1356000 | 10.7 | 4.42 | 3.00e-11 | 1.73E-10 |
| PBANKA_0921200 | 8.52 | 4.42 | 2.87e-03 | 5.25E-03 |
| PBANKA_1464600 | 8.81 | 4.4 | 7.61e-32 | 1.29E-30 |
| PBANKA_1310800 | 11.54 | 4.39 | 5.77e-26 | 7.47E-25 |
| PBANKA_0817900 | 9.69 | 4.38 | 1.11e-68 | 9.34E-67 |
| PBANKA_1231700 | 11.97 | 4.38 | 1.17e-40 | 2.92E-39 |
| PBANKA_1329300 | 11.42 | 4.38 | 2.56e-35 | 5.08E-34 |
| PBANKA_0926700 | 12.55 | 4.38 | 1.30e-14 | 9.63E-14 |
| PBANKA_1313400 | 9.24 | 4.37 | 5.71e-43 | 1.58E-41 |
| PBANKA_1140600 | 8.42 | 4.37 | 1.59e-21 | 1.70E-20 |
| PBANKA_1346100 | 8.3 | 4.35 | 5.07e-91 | 8.10E-89 |
| PBANKA_1037600 | 8.81 | 4.35 | 2.59e-69 | 2.24E-67 |
| PBANKA_1365500 | 11.7 | 4.35 | 1.70e-26 | 2.28E-25 |
| PBANKA_1446200 | 6.12 | 4.35 | 2.12e-16 | 1.72E-15 |
| PBANKA_1457400 | 8.91 | 4.34 | 3.27e-37 | 6.97E-36 |
| PBANKA_0412200 | 8.64 | 4.34 | 1.18e-35 | 2.36E-34 |
| PBANKA_0823100 | 11.22 | 4.34 | 4.02e-20 | 3.97E-19 |
| PBANKA_0700900 | 7.97 | 4.34 | 2.23e-18 | 2.01E-17 |
| PBANKA_0518800 | 8.3 | 4.33 | 9.26e-29 | 1.38E-27 |
| PBANKA_0206600 | 10.11 | 4.33 | 2.61e-18 | 2.34E-17 |
| PBANKA_0805100 | 9.32 | 4.32 | 4.51e-61 | 2.59E-59 |
| PBANKA_1008500 | 12.73 | 4.32 | 1.14e-21 | 1.22E-20 |
| PBANKA_1342800 | 7.77 | 4.31 | 6.61e-38 | 1.44E-36 |
| PBANKA_1120300 | 7.28 | 4.31 | 3.12e-21 | 3.28E-20 |
| PBANKA_0936600 | 8.9 | 4.3 | 3.48e-33 | 6.35E-32 |
| PBANKA_0613700 | 11.21 | 4.3 | 1.07e-15 | 8.46E-15 |
| PBANKA_0816400 | 10.86 | 4.29 | 1.71e-71 | 1.62E-69 |
| PBANKA_1010200 | 9.43 | 4.29 | 1.87e-47 | 6.43E-46 |
| PBANKA_0214900 | 6.49 | 4.29 | 1.84e-11 | 1.08E-10 |
| PBANKA_1226300 | 8.6 | 4.28 | 9.04e-30 | 1.39E-28 |
| PBANKA_1116000 | 10.99 | 4.28 | 2.26e-22 | 2.49E-21 |
| PBANKA_1305100 | 10.64 | 4.26 | 1.23e-31 | 2.05E-30 |
| PBANKA_1013100 | 10.1 | 4.26 | 2.74e-27 | 3.81E-26 |
| PBANKA_0923400 | 10.04 | 4.26 | 7.10e-19 | 6.59E-18 |
| PBANKA_0829700 | 6.98 | 4.25 | 1.28e-24 | 1.57E-23 |
| PBANKA_0316821 | 5.81 | 4.25 | 7.89e-18 | 6.92E-17 |
| PBANKA_1201900 | 9.85 | 4.24 | 3.80e-23 | 4.40E-22 |
| PBANKA_0600041 | 6.6 | 4.24 | 1.74e-04 | 4.10E-04 |
| PBANKA_1224200 | 10.9 | 4.23 | 3.97e-40 | 9.67E-39 |
| PBANKA_0405400 | 10.67 | 4.21 | 1.68e-26 | 2.26E-25 |
| PBANKA_1207900 | 10.1 | 4.21 | 3.75e-21 | 3.93E-20 |
| PBANKA_0201500 | 9.96 | 4.2 | 2.32e-40 | 5.75E-39 |
| PBANKA_1145700 | 7.33 | 4.2 | 1.82e-22 | 2.03E-21 |
| PBANKA_1234800 | 10.79 | 4.2 | 7.49e-20 | 7.28E-19 |
| PBANKA_1354600 | 7.72 | 4.2 | 3.46e-14 | 2.48E-13 |
| PBANKA_1347600 | 8.21 | 4.19 | 2.83e-42 | 7.61E-41 |
| PBANKA_0518700 | 10.69 | 4.19 | 3.38e-35 | 6.68E-34 |
| PBANKA_1125600 | 11.74 | 4.19 | 8.88e-26 | 1.14E-24 |
| PBANKA_0806100 | 8.66 | 4.18 | 2.52e-68 | 1.99E-66 |

| | | | | |
|---|---|---|---|---|
| PBANKA_1427200 | 8.15 | -4.52 | 8.60e-15 | 6.43E-14 |
| PBANKA_0206000 | 7.45 | -4.52 | 6.40e-12 | 3.91E-11 |
| PBANKA_0612400 | 11.94 | -4.52 | 8.82e-12 | 5.32E-11 |
| PBANKA_1312600 | 4.94 | -4.52 | 4.13e-03 | 7.32E-03 |
| PBANKA_0907500 | 5 | -4.51 | 5.72e-03 | 9.87E-03 |
| PBANKA_1334900 | 10.27 | -4.5 | 1.20e-08 | 5.32E-08 |
| PBANKA_1462300 | 5.15 | -4.5 | 4.05e-03 | 7.17E-03 |
| PBANKA_0607100 | 5.59 | -4.49 | 4.62e-03 | 8.09E-03 |
| PBANKA_1421500 | 10.15 | -4.47 | 2.65e-08 | 1.14E-07 |
| PBANKA_0601900 | 10.03 | -4.45 | 1.09e-27 | 1.54E-26 |
| PBANKA_0621100 | 6 | -4.43 | 5.13e-03 | 8.92E-03 |
| PBANKA_0711400 | 10.06 | -4.42 | 3.14e-03 | 5.70E-03 |
| PBANKA_1204700 | 5.96 | -4.42 | 3.68e-03 | 6.56E-03 |
| PBANKA_0616000 | 5.23 | -4.41 | 4.45e-03 | 7.81E-03 |
| PBANKA_0924600 | 5.54 | -4.38 | 4.88e-03 | 8.52E-03 |
| PBANKA_1404700 | 8.5 | -4.37 | 5.66e-17 | 4.77E-16 |
| PBANKA_0603100 | 5.36 | -4.33 | 5.11e-03 | 8.89E-03 |
| PBANKA_1319900 | 6.03 | -4.32 | 4.46e-03 | 7.83E-03 |
| PBANKA_1115000 | 10.68 | -4.29 | 1.78e-09 | 8.64E-09 |
| PBANKA_0916700 | 10.4 | -4.29 | 3.26e-08 | 1.38E-07 |
| PBANKA_0943400 | 10.42 | -4.26 | 4.25e-09 | 1.98E-08 |
| PBANKA_0820600 | 6.85 | -4.26 | 4.15e-03 | 7.34E-03 |
| PBANKA_1227400 | 8.43 | -4.25 | 1.15e-12 | 7.45E-12 |
| PBANKA_0908700 | 7.3 | -4.25 | 7.80e-09 | 3.56E-08 |
| PBANKA_0706800 | 8.12 | -4.24 | 1.79e-20 | 1.82E-19 |
| PBANKA_1021500 | 7.04 | -4.21 | 2.99e-08 | 1.27E-07 |
| PBANKA_1017400 | 5.72 | -4.2 | 1.78e-06 | 5.90E-06 |
| PBANKA_1133500 | 7.02 | -4.19 | 9.92e-09 | 4.48E-08 |
| PBANKA_0619000 | 7.62 | -4.15 | 1.62e-19 | 1.56E-18 |
| PBANKA_0522700 | 9.87 | -4.11 | 4.48e-16 | 3.57E-15 |
| PBANKA_0204500 | 8.9 | -4.11 | 1.47e-09 | 7.18E-09 |
| PBANKA_1334800 | 10.29 | -4.11 | 5.58e-09 | 2.58E-08 |
| PBANKA_0836700 | 6.53 | -4.11 | 1.17e-08 | 5.21E-08 |
| PBANKA_1210300 | 7.45 | -4.11 | 2.52e-07 | 9.56E-07 |
| PBANKA_0925900 | 8.18 | -4.09 | 1.24e-13 | 8.67E-13 |
| PBANKA_1240000 | 9.47 | -4.08 | 3.99e-35 | 7.79E-34 |
| PBANKA_0609500 | 11.69 | -4.04 | 5.62e-08 | 2.31E-07 |
| PBANKA_1204200 | 10.71 | -4.03 | 3.54e-09 | 1.66E-08 |
| PBANKA_1017300 | 9.6 | -4.01 | 1.27e-31 | 2.10E-30 |
| PBANKA_1036600 | 8.8 | -4 | 9.83e-14 | 6.93E-13 |
| PBANKA_0836300 | 9.24 | -3.96 | 3.57e-26 | 4.70E-25 |
| PBANKA_1027300 | 9.39 | -3.92 | 4.23e-25 | 5.27E-24 |
| PBANKA_1361400 | 7.2 | -3.9 | 6.72e-10 | 3.42E-09 |
| PBANKA_1335900 | 7.72 | -3.89 | 9.98e-13 | 6.53E-12 |
| PBANKA_1115800 | 7.11 | -3.89 | 6.56e-08 | 2.67E-07 |
| PBANKA_1450600 | 6.61 | -3.89 | 7.88e-07 | 2.79E-06 |
| PBANKA_1341300 | 6.8 | -3.89 | 1.23e-06 | 4.22E-06 |
| PBANKA_0830200 | 11.71 | -3.87 | 1.19e-33 | 2.23E-32 |
| PBANKA_0301600 | 10.23 | -3.85 | 1.19e-07 | 4.71E-07 |
| PBANKA_1235300 | 6.39 | -3.85 | 5.65e-07 | 2.03E-06 |
| PBANKA_0934300 | 8.02 | -3.83 | 1.18e-12 | 7.67E-12 |
| PBANKA_0207100 | 6.1 | -3.83 | 5.63e-06 | 1.75E-05 |
| PBANKA_1359700 | 6.59 | -3.83 | 1.13e-04 | 2.75E-04 |
| PBANKA_0925700 | 6.96 | -3.81 | 3.51e-08 | 1.48E-07 |
| PBANKA_1314000 | 6.71 | -3.8 | 3.19e-07 | 1.19E-06 |
| PBANKA_1225000 | 9.91 | -3.78 | 1.55e-30 | 2.46E-29 |
| PBANKA_1453700 | 10.09 | -3.78 | 8.86e-19 | 8.16E-18 |
| PBANKA_1302900 | 7.21 | -3.78 | 8.72e-10 | 4.37E-09 |
| PBANKA_1302700 | 7.48 | -3.78 | 3.58e-05 | 9.58E-05 |
| PBANKA_1411900 | 7.69 | -3.77 | 2.84e-24 | 3.46E-23 |
| PBANKA_0915200 | 9.53 | -3.77 | 6.70e-18 | 5.92E-17 |
| PBANKA_0411500 | 7.4 | -3.77 | 4.62e-10 | 2.39E-09 |
| PBANKA_0203750 | 7.23 | -3.77 | 7.66e-08 | 3.10E-07 |
| PBANKA_1354900 | 10.56 | -3.77 | 1.10e-06 | 3.81E-06 |
| PBANKA_0301300 | 8.92 | -3.76 | 3.82e-08 | 1.60E-07 |
| PBANKA_1364300 | 9.57 | -3.75 | 8.91e-12 | 5.38E-11 |
| PBANKA_1034800 | 7.08 | -3.75 | 5.77e-06 | 1.79E-05 |
| PBANKA_0608500 | 7.62 | -3.74 | 3.51e-08 | 1.48E-07 |
| PBANKA_0715100 | 6.27 | -3.72 | 1.10e-06 | 3.80E-06 |
| PBANKA_0812700 | 8.41 | -3.71 | 3.18e-19 | 3.01E-18 |
| PBANKA_0507800 | 7.28 | -3.7 | 8.52e-07 | 3.00E-06 |
| PBANKA_1454000 | 8.08 | -3.68 | 1.26e-20 | 1.29E-19 |
| PBANKA_1144400 | 7.32 | -3.67 | 3.61e-17 | 3.07E-16 |
| PBANKA_1431600 | 7.48 | -3.67 | 2.36e-13 | 1.62E-12 |

| | | | | |
|---|---|---|---|---|
| PBANKA_0112600 | 9.77 | 4.18 | 2.74e-41 | 7.07E-40 |
| PBANKA_0503600 | 9.73 | 4.18 | 5.26e-24 | 6.36E-23 |
| PBANKA_0521300 | 11.47 | 4.18 | 7.34e-10 | 3.72E-09 |
| PBANKA_0917700 | 8.69 | 4.17 | 9.93e-41 | 2.50E-39 |
| PBANKA_0831900 | 9.29 | 4.15 | 1.63e-62 | 1.03E-60 |
| PBANKA_1450100 | 8.2 | 4.14 | 8.67e-43 | 2.37E-41 |
| PBANKA_1022000 | 10.31 | 4.14 | 5.32e-20 | 5.24E-19 |
| PBANKA_0607300 | 9.74 | 4.13 | 5.34e-45 | 1.63E-43 |
| PBANKA_0713600 | 7.61 | 4.13 | 2.94e-40 | 7.26E-39 |
| PBANKA_1245800 | 10.01 | 4.12 | 1.88e-39 | 4.50E-38 |
| PBANKA_1329800 | 11.14 | 4.12 | 1.36e-35 | 2.72E-34 |
| PBANKA_1201700 | 6.62 | 4.12 | 4.41e-27 | 6.12E-26 |
| PBANKA_0315600 | 9.92 | 4.12 | 1.14e-16 | 9.41E-16 |
| PBANKA_1464500 | 10.32 | 4.11 | 1.21e-37 | 2.61E-36 |
| PBANKA_0500971 | 4.69 | 4.11 | 9.02e-06 | 2.70E-05 |
| PBANKA_0102200 | 9.56 | 4.1 | 4.27e-21 | 4.45E-20 |
| PBANKA_0803400 | 8.4 | 4.09 | 7.97e-13 | 5.27E-12 |
| PBANKA_0412800 | 8.53 | 4.08 | 3.51e-25 | 4.42E-24 |
| PBANKA_1445600 | 8.11 | 4.08 | 6.45e-24 | 7.75E-23 |
| PBANKA_1327300 | 10.25 | 4.08 | 4.19e-12 | 2.60E-11 |
| PBANKA_1416200 | 5.47 | 4.08 | 7.90e-11 | 4.41E-10 |
| PBANKA_1124100 | 8.47 | 4.07 | 2.03e-36 | 4.18E-35 |
| PBANKA_1440100 | 9.63 | 4.06 | 5.18e-44 | 1.52E-42 |
| PBANKA_0826900 | 11.27 | 4.06 | 6.77e-25 | 8.40E-24 |
| PBANKA_1425000 | 10.59 | 4.06 | 1.55e-09 | 7.55E-09 |
| PBANKA_0511900 | 11.69 | 4.05 | 3.02e-49 | 1.13E-47 |
| PBANKA_1212900 | 8.54 | 4.05 | 1.82e-34 | 3.49E-33 |
| PBANKA_0819900 | 11.03 | 4.05 | 1.28e-25 | 1.64E-24 |
| PBANKA_0905000 | 8.96 | 4.05 | 1.88e-20 | 1.91E-19 |
| PBANKA_0617200 | 10.4 | 4.05 | 1.18e-17 | 1.03E-16 |
| PBANKA_API00011 | 7.51 | 4.05 | 2.03e-12 | 1.29E-11 |
| PBANKA_1242800 | 9.5 | 4.04 | 5.33e-78 | 6.49E-76 |
| PBANKA_0931300 | 10.83 | 4.04 | 2.14e-62 | 1.32E-60 |
| PBANKA_0811100 | 8.71 | 4.04 | 2.24e-39 | 5.33E-38 |
| PBANKA_1018700 | 7.66 | 4.04 | 5.80e-22 | 6.29E-21 |
| PBANKA_0600091 | 5.16 | 4.04 | 3.65e-05 | 9.77E-05 |
| PBANKA_0414600 | 10.06 | 4.03 | 5.62e-22 | 6.10E-21 |
| PBANKA_1322700 | 9.06 | 4.02 | 5.31e-36 | 1.08E-34 |
| PBANKA_1424600 | 9.59 | 4.01 | 1.13e-55 | 5.48E-54 |
| PBANKA_0711900 | 15.37 | 4.01 | 1.48e-34 | 2.85E-33 |
| PBANKA_1363000 | 7.02 | 4.01 | 8.12e-19 | 7.52E-18 |
| PBANKA_0919300 | 9 | 4 | 1.55e-47 | 5.39E-46 |
| PBANKA_1323000 | 7.49 | 4 | 5.39e-25 | 6.70E-24 |
| PBANKA_1324400 | 10.22 | 4 | 1.03e-19 | 9.98E-19 |
| PBANKA_1227100 | 10.4 | 4 | 3.76e-18 | 3.36E-17 |
| PBANKA_1326600 | 8.81 | 3.99 | 4.88e-32 | 8.41E-31 |
| PBANKA_0610400 | 11.04 | 3.99 | 5.20e-32 | 8.90E-31 |
| PBANKA_0314500 | 10.09 | 3.99 | 2.75e-21 | 2.90E-20 |
| PBANKA_1352700 | 9.48 | 3.98 | 1.16e-44 | 3.47E-43 |
| PBANKA_0810600 | 10.73 | 3.98 | 2.93e-39 | 6.88E-38 |
| PBANKA_1346700 | 10.47 | 3.98 | 6.37e-12 | 3.89E-11 |
| PBANKA_1229200 | 11.17 | 3.97 | 1.53e-38 | 3.48E-37 |
| PBANKA_0416200 | 9.55 | 3.96 | 4.95e-52 | 2.02E-50 |
| PBANKA_0714800 | 7.51 | 3.96 | 1.19e-40 | 2.96E-39 |
| PBANKA_1354300 | 10.64 | 3.96 | 6.24e-27 | 8.55E-26 |
| PBANKA_1327251 | 8.87 | 3.96 | 6.56e-12 | 4.00E-11 |
| PBANKA_1355100 | 11.73 | 3.95 | 4.63e-42 | 1.23E-40 |
| PBANKA_0826700 | 10.87 | 3.95 | 9.29e-33 | 1.64E-31 |
| PBANKA_1356700 | 10.36 | 3.95 | 2.34e-30 | 3.67E-29 |
| PBANKA_1440400 | 7.17 | 3.95 | 4.94e-24 | 5.98E-23 |
| PBANKA_1223300 | 8.74 | 3.94 | 1.74e-28 | 2.54E-27 |
| PBANKA_1339200 | 7.59 | 3.94 | 6.07e-20 | 5.97E-19 |
| PBANKA_1237000 | 8.79 | 3.94 | 3.19e-19 | 3.02E-18 |
| PBANKA_1231600 | 12.06 | 3.94 | 1.71e-10 | 9.21E-10 |
| PBANKA_1363100 | 8.71 | 3.93 | 3.19e-30 | 4.96E-29 |
| PBANKA_0216741 | 5.68 | 3.93 | 1.10e-09 | 5.45E-09 |
| PBANKA_0943500 | 8.65 | 3.92 | 2.36e-68 | 1.88E-66 |
| PBANKA_1413900 | 9.51 | 3.92 | 1.09e-43 | 3.18E-42 |
| PBANKA_0921500 | 7.39 | 3.92 | 3.75e-14 | 2.68E-13 |
| PBANKA_0905600 | 9.2 | 3.91 | 2.63e-23 | 3.10E-22 |
| PBANKA_1309000 | 8.5 | 3.9 | 2.00e-36 | 4.14E-35 |
| PBANKA_1343750 | 8.07 | 3.9 | 2.23e-10 | 1.19E-09 |
| PBANKA_0521400 | 8.73 | 3.89 | 4.56e-62 | 2.71E-60 |
| PBANKA_1125900 | 11.37 | 3.89 | 8.83e-37 | 1.85E-35 |

| | | | | |
|---|---|---|---|---|
| PBANKA_0709800 | 8.22 | -3.66 | 4.41e-16 | 3.53E-15 |
| PBANKA_0410100 | 6.7 | -3.66 | 6.34e-06 | 1.95E-05 |
| PBANKA_1461300 | 10.84 | -3.66 | 7.73e-06 | 2.34E-05 |
| PBANKA_0201340 | 6.54 | -3.66 | 8.84e-06 | 2.66E-05 |
| PBANKA_1365200 | 9.6 | -3.63 | 2.95e-09 | 1.40E-08 |
| PBANKA_0620500 | 7.86 | -3.63 | 6.62e-06 | 2.02E-05 |
| PBANKA_1436600 | 10.79 | -3.62 | 1.22e-06 | 4.19E-06 |
| PBANKA_1347900 | 8.78 | -3.61 | 3.46e-23 | 4.03E-22 |
| PBANKA_0508500 | 6.88 | -3.61 | 1.68e-05 | 4.77E-05 |
| PBANKA_1137800 | 9.86 | -3.6 | 5.75e-21 | 5.92E-20 |
| PBANKA_1006100 | 8.77 | -3.6 | 1.01e-20 | 1.03E-19 |
| PBANKA_0518500 | 7.67 | -3.59 | 1.74e-06 | 5.79E-06 |
| PBANKA_0614100 | 6.53 | -3.59 | 2.98e-05 | 8.11E-05 |
| PBANKA_0412500 | 8.61 | -3.58 | 2.78e-13 | 1.89E-12 |
| PBANKA_0112100 | 10.73 | -3.58 | 1.20e-10 | 6.55E-10 |
| PBANKA_0935800 | 7.41 | -3.58 | 1.47e-09 | 7.17E-09 |
| PBANKA_1223900 | 6.59 | -3.57 | 7.06e-07 | 2.51E-06 |
| PBANKA_1244700 | 7.58 | -3.56 | 3.54e-20 | 3.51E-19 |
| PBANKA_0522000 | 8.14 | -3.56 | 4.54e-14 | 3.23E-13 |
| PBANKA_1348700 | 7.51 | -3.56 | 2.16e-13 | 1.48E-12 |
| PBANKA_1116200 | 7.31 | -3.56 | 3.78e-07 | 1.40E-06 |
| PBANKA_1136200 | 7.8 | -3.55 | 1.43e-13 | 1.00E-12 |
| PBANKA_0103500 | 7.07 | -3.53 | 6.72e-08 | 2.73E-07 |
| PBANKA_0514500 | 7.83 | -3.52 | 4.76e-15 | 3.63E-14 |
| PBANKA_0411900 | 5.97 | -3.52 | 1.71e-05 | 4.85E-05 |
| PBANKA_1214200 | 8.64 | -3.51 | 3.43e-11 | 1.96E-10 |
| PBANKA_0925100 | 7.27 | -3.51 | 1.64e-07 | 6.40E-07 |
| PBANKA_1301300 | 7.94 | -3.5 | 1.43e-10 | 7.77E-10 |
| PBANKA_0308000 | 6.39 | -3.47 | 1.82e-05 | 5.13E-05 |
| PBANKA_1352400 | 8.06 | -3.46 | 5.65e-24 | 6.81E-23 |
| PBANKA_1231300 | 9.84 | -3.46 | 2.82e-07 | 1.06E-06 |
| PBANKA_1244500 | 6.12 | -3.46 | 1.60e-05 | 4.55E-05 |
| PBANKA_1320700 | 9.58 | -3.45 | 2.04e-11 | 1.19E-10 |
| PBANKA_0416000 | 11.23 | -3.44 | 5.98e-17 | 5.03E-16 |
| PBANKA_1425600 | 6.97 | -3.43 | 9.93e-09 | 4.48E-08 |
| PBANKA_1108700 | 10.38 | -3.43 | 6.09e-07 | 2.18E-06 |
| PBANKA_0617300 | 6.95 | -3.42 | 1.81e-10 | 9.72E-10 |
| PBANKA_0835600 | 7.46 | -3.42 | 1.03e-08 | 4.63E-08 |
| PBANKA_1029400 | 9.37 | -3.41 | 7.69e-13 | 5.10E-12 |
| PBANKA_0902500 | 9.06 | -3.41 | 9.10e-12 | 5.48E-11 |
| PBANKA_0601000 | 7.09 | -3.41 | 5.48e-06 | 1.70E-05 |
| PBANKA_0930900 | 7.76 | -3.4 | 4.63e-11 | 2.63E-10 |
| PBANKA_0907100 | 9.82 | -3.4 | 5.57e-06 | 1.73E-05 |
| PBANKA_0600600 | 10.54 | -3.4 | 2.54e-05 | 6.98E-05 |
| PBANKA_1457200 | 9.32 | -3.39 | 3.67e-13 | 2.48E-12 |
| PBANKA_1143800 | 9.47 | -3.39 | 1.90e-07 | 7.30E-07 |
| PBANKA_1404900 | 8.75 | -3.38 | 3.52e-12 | 2.20E-11 |
| PBANKA_0825400 | 8.49 | -3.37 | 4.10e-21 | 4.29E-20 |
| PBANKA_1313600 | 6.3 | -3.37 | 6.77e-05 | 1.72E-04 |
| PBANKA_0416700 | 7.42 | -3.36 | 5.63e-08 | 2.31E-07 |
| PBANKA_0600800 | 6.74 | -3.36 | 2.18e-06 | 7.17E-06 |
| PBANKA_0622000 | 8.28 | -3.36 | 4.51e-03 | 7.90E-03 |
| PBANKA_0927600 | 11.15 | -3.35 | 3.29e-11 | 1.89E-10 |
| PBANKA_0105900 | 7.1 | -3.35 | 5.72e-07 | 2.06E-06 |
| PBANKA_1409600 | 8.74 | -3.35 | 6.97e-07 | 2.48E-06 |
| PBANKA_0830100 | 6.88 | -3.35 | 2.84e-06 | 9.20E-06 |
| PBANKA_0801800 | 10.15 | -3.34 | 7.52e-08 | 3.05E-07 |
| PBANKA_0312700 | 10.24 | -3.34 | 8.20e-07 | 2.90E-06 |
| PBANKA_1027200 | 7.18 | -3.33 | 6.49e-07 | 2.32E-06 |
| PBANKA_0523400 | 6.75 | -3.33 | 1.05e-06 | 3.63E-06 |
| PBANKA_0210700 | 7.35 | -3.32 | 5.37e-11 | 3.03E-10 |
| PBANKA_0612300 | 7.65 | -3.32 | 4.75e-10 | 2.46E-09 |
| PBANKA_1019200 | 7.61 | -3.31 | 5.88e-11 | 3.30E-10 |
| PBANKA_1322000 | 8.73 | -3.3 | 1.37e-20 | 1.40E-19 |
| PBANKA_1028800 | 7.25 | -3.29 | 2.32e-11 | 1.35E-10 |
| PBANKA_0603700 | 8.33 | -3.29 | 2.52e-10 | 1.34E-09 |
| PBANKA_0906200 | 7.8 | -3.29 | 1.77e-08 | 7.73E-08 |
| PBANKA_0210300 | 6.45 | -3.29 | 8.69e-06 | 2.62E-05 |
| PBANKA_0934700 | 6.71 | -3.29 | 1.39e-04 | 3.32E-04 |
| PBANKA_0941600 | 8.75 | -3.28 | 4.52e-21 | 4.69E-20 |
| PBANKA_0203500 | 8.63 | -3.28 | 1.10e-09 | 5.45E-09 |
| PBANKA_0523700 | 10.87 | -3.28 | 4.03e-08 | 1.68E-07 |
| PBANKA_1113500 | 8.04 | -3.28 | 1.81e-07 | 7.02E-07 |
| PBANKA_1228200 | 8.41 | -3.27 | 1.73e-12 | 1.11E-11 |

| | | | | |
|---|---|---|---|---|
| PBANKA_1021100 | 8.3 | 3.89 | 1.36e-26 | 1.84E-25 |
| PBANKA_1034400 | 11.7 | 3.89 | 1.89e-23 | 2.23E-22 |
| PBANKA_1125100 | 9.35 | 3.88 | 5.39e-15 | 4.09E-14 |
| PBANKA_0600061 | 6.3 | 3.88 | 1.25e-13 | 8.73E-13 |
| PBANKA_0932400 | 9.92 | 3.87 | 8.53e-27 | 1.16E-25 |
| PBANKA_1102200 | 11.84 | 3.86 | 5.18e-32 | 8.89E-31 |
| PBANKA_0720400 | 11.99 | 3.86 | 3.05e-28 | 4.44E-27 |
| PBANKA_0819300 | 9.32 | 3.85 | 6.00e-38 | 1.33E-36 |
| PBANKA_1039400 | 10.57 | 3.85 | 7.58e-18 | 6.68E-17 |
| PBANKA_1026900 | 9.43 | 3.84 | 2.05e-62 | 1.28E-60 |
| PBANKA_0817700 | 10.2 | 3.84 | 6.47e-38 | 1.42E-36 |
| PBANKA_1005600 | 7.52 | 3.84 | 6.81e-05 | 1.73E-04 |
| PBANKA_1309700 | 10.18 | 3.82 | 1.22e-27 | 1.72E-26 |
| PBANKA_0213100 | 6.19 | 3.82 | 7.30e-17 | 6.12E-16 |
| PBANKA_0524821 | 5.41 | 3.82 | 2.09e-05 | 5.80E-05 |
| PBANKA_1006700 | 9.23 | 3.81 | 1.87e-47 | 6.43E-46 |
| PBANKA_0715200 | 9.26 | 3.81 | 8.64e-20 | 8.37E-19 |
| PBANKA_0600051 | 6.19 | 3.81 | 8.44e-13 | 5.55E-12 |
| PBANKA_1401100 | 8.18 | 3.8 | 1.91e-54 | 8.71E-53 |
| PBANKA_1028500 | 9.66 | 3.8 | 7.36e-53 | 3.11E-51 |
| PBANKA_1002200 | 8.9 | 3.8 | 1.08e-05 | 3.18E-05 |
| PBANKA_1449400 | 9.63 | 3.79 | 1.04e-52 | 4.36E-51 |
| PBANKA_1245900 | 8.57 | 3.79 | 4.88e-28 | 7.04E-27 |
| PBANKA_1309500 | 9.87 | 3.78 | 1.40e-54 | 6.52E-53 |
| PBANKA_1018600 | 11.3 | 3.78 | 6.12e-46 | 1.97E-44 |
| PBANKA_0308500 | 8.63 | 3.78 | 5.11e-33 | 9.19E-32 |
| PBANKA_0711600 | 9.73 | 3.78 | 9.73e-19 | 8.94E-18 |
| PBANKA_1237800 | 11.14 | 3.78 | 2.42e-14 | 1.76E-13 |
| PBANKA_0602600 | 6.36 | 3.78 | 1.80e-12 | 1.15E-11 |
| PBANKA_0712900 | 11.19 | 3.78 | 6.91e-12 | 4.21E-11 |
| PBANKA_0514100 | 10.22 | 3.77 | 2.71e-96 | 5.33E-94 |
| PBANKA_0316300 | 9.4 | 3.77 | 2.76e-29 | 4.17E-28 |
| PBANKA_1008000 | 8.38 | 3.77 | 1.46e-24 | 1.79E-23 |
| PBANKA_0820900 | 7.35 | 3.77 | 7.23e-12 | 4.40E-11 |
| PBANKA_0700761 | 15.81 | 3.77 | 2.81e-10 | 1.48E-09 |
| PBANKA_0705100 | 8.93 | 3.76 | 8.78e-46 | 2.77E-44 |
| PBANKA_0925300 | 10.36 | 3.76 | 1.62e-16 | 1.32E-15 |
| PBANKA_1405600 | 8.37 | 3.75 | 1.13e-46 | 3.74E-45 |
| PBANKA_0823900 | 7.56 | 3.75 | 5.34e-28 | 7.68E-27 |
| PBANKA_0307000 | 10.32 | 3.74 | 3.33e-19 | 3.15E-18 |
| PBANKA_0412400 | 9.33 | 3.73 | 6.63e-15 | 4.99E-14 |
| PBANKA_0519900 | 11.4 | 3.71 | 5.14e-26 | 6.71E-25 |
| PBANKA_1037400 | 10.14 | 3.71 | 1.97e-15 | 1.53E-14 |
| PBANKA_1246100 | 5.99 | 3.71 | 3.37e-05 | 9.09E-05 |
| PBANKA_0614500 | 8.93 | 3.7 | 1.26e-23 | 1.50E-22 |
| PBANKA_0905700 | 7.21 | 3.7 | 9.43e-23 | 1.07E-21 |
| PBANKA_0919200 | 8.7 | 3.7 | 6.03e-22 | 6.52E-21 |
| PBANKA_1423500 | 11.17 | 3.69 | 8.17e-28 | 1.16E-26 |
| PBANKA_1439800 | 6.93 | 3.69 | 7.65e-18 | 6.73E-17 |
| PBANKA_0604800 | 10.52 | 3.68 | 1.38e-23 | 1.65E-22 |
| PBANKA_0605700 | 10.38 | 3.67 | 3.61e-20 | 3.58E-19 |
| PBANKA_1028400 | 11.14 | 3.66 | 3.27e-17 | 2.79E-16 |
| PBANKA_1204400 | 9.43 | 3.66 | 1.61e-11 | 9.55E-11 |
| PBANKA_1134000 | 9.15 | 3.65 | 1.46e-23 | 1.74E-22 |
| PBANKA_1463800 | 8.39 | 3.64 | 7.71e-47 | 2.58E-45 |
| PBANKA_1445400 | 10.63 | 3.64 | 2.47e-28 | 3.62E-27 |
| PBANKA_0317141 | 6.53 | 3.64 | 1.11e-14 | 8.27E-14 |
| PBANKA_1436400 | 6.62 | 3.64 | 1.84e-11 | 1.08E-10 |
| PBANKA_0818700 | 5.61 | 3.64 | 2.76e-10 | 1.46E-09 |
| PBANKA_1456100 | 11.23 | 3.63 | 2.34e-26 | 3.13E-25 |
| PBANKA_1012400 | 8.69 | 3.63 | 3.70e-23 | 4.29E-22 |
| PBANKA_1327600 | 7.81 | 3.63 | 4.19e-21 | 4.37E-20 |
| PBANKA_0800500 | 15.24 | 3.63 | 1.65e-12 | 1.06E-11 |
| PBANKA_0804000 | 9.68 | 3.62 | 1.47e-25 | 1.87E-24 |
| PBANKA_1410100 | 10.77 | 3.61 | 2.88e-32 | 4.99E-31 |
| PBANKA_1030700 | 7.94 | 3.61 | 2.50e-30 | 3.92E-29 |
| PBANKA_0807600 | 10.33 | 3.61 | 2.57e-20 | 2.58E-19 |
| PBANKA_0511200 | 10.04 | 3.61 | 1.90e-16 | 1.55E-15 |
| PBANKA_1435900 | 8.94 | 3.61 | 1.64e-13 | 1.14E-12 |
| PBANKA_1037800 | 12.55 | 3.61 | 1.32e-08 | 5.85E-08 |
| PBANKA_1006900 | 7.94 | 3.6 | 1.59e-33 | 2.95E-32 |
| PBANKA_1234200 | 11.02 | 3.6 | 4.25e-19 | 3.99E-18 |
| PBANKA_1211800 | 8.45 | 3.59 | 5.53e-26 | 7.18E-25 |
| PBANKA_1433100 | 9.02 | 3.58 | 4.01e-31 | 6.53E-30 |

| | | | | |
|---|---|---|---|---|
| PBANKA_0936500 | 7.25 | -3.27 | 1.24e-06 | 4.24E-06 |
| PBANKA_1223400 | 7.21 | -3.27 | 1.50e-06 | 5.04E-06 |
| PBANKA_1139200 | 6.26 | -3.27 | 1.51e-06 | 5.08E-06 |
| PBANKA_0803700 | 8.29 | -3.26 | 6.82e-20 | 6.67E-19 |
| PBANKA_0715600 | 9.17 | -3.26 | 2.78e-12 | 1.75E-11 |
| PBANKA_0911200 | 6.55 | -3.26 | 4.18e-04 | 9.11E-04 |
| PBANKA_0614000 | 10.51 | -3.25 | 3.80e-20 | 3.76E-19 |
| PBANKA_0408900 | 8.84 | -3.25 | 2.39e-17 | 2.06E-16 |
| PBANKA_0924500 | 9.47 | -3.25 | 5.69e-12 | 3.50E-11 |
| PBANKA_1416900 | 8.86 | -3.25 | 3.59e-08 | 1.51E-07 |
| PBANKA_0514800 | 6.86 | -3.25 | 1.61e-07 | 6.27E-07 |
| PBANKA_1023700 | 7.58 | -3.25 | 1.29e-06 | 4.41E-06 |
| PBANKA_0613300 | 7.91 | -3.24 | 2.25e-13 | 1.55E-12 |
| PBANKA_0203000 | 10.05 | -3.24 | 3.21e-12 | 2.01E-11 |
| PBANKA_0907300 | 7.24 | -3.24 | 3.45e-10 | 1.80E-09 |
| PBANKA_1239000 | 9.97 | -3.24 | 1.36e-05 | 3.92E-05 |
| PBANKA_1419000 | 8.37 | -3.23 | 6.45e-24 | 7.75E-23 |
| PBANKA_0919600 | 8.54 | -3.23 | 1.26e-10 | 6.88E-10 |
| PBANKA_0937900 | 6.91 | -3.23 | 2.14e-10 | 1.14E-09 |
| PBANKA_1226800 | 9.81 | -3.23 | 5.97e-10 | 3.06E-09 |
| PBANKA_0613000 | 7.49 | -3.23 | 3.61e-07 | 1.34E-06 |
| PBANKA_0814500 | 7.35 | -3.23 | 6.10e-06 | 1.88E-05 |
| PBANKA_0408500 | 7.52 | -3.22 | 1.82e-15 | 1.42E-14 |
| PBANKA_0615200 | 8.93 | -3.21 | 1.28e-12 | 8.28E-12 |
| PBANKA_1356500 | 8.25 | -3.21 | 3.08e-11 | 1.78E-10 |
| PBANKA_1033300 | 7.46 | -3.21 | 5.87e-09 | 2.70E-08 |
| PBANKA_1134900 | 11.83 | -3.21 | 8.48e-09 | 3.84E-08 |
| PBANKA_1350200 | 6.47 | -3.21 | 1.25e-04 | 3.00E-04 |
| PBANKA_1330700 | 8.93 | -3.2 | 3.30e-09 | 1.56E-08 |
| PBANKA_0917000 | 7.82 | -3.2 | 4.59e-07 | 1.67E-06 |
| PBANKA_1121900 | 8.01 | -3.2 | 4.33e-06 | 1.36E-05 |
| PBANKA_1358400 | 6.77 | -3.2 | 7.27e-06 | 2.21E-05 |
| PBANKA_0600900 | 6.5 | -3.2 | 5.79e-05 | 1.49E-04 |
| PBANKA_1419800 | 8.26 | -3.19 | 5.76e-16 | 4.56E-15 |
| PBANKA_1300700 | 10.25 | -3.19 | 2.06e-09 | 9.91E-09 |
| PBANKA_1036800 | 6.75 | -3.19 | 7.83e-05 | 1.96E-04 |
| PBANKA_0802200 | 8.52 | -3.18 | 2.70e-25 | 3.42E-24 |
| PBANKA_0907700 | 8.27 | -3.17 | 2.66e-17 | 2.29E-16 |
| PBANKA_1033400 | 8.64 | -3.17 | 1.78e-11 | 1.05E-10 |
| PBANKA_0913900 | 8.7 | -3.16 | 2.27e-19 | 2.17E-18 |
| PBANKA_1362300 | 10.2 | -3.16 | 5.50e-12 | 3.39E-11 |
| PBANKA_0802000 | 10.69 | -3.16 | 3.23e-07 | 1.21E-06 |
| PBANKA_1129400 | 8.61 | -3.15 | 3.66e-18 | 3.28E-17 |
| PBANKA_1015600 | 7.41 | -3.15 | 7.58e-10 | 3.83E-09 |
| PBANKA_0303300 | 7.34 | -3.15 | 2.32e-06 | 7.62E-06 |
| PBANKA_1205800 | 6.71 | -3.15 | 1.30e-05 | 3.78E-05 |
| PBANKA_0402100 | 8.94 | -3.14 | 3.61e-13 | 2.45E-12 |
| PBANKA_0608200 | 7.84 | -3.14 | 5.02e-08 | 2.07E-07 |
| PBANKA_0107000 | 7.21 | -3.14 | 4.39e-05 | 1.15E-04 |
| PBANKA_1332900 | 5.78 | -3.14 | 6.95e-05 | 1.76E-04 |
| PBANKA_1337600 | 8.95 | -3.13 | 2.32e-14 | 1.69E-13 |
| PBANKA_0915100 | 5.97 | -3.13 | 5.74e-08 | 2.35E-07 |
| PBANKA_0410200 | 6.61 | -3.13 | 3.39e-06 | 1.09E-05 |
| PBANKA_1441800 | 5.82 | -3.13 | 2.85e-05 | 7.77E-05 |
| PBANKA_1101600 | 8.86 | -3.12 | 9.17e-17 | 7.61E-16 |
| PBANKA_1417900 | 9.94 | -3.12 | 1.06e-16 | 8.82E-16 |
| PBANKA_0803000 | 7.93 | -3.12 | 2.78e-14 | 2.00E-13 |
| PBANKA_0506600 | 9.89 | -3.12 | 1.76e-13 | 1.22E-12 |
| PBANKA_0619200 | 13.06 | -3.12 | 8.99e-07 | 3.15E-06 |
| PBANKA_1112100 | 6.42 | -3.12 | 2.72e-06 | 8.84E-06 |
| PBANKA_1340400 | 6.06 | -3.12 | 1.33e-04 | 3.18E-04 |
| PBANKA_1232700 | 7.13 | -3.11 | 5.88e-12 | 3.60E-11 |
| PBANKA_1430100 | 7.57 | -3.11 | 6.20e-09 | 2.84E-08 |
| PBANKA_0813300 | 8.77 | -3.11 | 2.02e-08 | 8.76E-08 |
| PBANKA_1446700 | 7.92 | -3.11 | 1.29e-07 | 5.07E-07 |
| PBANKA_1017700 | 7.32 | -3.11 | 1.45e-06 | 4.91E-06 |
| PBANKA_1458800 | 8.13 | -3.11 | 1.35e-05 | 3.90E-05 |
| PBANKA_1422200 | 7.1 | -3.11 | 1.49e-04 | 3.53E-04 |
| PBANKA_0902300 | 7.57 | -3.11 | 5.16e-03 | 8.96E-03 |
| PBANKA_1016000 | 7.68 | -3.1 | 2.82e-10 | 1.48E-09 |
| PBANKA_0508600 | 7.63 | -3.1 | 4.02e-09 | 1.88E-08 |
| PBANKA_0405800 | 7.26 | -3.1 | 5.07e-07 | 1.84E-06 |
| PBANKA_0208000 | 7.2 | -3.1 | 6.31e-07 | 2.26E-06 |
| PBANKA_0939900 | 6.16 | -3.1 | 1.69e-04 | 3.98E-04 |

| | | | | |
|---|---|---|---|---|
| PBANKA_0513200 | 7.98 | 3.58 | 6.22e-30 | 9.64E-29 |
| PBANKA_0204100 | 7.89 | 3.58 | 1.22e-28 | 1.81E-27 |
| PBANKA_1216100 | 6.94 | 3.58 | 2.11e-21 | 2.23E-20 |
| PBANKA_0403000 | 9.66 | 3.58 | 5.96e-18 | 5.27E-17 |
| PBANKA_1128400 | 5.67 | 3.58 | 8.09e-11 | 4.50E-10 |
| PBANKA_0410600 | 10.07 | 3.56 | 1.15e-32 | 1.99E-31 |
| PBANKA_0717800 | 11 | 3.56 | 5.23e-22 | 5.70E-21 |
| PBANKA_0522800 | 10.85 | 3.56 | 2.20e-17 | 1.90E-16 |
| PBANKA_1342400 | 7.46 | 3.55 | 1.40e-28 | 2.07E-27 |
| PBANKA_0201300 | 8.46 | 3.55 | 8.23e-18 | 7.20E-17 |
| PBANKA_0610900 | 10.69 | 3.54 | 2.96e-53 | 1.28E-51 |
| PBANKA_0315100 | 8.62 | 3.54 | 4.36e-33 | 7.92E-32 |
| PBANKA_0941500 | 10.97 | 3.53 | 1.95e-20 | 1.97E-19 |
| PBANKA_0811400 | 11.74 | 3.53 | 1.15e-08 | 5.13E-08 |
| PBANKA_1146241 | 4.23 | 3.53 | 1.96e-03 | 3.72E-03 |
| PBANKA_0311300 | 6.31 | 3.52 | 7.90e-12 | 4.79E-11 |
| PBANKA_0407600 | 9.53 | 3.51 | 2.25e-35 | 4.48E-34 |
| PBANKA_0703700 | 10.14 | 3.51 | 3.42e-11 | 1.96E-10 |
| PBANKA_1327000 | 8.42 | 3.5 | 2.00e-43 | 5.72E-42 |
| PBANKA_0104800 | 9.48 | 3.5 | 3.79e-38 | 8.46E-37 |
| PBANKA_0304100 | 9.2 | 3.5 | 1.39e-17 | 1.21E-16 |
| PBANKA_1243300 | 6.48 | 3.5 | 3.85e-17 | 3.27E-16 |
| PBANKA_1407600 | 11 | 3.5 | 1.08e-16 | 8.98E-16 |
| PBANKA_0602800 | 9.55 | 3.5 | 1.29e-11 | 7.71E-11 |
| PBANKA_0714900 | 6.68 | 3.5 | 2.08e-08 | 9.02E-08 |
| PBANKA_0943600 | 10.41 | 3.48 | 9.73e-19 | 8.94E-18 |
| PBANKA_0709100 | 9.15 | 3.48 | 2.93e-17 | 2.52E-16 |
| PBANKA_1313500 | 7.15 | 3.48 | 2.63e-11 | 1.52E-10 |
| PBANKA_0524200 | 10.79 | 3.47 | 1.50e-10 | 8.13E-10 |
| PBANKA_1433300 | 8.8 | 3.46 | 3.21e-54 | 1.45E-52 |
| PBANKA_1213700 | 8.89 | 3.46 | 3.52e-49 | 1.30E-47 |
| PBANKA_0202900 | 8.97 | 3.46 | 1.02e-32 | 1.79E-31 |
| PBANKA_1443800 | 10.74 | 3.46 | 1.85e-22 | 2.06E-21 |
| PBANKA_0919100 | 11.83 | 3.45 | 4.97e-37 | 1.05E-35 |
| PBANKA_1003300 | 8.03 | 3.45 | 1.54e-28 | 2.27E-27 |
| PBANKA_0818600 | 8.44 | 3.45 | 1.64e-20 | 1.68E-19 |
| PBANKA_1302400 | 9.31 | 3.44 | 2.03e-24 | 2.47E-23 |
| PBANKA_0200600 | 7.51 | 3.44 | 3.47e-13 | 2.36E-12 |
| PBANKA_0618800 | 8.87 | 3.43 | 1.91e-30 | 3.02E-29 |
| PBANKA_1208800 | 7.87 | 3.43 | 1.26e-15 | 9.94E-15 |
| PBANKA_1243200 | 7.58 | 3.42 | 1.20e-19 | 1.16E-18 |
| PBANKA_1242200 | 7.21 | 3.42 | 6.62e-15 | 4.98E-14 |
| PBANKA_0610500 | 8.75 | 3.42 | 1.99e-12 | 1.27E-11 |
| PBANKA_1451200 | 8.47 | 3.42 | 8.52e-07 | 3.00E-06 |
| PBANKA_0622961 | 15.17 | 3.42 | 6.17e-05 | 1.58E-04 |
| PBANKA_0621400 | 10.15 | 3.41 | 1.16e-43 | 3.36E-42 |
| PBANKA_0817400 | 10.2 | 3.41 | 9.25e-38 | 2.01E-36 |
| PBANKA_1302500 | 7.38 | 3.41 | 2.05e-31 | 3.38E-30 |
| PBANKA_0404000 | 10.69 | 3.41 | 1.04e-18 | 9.55E-18 |
| PBANKA_0931200 | 11.38 | 3.41 | 1.62e-17 | 1.40E-16 |
| PBANKA_1364100 | 7.7 | 3.41 | 8.00e-13 | 5.29E-12 |
| PBANKA_1315800 | 5.46 | 3.41 | 6.20e-08 | 2.53E-07 |
| PBANKA_0407700 | 10.77 | 3.4 | 1.07e-25 | 1.37E-24 |
| PBANKA_1033900 | 11.65 | 3.4 | 7.88e-18 | 6.92E-17 |
| PBANKA_0928200 | 10.22 | 3.4 | 1.78e-15 | 1.39E-14 |
| PBANKA_0214100 | 5.48 | 3.4 | 5.07e-04 | 1.08E-03 |
| PBANKA_1405500 | 7.29 | 3.39 | 3.87e-23 | 4.47E-22 |
| PBANKA_API00280 | 3.94 | 3.38 | 5.14e-03 | 8.94E-03 |
| PBANKA_0405500 | 11.2 | 3.37 | 7.86e-32 | 1.33E-30 |
| PBANKA_1403400 | 5.73 | 3.37 | 1.78e-07 | 6.90E-07 |
| PBANKA_0805400 | 6.81 | 3.36 | 2.66e-13 | 1.82E-12 |
| PBANKA_0620000 | 7.82 | 3.34 | 8.82e-41 | 2.23E-39 |
| PBANKA_0409500 | 6.33 | 3.34 | 6.36e-07 | 2.27E-06 |
| PBANKA_1407400 | 10.74 | 3.33 | 4.21e-13 | 2.84E-12 |
| PBANKA_0304200 | 5.18 | 3.33 | 7.25e-09 | 3.31E-08 |
| PBANKA_0203300 | 7.6 | 3.32 | 4.43e-26 | 5.80E-25 |
| PBANKA_0315400 | 9.01 | 3.32 | 8.67e-25 | 1.07E-23 |
| PBANKA_0937200 | 11.51 | 3.31 | 5.52e-07 | 1.99E-06 |
| PBANKA_0604200 | 10.7 | 3.3 | 7.23e-45 | 2.18E-43 |
| PBANKA_1106400 | 8.73 | 3.3 | 1.42e-41 | 3.71E-40 |
| PBANKA_1117500 | 10.49 | 3.29 | 2.27e-16 | 1.84E-15 |
| PBANKA_1333400 | 7.06 | 3.28 | 1.89e-18 | 1.71E-17 |
| PBANKA_1019400 | 11.47 | 3.27 | 2.94e-36 | 6.01E-35 |
| PBANKA_1432500 | 10.5 | 3.27 | 3.03e-20 | 3.02E-19 |

| | | | | |
|---|---|---|---|---|
| PBANKA_1454200 | 11.22 | -3.1 | 2.35e-04 | 5.37E-04 |
| PBANKA_1307000 | 9.55 | -3.09 | 7.12e-17 | 5.99E-16 |
| PBANKA_1354100 | 8.04 | -3.09 | 3.38e-13 | 2.30E-12 |
| PBANKA_0921300 | 6.74 | -3.09 | 2.80e-04 | 6.33E-04 |
| PBANKA_0306700 | 8.68 | -3.08 | 1.21e-14 | 8.97E-14 |
| PBANKA_1339300 | 7.66 | -3.08 | 1.91e-08 | 8.28E-08 |
| PBANKA_0505500 | 7.11 | -3.08 | 1.98e-04 | 4.59E-04 |
| PBANKA_1310500 | 8.65 | -3.07 | 7.48e-12 | 4.54E-11 |
| PBANKA_1205600 | 6.55 | -3.07 | 4.93e-09 | 2.29E-08 |
| PBANKA_0605600 | 11.09 | -3.07 | 1.17e-05 | 3.44E-05 |
| PBANKA_0518400 | 6.17 | -3.07 | 1.28e-05 | 3.72E-05 |
| PBANKA_0102900 | 9.69 | -3.06 | 1.31e-22 | 1.48E-21 |
| PBANKA_1449000 | 7.85 | -3.06 | 1.06e-05 | 3.13E-05 |
| PBANKA_1117700 | 9.3 | -3.05 | 2.20e-09 | 1.06E-08 |
| PBANKA_0715300 | 7.75 | -3.05 | 1.80e-08 | 7.84E-08 |
| PBANKA_1338800 | 7.08 | -3.05 | 3.43e-08 | 1.45E-07 |
| PBANKA_1313100 | 9.89 | -3.04 | 8.24e-13 | 5.42E-12 |
| PBANKA_1304300 | 7.88 | -3.04 | 7.29e-10 | 3.70E-09 |
| PBANKA_1205200 | 8.1 | -3.04 | 2.81e-07 | 1.06E-06 |
| PBANKA_1403600 | 6.43 | -3.04 | 3.56e-06 | 1.14E-05 |
| PBANKA_0808000 | 10 | -3.03 | 3.96e-25 | 4.95E-24 |
| PBANKA_0715400 | 7.1 | -3.03 | 1.88e-11 | 1.10E-10 |
| PBANKA_1109700 | 9.61 | -3.02 | 2.08e-16 | 1.69E-15 |
| PBANKA_1017200 | 8.1 | -3.02 | 1.32e-12 | 8.51E-12 |
| PBANKA_1103100 | 8.79 | -3.02 | 2.24e-07 | 8.55E-07 |
| PBANKA_1308200 | 7.78 | -3.02 | 1.55e-06 | 5.20E-06 |
| PBANKA_0616300 | 6.9 | -3.02 | 4.62e-05 | 1.21E-04 |
| PBANKA_1322400 | 7.43 | -3.01 | 2.55e-08 | 1.10E-07 |
| PBANKA_1357700 | 8.17 | -3.01 | 2.79e-08 | 1.19E-07 |
| PBANKA_0816200 | 7.82 | -3.01 | 3.50e-05 | 9.39E-05 |
| PBANKA_0703300 | 8.29 | -3.01 | 3.97e-04 | 8.68E-04 |
| PBANKA_0502300 | 7.31 | -3 | 6.79e-08 | 2.75E-07 |
| PBANKA_1230100 | 11.11 | -3 | 1.56e-06 | 5.24E-06 |
| PBANKA_0819000 | 11.17 | -2.99 | 1.39e-14 | 1.03E-13 |
| PBANKA_0406300 | 6.6 | -2.99 | 1.80e-05 | 5.08E-05 |
| PBANKA_1102800 | 6.48 | -2.99 | 5.92e-05 | 1.52E-04 |
| PBANKA_1110200 | 9.07 | -2.99 | 2.83e-04 | 6.39E-04 |
| PBANKA_1338700 | 8.83 | -2.98 | 2.40e-15 | 1.86E-14 |
| PBANKA_1127800 | 8.42 | -2.98 | 1.93e-11 | 1.13E-10 |
| PBANKA_0914600 | 8.85 | -2.98 | 3.78e-11 | 2.16E-10 |
| PBANKA_0410300 | 8.14 | -2.98 | 2.15e-10 | 1.14E-09 |
| PBANKA_0411950 | 7.5 | -2.98 | 9.55e-10 | 4.77E-09 |
| PBANKA_1408500 | 6.55 | -2.98 | 2.73e-06 | 8.87E-06 |
| PBANKA_0716500 | 7.23 | -2.98 | 8.16e-05 | 2.04E-04 |
| PBANKA_1003600 | 8.5 | -2.97 | 2.11e-27 | 2.95E-26 |
| PBANKA_0401500 | 9.53 | -2.97 | 5.58e-15 | 4.23E-14 |
| PBANKA_0701800 | 7.58 | -2.97 | 1.93e-10 | 1.03E-09 |
| PBANKA_1025900 | 6.56 | -2.97 | 1.49e-08 | 6.59E-08 |
| PBANKA_0202700 | 7.28 | -2.97 | 2.74e-08 | 1.17E-07 |
| PBANKA_1213400 | 5.38 | -2.97 | 1.91e-04 | 4.46E-04 |
| PBANKA_0307700 | 5.88 | -2.97 | 4.40e-04 | 9.54E-04 |
| PBANKA_1120100 | 9.11 | -2.96 | 9.12e-18 | 7.96E-17 |
| PBANKA_0520300 | 7.2 | -2.96 | 4.81e-08 | 2.00E-07 |
| PBANKA_0903900 | 7.9 | -2.95 | 2.83e-19 | 2.69E-18 |
| PBANKA_1441400 | 7.92 | -2.95 | 4.08e-09 | 1.90E-08 |
| PBANKA_1015000 | 7.48 | -2.95 | 3.01e-06 | 9.73E-06 |
| PBANKA_1016100 | 6.05 | -2.95 | 2.66e-05 | 7.28E-05 |
| PBANKA_0611600 | 11.88 | -2.94 | 6.89e-20 | 6.71E-19 |
| PBANKA_0314200 | 9.17 | -2.94 | 3.10e-17 | 2.65E-16 |
| PBANKA_0506000 | 10.68 | -2.94 | 1.03e-08 | 4.62E-08 |
| PBANKA_1417500 | 7.29 | -2.94 | 4.54e-07 | 1.65E-06 |
| PBANKA_0402000 | 6.67 | -2.94 | 3.72e-06 | 1.18E-05 |
| PBANKA_1020000 | 9.29 | -2.93 | 5.55e-07 | 2.00E-06 |
| PBANKA_0926200 | 6.38 | -2.93 | 7.59e-05 | 1.91E-04 |
| PBANKA_1404800 | 6.69 | -2.93 | 3.90e-04 | 8.55E-04 |
| PBANKA_1008700 | 8.6 | -2.92 | 1.07e-14 | 7.94E-14 |
| PBANKA_1034100 | 8.35 | -2.92 | 9.93e-13 | 6.50E-12 |
| PBANKA_0106300 | 11.2 | -2.92 | 1.10e-09 | 5.44E-09 |
| PBANKA_1035200 | 8.94 | -2.92 | 1.13e-05 | 3.33E-05 |
| PBANKA_1018800 | 8.53 | -2.91 | 6.20e-09 | 2.84E-08 |
| PBANKA_0818000 | 8.69 | -2.91 | 1.74e-08 | 7.60E-08 |
| PBANKA_0922500 | 10.17 | -2.9 | 1.43e-11 | 8.53E-11 |
| PBANKA_1364000 | 9.31 | -2.9 | 1.71e-10 | 9.22E-10 |
| PBANKA_1129200 | 6.45 | -2.9 | 9.98e-07 | 3.48E-06 |

| | | | | |
|---|---|---|---|---|
| PBANKA_0942900 | 8.34 | 3.26 | 7.45e-31 | 1.19E-29 |
| PBANKA_0937300 | 7.85 | 3.26 | 1.41e-22 | 1.59E-21 |
| PBANKA_0939600 | 12.69 | 3.26 | 5.91e-08 | 2.42E-07 |
| PBANKA_0306000 | 9 | 3.26 | 1.16e-04 | 2.81E-04 |
| PBANKA_1131200 | 7.83 | 3.25 | 8.15e-17 | 6.79E-16 |
| PBANKA_1238200 | 10.2 | 3.24 | 2.21e-22 | 2.45E-21 |
| PBANKA_0503000 | 7.04 | 3.24 | 1.99e-19 | 1.90E-18 |
| PBANKA_0501051 | 6.25 | 3.24 | 2.87e-06 | 9.30E-06 |
| PBANKA_1460800 | 7.01 | 3.23 | 5.19e-34 | 9.80E-33 |
| PBANKA_1202600 | 11.64 | 3.23 | 5.24e-21 | 5.41E-20 |
| PBANKA_1407800 | 8.98 | 3.22 | 8.90e-36 | 1.80E-34 |
| PBANKA_1405400 | 8.59 | 3.22 | 6.73e-31 | 1.09E-29 |
| PBANKA_1324000 | 7.83 | 3.22 | 1.47e-06 | 4.97E-06 |
| PBANKA_0504200 | 8.16 | 3.21 | 1.07e-24 | 1.32E-23 |
| PBANKA_0110000 | 8.08 | 3.18 | 1.61e-08 | 7.08E-08 |
| PBANKA_1107900 | 9.19 | 3.17 | 6.06e-23 | 6.94E-22 |
| PBANKA_1437100 | 9.96 | 3.17 | 2.12e-16 | 1.72E-15 |
| PBANKA_1146261 | 5.18 | 3.17 | 5.28e-06 | 1.64E-05 |
| PBANKA_0927200 | 10.07 | 3.16 | 7.35e-13 | 4.89E-12 |
| PBANKA_1358000 | 8.2 | 3.15 | 2.76e-15 | 2.13E-14 |
| PBANKA_1028900 | 7.74 | 3.15 | 4.86e-15 | 3.70E-14 |
| PBANKA_0524400 | 7.5 | 3.13 | 3.59e-14 | 2.57E-13 |
| PBANKA_0508100 | 12.98 | 3.13 | 2.62e-09 | 1.25E-08 |
| PBANKA_0921400 | 8.87 | 3.12 | 8.66e-10 | 4.35E-09 |
| PBANKA_1037300 | 10.02 | 3.11 | 6.29e-27 | 8.61E-26 |
| PBANKA_1245861 | 13.33 | 3.11 | 5.05e-06 | 1.58E-05 |
| PBANKA_0801600 | 9.98 | 3.1 | 4.60e-09 | 2.14E-08 |
| PBANKA_1426900 | 10.34 | 3.09 | 4.22e-23 | 4.85E-22 |
| PBANKA_1207700 | 8.36 | 3.09 | 3.83e-19 | 3.60E-18 |
| PBANKA_1007400 | 9.73 | 3.08 | 4.95e-33 | 8.95E-32 |
| PBANKA_1359400 | 8.78 | 3.07 | 1.16e-28 | 1.72E-27 |
| PBANKA_0312400 | 10.27 | 3.07 | 3.08e-19 | 2.92E-18 |
| PBANKA_0813800 | 7.39 | 3.07 | 7.39e-17 | 6.18E-16 |
| PBANKA_1318100 | 11.08 | 3.06 | 2.87e-39 | 6.76E-38 |
| PBANKA_0213000 | 8.88 | 3.06 | 4.36e-31 | 7.08E-30 |
| PBANKA_1464000 | 7.33 | 3.06 | 2.15e-09 | 1.03E-08 |
| PBANKA_0509000 | 10.41 | 3.05 | 7.97e-28 | 1.13E-26 |
| PBANKA_1464800 | 8.46 | 3.05 | 2.39e-23 | 2.82E-22 |
| PBANKA_1401300 | 10.71 | 3.05 | 2.31e-20 | 2.33E-19 |
| PBANKA_1322500 | 7.61 | 3.04 | 1.09e-30 | 1.75E-29 |
| PBANKA_1357800 | 9.46 | 3.04 | 2.89e-21 | 3.04E-20 |
| PBANKA_1231800 | 10.49 | 3.04 | 8.52e-11 | 4.73E-10 |
| PBANKA_0822800 | 8.67 | 3.03 | 2.54e-33 | 4.68E-32 |
| PBANKA_0936700 | 9.22 | 3.03 | 9.74e-32 | 1.64E-30 |
| PBANKA_0401900 | 8.33 | 3.02 | 1.93e-33 | 3.57E-32 |
| PBANKA_1218800 | 9.07 | 3.02 | 1.37e-19 | 1.32E-18 |
| PBANKA_1462800 | 7.93 | 3.01 | 6.47e-33 | 1.15E-31 |
| PBANKA_1136300 | 8.33 | 3.01 | 7.23e-17 | 6.07E-16 |
| PBANKA_1141700 | 10.81 | 3.01 | 6.92e-11 | 3.88E-10 |
| PBANKA_1339000 | 8.86 | 3 | 7.03e-37 | 1.49E-35 |
| PBANKA_0415600 | 9.04 | 3 | 1.91e-22 | 2.12E-21 |
| PBANKA_1206100 | 8.97 | 2.99 | 4.71e-32 | 8.14E-31 |
| PBANKA_1347800 | 8.4 | 2.99 | 4.45e-14 | 3.18E-13 |
| PBANKA_0210200 | 9.47 | 2.98 | 2.50e-23 | 2.95E-22 |
| PBANKA_1423300 | 12.5 | 2.97 | 3.11e-12 | 1.95E-11 |
| PBANKA_0700721 | 14.15 | 2.97 | 1.76e-03 | 3.38E-03 |
| PBANKA_0825900 | 11.04 | 2.96 | 2.87e-16 | 2.31E-15 |
| PBANKA_0211200 | 7.76 | 2.96 | 1.68e-15 | 1.32E-14 |
| PBANKA_0925200 | 10.52 | 2.96 | 1.83e-07 | 7.05E-07 |
| PBANKA_1004900 | 8.4 | 2.95 | 5.79e-33 | 1.04E-31 |
| PBANKA_1215600 | 10.71 | 2.95 | 1.65e-21 | 1.76E-20 |
| PBANKA_1127900 | 7.76 | 2.95 | 2.10e-20 | 2.12E-19 |
| PBANKA_1238800 | 11.57 | 2.94 | 2.58e-22 | 2.84E-21 |
| PBANKA_1232500 | 10.51 | 2.94 | 3.29e-22 | 3.60E-21 |
| PBANKA_0100200 | 8.3 | 2.94 | 3.68e-19 | 3.47E-18 |
| PBANKA_1126400 | 9.62 | 2.94 | 4.22e-17 | 3.58E-16 |
| PBANKA_1140700 | 7.34 | 2.94 | 3.78e-09 | 1.77E-08 |
| PBANKA_0609900 | 9.55 | 2.93 | 2.00e-12 | 1.27E-11 |
| PBANKA_0620900 | 10.38 | 2.92 | 6.82e-20 | 6.67E-19 |
| PBANKA_1210200 | 12.48 | 2.92 | 5.45e-11 | 3.07E-10 |
| PBANKA_0832800 | 8.01 | 2.92 | 1.92e-08 | 8.35E-08 |
| PBANKA_1018000 | 8.77 | 2.91 | 2.13e-18 | 1.92E-17 |
| PBANKA_0807100 | 7.39 | 2.9 | 3.19e-26 | 4.20E-25 |
| PBANKA_0203200 | 9.16 | 2.9 | 2.15e-14 | 1.57E-13 |

| | | | | |
|---|---|---|---|---|
| PBANKA_0827700 | 6.41 | -2.9 | 9.56e-05 | 2.36E-04 |
| PBANKA_0411400 | 6.46 | -2.9 | 4.77e-04 | 1.02E-03 |
| PBANKA_1110100 | 8.71 | -2.89 | 6.07e-15 | 4.59E-14 |
| PBANKA_0601100 | 8.71 | -2.89 | 1.36e-08 | 6.02E-08 |
| PBANKA_0901500 | 6.16 | -2.89 | 1.58e-04 | 3.74E-04 |
| PBANKA_1230000 | 5.89 | -2.89 | 2.78e-04 | 6.28E-04 |
| PBANKA_0203650 | 6.93 | -2.89 | 1.49e-03 | 2.91E-03 |
| PBANKA_1438000 | 9.46 | -2.88 | 3.24e-16 | 2.59E-15 |
| PBANKA_1319200 | 8.75 | -2.88 | 6.00e-13 | 4.03E-12 |
| PBANKA_1442100 | 7.48 | -2.88 | 2.77e-12 | 1.74E-11 |
| PBANKA_0603900 | 8.73 | -2.88 | 1.31e-09 | 6.43E-09 |
| PBANKA_1028600 | 7.66 | -2.88 | 1.16e-07 | 4.62E-07 |
| PBANKA_0721800 | 6.91 | -2.88 | 1.14e-06 | 3.93E-06 |
| PBANKA_0932100 | 6.9 | -2.88 | 8.60e-06 | 2.59E-05 |
| PBANKA_0907900 | 11.09 | -2.88 | 1.25e-04 | 3.00E-04 |
| PBANKA_1115700 | 9.52 | -2.87 | 1.18e-30 | 1.88E-29 |
| PBANKA_1454600 | 9.27 | -2.87 | 2.37e-12 | 1.50E-11 |
| PBANKA_1409900 | 7.83 | -2.87 | 8.63e-10 | 4.34E-09 |
| PBANKA_1107600 | 11.1 | -2.87 | 3.75e-08 | 1.57E-07 |
| PBANKA_1213300 | 9.67 | -2.87 | 3.24e-07 | 1.21E-06 |
| PBANKA_0607700 | 11.38 | -2.86 | 7.81e-17 | 6.52E-16 |
| PBANKA_0703800 | 7.63 | -2.86 | 6.24e-10 | 3.19E-09 |
| PBANKA_1336300 | 7.41 | -2.86 | 3.81e-07 | 1.41E-06 |
| PBANKA_1360200 | 7.23 | -2.86 | 5.20e-05 | 1.35E-04 |
| PBANKA_1320400 | 8.51 | -2.85 | 1.49e-16 | 1.22E-15 |
| PBANKA_0828600 | 7.6 | -2.85 | 2.99e-08 | 1.27E-07 |
| PBANKA_0928400 | 6.75 | -2.85 | 3.35e-08 | 1.42E-07 |
| PBANKA_0109300 | 7.07 | -2.84 | 4.66e-09 | 2.17E-08 |
| PBANKA_0704300 | 7.29 | -2.84 | 3.59e-08 | 1.51E-07 |
| PBANKA_1458300 | 9.28 | -2.83 | 4.77e-18 | 4.24E-17 |
| PBANKA_0313700 | 6.15 | -2.83 | 2.10e-04 | 4.86E-04 |
| PBANKA_0301100 | 8.16 | -2.82 | 1.46e-15 | 1.15E-14 |
| PBANKA_0401200 | 7.55 | -2.82 | 1.05e-09 | 5.24E-09 |
| PBANKA_0816900 | 7.79 | -2.82 | 1.61e-09 | 7.81E-09 |
| PBANKA_0910300 | 9.42 | -2.82 | 8.31e-09 | 3.77E-08 |
| PBANKA_1322200 | 7.73 | -2.82 | 4.31e-07 | 1.58E-06 |
| PBANKA_1241200 | 6.24 | -2.82 | 5.65e-04 | 1.20E-03 |
| PBANKA_1403500 | 8.89 | -2.81 | 5.53e-14 | 3.93E-13 |
| PBANKA_1460500 | 8.32 | -2.81 | 6.34e-13 | 4.24E-12 |
| PBANKA_1024200 | 8.51 | -2.81 | 2.13e-10 | 1.14E-09 |
| PBANKA_1454800 | 8.98 | -2.81 | 1.27e-05 | 3.69E-05 |
| PBANKA_0608600 | 11.96 | -2.81 | 5.39e-05 | 1.40E-04 |
| PBANKA_1106300 | 5.66 | -2.81 | 1.20e-04 | 2.91E-04 |
| PBANKA_0721600 | 6.37 | -2.81 | 1.93e-04 | 4.50E-04 |
| PBANKA_1235700 | 5.7 | -2.81 | 5.70e-04 | 1.20E-03 |
| PBANKA_1304000 | 8.91 | -2.8 | 6.28e-21 | 6.45E-20 |
| PBANKA_1237300 | 7.45 | -2.8 | 3.48e-10 | 1.81E-09 |
| PBANKA_0925600 | 7.77 | -2.8 | 9.19e-09 | 4.16E-08 |
| PBANKA_0705600 | 10.16 | -2.8 | 6.13e-08 | 2.50E-07 |
| PBANKA_1311500 | 7.11 | -2.8 | 1.08e-07 | 4.30E-07 |
| PBANKA_0306500 | 7.21 | -2.8 | 1.06e-04 | 2.60E-04 |
| PBANKA_0927100 | 5.38 | -2.8 | 1.79e-03 | 3.42E-03 |
| PBANKA_0938300 | 8.85 | -2.79 | 2.37e-14 | 1.72E-13 |
| PBANKA_1438300 | 10.49 | -2.79 | 1.97e-12 | 1.26E-11 |
| PBANKA_1019100 | 7.94 | -2.79 | 2.97e-11 | 1.71E-10 |
| PBANKA_0916100 | 7.73 | -2.79 | 1.84e-07 | 7.08E-07 |
| PBANKA_1130400 | 8.43 | -2.79 | 3.39e-06 | 1.09E-05 |
| PBANKA_1331800 | 7.2 | -2.79 | 2.01e-05 | 5.60E-05 |
| PBANKA_1436900 | 6.52 | -2.79 | 3.03e-05 | 8.21E-05 |
| PBANKA_1220500 | 9.75 | -2.78 | 2.13e-13 | 1.47E-12 |
| PBANKA_0615300 | 10.01 | -2.78 | 8.02e-13 | 5.29E-12 |
| PBANKA_0202400 | 7.78 | -2.78 | 1.11e-11 | 6.65E-11 |
| PBANKA_1114300 | 7.59 | -2.78 | 9.84e-10 | 4.91E-09 |
| PBANKA_0307100 | 9.59 | -2.78 | 2.53e-09 | 1.21E-08 |
| PBANKA_0938400 | 10.65 | -2.78 | 5.02e-09 | 2.33E-08 |
| PBANKA_0935500 | 10.18 | -2.78 | 6.06e-09 | 2.79E-08 |
| PBANKA_0110200 | 6.59 | -2.78 | 1.41e-04 | 3.35E-04 |
| PBANKA_1016600 | 5.97 | -2.78 | 3.12e-04 | 7.00E-04 |
| PBANKA_1016200 | 8.62 | -2.77 | 2.71e-12 | 1.71E-11 |
| PBANKA_0907200 | 8.14 | -2.77 | 1.00e-08 | 4.51E-08 |
| PBANKA_1208200 | 9.98 | -2.77 | 1.38e-05 | 3.98E-05 |
| PBANKA_1424000 | 7.69 | -2.77 | 1.83e-05 | 5.16E-05 |
| PBANKA_1109300 | 8.87 | -2.77 | 8.14e-05 | 2.04E-04 |
| PBANKA_1237100 | 11.12 | -2.77 | 3.20e-04 | 7.17E-04 |

| | | | | |
|---|---|---|---|---|
| PBANKA_0514300 | 10.28 | 2.9 | 1.28e-09 | 6.32E-09 |
| PBANKA_0804800 | 7.15 | 2.89 | 2.59e-30 | 4.04E-29 |
| PBANKA_0811900 | 7.12 | 2.88 | 1.95e-15 | 1.52E-14 |
| PBANKA_1428700 | 12.06 | 2.88 | 8.43e-07 | 2.97E-06 |
| PBANKA_0906500 | 9.61 | 2.87 | 1.70e-13 | 1.18E-12 |
| PBANKA_0930300 | 10.77 | 2.87 | 1.94e-13 | 1.34E-12 |
| PBANKA_1442400 | 8.81 | 2.87 | 2.25e-11 | 1.31E-10 |
| PBANKA_1350500 | 8.5 | 2.87 | 4.80e-09 | 2.23E-08 |
| PBANKA_0620100 | 9.86 | 2.86 | 1.29e-22 | 1.46E-21 |
| PBANKA_1314100 | 8.91 | 2.86 | 1.41e-18 | 1.28E-17 |
| PBANKA_1117600 | 7.43 | 2.86 | 1.13e-06 | 3.91E-06 |
| PBANKA_0109200 | 10.07 | 2.86 | 1.68e-06 | 5.60E-06 |
| PBANKA_0600500 | 7.83 | 2.84 | 2.87e-23 | 3.35E-22 |
| PBANKA_0619900 | 8.81 | 2.84 | 5.41e-18 | 4.80E-17 |
| PBANKA_1457700 | 11.27 | 2.84 | 2.49e-05 | 6.84E-05 |
| PBANKA_0110900 | 7.83 | 2.82 | 2.59e-18 | 2.33E-17 |
| PBANKA_1036000 | 11.51 | 2.81 | 9.85e-08 | 3.94E-07 |
| PBANKA_1240300 | 7.12 | 2.8 | 2.42e-21 | 2.56E-20 |
| PBANKA_1216900 | 8.71 | 2.79 | 1.98e-27 | 2.78E-26 |
| PBANKA_0720300 | 8.34 | 2.79 | 3.77e-25 | 4.72E-24 |
| PBANKA_1025700 | 8.67 | 2.79 | 3.35e-10 | 1.75E-09 |
| PBANKA_0107800 | 7.07 | 2.79 | 3.80e-09 | 1.78E-08 |
| PBANKA_1023500 | 9.53 | 2.78 | 1.39e-37 | 3.00E-36 |
| PBANKA_1325000 | 9.54 | 2.78 | 3.21e-11 | 1.84E-10 |
| PBANKA_1006200 | 11.15 | 2.78 | 4.99e-04 | 1.07E-03 |
| PBANKA_0721900 | 7.51 | 2.77 | 5.05e-21 | 5.23E-20 |
| PBANKA_0112641 | 6.24 | 2.76 | 7.93e-13 | 5.25E-12 |
| PBANKA_1106900 | 10.09 | 2.76 | 3.70e-04 | 8.16E-04 |
| PBANKA_0815300 | 8.55 | 2.75 | 2.48e-31 | 4.07E-30 |
| PBANKA_0307800 | 9.7 | 2.75 | 5.26e-26 | 6.84E-25 |
| PBANKA_0504900 | 8.46 | 2.75 | 3.26e-20 | 3.24E-19 |
| PBANKA_0829800 | 8.47 | 2.75 | 2.58e-16 | 2.08E-15 |
| PBANKA_1351900 | 9.69 | 2.74 | 4.82e-21 | 5.00E-20 |
| PBANKA_1018500 | 9.5 | 2.73 | 3.78e-13 | 2.55E-12 |
| PBANKA_1143600 | 10.11 | 2.73 | 1.29e-11 | 7.71E-11 |
| PBANKA_1227800 | 10.22 | 2.73 | 1.01e-09 | 5.02E-09 |
| PBANKA_0605200 | 9.85 | 2.72 | 4.97e-27 | 6.84E-26 |
| PBANKA_0520200 | 9.34 | 2.72 | 1.62e-14 | 1.19E-13 |
| PBANKA_0721100 | 7.45 | 2.72 | 4.15e-11 | 2.36E-10 |
| PBANKA_1237500 | 6.21 | 2.72 | 1.30e-10 | 7.12E-10 |
| PBANKA_0804100 | 11.56 | 2.72 | 1.82e-09 | 8.77E-09 |
| PBANKA_1302000 | 8.79 | 2.71 | 2.61e-10 | 1.38E-09 |
| PBANKA_0405300 | 9.18 | 2.71 | 1.48e-05 | 4.25E-05 |
| PBANKA_1420900 | 8.84 | 2.7 | 2.62e-52 | 1.09E-50 |
| PBANKA_1007500 | 7.88 | 2.7 | 2.26e-11 | 1.32E-10 |
| PBANKA_0403200 | 13.6 | 2.7 | 8.38e-09 | 3.80E-08 |
| PBANKA_0524700 | 10.71 | 2.7 | 4.88e-08 | 2.02E-07 |
| PBANKA_0418000 | 10.46 | 2.7 | 7.65e-06 | 2.32E-05 |
| PBANKA_0618300 | 6.11 | 2.7 | 3.32e-05 | 8.97E-05 |
| PBANKA_1455300 | 9.12 | 2.69 | 4.17e-23 | 4.80E-22 |
| PBANKA_1451800 | 9.68 | 2.69 | 1.24e-14 | 9.19E-14 |
| PBANKA_1110000 | 7.82 | 2.69 | 3.03e-09 | 1.44E-08 |
| PBANKA_1445500 | 9.16 | 2.69 | 1.19e-07 | 4.71E-07 |
| PBANKA_1351500 | 8.16 | 2.68 | 1.82e-10 | 9.75E-10 |
| PBANKA_1357900 | 7.9 | 2.68 | 8.13e-06 | 2.46E-05 |
| PBANKA_0619800 | 9.25 | 2.67 | 6.50e-14 | 4.60E-13 |
| PBANKA_1334600 | 8.77 | 2.67 | 7.31e-14 | 5.16E-13 |
| PBANKA_1031100 | 8.81 | 2.67 | 1.71e-12 | 1.10E-11 |
| PBANKA_0316100 | 8.87 | 2.67 | 1.78e-10 | 9.60E-10 |
| PBANKA_1347700 | 9.08 | 2.66 | 2.25e-29 | 3.41E-28 |
| PBANKA_1349200 | 10.63 | 2.66 | 5.91e-19 | 5.51E-18 |
| PBANKA_1426200 | 11.53 | 2.66 | 1.99e-11 | 1.17E-10 |
| PBANKA_0413600 | 7.13 | 2.65 | 2.45e-16 | 1.98E-15 |
| PBANKA_1145100 | 8.45 | 2.65 | 2.75e-10 | 1.45E-09 |
| PBANKA_0912900 | 6.87 | 2.63 | 1.49e-13 | 1.04E-12 |
| PBANKA_1328800 | 8.58 | 2.63 | 2.23e-09 | 1.07E-08 |
| PBANKA_1400700 | 11.36 | 2.63 | 1.81e-08 | 7.88E-08 |
| PBANKA_1420800 | 10.03 | 2.62 | 4.05e-18 | 3.61E-17 |
| PBANKA_0923900 | 6.88 | 2.62 | 3.65e-13 | 2.47E-12 |
| PBANKA_1133300 | 12.98 | 2.62 | 8.51e-07 | 3.00E-06 |
| PBANKA_0822100 | 9.14 | 2.61 | 2.71e-20 | 2.72E-19 |
| PBANKA_1207300 | 7.02 | 2.61 | 2.00e-13 | 1.38E-12 |
| PBANKA_0521700 | 12.61 | 2.61 | 4.37e-07 | 1.60E-06 |
| PBANKA_1411000 | 9.22 | 2.61 | 3.25e-06 | 1.04E-05 |

| | | | | |
|---|---|---|---|---|
| PBANKA_1007900 | 8.85 | -2.77 | 8.74e-04 | 1.78E-03 |
| PBANKA_1227000 | 8.72 | -2.76 | 2.03e-14 | 1.48E-13 |
| PBANKA_1402500 | 8.82 | -2.76 | 4.54e-13 | 3.06E-12 |
| PBANKA_0504500 | 7.86 | -2.76 | 2.74e-07 | 1.04E-06 |
| PBANKA_0703500 | 9.73 | -2.76 | 9.34e-06 | 2.78E-05 |
| PBANKA_1333700 | 9.76 | -2.76 | 3.21e-05 | 8.67E-05 |
| PBANKA_1414700 | 6.13 | -2.76 | 6.85e-04 | 1.43E-03 |
| PBANKA_0717200 | 8.78 | -2.75 | 2.14e-11 | 1.25E-10 |
| PBANKA_1237200 | 6.24 | -2.75 | 1.51e-03 | 2.94E-03 |
| PBANKA_1129100 | 8.79 | -2.74 | 2.85e-15 | 2.19E-14 |
| PBANKA_0313800 | 9.74 | -2.74 | 1.42e-14 | 1.04E-13 |
| PBANKA_1029300 | 7.77 | -2.74 | 1.95e-08 | 8.47E-08 |
| PBANKA_0817000 | 8.39 | -2.73 | 2.29e-19 | 2.19E-18 |
| PBANKA_0932500 | 10.98 | -2.73 | 1.35e-18 | 1.23E-17 |
| PBANKA_0414300 | 9.12 | -2.73 | 1.01e-12 | 6.61E-12 |
| PBANKA_1313300 | 6.74 | -2.73 | 2.91e-10 | 1.53E-09 |
| PBANKA_1404500 | 6.64 | -2.73 | 8.99e-10 | 4.50E-09 |
| PBANKA_1118900 | 7.52 | -2.73 | 1.75e-08 | 7.67E-08 |
| PBANKA_1339700 | 6.85 | -2.73 | 3.11e-08 | 1.32E-07 |
| PBANKA_1012600 | 8.06 | -2.73 | 6.75e-06 | 2.06E-05 |
| PBANKA_0812100 | 6.52 | -2.73 | 2.67e-04 | 6.06E-04 |
| PBANKA_1451400 | 5.73 | -2.73 | 4.60e-04 | 9.94E-04 |
| PBANKA_1439100 | 10.69 | -2.72 | 6.22e-15 | 4.69E-14 |
| PBANKA_1358700 | 8.34 | -2.72 | 6.33e-10 | 3.23E-09 |
| PBANKA_1014100 | 7.17 | -2.72 | 2.19e-05 | 6.06E-05 |
| PBANKA_1402700 | 6.61 | -2.72 | 2.57e-05 | 7.04E-05 |
| PBANKA_0604900 | 11.49 | -2.72 | 3.75e-05 | 9.99E-05 |
| PBANKA_0105800 | 7.13 | -2.72 | 3.99e-05 | 1.06E-04 |
| PBANKA_0714000 | 10.04 | -2.72 | 1.17e-04 | 2.83E-04 |
| PBANKA_1214700 | 9.81 | -2.71 | 1.27e-21 | 1.36E-20 |
| PBANKA_0501900 | 7.4 | -2.71 | 2.91e-10 | 1.53E-09 |
| PBANKA_0503900 | 6.93 | -2.7 | 4.57e-12 | 2.83E-11 |
| PBANKA_1031800 | 7.31 | -2.7 | 1.22e-07 | 4.84E-07 |
| PBANKA_1229600 | 8.25 | -2.69 | 6.13e-19 | 5.71E-18 |
| PBANKA_1243900 | 10.15 | -2.69 | 8.48e-18 | 7.41E-17 |
| PBANKA_1337800 | 8.23 | -2.69 | 1.55e-13 | 1.08E-12 |
| PBANKA_1020100 | 10.52 | -2.69 | 1.98e-11 | 1.16E-10 |
| PBANKA_1358300 | 9.96 | -2.69 | 6.65e-10 | 3.40E-09 |
| PBANKA_1415300 | 6.86 | -2.69 | 1.93e-05 | 5.40E-05 |
| PBANKA_1413000 | 6.48 | -2.69 | 6.01e-04 | 1.26E-03 |
| PBANKA_1305300 | 8.02 | -2.68 | 1.50e-08 | 6.62E-08 |
| PBANKA_0719900 | 7.1 | -2.68 | 1.42e-06 | 4.81E-06 |
| PBANKA_0313500 | 7.31 | -2.68 | 1.65e-06 | 5.52E-06 |
| PBANKA_0215800 | 5.54 | -2.68 | 8.08e-04 | 1.66E-03 |
| PBANKA_1308000 | 8.92 | -2.68 | 2.51e-03 | 4.65E-03 |
| PBANKA_0937500 | 9.83 | -2.67 | 1.72e-12 | 1.10E-11 |
| PBANKA_0814200 | 9.99 | -2.67 | 6.17e-12 | 3.78E-11 |
| PBANKA_0101200 | 8.03 | -2.67 | 2.09e-09 | 1.00E-08 |
| PBANKA_0404100 | 7.71 | -2.67 | 2.83e-08 | 1.21E-07 |
| PBANKA_0207900 | 6.05 | -2.67 | 2.68e-04 | 6.08E-04 |
| PBANKA_1138800 | 7.93 | -2.66 | 2.62e-13 | 1.79E-12 |
| PBANKA_0515300 | 9.21 | -2.66 | 5.43e-12 | 3.35E-11 |
| PBANKA_0614900 | 8.14 | -2.66 | 8.99e-11 | 4.97E-10 |
| PBANKA_1332400 | 7.37 | -2.66 | 5.41e-08 | 2.23E-07 |
| PBANKA_0809200 | 6.48 | -2.66 | 1.43e-05 | 4.12E-05 |
| PBANKA_1209700 | 7.61 | -2.66 | 3.51e-05 | 9.41E-05 |
| PBANKA_1443500 | 10.32 | -2.66 | 3.67e-05 | 9.81E-05 |
| PBANKA_0807300 | 7.28 | -2.66 | 4.94e-05 | 1.29E-04 |
| PBANKA_1331200 | 9.79 | -2.65 | 2.22e-22 | 2.45E-21 |
| PBANKA_0832600 | 7.38 | -2.65 | 1.88e-05 | 5.27E-05 |
| PBANKA_1236600 | 8.04 | -2.65 | 6.85e-05 | 1.73E-04 |
| PBANKA_1322600 | 6.72 | -2.65 | 1.90e-04 | 4.44E-04 |
| PBANKA_0701200 | 8.54 | -2.65 | 1.46e-03 | 2.86E-03 |
| PBANKA_0927400 | 6.06 | -2.65 | 1.81e-03 | 3.46E-03 |
| PBANKA_1435500 | 8.48 | -2.64 | 1.40e-11 | 8.32E-11 |
| PBANKA_0312500 | 8.34 | -2.64 | 1.47e-11 | 8.75E-11 |
| PBANKA_0410400 | 8.05 | -2.64 | 3.91e-11 | 2.23E-10 |
| PBANKA_1365400 | 7.73 | -2.64 | 5.51e-11 | 3.11E-10 |
| PBANKA_0830450 | 7.21 | -2.64 | 2.26e-05 | 6.27E-05 |
| PBANKA_1318600 | 10.18 | -2.64 | 5.95e-04 | 1.25E-03 |
| PBANKA_0927500 | 5.65 | -2.64 | 1.32e-03 | 2.60E-03 |
| PBANKA_1442300 | 9.06 | -2.63 | 4.79e-22 | 5.22E-21 |
| PBANKA_1311700 | 7.15 | -2.63 | 5.85e-12 | 3.59E-11 |
| PBANKA_0832200 | 8.53 | -2.63 | 8.25e-11 | 4.59E-10 |

| | | | | |
|---|---|---|---|---|
| PBANKA_0711700 | 6.99 | 2.6 | 2.36e-11 | 1.37E-10 |
| PBANKA_1109000 | 10.69 | 2.58 | 1.03e-08 | 4.62E-08 |
| PBANKA_0201600 | 12.22 | 2.58 | 3.49e-06 | 1.12E-05 |
| PBANKA_1338600 | 10.92 | 2.57 | 3.29e-17 | 2.81E-16 |
| PBANKA_1441900 | 8.17 | 2.57 | 4.04e-12 | 2.51E-11 |
| PBANKA_0827500 | 9.51 | 2.57 | 6.07e-09 | 2.79E-08 |
| PBANKA_0501200 | 16.07 | 2.54 | 2.19e-08 | 9.45E-08 |
| PBANKA_0805300 | 6.53 | 2.53 | 9.72e-06 | 2.89E-05 |
| PBANKA_1319800 | 6.08 | 2.53 | 3.21e-05 | 8.67E-05 |
| PBANKA_0506800 | 8.61 | 2.52 | 1.67e-07 | 6.50E-07 |
| PBANKA_0524800 | 9.51 | 2.52 | 4.18e-05 | 1.10E-04 |
| PBANKA_0922900 | 8.96 | 2.51 | 1.78e-23 | 2.11E-22 |
| PBANKA_1022500 | 10.86 | 2.51 | 2.54e-17 | 2.19E-16 |
| PBANKA_0918100 | 8.53 | 2.51 | 6.12e-14 | 4.34E-13 |
| PBANKA_1362700 | 7.29 | 2.5 | 3.83e-04 | 8.42E-04 |
| PBANKA_0801400 | 6.96 | 2.49 | 1.13e-06 | 3.91E-06 |
| PBANKA_1239800 | 9.98 | 2.48 | 3.29e-15 | 2.52E-14 |
| PBANKA_0621900 | 7.12 | 2.48 | 7.10e-07 | 2.53E-06 |
| PBANKA_1114600 | 8.96 | 2.47 | 4.80e-23 | 5.51E-22 |
| PBANKA_1439300 | 8.81 | 2.47 | 4.69e-19 | 4.39E-18 |
| PBANKA_0108100 | 8.33 | 2.47 | 3.03e-15 | 2.33E-14 |
| PBANKA_0414800 | 8.67 | 2.47 | 2.54e-11 | 1.47E-10 |
| PBANKA_0604500 | 12.39 | 2.47 | 4.94e-06 | 1.55E-05 |
| PBANKA_0705700 | 8.93 | 2.46 | 3.65e-26 | 4.79E-25 |
| PBANKA_1339100 | 7.67 | 2.46 | 6.06e-22 | 6.54E-21 |
| PBANKA_1221200 | 10.51 | 2.45 | 4.44e-16 | 3.55E-15 |
| PBANKA_1309200 | 9.49 | 2.45 | 8.93e-14 | 6.30E-13 |
| PBANKA_0801200 | 10.41 | 2.45 | 1.96e-11 | 1.15E-10 |
| PBANKA_1034000 | 7.77 | 2.44 | 1.91e-18 | 1.73E-17 |
| PBANKA_0705000 | 5.33 | 2.44 | 9.26e-06 | 2.76E-05 |
| PBANKA_1010700 | 6.4 | 2.43 | 9.28e-08 | 3.73E-07 |
| PBANKA_0824100 | 6.6 | 2.43 | 1.82e-07 | 7.03E-07 |
| PBANKA_1203400 | 8.68 | 2.42 | 7.45e-15 | 5.58E-14 |
| PBANKA_1128300 | 7.34 | 2.42 | 1.27e-10 | 6.92E-10 |
| PBANKA_0912800 | 6.93 | 2.42 | 4.50e-07 | 1.64E-06 |
| PBANKA_0819100 | 6.81 | 2.42 | 1.10e-06 | 3.82E-06 |
| PBANKA_1132100 | 6.11 | 2.42 | 1.23e-05 | 3.58E-05 |
| PBANKA_0111100 | 8.41 | 2.41 | 2.72e-20 | 2.73E-19 |
| PBANKA_1354000 | 7.91 | 2.41 | 3.30e-11 | 1.89E-10 |
| PBANKA_0903200 | 7.73 | 2.4 | 3.55e-12 | 2.22E-11 |
| PBANKA_0401100 | 10.22 | 2.4 | 8.10e-08 | 3.27E-07 |
| PBANKA_0710800 | 10.34 | 2.39 | 1.83e-24 | 2.24E-23 |
| PBANKA_1347500 | 7.52 | 2.39 | 5.55e-16 | 4.41E-15 |
| PBANKA_0517100 | 5.55 | 2.39 | 1.10e-03 | 2.21E-03 |
| PBANKA_0523100 | 7.59 | 2.38 | 1.29e-16 | 1.06E-15 |
| PBANKA_0621200 | 7.09 | 2.38 | 3.51e-14 | 2.52E-13 |
| PBANKA_0108700 | 10.21 | 2.38 | 3.69e-06 | 1.18E-05 |
| PBANKA_1135200 | 7.47 | 2.37 | 1.97e-07 | 7.53E-07 |
| PBANKA_1456900 | 10.84 | 2.37 | 3.38e-06 | 1.08E-05 |
| PBANKA_1461000 | 8.59 | 2.36 | 1.18e-13 | 8.31E-13 |
| PBANKA_1433900 | 7.4 | 2.36 | 3.24e-08 | 1.37E-07 |
| PBANKA_0707800 | 7.74 | 2.35 | 3.87e-17 | 3.28E-16 |
| PBANKA_0937700 | 10.18 | 2.35 | 1.48e-12 | 9.53E-12 |
| PBANKA_1308900 | 8.37 | 2.35 | 2.91e-12 | 1.83E-11 |
| PBANKA_0707300 | 11.18 | 2.35 | 3.91e-11 | 2.23E-10 |
| PBANKA_0109700 | 8.99 | 2.34 | 8.27e-20 | 8.02E-19 |
| PBANKA_0938900 | 8.91 | 2.34 | 1.44e-10 | 7.81E-10 |
| PBANKA_0513300 | 7.47 | 2.34 | 1.55e-10 | 8.37E-10 |
| PBANKA_0939100 | 11.87 | 2.34 | 1.53e-07 | 5.98E-07 |
| PBANKA_0908200 | 8.92 | 2.33 | 6.88e-26 | 8.89E-25 |
| PBANKA_1230600 | 7.3 | 2.33 | 6.00e-07 | 2.15E-06 |
| PBANKA_0623100 | 12.88 | 2.33 | 3.40e-04 | 7.55E-04 |
| PBANKA_0819800 | 9.33 | 2.32 | 1.04e-36 | 2.16E-35 |
| PBANKA_0914400 | 11.23 | 2.32 | 2.48e-14 | 1.80E-13 |
| PBANKA_0505800 | 7.92 | 2.32 | 9.21e-11 | 5.09E-10 |
| PBANKA_0926100 | 7.24 | 2.31 | 1.83e-11 | 1.08E-10 |
| PBANKA_1364500 | 7.01 | 2.31 | 3.85e-05 | 1.02E-04 |
| PBANKA_0910000 | 9.43 | 2.31 | 4.88e-05 | 1.27E-04 |
| PBANKA_0402800 | 8.74 | 2.3 | 3.67e-08 | 1.54E-07 |
| PBANKA_0928800 | 8.3 | 2.3 | 2.05e-07 | 7.82E-07 |
| PBANKA_1310900 | 6.99 | 2.3 | 4.46e-07 | 1.63E-06 |
| PBANKA_1116400 | 9.59 | 2.3 | 6.46e-07 | 2.31E-06 |
| PBANKA_0314400 | 6.87 | 2.3 | 7.23e-06 | 2.20E-05 |
| PBANKA_0408400 | 9.36 | 2.29 | 1.74e-18 | 1.58E-17 |

| | | | | |
|---|---|---|---|---|
| PBANKA_1320800 | 10.91 | -2.63 | 8.09e-10 | 4.09E-09 |
| PBANKA_1403200 | 5.83 | -2.63 | 2.79e-03 | 5.11E-03 |
| PBANKA_1006600 | 7.17 | -2.62 | 3.68e-10 | 1.91E-09 |
| PBANKA_0914100 | 8.73 | -2.62 | 2.30e-07 | 8.75E-07 |
| PBANKA_0938000 | 10.69 | -2.62 | 2.25e-06 | 7.40E-06 |
| PBANKA_0904500 | 8.05 | -2.62 | 2.40e-05 | 6.61E-05 |
| PBANKA_0801300 | 6.23 | -2.62 | 1.24e-03 | 2.46E-03 |
| PBANKA_1437300 | 11.37 | -2.61 | 8.72e-19 | 8.05E-18 |
| PBANKA_0405900 | 7.49 | -2.61 | 6.16e-08 | 2.51E-07 |
| PBANKA_1401800 | 6.25 | -2.61 | 2.93e-04 | 6.60E-04 |
| PBANKA_1039300 | 7.56 | -2.6 | 3.77e-12 | 2.35E-11 |
| PBANKA_1428800 | 8.33 | -2.6 | 7.90e-11 | 4.41E-10 |
| PBANKA_1209800 | 8.83 | -2.6 | 1.03e-08 | 4.62E-08 |
| PBANKA_1345400 | 8.88 | -2.6 | 4.43e-07 | 1.62E-06 |
| PBANKA_1353300 | 7.29 | -2.6 | 5.66e-06 | 1.75E-05 |
| PBANKA_0417600 | 9.15 | -2.6 | 1.27e-05 | 3.69E-05 |
| PBANKA_1009900 | 9.09 | -2.6 | 1.40e-03 | 2.74E-03 |
| PBANKA_1425900 | 9.72 | -2.59 | 7.43e-17 | 6.21E-16 |
| PBANKA_0831200 | 9.17 | -2.59 | 9.55e-16 | 7.55E-15 |
| PBANKA_1400600 | 12.93 | -2.59 | 3.20e-09 | 1.51E-08 |
| PBANKA_0712600 | 11.06 | -2.59 | 1.59e-07 | 6.23E-07 |
| PBANKA_0619500 | 7.04 | -2.59 | 1.02e-04 | 2.50E-04 |
| PBANKA_0511700 | 8.21 | -2.59 | 2.67e-04 | 6.06E-04 |
| PBANKA_0602400 | 8.89 | -2.58 | 5.68e-14 | 4.03E-13 |
| PBANKA_0716700 | 8.39 | -2.58 | 1.07e-10 | 5.89E-10 |
| PBANKA_0519400 | 11.17 | -2.58 | 1.09e-09 | 5.42E-09 |
| PBANKA_0827200 | 8.92 | -2.58 | 4.52e-08 | 1.88E-07 |
| PBANKA_1039200 | 6.99 | -2.58 | 2.67e-06 | 8.72E-06 |
| PBANKA_0806700 | 10.39 | -2.58 | 1.34e-05 | 3.86E-05 |
| PBANKA_0622700 | 7.03 | -2.58 | 1.57e-05 | 4.48E-05 |
| PBANKA_0800900 | 7.95 | -2.57 | 3.58e-09 | 1.68E-08 |
| PBANKA_1211700 | 7.75 | -2.57 | 1.67e-08 | 7.32E-08 |
| PBANKA_1128700 | 7.41 | -2.57 | 1.67e-07 | 6.50E-07 |
| PBANKA_0608100 | 7.82 | -2.57 | 6.09e-07 | 2.18E-06 |
| PBANKA_1142400 | 7.33 | -2.57 | 7.34e-06 | 2.23E-05 |
| PBANKA_1418500 | 6.27 | -2.57 | 1.62e-03 | 3.13E-03 |
| PBANKA_0418100 | 9.34 | -2.56 | 1.29e-06 | 4.42E-06 |
| PBANKA_0721000 | 8.88 | -2.56 | 1.79e-04 | 4.20E-04 |
| PBANKA_0207300 | 7.91 | -2.55 | 5.71e-12 | 3.51E-11 |
| PBANKA_0108300 | 8.3 | -2.55 | 2.18e-07 | 8.31E-07 |
| PBANKA_0807800 | 6.4 | -2.55 | 4.32e-05 | 1.14E-04 |
| PBANKA_0827900 | 6.38 | -2.55 | 6.00e-04 | 1.26E-03 |
| PBANKA_1453000 | 9.53 | -2.54 | 5.05e-17 | 4.27E-16 |
| PBANKA_0815900 | 8.85 | -2.54 | 2.14e-14 | 1.57E-13 |
| PBANKA_1040100 | 9.72 | -2.54 | 2.39e-09 | 1.14E-08 |
| PBANKA_1334000 | 8.17 | -2.54 | 4.97e-08 | 2.05E-07 |
| PBANKA_0509600 | 7.95 | -2.54 | 6.73e-07 | 2.40E-06 |
| PBANKA_0917100 | 7.63 | -2.54 | 4.74e-05 | 1.24E-04 |
| PBANKA_0416400 | 6.77 | -2.54 | 9.46e-05 | 2.34E-04 |
| PBANKA_0104700 | 7.62 | -2.54 | 1.14e-04 | 2.76E-04 |
| PBANKA_0701500 | 7.17 | -2.54 | 1.08e-03 | 2.16E-03 |
| PBANKA_0416500 | 9.51 | -2.53 | 2.65e-15 | 2.05E-14 |
| PBANKA_0305700 | 7.88 | -2.53 | 1.17e-09 | 5.79E-09 |
| PBANKA_1444700 | 8.6 | -2.53 | 3.64e-09 | 1.71E-08 |
| PBANKA_1005400 | 9.19 | -2.53 | 3.17e-07 | 1.18E-06 |
| PBANKA_0923200 | 7.28 | -2.53 | 8.75e-06 | 2.63E-05 |
| PBANKA_0918900 | 8.28 | -2.53 | 4.59e-05 | 1.20E-04 |
| PBANKA_1233000 | 6.97 | -2.53 | 9.13e-05 | 2.26E-04 |
| PBANKA_1406400 | 8.37 | -2.52 | 4.24e-14 | 3.03E-13 |
| PBANKA_1112300 | 8.07 | -2.52 | 5.17e-11 | 2.92E-10 |
| PBANKA_1413200 | 7.7 | -2.52 | 1.01e-07 | 4.05E-07 |
| PBANKA_0705200 | 6.84 | -2.52 | 3.84e-04 | 8.43E-04 |
| PBANKA_1224800 | 5.8 | -2.52 | 1.01e-03 | 2.04E-03 |
| PBANKA_1039600 | 8.36 | -2.51 | 2.88e-10 | 1.52E-09 |
| PBANKA_1352800 | 8.12 | -2.51 | 4.19e-08 | 1.75E-07 |
| PBANKA_0506400 | 7.3 | -2.51 | 1.44e-07 | 5.68E-07 |
| PBANKA_1137900 | 8.89 | -2.51 | 9.43e-06 | 2.81E-05 |
| PBANKA_1104500 | 6.87 | -2.51 | 2.34e-03 | 4.38E-03 |
| PBANKA_1016400 | 8.05 | -2.5 | 1.63e-13 | 1.14E-12 |
| PBANKA_0835500 | 8.95 | -2.5 | 1.85e-10 | 9.92E-10 |
| PBANKA_0208100 | 8.29 | -2.5 | 2.17e-08 | 9.36E-08 |
| PBANKA_1421200 | 7.46 | -2.5 | 8.74e-08 | 3.51E-07 |
| PBANKA_0925800 | 7.11 | -2.5 | 9.00e-07 | 3.15E-06 |
| PBANKA_0912600 | 8.83 | -2.5 | 1.16e-05 | 3.41E-05 |

| | | | | |
|---|---|---|---|---|
| PBANKA_1318200 | 8.89 | 2.29 | 1.11e-05 | 3.28E-05 |
| PBANKA_0934500 | 9.41 | 2.28 | 5.34e-21 | 5.51E-20 |
| PBANKA_1032700 | 9.37 | 2.28 | 6.84e-20 | 6.67E-19 |
| PBANKA_0913800 | 8.92 | 2.27 | 3.69e-25 | 4.63E-24 |
| PBANKA_0502900 | 8.54 | 2.27 | 3.10e-15 | 2.38E-14 |
| PBANKA_0805700 | 13.63 | 2.27 | 8.06e-12 | 4.88E-11 |
| PBANKA_0813000 | 11.57 | 2.27 | 4.38e-10 | 2.27E-09 |
| PBANKA_1205900 | 10.59 | 2.26 | 1.30e-16 | 1.07E-15 |
| PBANKA_1203800 | 8.99 | 2.25 | 3.02e-16 | 2.43E-15 |
| PBANKA_1015200 | 9.36 | 2.25 | 4.60e-15 | 3.52E-14 |
| PBANKA_0702900 | 11.17 | 2.25 | 3.57e-05 | 9.57E-05 |
| PBANKA_0927800 | 7.82 | 2.24 | 5.78e-18 | 5.13E-17 |
| PBANKA_1114400 | 9.22 | 2.23 | 2.88e-17 | 2.47E-16 |
| PBANKA_0823500 | 8.3 | 2.23 | 2.23e-14 | 1.63E-13 |
| PBANKA_1305000 | 11.63 | 2.23 | 1.19e-08 | 5.30E-08 |
| PBANKA_1332800 | 10.87 | 2.23 | 4.15e-05 | 1.09E-04 |
| PBANKA_1429500 | 7.54 | 2.21 | 3.92e-07 | 1.44E-06 |
| PBANKA_0201250 | 7.04 | 2.21 | 2.00e-05 | 5.58E-05 |
| PBANKA_1232000 | 7.59 | 2.2 | 1.29e-04 | 3.08E-04 |
| PBANKA_0836600 | 6.03 | 2.2 | 4.68e-04 | 1.01E-03 |
| PBANKA_0831800 | 7.79 | 2.19 | 1.24e-12 | 8.04E-12 |
| PBANKA_0509200 | 10.1 | 2.19 | 3.32e-09 | 1.57E-08 |
| PBANKA_1407900 | 10.5 | 2.18 | 5.17e-09 | 2.40E-08 |
| PBANKA_0800400 | 9.23 | 2.18 | 8.21e-08 | 3.31E-07 |
| PBANKA_1411500 | 7.99 | 2.18 | 4.36e-07 | 1.59E-06 |
| PBANKA_1325900 | 5.46 | 2.18 | 1.34e-05 | 3.86E-05 |
| PBANKA_1328300 | 8.39 | 2.17 | 1.58e-16 | 1.30E-15 |
| PBANKA_1119400 | 5.51 | 2.17 | 5.11e-03 | 8.89E-03 |
| PBANKA_1037200 | 7.97 | 2.16 | 1.97e-19 | 1.89E-18 |
| PBANKA_1117100 | 11.42 | 2.16 | 2.69e-07 | 1.02E-06 |
| PBANKA_0110400 | 8.49 | 2.15 | 4.36e-19 | 4.09E-18 |
| PBANKA_1021000 | 8.65 | 2.15 | 2.00e-10 | 1.07E-09 |
| PBANKA_0105300 | 10.52 | 2.15 | 4.24e-06 | 1.34E-05 |
| PBANKA_0408300 | 10.51 | 2.14 | 7.39e-05 | 1.86E-04 |
| PBANKA_1414400 | 8.34 | 2.13 | 1.10e-22 | 1.25E-21 |
| PBANKA_1005000 | 5.18 | 2.13 | 1.06e-04 | 2.59E-04 |
| PBANKA_1240900 | 5.16 | 2.13 | 2.79e-03 | 5.11E-03 |
| PBANKA_1003700 | 9.88 | 2.12 | 6.43e-20 | 6.31E-19 |
| PBANKA_0622800 | 9.38 | 2.12 | 4.65e-16 | 3.70E-15 |
| PBANKA_1033500 | 6.55 | 2.12 | 3.69e-09 | 1.73E-08 |
| PBANKA_1132000 | 8.38 | 2.12 | 3.87e-09 | 1.81E-08 |
| PBANKA_0523200 | 6.65 | 2.12 | 3.84e-06 | 1.22E-05 |
| PBANKA_0103300 | 10.81 | 2.12 | 1.44e-05 | 4.13E-05 |
| PBANKA_1363700 | 9.12 | 2.11 | 1.44e-10 | 7.81E-10 |
| PBANKA_1102100 | 8.31 | 2.11 | 1.45e-07 | 5.68E-07 |
| PBANKA_1306500 | 5.97 | 2.11 | 1.08e-03 | 2.16E-03 |
| PBANKA_1452500 | 10.43 | 2.09 | 3.33e-14 | 2.40E-13 |
| PBANKA_0212100 | 10.22 | 2.09 | 8.20e-13 | 5.40E-12 |
| PBANKA_1003100 | 11.11 | 2.09 | 3.28e-10 | 1.72E-09 |
| PBANKA_1121700 | 10.49 | 2.09 | 1.18e-07 | 4.69E-07 |
| PBANKA_0209700 | 10.76 | 2.09 | 7.31e-05 | 1.84E-04 |
| PBANKA_1010500 | 10.53 | 2.08 | 5.32e-15 | 4.04E-14 |
| PBANKA_1327021 | 9.72 | 2.08 | 4.39e-07 | 1.60E-06 |
| PBANKA_1107400 | 9.51 | 2.07 | 2.21e-22 | 2.45E-21 |
| PBANKA_1010800 | 7.91 | 2.07 | 2.78e-07 | 1.05E-06 |
| PBANKA_0906600 | 8.1 | 2.06 | 2.49e-13 | 1.71E-12 |
| PBANKA_0833500 | 7.98 | 2.06 | 2.65e-12 | 1.67E-11 |
| PBANKA_0816600 | 8.05 | 2.06 | 2.52e-09 | 1.21E-08 |
| PBANKA_0524300 | 11.28 | 2.06 | 5.75e-08 | 2.36E-07 |
| PBANKA_1234500 | 9 | 2.04 | 1.39e-07 | 5.48E-07 |
| PBANKA_0205800 | 11.74 | 2.04 | 4.75e-05 | 1.24E-04 |
| PBANKA_1452200 | 8.12 | 2.03 | 1.46e-16 | 1.20E-15 |
| PBANKA_1013500 | 9.29 | 2.03 | 6.98e-13 | 4.65E-12 |
| PBANKA_0308600 | 9.87 | 2.03 | 2.98e-12 | 1.87E-11 |
| PBANKA_0404600 | 8.91 | 2.03 | 1.58e-07 | 6.20E-07 |
| PBANKA_0941800 | 13.62 | 2.03 | 8.38e-06 | 2.53E-05 |
| PBANKA_1359100 | 5.95 | 2.03 | 5.82e-05 | 1.49E-04 |
| PBANKA_0803600 | 9.3 | 2.02 | 1.15e-11 | 6.90E-11 |
| PBANKA_1002300 | 9.84 | 2.02 | 4.15e-07 | 1.53E-06 |
| PBANKA_1005700 | 8.91 | 2 | 4.27e-06 | 1.35E-05 |
| PBANKA_0511000 | 7.75 | 2 | 6.83e-05 | 1.73E-04 |

| | | | | |
|---|---|---|---|---|
| PBANKA_0609200 | 6.74 | -2.5 | 1.18e-04 | 2.85E-04 |
| PBANKA_0406100 | 7.95 | -2.49 | 9.32e-12 | 5.60E-11 |
| PBANKA_1424500 | 8.48 | -2.49 | 7.48e-11 | 4.18E-10 |
| PBANKA_1024900 | 9.31 | -2.49 | 8.97e-11 | 4.97E-10 |
| PBANKA_1328100 | 8.57 | -2.49 | 6.33e-08 | 2.58E-07 |
| PBANKA_0914300 | 7.83 | -2.49 | 1.51e-07 | 5.91E-07 |
| PBANKA_1218300 | 9.36 | -2.49 | 5.99e-06 | 1.85E-05 |
| PBANKA_0931000 | 6.94 | -2.49 | 1.93e-04 | 4.51E-04 |
| PBANKA_0409000 | 7.33 | -2.49 | 7.05e-04 | 1.46E-03 |
| PBANKA_1000011 | 6.91 | -2.49 | 2.58e-03 | 4.77E-03 |
| PBANKA_0722941 | 5.66 | -2.49 | 3.86e-03 | 6.85E-03 |
| PBANKA_1351200 | 7.98 | -2.48 | 3.33e-09 | 1.57E-08 |
| PBANKA_0414400 | 9.5 | -2.48 | 4.08e-08 | 1.70E-07 |
| PBANKA_1330600 | 7.65 | -2.48 | 2.19e-06 | 7.21E-06 |
| PBANKA_1444200 | 8.83 | -2.48 | 7.03e-06 | 2.14E-05 |
| PBANKA_1447400 | 6.24 | -2.48 | 2.58e-04 | 5.87E-04 |
| PBANKA_0834000 | 6.12 | -2.48 | 3.69e-04 | 8.13E-04 |
| PBANKA_0504100 | 5.98 | -2.48 | 6.62e-04 | 1.38E-03 |
| PBANKA_0303500 | 6.32 | -2.48 | 7.92e-04 | 1.64E-03 |
| PBANKA_0925400 | 8.64 | -2.48 | 2.21e-03 | 4.15E-03 |
| PBANKA_1318900 | 6.4 | -2.48 | 3.68e-03 | 6.56E-03 |
| PBANKA_0305000 | 12.49 | -2.47 | 3.92e-08 | 1.64E-07 |
| PBANKA_0508900 | 8.24 | -2.47 | 2.04e-07 | 7.79E-07 |
| PBANKA_1216200 | 7.37 | -2.47 | 1.05e-06 | 3.65E-06 |
| PBANKA_0110500 | 10.46 | -2.47 | 1.81e-04 | 4.26E-04 |
| PBANKA_1348000 | 6.72 | -2.47 | 1.03e-03 | 2.08E-03 |
| PBANKA_1453300 | 9.99 | -2.46 | 3.63e-23 | 4.22E-22 |
| PBANKA_1402000 | 9.24 | -2.46 | 1.33e-15 | 1.05E-14 |
| PBANKA_1219900 | 7.95 | -2.46 | 1.12e-12 | 7.32E-12 |
| PBANKA_0927000 | 8.25 | -2.46 | 1.89e-10 | 1.01E-09 |
| PBANKA_1457800 | 8.32 | -2.46 | 4.95e-10 | 2.55E-09 |
| PBANKA_0208300 | 9.35 | -2.46 | 1.58e-07 | 6.18E-07 |
| PBANKA_1442200 | 12.24 | -2.46 | 7.43e-07 | 2.63E-06 |
| PBANKA_0313600 | 6.32 | -2.46 | 1.19e-05 | 3.48E-05 |
| PBANKA_0616600 | 7.14 | -2.46 | 5.57e-03 | 9.62E-03 |
| PBANKA_1415400 | 9.91 | -2.45 | 2.53e-15 | 1.96E-14 |
| PBANKA_1434700 | 9.7 | -2.45 | 3.40e-12 | 2.12E-11 |
| PBANKA_1456400 | 6.78 | -2.45 | 7.39e-07 | 2.62E-06 |
| PBANKA_1144000 | 8.53 | -2.45 | 6.86e-06 | 2.09E-05 |
| PBANKA_1356800 | 11.16 | -2.45 | 7.72e-04 | 1.60E-03 |
| PBANKA_0106600 | 6.71 | -2.45 | 3.60e-03 | 6.45E-03 |
| PBANKA_1446800 | 9.45 | -2.44 | 6.24e-10 | 3.19E-09 |
| PBANKA_1320200 | 10.3 | -2.44 | 1.04e-08 | 4.67E-08 |
| PBANKA_1115300 | 12.1 | -2.44 | 7.89e-08 | 3.18E-07 |
| PBANKA_1410600 | 8.43 | -2.44 | 3.45e-07 | 1.28E-06 |
| PBANKA_1119300 | 6.96 | -2.44 | 4.32e-04 | 9.38E-04 |
| PBANKA_0936400 | 8.8 | -2.43 | 2.25e-10 | 1.19E-09 |
| PBANKA_1108100 | 8.81 | -2.43 | 1.31e-09 | 6.42E-09 |
| PBANKA_1018900 | 6.85 | -2.43 | 6.16e-07 | 2.21E-06 |
| PBANKA_0908000 | 7.83 | -2.43 | 8.62e-06 | 2.60E-05 |
| PBANKA_0618600 | 9.56 | -2.42 | 2.22e-11 | 1.29E-10 |
| PBANKA_0209000 | 8.84 | -2.42 | 3.33e-09 | 1.57E-08 |
| PBANKA_1206400 | 7.18 | -2.42 | 1.01e-07 | 4.03E-07 |
| PBANKA_1433700 | 8.31 | -2.42 | 9.77e-07 | 3.41E-06 |
| PBANKA_1305700 | 7.69 | -2.42 | 1.39e-04 | 3.32E-04 |
| PBANKA_1301100 | 7.1 | -2.42 | 2.27e-04 | 5.22E-04 |
| PBANKA_0717100 | 6.14 | -2.42 | 1.33e-03 | 2.62E-03 |
| PBANKA_0614700 | 6.28 | -2.42 | 3.40e-03 | 6.13E-03 |
| PBANKA_1365100 | 8.36 | -2.41 | 2.71e-12 | 1.71E-11 |
| PBANKA_0402300 | 7.41 | -2.41 | 1.67e-08 | 7.34E-08 |
| PBANKA_1463200 | 7.17 | -2.41 | 2.16e-08 | 9.36E-08 |
| PBANKA_1015900 | 7.53 | -2.41 | 3.34e-07 | 1.24E-06 |
| PBANKA_1441200 | 7.54 | -2.41 | 6.21e-07 | 2.22E-06 |
| PBANKA_1334100 | 8.23 | -2.41 | 5.23e-06 | 1.63E-05 |
| PBANKA_1302800 | 9.54 | -2.41 | 5.50e-05 | 1.42E-04 |
| PBANKA_0302400 | 7.16 | -2.41 | 8.13e-05 | 2.03E-04 |
| PBANKA_1353100 | 11.09 | -2.41 | 1.28e-04 | 3.07E-04 |
| PBANKA_1312000 | 6.81 | -2.41 | 2.60e-03 | 4.81E-03 |
| PBANKA_1241600 | 5.39 | -2.41 | 3.63e-03 | 6.48E-03 |
| PBANKA_1106500 | 10.46 | -2.4 | 2.46e-10 | 1.30E-09 |
| PBANKA_0601600 | 10.64 | -2.4 | 2.59e-07 | 9.80E-07 |
| PBANKA_1025000 | 7.71 | -2.4 | 4.31e-07 | 1.58E-06 |
| PBANKA_0912700 | 7.34 | -2.4 | 1.54e-06 | 5.16E-06 |
| PBANKA_1420000 | 7.27 | -2.4 | 1.14e-05 | 3.35E-05 |

| | | | | |
|---|---|---|---|---|
| PBANKA_1021300 | 6.65 | -2.4 | 2.40e-04 | 5.48E-04 |
| PBANKA_0404300 | 6.61 | -2.4 | 5.02e-04 | 1.07E-03 |
| PBANKA_0701300 | 5.87 | -2.4 | 3.29e-03 | 5.95E-03 |
| PBANKA_0937100 | 8.12 | -2.39 | 5.58e-12 | 3.44E-11 |
| PBANKA_0702800 | 11.36 | -2.39 | 1.17e-08 | 5.21E-08 |
| PBANKA_1135900 | 7.56 | -2.39 | 2.88e-08 | 1.23E-07 |
| PBANKA_0604300 | 9.67 | -2.39 | 3.32e-08 | 1.40E-07 |
| PBANKA_1134700 | 7.03 | -2.39 | 3.81e-05 | 1.02E-04 |
| PBANKA_1143900 | 4.82 | -2.39 | 2.98e-03 | 5.43E-03 |
| PBANKA_1105100 | 8.64 | -2.38 | 2.05e-16 | 1.67E-15 |
| PBANKA_1218600 | 6.51 | -2.38 | 1.06e-04 | 2.60E-04 |
| PBANKA_1455900 | 6.45 | -2.38 | 1.11e-04 | 2.71E-04 |
| PBANKA_0605100 | 8.28 | -2.38 | 4.29e-04 | 9.33E-04 |
| PBANKA_1216800 | 10.27 | -2.38 | 5.41e-04 | 1.15E-03 |
| PBANKA_0104000 | 7.26 | -2.38 | 6.57e-04 | 1.37E-03 |
| PBANKA_0612200 | 9.84 | -2.37 | 9.46e-30 | 1.45E-28 |
| PBANKA_0304800 | 10.08 | -2.37 | 9.45e-12 | 5.67E-11 |
| PBANKA_0416600 | 8.62 | -2.37 | 7.89e-10 | 3.99E-09 |
| PBANKA_1023000 | 8.81 | -2.37 | 1.44e-09 | 7.02E-09 |
| PBANKA_1228100 | 11.03 | -2.37 | 1.43e-06 | 4.84E-06 |
| PBANKA_1362400 | 7.25 | -2.37 | 3.22e-06 | 1.04E-05 |
| PBANKA_1401600 | 8.21 | -2.37 | 9.38e-06 | 2.79E-05 |
| PBANKA_0605500 | 8.66 | -2.37 | 6.32e-05 | 1.61E-04 |
| PBANKA_0608800 | 6.59 | -2.37 | 2.52e-04 | 5.75E-04 |
| PBANKA_0602000 | 9.35 | -2.36 | 4.48e-16 | 3.57E-15 |
| PBANKA_1342700 | 7.48 | -2.36 | 1.00e-12 | 6.55E-12 |
| PBANKA_0943100 | 10.28 | -2.36 | 1.13e-06 | 3.90E-06 |
| PBANKA_1243000 | 7.77 | -2.36 | 3.70e-05 | 9.87E-05 |
| PBANKA_1422100 | 6.71 | -2.36 | 1.90e-03 | 3.62E-03 |
| PBANKA_0721700 | 9.96 | -2.35 | 6.59e-11 | 3.70E-10 |
| PBANKA_1362900 | 9.07 | -2.35 | 2.43e-10 | 1.29E-09 |
| PBANKA_1323200 | 7.53 | -2.35 | 6.56e-08 | 2.67E-07 |
| PBANKA_1448900 | 7.8 | -2.35 | 1.21e-07 | 4.79E-07 |
| PBANKA_0800700 | 7.06 | -2.35 | 2.73e-06 | 8.87E-06 |
| PBANKA_0508400 | 6.72 | -2.35 | 2.05e-04 | 4.74E-04 |
| PBANKA_0520400 | 8.07 | -2.34 | 8.96e-13 | 5.88E-12 |
| PBANKA_1438500 | 8.55 | -2.34 | 4.93e-10 | 2.55E-09 |
| PBANKA_0510100 | 9.22 | -2.34 | 1.43e-09 | 6.99E-09 |
| PBANKA_1416100 | 7.57 | -2.34 | 1.09e-08 | 4.88E-08 |
| PBANKA_1008200 | 7.76 | -2.34 | 1.12e-06 | 3.86E-06 |
| PBANKA_1011700 | 6.83 | -2.34 | 6.04e-05 | 1.55E-04 |
| PBANKA_1439000 | 6.59 | -2.34 | 5.59e-04 | 1.18E-03 |
| PBANKA_1435100 | 6.66 | -2.34 | 1.90e-03 | 3.62E-03 |
| PBANKA_1225700 | 9.22 | -2.33 | 2.43e-15 | 1.88E-14 |
| PBANKA_0933000 | 9.53 | -2.33 | 1.80e-14 | 1.32E-13 |
| PBANKA_0503700 | 8.76 | -2.33 | 3.36e-10 | 1.76E-09 |
| PBANKA_1201200 | 8.01 | -2.33 | 8.30e-10 | 4.19E-09 |
| PBANKA_0308300 | 9.6 | -2.33 | 1.37e-09 | 6.72E-09 |
| PBANKA_1206600 | 9.53 | -2.33 | 1.29e-07 | 5.09E-07 |
| PBANKA_0309000 | 6.61 | -2.33 | 6.33e-05 | 1.61E-04 |
| PBANKA_0206400 | 6.76 | -2.33 | 1.75e-04 | 4.11E-04 |
| PBANKA_0926300 | 6.76 | -2.33 | 1.93e-04 | 4.49E-04 |
| PBANKA_0827800 | 6.09 | -2.33 | 7.19e-04 | 1.49E-03 |
| PBANKA_1429400 | 6.83 | -2.33 | 2.11e-03 | 3.97E-03 |
| PBANKA_0501400 | 8.9 | -2.32 | 8.13e-08 | 3.27E-07 |
| PBANKA_1303900 | 9.7 | -2.32 | 2.72e-06 | 8.84E-06 |
| PBANKA_1120400 | 10.96 | -2.32 | 1.54e-05 | 4.40E-05 |
| PBANKA_0906100 | 6.59 | -2.32 | 4.64e-05 | 1.21E-04 |
| PBANKA_1449300 | 9.16 | -2.32 | 3.08e-04 | 6.91E-04 |
| PBANKA_0828400 | 8.8 | -2.31 | 1.23e-18 | 1.13E-17 |
| PBANKA_1356300 | 9.42 | -2.31 | 7.11e-15 | 5.34E-14 |
| PBANKA_0102600 | 9.57 | -2.31 | 1.07e-14 | 7.94E-14 |
| PBANKA_1457900 | 8.31 | -2.31 | 4.45e-12 | 2.75E-11 |
| PBANKA_0908800 | 8.55 | -2.31 | 8.61e-10 | 4.34E-09 |
| PBANKA_1206900 | 10.82 | -2.31 | 1.32e-05 | 3.81E-05 |
| PBANKA_1426500 | 8.83 | -2.3 | 2.34e-12 | 1.48E-11 |
| PBANKA_1360500 | 8.72 | -2.3 | 3.41e-10 | 1.78E-09 |
| PBANKA_1145600 | 8.88 | -2.3 | 4.15e-10 | 2.15E-09 |
| PBANKA_0402400 | 7.41 | -2.3 | 2.86e-05 | 7.79E-05 |
| PBANKA_1232800 | 7.39 | -2.3 | 4.87e-05 | 1.27E-04 |
| PBANKA_1104400 | 10.14 | -2.3 | 9.66e-05 | 2.38E-04 |
| PBANKA_1017600 | 8.68 | -2.29 | 7.83e-08 | 3.16E-07 |
| PBANKA_0821100 | 7.96 | -2.29 | 3.69e-07 | 1.37E-06 |
| PBANKA_1101400 | 10.48 | -2.29 | 7.34e-07 | 2.61E-06 |

| | | | | |
|---|---|---|---|---|
| PBANKA_0523900 | 11.44 | -2.29 | 1.04e-06 | 3.63E-06 |
| PBANKA_0615800 | 9.26 | -2.29 | 3.01e-06 | 9.72E-06 |
| PBANKA_0835200 | 7.88 | -2.29 | 1.09e-05 | 3.21E-05 |
| PBANKA_0910700 | 9.17 | -2.29 | 6.73e-05 | 1.71E-04 |
| PBANKA_0828200 | 7.46 | -2.29 | 1.08e-04 | 2.64E-04 |
| PBANKA_1401500 | 7.3 | -2.29 | 2.20e-04 | 5.08E-04 |
| PBANKA_0702400 | 8.99 | -2.28 | 1.51e-22 | 1.69E-21 |
| PBANKA_1231400 | 8.82 | -2.28 | 3.81e-12 | 2.37E-11 |
| PBANKA_1336900 | 8.54 | -2.28 | 1.74e-11 | 1.03E-10 |
| PBANKA_1301000 | 7.94 | -2.28 | 8.73e-10 | 4.38E-09 |
| PBANKA_0715900 | 8.12 | -2.28 | 7.89e-09 | 3.59E-08 |
| PBANKA_0819500 | 7.87 | -2.28 | 1.65e-08 | 7.26E-08 |
| PBANKA_1038900 | 9.57 | -2.28 | 7.58e-07 | 2.69E-06 |
| PBANKA_1320000 | 6.84 | -2.28 | 1.70e-06 | 5.67E-06 |
| PBANKA_1244200 | 8.79 | -2.28 | 1.44e-05 | 4.15E-05 |
| PBANKA_1132300 | 9.79 | -2.28 | 1.86e-04 | 4.35E-04 |
| PBANKA_0936000 | 8.64 | -2.28 | 1.34e-03 | 2.64E-03 |
| PBANKA_0401600 | 7.98 | -2.27 | 2.29e-09 | 1.10E-08 |
| PBANKA_0815200 | 8.27 | -2.27 | 5.21e-09 | 2.41E-08 |
| PBANKA_1427100 | 9.23 | -2.27 | 4.95e-08 | 2.05E-07 |
| PBANKA_1242000 | 8.14 | -2.27 | 1.06e-07 | 4.21E-07 |
| PBANKA_1418900 | 8.27 | -2.27 | 3.35e-06 | 1.08E-05 |
| PBANKA_0802400 | 7.18 | -2.27 | 1.20e-05 | 3.50E-05 |
| PBANKA_1430700 | 8.06 | -2.27 | 1.41e-05 | 4.06E-05 |
| PBANKA_0213200 | 10.09 | -2.27 | 3.56e-05 | 9.53E-05 |
| PBANKA_0802600 | 6.21 | -2.27 | 5.03e-04 | 1.08E-03 |
| PBANKA_0301200 | 6.25 | -2.27 | 1.35e-03 | 2.65E-03 |
| PBANKA_0316400 | 5.62 | -2.27 | 1.78e-03 | 3.40E-03 |
| PBANKA_1108800 | 5.61 | -2.27 | 3.47e-03 | 6.25E-03 |
| PBANKA_0910200 | 9.38 | -2.26 | 1.89e-09 | 9.11E-09 |
| PBANKA_0614300 | 7.16 | -2.26 | 1.95e-06 | 6.46E-06 |
| PBANKA_1459900 | 7.6 | -2.26 | 3.88e-05 | 1.03E-04 |
| PBANKA_0822400 | 6.35 | -2.26 | 2.28e-04 | 5.24E-04 |
| PBANKA_0606800 | 6.76 | -2.26 | 1.06e-03 | 2.14E-03 |
| PBANKA_0519000 | 11.39 | -2.26 | 1.80e-03 | 3.44E-03 |
| PBANKA_0800800 | 6.31 | -2.26 | 2.13e-03 | 4.01E-03 |
| PBANKA_0613200 | 9.37 | -2.25 | 1.90e-15 | 1.48E-14 |
| PBANKA_0502600 | 8.85 | -2.25 | 1.91e-12 | 1.22E-11 |
| PBANKA_1025400 | 7.47 | -2.25 | 2.55e-11 | 1.48E-10 |
| PBANKA_1131500 | 8.7 | -2.25 | 5.81e-10 | 2.98E-09 |
| PBANKA_0822600 | 7.61 | -2.25 | 9.04e-10 | 4.52E-09 |
| PBANKA_1344100 | 9.64 | -2.25 | 3.17e-08 | 1.35E-07 |
| PBANKA_0310300 | 7.06 | -2.25 | 4.32e-04 | 9.39E-04 |
| PBANKA_1332200 | 5.67 | -2.25 | 6.51e-04 | 1.36E-03 |
| PBANKA_0409400 | 10.28 | -2.25 | 1.06e-03 | 2.13E-03 |
| PBANKA_1223600 | 6.58 | -2.25 | 2.67e-03 | 4.91E-03 |
| PBANKA_0405100 | 6.35 | -2.25 | 2.75e-03 | 5.06E-03 |
| PBANKA_1319100 | 6.69 | -2.25 | 4.37e-03 | 7.70E-03 |
| PBANKA_0111600 | 8.86 | -2.24 | 5.74e-11 | 3.23E-10 |
| PBANKA_1123300 | 9.42 | -2.24 | 1.10e-10 | 6.02E-10 |
| PBANKA_1438200 | 9.1 | -2.24 | 5.59e-09 | 2.58E-08 |
| PBANKA_0410800 | 7.85 | -2.24 | 2.20e-07 | 8.38E-07 |
| PBANKA_1219200 | 8.57 | -2.24 | 3.65e-07 | 1.35E-06 |
| PBANKA_0621300 | 10.15 | -2.24 | 1.94e-06 | 6.42E-06 |
| PBANKA_1233100 | 8.31 | -2.24 | 2.15e-06 | 7.08E-06 |
| PBANKA_0806300 | 7.46 | -2.24 | 6.49e-06 | 1.99E-05 |
| PBANKA_1306300 | 9.23 | -2.24 | 1.57e-04 | 3.72E-04 |
| PBANKA_1429200 | 10.13 | -2.24 | 5.16e-03 | 8.96E-03 |
| PBANKA_1031300 | 7.86 | -2.23 | 1.75e-12 | 1.12E-11 |
| PBANKA_0919400 | 8.07 | -2.23 | 7.56e-10 | 3.83E-09 |
| PBANKA_0612800 | 7.46 | -2.23 | 8.68e-10 | 4.36E-09 |
| PBANKA_1434600 | 10.1 | -2.23 | 6.37e-05 | 1.62E-04 |
| PBANKA_1039800 | 6.28 | -2.23 | 2.24e-04 | 5.15E-04 |
| PBANKA_0301000 | 10.05 | -2.23 | 4.12e-04 | 8.97E-04 |
| PBANKA_1136400 | 6.78 | -2.23 | 5.40e-04 | 1.15E-03 |
| PBANKA_1030400 | 6.89 | -2.23 | 5.89e-04 | 1.24E-03 |
| PBANKA_1029200 | 6.5 | -2.23 | 9.80e-04 | 1.98E-03 |
| PBANKA_0903800 | 11.26 | -2.22 | 8.35e-07 | 2.95E-06 |
| PBANKA_0803200 | 10.33 | -2.22 | 1.30e-06 | 4.44E-06 |
| PBANKA_1428500 | 7.38 | -2.22 | 1.62e-06 | 5.43E-06 |
| PBANKA_1452000 | 9.09 | -2.22 | 3.57e-06 | 1.14E-05 |
| PBANKA_1224600 | 8.5 | -2.22 | 5.42e-06 | 1.69E-05 |
| PBANKA_1019900 | 7.15 | -2.22 | 1.74e-05 | 4.93E-05 |
| PBANKA_0507300 | 9.07 | -2.22 | 5.30e-04 | 1.13E-03 |

| | | | | |
|---|---|---|---|---|
| PBANKA_0834400 | 9.23 | -2.21 | 3.69e-11 | 2.11E-10 |
| PBANKA_1305500 | 9.03 | -2.21 | 1.07e-08 | 4.78E-08 |
| PBANKA_0413200 | 7.84 | -2.21 | 2.18e-08 | 9.40E-08 |
| PBANKA_0415900 | 6.83 | -2.21 | 1.37e-06 | 4.67E-06 |
| PBANKA_1313200 | 9.78 | -2.21 | 1.66e-04 | 3.92E-04 |
| PBANKA_1336500 | 6 | -2.21 | 3.70e-04 | 8.16E-04 |
| PBANKA_1228600 | 6.58 | -2.21 | 4.55e-03 | 7.97E-03 |
| PBANKA_0211900 | 8.4 | -2.2 | 1.05e-14 | 7.87E-14 |
| PBANKA_1307700 | 8.08 | -2.2 | 4.18e-11 | 2.37E-10 |
| PBANKA_0932800 | 8.2 | -2.2 | 7.23e-07 | 2.57E-06 |
| PBANKA_1028200 | 10.72 | -2.2 | 1.38e-06 | 4.71E-06 |
| PBANKA_1312900 | 8.95 | -2.2 | 6.37e-06 | 1.95E-05 |
| PBANKA_0511800 | 7.15 | -2.2 | 2.67e-05 | 7.30E-05 |
| PBANKA_0418200 | 8.11 | -2.2 | 6.50e-05 | 1.65E-04 |
| PBANKA_1432400 | 7.95 | -2.2 | 1.13e-04 | 2.75E-04 |
| PBANKA_1220800 | 11.23 | -2.2 | 5.37e-04 | 1.14E-03 |
| PBANKA_1105200 | 6.84 | -2.2 | 1.51e-03 | 2.94E-03 |
| PBANKA_0212800 | 7.62 | -2.19 | 1.79e-09 | 8.67E-09 |
| PBANKA_0513800 | 8.42 | -2.19 | 1.50e-08 | 6.63E-08 |
| PBANKA_1206200 | 9.71 | -2.19 | 1.66e-08 | 7.30E-08 |
| PBANKA_1244600 | 7.78 | -2.19 | 1.70e-05 | 4.84E-05 |
| PBANKA_1449100 | 6.11 | -2.19 | 5.51e-03 | 9.52E-03 |
| PBANKA_1349700 | 7.48 | -2.18 | 1.78e-09 | 8.64E-09 |
| PBANKA_0405200 | 9.72 | -2.18 | 2.25e-07 | 8.58E-07 |
| PBANKA_0610200 | 9.65 | -2.18 | 1.18e-05 | 3.46E-05 |
| PBANKA_1125700 | 6.96 | -2.18 | 1.19e-04 | 2.89E-04 |
| PBANKA_0107100 | 8.16 | -2.18 | 1.57e-04 | 3.71E-04 |
| PBANKA_1352500 | 9.81 | -2.18 | 4.83e-04 | 1.04E-03 |
| PBANKA_1139500 | 6.61 | -2.18 | 8.48e-04 | 1.74E-03 |
| PBANKA_1418700 | 5.72 | -2.18 | 4.38e-03 | 7.70E-03 |
| PBANKA_1303300 | 9.39 | -2.17 | 1.81e-09 | 8.77E-09 |
| PBANKA_1415500 | 7.66 | -2.17 | 1.18e-08 | 5.23E-08 |
| PBANKA_0518300 | 7.3 | -2.17 | 6.43e-06 | 1.97E-05 |
| PBANKA_1441100 | 9.31 | -2.17 | 7.59e-06 | 2.30E-05 |
| PBANKA_1235200 | 7.29 | -2.17 | 6.86e-05 | 1.73E-04 |
| PBANKA_0211500 | 8.66 | -2.17 | 1.10e-04 | 2.67E-04 |
| PBANKA_1333600 | 6.83 | -2.17 | 3.74e-03 | 6.65E-03 |
| PBANKA_1206000 | 9.7 | -2.16 | 6.45e-09 | 2.95E-08 |
| PBANKA_0932000 | 10.21 | -2.16 | 1.01e-07 | 4.05E-07 |
| PBANKA_1407300 | 7.12 | -2.16 | 4.23e-07 | 1.55E-06 |
| PBANKA_1229400 | 6.87 | -2.16 | 1.72e-05 | 4.89E-05 |
| PBANKA_0204400 | 6.92 | -2.16 | 2.50e-04 | 5.71E-04 |
| PBANKA_0411300 | 7.94 | -2.16 | 4.70e-04 | 1.01E-03 |
| PBANKA_0509900 | 6.84 | -2.16 | 4.30e-03 | 7.59E-03 |
| PBANKA_0409800 | 8.99 | -2.15 | 1.31e-10 | 7.14E-10 |
| PBANKA_1343200 | 10.48 | -2.15 | 1.03e-08 | 4.62E-08 |
| PBANKA_1308400 | 8.73 | -2.15 | 4.53e-08 | 1.88E-07 |
| PBANKA_1342000 | 9.36 | -2.15 | 8.35e-07 | 2.95E-06 |
| PBANKA_1464700 | 6.84 | -2.15 | 3.76e-06 | 1.19E-05 |
| PBANKA_0512700 | 7.81 | -2.15 | 1.55e-05 | 4.42E-05 |
| PBANKA_1132700 | 7.31 | -2.15 | 1.03e-04 | 2.52E-04 |
| PBANKA_1124600 | 6.43 | -2.15 | 1.96e-04 | 4.57E-04 |
| PBANKA_1111200 | 6.08 | -2.15 | 7.95e-04 | 1.64E-03 |
| PBANKA_0933200 | 7.28 | -2.15 | 8.58e-04 | 1.76E-03 |
| PBANKA_1447200 | 8.07 | -2.14 | 2.56e-09 | 1.22E-08 |
| PBANKA_1038300 | 7.69 | -2.14 | 6.13e-09 | 2.82E-08 |
| PBANKA_0614800 | 7.92 | -2.14 | 7.72e-08 | 3.12E-07 |
| PBANKA_1033200 | 7.56 | -2.14 | 3.08e-07 | 1.16E-06 |
| PBANKA_1448800 | 8.97 | -2.14 | 4.77e-07 | 1.73E-06 |
| PBANKA_1115100 | 7.51 | -2.14 | 5.64e-05 | 1.45E-04 |
| PBANKA_1238400 | 6.61 | -2.14 | 1.34e-03 | 2.64E-03 |
| PBANKA_0112300 | 8.25 | -2.13 | 2.71e-09 | 1.29E-08 |
| PBANKA_0316000 | 9.56 | -2.13 | 2.87e-08 | 1.23E-07 |
| PBANKA_1456800 | 8.81 | -2.13 | 1.78e-05 | 5.02E-05 |
| PBANKA_0315900 | 7.68 | -2.13 | 1.33e-04 | 3.19E-04 |
| PBANKA_1457100 | 6.9 | -2.13 | 1.97e-04 | 4.58E-04 |
| PBANKA_1000800 | 7.22 | -2.13 | 1.09e-03 | 2.18E-03 |
| PBANKA_1208000 | 6.74 | -2.13 | 1.70e-03 | 3.26E-03 |
| PBANKA_0213700 | 6.79 | -2.13 | 2.81e-03 | 5.15E-03 |
| PBANKA_1225100 | 7.8 | -2.12 | 2.87e-09 | 1.36E-08 |
| PBANKA_1402600 | 9.13 | -2.12 | 1.00e-08 | 4.51E-08 |
| PBANKA_1302200 | 8.42 | -2.12 | 5.61e-08 | 2.31E-07 |
| PBANKA_1436100 | 7.84 | -2.12 | 1.91e-07 | 7.34E-07 |
| PBANKA_1436200 | 9.6 | -2.12 | 2.27e-07 | 8.63E-07 |

| | | | | |
|---|---|---|---|---|
| PBANKA_0716600 | 8 | -2.12 | 2.07e-05 | 5.78E-05 |
| PBANKA_0812800 | 8.23 | -2.12 | 2.28e-05 | 6.30E-05 |
| PBANKA_1326000 | 9.89 | -2.11 | 1.32e-09 | 6.47E-09 |
| PBANKA_1103500 | 7.75 | -2.11 | 5.82e-06 | 1.80E-05 |
| PBANKA_0913200 | 7.23 | -2.11 | 9.09e-06 | 2.72E-05 |
| PBANKA_0505200 | 8.61 | -2.11 | 1.08e-05 | 3.20E-05 |
| PBANKA_0832500 | 7.82 | -2.11 | 1.15e-05 | 3.38E-05 |
| PBANKA_0816700 | 7.35 | -2.11 | 1.36e-04 | 3.26E-04 |
| PBANKA_0523300 | 6.72 | -2.11 | 2.44e-03 | 4.54E-03 |
| PBANKA_1131600 | 9.26 | -2.1 | 8.30e-11 | 4.61E-10 |
| PBANKA_1406300 | 7.81 | -2.1 | 1.59e-09 | 7.75E-09 |
| PBANKA_1011100 | 8.89 | -2.1 | 8.19e-09 | 3.72E-08 |
| PBANKA_0807500 | 9.1 | -2.1 | 9.03e-07 | 3.16E-06 |
| PBANKA_0910100 | 7.04 | -2.1 | 4.60e-06 | 1.45E-05 |
| PBANKA_0417800 | 10.36 | -2.1 | 4.88e-06 | 1.53E-05 |
| PBANKA_1002100 | 9.99 | -2.1 | 9.21e-05 | 2.28E-04 |
| PBANKA_1141900 | 6.98 | -2.1 | 1.63e-03 | 3.15E-03 |
| PBANKA_1439600 | 8.07 | -2.09 | 8.24e-09 | 3.74E-08 |
| PBANKA_1118600 | 10.05 | -2.09 | 1.82e-05 | 5.11E-05 |
| PBANKA_1455100 | 8.47 | -2.09 | 2.41e-05 | 6.63E-05 |
| PBANKA_1358800 | 6.63 | -2.09 | 1.22e-03 | 2.42E-03 |
| PBANKA_0712100 | 9.69 | -2.08 | 4.81e-11 | 2.72E-10 |
| PBANKA_1444100 | 9.38 | -2.08 | 1.10e-08 | 4.91E-08 |
| PBANKA_0602700 | 9.83 | -2.08 | 5.67e-08 | 2.33E-07 |
| PBANKA_0102800 | 7.85 | -2.08 | 2.31e-07 | 8.76E-07 |
| PBANKA_0406400 | 8.73 | -2.08 | 2.42e-06 | 7.92E-06 |
| PBANKA_1450200 | 9.45 | -2.08 | 4.71e-06 | 1.48E-05 |
| PBANKA_1405100 | 7.73 | -2.08 | 6.16e-06 | 1.90E-05 |
| PBANKA_0311700 | 9.73 | -2.08 | 1.49e-05 | 4.28E-05 |
| PBANKA_1434000 | 7.26 | -2.08 | 1.56e-05 | 4.44E-05 |
| PBANKA_1438600 | 7.02 | -2.08 | 6.16e-05 | 1.58E-04 |
| PBANKA_0720100 | 8.54 | -2.08 | 5.33e-04 | 1.14E-03 |
| PBANKA_0622100 | 9.19 | -2.07 | 2.10e-13 | 1.45E-12 |
| PBANKA_0916600 | 8.33 | -2.07 | 8.98e-12 | 5.41E-11 |
| PBANKA_0404800 | 10.46 | -2.07 | 3.05e-10 | 1.60E-09 |
| PBANKA_0111900 | 8.08 | -2.07 | 1.19e-09 | 5.87E-09 |
| PBANKA_1133700 | 9.08 | -2.07 | 8.65e-08 | 3.48E-07 |
| PBANKA_0212200 | 9.56 | -2.07 | 2.20e-06 | 7.25E-06 |
| PBANKA_0411700 | 7.91 | -2.07 | 2.82e-06 | 9.13E-06 |
| PBANKA_1121200 | 7.31 | -2.07 | 8.51e-06 | 2.57E-05 |
| PBANKA_1326300 | 8.96 | -2.07 | 2.66e-05 | 7.28E-05 |
| PBANKA_1361600 | 7.46 | -2.07 | 2.22e-04 | 5.12E-04 |
| PBANKA_0415100 | 7.04 | -2.07 | 9.71e-04 | 1.97E-03 |
| PBANKA_1127300 | 6.33 | -2.07 | 1.91e-03 | 3.63E-03 |
| PBANKA_1316900 | 7.85 | -2.06 | 7.08e-11 | 3.97E-10 |
| PBANKA_0917900 | 8.73 | -2.06 | 3.24e-08 | 1.37E-07 |
| PBANKA_1032600 | 8.72 | -2.06 | 8.54e-08 | 3.44E-07 |
| PBANKA_1238100 | 7.48 | -2.06 | 1.73e-06 | 5.78E-06 |
| PBANKA_1341200 | 7.38 | -2.06 | 4.35e-05 | 1.14E-04 |
| PBANKA_1102300 | 7.22 | -2.06 | 2.11e-04 | 4.88E-04 |
| PBANKA_1363400 | 6.14 | -2.06 | 6.23e-04 | 1.30E-03 |
| PBANKA_0301700 | 6.41 | -2.06 | 2.45e-03 | 4.55E-03 |
| PBANKA_0104400 | 7.27 | -2.05 | 5.84e-09 | 2.69E-08 |
| PBANKA_0523800 | 7.44 | -2.05 | 1.78e-07 | 6.89E-07 |
| PBANKA_0715500 | 8.15 | -2.05 | 3.10e-07 | 1.16E-06 |
| PBANKA_1138300 | 7.24 | -2.05 | 1.22e-06 | 4.18E-06 |
| PBANKA_0307600 | 8.25 | -2.05 | 1.09e-05 | 3.23E-05 |
| PBANKA_1326800 | 7.65 | -2.05 | 5.66e-05 | 1.46E-04 |
| PBANKA_1315400 | 7.68 | -2.05 | 1.79e-04 | 4.19E-04 |
| PBANKA_1301400 | 8.09 | -2.05 | 1.98e-04 | 4.61E-04 |
| PBANKA_0609300 | 6.94 | -2.05 | 2.50e-03 | 4.63E-03 |
| PBANKA_0516800 | 6.87 | -2.05 | 2.78e-03 | 5.11E-03 |
| PBANKA_0302500 | 10.2 | -2.04 | 1.76e-06 | 5.84E-06 |
| PBANKA_0919000 | 8.08 | -2.04 | 5.26e-06 | 1.64E-05 |
| PBANKA_1223100 | 8.54 | -2.04 | 7.46e-05 | 1.88E-04 |
| PBANKA_0911600 | 6.54 | -2.04 | 1.58e-04 | 3.73E-04 |
| PBANKA_0202000 | 8.13 | -2.04 | 1.58e-04 | 3.74E-04 |
| PBANKA_1337700 | 7.36 | -2.04 | 3.46e-03 | 6.22E-03 |
| PBANKA_1359000 | 10.61 | -2.03 | 1.07e-09 | 5.31E-09 |
| PBANKA_0417300 | 8.27 | -2.03 | 2.28e-08 | 9.80E-08 |
| PBANKA_0300700 | 9.32 | -2.03 | 9.55e-08 | 3.83E-07 |
| PBANKA_0313200 | 7.46 | -2.03 | 1.16e-04 | 2.82E-04 |
| PBANKA_1450400 | 7.31 | -2.03 | 3.49e-04 | 7.74E-04 |
| PBANKA_1030100 | 7.6 | -2.03 | 4.19e-04 | 9.13E-04 |

| | | | | |
|---|---|---|---|---|
| PBANKA_1101800 | 6.61 | -2.03 | 4.92e-04 | 1.05E-03 |
| PBANKA_1406900 | 9.93 | -2.03 | 1.34e-03 | 2.64E-03 |
| PBANKA_0711100 | 7.6 | -2.03 | 1.63e-03 | 3.15E-03 |
| PBANKA_1329700 | 10.14 | -2.02 | 6.91e-13 | 4.61E-12 |
| PBANKA_0805800 | 6.47 | -2.02 | 2.68e-04 | 6.08E-04 |
| PBANKA_1117400 | 7.88 | -2.02 | 8.28e-04 | 1.70E-03 |
| PBANKA_0932600 | 8.22 | -2.02 | 1.20e-03 | 2.39E-03 |
| PBANKA_0415500 | 8.49 | -2.01 | 1.36e-09 | 6.69E-09 |
| PBANKA_1118200 | 8.62 | -2.01 | 1.06e-08 | 4.75E-08 |
| PBANKA_0501300 | 9.31 | -2.01 | 2.15e-06 | 7.08E-06 |
| PBANKA_0209200 | 7.97 | -2.01 | 3.09e-05 | 8.39E-05 |
| PBANKA_1039100 | 6.69 | -2.01 | 2.21e-04 | 5.09E-04 |
| PBANKA_1245500 | 8.08 | -2.01 | 2.96e-04 | 6.66E-04 |
| PBANKA_0806800 | 6.87 | -2.01 | 4.24e-03 | 7.49E-03 |
| PBANKA_0619700 | 8.63 | -2 | 1.68e-12 | 1.08E-11 |
| PBANKA_0906700 | 9.12 | -2 | 4.24e-12 | 2.63E-11 |
| PBANKA_0414000 | 8.14 | -2 | 1.39e-10 | 7.54E-10 |
| PBANKA_1349100 | 10.66 | -2 | 3.02e-07 | 1.14E-06 |
| PBANKA_0912100 | 9.49 | -2 | 5.92e-06 | 1.83E-05 |
| PBANKA_1224700 | 7.99 | -2 | 9.26e-06 | 2.76E-05 |
| PBANKA_0919900 | 8.57 | -2 | 2.09e-04 | 4.83E-04 |
| PBANKA_0818300 | 5.93 | -2 | 1.19e-03 | 2.36E-03 |
