## Supplementary material for "Transcriptome analysis of *Plasmodium berghei* during exo-erythrocytic development": Table S7

**Table S7:** **preferntially expressed in blood schizonts compared to exo-erythrocytic stages**

| | sporozoites | EEF_24h | EEF_48h | EEF_54h | EEF_60h | EEF_DC | EF_ring_4h | EF_trophozoite_16h | EF_schizont_22h | EF_gametocyte | ookinete_16h |
|---|---|---|---|---|---|---|---|---|---|---|---|
| PBANKA_0204500 | 5.708 | 6.277 | 5.588 | 5.836 | 5.825 | 2.485 | 7.302 | 5.386 | 8.159 | 11.772 | 10.036 |
| PBANKA_0301000 | 5.257 | 1.999 | 6.39 | 7.206 | 9.432 | 9.426 | 10.219 | 10.05 | 11.655 | 11.421 | 9.017 |
| PBANKA_0515000 | 10.534 | 6.226 | 5.585 | 5.233 | 5.309 | 3.341 | 10.914 | 6.545 | 10.105 | 15.895 | 15.001 |
| PBANKA_0518500 | 0 | 2.195 | 4.612 | 4.547 | 2.511 | 2.311 | 5.986 | 4.581 | 7.448 | 10.657 | 8.241 |
| PBANKA_0522000 | 6.176 | 7.235 | 6.334 | 7.261 | 7.704 | 6.341 | 7.728 | 7.839 | 9.879 | 9.113 | 8.859 |
| PBANKA_0601900 | 10.09 | 10.161 | 9.054 | 9.676 | 10.146 | 7.807 | 9.199 | 8.844 | 12.247 | 9.413 | 7.655 |
| PBANKA_0608600 | 6.222 | 2.476 | 4.444 | 1.991 | 3.942 | 5.421 | 9.849 | 6.342 | 8.466 | 14.814 | 13.842 |
| PBANKA_0612400 | 0 | 2.761 | 5.181 | 5.66 | 6.597 | 3.116 | 10.656 | 6.468 | 9.811 | 15.16 | 11.788 |
| PBANKA_0613300 | 2.997 | 5.83 | 7.181 | 7.384 | 7.451 | 6.604 | 7.563 | 8.258 | 9.821 | 8.359 | 8.194 |
| PBANKA_0700800 | 0 | 2.349 | 5.711 | 6.011 | 6.004 | 0 | 6.128 | 5.628 | 8.84 | 7.629 | 5.448 |
| PBANKA_0703500 | 6.347 | 2.195 | 5.338 | 5.082 | 6.481 | 3.448 | 9.212 | 6.304 | 8.611 | 12.9 | 8.592 |
| PBANKA_0714000 | 0 | 2.349 | 5.253 | 5.234 | 5.369 | 5.313 | 8.661 | 5.12 | 7.614 | 13.291 | 9.681 |
| PBANKA_0907900 | 5.46 | 1.999 | 4.726 | 5.41 | 6.208 | 6.026 | 9.7 | 6.204 | 8.92 | 13.769 | 9.676 |
| PBANKA_0915200 | 7.683 | 9.188 | 9.709 | 9.812 | 9.765 | 8.11 | 9.185 | 7.19 | 11.865 | 5.846 | 5.551 |
| PBANKA_1029400 | 0 | 1.877 | 4.379 | 4.276 | 6.653 | 6.307 | 9.25 | 5.167 | 9.64 | 12.367 | 8.875 |
| PBANKA_1103900 | 2.56 | 2.104 | 4.147 | 3.921 | 2.511 | 0 | 5.358 | 6.65 | 6.241 | 8.002 | 10.51 |
| PBANKA_1115000 | 0 | 1.877 | 4.036 | 4.835 | 5.341 | 2.311 | 9.125 | 4.976 | 8.071 | 13.835 | 11.375 |
| PBANKA_1138600 | 5.411 | 1.877 | 4.606 | 4.143 | 4.342 | 0 | 8.099 | 5.717 | 6.809 | 12.57 | 5.519 |
| PBANKA_1230100 | 12.043 | 10.155 | 9.958 | 10.445 | 10.488 | 9.491 | 11.064 | 10.637 | 12.496 | 10.762 | 11.227 |
| PBANKA_1236600 | 8.19 | 2.349 | 4.271 | 5.194 | 6.534 | 6.167 | 7.589 | 6.65 | 8.786 | 10.044 | 7.001 |
| PBANKA_1240000 | 10.031 | 9.063 | 9.122 | 9.036 | 8.86 | 7.32 | 8.811 | 9.768 | 11.364 | 9.038 | 7.902 |
| PBANKA_1315300 | 0 | 1.877 | 3.578 | 3.951 | 1.731 | 0 | 8.889 | 5.129 | 8.296 | 13.833 | 11.405 |
| PBANKA_1319500 | 0 | 2.416 | 4.713 | 4.304 | 5.151 | 0 | 9.693 | 5.393 | 8.908 | 14.682 | 11.2 |
| PBANKA_1334800 | 0 | 4.969 | 4.61 | 4.529 | 5.394 | 2.485 | 8.845 | 5.065 | 8.2 | 13.257 | 11.625 |
| PBANKA_1409600 | 5.08 | 2.476 | 6.285 | 5.849 | 6.104 | 3.116 | 8.774 | 5.6 | 8.732 | 11.393 | 7.044 |
| PBANKA_1443300 | 0 | 5.8 | 5.766 | 7.161 | 8.34 | 6.53 | 10.641 | 4.741 | 13.789 | 7.167 | 7.062 |
| PBANKA_1455800 | 0 | 1.877 | 0 | 3.203 | 1.63 | 0 | 5.809 | 7.824 | 6.283 | 9.071 | 4.853 |
| PBANKA_1461300 | 7.932 | 2.835 | 5.259 | 4.176 | 4.821 | 2.204 | 8.581 | 4.88 | 7.019 | 13.401 | 13.092 |
| PBANKA_1463000 | 2.52 | 2.584 | 5.227 | 5.246 | 6.434 | 3.178 | 8.23 | 8.39 | 10.518 | 11.777 | 13.413 |
| PBANKA_0201000 | 0 | 0 | 1.63 | 1.314 | 0 | 0 | 7.936 | 7.198 | 5.036 | 4.485 | 3.997 |
| PBANKA_0214550 | 0 | 0 | 3.314 | 4.359 | 3.488 | 0 | 6.813 | 7.89 | 6.72 | 5.289 | 0.867 |
| PBANKA_0216801 | 2.648 | 0 | 1.63 | 2.804 | 4.442 | 0 | 6.827 | 5.51 | 8.052 | 4.843 | 5.517 |
| PBANKA_0500781 | 0 | 0 | 1.264 | 1.808 | 4.345 | 0 | 7.937 | 7.658 | 9.991 | 6.024 | 3.039 |
| PBANKA_0504400 | 0 | 0 | 1.555 | 3.343 | 1.968 | 0 | 9.502 | 4.784 | 8.716 | 13.657 | 11.736 |
| PBANKA_0512000 | 0 | 0 | 3.682 | 1.841 | 1.731 | 0 | 6.369 | 2.854 | 5.462 | 10.224 | 7.815 |
| PBANKA_0514900 | 5.505 | 0 | 1.374 | 5.339 | 5.949 | 5.21 | 11.275 | 6.456 | 9.942 | 16.294 | 16.045 |
| PBANKA_0517600 | 0 | 0 | 3.338 | 3.157 | 3.728 | 0 | 8.175 | 4.025 | 6.868 | 12.955 | 9.157 |
| PBANKA_0606200 | 0 | 0 | 3.641 | 1.314 | 3.604 | 0 | 7.931 | 5.02 | 7.183 | 12.131 | 8.241 |
| PBANKA_0619200 | 2.326 | 0 | 3.847 | 5.411 | 6.456 | 5.313 | 8.051 | 4.728 | 8.339 | 11.212 | 16.514 |
| PBANKA_0623450 | 5.874 | 0 | 1.761 | 6.173 | 7.011 | 2.204 | 7.237 | 2.218 | 9.74 | 9.628 | 7.148 |
| PBANKA_0704700 | 0 | 0 | 4.793 | 4.44 | 4.961 | 0 | 6.669 | 3.291 | 7.261 | 10.454 | 8.246 |
| PBANKA_0704800 | 0 | 0 | 3.338 | 3.733 | 4.189 | 0 | 9.463 | 4.184 | 8.06 | 14.332 | 10.589 |
| PBANKA_0707100 | 0 | 0 | 3.614 | 0 | 1.897 | 0 | 9.288 | 4.706 | 8.44 | 13.989 | 14.839 |
| PBANKA_0719100 | 0 | 0 | 1.264 | 3.446 | 4.244 | 0 | 4.905 | 3.339 | 6.454 | 9.293 | 9.544 |
| PBANKA_0804300 | 0 | 0 | 1.47 | 1.705 | 0 | 0 | 5.374 | 4.517 | 5.588 | 9.009 | 5.144 |
| PBANKA_0812600 | 0 | 0 | 3.641 | 4.003 | 4.483 | 0 | 7.536 | 3.122 | 6.733 | 12.006 | 5.362 |
| PBANKA_0823200 | 9.61 | 0 | 3.727 | 1.841 | 3.995 | 0 | 7.412 | 3.795 | 6.112 | 11.971 | 9.441 |
| PBANKA_0927600 | 2.326 | 0 | 4.94 | 5.049 | 5.627 | 5.802 | 9.622 | 6.151 | 9.003 | 14.485 | 9.952 |
| PBANKA_1030400 | 0 | 0 | 1.264 | 1.753 | 4.822 | 2.743 | 6.796 | 5.1 | 6.935 | 9.935 | 3.882 |
| PBANKA_1035200 | 0 | 0 | 3.727 | 3.753 | 5.562 | 2.974 | 6.744 | 4.15 | 7.903 | 10.934 | 11.529 |
| PBANKA_1038800 | 0 | 0 | 3.973 | 3.653 | 2.203 | 0 | 6.009 | 2.36 | 7.408 | 10.597 | 8.582 |
| PBANKA_1109600 | 5.813 | 0 | 4.274 | 3.799 | 4.147 | 0 | 6.485 | 4.374 | 6.852 | 9.954 | 8.432 |
| PBANKA_1129600 | 0 | 0 | 2.053 | 1.596 | 5.352 | 2.079 | 8.294 | 4.621 | 8.164 | 10.667 | 7.387 |
| PBANKA_1204200 | 11.651 | 0 | 4.6 | 3.712 | 4.961 | 2.311 | 9.152 | 5.537 | 7.868 | 13.432 | 11.882 |
| PBANKA_1312700 | 15.376 | 0 | 1.555 | 4.175 | 3.871 | 0 | 6.121 | 2.777 | 7.594 | 9.498 | 12.884 |
| PBANKA_1326100 | 5.635 | 0 | 4.207 | 3.962 | 1.514 | 0 | 6.78 | 4.584 | 6.469 | 9.319 | 6.238 |
| PBANKA_1333700 | 2.452 | 0 | 1.374 | 3.966 | 4.895 | 2.89 | 8.433 | 4.945 | 7.533 | 12.887 | 10.615 |
| PBANKA_1334900 | 0 | 0 | 4.461 | 3.935 | 4.349 | 2.204 | 8.643 | 6.138 | 8.054 | 13.111 | 11.861 |
| PBANKA_1340400 | 0 | 0 | 4.621 | 3.529 | 4.219 | 2.403 | 5.7 | 4.18 | 7.06 | 8.606 | 4.922 |
| PBANKA_1414500 | 5.459 | 0 | 3.727 | 3.69 | 0 | 0 | 7.305 | 3.183 | 6.002 | 11.62 | 8.144 |
| PBANKA_1436600 | 0 | 0 | 3.023 | 3.476 | 4.773 | 2.845 | 9.403 | 4.685 | 8.184 | 14.165 | 8.398 |
| PBANKA_1449000 | 5.459 | 0 | 4.202 | 4.448 | 5.312 | 2.933 | 6.979 | 4.743 | 7.946 | 10.646 | 7.731 |
| PBANKA_1342300 | 0 | 0 | 1.759 | 3.343 | 0 | 0 | 6.894 | 2.89 | 6.265 | 11.329 | 8.598 |
| PBANKA_0605800 | 6.863 | 0 | 0 | 1.396 | 0 | 0 | 5.914 | 2.533 | 4.235 | 10.019 | 8.218 |
| PBANKA_1334700 | 0 | 0 | 0 | 3.394 | 4.785 | 0 | 7.119 | 3.359 | 6.721 | 12.031 | 10.343 |
| PBANKA_1419300 | 0 | 0 | 0 | 1.117 | 1.63 | 0 | 8.545 | 4.418 | 7.692 | 13.561 | 11.471 |
| PBANKA_1432200 | 5.568 | 0 | 0 | 1.535 | 0 | 0 | 7.834 | 3.336 | 9.114 | 11.784 | 11.638 |
| PBANKA_1436300 | 0 | 0 | 0 | 1.222 | 0 | 0 | 6.241 | 3.546 | 5.529 | 10.254 | 5.392 |
| PBANKA_0109800 | 7.463 | 0 | 0 | 0 | 1.819 | 0 | 6.843 | 2.574 | 6.117 | 11.57 | 7.52 |
| PBANKA_0105100 | 0 | 0 | 0 | 0 | 0 | 0 | 6.255 | 2.402 | 4.997 | 10.488 | 6.514 |
| PBANKA_0110800 | 0 | 0 | 0 | 0 | 0 | 0 | 6.476 | 3.053 | 5.589 | 11.229 | 8.525 |
| PBANKA_1000081 | 0 | 0 | 0 | 0 | 0 | 0 | 6.226 | 6.47 | 8.365 | 4.607 | 0.52 |
| PBANKA_0500721 | 0 | 0 | 0 | 0 | 0 | 0 | 6.724 | 6.9 | 4.688 | 4.204 | 0 |
