## Supplementary material for "Transcriptome analysis of *Plasmodium berghei* during exo-erythrocytic development": Table S8

Table S8: **preferentially expressed in blood schizonts compared to all other stages**

| | sporozoites | EEF_24h | EEF_48h | EEF_54h | EEF_60h | EEF_DC | EF_ring_4h | EF_trophozoite_16h | EF_schizont_22h | EF_gametocyte | ookinete_16h |
|---|---|---|---|---|---|---|---|---|---|---|---|
| PBANKA_0601900 | 10.09 | 10.161 | 9.054 | 9.676 | 10.146 | 7.807 | 9.199 | 8.844 | 12.247 | 9.413 | 7.655 |
| PBANKA_0915200 | 7.683 | 9.188 | 9.709 | 9.812 | 9.765 | 8.11 | 9.185 | 7.19 | 11.865 | 5.846 | 5.551 |
| PBANKA_1443300 | 0 | 5.8 | 5.766 | 7.161 | 8.34 | 6.53 | 10.641 | 4.741 | 13.789 | 7.167 | 7.062 |
| PBANKA_0500781 | 0 | 0 | 1.264 | 1.808 | 4.345 | 0 | 7.937 | 7.658 | 9.991 | 6.024 | 3.039 |
