## Supplementary material for "Transcriptome analysis of *Plasmodium berghei* during exo-erythrocytic development": Table S9

**Table S9:** **preferentially expressed in DC compared to all other stages**

| | sporozoites | EEF_24h | EEF_48h | EEF_54h | EEF_60h | EEF_DC | EF_ring_4h | EF_trophozoite_16h | EF_schizont_22h | EF_gametocyte | ookinete_16h |
|---|---|---|---|---|---|---|---|---|---|---|---|
| PBANKA_0100061 | 6.517100741 | 1.851885259 | 5.470715858 | 4.900525057 | 7.637799705 | 10.92688507 | 5.874227997 | 4.336515496 | 5.16521768 | 5.96391331 | 3.680888064 |
| PBANKA_0100700 | 0 | 2.389790243 | 6.918383365 | 7.684353643 | 11.71638361 | 15.96120371 | 11.10594067 | 9.895084454 | 10.0179693 | 7.953405894 | 5.530069025 |
| PBANKA_0100900 | 9.625227286 | 9.93783861 | 9.652052186 | 9.168763239 | 9.687322553 | 14.32832341 | 9.751820454 | 8.786708282 | 7.751988269 | 8.940175315 | 8.328944442 |
| PBANKA_0104800 | 6.220711687 | 5.91404089 | 6.211095803 | 6.843336285 | 8.441961 | 12.38107467 | 9.404743721 | 7.913228741 | 8.878139207 | 8.165529683 | 7.003662609 |
| PBANKA_0107700 | 8.240503367 | 8.202507772 | 7.073712515 | 6.118614083 | 7.617917783 | 12.31850266 | 8.83436216 | 8.008735342 | 5.953352158 | 8.449961367 | 9.972011491 |
| PBANKA_0111200 | 9.661418478 | 10.03535969 | 8.735422629 | 8.394188688 | 9.605159143 | 13.51423379 | 10.25091347 | 8.957174953 | 7.818744877 | 8.890831825 | 10.55517665 |
| PBANKA_0112600 | 0 | 7.084074368 | 7.786243955 | 8.036089567 | 8.378474287 | 12.64734904 | 9.902883473 | 9.566927571 | 8.417885714 | 6.367072475 | 4.399067186 |
| PBANKA_0200400 | 7.547879729 | 5.053301393 | 5.658869191 | 5.901224657 | 8.533052274 | 13.0575731 | 8.49726095 | 6.414862275 | 7.566157774 | 7.462961009 | 4.36717631 |
| PBANKA_0200600 | 6.1351972 | 2.506242444 | 6.359920328 | 5.829341655 | 5.930386478 | 9.913435896 | 7.603658021 | 6.807180252 | 6.468375981 | 7.767760029 | 6.685104829 |
| PBANKA_0201500 | 0 | 2.558102 | 5.186533708 | 4.960511368 | 7.77582935 | 13.02906134 | 9.659799275 | 9.143351155 | 8.885347606 | 5.795459264 | 4.290153182 |
| PBANKA_0201600 | 0 | 6.185672575 | 7.895872102 | 9.780805093 | 11.95939 | 15.00107798 | 11.51263734 | 11.5948513 | 12.44361712 | 9.23537141 | 9.217493699 |
| PBANKA_0203200 | 0 | 7.454737487 | 7.528608653 | 7.730523704 | 8.11064332 | 11.76056177 | 9.183779705 | 8.77731438 | 8.863245069 | 8.293706858 | 6.772807897 |
| PBANKA_0205000 | 7.22932316 | 9.620537013 | 7.733211878 | 7.328527367 | 9.893408114 | 14.97170974 | 11.65188815 | 8.085723302 | 8.599155414 | 9.045172413 | 8.586121606 |
| PBANKA_0205100 | 0 | 7.426598641 | 8.36270722 | 8.238051234 | 8.035203574 | 10.46011793 | 7.35414808 | 5.875881261 | 5.841443577 | 6.611287001 | 6.667626925 |
| PBANKA_0206600 | 6.546319144 | 8.768928274 | 8.27331144 | 7.79383144 | 9.09518312 | 12.88245742 | 10.24573909 | 9.088710156 | 8.556770441 | 7.697506178 | 8.669341918 |
| PBANKA_0206900 | 0 | 6.99844062 | 6.141509892 | 4.753476596 | 3.998671331 | 9.227354035 | 6.254628434 | 5.859718126 | 2.853586414 | 4.838565286 | 4.143197131 |
| PBANKA_0211200 | 7.890405584 | 3.226564082 | 6.60636664 | 6.748194431 | 7.076849466 | 9.892372211 | 7.044870699 | 7.212157414 | 6.942670518 | 7.332958372 | 7.847669043 |
| PBANKA_0213400 | 10.41323238 | 8.988875479 | 8.328395516 | 7.883697271 | 8.295166014 | 12.8246153 | 9.865885325 | 8.423648938 | 7.101737722 | 6.92499002 | 10.29988049 |
| PBANKA_0214800 | 4.919194461 | 7.85639295 | 8.083344668 | 8.084423809 | 8.252212913 | 12.99877566 | 10.01992068 | 7.676436599 | 8.108949952 | 9.928432306 | 6.097858343 |
| PBANKA_0301500 | 6.545922095 | 8.253465546 | 6.612029198 | 6.053401362 | 7.827346232 | 12.68140204 | 9.090211568 | 7.536322227 | 6.10476557 | 6.653138175 | 8.264522679 |
| PBANKA_0304300 | 6.9659839 | 7.34040968 | 7.561790813 | 6.535108909 | 6.934109283 | 12.13503751 | 7.00133805 | 6.843610979 | 4.623838181 | 5.302245783 | 8.083129506 |
| PBANKA_0305300 | 9.38646609 | 8.420184084 | 6.916900715 | 6.560951038 | 7.436572444 | 11.94226303 | 8.220928653 | 6.795042716 | 5.245622943 | 6.743029849 | 8.540362329 |
| PBANKA_0311200 | 8.894487082 | 8.135329939 | 7.325218772 | 7.070114047 | 7.419616884 | 12.51293538 | 9.875353861 | 8.1689377 | 6.829344628 | 7.78578785 | 8.406931714 |
| PBANKA_0311800 | 12.2771616 | 9.696187424 | 9.666506047 | 8.918305877 | 11.50521627 | 15.55320036 | 11.55296473 | 8.578308139 | 10.26028891 | 8.367926055 | 6.214745747 |
| PBANKA_0316100 | 7.252810113 | 5.580260138 | 7.039245535 | 7.607497268 | 8.929659906 | 11.38443649 | 8.886901548 | 8.325018337 | 8.578979655 | 7.334737633 | 5.087674091 |
| PBANKA_0316200 | 9.180036954 | 6.958121608 | 8.160807685 | 9.061737895 | 13.4969079 | 18.86671213 | 13.25093941 | 12.73976097 | 10.5576245 | 11.50566579 | 10.44296605 |
| PBANKA_0317141 | 6.182159528 | 2.450363758 | 5.608021724 | 5.31018897 | 4.871957384 | 9.047396349 | 5.970908597 | 5.204473807 | 5.454509247 | 6.425469248 | 4.84069675 |
| PBANKA_0404000 | 9.63888799 | 9.349595312 | 9.580173771 | 9.761549166 | 10.40174243 | 13.16134754 | 10.03045047 | 10.32964437 | 9.737117664 | 9.525262632 | 8.647253321 |
| PBANKA_0414700 | 9.065714328 | 8.547530361 | 7.547191206 | 7.376382577 | 9.193937576 | 13.61807156 | 10.10750214 | 8.62584427 | 8.069765113 | 8.875469438 | 10.97136848 |
| PBANKA_0416200 | 9.565355056 | 8.664603757 | 8.444117833 | 8.09302093 | 8.627509343 | 11.76517287 | 9.354706306 | 8.940500426 | 7.815462716 | 8.570857429 | 9.572657008 |
| PBANKA_0506300 | 6.919534815 | 6.724496251 | 5.757842575 | 5.673956762 | 6.269375562 | 10.13505436 | 7.260483954 | 6.450993361 | 5.204537942 | 7.291012032 | 7.663539638 |
| PBANKA_0508100 | 12.25023991 | 11.29351861 | 11.20486139 | 11.15480096 | 12.42352069 | 15.59205239 | 12.26634985 | 12.19079866 | 12.43336404 | 11.18241452 | 11.5337853 |
| PBANKA_0512400 | 7.218580602 | 8.1136934 | 7.17156008 | 6.280389865 | 7.36154663 | 12.33393362 | 9.289632913 | 7.522734741 | 6.217435976 | 8.678509993 | 9.702704268 |
| PBANKA_0514000 | 2.626607756 | 6.652710495 | 6.405821407 | 5.958098005 | 6.039244787 | 10.07598947 | 5.792509037 | 5.636536012 | 5.536619748 | 6.112377459 | 6.625764534 |
| PBANKA_0514100 | 6.749634803 | 8.904019231 | 8.981450777 | 8.910846981 | 9.310667021 | 12.77687613 | 10.27407141 | 9.612616574 | 9.003333127 | 8.559115196 | 9.950211295 |
| PBANKA_0517000 | 7.253041525 | 8.740899171 | 8.293740157 | 7.859538446 | 11.66502319 | 15.69067071 | 12.115417 | 10.98707954 | 10.03392073 | 9.823149991 | 8.481166721 |
| PBANKA_0517500 | 7.275556388 | 9.424555864 | 8.03966255 | 7.48913148 | 8.404396807 | 12.85519051 | 10.28163785 | 8.203002203 | 8.433666707 | 8.859451486 | 7.913785444 |
| PBANKA_0519900 | 8.770607937 | 10.89728115 | 10.26736617 | 10.08314044 | 9.99998495 | 13.85030812 | 11.4244413 | 10.78233672 | 10.10987609 | 10.07072692 | 11.15935459 |
| PBANKA_0521300 | 10.26503525 | 9.270762649 | 9.163517346 | 9.580814229 | 10.78904259 | 14.28967118 | 10.60577137 | 10.60847626 | 10.09372541 | 9.079710602 | 8.600373271 |
| PBANKA_0524800 | 0 | 5.053301393 | 5.908505125 | 6.597336517 | 8.178598392 | 12.08341035 | 9.818335588 | 9.701179838 | 9.543129557 | 7.460488567 | 5.678293423 |
| PBANKA_0601700 | 8.809387482 | 8.109190062 | 7.428286844 | 6.869696236 | 7.064801907 | 11.56965749 | 8.793132227 | 7.523087938 | 5.530367003 | 6.562495964 | 8.832567007 |
| PBANKA_0604200 | 8.130215578 | 7.910241616 | 8.548793558 | 9.31903231 | 10.64695461 | 13.27151082 | 9.950186838 | 9.747494534 | 9.967391145 | 10.12062464 | 10.24595672 |
| PBANKA_0604500 | 12.1465093 | 10.93802898 | 10.80621039 | 10.93262847 | 11.45787533 | 14.66412237 | 12.09526001 | 11.50341859 | 12.18926408 | 9.6530557 | 12.52513327 |
| PBANKA_0605700 | 7.811069914 | 8.958086236 | 7.964783501 | 7.257091736 | 8.440158134 | 12.93021654 | 10.81144687 | 10.05890465 | 9.264054155 | 9.764761837 | 9.574407302 |
| PBANKA_0609100 | 10.11224125 | 8.223469027 | 8.024628959 | 7.752541373 | 9.3982361 | 13.32214642 | 9.916149253 | 8.498788235 | 8.259527022 | 8.570023911 | 10.88890749 |
| PBANKA_0610500 | 5.248462499 | 7.24814005 | 7.126133069 | 7.836483902 | 8.736881063 | 11.47389221 | 7.189204134 | 7.267839945 | 7.995047742 | 6.873362707 | 6.726121806 |
| PBANKA_0613700 | 0 | 9.512597396 | 8.889862135 | 8.595039981 | 9.05876951 | 13.92145362 | 11.2703793 | 10.74220468 | 9.574278231 | 11.0566475 | 8.785468735 |
| PBANKA_0613800 | 10.21861499 | 9.080907369 | 8.407605493 | 7.726609526 | 9.435113779 | 12.91814802 | 10.23230545 | 9.555132318 | 8.420801049 | 7.63484508 | 7.278426449 |
| PBANKA_0613900 | 9.022578365 | 8.606997854 | 7.621039869 | 6.973032007 | 8.692253307 | 12.16364281 | 8.618964569 | 6.870722726 | 6.14806317 | 5.925318403 | 7.34368629 |
| PBANKA_0615100 | 7.191722124 | 8.774559697 | 7.578067177 | 7.02607876 | 7.699248275 | 11.6417022 | 8.843119505 | 7.838106729 | 7.161154075 | 8.210332279 | 8.183042493 |
| PBANKA_0617600 | 7.590079858 | 7.719134577 | 8.319699417 | 8.390237577 | 8.317309684 | 11.82597599 | 7.573372352 | 7.286426329 | 7.419746004 | 7.656946345 | 9.806724134 |
| PBANKA_0617700 | 6.916651639 | 6.588861902 | 5.974590457 | 6.629591089 | 7.363104024 | 9.812841128 | 5.308278998 | 5.149174658 | 5.451358824 | 5.290452254 | 7.835068114 |
| PBANKA_0620000 | 5.202116977 | 6.752685386 | 6.606958066 | 5.940082029 | 6.500788367 | 10.25242479 | 7.696628278 | 7.334433874 | 6.916567994 | 6.311063888 | 8.133299637 |
| PBANKA_0621400 | 6.676157114 | 8.833979622 | 9.058818412 | 8.99696988 | 9.179795524 | 12.77993678 | 10.05410344 | 9.418364728 | 9.390570385 | 8.388561748 | 8.80817441 |
| PBANKA_0623000 | 11.65690693 | 10.18409246 | 8.429363477 | 7.816249474 | 10.82020725 | 14.55695718 | 9.459431515 | 7.462699546 | 8.159997162 | 6.643702118 | 3.9511867 |
| PBANKA_0623200 | 2.492651273 | 6.111388554 | 4.972737433 | 5.411167942 | 8.913846051 | 14.08523234 | 10.7692523 | 9.207564122 | 7.841838622 | 8.408629854 | 5.487975204 |
| PBANKA_0623300 | 0 | 8.420387078 | 7.29081287 | 6.737456055 | 9.015762962 | 14.36159713 | 11.47579239 | 9.260641818 | 8.41540928 | 8.205977223 | 5.463003918 |
| PBANKA_0623500 | 0 | 1.974116226 | 6.071065794 | 5.593458822 | 8.445246365 | 12.73561183 | 8.56929384 | 7.897964234 | 7.542633762 | 6.617055824 | 6.082083223 |
| PBANKA_0700700 | 0 | 6.932583032 | 6.092500483 | 6.151887584 | 7.116003524 | 12.85434903 | 9.654507796 | 7.961568985 | 6.788224998 | 7.004372307 | 5.270436639 |
| PBANKA_0700900 | 0 | 1.851885259 | 5.133305121 | 5.726000722 | 7.87477997 | 10.99144289 | 7.096637197 | 6.345452579 | 6.624876088 | 5.420403983 | 1.614512163 |
| PBANKA_0701000 | 0 | 2.450363758 | 5.625773966 | 5.272392028 | 8.928513276 | 13.53103358 | 10.38227657 | 8.156246664 | 7.531337401 | 6.086189536 | 4.569286487 |
| PBANKA_0702900 | 2.492651273 | 8.385030845 | 9.928592954 | 11.29800681 | 11.35428295 | 13.65657551 | 10.2425824 | 9.468633472 | 11.37406428 | 9.373893788 | 8.045528298 |
| PBANKA_0707300 | 9.205258829 | 10.15450978 | 10.10318321 | 10.43895744 | 10.43755427 | 13.56274835 | 10.97557687 | 10.54782856 | 11.20077421 | 9.494699743 | 9.821548154 |
| PBANKA_0707400 | 7.684804965 | 9.942331074 | 9.053557871 | 8.903961892 | 9.307969621 | 13.6548469 | 8.873260393 | 8.600034005 | 7.966426473 | 8.622695278 | 7.880085002 |
| PBANKA_0714700 | 0 | 5.351752105 | 6.119009402 | 5.356917111 | 5.125158324 | 9.251130789 | 4.698632661 | 5.527893661 | 3.998785169 | 2.703831147 | 3.413887749 |
| PBANKA_0720800 | 8.104402492 | 8.089467049 | 7.798133659 | 6.700556406 | 8.620610945 | 13.07440969 | 8.768098448 | 7.573673667 | 6.692270357 | 7.591006799 | 7.908333883 |
| PBANKA_0804800 | 7.199316614 | 6.918200645 | 6.677159314 | 6.293146599 | 5.73895345 | 9.266748384 | 7.194356578 | 6.782833592 | 6.394951277 | 5.296722147 | 5.972285965 |
| PBANKA_0805100 | 8.998827413 | 8.934598908 | 8.220986746 | 7.861068813 | 8.295807535 | 11.68222261 | 9.285116626 | 8.289122905 | 7.393588337 | 7.087549737 | 9.212631993 |
| PBANKA_0805400 | 5.845917145 | 6.690668014 | 5.987426434 | 5.467783645 | 5.947700402 | 9.179707268 | 5.858238955 | 6.220965754 | 5.875238518 | 5.3905572 | 5.899970478 |
| PBANKA_0806000 | 0 | 3.57970646 | 6.102868259 | 5.467783645 | 6.578090595 | 10.65445117 | 6.872217372 | 6.045586829 | 4.527958872 | 6.290298799 | 5.681624073 |
| PBANKA_0808700 | 8.512026919 | 6.923322129 | 7.68826508 | 7.338843852 | 8.142596801 | 12.80379534 | 8.823278587 | 9.150240766 | 7.739732912 | 8.606081212 | 8.785452468 |
| PBANKA_0810500 | 5.439438827 | 8.340628256 | 7.482740324 | 6.901312409 | 6.45051366 | 10.68458774 | 7.775735874 | 6.880424453 | 5.903979392 | 7.663210715 | 6.889121234 |
| PBANKA_0810600 | 7.669054103 | 10.19364021 | 10.26987307 | 10.15692709 | 9.97336034 | 13.37337802 | 10.05410344 | 9.498330814 | 9.379460998 | 8.73623221 | 8.392508788 |
| PBANKA_0811100 | 7.04141429 | 8.678687574 | 8.533978448 | 7.758652596 | 7.402592214 | 11.12799393 | 8.404726773 | 8.271310329 | 7.100913789 | 6.918575046 | 6.813702531 |
| PBANKA_0814000 | 2.298831494 | 7.727106394 | 7.031690767 | 6.550534368 | 6.777660513 | 11.81112797 | 7.882208081 | 6.592710572 | 5.778293484 | 6.60970363 | 7.318712486 |
| PBANKA_0817900 | 2.620362967 | 7.426706822 | 7.954000397 | 8.593968042 | 9.594783894 | 12.66084164 | 7.424035952 | 5.78706407 | 8.294436669 | 5.37145385 | 8.45594345 |
| PBANKA_0818700 | 0 | 5.480938389 | 5.806363381 | 5.44894647 | 4.798411786 | 7.805032773 | 5.096022419 | 5.395285189 | 4.241088339 | 4.232654865 | 3.948462099 |
| PBANKA_0819900 | 8.741325041 | 11.58285211 | 10.20747424 | 9.79176883 | 9.383897483 | 13.53085927 | 10.31653088 | 10.22363266 | 9.5427889 | 9.774095156 | 7.499588687 |
| PBANKA_0822700 | 0 | 7.129040558 | 6.677014259 | 5.993622382 | 6.803291514 | 10.72526737 | 7.812153516 | 6.715028072 | 5.578308288 | 8.037704692 | 8.043557858 |
| PBANKA_0824800 | 8.527973093 | 8.866666602 | 8.022532636 | 7.582337314 | 8.179277148 | 11.1278532 | 7.998748601 | 7.707985697 | 6.434656895 | 7.433399927 | 7.524415994 |
| PBANKA_0826900 | 8.999621131 | 9.916243417 | 10.86693099 | 10.62523864 | 10.42786738 | 13.88420317 | 10.67957131 | 10.54943322 | 9.809106776 | 9.651967773 | 9.391480029 |
| PBANKA_0903000 | 7.700718991 | 9.097317695 | 8.0117933 | 7.346890041 | 7.525749816 | 12.48312008 | 9.109257839 | 8.187650552 | 6.656465681 | 8.838081091 | 6.227252511 |
| PBANKA_0908300 | 8.46990228 | 6.900857112 | 9.489857107 | 10.71074431 | 11.07701095 | 13.7573768 | 9.377682807 | 9.387353235 | 8.425874714 | 8.861825955 | 9.242440409 |
| PBANKA_0912000 | 6.929290455 | 9.556237227 | 8.462106719 | 8.038204025 | 8.791046528 | 13.18795239 | 10.61776872 | 9.62142197 | 7.786263392 | 9.052604675 | 9.519551077 |
| PBANKA_0913400 | 0 | 7.054579281 | 7.939605636 | 7.916670636 | 11.48035248 | 15.17954811 | 10.22761522 | 10.18960005 | 9.322162391 | 7.726493727 | 7.484348087 |
| PBANKA_0914200 | 9.758247863 | 9.426733115 | 8.827854403 | 8.535993611 | 9.674694085 | 14.13596486 | 10.35260693 | 9.052036243 | 8.166292667 | 8.43612212 | 8.88616942 |
| PBANKA_0915300 | 8.610471846 | 9.069708312 | 8.069662825 | 7.493137534 | 8.96116574 | 12.77992486 | 9.722564853 | 9.038525687 | 7.846390798 | 7.985458805 | 9.280260803 |
| PBANKA_0919100 | 7.457936585 | 9.132874755 | 9.189137965 | 9.667838625 | 11.23771886 | 14.71392301 | 11.3573033 | 10.16849679 | 11.27608374 | 10.70348514 | 8.178830221 |
| PBANKA_0919200 | 0 | 6.427822222 | 7.229084158 | 7.552103635 | 7.837291739 | 11.46157894 | 8.328206044 | 7.952949567 | 7.801835805 | 7.390790024 | 4.627707434 |
| PBANKA_0923500 | 7.398758186 | 8.431921828 | 7.157732337 | 6.821242381 | 7.66353661 | 12.01244146 | 9.418671997 | 7.547125096 | 6.867408824 | 7.702389058 | 7.949765779 |
| PBANKA_0924400 | 8.202302981 | 9.263571152 | 8.187439679 | 7.604806871 | 7.963384794 | 12.9132469 | 7.918404935 | 7.029603096 | 5.534119119 | 5.060903345 | 7.409194845 |
| PBANKA_0926700 | 6.886384173 | 11.75897952 | 11.10463675 | 10.90778156 | 12.79924146 | 15.38426768 | 10.47557436 | 10.66137168 | 10.8014267 | 8.481305081 | 6.684122553 |
| PBANKA_0927200 | 7.131095166 | 8.906004924 | 8.642529082 | 8.749358737 | 9.32286084 | 12.65199944 | 9.945931723 | 9.40737038 | 9.49159096 | 8.514061306 | 7.488975982 |
| PBANKA_0929600 | 9.3227259 | 10.13259862 | 9.508325207 | 8.977967165 | 9.042078448 | 13.08581973 | 10.31317212 | 9.229925899 | 7.647375826 | 8.480289564 | 9.777350385 |

| | | | | | | | | | | | |
|---|---|---|---|---|---|---|---|---|---|---|---|
| PBANKA_0934500 | 9.499887619 | 7.724312166 | 7.314200473 | 7.077399248 | 8.874696482 | 11.52119151 | 9.363724513 | 9.010399258 | 9.237713715 | 9.358574934 | 8.58722843 |
| PBANKA_0937800 | 8.53996909 | 9.040320676 | 7.608535099 | 7.189390868 | 7.568336161 | 12.57018394 | 9.670303718 | 7.546475732 | 6.822488478 | 8.12322005 | 9.989238999 |
| PBANKA_0939100 | 11.94886357 | 9.817072046 | 9.105762418 | 9.01327088 | 11.80416965 | 14.0933249 | 11.51481939 | 9.741610767 | 11.76420997 | 10.77294534 | 12.08452029 |
| PBANKA_0939200 | 3.071172369 | 8.35639337 | 7.5814776 | 6.908039677 | 7.718327737 | 12.23852167 | 9.57860436 | 8.480182533 | 6.478526296 | 7.317806332 | 8.474745421 |
| PBANKA_0939600 | 9.416603598 | 11.74758055 | 11.08830947 | 11.06795964 | 11.16357303 | 15.3099569 | 12.60039818 | 12.25266415 | 12.01683422 | 11.10240554 | 9.883643964 |
| PBANKA_0943500 | 7.444448704 | 8.80869245 | 8.457762972 | 7.532471445 | 7.189184358 | 11.11650809 | 8.229186403 | 8.430164254 | 7.194901428 | 6.298164454 | 3.807106989 |
| PBANKA_1002700 | 9.398446918 | 7.955474156 | 6.948951061 | 6.045416094 | 7.004447404 | 12.11135926 | 8.18086897 | 6.532568032 | 6.149262462 | 5.889464966 | 7.4610977 |
| PBANKA_1003200 | 5.875973058 | 7.672501528 | 6.591393192 | 6.512260152 | 6.10805022 | 10.84679279 | 6.3446365 | 5.57247732 | 5.285166571 | 5.785397218 | 6.576206783 |
| PBANKA_1003300 | 0 | 6.633094434 | 7.529440242 | 7.874640062 | 7.728423815 | 10.59144991 | 7.643630335 | 6.791170465 | 7.152152865 | 6.596023026 | 5.464859656 |
| PBANKA_1003700 | 9.87174954 | 7.900452838 | 7.607737223 | 8.133014208 | 9.140046574 | 12.11216697 | 9.737855274 | 8.478098664 | 9.989108768 | 9.345983794 | 9.60701465 |
| PBANKA_1006700 | 9.283753192 | 8.491038392 | 7.584358587 | 7.720629829 | 8.103399581 | 11.71665028 | 8.78174887 | 8.226884125 | 7.91788228 | 7.464686459 | 8.606899204 |
| PBANKA_1006800 | 9.278276604 | 7.798927593 | 6.746817662 | 6.625541376 | 7.755486267 | 11.90876455 | 8.660356094 | 7.213806622 | 6.58471573 | 6.252759602 | 8.207294936 |
| PBANKA_1007400 | 9.539963121 | 7.877787283 | 7.351315045 | 7.406393053 | 8.931658132 | 12.1619526 | 9.897586872 | 8.789654504 | 9.09452012 | 8.275922983 | 9.301050694 |
| PBANKA_1007500 | 7.913523091 | 6.091615376 | 6.126315976 | 5.806393914 | 7.366425999 | 10.09158881 | 7.325227451 | 7.032863483 | 7.266047617 | 7.869028868 | 7.515618687 |
| PBANKA_1008500 | 6.099637605 | 8.910783444 | 7.654826166 | 7.810641988 | 12.73092986 | 15.64863088 | 12.68443415 | 11.63982016 | 11.34654778 | 9.824122822 | 9.52041368 |
| PBANKA_1010300 | 10.97487827 | 11.79131836 | 10.92238965 | 10.47376323 | 10.84040037 | 14.16339917 | 10.64415361 | 10.51580339 | 9.00320006 | 9.78854181 | 11.05968657 |
| PBANKA_1012400 | 6.574373864 | 6.22080166 | 6.607691781 | 5.92022408 | 7.516248799 | 11.15610225 | 8.268688598 | 7.420144119 | 7.584437589 | 8.365624471 | 9.122252749 |
| PBANKA_1019500 | 10.13563176 | 12.82086318 | 10.84622418 | 10.37272849 | 9.876299138 | 14.80309209 | 11.57611069 | 10.60325315 | 9.183755053 | 9.685849559 | 10.66804318 |
| PBANKA_1019700 | 10.57525956 | 10.47061017 | 9.671621797 | 9.499423378 | 9.557584375 | 14.11788491 | 10.96394608 | 10.8495766 | 9.492397643 | 9.33576451 | 10.63879507 |
| PBANKA_1020400 | 6.684402608 | 7.239645078 | 6.852775345 | 6.439722146 | 7.272836188 | 11.89508664 | 8.557616754 | 7.175775059 | 7.106940492 | 7.085440754 | 8.380394022 |
| PBANKA_1024100 | 5.615922449 | 8.551057139 | 8.333045204 | 7.932596636 | 9.033519628 | 12.74796747 | 9.486797333 | 8.423206914 | 7.749951897 | 6.876428862 | 7.834665367 |
| PBANKA_1024300 | 8.941230176 | 8.202652382 | 6.709738934 | 6.339364475 | 7.946677551 | 13.00890355 | 9.200335258 | 8.313085828 | 6.969760683 | 8.387725504 | 9.588361847 |
| PBANKA_1025200 | 7.004720845 | 9.156955012 | 8.434105494 | 8.021842998 | 7.397540137 | 11.61756747 | 8.995161916 | 8.288937674 | 6.759689259 | 6.507471633 | 7.022431224 |
| PBANKA_1026200 | 10.06779617 | 9.462188382 | 11.06678296 | 10.4916346 | 10.37784096 | 13.2180414 | 9.738195418 | 9.019007868 | 6.699392931 | 7.850944607 | 9.924949694 |
| PBANKA_1026900 | 8.737918297 | 9.235154075 | 7.683807381 | 7.661648867 | 9.077644749 | 11.930368 | 9.116201739 | 7.424978444 | 8.103812797 | 7.697931121 | 8.853913885 |
| PBANKA_1027500 | 7.07752566 | 7.260638775 | 5.919009783 | 5.685084905 | 5.843749301 | 9.633518352 | 4.807904079 | 4.62754815 | 2.987945008 | 3.676202543 | 3.467532993 |
| PBANKA_1030700 | 0 | 6.582648369 | 6.332347758 | 5.950555602 | 6.074095583 | 10.86400614 | 7.460260359 | 6.438983636 | 7.236042842 | 5.767763935 | 4.800739 |
| PBANKA_1030800 | 0 | 5.425059703 | 4.982262981 | 4.982859447 | 5.897596058 | 8.919596325 | 6.58621909 | 5.931194511 | 3.386473708 | 4.356508828 | 1.416852553 |
| PBANKA_1031000 | 6.052346351 | 7.539208457 | 7.694173631 | 6.651705309 | 7.215337976 | 11.6452833 | 7.624952358 | 7.531285092 | 5.678253412 | 7.363531462 | 4.390945168 |
| PBANKA_1032300 | 5.202116977 | 9.162297096 | 7.721490501 | 6.374498476 | 7.138595557 | 11.78198163 | 9.655961793 | 8.673365907 | 5.9773677 | 9.279787017 | 7.773251519 |
| PBANKA_1033600 | 0 | 7.281423647 | 7.198682533 | 6.120546894 | 5.410713075 | 10.74093773 | 6.894672504 | 5.773063602 | 4.078636318 | 7.315996719 | 3.345430226 |
| PBANKA_1034300 | 6.137271365 | 7.849463066 | 8.539022283 | 8.017937555 | 8.998894226 | 13.75996772 | 9.372011941 | 8.967408749 | 8.654673571 | 9.945532104 | 4.815341013 |
| PBANKA_1034400 | 0 | 8.515071074 | 10.47322446 | 10.76943432 | 10.21148981 | 14.27196962 | 12.14888583 | 12.06719665 | 10.41631621 | 9.563314699 | 8.347752681 |
| PBANKA_1035900 | 9.160311293 | 10.08931067 | 9.055963691 | 8.531472732 | 8.840874612 | 13.1131868 | 9.250655591 | 7.804436763 | 7.792761663 | 9.935580407 | 8.459447986 |
| PBANKA_1036000 | 7.477063777 | 10.42051534 | 10.43356709 | 10.40592498 | 10.34605924 | 13.93048123 | 11.51650611 | 11.79416871 | 11.10509605 | 10.03850025 | 8.240105477 |
| PBANKA_1100860 | 0 | 2.389790243 | 5.40209029 | 4.922878101 | 7.370859002 | 11.73795251 | 7.771760286 | 6.979749754 | 5.914949235 | 8.921212844 | 6.953887101 |
| PBANKA_1101100 | 0 | 7.24234407 | 6.577078289 | 5.700652572 | 10.24639469 | 14.86516827 | 10.68062068 | 8.144882596 | 8.619478752 | 7.286764205 | 6.901328406 |
| PBANKA_1101300 | 0 | 2.506242444 | 5.562776681 | 5.626817737 | 9.808279901 | 14.9026501 | 10.80808886 | 9.8993165 | 9.60760966 | 8.236355165 | 4.280277964 |
| PBANKA_1102200 | 0 | 6.411070812 | 7.454665439 | 7.835781028 | 10.69696988 | 14.64123483 | 12.58583712 | 11.71293157 | 10.81963278 | 8.821510454 | 6.100724528 |
| PBANKA_1103000 | 2.492651273 | 9.917710767 | 8.544801021 | 7.878848315 | 8.685660422 | 12.68897764 | 10.58354373 | 9.686149566 | 8.062615511 | 9.233019322 | 6.600899594 |
| PBANKA_1104600 | 8.241793633 | 8.468112047 | 7.898003207 | 7.887638869 | 8.569513801 | 12.3042794 | 8.626794747 | 8.389681264 | 7.857606674 | 8.028942343 | 9.914687571 |
| PBANKA_1105700 | 7.302829793 | 9.409214624 | 7.590321739 | 6.990967868 | 7.477118845 | 11.48836292 | 9.375213521 | 7.4909142 | 5.861354153 | 8.071878992 | 6.78979651 |
| PBANKA_1105800 | 7.363400983 | 7.578002653 | 7.048596542 | 6.209539244 | 7.289311875 | 11.44025422 | 7.010724954 | 6.804752106 | 5.456721577 | 7.320698703 | 9.370694355 |
| PBANKA_1122500 | 7.947002482 | 7.204627972 | 8.349249948 | 8.126851526 | 8.649037232 | 11.66471468 | 8.051652218 | 7.733771358 | 7.036072762 | 7.692784254 | 9.533512427 |
| PBANKA_1122700 | 7.666537985 | 8.890519556 | 8.449275444 | 8.407643997 | 11.62787143 | 15.84205806 | 10.89961731 | 10.26122521 | 10.20980396 | 9.344586828 | 12.34206648 |
| PBANKA_1127000 | 5.355436369 | 10.57442916 | 10.04260859 | 10.71536602 | 12.27082676 | 15.70500595 | 10.83458323 | 10.03068736 | 10.85099808 | 8.359565977 | 8.3406791 |
| PBANKA_1129300 | 3.220574347 | 6.690273882 | 5.759333852 | 5.318887639 | 5.981692512 | 10.47674864 | 7.843932919 | 6.101613546 | 4.504339414 | 6.24487787 | 6.423848314 |
| PBANKA_1129700 | 9.480842655 | 7.208328158 | 7.534510217 | 7.481732385 | 8.434792376 | 12.44082484 | 7.63256057 | 6.821737053 | 6.775777715 | 6.614775599 | 9.408687435 |
| PBANKA_1130900 | 8.692926846 | 9.682226035 | 9.128451361 | 9.042737077 | 9.694342086 | 14.23497325 | 11.19729898 | 9.813805177 | 9.602478944 | 11.09674651 | 10.83470798 |
| PBANKA_1134800 | 7.245766245 | 8.311494414 | 6.910406491 | 6.073655239 | 6.443932422 | 10.61115353 | 8.275492856 | 7.063403578 | 5.278879425 | 6.619522952 | 5.525433747 |
| PBANKA_1135300 | 10.46999504 | 10.41516895 | 9.300595499 | 8.424586101 | 9.31538552 | 12.98072595 | 10.34838324 | 9.42748705 | 7.526371082 | 8.110240257 | 8.402396316 |
| PBANKA_1135700 | 10.48196454 | 11.26362823 | 10.71507371 | 10.35576127 | 10.19054301 | 14.80515587 | 11.30859685 | 10.22824834 | 9.445324558 | 9.169998493 | 9.428175136 |
| PBANKA_1136300 | 0 | 7.615795516 | 6.93998449 | 7.236576898 | 7.770131212 | 10.93215028 | 7.96899359 | 6.239016224 | 7.975567594 | 7.61862807 | 4.645947365 |
| PBANKA_1141100 | 6.471184576 | 6.582648369 | 5.91282231 | 7.059562991 | 7.86585827 | 12.56500761 | 7.878765388 | 6.282454206 | 6.322672321 | 6.866688749 | 7.082176175 |
| PBANKA_1143500 | 7.122564447 | 8.45062971 | 7.017729586 | 6.030104937 | 6.366165339 | 11.42119237 | 7.796978293 | 6.117679156 | 5.429946088 | 6.475112446 | 6.101187269 |
| PBANKA_1145400 | 0 | 6.771251674 | 6.817021391 | 6.490301747 | 10.85843218 | 15.31056886 | 10.87771387 | 7.939469565 | 9.805190696 | 7.202753965 | 4.723181493 |
| PBANKA_1145700 | 0 | 5.722047314 | 5.941989501 | 5.934077061 | 4.869607228 | 10.02848864 | 7.453250493 | 6.324825564 | 5.894087941 | 6.87533422 | 4.441384697 |
| PBANKA_1145800 | 0 | 1.851885259 | 4.381452081 | 5.39308555 | 8.163338385 | 14.97924314 | 12.67713969 | 11.5770858 | 9.977395393 | 9.522846793 | 6.963005834 |
| PBANKA_1145900 | 0 | 5.896596185 | 5.806363381 | 5.668152826 | 8.012744561 | 15.42421027 | 12.80908042 | 11.46779368 | 9.191873372 | 8.084700393 | 5.21990475 |
| PBANKA_1200600 | 8.350443234 | 5.858625722 | 5.54086659 | 6.390524773 | 9.092469975 | 15.20778183 | 9.827018259 | 7.900598755 | 10.70176213 | 7.366811773 | 7.991442977 |
| PBANKA_1200800 | 8.385862244 | 9.9900682 | 8.924226409 | 9.157645068 | 9.266611407 | 12.86511406 | 8.869874254 | 8.603858527 | 8.054787907 | 9.359714254 | 8.507836015 |
| PBANKA_1207000 | 10.35677365 | 9.859752 | 7.951614632 | 6.550514141 | 8.72310789 | 13.4980896 | 9.507223482 | 8.269150704 | 6.967658115 | 7.561657255 | 9.097233175 |
| PBANKA_1207100 | 8.502128065 | 6.258365047 | 6.420381926 | 6.66191504 | 10.11620401 | 14.38330814 | 7.706702128 | 7.146054299 | 7.014120799 | 8.175094553 | 7.103451068 |
| PBANKA_1207900 | 0 | 1.851885259 | 3.648173836 | 4.931784834 | 10.22687385 | 13.23875364 | 7.595427266 | 5.436710626 | 8.898305291 | 6.786895472 | 6.449876033 |
| PBANKA_1208700 | 0 | 7.310099011 | 6.225937589 | 6.241951336 | 7.237311419 | 12.41750073 | 8.163458786 | 6.039077139 | 6.336424719 | 7.066762206 | 6.695470397 |
| PBANKA_1211500 | 7.968143601 | 9.083663 | 8.207206403 | 7.468787463 | 7.976454176 | 11.85773399 | 8.944682151 | 8.29361818 | 6.652304598 | 7.119419563 | 8.300716511 |
| PBANKA_1216900 | 3.022133598 | 7.042417061 | 7.665377887 | 7.714604016 | 7.752390438 | 11.23134166 | 8.891528134 | 8.417136245 | 8.435610308 | 7.423284255 | 5.968000867 |
| PBANKA_1221700 | 5.894539599 | 2.694476423 | 6.463169494 | 5.928722312 | 6.565817825 | 10.1094449 | 6.888225595 | 5.858968053 | 4.801379426 | 4.771461446 | 7.5295043 |
| PBANKA_1223300 | 0 | 6.611312046 | 7.824552566 | 8.613156808 | 8.139115316 | 11.25977283 | 8.521286512 | 8.111527867 | 7.346682728 | 7.647897146 | 7.314217808 |
| PBANKA_1227100 | 8.382363913 | 8.814682398 | 7.511406865 | 6.6099076 | 8.013551952 | 13.03318819 | 10.78385031 | 9.447539745 | 9.03295462 | 9.148694426 | 10.38983589 |
| PBANKA_1227900 | 7.658477166 | 8.8849551 | 7.957252909 | 7.136961916 | 8.513320912 | 12.54781863 | 9.298460677 | 8.427047281 | 6.647475816 | 6.913692092 | 8.919801941 |
| PBANKA_1228700 | 7.983987441 | 7.703026962 | 6.875884316 | 6.283542835 | 7.680153891 | 11.54091383 | 8.320297094 | 7.104668523 | 6.581997622 | 7.47620838 | 9.118567359 |
| PBANKA_1229000 | 6.454946663 | 7.545593755 | 7.536147726 | 6.884540651 | 10.07183963 | 16.2447802 | 11.49006405 | 10.4397447 | 8.206890354 | 7.333045758 | 4.684193056 |
| PBANKA_1230500 | 5.977629114 | 8.296384895 | 7.51136446 | 7.478623122 | 7.146102492 | 10.85771668 | 7.271202143 | 6.845160983 | 5.300659597 | 6.752449268 | 6.164894308 |
| PBANKA_1231600 | 5.642496565 | 7.478553062 | 10.41262386 | 11.32554015 | 11.44692221 | 14.86801168 | 11.55437084 | 10.52841817 | 10.91595206 | 10.630101 | 9.297238364 |
| PBANKA_1232100 | 0 | 10.19574281 | 8.97281096 | 8.501392269 | 9.594107926 | 13.59945092 | 10.41735505 | 10.12084677 | 8.991078341 | 8.061577721 | 7.246749698 |
| PBANKA_1239700 | 8.176924223 | 8.225504832 | 8.094895846 | 7.140134124 | 6.980789209 | 10.3342582 | 8.262158843 | 7.967738775 | 5.821655222 | 6.361924168 | 5.291157039 |
| PBANKA_1302500 | 5.940184956 | 7.275648369 | 7.07901786 | 6.999574662 | 6.628846848 | 9.648199728 | 7.129417531 | 6.891839282 | 6.25865348 | 6.15912571 | 4.714830876 |
| PBANKA_1309500 | 10.03470736 | 10.00529941 | 8.970897237 | 8.612467928 | 8.073158074 | 12.09763775 | 10.03963119 | 9.774436973 | 8.312285525 | 7.390372026 | 7.545658141 |
| PBANKA_1311800 | 8.349635354 | 8.869078583 | 7.872133016 | 7.316574454 | 7.741917414 | 12.18648016 | 9.287324632 | 8.014379438 | 7.203539396 | 7.655168871 | 9.844201144 |
| PBANKA_1313000 | 7.397309627 | 7.645478425 | 7.75412349 | 7.232487098 | 7.933312553 | 12.47380279 | 7.875206199 | 6.917042164 | 6.66235707 | 7.686173955 | 8.558496827 |
| PBANKA_1313400 | 8.086914017 | 7.455040055 | 6.877081136 | 6.424946118 | 8.060074354 | 11.84141397 | 9.268378975 | 7.358577908 | 7.476544096 | 8.672485152 | 9.301406807 |
| PBANKA_1313500 | 6.993494099 | 2.876537653 | 5.950006125 | 6.088664598 | 7.254980192 | 9.521890997 | 5.256018165 | 5.543704019 | 6.067474348 | 6.027100407 | 7.110743838 |
| PBANKA_1314900 | 11.15915818 | 7.511069611 | 8.368368332 | 7.707959271 | 10.18475521 | 14.13230263 | 9.808048115 | 8.965173217 | 8.822318394 | 11.56729305 | 7.592869743 |
| PBANKA_1317600 | 9.982023963 | 8.220898326 | 8.287132134 | 7.757479377 | 8.640845055 | 12.87521666 | 8.050751608 | 7.29773059 | 7.492166867 | 7.655908643 | 10.37275037 |
| PBANKA_1317900 | 8.282650802 | 9.142218305 | 8.420453103 | 7.962654474 | 7.516138933 | 11.74741461 | 9.244248177 | 9.050317603 | 7.126529159 | 7.603156727 | 9.074732615 |
| PBANKA_1320500 | 6.722999319 | 9.967267685 | 9.641839097 | 8.741570134 | 11.24577549 | 14.79359811 | 11.49709471 | 10.82861079 | 9.751452337 | 9.113729477 | 7.584737963 |
| PBANKA_1320600 | 0 | 6.117478055 | 5.334245105 | 4.518951527 | 1.505964805 | 8.405752488 | 5.436510556 | 4.096104559 | 3.419087856 | 4.125988767 | 1.730419991 |
| PBANKA_1321400 | 9.640827823 | 9.732120957 | 8.361191445 | 8.010546608 | 8.902488153 | 13.5128366 | 10.88412247 | 9.487683914 | 8.164570729 | 8.284834362 | 9.881705286 |
| PBANKA_1323000 | 2.709465703 | 3.120026423 | 6.273475891 | 6.572997979 | 6.723523719 | 10.21953967 | 7.455202248 | 6.692849993 | 6.24158902 | 5.358144841 | 5.454812625 |
| PBANKA_1323700 | 0 | 6.422807762 | 6.54624079 | 6.036352928 | 6.720926906 | 11.03317517 | 7.19930765 | 6.60423104 | 5.803464517 | 6.097351687 | 6.023251342 |
| PBANKA_1324900 | 6.286709864 | 6.922900536 | 5.982958198 | 5.285460282 | 5.929378845 | 10.80103336 | 7.240093775 | 5.958883707 | 5.313516143 | 7.24165367 | 7.374055185 |
| PBANKA_1327021 | 6.821153401 | 7.584838564 | 7.717797294 | 8.452342826 | 9.68368012 | 12.06799462 | 9.697643769 | 9.81961898 | 10.04949283 | 7.20840196 | 5.233228544 |
| PBANKA_1327251 | 2.851146841 | 5.031175672 | 6.337903419 | 6.525291702 | 8.700651311 | 11.78557938 | 8.428750658 | 7.317669525 | 7.695018902 | 6.350661978 | 4.796855397 |
| PBANKA_1327300 | 0 | 8.074354246 | 8.209234254 | 8.300070175 | 10.30066613 | 13.11792217 | 9.591257692 | 9.146699299 | 9.009002098 | 6.669370984 | 2.040988026 |
| PBANKA_1329600 | 6.902286118 | 3.142629976 | 6.781313018 | 6.857788535 | 9.105996022 | 13.68416205 | 9.248472121 | 8.233223254 | 8.354355746 | 9.683794122 | 6.345993157 |

| | | | | | | | | | | | |
|---|---|---|---|---|---|---|---|---|---|---|---|
| PBANKA_1332500 | 5.514156064 | 7.291595727 | 6.338301257 | 5.457490658 | 6.370382652 | 11.14644519 | 8.378138761 | 7.340966908 | 5.979693538 | 7.056160655 | 7.570522113 |
| PBANKA_1334200 | 6.794720537 | 8.558050267 | 7.38925417 | 6.702352033 | 7.446735097 | 11.73920552 | 8.936003537 | 7.875737239 | 5.78318794 | 6.708046275 | 9.713382341 |
| PBANKA_1334300 | 6.142940992 | 10.69729555 | 9.70809786 | 8.992640136 | 11.4449651 | 15.20468089 | 11.15019946 | 10.44973082 | 9.0721892 | 7.340023379 | 6.365438251 |
| PBANKA_1340300 | 0 | 9.283107618 | 7.767086071 | 6.738457099 | 6.590814274 | 11.92641945 | 9.639326385 | 9.107534771 | 5.874994788 | 6.527693562 | 3.629207346 |
| PBANKA_1342400 | 2.298831494 | 6.668574156 | 6.684041095 | 5.988296171 | 6.940589448 | 9.895280455 | 7.692975515 | 6.442018274 | 6.370828912 | 7.278334428 | 4.944578417 |
| PBANKA_1342800 | 2.298831494 | 6.401585837 | 5.948383089 | 5.719392318 | 8.032743731 | 10.55890367 | 7.040302783 | 5.878689528 | 6.258764995 | 6.157112508 | 5.778579819 |
| PBANKA_1343000 | 0 | 9.726662464 | 8.684609736 | 7.704255411 | 8.120941134 | 13.42851107 | 9.131791719 | 8.973067535 | 6.057392688 | 8.302375979 | 6.564683101 |
| PBANKA_1344500 | 5.121927917 | 9.440044758 | 10.72550296 | 10.89744573 | 11.57584038 | 16.21644903 | 11.21050512 | 10.30961812 | 10.03378454 | 8.770740044 | 8.832952602 |
| PBANKA_1346100 | 5.512607788 | 7.534097091 | 7.515792373 | 6.771788015 | 6.780833918 | 10.79792008 | 8.636339037 | 8.118062917 | 6.477053707 | 7.212855563 | 6.989023564 |
| PBANKA_1348300 | 10.14256651 | 9.539008128 | 8.920449201 | 8.309481301 | 8.746932459 | 12.63601136 | 8.8954626 | 8.287862711 | 6.869719644 | 7.486955143 | 10.58418056 |
| PBANKA_1349200 | 5.533064502 | 8.367138902 | 7.224147863 | 7.684105338 | 8.471380708 | 13.23482889 | 11.19023911 | 10.18840524 | 10.60358625 | 10.09138283 | 7.633675146 |
| PBANKA_1356200 | 7.660068207 | 8.145141057 | 7.984004645 | 7.803364263 | 7.892349959 | 11.2477123 | 7.681110077 | 6.462193656 | 6.619017241 | 6.629651453 | 3.982060334 |
| PBANKA_1356700 | 0 | 7.804902059 | 8.617436437 | 8.805817446 | 9.043897233 | 13.28890955 | 9.896951216 | 8.955029254 | 9.38198794 | 8.22867258 | 5.473655444 |
| PBANKA_1358000 | 0 | 7.974420195 | 7.64933104 | 6.970654084 | 8.448100454 | 10.53591262 | 7.860786849 | 7.151622767 | 7.260982659 | 7.312141434 | 6.373577742 |
| PBANKA_1360100 | 2.492651273 | 7.772825412 | 8.447855924 | 8.125403922 | 8.589336879 | 12.74983248 | 10.03288425 | 10.49582449 | 7.914019837 | 6.557745834 | 4.811242448 |
| PBANKA_1363000 | 6.133331183 | 7.170954365 | 7.143623518 | 6.611268398 | 5.781202958 | 9.162661648 | 5.912089206 | 5.731683808 | 5.178173942 | 5.73615865 | 6.932143739 |
| PBANKA_1363100 | 8.181769887 | 7.25989222 | 6.77633264 | 6.503189817 | 7.029847332 | 11.16258726 | 8.950526676 | 7.283338336 | 7.242347804 | 7.982718623 | 8.72289083 |
| PBANKA_1365600 | 0 | 2.389790243 | 4.441637618 | 4.866833276 | 5.781202958 | 11.08955058 | 6.590661277 | 6.369291959 | 6.51837224 | 5.557818484 | 4.984228662 |
| PBANKA_1401100 | 7.749133065 | 8.136254931 | 7.740032251 | 7.336543853 | 7.249173022 | 10.48528243 | 8.05731619 | 7.629122109 | 6.678532391 | 5.376484094 | 6.499760186 |
| PBANKA_1401200 | 8.646103388 | 9.040820758 | 8.237336791 | 7.67315728 | 8.200031545 | 12.87156137 | 9.70267083 | 8.288193682 | 7.230559235 | 6.457558378 | 6.807555161 |
| PBANKA_1401900 | 6.476250349 | 8.18950741 | 8.004287206 | 7.508413879 | 7.533452034 | 11.73160264 | 8.120901764 | 6.933280899 | 6.240177251 | 5.952392811 | 5.536221232 |
| PBANKA_1406700 | 9.069783702 | 11.53126895 | 10.50492842 | 9.74845064 | 9.543336906 | 13.80506596 | 11.42707077 | 11.38483358 | 8.641948274 | 8.68129465 | 9.343048149 |
| PBANKA_1407400 | 9.357993433 | 9.343281539 | 9.345459589 | 9.21132057 | 9.406054716 | 13.12027738 | 10.45265949 | 10.36108607 | 9.769039466 | 10.55028994 | 10.17497335 |
| PBANKA_1407800 | 6.760406304 | 8.247188314 | 8.128540165 | 7.66830596 | 7.590373623 | 11.34111038 | 9.140979888 | 8.466361572 | 8.107974074 | 9.193383197 | 5.640798715 |
| PBANKA_1408800 | 6.443888621 | 2.250849715 | 5.64828461 | 5.739804368 | 5.547612861 | 9.085969171 | 5.638973053 | 5.004280774 | 4.256611912 | 5.984413512 | 6.505336475 |
| PBANKA_1409300 | 5.154194688 | 8.425982872 | 8.152644833 | 7.22167545 | 7.56368101 | 11.72001898 | 8.573312353 | 8.331639957 | 5.925040245 | 6.665049031 | 8.67731878 |
| PBANKA_1412100 | 0 | 5.992243651 | 5.66374429 | 4.500498262 | 3.971662036 | 9.296007856 | 7.117495478 | 7.302904338 | 4.718287668 | 4.299669596 | 2.744183862 |
| PBANKA_1416400 | 9.375966778 | 7.864234205 | 7.48230777 | 6.437371798 | 7.307124093 | 12.00375215 | 9.140099153 | 7.95977065 | 6.571592079 | 8.182561506 | 8.207551262 |
| PBANKA_1420500 | 7.908702984 | 11.1189962 | 10.15584452 | 9.744143289 | 10.0345186 | 14.97861529 | 11.57985374 | 11.1842417 | 9.792716485 | 9.852846413 | 9.954319615 |
| PBANKA_1423200 | 8.261561374 | 8.206681766 | 7.576443468 | 7.087706368 | 7.922042837 | 11.63938293 | 8.66029878 | 7.453447858 | 5.971614396 | 7.249498076 | 8.235647288 |
| PBANKA_1424300 | 10.18906208 | 12.63513166 | 11.26377693 | 10.71062892 | 11.21912203 | 15.75879384 | 11.68085914 | 11.36921732 | 9.755488438 | 10.15421835 | 13.20226696 |
| PBANKA_1424600 | 2.912756836 | 6.565017369 | 6.65376189 | 6.886742823 | 8.737602632 | 12.45220084 | 9.518868058 | 8.086477857 | 8.450534672 | 8.280960406 | 8.626266666 |
| PBANKA_1424800 | 0 | 6.310786671 | 6.018984744 | 5.29044971 | 4.970893001 | 9.184469204 | 6.33737023 | 6.308119183 | 4.586022048 | 5.934508682 | 4.810699529 |
| PBANKA_1424900 | 0 | 5.840490719 | 6.564845308 | 6.334131464 | 5.263967929 | 9.760541832 | 6.6959194 | 6.277395502 | 4.852210721 | 6.725110258 | 4.92678407 |
| PBANKA_1425000 | 0 | 9.529071868 | 9.288193697 | 9.278820716 | 8.876306274 | 13.25502562 | 10.70294841 | 10.6867635 | 9.052686725 | 7.429977862 | 6.053297847 |
| PBANKA_1428900 | 10.58512088 | 9.193934864 | 8.589482428 | 8.643792882 | 9.657630841 | 13.94776691 | 10.22373864 | 8.962356019 | 9.264429737 | 8.905129557 | 8.867084005 |
| PBANKA_1432800 | 8.395258429 | 8.907615316 | 7.80230708 | 7.465116821 | 8.379203594 | 12.11121306 | 9.806825016 | 8.185893014 | 7.397023292 | 7.687583232 | 9.991146098 |
| PBANKA_1433100 | 7.729338046 | 6.6698627 | 6.996351208 | 6.918722944 | 8.015157301 | 11.73803127 | 8.678611501 | 8.429038566 | 8.163482269 | 7.484207877 | 7.812241856 |
| PBANKA_1434100 | 7.425815614 | 7.63320657 | 8.287336258 | 8.137212067 | 8.673594296 | 12.91400234 | 8.792631832 | 8.872671528 | 8.339397128 | 8.175622124 | 6.642501931 |
| PBANKA_1437100 | 0 | 2.694476423 | 6.618882299 | 7.209654128 | 10.03527569 | 12.69864629 | 10.35237687 | 8.340752 | 9.531571882 | 8.517531852 | 5.266924626 |
| PBANKA_1439200 | 12.14857457 | 12.35168409 | 11.64686612 | 11.27483212 | 12.23788849 | 17.27871037 | 14.20867658 | 13.35250543 | 12.563789 | 12.85367827 | 12.05070885 |
| PBANKA_1439800 | 2.492651273 | 5.89334475 | 6.25523172 | 6.713690813 | 6.32389105 | 9.478146297 | 6.12462335 | 5.718062696 | 5.790488836 | 6.067281938 | 4.763885771 |
| PBANKA_1440100 | 7.581857915 | 8.129320881 | 8.136614933 | 8.261286786 | 8.306763426 | 12.45624497 | 8.977291317 | 8.521049282 | 8.427372476 | 8.073257647 | 6.395095726 |
| PBANKA_1440200 | 7.075517817 | 7.060581608 | 6.47101475 | 5.970382136 | 6.590812332 | 11.01679717 | 7.528256961 | 6.262289502 | 5.851829576 | 6.375373819 | 5.134628189 |
| PBANKA_1441500 | 8.768032595 | 8.858534792 | 7.856214261 | 7.409558353 | 7.866069663 | 12.02438248 | 8.772475595 | 7.737185816 | 6.865323071 | 7.173661826 | 9.175589468 |
| PBANKA_1441700 | 0 | 8.539921819 | 8.565969562 | 8.070750268 | 7.589857364 | 12.46865891 | 9.61925655 | 8.499069604 | 7.918998067 | 6.655317525 | 3.660396736 |
| PBANKA_1442400 | 0 | 7.050899332 | 6.311985829 | 5.988296171 | 8.344282106 | 11.71265249 | 7.762318003 | 6.514556059 | 8.848976736 | 6.195572818 | 4.216402973 |
| PBANKA_1442500 | 8.532059062 | 8.830164794 | 8.063642102 | 7.616176848 | 8.050088269 | 11.80981722 | 9.555334231 | 8.542581058 | 6.80320505 | 7.417226495 | 8.212221179 |
| PBANKA_1445200 | 7.087647518 | 3.461220738 | 6.717501423 | 6.139648148 | 6.384686652 | 10.34285228 | 7.229934812 | 6.489181349 | 4.382152628 | 5.278434843 | 7.698718106 |
| PBANKA_1445400 | 8.144123679 | 10.11326541 | 10.09680127 | 10.62571839 | 10.48175089 | 13.06366575 | 9.876819817 | 9.538738205 | 9.465924432 | 8.656680028 | 7.783925261 |
| PBANKA_1449400 | 7.458215194 | 8.558050267 | 9.219547394 | 8.774886949 | 8.546511933 | 11.95655519 | 9.601976674 | 9.409993084 | 8.146667631 | 8.930141493 | 9.007195769 |
| PBANKA_1452300 | 8.667019386 | 9.209933134 | 8.796410445 | 8.600532263 | 8.448261364 | 12.32196306 | 9.169357141 | 9.368559812 | 7.723446552 | 7.695332451 | 8.203676625 |
| PBANKA_1455200 | 9.339704773 | 9.240184594 | 8.782317076 | 8.011859647 | 8.997897209 | 12.7782371 | 9.297823779 | 8.060395419 | 7.680851652 | 6.801450689 | 8.753126425 |
| PBANKA_1455300 | 8.058174755 | 8.597756428 | 8.631310313 | 8.556849143 | 8.470841518 | 11.2246268 | 8.827276169 | 8.700927146 | 8.541526288 | 7.614024461 | 8.818447634 |
| PBANKA_1463800 | 0 | 7.068453971 | 7.45287554 | 8.417223591 | 8.246715731 | 10.98413714 | 8.222214115 | 7.01922434 | 7.354577459 | 5.251956674 | 6.25034171 |
| PBANKA_1464000 | 0 | 2.80912216 | 4.739364399 | 5.491779399 | 5.80677847 | 9.996953987 | 7.282539614 | 6.677388653 | 6.919924378 | 6.280300952 | 5.907382997 |
| PBANKA_1464500 | 8.380508851 | 9.131202939 | 8.210467167 | 7.763336875 | 9.421951739 | 13.19393523 | 9.82176461 | 8.400454376 | 9.0653583 | 8.542578876 | 8.216129771 |
| PBANKA_1464600 | 0 | 6.509381228 | 6.486947362 | 6.536980424 | 6.789200102 | 11.64969308 | 9.045451939 | 8.171256633 | 7.236633977 | 7.98636806 | 6.075002543 |
| PBANKA_0300600 | 4.918017349 | 3.084222589 | 4.972737433 | 6.026685954 | 10.20284397 | 15.93726823 | 11.6749983 | 10.37292784 | 9.189536968 | 9.135696811 | 8.07429696 |
| PBANKA_0418500 | 0 | 2.845509661 | 4.929222656 | 4.82521256 | 10.90668346 | 14.99096343 | 8.230245941 | 5.089908432 | 8.952017765 | 4.592858084 | 1.476346564 |
| PBANKA_0100300 | 0 | 0 | 1.357993684 | 4.63579707 | 1.959060306 | 9.202431579 | 6.713925507 | 5.076714069 | 4.603004488 | 4.590992573 | 0.849622504 |
| PBANKA_0100500 | 0 | 0 | 4.249638257 | 5.734624328 | 7.905283646 | 12.38402532 | 8.512055446 | 7.804481886 | 7.569953659 | 7.299822368 | 3.314581484 |
| PBANKA_0112661 | 0 | 0 | 1.248106325 | 0 | 7.422901945 | 11.24725001 | 6.433194257 | 4.181690836 | 6.384987088 | 4.172928626 | 1.011541306 |
| PBANKA_0112721 | 0 | 0 | 1.612918338 | 0 | 6.579905009 | 11.59726762 | 7.633671961 | 6.886260459 | 5.080485649 | 7.1286562 | 4.248886309 |
| PBANKA_0201300 | 0 | 0 | 6.254298117 | 5.31546899 | 8.724952504 | 11.35145394 | 7.061254713 | 6.341009911 | 7.793792209 | 5.507161967 | 5.110754235 |
| PBANKA_0214600 | 0 | 0 | 7.123396613 | 7.441368236 | 10.4257409 | 14.86301746 | 9.936945597 | 8.701820399 | 8.295792756 | 6.600778959 | 4.25422019 |
| PBANKA_0316300 | 0 | 0 | 3.545515798 | 4.818902379 | 7.246768935 | 12.48974922 | 8.911496008 | 6.469748587 | 8.747058466 | 7.02536296 | 7.085088672 |
| PBANKA_0524100 | 0 | 0 | 3.779075852 | 4.626607899 | 7.764416731 | 11.28379194 | 8.57246567 | 6.851150098 | 5.671159271 | 5.606665548 | 4.222141831 |
| PBANKA_0524200 | 2.823096423 | 0 | 6.392452432 | 7.434753606 | 9.885583254 | 13.77727919 | 10.08221031 | 9.761630758 | 10.17380585 | 9.035805155 | 7.217969907 |
| PBANKA_0600041 | 0 | 0 | 1.800899129 | 1.38110124 | 5.081320801 | 9.693207483 | 6.323601407 | 4.946291348 | 5.014370826 | 4.283361349 | 1.112855868 |
| PBANKA_0623150 | 0 | 0 | 1.248106325 | 4.046040668 | 5.974915446 | 11.51485416 | 6.940888121 | 5.352367635 | 4.838566099 | 7.823646099 | 5.001221838 |
| PBANKA_0700061 | 0 | 0 | 1.357993684 | 1.38110124 | 1.722664483 | 9.288387198 | 4.052214172 | 3.22588984 | 2.086783656 | 5.998863435 | 0.577163707 |
| PBANKA_0800300 | 0 | 0 | 4.655414553 | 4.8009773 | 6.237970719 | 11.47653976 | 9.128949732 | 6.682748432 | 5.3395839 | 8.080973896 | 8.157471471 |
| PBANKA_0836200 | 0 | 0 | 3.899806758 | 3.725902556 | 10.11109647 | 14.37774854 | 9.257534199 | 8.819201923 | 7.380575397 | 6.035155381 | 3.690009189 |
| PBANKA_0931200 | 6.684187788 | 0 | 5.090721873 | 5.86713801 | 11.78081057 | 14.24051874 | 11.10691537 | 10.34966193 | 10.66569571 | 9.199111778 | 6.507975859 |
| PBANKA_1000500 | 0 | 0 | 5.895287723 | 6.007259431 | 6.70187005 | 12.30646206 | 9.112717421 | 8.250994153 | 7.113839582 | 7.181197215 | 4.993661923 |
| PBANKA_1000600 | 0 | 0 | 4.512712846 | 3.959501477 | 9.418021451 | 14.950878 | 9.931904019 | 9.929258605 | 7.535225241 | 8.050066257 | 5.943091861 |
| PBANKA_1220200 | 2.728828613 | 0 | 4.347002694 | 5.118557365 | 9.418021451 | 13.73848437 | 10.95344384 | 10.03853146 | 7.839532106 | 7.26399523 | 4.168986524 |
| PBANKA_1246161 | 0 | 0 | 4.125974258 | 1.783138751 | 8.440685694 | 13.61601233 | 9.514599292 | 8.580793189 | 6.738956395 | 8.872713979 | 6.661550013 |
| PBANKA_1300031 | 0 | 0 | 1.800899129 | 1.38110124 | 4.602650265 | 9.527328592 | 4.22760063 | 2.821505574 | 2.225036267 | 2.595438894 | 1.476346564 |
| PBANKA_1351500 | 0 | 0 | 3.666246704 | 5.015835649 | 8.518505479 | 11.00010558 | 6.33267164 | 2.589036715 | 8.141849391 | 6.405708537 | 7.569344956 |
| PBANKA_1400700 | 0 | 0 | 6.879793412 | 9.980362243 | 10.9134289 | 14.02627651 | 11.60269786 | 10.40793652 | 11.40154439 | 9.225522005 | 9.067287404 |
| PBANKA_1465000 | 5.355436369 | 0 | 1.854052822 | 2.008240071 | 9.075380243 | 13.29206159 | 7.658921907 | 2.953023874 | 7.046865446 | 6.239502714 | 6.606340065 |
| PBANKA_1465600 | 0 | 0 | 3.898139195 | 4.275162073 | 5.5760658 | 9.088940689 | 4.614914786 | 3.674914365 | 1.496060917 | 4.901118246 | 0.507714685 |
| PBANKA_1466000 | 0 | 0 | 1.453333024 | 1.299810393 | 1.959806009 | 8.31953338 | 5.426136647 | 4.899786416 | 3.586209044 | 3.775689037 | 0 |
| PBANKA_0600021 | 0 | 0 | 0 | 1.520480378 | 5.809153315 | 10.91591296 | 5.714460503 | 4.65505758 | 3.717744015 | 3.802784261 | 0.507714685 |
| PBANKA_0600031 | 0 | 0 | 0 | 1.38110124 | 6.333692524 | 12.3756365 | 3.918140565 | 2.282853937 | 0.716602979 | 4.945174043 | 0.974340119 |
| PBANKA_0007501 | 0 | 0 | 0 | 0 | 4.068156159 | 9.957597894 | 2.010666266 | 0.336221246 | 0 | 0 | 0 |
| PBANKA_0008101 | 0 | 0 | 0 | 0 | 5.913103048 | 11.14088699 | 4.232942996 | 3.186401715 | 1.95375497 | 2.039398495 | 0.640509685 |
| PBANKA_0316500 | 0 | 0 | 0 | 0 | 5.924256102 | 12.57209826 | 7.3124546 | 3.496855009 | 3.532233783 | 4.209177051 | 1.046917684 |
| PBANKA_0316921 | 0 | 0 | 0 | 0 | 6.759187554 | 10.78272716 | 8.296199472 | 6.576111199 | 6.066491385 | 3.297037842 | 0 |
| PBANKA_0500600 | 0 | 0 | 0 | 0 | 5.857378227 | 10.00312339 | 6.316453805 | 4.714420436 | 4.563972766 | 1.897239464 | 0.507714685 |
| PBANKA_0500700 | 0 | 0 | 0 | 0 | 7.838598303 | 11.59486086 | 8.494378167 | 7.445335496 | 6.084167854 | 4.257558542 | 1.305293599 |
| PBANKA_0600061 | 0 | 0 | 0 | 0 | 3.705002078 | 9.237494012 | 6.870853209 | 5.642325738 | 5.350534614 | 4.763973549 | 1.229009792 |
| PBANKA_0700081 | 0 | 0 | 0 | 0 | 4.510683365 | 9.966091399 | 5.290144932 | 4.387952266 | 2.25961036 | 5.537686235 | 1.352007024 |
| PBANKA_1000051 | 0 | 0 | 0 | 0 | 6.120818973 | 9.2514581 | 6.27956149 | 5.451578762 | 4.109212388 | 1.629602026 | 0.640509685 |

| | | | | | | | | | | | |
|---|---|---|---|---|---|---|---|---|---|---|---|
| PBANKA_1246100 | 0 | 0 | 0 | 0 | 5.003380545 | 9.105635489 | 4.561461122 | 2.971162287 | 4.845704567 | 2.339563907 | 0 |
| PBANKA_1400051 | 0 | 0 | 0 | 0 | 1.62242156 | 9.070052629 | 7.013554102 | 5.610119136 | 2.848626379 | 3.757091343 | 0 |
| PBANKA_0837141 | 0 | 0 | 0 | 0 | 1.959060306 | 9.143891969 | 5.460410959 | 4.253752484 | 2.771030554 | 4.245398741 | 0.93511558 |
| PBANKA_0623651 | 0 | 0 | 0 | 0 | 0 | 7.421324023 | 4.088387924 | 4.447437245 | 2.920282212 | 4.672467309 | 0 |
| PBANKA_0944101 | 0 | 0 | 0 | 0 | 0 | 8.630660518 | 5.017440769 | 3.98418863 | 2.836969845 | 3.046760723 | 0 |
| PBANKA_1146721 | 0 | 0 | 0 | 0 | 0 | 8.10524887 | 5.610913622 | 4.328367112 | 2.94368453 | 4.692102195 | 3.750993775 |
| PBANKA_1300011 | 0 | 0 | 0 | 0 | 0 | 8.188038985 | 2.23642561 | 1.76874602 | 0 | 1.839055188 | 0 |
